# Effects of manganese dioxide on macrophages under different exposure schemes

**DOI:** 10.1101/2025.09.12.675790

**Authors:** Marianne Vitipon, Esther Akingbagbohun, Véronique Collin-Faure, Fabienne Devime, Hélène Diemer, Daphna Fenel, Christine Carapito, Stéphane Ravanel, Thierry Rabilloud

## Abstract

Manganese dioxide is a material that is more and more used in its particular form, for many industrial applications such as chemical catalysis or for batteries. Thus, workers can be exposed to this particulate chemical. It is known that overexposure to manganese leads to a brain disease called manganism. However, manganese is also known to impact the pulmonary function, which is important as pulmonary exposure is of prime importance for workers. We thus investigated the effects of manganese dioxide on macrophages, i.e. the scavenger cells that take particulates in charge in our bodies. To this purpose, we used a combination of proteomic and targeted approaches, in order to obtain a wide view of the cellular responses to manganese dioxide. We also used a repeated exposure mode, in order to better mimic occupational exposure. Our results point out the fact that manganese oxide nanoparticles are rather toxic for macrophages and induce mitochondrial dysfunction, oxidative stress and a pro-inflammatory response.

**Environmental significance:** Manganese dioxide is more and more used in batteries, so that workers in batteries factories are exposed to this metallic oxide. As always for particulate materials, macrophages are the first line of defense of the organism. We thus investigated the effects of manganese dioxide on macrophages, using a repeated exposure scheme to mimic occupational exposure, and the effects were documented by a combination of proteomic and targeted approaches. The functional effects include mitochondrial dysfunction, oxidative stress and inflammation.

## 1. Introduction

Manganese is a transition metal that can take various oxidation states, which confers to this metal outstanding redox properties. Industrially speaking, these properties are used in redox devices such as batteries, where manganese dioxide (MnO_2_) has been used for a long time because of its performances (e.g. in [1–4] ). It is used as a catalyst in redox reactions [5] , for example in environmental applications such as pollutants degradation [6] . It is also used in the ceramic industry both in glazing [7] and in high performances ceramics [8–11] . It has also been used for a long time in the glass industry to modify the color of glass [12, 13]. Finally, it is also used as a black pigment, and this use traces back to the paleolithic [14]. Because of its industrial use, there is a potential exposure to manganese dioxide for workers in the concerned industries. Besides, manganese is also an essential trace metal for living organisms, as it is a cofactor for several enzymes [15]. Nevertheless, as several other essential transition metals such as iron, copper or zinc, manganese becomes toxic when present in excess in the body [16]. This toxicity is attributed to the redox properties of manganese in conjunction with the biological presence of hydrogen peroxide [17], via the Fenton reaction. This leads to the production of hydroxyl radicals, which damages biological molecules [18, 19] , inducing in turn biological damages [18, 20, 21].

In manganese-exposed populations, the toxic symptoms are prevalent in the nervous system, and cause a disease called manganism, which is closely related in its symptoms with Parkinsonism [22]. For this reason, most studies devoted to manganese toxicity have been carried out on neural systems (e.g. in [23–31]). However, the brain is a highly protected organ that becomes exposed to manganese by secondary accumulation after initial exposure by other routes [32–34].

Because of this indirect mechanism, the regulatory agencies do not specify a given speciation of manganese oxides in their recommendations, and express the limits as elemental manganese present in the exposure medium, e.g. air [35].

Indeed, the toxic mechanisms at play with manganese are very generic [36–40] . As they are centered on cellular energetics, it is therefore not surprising that the effects are most visible in organs with high energy consumption such as the brain. However, other organs that are more directly exposed to manganese oxide particles, such as the lung, also show pathological responses to manganese [41]. Thus, the responses of both alveolar macrophages and lung epithelial cells to manganese have been studied [42–46] . However, in some of these studies, other manganese oxides of different oxidation states (compared to MnO_2_) such as Mn_3_O_4_ [43, 44] or Mn_2_O_3_ [45, 46] have been studied.

It is thus somewhat surprising that the toxicity of MnO_2_ on macrophages has not been specifically studied, regarding both its high use in industry and the fact that MnO_2_ is known to increase respiratory pathologies, even at low concentrations [47] . We thus decided to undertake this study, and to use a combination of proteomics and targeted experiments to increase the amount of gained biological knowledge. We also decided not to use the classical acute exposure scheme, where the effects are measured directly after exposure to a single dose [43–46] . Instead, we used a repeated exposure scheme, which was likely to better mimic the workers exposure, especially in the context of progressive particles dissolution in cells, as observed for silver nanoparticles [48]. We complemented this scheme by a recovery scheme, where the cells are exposed to a single dose and let to recover for some time before the experimental readouts. This scheme allows to investigate the persistence of the effects [49, 50] , in particular in view of the intracellular dissolution of the particles.

## 2. Material and methods

### 2.1 Pigment particles preparation and characterization

In order to work on a product as close as possible to the manganese dioxide dust encountered in working environments, we used Pigment Black 14 (Manganese dioxide, ref. 0999/100), which was purchased from Zecchi Colori (Florence, Italy). The pigment was received as a dry powder and dispersed at 10 mg/ml in an aqueous solution of arabic gum (10 mg/ml), previously sterilised overnight at 80°C in humid atmosphere. Pigment’s dispersions were then re-sterilised under the same conditions to minimise microbial contamination. To reduce agglomerates and aggregates, the dispersions were sonicated in a Vibra Cell VC 750 sonicator (VWR, Fontenay-sous-Bois, France) equipped with a Cup Horn probe. Sonication was carried out in pulse mode for 30 minutes (1 s ON/ 1 s OFF) at 60% amplitude, corresponding to 90W per pulse, in volume not exceeding 3 ml to ensure efficient energy transfer. Dispersions were stored at 4 °C and gently stirred before each experiment to avoid sediment accumulation on tube walls. Prior to use, pigment dispersions were sonicated for 15 minutes in an ultrasonic bath, then diluted in sterile water to intermediate concentrations as required. To limit potential sedimentation and microbial contamination over time, fresh pigment preparations were renewed at least once per month.

For the in-solution characterization of the pigment dispersions, Dynamic Light Scattering (DLS) and Electrophoretic Light Scattering (ELS) were performed using a Litesizer 200 instrument (Anton Paar, Les Ulis, France) equipped with an Omega reusable cuvette (225288, Anton Paar), suitable for both measurements. Before introducing the sample, the zeta potential of the cuvette filled with PBS 0.001X alone was measured to verify cleanliness and exclude background signal or instrument drift. Pigment dispersions were diluted to a final concentration of 10 µg/ml in 0.001X PBS to ensure adequate particle presence for detection while maintaining optimal optical transmittance and ionic strength for electrophoretic mobility analysis. DLS measurements were conducted at 25 °C, and two sequential measurements were recorded with a 1-minute interval to monitor potential sedimentation effects. The hydrodynamic size distribution was expressed as number-weighted intensity. ELS measurements were conducted at 25 °C, three measurements were taken, each separated by a 30-second delay. Optical parameters were automatically adjusted by the instrument to ensure accurate signal acquisition for each sample.

Transmission Electronic Microscopy (TEM) was performed as previously described [51]. Briefly, 10 µL of a 500µg/ml dispersion were added to a glow discharge grid coated with a carbon supporting film for 5 minutes. The excess solution was soaked off using a filter paper and the grid was air-dried. The images were taken under low dose conditions (<10 e-/Å2) with defocus values comprised between 1.2 and 2.5 mm on a Tecnai 12 LaB6 electron microscope at 120 kV accelerating voltage using a 4k x 4k CEMOS TVIPS F416 camera.

### 2.2. Cell culture and treatment

The mouse macrophage cell line J774A.1 was purchased from the European Cell Culture Collection (Salisbury, UK). For cell maintenance, cells were cultured in DMEM supplemented with 10% of fetal bovine serum (FBS) in non-adherent flasks (Cellstar flasks for suspension culture, Greiner Bio One, Les Ulis, France). Cells were split every 2 days at 200,000 cells/ml and harvested at 1 million cells/ml.

#### 2.2.1. Determination of Sublethal Dose (LD₂₀)

To determine the sublethal dose 20 (LD₂₀) for each pigment, cells were exposed to a range of pigment concentrations, and viability was assessed using the MTT assay with the modifications described in [52] after a 24-hour exposure period. Briefly, J774A.1 macrophages were seeded at 500,000 cells/ml in 24-well adherent plates in DMEM supplemented with 1% horse serum, 1% HEPES, and 10,000 units/ml penicillin-streptomycin as described before [49] to prevent cell proliferation and pigment dilution through time and let rest for 24 hours before pigment exposure. Absorbance readings were obtained using a BMG Labtech FLUOstar Omega® plate reader. The LD₂₀ value was defined as the concentration inducing approximately 20% reduction in viability compared to the untreated control.

#### 2.2.2. Recovery exposure settings

Cells were seeded at 500,000 cells/ml in DMEM supplemented with horse serum, as described above. Manganese dioxide was added at the selected concentration for 24 hours as for an acute exposure. Afterwards, the medium was changed in order to remove the remaining particles. The medium was then renewed every 24 hours until the end of the 72h-recovery period. At the end of the exposure, cells were treated and harvested depending on the functional test performed. For the viability test, the MTT assay was used as described above. In some experiments the viability was tested 24 or 48 hours after removal of the manganese dioxide-containing medium.

#### 2.2.3. Repeated exposure settings

Cells were seeded at 500,000 cells/ml in DMEM supplemented with horse serum, as described above. Manganese dioxide was added at the selected concentration daily for four days, with a medium change every 2 days. 24 hours after the last pigment addition, the cells were treated and harvested depending on the functional test performed. For the viability test, the MTT assay was used as described above.

### 2.3. Cellular quantification of metals by inductively coupled plasma-mass spectrometry (ICP-MS)

Cells were seeded as previously described into 6-well adherent plates. After the repeated exposure or the recovery period, cells were rinsed with PBS and harvested into 2 ml Eppendorf tubes. Samples were centrifuged for 5 minutes at 1200 rpm, the supernatant was carefully removed, and 200 µL of ICP lysis buffer (5mM Hepes NaOH pH 7.5, 0.75 mM spermine tetrahydrohloride, 0.1% (w/v) dodecyltrimethylammonio-propane-sulfonate (SB 3-12)) was added to the pellet. The cell pellet was thoroughly vortexed to ensure complete lysis. To separate the soluble and insoluble fractions, lysates were centrifuged at 15,000 g for 30 minutes. The supernatant was transferred into a fresh Eppendorf tube. Samples were stored at -20°C until analysis. The resulting supernatant (soluble fraction) was expected to contain soluble metal ions or degradation products, while the pellet (solid fraction) retained any intracellular, undissolved pigment particles. Both fractions, along with raw pigment suspensions at the working dilution, were analysed by Inductively Coupled Plasma Mass Spectrometry (ICP-MS) to quantify the elemental metal content. Protein concentration in each cell sample was determined using the Bradford assay to normalize metal content to total protein levels.

For the ICP-MS analysis, samples were dehydrated to dryness and digested at 90°C for 4 hours using 65% (w/v) nitric acid. Mineralized samples were analyzed using an iCAP RQ quadrupole mass spectrometer (Thermo Fisher Scientific). Concentrations were determined using standard curves made from serial dilutions of a multi-element solution (ICP multi-element standard IV, Merck) and corrected using an internal standard solution containing 45Sc and 103Rh, added online. Manganese was quantified using 55Mn data in both standard and helium collision modes. Data integration was done using the Qtegra software (Thermo Fisher Scientific).

### 2.4. Proteomics

Proteomics was carried out essentially as described previously [53]. However, the experimental details are given here for the sake of consistency

#### 2.4.1. Sample preparation

After exposure to the manganese dioxide particles and the recovery period (if necessary), the cells were harvested by flushing the 6 well plates. They were collected by centrifugation (300g, 5 minutes) and rinsed twice in PBS. The cell pellets were lysed in 100 µl of extraction buffer (4M urea, 2.5% cetyltrimethylammonium chloride, 100mM sodium phosphate buffer pH 3, 150µM methylene blue). The extraction was let to proceed at room temperature for 30 minutes, after which the lysate was centrifuged (15,000g, 15 minutes) to pellet the nucleic acids. The supernatants were then stored at -20°C until use.

#### 2.4.2. Shotgun proteomics

For the shotgun proteomic analysis, the samples were included in polyacrylamide plugs according to Muller et al. [54] with some modifications to downscale the process [55]. To this purpose, the photopolymerization system using methylene blue, toluene sulfinate and diphenyliodonium chloride was used [56].

As mentioned above, the methylene blue was included in the cell lysis buffer. The other initiator solutions consisted in a 1 M solution of sodium toluene sulfinate in water and in a saturated water solution of diphenyliodonium chloride. The ready-to-use polyacrylamide solution consisted of 1.2 ml of a commercial 40% acrylamide/bis solution (37.5/1) to which 100 µl of diphenyliodonium chloride solution, 100 µl of sodium toluene sulfinate solution and 100 µl of water were added.

To the protein samples (16 µl), 4 µl of acrylamide solution were added and mixed by pipetting in a 500µl conical polypropylene microtube. 100 µl of water-saturated butanol were then layered on top of the samples, and polymerization was carried out under a 1500 lumen 2700K LED lamp for 2 hours, during which the initially blue gel solution discolored. At the end of the polymerization period, the butanol was removed, and the gel plugs were fixed for 1 hr with 200 µl of 30% ethanol 2 % phosphoric acid, followed by 3x 15 minutes washes in 20% ethanol. The fixed gel plugs were then stored at -20°C until use.

Gel plug processing, digestion, peptide extraction and nanoLC-MS/MS was performed as previously described [53], using a nanoACQUITY Ultra-Performance-LC (Waters Corporation, Milford, USA) coupled to a Q-Exactive HF-X mass spectrometer (Thermo Fisher Scientific, Bremen, Germany).

For protein identification, the MS/MS data were interpreted using a local Mascot server with MASCOT 2.6.2 algorithm (Matrix Science, London, UK) against an in-house generated database containing all Mus musculus and Rattus norvegicus entries from UniProtKB/SwissProt (version 2025_01, 25,574 sequences) and their corresponding reverse entries. Spectra were searched with a mass tolerance of 10 ppm for MS and 0.05 Da for MS/MS data. Trypsin was selected as the enzyme and a maximum of one missed cleavage was allowed. Acetylation of protein N-termini, carbamidomethylation of cysteine residues and oxidation of methionine residues were specified as variable modifications. Identification results were imported into the Proline software [57] version 2.3 (http://proline.profiproteomics.fr/) for validation. Peptide Spectrum Matches (PSM) with pretty rank equal to one were retained. False Discovery Rate was then optimized to be below 1% at PSM level using Mascot Adjusted E-value and below 1% at Protein Level using Mascot Mudpit score.

Mass spectrometry data are available via ProteomeXchange with the identifier PXD065155.

#### 2.4.3. Label Free Quantification

Peptides abundances were extracted thanks to Proline software version 2.3 (http://proline.profiproteomics.fr/) using a m/z tolerance of 10 ppm. Alignment of the LC-MS runs was performed using Loess smoothing. Cross Assignment was performed within groups only. Protein abundances were computed by sum of peptides abundances (normalized using the median).

#### 2.4.4. Data analysis

For the global analysis of the protein abundances data, missing values were imputed with a low, non-null value. The complete abundance dataset was then analyzed by the PAST software [58].

Proteins were considered as significantly different if their p value in the Mann-Whitney U-test against control values was inferior to 0.05. The selected proteins were then submitted to pathway analysis using the DAVID tool [59], with a cutoff value set at a FDR of 0.2.

### 2.5 Assessment of functional parameters

#### 2.5.1. Mitochondrial transmembrane potential

The mitochondrial transmembrane potential assay was performed essentially as described previously [60]. After exposure to manganese dioxide under the recovery or repeated schemes, rhodamine 123 (Rh123) was added to the cultures at an 80 nM final concentration (to avoid quenching [61]), and the cultures were further incubated at 37°C for 30 minutes. At the end of this period, the cells were collected, washed in cold PBS containing 0.1% glucose, resuspended in PBS glucose and analyzed for the green fluorescence (excitation 488 nm emission 525nm) on a Melody flow cytometer. As a positive control, butanedione monoxime (BDM) was added at a 30mM final concentration together with the Rh123 [62]. As a negative control, carbonyl cyanide 4- (trifluoromethoxy)phenylhydrazone (FCCP) was added at a 5µM final concentration together with the Rh123 [62].

#### 2.5.2. Assay for oxidative stress

For the oxidative stress assay, a protocol based on the oxidation of dihydrorhodamine 123 (DHR123) was used, essentially as described previously [60]. After exposure to manganese dioxide under the recovery or repeated schemes, the cells were treated in PBS containing 500 ng/ml DHR123 for 20 minutes at 37°C. The cells were then harvested, washed in cold PBS containing 0.1% glucose, resuspended in PBS glucose and analyzed for the green fluorescence (same parameters as rhodamine 123) on a Melody flow cytometer. Menadione (applied on the cells for 2 hours prior to treatment with DHR123) was used as a positive control in a concentration range of 25-50µM.

#### 2.5.3. Reduced glutathione assay

Intracellular reduced glutathione levels were assessed using the monochlorobimane technique, with some modifications [63]. Briefly, after exposure to manganese dioxide under the recovery or repeated schemes, the cells were harvested, centrifuged during 5 min, and labeled with 70 µM monochlorobimane (diluated in warm PBS) for 5 min at 37 °C. The reaction was stopped via an incubation in ice for 5 min in the dark. The cells were washed twice with cold PBS and finally, they were analyzed via a Cytoflex flow cytometer (Beckman, Villepinte, France) using a laser excitation at 405 nm and an emission at 448 ± 45 nm, with exclusion of the dead cells from the analysis.

#### 2.5.4. Lysosomal assay

For the lysosomal function assay, the Lysosensor method was used, as described previously [60]. After exposure to manganese dioxide under the recovery or repeated schemes, the medium was removed, the cell layer was rinsed with complete culture medium and incubated with 1µM Lysosensor Green (Molecular Probes) diluted in warm (37°C) complete culture medium for 1 hour at 37°C. At the end of this period, the cells were collected, washed in cold PBS containing 0.1% glucose, resuspended in PBS glucose and analyzed for the green fluorescence (excitation 488 nm emission 540nm) on a FacsCalibur flow cytometer (BD Biosciences, Le Pont-de-Claix, France).

#### 2.5.5. Phagocytosis assay

For this assay, the cells were first exposed to manganese dioxide under the recovery or repeated schemes. After this exposure to pigment particles, the cells were then exposed to 0.5 µm latex beads (carboxylated surface, yellow green-labelled, from Polysciences excitation 488 nm emission 527/32 nm) for 3 hours at a concentration of 5µg/ml. After this second exposure, the cells were collected, rinsed twice with PBS, and analyzed for the two fluorescences (green and red) on a FacsCalibur flow cytometer (BD Biosciences, Le Pont-de-Claix, France).

#### 2.5.6. Measurement of Cytokines Secretion

After exposure to manganese dioxide under the recovery or repeated schemes, the culture medium was collected and analyzed for proinflammatory cytokines. Tumor Necrosis Factor, Monocyte Chemoattractant Protein and interleukin 6 levels were measured using the Cytometric Bead Array Mouse Inflammation Kit (catalog numbers 558299, 558301 and 558266, BD Biosciences, Le Pont de Claix), and analyzed with FCAP Array software (3.0, BD Biosciences) according to the manufacturer’s instructions. In some experiments lipopolysaccharide (LPS) was added at 50 ng/ml for the last 18 hours of culture prior to cell culture supernatant collection and analysis.

## 3. Results

### 3.1. Physicochemical characterisation of manganese dioxide particles

Manganese dioxide particles were first characterized using transmission electron microscopy. Representative electron microscopy images in **Figure 1** illustrate the morphological features of the MnO_2_ particles used in this work. The particles appear as nanoscale (100-200 nm) aggregates of primary rod-shaped particles that are 20-50 nm long and only a few nanometers wide. This structure is similar, although more compact, to the structure of synthetic amorphous silicas [50].

**Figure 1:**
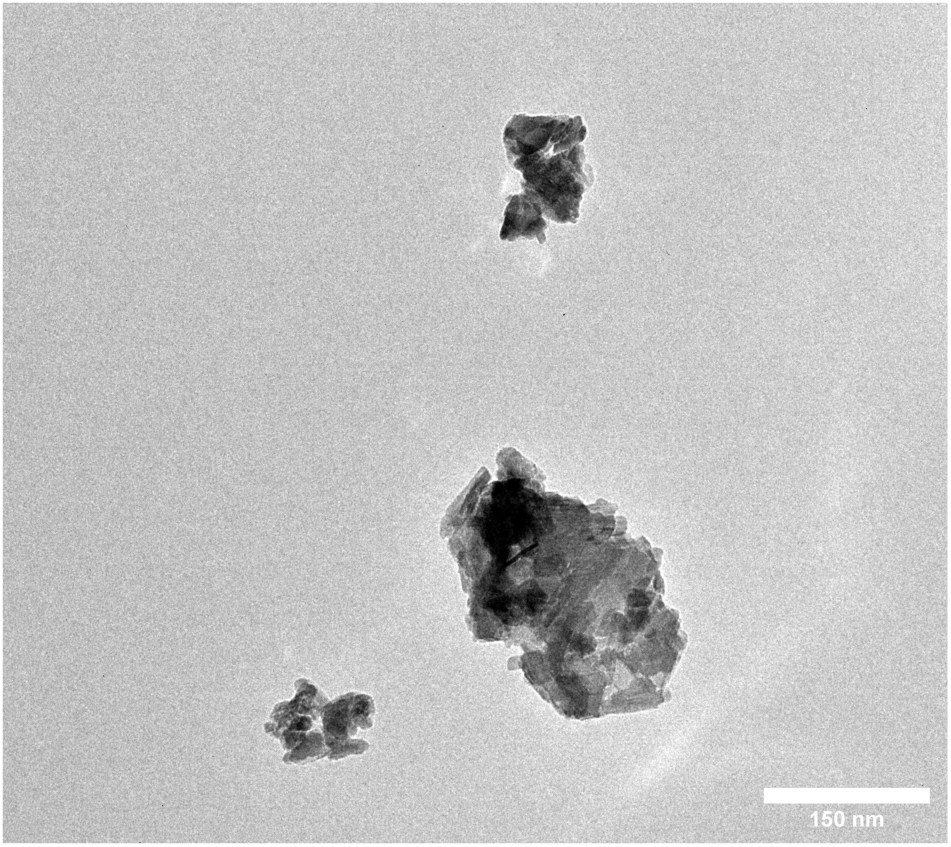
representative electron microscopy images of MnO2 (PBk14) particles at 23000x magnification. Particles were prepared as described in section 2.1, diluted at 500µg/ml in sterile water. A 10 µL aliquot was applied to a glow discharge grid coated with a carbon supporting film for 5 minutes. The excess solution was soaked off by a filter paper and the grid was air-dried. The images were taken under low dose conditions (<10 e-/Å2) with defocus values between 1.2 and 2.5 µm on a Tecnai 12 LaB6 electron microscope at 120 kV accelerating voltage using 4k x 4k CEMOS TVIPS F416 camera.

The in-solution behavior of the MnO_2_ nanoparticles was then studied using Dynamic Light Scattering (DLS) to assess their size distribution, polydispersity index (PI), and surface charge. In solution, the MnO_2_ particles appeared with a hydrodynamic diameter of 422±11 nm, with a polydispersity index of 26±2, and a zeta potential of -32±0.2 mV. This suggested that despite the sonication process, multiparticle agglomerates occurred in solution, at least in the saline solutions used for DLS/ELS.

### 3.2. Cell exposure and dose selection

For particles exposure, cells were treated as described in **Figure 2**. Macrophages were seeded at 500,000 cells/ml in DMEM supplemented with 1% horse serum and 1% streptomycin-penicillin in 24-well adherent plates. The following day, MnO_2_ was added for a 24-hour acute exposure.

**Figure 2.**
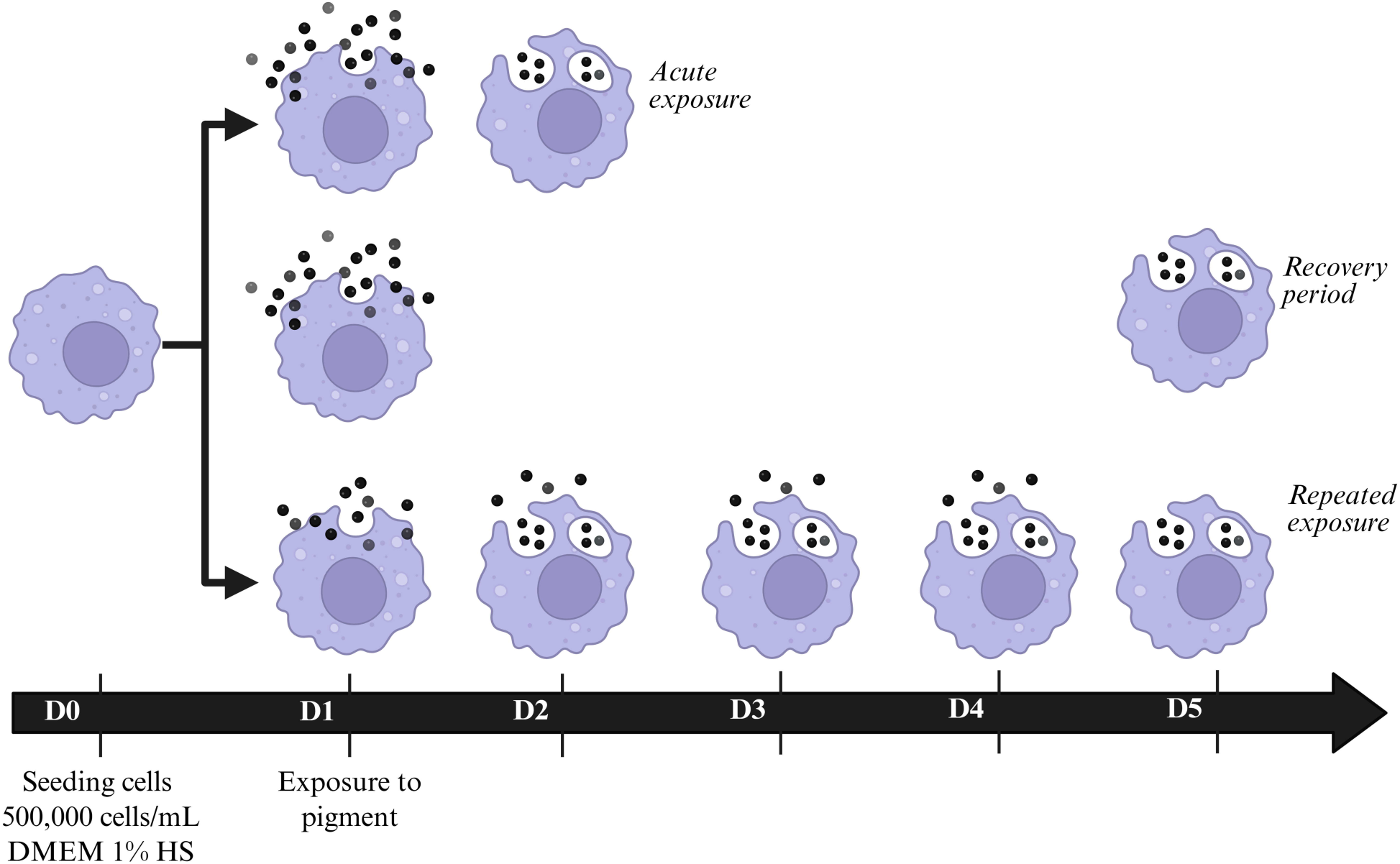
Exposure scheme. The exposure schemes used in this work are schematized in this figure. The acute exposure scheme was used only for a first determination of the adequate working concentrations

In a first series of experiments, we determined which concentrations of particles could be used on macrophages, by performing viability assays after the acute exposure. In **Figure 3**, the MTT assay showed a LD20 around 100µg/ml immediately after a 24 hours exposure (black curve). However, when cells were loaded with MnO_2_ particles for 24 hours and then left to recover in a particle-free medium, a delayed toxicity developed, as shown in **Figure 3** (blue curve). After a 48h recovery post-exposure, the viability of cells exposed to 40µg/ml MnO_2_ dropped from 89% immediately post -exposure to 57%. This drop in viability continued when the recovery period was extended to 72h, as shown in **Figure 3** (green curve), as the viability of cells exposed to 40µg/ml manganese dioxide dropped to 43%, and the viability of cells exposed to 20µg/ml manganese dioxide decreased to 84%.

**Figure 3:**
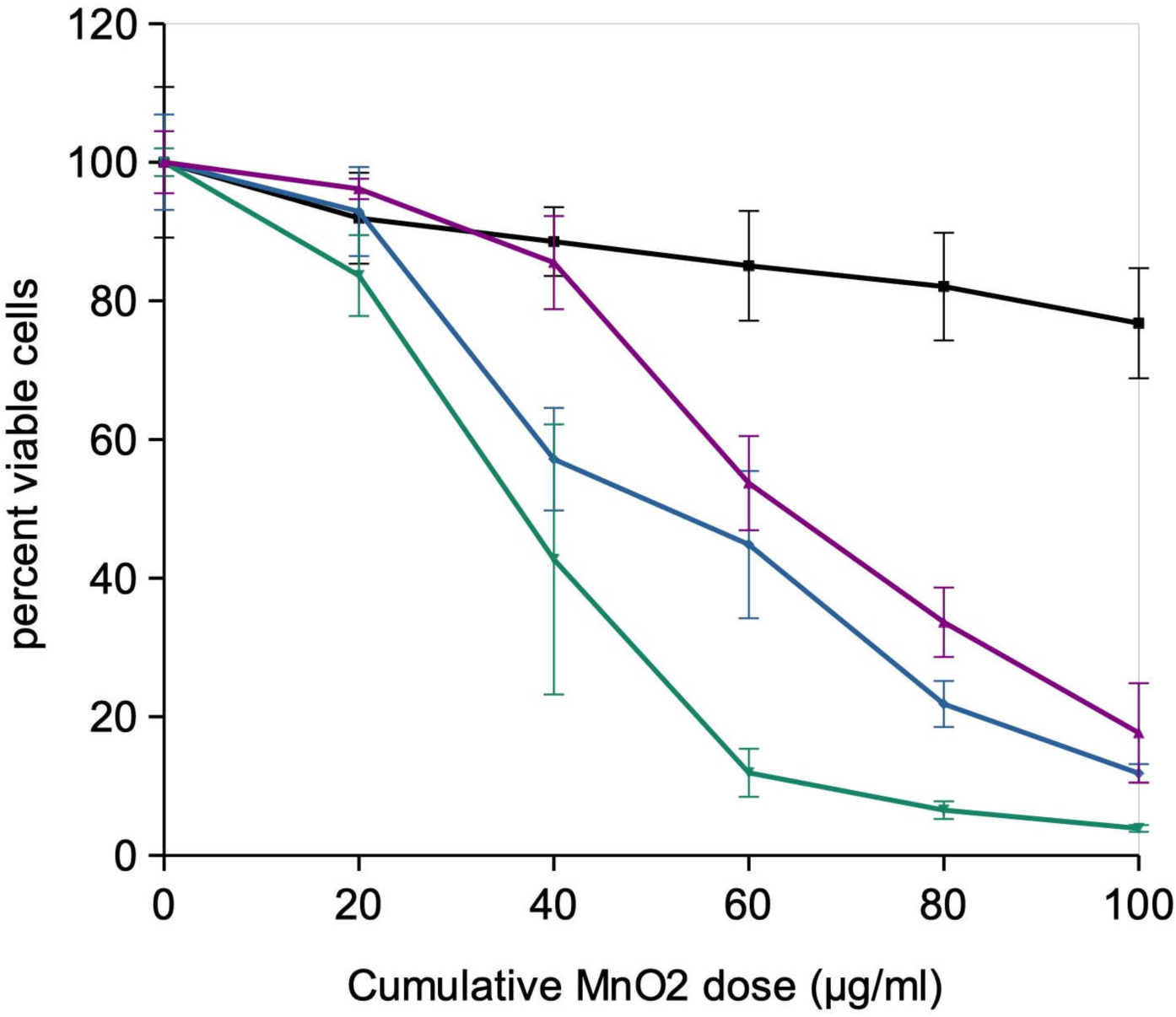
Cell viability after treatment with manganese dioxide nanoparticles. The cells were exposed to MnO2 particles under different schemes acute: exposure for 24h at the indicated concentrations, viability readout immediately after exposure delayed 48 h: exposure for 24h at the indicated concentrations, then removal of the particles-loaded culture medium and replacement by fresh medium. Viability readout 48h after this medium change delayed 72h: exposure for 24h at the indicated concentrations, then removal of the particles-loaded culture medium and replacement by fresh medium. Viability readout 3 days after the end of the exposure period repeated: cells are exposed every day for 4 days at a concentration that is one fourth of the total dose. Medium change every 48h. Viability readout 24 hours after the last particle addition.

Finally, in a repeated exposure mode, as shown in **Figure 3** (purple curve), the viability was above 90% for a 4x5 µg/ml exposure, 85% for a 4x10 µg/ml exposure, but dropped to18% for a 4x25 µg/ml exposure.

### 3.3. Particles uptake and dissolution by cells

To quantitatively assess pigment uptake after acute exposure and retention following the recovery period, mineralization experiments were conducted on the lysates of the MnO_2_ -loaded cells. The supernatant (soluble fraction) gave the proportion of soluble manganese, while the pellet (internalized particular fraction) gave the amount of pigment remaining as particular material after the recovery period.

The results, shown in **Table 1**, showed that manganese dioxide was efficiently internalized by the macrophages in these exposure schemes, with 30-45% of the theoretical input still present after 4 to 5 days.

**Table 1:** quantification of internalized and dissolved manganese in MnO_2_ -exposed macrophages. The results are expressed in µg manganese/culture well The results are expressed as mean±standard deviation ND: not detected

|  | control cells | 4x5µg/ml MnO<sub>2</sub><br>(4 days) | 4x10µg/ml MnO<sub>2</sub><br>(4 days) | 20 µg/ml MnO<sub>2</sub><br>24hrs<br>+ 72h recovery |
| --- | --- | --- | --- | --- |
| Manganese input<br>(measured) | ND | 11.38±2.7 | 22.13±7.6 | 11.38±2.7 |
| Total cell-associated<br>manganese | ND | 3.47±0.42 | 8.47±0.73 | 5.07±0.75 |
| Non-sedimentable,<br>cell-associated<br>manganese | ND | 0.024±0.001 | 0.041±0.002 | 0.019±0.001 |
| Mineralization<br>yield (%) | ND | 90±16 | 87.5±0.3 | 90±16 |
| Percent<br>manganese<br>internalized | ND | 30.5±3.7 | 38.3±3.3 | 44.6±6.6 |
| Percent<br>manganese<br>dissolved | ND | 0.7±0.1 | 0.5±0.1 | 0.4±0.1 |
| Non-sedimentable,<br>cell-associated<br>iron | 0.019±0.018 | 0.036±0.031 | 0.073±0.066 | 0.025±0.011 |

ND: not detected

### 3.4. Global analysis of the proteomic experiments

The shotgun proteomic analysis was able to detect and quantify 2286 proteins (**Table S1**). As a first analysis, we analyzed the complete dataset via the PAST software, through a principal coordinates analysis. The results, displayed in **Figure 4**, showed a good separation between groups, with the recovery group being closest to the control one, and the repeatedly-treated groups becoming more remote from the control group with increasing dose. These results correlated with the amount of cellular non-sedimentable manganese (Table 1).

**Figure 4:**
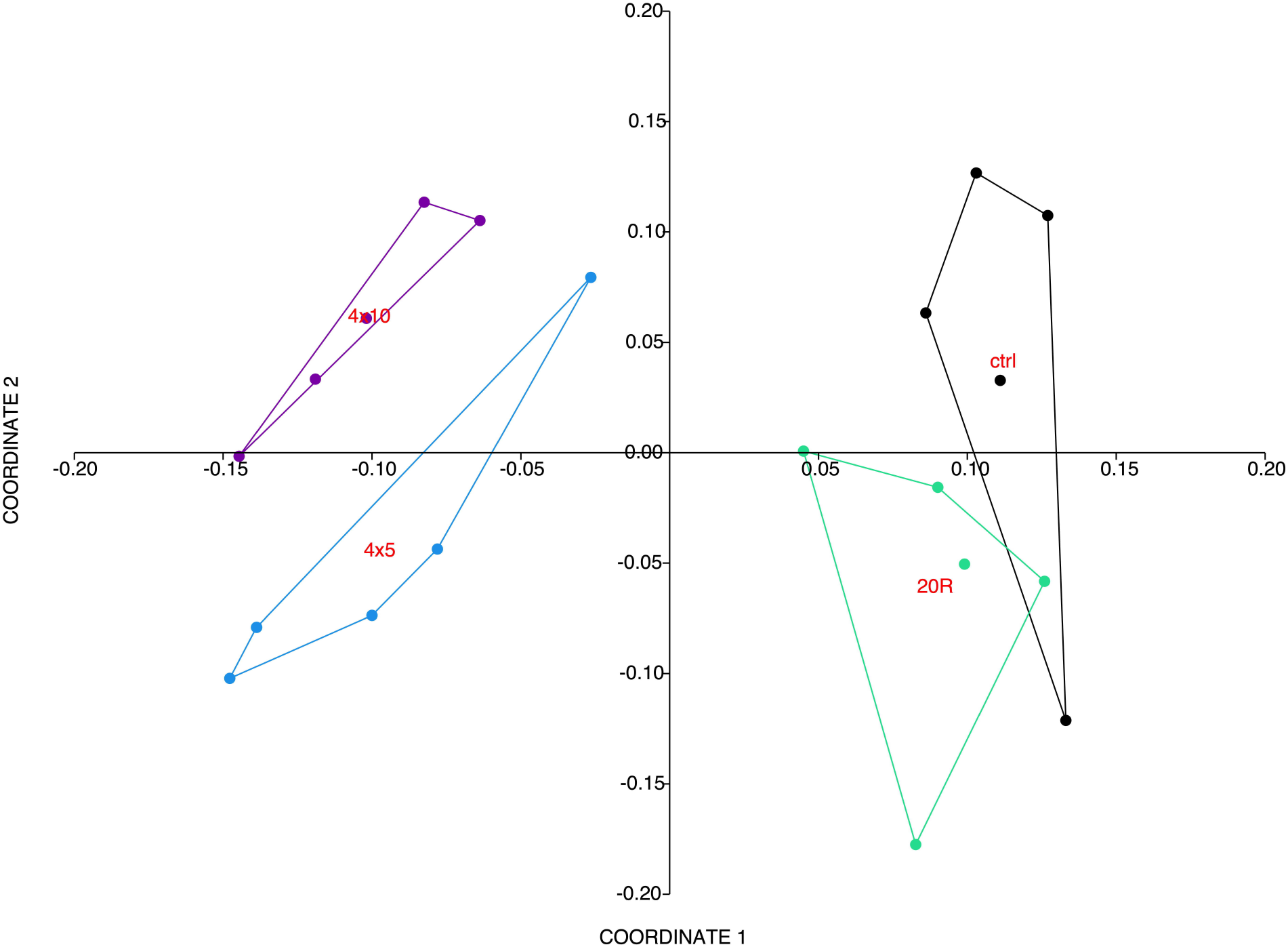
Global analysis of the proteomic data. The complete proteomic data table (2286 proteins) was analyzed by Principal Coordinates Analysis (Gower distance), using the PAST software. The results are represented as the X-Y diagram of the first two axes of the Principal Component Analysis, representing 40% of the total variance. Eigenvalue scale. This representation allows to figure out how, at the global proteome scale, the samples are related to each other. Samples grouped in such a diagram indicate similar proteomes, and the larger the distance between samples are, the more dissimilar their respective proteomes.

The extent of the proteomes modulations was confirmed by the ANalysis Of SIMilarity test [64]. This test showed a p-value (probability by random) of 0.015 for the 4x5 µg/ml vs. control comparison, 0.0154 for the 4x10 µg/ml vs. control comparison, and 0.0564 for the recovery vs. control comparison. The recovery group was significantly different from the repeatedly-treated ones, with p=0.009 in the recovery vs. 4x5 µg/ml comparison and p= 0.0088 in the recovery vs. 4x10 µg/ml comparison.

In order to select the proteins whose abundances are modulated by the treatments, we conducted a Mann-Whitney test (U ≤2, equivalent to p<0.05) for each of the treated vs control comparisons. The proteins which passed this threshold were considered significantly altered in their abundances and were selected (**Tables S2 to S4**).

To gain further insight into the significance of the observed changes, this list of modulated proteins was used to perform pathways analyses using the DAVID software, and the results are shown in **Tables S5 to S7**. The pathway analysis pointed out several cellular processes which were modulated upon treatment with manganese dioxide particles. Some were generic, e.g. mitochondria, lysosomes, while some were more specific to macrophages functions, such as immune responses.

### 3.5. Mitochondria and central carbon metabolism

Mitochondrial proteins represented an abundant class among the proteins modulated in response to manganese dioxide particles, with 107 proteins (**Table S8**). A detailed examination of the modulated proteins showed a decrease in 7 Complex I (NADH dehydrogenase) subunits out of the 12 that were detected in the proteomic screen, and also a decrease in the Complex I assembly factor (Q8JZN5) . One subunit (out of 2 detected) of Complex II (succinate dehydrogenase) was increased in response to repeated exposure to MnO_2_ (4x5 µg/ml) while only one subunit (out of 5 detected) of Complex III (ubiquinol cytochrome C oxidoreductase) was also modulated (decreased in response to manganese). Five subunits (out of 9 detected) of complex IV (cytochrome oxidase) were decreased in response to manganese, while only 2 subunits (out of 11 detected) of Complex V (ATP synthase) were decreased in response to manganese. Among the other mitochondrial complexes which respond to exposure to manganese, the MICOS complex can be noticed. Among the 5 subunits that were detected, 2 (Mic10 and Mic27) decreased in response to manganese, while 2 (Mic19 and MIC60) increased significantly in response to manganese and one (Mic25) increased but not statistically significantly.

These important changes prompted us to investigate the mitochondrial transmembrane potential, which is the result of the delicate balance between the activities of OXPHOS complexes I, III and IV, which pump protons from the mitochondrial matrix to the intermembrane space, and the activity of Complex V, which uses proton re-entry in the mitochondrial matrix to produce ATP by ADP phosphorylation. The results, displayed on **Figure 5**, showed a significant increase of the mitochondrial transmembrane potential in response to all conditions of exposure to manganese.

**Figure 5:**
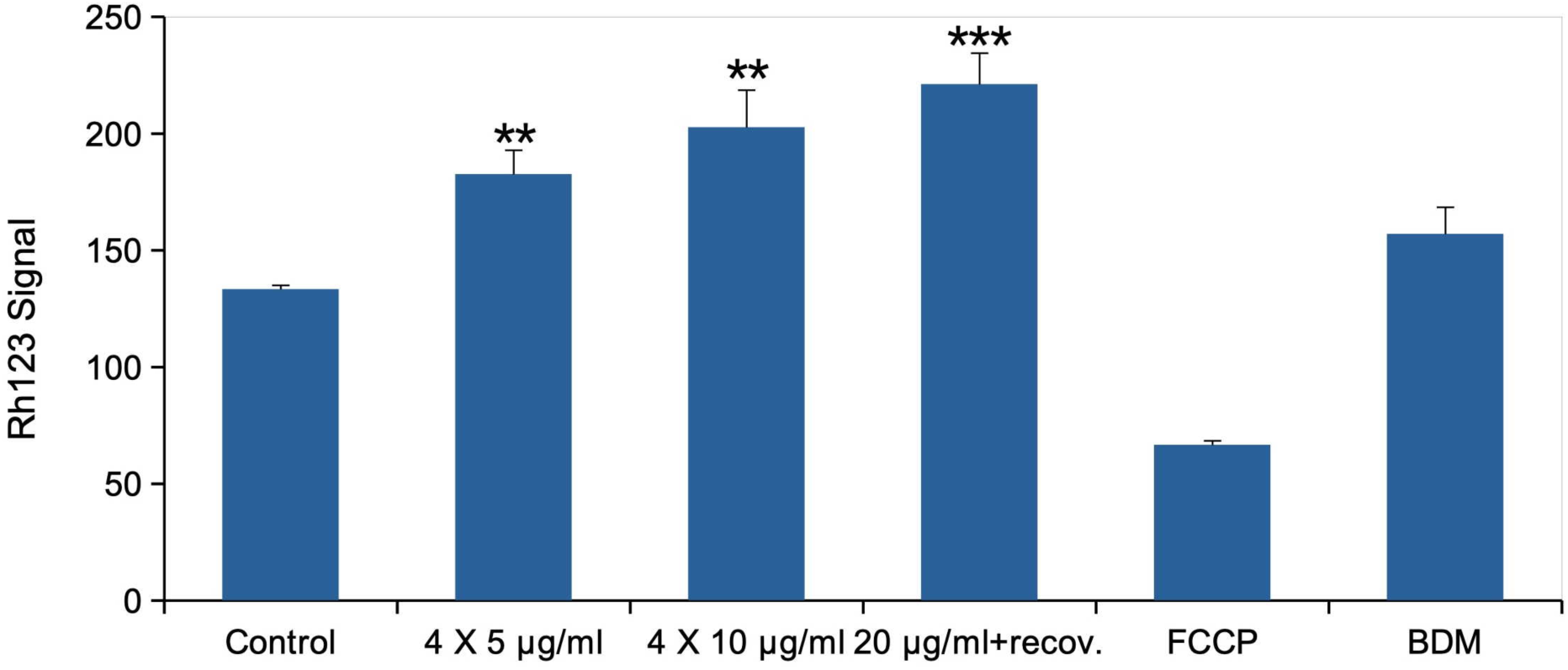
Mitochondria. The cells were exposed to MnO_2_ particles according to the three schemes used in this work. Mitochondrial transmembrane potential was then measured via the rhodamine 123 method. All cells were positive for rhodamine 123 internalization in mitochondria, and the mean fluorescence is the displayed parameter. Results are displayed as mean± standard deviation (N=4) Control: Unexposed cells 4x5 µg/ml: cells exposed to MnO2 at 5µg/ml for 4 consecutive days. Readout 24 h after last exposure 4x10 µg/ml: cells exposed to MnO2 at 10µg/ml for 4 consecutive days. Readout 24 h after last exposure 20µg/ml + recovery: cells exposed to MnO2 at 20µg/ml for 24 h, the left to recover in fresh medium without particles for 72h FCCP: Cells exposed for 30 minutes to FCCP (induces a decrease in the transmembrane potential) BDM: Cells exposed for 30 minutes to Butanedione monoxime (induces an increase in the transmembrane potential) **: statistically different from the unexposed control (p<0.01, Student T test) ***: statistically different from the unexposed control (p<0.001, Student T test)

Going further into the central metabolism, this cluster of proteins also appeared in the pathway analysis. The proteins of this pathway modulated in response to MnO_2_ are listed in **Table S9**. It appears that amon the total 26 proteins implicated in glycolysis and pentose phosphate pathway detected in the proteomic screen, 17 were modulated in response to manganese oxide. Amon them all were increased except cytosolic aconitase (P28271), which was decreased. In the glycolysis pathway, only phosphofructokinase and glucose 6-phosphate isomerase were not modulated in response to manganese oxide. All the other proteins of the pathway were induced. It is noticeable that pyruvate kinase R isoform (P53657) is not detected in the 4x5 µg/ml and recovery conditions, poorly detected in the control condition, but strongly detected and thus induced in the 4x10 µg/ml condition.

Regarding the pentose phosphate pathway, the situation was more balanced, with two unresponsive proteins representing the two first steps of the pathway (glucose 6 phosphate dehydrogenase and 6-phosphoglucolactonase), while transaldolase (Q93092) and transketolase (P40142) were increased.

### 3.6. Oxidative stress

As an increase in mitochondrial transmembrane potential has been correlated to oxidative stress [65, 66] , we probed the level of oxidative stress in pigment-exposed cells by the DHR123 method. This decision was further substantiated by the strong observed induction of the ROS producing protein ROMO1 (P60603) [67] , and by the fact that several proteins playing a major role in the handling of oxidative stress were increased in response to manganese (**Table S10**). The results, displayed on **Figure 6A**, showed a significant increase in the level of cellular oxidative stress. To investigate the cellular response to oxidative stress further, we measured the level of intracellular reduced glutathione by the monochlorobimane conjugation method. The results, displayed on **Figure 6B**, showed a significant increase in the level of intracellular reduced glutathione in the recovery condition and in cells repeated exposed to 4x10µg/ml MnO_2_ .

**Figure 6:**
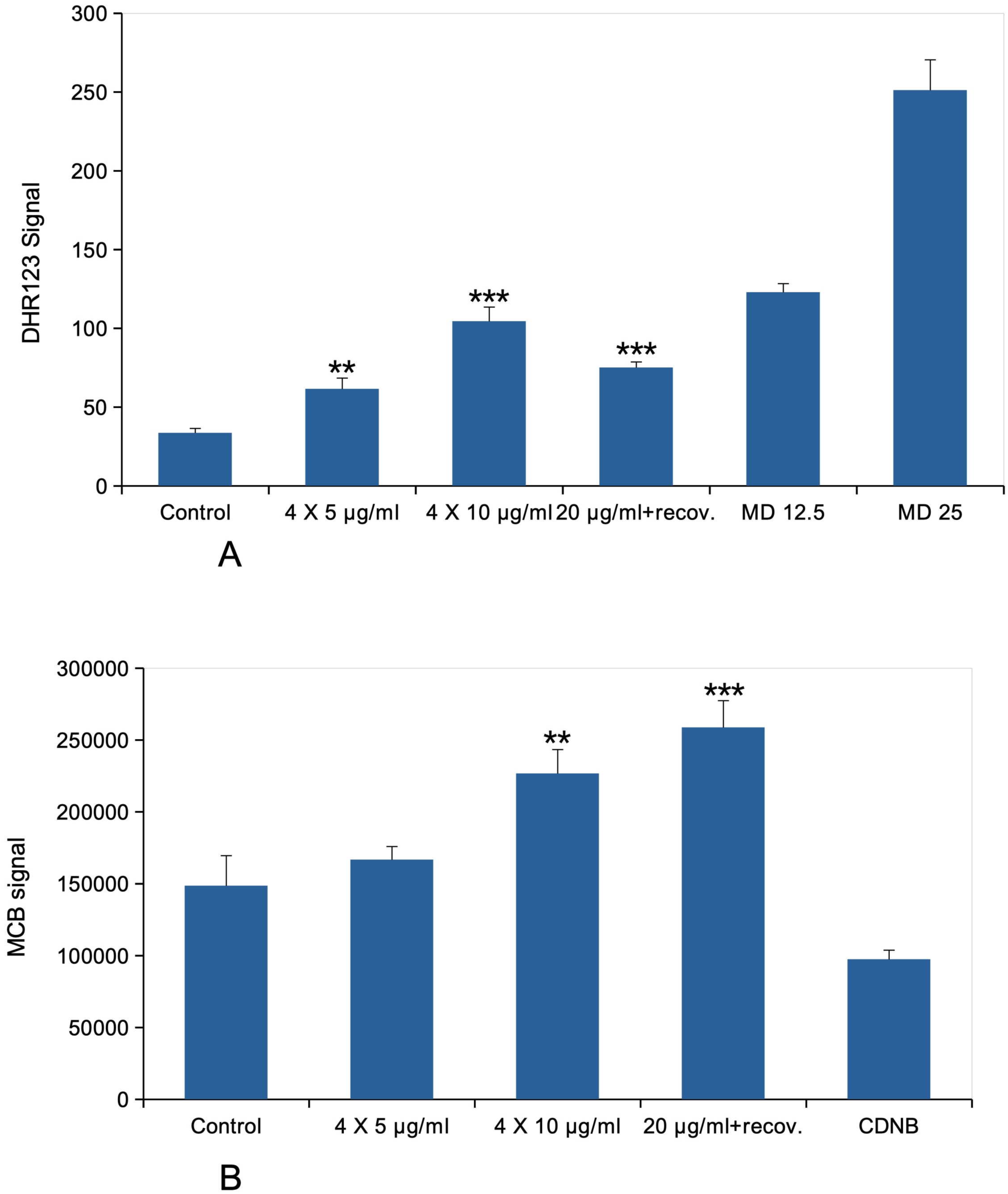
Oxidative stress, glutathione content, lysosome. Control: Unexposed cells 4x5 µg/ml: cells exposed to MnO_2_ at 5µg/ml for 4 consecutive days. Readout 24 h after last exposure 4x10 µg/ml: cells exposed to MnO_2_ at 10µg/ml for 4 consecutive days. Readout 24 h after last exposure 20µg/ml + recovery: cells exposed to MnO_2_ at 20µg/ml for 24 h, the left to recover in fresh medium without particles for 72h MD: menadione (added 2 hours before measurement) CDNB: chlorodinitrobenzene (added 30 minutes before measurement) Panel A: oxidative stress level The cells were exposed to MnO_2_ particles according to the three schemes used in this work.. The level of oxidative stress was measured via the DHR123 method. Results are displayed as mean± standard deviation (N=4). **: statistically different from the unexposed control (p<0.01, Student T test) ***: statistically different from the unexposed control (p<0.001, Student T test) Panel B: Reduced glutathione level The cells were exposed to MnO_2_ particles according to the three schemes used in this work. The level of reduced glutathione was measured via the monochlorobimane conjugation method. Results are displayed as mean± standard deviation (N=4). **: statistically different from the unexposed control (p<0.01, Student T test) ***: statistically different from the unexposed control (p<0.001, Student T test)

### 3.7. Lysosomes

Lysosomal proteins represented an important cluster among the modulated proteins, with 34 proteins (**Table S11**). Interestingly, all of these modulated but 7 were decreased in response to manganese. Among the few proteins that increased in response to manganese, we could find an aldose reductase (P45377), metallothionein 1 (P02802), CREG1 (O88668), VAMP-8 (O70404), the subunits G1 and H of the vacuolar proton ATPase (Q9CR51 and Q8BVE3), as well as the proton ATPase assembly factor (Q78T54). Oppositely, two other subunits of the proton ATPase were decreased in response to manganese (P50516, P63081) while 6 subunits were not significantly modified. This prompted us to investigate whether the lysosomal integrity or proton pumping activity was altered in response to manganese, which we measured by the Lysosensor pumping method. The results, displayed on **Figure 7**, showed a significant increase in the Lysosensor signal, meaning no significant leakage of the lysosomal membrane and thus no gross damages to the lysosomal integrity, and also an increase of the proton pumping activity, either by an increase in the number of active lysosomes or an increase in the proton pump activity at equal lysosomes number, or a combination of both.

**Figure 7:**
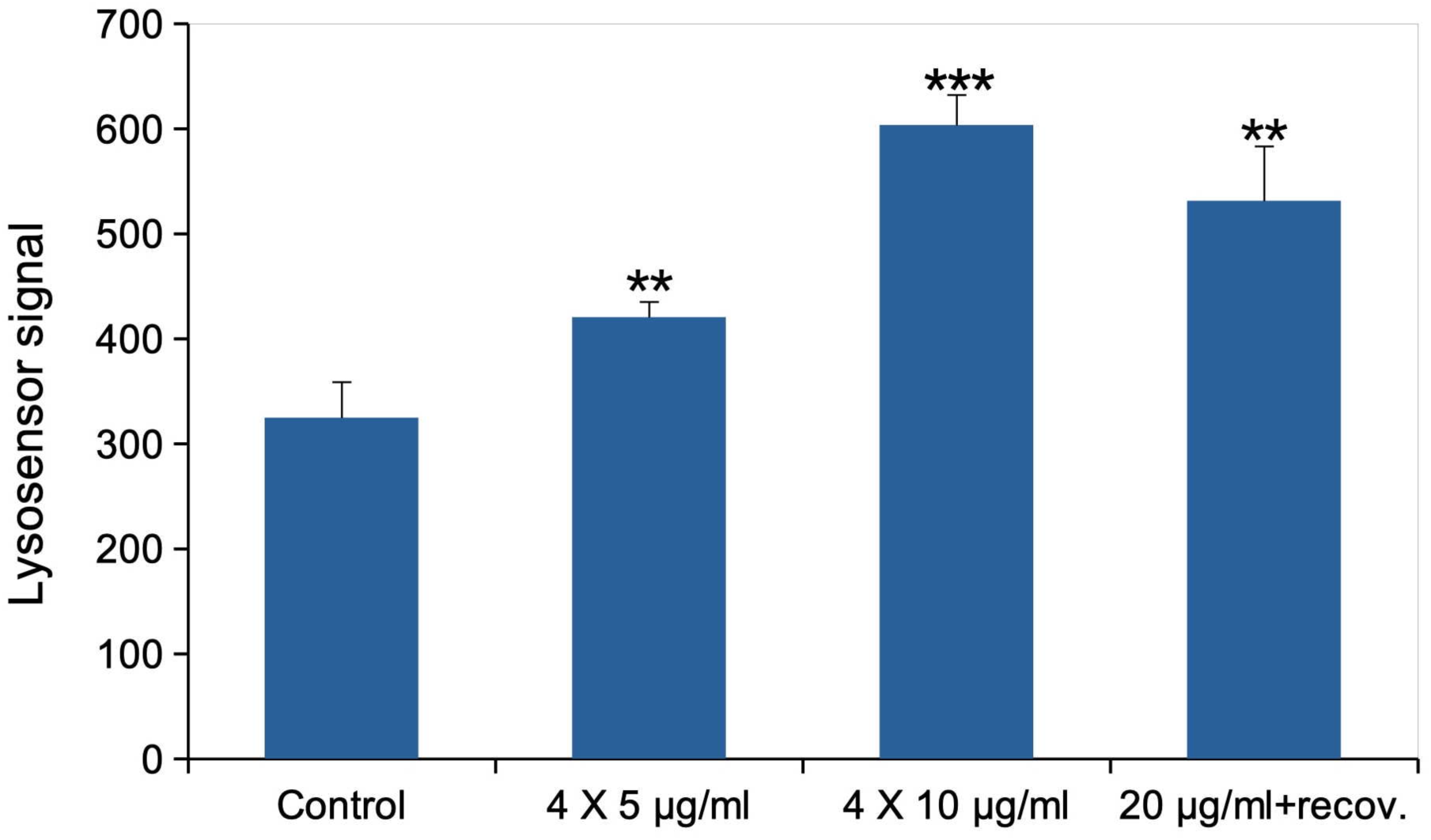
Lysosomal proton pumping (Lysosensor method). The cells were exposed to MnO_2_ particles according to the three schemes used in this work. All cells were positive for lysosensor internalization in lysosomes, and the mean fluorescence is the displayed parameter. Results are displayed as mean± standard deviation (N=4). **: statistically different from the unexposed control (p<0.01, Student T test) ***: statistically different from the unexposed control (p<0.001, Student T test)

### 3.7. Immune functions

As the lysosomal function was increased, we wondered whether phagocytosis would also be modulated in response to MnO_2_ . This working hypothesis was further reinforced by the fact that 47 proteins annotated as related to immunity were found modulated in response to MnO_2_ (**Table S12**). Among those manganese-induced changes, we could detect a decrease in the IgG receptor (P26151), in the inhibitory receptor PILRA (Q2YFS3), and in a subunit of complement receptor 3 (P05555) but an increase in the IgE receptor (P20491). Regarding the proteins known to be implicated in the control of inflammation we observed a strong induction of galectin-3 (P16110) [68] , a decrease in the pro-inflammatory protein TM106A (Q8VC04) [69] in the repeated exposure mode (but an increase in the recovery mode). We observed an increase in the down-regulator of the cGAS/STING pathway TMEM33 (Q9CR67) [70] , but an increase of the up-regulator of the cGAS/STING pathway G3BP1 [71] (P97855) in the repeated exposure mode, while a decrease was observed in the recovery mode. We also observed a discrepant modulation of the M1-polarizing protein Irgm1 (Q60766) [72], which was decreased in the repeated exposure mode but increased in the recovery mode. The wide-scope immune response deactivator Lyn (P25911) [73] was also decreased in response to manganese oxide.

On the endocytosis/phagocytosis side, a strong induction of sorting nexins 9 and 18 (respectively 91VH2 and Q91ZR2) as well as VAMP8 (O70404), but a decrease in the membrane fusion-associated proteins EHD4 (Q9EQP2) and NSF (P46460) was observed.

As this complex array of changes did not allow to make simple predictions, we tested the phagocytic function (**Figure 8**) and the inflammatory response (**Figure 9**).

**Figure 8:**
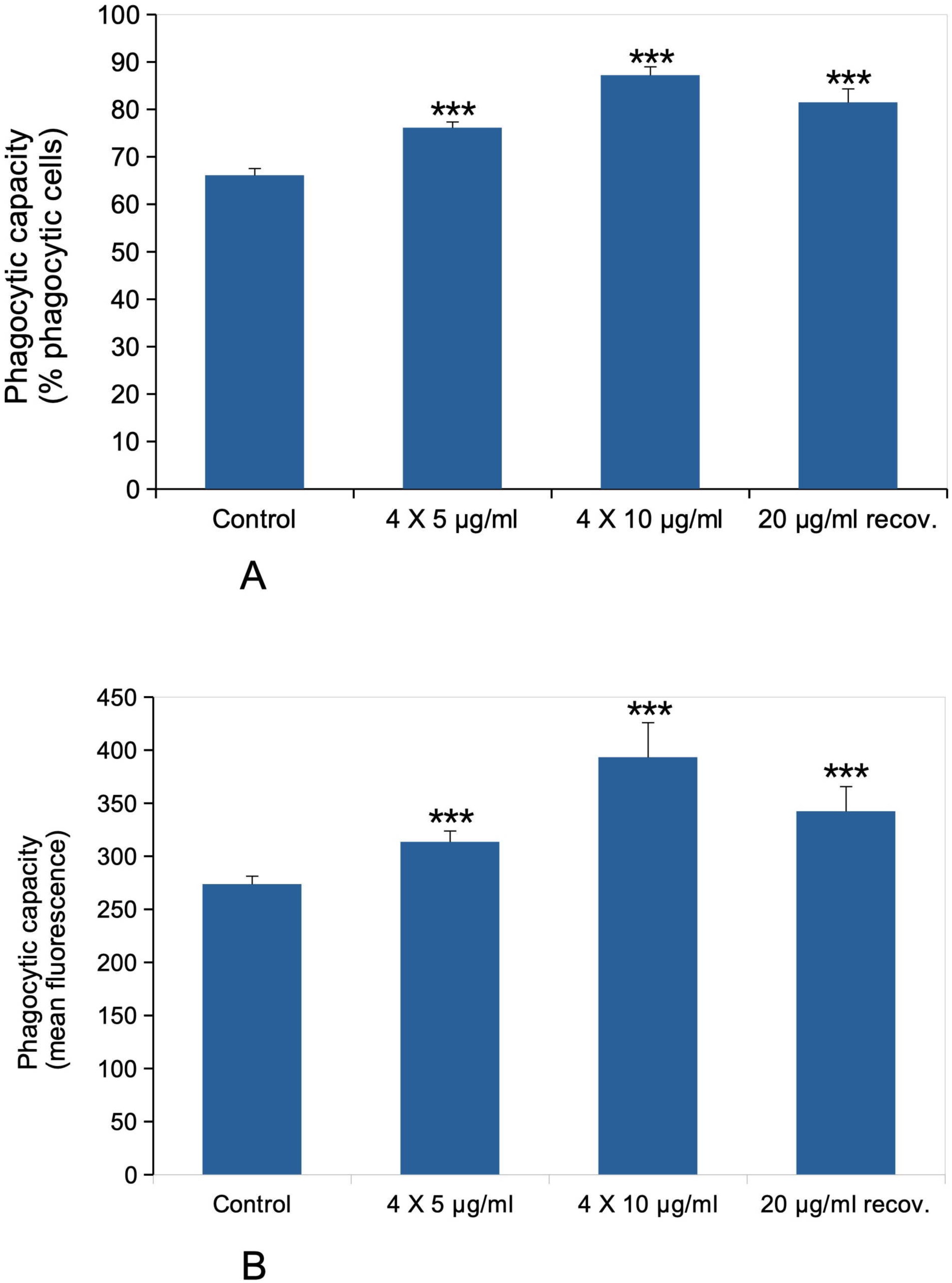
Phagocytic activity. The cells were exposed to MnO_2_ particles according to the three schemes used in this work. At the end of the exposure period, the cells were treated with green fluorophore labelled carboxylated latex beads for 3 hours. Panel A: The percentage of green fluorescence-positive cells, indicating the percentage of cells able to internalize the test beads in 3 hours, is the displayed parameter. Results are displayed as mean± standard deviation (N=4). Significance marks: ***: statistically different from the unexposed control (p<0.001, Student T test) Panel B: The mean fluorescence, indicating the amount of green beads internalized, is the displayed parameter. Results are displayed as mean± standard deviation (N=4). Significance marks: ***: statistically different from the unexposed control (p<0.001, Student T test)

**Figure 9:**
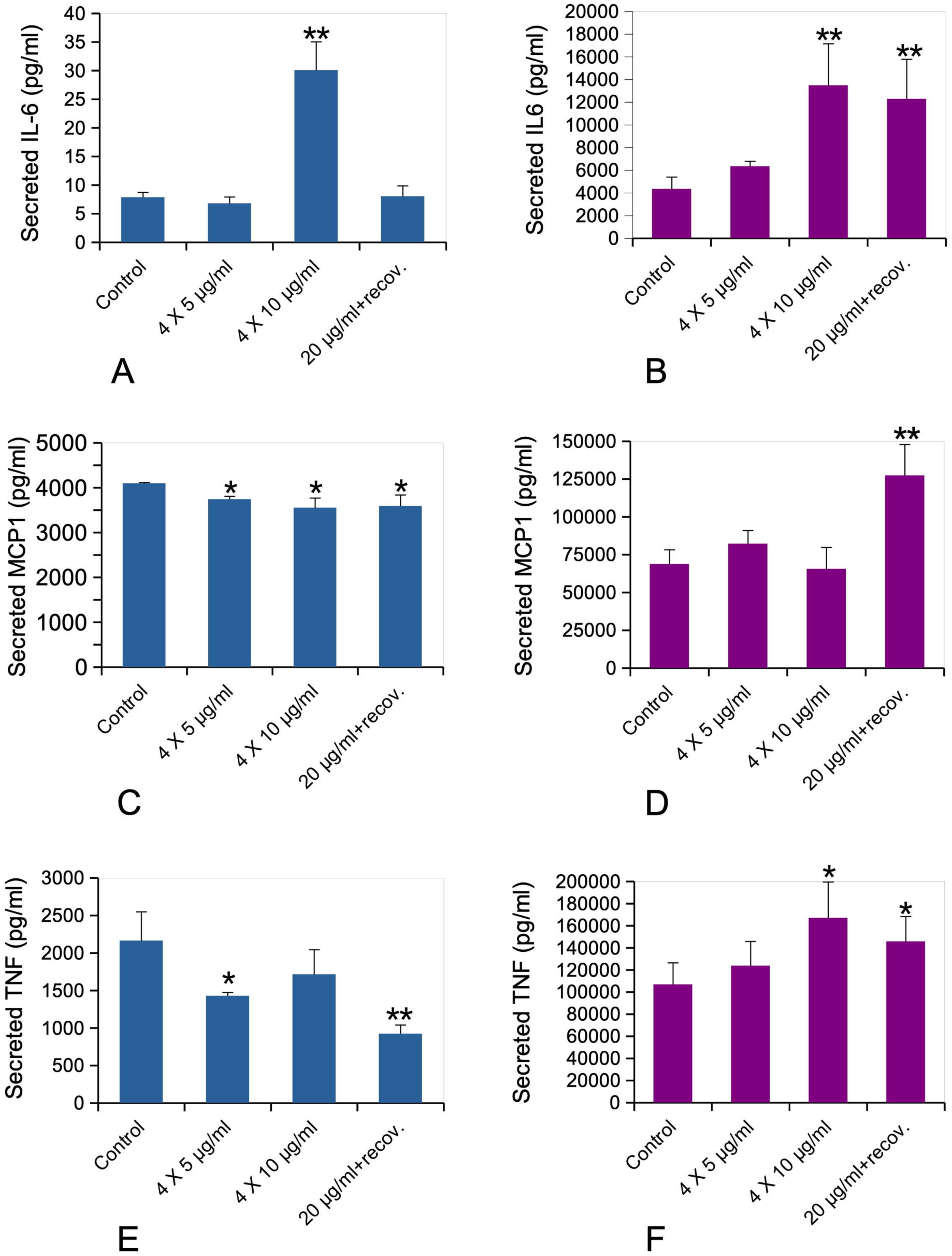
Cytokine release. The cells were exposed to MnO_2_ particles according to the three schemes used in this work. The cells were treated (or not) with 50ng/ml lipopolysaccharide in complete cell culture medium for the last 24 hours. The cell medium was then collected for secreted IL-6, MCP-1 and TNF alpha measurements. Panel A: IL-6 release without LPS stimulation Panel B: IL-6 release with LPS stimulation (50 ng/ml) Panel C: MCP-1 release without LPS stimulation Panel D: MCP-1 release with LPS stimulation (50 ng/ml) Panel E: TNF-alpha release without LPS stimulation Panel F: TNF-alpha release with LPS stimulation (50 ng/ml) *: statistically different from the unexposed control (p<0.05, Student T test) **: statistically different from the unexposed control (p<0.01, Student T test)

Testing of the phagocytic function showed a small but significant increase in both the proportion of phagocytic cells (**Figure 8A**) and the internalization capacity of the phagocytic cells (**Figure 8B**).

Regarding the secretion of pro-inflammatory cytokines, we tested two schemes. First the response to MnO_2_ only (**Figure 9 A, C, E**) and then the response to MnO_2_ , followed by a final stimulation with LPS (**Figure 9 B, D, F**).

The effects of MnO_2_ alone were dependent on the exposure conditions. A strong stimulation of IL-6 secretion was observed only in the 4x10 µg/ml condition (**Figure 9A**), while a small decrease in the secretion of the chimiokine MCP1 was observed under all exposure scenarios (**Figure 9C**). The situation was more complex for TNF, with a small but significant decrease in the 4x5 µg/ml and recovery scenarios, while the change was not significant for the 4x10 µg/ml condition (**Figure 9E**). The responses were significantly altered when cells were stimulated with LPS in addition to the MnO_2_ treatments. IL-6 secretion was strongly induced in the 4x10 µg/ml and recovery conditions, as was TNF secretion (but to a lesser extent). MCP1 was increased only in the recovery condition.

## 4. Discussion

Because of its use as a pigment, where sizing is known to alter the color and is thus controlled, Pigment Black 14 represented an easy and controlled source of nano-sized manganese dioxide. When macrophages were treated with these manganese dioxide nanoparticles, the first striking result was the onset of the delayed toxicity that we observed. It is likely due to the rather fast dissolution of MnO_2_ when internalized in the cells. Indeed this dissolution was intermediate in speed between the very fast dissolution of zinc oxide, which can produce its toxic effect directly as an extracellular ion [74] ,(although in highly phagocytic cells such as macrophages, intracellular zinc oxide can be observed [75] ), and the much slower dissolution of e.g. silver, where no delayed toxicity could be observed [76], probably because the cells can adapt to the low concentration of silver ion that is progressively liberated.

This delayed toxicity also explained why we could use a cumulated dose of 40µg/ml in repeated exposure, while it could not be used in the recovery scheme where the delayed toxicity fully developed. In the repeated exposure mode, the first low, non toxic doses are likely to induce a protective response in cells, which protects them against the following doses.

However, when dealing with supposed professional exposure using in vitro systems, it is important to determine relevant exposure doses. To this purpose, we used a model of lung exposure. A day of normal breathing is equivalent to the inhalation of 10 cubic meters of air. As manganese concentration up to 18 mg Mn/m^3^ in air have been observed in dry cell batteries factories [35] , this corresponds to up to 180 mg of inhaled manganese per day. If we assume that this manganese is taken up by the alveolar macrophages, these 180 mg are distributed between the ca. 7 billions alveolar macrophages present in the lungs (1.4 billion per lobe [77] x 5 lobes). This represents ca. 25 µg Mn/million macrophages per day, i.e. 40 µg MnO_2_/million macrophages and per day. As we work with macrophages concentrations of 1 million cells/ml, this would represent 40 µg MnO_2_/ml and per day. Of course these raw calculations are probably imprecise and give an upper threshold, but it means in turns that the concentrations that we have used are in an adequate range, even for the recovery exposure scheme.

In our work, we described a mitochondrial hyperpolarization (increased mitochondrial transmembrane potential) in response to MnO_2_ , while the opposite was described by Alhadlaq et al. on other cell lines [40] . However, this apparent discrepancy may be due to a questionable interpretation of their results by Alhadlaq et al. Indeed, they mention in their paper that they use the same rhodamine 123 concentration than in a previous paper of theirs [78] , i.e. 10µM. Unfortunately, this concentration is well above the quenching threshold [61], so that the mitochondrial transmembrane potential evolves in this case oppositely to the fluorescence signal [79] . In the absence of a negative control such as CCCP or FCCP [61, 62, 79], which we also used to determine the correlation between fluorescence intensity and mitochondrial transmembrane potential, this is the most likely hypothesis. Furthermore, manganese has been shown to bind to and inhibit ATP synthase [80], which is consistent with the concentration-dependent increase of the mitochondrial transmembrane potential observed in our study.

Within this scheme, the strong induction of the ATP synthase inhibitor (O35143) was somewhat puzzling. This protein is known to inhibit ATP synthase when the transmembrane potential is very low, so that ATP synthase works in the reverse direction to maintain the proton gradient [81]. However, by inhibiting the ATP synthase, manganese increases the transmembrane potential. This apparent discrepancy is solved by the fact that an induction of the amount of ATP synthase inhibitor protein has been shown to decrease the transmembrane potential [82]. Thus, the observed increase in the amount of ATP synthase inhibitor protein upon treatment with MnO_2_ can be seen as an attempt by the cells to normalize the transmembrane potential which is increased by the manganese-driven inhibition of ATP synthase [80].

The same perturbation in mitochondria has been described on the whole worm C. elegans in response to MnCl_2_ [83]. However, the authors also described an increase in the intracellular iron content in response to MnCl_2_ , while we did not observe this phenomenon in our case (Table 1). This may be due either to the different exposure scheme, as Angeli et al. used an acute exposure to a soluble manganese salt and observed this effect immediately after exposure, or to the fact that our study is centered on macrophages, which are known to have a specific iron metabolism [84].

More generally it appears that manganese disturbs the general energy metabolism of the cells, including the glycolysis and pentose phosphate pathways. This phenomenon has been described on other cell types, with an increase in the GAPDH activity [85] and a decrease in hexokinase, pyruvate kinase and lactate dehydrogenase activities [86]. Regarding GAPDH, the results described on astrocytes with zinc by Hazell et al. [85] coincide well with the results that we obtained on macrophages with zinc oxide [87]. As glycolysis is a highly conserved pathway, it is likely that the induction observed on astrocytes with manganese by Hazell et al. [85] will also occur in macrophages. Regarding the other glycolysis enzymes, we observed an increase in their amounts while Malthankar et al. observed a decrease in their activities [86]. This shall not be viewed as a discrepancy since 1) enzyme activities do not correlate so well with their amounts detected in proteomics [88, 89] , and 2) the increase in the protein amounts may be a compensatory mechanism for a not-so-working glycolysis process, as described for example for schizophrenia [90] . This suggests that on the whole, manganese disturbs the energy metabolism at both the cytosolic and mitochondrial levels.

However, this mechanism is different from the one observed in response to ZnO nanoparticles [87]. In this latter case, a decrease in mitochondrial potential was observed instead of the increase observed here with MnO_2_. For zinc oxide, a decrease in the GAPDH activity was observed, which induced in turn a response to toxic aldehydes such as methyglyoxal.

In the response to MnO_2_ nanoparticles we did not observed a response to alkyl aldehydes (**Table S13**) although a response to formaldehyde was observed through the induction of formylglutathione hydrolase (Q9R0P3). Indeed, it has been shown that formaldehyde production may be linked to oxidative stress through the 1C metabolism [91], which is in line with our observed induction of oxidative stress. This hypothesis, is further substantiated by the observed decrease of DHFR (P00375) in response to manganese, as DHFR produces tetrahydrofolate, which is itself the formaldehyde precursor [91].

As manganese is very close from iron in transition metal series of the periodic table, it was quite interesting to compare the responses induced by MnO_2_ to those induced by iron oxide pigments [92]. Indeed, in the recovery scheme that was also tested for iron pigments, MnO_2_ induced many common effects with Fe_2_O_3_ pigments, such as an increase in the mitochondrial transmembrane potential, oxidative stress and increase in glutathione level.

Despite these common outcomes, the molecular response at the proteome level was quite different between MnO_2_ and Fe_2_O_3_. For example, different subunits of the Oxphos and MICOS complexes are modulated in response to MnO_2_ and Fe_2_O_3_. Regarding the protective response, although formyl glutathione hydrolase is induced in response to both metallic oxides, ferritin and heme oxygenase are strongly induced in response to Fe_2_O_3_, and not in response to MnO_2_ (Table S13), which shows their specificity toward iron compared to the closely related manganese ion. The oxidative stress response is also strongly different: while Mn superoxide dismutase and the regulatory subunit of glutamate cysteine ligase (the limiting step in glutathione biosynthesis) are strongly induced in response to both metallic oxides, the response is different for glutaredoxin, catalase, peroxiredoxins and thioredoxin family proteins.

Regarding more specific functions of macrophages, such as phagocytosis and lysosomal activity, MnO_2_ induced an increase in phagocytosis under these precise exposure schemes, which we did not observe in response to other particles such as PET [93] or polystyrene [53] particles. However, an increase in the lysosomal activity was observed in response to MnO_2_ particles, as it was in response to Fe_2_O_3_ particles [92].

Moreover, the inflammatory response was quite different between the two pigments in this precise exposure scheme. While Fe_2_O_3_ alone induced an increase in the production of IL-6 and TNF, a stable IL-6 production and a decreased TNF production were observed in response to MnO_2_. Conversely, when a double treatment with metal oxides and LPS was carried out, MnO_2_ induced an increased production of IL-6 and TNF, while a decrease in the production of IL-6 and an increase in the production of TNF was observed in response to Fe_2_O_3_ .

By contrast, the repeated exposure to MnO_2_ at the 4x10 µg/ml dose induced an increase in IL-6 production with or without LPS stimulation, and an increase in TNF production with LPS stimulation. This indicated an increased pro-inflammatory response of macrophages induced by this repeated exposure, which is in line with the observed respiratory effects of MnO_2_ [47].

## Conclusions

By using non-classical, although relevant exposure schemes, this work showed strong effects of MnO_2_ on macrophages, substantiating at the proteomic level prior work on the pulmonary toxicity of manganese [33, 42, 47, 94]. Of note are the mitochondrial effects of MnO_2_, which induce in turn oxidative stress. Furthermore, a pro-inflammatory response is also induced. As the coupling of chronic inflammation and sustained oxidative stress has been implicated in chronic pathologies such as asbestosis [95] and silicosis [96] , this means in turn that the possible effects of manganese com-pounds and especially MnO_2_ at the pulmonary level should be more scrutinized, in addition to the now well-known cerebral effects of manganese.

## Acknowledgments

We thank Guy Schoehn for the establishment of the IBS/ISBG EM facility

## Data availability

The proteomic data are available via ProteomeXchange with the identifier PXD065155

The cell biology data are available through the BioStudies database under the DOI 10.6019/S-BSST2154

## Authors contributions

MV, EA, VCF, FD, HD, DF: investigation, formal analysis

SR, CC, TR: formal analysis, funding acquisition, project administration

MV, HD, SR, TR: Writing – original draft

TR: conceptualization

All co-authors: Writing – review and editing

