## Supplemental Table 1 for "Effects of manganese dioxide on macrophages under different exposure schemes"

Supplementary Table 1: list of proteins detected and quantified by the proteomic screen

| accession number | gene_name | description | protein_set _score | coverage | MW | #peptides | Ctrl 1 | Ctrl 2 | Ctrl 3 | Ctrl 4 | Ctrl 5 | PBK14 20 recov 1 | PBK14 20 recov 2 | PBK14 20 recov 3 | PBK14 20 recov 4 | PBK14 20 recov 5 | PBK14 4x5 rep1 | PBK14 4x5 rep2 | PBK14 4x5 rep3 | PBK14 4x5 rep4 | PBK14 4x5 rep5 | PBK14 4x10 rep1 | PBK14 4x10 rep2 | PBK14 4x10 rep3 | PBK14 4x10 rep4 | PBK14 4x10 rep5 |
|---|---|---|---|---|---|---|---|---|---|---|---|---|---|---|---|---|---|---|---|---|---|---|---|---|---|---|
| Q6GQT1 | A2m | Alpha-2-macroglobulin-P | 98.24 | 1.29 | 164353 | 2 | 910942.5 | 113359.84 | 1169541.62 | 956407.88 | 1067141.38 | 2 | 167421.91 | 174487.77 | 1353232.88 | 1247312.88 | 67622.38 | 2 | 774092.19 | 171814.47 | 1413823.12 | 2 | 226369.14 | 235224.86 | 162200.67 | 2 |
| P58742 | Aaas | Aladin | 242.87 | 10.62 | 59431 | 5 | 3154026.25 | 617844.38 | 1826723.75 | 2448212.5 | 2108634.5 | 2584982 | 2185668.5 | 1582836.25 | 805614.12 | 2102648.25 | 483099.38 | 506121.97 | 1658671.12 | 703082.94 | 1210714 | 715994.44 | 2 | 1185492.5 | 1546874 | 2 |
| Q9D2R0 | Aacs | Acetoacetyl-CoA synthetase | 208.85 | 7.29 | 75200 | 5 | 1574391.38 | 1233439.88 | 1725142.5 | 2058007.75 | 3789423.25 | 2 | 1116855.75 | 1785806.75 | 1152660.75 | 1774156 | 2710656.5 | 1508483.88 | 1291147.5 | 1919557.5 | 2310147.5 | 1038143.75 | 2682955.75 | 2109125.75 | 1504172 | 2174250 |
| Q8BGQ7 | Aars1 | Alanine--tRNA ligase, cytoplasmic | 1040.82 | 23.86 | 106909 | 19 | 14857398 | 15724374 | 22310118 | 16816902 | 32632782 | 28528192 | 24486130 | 24159734 | 27561500 | 29517566 | 37923506 | 35735354 | 40622742 | 37853658 | 40428814 | 41443006 | 46812290 | 47575250 | 44113666 | 47680474 |
| Q14CH7 | Aars2 | Alanine--tRNA ligase, mitochondrial | 38.01 | 1.02 | 106783 | 1 | 1787113.25 | 919898.62 | 1093745.12 | 1682108.5 | 2 | 1952142 | 1410547.75 | 2252466.75 | 557929.44 | 949567.19 | 2 | 1222469.5 | 1401350.12 | 890133.56 | 1354080.75 | 1178950.25 | 2025373.12 | 1973335.12 | 1520343.62 | 2 |
| Q3THG9 | Aarsd1 | Alanyl-tRNA editing protein Aarsd1 | 38.45 | 2.43 | 44971 | 1 | 2 | 809174.62 | 596254 | 400109.28 | 979074.56 | 2 | 802522.81 | 2 | 663535.25 | 817514 | 2 | 1000583.94 | 2 | 1004711.12 | 2 | 1368484.38 | 2 | 2 | 2 | 2 |
| Q9JKX4 | Aatf | Protein AATF | 110.05 | 4.75 | 59482 | 2 | 2 | 116404.89 | 61446.05 | 233374.38 | 69092.99 | 249812.06 | 2 | 2 | 2 | 2 | 2 | 96576.89 | 86655.72 | 180035.36 | 2 | 2 | 2 | 2 | 2 | 2 |
| P06795 | Abcb1b | ATP-dependent translocase ABCB1 | 130.3 | 3.84 | 140994 | 2 | 2 | 2 | 2 | 2 | 2 | 2 | 2 | 2 | 2 | 2 | 2 | 2 | 2 | 2 | 2 | 2 | 2 | 2 | 2 | 2 |
| Q61102 | Abcb7 | Iron-sulfur clusters transporter ABCB7, mitochonc | 167.68 | 4.79 | 82581 | 3 | 2724526 | 1439373.12 | 1777288 | 2230966 | 970211.81 | 197625.11 | 196476.97 | 1742181 | 2469451 | 1338488 | 2 | 1483278.12 | 2298618.75 | 2853671.75 | 1805382.75 | 2 | 1679857.25 | 2241865.5 | 1745019.5 | 468770.84 |
| O35379 | Abcc1 | Multidrug resistance-associated protein 1 | 139.88 | 2.55 | 171185 | 3 | 73606.3 | 116741.29 | 77272.27 | 292157.66 | 2 | 128686.99 | 195494.64 | 196670.14 | 222431.14 | 298854.47 | 104314.24 | 101222.12 | 149251.77 | 266666.16 | 341144.56 | 2 | 683839 | 232727.47 | 91093.26 | 2 |
| B2RX12 | Abcc3 | ATP-binding cassette sub-family C member 3 | 142.26 | 2.56 | 169223 | 3 | 746961.12 | 2 | 888048.25 | 937087.94 | 913032.5 | 626987.69 | 857365.75 | 1412054.12 | 1135170.25 | 1409628.62 | 781985.88 | 2 | 956828.06 | 747923.19 | 644769.31 | 946910.62 | 1000792.88 | 683978.38 | 2 | 2 |
| O88269 | Abcc6 | ATP-binding cassette sub-family C member 6 | 38.24 | 0.47 | 164995 | 1 | 2 | 2 | 2 | 2 | 2 | 695646.69 | 446254.91 | 2 | 2 | 238471.39 | 384541.78 | 347615.19 | 2 | 2 | 825468.56 | 2 | 2 | 2 | 590076.25 | 2 |
| P55096 | Abcd3 | ATP-binding cassette sub-family D member 3 | 83.76 | 3.34 | 75475 | 2 | 599392.06 | 1020571 | 513711.19 | 645539.75 | 688396 | 735577.19 | 1081702 | 638016.06 | 719588.44 | 781535.12 | 2 | 2 | 893601.94 | 706414.44 | 352195.5 | 556350.25 | 1202366.38 | 1120478.5 | 1279762.88 | 1033942.75 |
| P61222 | Abce1 | ATP-binding cassette sub-family E member 1 | 175.41 | 8.51 | 67314 | 4 | 2 | 2 | 104017.3 | 636664.5 | 314866.66 | 2 | 94439.93 | 1064107.62 | 199978.86 | 2 | 209665.58 | 2 | 456427.5 | 1127618.88 | 2 | 84242.33 | 2 | 467475.38 | 264361.88 | 2 |
| Q6MG08 | Abcf1 | ATP-binding cassette sub-family F member 1 | 65.05 | 2.38 | 95252 | 2 | 2 | 733240 | 2 | 643776.62 | 1587359.75 | 2 | 449785.06 | 907126.94 | 2 | 815864 | 659697 | 1264973 | 2 | 2 | 783709 | 2 | 2196948.25 | 2 | 3113020.75 | 1201255.88 |
| Q6PE15 | Abhd10 | Palmitoyl-protein thioesterase ABHD10, mitochon | 44.52 | 3.7 | 33040 | 1 | 2 | 2 | 2 | 2 | 2 | 2 | 2 | 2 | 2 | 2 | 2 | 2 | 2 | 2 | 237383.33 | 2 | 2 | 2 | 2 | 2 |
| Q6AYT7 | Abhd12 | Lysophosphatidylserine lipase ABHD12 | 210.14 | 14.57 | 45296 | 4 | 2 | 2 | 2 | 2 | 1329202 | 233385.81 | 162717.11 | 1035809.94 | 1738530.75 | 1535638.12 | 1327822.88 | 1268811.38 | 1163614.75 | 418129.75 | 132839.12 | 2 | 2 | 2 | 2 | 339556 |
| Q6DGG1 | Abhd14b | Putative protein-lysine deacylase ABHD14B | 55.66 | 5.24 | 22618 | 1 | 447004.19 | 2 | 487994.19 | 281489.12 | 311151.19 | 2 | 667523.44 | 2 | 2 | 415968.72 | 2 | 2 | 2 | 508769.91 | 2 | 2 | 2 | 2 | 2 | 2 |
| Q6MG55 | Abhd16a | Phosphatidylserine lipase ABHD16A | 94.32 | 3.76 | 63038 | 2 | 2 | 2 | 2 | 2 | 2 | 2 | 2 | 2 | 551708.19 | 2 | 2 | 2 | 2 | 539457.94 | 2 | 2 | 2 | 2 | 2 | 2 |
| B5DFK7 | Abhd17c | Alpha/beta hydrolase domain-containing protein 1 | 91.35 | 5.62 | 35098 | 2 | 2346624 | 876472.38 | 1575737 | 647012.56 | 2 | 2 | 1038585.88 | 4449084 | 2 | 983758.38 | 676623.31 | 761424.81 | 534403.69 | 670543.5 | 2356109 | 1936339 | 607525.81 | 3352568.5 | 713195.69 | 3993890.5 |
| Q6QA69 | Abhd5 | 1-acylglycerol-3-phosphate O-acyltransferase AB | 52.86 | 4.56 | 39104 | 1 | 2 | 2 | 99175.3 | 144968.28 | 62399.81 | 2 | 344955.62 | 136892.09 | 340515.06 | 539073.19 | 2 | 91393.94 | 2 | 374000.12 | 407251.38 | 2 | 2 | 2 | 2 | 368581.62 |
| Q8CBW3 | Abi1 | Abl interactor 1 | 121.21 | 5.61 | 52288 | 2 | 2910481.75 | 3416475.75 | 2762181 | 2067430.5 | 2404614 | 3419500.75 | 2889287.25 | 3350285 | 1423159.88 | 2294113.75 | 2216953.75 | 1897014.12 | 1907361.5 | 1904903.88 | 2364202.25 | 2585467.5 | 3149335.25 | 3517339.5 | 2543449.25 | 3275168.25 |
| P00520 | Abl1 | Tyrosine-protein kinase ABL1 | 55.55 | 0.71 | 122673 | 1 | 1397034.75 | 639634.62 | 2 | 970654.12 | 2 | 963209.31 | 2 | 2 | 313910.03 | 690571.19 | 1234880.38 | 2 | 1220023.62 | 1624548.75 | 3036790.5 | 1916466.5 | 3021854.25 | 2281575.75 | 2655601.5 | 3898689 |
| A0A0G2 | Abr | Active breakpoint cluster region-related protein | 75.41 | 1.98 | 97693 | 2 | 1478263.62 | 2632615.25 | 1817237.5 | 2046099.25 | 845149.38 | 2827181 | 2627192 | 4017348 | 840522.19 | 2158754.5 | 1732267.38 | 843833.31 | 939904.56 | 1055789.12 | 912121.19 | 2991135 | 2911798.5 | 4207900.5 | 1110420.88 | 3204388.5 |
| Q4KML4 | Abracl | Costars family protein ABRACL | 213.89 | 50.62 | 9030 | 4 | 484789.72 | 120272.49 | 2337638.5 | 867623.94 | 777305.31 | 751256.5 | 1476564.5 | 1210289.62 | 85934.16 | 1271148.25 | 962084.62 | 1046399.75 | 1345466.25 | 239288.27 | 2604042.75 | 605073 | 1501829.62 | 1172407.88 | 2 | 149558.11 |
| Q921H8 | Acaa1a | 3-ketoacyl-CoA thiolase A, peroxisomal | 873.27 | 46.23 | 43953 | 13 | 12110358 | 10982388 | 13554334 | 14228259 | 20047524 | 7530803 | 14262904 | 9776953 | 19382448 | 14465819 | 10789693 | 13384217 | 10824546 | 13473834 | 8976255 | 3574571.25 | 8340243 | 6947361 | 6106293 | 7106682 |
| Q8BWT1 | Acaa2 | 3-ketoacyl-CoA thiolase, mitochondrial | 215.66 | 9.57 | 41830 | 3 | 1685788.38 | 4060130.5 | 3647722 | 1710022.75 | 2401469.25 | 1965904.12 | 1585725.75 | 2290765.25 | 1949724.12 | 151453.09 | 1479511.38 | 1383788.12 | 1573694.62 | 1789998.62 | 1870887.75 | 1307730.38 | 1633184.62 | 1472161.88 | 1676436.5 | 1608282.38 |
| Q9D7B6 | Acad8 | Isobutyryl-CoA dehydrogenase, mitochondrial | 145.64 | 7.75 | 45020 | 3 | 1150982.88 | 3561105 | 3260042 | 1096432.75 | 1032790.38 | 1454686.88 | 2923181.25 | 1771341.62 | 2358628 | 2297220.25 | 895350.88 | 1406218 | 1618052.75 | 1908556.25 | 814624.19 | 933163.88 | 1095027.38 | 1251247 | 898567.94 | 3163562.5 |
| Q8JZN5 | Acad9 | Complex I assembly factor ACAD9, mitochondrial | 375.79 | 15.04 | 68722 | 8 | 5865435 | 4934719 | 8300346 | 6313488.5 | 7132175 | 4912872 | 2912914 | 5494705.5 | 5289403 | 4681942.5 | 1002076.81 | 2025586.75 | 3740948 | 2678594.5 | 3827852.5 | 690338.5 | 1965555.75 | 2042634.25 | 2621769.5 | 1648101.88 |
| P51174 | Acadl | Long-chain specific acyl-CoA dehydrogenase, mi | 777.47 | 31.63 | 47908 | 13 | 461522.19 | 303916.34 | 962936.25 | 545305.69 | 327519.66 | 263488.84 | 407188.03 | 212387.72 | 693438.5 | 718974.25 | 357995.16 | 282784.44 | 278867.5 | 482169.16 | 1328076.88 | 242989.67 | 2 | 2 | 2 | 697590.19 |
| P45952 | Acadm | Medium-chain specific acyl-CoA dehydrogenase, | 275.9 | 15.68 | 46481 | 6 | 7691107.5 | 6979820 | 14773330 | 11226067 | 14993330 | 11113062 | 9802073 | 11218961 | 12901425 | 16495349 | 15883484 | 14724453 | 16898546 | 15326938 | 11504769 | 13162482 | 17629214 | 20324322 | 13927075 | 18865074 |
| P15651 | Acads | Short-chain specific acyl-CoA dehydrogenase, mi | 259.74 | 16.75 | 44765 | 5 | 3051880 | 3058870.75 | 4769671 | 1578676.25 | 3866610.5 | 2 | 2209676.25 | 1459848.88 | 2701932 | 2216938.25 | 2022219.12 | 2032165.88 | 1846466.5 | 1558575.38 | 2303273 | 2 | 886844.56 | 2066879.75 | 1369988.75 | 2 |
| Q9DBL1 | Acadsb | Short/branched chain specific acyl-CoA dehydrog | 207.01 | 11.57 | 47874 | 4 | 7018289.5 | 14215144 | 9009966 | 9093174 | 6199821 | 8367582 | 7702696.5 | 8207247 | 6950683 | 7949475 | 5699357 | 3853466 | 4785689.5 | 6292184.5 | 3182772.5 | 4583921.5 | 3738578.75 | 3548650.25 | 5525142 | 3196016.25 |
| P50544 | Acadvl | Very long-chain specific acyl-CoA dehydrogenase | 284.45 | 14.02 | 70875 | 6 | 2 | 70275.31 | 79263.61 | 59675.61 | 810189.5 | 2 | 136818.83 | 63004.89 | 628605.94 | 797179.38 | 1178319.12 | 1252342.5 | 619003.94 | 3071794.75 | 583562.06 | 1103071.62 | 176685.84 | 558532.25 | 886448.75 | 120626 |
| Q8QZT1 | Acat1 | Acetyl-CoA acetyltransferase, mitochondrial | 663.07 | 33.25 | 44816 | 10 | 4057124.5 | 6374660.5 | 3994501.75 | 6695482 | 18768068 | 5695397 | 7453870 | 7394994.5 | 14615872 | 11774977 | 5706702 | 6678197 | 7156683 | 6850859.5 | 7459854 | 2151751 | 4683642 | 4347943 | 2942792.5 | 3759518 |
| Q8CAY6 | Acat2 | Acetyl-CoA acetyltransferase, cytosolic | 254.6 | 17.88 | 41298 | 5 | 1392575.5 | 1746129.25 | 240530.69 | 95975.54 | 2681284.5 | 4303969.5 | 3356039.5 | 3578856.25 | 1851600.12 | 1124336 | 2676739 | 3018225 | 1391103 | 2905143 | 1842299.88 | 1449582.38 | 2139814.25 | 1263280.12 | 1456947.12 | 2540682.75 |
| Q8BMP6 | Acbd3 | Golgi resident protein GCP60 | 85.45 | 3.05 | 60181 | 1 | 2 | 2 | 2 | 2 | 2 | 2 | 2 | 2 | 2 | 2 | 773232.31 | 2 | 506758.19 | 617358.69 | 2 | 2 | 2 | 2 | 2 | 2 |
| Q9JIX8 | Acin1 | Apoptotic chromatin condensation inducer in the n | 319.07 | 6.05 | 150719 | 7 | 606928.56 | 360793.91 | 430008.47 | 2738812.25 | 3932057 | 3142642.75 | 1704834.75 | 4061592.5 | 3574682.25 | 1994209 | 5102157.5 | 4063078.75 | 4961646 | 3748733.5 | 1129638.88 | 4681146.5 | 2851197 | 2814875.5 | 5308140 | 2034453.12 |
| P16638 | Acly | ATP-citrate synthase | 1267.55 | 20.82 | 120636 | 19 | 10764280 | 12073920 | 12577072 | 25875580 | 33139886 | 24824086 | 24732826 | 32984408 | 21849850 | 17316538 | 21090794 | 21959984 | 19965678 | 20014532 | 9839844 | 18106666 | 17511744 | 16071986 | 15848867 | 15717522 |
| P28271 | Aco1 | Cytoplasmic aconitate hydratase | 41.59 | 1.35 | 98126 | 1 | 1090358 | 1434811.5 | 927537 | 960506.75 | 831564.44 | 913840.81 | 726633 | 1103833.62 | 542521.94 | 2 | 654063.81 | 627699.31 | 800134.31 | 625394 | 2 | 948901.38 | 2 | 2 | 2 | 2 |
| Q99KI0 | Aco2 | Aconitate hydratase, mitochondrial | 1222.86 | 30 | 85464 | 21 | 25205842 | 24182628 | 23407314 | 17610272 | 32737620 | 32467474 | 30172486 | 29234874 | 43662334 | 37533214 | 31377408 | 30659816 | 33336246 | 30630990 | 18213444 | 31969666 | 27576552 | 19485582 | 31092118 | 12738785 |
| Q32MW3 | Acot10 | Acyl-coenzyme A thioesterase 10, mitochondrial | 106.69 | 4.1 | 50552 | 3 | 2 | 2 | 2 | 2 | 2 | 2 | 293558.69 | 2 | 606981.75 | 2 | 890970.25 | 531893.5 | 2 | 2 | 2 | 2 | 2 | 2 | 2 | 2 |
| Q9CQR4 | Acot13 | Acyl-coenzyme A thioesterase 13 | 148.54 | 28.57 | 15183 | 3 | 1088837.5 | 2074729.5 | 1445821 | 1207452 | 1825490 | 2 | 1126604.25 | 2120393 | 1531588.12 | 1269935.5 | 2520625 | 2613735.75 | 2764020.75 | 2564028.5 | 1266258.5 | 1332685.88 | 2051726 | 2897519.5 | 2243210.5 | 3308565.75 |
| O55171 | Acot2 | Acyl-coenzyme A thioesterase 2, mitochondrial | 150.52 | 9.27 | 49701 | 3 | 2 | 2 | 2 | 513100.56 | 1294147 | 548905 | 814313.81 | 1721972.25 | 69583.21 | 1502437.88 | 494270.38 | 779888.44 | 2 | 2 | 2 | 2 | 2 | 2 | 2 | 2 |
| Q91V12 | Acot7 | Cytosolic acyl coenzyme A thioester hydrolase | 238.91 | 16.27 | 42537 | 5 | 2703190 | 3356388 | 3565842 | 2 | 4401150 | 3701512.25 | 3835209.25 | 4236185 | 1242605.62 | 1431446.25 | 6084909.5 | 6206951.5 | 4127295 | 4740934 | 2971209.5 | 4057481.5 | 5491791 | 4560275 | 4650192 | 4416386 |
| P58137 | Acot8 | Acyl-coenzyme A thioesterase 8 | 140.96 | 12.5 | 35827 | 3 | 472621.66 | 2 | 372975.59 | 342082.69 | 604148.38 | 2 | 461638.75 | 2 | 2 | 2 | 2 | 2 | 348685.94 | 2 | 2 | 2 | 2 | 2 | 2 | 2 |
| Q63448 | Acox3 | Peroxisomal acyl-coenzyme A oxidase 3 | 100.25 | 3.71 | 78446 | 2 | 2 | 788665.62 | 2 | 2 | 548009.12 | 2 | 2 | 2 | 801669.5 | 239207.03 | 2 | 2 | 2 | 245683.58 | 150289.38 | 2 | 2 | 2 | 2 | 291440.38 |
| P41498 | Acp1 | Low molecular weight phosphotyrosine protein ph | 168.01 | 13.92 | 18152 | 3 | 7733582 | 5949479 | 6473837 | 4231382.5 | 7086816 | 4303485.5 | 4753398 | 4807112 | 4046285 | 5353144 | 8176747 | 8014165.5 | 10081861 | 7470591.5 | 12888423 | 9088131 | 9813065 | 8269658 | 9215791 | 7692987 |
| P20611 | Acp2 | Lysosomal acid phosphatase | 89.05 | 4.26 | 48320 | 2 | 1293806.38 | 1468385.88 | 1497668.12 | 4111840 | 864008.31 | 1475663.75 | 3618455.25 | 3410732.5 | 1130844.88 | 2086831.5 | 2034910.5 | 586143.75 | 1384755.88 | 844689 | 896655.94 | 978879.25 | 1014803.69 | 1184719.25 | 573371 | 1358322.25 |
| Q8BP40 | Acp6 | Lysophosphatidic acid phosphatase type 6 | 161.23 | 9.81 | 47625 | 3 | 1265285.25 | 77633.34 | 2296400 | 872767.5 | 2723702.75 | 2 | 980794.31 | 1433239.75 | 1428926.38 | 2710342 | 717916.88 | 1797177.12 | 910706.69 | 1857611.12 | 2 | 923343.19 | 2 | 2 | 898265.25 | 2 |
| Q8VCW8 | Acsf2 | Medium-chain acyl-CoA ligase ACSF2, mitochonc | 124.25 | 9.59 | 67951 | 3 | 1972319.38 | 1903465.5 | 1965640.62 | 936469.06 | 1827733.12 | 1265208.75 | 1153289.75 | 1553528.5 | 2262132.75 | 1651313.12 | 1430726.75 | 1196965.88 | 1894535.12 | 1292320.38 | 415904.62 | 2 | 2 | 1460940.38 | 773044.94 | 2 |
| P41216 | Acsl1 | Long-chain-fatty-acid--CoA ligase 1 | 748.87 | 20.46 | 77951 | 13 | 15011382 | 7030842.5 | 15467761 | 9452538 | 12973224 | 8044199 | 9666626 | 9527546 | 13460395 | 16206658 | 4069211 | 3346427.75 | 6735819 | 3750970.25 | 7076272.5 | 4907084 | 3824109 | 8155899.5 | 1758339 | 6194244 |
| Q9CZW4 | Acsl3 | Fatty acid CoA ligase Acsl3 | 414.45 | 19.17 | 80492 | 9 | 604833.38 | 163640 | 898090.12 | 1054112.5 | 2606169 | 2 | 855332.31 | 1487423 | 3367468.75 | 1932792.25 | 126500.71 | 1070403.38 | 2877102 | 2848384 | 471670.5 | 1279034.25 | 2 | 2 | 1410819 | 2 |
| Q9QUJ7 | Acsl4 | Long-chain-fatty-acid--CoA ligase 4 | 704.09 | 20.25 | 79077 | 13 | 8631226 | 15733784 | 6813302 | 12792210 | 10441894 | 18128884 | 17248188 | 16297322 | 13496159 | 9729802 | 6783487.5 | 6693440 | 6636458.5 | 8391071 | 6725141 | 5313456.5 | 6653838 | 8528710 | 5275641 | 4391894.5 |
| Q8JZR0 | Acsl5 | Long-chain-fatty-acid--CoA ligase 5 | 442.28 | 17.57 | 76206 | 10 | 1590623 | 1377870.75 | 2888586.25 | 2468746.5 | 5118126.5 | 1112959.38 | 1982411.5 | 3642007.5 | 3120942.75 | 3590626.5 | 2430864.5 | 1299556.25 | 1430763.25 | 1384326 | 2504015.5 | 2314255.25 | 1791126.38 | 1758562.88 | 2254611.5 | 1991893.12 |
| P60710 | Actb | Actin, cytoplasmic 1 | 2631.91 | 91.73 | 41737 | 36 | 149365.02 | 538855.12 | 448502.16 | 2 | 326613.28 | 2 | 2 | 2 | 208345.31 | 2 | 349789.56 | 799870.38 | 860897.06 | 1983560.5 | 1520018.38 | 2 | 1384757.12 | 1591499.75 | 2 | 1315200.38 |
| Q8BFZ3 | Actbl2 | Beta-actin-like protein 2 | 736.96 | 39.36 | 42004 | 14 | 1645875.62 | 1728979.25 | 1996273.38 | 2 | 1651825.25 | 2 | 726752.31 | 848372.12 | 402927.16 | 1074791 | 2 | 2 | 2 | 2 | 2 | 2 | 2 | 2 | 2 | 2 |
| P63259 | Actg1 | Actin, cytoplasmic 2 | 2642.15 | 91.73 | 41793 | 36 | 131565.94 | 446769.88 | 449241.94 | 2 | 118483.15 | 339154.06 | 2 | 503080.47 | 345015 | 407441.69 | 206246.73 | 653686.12 | 738996.31 | 2158185.5 | 1957427.38 | 2 | 1238970.88 | 1530309.5 | 2 | 715256 |
| Q9Z2N8 | Actl6a | Actin-like protein 6A | 410.94 | 20.75 | 47448 | 7 | 4412237 | 1282945.25 | 3303206.25 | 2933468.75 | 3265360 | 2698187.5 | 1302338.5 | 2 | 2658945 | 1178042.5 | 4513018 | 32371100 | 1324520 | 4744370 | 5851393 | 3170515.25 | 6346483 | 11645773 | 5269196 | 4028902 |
| Q7TPR4 | Actn1 | Alpha-actinin-1 | 611.02 | 17.15 | 103068 | 11 | 3943575 | 3686295.75 | 3464208.5 | 2745882.5 | 2673903 | 4004183 | 3343091 | 1429344.5 | 2060494 | 2297425 | 4206290.5 | 4138999 | 4806970 | 5031682 | 5318688 | 3874512.5 | 4417756.5 | 6480856 | 5887459 | 4943698.5 |
| P57780 | Actn4 | Alpha-actinin-4 | 968.27 | 23.25 | 104977 | 16 | 12227532 | 14896407 | 17742344 | 10474767 | 13127143 | 10580814 | 9155562 | 10228953 | 11889627 | 14012311 | 11984667 | 13590545 | 12303492 | 10247027 | 18104354 | 8576298 | 8141402 | 11213524 | 8454027 | 6705712.5 |
| P61164 | Actr1a | Alpha-centractin | 268.44 | 17.02 | 42614 | 5 | 770763.19 | 2 | 1839334.88 | 2 | 33094754 | 598727.25 | 1096417 | 480235 | 2 | 364868.78 | 2 | 2 | 2 | 2 | 2 | 2 | 1229802.88 | 403550.66 | 2 | 2 |
| Q8R5C5 | Actr1b | Beta-centractin | 283.82 | 13.03 | 42281 | 5 | 1050723 | 2179030.5 | 1479127.38 | 1036105.62 | 1085952 | 1009815.06 | 1047446.25 | 157646.72 | 1000024.25 | 904454 | 1042687.69 | 1526150 | 1053810.62 | 1029391.56 | 2 | 922352.62 | 666775.38 | 606604.31 | 916646 | 2 |
| P61161 | Actr2 | Actin-related protein 2 | 646.77 | 36.55 | 44761 | 13 | 14065531 | 7363834.5 | 35632610 | 23854460 | 20848406 | 19260138 | 30082664 | 50179394 | 19852936 | 33947086 | 20818046 | 16668703 | 19825948 | 15481399 | 38188254 | 6952711 | 12512247 | 33450624 | 6713018.5 | 9244954 |
| Q4V7C7 | Actr3 | Actin-related protein 3 | 1033.54 | 49.28 | 47357 | 20 | 41047474 | 36815662 | 53018906 | 42904142 | 60847126 | 39360230 | 43171826 | 56384350 | 45925358 | 61353738 | 65656870 | 62611442 | 54860170 | 55348958 | 49236886 | 43453878 | 33881946 | 32908444 | 41918226 | 27207962 |
| Q99JW2 | Acy1 | Aminoacylase-1 | 178.91 | 12.75 | 45781 | 3 | 2 | 2 | 2 | 2 | 551007.5 | 2 | 2 | 2 | 2 | 2 | 1637119.62 | 1302370 | 1642208.75 | 949585.38 | 2 | 2094602.12 | 2 | 970662.81 | 1319936.5 | 2 |
| O88839 | Adam15 | Disintegrin and metalloproteinase domain-contain | 60.96 | 1.39 | 92664 | 1 | 734685.81 | 1109270.75 | 2 | 720244.31 | 995501.12 | 615540.75 | 2 | 158888.95 | 748540.06 | 724582.25 | 869601.94 | 912422.06 | 924731.88 | 1347082.75 | 764735.44 | 2 | 2 | 1225998.25 | 1046562.12 | 2 |
| Q9Z0F8 | Adam17 | Disintegrin and metalloproteinase domain-contain | 38.48 | 1.57 | 93056 | 1 | 2 | 2 | 2 | 2 | 2 | 2 | 2 | 2 | 2 | 2 | 2 | 494984.03 | 514420 | 513041.91 | 2 | 2 | 2 | 2 | 2 | 2 |
| Q05910 | Adam8 | Disintegrin and metalloproteinase domain-contain | 281.92 | 10.77 | 90046 | 5 | 756145.75 | 422430.5 | 680153.19 | 2 | 2 | 2 | 2196302.75 | 271570.22 | 2278230.25 | 1075510.5 | 2 | 758826.44 | 927212.19 | 1010767.12 | 2 | 848237.12 | 1723943 | 2 | 1921231 | 278060.75 |
| Q61072 | Adam9 | Disintegrin and metalloproteinase domain-contain | 164.58 | 7.46 | 92079 | 4 | 1861648.25 | 2290644 | 1780622.75 | 1411145.25 | 873784.44 | 2 | 502634.91 | 120830.77 | 1226207.25 | 173120.5 | 108338.09 | 2 | 881054 | 1160993 | 741829 | 643923 | 429908.94 | 1185651.38 | 2 | 819297.75 |
| Q9D0L4 | Adck1 | AarF domain-containing protein kinase 1 | 36.85 | 1.71 | 59736 | 1 | 2 | 963811.12 | 5493717.5 | 1028087.5 | 1148851.5 | 759092.56 | 709312.5 | 1247999.75 | 756582.56 | 1146571.12 | 2914060.25 | 532902.88 | 820197.75 | 691931 | 973741.88 | 735491.88 | 6193713 | 868087.31 | 7631996.5 | 8054816 |
| P28474 | Adh5 | Alcohol dehydrogenase class-3 | 377.67 | 32.89 | 39548 | 8 | 4838420.5 | 4388677.5 | 4478782 | 4545309 | 3089322 | 3643802 | 3788089.25 | 4764370.5 | 3578659.5 | 3866651 | 2928195.5 | 5680606 | 6346668.5 | 5780611.5 | 8703673 | 3324662 | 7905537 | 6551630 | 4772998 | 5789241.5 |
| P55264 | Adk | Adenosine kinase | 42.87 | 2.49 | 40149 | 1 | 2 | 2 | 2 | 2 | 65286.82 | 172367.59 | 380458.03 | 151018.31 | 1771826.75 | 2 | 169077.12 | 1324095 | 2 | 2 | 2 | 2 | 2 | 2 | 2 | 2 |
| Q9JKL8 | Adnp | Activity-dependent neuroprotector homeobox prot | 62.28 | 2.09 | 123626 | 2 | 2 | 2 | 2 | 2 | 2 | 2 | 2 | 2 | 2 | 2 | 2 | 2 | 2 | 2 | 2 | 2 | 2 | 2 | 2 | 2 |
| Q6PDY2 | Ado | 2-aminoethanethiol dioxygenase | 119.6 | 9.38 | 28372 | 3 | 2296517.5 | 4440396 | 2562323.25 | 2529297.5 | 1060034.5 | 2177544.25 | 2139248.75 | 192391.39 | 2271212.75 | 1320264.5 | 3744230 | 3565408 | 4337371 | 4817470 | 895368.56 | 4379157 | 3597892.75 | 1754633 | 3941097 | 3831800 |
| Q8VDL4 | Adpgk | ADP-dependent glucokinase | 225.31 | 9.27 | 53902 | 3 | 4469715.5 | 4528430 | 3451894.5 | 4140623 | 4890851 | 4064973.5 | 3164527.25 | 4282185.5 | 3901249.25 | 3951973 | 2485890.25 | 2583490 | 3343616.25 | 4054239.75 | 2907139.75 | 3467294.5 | 3604531 | 4875579 | 1220416.12 | 4360444.5 |
| Q8CG72 | Adprs | ADP-ribosylhydrolase ARH3 | 37.46 | 2.43 | 39414 | 1 | 2 | 699564.44 | 2 | 936984.75 | 950379 | 2 | 2 | 1548522.12 | 937511.56 | 1197380.12 | 2 | 2 | 899828.5 | 2 | 2 | 2 | 2 | 2 | 2 | 578137.25 |
| Q9JKV1 | Adrm1 | Proteasomal ubiquitin receptor ADRM1 | 116.16 | 7.62 | 42060 | 2 | 887546.06 | 6818080 | 316492.59 | 3467398 | 400803.53 | 1439776.62 | 1942169 | 1244364 | 4517797.5 | 489109.91 | 2442255.75 | 5480902.5 | 2309017.75 | 4998656.5 | 4375447 | 2907012.25 | 4012724.75 | 1985470.62 | 2 | 3801738.25 |
| P54822 | Adsl | Adenylosuccinate lyase | 198.3 | 8.88 | 54866 | 4 | 1115374 | 2091315.5 | 1606408.5 | 3878650.25 | 8045398 | 5801379 | 4653062 | 7922854 | 7258304 | 2580748.5 | 5544083 | 6864935.5 | 2073424.75 | 1497188.62 | 1530340.75 | 5538483.5 | 1091927.88 | 2 | 1954921.5 | 1798978 |
| P28650 | Adss1 | Adenylosuccinate synthetase isozyme 1 | 57.1 | 3.06 | 50254 | 1 | 2 | 129742.62 | 2 | 2 | 696773.38 | 2 | 632175.25 | 98938.09 | 2 | 975611.75 | 244390.44 | 1039360.19 | 730880.75 | 703297.56 | 565399.81 | 339909.19 | 933117 | 298355.16 | 2 | 263731.34 |
| P46664 | Adss2 | Adenylosuccinate synthetase isozyme 2 | 420.25 | 21.71 | 50021 | 8 | 15693196 | 6199466.5 | 11826991 | 11162127 | 7400006.5 | 9440047 | 6046982.5 | 10644631 | 13465497 | 9034901 | 3933560 | 6725660 | 4199166.5 | 7290295.5 | 14703776 | 6190972.5 | 14061264 | 6242935 | 13274615 | 11640328 |
| Q920A7 | Afg3l1 | Mitochondrial inner membrane m-AAA protease | 321.24 | 10.65 | 87047 | 7 | 760866.19 | 1823952.38 | 2623305 | 4445868 | 3443800 | 2765147 | 3175086.75 | 2258758.5 | 3630095.5 | 4180383.75 | 1542988.5 | 812021.06 | 4020275.5 | 2973418 | 2235014.5 | 3635827.5 | 3309711 | 3865845.75 | 3569742.5 | 3679443.75 |
| Q8JZQ2 | Afg3l2 | Mitochondrial inner membrane m-AAA protease | 324.67 | 9.73 | 89519 | 7 | 6055052 | 6920704 | 5515274 | 2532091.5 | 3973665.75 | 7767280 | 7863292 | 6718208 | 9536636 | 6949773.5 | 4596736 | 3443136 | 3800383.5 | 4462960.5 | 4225370.5 | 7364634.5 | 4678547.5 | 4976635 | 5712268.5 | 3025861.5 |
| Q64191 | Aga | N(4)-(beta-N-acetylglucosaminyl)-L-asparaginase | 196.33 | 13.58 | 37022 | 4 | 5603283 | 7575216 | 2231769.5 | 4589697 | 1265539.62 | 4752937 | 4774203 | 2059993.75 | 4798416 | 5967330 | 3011998.75 | 2214186 | 2631005 | 2199155.5 | 2492699 | 2760377.25 | 4987954.5 | 4965557 | 3895319 | 4999917 |
| Q9ESW4 | Agk | Acylglycerol kinase, mitochondrial | 49.76 | 3.33 | 46976 | 1 | 1116416.75 | 1494834.62 | 2 | 2 | 1213669.5 | 2 | 872474.88 | 1030942.62 | 895007.81 | 2 | 2 | 828873.75 | 761793.94 | 649206.62 | 2 | 2 | 2 | 2 | 925553.12 | 2 |
| Q8C0I1 | Agps | Alkyldihydroxyacetonephosphate synthase, perox | 567.62 | 20.16 | 71684 | 11 | 6333586 | 12840378 | 4893082.5 | 8723466 | 8498652 | 6495956.5 | 5325704 | 7814936.5 | 6569041 | 3946182 | 6330360.5 | 5387757.5 | 6393187.5 | 7333592 | 3315849.5 | 6473814.5 | 7081093.5 | 3515731.25 | 8115786 | 5688232.5 |
| P10760 | Ahcy | Adenosylhomocysteinase | 867.81 | 33.8 | 47538 | 15 | 91273714 | 124313010 | 58233554 | 78015018 | 48621862 | 72116410 | 70969882 | 64060210 | 56488526 | 45720814 | 64944106 | 74308162 | 62162490 | 57795622 | 58655914 | 56626154 | 64818762 | 70728146 | 67711098 | 73039234 |
| B5DFN2 | Ahcyl1 | S-adenosylhomocysteine hydrolase-like protein 1 | 279.3 | 12.42 | 53753 | 6 | 3427070 | 5728816 | 3865222 | 4228606.5 | 4206938 | 3360969 | 2228797.5 | 902458.88 | 3149210.75 | 2572746.75 | 4335644.5 | 2856275.75 | 3407000 | 3273964 | 2259869.25 | 2 | 3519145.5 | 2339347.75 | 4373719 | 2 |
| Q68FL4 | Ahcyl2 | Putative adenosylhomocysteinase 3 | 226.49 | 7.01 | 66899 | 5 | 460728.03 | 833272.5 | 2 | 2 | 269379.44 | 1667447.12 | 1700869.88 | 1796665.75 | 1535112.62 | 2 | 2229230.25 | 1802219.88 | 1796124 | 2060844.75 | 2 | 2030279.25 | 1341974.62 | 1047827.25 | 1664445 | 2 |
| Q8BK64 | Ahsa1 | Activator of 90 kDa heat shock protein ATPase hc | 103.71 | 7.1 | 38117 | 2 | 2098269.5 | 3661412.5 | 2899471.75 | 4391072.5 | 3526008.75 | 4583782.5 | 3879925 | 1454635.25 | 4062477.5 | 458614.44 | 6156676 | 6984405 | 7003834.5 | 7022895 | 5911192 | 8303389.5 | 9815597 | 11243295 | 7981015 | 10530410 |

| | | | | | | | | | | | | | | | | | | | | | | | | | | |
|---|---|---|---|---|---|---|---|---|---|---|---|---|---|---|---|---|---|---|---|---|---|---|---|---|---|---|
| Q9Z0X1 | Aifm1 | Apoptosis-inducing factor 1, mitochondrial | 773.7 | 29.58 | 66765 | 14 | 3788268.75 | 2802908.5 | 3033380.5 | 2710384.75 | 3952886.25 | 5942557 | 5319947.5 | 4914535 | 11501732 | 3378811.5 | 14513321 | 14349574 | 8970858 | 10582814 | 9016713 | 11157832 | 5260577.5 | 7356709.5 | 9470906 | 7630630 |
| P31230 | Aimp1 | Aminoacyl tRNA synthase complex-interacting mu | 152.75 | 12.26 | 33997 | 3 | 2 | 2 | 2 | 1287731.38 | 2 | 2 | 2 | 2 | 1386381.25 | 2 | 3275002.5 | 3190385 | 1069037.62 | 1751021.62 | 2 | 1911813.62 | 697621.31 | 2 | 2992088.5 | 2 |
| Q32PX2 | Aimp2 | Aminoacyl tRNA synthase complex-interacting mu | 48.95 | 2.81 | 35443 | 1 | 2 | 42311.3 | 109660.81 | 148876.75 | 72458.54 | 2 | 184229.22 | 2 | 104898.42 | 143319.69 | 2 | 84452.53 | 90802.02 | 2 | 757481.69 | 2 | 107147.17 | 2 | 2 | 128825.36 |
| O08915 | Aip | AH receptor-interacting protein | 59.58 | 4.24 | 37605 | 1 | 2 | 2 | 2 | 2 | 2009211.25 | 394908.28 | 934558.62 | 581722 | 1542849.5 | 2 | 2 | 2 | 2 | 2 | 2 | 2 | 2 | 2 | 2 | 394744.97 |
| Q9WTP6 | Ak2 | Adenylate kinase 2, mitochondrial | 301.22 | 23.85 | 26469 | 6 | 8026213.5 | 20289658 | 5081166.5 | 13721287 | 12178465 | 19437234 | 16732800 | 16228528 | 21647006 | 12905166 | 19028202 | 20311212 | 15597031 | 17610926 | 11706063 | 18504474 | 14629627 | 11558658 | 13325951 | 15661244 |
| Q9WTP7 | Ak3 | GTP:AMP phosphotransferase AK3, mitochondria | 98.74 | 18.5 | 25426 | 2 | 521844.66 | 613491.19 | 297366.12 | 1226404.38 | 968161.75 | 2 | 150859.41 | 1372263.5 | 885752.75 | 1542871.75 | 779254.56 | 2 | 227683.58 | 179257.98 | 2 | 2 | 2 | 2 | 2 | 2 |
| O88845 | Akap10 | A-kinase anchor protein 10, mitochondrial | 48.53 | 1.36 | 73632 | 1 | 2 | 694748.75 | 547950.69 | 2 | 1012751.56 | 2 | 2 | 2 | 2 | 2 | 873755.62 | 717045.38 | 516837.62 | 2 | 2 | 2 | 759364.12 | 2 | 2 | 2 |
| Q9DBR0 | Akap8 | A-kinase anchor protein 8 | 43.88 | 1.89 | 76294 | 1 | 2 | 2 | 2 | 636885.25 | 2 | 2 | 588018.56 | 63897.02 | 2 | 722966.81 | 2 | 707365.5 | 669790.62 | 642966 | 434615.09 | 725093.94 | 2 | 2 | 2 | 2 |
| Q9JII6 | Akr1a1 | Aldo-keto reductase family 1 member A1 | 411.4 | 24.31 | 36587 | 9 | 7119431 | 9210204 | 17224076 | 7946716.5 | 7848720 | 8816801 | 11161597 | 15344642 | 6971672.5 | 11833245 | 13086845 | 13214460 | 11213242 | 11956523 | 17647110 | 8892900 | 11881400 | 14419430 | 11972482 | 10729342 |
| P45376 | Akr1b1 | Aldo-keto reductase family 1 member B1 | 583.58 | 42.72 | 35732 | 13 | 36027502 | 22539574 | 35663510 | 17784270 | 41214042 | 21286462 | 17430054 | 21788014 | 31817478 | 34607562 | 66405622 | 67040726 | 50373734 | 44175058 | 46622906 | 53113482 | 36860802 | 37624078 | 50000858 | 35500738 |
| P45377 | Akr1b8 | Aldose reductase-related protein 2 | 599.28 | 32.28 | 36121 | 11 | 1129529 | 3535255.25 | 477839.09 | 1352539.5 | 1671802.38 | 3456705.5 | 3381970.75 | 4171152.25 | 8702617 | 1112073.75 | 13093079 | 14349886 | 3208626.75 | 6125177 | 1847781.5 | 13690011 | 3773392.5 | 3245477.5 | 5282296 | 4849171 |
| Q8CG76 | Akr7a2 | Aflatoxin B1 aldehyde reductase member 2 | 180.37 | 12.81 | 40612 | 4 | 2 | 1342043.38 | 2 | 1417955.75 | 1532417 | 2 | 1836531.88 | 841564.5 | 1245708.5 | 1942688 | 2633762.5 | 2638105.75 | 2029602.25 | 2469918.25 | 1491462.5 | 1890017.25 | 615620.94 | 1384285.88 | 1285391.62 | 1171690.88 |
| P10518 | Alad | Delta-aminolevulinic acid dehydratase | 174.29 | 16.06 | 36024 | 4 | 1811783.75 | 837386.25 | 1961853 | 971904.31 | 1329899.75 | 2042950.75 | 1581957.25 | 3328596.5 | 1378840.62 | 1121115.88 | 2469096.5 | 1003828.62 | 1032746.5 | 1610162.62 | 1670802 | 865243.88 | 557634.62 | 1191387 | 2575795.75 | 2303444 |
| Q61490 | Alcam | CD166 antigen | 479.53 | 22.64 | 65092 | 9 | 2979135.5 | 3919226.25 | 4262604.5 | 3312755 | 3876958.5 | 2985769 | 4523581 | 6990898 | 5099057.5 | 5204156 | 4327671 | 8769341 | 6444693 | 9102001 | 1734912.38 | 6839106 | 3251541 | 5608044.5 | 6477749.5 | 3258465.75 |
| Q3T1L0 | Aldh16a1 | Aldehyde dehydrogenase family 16 member A1 | 137.48 | 3.24 | 85414 | 2 | 2 | 2 | 2 | 2 | 717292.12 | 2 | 82538.82 | 2 | 867008.38 | 202610.25 | 631036.12 | 2 | 2 | 2 | 518911.06 | 2 | 2 | 2 | 2 | 2 |
| Q9Z110 | Aldh18a1 | Delta-1-pyrroline-5-carboxylate synthase | 937.92 | 22.64 | 87266 | 18 | 2298667.75 | 6509775 | 3835623.75 | 5044323 | 8296683 | 8729280 | 8873910 | 9654278 | 12390760 | 9830626 | 8831375 | 5642840 | 6711214 | 8842139 | 7579126.5 | 9527506 | 2052646.75 | 4290168 | 5805551 | 3643720.75 |
| P28037 | Aldh1l1 | Cytosolic 10-formyltetrahydrofolate dehydrogenas | 40.26 | 1.66 | 98874 | 1 | 2 | 2 | 2 | 2 | 993860.06 | 2 | 2 | 2 | 2 | 2 | 2 | 2 | 757865.38 | 2 | 2 | 2 | 2 | 2 | 160433.22 | 2 |
| P47738 | Aldh2 | Aldehyde dehydrogenase, mitochondrial | 1236.71 | 43.74 | 56538 | 23 | 41479202 | 39487362 | 62696750 | 39923990 | 51069782 | 39791474 | 34318890 | 38395702 | 43847130 | 61077410 | 58440954 | 52779854 | 61895882 | 56116566 | 65195586 | 49744534 | 47677718 | 50848014 | 52367658 | 40439558 |
| P47740 | Aldh3a2 | Aldehyde dehydrogenase family 3 member A2 | 129.77 | 7.85 | 53971 | 3 | 470886.78 | 577645.44 | 555257.5 | 2 | 2 | 977919.56 | 2 | 891817.69 | 573602.75 | 1157076.88 | 806568.12 | 647970.25 | 2 | 506964.03 | 2 | 2 | 2 | 2 | 2 | 2 |
| Q5XI42 | Aldh3b1 | Aldehyde dehydrogenase family 3 member B1 | 137.56 | 7.05 | 52147 | 3 | 2143923.25 | 1562861.38 | 3865812.5 | 756363.5 | 1972246 | 1493808.62 | 1677865.62 | 2276118.5 | 2471453 | 2513287.25 | 1259492.5 | 2086215.5 | 2125746.25 | 2080103 | 1959583.88 | 1123901.12 | 1864509.25 | 1177948.62 | 1917906.5 | 2 |
| Q9JLJ2 | Aldh9a1 | 4-trimethylaminobutyraldehyde dehydrogenase | 945.51 | 39.88 | 53515 | 16 | 3612795.75 | 4110103.5 | 6632155 | 3853619 | 13794334 | 4164881.5 | 5256696 | 10912960 | 11763159 | 7482808.5 | 11214686 | 9975535 | 10785041 | 9284123 | 6174317 | 2882112.25 | 5250862 | 5049568.5 | 7159745 | 5587589.5 |
| P05064 | Aldoa | Fructose-bisphosphate aldolase A | 2239.29 | 87.36 | 39356 | 36 | 29702812 | 51170450 | 24810022 | 23877682 | 39785570 | 23068214 | 19911356 | 19880840 | 6046355 | 31403190 | 18647542 | 24111222 | 17180686 | 17314482 | 22092306 | 52906678 | 25868152 | 46411154 | 53371602 | 53132102 |
| P05063 | Aldoc | Fructose-bisphosphate aldolase C | 1252.38 | 61.16 | 39395 | 21 | 13274122 | 21123936 | 14549677 | 11569550 | 24268408 | 22008330 | 24583246 | 24554790 | 23588642 | 18820232 | 38415966 | 41127062 | 34667174 | 38105294 | 21977542 | 29246642 | 36152190 | 26580400 | 31875758 | 25395772 |
| Q9DB25 | Alg5 | Dolichyl-phosphate beta-glucosyltransferase | 64.73 | 6.17 | 36791 | 2 | 2 | 1236291.5 | 1012239.12 | 945478.94 | 2 | 1992717.5 | 1135657.25 | 2 | 2065426.62 | 976652.44 | 1569033.38 | 635094.31 | 974172.5 | 933819.56 | 2 | 1079575.62 | 2 | 1629877 | 1499981.25 | 1444380 |
| P12527 | Alox5 | Polyunsaturated fatty acid 5-lipoxygenase | 241.01 | 5.2 | 78087 | 4 | 1699855 | 1432413 | 2061773.12 | 5336951.5 | 3887440.5 | 276313.5 | 1406784.75 | 398087.81 | 1977974.75 | 1111439.88 | 1165149 | 484581.97 | 2188247.5 | 1599144.62 | 2835594.5 | 166112.67 | 875472 | 2325642.5 | 1407697 | 2657502.5 |
| P20291 | Alox5ap | Arachidonate 5-lipoxygenase-activating protein | 177.28 | 18.63 | 18070 | 4 | 4542682 | 4309182 | 3095317 | 5150982 | 8571666 | 11633614 | 5258592 | 17239840 | 17905152 | 7383497.5 | 7620363 | 2876846.5 | 3145408 | 2885443.75 | 3378766.25 | 7705565 | 5270311.5 | 7180145.5 | 4226630 | 6673684.5 |
| Q5BJY6 | Amdhd2 | N-acetylglucosamine-6-phosphate deacetylase | 68.67 | 2.93 | 43539 | 1 | 490730.38 | 75753.05 | 777316.56 | 2 | 196652.94 | 2 | 2 | 2 | 465310.75 | 414555.03 | 328492.56 | 366856.91 | 1121562.25 | 503024.62 | 859137.06 | 279892.53 | 550769.81 | 371023.97 | 208456.95 | 329323.12 |
| O08739 | Ampd3 | AMP deaminase 3 | 44.69 | 0.91 | 88652 | 1 | 2 | 2 | 2 | 2 | 2 | 2 | 2 | 2 | 980511.69 | 2 | 476901.34 | 755235.06 | 891106.88 | 826477.5 | 2 | 2 | 1566766.25 | 1428571.75 | 2 | 2 |
| Q810B6 | Ankfy1 | Rabankyrin-5 | 57.88 | 0.77 | 128653 | 1 | 2355393.5 | 1792635.75 | 2053253 | 2208419.25 | 1463715.62 | 2060796.25 | 1647325.12 | 1949957.88 | 1105956.62 | 993583.19 | 1134123.75 | 1075707.25 | 1349676.75 | 1102966.38 | 1412295.62 | 908798.31 | 1413496.5 | 2187236.5 | 1058798.88 | 1838806.25 |
| Q9EST5 | Anp32b | Acidic leucine-rich nuclear phosphoprotein 32 fan | 136.23 | 9.93 | 31079 | 3 | 2 | 2 | 2 | 708868.62 | 635684.25 | 245871.48 | 552093.19 | 572314.81 | 646190.06 | 581817.94 | 312149.41 | 323141.19 | 2 | 2 | 2 | 2 | 944254.5 | 2 | 2 | 2 |
| P97822 | Anp32e | Acidic leucine-rich nuclear phosphoprotein 32 fan | 53.98 | 5.38 | 29622 | 1 | 1555235.5 | 2 | 899911.19 | 979446 | 922703.06 | 932280.75 | 2 | 702264.69 | 517301.75 | 865650 | 2 | 2 | 2 | 2 | 2 | 2 | 2 | 2 | 2 | 2 |
| P97449 | Anpep | Aminopeptidase N | 257.49 | 4.66 | 109651 | 5 | 1114928.62 | 1055174 | 229718.31 | 2 | 2 | 672719.88 | 217784.62 | 427657.06 | 282716 | 244936.14 | 950020 | 1813020 | 1047923.25 | 88379 | 2 | 2685281.25 | 1572208.88 | 1258778.75 | 68199.94 | 2 |
| Q501J2 | Antkmt | Adenine nucleotide translocase lysine N-methyltra | 92.53 | 9.61 | 24740 | 2 | 2 | 2 | 2 | 2 | 2 | 2 | 2 | 2 | 2 | 257188.22 | 2 | 2 | 2 | 2 | 2 | 2 | 2 | 2 | 2 | 2 |
| P10107 | Anxa1 | Annexin A1 | 2096.89 | 77.75 | 38734 | 33 | 68152450 | 53194074 | 68094202 | 57864230 | 62660078 | 46651882 | 62137354 | 58418646 | 53152146 | 66886970 | 58387310 | 56864686 | 52579758 | 47998998 | 55097390 | 34424622 | 48305362 | 40843458 | 30044642 | 39354562 |
| P97384 | Anxa11 | Annexin A11 | 278.48 | 14.91 | 54079 | 6 | 3517906.25 | 2470020 | 1342807 | 3165342.5 | 2914366.5 | 942905.88 | 1831516.75 | 1711469 | 1272627.5 | 2304000 | 1309748.38 | 2282794.75 | 2235426 | 1604209.12 | 1148566 | 977605.56 | 1810177.5 | 1439584.5 | 1636939.12 | 1537745.25 |
| P07356 | Anxa2 | Annexin A2 | 1537.91 | 62.83 | 38676 | 24 | 142672770 | 112233330 | 194697618 | 70090050 | 106000962 | 85119722 | 100316978 | 101476650 | 82270466 | 187931426 | 146838130 | 152259634 | 145981234 | 130458386 | 163193666 | 120963106 | 157252834 | 150507298 | 135496674 | 126076122 |
| O35639 | Anxa3 | Annexin A3 | 969.06 | 45.82 | 36384 | 16 | 27259306 | 26523176 | 25036304 | 23828364 | 23017720 | 31858110 | 25662670 | 30153740 | 23101416 | 27227932 | 41935034 | 39311054 | 30589982 | 29669408 | 25001920 | 37454066 | 32210188 | 33978170 | 38195906 | 26062976 |
| P97429 | Anxa4 | Annexin A4 | 1547.45 | 56.74 | 35916 | 19 | 116795306 | 161584834 | 96054538 | 75794178 | 54846330 | 131658482 | 126236218 | 129608298 | 81397642 | 88745226 | 137044930 | 155682770 | 165856370 | 138351458 | 137834274 | 146368386 | 188628162 | 179039010 | 208931874 | 145825282 |
| P48036 | Anxa5 | Annexin A5 | 1841.31 | 77.43 | 35752 | 27 | 74433442 | 65635002 | 83920434 | 60786442 | 38640962 | 100408762 | 97151242 | 65095842 | 60455482 | 108914722 | 97273666 | 98034242 | 106528226 | 93018850 | 89997202 | 55797974 | 103943450 | 94801330 | 85441970 | 92453482 |
| P14824 | Anxa6 | Annexin A6 | 607.14 | 20.06 | 75885 | 13 | 12243927 | 5973918.5 | 19104462 | 3273527 | 17450170 | 2381533.5 | 3858216.25 | 6297915 | 5803733.5 | 13511268 | 16981508 | 17358620 | 20450712 | 15774086 | 27271420 | 23973778 | 20022402 | 32962934 | 19579356 | 18819134 |
| Q07076 | Anxa7 | Annexin A7 | 543.49 | 19.65 | 49925 | 9 | 8731718 | 10115022 | 8965135 | 7079648 | 6936030 | 5356104.5 | 6052643 | 7622528 | 11759929 | 6648532.5 | 14224084 | 16506326 | 12418379 | 12174210 | 9414593 | 13054605 | 12110024 | 12165517 | 9233605 | 13062747 |
| O35643 | Ap1b1 | AP-1 complex subunit beta-1 | 588.49 | 14.1 | 103935 | 10 | 3084675 | 3088713.25 | 2248709 | 5270204 | 4313089 | 7187583 | 5247875 | 6968444 | 6767137.5 | 4719072.5 | 5888561 | 5311995 | 5200669.5 | 4308224 | 3157457.75 | 5818119 | 5099759.5 | 5622210 | 4680265 | 4369734.5 |
| P22892 | Ap1g1 | AP-1 complex subunit gamma-1 | 208.17 | 6.81 | 91350 | 4 | 628317.62 | 126829.19 | 462788.62 | 531519.56 | 384981.28 | 456587 | 200023.8 | 84202.86 | 777965.81 | 505399.94 | 2 | 2 | 231853.38 | 2 | 1321166.12 | 501453.78 | 98859.71 | 2 | 129965.77 | 2 |
| P35585 | Ap1m1 | AP-1 complex subunit mu-1 | 175.62 | 8.98 | 48543 | 3 | 2952893 | 4074698 | 1856955.62 | 3381051.5 | 3914457 | 5203302 | 2577642.25 | 3970970.25 | 3672984 | 2982411.5 | 4716970 | 3329461.75 | 4737647 | 4393224 | 2683972.75 | 4200117 | 2853713.25 | 3975690.5 | 4815235.5 | 4271534 |
| Q7TN05 | Ap1s3 | AP-1 complex subunit sigma-3 | 36.24 | 5.84 | 18440 | 1 | 2 | 2 | 2 | 123981.8 | 2 | 2 | 502741.94 | 2 | 2 | 658105 | 2 | 2 | 2 | 2 | 2 | 2 | 2 | 2 | 2 | 2 |
| P17426 | Ap2a1 | AP-2 complex subunit alpha-1 | 747.32 | 19.96 | 107664 | 17 | 1874738.75 | 3138201.5 | 1482878.5 | 1485309.88 | 2664180.25 | 3549761.5 | 4617316 | 4699407 | 3368072 | 1837396.5 | 3379391.25 | 1961350 | 3207430.5 | 2260954.25 | 2498707.5 | 2522117.5 | 3746110 | 4523029.5 | 4109693.5 | 3924743.5 |
| P62944 | Ap2b1 | AP-2 complex subunit beta | 466.89 | 10.99 | 104553 | 9 | 2411310.25 | 2033997.75 | 1746355.12 | 3509958.5 | 4102000 | 2337627.25 | 2286224.75 | 4070818.25 | 2912285 | 3302223 | 1602881.5 | 1542106.75 | 1089845.88 | 1233824.88 | 2 | 1537738.25 | 2 | 2 | 1063423.5 | 2 |
| P84091 | Ap2m1 | AP-2 complex subunit mu | 313.35 | 22.3 | 49655 | 8 | 696861.56 | 1612514.5 | 2142086.75 | 1601551.5 | 1967952 | 1174223.62 | 2087901.5 | 733981.69 | 6226968 | 2398313.25 | 4430581 | 3453156.75 | 1252507.38 | 3953238 | 612530.06 | 2898633.5 | 3607123 | 1006597.5 | 1647875.12 | 366348.38 |
| Q9Z1T1 | Ap3b1 | AP-3 complex subunit beta-1 | 178.56 | 3.98 | 122740 | 4 | 3847252.75 | 3624042.5 | 3486212.5 | 2933159.5 | 2237082 | 3118078.5 | 3562562.25 | 4822365 | 3132574 | 3052898.5 | 2033257.25 | 1209749.25 | 3398552.75 | 3918246.5 | 1800161 | 3405127 | 4468898 | 6299555 | 3275009.5 | 1964261.5 |
| Q9DCR2 | Ap3s1 | AP-3 complex subunit sigma-1 | 46.62 | 5.7 | 21732 | 1 | 2 | 2 | 2 | 2 | 2 | 2 | 2 | 2 | 760315.88 | 2 | 949730.12 | 1174466.25 | 1064903.62 | 1068920 | 779012.56 | 1358409.62 | 1371300.75 | 1277988.12 | 883522.12 | 1549241.25 |
| Q8R146 | Apeh | Acylamino-acid-releasing enzyme | 37.74 | 1.09 | 81581 | 1 | 4977963 | 5092038.5 | 5290853.5 | 1476731.75 | 2520435.5 | 1775069.88 | 1294977.12 | 2529241 | 1609181.12 | 2424875.5 | 1016858.25 | 1153205 | 827822.19 | 1046949.12 | 631601.19 | 1200745.75 | 2283224.75 | 991735.19 | 2503434.75 | 2 |
| O35841 | Api5 | Apoptosis inhibitor 5 | 315.68 | 12.3 | 56785 | 6 | 6659129 | 7436401.5 | 7524829 | 7021067 | 9185429 | 7465720.5 | 5749701 | 9631847 | 6636399.5 | 5840428 | 7585851 | 6494775 | 5612265.5 | 5749233 | 5126585 | 6101737 | 5430676 | 6879302.5 | 5930869 | 4225924.5 |
| Q9D7N9 | Apmap | Adipocyte plasma membrane-associated protein | 405.24 | 19.04 | 46434 | 7 | 2 | 82303.55 | 166378.55 | 331876.44 | 587064.88 | 88847.07 | 287085.53 | 2 | 324844.09 | 201345.48 | 2 | 91287.54 | 377837.56 | 1124326.25 | 804681.5 | 2 | 476109.5 | 287401.62 | 90027.2 | 179321.61 |
| Q8VBT6 | Apobr | Apolipoprotein B receptor | 350.48 | 9.02 | 102705 | 7 | 1926176.62 | 3188272.75 | 641726.31 | 3315275 | 2390578 | 829113.81 | 1897017.25 | 1008668.75 | 3117714 | 2703022.75 | 6949761 | 4067970.5 | 4110096.5 | 3676137.25 | 1625616 | 1386489.38 | 2365862 | 710166.06 | 3030457.5 | 2 |
| Q78IK4 | Apool | MICOS complex subunit Mic27 | 240.83 | 20.75 | 29261 | 4 | 1897305.62 | 1302169.75 | 1464236 | 1635550.5 | 1499614.75 | 2 | 1562425.38 | 872966.44 | 2682531 | 2965031 | 1129232.75 | 1017179 | 845855.31 | 1628721 | 1443955.38 | 1122463.5 | 915354.19 | 2 | 1177567.12 | 2 |
| P08592 | App | Amyloid-beta precursor protein | 35.25 | 1.04 | 86704 | 1 | 2 | 660033.44 | 2 | 2 | 2 | 2 | 2 | 2 | 2 | 970340.69 | 2 | 2 | 2 | 664762.88 | 485738.22 | 2 | 553143.44 | 2 | 2 | 2 |
| B4F779 | Appl2 | DCC-interacting protein 13-beta | 45.45 | 1.06 | 74117 | 1 | 2 | 2 | 524417.81 | 326409.72 | 548741.94 | 2 | 1098701.25 | 1739055.12 | 1056911.12 | 802328.69 | 2 | 2 | 453647.31 | 371785.81 | 2 | 2 | 2 | 2 | 715693.81 | 2 |
| P08030 | Aprt | Adenine phosphoribosyltransferase | 798.36 | 78.89 | 19724 | 13 | 2989605.75 | 5781460 | 7919466.5 | 9042016 | 9310141 | 13760296 | 12926512 | 12873876 | 7951732.5 | 11989038 | 11692195 | 10646377 | 12062066 | 8760499 | 25475784 | 9417290 | 9771110 | 12319301 | 7725669.5 | 10235680 |
| Q4LDD4 | Arap1 | Arf-GAP with Rho-GAP domain, ANK repeat and I | 151.04 | 2.34 | 162276 | 3 | 2 | 2 | 2 | 2 | 1190503.62 | 860328.5 | 1887876.25 | 1992832 | 1778480.25 | 560440.06 | 860140.56 | 2 | 2 | 2 | 2 | 2 | 2 | 2 | 2 | 2 |
| Q8R5G7 | Arap3 | Arf-GAP with Rho-GAP domain, ANK repeat and I | 37.24 | 0.46 | 169741 | 1 | 1470462.88 | 1943104.25 | 981210.88 | 1248720 | 585809.06 | 746804.75 | 1108202.38 | 2 | 431217.88 | 755556 | 2 | 2 | 2 | 1681334.88 | 1188288.25 | 1596068.5 | 1385144.25 | 1395255.38 | 2 | 854702.88 |
| Q5XJY5 | Arcn1 | Coatomer subunit delta | 531.35 | 19.37 | 57229 | 11 | 2477503 | 1850133.75 | 3494991.5 | 3059532 | 4569527 | 4095419 | 4870941 | 2931953.75 | 5621101 | 3059620.5 | 4136920.5 | 5730033.5 | 3820627 | 6494790 | 5040120.5 | 5253984 | 2856162.75 | 905628 | 4523129 | 1387585.38 |
| P61205 | Arf3 | ADP-ribosylation factor 3 | 479.18 | 45.3 | 20601 | 8 | 1970583.88 | 3455025.75 | 8473084 | 3215017.5 | 5992981.5 | 8790840 | 8074858 | 11519033 | 3731770.25 | 6259883 | 12392635 | 6932315 | 7370494 | 4411302.5 | 9695548 | 4218229.5 | 7888651 | 6809679.5 | 5889520 | 7617100 |
| P61750 | Arf4 | ADP-ribosylation factor 4 | 387.87 | 31.67 | 20397 | 6 | 4864725.5 | 4454073 | 4612127 | 4118566.5 | 4334318.5 | 4858068 | 4306347.5 | 5061410 | 3212091.5 | 2721519.75 | 5215312 | 3419188.75 | 4122268.5 | 4210737 | 4776318 | 4263689.5 | 3822842 | 4886203.5 | 4634482 | 4035415.25 |
| P84083 | Arf5 | ADP-ribosylation factor 5 | 398.46 | 33.33 | 20530 | 6 | 591706.56 | 735106.5 | 2 | 676942.31 | 2 | 2 | 714756.19 | 2 | 732713.75 | 1488876.38 | 1200409 | 1184836.75 | 995314.81 | 910760 | 1287329.62 | 802542.56 | 1214126.75 | 895252.62 | 487480 | 1029457.81 |
| P62331 | Arf6 | ADP-ribosylation factor 6 | 222.55 | 32 | 20082 | 5 | 4093128 | 3555610 | 1232045.62 | 2815701.25 | 2284326.5 | 2915547.25 | 3160483.5 | 4114031 | 1184722 | 2846289.5 | 2 | 2426014.75 | 2807746 | 2874011.5 | 5138942 | 2692025.75 | 3211273.25 | 4360217 | 1151302.5 | 3984406 |
| A2A5R2 | Arfgef2 | Brefeldin A-inhibited guanine nucleotide-exchang | 193.33 | 2.85 | 202240 | 4 | 2015456.25 | 1226551.88 | 2048754.25 | 1024803.88 | 1280350.25 | 1358076.12 | 1153857.88 | 67945.49 | 724778.44 | 2 | 2 | 2 | 982825 | 753398.56 | 279739.66 | 2 | 800008.19 | 1149364.62 | 2 | 1405308.75 |
| Q3UIA2 | Arhgap17 | Rho GTPase-activating protein 17 | 45.23 | 1.06 | 92202 | 1 | 1606328.12 | 1761663.62 | 705776.38 | 902074.44 | 840755.69 | 1164809.38 | 2 | 784261.38 | 513321.28 | 463219.31 | 568867.38 | 540131.12 | 461600.47 | 502356.47 | 484372.94 | 2 | 2 | 451081.47 | 2 | 2 |
| Q8BYW1 | Arhgap25 | Rho GTPase-activating protein 25 | 186.78 | 6.94 | 73383 | 4 | 2862671.5 | 3565716.5 | 1568283 | 5475207.5 | 3601619 | 4282189.5 | 4957916.5 | 4962829 | 5305033.5 | 2689926.25 | 2782466.25 | 3920788.75 | 2794025.5 | 3534954 | 972759.38 | 2267012.5 | 3549324.5 | 2269530.25 | 2498251 | 4012388.5 |
| A2AB59 | Arhgap27 | Rho GTPase-activating protein 27 | 50.33 | 1.15 | 97049 | 1 | 742253.81 | 1028700.62 | 2 | 1137055.38 | 1503044.12 | 2 | 2 | 2 | 981700.75 | 774268.44 | 2 | 2 | 2 | 2 | 575571.5 | 2 | 2 | 2 | 2 | 2 |
| Q640N3 | Arhgap30 | Rho GTPase-activating protein 30 | 140.95 | 2.82 | 120114 | 3 | 2 | 2 | 2 | 2 | 2 | 2 | 2 | 2 | 2 | 2 | 426112.12 | 2 | 2 | 2 | 2 | 2 | 2 | 2 | 2 | 2 |
| Q5XI73 | Arhgdia | Rho GDP-dissociation inhibitor 1 | 503.32 | 49.02 | 23407 | 8 | 20611972 | 48225806 | 42974394 | 42857042 | 79546434 | 48252582 | 51338314 | 78882826 | 50282574 | 72779930 | 94013306 | 85219514 | 67931658 | 64423098 | 50806754 | 73836290 | 53238970 | 39812978 | 79411490 | 39051794 |
| Q61599 | Arhgdib | Rho GDP-dissociation inhibitor 2 | 315.88 | 41.5 | 22851 | 6 | 12085706 | 8637191 | 14381859 | 10979852 | 17308962 | 9747859 | 11646135 | 16862480 | 13793395 | 9501614 | 21818246 | 27089976 | 24190888 | 20165530 | 19982184 | 14207953 | 21622128 | 14310694 | 25044748 | 13554568 |
| Q61210 | Arhgef1 | Rho guanine nucleotide exchange factor 1 | 208.41 | 7.5 | 102805 | 4 | 892280 | 564450.19 | 113928.69 | 677823.38 | 2238951.5 | 141678.97 | 2 | 2 | 691670.25 | 714957.94 | 2567698.5 | 1807689.12 | 962685.25 | 1459579 | 1104396.5 | 2 | 1468746.12 | 1354684 | 2 | 2 |
| Q60875 | Arhgef2 | Rho guanine nucleotide exchange factor 2 | 637.51 | 16.24 | 111974 | 14 | 1510922.88 | 2733868.5 | 2547543.5 | 2325896.75 | 3608813.5 | 4840334.5 | 3543614.25 | 2995796.25 | 9012949 | 2712715.25 | 6179504 | 9439668 | 4405860.5 | 9257214 | 3771255.5 | 5120592.5 | 4070074.75 | 3682908 | 3869355.5 | 11118500 |
| A2BH40 | Arid1a | AT-rich interactive domain-containing protein 1A | 132.86 | 1.45 | 242092 | 3 | 2 | 2 | 2 | 2 | 2 | 2 | 2 | 2 | 2 | 2 | 2 | 2 | 2 | 2 | 2 | 2 | 2 | 2 | 2 | 2 |
| P61211 | Arl1 | ADP-ribosylation factor-like protein 1 | 119.44 | 18.78 | 20412 | 2 | 1955424.62 | 232071.48 | 1111076.12 | 497388.81 | 887798.69 | 382902.91 | 670520.69 | 289705.25 | 1319433.12 | 801251.88 | 245937.22 | 290387.41 | 1019834.19 | 810144.88 | 1401095.38 | 291718.91 | 830573.25 | 613499.06 | 587967.25 | 891566.56 |
| P37996 | Arl3 | ADP-ribosylation factor-like protein 3 | 48.69 | 5.49 | 20457 | 1 | 1035682.69 | 757026.5 | 789468.12 | 726200.62 | 651685.88 | 779741 | 2 | 2 | 587319.19 | 2 | 2 | 916530.25 | 589204.38 | 561637.94 | 2 | 2 | 2 | 2 | 2 | 2 |
| Q8R5J9 | Arl6ip5 | PRA1 family protein 3 | 80.16 | 10.64 | 21558 | 2 | 1938221.62 | 2 | 3499930 | 1969192.25 | 661624 | 1250375.12 | 1242387.25 | 3170000.5 | 1246112.62 | 1621319.38 | 524825.81 | 530187.44 | 892421.19 | 548318.12 | 1630810.75 | 1038355.25 | 1461964.25 | 1062310.88 | 767024.75 | 1633933.12 |
| Q66HA6 | Arl8b | ADP-ribosylation factor-like protein 8B | 353.84 | 34.95 | 21539 | 7 | 6990260 | 6658908.5 | 8328986 | 5877835 | 5053010 | 9786702 | 7700510 | 9171300 | 4816356.5 | 4755839.5 | 5613072.5 | 5161671 | 6602783.5 | 6082013.5 | 7510583 | 6360482.5 | 8721134 | 9133912 | 6338864 | 7631375 |
| Q99M08 | Arln | Sarcoplasmic/endoplasmic reticulum calcium ATF | 79 | 21.54 | 7404 | 1 | 2 | 2 | 2 | 2 | 566790.94 | 2 | 2 | 2 | 639497.12 | 604841.5 | 722542.81 | 2 | 691029.12 | 619135.31 | 2 | 2 | 924998.56 | 2 | 710347.5 | 2 |
| Q8BNU0 | Armc6 | Armadillo repeat-containing protein 6 | 92.46 | 4.49 | 50683 | 1 | 39106.47 | 162312.89 | 2 | 2 | 64262.4 | 2 | 170653.23 | 2 | 156436.75 | 95361.9 | 2 | 2 | 83254.73 | 114253.41 | 2 | 2 | 2 | 2 | 2 | 2 |
| Q9DBR3 | Armc8 | Armadillo repeat-containing protein 8 | 44.23 | 1.78 | 75364 | 1 | 387556.09 | 620073.25 | 330131.06 | 361840 | 432933.97 | 2 | 384789.06 | 2 | 2 | 2 | 712553.31 | 731087.81 | 463538.53 | 459387.19 | 2 | 900430.81 | 2 | 632032.69 | 662165.5 | 746739.56 |
| O88656 | Arpc1b | Actin-related protein 2/3 complex subunit 1B | 542.89 | 29.84 | 41057 | 10 | 3950297.5 | 6621550 | 3591698 | 13135074 | 23102742 | 24549932 | 20007496 | 23061116 | 30562190 | 18848198 | 29356634 | 34373086 | 20006690 | 19579284 | 11147530 | 27876422 | 16717224 | 8744902 | 22187158 | 6843695 |
| Q9CVB6 | Arpc2 | Actin-related protein 2/3 complex subunit 2 | 376.03 | 29 | 34357 | 7 | 2 | 867033.31 | 5604529 | 2225375.5 | 1713270.25 | 1343543.88 | 5879761 | 9874703 | 2940207.5 | 6583589.5 | 1990039.25 | 3758356.25 | 6305431.5 | 4418616 | 9939291 | 670710.88 | 2399613 | 6979539 | 834076.69 | 991376.69 |
| Q9JM76 | Arpc3 | Actin-related protein 2/3 complex subunit 3 | 240.64 | 38.2 | 20525 | 5 | 36901054 | 19186378 | 35692566 | 31604164 | 26978390 | 21037770 | 32416528 | 30253432 | 37171606 | 33818910 | 14707474 | 20872154 | 14221682 | 16664045 | 31327250 | 10355558 | 12206244 | 13244689 | 5859636.5 | 10752709 |
| P59999 | Arpc4 | Actin-related protein 2/3 complex subunit 4 | 387.53 | 49.4 | 19667 | 8 | 60034538 | 93693834 | 78313666 | 82753746 | 80929082 | 84223346 | 72674538 | 96685178 | 66755446 | 63990694 | 66816402 | 86081090 | 81723338 | 76485266 | 38343918 | 57356946 | 52702762 | 34119898 | 40217626 | 38384474 |
| Q4KLF8 | Arpc5 | Actin-related protein 2/3 complex subunit 5 | 352.51 | 43.71 | 16320 | 6 | 19920314 | 24855742 | 28453406 | 27251604 | 47787850 | 32577404 | 27722562 | 52983170 | 29333804 | 37972170 | 29931568 | 30670582 | 33747350 | 27191622 | 27761202 | 25939312 | 23505250 | 25177570 | 32623336 | 27261646 |
| A1L108 | Arpc5l | Actin-related protein 2/3 complex subunit 5-like pr | 62.43 | 7.84 | 17010 | 1 | 569147.94 | 115866.55 | 750659.38 | 424665.06 | 1737391.5 | 194161.64 | 1060367 | 192341.23 | 1312679.88 | 1366763.25 | 2 | 852520.75 | 563906.5 | 626563.25 | 645310.69 | 200141.22 | 804294.38 | 262844.81 | 2 | 431991.59 |
| P50428 | Arsa | Arylsulfatase A | 120.15 | 7.31 | 53748 | 2 | 2 | 160870.12 | 2 | 209128.88 | 185090.72 | 2 | 318234.06 | 2 | 171432.95 | 772256.06 | 2 | 104712.91 | 2 | 93098.1 | 2 | 2 | 151855.23 | 2 | 2 | 2 |
| P50429 | Arsb | Arylsulfatase B | 225.44 | 11.24 | 59647 | 5 | 11052510 | 41677662 | 9668609 | 29691966 | 24300826 | 19659024 | 23897950 | 19531650 | 26512258 | 25551388 | 11860489 | 12583316 | 12845280 | 13548618 | 21095094 | 13753672 | 22359338 | 25133506 | 16701349 | 24938332 |
| Q9WV54 | Asah1 | Acid ceramidase | 816.62 | 33.5 | 44670 | 13 | 6280424 | 3044995 | 9448827 | 3993270.25 | 7702790.5 | 2477358.5 | 5963602.5 | 4072677 | 11172639 | 16548362 | 5006175.5 | 5897261.5 | 8671707 | 6948483 | 10042950 | 3303813.5 | 8225957.5 | 5988163 | 5177254.5 | 4725115 |
| Q1AAU6 | Asap1 | Arf-GAP with SH3 domain, ANK repeat and PH do | 110.84 | 2.8 | 127088 | 2 | 2 | 2 | 730088.5 | 823658.06 | 254597.53 | 2 | 81846.97 | 2 | 900507 | 738319.44 | 200507.95 | 271318.59 | 465892.97 | 433824.91 | 714877.75 | 170668.11 | 2 | 1382230.38 | 130081.27 | 276255.69 |
| Q91YI0 | Asl | Argininosuccinate lyase | 415.74 | 22.41 | 51739 | 9 | 3509090.5 | 1034505.12 | 6648063 | 277619.41 | 5192810 | 6284850 | 8120261 | 7353397 | 5600579 | 5908850 | 8500060 | 7979534 | 3621716.25 | 4370568 | 7829727.5 | 3646923 | 1375118.62 | 9023383 | 4894597 | 4326188 |

| | | | | | | | | | | | | | | | | | | | | | | | | | | |
|---|---|---|---|---|---|---|---|---|---|---|---|---|---|---|---|---|---|---|---|---|---|---|---|---|---|---|
| Q61024 | Asns | Asparagine synthetase [glutamine-hydrolyzing] | 414.96 | 17.29 | 64283 | 8 | 776115.56 | 2071921.25 | 2247418 | 1368682.5 | 7386565 | 4865316 | 3601033.25 | 5986780.5 | 4272290 | 4938914 | 5354357 | 5325321 | 7439305.5 | 5578233 | 3583059 | 4516983 | 5538621.5 | 5720265 | 6325166.5 | 4380770 |
| Q8BSY0 | Asph | Aspartyl/asparaginyl beta-hydroxylase | 317.83 | 17 | 83042 | 7 | 307971.28 | 745287.31 | 1068580.12 | 238948.28 | 1055370 | 2 | 154460.08 | 226008.36 | 942368.88 | 1131256.38 | 1573197.25 | 313651.94 | 845549.62 | 646811.5 | 1010784.38 | 1277534.5 | 1101618.75 | 1116888.5 | 2 | 414422.12 |
| Q8C0M9 | Asrgl1 | Isoaspartyl peptidase/L-asparaginase | 83.98 | 5.52 | 33950 | 2 | 1690916 | 2894471 | 1430030.62 | 4321894 | 4126894.75 | 2343946 | 2312596.25 | 2 | 2258820.5 | 2678589.25 | 2003191.62 | 2531290.5 | 1622141.75 | 2324340 | 1912839.75 | 1683306.5 | 2036858.12 | 2154148.5 | 2480858 | 2979514.75 |
| P16460 | Ass1 | Argininosuccinate synthase | 297.31 | 16.99 | 46584 | 6 | 419610 | 1442446 | 2260190 | 2549590.25 | 2535476.25 | 4784286 | 5565525 | 5159064.5 | 8680728 | 2981132.5 | 525744.88 | 4577109.5 | 1554590.5 | 4247562 | 1522642.12 | 2709551.5 | 1632722.12 | 1146435.75 | 1927852.88 | 2482161.25 |
| Q9D906 | Atg7 | Ubiquitin-like modifier-activating enzyme ATG7 | 141.96 | 4.3 | 77520 | 3 | 4201814.5 | 3761543.75 | 4630190.5 | 3653197.25 | 3896952.5 | 93426.41 | 2989151.25 | 1120287 | 2885658.5 | 3649918.25 | 1759115.75 | 1743334.88 | 2780771.75 | 2936376.25 | 2306131 | 1619819.5 | 3279527.75 | 2674177 | 2116290 | 2440309.5 |
| Q9CWJ9 | Atic | Bifunctional purine biosynthesis protein ATIC | 1178.61 | 46.11 | 64217 | 21 | 20026352 | 17812814 | 49520446 | 30595938 | 31341382 | 19418430 | 20263088 | 21766270 | 22938724 | 24909858 | 18974720 | 18253922 | 23863616 | 19355820 | 47547734 | 14300931 | 15431147 | 19764202 | 12588296 | 16184157 |
| Q6PA06 | Atl2 | Atlastin-2 | 213.8 | 10.63 | 66224 | 4 | 27624.37 | 73855.05 | 222084.8 | 2 | 2 | 2 | 2 | 2 | 2 | 309574.97 | 2 | 2 | 2 | 2 | 255863.09 | 1096298.25 | 278949.28 | 252315.45 | 2 | 2 |
| O08997 | Atox1 | Copper transport protein ATOX1 | 245.32 | 67.65 | 7338 | 5 | 9355946 | 5643894 | 10550204 | 5045698 | 18982864 | 3591144.5 | 2980069.5 | 7162022 | 2933443.5 | 18742262 | 18239776 | 23627698 | 23063174 | 24252604 | 16685044 | 18242782 | 16819030 | 12633327 | 21913490 | 19017740 |
| P98197 | Atp11a | Phospholipid-transporting ATPase IH | 41.4 | 0.59 | 135502 | 1 | 2 | 864567.38 | 2 | 389828.25 | 2 | 2 | 2 | 857743.19 | 429865.84 | 378235.56 | 2 | 2 | 2 | 2 | 2 | 2 | 2 | 2 | 2 | 451755.34 |
| Q9EPE9 | Atp13a1 | Endoplasmic reticulum transmembrane helix trans | 337.08 | 7.58 | 132388 | 7 | 1862249.5 | 2614439.25 | 780993.12 | 3449282.5 | 3115770.75 | 1189986.25 | 2248885 | 1621355.25 | 3885627.5 | 2266202.75 | 113702.87 | 1457265.38 | 1412911.12 | 1926099.88 | 408893.19 | 770737 | 1122305.25 | 1391580.88 | 703694.38 | 862439.69 |
| P06685 | Atp1a1 | Sodium/potassium-transporting ATPase subunit a | 1911.99 | 28.74 | 113054 | 27 | 51683118 | 58733018 | 37163354 | 68387410 | 73034482 | 60888610 | 48991602 | 72058594 | 59910234 | 55531886 | 45732318 | 42094854 | 51719846 | 51962398 | 51220970 | 41268482 | 47382974 | 61480710 | 47936278 | 54439542 |
| P06687 | Atp1a3 | Sodium/potassium-transporting ATPase subunit a | 905.11 | 16.78 | 111692 | 13 | 2391573 | 2419511 | 2331271.25 | 1166807.12 | 999970.25 | 2187376.5 | 1800446.25 | 3521716.75 | 1410381.75 | 2316612.75 | 815914.19 | 853971.25 | 1347429 | 1081042.62 | 1663449.25 | 908186.5 | 1011369.31 | 1821785.25 | 909326.06 | 1321809.12 |
| Q9WV27 | Atp1a4 | Sodium/potassium-transporting ATPase subunit a | 457.91 | 9.79 | 114887 | 8 | 1163305.5 | 2 | 2 | 2 | 2 | 2 | 2 | 2 | 2 | 2 | 2 | 2 | 410954.69 | 2 | 2 | 2 | 2 | 2 | 2 | 2 |
| P97370 | Atp1b3 | Sodium/potassium-transporting ATPase subunit b | 252.7 | 21.58 | 31776 | 5 | 5617668.5 | 8129465 | 9098930 | 10026467 | 7901066.5 | 3871233 | 4330180.5 | 6829714.5 | 8033799 | 9435028 | 3880851.5 | 7943046 | 9261379 | 6296454 | 4811753 | 10667640 | 5113354 | 1609853.88 | 3176620 | 1347620.12 |
| O55143 | Atp2a2 | Sarcoplasmic/endoplasmic reticulum calcium ATF | 1427.38 | 30.65 | 114858 | 25 | 10826347 | 18011260 | 9000194 | 10509625 | 14556440 | 17229276 | 18068524 | 16416002 | 16608979 | 13576443 | 16757226 | 21833882 | 20523892 | 21679550 | 16254305 | 18096568 | 18989466 | 21219398 | 18596494 | 18617870 |
| Q64518 | Atp2a3 | Sarcoplasmic/endoplasmic reticulum calcium ATF | 548.09 | 14.01 | 109530 | 10 | 302150.16 | 1849095.5 | 2 | 1776137.75 | 1507069.62 | 1673561.88 | 738210.56 | 230114.34 | 2885349.5 | 2 | 751637.25 | 503041.09 | 473042.81 | 1921527 | 2898360.25 | 1003264.5 | 1328229.5 | 2197889.75 | 2 | 713653.88 |
| G5E829 | Atp2b1 | Plasma membrane calcium-transporting ATPase | 822.81 | 15.57 | 134747 | 15 | 4135646 | 7195983.5 | 4185013.5 | 6333272 | 7697583.5 | 7091795.5 | 5926226 | 8207058 | 7190165 | 9122702 | 5759155 | 3596408.5 | 2503848.25 | 6012841 | 3963900.5 | 6566650 | 8816234 | 14002555 | 8880544 | 10043484 |
| Q03265 | Atp5f1a | ATP synthase subunit alpha, mitochondrial | 2364.39 | 60.94 | 59753 | 36 | 4625405 | 3688032 | 9469738 | 2389386.25 | 11274999 | 3667797.25 | 5017787 | 7191877 | 5053144.5 | 24299220 | 3058582.5 | 2759760.25 | 6352849 | 4288752 | 13676753 | 1572030.88 | 1886899.38 | 2830265 | 1351538.25 | 2107735 |
| P56480 | Atp5f1b | ATP synthase subunit beta, mitochondrial | 1954.19 | 69.94 | 56300 | 30 | 105220114 | 121294354 | 132061626 | 65237022 | 151038066 | 108016986 | 119977018 | 131042450 | 162751698 | 130889842 | 74738698 | 85224322 | 92242922 | 105030346 | 89526314 | 65165866 | 115369034 | 111750890 | 82236538 | 94639818 |
| Q91VR2 | Atp5f1c | ATP synthase subunit gamma, mitochondrial | 473.88 | 30.87 | 32886 | 7 | 8763980 | 328422.34 | 5362742 | 10908781 | 9171022 | 7529054 | 8532879 | 8048202.5 | 10123348 | 10981390 | 8399199 | 6682774.5 | 6934425 | 8692880 | 4950455 | 6328607 | 5789479.5 | 8384286.5 | 924117.75 | 7882051 |
| P35434 | Atp5f1d | ATP synthase subunit delta, mitochondrial | 179.87 | 16.67 | 17595 | 3 | 26766724 | 23135722 | 22454434 | 24157306 | 23628190 | 20813266 | 16717189 | 25370594 | 21684144 | 19520030 | 24777538 | 18879430 | 26171014 | 20788810 | 16401975 | 20637610 | 18656792 | 27219738 | 23154854 | 22755752 |
| P29418 | Atp5f1e | ATP synthase subunit epsilon, mitochondrial | 57.46 | 15.69 | 5767 | 1 | 2 | 884121.81 | 2 | 888455.31 | 2412181.25 | 3610428.75 | 830436.31 | 4632722 | 3818656.25 | 1167003.5 | 3412997.75 | 2 | 2 | 2 | 2 | 2 | 2 | 1626268.62 | 2 | 1844002.62 |
| O35143 | Atp5if1 | ATPase inhibitor, mitochondrial | 56.36 | 8.49 | 12159 | 1 | 2 | 2 | 2 | 638228.69 | 1262100 | 921967.94 | 2 | 2 | 877908.44 | 2 | 2098631 | 3314753.75 | 1882607 | 1902794.12 | 1274086.25 | 5901235.5 | 3008104.5 | 1422321.12 | 4440892 | 1494093.12 |
| Q06185 | Atp5me | ATP synthase subunit e, mitochondrial | 183.09 | 60.56 | 8236 | 4 | 2 | 2 | 1655193.25 | 2 | 1176174.38 | 905054.25 | 652273.12 | 2398256.25 | 572174.5 | 1143396 | 2 | 1129106 | 1013834.19 | 853593.88 | 2 | 2 | 2 | 1354453.25 | 2791849.75 | 2 |
| Q6PDU7 | Atp5mg | ATP synthase subunit g, mitochondrial | 70.51 | 14.56 | 11433 | 1 | 2068122.62 | 7256615.5 | 9648442 | 2 | 11604878 | 6226397 | 6392728 | 12379213 | 8355344.5 | 2 | 5233444 | 4393209 | 5859957 | 6071376 | 2 | 1888102.75 | 2 | 2 | 2 | 2 |
| Q78IK2 | Atp5mk | ATP synthase membrane subunit K, mitochondria | 285.01 | 44.83 | 6382 | 4 | 3405935.25 | 7228626 | 3425389.5 | 5696569 | 12159346 | 9966026 | 10584700 | 14478956 | 8289979 | 8343646.5 | 3138785.75 | 4302186.5 | 5350026 | 4130560.75 | 2443257.25 | 4152667.5 | 4103461.5 | 5075045.5 | 5330298 | 3105984 |
| Q9CQQ7 | Atp5pb | ATP synthase F(0) complex subunit B1, mitochon | 219.65 | 20.7 | 28949 | 4 | 6840195.5 | 5844941 | 6928086.5 | 7313789 | 9111476 | 1827050.5 | 2142394.75 | 5548358.5 | 5265286.5 | 7746120 | 3637253 | 3893909.5 | 5252252 | 4264695 | 5193827.5 | 4000270.5 | 1617523.75 | 7188362 | 4962959 | 6273238 |
| Q9DCX2 | Atp5pd | ATP synthase subunit d, mitochondrial | 219.72 | 40.99 | 18749 | 5 | 3881259 | 5694253 | 6281505.5 | 2615428 | 7635498 | 5472231.5 | 4028929.75 | 4423016 | 6099726 | 8547936 | 6685845.5 | 7126541.5 | 7656401 | 8330359.5 | 4725985 | 5729698 | 5859278.5 | 4425733 | 5491643.5 | 8453325 |
| Q9DB20 | Atp5po | ATP synthase subunit O, mitochondrial | 432.27 | 38.5 | 23364 | 7 | 6458527.5 | 14510084 | 10689834 | 1997533 | 15415220 | 12327827 | 10460673 | 9494218 | 12454487 | 11318398 | 8053922 | 9163450 | 11650329 | 9329042 | 6712365 | 9058306 | 7414272.5 | 5918900.5 | 9348206 | 6007372 |
| P25286 | Atp6v0a1 | V-type proton ATPase 116 kDa subunit a 1 | 69.05 | 1.19 | 96328 | 1 | 2 | 1219348.25 | 2 | 1677194 | 1885745.25 | 2596508 | 1655302.88 | 1928735.38 | 2318678.75 | 1726708.88 | 1189518.75 | 786211.5 | 2 | 939038.88 | 2 | 812203 | 2 | 2 | 801053.06 | 2 |
| P63081 | Atp6v0c | V-type proton ATPase 16 kDa proteolipid subunit | 453.76 | 78.06 | 15808 | 8 | 75299250 | 77029386 | 67346314 | 65527334 | 51232546 | 69549162 | 53896914 | 85451954 | 72418826 | 62634422 | 34678934 | 28890156 | 61342130 | 53516830 | 46498514 | 25316982 | 48405414 | 89260498 | 71275066 | 69942754 |
| P51863 | Atp6v0d1 | V-type proton ATPase subunit d 1 | 324.48 | 17.66 | 40301 | 6 | 17824448 | 11490044 | 14576622 | 9431236 | 11415456 | 10631918 | 9244979 | 11038934 | 10491650 | 12488004 | 8094022.5 | 5722948.5 | 9429027 | 8131502.5 | 12817683 | 9330852 | 11026087 | 10188483 | 8231695 | 11289980 |
| P50516 | Atp6v1a | V-type proton ATPase catalytic subunit A | 1285.75 | 49.27 | 68326 | 21 | 40417418 | 63318370 | 55133682 | 46317678 | 52444506 | 52934786 | 55546662 | 66028462 | 47203498 | 51957766 | 45255078 | 41091898 | 23128326 | 37875298 | 25712602 | 43724562 | 40579890 | 26422168 | 40831694 | 47895014 |
| P62814 | Atp6v1b2 | V-type proton ATPase subunit B, brain isoform | 1486.14 | 58.32 | 56551 | 25 | 58114250 | 78147642 | 58250134 | 40014958 | 39076658 | 72711946 | 57090542 | 67267538 | 60514182 | 43185926 | 57585450 | 51348714 | 48837598 | 51001090 | 43737698 | 58504826 | 57897834 | 64704834 | 76744450 | 67141170 |
| Q5FVI6 | Atp6v1c1 | V-type proton ATPase subunit C 1 | 178.45 | 6.28 | 43901 | 4 | 3771200.5 | 4233084 | 4108732.75 | 6903256.5 | 7457326.5 | 10687660 | 8610493 | 4393404 | 10619413 | 6793600 | 6736189.5 | 4295677.5 | 2947538.5 | 1987255 | 5970034.5 | 2482885.5 | 3170163.5 | 649977.81 | 2589620.5 | 4958885.5 |
| P57746 | Atp6v1d | V-type proton ATPase subunit D | 120.16 | 12.96 | 28369 | 2 | 2 | 2 | 2 | 2 | 1052252.38 | 2 | 2 | 1379327.62 | 1181317 | 2 | 1398127.75 | 1398636.75 | 1347891.5 | 1198767.75 | 2 | 855253.62 | 2 | 1340582.25 | 544499.81 | 2 |
| P50518 | Atp6v1e1 | V-type proton ATPase subunit E 1 | 421.88 | 25.66 | 26157 | 7 | 13937820 | 10575378 | 7960529.5 | 6459275.5 | 6171211 | 18544002 | 30127150 | 13299110 | 12984128 | 8695880 | 8912411 | 11071422 | 18954148 | 10115088 | 17629484 | 12144979 | 19593662 | 12672636 | 12802992 | 9741626 |
| Q9CR51 | Atp6v1g1 | V-type proton ATPase subunit G 1 | 295.68 | 34.75 | 13724 | 4 | 2 | 542929.56 | 2 | 995452.44 | 1445024.25 | 2 | 834100.56 | 2136486.75 | 1468424 | 1448848.75 | 2383006.75 | 2891978.5 | 1653574.12 | 1529986.25 | 713232.25 | 2675486 | 1306616.5 | 1342587.12 | 2221189 | 2 |
| Q8BVE3 | Atp6v1h | V-type proton ATPase subunit H | 311.71 | 16.36 | 55855 | 7 | 697950.5 | 592864.12 | 538543.75 | 1240671.38 | 717263.56 | 2976704.75 | 3007390.5 | 3972842 | 2819988 | 1650369.75 | 4564762 | 3849971 | 4224204.5 | 5540296 | 5296571.5 | 4642321 | 4408350 | 4816911 | 5232386 | 3798193 |
| Q64430 | Atp7a | Copper-transporting ATPase 1 | 205.08 | 2.62 | 161959 | 5 | 2921221.5 | 5792189 | 4992709 | 4142166.5 | 3116394.25 | 5635956.5 | 6622747 | 9643801 | 1927968.75 | 3615122.5 | 3077435.25 | 3323434.25 | 2590063.25 | 2703677.75 | 3516664.5 | 2482957.75 | 3131783 | 4763616.5 | 3858894 | 3699361.25 |
| P28658 | Atxn10 | Ataxin-10 | 174.79 | 13.26 | 53707 | 4 | 1353991 | 1317362.25 | 1253093.25 | 1483892.75 | 2182346.75 | 1349542.25 | 1350949.88 | 2009942 | 1133995.62 | 726975.94 | 1285113.62 | 977574.94 | 1612120.12 | 817473.88 | 1156894.88 | 1371654.25 | 944279.5 | 1646263.62 | 941803.75 | 1441520.88 |
| Q7TQH0 | Atxn2l | Ataxin-2-like protein | 63.65 | 1.62 | 110649 | 1 | 2 | 2 | 2 | 2 | 2 | 2 | 2 | 2 | 2 | 614708.94 | 1284224.12 | 2 | 2 | 2 | 2 | 1005087.38 | 2 | 2 | 2 | 2 |
| P70295 | Aup1 | Lipid droplet-regulating VLDL assembly factor AU | 122.99 | 11.46 | 46121 | 3 | 542371.44 | 507927.88 | 896573.56 | 876651.25 | 28996.52 | 759512.5 | 2 | 2 | 741123.81 | 1044502.94 | 913019.69 | 2 | 82496.77 | 710788.75 | 2 | 598094.19 | 2 | 861390.69 | 2 | 2 |
| P01887 | B2m | Beta-2-microglobulin | 97.92 | 15.97 | 13779 | 2 | 1504643.88 | 458373.53 | 8252257 | 2 | 4225056 | 2183554.75 | 1421758.38 | 5511220 | 685105.62 | 4080974 | 3342360.25 | 1769961 | 2 | 2 | 572968 | 2 | 2 | 2 | 2 | 2 |
| Q5XIJ6 | Babam1 | BRISC and BRCA1-A complex member 1 | 59.19 | 3.29 | 36878 | 1 | 2 | 2 | 2 | 2 | 2 | 2 | 2 | 2 | 536083.44 | 2 | 2188701.5 | 2072568.75 | 1626351.75 | 1923385.75 | 2 | 2 | 2 | 2 | 1885959.88 | 2 |
| Q6P7Q1 | Babam2 | BRISC and BRCA1-A complex member 2 | 43.99 | 3.66 | 43559 | 1 | 10639552 | 294708.06 | 866325.06 | 329391.03 | 1277591.5 | 2384929.75 | 223653.64 | 254666.89 | 726267.94 | 1003388.44 | 273820.69 | 783656.12 | 957470.06 | 583890.06 | 749748.12 | 826456.5 | 1774745.75 | 1787554.12 | 2035518.62 | 620086.94 |
| O08734 | Bak1 | Bcl-2 homologous antagonist/killer | 125.45 | 10.05 | 23295 | 2 | 4226833 | 4227369.5 | 3893848.25 | 3498452.5 | 3758558 | 3361577 | 5627437.5 | 3223317.5 | 4395613.5 | 2 | 3898690 | 3699351.5 | 2193547.5 | 3442582.5 | 2769250.5 | 5951657 | 7392023 | 6606081 | 5576039.5 | 5810183 |
| O54962 | Banf1 | Barrier-to-autointegration factor | 152.17 | 44.94 | 10103 | 3 | 682882.12 | 171385.73 | 762515.69 | 466400.09 | 1603738.5 | 608168.38 | 602515.44 | 438178.56 | 737951.25 | 1239810 | 248304.06 | 148970.89 | 877014.38 | 485513.62 | 791956.25 | 2 | 510250.06 | 133331.09 | 336755.56 | 419320.06 |
| Q91XV3 | Basp1 | Brain acid soluble protein 1 | 1220.93 | 84.07 | 22087 | 16 | 4899134.5 | 21497860 | 3050009.5 | 24640508 | 4990265.5 | 13939046 | 13217033 | 12827260 | 47582938 | 11964437 | 15036685 | 31521686 | 16285452 | 33491336 | 16662203 | 39463950 | 32998966 | 22803346 | 52125850 | 43376354 |
| Q07813 | Bax | Apoptosis regulator BAX | 332.54 | 30.21 | 21395 | 5 | 19227790 | 18436656 | 14229237 | 11201260 | 10010947 | 13355708 | 11766482 | 16605920 | 10197946 | 7707765 | 11828392 | 10679925 | 9664730 | 8240736.5 | 5963051 | 8193922 | 15595249 | 12436076 | 11791528 | 9039696 |
| Q61335 | Bcap31 | B-cell receptor-associated protein 31 | 42.63 | 4.08 | 27957 | 1 | 35601.73 | 536424.69 | 2 | 2 | 71296.44 | 2 | 84164.8 | 2 | 585349.5 | 21483.98 | 2 | 2 | 108011.04 | 564123.94 | 367869.06 | 358554.38 | 497929.72 | 146203.69 | 2 | 1141248.5 |
| P24288 | Bcat1 | Branched-chain-amino-acid aminotransferase, cy | 45.09 | 3.37 | 42791 | 1 | 2 | 2 | 2 | 403776.91 | 604793.94 | 2 | 2 | 998122.5 | 471037.91 | 601069.25 | 2 | 2 | 2 | 2 | 2 | 847200.06 | 2 | 2 | 138389.5 | 2 |
| O35855 | Bcat2 | Branched-chain-amino-acid aminotransferase, mi | 212.27 | 8.4 | 44127 | 3 | 7601986 | 7804404 | 7064165 | 7709301 | 6745147 | 6533809 | 7151395 | 7941665 | 6263648 | 6091324 | 10133894 | 7639865 | 6785712.5 | 6506901.5 | 6503449 | 7927402.5 | 6670253 | 6242087 | 6626725.5 | 6670542.5 |
| P11960 | Bckdha | 2-oxoisovalerate dehydrogenase subunit alpha, n | 66.65 | 3.63 | 50164 | 2 | 2 | 2 | 762155.88 | 2 | 1120714.38 | 2 | 2 | 697430.12 | 407079.38 | 2 | 1141771.38 | 1313179.25 | 2 | 2 | 679137.25 | 389960.25 | 796496 | 874014.06 | 2 | 2 |
| P53563 | Bcl2l1 | Bcl-2-like protein 1 | 120.13 | 9.87 | 26158 | 3 | 872738.94 | 702604.5 | 1064574 | 2 | 79372.86 | 2 | 99428.72 | 81143.65 | 2 | 935705 | 2 | 50928.7 | 495334.81 | 87571.73 | 1357406.88 | 2 | 2 | 754667.31 | 2 | 374424.81 |
| P59017 | Bcl2l13 | Bcl-2-like protein 13 | 141.13 | 12.21 | 46719 | 3 | 2 | 529414.06 | 2 | 2 | 55342.47 | 2 | 396319.91 | 2 | 1066284.88 | 948859.44 | 1103659.12 | 1323844.88 | 1365154.75 | 1207360.25 | 2 | 1207189.12 | 1128477.12 | 860885.56 | 1117890.12 | 1163538.75 |
| Q8K019 | Bclaf1 | Bcl-2-associated transcription factor 1 | 40.68 | 0.98 | 106002 | 1 | 1535847.75 | 2460749.5 | 170131.84 | 3130328.5 | 377198 | 2989436.5 | 2721738.5 | 2 | 3421658.25 | 2263511 | 2612928 | 3256075.5 | 2506674 | 2407824.25 | 1577483.75 | 2601022.5 | 2732228.75 | 2111706 | 3008929 | 2562212.25 |
| Q9CZP5 | Bcs1l | Mitochondrial chaperone BCS1 | 115.28 | 6.94 | 47406 | 2 | 2 | 2 | 56555.04 | 2 | 30001.29 | 51218.16 | 75702.81 | 2 | 54625.32 | 462738.38 | 2 | 39154.84 | 284754.25 | 38872.45 | 212603.17 | 2 | 65933.13 | 2 | 2 | 2 |
| D3Z6Q9 | Bin2 | Bridging integrator 2 | 251.4 | 10.22 | 52554 | 4 | 2095120.12 | 930018.38 | 2 | 1553352.75 | 2885104 | 2384462.5 | 2110697.5 | 1218321.75 | 3162429.75 | 1208974.38 | 4481338 | 5000398.5 | 3642050 | 4737144 | 2269586.5 | 4356563 | 4434156 | 3362007 | 4922474.5 | 3540454.75 |
| O88738 | Birc6 | Dual E2 ubiquitin-conjugating enzyme/E3 ubiquiti | 46.51 | 0.23 | 532170 | 1 | 68989.83 | 2 | 155151.41 | 2 | 82147.55 | 2 | 2 | 2 | 144146.98 | 96737.19 | 116538.2 | 219438.5 | 189859.17 | 237339.33 | 478516.47 | 2 | 229710.17 | 284162.19 | 2 | 212691.83 |
| Q9CY64 | Blvra | Biliverdin reductase A | 288.7 | 23.73 | 33525 | 6 | 4348333 | 3045923.5 | 4052789 | 2906051 | 5641208 | 5624017 | 5893045 | 6941539 | 7059546 | 7843747.5 | 5619433.5 | 5401673.5 | 3834120.25 | 3170150.25 | 4828707 | 5138730 | 3684769.5 | 5466146.5 | 3318415 | 2865863.25 |
| Q923D2 | Blvrb | Flavin reductase (NADPH) | 368.46 | 42.72 | 22197 | 5 | 2 | 744125.31 | 74026.41 | 916532.38 | 909614.75 | 2 | 126051.77 | 276664 | 666382.75 | 1300019.12 | 4655565 | 3433830.25 | 5021616.5 | 4789113 | 3993346.25 | 11786431 | 6163093 | 4988602 | 9568331 | 2956711 |
| Q6QD59 | Bnip1 | Vesicle transport protein SEC20 | 38.31 | 4.82 | 26175 | 1 | 862904.19 | 916113.56 | 2 | 2 | 1218534.12 | 2 | 2 | 648321.56 | 458044.56 | 638689.69 | 2 | 1301798.25 | 1510980.5 | 2 | 2 | 2 | 2 | 2 | 2 | 2 |
| Q8BGS2 | Bola2 | BolA-like protein 2 | 58.37 | 18.6 | 10215 | 2 | 2 | 2 | 2 | 2 | 2 | 408182.91 | 2 | 2 | 2 | 2 | 1621187.5 | 1576989.75 | 509885.84 | 1269350.38 | 864614.38 | 2 | 2 | 854409.06 | 454425.66 | 963789.69 |
| Q8R164 | Bphl | Valacyclovir hydrolase | 35.17 | 4.12 | 32851 | 1 | 1162641.12 | 2 | 1625179.75 | 687468.12 | 2 | 2 | 1166695.25 | 1542122.62 | 2 | 570700.44 | 2 | 2 | 874105.12 | 2 | 2 | 2311647.75 | 2 | 923243.5 | 2 | 2 |
| Q9Z0S1 | Bpnt1 | 3'(2'),5'-bisphosphate nucleotidase 1 | 132.15 | 10.39 | 33196 | 3 | 1600691.5 | 3531484 | 3168758.25 | 2996333.5 | 4741087.5 | 2989963.25 | 4794500 | 4338144 | 3408451.25 | 4329267 | 3446251.5 | 4744150 | 2231682.5 | 4502282 | 3972045.5 | 4055036 | 2508818.5 | 1873673.25 | 4398467 | 620853.12 |
| Q6MGA9 | Brd2 | Bromodomain-containing protein 2 | 39.84 | 1.63 | 88051 | 1 | 2 | 2 | 2 | 431901.22 | 789414 | 2 | 2 | 2 | 639081.69 | 2 | 2 | 728095.75 | 677281.19 | 509002.41 | 572667.31 | 718327.81 | 818005.62 | 655771.25 | 2 | 2 |
| Q4QQT6 | Brix1 | Ribosome biogenesis protein BRX1 homolog | 110.26 | 8.81 | 41115 | 3 | 2 | 2 | 50965.27 | 2 | 2 | 2 | 2 | 2 | 2 | 2 | 2 | 2 | 61008.66 | 107527.67 | 313215.47 | 2 | 2 | 2 | 111967.73 | 2 |
| Q91VR8 | Brk1 | Protein BRICK1 | 117.96 | 17.33 | 8761 | 2 | 6540790.5 | 6466445 | 1854056.25 | 5605246 | 344074.94 | 5068431 | 4664222.5 | 376026.69 | 3873147.75 | 3459649.75 | 1345984.25 | 2 | 3705036 | 2488386.5 | 1185375.75 | 4055927.5 | 3241623.75 | 5740787 | 3338742.5 | 6424408 |
| Q4V8K5 | Brox | BRO1 domain-containing protein BROX | 260.33 | 14.6 | 46192 | 4 | 2546721.25 | 3248494.5 | 798601.38 | 2204525.5 | 950077.12 | 1847293 | 1485935 | 2477972.5 | 2311213.75 | 997439.69 | 259257.86 | 2 | 412646.19 | 1433311.5 | 706209.88 | 2 | 3138416.25 | 2363103 | 964679.75 | 903114.94 |
| P18572 | Bsg | Basigin | 152.76 | 8.74 | 42445 | 3 | 6967153.5 | 6870972.5 | 5594994 | 8548620 | 7067159.5 | 4104970.75 | 3037822.5 | 6394070 | 3848687 | 4788228 | 6456850 | 5771997.5 | 7687915 | 6458344 | 5318201 | 7780640 | 6361833.5 | 11540501 | 9301632 | 8822195 |
| Q8R2Q8 | Bst2 | Bone marrow stromal antigen 2 | 130.69 | 6.4 | 19152 | 2 | 316569.66 | 2078147 | 39158.73 | 1355576.38 | 2 | 512175.16 | 145770.34 | 2 | 744231.56 | 53335.88 | 2 | 2 | 805826.06 | 2 | 2 | 2 | 2 | 2 | 2 | 2 |
| Q9CQH7 | Btf3l4 | Transcription factor BTF3 homolog 4 | 37.25 | 4.43 | 17271 | 1 | 1267602.88 | 934343.5 | 1885596 | 1058577.12 | 1145849.75 | 547360.44 | 899888.44 | 1865359.62 | 761914.94 | 1646934.25 | 1376301.12 | 905703.38 | 1207683.62 | 1107162.88 | 1436030 | 1754806.88 | 1481164.38 | 1184278.12 | 1314766.75 | 1641033.88 |
| Q9WVA3 | Bub3 | Mitotic checkpoint protein BUB3 | 464.16 | 26.99 | 36955 | 9 | 8015411.5 | 7251370 | 4369169 | 9499091 | 13199837 | 6591767 | 8351545 | 10170387 | 10168624 | 7232028 | 10513295 | 9294272 | 8597819 | 6260941 | 4749236.5 | 2060890.25 | 6629213 | 7771014 | 5815921 | 8406151 |
| O54825 | Bysl | Bystin | 44.45 | 2.06 | 49784 | 1 | 2 | 2 | 2 | 2 | 2 | 2 | 2 | 2 | 2 | 2 | 2 | 2 | 2 | 2 | 2 | 2 | 2 | 2 | 2 | 2 |
| P30993 | C5ar1 | C5a anaphylatoxin chemotactic receptor 1 | 228.58 | 9.4 | 39023 | 5 | 4520583 | 14449967 | 5509932 | 4057392 | 7032749 | 19437082 | 17591396 | 8640233 | 13365501 | 9205211 | 6496543.5 | 6076563.5 | 9852477 | 10446799 | 3816359.5 | 6217864.5 | 8882338 | 7205486 | 9360184 | 4651491.5 |
| Q9CXW3 | Cacybp | Calcyclin-binding protein | 185.42 | 21.83 | 26510 | 4 | 1286167.5 | 862311.81 | 2693724.75 | 4414041 | 899383.88 | 657630.19 | 115789.46 | 2 | 1458347.38 | 1364132.5 | 2188627.25 | 3470379.5 | 2051232.5 | 1657603.62 | 4081677.5 | 3916888 | 1125808.12 | 436406.16 | 2060123.75 | 160914.81 |
| B2RQC6 | Cad | Multifunctional protein CAD | 408.88 | 4.36 | 243238 | 8 | 2409349.5 | 3725694.5 | 4811599.5 | 5443382 | 4212431 | 3084986.75 | 3571845.75 | 2849377.25 | 3038123.25 | 2903305.5 | 5020576 | 3948916.75 | 5562407.5 | 6432562 | 7267628.5 | 3428167.5 | 4961402 | 6103798.5 | 2660789.75 | 4859092.5 |
| Q8R5M8 | Cadm1 | Cell adhesion molecule 1 | 31.83 | 1.97 | 49788 | 1 | 1131991.12 | 1296511 | 244147.12 | 764691.44 | 133796.69 | 2 | 2 | 38663.09 | 1071606.12 | 2 | 1288167 | 2115178 | 1716078.38 | 1947441.12 | 1947803.25 | 929565.75 | 2 | 2194545.75 | 1724170.62 | 2845871.25 |
| P0DP26 | Calm1 | Calmodulin-1 | 264.75 | 42.95 | 16838 | 6 | 55610166 | 53260054 | 1063857.62 | 50211966 | 378542.94 | 37511230 | 30686472 | 321719.88 | 29622964 | 39908302 | 40284090 | 47159166 | 44892838 | 36251858 | 39740718 | 41865374 | 59043142 | 65540462 | 49318286 | 82821786 |
| P14211 | Calr | Calreticulin | 1099.48 | 53.85 | 47995 | 19 | 31973964 | 29503016 | 27311712 | 48223334 | 80996402 | 50165110 | 52503486 | 105097554 | 42894658 | 80154810 | 35949894 | 34287078 | 46122494 | 36756802 | 31423522 | 41008650 | 28735426 | 24969246 | 44711278 | 19450806 |
| O35887 | Calu | Calumenin | 222.13 | 18.1 | 37064 | 5 | 3512765.25 | 4007345.25 | 6034832 | 4976201.5 | 3413002.5 | 1920985.25 | 3177052.75 | 3910881 | 2894053.75 | 5764517 | 3868048.75 | 4837302 | 6190252 | 3965133 | 4681704 | 5528765.5 | 3909356.75 | 5388572 | 2133857.5 | 3474420 |
| Q6PHZ2 | Camk2d | Calcium/calmodulin-dependent protein kinase typ | 228.18 | 10.42 | 56369 | 5 | 3551099.5 | 3338733 | 2477870.5 | 1647751.12 | 3625732.5 | 2438797 | 2744452.5 | 2664616 | 5531193.5 | 6317036.5 | 3663084 | 4007589 | 3778555 | 4857742.5 | 4336448 | 1555611.75 | 3377590.25 | 3646485.75 | 1455641.88 | 3270009 |
| P97536 | Cand1 | Cullin-associated NEDD8-dissociated protein 1 | 674.58 | 13.09 | 136362 | 13 | 11176378 | 21132404 | 9445359 | 10732416 | 8819796 | 10099260 | 11191298 | 4016789.5 | 8692355 | 8108085 | 6553329.5 | 12274046 | 12064387 | 8743069 | 6266695.5 | 11321927 | 12859224 | 13774437 | 11293430 | 13745200 |
| P35564 | Canx | Calnexin | 965.5 | 30.12 | 67278 | 18 | 11680755 | 10880212 | 12214000 | 15951912 | 12449346 | 10106980 | 11162696 | 9945812 | 14933893 | 11800377 | 9814306 | 9192345 | 10559762 | 10239144 | 9637266 | 10700595 | 9853512 | 9222194 | 6904150 | 10544959 |
| P40124 | Cap1 | Adenylyl cyclase-associated protein 1 | 1697.28 | 55.06 | 51565 | 23 | 63808562 | 74907690 | 60009618 | 94760506 | 69672642 | 87411826 | 85057282 | 104806162 | 65613878 | 60737974 | 104263618 | 103568586 | 91174018 | 84373210 | 103075074 | 82604674 | 94583426 | 81479138 | 84232954 | 87148554 |
| P24452 | Capg | Macrophage-capping protein | 832.16 | 42.33 | 39240 | 14 | 2 | 2 | 1897432.12 | 2 | 797125.56 | 2 | 2 | 2 | 2 | 1754610.12 | 2987200.75 | 2615674.5 | 7843720.5 | 3403464.5 | 1971825.62 | 400574.81 | 1274019.75 | 966181.56 | 2132503.75 | 2 |
| O08529 | Capn2 | Calpain-2 catalytic subunit | 162.4 | 5.14 | 79872 | 3 | 1775949.62 | 1635024.88 | 3675185.25 | 995065.44 | 987664.12 | 1171778.5 | 1312556.5 | 937368 | 1095980.5 | 1438856.38 | 1760669.88 | 2914380.5 | 3828723.25 | 3376346.25 | 4717351.5 | 1525086.12 | 2681595.25 | 3873440.25 | 2009967.62 | 2038465.75 |
| O88456 | Capns1 | Calpain small subunit 1 | 160.51 | 16.04 | 28406 | 3 | 1162280.75 | 1316239.75 | 1902128.5 | 1207570 | 1200935.75 | 1212966.5 | 1860974.75 | 1407135.25 | 1126501.88 | 1796750 | 503185.31 | 995577.25 | 1028997.88 | 2631913 | 4574847 | 162320.62 | 1897868.25 | 1593358.5 | 355385.28 | 2164132.25 |
| Q5M9G3 | Caprin1 | Caprin-1 | 128.89 | 4.81 | 78121 | 3 | 943446.62 | 2 | 582356.25 | 2 | 449776.47 | 669854.88 | 498388.97 | 179587.69 | 877046.38 | 701061.62 | 1930102.62 | 1004292.75 | 2471390.5 | 2106798.5 | 1648500.12 | 977938 | 2 | 2 | 2 | 1124780.75 |
| P47753 | Capza1 | F-actin-capping protein subunit alpha-1 | 325.51 | 20.98 | 32940 | 6 | 4453168 | 1665168.38 | 7828075.5 | 3809537 | 2646451.75 | 563391.56 | 1562906.38 | 1291930.5 | 3028700.5 | 1794140.62 | 532035.88 | 2781435 | 3406923.25 | 3625206 | 4446394 | 100709.66 | 3237609.5 | 2591567 | 1217628.5 | 3527774 |
| P47754 | Capza2 | F-actin-capping protein subunit alpha-2 | 239.39 | 20.28 | 32967 | 5 | 15324306 | 12337470 | 22295166 | 7149418 | 15259538 | 7604127 | 8064184 | 3941744 | 6652318 | 11802642 | 7084455 | 5551150 | 7799050 | 7179145 | 12489584 | 2544906.25 | 4327893 | 6577534 | 3199923 | 4539053.5 |
| P47757 | Capzb | F-actin-capping protein subunit beta | 314.26 | 22.74 | 31345 | 5 | 39950358 | 48412102 | 25477404 | 35014570 | 46382710 | 36411250 | 39819450 | 46205954 | 53296538 | 39834610 | 44858850 | 50486186 | 34958050 | 39342986 | 24532458 | 17643928 | 13944169 | 21228166 | 21062962 | 21257944 |

| | | | | | | | | | | | | | | | | | | | | | | | | | | |
|---|---|---|---|---|---|---|---|---|---|---|---|---|---|---|---|---|---|---|---|---|---|---|---|---|---|---|
| Q4AE70 | Carm1 | Histone-arginine methyltransferase CARM1 | 47.15 | 2 | 70341 | 1 | 2 | 91145.7 | 2 | 101654.23 | 234847.55 | 2 | 2 | 2 | 185837.86 | 2 | 83549.87 | 145656.7 | 2 | 308058.31 | 2 | 68411.55 | 223994.7 | 2 | 2 | 137205.36 |
| Q9ER72 | Cars1 | Cysteine--tRNA ligase, cytoplasmic | 256.06 | 5.9 | 94860 | 4 | 2 | 1678257.12 | 2 | 2 | 3904268.5 | 2056942.75 | 1609134.25 | 2894205.25 | 3116892.5 | 3268230 | 7720486 | 6098886 | 6003927 | 5086881 | 4414969 | 8308785.5 | 6971766 | 5075053 | 9450274 | 4149784.25 |
| P29452 | Casp1 | Caspase-1 | 247.73 | 10.95 | 45640 | 5 | 9711280 | 7807474 | 9343158 | 9227693 | 13095631 | 13083000 | 12511570 | 8719707 | 9938592 | 12338489 | 11269922 | 14702326 | 13120986 | 9414579 | 13147394 | 9626244 | 13656546 | 9628586 | 11061056 | 15483574 |
| P70677 | Casp3 | Caspase-3 | 97.74 | 6.86 | 31475 | 2 | 831058.94 | 1844383.12 | 512194.91 | 4198423.5 | 2 | 6075640 | 3694750.75 | 5983991 | 1062650 | 1014546.25 | 2660061 | 2443687.75 | 3067982.25 | 2311632 | 2 | 1390918.25 | 2522472.25 | 2629002.25 | 1976722.5 | 2047996.12 |
| O89110 | Casp8 | Caspase-8 | 43.58 | 2.29 | 55357 | 1 | 2 | 1322800.38 | 1033700.06 | 2 | 908009.25 | 2 | 1065988.75 | 1031803.12 | 2 | 746697.94 | 2 | 872152.12 | 2 | 2 | 2 | 825776.69 | 2 | 2 | 2 | 2 |
| P51125 | Cast | Calpastatin | 56.49 | 2.92 | 84922 | 1 | 2 | 2 | 2 | 2 | 83398.97 | 2 | 2 | 2 | 106637.91 | 75580.57 | 2 | 151925.41 | 2 | 118288.98 | 2 | 2 | 2 | 2 | 2 | 2 |
| P04762 | Cat | Catalase | 335.45 | 13.09 | 59757 | 6 | 3188444.75 | 2783376.5 | 4354390.5 | 183842.34 | 3192236 | 3034477 | 1768582.88 | 4294577 | 1083005.5 | 2674074.75 | 14609217 | 18908598 | 7158840 | 14648034 | 5916564 | 15377904 | 9396479 | 4177789.5 | 13383325 | 5859010 |
| P48758 | Cbr1 | Carbonyl reductase [NADPH] 1 | 164.41 | 9.03 | 30641 | 3 | 2441443.5 | 1501072.25 | 1516362 | 2428667.5 | 1450550 | 3856318 | 3577368 | 4228679 | 3718460.75 | 1998752.25 | 3401113.5 | 3490770.75 | 1662674.75 | 1962451.5 | 2738676 | 4345914 | 5630500 | 3725519 | 3257181.5 | 9048481 |
| Q8K354 | Cbr3 | Carbonyl reductase [NADPH] 3 | 134.38 | 9.75 | 30953 | 3 | 2 | 469415.56 | 2 | 2 | 705827.56 | 587063.81 | 704904.25 | 2 | 407715 | 2 | 1301904.62 | 710783.25 | 979061.56 | 1518611.25 | 903919.31 | 1294165.75 | 733597.44 | 1036044.19 | 1093878.88 | 2 |
| Q91VT4 | Cbr4 | 3-oxoacyl-[acyl-carrier-protein] reductase | 36.69 | 4.24 | 25415 | 1 | 2 | 2 | 2 | 2 | 2 | 2 | 2 | 2 | 1393500.5 | 2 | 2 | 2 | 2 | 2 | 2 | 2 | 2 | 2 | 2 | 2 |
| P23198 | Cbx3 | Chromobox protein homolog 3 | 333.92 | 32.24 | 20855 | 5 | 7450226.5 | 5331500 | 7755827.5 | 4312922.5 | 8388201 | 3932927.75 | 3517973.5 | 6122909 | 4170302.75 | 4952925 | 5690278 | 5600859 | 6489706.5 | 3929435 | 5859210.5 | 5590375 | 7026068.5 | 6131924 | 5675097.5 | 1866288.38 |
| Q61686 | Cbx5 | Chromobox protein homolog 5 | 146.66 | 12.57 | 22186 | 3 | 1442242 | 1265525.88 | 700783.69 | 2334268 | 2510495.25 | 734041.75 | 771620.25 | 2 | 1634009 | 1055435.62 | 784650.94 | 1315928.25 | 1469804.75 | 1201099.38 | 926160.38 | 1717345.88 | 1167665.88 | 456122.88 | 1550929.12 | 2 |
| Q8VDP4 | Ccar2 | Cell cycle and apoptosis regulator protein 2 | 84.21 | 2.6 | 103002 | 2 | 2 | 2 | 2 | 2 | 2 | 2 | 2 | 2 | 2 | 2 | 739268.75 | 2 | 2 | 461515.91 | 2 | 2 | 705637.69 | 2 | 2 | 2 |
| Q8VE99 | Ccdc115 | Coiled-coil domain-containing protein 115 | 187.89 | 23.89 | 19743 | 4 | 1381828.88 | 664761.75 | 2 | 2255037.75 | 1897466.25 | 2 | 730055.75 | 2 | 2343053 | 1226192 | 2 | 2788373.5 | 2799271.5 | 2528529 | 647823.69 | 426758.75 | 802642.69 | 2 | 2 | 926353 |
| Q5BK07 | Ccdc43 | Coiled-coil domain-containing protein 43 | 36.53 | 4.05 | 25061 | 1 | 1482455.88 | 1305186.5 | 1784395 | 1051406.25 | 257336.77 | 2 | 664296.06 | 425242.25 | 1029527.31 | 734621 | 1125230.5 | 1401449.62 | 1289341.62 | 1686817.12 | 1278412 | 765058 | 2 | 1767404.25 | 914037.19 | 928429.81 |
| Q5U2X6 | Ccdc47 | PAT complex subunit CCDC47 | 221.74 | 10.14 | 55733 | 5 | 909126.62 | 832468.19 | 171323.95 | 531475.06 | 884316.81 | 2 | 821839.25 | 2 | 528263.75 | 345788.28 | 2 | 403047.19 | 797971.44 | 390353.34 | 2 | 2 | 2 | 2 | 2 | 2 |
| Q5XIB5 | Ccdc86 | Coiled-coil domain-containing protein 86 | 155.48 | 9.09 | 38585 | 3 | 2 | 2 | 2 | 2 | 2 | 2 | 2 | 2 | 2 | 2 | 2 | 778589.56 | 2 | 2 | 2 | 578017.69 | 2 | 2 | 2 | 2 |
| Q8VC31 | Ccdc9 | Coiled-coil domain-containing protein 9 | 39.77 | 1.29 | 61445 | 1 | 2 | 2 | 2 | 2 | 2 | 2 | 2 | 2 | 2 | 2 | 229635.08 | 2 | 2 | 2 | 2 | 2 | 2 | 2 | 2 | 2 |
| P80314 | Cct2 | T-complex protein 1 subunit beta | 1371.55 | 54.95 | 57477 | 22 | 17484746 | 26218186 | 29614788 | 22148408 | 28373718 | 23819830 | 25087054 | 16678305 | 29117518 | 23540690 | 34388082 | 42108058 | 36619170 | 42793866 | 34957058 | 26175418 | 29551102 | 30537646 | 18817074 | 30194774 |
| P80318 | Cct3 | T-complex protein 1 subunit gamma | 1716.17 | 48.99 | 60630 | 29 | 66344106 | 97237906 | 66587698 | 86394674 | 90671194 | 69727362 | 58835178 | 68214314 | 64070782 | 58366122 | 67145322 | 62053666 | 63675350 | 81119754 | 66888022 | 47032462 | 58482254 | 70419626 | 47495442 | 60043898 |
| P80315 | Cct4 | T-complex protein 1 subunit delta | 1593.52 | 58.63 | 58066 | 27 | 91069810 | 82091754 | 77193618 | 78976786 | 68691130 | 68406170 | 60557294 | 46598494 | 72998562 | 73902290 | 63087982 | 64560522 | 75206482 | 70540538 | 71134722 | 64484190 | 75444570 | 85292042 | 74495482 | 71927938 |
| P80316 | Cct5 | T-complex protein 1 subunit epsilon | 974.92 | 26.62 | 59624 | 14 | 29297386 | 38452862 | 27944014 | 43502854 | 36790206 | 35202594 | 36337890 | 29997378 | 30209028 | 28858722 | 26075702 | 29853908 | 28940472 | 31086478 | 32677602 | 29186024 | 29350458 | 25998950 | 31530438 | 32507142 |
| P80317 | Cct6a | T-complex protein 1 subunit zeta | 1122.09 | 38.79 | 58004 | 21 | 19999806 | 12421649 | 20511824 | 24611014 | 22871182 | 15782502 | 20979636 | 22610578 | 20913246 | 18462184 | 20267894 | 23888950 | 27360376 | 25058048 | 31391298 | 25295708 | 27180668 | 34722510 | 23671688 | 28982024 |
| P80313 | Cct7 | T-complex protein 1 subunit eta | 1455.54 | 54.41 | 59652 | 26 | 37496190 | 48713858 | 39065046 | 51724586 | 55765450 | 43689794 | 47472762 | 50487346 | 41520718 | 39513906 | 31579980 | 37564686 | 36647198 | 38271866 | 27922018 | 35419078 | 26713512 | 34341098 | 34661314 | 41800722 |
| P42932 | Cct8 | T-complex protein 1 subunit theta | 1255.53 | 40.15 | 59555 | 23 | 65590282 | 81012210 | 73498890 | 61645954 | 75480818 | 58934134 | 51355138 | 66749206 | 50582990 | 52395242 | 54396142 | 51981442 | 51978014 | 54507254 | 51336910 | 45046614 | 38367490 | 52167938 | 49501530 | 49083430 |
| P10810 | Cd14 | Monocyte differentiation antigen CD14 | 706.03 | 34.7 | 39204 | 11 | 10942571 | 10061821 | 15048407 | 11770358 | 10768785 | 6218570 | 7596580.5 | 8346176 | 6411814.5 | 14659994 | 1973751.75 | 1787679.5 | 1974995.75 | 3591940.5 | 3960309 | 1611017.12 | 2270443.5 | 2640530.25 | 1191127.88 | 2485951 |
| Q62192 | Cd180 | CD180 antigen | 191.11 | 10.14 | 74302 | 4 | 69080.31 | 511586.88 | 1441044.5 | 2 | 2 | 2 | 867188 | 1808448.12 | 185391.73 | 872503 | 2 | 2 | 72187.95 | 94746.44 | 2 | 2 | 212177.36 | 2 | 93984.89 | 199994.28 |
| Q9JLQ0 | Cd2ap | CD2-associated protein | 65.88 | 2.04 | 70450 | 1 | 2 | 2 | 2 | 2 | 102048.27 | 73780.06 | 2 | 88566.72 | 241962.91 | 85625.08 | 44772.95 | 2 | 127520.14 | 2 | 154264.75 | 2 | 154555.22 | 2 | 2 | 2 |
| P15379 | Cd44 | CD44 antigen | 111.76 | 4.88 | 85617 | 2 | 17653470 | 19472118 | 14617712 | 11250241 | 10323601 | 5291979.5 | 15385374 | 16759404 | 15350580 | 11338676 | 16319432 | 1241387.75 | 13994509 | 15466909 | 10715787 | 25429046 | 22013830 | 19005000 | 5357116 | 21476132 |
| P13379 | Cd5 | T-cell surface glycoprotein CD5 | 278.94 | 12.15 | 53849 | 4 | 1926688.25 | 1240782.25 | 820710.69 | 1731334.75 | 3880314 | 2 | 2 | 2 | 1801055.38 | 1791979.5 | 2 | 2 | 2 | 2 | 2 | 2 | 473087.78 | 2 | 2 | 2 |
| P31996 | Cd68 | Macrosialin | 117.89 | 8.9 | 34818 | 3 | 165959250 | 207126978 | 138495938 | 95581930 | 82793746 | 124288546 | 107422362 | 147383826 | 98917250 | 82176634 | 80425250 | 76476426 | 74132970 | 77241010 | 63619786 | 120026946 | 131139322 | 108484226 | 3420141.25 | 105867514 |
| P21855 | Cd72 | B-cell differentiation antigen CD72 | 101.34 | 3.67 | 40347 | 2 | 2 | 2 | 870634.69 | 2 | 548804.25 | 1391849.62 | 2 | 1202545.62 | 593944.94 | 2 | 3400129.5 | 924282.88 | 1011238.31 | 2 | 616115.19 | 1058070 | 1160945.12 | 2 | 1735224.38 | 2918023.25 |
| P40237 | Cd82 | CD82 antigen | 49.91 | 4.14 | 29629 | 1 | 2 | 2 | 2 | 2 | 768567.06 | 2 | 2 | 2 | 2 | 663067.88 | 2 | 2 | 499663.62 | 2 | 2 | 2 | 893503.62 | 2 | 2 | 2 |
| P40240 | Cd9 | CD9 antigen | 77.06 | 11.06 | 25258 | 1 | 2 | 2 | 2 | 2 | 2 | 2 | 2 | 2 | 144116.47 | 647923.75 | 84141.55 | 2 | 2 | 2 | 2 | 2 | 2 | 218442.92 | 2 | 2 |
| O89103 | Cd93 | Complement component C1q receptor | 38.73 | 1.71 | 69354 | 1 | 2 | 2 | 2 | 2 | 2 | 2 | 2 | 2 | 2 | 2 | 2 | 2 | 2 | 2 | 2 | 2 | 2 | 2 | 2 | 2 |
| Q61081 | Cdc37 | Hsp90 co-chaperone Cdc37 | 170.61 | 12.93 | 44593 | 4 | 845159.25 | 222068.08 | 715776.12 | 868678.31 | 1865623.5 | 2 | 508371.69 | 1101593.88 | 591183.81 | 62151.34 | 785089.44 | 1003428.06 | 1520544 | 295437.03 | 944693 | 1286293.75 | 709945.19 | 163847.92 | 1262627.88 | 1060390.25 |
| P60766 | Cdc42 | Cell division control protein 42 homolog | 397.38 | 52.88 | 21259 | 7 | 3311589 | 9642758 | 2980467 | 9994476 | 6508458.5 | 4819906.5 | 9451364 | 4506138 | 11199434 | 7563479 | 7283191 | 12266770 | 11891869 | 12841441 | 12267923 | 11389411 | 14098203 | 6780438.5 | 4888620 | 11313770 |
| O08837 | Cdc5l | Cell division cycle 5-like protein | 346.24 | 9.6 | 92218 | 6 | 2511668 | 3042240.75 | 1703384 | 3299077.5 | 3472249 | 1828458 | 1792628.5 | 1970645.38 | 3201020 | 1705721.5 | 2300372.25 | 1941310.62 | 1880023.75 | 2301621.75 | 1497935 | 2125652.25 | 2277406 | 2850383.25 | 2692081.25 | 3289145 |
| Q8VDP6 | Cdipt | CDP-diacylglycerol--inositol 3-phosphatidyltransfe | 224.95 | 29.58 | 23599 | 5 | 811432.38 | 1736875.12 | 181852.84 | 2 | 235495.7 | 3642838.5 | 2299277 | 269238.75 | 4303924.5 | 2586288 | 2215709.75 | 147121.11 | 3322537.5 | 3475057.75 | 2991529 | 2138274 | 3933958 | 4637726.5 | 2483439.25 | 2760026.25 |
| Q64261 | Cdk6 | Cyclin-dependent kinase 6 | 81.56 | 6.75 | 37028 | 2 | 2 | 2 | 2 | 2 | 2 | 2 | 2 | 2 | 2 | 2 | 2 | 890122.06 | 889137.31 | 1085941.25 | 2 | 926664.38 | 2 | 2 | 2 | 486116.06 |
| Q8BI72 | Cdkn2aip | CDKN2A-interacting protein | 80.55 | 4.62 | 59745 | 2 | 2 | 2 | 2 | 2 | 33651.82 | 2 | 2 | 2 | 491074.66 | 2 | 2 | 369982.66 | 2 | 2 | 2 | 2 | 2 | 2 | 2 | 2 |
| Q5RK03 | Cdkn2aipnl | CDKN2AIP N-terminal-like protein | 39.25 | 8.62 | 13197 | 1 | 946666 | 1012831.94 | 1228672.62 | 567696.88 | 856297.94 | 681362.62 | 773188.19 | 1137079.75 | 476785.09 | 761563.44 | 692817.31 | 838705.12 | 918902.5 | 571811.62 | 1235595 | 869396.88 | 688560.69 | 1036078.94 | 2 | 823710.94 |
| Q99L43 | Cds2 | Phosphatidate cytidylyltransferase 2 | 74.11 | 4.28 | 51314 | 1 | 2 | 2 | 2 | 2 | 2 | 2 | 2 | 1062724.75 | 819748.06 | 589795.19 | 2 | 2 | 2 | 2 | 2 | 2 | 2 | 2 | 2 | 2 |
| Q4VAA2 | Cdv3 | Protein CDV3 | 53.5 | 10.68 | 29729 | 1 | 2 | 1712873 | 2 | 2219471.5 | 384129.78 | 2968753 | 1443390.25 | 498945.81 | 1665730.62 | 1206063.5 | 3441231.5 | 2800357.75 | 1721035.5 | 2730475.75 | 2461540.5 | 1078548.5 | 1761052.62 | 1371977.62 | 2941863.5 | 961789.81 |
| Q792H5 | Celf2 | CUGBP Elav-like family member 2 | 106.54 | 5.31 | 54271 | 2 | 2 | 1974108.88 | 1893660.38 | 1919124.12 | 3012231.75 | 1425361.5 | 760636.81 | 1283874.5 | 660643 | 1052506.88 | 890880.88 | 1416583.62 | 1266816.62 | 973876.19 | 1253296.25 | 1050261.5 | 1454344.88 | 1447670.88 | 1454315.38 | 1635111 |
| Q924Z4 | Cers2 | Ceramide synthase 2 | 45.93 | 2.37 | 45024 | 1 | 2 | 2 | 2 | 553219.19 | 320759.75 | 2 | 2 | 862072.94 | 1014169.69 | 546164.38 | 443347.75 | 2 | 381799.25 | 759043.12 | 2 | 462624.66 | 1160628.38 | 2 | 1342733 | 2 |
| Q9D6J1 | Cers4 | Ceramide synthase 4 | 34.31 | 2.04 | 46017 | 1 | 67388.45 | 810709.44 | 2 | 351906.12 | 2 | 1153934.5 | 139888.66 | 2 | 2086187.62 | 2 | 141253.31 | 2 | 2 | 1214665.38 | 2 | 1689468.62 | 1646471.5 | 2 | 2 | 1383018.75 |
| B2RW38 | Cfap58 | Cilia- and flagella-associated protein 58 | 38.86 | 1.03 | 103524 | 1 | 2 | 877051 | 2 | 891536 | 85920.46 | 2 | 922492.31 | 2 | 709786 | 2 | 4759673 | 2 | 580996.12 | 3876378.25 | 2 | 6792919.5 | 2 | 2450950 | 1839367.75 | 2461385 |
| P18760 | Cfl1 | Cofilin-1 | 674.92 | 74.1 | 18560 | 10 | 15788739 | 14159229 | 26441510 | 19707402 | 53186534 | 31805398 | 21150924 | 40351130 | 66201578 | 44425730 | 118301770 | 108682354 | 71478322 | 67099610 | 67825754 | 116420698 | 70246474 | 78065674 | 59094290 | 73162114 |
| Q9D1L0 | Chchd2 | Coiled-coil-helix-coiled-coil-helix domain-containi | 69.34 | 15.69 | 15661 | 1 | 2 | 2 | 2 | 2 | 635389.25 | 2 | 2 | 2 | 1834031.88 | 497813.38 | 3066688 | 1898014.75 | 611997.06 | 1354658.75 | 2 | 763511.56 | 2 | 2 | 960059.12 | 2 |
| Q9CRB9 | Chchd3 | MICOS complex subunit Mic19 | 145.24 | 7.49 | 26335 | 2 | 757008.56 | 2 | 2 | 2 | 2469580.5 | 529840.06 | 2 | 2 | 821619.19 | 1906321.88 | 4496131 | 5620043 | 2415838.75 | 3734703 | 1129131.38 | 5602538 | 2721476.25 | 1046669.44 | 3518311.25 | 1537564.12 |
| D4A7N1 | Chchd6 | MICOS complex subunit Mic25 | 104.05 | 8.81 | 29211 | 2 | 1812383.62 | 2 | 2256170.25 | 255264.31 | 2 | 2170651.5 | 2 | 2 | 2794185.5 | 2 | 2 | 2 | 2 | 3537943.25 | 2 | 1734246.25 | 580543.12 | 1660560 | 3348455 | 1582975.38 |
| Q6PDQ2 | Chd4 | Chromodomain-helicase-DNA-binding protein 4 | 321.53 | 4.49 | 217751 | 8 | 2056993.75 | 2052732.75 | 2779898 | 3695542.5 | 6919680.5 | 2492817 | 2996170.25 | 3419165.5 | 4804269.5 | 3809689.5 | 4182584 | 4070458.5 | 3757171.5 | 1247148.25 | 2730645.5 | 3318058.25 | 1230369.75 | 2843932.25 | 1077230.12 | 1340209.12 |
| Q8CGZ0 | Cherp | Calcium homeostasis endoplasmic reticulum prot | 41.43 | 1.07 | 106169 | 1 | 2 | 2 | 832860.88 | 2 | 798004.62 | 2 | 2 | 2 | 2 | 2 | 2 | 2 | 914184.69 | 2 | 2 | 2 | 2 | 2 | 2 | 2 |
| Q9D8B3 | Chmp4b | Charged multivesicular body protein 4b | 164.67 | 14.73 | 24936 | 3 | 1597388.12 | 2477920.25 | 2048380.5 | 1730587.75 | 4172110.75 | 3856480 | 4818101 | 137097.73 | 2176711 | 2833244.25 | 4628973 | 5113061 | 4478849 | 4513796 | 4442939 | 6408556 | 2395529 | 3169040 | 6794688 | 6147525 |
| Q9CY57 | Chtop | Chromatin target of PRMT1 protein | 84.15 | 5.22 | 26585 | 1 | 2 | 2 | 2 | 2 | 2 | 2 | 2 | 2 | 2 | 2 | 2 | 2 | 2 | 2 | 2 | 2 | 2 | 2 | 2 | 2 |
| Q9DCL2 | Ciao2a | Cytosolic iron-sulfur assembly component 2A | 42 | 5.62 | 18418 | 1 | 732170.88 | 114519.24 | 928729.5 | 267938.38 | 572224.75 | 67372.38 | 2 | 170072.39 | 2 | 574357.56 | 2 | 2 | 2 | 451500.47 | 497050.34 | 2 | 2 | 266625.38 | 2 | 341262 |
| Q8WTY4 | Ciapin1 | Anamorsin | 116.81 | 8.74 | 33429 | 2 | 871341.88 | 1813822 | 744100.31 | 2628978 | 1930680 | 595794.75 | 1030829.56 | 2082379.5 | 1303724.12 | 1030217.38 | 1918704.75 | 2508809.5 | 2038417 | 1716986.88 | 1644918.75 | 1857273.25 | 1691048.5 | 1468785.62 | 2254294.5 | 1528975.75 |
| Q9CQB5 | Cisd2 | CDGSH iron-sulfur domain-containing protein 2 | 82.12 | 20 | 15242 | 2 | 2 | 166791.09 | 2 | 316151.62 | 1268334.5 | 2 | 269513.12 | 2 | 788509.31 | 2 | 2 | 741994.38 | 822327.12 | 629320.56 | 628401.06 | 2 | 365191.19 | 2 | 2 | 2 |
| Q8BMK4 | Ckap4 | Cytoskeleton-associated protein 4 | 897.48 | 30.26 | 63692 | 14 | 2045079.12 | 1276170.88 | 2805130.25 | 4041542.75 | 4977199 | 183101.31 | 3188411.25 | 4098052 | 4076896.75 | 7389311.5 | 391244.62 | 1179653.5 | 1516944 | 979984.5 | 1517138.38 | 2 | 2 | 1215795.38 | 2 | 753378.94 |
| A2AGT5 | Ckap5 | Cytoskeleton-associated protein 5 | 244.93 | 3.2 | 225635 | 5 | 2 | 405563.78 | 44594.29 | 1180675.5 | 208489.14 | 2 | 656123.94 | 154503.33 | 1954642.12 | 93655.49 | 781723.56 | 1639799.25 | 2 | 481229 | 144371.23 | 2 | 350555.59 | 2 | 2 | 2 |
| Q04447 | Ckb | Creatine kinase B-type | 945.91 | 40.42 | 42713 | 13 | 57016830 | 72079594 | 47728022 | 47361862 | 46051482 | 43721730 | 50847670 | 39551214 | 49061258 | 50364234 | 62532250 | 59027898 | 74543522 | 73024210 | 65546350 | 72081626 | 70585530 | 79249106 | 24684706 | 77560458 |
| P00564 | Ckm | Creatine kinase M-type | 34.52 | 8.4 | 43045 | 1 | 2 | 2 | 108148.75 | 2 | 120064.53 | 2 | 182938.78 | 2 | 225838.83 | 144456.66 | 144608.62 | 2 | 2 | 2 | 2 | 2 | 2 | 2 | 279714.72 | 2 |
| Q8BRT1 | Clasp2 | CLIP-associating protein 2 | 114.8 | 1.79 | 140739 | 3 | 702967.44 | 1062777.38 | 647279.88 | 502298.75 | 780302.75 | 2 | 142333.39 | 62377.88 | 142578.33 | 2 | 2 | 2 | 2 | 237896.53 | 635456.62 | 568163.38 | 2 | 737242.12 | 2 | 670829.88 |
| P49300 | Clec10a | C-type lectin domain family 10 member A | 308.24 | 23.36 | 34596 | 6 | 5976716.5 | 1849346.75 | 7151406 | 3444021.75 | 3082026.5 | 4734295.5 | 1419452 | 812038.25 | 4667197 | 4163329.25 | 7551285 | 10072427 | 9855357 | 7629128 | 5815012.5 | 9086895 | 7537267 | 7112551.5 | 7413034 | 6110689 |
| Q9R0Q8 | Clec4e | C-type lectin domain family 4 member E | 47.26 | 6.07 | 24431 | 1 | 2 | 2 | 2 | 2 | 2 | 2 | 2 | 2 | 741089.44 | 2 | 2 | 2 | 2 | 2 | 2 | 2 | 2 | 2 | 2 | 2 |
| Q6MG61 | Clic1 | Chloride intracellular channel protein 1 | 974.04 | 65.56 | 26981 | 16 | 26968158 | 18304856 | 37844354 | 28932872 | 35251702 | 23901558 | 25802932 | 39489206 | 35451110 | 46647030 | 39502898 | 39852202 | 46347422 | 45278302 | 47106122 | 28934484 | 34945310 | 28225012 | 32661868 | 28450174 |
| Q9QYB1 | Clic4 | Chloride intracellular channel protein 4 | 485.48 | 41.9 | 28729 | 9 | 19695930 | 9273755 | 32826244 | 23145780 | 14716832 | 7359122 | 18941076 | 10768393 | 5155896 | 20794724 | 13639570 | 15342562 | 18691294 | 18547624 | 32383716 | 4265395 | 11416401 | 19781366 | 6708831.5 | 8531934 |
| Q922J3 | Clip1 | CAP-Gly domain-containing linker protein 1 | 43.94 | 0.79 | 155814 | 1 | 2 | 706703.19 | 66144.17 | 909883.94 | 106602.25 | 2 | 2 | 2 | 668490.31 | 2 | 772545.94 | 1236496 | 1081512.38 | 717420.81 | 2 | 2 | 2 | 2 | 2 | 2 |
| Q04753 | Clns1a | Methylosome subunit pICln | 52.84 | 5.51 | 26092 | 1 | 834791.12 | 1183008.38 | 637328.44 | 1173457.62 | 988731.31 | 2 | 628352.81 | 2 | 557902.5 | 916508.94 | 517390.38 | 2 | 898115.69 | 484148.34 | 2 | 2 | 2 | 762540.38 | 2 | 2 |
| Q8VBZ3 | Clptm1 | Putative lipid scramblase CLPTM1 | 296.51 | 12.05 | 75291 | 5 | 1385288.75 | 2203916.5 | 771805.5 | 3569884.75 | 1969743.5 | 2290605.25 | 2470595.75 | 2718709.75 | 3852861.5 | 2740750.75 | 2183536.5 | 1983483.5 | 2975670.5 | 2507126.5 | 3116336.75 | 2421306.5 | 2928435.5 | 4579359.5 | 5038727.5 | 3635243.5 |
| Q5U2U0 | Clpx | ATP-dependent clpX-like chaperone, mitochondri | 107.87 | 5.21 | 69208 | 3 | 1263421.62 | 1593959.12 | 1282999.88 | 918913.88 | 1201790.62 | 2 | 1241799.25 | 2 | 611453.06 | 726342.38 | 2 | 811907.25 | 1098744.25 | 893415.31 | 2 | 659967.94 | 2 | 2 | 699500.69 | 2 |
| O08585 | Clta | Clathrin light chain A | 145.32 | 15.32 | 25604 | 4 | 7078645 | 4337204 | 9287254 | 10751168 | 7173574 | 6786876 | 5897981.5 | 10337619 | 7844306 | 7909092.5 | 7073943.5 | 5468767 | 6739486 | 2904535.75 | 5117806 | 9042660 | 6967282 | 4964143 | 6589053 | 5738109 |
| P08082 | Cltb | Clathrin light chain B | 41.19 | 3.93 | 25117 | 1 | 1630371.5 | 2 | 2562463.75 | 1754806.5 | 3389426.25 | 2 | 2 | 2 | 1193048.62 | 1807547.25 | 3919720.5 | 3777096 | 578854.94 | 2 | 1441953.12 | 4694506.5 | 3504893.25 | 2137469.25 | 4091038.5 | 2178131.75 |
| Q68FD5 | Cltc | Clathrin heavy chain 1 | 3318.36 | 37.73 | 191557 | 52 | 52039770 | 42788902 | 56770214 | 52075530 | 61955198 | 41645902 | 47704302 | 46006426 | 54717390 | 61028994 | 46237766 | 46263614 | 54939618 | 49549350 | 65121350 | 40975466 | 46805358 | 56138958 | 41429610 | 45722014 |
| Q8R4N0 | Clybl | Citramalyl-CoA lyase, mitochondrial | 161.21 | 16.27 | 37549 | 3 | 129069.1 | 288384.44 | 165949.36 | 235270.31 | 124284.28 | 276275.78 | 360688 | 121069.62 | 257675.73 | 193703.5 | 214392.25 | 168348.08 | 55044.22 | 216480.77 | 273410.28 | 35121.78 | 250904.73 | 214899.98 | 71981.32 | 263923.5 |
| Q99KK2 | Cmas | N-acylneuraminate cytidylyltransferase | 177.62 | 12.04 | 48058 | 4 | 3873425 | 7227545 | 5650022 | 3021965.25 | 4084372 | 6384452 | 5653673.5 | 4020227.25 | 5985851.5 | 5922819 | 4700457 | 4221574 | 6824387 | 6841621 | 4916592 | 5123299.5 | 6782083 | 7866435.5 | 7388671.5 | 6250206 |
| Q4KM73 | Cmpk1 | UMP-CMP kinase | 175.47 | 25.51 | 22169 | 4 | 2 | 2 | 2 | 78223.92 | 720241.81 | 2 | 254394.02 | 228135.98 | 877185.12 | 845820.62 | 475871.12 | 1205477.12 | 1268929.75 | 1454895 | 1478276.25 | 244149.39 | 506205.94 | 155628 | 161835.38 | 501818.44 |
| Q3U5Q7 | Cmpk2 | UMP-CMP kinase 2, mitochondrial | 372.89 | 16.78 | 50036 | 7 | 1301941.75 | 110944.92 | 1007595.31 | 2 | 383668.5 | 4200025 | 6962541 | 3868402.5 | 3923066.25 | 4164663.5 | 709206.31 | 581709 | 1453868 | 416213.47 | 2 | 1451886.75 | 1322394.5 | 2197981.5 | 1498587.75 | 560031.88 |
| P53996 | Cnbp | CCHC-type zinc finger nucleic acid binding protei | 124.01 | 12.92 | 19592 | 2 | 2 | 349616.12 | 2 | 1061163.38 | 2357715.5 | 2 | 90130.98 | 186090.42 | 3124323.5 | 2 | 343610.25 | 9185783 | 3735105 | 6102546.5 | 4144444 | 2058872.5 | 2547602.75 | 239675.92 | 2 | 3784328.25 |
| Q9D1A2 | Cndp2 | Cytosolic non-specific dipeptidase | 755.17 | 29.89 | 52767 | 14 | 6382046.5 | 12716889 | 2406178.5 | 6861020 | 8545615 | 10674824 | 8897658 | 4956182.5 | 13585741 | 14416269 | 14913895 | 16108140 | 11409611 | 12398761 | 10696110 | 7022303 | 13365822 | 14267500 | 9372979 | 15346324 |
| Q9CX13 | Cnih4 | Protein cornichon homolog 4 | 52.68 | 14.39 | 16089 | 1 | 2 | 2 | 2 | 648318.94 | 845318.44 | 2 | 777316.31 | 1758350.25 | 826558.38 | 865881.31 | 2 | 2 | 2 | 2 | 2 | 2 | 2 | 897346.19 | 329691.78 | 2 |
| Q9QXT0 | Cnpy2 | Protein canopy homolog 2 | 83.99 | 8.79 | 20767 | 1 | 2061376.12 | 1992621.25 | 1703973.5 | 2076097 | 1623956.38 | 1323680.75 | 1469686.38 | 2620545.75 | 1204640.5 | 2 | 1639381.12 | 1550820.88 | 1499650.62 | 1580990.25 | 1199752.75 | 2695128.25 | 1791855 | 3256960 | 2901415 | 2702003 |
| Q9DAU1 | Cnpy3 | Protein canopy homolog 3 | 44.65 | 3.99 | 30538 | 1 | 1871318.88 | 1140381.25 | 1410310.88 | 1925868.25 | 1237352.38 | 970706.81 | 483912.25 | 910698 | 665417.5 | 624595.81 | 1629011.12 | 866482 | 1616993.88 | 850250.5 | 2102237.25 | 2045644.38 | 1906464.62 | 2136629.25 | 1598829.88 | 2199510.25 |
| Q9DBL7 | Coasy | Bifunctional coenzyme A synthase | 74.49 | 4.97 | 62023 | 2 | 2 | 2 | 129622.33 | 191356.17 | 52311.06 | 2 | 276016.84 | 2 | 213791.58 | 267120.62 | 2 | 2 | 2 | 279602.25 | 246069.91 | 2 | 93161.65 | 2 | 2 | 2 |
| Q8K4Q8 | Colec12 | Collectin-12 | 377.28 | 10.92 | 81304 | 7 | 5276827 | 5270448.5 | 3842563.5 | 2000265.75 | 1258247 | 5135617 | 5263651 | 2414745.5 | 3671383 | 4958589 | 1926394.75 | 3975187.25 | 4834398 | 5384093.5 | 5596976.5 | 3668358 | 3694810.5 | 3236687.5 | 3520010.5 | 1300697.12 |
| Q8K297 | Colgalt1 | Procollagen galactosyltransferase 1 | 478.56 | 17.34 | 71061 | 10 | 3446689.5 | 2142996.5 | 1756280 | 3572467.5 | 4056921.25 | 976107.81 | 2390221.75 | 1038437.06 | 2562549.75 | 1670730.38 | 2313414.25 | 1546232 | 1900175 | 3655418.75 | 2401069 | 2188510 | 1022942.38 | 919965.31 | 1894138.62 | 672400 |
| Q8BXC6 | Commd2 | COMM domain-containing protein 2 | 48.82 | 3.52 | 22848 | 1 | 715110.56 | 2 | 1085068.12 | 1058754.38 | 526906.81 | 2 | 613236.12 | 495310 | 2 | 528414.81 | 471272.88 | 2 | 2 | 2 | 1097097.25 | 2 | 2 | 1932410.5 | 2 | 3055247.5 |
| O88587 | Comt | Catechol O-methyltransferase | 90.49 | 12.08 | 29486 | 2 | 53179.45 | 2 | 165769.89 | 2 | 137782.53 | 2 | 2 | 136826.61 | 2 | 309277.16 | 128755.41 | 237570.7 | 188411.55 | 2 | 492327.59 | 2 | 2 | 2 | 58956.62 | 216771.02 |
| Q8BIG7 | Comtd1 | Catechol O-methyltransferase domain-containing | 249.57 | 29.01 | 28961 | 4 | 885121.19 | 749804.56 | 1262853.5 | 692188.12 | 1466603.88 | 1051744.5 | 1064000.5 | 57168.59 | 1240829.25 | 1649564.5 | 692593.5 | 657157.62 | 956760.81 | 638682.75 | 1737067.88 | 2 | 490647.53 | 1071415.12 | 2 | 129438.45 |
| Q8CIE6 | Copa | Coatomer subunit alpha | 590.18 | 11.44 | 138432 | 12 | 9957043 | 11224716 | 12177465 | 13627816 | 20162010 | 10711556 | 13496043 | 15140798 | 14169226 | 16696027 | 9644448 | 8538228 | 12410765 | 13524690 | 10359634 | 8661436 | 10247962 | 5128620 | 6894836 | 5790050.5 |
| P23514 | Copb1 | Coatomer subunit beta | 332.83 | 9.55 | 107011 | 6 | 4289806.5 | 8049958 | 6051631 | 10380670 | 10505427 | 6166092 | 7641422 | 8017075 | 5384940.5 | 5908088 | 4336440 | 3990363 | 5847971 | 4576215 | 7673610 | 4934612 | 4457491 | 5558401.5 | 5115665 | 9906241 |
| O35142 | Copb2 | Coatomer subunit beta' | 630.18 | 14.03 | 102551 | 11 | 10128057 | 14578192 | 15414304 | 14487480 | 15957242 | 16669212 | 15250814 | 21487808 | 13752553 | 12137928 | 14882119 | 15106767 | 16055719 | 15177573 | 16557099 | 16408677 | 14234876 | 15852869 | 18272416 | 10104255 |

| | | | | | | | | | | | | | | | | | | | | | | | | | | |
|---|---|---|---|---|---|---|---|---|---|---|---|---|---|---|---|---|---|---|---|---|---|---|---|---|---|---|
| O89079 | Cope | Coatomer subunit epsilon | 197.67 | 16.56 | 34567 | 4 | 463280.31 | 4973848.5 | 856377.25 | 3836481 | 98250.09 | 1047811.81 | 3175708.5 | 1020277.62 | 2531526.75 | 1395998.5 | 2188892.25 | 2859701 | 4078439 | 2697241.5 | 3207155.25 | 2930332.5 | 3524882.75 | 4575811.5 | 59721.71 | 3622909.5 |
| Q4AEF8 | Copg1 | Coatomer subunit gamma-1 | 476.55 | 15.22 | 97614 | 10 | 1605749.75 | 1161831 | 2369897 | 1543197.88 | 3551608.75 | 2210841 | 2000021 | 4488575.5 | 2894100.75 | 2453982 | 4180687.75 | 3757736 | 2721097.5 | 3095165.25 | 1818870.12 | 4256860 | 2260269 | 3212452 | 4620534 | 2706642.5 |
| Q9QXK3 | Copg2 | Coatomer subunit gamma-2 | 656.14 | 20.21 | 97681 | 14 | 4311171.5 | 1014506.19 | 2658883 | 4211877 | 5610105.5 | 2886373.75 | 2347811.5 | 2583550.75 | 5055638.5 | 5137048.5 | 5504077 | 6057466.5 | 7519365.5 | 8650993 | 2341388.25 | 6013966 | 5879646 | 3653367.5 | 6868010 | 5853466 |
| O88543 | Cops3 | COP9 signalosome complex subunit 3 | 196.16 | 9.46 | 47832 | 3 | 2440046.75 | 1931010 | 3329929 | 2440862.5 | 4046407 | 1023144.12 | 2803803.25 | 784634 | 3100308.25 | 2501637 | 244537.41 | 1209276.62 | 1678234.25 | 2505780.25 | 2518451.25 | 2 | 535721.62 | 757829.06 | 355072.88 | 1023719.5 |
| O88544 | Cops4 | COP9 signalosome complex subunit 4 | 203.13 | 13.3 | 46285 | 4 | 683027.62 | 2 | 852563.25 | 933452.25 | 2 | 1032318.5 | 2 | 643484.62 | 2 | 952935.5 | 653765.62 | 568514.75 | 749171.75 | 2 | 700151.56 | 649791.94 | 514381.97 | 608499.81 | 707995.44 | 2 |
| O35864 | Cops5 | COP9 signalosome complex subunit 5 | 122.66 | 8.68 | 37549 | 3 | 4030863.25 | 4898409.5 | 3202776.5 | 2447410.5 | 1245726.75 | 1240920.38 | 1060082.12 | 2 | 1879803.5 | 2950018.25 | 1701907.25 | 1593317.5 | 1201549.38 | 978195.88 | 1175422.88 | 1931377.88 | 2 | 926072.56 | 340138.69 | 864711.12 |
| Q9CZ04 | Cops7a | COP9 signalosome complex subunit 7a | 74.93 | 5.09 | 30224 | 1 | 2 | 2 | 2 | 2 | 2 | 2 | 2 | 753280.44 | 895380.81 | 2 | 2 | 2 | 2 | 2 | 2 | 2 | 2 | 2 | 2 | 2 |
| Q8VBV7 | Cops8 | COP9 signalosome complex subunit 8 | 83.01 | 6.22 | 23256 | 1 | 2 | 2 | 1347400.12 | 2699991.25 | 2771469.25 | 1733489.75 | 2 | 2578664.75 | 1817896.25 | 1932161.38 | 2 | 1397416.88 | 1406596 | 1484994.25 | 939408.19 | 531714.94 | 966808.31 | 1952001.38 | 2087673 | 1459512 |
| P61924 | Copz1 | Coatomer subunit zeta-1 | 55.67 | 7.91 | 20198 | 1 | 2 | 83204.16 | 307026.72 | 780900.19 | 409294.31 | 2 | 2 | 2 | 561587.75 | 471005.06 | 258319.23 | 2 | 2 | 284971.5 | 2 | 72732.75 | 2 | 527005.75 | 2 | 396242.59 |
| Q566J8 | Coq8b | Atypical kinase COQ8B, mitochondrial | 60.47 | 4.13 | 59232 | 1 | 2 | 2 | 2 | 61702.93 | 30678.04 | 2 | 95516.27 | 2 | 90226.48 | 2 | 2 | 2 | 2 | 2 | 2 | 2 | 2 | 2 | 2 | 2 |
| O89053 | Coro1a | Coronin-1A | 1041.71 | 45.99 | 50989 | 19 | 22050712 | 15196892 | 26321714 | 34182190 | 71280666 | 39440066 | 32595166 | 52175746 | 100995586 | 50787218 | 92840818 | 84727258 | 48154950 | 51512678 | 54722070 | 50078426 | 34016918 | 29266274 | 40845486 | 28288204 |
| Q9WUM3 | Coro1b | Coronin-1B | 167.97 | 8.26 | 53912 | 4 | 2 | 2 | 1292125.12 | 876598.44 | 2298199 | 2206266 | 1121099.75 | 1830067.12 | 3917753 | 1591490.25 | 1525166.75 | 1762850 | 699480.38 | 1057308.38 | 1139035.62 | 1589792.25 | 2 | 2 | 817508.62 | 2 |
| Q9WUM4 | Coro1c | Coronin-1C | 505.53 | 21.31 | 53121 | 9 | 23816082 | 9301081 | 22466392 | 7706058 | 15700172 | 21137138 | 19503602 | 15508272 | 10631011 | 8650512 | 10640959 | 13485459 | 10214477 | 9695328 | 6841066 | 8834863 | 11623583 | 21145490 | 8412299 | 7825319 |
| Q9D2V7 | Coro7 | Coronin-7 | 234.06 | 5.53 | 100812 | 5 | 4420868.5 | 7630926 | 3522103.25 | 5555716.5 | 7855443 | 8445051 | 6736682 | 955939.88 | 6847301 | 5425578 | 2499342.25 | 3525526 | 3359746.5 | 6387728.5 | 3410937.75 | 4888637 | 3705301 | 3025751.25 | 7578852 | 2502569 |
| B0BNA5 | Cotl1 | Coactosin-like protein | 516.27 | 52.11 | 15932 | 9 | 37487218 | 28739548 | 43441658 | 33445276 | 27568018 | 25740864 | 30831406 | 46387942 | 22875328 | 26932342 | 29212108 | 40983942 | 35966766 | 36978218 | 60334270 | 30435038 | 41104034 | 50451034 | 29684566 | 49392054 |
| Q8BJ03 | Cox15 | Heme A synthase COX15 | 49.84 | 3.63 | 45853 | 1 | 2 | 2 | 194378.09 | 317368.38 | 2 | 237290.69 | 234121.84 | 2 | 352543.38 | 2 | 2 | 2 | 158650.89 | 2 | 790452.94 | 2 | 2 | 2 | 2 | 2 |
| P56394 | Cox17 | Cytochrome c oxidase copper chaperone | 73.75 | 25.4 | 6784 | 1 | 2 | 2 | 2 | 2 | 2 | 2 | 2 | 1645458.88 | 1377222 | 2 | 1487951.5 | 846729.44 | 641215.44 | 714065.62 | 2 | 2 | 2 | 2 | 2 | 2 |
| P19783 | Cox4i1 | Cytochrome c oxidase subunit 4 isoform 1, mitoch | 279.76 | 38.46 | 19530 | 6 | 23468890 | 20638818 | 24356822 | 16898870 | 26463962 | 17498326 | 14633492 | 20751230 | 24990894 | 19614238 | 18894994 | 18847298 | 26638646 | 19477798 | 17288550 | 15868822 | 20232414 | 26010396 | 22375658 | 20566546 |
| P11240 | Cox5a | Cytochrome c oxidase subunit 5A, mitochondrial | 289.11 | 27.4 | 16130 | 5 | 65235450 | 70873722 | 64491278 | 75663674 | 51594214 | 47910514 | 41954354 | 57179606 | 38602382 | 46892634 | 29205790 | 27138892 | 34655898 | 34196502 | 42438350 | 34523678 | 44207730 | 66395698 | 42596682 | 60365170 |
| P19536 | Cox5b | Cytochrome c oxidase subunit 5B, mitochondrial | 130.76 | 28.91 | 13813 | 3 | 16315440 | 24130888 | 14095708 | 25575058 | 22468246 | 36597558 | 32901140 | 43252610 | 37052070 | 19826180 | 25851700 | 23346860 | 18389206 | 21944142 | 12447065 | 11294373 | 22146450 | 21905494 | 18943784 | 24055350 |
| P10818 | Cox6a1 | Cytochrome c oxidase subunit 6A1, mitochondrial | 54.71 | 18.92 | 12301 | 1 | 2 | 2 | 2 | 2 | 2 | 2 | 2 | 2 | 2 | 2 | 2 | 2 | 55258.38 | 2 | 2 | 2 | 2 | 2 | 2 | 2 |
| P56391 | Cox6b1 | Cytochrome c oxidase subunit 6B1 | 240.41 | 50 | 10071 | 4 | 4305351 | 4282084.5 | 3941436.5 | 4241617 | 2150465.5 | 1820962.12 | 1739984.5 | 2265527.5 | 2294709 | 3133573.5 | 2 | 2 | 2 | 2 | 1949332.25 | 1294350.62 | 2 | 1680758 | 2238895.25 | 2 |
| P11951 | Cox6c2 | Cytochrome c oxidase subunit 6C-2 | 95.88 | 25 | 8455 | 2 | 2 | 899062.75 | 1442438.12 | 759225.81 | 3255556 | 1750657.38 | 1311954.12 | 2762695.25 | 5398744.5 | 1823900 | 2112891.75 | 2701211 | 2158762 | 2665382.25 | 2761117.25 | 2 | 2777524.25 | 2063424.38 | 2605933.25 | 2551206.75 |
| P48771 | Cox7a2 | Cytochrome c oxidase subunit 7A2, mitochondrial | 63.11 | 12.05 | 9291 | 1 | 4008257.25 | 4249798.5 | 2589541 | 4579443 | 5480141 | 3044305.75 | 2658512.25 | 3710399.75 | 2716044.75 | 4599937 | 1573091.25 | 1710618.75 | 1420000.12 | 1415482.12 | 1837799.38 | 2014566.88 | 1824436.12 | 1582048 | 1746673.75 | 1966623.75 |
| P17665 | Cox7c | Cytochrome c oxidase subunit 7C, mitochondrial | 93.59 | 50.79 | 7333 | 2 | 1013006.81 | 862747.12 | 1518389.25 | 1606879.88 | 953151.75 | 2 | 705430.94 | 855514.94 | 759492.12 | 1224121.75 | 2 | 453234.91 | 802563.44 | 680668.94 | 998831.81 | 34012.66 | 101052.56 | 947053.62 | 84409.65 | 95320.48 |
| O89001 | Cpd | Carboxypeptidase D | 86.01 | 1.45 | 152406 | 2 | 2152882.5 | 1313257.25 | 1658038.5 | 447445.75 | 635161.75 | 1296254 | 717120.56 | 1060724.5 | 582647.88 | 1175258.5 | 1015470.81 | 1004772.75 | 1082302.62 | 1412347.25 | 2 | 1307674.25 | 1707899.38 | 1234252 | 2 | 974433.12 |
| Q8C166 | Cpne1 | Copine-1 | 296.25 | 11.38 | 58887 | 5 | 2998588.5 | 9863400 | 4754930.5 | 5503755 | 2891441.5 | 2332574 | 5052227 | 1070665.25 | 6225223 | 3240091.75 | 9802791 | 17544012 | 9378475 | 13437070 | 5797147.5 | 10802596 | 7430160 | 5630725 | 6996298 | 10651492 |
| Q8BT60 | Cpne3 | Copine-3 | 120.47 | 6 | 59585 | 3 | 697019.25 | 680135.62 | 880225.19 | 882320.12 | 986491.25 | 298937.94 | 407248.5 | 184662.38 | 626823.62 | 1119291.5 | 235830.98 | 713199.25 | 248319.69 | 345309.31 | 1023066.44 | 2 | 328182.53 | 259579.23 | 2 | 257776.48 |
| Q9WVJ3 | Cpq | Carboxypeptidase Q | 247.46 | 12.98 | 51813 | 5 | 952888.88 | 1670381.75 | 1302459 | 1054140.5 | 1634993.75 | 2 | 145504.2 | 621619.94 | 529590.56 | 1438468.62 | 2 | 2 | 2 | 2 | 2 | 2 | 2 | 2 | 2 | 2 |
| Q9EPU4 | Cpsf1 | Cleavage and polyadenylation specificity factor su | 93.42 | 1.25 | 160818 | 2 | 2 | 2 | 2 | 2 | 2 | 2 | 2 | 2 | 2 | 2 | 2 | 2 | 30978.86 | 69380.15 | 2 | 2 | 2 | 2 | 2 | 2 |
| Q6NVF9 | Cpsf6 | Cleavage and polyadenylation specificity factor su | 368.72 | 16.33 | 59153 | 7 | 9715491 | 11815210 | 4055823 | 10476104 | 4395805 | 7186493 | 6554883 | 3950498 | 7050918.5 | 6460988 | 6361674 | 8626421 | 6470563.5 | 8493272 | 5822369 | 9693541 | 9473805 | 10334086 | 10728229 | 12707224 |
| Q5XI29 | Cpsf7 | Cleavage and polyadenylation specificity factor su | 187.95 | 10.39 | 51073 | 4 | 2 | 2 | 2 | 2 | 905423 | 2 | 2 | 2 | 664349.88 | 560755.75 | 495922.09 | 1013587.06 | 2 | 777103.25 | 665096.12 | 2 | 709047.88 | 2 | 1078673.5 | 2 |
| P18886 | Cpt2 | Carnitine O-palmitoyltransferase 2, mitochondrial | 47.37 | 1.52 | 74110 | 1 | 1313214.62 | 1845231.38 | 1696600.88 | 1516537.25 | 1425697.62 | 2 | 2 | 1824995.12 | 2 | 2 | 546640.69 | 2 | 2 | 2 | 2 | 590470.06 | 1307845.62 | 2 | 2 | 2 |
| Q64735 | Cr1l | Complement component receptor 1-like protein | 44.13 | 2.07 | 53763 | 1 | 2801565.5 | 3008579.5 | 2720423.25 | 2689988.5 | 1286312.75 | 1444023.62 | 1607272.5 | 2 | 2244076.75 | 2399374.5 | 1378454.5 | 1626061 | 1645643 | 2 | 2047335.12 | 1048882.75 | 1881184.12 | 1849950.12 | 2 | 2046875.5 |
| O88668 | Creg1 | Protein CREG1 | 147.04 | 9.55 | 24452 | 3 | 197486.19 | 497987.09 | 478435.28 | 679791 | 302760.25 | 927307.75 | 1772237.88 | 396012.91 | 1298007.38 | 848575.62 | 566873.31 | 941771.75 | 673787 | 1343447.88 | 2703111.75 | 2 | 1775302.75 | 1732937.25 | 132424.36 | 1092411.75 |
| Q91XD7 | Creld1 | Protein disulfide isomerase Creld1 | 114.94 | 5.48 | 45718 | 2 | 4829908 | 2189575.5 | 2257698.75 | 2463995.25 | 364393.97 | 3702490.5 | 1033065.12 | 2025745.12 | 3655823 | 2448002.25 | 3149795.75 | 1757626.25 | 2281230.75 | 3266891.25 | 4068856.5 | 998143.81 | 4388310.5 | 5223486 | 2638674.25 | 2787919 |
| Q4G063 | Creld2 | Protein disulfide isomerase Creld2 | 57.07 | 4.3 | 38251 | 1 | 2 | 734769.31 | 637293.44 | 2 | 758368.06 | 2 | 2 | 2 | 2 | 2 | 2 | 2 | 565916.19 | 2 | 2 | 2 | 2 | 2 | 2 | 2 |
| P63254 | Crip1 | Cysteine-rich protein 1 | 57.78 | 12.99 | 8550 | 1 | 2 | 2 | 2 | 2 | 2 | 2 | 2 | 2 | 2 | 2 | 4247174.5 | 3538392.25 | 1130459 | 1207162 | 2 | 1510186.12 | 2 | 2 | 2 | 2 |
| Q99KP3 | Cryl1 | Lambda-crystallin homolog | 115.65 | 7.84 | 35209 | 2 | 2 | 2207033 | 2 | 1399858 | 2 | 474510.66 | 2 | 2 | 2008641.5 | 2 | 2 | 502310.25 | 2 | 1044471.25 | 2 | 2 | 2 | 647392.62 | 186326.66 | 399958.12 |
| P47199 | Cryz | Quinone oxidoreductase | 35.34 | 2.42 | 35269 | 1 | 652807.56 | 664400.31 | 2 | 509164.84 | 908319.69 | 1039004.38 | 320627.41 | 830478.06 | 787337.44 | 552563.31 | 1059260.75 | 973072.81 | 606698.88 | 686086.5 | 428447.97 | 348021.12 | 2 | 791180.44 | 852586 | 1261843.38 |
| Q9CZU6 | Cs | Citrate synthase, mitochondrial | 683.77 | 32.33 | 51737 | 13 | 39932634 | 42029234 | 65507578 | 55162842 | 40203898 | 32449868 | 54560418 | 73072146 | 47358458 | 54639170 | 22206608 | 20020954 | 36907074 | 44111322 | 40753478 | 30543878 | 44359926 | 53590410 | 29695570 | 24778290 |
| P18395 | Csde1 | Cold shock domain-containing protein E1 | 177.32 | 7.89 | 88895 | 4 | 2532412.75 | 1696915 | 1335742.12 | 1716086 | 2368621.5 | 433744.25 | 506747.19 | 568499.69 | 490966.91 | 790579.75 | 2614241.25 | 3138766.5 | 3546241.5 | 3071236.75 | 1294135 | 1863632.25 | 1967614.12 | 2669362 | 1299263.25 | 2578365.75 |
| Q9ERK4 | Cse1l | Exportin-2 | 355.02 | 11.84 | 110455 | 9 | 13702212 | 15521284 | 9073197 | 8767833 | 11152760 | 5889202.5 | 6216690 | 1414926 | 3816168.5 | 5343307.5 | 10120087 | 2389382.25 | 9738372 | 9004469 | 10485803 | 3123076 | 9814846 | 15818800 | 10980938 | 12472340 |
| P09581 | Csf1r | Macrophage colony-stimulating factor 1 receptor | 77.34 | 1.74 | 109179 | 2 | 1769150.25 | 3220037.5 | 2556959 | 2367380.75 | 4335557 | 1899541.75 | 1447376.88 | 638239.19 | 1873098 | 2488783.25 | 1497542 | 1429790.5 | 1310795.38 | 1335856.62 | 2 | 1212014.12 | 1254697.38 | 1418438.88 | 1076713.12 | 1134834 |
| Q00941 | Csf2ra | Granulocyte-macrophage colony-stimulating facto | 33.97 | 2.58 | 41769 | 1 | 2 | 797844.5 | 833551.25 | 1055645 | 795426.81 | 2 | 583862.19 | 211464.19 | 672580.88 | 973895.88 | 2 | 2 | 2 | 512715.06 | 2 | 2 | 994292.5 | 2 | 903778.69 | 2 |
| P32577 | Csk | Tyrosine-protein kinase CSK | 183.41 | 9.78 | 50746 | 4 | 4502497 | 5576313 | 3487286.5 | 6605307 | 5559377 | 3901235.5 | 3947326 | 5013808.5 | 3559012.5 | 3115808.5 | 2683565.75 | 1975767.88 | 2773500.5 | 2634539 | 2565015.25 | 3493781.5 | 2602728 | 4366455 | 2659443 | 2845839.5 |
| P19139 | Csnk2a1 | Casein kinase II subunit alpha | 137.31 | 8.95 | 45073 | 2 | 2391625.5 | 1494863.5 | 1569905.75 | 121387.17 | 1431137.5 | 2 | 1169687 | 98105.86 | 732821.75 | 742751.06 | 1097113 | 1417887.88 | 1673073.12 | 1511913.25 | 2215368.5 | 1333061.88 | 2898039.5 | 1866324.25 | 1492170 | 1731088 |
| P67871 | Csnk2b | Casein kinase II subunit beta | 54.89 | 5.12 | 24942 | 1 | 1526465.62 | 1589488 | 1894390.75 | 1304467.5 | 1124134 | 2 | 885596.88 | 1457158.75 | 2 | 1096922.62 | 1451257.88 | 1336205.88 | 1477540.38 | 1437904 | 1635350 | 848538.5 | 1274875.5 | 1258747.75 | 1112910.12 | 1121203.5 |
| Q62426 | Cstb | Cystatin-B | 155.67 | 44.9 | 11046 | 3 | 7244759 | 6622262 | 7574419 | 5557882.5 | 7828922 | 7157263 | 8995716 | 13235120 | 8885087 | 7127745 | 16487954 | 18527734 | 11732268 | 8017994 | 14924616 | 16760776 | 14373458 | 10242861 | 7963583 | 7664079 |
| Q5BJQ6 | Cstf1 | Cleavage stimulation factor subunit 1 | 35.9 | 2.09 | 48382 | 1 | 2 | 858048.56 | 2 | 936213.94 | 72137.78 | 437859.69 | 959195.12 | 118312.38 | 1191674.5 | 918143.25 | 2 | 1081272.12 | 710352.5 | 641387.94 | 2 | 2 | 2 | 697899.88 | 2 | 1223530.12 |
| Q8BIQ5 | Cstf2 | Cleavage stimulation factor subunit 2 | 115.65 | 8.79 | 61341 | 2 | 2 | 2 | 2 | 113048.31 | 47526.37 | 2 | 2 | 2 | 2 | 2 | 99244.47 | 113327.56 | 2 | 211656.09 | 2 | 45726.68 | 2 | 2 | 2 | 2 |
| Q99LI7 | Cstf3 | Cleavage stimulation factor subunit 3 | 189.62 | 5.3 | 82877 | 3 | 2 | 35541.3 | 40810.74 | 88212.97 | 239633.67 | 85679.53 | 123621.74 | 28038.53 | 93828.95 | 189426.94 | 2 | 65966.11 | 53643.18 | 112424.98 | 2 | 60352.66 | 2 | 135845.02 | 2 | 2 |
| O88712 | Ctbp1 | C-terminal-binding protein 1 | 212.28 | 11.11 | 47745 | 5 | 2 | 2 | 2 | 460541.97 | 971752.88 | 1138458.75 | 813006.19 | 874000.12 | 713717.69 | 538521.88 | 1099521.62 | 605675.12 | 633201.12 | 712481.88 | 2 | 915464.69 | 2 | 2 | 2 | 2 |
| P56546 | Ctbp2 | C-terminal-binding protein 2 | 291.78 | 15.96 | 48957 | 6 | 1384937.38 | 1728993.25 | 813619.38 | 984581.62 | 3370692.75 | 819265.88 | 719818.12 | 4017390.75 | 1774820.5 | 2236464.75 | 2237285.5 | 3820392.25 | 3400154 | 4005842.5 | 1230954.62 | 2683839.25 | 189172.12 | 264715.81 | 378232.47 | 1668184 |
| Q8VCN5 | Cth | Cystathionine gamma-lyase | 497.8 | 27.14 | 43567 | 10 | 9377385 | 11501992 | 20802036 | 13511600 | 17593928 | 23467510 | 23925028 | 22619538 | 21611462 | 23167656 | 25079238 | 26935692 | 23539544 | 22674670 | 26509684 | 26265474 | 32094046 | 23433328 | 23389266 | 24455162 |
| P70698 | Ctps1 | CTP synthase 1 | 43.7 | 2.03 | 66682 | 1 | 2 | 2 | 2 | 2 | 2 | 2 | 2 | 2 | 328691.41 | 2 | 2 | 817919.94 | 2 | 559575.94 | 828224.94 | 2 | 2 | 2 | 2 | 2 |
| P16675 | Ctsa | Lysosomal protective protein | 900.07 | 30.17 | 53844 | 14 | 60496610 | 63021898 | 65243974 | 33514098 | 53683686 | 45084330 | 43242146 | 54966158 | 45749890 | 60867682 | 36576238 | 32424966 | 35394314 | 34484194 | 31973214 | 22223014 | 20966140 | 27274796 | 27781034 | 23042262 |
| P10605 | Ctsb | Cathepsin B | 1014.1 | 60.18 | 37280 | 15 | 72995490 | 24111864 | 90245346 | 28728632 | 16502801 | 33186804 | 47441650 | 57439558 | 26793874 | 44432298 | 22486784 | 25229406 | 32891094 | 17492196 | 62794622 | 15591084 | 31778506 | 31304148 | 17033588 | 22549392 |
| P97821 | Ctsc | Dipeptidyl peptidase 1 | 347.78 | 11.04 | 52376 | 5 | 9966582 | 10057206 | 11552610 | 9946583 | 1859763.12 | 7648097.5 | 7905754 | 10865365 | 7313495 | 10695562 | 2979265.5 | 3344654.25 | 4061609.75 | 3669782.75 | 4383450 | 1124639.38 | 2812850 | 3138291 | 2359982.75 | 2088358.62 |
| P18242 | Ctsd | Cathepsin D | 1137.54 | 48.78 | 44954 | 18 | 154405586 | 189911314 | 153709570 | 98533914 | 88569002 | 123585466 | 139359426 | 125183786 | 119250498 | 163430354 | 87023042 | 83809050 | 93645130 | 88233130 | 85334074 | 95319674 | 91113586 | 104785394 | 79968218 | 108342714 |
| P06797 | Ctsl | Procathepsin L | 229.11 | 27.54 | 37547 | 5 | 2572353.5 | 4053246 | 4163844.5 | 2752805.5 | 3479982 | 7705782 | 11124732 | 8979567 | 3654417 | 5710457 | 2467147 | 1096804.12 | 2535969.5 | 2235813.5 | 4256049.5 | 584025.62 | 4600990 | 5319623.5 | 2588154 | 3964336.5 |
| O70370 | Ctss | Cathepsin S | 602.93 | 32.94 | 38475 | 10 | 32769674 | 13329167 | 38621730 | 15359048 | 13989004 | 13682371 | 15477957 | 24647312 | 20715630 | 25611748 | 8828352 | 6007317.5 | 13223215 | 7078028.5 | 19334650 | 9084605 | 18196578 | 13329218 | 8303492.5 | 16853626 |
| Q9WUU7 | Ctsz | Cathepsin Z | 759.59 | 47.06 | 33996 | 13 | 283587170 | 356323202 | 202208258 | 191444130 | 102252082 | 236393378 | 271454978 | 196376322 | 162459650 | 195841762 | 244770226 | 253637522 | 232322002 | 225858482 | 251332786 | 215168370 | 331994754 | 284058210 | 191697634 | 279024162 |
| Q9WTX6 | Cul1 | Cullin-1 | 93.96 | 2.19 | 89692 | 2 | 2 | 2 | 917832.69 | 2 | 2 | 2 | 2 | 2 | 312397.34 | 2 | 611424.75 | 664805.38 | 870010.38 | 927142.94 | 541407.31 | 1006347.56 | 2 | 2 | 1094393 | 2 |
| A2A432 | Cul4b | Cullin-4B | 36.9 | 1.03 | 110699 | 1 | 2 | 1336532.75 | 707095.69 | 2 | 2 | 1090171.25 | 694747.69 | 2 | 554620.81 | 2 | 1300157.38 | 1009184.5 | 1039473.5 | 994183.5 | 2 | 1103576.75 | 680651.25 | 2 | 2 | 2 |
| Q6MGD0 | Cuta | Protein CutA | 33.07 | 7.91 | 18659 | 1 | 1373587.75 | 200078.66 | 398775.09 | 312388.62 | 353372.47 | 2 | 2 | 351151.12 | 437511.75 | 490268.69 | 119162.31 | 2 | 173892.3 | 254682.28 | 1480043 | 2 | 2 | 110564.4 | 188906.61 | 289753.66 |
| Q5BJP2 | Cwc15 | Spliceosome-associated protein CWC15 homolog | 111.27 | 11.35 | 26638 | 2 | 2 | 2 | 120877.98 | 129598.01 | 2 | 2 | 2 | 2 | 2 | 2 | 2 | 324715.03 | 2 | 312519 | 2 | 2 | 290940.06 | 2 | 473357.5 | 2 |
| Q9CQX2 | Cyb5b | Cytochrome b5 type B | 192.13 | 15.75 | 16318 | 2 | 915671.38 | 718192.81 | 1818190 | 585069.75 | 2019306 | 586687 | 2 | 1921275.25 | 1870178.75 | 2225749.75 | 3277430.5 | 1616061.88 | 2948501 | 2703744.5 | 3447611.5 | 2964692 | 2790975 | 4193625 | 4072544.5 | 3269958.75 |
| Q9DB73 | Cyb5r1 | NADH-cytochrome b5 reductase 1 | 253.07 | 22.95 | 34135 | 5 | 93336 | 2 | 2 | 2 | 529963.38 | 2 | 109964.15 | 469209.91 | 2596702.25 | 657124.5 | 2 | 2 | 2 | 2 | 442642.03 | 2 | 2 | 2 | 2 | 2 |
| Q9DCN2 | Cyb5r3 | NADH-cytochrome b5 reductase 3 | 616.91 | 32.89 | 34128 | 10 | 7118848 | 9929315 | 4224130 | 7929098 | 13780909 | 14013415 | 13932626 | 14568366 | 25970378 | 14913076 | 11151909 | 9860934 | 9020905 | 8622837 | 7362230 | 5090384.5 | 6342765.5 | 11099776 | 2356297.75 | 7148707 |
| Q61462 | Cyba | Cytochrome b-245 light chain | 50.9 | 5.21 | 20748 | 1 | 2465377.75 | 3670010 | 2 | 4600142 | 26856.51 | 4544493.5 | 3518501.25 | 248899.5 | 4897103.5 | 3663091 | 3785782.75 | 3445789.25 | 2138252 | 3293907.25 | 2302655 | 3587391.25 | 4006995 | 2584069.5 | 1655300.75 | 2748392.25 |
| Q61093 | Cybb | NADPH oxidase 2 | 97.22 | 3.86 | 65305 | 2 | 2 | 165377.86 | 102919.84 | 111421.06 | 743832.44 | 429791.66 | 430124 | 1366225.88 | 2322957 | 675046 | 2 | 665717.38 | 2 | 384656.62 | 2 | 2 | 2 | 2 | 489020.28 | 248475.17 |
| Q9D0M3 | Cyc1 | Cytochrome c1, heme protein, mitochondrial | 220.08 | 21.23 | 35328 | 5 | 1609983.12 | 2611508.5 | 731450.31 | 2735437.75 | 2969240.75 | 2991104.75 | 2533706.75 | 3948920.5 | 2495140.25 | 2801165.25 | 2746327.5 | 2216850.75 | 2 | 3316280.5 | 3307633.5 | 2667254.25 | 3823782 | 4227659.5 | 2981094.25 | 3458064.5 |
| P62897 | Cycs | Cytochrome c, somatic | 429.23 | 52.38 | 11605 | 7 | 7632678.5 | 3735869.25 | 17590750 | 3471805.75 | 4405584 | 10279281 | 7615650 | 9514301 | 9664836 | 6216820 | 8086602.5 | 9753988 | 9325436 | 5130980.5 | 7481019.5 | 10743440 | 9966158 | 6680461 | 8128925 | 8060983.5 |
| Q7TMB8 | Cyfip1 | Cytoplasmic FMR1-interacting protein 1 | 751.51 | 13.01 | 145241 | 15 | 6839314 | 7714240.5 | 8690557 | 7099803 | 8213055.5 | 11880539 | 9639829 | 16722741 | 12318325 | 9600170 | 16534412 | 13017145 | 15349741 | 14831595 | 13903786 | 11246515 | 11660331 | 16479392 | 11029072 | 10603626 |
| Q64654 | Cyp51a1 | Lanosterol 14-alpha demethylase | 43.18 | 1.99 | 56707 | 1 | 2 | 2 | 2 | 434339.94 | 1032993.44 | 2 | 2 | 2 | 2 | 2 | 2 | 2 | 2 | 2 | 2 | 2 | 2 | 2 | 2 | 2 |
| Q921M7 | Cyrib | CYFIP-related Rac1 interactor B | 569.19 | 23.15 | 36776 | 7 | 3460429 | 3837350.5 | 3078074.5 | 4561986 | 8028799.5 | 5232979 | 5709823.5 | 6515968.5 | 7320806 | 6863387 | 4089260.25 | 3996477.5 | 1968951.38 | 3991010 | 2921961.25 | 1665770.12 | 2420803.5 | 4337324 | 3379288 | 1737259.5 |
| P98078 | Dab2 | Disabled homolog 2 | 658.28 | 22.45 | 82312 | 13 | 4214928 | 6248769.5 | 2208279.25 | 6501170 | 5655048 | 1973320.12 | 2980675.25 | 5100780.5 | 7580857.5 | 2785238.75 | 4182532 | 6909883 | 1974296.75 | 2921055 | 4328582 | 5187411 | 2070847.38 | 2301310.5 | 2158352.75 | 5032431 |
| P61804 | Dad1 | Dolichyl-diphosphooligosaccharide--protein glyco | 233.91 | 28.32 | 12497 | 4 | 2654903.75 | 3176288 | 4159078.5 | 4000526.75 | 3198598.5 | 4849354 | 4303510.5 | 6413080 | 4563749.5 | 4911875 | 2227366.25 | 2716564 | 3881935 | 3760852.5 | 9782006 | 2537953.5 | 5798414 | 5273270.5 | 3806530 | 6960222 |
| Q9ER88 | Dap3 | Small ribosomal subunit protein mS29 | 99.84 | 6.39 | 44699 | 2 | 502544 | 2 | 1137965.88 | 1641574.12 | 1113614.25 | 947314.38 | 518148.66 | 1089811.75 | 315649.5 | 2 | 1271784.12 | 747442.31 | 1224909.75 | 558061 | 590279.75 | 2 | 2 | 2 | 188718.64 | 2 |
| Q922B2 | Dars1 | Aspartate--tRNA ligase, cytoplasmic | 869.59 | 37.13 | 57147 | 17 | 11624502 | 16994082 | 19068650 | 12116514 | 22311876 | 13040126 | 13883816 | 15926322 | 12534238 | 11261261 | 13193452 | 11868038 | 15662639 | 12045594 | 20345900 | 13204401 | 15334048 | 17548486 | 10929534 | 16018922 |
| Q8BIP0 | Dars2 | Aspartate--tRNA ligase, mitochondrial | 148.2 | 6.28 | 74102 | 4 | 1149143.25 | 617746.56 | 647114.25 | 602328.69 | 1085097 | 883530.12 | 2 | 1241942.12 | 886511.56 | 1685919.75 | 978142.88 | 789979.12 | 1285189.62 | 1489511 | 750079.56 | 2 | 2 | 1122313.5 | 2 | 2 |
| Q9JII5 | Dazap1 | DAZ-associated protein 1 | 36.32 | 3.69 | 43214 | 1 | 751451.88 | 2 | 158279.14 | 2 | 73793.37 | 2 | 2 | 2 | 573320.38 | 271167.41 | 204524.88 | 324783.72 | 2 | 461146.84 | 932472.31 | 2 | 2 | 2 | 2 | 2 |
| P31786 | Dbi | Acyl-CoA-binding protein | 193.65 | 47.13 | 10000 | 3 | 2 | 826058.38 | 915903.38 | 2 | 1696471.38 | 3686650 | 769765.12 | 3290002 | 1510455.25 | 1131934.5 | 3593361.75 | 4217579.5 | 3167867.75 | 2436100.25 | 3387703.5 | 4398622.5 | 4887994 | 1963294.38 | 5033773 | 3547265.5 |
| Q62418 | Dbnl | Drebrin-like protein | 220.25 | 10.32 | 48700 | 4 | 2 | 935334.19 | 1055620.88 | 592149.06 | 1687568 | 2241108.5 | 2 | 1853245.38 | 1138046.12 | 2 | 3730809.75 | 4507911 | 2232920.25 | 2870302.5 | 1895743.5 | 3501083 | 1248764.62 | 1194395.62 | 138780.22 | 1504003.75 |
| P53395 | Dbt | Lipoamide acyltransferase component of branche | 193.77 | 18.46 | 53247 | 5 | 632116.81 | 92005.46 | 823346 | 2 | 873944.38 | 2 | 599693.69 | 75617.88 | 641712.44 | 1120192.62 | 71278.73 | 111344.64 | 477808.56 | 129992.18 | 586422 | 2 | 2 | 138534.36 | 2 | 139377.25 |
| Q6AY55 | Dcakd | Dephospho-CoA kinase domain-containing protei | 73.78 | 5.83 | 27352 | 2 | 49519.7 | 2 | 140009.83 | 121597.67 | 92570.44 | 2 | 2 | 2 | 40184.57 | 611118.12 | 2 | 2 | 2 | 181454.97 | 636859.5 | 65029.71 | 143474.94 | 2 | 2 | 2 |
| Q8K4F7 | Dcps | m7GpppX diphosphatase | 120.03 | 8.04 | 38714 | 3 | 1652880.62 | 2009057.88 | 844786.38 | 1627218 | 1214186.25 | 2836893 | 1607809.12 | 1389388.62 | 3221535 | 2488184.75 | 4660958 | 4302133 | 3127991.25 | 2422138 | 1318319.25 | 2480024.25 | 1519943.25 | 1065134.38 | 1563875.25 | 663066.06 |
| O08788 | Dctn1 | Dynactin subunit 1 | 255.96 | 3.12 | 141676 | 4 | 377380.41 | 1297145.12 | 1630998.25 | 2 | 1638350.25 | 2 | 1851527.62 | 2536811.75 | 1454082.25 | 800722.81 | 847324.88 | 483088.47 | 1242989.5 | 1329250.75 | 1594324.38 | 2 | 592015.31 | 2 | 594388.75 | 1963178.75 |
| Q6AYH5 | Dctn2 | Dynactin subunit 2 | 86.64 | 3.98 | 44148 | 2 | 1761509.75 | 1089012.12 | 1640750 | 759896.62 | 1497888.62 | 891362.06 | 617492.38 | 2 | 858954.88 | 1166443.75 | 1018521.19 | 2009283.25 | 994618.44 | 1769697.75 | 1925154 | 1095022.25 | 1926627.38 | 1149575.12 | 1624264.62 | 1047082.19 |
| Q9Z0Y1 | Dctn3 | Dynactin subunit 3 | 49.84 | 4.3 | 20978 | 1 | 2 | 1913965.38 | 2 | 1526840.25 | 884301 | 2 | 1274132.38 | 2 | 735944.56 | 994955.25 | 2 | 2 | 2 | 2 | 2 | 324364.22 | 600127.38 | 1539035.38 | 887926.44 | 2 |
| Q91X52 | Dcxr | L-xylulose reductase | 102.39 | 9.02 | 25746 | 2 | 881139.62 | 578359.81 | 1532423.38 | 2 | 536282.25 | 2 | 2 | 573599.56 | 583108.5 | 2 | 1048352.81 | 1064883.5 | 1070155.5 | 1480067.5 | 1235769.38 | 887516.19 | 928220.81 | 2 | 1260739.5 | 1437128.12 |
| Q6MG60 | Ddah2 | Putative hydrolase DDAH2 | 57.23 | 4.21 | 29688 | 1 | 2 | 2 | 2 | 2 | 2 | 2 | 2 | 2 | 2 | 2 | 2 | 620267.44 | 2 | 519638.72 | 176066.17 | 367734 | 2 | 2 | 2 | 197693.38 |

| | | | | | | | | | | | | | | | | | | | | | | | | | | |
|---|---|---|---|---|---|---|---|---|---|---|---|---|---|---|---|---|---|---|---|---|---|---|---|---|---|---|
| Q3U1J4 | Ddb1 | DNA damage-binding protein 1 | 798.77 | 18.86 | 126853 | 16 | 1331627.38 | 385046.47 | 2019138.88 | 1871846.38 | 2137897.25 | 784149.62 | 289826.41 | 1039297.69 | 1744302.75 | 1224534.38 | 3464360.25 | 2194599.5 | 2013639.25 | 1625325.12 | 726071.38 | 3020492 | 1502917.25 | 596281.88 | 3055690.5 | 717622.5 |
| A2ADY9 | Ddi2 | Protein DDI1 homolog 2 | 116.99 | 8.52 | 44591 | 3 | 2 | 559773.69 | 2 | 2 | 2 | 2 | 2 | 2 | 664471.19 | 2 | 1633155.62 | 1699669.62 | 1126665.12 | 1194470.75 | 582048.44 | 2 | 2 | 901158.25 | 2 | 1432063.75 |
| O54734 | Ddost | Dolichyl-diphosphooligosaccharide--protein glyco | 542.13 | 32.2 | 49028 | 10 | 988871.19 | 793772.88 | 158896.59 | 5857652 | 4563543.5 | 3933883.25 | 4663567 | 5578807.5 | 5215335.5 | 2999775.5 | 5109761.5 | 3699744.5 | 3315650.5 | 4233974.5 | 1295737.75 | 4423557 | 3904115.75 | 1116056.25 | 3592805.5 | 1049277.25 |
| Q80WW9 | Ddrgk1 | DDRGK domain-containing protein 1 | 32.91 | 6.03 | 35977 | 1 | 2 | 2 | 112686.25 | 2 | 2 | 2 | 2 | 2 | 2 | 63175.1 | 2 | 2 | 2 | 2 | 2 | 2 | 2 | 2 | 2 | 2 |
| O35215 | Ddt | D-dopachrome decarboxylase | 50.82 | 11.02 | 13077 | 1 | 1328448.25 | 2 | 2 | 2 | 2 | 2 | 2 | 2 | 462074.31 | 761013.81 | 2673077.75 | 2518914.75 | 2023755.5 | 2096887.88 | 2026025.62 | 3760333.25 | 2855388.75 | 2154484.75 | 1177419.62 | 2557334.75 |
| Q641Y8 | Ddx1 | ATP-dependent RNA helicase DDX1 | 413.21 | 15.68 | 82497 | 8 | 1441201.88 | 1416686.38 | 3692550 | 1124458 | 2651993.75 | 1009614.31 | 3116001.75 | 1311853 | 1272439.62 | 2361675.75 | 2487210 | 3118347.75 | 2806815 | 2818438 | 2458834.5 | 1210894.75 | 3579602.75 | 2562932.5 | 3245978 | 1410223.88 |
| Q80Y44 | Ddx10 | Probable ATP-dependent RNA helicase DDX10 | 45.92 | 1.37 | 100739 | 1 | 2 | 2 | 2 | 2 | 2 | 2 | 2 | 2 | 2 | 2 | 2 | 545771 | 2 | 2 | 2 | 2 | 2 | 2 | 2 | 2 |
| Q501J6 | Ddx17 | Probable ATP-dependent RNA helicase DDX17 | 873.51 | 22.15 | 72399 | 16 | 5375434 | 4466246 | 6413443 | 5826924 | 9431389 | 5613790.5 | 7155625 | 7511244.5 | 6912485 | 8018618 | 6001200.5 | 5181111.5 | 2345533.25 | 3859021.25 | 4644957 | 2544911.5 | 3525124.75 | 3791416 | 2796651.5 | 1642423 |
| Q8K363 | Ddx18 | ATP-dependent RNA helicase DDX18 | 391.18 | 13.94 | 74181 | 7 | 2299865 | 6645848.5 | 1592590.62 | 4430764 | 3643527 | 1183985.75 | 3736056 | 986316.25 | 2682083.25 | 3686101 | 2877107.5 | 3190479.5 | 5846776 | 3004100 | 3440334 | 4203383.5 | 3308428.5 | 2424459.75 | 1656854.25 | 2051049.88 |
| Q61655 | Ddx19a | ATP-dependent RNA helicase DDX19A | 153.04 | 7.74 | 53933 | 3 | 5206568 | 2 | 3733487.5 | 8794181 | 779624.81 | 2 | 1520118.88 | 10816044 | 506753.78 | 8691126 | 2 | 1573127.5 | 1329928.5 | 7139836.5 | 5876803 | 5014062 | 6736921.5 | 1697619.75 | 6842722.5 | 4790828 |
| Q9JIK5 | Ddx21 | Nucleolar RNA helicase 2 | 1382.49 | 34.2 | 93551 | 27 | 14660929 | 19725980 | 7447279 | 20807374 | 30574464 | 9520805 | 14202333 | 18484526 | 18858874 | 13471442 | 13095949 | 14587220 | 15299324 | 15790336 | 8172312 | 10062981 | 13197494 | 11177677 | 9889287 | 9561682 |
| Q9ESV0 | Ddx24 | ATP-dependent RNA helicase DDX24 | 101.35 | 3.85 | 96429 | 3 | 2 | 2 | 2 | 2 | 2 | 2 | 2 | 2 | 2 | 2 | 2 | 2 | 2 | 2 | 2 | 2 | 2 | 2 | 2 | 2 |
| Q921N6 | Ddx27 | Probable ATP-dependent RNA helicase DDX27 | 129.44 | 3.82 | 85939 | 3 | 1476115 | 2 | 675239.44 | 480835.78 | 502128.66 | 821956.62 | 4400984 | 1287927.62 | 123141.02 | 6019432 | 671347.44 | 1509798.62 | 1866166 | 1671815.88 | 5873804 | 2 | 1484265 | 1916440.12 | 2 | 2471833.5 |
| Q5U216 | Ddx39a | ATP-dependent RNA helicase DDX39A | 429.88 | 15.93 | 49109 | 8 | 1625407.75 | 1222585.25 | 2933083.5 | 1152070.88 | 2319994.25 | 893163.81 | 1183292.38 | 1913035.88 | 1696251.5 | 1561566.12 | 1480788.88 | 1512951.5 | 575010.88 | 209403.33 | 460191.94 | 1904587.25 | 232997.36 | 125679.16 | 2 | 2 |
| Q63413 | Ddx39b | Spliceosome RNA helicase Ddx39b | 563.56 | 22.9 | 49035 | 10 | 5381199 | 2053815.75 | 8287410 | 6378374 | 5546578.5 | 933094.38 | 1392225.25 | 6251334 | 4639227 | 4368186.5 | 4339041 | 3156720.25 | 4329065.5 | 3535299.5 | 3412934.75 | 3723622.25 | 1541691.25 | 1660099.75 | 840798.75 | 2036792.5 |
| Q62167 | Ddx3x | ATP-dependent RNA helicase DDX3X | 798.15 | 25.83 | 73101 | 15 | 2756914 | 4817388.5 | 1882745.5 | 4409716 | 9113862 | 1573549 | 2481869.75 | 2652799.25 | 6385099 | 3344392.25 | 5449289.5 | 5766802 | 5196728 | 5323774.5 | 2744434.5 | 4423331.5 | 4344011.5 | 1923125 | 5459253 | 2704121.25 |
| Q810A7 | Ddx42 | ATP-dependent RNA helicase DDX42 | 33.7 | 0.97 | 101965 | 1 | 2 | 2 | 505583.03 | 674433.56 | 819654 | 865213 | 645262.06 | 2 | 607758.69 | 592162.81 | 1072049.62 | 673059.94 | 721564.25 | 658197.69 | 708024.5 | 2 | 1197792 | 747259.06 | 2 | 2 |
| Q569Z5 | Ddx46 | Probable ATP-dependent RNA helicase DDX46 | 376.33 | 7.46 | 117448 | 7 | 2811301.25 | 4895548.5 | 3641596 | 3171037.5 | 8975891 | 3655009.5 | 4168928 | 5285415 | 6178812.5 | 5274095 | 4773318 | 6236200.5 | 3255317.25 | 4716889 | 3178219 | 4740643 | 1796028.12 | 3769890 | 3384032 | 1917074 |
| Q9CWX9 | Ddx47 | Probable ATP-dependent RNA helicase DDX47 | 115.26 | 7.47 | 50639 | 3 | 2 | 2 | 603123.5 | 2 | 144576.19 | 2 | 2 | 2 | 47010.39 | 366677.78 | 2 | 2 | 2 | 2 | 352917 | 2 | 2 | 2 | 2 | 2 |
| Q4FZF3 | Ddx49 | Probable ATP-dependent RNA helicase DDX49 | 45.07 | 2.08 | 54094 | 1 | 984464.12 | 2 | 698263.81 | 418737.91 | 2 | 2 | 541988.56 | 1025772 | 2 | 905091.75 | 2 | 2 | 526620.38 | 2 | 1906054.25 | 2 | 1294583 | 2 | 466560.03 | 708975 |
| Q61656 | Ddx5 | Probable ATP-dependent RNA helicase DDX5 | 1469.85 | 43.97 | 69290 | 26 | 10820591 | 11027891 | 17058738 | 15185129 | 18614352 | 8678098 | 8961564 | 12671051 | 12483630 | 13531575 | 9202171 | 9096006 | 9488320 | 12148721 | 13258819 | 10617940 | 8327026 | 8267685 | 2896182.5 | 6609962.5 |
| Q9D0R4 | Ddx56 | Probable ATP-dependent RNA helicase DDX56 | 187.11 | 12.27 | 61212 | 5 | 511709.69 | 607856.69 | 2 | 151845.48 | 1086059.12 | 2 | 67791.89 | 80008.64 | 2 | 667825.5 | 2 | 2 | 2 | 642371.62 | 233634.5 | 534071.38 | 2 | 606559.81 | 690544.12 | 680209.62 |
| P54823 | Ddx6 | Probable ATP-dependent RNA helicase DDX6 | 272.27 | 13.66 | 54192 | 4 | 996414.94 | 786558.12 | 1063220.12 | 842804.44 | 2374853.25 | 1539532.5 | 1779678.75 | 1885490.5 | 1217537.5 | 2718213.5 | 899238.5 | 2019959.25 | 1033618.31 | 782486.25 | 2 | 740215.69 | 1133890.12 | 528082.69 | 2 | 1802754 |
| Q9CQ62 | Decr1 | 2,4-dienoyl-CoA reductase [(3E)-enoyl-CoA-produ | 227.99 | 17.61 | 36214 | 4 | 7909426 | 11676932 | 6703230.5 | 2400037.5 | 6852787.5 | 3609069.5 | 7474065 | 5377580 | 5153732 | 6822038.5 | 4233362 | 4501077 | 6585457 | 5507950 | 1902163.12 | 5799764 | 2847092.75 | 2366610.5 | 2842232 | 2524776.75 |
| Q9WV68 | Decr2 | Peroxisomal 2,4-dienoyl-CoA reductase [(3E)-enc | 38.79 | 5.82 | 31300 | 1 | 2 | 440195.09 | 2 | 2 | 1149271.88 | 2 | 2 | 2 | 906220.56 | 2 | 2 | 2 | 785530.25 | 2 | 406918.38 | 887682.56 | 2 | 2 | 2 | 2 |
| Q7TNV0 | Dek | Protein DEK | 215.56 | 11.05 | 43159 | 5 | 3961927 | 5561773 | 1647507 | 7332311 | 3303278 | 7181484.5 | 9537751 | 5962845.5 | 3682802.5 | 3563841.5 | 3541313.75 | 1870320.75 | 3154708.5 | 2067077 | 7414833.5 | 1711710.12 | 6849680.5 | 5535017.5 | 2772607 | 5395717 |
| A6H8H2 | Dennd4c | DENN domain-containing protein 4C | 58.01 | 1.99 | 211458 | 2 | 907617.25 | 90579.32 | 736048.56 | 2 | 642784.5 | 2 | 2 | 173010.3 | 344994.75 | 695540.06 | 2 | 2 | 419230.56 | 99867.75 | 184739.7 | 2 | 87708.19 | 2 | 2 | 2 |
| Q91YP3 | Dera | Deoxyribose-phosphate aldolase | 65.78 | 3.77 | 34975 | 1 | 2 | 2 | 2 | 2 | 577671.75 | 2 | 2 | 2 | 2 | 654190.12 | 2 | 2 | 2 | 2 | 2 | 2 | 2 | 2 | 2 | 2 |
| Q8BH86 | Dglucy | D-glutamate cyclase, mitochondrial | 36.59 | 1.94 | 66366 | 1 | 2 | 877416.75 | 2 | 2 | 2 | 2 | 816341.56 | 2 | 615584 | 830196.38 | 2 | 2 | 2 | 2 | 868359.69 | 2 | 2 | 2 | 2 | 2 |
| O88455 | Dhcr7 | 7-dehydrocholesterol reductase | 83.97 | 4.03 | 53919 | 2 | 2384252 | 3667086.5 | 374852.22 | 2470925.25 | 4113535.75 | 3945150 | 3047058.5 | 5303026.5 | 4322313.5 | 2758480.75 | 1489333.12 | 850303.19 | 2140464 | 2477889.25 | 2019155.5 | 1715643 | 2501226 | 3219291.75 | 1963756.5 | 2285889.5 |
| Q9DBB8 | Dhdh | Trans-1,2-dihydrobenzene-1,2-diol dehydrogenas | 40.86 | 3.6 | 36301 | 1 | 2846774.5 | 2 | 2999500.25 | 2726958.75 | 1387256.25 | 1472728 | 2069980 | 2340339 | 812665.69 | 2155907.75 | 1609892.75 | 1666710.5 | 2295097.75 | 1389391.75 | 1173599.88 | 2 | 1380553 | 2113624.25 | 1001620.88 | 4265782 |
| P00375 | Dhfr | Dihydrofolate reductase | 41.6 | 3.74 | 21606 | 1 | 2724639 | 2976084.5 | 4816422.5 | 2806103.25 | 3378183 | 2353422.75 | 2690691 | 3733866.25 | 423279.97 | 3794531 | 646227.12 | 433386.78 | 2325262.5 | 2526174.25 | 3332033.25 | 2080717.5 | 1921578.5 | 1766163.75 | 1673888.62 | 2254035 |
| O35435 | Dhodh | Dihydroorotate dehydrogenase (quinone), mitoch | 53.27 | 2.28 | 42700 | 1 | 2 | 2 | 2 | 275087.25 | 332475.47 | 2 | 83383.91 | 2 | 454920.09 | 645054.25 | 2 | 114952.87 | 479912.09 | 239309.09 | 338735.78 | 2 | 241949.44 | 306739.06 | 2 | 259515.19 |
| Q99L04 | Dhrs1 | Dehydrogenase/reductase SDR family member 1 | 242.89 | 16.61 | 34005 | 4 | 4151555.75 | 9677468 | 3599233.25 | 2299553 | 4747139.5 | 11220348 | 9663236 | 1871972.75 | 11582947 | 7635485.5 | 7861526 | 6296899 | 5712195 | 7364894.5 | 1956686.12 | 6242249 | 5877738.5 | 4171303.25 | 1426413.62 | 3595906 |
| O88876 | Dhrs3 | Short-chain dehydrogenase/reductase 3 | 34.84 | 3.31 | 33652 | 1 | 652819.12 | 659943.44 | 2 | 785387 | 915550.19 | 1799909.5 | 702718.06 | 2 | 1160150.25 | 953881.44 | 711278.25 | 1060097.88 | 668102 | 705703.31 | 955153.5 | 399562 | 938871.69 | 677820.69 | 2 | 2 |
| Q99LB2 | Dhrs4 | Dehydrogenase/reductase SDR family member 4 | 88.19 | 9.68 | 29885 | 3 | 2 | 1384536.38 | 734275.5 | 335406.75 | 2062313 | 2 | 635616.31 | 2089196.75 | 2090351.38 | 2047995.12 | 2 | 2 | 2 | 2 | 2 | 2 | 2 | 2 | 2 | 2 |
| Q99J47 | Dhrs7b | Dehydrogenase/reductase SDR family member 7I | 140.65 | 12.07 | 34987 | 3 | 1084185.12 | 845515.62 | 782108.88 | 2 | 1065459.5 | 1072482.62 | 956901.25 | 1414734.88 | 1123041.5 | 417966.28 | 942625.19 | 727917.94 | 860612.81 | 718750.81 | 2 | 805130.88 | 819989.88 | 2 | 1230814.12 | 2 |
| Q58NB6 | Dhrs9 | Dehydrogenase/reductase SDR family member 9 | 38.76 | 3.13 | 35242 | 1 | 1658618.38 | 2 | 506758.19 | 2 | 546151.44 | 2 | 2 | 2 | 1006211.88 | 420361.28 | 2 | 2 | 2 | 263712.06 | 2 | 2 | 2 | 2 | 2 | 2 |
| O35286 | Dhx15 | ATP-dependent RNA helicase DHX15 | 885.64 | 22.01 | 91007 | 19 | 10356263 | 6667808.5 | 7292084.5 | 19089526 | 9522261 | 9814779 | 7689165 | 10851987 | 14616590 | 7605344.5 | 10966342 | 10628593 | 11881156 | 12045459 | 12307624 | 6726859 | 10439413 | 13792273 | 1050933.38 | 10523434 |
| D4A2Z8 | Dhx36 | ATP-dependent DNA/RNA helicase DHX36 | 54.71 | 1 | 113843 | 1 | 604662.75 | 2 | 2 | 256693.17 | 548055.06 | 2 | 83686.76 | 109274.23 | 394923.19 | 558372.44 | 55624.84 | 2 | 2 | 135785.05 | 138862.05 | 2 | 154388.91 | 2 | 2 | 101600.53 |
| O70133 | Dhx9 | ATP-dependent RNA helicase A | 966.91 | 15.22 | 149475 | 20 | 12296841 | 25595374 | 14642867 | 22321210 | 19093802 | 16504436 | 14519060 | 13204253 | 18110976 | 15829535 | 16459609 | 19301276 | 18026124 | 19309538 | 10822664 | 15540193 | 10347514 | 11487431 | 9355492 | 9744599 |
| F1M775 | Diaph1 | Protein diaphanous homolog 1 | 38.3 | 0.79 | 140409 | 1 | 1289337.62 | 2 | 806902.94 | 890822.94 | 1161253.88 | 2 | 1547050.62 | 1097259.62 | 609201.38 | 538483.62 | 895051.88 | 1511037.88 | 2 | 521736.12 | 530176.31 | 2 | 1454353 | 992082.94 | 1397759.12 | 2 |
| Q3UH60 | Dip2b | Disco-interacting protein 2 homolog B | 184.29 | 4.07 | 171126 | 4 | 2 | 678241.44 | 2 | 332443.78 | 1291017.38 | 2 | 2 | 2 | 259264.62 | 49410.25 | 669620.44 | 694035.38 | 2174179.5 | 817137.88 | 1057092.88 | 679018.25 | 808820.38 | 1443287.75 | 857597.12 | 759810.69 |
| Q9ESX5 | Dkc1 | H/ACA ribonucleoprotein complex subunit DKC1 | 213.44 | 13.16 | 57402 | 4 | 649008.12 | 199357.58 | 493166.44 | 210426.92 | 470222.88 | 2 | 546547.62 | 75193.27 | 497015.91 | 498540.25 | 2 | 123690.24 | 2 | 487297.72 | 2 | 156640.59 | 1045381.88 | 247032.88 | 121895.05 | 460948.62 |
| Q8BMF4 | Dlat | Dihydrolipoyllysine-residue acetyltransferase co | 505.14 | 16.67 | 67942 | 9 | 19587984 | 15321852 | 35933482 | 13986608 | 18270518 | 19833712 | 15017975 | 32152414 | 18983468 | 26333374 | 24342962 | 19963064 | 23889270 | 23824966 | 27150124 | 27057662 | 35037230 | 45361118 | 32903870 | 37266826 |
| O08749 | Dld | Dihydrolipoyl dehydrogenase, mitochondrial | 743.84 | 33.2 | 54272 | 13 | 20545326 | 25545548 | 26905368 | 14114178 | 16944786 | 24357402 | 23785566 | 26946452 | 17377676 | 22195810 | 14554430 | 15731399 | 21012594 | 17102224 | 20122074 | 15721637 | 16596246 | 18312358 | 19260876 | 17352710 |
| Q01205 | Dlst | Dihydrolipoyllysine-residue succinyltransferase | 772.62 | 30.4 | 48925 | 14 | 22260872 | 41642378 | 26501730 | 25538066 | 20896080 | 25262490 | 25008292 | 30052748 | 25260500 | 35489654 | 26782988 | 21445362 | 32664818 | 43344182 | 20662486 | 25425142 | 25991512 | 37540174 | 23572110 | 39299554 |
| Q9QYJ0 | Dnaja2 | DnaJ homolog subfamily A member 2 | 244.48 | 22.09 | 45746 | 5 | 711557.75 | 906188.44 | 2 | 1072374.62 | 1040374.25 | 1176098.62 | 127951.31 | 224342.69 | 792002.12 | 904091.25 | 692064.38 | 3014897.25 | 148894.56 | 2631479.25 | 275388.97 | 2324135 | 331817.91 | 182231.8 | 121824.21 | 389914.31 |
| Q99M87 | Dnaja3 | DnaJ homolog subfamily A member 3, mitochond | 78.91 | 7.71 | 52443 | 2 | 2 | 2 | 2 | 843718.38 | 617236.69 | 2 | 2 | 2 | 730760 | 2 | 2 | 2 | 2 | 2 | 2 | 2 | 2 | 2 | 715520 | 2 |
| Q9QYJ3 | Dnajb1 | DnaJ homolog subfamily B member 1 | 36.82 | 3.24 | 38167 | 1 | 453459.16 | 626159.5 | 526263.25 | 2 | 990360 | 2 | 2 | 2 | 400953.09 | 356465.88 | 672153.31 | 2 | 2 | 733725.25 | 2 | 2 | 2 | 823860.5 | 2 | 1049788.88 |
| Q6TUG0 | Dnajb11 | DnaJ homolog subfamily B member 11 | 241.05 | 14.53 | 40495 | 4 | 1289898.25 | 2 | 1573420.38 | 1268236.5 | 2946835 | 1221611.62 | 770790.56 | 2268757.25 | 1400435.38 | 1807023.5 | 1087015.38 | 1106782.5 | 1378547.75 | 1213902.62 | 1158212.62 | 1548969.75 | 1313954.75 | 1434336.62 | 1218063.88 | 820106.38 |
| Q5U458 | Dnajc11 | DnaJ homolog subfamily C member 11 | 91.14 | 4.65 | 63233 | 2 | 2 | 2 | 2 | 2 | 691453.19 | 2 | 2 | 2 | 905416.31 | 358191.19 | 2 | 2 | 568897.56 | 2 | 2 | 2 | 140742.06 | 2 | 2 | 2 |
| P54103 | Dnajc2 | DnaJ homolog subfamily C member 2 | 89.61 | 3.38 | 71722 | 2 | 2 | 2 | 50108.62 | 517769.44 | 1674209.38 | 1585972.62 | 1845382 | 2 | 1094091.25 | 871660 | 1244544.88 | 936997.38 | 2 | 2 | 1206647.62 | 2 | 2 | 2 | 2138815.25 | 2 |
| Q91YW3 | Dnajc3 | DnaJ homolog subfamily C member 3 | 31.97 | 1.79 | 57464 | 1 | 2 | 2 | 2 | 2 | 2 | 2 | 496245.94 | 2 | 2 | 2 | 445175.84 | 2 | 2 | 2 | 2 | 2 | 2 | 2 | 2 | 2 |
| Q9QYI3 | Dnajc7 | DnaJ homolog subfamily C member 7 | 194.82 | 9.72 | 56476 | 4 | 822425.25 | 1467690.5 | 2 | 2675858.5 | 3109429.75 | 970735.75 | 1002279.62 | 1188447.88 | 465398.97 | 512726.5 | 812758.12 | 1022292 | 850311.25 | 1101318 | 491182.19 | 1202886.88 | 577744.62 | 2 | 1421142.5 | 674082.44 |
| P56542 | Dnase2 | Deoxyribonuclease-2-alpha | 96.25 | 5.38 | 38811 | 1 | 2 | 157091.77 | 2 | 73371.59 | 112706.55 | 149906.23 | 188959.11 | 61216.95 | 136254.56 | 115954.48 | 54441.09 | 2 | 2 | 71044.97 | 104949.8 | 2 | 102227.35 | 2 | 2 | 2 |
| O35303 | Dnm1l | Dynamin-1-like protein | 105.23 | 3.97 | 83908 | 2 | 51488.3 | 2 | 2 | 2 | 1152263.62 | 2 | 936895.38 | 1112222.62 | 1989338.75 | 931891.88 | 2 | 2 | 2 | 156249.11 | 637020.44 | 2 | 153506.16 | 2 | 2 | 86257.29 |
| P39052 | Dnm2 | Dynamin-2 | 663.16 | 16.21 | 98230 | 14 | 13263781 | 13478282 | 11996636 | 12278042 | 12508248 | 15012535 | 13241766 | 13945679 | 11337590 | 10613720 | 12816476 | 10209760 | 13292963 | 12121189 | 13532649 | 17006294 | 14570324 | 17080540 | 16687280 | 18865232 |
| P13864 | Dnmt1 | DNA (cytosine-5)-methyltransferase 1 | 365.76 | 5.93 | 183189 | 8 | 794419.38 | 1027196.06 | 671121.75 | 1295460.12 | 1833908.38 | 2 | 2 | 2 | 876366 | 2 | 475003 | 2 | 2 | 2 | 2 | 2 | 2 | 2 | 2 | 2 |
| O88508 | Dnmt3a | DNA (cytosine-5)-methyltransferase 3A | 296.51 | 5.62 | 101672 | 5 | 971561.38 | 549706.75 | 110232.73 | 1115566.12 | 546426.75 | 2 | 2 | 2 | 408539.62 | 767546.38 | 1417473.5 | 923991.12 | 1462814.75 | 1446965.75 | 1065983 | 558657.88 | 807051.38 | 607103.44 | 717015 | 2 |
| Q8R2M2 | Dnttip2 | Deoxynucleotidyltransferase terminal-interacting | | 44.82 | 1.32 | 84277 | 1 | 601587.62 | 1154778.25 | 2 | 3617140.5 | 2 | 3338981.5 | 3240471.5 | 2 | 1254413.38 | 2 | 810724.81 | 766662.56 | 2546607.25 | 984145.88 | 1903160.5 | 1209254.88 | 856153 | 2 | 668892.06 | 2334931.75 |
| Q8BZN6 | Dock10 | Dedicator of cytokinesis protein 10 | 72.52 | 1.65 | 249576 | 2 | 2 | 2 | 2 | 682002.5 | 2 | 2 | 2 | 2 | 2 | 586307.62 | 2 | 2 | 931587.56 | 540198.94 | 1211701.75 | 1034300.19 | 2 | 2 | 2 | 2 |
| Q8C3J5 | Dock2 | Dedicator of cytokinesis protein 2 | 303.39 | 4.6 | 211704 | 6 | 760057 | 676735.75 | 1636756 | 2395105 | 838721.81 | 1049229.75 | 1279967.88 | 1899265.12 | 1399970.25 | 1925301.5 | 1427600.25 | 661648.81 | 1276214.12 | 1373077.25 | 2786278.25 | 94422.57 | 1154339 | 995532.94 | 1505613.75 | 2626678.25 |
| Q8C147 | Dock8 | Dedicator of cytokinesis protein 8 | 91.29 | 1.1 | 238978 | 2 | 2653438.5 | 2026465.25 | 2961496 | 2843847.5 | 2720138 | 2585858.5 | 1289854.75 | 2 | 1773110.25 | 1349753.75 | 675902.75 | 605989.25 | 2 | 2 | 2003271.75 | 909702 | 2 | 1123261.5 | 1156242.38 | 1304775.38 |
| O70152 | Dpm1 | Dolichol-phosphate mannosyltransferase subunit | 32.73 | 8.46 | 29175 | 1 | 2 | 2 | 343657.75 | 2 | 331968.75 | 201520.62 | 706037.25 | 217647.58 | 301428.78 | 484543.69 | 68886.53 | 2 | 2 | 2 | 338690.94 | 2 | 2 | 2 | 53020.79 | 181976.73 |
| Q9D1Q4 | Dpm3 | Dolichol-phosphate mannosyltransferase subunit | 44.45 | 10.87 | 10139 | 1 | 3945883.25 | 2948939.5 | 3786439.5 | 3286844.25 | 2369528.75 | 3319600 | 2792770.75 | 4626073 | 1710608 | 2349878.5 | 1862478.38 | 1529527.75 | 2017991.75 | 1664096.88 | 2142684.25 | 1919664.88 | 3174364 | 4153257 | 2097101 | 3610791.75 |
| Q99KK7 | Dpp3 | Dipeptidyl peptidase 3 | 156.56 | 6.91 | 82898 | 3 | 962061.69 | 1363304.38 | 1349153 | 923749.69 | 3251290.5 | 3118119 | 3016987 | 2112753 | 2426007.5 | 1615922.25 | 1147504.75 | 2716079.75 | 2021257.5 | 2632604.5 | 2337838.75 | 2761944.5 | 1066895.25 | 1954686.75 | 773891.31 | 1741632.12 |
| Q9ET22 | Dpp7 | Dipeptidyl peptidase 2 | 103.2 | 8.1 | 56254 | 2 | 219521.67 | 427215.03 | 1206399 | 230317.77 | 111702.36 | 281315.66 | 2 | 122926 | 211241.08 | 209527.14 | 1475270.12 | 129042.91 | 304643.41 | 1109936.12 | 2560766 | 2 | 1531263.75 | 2 | 3231001.75 | 453801.97 |
| O08553 | Dpysl2 | Dihydropyrimidinase-related protein 2 | 1213.69 | 45.45 | 62278 | 18 | 37916970 | 61949206 | 36791490 | 57069602 | 81409818 | 57630126 | 50552214 | 74241450 | 62825138 | 53524082 | 64908898 | 68809138 | 53034934 | 55139554 | 45906418 | 83832890 | 60697826 | 59619330 | 76217482 | 60155534 |
| P32233 | Drg1 | Developmentally-regulated GTP-binding protein 1 | 175.45 | 12.81 | 40512 | 4 | 2 | 1028853.94 | 1470820.38 | 985166.31 | 417233.94 | 75580.82 | 130172.73 | 2 | 2 | 526585.62 | 1142197 | 1480585.62 | 2542329 | 2435134.5 | 2250585.25 | 1115609.88 | 1227403 | 1277250.88 | 1265857.75 | 1428072.12 |
| Q9QXB9 | Drg2 | Developmentally-regulated GTP-binding protein 2 | 197.97 | 14.01 | 40718 | 4 | 2 | 2 | 2 | 336463.69 | 288797.31 | 151012.02 | 628972.88 | 73437.12 | 492612.25 | 2 | 104563.53 | 136479.3 | 938354.69 | 937262.5 | 534418.25 | 2 | 138872.48 | 117800.88 | 2 | 421481.81 |
| E9Q557 | Dsp | Desmoplakin | 110.2 | 0.9 | 332912 | 3 | 332594.94 | 1274575.38 | 2 | 2795645.5 | 2 | 617669.81 | 743440.69 | 504469.06 | 2 | 1823220.88 | 1683168.5 | 2 | 2 | 234995.5 | 2 | 2 | 2 | 2 | 1517889.88 | 2 |
| Q7M0E3 | Dstn | Destrin | 264.71 | 37.58 | 18534 | 5 | 4196600.5 | 3060336.25 | 4590758 | 4310259 | 4519185 | 7666249 | 4288128 | 6777347 | 3951379 | 3282883.75 | 18839586 | 14956345 | 20056462 | 15165702 | 17590094 | 21653850 | 26320282 | 22759698 | 23829482 | 20930826 |
| Q9D7X3 | Dusp3 | Dual specificity protein phosphatase 3 | 338.48 | 28.11 | 20472 | 4 | 6790830 | 11850266 | 6409896.5 | 8638889 | 7945000 | 8845125 | 7341637 | 10241188 | 9318432 | 8444439 | 5625802 | 6012172 | 4370785 | 5446378 | 4121944 | 4924042 | 3329517.75 | 4188902 | 4017292.5 | 3708265.75 |
| Q9CQ43 | Dut | Deoxyuridine 5'-triphosphate nucleotidohydrolase | 220.22 | 34.57 | 17385 | 4 | 2 | 2 | 2 | 249950.97 | 2234611.5 | 2 | 2 | 2 | 2 | 2 | 1712360.25 | 1147168 | 2 | 307786.78 | 2 | 2 | 489782.94 | 2 | 2 | 2 |
| Q9JHU4 | Dync1h1 | Cytoplasmic dynein 1 heavy chain 1 | 3268.821 | 15.7 | 532045 | 63 | 48216002 | 51349602 | 72903882 | 47213374 | 57321802 | 46383662 | 41602634 | 56363270 | 38491654 | 52130802 | 49494186 | 47506494 | 55017866 | 48757254 | 56971734 | 40366078 | 40930726 | 47820138 | 36628994 | 32126934 |
| O88487 | Dync1i2 | Cytoplasmic dynein 1 intermediate chain 2 | 43.49 | 1.63 | 68394 | 1 | 2 | 2 | 103584.25 | 2 | 2 | 2 | 2 | 2 | 146130.98 | 2 | 2 | 2 | 2 | 54983.18 | 2 | 2 | 2 | 2 | 2 | 2 |
| Q8R1Q8 | Dync1li1 | Cytoplasmic dynein 1 light intermediate chain 1 | 196.89 | 7.07 | 56614 | 3 | 2 | 2 | 2 | 2 | 572640.12 | 1033193.12 | 2 | 1105852.12 | 729851.94 | 585884 | 1620736.88 | 1790539 | 1380740.62 | 1500818.12 | 628672.25 | 2 | 384929.12 | 2 | 1479280.62 | 261687.52 |
| P62627 | Dynlrb1 | Dynein light chain roadblock-type 1 | 86.54 | 21.88 | 10990 | 2 | 638098.94 | 1458284 | 822269.06 | 693453.31 | 1465202.75 | 1371334.38 | 874332.94 | 1647039.25 | 783915.25 | 822466 | 3148254.5 | 2631294.5 | 2535769 | 2852504.25 | 1696800.88 | 2851766 | 2721931.75 | 2206940 | 2968842 | 3380210.75 |
| Q9D903 | Ebna1bp2 | Probable rRNA-processing protein EBP2 | 71.43 | 3.92 | 34703 | 1 | 2 | 2 | 657359.44 | 2 | 919740.38 | 2 | 2 | 592636 | 2 | 620188.06 | 2 | 2 | 2 | 2 | 2 | 2 | 2 | 2 | 2 | 2 |
| P70245 | Ebp | 3-beta-hydroxysteroid-Delta(8),Delta(7)-isomeras | 39 | 4.35 | 26215 | 1 | 1458888.75 | 1253762.62 | 1372635.25 | 1448347.38 | 660188.38 | 2340941.25 | 1890605.38 | 3047401.75 | 1556552.62 | 1001341.44 | 1985055.38 | 1303740.62 | 1441539.5 | 1429502.12 | 1062944.12 | 3799011.5 | 2099697.5 | 2112185 | 2574937.25 | 1613916.5 |
| O35459 | Ech1 | Delta(3,5)-Delta(2,4)-dienoyl-CoA isomerase, mit | 276.35 | 16.21 | 36118 | 4 | 8338462 | 8295967 | 7909071.5 | 5378734 | 7024098 | 1339023.5 | 2326403.5 | 3594912.5 | 4821097 | 3781058 | 5632368 | 5147330 | 7614112 | 5267336.5 | 4753896 | 4370487 | 4753312 | 5710850.5 | 4788116 | 6006393 |
| P14604 | Echs1 | Enoyl-CoA hydratase, mitochondrial | 203.73 | 15.86 | 31516 | 3 | 2 | 958898.12 | 2 | 1298543.38 | 4483370.5 | 4647332 | 2999480 | 3193655.5 | 2766131 | 3656071.75 | 1356210.25 | 1435149 | 315711.09 | 248051.05 | 2 | 3118906 | 265329 | 2 | 2 | 459306.69 |
| P42125 | Eci1 | Enoyl-CoA delta isomerase 1, mitochondrial | 243.58 | 17.99 | 32250 | 4 | 91506.73 | 342071 | 488017.53 | 386377.5 | 493984.72 | 331076.34 | 530816.38 | 471024.25 | 722603.31 | 857499.81 | 1563667.75 | 1875117.38 | 1010825.12 | 1778161.38 | 2323970.25 | 52746.92 | 954491.12 | 790500.75 | 2 | 865943.75 |
| Q9WUR2 | Eci2 | Enoyl-CoA delta isomerase 2 | 87.37 | 7.93 | 43268 | 2 | 2 | 1114586.12 | 927425.81 | 1533723.12 | 698365.19 | 2 | 854756.5 | 1078548.25 | 1004109.69 | 1321046.75 | 719105.75 | 2 | 1666261.38 | 1310907 | 5513411 | 2901443.5 | 2 | 2 | 511523.66 | 2456691 |
| Q6PDI5 | Ecpas | Proteasome adapter and scaffold protein ECM29 | 150.09 | 1.63 | 203915 | 2 | 61495.48 | 62395.95 | 627383.44 | 2 | 539180.81 | 222733.95 | 76136.41 | 115455.01 | 2 | 550267.69 | 89484.35 | 2 | 97001.63 | 776253.88 | 2 | 2 | 617333.44 | 2 | 2 | 2 |
| Q3UJB9 | Edc4 | Enhancer of mRNA-decapping protein 4 | 229.39 | 4.27 | 152484 | 4 | 682731.44 | 1134807.5 | 1302500.25 | 1118666.5 | 2093876.75 | 100394.34 | 1720722.62 | 2531048.5 | 1536827.25 | 2485260.5 | 615591.75 | 2 | 776018.81 | 1558078.5 | 1071635.62 | 751530.12 | 164770.02 | 948482.81 | 604275.38 | 493288.75 |
| Q8BL66 | Eea1 | Early endosome antigen 1 | 129.17 | 2.2 | 160915 | 3 | 2 | 2 | 2 | 2 | 1178439.62 | 2 | 2 | 2 | 2 | 301107.91 | 2958061.5 | 3319228.5 | 1242859.5 | 639464.38 | 590381.06 | 2924144.5 | 928224.56 | 1139135.62 | 1824110.5 | 824188.31 |
| P10126 | Eef1a1 | Elongation factor 1-alpha 1 | 1553.77 | 47.62 | 50114 | 25 | 235127010 | 368272994 | 103546554 | 504683650 | 535634114 | 347666114 | 210210274 | 372848194 | 244765074 | 510729762 | 798043010 | 793966850 | 589089090 | 679891202 | 458987650 | 779988098 | 249989506 | 462731010 | 591355650 | 417772162 |
| O70251 | Eef1b | Elongation factor 1-beta | 133.94 | 9.33 | 24694 | 3 | 24819446 | 32500400 | 18243986 | 33297950 | 20686492 | 23362874 | 19326118 | 27289778 | 21648878 | 14441360 | 12696932 | 18720810 | 19401668 | 19631648 | 13200890 | 19176992 | 17034398 | 17035716 | 20108024 | 22651642 |
| P57776 | Eef1d | Elongation factor 1-delta | 669.69 | 36.3 | 31293 | 10 | 55972342 | 56603278 | 49338410 | 54788842 | 53468642 | 42996234 | 41529478 | 47330046 | 34967330 | 43419870 | 29211538 | 30174042 | 34276694 | 32177534 | 27318584 | 26016820 | 27842364 | 25829278 | 29964362 | 31675226 |
| Q9D1M4 | Eef1e1 | Eukaryotic translation elongation factor 1 epsilon- | 48.99 | 4.02 | 19859 | 1 | 2 | 408585.62 | 2 | 263159.56 | 957276.94 | 2082507.12 | 1313865.5 | 2 | 1218317.75 | 1010542.19 | 1199568.62 | 1288425.12 | 594759.06 | 1078047.25 | 2 | 1071801.25 | 2 | 2 | 952162.44 | 2 |
| Q9D8N0 | Eef1g | Elongation factor 1-gamma | 777.49 | 34.1 | 50061 | 14 | 33165534 | 47718378 | 42995058 | 45807882 | 50173314 | 43774034 | 41547794 | 56753538 | 41285366 | 39635074 | 39371550 | 46991454 | 40413354 | 41727050 | 42073010 | 53877742 | 43776738 | 46687426 | 48268434 | 44974822 |

| | | | | | | | | | | | | | | | | | | | | | | | | | | |
|---|---|---|---|---|---|---|---|---|---|---|---|---|---|---|---|---|---|---|---|---|---|---|---|---|---|---|
| P58252 | Eef2 | Elongation factor 2 | 2380.61 | 51.05 | 95314 | 39 | 31392468 | 140272562 | 63522438 | 52072334 | 121441002 | 108663418 | 112441618 | 96731738 | 100908458 | 77359082 | 126938458 | 128735154 | 95301002 | 96078858 | 54199662 | 76776666 | 59538882 | 39720238 | 59064954 | 37246422 |
| Q9JHW4 | Eefsec | Selenocysteine-specific elongation factor | 108.81 | 4.8 | 63539 | 2 | 2 | 575482.12 | 2 | 844075.25 | 961347.19 | 827221.75 | 2 | 2 | 839834.19 | 875189.88 | 2067091.88 | 1987311.62 | 2 | 2 | 557989.19 | 1101233.12 | 1070689.12 | 2 | 2 | 2 |
| Q4FZY0 | Efhd2 | EF-hand domain-containing protein D2 | 923.07 | 54.81 | 26759 | 16 | 27441146 | 30060710 | 40557162 | 28768302 | 43913890 | 26766554 | 24984000 | 35640258 | 27070610 | 35918846 | 34234806 | 34456370 | 36509866 | 27057910 | 47368010 | 45285722 | 31374786 | 46245714 | 38968082 | 37293930 |
| O08810 | Eftud2 | 116 kDa U5 small nuclear ribonucleoprotein comp | 324.33 | 6.8 | 109361 | 6 | 1687209.25 | 6091546.5 | 2031990.88 | 5727883 | 12020878 | 6639329 | 4418242 | 8894335 | 7520296.5 | 7625877 | 4730162 | 1726124.25 | 4439691 | 5790952 | 1773366.25 | 5040510.5 | 4095159.5 | 3472898 | 5972961 | 2635821.5 |
| Q99MS7 | Ehbp1l1 | EH domain-binding protein 1-like protein 1 | 52.4 | 0.76 | 184834 | 1 | 746429.62 | 2 | 729282.5 | 2 | 627420.38 | 151211.81 | 648485.31 | 210187.55 | 511596.62 | 434878.41 | 2 | 407742.69 | 491908.56 | 2 | 2 | 88428.33 | 2 | 2 | 2 | 192061.28 |
| Q641Z6 | Ehd1 | EH domain-containing protein 1 | 355.52 | 14.61 | 60603 | 6 | 4805933 | 2401201.25 | 3549479.75 | 4232502.5 | 6519411 | 5149916 | 5525972.5 | 4818188 | 6126069 | 5602845 | 2091652.25 | 3060500.5 | 4505352 | 4612000 | 2028395.75 | 2839914.5 | 2917296 | 6704579.5 | 1543373.25 | 5210615 |
| Q9EQP2 | Ehd4 | EH domain-containing protein 4 | 397.99 | 15.53 | 61481 | 8 | 3857906.5 | 4967847 | 5644242 | 3550824.5 | 10134876 | 4227047 | 7004378.5 | 4080913.5 | 7819204.5 | 11722443 | 943233.81 | 2586093 | 2960140.75 | 4725046 | 3202818.5 | 894422.75 | 1547067.75 | 2744305.75 | 544961.44 | 2734872.75 |
| Q60872 | Eif1a | Eukaryotic translation initiation factor 1A | 59.88 | 11.11 | 16502 | 1 | 468152.47 | 1139875 | 577039.5 | 1280119.62 | 1447810 | 1743473.88 | 1313507.62 | 1323046.12 | 887667.69 | 718357.94 | 655403.38 | 842482.31 | 417214.56 | 2 | 2 | 2 | 769518.81 | 865724.75 | 553352.31 | 1067576.12 |
| Q8BJW6 | Eif2a | Eukaryotic translation initiation factor 2A | 180.76 | 10.33 | 64403 | 5 | 1601284.75 | 1439726.12 | 91780.69 | 3884518.25 | 2937196.25 | 944455.44 | 3440685 | 1492725.62 | 2261429.5 | 1016754 | 1378484 | 6439541 | 1112030 | 2093338 | 1636502 | 6294336 | 1145088.12 | 3724049.25 | 1139483.75 | 1839304.88 |
| Q03963 | Eif2ak2 | Interferon-induced, double-stranded RNA-activate | 84.96 | 4.27 | 58280 | 2 | 113844.98 | 1406506.75 | 2 | 1251380.62 | 2 | 1814995.38 | 2039660.75 | 120781.25 | 1248917.5 | 2 | 1037266.56 | 1602601.62 | 575521.69 | 943089.81 | 2 | 1848189.12 | 2114956.5 | 2 | 184235.06 | 2 |
| P70541 | Eif2b3 | Translation initiation factor eIF2B subunit gamma | 95.76 | 4.65 | 50435 | 2 | 820850.56 | 150481.12 | 692254.19 | 2 | 915067.12 | 209918.22 | 526455.38 | 113517.87 | 688907.62 | 417285.69 | 94837.93 | 533514.31 | 824625.62 | 392010.72 | 261188.92 | 2 | 2 | 2 | 2 | 2 |
| Q61749 | Eif2b4 | Translation initiation factor eIF2B subunit delta | 225.13 | 8.59 | 57624 | 4 | 2368358 | 1109515.25 | 3271612.5 | 607534.19 | 3657095 | 1938666 | 1815537.75 | 6323597.5 | 2396646.75 | 1863831.5 | 1399339.62 | 1428178.62 | 2340696.75 | 2128744 | 1138847.5 | 1064318.5 | 2 | 1452736.5 | 2 | 2 |
| P68101 | Eif2s1 | Eukaryotic translation initiation factor 2 subunit 1 | 396.54 | 27.3 | 36108 | 8 | 9737491 | 11147962 | 13358570 | 11767718 | 21204608 | 20977982 | 18835206 | 27986656 | 17727166 | 15158821 | 29855744 | 23828778 | 26518606 | 24949358 | 14651443 | 22947946 | 15905980 | 13793750 | 22748670 | 5482553.5 |
| Q99L45 | Eif2s2 | Eukaryotic translation initiation factor 2 subunit 2 | 244.87 | 8.46 | 38092 | 5 | 4167082.5 | 4643582 | 2449438 | 6060031.5 | 7350135 | 8797467 | 9735414 | 14109189 | 12231153 | 5757658.5 | 11731560 | 9318759 | 10532087 | 9323444 | 6594662.5 | 10411003 | 9204718 | 9059762 | 13275035 | 8546372 |
| Q9Z0N1 | Eif2s3x | Eukaryotic translation initiation factor 2 subunit 3, | 751.19 | 33.26 | 51065 | 13 | 3247721 | 4302019 | 6345901 | 5795777.5 | 10736560 | 7081690.5 | 8304664 | 11182894 | 7308629 | 8965900 | 22658074 | 26041514 | 16102489 | 16729072 | 9312756 | 16178032 | 12088381 | 9674222 | 17527564 | 14084660 |
| P23116 | Eif3a | Eukaryotic translation initiation factor 3 subunit A | 601.01 | 9.75 | 161936 | 13 | 4492523.5 | 5516413 | 4311230.5 | 3401262 | 4692750 | 2638923.75 | 4322498 | 6232332.5 | 5873450.5 | 4500970.5 | 11102821 | 9795734 | 10696893 | 11979127 | 7383088.5 | 9240019 | 8334341 | 7508762 | 8914103 | 7389175 |
| Q8JZQ9 | Eif3b | Eukaryotic translation initiation factor 3 subunit B | 430.23 | 16.69 | 91370 | 8 | 9718182 | 11081439 | 7008297 | 9498532 | 4613059.5 | 6968340 | 7184258 | 5523813 | 8801638 | 4621960 | 8290778 | 8790962 | 11475785 | 10307301 | 4603188 | 8750947 | 10792210 | 6862862 | 10687162 | 4684503.5 |
| B5DFC8 | Eif3c | Eukaryotic translation initiation factor 3 subunit C | 373.81 | 8.34 | 105435 | 8 | 7501966 | 4810217 | 7973480.5 | 7757198.5 | 5960281.5 | 1273142.88 | 6716745 | 3186139.25 | 5445673 | 5462474 | 3980322.25 | 5058765.5 | 3837151.5 | 4776934 | 5655003.5 | 3713854.5 | 2466684.5 | 5199586.5 | 3294042.5 | 4888435 |
| O70194 | Eif3d | Eukaryotic translation initiation factor 3 subunit D | 347.33 | 14.96 | 63989 | 6 | 870616.75 | 6633475 | 1223850.5 | 4293388.5 | 3775019.75 | 2682591.75 | 3605745.5 | 3394510.25 | 4373787 | 2072952.25 | 1595879.38 | 3900963.25 | 2693783 | 4012594 | 593191.31 | 713361.62 | 1035198 | 1835027.75 | 2103004.5 | 2648227 |
| P60229 | Eif3e | Eukaryotic translation initiation factor 3 subunit E | 369.64 | 11.69 | 52221 | 5 | 16118402 | 19712942 | 17804114 | 22389140 | 34142374 | 12629628 | 12169330 | 20120472 | 11413941 | 20813110 | 10963845 | 9694226 | 11030013 | 11987529 | 9341992 | 8119557 | 8050870 | 10818258 | 7636755.5 | 7956243 |
| Q9DCH4 | Eif3f | Eukaryotic translation initiation factor 3 subunit F | 384.57 | 22.99 | 37984 | 7 | 14369273 | 22208262 | 6738653.5 | 26393614 | 21863214 | 15409306 | 19125446 | 24053910 | 19119418 | 14968120 | 14282862 | 8573835 | 13062416 | 10154292 | 7833597.5 | 8106138 | 9167058 | 10332428 | 10971346 | 7900344 |
| Q9Z1D1 | Eif3g | Eukaryotic translation initiation factor 3 subunit G | 267.82 | 22.5 | 35638 | 6 | 813718.94 | 645834.88 | 1068401.25 | 1489206.38 | 1473741.88 | 2024314.62 | 3544878.5 | 679236.44 | 2623053 | 1785731.62 | 3110392.5 | 3249961.5 | 3699167.5 | 3425830 | 5599095.5 | 2150807.5 | 4024842 | 5127011.5 | 159070.23 | 2042310.25 |
| Q91WK2 | Eif3h | Eukaryotic translation initiation factor 3 subunit H | 182.9 | 7.95 | 39832 | 3 | 94022.71 | 277678.41 | 157881.14 | 288477.88 | 254432.16 | 82897.69 | 43657.49 | 369548.25 | 821735.62 | 58627.43 | 118537.26 | 669953 | 158209.44 | 687217.19 | 261040.91 | 500065.62 | 2 | 2 | 132959.86 | 481963.5 |
| B0BNA7 | Eif3i | Eukaryotic translation initiation factor 3 subunit I | 172.75 | 11.69 | 36461 | 4 | 29398016 | 3801085 | 30653224 | 4850883.5 | 27350562 | 4235234 | 29940402 | 3139829.75 | 3701199.5 | 4153632.75 | 6618297 | 8789262 | 9933462 | 8919910 | 7780716.5 | 7491304.5 | 8128444.5 | 8576985 | 8626110 | 8295092.5 |
| A0JPM9 | Eif3j | Eukaryotic translation initiation factor 3 subunit J | 62.78 | 5.02 | 29187 | 1 | 550046.69 | 743268.56 | 835484.25 | 1031586.94 | 491166.31 | 2 | 2 | 1106442.25 | 722213.5 | 859801.31 | 710534.31 | 2 | 1044826.88 | 668109.5 | 1000757.69 | 927135.81 | 2 | 988481.12 | 827719.19 | 2 |
| Q9DBZ5 | Eif3k | Eukaryotic translation initiation factor 3 subunit K | 48.02 | 5.05 | 25087 | 1 | 2 | 1064259.25 | 719059.44 | 2 | 691208.75 | 2 | 632382.88 | 857385.19 | 2 | 873066.38 | 2 | 2 | 2 | 2 | 1165680.5 | 2 | 2 | 2 | 2 | 2 |
| Q8QZY1 | Eif3l | Eukaryotic translation initiation factor 3 subunit L | 325.45 | 11.52 | 66613 | 6 | 2225394.5 | 3086551 | 2302216 | 2182133 | 1281392.12 | 813082.06 | 474588.06 | 503564.09 | 1988165.25 | 2983264.25 | 2014352.5 | 1748102.5 | 3635363 | 3587046 | 2821920 | 1316842.5 | 2698696.75 | 3472049.75 | 2631345.25 | 3439474.5 |
| Q99JX4 | Eif3m | Eukaryotic translation initiation factor 3 subunit M | 159.29 | 12.83 | 42517 | 3 | 769050 | 637726.88 | 1035220.56 | 2086154.25 | 1062012.38 | 849435.44 | 2 | 1634080.25 | 1360891.38 | 662229.62 | 843225.81 | 605761.69 | 1432683.38 | 2 | 2 | 2 | 2 | 1090515 | 469843.91 | 2 |
| P60843 | Eif4a1 | Eukaryotic initiation factor 4A-I | 1184.98 | 53.94 | 46154 | 18 | 3275947 | 2629167.75 | 6765980 | 3103604.5 | 5576520 | 2458810.5 | 3121775 | 4032822.75 | 1917453.5 | 5094446.5 | 2450592.75 | 1340812.12 | 2173055 | 1289845 | 5445591.5 | 1534033.75 | 1091110.25 | 1629938 | 1270441.25 | 2532110.5 |
| Q91VC3 | Eif4a3 | Eukaryotic initiation factor 4A-III | 616.41 | 26.28 | 46840 | 10 | 3054452.75 | 6760473 | 3182738.5 | 3949801.5 | 4715227.5 | 3386967.5 | 4413072 | 1828835.88 | 10273836 | 2731342.75 | 6137024.5 | 7396773.5 | 3106662.5 | 6151579 | 3090563.5 | 2090045.25 | 2328141 | 2196940 | 1150477.88 | 2755944 |
| Q8BGD9 | Eif4b | Eukaryotic translation initiation factor 4B | 606.48 | 24.71 | 68840 | 11 | 5471462 | 8752870 | 3847775.75 | 6176553 | 6746817.5 | 4252517.5 | 3177989.5 | 5791324 | 8221768 | 4665329 | 17471482 | 17679420 | 8943103 | 12901642 | 4497128.5 | 14066219 | 7615061.5 | 5568347 | 11003616 | 7491857.5 |
| Q6NZJ6 | Eif4g1 | Eukaryotic translation initiation factor 4 gamma 1 | 539.48 | 8.56 | 176077 | 11 | 1183375.38 | 4206122.5 | 3073318.25 | 4049755 | 4895371 | 2623924.25 | 3864915.75 | 5631895 | 3101459 | 3806596.25 | 8121578.5 | 8778624 | 7193809 | 9520462 | 5708514.5 | 7098221.5 | 3593569 | 5049929 | 7026229 | 4477652 |
| Q62448 | Eif4g2 | Eukaryotic translation initiation factor 4 gamma 2 | 226.32 | 6.29 | 102106 | 5 | 2 | 2 | 2 | 377597.38 | 811663.25 | 319010.91 | 2 | 2 | 626304.31 | 2 | 1701719.25 | 1118285.25 | 415981.38 | 898113 | 2 | 671024.38 | 2 | 2 | 518289.22 | 782418.31 |
| Q80XI3 | Eif4g3 | Eukaryotic translation initiation factor 4 gamma 3 | 271.43 | 4.43 | 174890 | 6 | 551762.5 | 1148956.62 | 1353157.12 | 610006.31 | 1307916.38 | 94152.57 | 656862.62 | 141346.08 | 702728.06 | 251572.77 | 2205656.75 | 2798218.5 | 1735903.12 | 2367263.25 | 898998.31 | 2558340.5 | 2390761 | 840707.38 | 3345698.5 | 1848001.88 |
| P59325 | Eif5 | Eukaryotic translation initiation factor 5 | 161.56 | 8.39 | 48968 | 4 | 2 | 995849.88 | 1382684.5 | 2 | 1492343.38 | 1391790 | 1202094.5 | 1308877.75 | 877351 | 1482653.25 | 1827858.25 | 1831967.5 | 3076929 | 1890178.75 | 1359873.5 | 1275414.88 | 1222210.62 | 1034740.81 | 1568585.5 | 924941.81 |
| P63242 | Eif5a | Eukaryotic translation initiation factor 5A-1 | 454.02 | 36.36 | 16832 | 6 | 6262015.5 | 5722508 | 14606396 | 11419841 | 13472228 | 12225256 | 13059533 | 10451734 | 16719640 | 8481649 | 9715258 | 8496223 | 6835777 | 8116053 | 17703304 | 6333765 | 8806931 | 6820018 | 7034282.5 | 6622637.5 |
| B2GUV7 | Eif5b | Eukaryotic translation initiation factor 5B | 134.42 | 4.52 | 137686 | 3 | 1088965.25 | 2 | 761629.5 | 225619.33 | 1908740 | 202106.62 | 177335.77 | 305126.25 | 1483288.62 | 743271 | 790192.38 | 827222.81 | 2153009.75 | 1329438.25 | 1035589.56 | 585664.06 | 906818.12 | 1079347.75 | 1132401.75 | 1725894 |
| O55135 | Eif6 | Eukaryotic translation initiation factor 6 | 276.19 | 19.59 | 26511 | 3 | 7538163.5 | 9948550 | 9240904 | 8452255 | 10800786 | 7323474.5 | 7117014 | 11026257 | 6045598.5 | 6752291 | 10245484 | 9418166 | 8258942.5 | 7098880.5 | 8002164.5 | 8959047 | 8692982 | 8629110 | 11200599 | 9098348 |
| Q80Y81 | Elac2 | Zinc phosphodiesterase ELAC protein 2 | 54.74 | 0.96 | 92719 | 1 | 2188174.5 | 1917915 | 478612.25 | 2766630.5 | 1154082.75 | 2782130.25 | 2824285.75 | 2444696 | 2 | 1154281.12 | 1866317.62 | 2121689 | 1959001.62 | 2 | 1689062.75 | 1792127.38 | 1109059.25 | 2 | 2 | 2111450.5 |
| B5DF91 | Elavl1 | ELAV-like protein 1 | 488.16 | 27.3 | 36169 | 9 | 21195206 | 24887008 | 19642276 | 26699298 | 31745138 | 24547870 | 24290250 | 33821718 | 21071814 | 21814548 | 16274388 | 15319762 | 13510757 | 9994121 | 15633048 | 9066060 | 11128088 | 10306440 | 9135048 | 9834043 |
| Q8BPU7 | Elmo1 | Engulfment and cell motility protein 1 | 86.86 | 2.75 | 83936 | 2 | 1261092.5 | 2677358 | 888784.38 | 2019837.5 | 1862457.88 | 569955 | 1050510.88 | 747948.94 | 568839.75 | 2504942 | 734539.94 | 2 | 2 | 762506.5 | 1646194.75 | 2 | 2294865.25 | 2115635.75 | 866796.88 | 1160407.62 |
| P62869 | Elob | Elongin-B | 100.81 | 16.1 | 13170 | 2 | 2 | 1430083.5 | 1271782.88 | 2 | 3174588 | 1914904.62 | 2400366 | 2564377 | 1348510.5 | 2764509 | 3293361 | 2464179.5 | 1062854.75 | 2417170.75 | 4218186 | 1824549.62 | 2265853.25 | 3238447.25 | 1174368.25 | 1963015.75 |
| P83940 | Eloc | Elongin-C | 167.27 | 29.46 | 12473 | 2 | 2 | 2 | 2 | 2 | 991887.62 | 2 | 694817.06 | 1007463.06 | 735680.88 | 591223.5 | 407574.47 | 774484.81 | 551533.19 | 1375435.88 | 319337.62 | 2 | 582546.62 | 873253.12 | 946587.5 | 880469.88 |
| Q8C7X2 | Emc1 | ER membrane protein complex subunit 1 | 582.9 | 17.55 | 111605 | 12 | 3029162 | 7569389 | 3784771 | 4879401.5 | 2587064.75 | 4447169 | 5111298.5 | 5576943 | 5293076 | 3686880 | 6236766 | 3671591 | 6994168.5 | 5080568 | 3332956.75 | 2332676.25 | 6046814 | 9272906 | 1968889.25 | 7232666.5 |
| B0BNG0 | Emc2 | ER membrane protein complex subunit 2 | 48.06 | 4.04 | 34870 | 1 | 399889.25 | 498415.53 | 2 | 470525.28 | 687011.44 | 710401.25 | 1011525.31 | 3221412 | 910349.75 | 1411488.25 | 808292.56 | 2 | 735148.75 | 678884.75 | 1821959.25 | 2 | 750811.62 | 970587.31 | 559651.81 | 2 |
| Q5U2V8 | Emc3 | ER membrane protein complex subunit 3 | 139.38 | 16.09 | 29980 | 3 | 2 | 2 | 2 | 2 | 62549.83 | 114377.25 | 216315.56 | 82831.26 | 529238.75 | 2 | 131252.95 | 2 | 104548.59 | 2 | 2 | 2 | 2 | 2 | 2 | 2 |
| Q9CZX9 | Emc4 | ER membrane protein complex subunit 4 | 45.42 | 5.46 | 20117 | 1 | 2 | 2 | 527464.19 | 2 | 2 | 2 | 2 | 904676.94 | 537893.38 | 346392.59 | 2 | 2 | 2 | 464085.19 | 2 | 1072215 | 625486.5 | 2 | 2 | 2 |
| Q9CQW0 | Emc6 | ER membrane protein complex subunit 6 | 34.33 | 6.36 | 12017 | 1 | 262444 | 1414839.12 | 2 | 2 | 2 | 2 | 1133251.25 | 2 | 1665914.38 | 1078678.25 | 2 | 2 | 913380.19 | 2102225.5 | 605245.06 | 2 | 1060937.88 | 1655878.12 | 2 | 1493074 |
| O70378 | Emc8 | ER membrane protein complex subunit 8 | 203.37 | 18.36 | 23348 | 3 | 3672414 | 1596709.88 | 3345747 | 2985693.5 | 2930835 | 3038294.5 | 2025328.88 | 2900971.75 | 1646451 | 1949447.75 | 2040335.75 | 72591.63 | 1908215.38 | 2378224 | 1863712.25 | 3281090 | 2197201.25 | 1400702 | 1888443.25 | 2 |
| O08579 | Emd | Emerin | 128.04 | 12.74 | 29436 | 3 | 2 | 1614465 | 1850077.75 | 2087081.25 | 2759155.5 | 2 | 469035.59 | 73369.82 | 2286075.75 | 2899406 | 2335935.25 | 2383991 | 1925649.75 | 1879203.25 | 2 | 1375442.75 | 1695607.5 | 1050035 | 1348275.75 | 2 |
| O35130 | Emg1 | Ribosomal RNA small subunit methyltransferase | 52.03 | 4.1 | 26974 | 1 | 2 | 2 | 762485.31 | 2 | 2 | 2 | 2 | 644649.06 | 2 | 2 | 2 | 2 | 2 | 264165.34 | 2 | 378719.38 | 2 | 2 | 2 | 645794.69 |
| Q6P6T4 | Eml2 | Echinoderm microtubule-associated protein-like 2 | 67.14 | 3.24 | 70711 | 1 | 718982 | 410968.59 | 980690.81 | 2 | 1235409.25 | 619532.25 | 652241.69 | 501691.12 | 1879890.25 | 1320338.62 | 500941.62 | 687030.69 | 1742531 | 762114.38 | 899776.31 | 318621.62 | 806246.38 | 764060.19 | 2 | 2 |
| Q8VC03 | Eml3 | Echinoderm microtubule-associated protein-like 3 | 34.26 | 1.56 | 95695 | 1 | 1289132.5 | 2 | 404504.06 | 2 | 390514.66 | 2 | 2 | 646904.94 | 2 | 872126.5 | 2 | 2 | 1003925.38 | 633878.81 | 866441.75 | 2 | 2 | 1047452.88 | 2 | 2 |
| P17182 | Eno1 | Alpha-enolase | 1996.97 | 65.21 | 47141 | 28 | 18871678 | 20500014 | 31773610 | 13039551 | 31881922 | 24144164 | 30472144 | 38017822 | 39410174 | 51412194 | 53396754 | 57394990 | 46220606 | 38302958 | 50591174 | 65843842 | 43127166 | 35574562 | 41043554 | 31954728 |
| P17183 | Eno2 | Gamma-enolase | 509.23 | 20.05 | 47297 | 6 | 2 | 2 | 2 | 2 | 2 | 2 | 2 | 1131578.12 | 865662.12 | 2 | 2 | 1364816.5 | 784846.81 | 2 | 2 | 2 | 2 | 2 | 2 | 2 |
| Q8BYW9 | Eogt | EGF domain-specific O-linked N-acetylglucosami | 132.31 | 5.31 | 61446 | 3 | 374883.53 | 688966.31 | 104459.12 | 612042.75 | 2 | 749919.62 | 485618.81 | 2 | 2 | 2 | 641993.75 | 562650.25 | 644598.38 | 2 | 2 | 2 | 2 | 2 | 2 | 2 |
| O70318 | Epb41l2 | Band 4.1-like protein 2 | 298.75 | 10.43 | 109940 | 6 | 852296 | 4730681 | 7958770 | 2 | 2418073 | 2 | 2 | 1250615.25 | 2441056.5 | 431105.09 | 5041017 | 5139002.5 | 11135032 | 7011486 | 395793.78 | 9072128 | 2314726 | 2 | 10893033 | 2 |
| Q8CGC7 | Eprs1 | Bifunctional glutamate/proline--tRNA ligase | 1856.19 | 25.73 | 170079 | 32 | 14743673 | 25172832 | 21510540 | 32631328 | 51516306 | 27958166 | 26187834 | 28165730 | 39155294 | 45221974 | 54631246 | 53252894 | 40964334 | 45416786 | 22752440 | 49331074 | 35605386 | 31617296 | 41877726 | 34070370 |
| P42567 | Eps15 | Epidermal growth factor receptor substrate 15 | 57.54 | 1.34 | 98471 | 1 | 478077.12 | 170217.27 | 593845.94 | 251958.88 | 308871.16 | 78573.1 | 2 | 2 | 557743.19 | 2 | 2 | 2 | 2 | 2 | 2 | 2 | 373569.22 | 2 | 2 | 210251.42 |
| Q60902 | Eps15l1 | Epidermal growth factor receptor substrate 15-like | 35.09 | 0.88 | 99309 | 1 | 430351.59 | 1008090.62 | 741108.12 | 753788.5 | 1225748.62 | 703667.38 | 773916.19 | 2 | 2 | 1043974.62 | 2 | 2 | 2 | 2 | 987447.94 | 977157.69 | 2 | 522861 | 2 | 384905.62 |
| Q9EQH2 | Erap1 | Endoplasmic reticulum aminopeptidase 1 | 429.82 | 10.22 | 106599 | 9 | 4172599.5 | 2289690.75 | 2976944 | 6837048.5 | 8261080 | 3263968.75 | 2421891.25 | 3739057 | 7065165 | 5880967.5 | 1837220.5 | 2720313 | 3605527.25 | 3305518 | 634957.88 | 2152775.75 | 2318369 | 1171257 | 2357924.25 | 642357.25 |
| Q9DC16 | Ergic1 | Endoplasmic reticulum-Golgi intermediate compa | 42.45 | 3.45 | 32562 | 1 | 697680.62 | 671538.06 | 411739.53 | 550431.94 | 794284.56 | 1662543.5 | 1299834.62 | 3129918.25 | 888176 | 2 | 1210114 | 1181084.38 | 1222404.12 | 940654.5 | 734632.19 | 1225969 | 950859.62 | 940519.69 | 961146.62 | 1002520.81 |
| Q9CQE7 | Ergic3 | Endoplasmic reticulum-Golgi intermediate compa | 77.33 | 6.01 | 43208 | 2 | 2 | 497838.16 | 1708803.12 | 2 | 225254.61 | 2 | 911523.62 | 814794.56 | 460935.22 | 565383.56 | 963720.31 | 794002.62 | 904201.12 | 741500.5 | 476723.31 | 710166.31 | 791905.94 | 472389.62 | 468892 | 938396.62 |
| B5DEH2 | Erlin2 | Erlin-2 | 247.74 | 12.68 | 37710 | 4 | 2027174.75 | 1370976.5 | 1819136.5 | 1351364.5 | 2319287.5 | 862299.88 | 713583.5 | 775014.06 | 981843.25 | 2322503 | 563624.38 | 2072434.75 | 1284852.25 | 704813.88 | 2 | 782337.31 | 2 | 1904130.25 | 2 | 1474409.75 |
| Q8R180 | Ero1a | ERO1-like protein alpha | 859.21 | 31.25 | 54084 | 16 | 3717368.75 | 2176000.5 | 5885350.5 | 4745550.5 | 4152475 | 9543592 | 17777838 | 16568958 | 13682502 | 24260640 | 5008791 | 3599334.75 | 3860243.75 | 4575956 | 8753255 | 5950897.5 | 10286246 | 9318237 | 6567005.5 | 8104663 |
| P57759 | Erp29 | Endoplasmic reticulum resident protein 29 | 363.96 | 32.82 | 28823 | 8 | 5431505 | 12407815 | 5713145 | 8786215 | 6681324 | 4454186 | 3798938.75 | 4711707.5 | 6048345 | 5583751.5 | 5509057.5 | 6566905 | 6109407.5 | 5290510 | 3273372 | 5159876 | 3681351.75 | 4101669 | 4248636 | 2532350 |
| Q9D1Q6 | Erp44 | Endoplasmic reticulum resident protein 44 | 59.79 | 3.45 | 46853 | 1 | 2 | 2 | 2 | 2 | 2 | 2 | 2 | 2 | 2 | 2 | 2 | 2 | 2 | 2 | 2 | 2 | 2 | 2 | 2 | 2 |
| Q9R0P3 | Esd | S-formylglutathione hydrolase | 944.79 | 56.38 | 31320 | 13 | 25940540 | 26521614 | 25612578 | 20582998 | 29878124 | 51622646 | 40259646 | 62494562 | 48655582 | 44654690 | 56858138 | 55370386 | 65995338 | 55665290 | 47097394 | 97382994 | 77140818 | 65375730 | 94525538 | 61382050 |
| Q3U7R1 | Esyt1 | Extended synaptotagmin-1 | 480.21 | 12 | 121554 | 8 | 1244597.25 | 630825.94 | 615018.06 | 1669567.75 | 1273106.88 | 1853735.62 | 2335907.25 | 569408.38 | 926064.94 | 1979313.25 | 511136.19 | 978330.25 | 1350477.12 | 2685980.5 | 2536759 | 615425.75 | 2518874 | 3253591.75 | 905112.5 | 4127044.5 |
| Q3TZZ7 | Esyt2 | Extended synaptotagmin-2 | 105.4 | 3.31 | 94139 | 2 | 2 | 536690.69 | 81658.23 | 2 | 2 | 2 | 710881.94 | 2 | 2 | 442886.53 | 2 | 2 | 91195.18 | 118583.01 | 182174.75 | 2 | 2 | 2 | 2 | 2 |
| Q5U2Q7 | Etf1 | Eukaryotic peptide chain release factor subunit 1 | 227.15 | 11.44 | 49031 | 4 | 3757929 | 3953695.25 | 5297127.5 | 4717134.5 | 6760934.5 | 2989044 | 4771884.5 | 3185173.5 | 3824166 | 3867261.25 | 2439071.25 | 3950169.5 | 3343839.75 | 2787527.75 | 3521739 | 1511522.75 | 2352325.75 | 2378620.5 | 977832.44 | 2471458 |
| Q99LC5 | Etfa | Electron transfer flavoprotein subunit alpha, mitoc | 879.7 | 60.96 | 35009 | 14 | 19234188 | 21016160 | 25011634 | 13869950 | 28862888 | 8286004 | 13759056 | 10803474 | 23768692 | 27181398 | 14614708 | 15529617 | 17529696 | 15859666 | 17909524 | 12465851 | 18049480 | 17697652 | 11540176 | 15831889 |
| Q9DCW4 | Etfb | Electron transfer flavoprotein subunit beta | 755.97 | 52.16 | 27623 | 14 | 2698588.25 | 1421493 | 3948559.5 | 2315896.5 | 4298695 | 506458.75 | 1770016 | 455143.06 | 1906440.62 | 4488118.5 | 1020515.31 | 1340219 | 1940479.75 | 1622355.75 | 2890063 | 449466.47 | 1667629.5 | 922056.75 | 4551244.5 | 1005520 |
| Q921G7 | Etfdh | Electron transfer flavoprotein-ubiquinone oxidore | 291.02 | 11.85 | 68091 | 6 | 2509600.75 | 1083471 | 642800.88 | 952283.62 | 2 | 694799.88 | 135267.56 | 317937.66 | 1563220.5 | 2498943.5 | 2916740.25 | 2980252.75 | 1763176 | 3357184.5 | 1178410.25 | 2571413.75 | 1573560.12 | 1434587.62 | 180469.5 | 3557995 |
| Q9DCM0 | Ethe1 | Persulfide dioxygenase ETHE1, mitochondrial | 220.14 | 21.26 | 27739 | 4 | 2 | 947891.25 | 1885262.75 | 440054.94 | 459103.72 | 873082.25 | 446937.44 | 1021857.31 | 507680.41 | 398993.56 | 4417527.5 | 4371790 | 1240820.12 | 4079207.5 | 531892.19 | 2820368 | 2 | 1913820.5 | 4533672 | 2 |
| P20934 | Evi2a | Protein EVI2A | 74 | 9.42 | 24087 | 1 | 2 | 2 | 2 | 2 | 263744.59 | 2 | 2 | 2 | 2 | 2 | 2 | 2 | 2 | 2 | 2 | 2 | 2 | 2 | 2 | 2 |
| Q61545 | Ewsr1 | RNA-binding protein EWS | 163.55 | 7.79 | 68462 | 4 | 2516684 | 5769972.5 | 3893577 | 5500519.5 | 5770154.5 | 2943024.75 | 3337977.75 | 5527041.5 | 5005994 | 2632041.5 | 7841040.5 | 8740657 | 3308176 | 11424795 | 6100432.5 | 5448617.5 | 6211537 | 5226900 | 4811289 | 7632509 |
| Q9D4H1 | Exoc2 | Exocyst complex component 2 | 77.85 | 1.84 | 103959 | 2 | 2 | 2 | 775453.19 | 315386.62 | 1258085.12 | 2 | 2 | 114482.5 | 1080840.5 | 417258.75 | 2 | 319927.78 | 459108.75 | 410139.12 | 367205.62 | 2 | 2 | 2 | 2 | 173072.27 |
| O54924 | Exoc8 | Exocyst complex component 8 | 74.19 | 3.35 | 81043 | 2 | 1017636.62 | 2345197 | 1041333.56 | 839897.62 | 1686102 | 2 | 2040086.88 | 2976509.5 | 1638862.5 | 1518030.12 | 2194252.75 | 1826530.5 | 1766561.25 | 1611301.25 | 815278.75 | 2122011 | 936613.81 | 2 | 1359206.88 | 2 |
| P56960 | Exosc10 | Exosome complex component 10 | 42.58 | 1.47 | 100942 | 1 | 1044002.44 | 840726.56 | 928552.62 | 1003989.94 | 877234.62 | 457106.22 | 1213071.5 | 717949.81 | 415453.72 | 598944.25 | 363504.47 | 575011.06 | 750764.75 | 2127314.75 | 899195 | 2824528.25 | 1938223.88 | 1557320.88 | 3020617 | 1147283.38 |
| Q8VBV3 | Exosc2 | Exosome complex component RRP4 | 111.12 | 5.46 | 32632 | 2 | 2 | 2 | 2 | 2 | 2 | 2 | 2 | 2 | 337273.06 | 2 | 2 | 2 | 2 | 2 | 2 | 2 | 2 | 2 | 2 | 2 |
| Q8BTW3 | Exosc6 | Exosome complex component MTR3 | 69.73 | 5.86 | 28370 | 1 | 2 | 371385.78 | 2 | 680378.12 | 2 | 951921 | 2 | 2 | 685672.31 | 2 | 673544.44 | 2 | 2 | 2 | 632621.25 | 2 | 2 | 2 | 2 | 2 |
| Q9D753 | Exosc8 | Exosome complex component RRP43 | 42.54 | 6.88 | 29949 | 1 | 1228962.75 | 794661 | 2 | 1132313 | 1475762.5 | 1165836.5 | 2 | 974935.88 | 853291.75 | 2 | 2 | 2 | 800317.25 | 2 | 2 | 1511911.12 | 2 | 2 | 1322586.38 | 2055975.62 |
| P26040 | Ezr | Ezrin | 659.1 | 17.24 | 69407 | 12 | 2 | 2 | 2 | 2 | 2 | 2 | 2 | 2 | 2 | 2 | 1154697.5 | 2 | 2 | 584392.44 | 2 | 2 | 2 | 2 | 2 | 2 |
| P18292 | F2 | Prothrombin | 88.51 | 2.59 | 70412 | 2 | 2 | 2 | 2 | 1593031.75 | 737138.25 | 2 | 2 | 95171.05 | 752121.56 | 663502.75 | 2 | 2 | 397535.91 | 2 | 2 | 2 | 2 | 2 | 2 | 2 |
| Q05816 | Fabp5 | Fatty acid-binding protein 5 | 670.01 | 64.44 | 15137 | 10 | 34245050 | 34500042 | 50742414 | 9556888 | 10992836 | 39574022 | 37961902 | 48729858 | 5096551 | 6893748 | 79571738 | 91973610 | 85874338 | 68784394 | 83715738 | 54366678 | 84873338 | 70096202 | 73705058 | 54591358 |
| Q3TDN2 | Faf2 | FAS-associated factor 2 | 51.71 | 2.7 | 52471 | 1 | 2 | 375419.66 | 2 | 764058.25 | 2 | 630737.75 | 2 | 2 | 2 | 485179.53 | 794711.19 | 584989.44 | 2 | 2 | 2 | 2 | 2 | 2 | 2 | 2 |
| P35505 | Fah | Fumarylacetoacetase | 55.4 | 2.86 | 46176 | 1 | 2 | 2 | 2 | 2 | 1011573 | 2 | 2 | 2 | 843195.94 | 2 | 1874743.25 | 1458046.12 | 2 | 1185484.88 | 2 | 1209699.5 | 2 | 2 | 735634.19 | 2 |
| Q3TC72 | Fahd2a | Oxaloacetate tautomerase Fahd2a, mitochondrial | 39.87 | 3.51 | 34676 | 1 | 2 | 2 | 2 | 2 | 326108.38 | 2 | 2 | 2 | 2 | 2 | 754840.44 | 2 | 2 | 763890.38 | 2 | 2 | 2 | 2 | 2 | 2 |
| Q8VE88 | Fam114a2 | Protein FAM114A2 | 51.33 | 1.81 | 54045 | 1 | 2 | 2 | 2 | 2 | 440594.12 | 2 | 2 | 2 | 503267.75 | 2 | 2 | 2 | 2 | 2 | 2 | 2 | 2 | 2 | 2 | 2 |
| Q6A0A9 | FAM120A | Constitutive coactivator of PPAR-gamma-like prot | 46.93 | 0.99 | 121646 | 1 | 2 | 2 | 2 | 2 | 160986.61 | 2 | 396864.84 | 82297.96 | 190614.39 | 2 | 125421.01 | 2 | 481829.94 | 293867.56 | 2 | 2 | 2 | 2 | 2 | 2 |
| Q9CR98 | Fam136a | Protein FAM136A | 198.25 | 25.36 | 15674 | 4 | 5526609.5 | 13855725 | 4868453 | 5712001 | 3245322.5 | 14823904 | 13813124 | 4842778 | 11233259 | 7938137 | 13952224 | 15466217 | 10882027 | 14375035 | 7422468 | 10991867 | 15795834 | 13205040 | 13374438 | 12085440 |
| Q810F4 | Fam3c | Protein FAM3C | 94.23 | 11.01 | 24714 | 2 | 2 | 758772.19 | 2 | 3657105 | 146601.3 | 1072585.12 | 2 | 1057453.38 | 3684981.5 | 2 | 2 | 2 | 3846548.25 | 4207430.5 | 2 | 2 | 2 | 911421.62 | 2 | 2 |

| | | | | | | | | | | | | | | | | | | | | | | | | | | |
|---|---|---|---|---|---|---|---|---|---|---|---|---|---|---|---|---|---|---|---|---|---|---|---|---|---|---|
| Q66H50 | Far1 | Fatty acyl-CoA reductase 1 | 465.82 | 19.22 | 59268 | 9 | 7513762 | 3707635 | 6519636 | 5714426.5 | 3053391.5 | 2548047.5 | 1694603.5 | 4584293 | 2635502.75 | 3404838.5 | 516316.78 | 2 | 4560184.5 | 2672104.5 | 2263162 | 2 | 1210468.88 | 4050888 | 866485.19 | 1166498.38 |
| Q8C0C7 | Farsa | Phenylalanine--tRNA ligase alpha subunit | 215.64 | 7.68 | 57599 | 4 | 3600885.75 | 2811630.25 | 4019623.25 | 1439490.62 | 3243220 | 1472546 | 991522.06 | 3088438.5 | 1013014.31 | 3128988.75 | 1834197 | 535715.38 | 1501343.38 | 1303359.62 | 2087507.75 | 660146.69 | 360409.69 | 775595.56 | 821682.5 | 2 |
| Q9WUA2 | Farsb | Phenylalanine--tRNA ligase beta subunit | 397.74 | 14.77 | 65697 | 9 | 8228052.5 | 9936640 | 10939334 | 12441193 | 27447930 | 13104995 | 7810103.5 | 15776366 | 12783254 | 16009206 | 12624519 | 16383986 | 11696362 | 16390624 | 11321425 | 13012436 | 14303867 | 14585278 | 15916648 | 16891354 |
| P19096 | Fasn | Fatty acid synthase | 1767 | 16.57 | 272428 | 36 | 17312400 | 24271658 | 38467318 | 15947310 | 55195218 | 17710234 | 17258050 | 48926214 | 10636876 | 23792868 | 16955904 | 17329470 | 19926082 | 20126160 | 37415934 | 18805980 | 15017302 | 13284740 | 17497128 | 14697933 |
| P62862 | Fau | Ubiquitin-like FUBI-ribosomal protein eS30 fusion | 40.7 | 7.52 | 14416 | 1 | 2 | 2 | 2 | 2 | 781201 | 2 | 2 | 764022.31 | 1195101.75 | 2 | 2 | 2 | 2 | 559601.62 | 2 | 2 | 2 | 2 | 2 | 2 |
| P22509 | Fbl | rRNA 2'-O-methyltransferase fibrillarin | 507.67 | 39.45 | 34222 | 9 | 939790.69 | 3651883.5 | 1039457.06 | 2869129 | 9344312 | 1652853.25 | 5250978 | 2713877 | 11360373 | 5593778 | 11051714 | 13313134 | 10059814 | 11597807 | 4485952.5 | 12271223 | 9423556 | 4107783.25 | 5634924 | 5668565.5 |
| Q78JE5 | Fbxo22 | F-box only protein 22 | 56.19 | 2.24 | 44203 | 1 | 513828.25 | 2 | 2 | 2 | 510410.97 | 670892.31 | 655319.5 | 480013.31 | 1239180.25 | 457518.94 | 923267.75 | 747837.06 | 969258.12 | 2 | 821296.38 | 1092166.12 | 1689552.25 | 1506619.62 | 1130237.62 | 1725798.38 |
| Q9QZN4 | Fbxo6 | F-box only protein 6 | 48.17 | 3.73 | 34493 | 1 | 2 | 631808.06 | 2 | 2 | 354971.72 | 2 | 239741.08 | 2 | 2 | 276621.09 | 2 | 2 | 2 | 2 | 244209.09 | 643411.38 | 2 | 673071.12 | 518347.91 | 796192.62 |
| P20491 | Fcer1g | High affinity immunoglobulin epsilon receptor sub | 153.72 | 23.26 | 9652 | 3 | 2 | 931609.12 | 2 | 1123922.88 | 2157792.5 | 3271061.5 | 1921324.88 | 3008396.5 | 5067920.5 | 1606688.75 | 4555421 | 5484753 | 4560800.5 | 4709836 | 2091373.12 | 4432980 | 3293572.75 | 2238744.25 | 4874449 | 2163969.5 |
| P26151 | Fcgr1 | High affinity immunoglobulin gamma Fc receptor I | 130.07 | 8.66 | 44888 | 3 | 1108650.38 | 1011444.88 | 1563863.75 | 1690631.62 | 1458674.88 | 1637133.38 | 1676078.12 | 2194248.25 | 1163304.12 | 1290947.12 | 933770.94 | 828623.44 | 2 | 371523.06 | 683253.31 | 2 | 2 | 2 | 2 | 2 |
| P08101 | Fcgr2 | Low affinity immunoglobulin gamma Fc region rec | 136.91 | 12.12 | 36695 | 3 | 775619.88 | 2 | 2056802 | 2 | 775474 | 2 | 2 | 1303259.75 | 2199028.25 | 2056051.88 | 817119 | 1038085.06 | 1072622.75 | 127375.96 | 155823.17 | 2086079.88 | 2 | 2 | 2 | 2 |
| P08508 | Fcgr3 | Low affinity immunoglobulin gamma Fc region rec | 105.92 | 8.05 | 30036 | 2 | 2566354.25 | 1890575.12 | 2654166.25 | 2185767.75 | 2148224 | 3068961.5 | 1595041 | 2115634 | 2448457.25 | 2290078.5 | 2947196.75 | 3321333.5 | 2816845.5 | 3080279 | 3746763.75 | 3012151 | 2572532.25 | 2589931.75 | 525771.38 | 4302111 |
| A0A0B4 | Fcgr4 | Low affinity immunoglobulin gamma Fc region rec | 36.81 | 4.02 | 28398 | 1 | 947983.94 | 2 | 447056.06 | 637118.75 | 491916.41 | 1068709.5 | 669145 | 2 | 342697.09 | 2 | 664022.19 | 2 | 2 | 2 | 2 | 797233.62 | 2 | 691152.75 | 662073.12 | 2 |
| Q920E5 | Fdps | Farnesyl pyrophosphate synthase | 742.65 | 35.69 | 40582 | 11 | 2788142.25 | 1404335 | 8257495.5 | 5937261 | 7903806 | 3471377.25 | 6351176 | 2878236.75 | 3018149.25 | 8617563 | 3085041.5 | 3056091.5 | 4798622 | 4415102.5 | 11790018 | 2186025.25 | 4054210 | 3343156.25 | 2608239 | 1528961.62 |
| Q61578 | Fdxr | NADPH:adrenodoxin oxidoreductase, mitochondr | 212.12 | 12.96 | 54202 | 5 | 2 | 2 | 2 | 2 | 2 | 748379.62 | 2 | 2 | 662311.81 | 397170.91 | 2 | 491595.25 | 2 | 2 | 2 | 1644353 | 2 | 562433.56 | 4214871 | 2277529.25 |
| P39749 | Fen1 | Flap endonuclease 1 | 254.77 | 7.67 | 42315 | 3 | 2 | 216223.81 | 1905165.62 | 679281.31 | 1593438.88 | 197147.48 | 562011.12 | 94704.61 | 524055.94 | 1108385.5 | 122188.42 | 2 | 824779.12 | 635463.38 | 1304063.25 | 2 | 2 | 118383.57 | 2 | 150763.84 |
| Q8K1B8 | Fermt3 | Fermitin family homolog 3 | 747.53 | 27.82 | 75635 | 13 | 20451748 | 13721778 | 20066230 | 16604394 | 19826126 | 15104583 | 21203478 | 21247850 | 15783103 | 16276047 | 7946444.5 | 10412487 | 17001436 | 12269220 | 19921732 | 6981652 | 17846418 | 17006730 | 9807466 | 17088752 |
| P97807 | Fh | Fumarate hydratase, mitochondrial | 1048.52 | 50.89 | 54357 | 18 | 28124744 | 48843710 | 32662490 | 44863306 | 50119422 | 34446382 | 30612378 | 28356746 | 40579486 | 36512094 | 32292050 | 26534302 | 31853222 | 33343596 | 22832628 | 26352352 | 25246714 | 32258082 | 31258660 | 28148224 |
| Q9R059 | Fhl3 | Four and a half LIM domains protein 3 | 39.38 | 3.46 | 31795 | 1 | 2 | 2 | 2 | 2 | 658314.12 | 2 | 2 | 2 | 816534.38 | 2 | 2 | 824699.88 | 432544.28 | 2 | 2 | 999118.44 | 589060.94 | 2 | 990300.81 | 1065048.25 |
| Q5U317 | Fip1l1 | Pre-mRNA 3'-end-processing factor FIP1 | 206.97 | 9.51 | 60188 | 4 | 2 | 2 | 2 | 2 | 2 | 2 | 2 | 2 | 2 | 2 | 106809.36 | 99565.51 | 2 | 684300.81 | 2 | 310581.06 | 2 | 2 | 2 | 2 |
| P84817 | Fis1 | Mitochondrial fission 1 protein | 187.35 | 21.05 | 16995 | 2 | 133246.78 | 541411.38 | 335517.06 | 770010.62 | 242022.56 | 193991.62 | 324281.44 | 176011.06 | 903861.38 | 183449.81 | 2 | 54500.27 | 273708.59 | 277534.69 | 536543.12 | 2 | 385358.5 | 278508.66 | 2 | 543697.56 |
| P26883 | Fkbp1a | Peptidyl-prolyl cis-trans isomerase FKBP1A | 164 | 34.26 | 11923 | 3 | 1820123.38 | 1370842.75 | 1429936.5 | 1151773.5 | 1995983.62 | 2 | 2 | 2217760 | 707807.06 | 765213.62 | 14284028 | 16163495 | 10632491 | 10729690 | 6113886 | 5814974 | 6636910.5 | 6188723 | 8938636 | 6504553 |
| P45878 | Fkbp2 | Peptidyl-prolyl cis-trans isomerase FKBP2 | 148.08 | 23.57 | 15344 | 3 | 2389241 | 4266183 | 2757367.5 | 971771.75 | 2312398.5 | 1720126 | 1019130.19 | 2 | 900704.94 | 1700529.38 | 3945045 | 4463114.5 | 1715068.75 | 3911644 | 1309687.88 | 6121918 | 2584167.5 | 5057554.5 | 4174388.25 | 5385950.5 |
| P30416 | Fkbp4 | Peptidyl-prolyl cis-trans isomerase FKBP4 | 490.44 | 27.73 | 51572 | 11 | 9026196 | 15987230 | 9500416 | 14165276 | 29807284 | 7154258.5 | 8932034 | 16707499 | 11599939 | 11551719 | 7792342.5 | 8838470 | 7062017 | 9838070 | 10124208 | 3712817.75 | 8358066.5 | 8754953 | 7100808 | 5947055 |
| Q64378 | Fkbp5 | Peptidyl-prolyl cis-trans isomerase FKBP5 | 271.97 | 14.25 | 50966 | 7 | 2 | 945829.69 | 2 | 4136526 | 9293367 | 1959838.25 | 91566.31 | 3055993.75 | 7751744 | 5343818 | 7512550 | 6334003 | 7396227.5 | 8466415 | 3896969.25 | 6404583 | 3971553 | 4070388.25 | 9069918 | 3343920.5 |
| O35465 | Fkbp8 | Peptidyl-prolyl cis-trans isomerase FKBP8 | 392.72 | 16.42 | 43529 | 7 | 1099362.5 | 1504291.12 | 540495.12 | 2 | 5272758 | 646357.38 | 113416.18 | 634406.88 | 2367571.75 | 1924173.5 | 1901796.25 | 854775.31 | 2 | 1413535.62 | 664305.12 | 1091670.25 | 2 | 116595.34 | 2 | 492936.47 |
| Q9JJ28 | Flii | Protein flightless-1 homolog | 668.81 | 14.48 | 144803 | 13 | 3291807.5 | 1568800.5 | 3144041.5 | 3671381.5 | 5718925.5 | 3305109.5 | 3163912 | 5612757.5 | 7782939.5 | 3629112.75 | 2125260.25 | 3158985.5 | 4067326 | 3433852.75 | 3198227.25 | 3190101.5 | 3118076.75 | 2066691 | 2587004.25 | 2417998.25 |
| Q8BTM8 | Flna | Filamin-A | 4042.23 | 33.36 | 281222 | 69 | 67233162 | 91182226 | 83583186 | 30546066 | 42653442 | 105562506 | 105359938 | 160628706 | 82822562 | 75044834 | 128851098 | 143336658 | 99317506 | 108401874 | 90491330 | 174092962 | 126538906 | 102280722 | 133019034 | 118139626 |
| Q80X90 | Flnb | Filamin-B | 386.48 | 4.38 | 277825 | 9 | 1131113.5 | 3068476.25 | 1578489.5 | 1299448.12 | 1630904 | 5051342.5 | 3173963.25 | 5835995 | 2369210 | 1863617.75 | 1922865.38 | 1705994.5 | 2372185 | 1040371.56 | 898243.12 | 2723106.5 | 3625364 | 2130807 | 3131857 | 2485479.75 |
| O08917 | Flot1 | Flotillin-1 | 203.81 | 10.28 | 47513 | 4 | 880823.19 | 2 | 751238.25 | 203121.92 | 371118.22 | 2 | 109398.95 | 2 | 427738.22 | 255092.84 | 1676264 | 2 | 1068739 | 1212267.88 | 272825.38 | 2328233.75 | 1103875.75 | 2 | 1318183.5 | 237808.22 |
| Q9JL26 | Fmnl1 | Formin-like protein 1 | 875.4 | 18.1 | 122060 | 17 | 11775747 | 13101773 | 15454244 | 14395907 | 17995738 | 9846187 | 8344708.5 | 6498806 | 12363955 | 12501307 | 14271600 | 13471452 | 12837996 | 12017207 | 8079108 | 8110180.5 | 8716334 | 11555913 | 7678054 | 9919609 |
| Q6ZPF4 | Fmnl3 | Formin-like protein 3 | 208.05 | 4.18 | 117169 | 4 | 56726.83 | 116361.52 | 88101.53 | 357481.81 | 127981.75 | 2 | 163685.3 | 2 | 221505.67 | 161552.89 | 99536.05 | 75011.66 | 95282.38 | 168056.34 | 2 | 2 | 137750.98 | 195887.95 | 2 | 197539.77 |
| P35922 | Fmr1 | Fragile X messenger ribonucleoprotein 1 | 55.5 | 2.28 | 68989 | 1 | 2 | 59840.74 | 177555.25 | 2 | 117576.31 | 2 | 2 | 2 | 163269.72 | 2 | 78963.84 | 97324.18 | 117820.79 | 245184.12 | 2 | 2 | 256910.16 | 280461.5 | 158066.94 | 2 |
| Q80TY0 | Fnbp1 | Formin-binding protein 1 | 242.96 | 8.93 | 71344 | 6 | 1092009.38 | 1223677.5 | 2 | 744231.5 | 1045396.5 | 1015549.69 | 2 | 1836836.88 | 1055742.5 | 822667.25 | 1259399.75 | 2026984.25 | 1266639.75 | 2015013.88 | 661937.19 | 2 | 997421.12 | 2 | 2 | 636295.62 |
| Q6NWW9 | Fndc3b | Fibronectin type III domain-containing protein 3B | 98.67 | 2.98 | 132764 | 2 | 2 | 2 | 2 | 2 | 2 | 2 | 2 | 159719.3 | 2 | 664641.19 | 2 | 2 | 2 | 2 | 2 | 67932.3 | 2 | 2 | 2 | 2 |
| P97376 | Frg1 | Protein FRG1 | 38.13 | 5.04 | 29127 | 1 | 2 | 2 | 2 | 2 | 456082.84 | 2 | 525140.56 | 2 | 595037.06 | 2 | 2 | 727802.81 | 437270.25 | 439005.03 | 2 | 817224.81 | 2 | 2 | 2 | 2 |
| P09528 | Fth1 | Ferritin heavy chain | 87.34 | 8.79 | 21067 | 2 | 21016034 | 26122112 | 8789650 | 7839957.5 | 2467544 | 33741338 | 32217188 | 9495758 | 10437348 | 14162244 | 13423802 | 11080311 | 10667943 | 9986929 | 3063430.25 | 9194916 | 12454194 | 11041384 | 8616868 | 9829627 |
| P29391 | Ftl1 | Ferritin light chain 1 | 646.51 | 57.92 | 20802 | 10 | 69129738 | 69147330 | 63428554 | 20708536 | 17695390 | 40861622 | 49445170 | 39498018 | 18479588 | 30884226 | 25487714 | 19431178 | 25413020 | 23620834 | 26720972 | 29735662 | 29694076 | 34582122 | 24641600 | 25988994 |
| Q5RJT2 | Ftsj3 | pre-rRNA 2'-O-ribose RNA methyltransferase FTS | 160.83 | 7 | 94767 | 3 | 57163.5 | 70092.2 | 2 | 310705.47 | 173215.77 | 2 | 2 | 242184.97 | 189459.47 | 246083.64 | 2 | 337634.75 | 243592.42 | 281504.47 | 557750.31 | 2 | 258004.05 | 140738.78 | 2 | 2 |
| Q32PX7 | Fubp1 | Far upstream element-binding protein 1 | 611.73 | 20.97 | 67197 | 13 | 4716346 | 4036515 | 2221352.5 | 2167109.5 | 5243058 | 5748444.5 | 3232238.5 | 5802619 | 1983143.88 | 2286603.5 | 3739216.75 | 6782242.5 | 5193938.5 | 2883986.25 | 2455527.75 | 5874498 | 3253761.75 | 3650152.5 | 5294242.5 | 5589919.5 |
| Q8R2K1 | Fuom | Fucose mutarotase | 61.23 | 7.84 | 16805 | 1 | 2 | 2 | 204250 | 2 | 226592.38 | 2 | 2 | 2 | 2 | 222760.53 | 2 | 71943.52 | 209200.94 | 501048.19 | 568715.06 | 2 | 349889.97 | 2 | 165525.36 | 206795.44 |
| P56959 | Fus | RNA-binding protein FUS | 277.57 | 9.46 | 52673 | 5 | 4722045.5 | 2623942.75 | 3459410.5 | 6689119.5 | 7706957 | 4450891 | 2467391 | 4798334 | 7134277 | 2405473 | 7504534.5 | 4856388.5 | 8097868 | 4483741 | 5486300 | 6884028.5 | 6614484 | 11005154 | 10234220 | 7049801 |
| Q61584 | Fxr1 | RNA-binding protein FXR1 | 406.55 | 15.66 | 76222 | 9 | 8808032 | 8497144 | 5026144.5 | 11200847 | 10557697 | 7564829 | 6638563 | 7644761 | 5718660 | 6321714 | 9245533 | 6600738.5 | 7887739.5 | 6220573.5 | 4660092.5 | 7512621 | 6998661 | 10730890 | 6878676.5 | 7297158.5 |
| Q9WVR4 | Fxr2 | RNA-binding protein FXR2 | 108.63 | 5.35 | 73743 | 2 | 2 | 2679955.5 | 2 | 583827.19 | 1767519 | 2 | 2 | 812980.69 | 1702662.12 | 2660214.75 | 2 | 620730.25 | 2 | 2345360.25 | 1683995.12 | 2048425.25 | 1562708 | 2 | 1637007.12 | 2 |
| P97808 | Fxyd5 | FXYD domain-containing ion transport regulator 5 | 52.18 | 5.06 | 19454 | 1 | 5102960.5 | 5245896.5 | 3498225 | 5173172 | 2297217.5 | 5844122 | 5089434 | 8485950 | 5437924.5 | 307377.34 | 2659923.75 | 2663847.75 | 3128726.25 | 2925531.5 | 2691481.75 | 4619991 | 5138508 | 3590548 | 6272845.5 | 4680410.5 |
| D3ZIE4 | Fyb1 | FYN-binding protein 1 | 67.93 | 1.08 | 91412 | 1 | 437376.72 | 449949.84 | 2 | 1104212 | 2 | 897346.12 | 928093.06 | 829328.56 | 1742339.62 | 2 | 734331.31 | 890967 | 1182367.62 | 649541.88 | 628818 | 917970.81 | 802852.19 | 860394.31 | 795286.06 | 2 |
| P97855 | G3bp1 | Ras GTPase-activating protein-binding protein 1 | 416.46 | 21.94 | 51829 | 8 | 2828582.5 | 3678635.5 | 3172076.25 | 2826818.75 | 2906179 | 2091237.88 | 1358502.25 | 1242791.25 | 2259942.5 | 2201344 | 4617295 | 5067650.5 | 5027289 | 4846177.5 | 3790235.5 | 5474398 | 4820392.5 | 2039262.38 | 5806616.5 | 3681098 |
| P97379 | G3bp2 | Ras GTPase-activating protein-binding protein 2 | 140.14 | 9.75 | 54088 | 3 | 295010 | 2 | 153760.98 | 68832.12 | 98367.58 | 2 | 2 | 275132.19 | 1735140.38 | 93032.45 | 1518125.75 | 1703441.25 | 1470523.62 | 1702373.12 | 485916.84 | 2053765.25 | 2142269.5 | 185319.78 | 2867176.25 | 2 |
| Q00612 | G6pdx | Glucose-6-phosphate 1-dehydrogenase X | 909.04 | 38.64 | 59263 | 18 | 25215466 | 33521372 | 27085950 | 24584892 | 37825638 | 53109550 | 45463218 | 45528342 | 36914922 | 37657726 | 61965706 | 65615970 | 45767062 | 45735122 | 40423842 | 60551838 | 53268030 | 37265554 | 59610586 | 45628554 |
| P70699 | Gaa | Lysosomal alpha-glucosidase | 550.97 | 16.79 | 106248 | 10 | 3150447 | 4555249.5 | 2791608.5 | 2046871 | 2295129.75 | 1452528.75 | 2434992.25 | 1140688.62 | 2324658.25 | 1778663 | 454174.06 | 2184825.25 | 1292401 | 3200160.75 | 2851640.75 | 1192431.12 | 1538123 | 2043327.75 | 1303333.12 | 3120684.75 |
| P60517 | Gabarap | Gamma-aminobutyric acid receptor-associated pr | 108.35 | 15.38 | 13918 | 2 | 2 | 70469.18 | 166450.73 | 84299.91 | 123186.77 | 106486.11 | 357737.12 | 39801.54 | 378841.28 | 178983.41 | 2 | 2 | 235144.97 | 85997.6 | 438716.5 | 2 | 172479.59 | 2 | 2 | 2 |
| Q9CR59 | Gadd45gip1 | Large ribosomal subunit protein mL64 | 63.67 | 7.66 | 25820 | 2 | 2 | 2 | 2 | 2 | 2 | 2 | 2 | 2 | 2 | 2 | 880363.56 | 814936 | 2 | 1854396.62 | 2 | 807515.75 | 2 | 2 | 2 | 2 |
| P18645 | Gale | UDP-glucose 4-epimerase | 44.95 | 3.17 | 38225 | 1 | 2 | 2 | 2 | 2 | 2 | 2 | 2 | 2 | 2 | 2 | 503196.28 | 838057.94 | 493855.56 | 2 | 2 | 2 | 2 | 2 | 2 | 2 |
| Q9R0N0 | Galk1 | Galactokinase | 315.15 | 14.54 | 42295 | 7 | 2 | 1757560.62 | 2253764.75 | 1047678.38 | 4286003 | 2638388 | 3953308 | 6729657 | 5417976 | 4555444.5 | 9983930 | 8559553 | 7913805.5 | 5801622.5 | 5493793 | 4574603 | 2306349.5 | 3694550 | 4247227 | 2753370.5 |
| Q5XIG6 | Galk2 | N-acetylgalactosamine kinase | 49.59 | 1.75 | 50198 | 1 | 1790036.12 | 2169998.25 | 1881155.62 | 566085.94 | 1083897.5 | 1514201.25 | 1089314.62 | 1211159.5 | 1218538.88 | 1004982.88 | 1266486.75 | 1310556.38 | 1157307 | 1113477.62 | 1330901.75 | 1150338.5 | 1252012.12 | 1950160.38 | 190331.12 | 1605081.88 |
| Q571E4 | Galns | N-acetylgalactosamine-6-sulfatase | 290.25 | 14.23 | 57673 | 7 | 5198303 | 5460391.5 | 4994883.5 | 3325586.5 | 3415042 | 4418275 | 4112612 | 2915524 | 3378284.5 | 4786504 | 2335514.25 | 2095480.12 | 1110362.75 | 2764975.75 | 6015533 | 1913348 | 3933784.5 | 4620054.5 | 1216433 | 2979700 |
| Q80VA0 | Galnt7 | N-acetylgalactosaminyltransferase 7 | 129 | 4.87 | 75419 | 3 | 824577.31 | 2 | 296602.88 | 692481.56 | 2 | 1729049.5 | 2 | 1033933.62 | 1644264.5 | 775466.94 | 1611618.12 | 1629747.75 | 1179860.75 | 2997367 | 892947.94 | 1080482.75 | 1563489 | 1807969.12 | 1315545.88 | 1481727.12 |
| Q8BHN3 | Ganab | Neutral alpha-glucosidase AB | 859.4 | 21.5 | 106911 | 17 | 4031129 | 14773727 | 6098189.5 | 9108546 | 18033066 | 8371752 | 5637241 | 10829484 | 13104222 | 10956927 | 13095488 | 12051952 | 10079562 | 10730956 | 7093461.5 | 10020612 | 9914376 | 9797724 | 9180864 | 3971275 |
| P16858 | Gapdh | Glyceraldehyde-3-phosphate dehydrogenase | 1525.11 | 74.17 | 35810 | 23 | 1654572.88 | 3804337.5 | 48558.84 | 4208651.5 | 686905.19 | 3212006.75 | 2552205.75 | 304902.38 | 4247690 | 3352505 | 4327892 | 5021248 | 4530653.5 | 5400100 | 2362159 | 5685754 | 6131418 | 3387269.5 | 1934048 | 5105123 |
| Q6PAR5 | Gapvd1 | GTPase-activating protein and VPS9 domain-con | 109.62 | 2.4 | 162402 | 2 | 1377230 | 292540.06 | 913777.69 | 405275.41 | 903703.88 | 214007.91 | 1063378.75 | 141528.53 | 479522.91 | 861756.75 | 362441.69 | 2 | 1122682.38 | 671313.75 | 958478.06 | 91792.41 | 2 | 214235.33 | 2 | 503630.03 |
| Q6AYA1 | Gar1 | H/ACA ribonucleoprotein complex subunit 1 | 91.84 | 9.73 | 23011 | 2 | 54755.03 | 271201.22 | 107710.47 | 321889 | 164889.19 | 154877.36 | 586386.62 | 210863.12 | 395834.09 | 196718.22 | 166377.98 | 882268.81 | 278855.69 | 399300.62 | 571406.38 | 2 | 311752.72 | 147318.69 | 2 | 2 |
| Q5I0G4 | Gars1 | Glycine--tRNA ligase | 1025.97 | 29.81 | 81793 | 18 | 11906081 | 7679214 | 18289832 | 16285279 | 18419070 | 10560660 | 13317587 | 17404582 | 14781104 | 14040758 | 16726976 | 18604314 | 20758266 | 19599232 | 32421080 | 22169148 | 18116376 | 18279184 | 18577508 | 15983027 |
| Q64737 | Gart | Trifunctional purine biosynthetic protein adenosin | 575.17 | 18.02 | 107503 | 10 | 574757.38 | 8463467 | 1824071.75 | 4809965 | 1373617.62 | 1743791.75 | 3638425.25 | 1485679.5 | 3483740.25 | 3155581.25 | 6026824.5 | 2366141.5 | 6069589.5 | 7255567.5 | 7431293 | 3753578.25 | 5396053.5 | 5830079.5 | 2704956.5 | 6887531.5 |
| O55148 | Gas7 | Growth arrest-specific protein 7 | 40.57 | 1.66 | 48275 | 1 | 954719.88 | 684716.38 | 2 | 1891482.12 | 2 | 1021380.75 | 577056.25 | 2 | 2 | 2 | 2 | 590069.25 | 2 | 634312.38 | 1151899.38 | 993184 | 820279.5 | 1101381 | 2 | 1071035.5 |
| Q8BFQ8 | Gatd1 | Glutamine amidotransferase-like class 1 domain-c | 38.35 | 5.91 | 23277 | 1 | 960315.19 | 2 | 434372.62 | 2 | 635976.06 | 2 | 2 | 2 | 391473.69 | 507641.5 | 688421.56 | 2 | 2 | 2 | 619119.19 | 2 | 2 | 2 | 2 | 2 |
| P50442 | Gatm | Glycine amidinotransferase, mitochondrial | 806.33 | 35.46 | 48242 | 15 | 20488936 | 39939994 | 20828474 | 28084676 | 35189606 | 13966190 | 13706354 | 20622212 | 27369850 | 26238050 | 25048970 | 25162272 | 31868026 | 28221572 | 17812358 | 19040404 | 18233614 | 23397176 | 23790266 | 21549418 |
| P17439 | Gba1 | Lysosomal acid glucosylceramidase | 303.3 | 16.7 | 57622 | 7 | 410160.22 | 236287.12 | 1199925.75 | 649997.31 | 2272681 | 322087.16 | 439174.69 | 645179.56 | 2260296.75 | 3202660 | 66594.66 | 1418827.75 | 2124230.25 | 1278292 | 1555599.38 | 2 | 175450.5 | 553888.81 | 548373 | 222066.53 |
| Q9Z0E6 | Gbp2 | Guanylate-binding protein 2 | 45 | 2.04 | 66739 | 1 | 4477860 | 4877511 | 4182352.75 | 4710503.5 | 6130350 | 741177.5 | 556665.44 | 5856077 | 474604.62 | 402787.31 | 2457598 | 1918367.12 | 2 | 1739172.88 | 2281119.25 | 809392.12 | 1346131.62 | 2245475.25 | 1211270.12 | 2 |
| Q61107 | Gbp3 | Guanylate-binding protein 3 | 95.68 | 4.19 | 70801 | 2 | 2 | 2 | 2 | 448481.12 | 437770 | 2 | 2 | 332856 | 1405585.62 | 843012.25 | 2 | 2 | 2 | 2 | 2 | 2 | 2 | 214458.27 | 2 | 2 |
| O88986 | Gcat | 2-amino-3-ketobutyrate coenzyme A ligase, mitoc | 237.38 | 17.79 | 44931 | 4 | 515595.69 | 219652.44 | 101391.18 | 241532.59 | 253368.88 | 2 | 204691.16 | 2 | 176482.3 | 88069.7 | 121288.48 | 398855.25 | 217535.27 | 114124.48 | 197662.53 | 2 | 341033.16 | 189321.05 | 2 | 2 |
| Q60759 | Gcdh | Glutaryl-CoA dehydrogenase, mitochondrial | 83.14 | 4.79 | 48606 | 2 | 1752240.62 | 1415532.5 | 1503616.25 | 1680465 | 2 | 1767897.62 | 1304576.62 | 1605768.25 | 1529796.25 | 1083321.88 | 893350 | 705440.38 | 902946.31 | 830659.81 | 870196.75 | 987682.69 | 1007520.12 | 1118014.88 | 458342.06 | 1262166.25 |
| P19468 | Gclc | Glutamate--cysteine ligase catalytic subunit | 50.47 | 1.41 | 72619 | 1 | 2 | 2 | 2 | 2 | 349672.56 | 2 | 2 | 2 | 650258.69 | 440876.53 | 2 | 2 | 373134.59 | 142083.77 | 167409.59 | 2 | 2 | 2 | 2 | 2 |
| O09172 | Gclm | Glutamate--cysteine ligase regulatory subunit | 120.51 | 7.66 | 30535 | 2 | 557746.31 | 1343189 | 2 | 1719910.25 | 2426111.25 | 4069312 | 3859213.75 | 5964217 | 4333771.5 | 3143511.25 | 6221283.5 | 5728182 | 5130086.5 | 5111438.5 | 1620347.75 | 8920770 | 3473076.75 | 1971533.38 | 6062396.5 | 1696200.12 |
| E9PVA8 | Gcn1 | Stalled ribosome sensor GCN1 | 723.97 | 8.01 | 293021 | 17 | 5853250 | 7840433 | 10032417 | 4507124.5 | 7489649 | 4123665 | 3358379 | 5055318 | 3693987 | 6361358 | 2810361.75 | 3184800.5 | 7161292.5 | 6457730.5 | 4950020.5 | 2061106.38 | 2761321.25 | 9588999 | 4162959.5 | 5492215.5 |
| P50396 | Gdi1 | Rab GDP dissociation inhibitor alpha | 355.17 | 12.53 | 50522 | 5 | 1273546.62 | 1611396.38 | 2110220.75 | 1214193 | 1784088.62 | 2180317.25 | 1975787 | 2977043 | 1384478.5 | 1656203.88 | 2770104.25 | 4006699 | 3551606.5 | 1362352.38 | 2128957 | 2553054.25 | 3758165 | 3013912.25 | 4966340 | 2278581 |
| P50399 | Gdi2 | Rab GDP dissociation inhibitor beta | 929.5 | 50.11 | 50537 | 16 | 22538074 | 25860356 | 29365346 | 15215403 | 21359916 | 17413382 | 16499529 | 21090406 | 10734811 | 16415096 | 18544224 | 18862462 | 24014622 | 21244686 | 18754146 | 17943188 | 20045954 | 21243534 | 19475504 | 14693805 |
| G3V9T7 | Get3 | ATPase Get3 | 107.59 | 5.75 | 38823 | 2 | 783101.81 | 719937.25 | 2 | 1539860.12 | 3761926.5 | 1412206.25 | 1623259.25 | 771226.94 | 3307472 | 2872747 | 241180.03 | 142483.2 | 641570.81 | 253432.94 | 2 | 2 | 757128.88 | 2 | 2 | 2 |
| Q8K0D5 | Gfm1 | Elongation factor G, mitochondrial | 345.25 | 10.79 | 83550 | 8 | 1362288.5 | 523055.44 | 1928451.25 | 726433.38 | 1865306 | 1541938.25 | 1179749.75 | 2723652.25 | 1240703.75 | 5314760 | 4159827.5 | 3222339.75 | 5377717 | 2208032.5 | 2313085.5 | 3634126.5 | 2185122.75 | 2274656.75 | 3299990.25 | 1581623 |
| Q8R0H9 | Gga1 | ADP-ribosylation factor-binding protein GGA1 | 72.35 | 2.99 | 69972 | 2 | 1236234.75 | 2041166.25 | 1794449.75 | 2 | 1599505.5 | 1879470.62 | 1488434 | 2630487 | 1139897.75 | 914716.62 | 2 | 2 | 1307120.25 | 2061889.75 | 853382.19 | 2 | 2 | 2 | 2 | 1939327 |
| Q99J23 | Ghdc | GH3 domain-containing protein | 78.07 | 2.44 | 58508 | 1 | 41922.75 | 81251.31 | 2 | 121425.19 | 35484.57 | 162311.53 | 218443.12 | 2 | 190011.25 | 92191.73 | 2 | 2 | 72310.95 | 142275.3 | 2 | 2 | 126976.59 | 2 | 2 | 2 |
| Q91VC9 | Ghitm | Growth hormone-inducible transmembrane protei | 224.43 | 19.36 | 37275 | 4 | 1090722.25 | 1184988.25 | 189889.14 | 757286.12 | 128793.72 | 2 | 1299921 | 184555.45 | 796890.69 | 139183.38 | 2486594.75 | 2007378.25 | 2660921.75 | 2 | 443822.66 | 2 | 241627.03 | 3102005.75 | 2 | 608735 |
| Q6Y7W8 | Gigyf2 | GRB10-interacting GYF protein 2 | 85.24 | 2.63 | 149193 | 2 | 750180.5 | 1007115.94 | 703421.56 | 771270.62 | 1046175.19 | 682501.06 | 2 | 2 | 827074 | 542113.38 | 1460790.25 | 1124892.12 | 2 | 1293086.5 | 774791.94 | 522085.53 | 769167.69 | 2 | 2 | 112479.73 |
| P51569 | Gla | Alpha-galactosidase A | 423.28 | 21.24 | 47643 | 6 | 26989780 | 37989886 | 21566474 | 12942791 | 12983823 | 15090594 | 21304952 | 19669248 | 6012070.5 | 17250508 | 20551922 | 16716627 | 19833078 | 15523125 | 18472344 | 28096182 | 29351796 | 32446884 | 27744742 | 33611466 |
| P23780 | Glb1 | Beta-galactosidase | 595.47 | 15.3 | 73121 | 9 | 23688268 | 30222378 | 22891512 | 27925610 | 24686742 | 23420534 | 23562538 | 23872482 | 27077046 | 24791224 | 11714308 | 14315416 | 19810458 | 16135974 | 27148040 | 18623072 | 29055778 | 20520362 | 14513610 | 26881006 |
| Q8BMF8 | Gldn | Gliomedin | 34.53 | 1.28 | 59137 | 1 | 6824907.5 | 359485.56 | 5022631.5 | 4859995.5 | 553862.94 | 4884347.5 | 495199.34 | 5317460 | 1918824 | 486171.84 | 4128202.5 | 7111019.5 | 854035.94 | 3837324.25 | 3098674.5 | 761763.94 | 877784 | 4267332.5 | 5050425.5 | 12775542 |
| Q61543 | Glg1 | Golgi apparatus protein 1 | 761.99 | 15.23 | 133734 | 13 | 11023077 | 10381269 | 9165269 | 8765269 | 4318496 | 2416385 | 2097386 | 3033592.5 | 4057770.5 | 5895341.5 | 1991179 | 4741745 | 2408291.25 | 1255900.25 | 2680965.5 | 2015167.5 | 2383715.75 | 3073398.75 | 1749435.12 | 1896130.75 |
| Q9CWG1 | Glipr1 | Glioma pathogenesis-related protein 1 | 52.43 | 3.92 | 29129 | 1 | 794861.38 | 405147.88 | 615461.5 | 757752.38 | 645906.31 | 2 | 2 | 2 | 653752.5 | 373465.47 | 2 | 2 | 2 | 2 | 743013.88 | 2 | 2 | 2 | 2 | 2 |
| Q8BZM1 | Glmn | Glomulin | 56.34 | 1.85 | 67756 | 1 | 791166.56 | 133686.56 | 439505.53 | 232825.61 | 564132.19 | 2 | 151746.5 | 2 | 257009.48 | 421766.03 | 2 | 2 | 2 | 210056.2 | 2 | 2 | 191477.7 | 2 | 2 | 2 |
| Q9JHJ3 | Glmp | Glycosylated lysosomal membrane protein | 54.82 | 3.22 | 43804 | 1 | 2 | 2 | 2 | 231248.09 | 2 | 2 | 2 | 2 | 2 | 2 | 2 | 76763.63 | 2 | 126056.24 | 2 | 306566.62 | 225718.92 | 181727.16 | 225523.55 | 2 |
| Q9CPU0 | Glo1 | Lactoylglutathione lyase | 97.94 | 8.7 | 20810 | 2 | 54189.61 | 69940.69 | 1396868.75 | 2 | 746744 | 2 | 206380.53 | 271835.47 | 2 | 1167698.62 | 90959.31 | 270888.66 | 637327 | 884880.88 | 3695179.25 | 2 | 221185.28 | 357254.41 | 2 | 152986.41 |
| Q9CPV4 | Glod4 | Glyoxalase domain-containing protein 4 | 291.36 | 26.51 | 33317 | 6 | 2145665.5 | 1357131.88 | 3015312.75 | 2588605 | 5361060 | 1866779.75 | 250127.17 | 2165025.5 | 2530534 | 4822941.5 | 2722495.5 | 2251564 | 1604367.12 | 2583202.25 | 2203231.75 | 2292704.25 | 2195379 | 1994068.12 | 2161480.25 | 2819696.5 |
| Q9QUH0 | Glrx | Glutaredoxin-1 | 155.5 | 44.86 | 11871 | 3 | 13159430 | 24741158 | 10146403 | 15153611 | 13293549 | 32381568 | 35286842 | 25786606 | 25500854 | 19932350 | 17560038 | 16522744 | 13678486 | 14231838 | 9859014 | 13904983 | 15139144 | 11632714 | 16845166 | 13664688 |
| Q80Y14 | Glrx5 | Glutaredoxin-related protein 5, mitochondrial | 147.62 | 20.39 | 16292 | 2 | 933031.19 | 983211.12 | 706816.44 | 250712.92 | 289239.72 | 225616.69 | 168701.95 | 117217.07 | 308585.84 | 390916.81 | 277660.88 | 180294.94 | 390664.03 | 2 | 555463.12 | 2 | 188788.33 | 2 | 2 | 2 |

| | | | | | | | | | | | | | | | | | | | | | | | | | | |
|---|---|---|---|---|---|---|---|---|---|---|---|---|---|---|---|---|---|---|---|---|---|---|---|---|---|---|
| D3Z7P3 | Gls | Glutaminase kidney isoform, mitochondrial | 384.18 | 16.91 | 73964 | 7 | 11983941 | 14366878 | 12168808 | 12914043 | 15878416 | 14035675 | 11767126 | 18758262 | 13067364 | 13960429 | 10759921 | 8440191 | 10511498 | 10144089 | 7605875 | 11812026 | 13634850 | 18923092 | 15406768 | 15721714 |
| Q8K0R6 | Gltpd2 | Glycolipid transfer protein domain-containing prot | 45.85 | 2.18 | 34453 | 1 | 4691014.5 | 2 | 629139.62 | 3399903.5 | 2215430.5 | 3537215.25 | 3960289.75 | 3583997.75 | 2127558.75 | 2675442.5 | 1703943.38 | 2614212.5 | 2367811.5 | 2 | 3284285 | 1983231.5 | 2 | 1253580.38 | 2 | 2 |
| P26443 | Glud1 | Glutamate dehydrogenase 1, mitochondrial | 1210.98 | 49.1 | 61337 | 20 | 26388142 | 53733810 | 57954098 | 44305906 | 60398218 | 40474486 | 42727042 | 64773746 | 46134430 | 63967530 | 33807906 | 30448242 | 37978150 | 41521714 | 42487786 | 31071054 | 22433482 | 48373978 | 28067538 | 31274426 |
| Q5RKH0 | Glyr1 | Cytokine-like nuclear factor N-PAC | 237.65 | 10.33 | 60421 | 4 | 1365173.5 | 1882605.25 | 1132758.38 | 1722122.25 | 2907904.75 | 1296961.25 | 1165448 | 1338204.25 | 1249441.88 | 1632180 | 725383.38 | 2 | 962743.19 | 289594.06 | 2 | 973932.44 | 625086.25 | 1969732.38 | 253835.92 | 962293.38 |
| Q60648 | Gm2a | Ganglioside GM2 activator | 49.93 | 11.4 | 20824 | 1 | 2 | 2 | 166407.61 | 179616.06 | 2 | 2 | 189242.59 | 2 | 83998.81 | 2 | 217909.89 | 136029.22 | 2 | 105012.73 | 2 | 2 | 130564.09 | 2 | 2 | 2 |
| Q8K0C9 | Gmds | GDP-mannose 4,6 dehydratase | 67.14 | 5.91 | 41985 | 2 | 837407.31 | 934362.62 | 773000.19 | 2 | 453222.12 | 2 | 2 | 2 | 2 | 2 | 664963.56 | 807964 | 816751.06 | 500143.53 | 2 | 2 | 2 | 2 | 1214999.12 | 2 |
| Q9ERL7 | Gmfg | Glia maturation factor gamma | 158.12 | 22.54 | 16748 | 3 | 783831.19 | 2 | 469933.69 | 1498524.38 | 9201175 | 2281648.5 | 1434031 | 3626422.5 | 4732208.5 | 3038634 | 2960222.25 | 2691981.5 | 1457360 | 2678469.5 | 214162.25 | 2323639.5 | 215777.81 | 2 | 1070887.88 | 2 |
| Q6PGG2 | Gmip | GEM-interacting protein | 66.24 | 1.34 | 107545 | 1 | 2 | 2 | 88607.34 | 2 | 62767.18 | 2 | 148127.89 | 53139.3 | 2 | 82716.31 | 2 | 2 | 54705.69 | 2 | 404416.81 | 2 | 2 | 2 | 2 | 2 |
| Q5XIC1 | Gmppa | Mannose-1-phosphate guanylyltransferase regula | 57.93 | 3.57 | 46191 | 1 | 2 | 2 | 2 | 2 | 2 | 2 | 2 | 2 | 981383.62 | 2 | 1034577.25 | 1049482.25 | 2 | 2 | 2 | 954922.81 | 2 | 2 | 2 | 2 |
| Q8BTZ7 | Gmppb | Mannose-1-phosphate guanylyltransferase cataly | 77.92 | 5 | 39917 | 1 | 1355008.62 | 182708.55 | 1270536.75 | 1208448 | 1350790.38 | 2 | 168152.03 | 305658.19 | 1473389.62 | 985050.5 | 468150.22 | 438642.44 | 1322656.75 | 881427.44 | 634864.75 | 416240.84 | 515142.91 | 2 | 696422.31 | 1043799.31 |
| Q9DCZ1 | Gmpr | GMP reductase 1 | 45.57 | 6.38 | 37482 | 1 | 2 | 2 | 84850.61 | 2 | 79619.46 | 2 | 60991.45 | 2 | 45887.22 | 72922.44 | 2 | 87379.16 | 46587.17 | 65933.96 | 335567.97 | 2 | 132779.34 | 63683.49 | 2 | 2 |
| Q3THK7 | Gmps | GMP synthase [glutamine-hydrolyzing] | 384.7 | 12.7 | 76723 | 7 | 1790518.5 | 1742860 | 2035929 | 2145215.75 | 2654113.25 | 628116 | 1662653.38 | 1179275.25 | 1678558.88 | 2894089.5 | 645404.62 | 994280.5 | 1008670 | 1722025.5 | 3172380.5 | 552581.56 | 891843.19 | 1202097.25 | 2 | 1023859.62 |
| P27601 | Gna13 | Guanine nucleotide-binding protein subunit alpha | 104.78 | 5.57 | 44055 | 2 | 639408.81 | 155423.39 | 951829.81 | 152749.5 | 261820.06 | 2 | 392267.47 | 114537.58 | 666182.56 | 797927.25 | 149034.27 | 191704.81 | 493367.38 | 208535.47 | 2 | 145703.8 | 330968.53 | 198483.42 | 2 | 2 |
| P08752 | Gnai2 | Guanine nucleotide-binding protein G(i) subunit a | 832.73 | 38.31 | 40489 | 12 | 38154990 | 37151450 | 43610278 | 33984730 | 42021894 | 34239930 | 28910452 | 37368082 | 31825398 | 36520238 | 22027290 | 16357914 | 20013668 | 17289804 | 28910366 | 13794714 | 15245395 | 13513181 | 15860340 | 13804663 |
| Q9DC51 | Gnai3 | Guanine nucleotide-binding protein G(i) subunit a | 341.75 | 15.54 | 40538 | 5 | 4551120 | 5159646 | 1901362.38 | 3842177.5 | 1557731.5 | 4022973.5 | 5181303.5 | 5574130 | 3067184.5 | 3896231.5 | 3520737.75 | 1902296.5 | 3481367 | 3380558.5 | 4634676.5 | 4176531.5 | 3629260.75 | 5385406 | 3744826 | 3657996 |
| P21279 | Gnaq | Guanine nucleotide-binding protein G(q) subunit a | 234.09 | 13.09 | 42158 | 4 | 1752609.25 | 808115.06 | 3920693.75 | 1082933.38 | 586319.25 | 450407.19 | 778008.94 | 1900967 | 472739.53 | 2383170.5 | 149314.28 | 628295.25 | 653289.81 | 1051854.62 | 1525832.25 | 149404.42 | 1473481.38 | 994037.94 | 406811.34 | 1086891.38 |
| P63094 | Gnas | Guanine nucleotide-binding protein G(s) subunit a | 229.42 | 15.74 | 45664 | 5 | 1186374 | 595689.62 | 1928718.38 | 2 | 1443551.62 | 774078.75 | 2 | 2 | 1065774.75 | 538684.75 | 609747.06 | 1027515.75 | 529044.81 | 436170.44 | 990871.75 | 854307.06 | 799679.75 | 2593235.5 | 1200942.25 | 2 |
| P54311 | Gnb1 | Guanine nucleotide-binding protein G(I)/G(S)/G(T | 382.11 | 20.88 | 37377 | 7 | 76517.09 | 1188626.38 | 1115604.88 | 2 | 162681.81 | 990081.88 | 231177.19 | 222230.75 | 126882.41 | 889543.31 | 180075.75 | 2276775.25 | 2310264.25 | 2272620.25 | 2285550 | 2901582 | 2648589.25 | 171587.8 | 4178300 | 1190268.12 |
| P54313 | Gnb2 | Guanine nucleotide-binding protein G(I)/G(S)/G(T | 725.56 | 32.35 | 37331 | 13 | 4528335 | 2856450 | 5906525 | 2720731 | 5751241.5 | 2175717.25 | 2495816 | 5175966 | 2898574.5 | 3819373.5 | 4977109 | 3812502.75 | 7683495 | 5954461 | 3037439 | 4226700 | 6984495.5 | 6034653 | 6843365.5 | 6884511 |
| Q9DAS9 | Gng12 | Guanine nucleotide-binding protein G(I)/G(S)/G(O | 64.67 | 15.28 | 7997 | 1 | 75157.62 | 2 | 2 | 2 | 67926.49 | 559734.94 | 2 | 241533.22 | 1899181.38 | 154019.72 | 751748.06 | 1018766.94 | 2 | 2 | 2 | 533182.31 | 2 | 2 | 2 | 2 |
| P63213 | Gng2 | Guanine nucleotide-binding protein G(I)/G(S)/G(O | 55.27 | 16.9 | 7850 | 1 | 2840727.75 | 4342172 | 1985322.25 | 3113298.75 | 2776253.75 | 3885324.5 | 2904977.5 | 3630422 | 2004480 | 2016909.5 | 2145472 | 2150965.25 | 1842302.75 | 2200243.75 | 2189579.5 | 2404227.25 | 2268590.5 | 3014101.75 | 2921863.75 | 2737714.5 |
| P36916 | Gnl1 | Guanine nucleotide-binding protein-like 1 | 78.68 | 2.64 | 68771 | 2 | 2160702.25 | 3770719 | 1046130.25 | 1234449 | 1144398.62 | 3720172.5 | 1252921 | 1889083.25 | 2209762.5 | 894351.06 | 846755.12 | 1414294.62 | 565612.19 | 676904 | 726549.5 | 1338315.38 | 759261.25 | 2 | 1167256.38 | 2 |
| Q8CI11 | Gnl3 | Guanine nucleotide-binding protein-like 3 | 242.25 | 15.8 | 60786 | 5 | 2 | 2 | 535805.69 | 2 | 980545.38 | 519682.94 | 438588.06 | 90197 | 2 | 2 | 2 | 116299.68 | 2 | 571072.19 | 297565.81 | 2 | 2 | 2 | 459433.5 | 2 |
| O88958 | Gnpda1 | Glucosamine-6-phosphate deaminase 1 | 212.99 | 9 | 32549 | 3 | 4410893.5 | 4781867.5 | 5978829.5 | 1693682.12 | 2626266 | 1631027 | 2584914.5 | 3078493 | 1904788 | 2395731 | 3745373.5 | 4151411.5 | 2821306.5 | 3185772 | 2581372.5 | 5023229.5 | 3848980.25 | 4635546 | 3877529 | 3271549.5 |
| Q8BFR4 | Gns | N-acetylglucosamine-6-sulfatase | 477.23 | 17.65 | 61175 | 8 | 27891990 | 41338570 | 24056404 | 20303390 | 21449938 | 26083300 | 25711392 | 26993846 | 21171752 | 22315932 | 18783092 | 18641872 | 20313132 | 18097350 | 20170512 | 19881048 | 21256854 | 25787610 | 23036426 | 27720032 |
| Q9CRA5 | Golph3 | Golgi phosphoprotein 3 | 88.36 | 5.37 | 33752 | 1 | 184217.38 | 148947.45 | 165253.91 | 216823.31 | 645104.25 | 258551.7 | 97226.88 | 32230.03 | 95658.4 | 203858.19 | 146304.05 | 93218.91 | 46519.13 | 126426.48 | 324017.12 | 91722.2 | 214645.58 | 43212.68 | 319656.47 | 567494.5 |
| Q9CR60 | Golt1b | Vesicle transport protein GOT1B | 86.94 | 10.87 | 15422 | 2 | 36930.93 | 70095.91 | 52566.57 | 124353.3 | 34206.23 | 402356.62 | 531813.56 | 311920.12 | 545582.62 | 345665.38 | 114545.47 | 265599.25 | 225204.02 | 456010.72 | 477998.69 | 2 | 1052988.25 | 570902.38 | 150759.03 | 384939.88 |
| Q99JX3 | Gorasp2 | Golgi reassembly-stacking protein 2 | 121.45 | 5.32 | 47038 | 2 | 687307.81 | 783291.5 | 766412.56 | 1012279.19 | 1111869.25 | 1096108 | 919737.44 | 1170524.25 | 924634.12 | 827632.75 | 2184248.75 | 1965609.88 | 2278919.5 | 1597432.62 | 1374193.25 | 2693940.25 | 2669931.5 | 3110761.75 | 3844427.75 | 2783130.5 |
| O88630 | Gosr1 | Golgi SNAP receptor complex member 1 | 38.21 | 2.8 | 28489 | 1 | 203459.81 | 2 | 2 | 234129.75 | 2 | 2 | 2 | 994283.94 | 2 | 2 | 2 | 2 | 925343.75 | 630330.75 | 2 | 2 | 2 | 2 | 2 | 533905.38 |
| P05201 | Got1 | Aspartate aminotransferase, cytoplasmic | 964.18 | 40.44 | 46248 | 17 | 123316370 | 143328114 | 104480994 | 76538338 | 67794346 | 88768746 | 79902450 | 78955834 | 61168018 | 83637570 | 150538642 | 147735762 | 136639330 | 117523442 | 92255458 | 117431370 | 112922802 | 121927818 | 98044714 | 122358738 |
| P05202 | Got2 | Aspartate aminotransferase, mitochondrial | 1165.76 | 47.91 | 47411 | 21 | 119601050 | 220173426 | 187098018 | 182662338 | 202095650 | 206447570 | 172067442 | 195485906 | 224075234 | 177840114 | 258291330 | 263498082 | 279758434 | 228734994 | 143151234 | 220744098 | 175432578 | 178919602 | 196247394 | 155445426 |
| Q9WTK3 | Gpaa1 | Glycosylphosphatidylinositol anchor attachment 1 | 139.14 | 4.83 | 67949 | 3 | 1439940.38 | 2 | 482193.31 | 532773.38 | 713117.69 | 2 | 406518.88 | 1152078.12 | 1305717.38 | 461838.09 | 1186945.88 | 1280868.38 | 539692.44 | 710415.75 | 2 | 953074.19 | 811865.69 | 2 | 684540.94 | 773334.62 |
| Q8C0N2 | Gpat3 | Glycerol-3-phosphate acyltransferase 3 | 52.78 | 2.51 | 49000 | 1 | 432245.59 | 2 | 2 | 2 | 2 | 1358715.75 | 990220.5 | 1147605.12 | 643576.81 | 434529.34 | 2 | 2 | 2 | 980599.88 | 703473.88 | 2 | 2 | 943908.75 | 1271862.38 | 1127165.12 |
| Q3ULJ0 | Gpd1l | Glycerol-3-phosphate dehydrogenase 1-like prote | 94.81 | 6.55 | 38226 | 2 | 3365666 | 2290730 | 3489163 | 2528300.75 | 1783779 | 1078483.75 | 1731017 | 2416357 | 1146977.75 | 1832222.25 | 1902682 | 672196.75 | 1765201.38 | 1342646 | 2110936.75 | 2 | 1677502 | 1195664.38 | 2093463.25 | 1268181.88 |
| Q64521 | Gpd2 | Glycerol-3-phosphate dehydrogenase, mitochond | 534.55 | 14.03 | 80954 | 10 | 3763481 | 3913397.25 | 3148935.25 | 4037330.5 | 2757806.25 | 1220497.62 | 2179101 | 2286991 | 2751843.5 | 2871082.5 | 1070887.25 | 690892.88 | 1388800.62 | 1018378.94 | 4934957 | 2198288 | 5401137 | 3145866 | 4942316 | 905327.75 |
| P06745 | Gpi | Glucose-6-phosphate isomerase | 754.68 | 34.95 | 62767 | 15 | 18399652 | 17661602 | 19883522 | 16281490 | 12329454 | 17578774 | 19705344 | 9490201 | 20169952 | 26197856 | 18823020 | 20916678 | 18839154 | 15300674 | 36636870 | 23549276 | 19351234 | 19481718 | 19359552 | 16123813 |
| Q8CIM5 | Gpr84 | G-protein coupled receptor 84 | 340.35 | 14.39 | 43717 | 5 | 1892799.5 | 1617385.25 | 2 | 3311343.5 | 1477818.25 | 1552145.5 | 470775.91 | 656183.12 | 739408.44 | 3025175.5 | 2 | 908286.56 | 2 | 2 | 2 | 1154470.75 | 1791286.25 | 1533583.25 | 2 | 2 |
| P97834 | Gps1 | COP9 signalosome complex subunit 1 | 46.77 | 2.34 | 53428 | 1 | 954119.25 | 955151.38 | 1752671.38 | 1131733.75 | 932492.19 | 909612.5 | 1045888.94 | 37160.68 | 704250.5 | 717611.75 | 611200 | 2 | 709636.44 | 368466.41 | 1406588.75 | 497402.81 | 745334.56 | 2 | 656834.06 | 2 |
| P11352 | Gpx1 | Glutathione peroxidase 1 | 319.34 | 42.29 | 22329 | 7 | 11995194 | 10725700 | 18087034 | 6604270.5 | 12042409 | 6138743.5 | 6281564 | 13441629 | 10302670 | 16648380 | 13450772 | 11877240 | 14720146 | 11337117 | 15608672 | 11681104 | 9893870 | 10397275 | 9677882 | 8110383 |
| O70325 | Gpx4 | Phospholipid hydroperoxide glutathione peroxida | 178.26 | 24.87 | 22229 | 4 | 2045352 | 4532702 | 1433915.75 | 1856456.62 | 3363583 | 5683012.5 | 3246732.5 | 3899931.5 | 3444215.5 | 4097661 | 3301795.5 | 3451769.25 | 3282645.5 | 3190018 | 7249973 | 1543101.75 | 3161058.25 | 3579574 | 1932396 | 5787221 |
| P62994 | Grb2 | Growth factor receptor-bound protein 2 | 39.29 | 3.23 | 25206 | 1 | 3268615.25 | 2947037 | 3935995.5 | 2591057 | 4442747.5 | 4505027.5 | 4745182.5 | 5766173.5 | 2803413.75 | 3608225 | 4932991.5 | 4445115.5 | 2913034 | 2206395.25 | 2744589.5 | 3193843 | 2681442.5 | 1543626.25 | 2366519.25 | 1418345.25 |
| O35127 | Grcc10 | Protein C10 | 118.59 | 28.57 | 13194 | 2 | 2 | 36460.06 | 169661.28 | 2 | 136120.59 | 2 | 2 | 2 | 2 | 341403.19 | 44519.49 | 238722.91 | 50954.82 | 79465.16 | 566369.31 | 2 | 223203.5 | 2 | 2 | 2 |
| Q91Z53 | Grhpr | Glyoxylate reductase/hydroxypyruvate reductase | 206.43 | 14.94 | 35329 | 5 | 2995449.75 | 2979767.25 | 1463650.12 | 1211948.5 | 4621316 | 3089754 | 3549790.25 | 2 | 1625587.75 | 3439841.5 | 5461698.5 | 4513914.5 | 1928531.75 | 2235647.25 | 1198556.25 | 4119174.5 | 3331800 | 1687748.75 | 4453811.5 | 2230468 |
| P28798 | Grn | Progranulin | 831.59 | 28.86 | 63458 | 15 | 60492282 | 37321506 | 31341604 | 22637978 | 29928626 | 42698530 | 20145478 | 30984380 | 37691046 | 29619368 | 41440146 | 27162906 | 16986822 | 28210688 | 36897242 | 19570606 | 36746994 | 39195438 | 33897054 | 41303290 |
| P97576 | Grpel1 | GrpE protein homolog 1, mitochondrial | 121.23 | 14.29 | 24297 | 3 | 2 | 2 | 2 | 2 | 2 | 2 | 2 | 73635.1 | 2 | 2 | 2 | 142603.08 | 2 | 2 | 2 | 2 | 2 | 2 | 2 | 2 |
| Q8C5Q4 | Grsf1 | G-rich sequence factor 1 | 103.97 | 7.93 | 53076 | 2 | 2 | 2 | 2 | 653917.81 | 430163.75 | 2 | 404720.03 | 616415.31 | 372455.41 | 408386.16 | 38833.43 | 2 | 2 | 629565.88 | 2 | 2 | 104637.65 | 2 | 2 | 2 |
| Q9D8T2 | Gsdmd | Gasdermin-D | 81.5 | 5.75 | 53238 | 2 | 466438.44 | 2 | 2334604 | 4269924 | 2 | 2 | 482132.59 | 2 | 442426.03 | 2 | 1386908.75 | 952114.88 | 1501980.38 | 1088231.88 | 1886354.75 | 765018.56 | 1155700.62 | 978280.31 | 956250.19 | 993194.69 |
| P13020 | Gsn | Gelsolin | 1055.14 | 31.15 | 85942 | 19 | 18004524 | 14594700 | 21143894 | 30874014 | 47171374 | 31555634 | 20980570 | 32073128 | 41856894 | 36987374 | 42322898 | 43732490 | 37335002 | 37526546 | 37941938 | 43378230 | 36620398 | 29174428 | 34736734 | 32724968 |
| Q8R050 | Gspt1 | Eukaryotic peptide chain release factor GTP-bind | 177.63 | 8.33 | 68625 | 4 | 361570.44 | 477075.25 | 2 | 2 | 2 | 2 | 2 | 2 | 744064.19 | 321261.47 | 3959867.25 | 4096682.75 | 3440485.5 | 3137756.5 | 2 | 4266507.5 | 2988867.25 | 2672725.75 | 540514.06 | 1097766.38 |
| P47791 | Gsr | Glutathione reductase, mitochondrial | 317.81 | 10.4 | 53663 | 6 | 8307580 | 12517123 | 9198177 | 6652179.5 | 9185014 | 8983948 | 9032163 | 6930154 | 8960204 | 7774349.5 | 21388952 | 20359132 | 15323620 | 14508521 | 9236604 | 28001674 | 14368759 | 13363396 | 4995836 | 10838600 |
| P46413 | Gss | Glutathione synthetase | 81.82 | 7.81 | 52345 | 2 | 2 | 2 | 2 | 121357.03 | 177017.22 | 273630.78 | 224921.39 | 117314.7 | 2 | 2 | 234088.2 | 263189.12 | 308267.78 | 207394.84 | 2 | 162060.69 | 2 | 2 | 225615.92 | 2 |
| P10649 | Gstm1 | Glutathione S-transferase Mu 1 | 204.5 | 15.14 | 25970 | 4 | 2 | 1709555.38 | 649286.44 | 2 | 934347.31 | 2 | 2 | 578168.12 | 648103.38 | 736466.81 | 1231490.25 | 1570493.88 | 965227.69 | 1015168.94 | 777305 | 2 | 714815.12 | 2 | 109416.55 | 2 |
| P48774 | Gstm5 | Glutathione S-transferase Mu 5 | 123.26 | 11.16 | 26635 | 3 | 2 | 549747.81 | 2 | 1429050 | 1190204.12 | 984227.94 | 717092.12 | 1067435.88 | 1268109.5 | 679128.81 | 2063216.38 | 1471445.38 | 2163423.5 | 1559212.5 | 1296197.5 | 1229201.38 | 2638258.5 | 2185829.25 | 1888519.62 | 2039153.25 |
| O09131 | Gsto1 | Glutathione S-transferase omega-1 | 72.19 | 8.33 | 27498 | 2 | 2 | 2 | 2 | 2 | 2 | 2 | 2 | 2 | 74492.48 | 2 | 85065.26 | 147918.61 | 2 | 130867.05 | 2 | 2 | 2 | 2 | 2 | 2 |
| Q99ME9 | Gtpbp4 | GTP-binding protein 4 | 115.06 | 4.42 | 74113 | 3 | 2 | 2 | 2 | 2 | 2 | 2 | 2 | 2 | 2 | 2 | 2 | 2 | 2 | 2 | 2 | 2 | 2 | 2 | 2 | 2 |
| Q5BK22 | Gtpbp8 | GTP-binding protein 8 | 33.56 | 3.51 | 32232 | 1 | 1069907.75 | 2 | 2 | 2 | 570309.56 | 2 | 2 | 1029049.12 | 2 | 2 | 2 | 943311 | 350008.66 | 2 | 2 | 2 | 2 | 2 | 575019.69 | 443339.25 |
| P12265 | Gusb | Beta-glucuronidase | 858.58 | 26.54 | 74195 | 15 | 27818886 | 100815442 | 30579610 | 15711704 | 49686566 | 39637174 | 51868046 | 53371366 | 52303194 | 45559826 | 36246350 | 38085802 | 46750746 | 42055230 | 60368226 | 40852182 | 52229046 | 56149062 | 43666218 | 67819322 |
| A2RRU1 | Gys1 | Glycogen [starch] synthase, muscle | 77.44 | 2.98 | 84072 | 2 | 2 | 2 | 2 | 119116.51 | 1132814.5 | 80024.29 | 1587905.62 | 1251851 | 1846517.25 | 320503.84 | 2 | 1018496.12 | 776057.81 | 491020.19 | 1104797.38 | 2 | 584982.69 | 2 | 679050.62 | 674523.56 |
| P10922 | H1-0 | Histone H1.0 | 378.58 | 29.38 | 20861 | 6 | 1285371.25 | 3029025 | 1091846.62 | 4531133 | 1639985.25 | 3297928.25 | 2156175.5 | 2085060 | 10926613 | 2953339 | 3644561 | 6889646 | 3386781.5 | 6143003 | 891549 | 4197003 | 1761828.12 | 1796149.5 | 652778.19 | 1298171.88 |
| P43275 | H1-1 | Histone H1.1 | 619.63 | 41.78 | 21785 | 10 | 5052985 | 5968743.5 | 4382525 | 10371564 | 18441712 | 2872929 | 3317860.5 | 4463485 | 7513271.5 | 6271846.5 | 5997003 | 6853898 | 8245523 | 9300256 | 7468098.5 | 8000306 | 8211499 | 6276098 | 8373700.5 | 6464207 |
| P15864 | H1-2 | Histone H1.2 | 933.2 | 48.11 | 21267 | 15 | 4448240 | 4867813.5 | 2139827.75 | 10203490 | 10158100 | 4417790 | 2541013.5 | 6562383 | 5734103 | 4301877.5 | 4427051.5 | 6785330.5 | 5437286 | 6078030 | 3069719.75 | 9357688 | 6561468.5 | 2972445.5 | 8313986 | 6551971 |
| P43277 | H1-3 | Histone H1.3 | 800.18 | 41.18 | 22100 | 13 | 2 | 1465921.5 | 2 | 1851447.62 | 1741147.75 | 2 | 2 | 1626947.25 | 1584184 | 459655.03 | 2 | 1576426.5 | 1354392.38 | 2503083.5 | 2 | 7998961 | 4952878.5 | 4361622.5 | 6008979.5 | 4911757 |
| P43274 | H1-4 | Histone H1.4 | 775.06 | 37.44 | 21977 | 12 | 2 | 6241283 | 2 | 15579810 | 20757194 | 7134552.5 | 1110585.75 | 10723231 | 16609864 | 11558419 | 9738746 | 15909549 | 13495878 | 13636937 | 4593466.5 | 19388694 | 2072258.62 | 947803.19 | 15325284 | 4280483 |
| P43276 | H1-5 | Histone H1.5 | 667.17 | 36.77 | 22576 | 11 | 2908384 | 3094120.25 | 1730292.62 | 7817264 | 9492114 | 6186697.5 | 2921940.5 | 9593600 | 13001385 | 4197166 | 10845643 | 11092192 | 6453506.5 | 7455290 | 4222432.5 | 9826282 | 5961804 | 3883543.5 | 7197047.5 | 3802110.5 |
| P01900 | H2-D1 | H-2 class I histocompatibility antigen, D-D alpha c | 747.34 | 41.92 | 41110 | 14 | 2 | 2996176 | 204206.59 | 2679340.5 | 2912626.5 | 2149102.5 | 1246608.75 | 2651931.75 | 5414471 | 1932304.5 | 13955162 | 8535286 | 5678662 | 5773805 | 5734027.5 | 5464953.5 | 5375568 | 3252325 | 6184366 | 3924042.75 |
| P01902 | H2-K1 | H-2 class I histocompatibility antigen, K-D alpha c | 738.94 | 38.32 | 41490 | 14 | 11048726 | 12331325 | 12797458 | 25265732 | 23074630 | 7366601.5 | 11446239 | 22128440 | 17694486 | 22448316 | 11022252 | 11706928 | 11723088 | 11349006 | 9424982 | 11346556 | 11987522 | 11736058 | 12886140 | 9643701 |
| P01897 | H2-L | H-2 class I histocompatibility antigen, L-D alpha c | 327.25 | 21.27 | 40711 | 6 | 552173.75 | 2 | 2 | 553583.5 | 1461362.62 | 1057907.25 | 738171.62 | 1113074.62 | 2279357.5 | 1638222 | 4535230.5 | 3432047.5 | 3455267.25 | 3777174.75 | 3859601.25 | 4108263 | 4436619.5 | 6588569 | 3720116.25 | 3317544.25 |
| C0HKE1 | H2ac4 | Histone H2A type 1-B | 536.63 | 63.85 | 14135 | 8 | 2 | 2 | 2 | 116539.45 | 125289.62 | 58148.67 | 80372.19 | 298514.91 | 360950 | 2 | 141785.36 | 159597.83 | 77683.26 | 77242.51 | 2 | 2 | 2 | 2 | 78787.46 | 2 |
| P0C0S6 | H2az1 | Histone H2A.Z | 360.38 | 48.44 | 13553 | 5 | 824305.81 | 2 | 2 | 1255289.5 | 2093726.38 | 1364911.62 | 754336.31 | 1237728.75 | 2571369.75 | 1819550 | 2 | 2 | 2 | 471652.47 | 2 | 33641.06 | 90808.16 | 2 | 2 | 2 |
| Q64525 | H2bc18 | Histone H2B type 2-B | 1046.73 | 72.22 | 13920 | 17 | 1245315.12 | 13559132 | 2 | 1256485.38 | 5144135.5 | 5776045.5 | 2 | 129406.91 | 2568342.5 | 2 | 1965249.12 | 1280310.75 | 2 | 624309 | 632983.31 | 1371391 | 2257939.5 | 486133.72 | 4238733 | 28507954 |
| Q64524 | H2bc21 | Histone H2B type 2-E | 573.95 | 45.24 | 13993 | 9 | 338210.47 | 1488216 | 2 | 1210195.12 | 498246.94 | 1296698.5 | 227113.03 | 2 | 615642.62 | 2 | 248430.61 | 803815.06 | 126061.98 | 414286.44 | 2 | 116469.02 | 1694681.38 | 190409.34 | 1325673 | 4967460.5 |
| P10853 | H2bc7 | Histone H2B type 1-F/J/L | 970.22 | 63.49 | 13936 | 15 | 1806103.88 | 12841924 | 2 | 2281577 | 7010302 | 5794536 | 312753.03 | 2 | 2655855.5 | 2 | 1371214.88 | 1732918.5 | 133244.42 | 649286.88 | 2 | 1000141.31 | 2544852.25 | 346873.72 | 7510851.5 | 7622505.5 |
| P02301 | H3-5 | Histone H3.3C | 391.76 | 33.82 | 15315 | 8 | 275185058 | 577245826 | 187367090 | 949283394 | 1013308930 | 620586626 | 489224866 | 574521602 | 805731074 | 522402498 | 713843778 | 701416450 | 396806818 | 538909250 | 139054850 | 590526466 | 207720338 | 142749698 | 353498082 | 125868146 |
| P62804 | H4c2 | Histone H4 | 870.93 | 58.25 | 11367 | 13 | 82926906 | 136814050 | 83739866 | 593581826 | 558733314 | 320808194 | 329455202 | 533959202 | 654487874 | 317823906 | 147326210 | 154837682 | 132662274 | 119125426 | 201914706 | 167211298 | 128124258 | 139661698 | 149967442 | 108717506 |
| Q61425 | Hadh | Hydroxyacyl-coenzyme A dehydrogenase, mitoch | 135.32 | 9.24 | 34464 | 3 | 607289.75 | 662230.75 | 1256944.25 | 1385734.38 | 1741114.62 | 1064176 | 1132814 | 87011.98 | 2467141 | 2666716.25 | 2212798 | 3129296.75 | 2467107 | 2316319.5 | 1994959.12 | 1790657.38 | 959847.25 | 1049036 | 1795563.62 | 898702.94 |
| Q8BMS1 | Hadha | Trifunctional enzyme subunit alpha, mitochondria | 1022.83 | 33.55 | 82670 | 19 | 8553474 | 6586773 | 12173214 | 7117688 | 12133054 | 5131094 | 6313278 | 8996370 | 8415125 | 9746682 | 5014700.5 | 7983497.5 | 7068162.5 | 7483063.5 | 12859570 | 5765869.5 | 7869768 | 4330978.5 | 6133502.5 | 5076732 |
| Q99JY0 | Hadhb | Trifunctional enzyme subunit beta, mitochondrial | 294.17 | 18.95 | 51386 | 7 | 2 | 59699.95 | 1029154.25 | 861854 | 1556145.12 | 1254267.25 | 160917.64 | 174199.06 | 466533.69 | 3377007.75 | 3009793 | 2256596.75 | 2033677.25 | 756480.25 | 960892 | 4107254 | 858662.12 | 1453626.75 | 1342473.12 | 1168064.62 |
| P21213 | Hal | Histidine ammonia-lyase | 63.93 | 2.74 | 72284 | 1 | 2 | 2 | 60991.9 | 2 | 2 | 2 | 2 | 2 | 70194.73 | 2 | 27678.58 | 139588.62 | 2 | 2 | 2 | 2 | 287192.06 | 119641.04 | 2 | 2 |
| Q61035 | Hars1 | Histidine--tRNA ligase, cytoplasmic | 372.13 | 15.32 | 57432 | 9 | 6821393.5 | 7451798 | 8672876 | 7643621 | 8303312.5 | 7896392.5 | 6994640 | 9265368 | 4968031.5 | 6022421.5 | 6302106 | 7744583 | 7707169 | 8356624.5 | 11229619 | 6781912.5 | 7410672 | 10689681 | 6254168 | 9687044 |
| Q8BY71 | Hat1 | Histone acetyltransferase type B catalytic subunit | 68.19 | 3.12 | 49278 | 1 | 2 | 796576 | 930114.69 | 1438100.12 | 1164984.38 | 2 | 505121.44 | 2 | 398031.44 | 2 | 2 | 561177.75 | 744582.69 | 799957.94 | 687254.69 | 2 | 2 | 1048223.19 | 2 | 2 |
| O35387 | Hax1 | HCLS1-associated protein X-1 | 39.15 | 3.21 | 31654 | 1 | 775025.69 | 2 | 429363.62 | 272238.41 | 2 | 2 | 106687.94 | 2 | 491705.22 | 498855.56 | 2 | 2 | 2 | 376040.56 | 335109.97 | 2 | 96184.21 | 2 | 2 | 287751.16 |
| P53702 | Hccs | Holocytochrome c-type synthase | 31.22 | 4.78 | 30978 | 1 | 2 | 2 | 2 | 2 | 2 | 2 | 2 | 2 | 2 | 2 | 2 | 2 | 758453.5 | 597143.69 | 2 | 2 | 2 | 690492.56 | 2 | 549053.19 |
| D3ZN95 | Hcfc1 | Host cell factor 1 | 334.83 | 5.56 | 209153 | 8 | 2310467.5 | 2341547 | 2788222.5 | 2822241.5 | 2566342 | 510359.06 | 1858113.5 | 1787471.88 | 737341.88 | 2647640 | 5315207.5 | 4730599 | 1521441.88 | 3045461.75 | 1636776 | 4776912 | 970201.69 | 2 | 763179.56 | 2 |
| P08103 | Hck | Tyrosine-protein kinase HCK | 150.33 | 7.63 | 59129 | 3 | 2 | 2 | 2 | 2 | 1420908.75 | 2 | 2 | 877668.69 | 1369271.75 | 457328.25 | 2 | 435183.72 | 2 | 2 | 2 | 862718.75 | 2 | 2 | 571154 | 700939.62 |
| P49710 | Hcls1 | Hematopoietic lineage cell-specific protein | 331.7 | 12.14 | 54240 | 5 | 2 | 2 | 2 | 2 | 657500.44 | 2 | 2 | 2 | 1656552.25 | 431413.28 | 1713799.5 | 6783646.5 | 1696582.75 | 3450192 | 927273.25 | 1275204.25 | 897277.81 | 2 | 1687206.75 | 1107950.38 |
| O09106 | Hdac1 | Histone deacetylase 1 | 94.87 | 6.02 | 55075 | 2 | 770184.94 | 282217.19 | 270335.94 | 569360.81 | 348383.53 | 166106.81 | 319236.78 | 369468.44 | 844082 | 378217.38 | 2 | 239766.56 | 537499.56 | 629998.81 | 630378.81 | 2 | 2 | 234616.89 | 2 | 676618 |
| P70288 | Hdac2 | Histone deacetylase 2 | 112.23 | 5.94 | 55302 | 2 | 2 | 2 | 2 | 644874.06 | 681028.31 | 673017.62 | 450580.47 | 2 | 180290.78 | 508127.16 | 2 | 2 | 192039.83 | 377775.81 | 298943.78 | 2 | 2 | 98171.52 | 2 | 2 |
| P51859 | Hdgf | Hepatoma-derived growth factor | 275.83 | 28.69 | 26269 | 6 | 4579929 | 7275435.5 | 4926937.5 | 3396626.5 | 14147392 | 6760271.5 | 5775285 | 4802708 | 8461824 | 5649474.5 | 13440667 | 17661998 | 13602244 | 11135436 | 6351667.5 | 13529121 | 14034460 | 4621100 | 11375784 | 10690850 |
| Q8VDJ3 | Hdlbp | Vigilin | 458.18 | 9.38 | 141743 | 10 | 2512035 | 1492917.62 | 2756468.75 | 232590.34 | 3539645.25 | 2 | 745216.44 | 2323438 | 2759139.5 | 1650937.25 | 1938252.25 | 2776141.75 | 3953943 | 3845122.75 | 1872110.62 | 4525798 | 4088758.5 | 1719850.12 | 1205821.25 | 919447.56 |
| Q8BQM4 | Heatr3 | HEAT repeat-containing protein 3 | 33.78 | 1.91 | 74305 | 1 | 2 | 2 | 2 | 2 | 133553.69 | 2 | 132760.5 | 130832.45 | 214513.45 | 186065.62 | 2 | 2 | 81858.62 | 148570.47 | 2 | 2 | 2 | 2 | 2 | 2 |
| Q60848 | Hells | Lymphocyte-specific helicase | 37.56 | 1.22 | 95126 | 1 | 2 | 2 | 2 | 2 | 2 | 2 | 2 | 2 | 2 | 2 | 2 | 588570.69 | 2 | 2 | 2 | 2 | 2 | 2 | 2 | 2 |
| E9QAM5 | Helz2 | 3'-5' exoribonuclease HELZ2 | 95.52 | 0.71 | 331563 | 2 | 2 | 2 | 2 | 2 | 353925.38 | 2 | 2 | 570051.75 | 345391.47 | 2 | 2 | 2 | 2 | 2 | 2 | 2 | 2 | 2 | 2 | 2 |
| P29416 | Hexa | Beta-hexosaminidase subunit alpha | 430.36 | 20.45 | 60613 | 9 | 11239542 | 9282127 | 9346251 | 12898934 | 8747906 | 3946593.75 | 5545161.5 | 6881814.5 | 7755726 | 7295286 | 2764089.5 | 2039271.25 | 792291.81 | 2729312.5 | 3822945.5 | 4363559.5 | 3434400 | 6897750 | 4884175 | 4994353.5 |

| | | | | | | | | | | | | | | | | | | | | | | | | | | |
|---|---|---|---|---|---|---|---|---|---|---|---|---|---|---|---|---|---|---|---|---|---|---|---|---|---|---|
| P20060 | Hexb | Beta-hexosaminidase subunit beta | 353.4 | 17.72 | 61116 | 8 | 19787152 | 25215262 | 21984138 | 24970668 | 20180386 | 11218148 | 12726519 | 7381907 | 17365548 | 17356838 | 8912201 | 7231532.5 | 8537964 | 8499397 | 11826519 | 6433805 | 10468561 | 12581139 | 9981005 | 15225826 |
| Q99L13 | Hibadh | 3-hydroxyisobutyrate dehydrogenase, mitochondi | 322.32 | 28.36 | 35440 | 6 | 2 | 2621118 | 228263.44 | 1843738.75 | 4913798 | 5090706 | 4773189 | 6555788 | 6295202 | 6012854 | 3418446 | 2449322.75 | 1502430.75 | 2331817 | 2065595.25 | 1363039 | 752755 | 309442.44 | 1550039 | 2 |
| Q8QZS1 | Hibch | 3-hydroxyisobutyryl-CoA hydrolase, mitochondria | 222.23 | 14.29 | 43038 | 4 | 1248424.5 | 1439076.25 | 1369824.75 | 2 | 1942068 | 1525298.12 | 2963168.5 | 1438493.38 | 2135557.25 | 1676540.75 | 1235569.88 | 1479652.88 | 1123783.38 | 108911.26 | 801324.38 | 807870.75 | 2 | 2 | 2 | 2 |
| Q9JLZ6 | Hic2 | Hypermethylated in cancer 2 protein | 33.92 | 1.62 | 66766 | 1 | 2 | 2 | 2 | 2 | 2 | 2 | 2 | 2 | 2 | 2 | 2 | 2 | 2 | 2 | 2 | 2 | 2 | 2 | 2 | 2 |
| Q9JLR9 | Higd1a | HIG1 domain family member 1A, mitochondrial | 219.15 | 46.32 | 10425 | 4 | 25770.93 | 2 | 154956.16 | 230581.56 | 204226.03 | 421279.34 | 385193.78 | 190912.72 | 579368.12 | 140592.53 | 31739.14 | 69905.26 | 180482.31 | 614238.31 | 707762.25 | 2 | 505764.78 | 219505.78 | 2 | 328058.34 |
| P62959 | Hint1 | Adenosine 5'-monophosphoramidase HINT1 | 100.33 | 18.25 | 13777 | 2 | 5476761 | 9991302 | 6193847.5 | 5337748 | 15595922 | 5710656 | 5679682.5 | 8666787 | 7546599.5 | 12919569 | 18494190 | 21858152 | 18282924 | 17580766 | 18741080 | 27873396 | 24914500 | 24222898 | 33652446 | 20117758 |
| Q9D0S9 | Hint2 | Adenosine 5'-monophosphoramidase HINT2 | 43.74 | 7.98 | 17320 | 1 | 2 | 2 | 2 | 2 | 1039813.69 | 3880632.25 | 1666347.25 | 2713246 | 2764918.5 | 1002608.75 | 3325129.25 | 2115428.5 | 1857201.75 | 1952721.88 | 690175.44 | 2151002 | 1213091 | 2 | 2787937.25 | 1493163.25 |
| Q8VD75 | Hip1 | Huntingtin-interacting protein 1 | 100.57 | 2.62 | 115202 | 2 | 2 | 549195.38 | 2 | 2 | 992629.12 | 2 | 2 | 2 | 482044 | 2 | 1330436.25 | 2 | 1007411.44 | 762428.44 | 729034.88 | 2 | 2 | 680857.94 | 686411.62 | 2 |
| Q8BLH7 | Hirip3 | HIRA-interacting protein 3 | 40.47 | 3.66 | 65215 | 1 | 73038.48 | 2 | 2 | 2 | 55096.05 | 2 | 2 | 2 | 89732.72 | 74670.57 | 2 | 192380.08 | 2 | 2 | 2 | 2 | 2 | 2 | 2 | 2 |
| P17710 | Hk1 | Hexokinase-1 | 944.78 | 20.74 | 108303 | 18 | 13060458 | 30745764 | 24910678 | 17082470 | 34166966 | 22835026 | 22992392 | 17073234 | 29715316 | 26320124 | 16544660 | 28764784 | 25086312 | 23532986 | 13279902 | 14515094 | 14482957 | 17428220 | 28088060 | 10454567 |
| O08528 | Hk2 | Hexokinase-2 | 717.61 | 18.76 | 102535 | 15 | 512493.81 | 265491.81 | 783772.19 | 522957.19 | 2402984 | 2316517 | 2888890 | 1622379.75 | 4856330.5 | 3623743.75 | 2939062.5 | 2291841.25 | 1815647.25 | 965715.75 | 1405528.5 | 2625344.75 | 4722898 | 4279405 | 3108770.75 | 3588519 |
| Q3TRM8 | Hk3 | Hexokinase-3 | 1424.12 | 33.19 | 100101 | 23 | 23562036 | 27690690 | 31346234 | 15946866 | 19989834 | 41054846 | 40949226 | 38174998 | 37476278 | 27504036 | 27184270 | 22011450 | 25849816 | 26634206 | 25436138 | 20303106 | 29758342 | 28309516 | 25037410 | 32954806 |
| Q9D8V0 | Hm13 | Minor histocompatibility antigen H13 | 217.29 | 8.2 | 41748 | 3 | 2455394.5 | 3476177.75 | 898065.25 | 3072537.75 | 270824 | 2737621.75 | 3083517.25 | 428842.12 | 3717952.75 | 1899122.25 | 1626031.12 | 2650128.5 | 3612896 | 4022900.75 | 3503772.25 | 2124906.25 | 4759831.5 | 6173265.5 | 3275465 | 6051317 |
| P19356 | Hmbs | Porphobilinogen deaminase | 115.77 | 6.37 | 39361 | 2 | 43372.33 | 117652.48 | 289625.75 | 151480.09 | 137607.19 | 134099.14 | 211582.59 | 62694.96 | 212505.95 | 219082.88 | 89699.34 | 280996 | 162981.5 | 499531.38 | 1320207.75 | 2 | 2 | 101025.65 | 160243.25 | 76396.27 |
| P63158 | Hmgb1 | High mobility group protein B1 | 183.08 | 23.26 | 24894 | 4 | 3850828.5 | 5465282 | 2869491.25 | 9230905 | 12269052 | 1990566.62 | 1902141.5 | 5558779 | 5795475.5 | 6177003.5 | 2400657.5 | 3852208.5 | 2761465 | 2077118.62 | 2465305.25 | 9592140 | 4183629.5 | 3578681.5 | 8142885.5 | 5488916 |
| P30681 | Hmgb2 | High mobility group protein B2 | 269.84 | 32.86 | 24162 | 6 | 3082232.5 | 7183966.5 | 3336177.25 | 18103126 | 24251446 | 4705446 | 4737182 | 1613351.25 | 11207679 | 7761622 | 14228186 | 19717074 | 13372695 | 13959133 | 13469926 | 18534950 | 15273206 | 9408636 | 20038068 | 10758207 |
| P38060 | Hmgcl | Hydroxymethylglutaryl-CoA lyase, mitochondrial | 490.55 | 35.08 | 34239 | 8 | 2713818.75 | 4619631 | 4973925 | 3492790 | 5131860 | 4572965.5 | 4326210.5 | 4349983 | 6684221 | 4779825.5 | 10180738 | 10605768 | 10540576 | 11068789 | 8515280 | 10566430 | 8757616 | 9084900 | 7901771 | 7933410 |
| P17425 | Hmgcs1 | Hydroxymethylglutaryl-CoA synthase, cytoplasmi | 78.55 | 4.04 | 57434 | 2 | 2 | 2 | 2 | 408224.91 | 1463533.5 | 2 | 2 | 421242.44 | 2 | 2 | 633482.81 | 2 | 2 | 2 | 2 | 2 | 2 | 2 | 2 | 2 |
| B4F777 | Hmgn5 | High mobility group nucleosome-binding domain- | 44.59 | 3.5 | 48605 | 1 | 2 | 2 | 2 | 2 | 2 | 2 | 2 | 2 | 293763.5 | 2 | 2 | 2 | 514935.41 | 2 | 2 | 2 | 2 | 2 | 2 | 630210.69 |
| P14901 | Hmox1 | Heme oxygenase 1 | 176.91 | 14.19 | 32929 | 3 | 2 | 2 | 2 | 2 | 2 | 2 | 2 | 2 | 1132158 | 1119616 | 848793.31 | 743616.81 | 2 | 722943.5 | 2 | 921283.12 | 606736.31 | 2 | 1322747.62 | 2 |
| O70252 | Hmox2 | Heme oxygenase 2 | 342.55 | 18.73 | 35739 | 6 | 5140745.5 | 7255867.5 | 5456864 | 3549320.25 | 7016504 | 7399380.5 | 3617681 | 9453534 | 2718303.5 | 5579456.5 | 2750839.5 | 4995264.5 | 8226347.5 | 7458900.5 | 4639691 | 2727896.5 | 2966126 | 2066780 | 10417049 | 2059548 |
| Q9CX86 | Hnrnpa0 | Heterogeneous nuclear ribonucleoprotein A0 | 378.53 | 23.61 | 30530 | 7 | 2436725 | 790609.25 | 2454423.75 | 1421463 | 2207557 | 453599.81 | 715436.25 | 358507.16 | 3464387 | 1875700.25 | 1743464.75 | 3419569.25 | 3824330 | 3411377 | 3645874.75 | 2031702.25 | 3052354.25 | 2806693 | 752510.12 | 3271676.25 |
| P04256 | Hnrnpa1 | Heterogeneous nuclear ribonucleoprotein A1 | 922.24 | 42.5 | 34212 | 13 | 10459879 | 8020497.5 | 9060255 | 6834988 | 9241633 | 6145070 | 6693154.5 | 5860919.5 | 12310248 | 10629502 | 11583894 | 13928928 | 11371166 | 9031798 | 18682666 | 13406964 | 8257569 | 11159551 | 6544095 | 9789867 |
| O88569 | Hnrnpa2b1 | Heterogeneous nuclear ribonucleoproteins A2/B1 | 1006.83 | 46.46 | 37403 | 15 | 6558997.5 | 9216480 | 5543396.5 | 13811721 | 22830874 | 13345935 | 12015672 | 15930502 | 19338902 | 13158522 | 50640878 | 49511358 | 20395110 | 25400744 | 12787041 | 33485988 | 15608739 | 7200536 | 30131604 | 9683344 |
| Q6URK4 | Hnrnpa3 | Heterogeneous nuclear ribonucleoprotein A3 | 846.37 | 28.23 | 39652 | 12 | 5360409.5 | 2552453.75 | 20511932 | 2967093 | 11096946 | 4304718 | 4387730 | 8741099 | 8841004 | 13837883 | 13461314 | 13992212 | 16005755 | 11111562 | 22547284 | 12904070 | 8409687 | 11104448 | 8298703.5 | 8564512 |
| Q99020 | Hnrnpab | Heterogeneous nuclear ribonucleoprotein A/B | 401.98 | 22.81 | 30831 | 8 | 1065221.25 | 1798585.62 | 1505761.38 | 6333071 | 2370979 | 5681750 | 6311393 | 8445076 | 4849376.5 | 6148774.5 | 5830914 | 8269768.5 | 3026429.75 | 3972671.25 | 2297546 | 4054620.5 | 2519935.5 | 1691347.62 | 2146369.75 | 6835261 |
| G3V9R8 | Hnrnpc | Heterogeneous nuclear ribonucleoprotein C | 862.31 | 47.65 | 32857 | 15 | 8545651 | 19419796 | 14325024 | 13031076 | 14694468 | 10859555 | 8183293.5 | 11668376 | 19432452 | 20242870 | 12242507 | 18303082 | 20935620 | 25126764 | 11950789 | 8134016 | 17814424 | 8781343 | 9146659 | 11123706 |
| Q60668 | Hnrnpd | Heterogeneous nuclear ribonucleoprotein D0 | 265.89 | 16.62 | 38354 | 5 | 55863.88 | 2 | 427055.19 | 590160.81 | 461078.22 | 844248.62 | 1147341.12 | 1350896.88 | 672583.44 | 407706.06 | 2 | 208132.94 | 390894.94 | 2 | 645556.69 | 56532.18 | 2 | 2 | 83240.72 | 2 |
| Q794E4 | Hnrnpf | Heterogeneous nuclear ribonucleoprotein F | 788.38 | 27.47 | 45730 | 10 | 10841449 | 14709660 | 9357838 | 17219558 | 11303217 | 10977457 | 9167369 | 9999884 | 7280592 | 7521879 | 8846536 | 11484136 | 8916365 | 9449960 | 9588626 | 10587237 | 10126870 | 10852775 | 14276218 | 14015202 |
| O35737 | Hnrnph1 | Heterogeneous nuclear ribonucleoprotein H | 637.01 | 31.85 | 49199 | 10 | 949454.06 | 1723862.25 | 1234388.62 | 1692359.75 | 2521066 | 879649.06 | 700714.12 | 202986.61 | 618831.38 | 663010.88 | 1988888.75 | 2226868 | 2310739.5 | 1495003.5 | 2023818.75 | 1448498 | 287894.69 | 987214 | 454018.28 | 2 |
| P70333 | Hnrnph2 | Heterogeneous nuclear ribonucleoprotein H2 | 338.03 | 17.59 | 49280 | 5 | 117122.43 | 2 | 211868.23 | 312136.69 | 2 | 2 | 325923.41 | 303463.16 | 231375.95 | 373507.75 | 2 | 2 | 2 | 2 | 243597.8 | 2 | 2 | 2 | 2 | 2 |
| P61979 | Hnrnpk | Heterogeneous nuclear ribonucleoprotein K | 996.27 | 37.8 | 50976 | 17 | 7852576 | 18073958 | 17409362 | 30763140 | 28724942 | 18411612 | 19099608 | 25529170 | 28681204 | 27520128 | 37159174 | 39831810 | 37411566 | 43743558 | 31724326 | 33389050 | 30941152 | 33351522 | 42603974 | 32481794 |
| F1LQ48 | Hnrnpl | Heterogeneous nuclear ribonucleoprotein L | 702.06 | 26.65 | 67903 | 12 | 8078394 | 11682807 | 4391476 | 12318553 | 5053131 | 9879114 | 8760970 | 5351971 | 18972768 | 11996494 | 15704475 | 25198792 | 17523854 | 19698384 | 12949119 | 17124440 | 15093348 | 12263827 | 17884082 | 13788538 |
| Q921F4 | Hnrnpll | Heterogeneous nuclear ribonucleoprotein L-like | 103.29 | 5.25 | 64125 | 2 | 2 | 2 | 2 | 2 | 2 | 2 | 2 | 2 | 664092.75 | 55911.63 | 79104.88 | 526119.5 | 163192.05 | 2 | 2 | 40934.68 | 2 | 2 | 2 | 2 |
| Q9D0E1 | Hnrnpm | Heterogeneous nuclear ribonucleoprotein M | 1784.12 | 44.72 | 77649 | 31 | 7878616 | 46795074 | 11486768 | 27636146 | 62377458 | 51131622 | 40563206 | 57236146 | 83394586 | 34916734 | 54857442 | 54895574 | 30826768 | 45772246 | 31779582 | 42702278 | 29878382 | 25979770 | 33275916 | 29419130 |
| Q6IMY8 | Hnrnpu | Heterogeneous nuclear ribonucleoprotein U | 1445.71 | 32.71 | 87732 | 23 | 5522746 | 5998203.5 | 6943782 | 12461687 | 17619528 | 5581190 | 5510473.5 | 11481591 | 12676835 | 10846247 | 15979684 | 16191684 | 16072622 | 16118166 | 17914556 | 14062087 | 13087083 | 10635705 | 15987252 | 12866124 |
| Q8VDM6 | Hnrnpul1 | Heterogeneous nuclear ribonucleoprotein U-like p | 167.7 | 6.17 | 96002 | 4 | 1370033.75 | 2 | 2 | 2 | 55364.53 | 2 | 120899.16 | 910225.25 | 144267.31 | 255116.22 | 584656.44 | 2 | 766887.69 | 300848.03 | 1320530 | 2 | 158826.3 | 2 | 2 | 2 |
| Q00PI9 | Hnrnpul2 | Heterogeneous nuclear ribonucleoprotein U-like p | 531.35 | 12.75 | 84940 | 11 | 10308112 | 9392481 | 9700701 | 11237351 | 11179613 | 12924202 | 10077641 | 11027016 | 10390541 | 8395536 | 11626056 | 11625682 | 15840958 | 14011494 | 8366131.5 | 9351326 | 11196371 | 13046958 | 11826352 | 12429449 |
| Q3TEA8 | Hp1bp3 | Heterochromatin protein 1-binding protein 3 | 695.71 | 26.9 | 60867 | 14 | 7017691 | 8743677 | 4710205 | 8029991 | 12240141 | 5102715 | 8237344.5 | 3382355.5 | 9300695 | 7529281 | 1809590.75 | 5209964.5 | 4447484 | 3023248 | 5152734.5 | 3807166 | 2187949.75 | 1009430.62 | 4658403 | 1720478.5 |
| P00493 | Hprt1 | Hypoxanthine-guanine phosphoribosyltransferase | 126.49 | 8.26 | 24570 | 3 | 6627092 | 15152750 | 14653589 | 13976667 | 13879412 | 8680914 | 7890257 | 9818608 | 7660291.5 | 7672655 | 8157725 | 10190010 | 6757119.5 | 6642590 | 3641746 | 11204917 | 5331092.5 | 4193182 | 9311485 | 4898591 |
| Q3TC93 | Hs1bp3 | HCLS1-binding protein 3 | 49.36 | 4.81 | 43691 | 1 | 861790.12 | 1053739.38 | 974514.62 | 1240309.38 | 809862.62 | 1044180.62 | 618456.31 | 1195642.5 | 638137.44 | 529090.94 | 1311798.38 | 1414839.5 | 1148283.12 | 994599.25 | 665566.44 | 1191153.88 | 1139490.62 | 1038954.25 | 1471851.5 | 1211891.25 |
| Q8R3H7 | Hs2st1 | Heparan sulfate 2-O-sulfotransferase 1 | 43.3 | 2.53 | 41813 | 1 | 2 | 2 | 973951.44 | 368905.66 | 730519.81 | 130584.73 | 2 | 2 | 485938.09 | 550506.62 | 2 | 2 | 144406.98 | 334251.81 | 2 | 2 | 141590.77 | 2 | 2 | 2 |
| O08756 | Hsd17b10 | 3-hydroxyacyl-CoA dehydrogenase type-2 | 473.01 | 34.48 | 27419 | 6 | 806355.25 | 2 | 363963.47 | 2 | 256350.61 | 1248042.88 | 1757767.12 | 128835.84 | 1396878 | 2291441 | 835074.75 | 751220.5 | 1304254.75 | 178072.88 | 2 | 1271165 | 1339998.75 | 1070183.38 | 1264898.88 | 2 |
| Q9EQ06 | Hsd17b11 | Estradiol 17-beta-dehydrogenase 11 | 125.95 | 9.4 | 32881 | 2 | 985710.25 | 1361469.88 | 824658.75 | 481430.25 | 784571.25 | 1359958.38 | 1593604.62 | 720085.56 | 1831164.75 | 1504905.75 | 345350.59 | 2 | 74769.93 | 2 | 417029.38 | 938852.94 | 2 | 2 | 2 | 282193.94 |
| O70503 | Hsd17b12 | Very-long-chain 3-oxoacyl-CoA reductase | 862.82 | 56.73 | 34742 | 16 | 11685065 | 13341166 | 7106831 | 16014349 | 11538698 | 14953956 | 15093060 | 7045533.5 | 17224804 | 17645932 | 18152814 | 18317658 | 28618138 | 24939064 | 20177930 | 18462868 | 27264064 | 36997754 | 28010094 | 32055632 |
| P51660 | Hsd17b4 | Peroxisomal multifunctional enzyme type 2 | 823.67 | 24.08 | 79482 | 14 | 9412022 | 12243839 | 4974717.5 | 13527073 | 19139908 | 5706372 | 4344616.5 | 8310890 | 8382862.5 | 7739732 | 9557995 | 10067171 | 6230256 | 6954089 | 4753569.5 | 9694496 | 4688102.5 | 2231365.5 | 2970547.5 | 2730009.5 |
| O88736 | Hsd17b7 | 3-keto-steroid reductase/17-beta-hydroxysteroid | 221.56 | 14.37 | 37317 | 3 | 504994.44 | 2 | 2 | 1419971.5 | 1083621 | 2 | 1127527.62 | 258421.61 | 1436447.5 | 2086647.12 | 2 | 132941.36 | 833657.12 | 260795.7 | 2 | 701911.12 | 258559.95 | 119354.96 | 1442912 | 2 |
| P50171 | Hsd17b8 | (3R)-3-hydroxyacyl-CoA dehydrogenase | 246.92 | 25.1 | 26588 | 5 | 1987736.25 | 5179427 | 3279023.5 | 3722370.25 | 7928870 | 7176205.5 | 6310508.5 | 6383840.5 | 8740154 | 6269748.5 | 5615901 | 3387573.5 | 4580250.5 | 4770076 | 1447891.25 | 3470275 | 14431367 | 10378049 | 4269041 | 9106470 |
| Q4V8B7 | Hsdl1 | Inactive hydroxysteroid dehydrogenase-like prote | 50.98 | 3.33 | 36908 | 1 | 29243.71 | 2 | 112473.22 | 2 | 47639.64 | 2 | 225342.64 | 2 | 2 | 128354.25 | 2 | 2 | 2 | 2 | 2 | 2 | 146989.69 | 92514.53 | 2 | 134429.25 |
| P07901 | Hsp90aa1 | Heat shock protein HSP 90-alpha | 1417.43 | 33.56 | 84788 | 26 | 23226138 | 22802902 | 31683052 | 19700340 | 27727098 | 23346550 | 21657966 | 24514866 | 20062826 | 22863532 | 38366622 | 39885490 | 42053810 | 40173150 | 27697450 | 26814400 | 28390496 | 29914310 | 24680134 | 19510208 |
| P11499 | Hsp90ab1 | Heat shock protein HSP 90-beta | 2310.58 | 50.41 | 83281 | 38 | 89718114 | 93755842 | 103388802 | 114726002 | 146057618 | 70864202 | 70147162 | 112504674 | 87233026 | 121164938 | 105517818 | 104953474 | 146953506 | 141412642 | 137717986 | 106797906 | 93954842 | 120082194 | 116033594 | 90480122 |
| P08113 | Hsp90b1 | Endoplasmin | 2649.79 | 51.75 | 92476 | 45 | 95544482 | 103226754 | 106415474 | 117562722 | 126410458 | 95517858 | 80913866 | 133763882 | 104901946 | 110233634 | 116723962 | 99557090 | 124294570 | 121970034 | 103052850 | 124766106 | 108889522 | 102907810 | 98570890 | 102182346 |
| Q6AYB4 | Hspa14 | Heat shock 70 kDa protein 14 | 38.21 | 3.14 | 54463 | 1 | 2 | 492732.88 | 2 | 807909.62 | 1143635.88 | 2 | 597046.31 | 2 | 738204.94 | 475463.19 | 2 | 2 | 544739.56 | 2 | 2 | 896742.94 | 2 | 2 | 2 | 2 |
| P14659 | Hspa2 | Heat shock-related 70 kDa protein 2 | 1022.49 | 23.22 | 69642 | 16 | 3608016.75 | 2 | 4945848 | 2678544 | 4165102 | 1685509.75 | 1429633.12 | 3553170 | 1331129.25 | 2118857.75 | 3032791.75 | 2 | 2375560.25 | 1844850.88 | 1692340.25 | 899591 | 991360.19 | 1431126 | 2 | 1678671.12 |
| Q61316 | Hspa4 | Heat shock 70 kDa protein 4 | 1274.29 | 28.54 | 94133 | 21 | 4907650.5 | 6197434 | 4315949 | 6441114.5 | 9485550 | 4693512 | 4998828 | 6856160 | 7299876 | 6586147 | 6877633 | 7760550 | 7106261 | 5599524 | 4793567.5 | 6851360 | 7097180 | 5962984.5 | 8602663 | 5287595.5 |
| P06761 | Hspa5 | Endoplasmic reticulum chaperone BiP | 2764.99 | 53.52 | 72347 | 42 | 346360226 | 334654530 | 306070562 | 220382018 | 304953154 | 291061922 | 263029602 | 323999938 | 268681154 | 330389986 | 407069858 | 360238050 | 393813954 | 351305218 | 257148962 | 407122146 | 334416578 | 359918754 | 388411842 | 340466754 |
| P63017 | Hspa8 | Heat shock cognate 71 kDa protein | 3151.25 | 68.27 | 70871 | 47 | 421192898 | 431482370 | 373439234 | 421580098 | 450136930 | 284628066 | 257253202 | 317597506 | 297581762 | 362076450 | 358839650 | 359159010 | 313883746 | 310589186 | 372030082 | 269497954 | 320537218 | 317056450 | 216600722 | 301734146 |
| P38647 | Hspa9 | Stress-70 protein, mitochondrial | 2285.94 | 52.43 | 73461 | 32 | 72239962 | 83420410 | 65089530 | 65672986 | 90745834 | 99586074 | 82172226 | 98045706 | 114207554 | 110264730 | 140855410 | 128422482 | 117922594 | 127593634 | 112759482 | 135692866 | 136480514 | 125622346 | 116359058 | 110647066 |
| Q6IMX7 | Hspbp1 | Hsp70-binding protein 1 | 163.31 | 12.61 | 39191 | 3 | 2276276 | 2767235 | 2667044.25 | 4319962 | 3298623.75 | 336641.5 | 1060189.5 | 602820.38 | 177643.69 | 2344810.5 | 2 | 1270804.12 | 1660076.38 | 1809124 | 2272670.75 | 2 | 955965.06 | 2 | 1231669.25 | 2074187.5 |
| P63038 | Hspd1 | 60 kDa heat shock protein, mitochondrial | 3091.51 | 74.52 | 60955 | 46 | 227914146 | 257834194 | 190733042 | 196728514 | 184257490 | 197493762 | 197040962 | 175385682 | 189324306 | 193010818 | 168421026 | 177226866 | 192883250 | 197512018 | 187562658 | 177580770 | 203482674 | 228764610 | 178564578 | 213139298 |
| Q64433 | Hspe1 | 10 kDa heat shock protein, mitochondrial | 356.76 | 58.82 | 10963 | 6 | 10439635 | 8318584 | 10826918 | 8130042.5 | 13182447 | 8634696 | 7068411 | 9117255 | 9908042 | 11227066 | 13994695 | 12701828 | 9393244 | 8233345 | 8043454 | 22809618 | 9476530 | 9899103 | 11359955 | 10804694 |
| Q61699 | Hsph1 | Heat shock protein 105 kDa | 222.94 | 7.46 | 96407 | 4 | 779539.62 | 839142.06 | 78498.63 | 279943.88 | 424913.59 | 149567.8 | 322768.25 | 324333.03 | 2 | 383367.75 | 105892.6 | 96328.48 | 2 | 286543.56 | 642057.75 | 2 | 2 | 286402.28 | 133961.84 | 2 |
| Q9Z2G9 | Htatip2 | Oxidoreductase HTATIP2 | 213.75 | 24.79 | 26870 | 5 | 2967663.75 | 2058720 | 2879740.5 | 1757414 | 2519933.75 | 4710595.5 | 2793840.25 | 2269217.5 | 2118901.5 | 2283326.5 | 1985841.12 | 1219077.75 | 2395411.75 | 2964929 | 2421348 | 390380.5 | 2325389.75 | 1661721.38 | 1874837.75 | 5496863 |
| Q9JIY5 | Htra2 | Serine protease HTRA2, mitochondrial | 327.29 | 24.24 | 49348 | 7 | 1262615.88 | 2 | 58630.38 | 2 | 1007276.25 | 153194.03 | 146968.27 | 63749.01 | 2 | 581959.44 | 1485032 | 628318.12 | 603383.38 | 894101.69 | 958396.44 | 2 | 258828.73 | 2 | 156604.81 | 415012.97 |
| Q7TMY8 | Huwe1 | E3 ubiquitin-protein ligase HUWE1 | 226.28 | 1.64 | 482635 | 6 | 928049.19 | 1128819.88 | 945011.19 | 1342788.25 | 1402715 | 910651.38 | 544374.38 | 972533.81 | 692609.38 | 808628.5 | 780228.44 | 1364424.62 | 1440775.75 | 1975544.75 | 1398590.75 | 1226988.62 | 1042421.44 | 1789290.5 | 1382888.62 | 2061584.88 |
| Q8R1F5 | Hyi | Putative hydroxypyruvate isomerase | 207.87 | 17.69 | 30449 | 4 | 8325826 | 9724301 | 7996059 | 1643181 | 5490223.5 | 1619728.12 | 1407834 | 2940198 | 955364.75 | 3381309 | 7881924 | 6480740.5 | 6322256.5 | 5693454 | 3507237 | 10113052 | 4366669.5 | 4503192 | 4029066 | 4717910.5 |
| Q9JKR6 | Hyou1 | Hypoxia up-regulated protein 1 | 1559.98 | 32.03 | 111181 | 29 | 14034649 | 15869185 | 18684938 | 16594852 | 23663950 | 11755493 | 9986856 | 18485838 | 15746182 | 22933858 | 15742639 | 15060052 | 18559676 | 16435358 | 18230654 | 16747741 | 14110181 | 15847406 | 14257796 | 17872962 |
| Q9DB29 | Iah1 | Isoamyl acetate-hydrolyzing esterase 1 homolog | 52.63 | 4.42 | 27974 | 1 | 670690.75 | 701784.06 | 2 | 396089.5 | 832990.56 | 2 | 852807.31 | 763483.44 | 769763.44 | 663899.75 | 1778268.38 | 1640602.12 | 1448604.62 | 975405.25 | 2 | 2 | 677659.75 | 2 | 660174.31 | 2 |
| P03975 | Iap | IgE-binding protein | 596.92 | 21.72 | 62747 | 10 | 3158270.5 | 8585078 | 1157145.62 | 8936034 | 3726811 | 2283924.5 | 1138331.5 | 2300628 | 3192655.25 | 2101543.75 | 3051770 | 6191416 | 2933300.75 | 7688233.5 | 4390638.5 | 3076217.5 | 2447235 | 1125301.88 | 1234312.25 | 1967349.88 |
| Q8BU30 | Iars1 | Isoleucine--tRNA ligase, cytoplasmic | 358.88 | 5.31 | 144271 | 8 | 3628220 | 4470157 | 7016193 | 5277355.5 | 6110085 | 4526498 | 5788787.5 | 7741787 | 4927516.5 | 8039221.5 | 2212312.75 | 2340377.5 | 4766107.5 | 4669551.5 | 6190179 | 3706931.5 | 3955375.5 | 3403219 | 3087055.5 | 2975305.75 |
| Q8BIJ6 | Iars2 | Isoleucine--tRNA ligase, mitochondrial | 36.7 | 1.58 | 112804 | 1 | 1544536.62 | 1239451.38 | 1014972.94 | 2 | 1160876 | 1344363.25 | 864929.88 | 1871212.5 | 1268032.75 | 1022977.25 | 2 | 2 | 839964.38 | 1027827.94 | 2 | 2 | 2 | 1320939.12 | 2 | 2 |
| P13597 | Icam1 | Intercellular adhesion molecule 1 | 212.3 | 8.75 | 58844 | 4 | 863116.69 | 835744.38 | 635151.62 | 1105764.38 | 716141 | 2 | 2 | 690183 | 869789.06 | 1334654.25 | 623308.38 | 574125.69 | 87507.64 | 1051055.88 | 1285167.25 | 1391707.25 | 1490174.75 | 1601001.62 | 793571.81 | 2 |
| P35559 | Ide | Insulin-degrading enzyme | 55.19 | 1.37 | 117710 | 1 | 2 | 2 | 2 | 2 | 2 | 2 | 2 | 2 | 2 | 2 | 2 | 2 | 1110882.5 | 2 | 2 | 2 | 2 | 2 | 1343344 | 2 |
| O88844 | Idh1 | Isocitrate dehydrogenase [NADP] cytoplasmic | 644.15 | 31.16 | 46674 | 11 | 20527038 | 14397177 | 17684364 | 17129154 | 15659572 | 15205847 | 16632154 | 8955653 | 12934239 | 10098615 | 15787068 | 17975946 | 18152646 | 19246306 | 12473038 | 17075798 | 18199240 | 18784320 | 15225931 | 21651568 |
| P54071 | Idh2 | Isocitrate dehydrogenase [NADP], mitochondrial | 746.43 | 34.96 | 50906 | 14 | 12789399 | 9806794 | 9344818 | 9947449 | 13101064 | 3825227 | 7784194 | 11129286 | 8093268.5 | 11195578 | 16654475 | 12904393 | 12621396 | 14847141 | 12017250 | 11611859 | 9269064 | 12455971 | 12174978 | 8720933 |
| Q99NA5 | Idh3a | Isocitrate dehydrogenase [NAD] subunit alpha, m | 418.51 | 25.96 | 39614 | 8 | 4581385 | 4815212 | 10644220 | 2813986 | 12128070 | 3241415.5 | 4844836.5 | 7462025 | 8166586 | 9396930 | 9410803 | 7641888 | 9908200 | 9958612 | 10767001 | 12155461 | 8559182 | 11806556 | 5351262 | 11783401 |
| Q68FX0 | Idh3B | Isocitrate dehydrogenase [NAD] subunit beta, mit | 169.05 | 11.95 | 42354 | 4 | 2 | 605165.38 | 2 | 2 | 2 | 2 | 2 | 2 | 2 | 2 | 2 | 378894.84 | 663085 | 284883.78 | 2 | 1439013.12 | 545379.81 | 721086.25 | 1000910 | 901698.88 |
| P41565 | Idh3g | Isocitrate dehydrogenase [NAD] subunit gamma 1 | 209.09 | 10.94 | 42851 | 4 | 7617618.5 | 4496663 | 2441692.5 | 4935940.5 | 6113410.5 | 5405264 | 4569616 | 8516608 | 4424851 | 4244958.5 | 6291270 | 5641899.5 | 5819643 | 5072660 | 3225908.5 | 8551313 | 5337318.5 | 6503307 | 8736024 | 5051814 |
| P58044 | Idi1 | Isopentenyl-diphosphate Delta-isomerase 1 | 195.18 | 22.91 | 26289 | 4 | 2 | 100884.09 | 2 | 2 | 1087807 | 2 | 1178474.75 | 2 | 251541.09 | 589479.44 | 67025.71 | 2 | 130177.34 | 131243.45 | 2 | 2 | 2 | 2 | 402116.5 | 2 |
| Q9R002 | Ifi202 | Interferon-activable protein 202 | 124.9 | 4.27 | 50466 | 1 | 2 | 2 | 2 | 2 | 2 | 1825709 | 1069292.12 | 1148803.75 | 620579.81 | 2 | 2 | 2 | 2 | 2 | 2 | 2 | 2 | 2 | 753406.12 | 616056.44 |
| P0DOV2 | Ifi204 | Interferon-activable protein 204 | 185.47 | 7.43 | 69438 | 5 | 514680 | 557228 | 2 | 1441750 | 2120659.5 | 2045103.75 | 4366755 | 6472017 | 8597468 | 3732020 | 338369.5 | 1101439.62 | 1280522.5 | 2 | 2 | 523101.03 | 822663.5 | 1012132.81 | 590420 | 2 |
| Q9ESY9 | Ifi30 | Gamma-interferon-inducible lysosomal thiol reduc | 248.78 | 20.16 | 27784 | 5 | 8190037.5 | 7046190.5 | 7372462.5 | 5182271 | 4723032.5 | 4666918 | 3878838.5 | 5109326 | 3600058 | 3031614 | 3755892 | 4130874 | 1635717.5 | 4664179.5 | 4194713 | 1341894.12 | 3011742 | 5898206 | 3000273.5 | 3622252.5 |
| Q9D8C4 | Ifi35 | Interferon-induced 35 kDa protein homolog | 37.95 | 5.94 | 31876 | 1 | 2 | 2 | 2 | 2 | 43802.77 | 2 | 102307.15 | 2 | 86027.57 | 2 | 66633.06 | 2 | 2 | 2 | 150615.12 | 2 | 2 | 2 | 2 | 327371 |
| Q8BV66 | Ifi44 | Interferon-induced protein 44 | 49.62 | 2.61 | 47852 | 1 | 2 | 163905.47 | 2 | 2 | 950171.38 | 852769.19 | 4290667 | 456487.66 | 3013440.75 | 2343628.25 | 2 | 2 | 2 | 2 | 2 | 2 | 2 | 2 | 2 | 2 |
| Q9BDB7 | Ifi44l | Interferon-induced protein 44-like | 160.87 | 8.5 | 49680 | 3 | 2 | 2 | 2 | 2 | 892866.5 | 3487201.75 | 3250437 | 3870667.75 | 2028524.75 | 1905512.88 | 2 | 2 | 2 | 2 | 2 | 2 | 2 | 130491.77 | 2 | 2 |
| Q99J93 | Ifitm2 | Interferon-induced transmembrane protein 2 | 252.04 | 17.36 | 15743 | 3 | 3165886 | 3724648.5 | 2567999.25 | 1737459.12 | 1044223.56 | 3241124 | 2254297.75 | 4923385 | 1606223.25 | 626365.62 | 2 | 2 | 511821.19 | 438021.78 | 2 | 2 | 2 | 789722.88 | 916156.06 | 2 |
| Q9CQW9 | Ifitm3 | Interferon-induced transmembrane protein 3 | 478.05 | 48.91 | 14954 | 5 | 6990885 | 7555690.5 | 6112185.5 | 5423869 | 5122507 | 8250985 | 5130915.5 | 17265762 | 5649892.5 | 3963981.5 | 2 | 2 | 1380408.75 | 463861.12 | 807713.25 | 2014485.25 | 2664617.75 | 5119891.5 | 4826195 | 2118100.5 |
| Q9CPN8 | Igf2bp3 | Insulin-like growth factor 2 mRNA-binding protein | 277.75 | 12.44 | 63575 | 6 | 703394 | 1207648.38 | 617548.56 | 1184910.75 | 1431685.62 | 1331637.5 | 991882.56 | 2 | 1690719.25 | 1232161 | 1874246.25 | 1143705.62 | 629260.38 | 1705438.38 | 510513.88 | 1304975.75 | 1107877.25 | 1332208.25 | 1104272.88 | 1461126.75 |
| Q8R366 | Igsf8 | Immunoglobulin superfamily member 8 | 54.73 | 1.64 | 65011 | 1 | 2 | 456170.06 | 463440.56 | 476403.78 | 2 | 2 | 559354.81 | 2 | 2 | 492764.34 | 872994.5 | 958789.31 | 586208.81 | 426174.09 | 2 | 2 | 611315.06 | 515354.88 | 2 | 2 |
| Q9DCE9 | Igtp | Immunity-related GTPase family M protein 3 | 37.38 | 2.13 | 48494 | 1 | 626401.06 | 114323.32 | 471711.84 | 521948.31 | 738063.31 | 690533.44 | 1650147 | 227845.33 | 2173068 | 1790851.25 | 2 | 2 | 2 | 2 | 2 | 2 | 2 | 2 | 2 | 2 |
| O88351 | Ikbkb | Inhibitor of nuclear factor kappa-B kinase subunit | 97.13 | 3.57 | 86690 | 2 | 2 | 873069.06 | 2 | 2 | 888978.56 | 1306493.25 | 1181910.38 | 951296.25 | 1207512.25 | 418248.03 | 1337411.5 | 1062161.25 | 1058185.25 | 1237493.5 | 2 | 981625.88 | 1170010.5 | 716052 | 2 | 1083021.88 |
| O88522 | Ikbkg | NF-kappa-B essential modulator | 38.21 | 2.67 | 47972 | 1 | 1254715.75 | 2 | 2 | 2 | 2 | 2 | 1088615.88 | 1470450.38 | 2 | 2 | 1954649.62 | 1304034.75 | 895461.12 | 916643.75 | 795657.5 | 1302565.25 | 960578.69 | 2 | 2 | 2 |
| Q9CXY6 | Ilf2 | Interleukin enhancer-binding factor 2 | 213.99 | 13.08 | 43062 | 5 | 2 | 2 | 553533.12 | 2 | 2 | 2 | 2 | 2 | 2 | 2 | 2 | 2 | 2 | 2 | 2 | 2 | 2 | 2 | 2 | 2 |

| | | | | | | | | | | | | | | | | | | | | | | | | | | |
|---|---|---|---|---|---|---|---|---|---|---|---|---|---|---|---|---|---|---|---|---|---|---|---|---|---|---|
| Q9Z1X4 | Ilf3 | Interleukin enhancer-binding factor 3 | 286.64 | 7.13 | 96021 | 6 | 2 | 2 | 2123925.25 | 2 | 1230544.25 | 1149856.12 | 2 | 2 | 523658.31 | 2 | 2 | 2 | 2 | 470648.41 | 2 | 2 | 2 | 1309865.12 | 385761 | 2 |
| O55222 | Ilk | Scaffold protein ILK | 72.09 | 4.2 | 51373 | 2 | 1704309.75 | 3210618.75 | 1674992.75 | 2475245 | 2593341 | 2130170 | 2098161.75 | 1719687.12 | 2285113 | 1508791.5 | 1634618.75 | 610386.81 | 1484026.25 | 2035228.25 | 1130897.38 | 680246.62 | 1417336.75 | 1635344.25 | 727841.44 | 855926.44 |
| Q8R0F6 | Ilkap | Integrin-linked kinase-associated serine/threonin | 38.98 | 2.3 | 42774 | 1 | 1338126.12 | 1132639.38 | 1268474.88 | 2045567 | 534302.56 | 2 | 362474.97 | 1190728.5 | 479854.75 | 1423334.38 | 2 | 682640.62 | 2 | 292527.06 | 677218 | 1445874.62 | 1657035.62 | 1485867.62 | 521472.88 | 1588572.38 |
| Q8BU33 | Ilvbl | 2-hydroxyacyl-CoA lyase 2 | 65.31 | 3.48 | 68156 | 1 | 2 | 2 | 2 | 2 | 39434.28 | 2 | 2 | 2 | 144916.05 | 2 | 2 | 2 | 2 | 2 | 2 | 2 | 2 | 2 | 2 | 2 |
| Q8CAQ8 | Immt | MICOS complex subunit Mic60 | 1805.59 | 39.1 | 83900 | 29 | 15482612 | 23295006 | 17427990 | 24174128 | 21498916 | 23632562 | 23471872 | 21145300 | 34057606 | 30306002 | 39647094 | 34344882 | 35009886 | 38469806 | 26522260 | 41791870 | 32802866 | 37846578 | 31860216 | 31742002 |
| Q5PQR5 | Imp4 | U3 small nucleolar ribonucleoprotein protein IMP4 | 59.03 | 7.22 | 33656 | 2 | 2 | 590944 | 2 | 2 | 229983.75 | 2 | 2 | 2 | 2 | 2 | 2 | 2 | 2 | 2 | 2 | 2 | 2 | 2 | 2 | 2 |
| O55023 | Impa1 | Inositol monophosphatase 1 | 479.53 | 27.08 | 30436 | 10 | 10867726 | 10868682 | 9837946 | 3703185.25 | 8028560 | 4991466 | 6187442 | 8622216 | 5581928.5 | 4602105 | 9779748 | 9818879 | 8875365 | 9268047 | 7563485 | 11730638 | 10468713 | 14370735 | 14189176 | 12830838 |
| Q8CIN7 | Impa2 | Inositol monophosphatase 2 | 38.29 | 3.45 | 31796 | 1 | 907096.62 | 2113842.5 | 473480.91 | 2 | 720545.19 | 2 | 2 | 2 | 2 | 685412.75 | 1171977.62 | 1195013.12 | 599790.5 | 501034.91 | 2 | 2 | 2 | 2 | 2 | 2 |
| P24547 | Impdh2 | Inosine-5'-monophosphate dehydrogenase 2 | 823.83 | 26.26 | 55815 | 13 | 17446900 | 16179832 | 17789444 | 14903950 | 28514204 | 5449529 | 6618694.5 | 10749625 | 5516218.5 | 10050361 | 15306482 | 14230199 | 14013454 | 13951523 | 12153343 | 10350058 | 6980839.5 | 6417638.5 | 5090295 | 4867009 |
| Q9WU62 | Incenp | Inner centromere protein | 109.99 | 4.2 | 101209 | 3 | 2 | 2 | 2 | 2 | 503581.09 | 486744.53 | 431196.41 | 2 | 2 | 2 | 2 | 2 | 2 | 2 | 2 | 2 | 2 | 2 | 2 | 2 |
| Q9ES52 | Inpp5d | Phosphatidylinositol 3,4,5-trisphosphate 5-phospl | 433.67 | 6.8 | 133542 | 9 | 4920945 | 8811288 | 3866741.5 | 7274402.5 | 5717404 | 4282676 | 6107134.5 | 5960528 | 9653415 | 4625890 | 5258778 | 8791083 | 6909862 | 9875524 | 4563017.5 | 7894830 | 5141379 | 2077146.25 | 2648150.25 | 3514385.5 |
| Q6P549 | Inppl1 | Phosphatidylinositol 3,4,5-trisphosphate 5-phospl | 198.93 | 3.5 | 138973 | 4 | 513337.97 | 2602718 | 2 | 1349229.75 | 2003154.25 | 1513524.88 | 1103486 | 2 | 1387282.75 | 1353235.25 | 667118.69 | 2 | 2 | 1011313.5 | 884925.81 | 622936.31 | 2 | 691812.31 | 703463.56 | 1999841.38 |
| Q3MHC2 | Ints11 | Integrator complex subunit 11 | 43.79 | 1.67 | 67841 | 1 | 2 | 2 | 70072.14 | 389436.88 | 140705.73 | 2 | 136732.59 | 86235.33 | 322500.16 | 195767.98 | 48763.33 | 106580.53 | 101640.85 | 201461.83 | 2 | 2 | 314656.44 | 202449.38 | 2 | 252524.05 |
| Q8VI75 | Ipo4 | Importin-4 | 46.08 | 1.57 | 119275 | 1 | 2 | 2 | 60303.5 | 2 | 33030.78 | 102567.49 | 122518.11 | 58072.23 | 49993.58 | 72882.61 | 2 | 77820.64 | 49839.71 | 61975.25 | 91679.59 | 2 | 2 | 2 | 2 | 2 |
| Q8BKC5 | Ipo5 | Importin-5 | 366.51 | 9.12 | 123591 | 7 | 2166792.5 | 1750032.5 | 3342330.25 | 180118.36 | 2687084.5 | 1073976.5 | 2 | 213101.34 | 576055 | 1524193.25 | 2495431.5 | 2078684.88 | 4234558 | 2430324.5 | 2475816 | 1984055.25 | 2163947.5 | 2360061.5 | 2 | 2 |
| Q9EPL8 | Ipo7 | Importin-7 | 38.81 | 1.06 | 119486 | 1 | 481251.06 | 2 | 2 | 100294.14 | 31611.03 | 208238.64 | 137164.77 | 2 | 167087.64 | 368852.69 | 82011.79 | 77560.66 | 146778.97 | 93730.05 | 315523.75 | 2 | 2 | 2 | 2 | 2 |
| Q91YE6 | Ipo9 | Importin-9 | 292.36 | 8.07 | 116052 | 5 | 5661126.5 | 5772475 | 4431324 | 4939674.5 | 5068063.5 | 2326718 | 2362680.5 | 2742037.5 | 2651297.5 | 1563304.5 | 4272772.5 | 3232113.25 | 4295584.5 | 3404548.25 | 2630744.25 | 4011597 | 3580901.75 | 3850355.5 | 3650960.25 | 2961686.75 |
| Q9JKF1 | Iqgap1 | Ras GTPase-activating-like protein IQGAP1 | 2860 | 40.13 | 188742 | 54 | 45486250 | 40236554 | 50791450 | 42160362 | 51043814 | 50010554 | 53459950 | 67535906 | 59378278 | 57945302 | 49686486 | 47107738 | 51163022 | 53226834 | 52519630 | 52345450 | 39549406 | 53170506 | 40452178 | 45854762 |
| Q60664 | Irag2 | Inositol 1,4,5-triphosphate receptor associated 2 | 174.62 | 9.09 | 59588 | 3 | 1000608.44 | 1222429 | 2 | 853738.88 | 587643.75 | 737718.56 | 955276 | 2 | 587610.25 | 2 | 2114071 | 471236.03 | 780434.81 | 2 | 2 | 2 | 1243926.88 | 2 | 1176032.62 | 2 |
| E9Q1P8 | Irf2bp2 | Interferon regulatory factor 2-binding protein 2 | 102.63 | 3.33 | 59292 | 2 | 2 | 776193.19 | 2 | 2041668.25 | 1247223.25 | 2 | 1280449 | 1335833.62 | 929004.56 | 2 | 2260324.75 | 1033999.25 | 625590.94 | 1245563 | 2 | 1315318.5 | 2 | 2 | 1472488.62 | 2 |
| P56477 | Irf5 | Interferon regulatory factor 5 | 52.19 | 2.41 | 56005 | 1 | 2 | 2 | 674068.19 | 744049.19 | 944897.62 | 2 | 2 | 735874.38 | 475758.91 | 504316.41 | 2 | 2 | 2 | 575177.06 | 2 | 2 | 2 | 2 | 2 | 2 |
| Q60766 | Irgm1 | Immunity-related GTPase family M protein 1 | 260.48 | 15.16 | 46552 | 5 | 817056.56 | 511029.62 | 502377.16 | 1397889.12 | 1237583.62 | 2139113 | 2168098.75 | 3554083 | 3490261.25 | 896273.75 | 591074.69 | 69662.84 | 744142.5 | 71264.24 | 152039.31 | 2 | 1177696.38 | 2 | 765447.38 | 179397.58 |
| Q64339 | Isg15 | Ubiquitin-like protein ISG15 | 40.09 | 11.18 | 17898 | 1 | 2 | 2 | 46701.96 | 2 | 2 | 2 | 2 | 2 | 142907.44 | 159444.38 | 810289.06 | 744299.44 | 710402.06 | 1278398.5 | 1048624.62 | 381620.69 | 1937971.62 | 793687.69 | 1464504.88 | 1004523.5 |
| Q6I7R3 | Isoc1 | Isochorismatase domain-containing protein 1 | 80.87 | 8.42 | 32002 | 2 | 498955.91 | 831354.44 | 305753.94 | 992868 | 971512.31 | 579335.38 | 508858.31 | 1025442.75 | 697822.19 | 821014.38 | 279246.12 | 285199.72 | 669245.12 | 503030.16 | 607500.06 | 2 | 266926.25 | 218329.69 | 127783.14 | 2 |
| P85094 | Isoc2a | Isochorismatase domain-containing protein 2A | 226.34 | 33.98 | 22417 | 4 | 1123662.62 | 38672.56 | 1115406 | 262055.09 | 184989.12 | 82801.46 | 558390.62 | 299391.91 | 664725.44 | 1003151.94 | 2 | 242044.41 | 630344.31 | 675390.88 | 1285398.5 | 53972.43 | 277697.19 | 419685.81 | 95664.51 | 325657.53 |
| Q6AYK3 | Isyna1 | Inositol-3-phosphate synthase 1 | 326.71 | 10.59 | 60884 | 7 | 7014309 | 8153096 | 4548130.5 | 5811186 | 4295313.5 | 5693363 | 4435100.5 | 4835266.5 | 3437322.25 | 3552967.25 | 14559522 | 12927007 | 9879809 | 10095990 | 9833470 | 12305065 | 11576954 | 3107036 | 9374090 | 7516114.5 |
| P11688 | Itga5 | Integrin alpha-5 | 756.91 | 20.51 | 115043 | 16 | 6675440.5 | 7377218 | 7774690.5 | 7683086 | 15388904 | 9210694 | 5573587 | 12747213 | 16604475 | 14890976 | 13040438 | 11304105 | 10397728 | 11080205 | 5912746.5 | 9459088 | 2399423.5 | 2654680.75 | 4634072.5 | 1925010.88 |
| P24063 | Itgal | Integrin alpha-L | 44.54 | 0.95 | 128314 | 1 | 453825.03 | 667891.94 | 405872.12 | 787520.19 | 884851.75 | 2 | 426510.38 | 2 | 768646.62 | 627547.31 | 469022.25 | 2 | 2 | 2 | 2 | 2 | 703817.06 | 739349.94 | 583936.31 | 599119.06 |
| P05555 | Itgam | Integrin alpha-M | 966.76 | 18.39 | 127481 | 16 | 13079176 | 11892058 | 15496410 | 14781898 | 26167896 | 13002111 | 13304308 | 19713490 | 25160576 | 24379834 | 9109539 | 8581599 | 8856685 | 7429529 | 4696345 | 7407742 | 2829743.25 | 2151679.25 | 5894723 | 4603535 |
| P09055 | Itgb1 | Integrin beta-1 | 859.43 | 23.68 | 88231 | 17 | 26827254 | 24903076 | 25213880 | 34015998 | 42076114 | 19639934 | 18380142 | 35698314 | 33658262 | 28997768 | 17964546 | 21100814 | 17282256 | 18028760 | 21999884 | 14219216 | 21699568 | 20745810 | 18961220 | 20015696 |
| P11835 | Itgb2 | Integrin beta-2 | 1369.22 | 34.63 | 85026 | 25 | 34598550 | 41012110 | 30867802 | 29820060 | 28389790 | 28077724 | 22246126 | 40234966 | 50593506 | 30388658 | 21049082 | 19726968 | 16605289 | 20482824 | 12470584 | 19564462 | 22706786 | 22723816 | 21980834 | 13978626 |
| O89051 | Itm2b | Integral membrane protein 2B | 51.18 | 4.51 | 30260 | 1 | 2 | 2 | 2 | 2 | 2 | 2 | 2 | 2 | 2 | 2 | 2 | 2 | 2 | 2 | 2 | 2 | 2 | 2 | 2 | 2 |
| Q9D892 | Itpa | Inosine triphosphate pyrophosphatase | 129.93 | 14.14 | 21897 | 3 | 1457228.38 | 1331999.88 | 450398.84 | 961818.62 | 2143054 | 2252746.5 | 2171167.5 | 1974389.38 | 1377916 | 1043222.69 | 874691.62 | 1648771.38 | 677247 | 810205.06 | 698683.06 | 2 | 601340.19 | 625035.38 | 2 | 2 |
| P29995 | Itpr2 | Inositol 1,4,5-trisphosphate-gated calcium channe | 84.9 | 0.59 | 307058 | 2 | 257495.52 | 662316.56 | 2 | 2507555 | 2 | 741941.62 | 2202847.5 | 231457.66 | 503585.09 | 2 | 2 | 1643033.25 | 1618454.38 | 649214.81 | 2 | 2693816.75 | 3785702.75 | 1013701.75 | 94802.89 | 2 |
| P70227 | Itpr3 | Inositol 1,4,5-trisphosphate-gated calcium channe | 91.98 | 0.6 | 304275 | 2 | 2016657.88 | 4048461.75 | 2300514.5 | 2038787.62 | 9579498 | 7074920 | 4463379 | 10200817 | 7178640 | 6831597 | 3563906.75 | 2 | 2303629 | 2444522 | 2077095.75 | 2 | 2982425.5 | 2530269.5 | 3770508.25 | 2860315.5 |
| Q9JHI5 | Ivd | Isovaleryl-CoA dehydrogenase, mitochondrial | 169.63 | 10.38 | 46325 | 4 | 3247137.5 | 4195867 | 3298377.75 | 3265066.5 | 3435865.5 | 3498016.25 | 2936365 | 3610277 | 2584817 | 1456980.5 | 1817727.5 | 1643076.88 | 2000313 | 1599960 | 1280573 | 2306199.5 | 2260022.75 | 2661476.25 | 2226282.25 | 2168133 |
| P97825 | Jpt1 | Jupiter microtubule associated homolog 1 | 40.54 | 9.74 | 16081 | 1 | 2 | 186149.55 | 2 | 81942.3 | 246853.66 | 2 | 2 | 2 | 90371.71 | 82318.56 | 2 | 90575.14 | 144599.5 | 206767.44 | 893351.06 | 289027 | 269024.91 | 239985.27 | 344037.25 | 1533605.88 |
| Q5BK20 | Jpt2 | Jupiter microtubule associated homolog 2 | 99.93 | 14.21 | 20070 | 2 | 1466648.25 | 1573536.75 | 665057.81 | 3362399.25 | 757206.25 | 3674476.5 | 3370277.5 | 2169103.25 | 2101680.75 | 2 | 3595205.5 | 3592477 | 2264140 | 2087067.75 | 1567109.75 | 2106154 | 2296839 | 647897.06 | 2354693.25 | 779488.12 |
| Q02257 | Jup | Junction plakoglobin | 367.57 | 13.42 | 81801 | 7 | 486167.47 | 350257.09 | 2 | 2375681.5 | 755669.5 | 1970374.75 | 1318673.5 | 1977063 | 1521604.88 | 2382981.25 | 2362381.25 | 2 | 2 | 2 | 529674.62 | 2 | 2 | 2 | 1889150.5 | 2 |
| Q99MN1 | Kars1 | Lysine--tRNA ligase | 298.75 | 11.26 | 67840 | 6 | 3159663.75 | 3401951.25 | 7530557.5 | 4091015.75 | 6179307.5 | 2185044 | 4335543 | 4807106.5 | 3891169 | 4828323 | 5070385.5 | 5814482 | 7534855 | 7005761.5 | 7087850 | 6368404 | 6402424 | 7053089 | 7052711 | 6454910 |
| Q6WVG3 | Kctd12 | BTB/POZ domain-containing protein KCTD12 | 315.24 | 18.96 | 35892 | 6 | 12187814 | 13982342 | 4718145.5 | 7368880.5 | 6173133 | 5678404 | 7492858 | 7426598 | 3927506 | 5390041.5 | 12317185 | 10709067 | 9959853 | 9310034 | 9793183 | 11102153 | 8966264 | 8715608 | 13391616 | 8630293 |
| Q6ZQ88 | Kdm1a | Lysine-specific histone demethylase 1A | 57.11 | 1.76 | 92851 | 1 | 2 | 2 | 2 | 2 | 2 | 2 | 2 | 2 | 2 | 2 | 2 | 2 | 2 | 2 | 2 | 2 | 2 | 2 | 2 | 2 |
| Q9CXT6 | Kdm8 | Bifunctional peptidase and arginyl-hydroxylase J | 34.27 | 2.17 | 47145 | 1 | 587004.81 | 741062.12 | 233807.59 | 1112596 | 239098.09 | 2 | 2 | 94136.98 | 872458.88 | 1492081.62 | 5061917.5 | 5748963 | 5361124.5 | 5354169 | 3723322.5 | 6701942.5 | 7829952.5 | 8186082 | 7989992 | 8723644 |
| Q60749 | Khdrbs1 | KH domain-containing, RNA-binding, signal trans | 113.49 | 4.74 | 48371 | 3 | 2 | 2 | 2 | 1419626.38 | 2695246.25 | 2 | 1912155.5 | 1552685.75 | 2292121 | 1786967.75 | 2566521 | 1841689.62 | 2 | 1445221.75 | 2 | 1770499.62 | 2 | 2 | 889321.94 | 2 |
| Q3U0V1 | Khsrp | Far upstream element-binding protein 2 | 1094.47 | 30.75 | 76775 | 20 | 5712495.5 | 20302098 | 13031439 | 11912410 | 19202644 | 12633669 | 9263024 | 12385738 | 13120324 | 12533748 | 36732502 | 37390714 | 26217434 | 32992780 | 24233652 | 28538248 | 36204358 | 20755536 | 41699170 | 21183100 |
| Q6P9P6 | Kif11 | Kinesin-like protein KIF11 | 68.9 | 2.19 | 118027 | 2 | 2 | 902715.19 | 2 | 1435365.25 | 773950.19 | 608150.44 | 2 | 2 | 2 | 2 | 2 | 2 | 970352.12 | 1190545.75 | 2 | 1006158.81 | 2 | 2 | 2 | 2 |
| P28741 | Kif3a | Kinesin-like protein KIF3A | 41.78 | 1.14 | 80170 | 1 | 2 | 755197.94 | 874194.94 | 2 | 909590.94 | 2 | 680464.31 | 96140.14 | 727756.56 | 2 | 2 | 2 | 807602.31 | 858913.25 | 455926.03 | 2 | 2 | 583674.19 | 2 | 2 |
| Q2PQA9 | Kif5b | Kinesin-1 heavy chain | 329.74 | 8.52 | 109531 | 7 | 336752.34 | 2 | 2 | 1196363.5 | 2771889 | 1241017.88 | 2727864.75 | 4118916 | 1804185.75 | 2509107.5 | 1013368.69 | 2 | 1785132.12 | 300910.41 | 344957.25 | 1996148.25 | 1349515.5 | 1584325.88 | 1080194 | 1347221.25 |
| Q9QWT9 | Kifc1 | Kinesin-like protein KIFC1 | 111.94 | 5.34 | 74153 | 2 | 2 | 2 | 2 | 2 | 2 | 2 | 2 | 2 | 2 | 2 | 2 | 2 | 2 | 2 | 2 | 2 | 2 | 2 | 2 | 2 |
| Q9DBS5 | Klc4 | Kinesin light chain 4 | 113.27 | 5.65 | 68613 | 2 | 1184605.75 | 792055.12 | 937761.88 | 2 | 2 | 2 | 682824.69 | 1089125.38 | 437977 | 1078119.75 | 711627.25 | 2 | 2 | 576832.56 | 2 | 2 | 2 | 627482.38 | 2 | 2 |
| P52293 | Kpna2 | Importin subunit alpha-1 | 438.36 | 18.34 | 57928 | 8 | 892339.06 | 1088957.62 | 2143986.75 | 814192.25 | 3481134.75 | 2 | 158659.84 | 894056.69 | 1451458.62 | 1626852.88 | 1143916.25 | 1066026.25 | 852804.62 | 1735035.75 | 1363279.75 | 2 | 2 | 2 | 359489.72 | 2 |
| O35343 | Kpna4 | Importin subunit alpha-3 | 75.9 | 3.26 | 57923 | 2 | 2 | 2 | 2 | 2 | 2 | 2 | 2 | 2 | 2 | 2 | 2 | 2 | 2 | 2 | 2 | 2 | 2 | 2 | 2 | 2 |
| P70168 | Kpnb1 | Importin subunit beta-1 | 613.98 | 15.18 | 97184 | 11 | 18024284 | 18542902 | 18828454 | 20989976 | 22986042 | 12895281 | 14158646 | 18709666 | 12434120 | 17518644 | 13247630 | 12490846 | 14452863 | 14233406 | 18420588 | 11314284 | 9490507 | 12631413 | 11150694 | 13587731 |
| Q8BGA5 | Krr1 | KRR1 small subunit processome component hom | 42.71 | 2.37 | 43538 | 1 | 2 | 2 | 390222.56 | 174732.91 | 345109.66 | 2 | 2 | 118947.33 | 2 | 583036.62 | 86528.86 | 82546.09 | 2 | 86730.58 | 2 | 2 | 2 | 2 | 2 | 2 |
| B2RZC9 | Krtcap2 | Dolichyl-diphosphooligosaccharide--protein glyco | 65.51 | 19.85 | 14662 | 1 | 2 | 866284.75 | 472157.84 | 1383226.12 | 94701.45 | 1079993.88 | 1915256.25 | 1990553.25 | 2937358.75 | 825837.5 | 120960.31 | 2 | 540268.31 | 983681.25 | 833889.81 | 2 | 638481.31 | 2 | 2 | 2 |
| Q58FK9 | Kyat3 | Kynurenine--oxoglutarate transaminase 3 | 48.82 | 2.64 | 51044 | 1 | 2 | 2 | 2 | 2 | 2 | 171284.36 | 2 | 2 | 810113.25 | 573465.88 | 155500.16 | 2 | 2 | 2 | 2 | 2 | 282769.41 | 148014.27 | 103563.77 | 2 |
| P11438 | Lamp1 | Lysosome-associated membrane glycoprotein 1 | 582.22 | 19.95 | 43865 | 8 | 6330820 | 3107480 | 6954134 | 5061107 | 727098.81 | 4503363 | 5053910.5 | 4073976 | 2472249.5 | 2745470.75 | 1192060.25 | 2077354.38 | 2373253 | 1296478 | 5368394 | 3056312 | 5204221 | 4010351 | 2302312.5 | 4414717.5 |
| P17047 | Lamp2 | Lysosome-associated membrane glycoprotein 2 | 317.8 | 14.22 | 45681 | 7 | 56947722 | 43788750 | 45855514 | 33826986 | 22624090 | 38065446 | 40730018 | 38625830 | 33146262 | 37056942 | 18400476 | 16979212 | 17309448 | 15681099 | 24729204 | 20627916 | 28342646 | 21965152 | 3258490.5 | 23858418 |
| Q9CQ22 | Lamtor1 | Ragulator complex protein LAMTOR1 | 178.38 | 23.6 | 17749 | 3 | 602626.31 | 113806.23 | 672737.12 | 141168.66 | 882906.62 | 286843.78 | 205106.03 | 229184.73 | 373496.31 | 761756.12 | 98325.8 | 88993.02 | 630381.06 | 217988.72 | 2 | 2 | 250080.88 | 2 | 110911.59 | 2 |
| Q9JHS3 | Lamtor2 | Ragulator complex protein LAMTOR2 | 43.02 | 8.8 | 13480 | 1 | 2 | 2 | 1203445.25 | 460301.78 | 616191.69 | 2 | 2 | 992607.62 | 785435 | 793454.12 | 1004879.38 | 2 | 623846.5 | 404215.56 | 1293733.38 | 2 | 2 | 615837.94 | 2 | 2 |
| Q9CPY7 | Lap3 | Cytosol aminopeptidase | 358.2 | 19.65 | 56141 | 8 | 660336.38 | 3107061 | 3090968 | 1423083 | 3455596.5 | 1938068.25 | 3713015.25 | 3267388.5 | 4673844.5 | 3421197.25 | 1880793.38 | 3211309.75 | 3918387.5 | 5509800.5 | 4868407.5 | 3463354 | 835267.5 | 3788409.75 | 255791.17 | 2568184 |
| Q8BWW4 | Larp4 | La-related protein 4 | 213.06 | 5.7 | 79763 | 4 | 680021.69 | 1343232.75 | 369664.75 | 515094.84 | 2433230.75 | 2 | 2 | 702442.25 | 273196 | 1096200.12 | 3048434.25 | 2490558.5 | 1369187.75 | 958279.38 | 757073.31 | 2584939.5 | 2 | 1115129.75 | 2035226.5 | 1548661.5 |
| Q05CL8 | Larp7 | La-related protein 7 | 108.97 | 4.74 | 64802 | 2 | 1846141.25 | 655038.44 | 1274829.88 | 2513206 | 3134402.5 | 2 | 786017.69 | 1016477.06 | 1760145.12 | 819524.5 | 2 | 2 | 1970279.5 | 692084.12 | 767396.5 | 1405361.5 | 2626531.5 | 1254619.88 | 1381517.5 | 2052266.75 |
| Q8BMJ2 | Lars1 | Leucine--tRNA ligase, cytoplasmic | 442.31 | 9.17 | 134192 | 9 | 596488.12 | 1541827.62 | 2873427.75 | 2252988.5 | 3986591.75 | 2 | 1276140 | 2 | 2081350 | 2610340.75 | 6691633 | 4431670 | 2218212 | 3147778 | 2397936.75 | 2887659 | 1335867.75 | 2 | 3874827.5 | 484101.16 |
| Q61792 | Lasp1 | LIM and SH3 domain protein 1 | 480.51 | 33.46 | 29994 | 9 | 3079353.25 | 2501092.5 | 2487438.75 | 9394012 | 7429395.5 | 19475224 | 17196160 | 4064809 | 14479289 | 5231668.5 | 24225150 | 31162676 | 12979984 | 18550266 | 13604862 | 22630266 | 21510112 | 19416878 | 17571968 | 26931716 |
| Q3U9G9 | Lbr | Delta(14)-sterol reductase LBR | 726.91 | 16.93 | 71440 | 12 | 2454437.25 | 1493452.75 | 1806088.62 | 4156904.5 | 2222882.25 | 2182053 | 1615546.12 | 3042618.25 | 3131925.75 | 1806102.25 | 1368570 | 2221228.75 | 3071014 | 2562455.5 | 2168040.5 | 1926677.88 | 3362708 | 2530550.5 | 2722118.75 | 2891362.75 |
| Q3UN02 | Lclat1 | Lysocardiolipin acyltransferase 1 | 37.28 | 3.46 | 44400 | 1 | 2 | 2 | 2 | 2 | 2 | 2 | 2 | 2 | 2 | 536793.75 | 2 | 2 | 2 | 2 | 2 | 2 | 2 | 2 | 854649.88 | 2 |
| Q6P4Z6 | Lcmt1 | Leucine carboxyl methyltransferase 1 | 40.58 | 2.71 | 38330 | 1 | 2 | 710461.75 | 651559.56 | 1892093.88 | 1440344.5 | 2506589.75 | 2158924.75 | 2 | 2 | 255416.25 | 1037906.12 | 436063.81 | 405522.12 | 2 | 2 | 911304.38 | 2 | 755983.69 | 2 | 2 |
| Q61233 | Lcp1 | Plastin-2 | 2754.9 | 68.26 | 70149 | 44 | 282716802 | 154179330 | 285835394 | 153986370 | 289649922 | 299847874 | 174975186 | 279381154 | 268740226 | 300474498 | 319804226 | 349517634 | 292643810 | 279633986 | 275633698 | 279517954 | 293071074 | 122616850 | 316699394 | 235950802 |
| P06151 | Ldha | L-lactate dehydrogenase A chain | 1308.42 | 62.95 | 36499 | 21 | 107720930 | 114408562 | 113041034 | 111911970 | 185814290 | 178791042 | 179128546 | 224662658 | 257644754 | 209821682 | 378680290 | 430477794 | 234942530 | 269703106 | 193557730 | 322319906 | 219004674 | 167771970 | 288724514 | 154744930 |
| P35951 | Ldlr | Low-density lipoprotein receptor | 158.32 | 5.57 | 94947 | 4 | 886900.88 | 757575.06 | 546568.69 | 4491406 | 2364405 | 2 | 2 | 108581.01 | 3211197.25 | 715026.06 | 383655.53 | 821627.06 | 613677.31 | 871592.56 | 2 | 936667.88 | 2 | 2 | 2 | 2 |
| Q9Z2I0 | Letm1 | Mitochondrial proton/calcium exchanger protein | 44.39 | 3.25 | 82989 | 1 | 2 | 2 | 2 | 201700.94 | 2 | 2 | 2 | 272414.06 | 149626.73 | 152320.19 | 2 | 2 | 105822.29 | 2 | 396847.84 | 2 | 251489.61 | 2 | 2 | 2 |
| P11762 | Lgals1 | Galectin-1 | 467.38 | 41.48 | 14857 | 6 | 49418446 | 54335358 | 40190382 | 49369270 | 63991886 | 70464026 | 71938010 | 81416442 | 57921298 | 62958006 | 82270634 | 85292090 | 77860538 | 74018418 | 69789218 | 60154234 | 79010378 | 58068374 | 45451058 | 66530626 |
| P16110 | Lgals3 | Galectin-3 | 533.1 | 38.64 | 27515 | 9 | 3127265 | 5263680 | 5333368.5 | 4447551 | 19641258 | 20913506 | 11014831 | 27253018 | 28267058 | 18261660 | 104838130 | 112329122 | 53973854 | 64469754 | 27105718 | 112963626 | 55047658 | 31998546 | 130974898 | 27602866 |
| Q07797 | Lgals3bp | Galectin-3-binding protein | 1071.16 | 28.77 | 64491 | 15 | 14545500 | 15074563 | 14051970 | 16025310 | 9934597 | 18567562 | 19561450 | 12657042 | 19593790 | 21898162 | 2355002.75 | 6603900.5 | 5928308.5 | 5573111.5 | 4431728 | 4639060 | 6752272.5 | 12554632 | 8493354 | 10972786 |
| O08573 | Lgals9 | Galectin-9 | 54.64 | 2.27 | 40036 | 1 | 1424258.88 | 3599328 | 744543.56 | 5550667.5 | 6586659 | 5355388 | 4426052.5 | 7327019 | 3947723.25 | 2198478.25 | 2 | 350567.38 | 746666.88 | 2 | 1163614.25 | 1121253.5 | 2 | 1235750.12 | 2 | 843971.19 |
| P37913 | Lig1 | DNA ligase 1 | 88.93 | 3.28 | 102290 | 2 | 823656.38 | 127370.51 | 2 | 2 | 2 | 2 | 2 | 2 | 2 | 2 | 2 | 2 | 2 | 72181.85 | 2 | 2 | 2 | 2 | 2 | 2 |
| Q64281 | Lilrb4a | Leukocyte immunoglobulin-like receptor subfamil | 119.24 | 8.66 | 37544 | 2 | 1134973.38 | 2 | 1063177 | 2335450.5 | 1652565 | 2 | 1807207 | 2362030.75 | 2809913.5 | 2348706.75 | 1699496.38 | 2 | 1409432 | 1647965.5 | 2 | 2 | 2 | 1465591.5 | 689566.31 | 1959527.62 |
| Q9ERG0 | Lima1 | LIM domain and actin-binding protein 1 | 338.87 | 9.83 | 84060 | 7 | 6374886 | 7938304.5 | 2 | 4734849 | 1405553.12 | 2953555.25 | 604411.44 | 1670986.25 | 1603965.25 | 892129 | 13107990 | 5254336 | 3664395.75 | 9057916 | 5024596 | 4705284.5 | 3940388.75 | 8139275 | 3501655.25 | 5202580 |
| Q9D0F3 | Lman1 | Protein ERGIC-53 | 387.25 | 16.63 | 57789 | 8 | 6471760 | 9967613 | 6017640.5 | 8241291 | 11374695 | 8688213 | 8293158.5 | 10977572 | 8470292 | 11254372 | 11458425 | 10007521 | 10427032 | 12079297 | 8214031.5 | 12342276 | 9490184 | 8232485.5 | 14733898 | 8011833 |
| Q9DBH5 | Lman2 | Vesicular integral-membrane protein VIP36 | 33.89 | 3.35 | 40430 | 1 | 2 | 2 | 2 | 2 | 383405.16 | 2 | 2 | 2 | 477859.09 | 618117.12 | 2 | 2 | 559782.06 | 563865.19 | 2 | 2 | 2 | 2 | 2 | 2 |
| P59481 | Lman2l | VIP36-like protein | 45.85 | 4.32 | 39881 | 1 | 729340.56 | 555897.12 | 612265.56 | 2 | 436525.16 | 773208.19 | 2 | 2 | 640565.06 | 579522.44 | 498670.22 | 456803.97 | 605638.94 | 467388.28 | 652936.69 | 2 | 852728.62 | 1418580.38 | 659158.06 | 897624.94 |
| Q3U3R4 | Lmf1 | Lipase maturation factor 1 | 107.77 | 4.18 | 65878 | 2 | 2 | 2 | 2 | 1521914.75 | 2 | 2458131.5 | 2 | 2521072.25 | 836621 | 989659.31 | 706263.12 | 1061436.62 | 2817876.5 | 731677 | 2009238.88 | 2601880.5 | 1731799 | 1296884.25 | 2885979.5 | 2 |
| P48678 | Lmna | Prelamin-A/C | 2793.48 | 64.51 | 74238 | 47 | 186118754 | 252908290 | 148856210 | 170201762 | 196911938 | 217218674 | 204236594 | 218828130 | 217859138 | 205597266 | 293497250 | 306827426 | 230333074 | 240524578 | 179006322 | 280767234 | 197092050 | 194189346 | 200509810 | 188474674 |
| P14733 | Lmnb1 | Lamin-B1 | 1693.8 | 50 | 66786 | 32 | 20599522 | 16295477 | 10162278 | 16120838 | 18020970 | 30831242 | 12080993 | 11945370 | 12040372 | 24748378 | 16461793 | 11028343 | 9449685 | 10029842 | 10544206 | 12603257 | 11096241 | 9537561 | 17884888 | 6706702.5 |
| P21619 | Lmnb2 | Lamin-B2 | 379.17 | 13.09 | 67318 | 8 | 4169303.5 | 1628607 | 2943413.5 | 2732309 | 1342188.75 | 2515277.5 | 3509571.75 | 2480076.5 | 1207378.12 | 2 | 753934.44 | 776071 | 1597652.25 | 1182075.25 | 1788158.5 | 2691811.5 | 1302216.38 | 1010086 | 549936.88 | 932836.56 |
| Q8C129 | Lnpep | Leucyl-cystinyl aminopeptidase | 445.89 | 8.78 | 117304 | 9 | 1983611.75 | 4761126.5 | 1151500.88 | 4171673.75 | 6015312 | 1574130.5 | 6639135 | 8819496 | 8457968 | 4503275 | 6513262 | 6205095.5 | 5754197 | 5730634.5 | 2 | 6997071 | 2599078.5 | 2 | 4977944 | 2145955 |
| Q8CGK3 | Lonp1 | Lon protease homolog, mitochondrial | 844.89 | 25.08 | 105843 | 17 | 6567353.5 | 4798552.5 | 5443349.5 | 4223956.5 | 7961036.5 | 11063631 | 10650989 | 9479042 | 14065289 | 13852630 | 11698115 | 9505348 | 12599022 | 13572474 | 9914388 | 13300126 | 13773298 | 14880314 | 16567199 | 12039036 |
| P0C1Q3 | Lpcat2 | Lysophosphatidylcholine acyltransferase 2 | 184.74 | 11.21 | 59829 | 4 | 24546.79 | 2 | 2 | 2 | 2 | 2 | 2 | 2 | 118184.4 | 77406.05 | 2 | 132129.88 | 2 | 2 | 2 | 2 | 2 | 2 | 2 | 2 |
| Q8BYI6 | Lpcat2 | Lysophosphatidylcholine acyltransferase 2 | 349.02 | 18.75 | 60254 | 7 | 604018.69 | 1097276.38 | 92632.55 | 704812.5 | 44169.73 | 121768.02 | 398620.41 | 2 | 1536761.38 | 859650.25 | 368829.88 | 75224.69 | 1440277.25 | 123832.88 | 366040.28 | 1844387.25 | 2963896.5 | 2 | 2 | 2552038 |
| Q99N69 | Lpxn | Leupaxin | 97.42 | 5.7 | 43478 | 2 | 3449954.75 | 4921646 | 2246346 | 10380922 | 7231579 | 16328573 | 11070160 | 3368663.5 | 6890972 | 11908918 | 11452689 | 12564826 | 6624395.5 | 4966429 | 2233405 | 12761656 | 7732851 | 2691573.25 | 10289313 | 2739281 |
| Q921G6 | Lrch4 | Leucine-rich repeat and calponin homology doma | 55.43 | 2.5 | 73163 | 1 | 2 | 2 | 2 | 2 | 2 | 2 | 2 | 2 | 2 | 32411.53 | 2 | 2 | 2 | 2 | 2 | 2 | 2 | 2 | 2 | 148716.58 |
| Q91ZX7 | Lrp1 | Prolow-density lipoprotein receptor-related protei | 1964.86 | 10.03 | 504742 | 38 | 59843758 | 56183086 | 52469482 | 21953134 | 20906382 | 26477534 | 23861280 | 30367874 | 27461818 | 24147710 | 48515506 | 37448442 | 41971162 | 39814874 | 30783598 | 33513182 | 32547198 | 29584292 | 38614550 | 28688740 |

| | | | | | | | | | | | | | | | | | | | | | | | | | | |
|---|---|---|---|---|---|---|---|---|---|---|---|---|---|---|---|---|---|---|---|---|---|---|---|---|---|---|
| P55302 | Lrpap1 | Alpha-2-macroglobulin receptor-associated protei | 45.47 | 3.06 | 42215 | 1 | 1310670.88 | 944931.31 | 1683833.12 | 2 | 616399.31 | 2 | 610690.19 | 1272537.25 | 554617.31 | 1172855.88 | 2 | 762287.25 | 473910.72 | 2 | 1330654.5 | 853843.5 | 901698.5 | 1299037.75 | 2 | 2 |
| Q6PB66 | Lrpprc | Leucine-rich PPR motif-containing protein, mitoch | 779.99 | 13.86 | 156615 | 16 | 3194229.25 | 1025297.75 | 2872567.25 | 2433918 | 1990722.25 | 755490.31 | 590910.69 | 1428126.62 | 1197390.75 | 2450604.25 | 959433.25 | 925052.12 | 2306404.5 | 1870507.38 | 861150.12 | 1372832.5 | 2542339 | 2278362.5 | 53081.96 | 1512830.62 |
| D3ZXS4 | Lrrc39 | Leucine-rich repeat-containing protein 39 | 40.83 | 2.1 | 38716 | 1 | 1052112.25 | 449771.78 | 751369.75 | 649557.31 | 2 | 2 | 1004614.69 | 2 | 627725.56 | 531910.81 | 2 | 732109.06 | 2 | 2 | 1253055.25 | 2 | 2 | 481660.09 | 2 | 2 |
| Q505F5 | Lrrc47 | Leucine-rich repeat-containing protein 47 | 57.11 | 3.1 | 63590 | 1 | 2 | 2 | 2 | 2 | 2 | 2 | 2 | 2 | 166222.72 | 2 | 2 | 2 | 2 | 2 | 2 | 2 | 2 | 2 | 2 | 2 |
| Q5RJR8 | Lrrc59 | Leucine-rich repeat-containing protein 59 | 210.08 | 19.87 | 34869 | 5 | 1734386.38 | 1295067.75 | 4483953 | 338891.03 | 4258369 | 294676.19 | 1906291 | 4065258.5 | 2688617.75 | 5815032.5 | 1033262.38 | 154654.81 | 169477.89 | 1485401.38 | 4021226 | 2 | 236306.22 | 790257.19 | 2 | 2 |
| Q8R502 | Lrrc8c | Volume-regulated anion channel subunit LRRC8C | 51.43 | 1.74 | 92372 | 1 | 2 | 2 | 2 | 892325.12 | 748198.44 | 2 | 680809.62 | 2 | 741968.31 | 692979 | 2 | 2 | 2 | 519141.59 | 2 | 2 | 2 | 772457.94 | 731650.06 | 2 |
| Q3UZ39 | Lrrfip1 | Leucine-rich repeat flightless-interacting protein 1 | 546.07 | 18.52 | 79249 | 8 | 2625397.5 | 5709532.5 | 488388.03 | 6881398 | 5834521 | 4324096 | 6043938 | 8378474.5 | 9619907 | 4985294 | 6705957.5 | 11344484 | 8392904 | 8992784 | 7395230 | 9144341 | 8390963 | 6001216 | 7639786.5 | 6415298 |
| Q4V7E8 | Lrrfip2 | Leucine-rich repeat flightless-interacting protein 2 | 158.38 | 9.15 | 49773 | 3 | 2 | 566235.88 | 99642.81 | 975157.12 | 152176.89 | 712488.56 | 2 | 136110.95 | 719622.25 | 2 | 1395980.25 | 522120.72 | 992723.38 | 769369.75 | 712426.56 | 859349.19 | 881323.38 | 2 | 965887.56 | 2 |
| Q9D0R8 | Lsm12 | Protein LSM12 | 70.44 | 6.67 | 21701 | 1 | 2 | 2 | 2 | 2 | 2 | 2 | 2 | 2 | 2 | 2 | 2079250.12 | 1835252.25 | 876052.56 | 992875.75 | 2 | 1424545.62 | 1042521.94 | 2 | 1725788.62 | 2 |
| O35900 | Lsm2 | U6 snRNA-associated Sm-like protein LSm2 | 90.38 | 20 | 10835 | 1 | 2 | 208257.94 | 1463995 | 2 | 626805.88 | 2 | 2 | 2 | 2 | 648672.5 | 366660.06 | 246444.05 | 2 | 258641.25 | 2 | 2 | 2 | 2 | 2 | 2 |
| Q9QXA5 | Lsm4 | U6 snRNA-associated Sm-like protein LSm4 | 35.34 | 7.3 | 15076 | 1 | 4228523 | 2575980 | 4531583 | 1808680.38 | 1787296.25 | 5435069.5 | 3957563 | 3956109.25 | 2256694.25 | 2790208.25 | 2014468.62 | 1952771 | 2761591.5 | 1730145.38 | 3526862 | 2185815.75 | 2342952.5 | 2137582.75 | 1119057.12 | 3001498.5 |
| Q9CQQ8 | Lsm7 | U6 snRNA-associated Sm-like protein LSm7 | 99.87 | 15.53 | 11636 | 2 | 2178610.5 | 1354643.62 | 2680846.25 | 606957.31 | 840858.94 | 585085.44 | 315877.31 | 665404.69 | 363822.34 | 684464.06 | 1628829.38 | 1468809.38 | 1283933.25 | 1097639.62 | 1795357.5 | 882415.19 | 1910421.25 | 2 | 1197170.75 | 1654181.5 |
| Q6ZWM4 | Lsm8 | U6 snRNA-associated Sm-like protein LSm8 | 258.41 | 40.62 | 10403 | 4 | 6243289 | 1983133.75 | 6020018.5 | 6628222 | 7567423 | 3974720.25 | 3569950.75 | 5703832.5 | 3419935.75 | 3455295 | 4982778 | 4183651 | 3750944.25 | 3823411.5 | 1855136.38 | 3552554 | 4323715.5 | 4483218 | 3887779 | 5068622.5 |
| P19973 | Lsp1 | Lymphocyte-specific protein 1 | 535.53 | 29.7 | 36714 | 8 | 14110519 | 17164270 | 11376011 | 13671353 | 11952719 | 10026948 | 7991208 | 5507593.5 | 11162857 | 14086948 | 78403722 | 83259778 | 52048194 | 56271646 | 31221914 | 67572890 | 47556506 | 26809206 | 51803542 | 33848426 |
| Q8BLN5 | Lss | Lanosterol synthase | 209.45 | 6.41 | 83141 | 4 | 2 | 2 | 103244.32 | 467896.16 | 2 | 827457.12 | 665304 | 95375.59 | 759424.19 | 421871.88 | 215322.97 | 940441.75 | 1142981.5 | 1329781.75 | 1807262.88 | 1230484.88 | 1394468.5 | 1532039.12 | 1531865.88 | 1616098.25 |
| P24527 | Lta4h | Leukotriene A-4 hydrolase | 223.42 | 9.82 | 69051 | 4 | 2036869.75 | 1020593.12 | 1820357.75 | 609087.75 | 1719378.25 | 212814.55 | 1679254.75 | 2380851.5 | 1540015 | 1869862.75 | 1573164 | 2000940.5 | 3354102 | 1709402 | 1095291.88 | 1021245.31 | 2 | 682014.81 | 862657.62 | 1406496.25 |
| Q7TNC4 | Luc7l2 | Putative RNA-binding protein Luc7-like 2 | 124.21 | 7.65 | 46583 | 3 | 39366010 | 9667734 | 20542664 | 9372468 | 4557080.5 | 1290556.88 | 4249531.5 | 5341372 | 5224170 | 4593896.5 | 6681122.5 | 7028992 | 7142603.5 | 9114070 | 17385852 | 6426350.5 | 12322592 | 25041840 | 8826499 | 28069978 |
| Q5SUF2 | Luc7l3 | Luc7-like protein 3 | 212.25 | 8.56 | 51450 | 3 | 518779.44 | 549127.81 | 923922.25 | 941653.94 | 2675796 | 2 | 837703.06 | 1361167.38 | 900765.75 | 780294 | 677754.88 | 641636.31 | 1168980 | 501099.88 | 1512084.75 | 1194845.12 | 2 | 221851.95 | 108119.1 | 132290.33 |
| O88188 | Ly86 | Lymphocyte antigen 86 | 105.36 | 12.35 | 17811 | 2 | 5962006 | 3942528 | 6847746 | 4621028 | 1492208 | 3556491.25 | 3341332.25 | 5555572.5 | 1765903.62 | 996857.44 | 1665492.88 | 1668475.5 | 1450068.5 | 1049544.12 | 1920573.62 | 1602359.12 | 1811834 | 2987291.25 | 2288644.75 | 2369450.75 |
| Q08288 | Lyar | Cell growth-regulating nucleolar protein | 95.82 | 4.12 | 43736 | 2 | 2 | 99567.28 | 2 | 2 | 1037746 | 2 | 2 | 2 | 2 | 2 | 2 | 2 | 2 | 231392.72 | 2 | 2 | 292346.91 | 2 | 2 | 2 |
| P25911 | Lyn | Tyrosine-protein kinase Lyn | 553.23 | 24.02 | 58812 | 9 | 7142423.5 | 6123755.5 | 6448264 | 4608900.5 | 6531477.5 | 4185935.5 | 4722080 | 2879500.5 | 5550315.5 | 6349535.5 | 3081475.25 | 3749627.5 | 5235096 | 4034160 | 3497931 | 3521082.5 | 2951183 | 4583672.5 | 3326545 | 665942 |
| P97823 | Lypla1 | Acyl-protein thioesterase 1 | 254.91 | 27.39 | 24688 | 5 | 1976983.38 | 1847292.5 | 3186613.5 | 2871296.5 | 5703772.5 | 3649371.75 | 3567566.25 | 5554238 | 3702374.5 | 4623683 | 3834090.25 | 3758392.25 | 2919558 | 2831489 | 3417850.5 | 1708077.12 | 2688875.75 | 2219295.25 | 3423399.25 | 2265902.75 |
| P08905 | Lyz2 | Lysozyme C-2 | 539.68 | 67.57 | 16689 | 8 | 21099438 | 21372710 | 14986886 | 7911338 | 3895966.75 | 9630939 | 7339043 | 12571372 | 5710584.5 | 6615426.5 | 4716345 | 5142657 | 6084753 | 4160403 | 4950248 | 2864506.25 | 399978.75 | 6272601.5 | 4116865 | 4273050.5 |
| P24668 | M6pr | Cation-dependent mannose-6-phosphate recepto | 506.57 | 37.05 | 31172 | 8 | 24228282 | 26947624 | 27690430 | 26603188 | 23629946 | 21542216 | 19402504 | 34811482 | 15435472 | 25723824 | 12102146 | 12758232 | 15710543 | 12905508 | 19714754 | 13549334 | 16554526 | 20038508 | 16413667 | 17525036 |
| Q9QZQ8 | Macroh2a1 | Core histone macro-H2A.1 | 317.04 | 21.41 | 39290 | 5 | 2 | 708878.5 | 2 | 2 | 994500.06 | 2 | 2 | 2 | 1705542.25 | 689217.69 | 2 | 2 | 2 | 659737.88 | 2 | 1184817.25 | 2 | 2 | 2 | 2 |
| O54790 | Mafg | Transcription factor MafG | 80.97 | 15.43 | 17877 | 2 | 2 | 2 | 2 | 2 | 2 | 2 | 2 | 2 | 2 | 2 | 2 | 2 | 2 | 2 | 2 | 2 | 2 | 2 | 2 | 2 |
| Q6AYQ0 | Magix | PDZ domain-containing protein MAGIX | 36.76 | 3.07 | 34743 | 1 | 2 | 2473175.25 | 2 | 2 | 531263.06 | 1798487.75 | 1622262.25 | 2 | 459339.81 | 840886.88 | 1525048.75 | 1635678.12 | 2 | 2 | 2 | 2 | 2 | 315913.22 | 2 | 2 |
| P61327 | Magoh | Protein mago nashi homolog | 121.63 | 22.6 | 17164 | 3 | 2348122 | 2082727.88 | 1903218 | 3845321.25 | 3150051 | 1204063.5 | 2148695.75 | 2458895.25 | 2625433.5 | 2866454.25 | 1359911.5 | 2279630.75 | 2677077.5 | 1232353.38 | 2350264.75 | 1216546.88 | 1936521.88 | 1006958.12 | 237932.48 | 2170874.5 |
| Q9CQY5 | Magt1 | Dolichyl-diphosphooligosaccharide--protein glyco | 35.56 | 2.39 | 37970 | 1 | 1024919.69 | 545759.69 | 627943.12 | 2 | 818830.12 | 2 | 625248.25 | 193443.89 | 422855.62 | 2 | 2 | 2 | 735982.88 | 676701.06 | 808275.75 | 2 | 687051.19 | 746696.75 | 2 | 2 |
| Q8BGS0 | Mak16 | Protein MAK16 homolog | 74.56 | 8.45 | 35185 | 2 | 2 | 2 | 2 | 2 | 2 | 2 | 2 | 2 | 2 | 2 | 2 | 2 | 2 | 2 | 2 | 2 | 2 | 2 | 2 | 2 |
| O09159 | Man2b1 | Lysosomal alpha-mannosidase | 797.5 | 18.85 | 114648 | 14 | 14770489 | 11099904 | 20591526 | 6421226.5 | 8288575.5 | 3331001 | 3827680.5 | 2870970 | 7657755 | 10505126 | 13590139 | 9726657 | 14436526 | 11857461 | 10291874 | 7943811.5 | 6521047.5 | 4898468 | 4122349.25 | 6573316 |
| O54782 | Man2b2 | Epididymis-specific alpha-mannosidase | 168.71 | 3.44 | 115610 | 3 | 1241992.12 | 4213536 | 2525742 | 3346486.5 | 2765550 | 1951625.38 | 1364346.62 | 3251651 | 2248726.75 | 2187230.5 | 3122501 | 2300120.5 | 3317048.75 | 2897982.25 | 3471957.25 | 4561612.5 | 1548769.25 | 4752370.5 | 4554480.5 | 5206927.5 |
| Q8K2I4 | Manba | Beta-mannosidase | 237.79 | 8.65 | 100831 | 5 | 1753705.88 | 328097.62 | 2289352.5 | 86504.67 | 1642503.38 | 549004.25 | 172688.33 | 123964.67 | 1023346.5 | 756806.75 | 717573.25 | 260468.25 | 766597.62 | 730029.69 | 399209.88 | 719809.75 | 282660.56 | 293462.5 | 129826.73 | 329396.38 |
| P0C5H9 | Manf | Mesencephalic astrocyte-derived neurotrophic fac | 160.12 | 11.17 | 20388 | 3 | 272601.34 | 77017.85 | 708973.62 | 395196.38 | 63035.01 | 2 | 309889.09 | 2 | 2 | 291405.91 | 2 | 2 | 79823.38 | 196906.75 | 1517127.62 | 2 | 128373.77 | 375615.72 | 2 | 366257.94 |
| Q62625 | Map1lc3b | Microtubule-associated protein 1 light chain 3 bet | 33.77 | 4.93 | 16394 | 1 | 346488.78 | 2 | 585293.25 | 2 | 700468.75 | 2 | 2 | 2 | 2 | 391505.75 | 674483.06 | 580965.62 | 450225.16 | 551394.06 | 648925.69 | 2 | 789843.38 | 2 | 2 | 2 |
| Q8C052 | Map1s | Microtubule-associated protein 1S | 117.49 | 3.29 | 102939 | 3 | 2 | 137191.28 | 2 | 551549.56 | 917006.56 | 327711.78 | 1568519.62 | 185005.69 | 1671646.25 | 979497.44 | 154233.16 | 996590 | 697900.94 | 749306.88 | 2 | 2 | 2 | 2 | 2 | 2 |
| P31938 | Map2k1 | Dual specificity mitogen-activated protein kinase I | 109.5 | 6.62 | 43474 | 3 | 867555.69 | 341412.44 | 1072727.38 | 1263354.12 | 805438.81 | 755206.31 | 1479274.75 | 1442884.25 | 505284.97 | 1917824.75 | 179066.56 | 2 | 845546.5 | 575537.81 | 1609287.88 | 392284.94 | 818262.44 | 1453279.75 | 2 | 1627193.88 |
| P27546 | Map4 | Microtubule-associated protein 4 | 350.39 | 8 | 117429 | 6 | 947786.44 | 1419360.88 | 1718975.5 | 1831214.75 | 2042578.38 | 1475022.12 | 892676.31 | 1649767.25 | 4878722.5 | 2146993.5 | 6506036 | 6546057.5 | 3477113.5 | 4350250 | 2579878.25 | 2856013.25 | 3470272.5 | 1964331.75 | 3886989 | 2307908.75 |
| Q8BPM2 | Map4k5 | Mitogen-activated protein kinase kinase kinase ki | 39 | 0.83 | 95045 | 1 | 2 | 2 | 2 | 273518.59 | 948144.12 | 285334.38 | 217008.53 | 615201.81 | 533325.62 | 485667.25 | 1050381.38 | 957062.44 | 657652.81 | 649316.25 | 2 | 1260629.25 | 1007046.19 | 2 | 715167.12 | 2 |
| A2AJI0 | Map7d1 | MAP7 domain-containing protein 1 | 149.9 | 5.79 | 93276 | 4 | 1001440 | 3652218.5 | 622046.12 | 3507412.5 | 2068320 | 2756408.25 | 1894549.38 | 2710558.75 | 2586797.75 | 914188.25 | 3841039.5 | 4227787.5 | 2334640.25 | 3371614.25 | 2378982.75 | 3605738.5 | 2536826.5 | 2230674.25 | 2617280.75 | 3083633.5 |
| P63085 | Mapk1 | Mitogen-activated protein kinase 1 | 88.52 | 5.87 | 41276 | 2 | 1959082.75 | 6205382 | 4095243.25 | 3247900.5 | 4831905 | 6340620 | 5107902.5 | 6437119.5 | 6412637.5 | 7510327 | 10957967 | 10454359 | 6886947 | 8918702 | 5337382.5 | 9006197 | 5355044 | 6571626 | 9020926 | 6046449 |
| P47811 | Mapk14 | Mitogen-activated protein kinase 14 | 76.26 | 5.56 | 41287 | 1 | 2 | 2 | 190717.5 | 66739.32 | 81493.58 | 2 | 2 | 45602.15 | 2 | 191066.14 | 41100.46 | 61095.31 | 141496.08 | 200757.2 | 769617 | 2 | 2 | 91830.11 | 31346.47 | 68561.93 |
| P21708 | Mapk3 | Mitogen-activated protein kinase 3 | 258.01 | 18.16 | 43081 | 6 | 7988203 | 12212599 | 4484275 | 14996888 | 12413246 | 11647099 | 12385705 | 10784908 | 13102780 | 11458022 | 11937735 | 10614496 | 9015493 | 8826150 | 6003479 | 8759267 | 3753540.5 | 7273444.5 | 7688330.5 | 8138258 |
| Q61166 | Mapre1 | Microtubule-associated protein RP/EB family mer | 269.41 | 23.88 | 30016 | 5 | 6272556 | 15935866 | 5912440 | 8280878 | 13695462 | 10312074 | 10533020 | 11421521 | 9838128 | 7224735 | 8860180 | 10310043 | 6693497.5 | 7435832 | 3378132.75 | 7717715.5 | 4566370 | 3492483.5 | 4774758.5 | 2585413.25 |
| P26645 | Marcks | Myristoylated alanine-rich C-kinase substrate | 129.21 | 13.27 | 29661 | 3 | 1038699.94 | 2352934.5 | 662563.19 | 1498296 | 1080071.5 | 3210994.5 | 3774237.5 | 1435101.62 | 4622386.5 | 670658.69 | 2404413.75 | 3860567.5 | 3453738.5 | 3277946.25 | 2209690 | 2727933 | 4614228 | 2069344.88 | 3947676 | 2844894.5 |
| P28667 | Marcksl1 | MARCKS-related protein | 208.81 | 18 | 20165 | 4 | 1872308.62 | 5932740 | 1802210.88 | 3099292.5 | 2720566 | 2336681.75 | 1722704.75 | 2613212.5 | 8668759 | 1026667 | 1945906.38 | 5561216.5 | 1542841.62 | 4375545 | 1900030.12 | 3641049.75 | 3221170.5 | 1779688.5 | 2898264.25 | 5558318 |
| Q68FL6 | Mars1 | Methionine--tRNA ligase, cytoplasmic | 123.43 | 3.1 | 101431 | 3 | 9452807 | 10905404 | 4055911.75 | 10232485 | 6852026 | 9104686 | 4040471.5 | 9051929 | 5143307 | 4654351 | 6581833 | 5453263 | 5808749 | 6894030 | 3519835.25 | 4276063 | 3781241 | 5141325.5 | 6849196.5 | 4137332.25 |
| Q5U2R0 | Mat2b | Methionine adenosyltransferase 2 subunit beta | 108.88 | 7.49 | 37375 | 2 | 2 | 1716056.75 | 803014.5 | 2 | 3389206.5 | 1811199.75 | 2099367.75 | 3358115.25 | 2641574.25 | 2314513.75 | 2953987.75 | 2736519.75 | 1984693.88 | 2177247.25 | 2 | 992503.81 | 613808.44 | 2 | 2 | 2 |
| Q8K310 | Matr3 | Matrin-3 | 426.48 | 14.3 | 94630 | 8 | 917155.94 | 129395.91 | 531004.94 | 512322.78 | 1152094.5 | 2 | 157048.72 | 490660.72 | 2254393.5 | 1327784.12 | 1211262.75 | 1810742.88 | 935437.31 | 1177049.88 | 995220.5 | 2422343.5 | 1566792.25 | 717496.69 | 1434027.62 | 592373.88 |
| Q8VCF0 | Mavs | Mitochondrial antiviral-signaling protein | 188.87 | 11.53 | 53399 | 3 | 2 | 2 | 90639.56 | 2 | 58309.57 | 2 | 2 | 2 | 1203923.12 | 1933598 | 1172519.12 | 1146551 | 48353.05 | 802069.25 | 2 | 753591.44 | 2 | 2 | 2 | 2 |
| Q9Z2E1 | Mbd2 | Methyl-CpG-binding domain protein 2 | 75.03 | 6.76 | 43501 | 2 | 915605 | 1587771 | 525811.5 | 1760914.12 | 1739429.62 | 2084051.62 | 1181729.88 | 2747380 | 1638055 | 1239668.25 | 1443873.12 | 1103179.88 | 1279957.88 | 1263806.38 | 1543722.75 | 966848.5 | 1842699.38 | 2032666 | 1157218.38 | 1872437.62 |
| Q9JKP5 | Mbnl1 | Muscleblind-like protein 1 | 108.95 | 12.32 | 36976 | 2 | 2 | 2 | 2 | 1655456.62 | 1685138.88 | 2 | 469800.03 | 1005393.06 | 2 | 860270.31 | 3133360 | 2736094.25 | 2 | 1743546.12 | 2658506 | 2214198.25 | 2345656 | 2 | 2273211 | 1929497.25 |
| Q8CHK3 | Mboat7 | Lysophospholipid acyltransferase 7 | 42.76 | 2.54 | 53436 | 1 | 41252.16 | 68847.17 | 344942.34 | 266597.03 | 79614.24 | 587570.75 | 870020.81 | 107007.1 | 845666.06 | 289933.19 | 46003.07 | 60196 | 51059.97 | 337258.12 | 696364.31 | 2 | 544802.25 | 102483.52 | 2 | 2 |
| Q3ULD5 | Mccc2 | Methylcrotonoyl-CoA carboxylase beta chain, mit | 199.95 | 9.59 | 61379 | 4 | 2 | 2 | 471314.84 | 2 | 763593.31 | 1439555.25 | 592024.38 | 595665.69 | 2 | 259584.7 | 2 | 1126595.88 | 2 | 728894.75 | 3444664.5 | 2 | 120155.01 | 2 | 2 | 2 |
| P97310 | Mcm2 | DNA replication licensing factor MCM2 | 466.75 | 12.83 | 102078 | 9 | 5741649 | 7951722.5 | 6668155 | 7418037 | 17108018 | 953887.75 | 1890378 | 4415230 | 5000670.5 | 5865013 | 8469970 | 8531764 | 6544230 | 7781109 | 2907047 | 3600080.75 | 3029859.5 | 2791744.25 | 4273139.5 | 2 |
| P25206 | Mcm3 | DNA replication licensing factor MCM3 | 766.74 | 24.51 | 91546 | 15 | 7432625.5 | 5386140.5 | 6225632 | 8960733 | 12890711 | 1398071.75 | 2233366.25 | 4530676 | 6995796.5 | 4720434.5 | 6207410 | 8296383 | 10348992 | 7304144 | 3754682 | 9775681 | 3872019.5 | 3008544 | 3558319.25 | 2308282 |
| P49717 | Mcm4 | DNA replication licensing factor MCM4 | 485.28 | 14.15 | 96736 | 10 | 3512900.25 | 2208974 | 4835831.5 | 7280042 | 8152376.5 | 2129259.5 | 42740.65 | 2061931.38 | 2843059.75 | 3702196 | 3411350 | 2382884 | 2868465.5 | 4836575 | 2269823.75 | 2112699.5 | 2032690 | 2849955.5 | 280835.56 | 2920040.5 |
| P49718 | Mcm5 | DNA replication licensing factor MCM5 | 454.31 | 16.21 | 82407 | 11 | 14334609 | 23994526 | 12389125 | 23668398 | 18101840 | 5167847 | 2130634 | 10794580 | 8285995.5 | 7981872 | 8040220 | 7604583 | 8059879 | 10139252 | 6163571.5 | 6093196 | 5124019 | 2573817.75 | 7862198.5 | 9596300 |
| P97311 | Mcm6 | DNA replication licensing factor MCM6 | 709.91 | 19.24 | 92867 | 16 | 7016374 | 4972939.5 | 4528599 | 8230403 | 8945009 | 2668162.25 | 2204913.5 | 5255236 | 3732715.5 | 3387898.25 | 5954172.5 | 8142834.5 | 6939933 | 6337963.5 | 7721257 | 6487940.5 | 5803005 | 3423902 | 7762567 | 5478217 |
| Q61881 | Mcm7 | DNA replication licensing factor MCM7 | 554.39 | 18.5 | 81211 | 11 | 10809012 | 8981728 | 7440702.5 | 17969792 | 20951174 | 4286497.5 | 6001006 | 7453437 | 7134752 | 6537113 | 4556683 | 8747308 | 6959521 | 8936952 | 5406111 | 3390765 | 5366053.5 | 5655188 | 3016416 | 6699942 |
| B1H268 | Mcmbp | Mini-chromosome maintenance complex-binding | 61.43 | 1.71 | 73007 | 1 | 705141.25 | 1091027.75 | 107445.52 | 1137816.88 | 153762.58 | 766078.06 | 2 | 2 | 791844.25 | 893418.19 | 1029017.62 | 1130249.25 | 920719.44 | 965537.69 | 870311.69 | 1234780.88 | 1166791.12 | 979340.56 | 1169090.62 | 1151531.5 |
| Q4G009 | Mcts1 | Malignant T-cell-amplified sequence 1 | 45.11 | 4.95 | 20550 | 1 | 2 | 2 | 2 | 704045.31 | 1103538.75 | 1194346.5 | 1039913 | 1732684 | 1017505 | 773861.25 | 2014772.88 | 2100043.75 | 1280917.88 | 1501032.25 | 2 | 1153179.38 | 2 | 2 | 1294531.38 | 998120.75 |
| Q3UMR5 | Mcu | Calcium uniporter protein, mitochondrial | 58.92 | 2.86 | 39682 | 1 | 1130043.5 | 2 | 1072148.38 | 1204563.12 | 1334077.5 | 1261093.38 | 945305.62 | 1170887.25 | 2 | 1430092.12 | 442773.72 | 1117225.25 | 1327553.62 | 1459718.38 | 657223 | 1951256.75 | 1848635.75 | 1809116.75 | 1818235 | 1398138.75 |
| O88989 | Mdh1 | Malate dehydrogenase, cytoplasmic | 154.24 | 11.08 | 36483 | 3 | 3530576 | 713517.06 | 4389021.5 | 1880546.62 | 2019421.38 | 583458.75 | 1761720.25 | 564740.06 | 1678117 | 3438629 | 2760090 | 4859991 | 2086655.5 | 2170330.25 | 2392792.25 | 1388667.25 | 3123974.5 | 739411.25 | 1071418.25 | 1098312.62 |
| P08249 | Mdh2 | Malate dehydrogenase, mitochondrial | 1716.16 | 71.3 | 35611 | 26 | 264461570 | 311500418 | 294468802 | 133238802 | 162893506 | 173350834 | 174263090 | 119196994 | 178392162 | 238523922 | 195409314 | 220169714 | 204009730 | 181295010 | 175644626 | 172871426 | 236147058 | 220574242 | 237631170 | 250791458 |
| Q9D967 | Mdp1 | Magnesium-dependent phosphatase 1 | 48.06 | 7.93 | 18582 | 1 | 2 | 1221572 | 2 | 606455.75 | 1545051.75 | 1037199 | 999341.5 | 1311244.5 | 908544.19 | 843024 | 1297094 | 967053.19 | 2 | 883843 | 2 | 947712.31 | 777299.94 | 691733.94 | 1216719.5 | 2 |
| P06801 | Me1 | NADP-dependent malic enzyme | 42.61 | 3.15 | 63954 | 1 | 2 | 2 | 2 | 2 | 2 | 2 | 170552.17 | 2 | 554349.38 | 359083.41 | 2 | 2 | 2 | 2 | 2 | 2 | 2 | 2 | 2 | 2 |
| Q99KE1 | Me2 | NAD-dependent malic enzyme, mitochondrial | 315.61 | 13.41 | 65799 | 6 | 1294914.25 | 1275164.25 | 1395914 | 1339131 | 842166.38 | 1835338.25 | 2003958 | 2 | 2065349.5 | 2827799.75 | 1706155.5 | 1475398.5 | 1187609.88 | 113823.76 | 2135840 | 1626715.5 | 1309731.12 | 1953429 | 1381298.75 | 1478684.88 |
| Q9DCS3 | Mecr | Enoyl-[acyl-carrier-protein] reductase, mitochondr | 99.31 | 4.83 | 40343 | 2 | 2164193.25 | 3852965 | 1756987.38 | 591191.81 | 1942772.25 | 2000397.12 | 1865646 | 2123958.5 | 1677725.5 | 1372588.25 | 2000102 | 828592.62 | 558581.25 | 1126513.25 | 2 | 2016318.25 | 1112403.88 | 1567018.5 | 1562085.38 | 1745532 |
| Q8VCD5 | Med17 | Mediator of RNA polymerase II transcription subu | 37.04 | 1.08 | 72462 | 1 | 2 | 2 | 2 | 2 | 2 | 2 | 2 | 2 | 365680.28 | 2 | 2 | 2 | 2 | 2 | 2 | 2 | 2 | 2 | 2 | 2 |
| Q9ERE7 | Mesd | LRP chaperone MESD | 69.76 | 7.59 | 25207 | 1 | 392285.72 | 143627.83 | 102569.26 | 189929.84 | 69701.65 | 2 | 103089.47 | 2 | 229827.56 | 98878 | 68120.53 | 2 | 182258.27 | 175395.84 | 218755.98 | 2 | 227301.08 | 140503.45 | 2 | 2 |
| Q8BP48 | Metap1 | Methionine aminopeptidase 1 | 45.71 | 2.85 | 43221 | 1 | 1687266.75 | 1837850.25 | 1723755 | 1279430.5 | 1427339 | 1951496.75 | 1447192.5 | 1987489.38 | 965279.31 | 1050735 | 2015526.25 | 1447223.25 | 1776399.12 | 1686912.12 | 2824133.5 | 2 | 2 | 1178848.5 | 496309.91 | 1715664 |
| Q91YR5 | Mettl13 | eEF1A lysine and N-terminal methyltransferase | 44.12 | 1.72 | 78757 | 1 | 2 | 2 | 2 | 2 | 2 | 2 | 109289.67 | 2 | 42077.92 | 2 | 2 | 2 | 89783.9 | 143498.81 | 2 | 2 | 2 | 2 | 2 | 2 |
| Q9DCS2 | Mettl26 | Methyltransferase-like 26 | 84.54 | 9.8 | 22687 | 2 | 2 | 515798.03 | 1218049.62 | 953995.69 | 309110.12 | 996580.62 | 2 | 2 | 2 | 2 | 1855916.62 | 2057016.25 | 1703877.25 | 1499359.75 | 860893.19 | 741331.5 | 702133.69 | 806424.62 | 885955.19 | 2 |
| Q8C3P7 | Mettl3 | N(6)-adenosine-methyltransferase catalytic subur | 42.03 | 1.38 | 64616 | 1 | 893294.69 | 682119.19 | 2 | 1065219.62 | 2 | 2 | 2 | 2 | 2 | 2 | 2 | 595258.88 | 2 | 2 | 2 | 464562.66 | 2 | 1709955.88 | 2 | 2 |
| P21956 | Mfge8 | Lactadherin | 259.54 | 14.9 | 51241 | 5 | 8325948 | 8427169 | 11303059 | 10851336 | 9635244 | 5658167 | 5604066 | 14138725 | 7612923 | 9277971 | 8553494 | 6492346.5 | 8446707 | 6773229 | 2761362.75 | 5638457 | 8270068.5 | 8994106 | 7931550 | 7251746.5 |
| Q9D2V8 | Mfsd10 | Major facilitator superfamily domain-containing pr | 98.14 | 7.46 | 49369 | 2 | 316962.59 | 70132.71 | 2 | 686460.25 | 13778.24 | 467313.97 | 923879.56 | 93966.58 | 765547.31 | 2 | 42399.85 | 2 | 431218.09 | 2 | 2 | 2 | 2 | 540544.06 | 137218.06 | 113461.99 |
| Q9CPU4 | Mgst3 | Glutathione S-transferase 3, mitochondrial | 145.65 | 18.3 | 16958 | 2 | 2 | 530237.19 | 931216.88 | 937457.56 | 1297510.88 | 2 | 1097217 | 118102.36 | 824737.38 | 1002687.75 | 2 | 2 | 232483.95 | 375737.91 | 1088754.88 | 2 | 337245.06 | 327325.81 | 2 | 2 |
| Q8BI84 | Mia3 | Transport and Golgi organization protein 1 homol | 82.41 | 1.14 | 213675 | 2 | 696073.12 | 3275330.5 | 2 | 447391.34 | 441869.88 | 3259741.75 | 2055840.75 | 970834.69 | 2130853 | 2360829 | 5183737 | 5518998 | 4357785.5 | 3536025.75 | 393388.31 | 8258376.5 | 6753916 | 3244943.5 | 7403410 | 6951949 |
| D3ZBP4 | Mical1 | [F-actin]-monooxygenase MICAL1 | 118.73 | 2.48 | 116672 | 2 | 693870.88 | 2 | 2 | 791337.44 | 583115.94 | 2 | 2 | 1606635.25 | 495899.56 | 2 | 988599.88 | 1308112.12 | 1207612.12 | 308454.53 | 794121.88 | 815475.75 | 1108760.88 | 1688028.5 | 2 | 862450.38 |
| Q7TNS2 | Micos10 | MICOS complex subunit Mic10 | 39.21 | 9.21 | 8567 | 1 | 3443558.5 | 2122458.75 | 1942349.25 | 2219467.5 | 1415675.75 | 2 | 1602155.38 | 2 | 1478580.75 | 1664193 | 1379795.62 | 1465682 | 1327136.25 | 1273000.5 | 1483594.62 | 1959406 | 2181642 | 2766175.5 | 2067301.38 | 2642807.5 |
| P34884 | Mif | Macrophage migration inhibitory factor | 200.93 | 77.39 | 12504 | 5 | 3547754.75 | 5329339 | 6393131 | 3578547 | 9443750 | 5390170 | 4816335.5 | 8555480 | 4811239.5 | 8234505 | 6806805.5 | 6114621 | 8216556 | 5691512 | 4206916 | 5356232 | 4620034.5 | 2580844.75 | 5962440 | 611322.94 |
| Q5BJQ2 | Mindy1 | Ubiquitin carboxyl-terminal hydrolase MINDY-1 | 67.03 | 2.07 | 52729 | 1 | 3515773 | 2607064.25 | 2788941.5 | 1216116.25 | 1602055 | 3101769 | 2060990.75 | 3206017.5 | 1557468.75 | 1185543 | 1762381.12 | 1984166.75 | 2670223.5 | 1936167.62 | 1443431.38 | 2592722.25 | 2441077.5 | 3364664.75 | 2623400.25 | 3237281.5 |
| E9PVX6 | Mki67 | Proliferation marker protein Ki-67 | 1637.19 | 15.17 | 350864 | 34 | 5183721.5 | 3834775.25 | 10173954 | 6311470.5 | 7547546.5 | 3056281.25 | 4075171.25 | 5662478 | 3956328.5 | 7208517 | 10365986 | 12512936 | 9127081 | 9239927 | 8371798 | 8986108 | 9969296 | 10946419 | 13097530 | 6585433 |
| Q5FVQ4 | Mlec | Malectin | 167.34 | 12.71 | 32418 | 3 | 1167799.38 | 1034150.56 | 1788965 | 1418664 | 2823636.5 | 2 | 960279.38 | 3345538.25 | 2493523.75 | 1633892.25 | 2431380 | 1856269.38 | 1653271.5 | 2506933 | 602433.31 | 2141679 | 1305928.38 | 1187897.38 | 2282010 | 947350.5 |
| Q9D2Y4 | Mlkl | Mixed lineage kinase domain-like protein | 136.02 | 5.3 | 54317 | 2 | 544209.94 | 762482.5 | 2 | 1179105.12 | 1283127.62 | 3330024.5 | 2616151.5 | 4385413 | 2529525.25 | 1547834.62 | 1221413 | 1961016.12 | 1038784.75 | 1183699.62 | 1026932.38 | 2709914 | 900389.38 | 1170276.62 | 1957982.88 | 2 |
| P34960 | Mmp12 | Macrophage metalloelastase | 227.05 | 11.21 | 54971 | 5 | 1747290.25 | 1524405.88 | 3815198.75 | 2520478.25 | 2615260 | 5440064 | 4462009 | 7020914.5 | 3938979.25 | 5132736.5 | 7922356 | 5463830 | 8076679 | 8823701 | 10555502 | 8275962 | 3671428 | 11374950 | 9379204 | 6645517 |
| Q9D071 | Mms19 | MMS19 nucleotide excision repair protein homolo | 77.08 | 2.04 | 113089 | 2 | 2 | 2 | 686031.44 | 1234332.12 | 1100751.38 | 2 | 1366915.25 | 797902.44 | 678989.94 | 476941.12 | 2 | 856144.25 | 460809.97 | 300826 | 2 | 2 | 2 | 2 | 2 | 2 |
| P16332 | Mmut | Methylmalonyl-CoA mutase, mitochondrial | 214.38 | 8.42 | 82844 | 4 | 2548759.5 | 296886.38 | 726124 | 634725.88 | 1619609 | 346246.09 | 860963.56 | 1211661.62 | 1635524.38 | 1187291.38 | 1252181.62 | 893375.19 | 1344966.25 | 1493271.5 | 607778.31 | 1517951 | 1598270.62 | 689067.88 | 1720376.75 | 1057402.5 |
| D0QMC3 | Mndal | Myeloid cell nuclear differentiation antigen-like pr | 61.62 | 2.04 | 60526 | 1 | 2 | 645115.06 | 2 | 522515.84 | 1437640.38 | 690047.25 | 1289565.5 | 1618742 | 1035012.5 | 971116.44 | 2 | 2 | 434693.41 | 2 | 2 | 2 | 2 | 2 | 2 | 2 |
| Q3T1J9 | Mob1a | MOB kinase activator 1A | 68.83 | 8.33 | 25080 | 2 | 2750918.5 | 2277207.25 | 2351624.5 | 3883126.25 | 3112418.25 | 1804570.12 | 2368626 | 2132304 | 2017278.12 | 2473419.5 | 1210335.38 | 1454878.12 | 1510869.38 | 1467715 | 2223966 | 1416230.12 | 2307772.5 | 2924570.5 | 1929644.5 | 2895978.75 |
| Q6PEB6 | Mob4 | MOB-like protein phocein | 54.45 | 4.44 | 26032 | 1 | 810107.06 | 1470780.62 | 2 | 1133186.38 | 2 | 886716.44 | 1335083.5 | 2 | 1194137.38 | 2 | 2 | 2 | 903200.5 | 925972.75 | 2 | 698287.44 | 1153600.12 | 2 | 358966.97 | 2 |
| Q80UM7 | Mogs | Mannosyl-oligosaccharide glucosidase | 446.04 | 11.51 | 91831 | 8 | 2 | 2 | 2 | 2 | 1228963.88 | 203476.09 | 2 | 2 | 158971.36 | 2 | 2 | 128764.97 | 136777.91 | 120672.04 | 232455.92 | 2 | 2 | 244070.28 | 2 | 2 |

| | | | | | | | | | | | | | | | | | | | | | | | | | | |
|---|---|---|---|---|---|---|---|---|---|---|---|---|---|---|---|---|---|---|---|---|---|---|---|---|---|---|
| Q80TL7 | Mon2 | Protein MON2 homolog | 36.5 | 0.41 | 189078 | 1 | 390078.88 | 1069978.12 | 457646.88 | 707330.94 | 539219.62 | 963842.75 | 539615.56 | 922933.81 | 278393.97 | 2 | 330830 | 2 | 2 | 2 | 2 | 2 | 2 | 2 | 674696.94 | 2 |
| P23249 | Mov10 | Putative helicase MOV-10 | 75.68 | 1.79 | 113583 | 2 | 1101826.12 | 911823.44 | 1500767.38 | 1024390.88 | 1307391.5 | 96269.71 | 195562.91 | 1200971.75 | 883958.69 | 1172227.25 | 2 | 2 | 836688.88 | 969424 | 197143.97 | 626249.38 | 1246640.12 | 2 | 733190.69 | 1294621.12 |
| P63030 | Mpc1 | Mitochondrial pyruvate carrier 1 | 44.56 | 8.26 | 12455 | 1 | 1665262.12 | 636604.94 | 336553.59 | 1148037.62 | 103680.05 | 915608.56 | 896569.44 | 2 | 964987.19 | 1454985.88 | 857705.12 | 2 | 1083910.62 | 1013700 | 1265170 | 1480499.38 | 1076869.38 | 1727496.12 | 1549502.62 | 1564760.62 |
| P38718 | Mpc2 | Mitochondrial pyruvate carrier 2 | 220.02 | 37.01 | 14258 | 4 | 3153350.5 | 2311629.25 | 2101577 | 4641093 | 1375428.25 | 1760125.12 | 1926766.62 | 1758354.25 | 1735232.25 | 1374559.62 | 898991.81 | 304681.75 | 1246366.25 | 502657.31 | 2824356 | 1290914.75 | 2276919.25 | 1987371.38 | 380316.88 | 1559666.25 |
| A1L314 | Mpeg1 | Macrophage-expressed gene 1 protein | 665.85 | 19.35 | 78390 | 12 | 4092371.75 | 7790929 | 3064948.5 | 6733613.5 | 6004197.5 | 6303170 | 6677600.5 | 8248892 | 14188936 | 10183316 | 2868876 | 3017573 | 2500374.5 | 2445594.25 | 1998521.88 | 3412769.25 | 3295437 | 2346290.25 | 3529577.25 | 3842626 |
| P70290 | Mpp1 | 55 kDa erythrocyte membrane protein | 105.19 | 5.36 | 52227 | 3 | 2 | 2 | 2 | 2 | 410428.38 | 2 | 2 | 2 | 2 | 510128.09 | 2 | 99687.36 | 2 | 145380.58 | 2 | 2 | 127962.19 | 2 | 2 | 2 |
| Q99J99 | Mpst | 3-mercaptopyruvate sulfurtransferase | 210.24 | 8.42 | 33097 | 3 | 640321.12 | 1086336 | 2 | 2 | 438552.09 | 161688.08 | 154747.05 | 129216.96 | 1296097.38 | 1546859 | 2398695.25 | 2829877.75 | 2307712.25 | 1839291.75 | 473078.47 | 1950758.75 | 297180.53 | 2 | 243593.41 | 830080.38 |
| Q9CQT1 | Mri1 | Methylthioribose-1-phosphate isomerase | 315.46 | 17.07 | 39411 | 6 | 1047406.75 | 1068128 | 2008525.25 | 2113934 | 1061798.25 | 2 | 1180488.25 | 1706312.75 | 1329566.88 | 1134714.75 | 1007050.12 | 1472752.25 | 1664887.75 | 580751.19 | 2556196 | 877921.31 | 1074951 | 2792768.5 | 2 | 2077693 |
| Q99N96 | Mrpl1 | Large ribosomal subunit protein uL1m | 92.87 | 6.55 | 37597 | 2 | 617617 | 2 | 2 | 2 | 896280 | 2 | 2 | 672135.06 | 2 | 424669.88 | 983106.25 | 525942.69 | 2 | 514818.06 | 2 | 2 | 610533.62 | 2 | 2 | 2 |
| Q3TBW2 | Mrpl10 | Large ribosomal subunit protein uL10m | 50.94 | 4.2 | 29396 | 1 | 2 | 1169113.88 | 799396.31 | 2 | 869144.12 | 2 | 830591 | 2 | 573764.19 | 2 | 2 | 2 | 685596.44 | 384868.16 | 2 | 686256.81 | 2 | 2 | 2 | 2 |
| Q9CQF0 | Mrpl11 | Large ribosomal subunit protein uL11m | 41.92 | 9.38 | 20680 | 1 | 1462743.5 | 1465318.38 | 856885.69 | 512007.75 | 958565 | 2 | 597271.38 | 1387564.88 | 777314.12 | 870212.88 | 697581.88 | 985976.06 | 669662.56 | 746020.56 | 654610 | 2 | 678664.94 | 2 | 2 | 2 |
| Q9DB15 | Mrpl12 | Large ribosomal subunit protein bL12m | 40.73 | 6.97 | 21708 | 1 | 374326.25 | 2 | 369464.09 | 2 | 2 | 2 | 2 | 2 | 2 | 2 | 2 | 2 | 2 | 2 | 2 | 2 | 2 | 2 | 2 | 2 |
| Q7M0E7 | Mrpl14 | Large ribosomal subunit protein uL14m | 58.86 | 6.9 | 15913 | 1 | 2 | 2 | 2 | 102811.54 | 2 | 2 | 2 | 2 | 2 | 2 | 2 | 2 | 2 | 2 | 2 | 2 | 2 | 2 | 2 | 2 |
| Q9CPR5 | Mrpl15 | Large ribosomal subunit protein uL15m | 180.13 | 23.05 | 33542 | 5 | 1027635.94 | 1525119.62 | 716517.38 | 707966.25 | 856651.56 | 587277.06 | 605917.88 | 1541466.62 | 1665088.62 | 470134.16 | 2 | 2 | 63511.32 | 507334.44 | 2 | 2 | 2 | 343751.91 | 2 | 2 |
| Q5M818 | Mrpl16 | Large ribosomal subunit protein uL16m | 39.66 | 3.98 | 28895 | 1 | 2 | 330570.88 | 2 | 315317.38 | 673843.5 | 371523.81 | 238626.05 | 727351.25 | 496809.09 | 443995.5 | 517678.75 | 374803.41 | 631024.06 | 688799.38 | 2 | 519185.09 | 522021.06 | 474479.72 | 494181.59 | 2 |
| Q9D338 | Mrpl19 | Large ribosomal subunit protein bL19m | 176.21 | 12.67 | 33578 | 4 | 575289.62 | 1003186.94 | 502012.88 | 769077.5 | 832785.31 | 139775.58 | 2 | 2 | 227855.81 | 277760.75 | 2 | 2 | 100854.18 | 292543.81 | 2 | 640350.12 | 2 | 763403.31 | 702465.81 | 723937.06 |
| Q9D1N9 | Mrpl21 | Large ribosomal subunit protein bL21m | 139.29 | 17.7 | 23366 | 3 | 2 | 2 | 2 | 2 | 33420.17 | 2 | 2 | 2 | 123480.54 | 72996.34 | 2 | 103763.27 | 97874.83 | 173428.42 | 2 | 2 | 99401.3 | 2 | 2 | 2 |
| P0C2C0 | Mrpl22 | Large ribosomal subunit protein uL22m | 39.59 | 5.83 | 24039 | 1 | 2 | 2 | 2 | 2 | 2 | 2 | 2 | 2 | 2 | 2 | 2 | 2 | 2 | 2 | 2 | 2 | 2 | 2 | 2 | 2 |
| O35972 | Mrpl23 | Large ribosomal subunit protein uL23m | 58.65 | 13.01 | 17122 | 1 | 2 | 145619.5 | 2 | 124489.6 | 118540.95 | 195222.88 | 168373.88 | 2 | 247590.94 | 2 | 2 | 2 | 87422.24 | 145393.91 | 2 | 2 | 2 | 2 | 2 | 2 |
| Q99N92 | Mrpl27 | Large ribosomal subunit protein bL27m | 65.02 | 8.78 | 15945 | 1 | 2 | 2 | 2 | 2 | 840060.12 | 2 | 2 | 972234.62 | 2 | 2 | 2 | 2 | 2 | 2 | 2 | 2 | 2 | 2 | 2 | 2 |
| Q5PQN9 | Mrpl38 | Large ribosomal subunit protein mL38 | 45.33 | 2.63 | 44838 | 1 | 2 | 1111864.75 | 943492.88 | 453916.56 | 857129.12 | 2 | 2 | 2 | 630855.31 | 802992 | 1304391.12 | 970550.44 | 1245849.88 | 1205087 | 571852 | 2248741 | 1442691.75 | 1227250.38 | 1439595.38 | 1128540.25 |
| Q9JKF7 | Mrpl39 | Large ribosomal subunit protein mL39 | 69.91 | 5.36 | 38549 | 2 | 2 | 2 | 2 | 2 | 553128.31 | 2 | 453976.06 | 2 | 627750.31 | 231545.14 | 2 | 458751.41 | 351654.94 | 2 | 2 | 2 | 2 | 2 | 2 | 737706.25 |
| Q9DCU6 | Mrpl4 | Large ribosomal subunit protein uL4m | 41.83 | 3.74 | 33073 | 1 | 991142 | 205933.67 | 2 | 345830.09 | 2 | 2 | 2 | 2 | 2 | 2 | 2 | 2 | 2 | 2 | 2 | 2 | 2 | 2 | 2 | 2 |
| Q99N89 | Mrpl43 | Large ribosomal subunit protein mL43 | 43.03 | 3.83 | 20203 | 1 | 2 | 1688560.5 | 1011277.5 | 2 | 980117.19 | 1072420.5 | 1043970.44 | 1473773.38 | 1088763 | 1068710.88 | 2 | 2 | 658654.5 | 1020916.38 | 2 | 2 | 336871.62 | 2 | 577565.38 | 2 |
| Q9CY73 | Mrpl44 | Large ribosomal subunit protein mL44 | 120.75 | 8.71 | 37527 | 2 | 2 | 2 | 2 | 95626.23 | 2 | 2 | 2 | 2 | 203459.03 | 472933.84 | 2 | 2 | 2 | 86417.7 | 275655.97 | 2 | 2 | 2 | 2 | 2 |
| Q8R035 | Mrpl58 | Large ribosomal subunit protein mL62 | 51.23 | 5.83 | 23477 | 1 | 2 | 562101.62 | 394595.44 | 2 | 502196.31 | 2 | 806896.81 | 1018419.81 | 762878.12 | 2 | 789567.25 | 2 | 2 | 2 | 427945.41 | 2 | 2 | 2 | 2 | 954744.69 |
| Q9DCA2 | Mrps11 | Small ribosomal subunit protein uS11m | 74.06 | 8.9 | 20208 | 1 | 2 | 2 | 2 | 2 | 909801.75 | 2 | 2 | 2 | 652920.56 | 2 | 2 | 2 | 2 | 2 | 2 | 2 | 2 | 2 | 591446.62 | 2 |
| Q9CPX7 | Mrps16 | Small ribosomal subunit protein bS16m | 37.49 | 5.93 | 15192 | 1 | 1054757.88 | 607917.94 | 732618.56 | 725669.88 | 730625.19 | 7500410 | 7345684 | 960183.12 | 446655.47 | 890777.94 | 4314219.5 | 4876541 | 5346954 | 312793.69 | 597348.44 | 4020537.75 | 693989.19 | 587932.31 | 3746562.5 | 13807060 |
| Q924T2 | Mrps2 | Small ribosomal subunit protein uS2m | 147.05 | 8.59 | 32313 | 3 | 572693.81 | 157886.75 | 736555.44 | 386538 | 485594.34 | 2 | 2 | 403678.19 | 2 | 460902.09 | 2 | 395461.28 | 2 | 520097.56 | 335320.09 | 2 | 399536.28 | 2 | 2 | 2 |
| P58059 | Mrps21 | Small ribosomal subunit protein bS21m | 79.03 | 14.94 | 10561 | 1 | 2 | 2 | 2 | 2 | 270432.47 | 2 | 2 | 2 | 2 | 2 | 2 | 2 | 558826.44 | 2 | 2 | 2 | 2 | 2 | 2 | 2 |
| Q9CXW2 | Mrps22 | Small ribosomal subunit protein mS22 | 37.54 | 2.23 | 41192 | 1 | 2 | 2 | 2 | 2 | 2 | 2 | 2 | 2 | 2 | 2 | 2 | 2 | 2 | 2 | 2 | 2 | 2 | 2 | 2 | 2 |
| Q8VE22 | Mrps23 | Small ribosomal subunit protein mS23 | 79.16 | 11.3 | 20348 | 2 | 2 | 2 | 499731.59 | 191236.45 | 411779.06 | 2 | 2 | 68280.87 | 731638.38 | 598477.12 | 2 | 2 | 359359.59 | 188772.08 | 2 | 2 | 2 | 2 | 2 | 2 |
| Q80ZS3 | Mrps26 | Small ribosomal subunit protein mS26 | 39.3 | 3.5 | 23444 | 1 | 3996210.5 | 3562764.75 | 751150.56 | 4215843.5 | 203659.52 | 3591588.5 | 3481489.5 | 198001.48 | 2681446.25 | 2829682.75 | 3171956.25 | 3766047.5 | 4418395 | 2885196.25 | 5808528.5 | 3457521.75 | 5243349 | 5263895.5 | 4577651.5 | 6143227 |
| Q8BK72 | Mrps27 | Small ribosomal subunit protein mS27 | 76.15 | 5.54 | 47779 | 2 | 2197450.5 | 3423821.75 | 2108926.25 | 1879147 | 1451478.38 | 2 | 163035.25 | 951940.31 | 2 | 2172462.25 | 2 | 1109180.25 | 1561065.38 | 1461012.38 | 431436.41 | 2 | 1772355.88 | 2822044 | 2720152.5 | 2 |
| Q9CY16 | Mrps28 | Small ribosomal subunit protein bS1m | 123.82 | 24.73 | 20520 | 3 | 2 | 2 | 594156.38 | 2 | 592269.12 | 2 | 2 | 92127.45 | 507840 | 2 | 523197.44 | 69907.99 | 2 | 533357.12 | 2 | 2 | 2 | 2 | 2 | 2 |
| Q9D0G0 | Mrps30 | Large ribosomal subunit protein mL65 | 93.61 | 7.47 | 49939 | 2 | 2 | 2 | 2 | 2 | 205344.12 | 2 | 2 | 2 | 2 | 2 | 2 | 2 | 2 | 2 | 2 | 2 | 2 | 2 | 2 | 2 |
| Q9JIK9 | Mrps34 | Small ribosomal subunit protein mS34 | 68.92 | 6.88 | 25827 | 2 | 2 | 58794.62 | 2 | 192236 | 237450.75 | 200955.5 | 143106.11 | 46442.38 | 1017142.69 | 468199.12 | 2 | 86079.62 | 806180.31 | 375347.03 | 131282.91 | 2 | 87899.02 | 2 | 2 | 2 |
| Q8BJZ4 | Mrps35 | Small ribosomal subunit protein mS35 | 34.88 | 4.69 | 35975 | 1 | 2 | 467031.78 | 1443396.62 | 2 | 780257.56 | 196369.05 | 2 | 2 | 2 | 2 | 2 | 294007.69 | 1039766.12 | 110727.24 | 690045.94 | 2 | 2 | 2 | 2 | 335318.06 |
| Q99N87 | Mrps5 | Small ribosomal subunit protein uS5m | 109.68 | 5.09 | 48207 | 2 | 2 | 2 | 2 | 813450.38 | 2 | 2 | 2 | 997411.5 | 2034593 | 2 | 2 | 542644.75 | 2108050 | 2044901.12 | 2 | 6841002.5 | 1727886.88 | 2967177.75 | 5327282 | 2 |
| Q9D0I8 | Mrto4 | mRNA turnover protein 4 homolog | 117.99 | 14.23 | 27546 | 3 | 757628.06 | 1084293.88 | 1382130 | 1354883.62 | 2210032.5 | 810601.06 | 642132.06 | 1956985.38 | 1604708.25 | 1226849.25 | 1279479 | 567848.5 | 1068001.62 | 1481956.5 | 552462.56 | 846133.75 | 1015396.06 | 2 | 84910.86 | 97440.95 |
| P43247 | Msh2 | DNA mismatch repair protein Msh2 | 55.58 | 1.6 | 104151 | 1 | 1027629 | 256094.42 | 800181.19 | 607052.94 | 569577.81 | 183061.23 | 388186.94 | 66306.38 | 2 | 344784.75 | 207721.11 | 2 | 499754.47 | 482945.34 | 560570.19 | 148123.75 | 2 | 2 | 2 | 2 |
| P54276 | Msh6 | DNA mismatch repair protein Msh6 | 34.26 | 0.88 | 151084 | 1 | 2 | 536656.81 | 2 | 755682.94 | 29284.93 | 140571.69 | 365955.03 | 78213.17 | 698561.94 | 2 | 289242.03 | 980860.69 | 200382.52 | 2 | 462654.38 | 951615.62 | 451982.91 | 2 | 94480.84 | 2 |
| P26041 | Msn | Moesin | 2471.44 | 56.67 | 67767 | 41 | 36159634 | 47561142 | 33869290 | 64798106 | 59174970 | 66666686 | 60445490 | 70021162 | 100437402 | 92483298 | 171885314 | 194163906 | 118448938 | 131269890 | 138163378 | 143685234 | 127130450 | 126115922 | 122653026 | 123973450 |
| P30204 | Msr1 | Macrophage scavenger receptor types I and II | 152.05 | 9.17 | 50170 | 3 | 6066337 | 7348997 | 4959982.5 | 6351289 | 5461674 | 6122281.5 | 6893375.5 | 8855820 | 6257679 | 4531502 | 4474491.5 | 3600836.5 | 4323919 | 3884632 | 3759890 | 5172127 | 7432259 | 7291486 | 5659761 | 7746067 |
| P02802 | Mt1 | Metallothionein-1 | 58.2 | 19.67 | 6018 | 1 | 2 | 2 | 2 | 2 | 50731.34 | 2 | 2 | 2 | 615853.06 | 2 | 403821.53 | 2120836.75 | 90694.78 | 1419229.88 | 717447.25 | 2 | 138730.78 | 2 | 2 | 2 |
| Q9R190 | Mta2 | Metastasis-associated protein MTA2 | 452.98 | 16.02 | 75030 | 9 | 381540.62 | 2 | 974644.25 | 760620.44 | 2062637 | 1947519.75 | 847568.38 | 676517.19 | 1717542.88 | 2 | 2054906.62 | 913251.38 | 591792.31 | 1137398.38 | 1362916 | 3890159 | 588072.5 | 1484753.12 | 2913733 | 2 |
| Q924K8 | Mta3 | Metastasis-associated protein MTA3 | 180.62 | 7.61 | 67077 | 4 | 2 | 2 | 2 | 2 | 767301.12 | 2 | 2 | 2 | 2 | 2 | 895333.69 | 1426428.88 | 2 | 782207.94 | 2 | 2088873.38 | 2 | 2 | 2 | 2 |
| Q9CQ65 | Mtap | S-methyl-5'-thioadenosine phosphorylase | 381.16 | 33.22 | 31062 | 7 | 502744.09 | 429661.56 | 2443655.25 | 2040539.75 | 5129736 | 1782007 | 2370494.5 | 2437760.75 | 4396504 | 2630537 | 2953582.5 | 4400160 | 2439918.5 | 2107717 | 2010892.75 | 3277078 | 1899524.25 | 663495 | 1687828.5 | 522125.31 |
| Q922Q1 | Mtarc2 | Mitochondrial amidoxime reducing component 2 | 97.31 | 7.69 | 38194 | 2 | 2 | 2 | 1368279.12 | 2 | 1013685.69 | 2 | 2 | 2 | 2 | 690800 | 2 | 2 | 773364.19 | 2 | 976264.69 | 2 | 618029.31 | 2 | 2 | 622462.56 |
| P03930 | Mtatp8 | ATP synthase protein 8 | 50.61 | 19.4 | 7766 | 1 | 2 | 2 | 2 | 2 | 759249.94 | 768482.31 | 2 | 2 | 781713.06 | 2 | 2 | 588447.94 | 2 | 2 | 2 | 2 | 2 | 2 | 2 | 2 |
| Q791T5 | Mtch1 | Mitochondrial carrier homolog 1 | 233.51 | 10.54 | 41565 | 5 | 389802.81 | 1816145.75 | 584984 | 1368942.62 | 4531230 | 3706207.5 | 3091801.5 | 5969160.5 | 6969079 | 3931558.5 | 3713968.75 | 4547620 | 2995967.25 | 4724933 | 220750.84 | 2961209.5 | 1196300.25 | 2 | 1069942.88 | 2364991.75 |
| Q791V5 | Mtch2 | Mitochondrial carrier homolog 2 | 252 | 24.09 | 33499 | 5 | 4635596 | 8668499 | 5592020.5 | 2016270.5 | 4447500.5 | 7465036.5 | 7869922 | 5315001 | 6667137.5 | 5717997 | 4655684.5 | 3969950.75 | 5603046.5 | 5468796.5 | 2815370 | 3525576.5 | 3583461.5 | 2579359.25 | 3087931.5 | 1468473.5 |
| P00405 | Mtco2 | Cytochrome c oxidase subunit 2 | 373.37 | 26.43 | 25976 | 7 | 15704052 | 13871906 | 16031726 | 14906254 | 15910260 | 21134662 | 19275434 | 12499246 | 33932514 | 29490410 | 20138232 | 14153782 | 19011422 | 16089322 | 19054106 | 17084664 | 17970012 | 17748618 | 7809210.5 | 12964966 |
| Q80WJ7 | Mtdh | Protein LYRIC | 260.35 | 12.95 | 63846 | 5 | 863684.75 | 846007.81 | 651149.69 | 1122301.75 | 1200430.75 | 1500852.38 | 538237.38 | 1022016.75 | 3146888 | 2010927.25 | 2663551 | 2893410.25 | 1613111.12 | 2328980 | 122602.3 | 4395833 | 2 | 1591355.25 | 2352877.5 | 959698 |
| Q922D8 | Mthfd1 | C-1-tetrahydrofolate synthase, cytoplasmic | 429.61 | 12.83 | 101200 | 10 | 10855043 | 12847813 | 7876901.5 | 12099069 | 13685572 | 5068405 | 5026814.5 | 6737143 | 7560533.5 | 4759556.5 | 7925712 | 7086847 | 10204826 | 9285212 | 8244720 | 6200995 | 9569476 | 12682532 | 11670307 | 9005314 |
| Q3V3R1 | Mthfd1l | Monofunctional C1-tetrahydrofolate synthase, mit | 967.98 | 17.09 | 105729 | 17 | 6780439 | 6671691.5 | 5859803 | 7799267.5 | 9059728 | 5612687 | 6704728 | 6014438 | 16566150 | 8453444 | 6834477.5 | 4429546 | 6317811 | 5699434.5 | 4446788.5 | 4460335 | 4066388 | 5446905 | 3904819 | 7417998.5 |
| P18155 | Mthfd2 | Bifunctional methylenetetrahydrofolate dehydroge | 413.85 | 35.14 | 37863 | 9 | 1463138.88 | 2080885.62 | 72582.38 | 5298919.5 | 1553316.5 | 3716210 | 3156391.5 | 1760591 | 3841450.5 | 4393582 | 3265753.5 | 4763165 | 4101036.5 | 4387004 | 3526630.5 | 6137826 | 5517140 | 4750741 | 1637926.5 | 4914986 |
| P03921 | Mtnd5 | NADH-ubiquinone oxidoreductase chain 5 | 109.49 | 5.44 | 68475 | 2 | 2 | 2 | 181170.61 | 142938.64 | 45948.41 | 2 | 101711.7 | 37773.57 | 309550.03 | 155209.55 | 2 | 2 | 2 | 2 | 637448.44 | 2 | 2 | 107877.79 | 2 | 2 |
| P62774 | Mtpn | Myotrophin | 245.57 | 26.27 | 12861 | 3 | 2 | 1095913.5 | 2 | 1919900.5 | 2840651 | 4516728.5 | 7197681 | 7915985 | 7837965 | 3481241 | 8185089 | 8744912 | 9448653 | 6625326 | 4733332 | 1573543.62 | 4006832.75 | 3397590.75 | 5039611.5 | 1852668.62 |
| Q9CZU3 | Mtrex | Exosome RNA helicase MTR4 | 238.78 | 5 | 117636 | 5 | 1369454.25 | 1656076.25 | 2320006 | 198553.06 | 1664473.88 | 176903.86 | 398102.38 | 105758.35 | 1718006.75 | 1572978.12 | 2655701.75 | 1727701.5 | 1454283 | 1705959 | 3001103.5 | 1617570.25 | 2041132.88 | 1317010 | 2 | 1139113.38 |
| Q8R1S4 | Mtss1 | Protein MTSS 1 | 317.33 | 8.96 | 82408 | 5 | 1058211.75 | 1119015.88 | 1308424.75 | 475508.38 | 2937052.25 | 2685461.75 | 1191398 | 2005855.5 | 5052270 | 1902276 | 4757602.5 | 4277759.5 | 2236723 | 2733115.75 | 3122340 | 3374362.75 | 3485084.25 | 3877317 | 81927.39 | 3497870 |
| P47802 | Mtx1 | Metaxin-1 | 56.72 | 5.68 | 35624 | 1 | 116579.55 | 91164.46 | 147000.53 | 82939.05 | 103865.27 | 2 | 2 | 55171.68 | 102821.57 | 149783.92 | 2 | 52770.87 | 2 | 136581.08 | 354581.97 | 2 | 2 | 2 | 2 | 2 |
| O88441 | Mtx2 | Metaxin-2 | 42.99 | 4.18 | 29758 | 1 | 2 | 2 | 83846.53 | 2 | 2 | 2 | 2 | 2 | 256216.02 | 2 | 2 | 2 | 2 | 64869.79 | 2 | 2 | 2 | 2 | 2 | 2 |
| Q99JF5 | Mvd | Diphosphomevalonate decarboxylase | 33.31 | 2 | 44072 | 1 | 2 | 276890.44 | 2 | 1073835.75 | 2 | 2 | 376409.41 | 2 | 2 | 389607.81 | 649265.94 | 2 | 1994622.88 | 2 | 914533.62 | 2 | 1455176 | 2078776.75 | 1427127.75 | 1508338.12 |
| Q9EQK5 | Mvp | Major vault protein | 957.51 | 22.76 | 95924 | 16 | 16983506 | 17974464 | 16716660 | 13496643 | 18498924 | 11739071 | 8994401 | 13201104 | 13138908 | 16165552 | 15276967 | 13669997 | 14399027 | 8777938 | 11883016 | 13522547 | 5367795.5 | 11687463 | 10817583 | 7591192 |
| O35682 | Myadm | Myeloid-associated differentiation marker | 302.69 | 21.25 | 35285 | 4 | 653041.06 | 1371613.12 | 722949.25 | 2607943.5 | 551924.69 | 2252708 | 937192.75 | 1201900.38 | 2129349 | 933837.88 | 2 | 2 | 2 | 2 | 2 | 1874527.75 | 2 | 2 | 2699133.5 | 2 |
| Q7TPV4 | Mybbp1a | Myb-binding protein 1A | 622 | 10.34 | 152037 | 13 | 11748572 | 15640232 | 8310070.5 | 12372192 | 11107584 | 9630485 | 8620172 | 12837765 | 7896032 | 9188104 | 10172424 | 9050800 | 13352510 | 14492735 | 6192767 | 19384078 | 18805010 | 24080896 | 23489408 | 18923574 |
| Q9CPT4 | Mydgf | Myeloid-derived growth factor | 111.13 | 16.27 | 17982 | 2 | 1598845.38 | 1729071.88 | 2969832.75 | 2120403 | 2586898.5 | 632000.88 | 665668.62 | 1914178.75 | 637530.94 | 2324775.25 | 4135803.75 | 2894894.75 | 4781744.5 | 3239149 | 2392328.25 | 3348804 | 3571744 | 3118410.5 | 2684700.5 | 3171564.5 |
| Q8C854 | Myef2 | Myelin expression factor 2 | 464.28 | 22.84 | 63295 | 10 | 2822063 | 7629188 | 3985145 | 3803605.5 | 11793214 | 9219143 | 10906660 | 11696676 | 12391065 | 6788964 | 6643222 | 6472051.5 | 6585379 | 8495028 | 3804650 | 4253688 | 6255369.5 | 4850522.5 | 8249418 | 5244303 |
| Q9JK81 | Myg1 | MYG1 exonuclease | 300.77 | 19.74 | 42723 | 6 | 8712497 | 8888581 | 7237035 | 6261198 | 7507057.5 | 6748610 | 5363955 | 7250482 | 6291544 | 7404553 | 9903092 | 8733480 | 5613204.5 | 4516756 | 6486083 | 8878641 | 7981093.5 | 8663368 | 8580462 | 8312949 |
| A2AQP0 | Myh7b | Myosin-7B | 83.49 | 0.88 | 221497 | 2 | 9182563 | 13507803 | 75768.74 | 15127328 | 2622858.75 | 18483606 | 10243457 | 1694266.62 | 30398210 | 20142752 | 184951922 | 202230274 | 127388786 | 187972418 | 91578922 | 340920322 | 237405666 | 126010874 | 83090978 | 143220674 |
| Q8VDD5 | Myh9 | Myosin-9 | 6340.121 | 49.54 | 226372 | 99 | 124756418 | 164666674 | 108607530 | 129897722 | 160523282 | 129501858 | 148523074 | 179642546 | 171053250 | 165409826 | 142028322 | 167512418 | 179914370 | 172526898 | 102423258 | 166127666 | 117964578 | 119570746 | 143176210 | 125290210 |
| Q60605 | Myl6 | Myosin light polypeptide 6 | 671.06 | 75.5 | 16930 | 11 | 102741986 | 192252562 | 123232466 | 158589266 | 128623970 | 130729938 | 118422634 | 154811186 | 95665810 | 122879106 | 87240922 | 83216290 | 97496970 | 107707954 | 84022106 | 93629010 | 107849362 | 138046114 | 127537986 | 130677962 |
| Q63355 | Myo1c | Unconventional myosin-Ic | 1088.89 | 20.21 | 119811 | 18 | 2 | 2 | 2 | 2 | 2 | 2 | 510500.81 | 2 | 2 | 2 | 2 | 2 | 2 | 2 | 2 | 2 | 2 | 2 | 2 | 2 |
| Q9WTI7 | Myo1c | Unconventional myosin-Ic | 1257 | 22.95 | 121944 | 21 | 33232.53 | 2 | 2 | 2 | 2 | 160580.95 | 2 | 70762.62 | 1706947.25 | 2 | 144366.97 | 2 | 478593.88 | 2 | 2 | 239022.05 | 2 | 2 | 2 | 511791.91 |
| E9Q634 | Myo1e | Unconventional myosin-Ie | 445.54 | 12.56 | 126818 | 10 | 2019042.75 | 2130513.25 | 2325734.25 | 1575068.5 | 2006487.88 | 1591630.75 | 1209055.5 | 224490.03 | 3408237 | 1843765.88 | 2322521.25 | 2138297.75 | 2922257.5 | 2353025.25 | 1531348.5 | 1535794.25 | 536508.62 | 1902320 | 855713.5 | 2 |
| P70248 | Myo1f | Unconventional myosin-If | 192.09 | 3.55 | 125947 | 4 | 2 | 2 | 344364.84 | 2 | 1077788.88 | 2 | 2 | 917533.56 | 73117.61 | 134505.86 | 2 | 881149.31 | 2 | 2 | 746963.56 | 2 | 2 | 2 | 2 | 2 |
| Q5SUA5 | Myo1g | Unconventional myosin-Ig | 89.79 | 2.64 | 117227 | 2 | 2 | 571411.56 | 2 | 2 | 2 | 2 | 2 | 2 | 365638.69 | 2 | 2 | 2 | 2 | 2 | 2 | 2 | 2 | 2 | 2 | 559276.12 |
| Q99104 | Myo5a | Unconventional myosin-Va | 99.26 | 1.67 | 215538 | 3 | 1682476.88 | 2 | 2013907.75 | 985074.56 | 1613183.25 | 516435.78 | 501387.28 | 1125769.75 | 671883.5 | 1125578.62 | 1741747.62 | 296087.03 | 628005.81 | 499792.22 | 1374276 | 2610861.25 | 422243.62 | 521588.75 | 502711.03 | 398383.5 |
| Q69ZN7 | Myof | Myoferlin | 581.43 | 6.69 | 233324 | 11 | 5031534 | 4741635 | 2464479.75 | 1675938.75 | 1839702.75 | 4447420 | 4947511 | 7072817 | 3494810 | 4579746.5 | 9010014 | 9206379 | 6817130 | 9112358 | 2766071.5 | 12003277 | 11177555 | 8210976 | 11820012 | 8655194 |
| Q80UM3 | Naa15 | N-alpha-acetyltransferase 15, NatA auxiliary subu | 145.59 | 4.16 | 100961 | 3 | 1582372.62 | 72135.76 | 518811.69 | 115409.57 | 1365965.12 | 2 | 152682.05 | 2 | 2 | 2 | 2 | 90276.83 | 109890.77 | 413559.53 | 476779.97 | 2153122.75 | 943140.25 | 2 | 703203.81 | 1627155.88 |
| Q8VE10 | Naa40 | N-alpha-acetyltransferase 40 | 60.3 | 5.06 | 27229 | 1 | 2 | 2 | 2 | 2 | 63379.79 | 2 | 2 | 2 | 42800.21 | 80294.31 | 84679.04 | 2 | 63393.28 | 2 | 288810.25 | 2 | 2 | 128094.05 | 2 | 2 |
| P70670 | Naca | Nascent polypeptide-associated complex subunit | 379.16 | 3.34 | 220499 | 5 | 3288299.25 | 1399716 | 11023224 | 3080776.25 | 4828937.5 | 2953856 | 3810657.25 | 5418185 | 1973876 | 5182418.5 | 3800566 | 3764466.25 | 5534547.5 | 4338622 | 4647171.5 | 2380060.75 | 3539511 | 1794659.62 | 1086184.5 | 2528630 |
| Q8VBW6 | Nae1 | NEDD8-activating enzyme E1 regulatory subunit | 47.09 | 1.87 | 60274 | 1 | 941643.75 | 199879.64 | 172591.36 | 187883.06 | 75477.48 | 104805.47 | 311894.97 | 307387.41 | 389392.97 | 278488.06 | 95502.22 | 195449.42 | 324268.31 | 281874.59 | 443547.88 | 2 | 142548.69 | 108748.75 | 2 | 2 |
| Q9QWR8 | Naga | Alpha-N-acetylgalactosaminidase | 391.34 | 21.45 | 47235 | 6 | 16937934 | 21701574 | 13747993 | 3549274.25 | 13916981 | 11739228 | 15029969 | 16961376 | 10558998 | 12611924 | 7970166.5 | 6973715.5 | 10817940 | 10850923 | 2414233.5 | 6060424 | 4817002 | 2018677.75 | 5581674.5 | 3337258.5 |
| P81799 | Nagk | N-acetyl-D-glucosamine kinase | 62.07 | 2.92 | 37196 | 1 | 1261938 | 1158371 | 1011727.94 | 659119.56 | 1081482.75 | 755176 | 742635.81 | 2 | 792218 | 1050637.38 | 951833.5 | 740815.44 | 802238 | 753407.06 | 1022700.12 | 867598.5 | 1274477.75 | 1327315.5 | 961407.44 | 2 |
| Q8BJ48 | Nagpa | N-acetylglucosamine-1-phosphodiester alpha-N-a | 260.02 | 10.06 | 56044 | 4 | 874187.12 | 2732565 | 2 | 2681817.25 | 958009.56 | 2 | 2 | 1364755.88 | 1168209.25 | 2 | 1308329.12 | 787417.12 | 2 | 1131368.88 | 2057773.38 | 436017.19 | 1164386.5 | 1439080.38 | 1456397 | 972872.5 |
| Q80Z29 | Nampt | Nicotinamide phosphoribosyltransferase | 211.51 | 12.22 | 55438 | 5 | 3552803.5 | 3864766.5 | 3482315 | 5793184.5 | 5809184.5 | 7453220 | 5718451.5 | 8704219 | 8856752 | 7915454 | 6698536.5 | 5358302 | 5385704 | 5975802 | 3938434 | 7612761.5 | 5884829.5 | 7763420.5 | 7915381 | 7467488 |
| Q99J77 | Nans | N-acetylneuraminate-9-phosphate synthase | 211.17 | 15.32 | 40024 | 4 | 1865656.88 | 3269629.5 | 2632050 | 2145249.5 | 3250990.25 | 2575700 | 3046595.5 | 3255095 | 3057282.25 | 2819185.25 | 3518802.25 | 3523216.75 | 3346991.75 | 3165028.5 | 2021831.62 | 3511078.25 | 2442557.25 | 1937048.62 | 378662.81 | 1802011.25 |
| P28656 | Nap1l1 | Nucleosome assembly protein 1-like 1 | 179.77 | 12.02 | 45345 | 3 | 1951933.5 | 295461.94 | 5139031.5 | 5490414 | 5357791.5 | 507028.94 | 2380314.75 | 765114.94 | 2709621 | 4008197.5 | 373985.09 | 2 | 1087174.38 | 1000198.5 | 2 | 2 | 2 | 2 | 2 | 2 |
| Q78ZA7 | Nap1l4 | Nucleosome assembly protein 1-like 4 | 218.24 | 11.47 | 42679 | 4 | 3800152.75 | 1854539.62 | 3364547.75 | 4354118 | 3889170.25 | 2292509 | 2205306.75 | 3247710.5 | 2295402 | 2856551.5 | 2306022.25 | 1254050.88 | 2709665.5 | 2004064.38 | 2706094.75 | 3257473.75 | 1972355.75 | 3029387.5 | 2737314.5 | 2958763.5 |
| P54921 | Napa | Alpha-soluble NSF attachment protein | 426.67 | 33.56 | 33193 | 8 | 5305131 | 468511.75 | 6291800 | 4541288.5 | 7333117 | 2807310.75 | 5320334.5 | 4893088.5 | 4524527 | 6427165 | 2447004.25 | 2007481.75 | 3744235.5 | 2394148 | 2711706.25 | 2328015.5 | 1480392.25 | 2199435.5 | 223612.33 | 1910251.88 |
| Q9CWZ7 | Napg | Gamma-soluble NSF attachment protein | 92.35 | 4.49 | 34732 | 2 | 2662191 | 1130505.62 | 2707062.25 | 2567666 | 2077869.75 | 889388.75 | 732696 | 935287.88 | 2573530.25 | 2897854.75 | 161494.03 | 227488.69 | 1818851.38 | 1402031.38 | 910754.56 | 2 | 472307.56 | 268797.47 | 249535.89 | 470370.53 |

| | | | | | | | | | | | | | | | | | | | | | | | | | | |
|---|---|---|---|---|---|---|---|---|---|---|---|---|---|---|---|---|---|---|---|---|---|---|---|---|---|---|
| O09043 | Napsa | Napsin-A | 169.67 | 13.84 | 45544 | 3 | 16189915 | 18799612 | 13301041 | 13880866 | 11503477 | 13429767 | 10915863 | 15108324 | 14470656 | 12683789 | 10354294 | 9532162 | 11092173 | 9808202 | 4511010.5 | 12049505 | 6660784 | 8497764 | 8948697 | 5883593 |
| Q8BP47 | Nars1 | Asparagine--tRNA ligase, cytoplasmic | 467.25 | 20.57 | 64279 | 9 | 1869360 | 6212728 | 2245978.75 | 3153781.25 | 5512128 | 7168069 | 8152496 | 11061540 | 10577510 | 7196997 | 11909907 | 13311732 | 5311852.5 | 8530780 | 3170412 | 8243874 | 2963372.5 | 3807652 | 6954885 | 1379614 |
| Q66HD3 | Nasp | Nuclear autoantigenic sperm protein | 146.24 | 5.67 | 84200 | 3 | 1761130.88 | 3454937.5 | 3083415.75 | 2987387.75 | 5566836.5 | 1896204.88 | 899731 | 88893.5 | 1083178.12 | 2085383.75 | 3659502.75 | 7677876.5 | 4409363.5 | 1772932.5 | 1208114.38 | 5535971.5 | 1401823.25 | 1784261.5 | 4943595.5 | 1279893.5 |
| Q8K224 | Nat10 | RNA cytidine acetyltransferase | 278.16 | 8.89 | 115419 | 6 | 799520.38 | 1759003.5 | 1545337 | 1106501.75 | 3669982.75 | 1555737.25 | 1599447.62 | 705511.25 | 1700880 | 1303565.75 | 2429458.75 | 2 | 1021278.38 | 2846246.5 | 1191337.25 | 463843.69 | 2947404.5 | 1772935.12 | 3794742.75 | 1558110.88 |
| P50295 | Nat2 | Arylamine N-acetyltransferase 2 | 46.52 | 2.41 | 33701 | 1 | 2 | 1023469.19 | 2 | 751146.12 | 55810.73 | 176188.02 | 117170.77 | 140426.61 | 243938.45 | 2 | 2 | 492981.78 | 2 | 817303.94 | 477660.75 | 641774.75 | 682035.19 | 660375.25 | 2 | 884089.06 |
| Q8C156 | Ncaph | Condensin complex subunit 2 | 75.35 | 2.46 | 82303 | 2 | 351273.25 | 1535207.12 | 972978.56 | 1715814.38 | 3508738 | 1083882.25 | 939058.56 | 2 | 1016152.75 | 1190568.62 | 2881668.5 | 2112692.5 | 1719144.25 | 1742198.5 | 1166320.12 | 2503369.25 | 2460475.5 | 1723927.38 | 1649259.88 | 2745133.75 |
| Q8BLF1 | Nceh1 | Neutral cholesterol ester hydrolase 1 | 834.11 | 31.86 | 45740 | 11 | 2452681 | 6522655 | 2121697 | 4225174.5 | 2681418.75 | 3239953.25 | 5333365.5 | 3613047.25 | 6247228 | 2576231 | 4450250 | 3017610.5 | 4510605 | 5222817 | 3475628 | 6416176 | 5663941 | 5113157 | 4328447 | 2135955.25 |
| Q09014 | Ncf1 | Neutrophil cytosol factor 1 | 250.29 | 12.31 | 44667 | 3 | 2981441.75 | 1262007.25 | 2668638 | 3852038.5 | 2751197 | 405663.06 | 3659283 | 3074873.25 | 3543981.5 | 3952931 | 2460563.25 | 2382914 | 3993716.75 | 2846724.5 | 3269233 | 1983086.5 | 2867906.5 | 3171312.5 | 1043373 | 2511184.5 |
| O70145 | Ncf2 | Neutrophil cytosol factor 2 | 242.78 | 10.29 | 59485 | 5 | 10453740 | 7764297 | 9135245 | 11035356 | 6368168 | 4601927 | 3864192 | 3500713 | 7188726 | 4653514 | 7071908 | 7112322.5 | 6191843 | 4788212.5 | 3495660.25 | 4296729.5 | 1761989 | 1753394.75 | 7163649.5 | 2913393.75 |
| P97369 | Ncf4 | Neutrophil cytosol factor 4 | 316.41 | 23.3 | 38707 | 7 | 3633692 | 3575241 | 5861834.5 | 3407018 | 11144370 | 6191383.5 | 3590122.5 | 4281676.5 | 6094370.5 | 4438619 | 9568875 | 4200926 | 2913263.25 | 1186873 | 3963525 | 5431234.5 | 5177531 | 5844824 | 4580238.5 | 1839175.12 |
| Q8K1X4 | Nckap1l | Nck-associated protein 1-like | 493.11 | 13.23 | 128905 | 9 | 6614639.5 | 5232776 | 6338283 | 4432460 | 7198865 | 2959159.5 | 5334987 | 8033285.5 | 4792920 | 5951388.5 | 3447525 | 4333870 | 6319404.5 | 5182330.5 | 4593691.5 | 2707175 | 3534296.5 | 4218610.5 | 4997550 | 2021312.25 |
| P09405 | Ncl | Nucleolin | 2415.4 | 39.75 | 76723 | 38 | 70650722 | 93761978 | 82491282 | 101139562 | 81065178 | 82090466 | 94629650 | 81875850 | 105684778 | 89979298 | 68862890 | 80568834 | 92121682 | 86447858 | 83022290 | 83494226 | 83951858 | 75201050 | 82587498 | 74220754 |
| Q5XIA1 | Ncln | BOS complex subunit NCLN | 214.01 | 7.46 | 62992 | 4 | 1702517.12 | 3378343.75 | 2907789 | 3512092 | 3339495.75 | 1053771.38 | 2326700.25 | 2832336.5 | 3639879.75 | 1541303.75 | 3284254.75 | 2561464.25 | 2761385.5 | 2675812 | 2436216 | 2449422.25 | 1318321.88 | 3226733 | 1626822 | 3312990.5 |
| Q91W39 | Ncoa5 | Nuclear receptor coactivator 5 | 49.57 | 1.73 | 65319 | 1 | 2 | 1759668.75 | 2 | 2 | 788468.31 | 754566.19 | 2 | 2 | 2 | 2 | 994627.88 | 2 | 1137483.88 | 1043244.62 | 2 | 2 | 430424.03 | 2 | 2 | 2 |
| P57716 | Ncstn | Nicastrin | 485.97 | 15.11 | 78492 | 9 | 12442723 | 15361036 | 4259427 | 10574167 | 2662665.5 | 11364729 | 11739333 | 6599812.5 | 7678664.5 | 8516665 | 7251463.5 | 8844271 | 8743596 | 7331616 | 9180587 | 8732785 | 12309859 | 11559210 | 1489123 | 14678874 |
| Q8VCB1 | Ndc1 | Nucleoporin NDC1 | 290.79 | 11.89 | 75409 | 5 | 982558.75 | 174492.16 | 586051.88 | 2770785.5 | 77365.73 | 2 | 2 | 654333.44 | 87040.63 | 2 | 199676.94 | 333644.56 | 2 | 2 | 757536.12 | 2 | 2 | 166247.56 | 257316.3 | 2241843.25 |
| Q62433 | Ndrg1 | Protein NDRG1 | 431.7 | 21.32 | 43009 | 6 | 2195349.25 | 3200801.75 | 4223435.5 | 1512579.75 | 4214666.5 | 7044810 | 5368003.5 | 7987352.5 | 5291346 | 7852961.5 | 12203046 | 9599240 | 7615808 | 4357332 | 4164202 | 6741291.5 | 5685017 | 6320821 | 7349623 | 7718850 |
| Q80W89 | Ndufa11 | NADH dehydrogenase [ubiquinone] 1 alpha subc | 88.35 | 16.31 | 14854 | 1 | 213339.84 | 512669.16 | 221170.66 | 725629.94 | 286684.91 | 2 | 779867.19 | 345511.5 | 522191.84 | 206946.08 | 112930.09 | 221410 | 77770.42 | 361474.5 | 437564.69 | 53144.18 | 460175.22 | 403758.69 | 231843.66 | 2 |
| Q7TMF3 | Ndufa12 | NADH dehydrogenase [ubiquinone] 1 alpha subc | 54.8 | 6.21 | 17086 | 1 | 1292617 | 439161.16 | 1355120 | 819242.31 | 1277067.88 | 276542.47 | 1210296.38 | 315515.75 | 1354448 | 1262708.12 | 2 | 606499 | 729440.44 | 684910.94 | 2 | 2 | 2 | 2 | 2 | 2 |
| Q9ERS2 | Ndufa13 | NADH dehydrogenase [ubiquinone] 1 alpha subc | 128.28 | 22.92 | 16860 | 3 | 5489364.5 | 5925315.5 | 4368043.5 | 4973944.5 | 3485926.5 | 4244008.5 | 4244567.5 | 3880203 | 2953882.5 | 2774555.75 | 2627483.5 | 1704241 | 2391005.75 | 2071138.25 | 2053243.38 | 1673015.88 | 1649582.38 | 3156083.5 | 391454.22 | 1767813.75 |
| Q9CQ75 | Ndufa2 | NADH dehydrogenase [ubiquinone] 1 alpha subc | 76.85 | 14.14 | 10916 | 1 | 1073813.75 | 2456893.75 | 1108872.25 | 2 | 1503855.38 | 1942883.5 | 1271327 | 1851692.88 | 1322975.75 | 837980.56 | 1125335.38 | 951566.06 | 959188.69 | 917084.25 | 2 | 2 | 2 | 2 | 2 | 2 |
| Q62425 | Ndufa4 | Cytochrome c oxidase subunit NDUFA4 | 269.19 | 78.05 | 9327 | 6 | 9717470 | 2558929 | 7266871 | 5583622.5 | 7485388 | 4173039.75 | 3926813.75 | 2853380.25 | 5041305 | 4117846.5 | 3415543 | 2682483.5 | 7207803.5 | 2604042.5 | 6057281 | 1861697.75 | 1706565.25 | 4448245 | 2590307.5 | 1613801.25 |
| Q5BK63 | Ndufa9 | NADH dehydrogenase [ubiquinone] 1 alpha subc | 190.52 | 11.14 | 42559 | 3 | 1812201.12 | 1321541.12 | 514055.75 | 1666522.25 | 217506.06 | 986630.25 | 405165.53 | 102012.91 | 220452.14 | 147467.5 | 68012.62 | 88906.38 | 250413.31 | 415854.12 | 409671.78 | 2 | 423691.94 | 141351.53 | 113765.64 | 200320.64 |
| Q9DCS9 | Ndufb10 | NADH dehydrogenase [ubiquinone] 1 beta subco | 179.29 | 33.52 | 21024 | 4 | 601239.88 | 108412.8 | 837900.81 | 1531677.62 | 1649202.75 | 103629.08 | 1233313.62 | 1087556.88 | 1266945.25 | 1902135.5 | 2 | 761488.12 | 2 | 910901.62 | 507719.94 | 2 | 637366.94 | 2 | 521525.16 | 1699062 |
| Q9CQH3 | Ndufb5 | NADH dehydrogenase [ubiquinone] 1 beta subco | 123.36 | 14.81 | 21710 | 3 | 567225.88 | 2 | 1519570.5 | 2 | 581673.5 | 2 | 2 | 2 | 523490.75 | 743326.12 | 2 | 2 | 2 | 157732.42 | 1248387.88 | 2 | 2 | 502505.28 | 2 | 2 |
| Q91VD9 | Ndufs1 | NADH-ubiquinone oxidoreductase 75 kDa subuni | 643.64 | 22.01 | 79777 | 14 | 6022782.5 | 5873129 | 7168196.5 | 7931593 | 10503735 | 5008654 | 2887670.5 | 5779651 | 4063099.25 | 5965970.5 | 3034090.25 | 1794975.25 | 5258641 | 4598194 | 7806179 | 3246227.75 | 8788156 | 5720888 | 998356.62 | 10603481 |
| Q91WD5 | Ndufs2 | NADH dehydrogenase [ubiquinone] iron-sulfur pro | 369.62 | 15.33 | 52626 | 6 | 2812738.5 | 3707192.75 | 1914599.5 | 3511760 | 2510454.25 | 1251882.88 | 2417935 | 129536.37 | 1333782 | 1083877.75 | 93646.58 | 157165.88 | 1322492.12 | 1456137.88 | 355400.22 | 2 | 326249.25 | 234073.33 | 70574.06 | 645298.69 |
| Q9DCT2 | Ndufs3 | NADH dehydrogenase [ubiquinone] iron-sulfur pro | 436.46 | 25.48 | 30149 | 8 | 1685487.75 | 1535637.25 | 5006471.5 | 859033.69 | 3108512.75 | 1960729.62 | 574041.56 | 1766878.25 | 1732995 | 3568086 | 1039460.19 | 1604375.62 | 1386423.25 | 1733199.25 | 2657121.25 | 2653988.75 | 3617822.25 | 4111867 | 2332506.75 | 2224756.5 |
| Q9DC70 | Ndufs7 | NADH dehydrogenase [ubiquinone] iron-sulfur pro | 150.43 | 10.27 | 24683 | 2 | 2689957.75 | 7029256 | 3418484.75 | 6247232 | 3656251.75 | 7954897 | 6373283 | 3911064.75 | 6434464 | 4593360.5 | 3294419 | 2769830.75 | 2286672.75 | 1853332.5 | 1060118.62 | 2335182 | 1106725.38 | 1030336.56 | 1192718.25 | 2 |
| Q91YT0 | Ndufv1 | NADH dehydrogenase [ubiquinone] flavoprotein 1 | 147.55 | 10.13 | 50834 | 3 | 1921741.5 | 1833127.25 | 1783156.75 | 2086493.38 | 1671178.38 | 1571856.75 | 1208416.62 | 1748422.25 | 1242223 | 1251929.12 | 940153.38 | 681869.38 | 1187536.5 | 1028157.44 | 1293136.12 | 561737.31 | 766497.5 | 764194.56 | 796410.06 | 804999.81 |
| P69682 | Necap1 | Adaptin ear-binding coat-associated protein 1 | 39.24 | 6.14 | 29792 | 1 | 2 | 2 | 2 | 2 | 2 | 2 | 2 | 2 | 2 | 2 | 2 | 2 | 2 | 2 | 2 | 2 | 2 | 2 | 2 | 2 |
| Q9D1J1 | Necap2 | Adaptin ear-binding coat-associated protein 2 | 127.91 | 18.42 | 28598 | 3 | 2 | 2 | 2 | 245752.5 | 322436.03 | 2 | 303895.09 | 2 | 265070.53 | 2 | 812674.5 | 600869.25 | 367753.12 | 351569.84 | 827695.62 | 2 | 826176.56 | 2 | 2 | 197292.52 |
| Q6P756 | Necap2 | Adaptin ear-binding coat-associated protein 2 | 95.27 | 17.49 | 28405 | 2 | 2 | 2 | 2 | 2 | 421139.38 | 2 | 525522.19 | 1255664.62 | 969670.56 | 270099 | 1388071.12 | 545412.19 | 2 | 715082.94 | 2 | 2 | 2 | 2 | 612620.56 | 2 |
| Q8K1R7 | Nek9 | Serine/threonine-protein kinase Nek9 | 58.25 | 1.52 | 107143 | 1 | 1226433 | 2 | 2 | 792325.81 | 1250235.25 | 938198.88 | 805515.56 | 1157283.88 | 975049.44 | 906374.88 | 1024090.81 | 820535.62 | 726733.88 | 2 | 2 | 2 | 628466.19 | 2 | 2 | 2 |
| Q6P5H2 | Nes | Nestin | 2237.76 | 25.48 | 207124 | 36 | 26189828 | 31193050 | 24225834 | 28153884 | 37743018 | 19674764 | 19417432 | 30854592 | 24378938 | 22886950 | 38771094 | 31888108 | 28596596 | 29926854 | 13418362 | 29382728 | 32196446 | 18901252 | 33166466 | 23161558 |
| Q60591 | Nfatc2 | Nuclear factor of activated T-cells, cytoplasmic 2 | 75.72 | 4.42 | 100020 | 2 | 2 | 844871.69 | 2 | 2 | 1099598 | 2 | 2 | 61489.83 | 725365.06 | 2 | 760427.88 | 792942.56 | 597161.69 | 863110.31 | 496774 | 2 | 755420.69 | 2 | 1197627.75 | 612732.06 |
| P25799 | Nfkb1 | Nuclear factor NF-kappa-B p105 subunit | 174.98 | 5.05 | 105615 | 4 | 2 | 2 | 465550.12 | 2 | 1073862.38 | 2 | 2 | 2 | 1333582 | 390088.5 | 928058.56 | 500831.5 | 214398.95 | 1247930.38 | 595043.5 | 2 | 115659.03 | 827724.62 | 222627.81 | 802648.25 |
| Q9WTK5 | Nfkb2 | Nuclear factor NF-kappa-B p100 subunit | 63.96 | 1.56 | 96832 | 1 | 2 | 2 | 2 | 2 | 97281.12 | 2 | 247675.55 | 2 | 214185.94 | 172541.45 | 2 | 2 | 103126.85 | 151755.59 | 105578.68 | 2 | 2 | 2 | 2 | 2 |
| O54910 | Nfkbie | NF-kappa-B inhibitor epsilon | 65.83 | 3.3 | 39200 | 1 | 2 | 2 | 130388.29 | 143955.3 | 97709.3 | 2 | 296837.78 | 124408.42 | 177206.12 | 127735.96 | 2 | 2 | 86888.17 | 242976.22 | 2 | 2 | 2 | 2 | 2 | 2 |
| Q9QZ23 | Nfu1 | NFU1 iron-sulfur cluster scaffold homolog, mitoch | 220.08 | 27.06 | 28568 | 5 | 1808615.25 | 1808231.75 | 1637104.62 | 2433262.75 | 2898107 | 946515.81 | 866051.38 | 2434337.5 | 4142326.5 | 2687623.5 | 3750257 | 388281.19 | 2744135 | 3283885.25 | 2575385.75 | 2 | 4420663.5 | 3945616.5 | 1040846.38 | 2401330.25 |
| Q8CCH2 | Nhlrc3 | NHL repeat-containing protein 3 | 323.61 | 23.63 | 38195 | 5 | 1681669.5 | 1356306.5 | 2495405.25 | 876724.5 | 371112.91 | 477529.34 | 869653.88 | 473436.44 | 1616215.38 | 2107578 | 317980.81 | 306703.84 | 1043262.62 | 600024 | 1496533.5 | 1068673.5 | 1260676.75 | 365807.09 | 1214046.12 | 455557.44 |
| Q9CRB2 | Nhp2 | H/ACA ribonucleoprotein complex subunit 2 | 150.45 | 12.42 | 17247 | 2 | 2 | 96022.74 | 2 | 74720.59 | 313835.84 | 177314.09 | 2 | 2 | 2 | 465464.22 | 2 | 93769.99 | 2 | 2 | 2 | 2 | 213495.61 | 2 | 2 | 315803.41 |
| Q3UW53 | Niban1 | Protein Niban 1 | 578.63 | 14.04 | 102649 | 12 | 11146725 | 14968255 | 9242685 | 14001358 | 19531028 | 13637630 | 13273816 | 14002027 | 20048890 | 17458496 | 18962850 | 20228632 | 19950640 | 17562650 | 12255116 | 21490206 | 18509316 | 19358846 | 22651332 | 18486924 |
| Q8R1F1 | Niban2 | Protein Niban 2 | 557.52 | 16.42 | 84819 | 11 | 4329201.5 | 3986480 | 6033422.5 | 2105416.5 | 2302049 | 7537920 | 7720119.5 | 6796071.5 | 4388783 | 4490143.5 | 7660116 | 3267919.75 | 6402190 | 5034158.5 | 4183421 | 6566265 | 6023073 | 6086367 | 4788129 | 4795792 |
| Q9EQ80 | Nif3l1 | NIF3-like protein 1 | 197.98 | 12.23 | 41746 | 3 | 45143.2 | 1878861.75 | 410040.75 | 2436397 | 3451036 | 2558898.5 | 2845066.75 | 3480456.5 | 1940987.62 | 2251500.25 | 3735775.25 | 3613045.25 | 3682550.75 | 3300895.5 | 2049405 | 2602649.75 | 2403908.25 | 1651714.12 | 1862460.12 | 1481720.38 |
| Q91VE6 | Nifk | MKI67 FHA domain-interacting nucleolar phospho | 162.77 | 14.51 | 36265 | 3 | 2 | 2 | 2 | 100134.38 | 2 | 2 | 2 | 2 | 854035.25 | 2 | 2 | 2 | 2 | 407321.72 | 411561.72 | 2 | 2 | 151078.98 | 178792.2 | 355832.97 |
| Q9CXK8 | Nip7 | 60S ribosome subunit biogenesis protein NIP7 ho | 104.25 | 23.33 | 20452 | 2 | 2 | 160394.58 | 2 | 242830.75 | 82458.59 | 2 | 2 | 279423.34 | 245052.58 | 165852.92 | 2 | 2 | 2 | 186890.2 | 2 | 2 | 2 | 2 | 325541.56 | 2 |
| O55125 | Nipsnap1 | Protein NipSnap homolog 1 | 51.35 | 3.17 | 33363 | 1 | 970621.06 | 1782762 | 1352573 | 759483.62 | 2074820.12 | 2043409.38 | 625542.12 | 2271984 | 2905218.75 | 2850992.5 | 6771081.5 | 4350709.5 | 4932520 | 5264809.5 | 1971781.62 | 6145674.5 | 4341445 | 897802.75 | 5653898 | 2991918 |
| O55126 | Nipsnap2 | Protein NipSnap homolog 2 | 58.33 | 3.2 | 32933 | 1 | 367098.19 | 2 | 2 | 444706.88 | 2 | 2 | 499284.47 | 756903.31 | 378200.5 | 2 | 2 | 2 | 2 | 2 | 2 | 2 | 2 | 2 | 2 | 2 |
| Q80TM9 | Nisch | Nischarin | 142.01 | 2.39 | 175012 | 3 | 2 | 441973.53 | 2 | 2 | 2 | 562234.38 | 2 | 2 | 113393.41 | 362327.31 | 705461.94 | 841086.06 | 560392.25 | 807464.56 | 2 | 885243.44 | 1033214.56 | 941515 | 2 | 2 |
| Q9JHW2 | Nit2 | Omega-amidase NIT2 | 126.72 | 8.33 | 30502 | 2 | 2 | 2 | 1104212.38 | 2 | 553426.75 | 2 | 2 | 2 | 2 | 833111.75 | 2 | 109531.66 | 2 | 2 | 2 | 2 | 2 | 2 | 2 | 2 |
| Q91YP2 | Nln | Neurolysin, mitochondrial | 38.2 | 3.27 | 80429 | 1 | 2 | 2 | 2 | 175156.61 | 2 | 2 | 2 | 2 | 279311.12 | 82296.47 | 2 | 2 | 2 | 2 | 2 | 2 | 2 | 2 | 2 | 2 |
| P19804 | Nme2 | Nucleoside diphosphate kinase B | 414.8 | 57.89 | 17283 | 8 | 16477504 | 32974314 | 30113970 | 36672374 | 76358810 | 39489710 | 34157258 | 67200658 | 41729402 | 41599078 | 78291994 | 77456554 | 52862246 | 47414762 | 24199374 | 40897022 | 33984042 | 23139646 | 41605106 | 9681004 |
| Q9WV85 | Nme3 | Nucleoside diphosphate kinase 3 | 41.05 | 10.06 | 19099 | 1 | 2 | 726311.38 | 678940.88 | 2 | 824940.31 | 2 | 2 | 583023.88 | 2 | 2 | 1487507.75 | 772405.56 | 824459.88 | 883563.06 | 1122998.38 | 826387.12 | 912780.06 | 1063492.5 | 939747.44 | 1005704.38 |
| O70310 | Nmt1 | Glycylpeptide N-tetradecanoyltransferase 1 | 375.74 | 15.93 | 56888 | 7 | 2 | 2343480.5 | 51883.49 | 1748806 | 1846380.5 | 909011.69 | 2507465.25 | 1907166.5 | 1680328.38 | 1181851.88 | 3194476.5 | 3607014.5 | 5019100 | 3980696.5 | 3052037 | 2328499.75 | 1875573.25 | 1204922.5 | 1308099.88 | 4269907 |
| Q61941 | Nnt | NAD(P) transhydrogenase, mitochondrial | 222.31 | 6.08 | 113838 | 5 | 2 | 851316.62 | 922663 | 2 | 1996141.88 | 777503.56 | 1070010.62 | 1855208.75 | 1593687.25 | 1675603.25 | 1501937.25 | 1667900.88 | 399041.75 | 974361.62 | 2 | 2 | 2 | 2 | 944102.06 | 2 |
| Q6VEU1 | Nob1 | RNA-binding protein NOB1 | 38.45 | 1.71 | 46425 | 1 | 2 | 318997.75 | 469258.75 | 448310.91 | 2 | 2 | 2 | 1964381.62 | 2 | 1492998 | 2 | 284762.34 | 456352.16 | 308433.5 | 788490 | 2 | 415186.19 | 870544 | 534841.5 | 710287.06 |
| Q5RJG1 | Nol10 | Nucleolar protein 10 | 36.69 | 1.02 | 80077 | 1 | 685984.56 | 4185082.5 | 12798194 | 1186730.5 | 2 | 1859969.25 | 2191054.75 | 2 | 1105107.5 | 8528444 | 1326282.12 | 1712100.5 | 2 | 650003.81 | 2 | 2 | 591487 | 2 | 2 | 2 |
| Q8BJW5 | Nol11 | Nucleolar protein 11 | 33.93 | 1.11 | 80818 | 1 | 2 | 2 | 2 | 2 | 2 | 2 | 2 | 2 | 2 | 2 | 389395.09 | 2 | 2 | 2 | 2 | 2 | 2 | 2 | 2 | 2 |
| E9Q5C9 | Nolc1 | Nucleolar and coiled-body phosphoprotein 1 | 239.12 | 7.26 | 73698 | 5 | 2 | 2 | 2 | 255217.48 | 2 | 2 | 2 | 2 | 2 | 2 | 2 | 2 | 28727.28 | 2 | 2 | 2 | 2 | 2 | 2 | 2 |
| Q6GQT9 | Nomo1 | BOS complex subunit NOMO1 | 392.61 | 9.64 | 133420 | 9 | 1034975.94 | 944385.38 | 474621.53 | 2092156.62 | 4431672 | 2600783.25 | 1696287 | 3599723.75 | 6492723 | 4274662 | 3476266.75 | 3403310 | 6711838 | 5873648 | 1952774.12 | 5893869.5 | 1860668.62 | 2480465.75 | 6755034 | 2 |
| Q99K48 | Nono | Non-POU domain-containing octamer-binding pro | 760.23 | 36.15 | 54541 | 15 | 5952172.5 | 8559130 | 6863387 | 14518457 | 25633350 | 12459536 | 12254834 | 20248524 | 19570134 | 14443554 | 12016748 | 10664952 | 8988622 | 8115527 | 6231924 | 8070245.5 | 9551312 | 9214575 | 8294721 | 5476815 |
| Q9CQS2 | Nop10 | H/ACA ribonucleoprotein complex subunit 3 | 40.06 | 17.19 | 7706 | 1 | 2 | 2 | 2 | 2 | 2 | 2 | 2 | 2 | 2 | 2 | 2 | 2 | 2 | 2 | 2 | 2 | 2 | 2 | 2 | 2 |
| Q1RP77 | Nop16 | Nucleolar protein 16 | 38.04 | 3.93 | 21117 | 1 | 2 | 557537.31 | 2 | 2 | 719970.44 | 2 | 2 | 2 | 2 | 2 | 2 | 2 | 2 | 2 | 2 | 2 | 462265.91 | 2 | 2 | 2 |
| Q922K7 | Nop2 | 28S rRNA (cytosine-C(5))-methyltransferase | 532.1 | 18.16 | 86752 | 11 | 671921.12 | 1271144.88 | 2034813.5 | 1707568 | 5741872 | 2334714.5 | 1703113.88 | 1488198.62 | 2837061.25 | 2409536.5 | 1949100.62 | 2523757.75 | 2280078 | 2635238.5 | 1395313.75 | 1059510.25 | 1580114 | 1008282.31 | 1636677.75 | 787573.19 |
| Q9D6Z1 | Nop56 | Nucleolar protein 56 | 803.37 | 26.03 | 64464 | 14 | 2101565.5 | 4271047.5 | 7835113 | 2829558.5 | 5662746 | 5811211.5 | 6250350 | 7556915 | 8255882 | 6105906 | 7831085 | 8060398.5 | 10093548 | 10671801 | 12949663 | 11062607 | 8238627.5 | 15962203 | 12282738 | 10925965 |
| Q6DFW4 | Nop58 | Nucleolar protein 58 | 504.86 | 25.37 | 60343 | 9 | 7846229.5 | 10351596 | 10421583 | 8862480 | 7747698.5 | 6737081 | 5246814 | 5930571 | 7066720 | 7190264 | 3015261.75 | 3000159.5 | 6669755 | 4664555.5 | 6073845 | 5083554 | 5514721 | 7353035 | 7480814.5 | 8022738 |
| O35516 | Notch2 | Neurogenic locus notch homolog protein 2 | 45.06 | 0.53 | 265633 | 1 | 679987.81 | 2 | 2 | 482415.34 | 2 | 2 | 2 | 2 | 2 | 368828.97 | 2 | 423561.69 | 2 | 2 | 554936.19 | 2 | 2 | 2 | 2 | 2 |
| O35604 | Npc1 | NPC intracellular cholesterol transporter 1 | 148.6 | 2.74 | 142885 | 3 | 3384036.75 | 3165190 | 2934944.5 | 2 | 2 | 2526286.75 | 985349.5 | 1506283 | 939973.75 | 2137087.5 | 2 | 959943.19 | 1365510 | 561051.31 | 2 | 756610.62 | 1084395.12 | 992818.12 | 1051847.88 | 1476021.88 |
| Q9Z0J0 | Npc2 | NPC intracellular cholesterol transporter 2 | 224.79 | 36.91 | 16442 | 5 | 12622767 | 6676157 | 6984917 | 5793348.5 | 3422078 | 5129394 | 4621308 | 5907170 | 4392889 | 10499966 | 887195.88 | 602002.12 | 2319336.5 | 2532777.5 | 996490.06 | 1730604.62 | 2 | 3035582.75 | 1350598.25 | 2546385 |
| Q6NSR8 | Npepl1 | Probable aminopeptidase NPEPL1 | 67.5 | 2.29 | 55940 | 1 | 2 | 2 | 890333.69 | 317294.38 | 486473.44 | 1083593.38 | 619240.38 | 63026.04 | 2 | 700344 | 1203304.5 | 1178328.88 | 1224975.25 | 1205976.75 | 850227.94 | 1212360.88 | 1614224.62 | 913143.62 | 1589625.5 | 1498088.12 |
| Q11011 | Npepps | Puromycin-sensitive aminopeptidase | 391.25 | 8.04 | 103325 | 7 | 2606534.75 | 3479656 | 2591371.25 | 1110200.12 | 3054948 | 3938569.5 | 2762086.5 | 2776830.25 | 5591779 | 3743977 | 7148321 | 6258970.5 | 5569901.5 | 3661506 | 4574184 | 7530916.5 | 4546659 | 2958738 | 5067262.5 | 2472954.75 |
| P13084 | Npm1 | Nucleophosmin | 747.82 | 43.15 | 32560 | 13 | 41174266 | 63659422 | 10496517 | 66429242 | 71401506 | 58287090 | 51198726 | 17708720 | 61331954 | 66373738 | 30846416 | 39087830 | 26301242 | 19145678 | 7123846 | 35281574 | 49829146 | 24000414 | 52645930 | 31659326 |
| Q9CPP0 | Npm3 | Nucleoplasmin-3 | 110.87 | 13.71 | 19023 | 2 | 24066.06 | 86731.16 | 202408.05 | 174175.86 | 503986.03 | 149515.91 | 2 | 2 | 1036893.06 | 783166.44 | 98479.7 | 734129.12 | 2 | 1297275.12 | 4331080.5 | 2 | 2 | 411783.53 | 1030425.25 | 3974586.25 |
| P97300 | Nptn | Neuroplastin | 142.52 | 8.56 | 44373 | 3 | 2389768 | 1796373.88 | 2844940 | 741108.06 | 1705180 | 1078663.75 | 1014743.19 | 1579725.5 | 916248.19 | 1592566.25 | 2587972.75 | 1781830.25 | 2068171 | 2278914.75 | 1854013.75 | 1921276.5 | 2128710 | 2928447 | 2485916.5 | 1660621.5 |
| Q9JI75 | Nqo2 | Ribosyldihydronicotinamide dehydrogenase [quin | 34.95 | 4.33 | 26248 | 1 | 1507739.25 | 1286004.5 | 1514669.88 | 1112830.5 | 1040020.69 | 2 | 1151715 | 1040033.25 | 578664.38 | 810650.25 | 2 | 2 | 765760 | 2 | 2 | 2 | 2 | 2 | 2 | 2 |
| P08556 | Nras | GTPase NRas | 159.74 | 18.52 | 21199 | 3 | 1598634.25 | 920045.88 | 2732995.5 | 1056122.25 | 2307792.5 | 1264861.12 | 1399229.38 | 229032.36 | 2440603 | 2457809 | 6250669 | 6986914.5 | 7202741 | 7389468 | 4437622 | 7290779.5 | 7415979 | 4871740 | 10011688 | 5187667.5 |
| P97333 | Nrp1 | Neuropilin-1 | 153.85 | 6.28 | 103000 | 3 | 2 | 220197.41 | 933915.88 | 170908.11 | 2 | 2 | 260848.02 | 207796.34 | 627135.19 | 657508.81 | 2 | 2 | 156560.11 | 181517.02 | 294842.81 | 2 | 2 | 2 | 285444.81 | 2 |
| Q07310 | Nrxn3 | Neurexin-3 | 44.63 | 0.44 | 173994 | 1 | 3413184.75 | 3220411 | 4263839 | 2995368.25 | 725782.25 | 3163383 | 2550172.5 | 79437.3 | 2765911.75 | 2900434 | 2427026.5 | 2754336 | 2567139.25 | 2774628.5 | 2382347 | 2829615 | 2048608 | 1577035.25 | 2250702.75 | 1836746.5 |
| Q9R1J0 | Nsdhl | Sterol-4-alpha-carboxylate 3-dehydrogenase, dec | 195.14 | 15.75 | 40686 | 5 | 1230273.62 | 2970973.75 | 1461485.62 | 3171597 | 4235039.5 | 4515402 | 4163744.5 | 5627847.5 | 3793358.25 | 851952.38 | 3807373.75 | 3545444 | 1181602.75 | 3105789.75 | 1074891.38 | 2345853 | 1322283.62 | 655278.81 | 2960217.75 | 11927326 |
| P46460 | Nsf | Vesicle-fusing ATPase | 565.22 | 18.41 | 82613 | 12 | 7657873.5 | 8285418.5 | 8381081 | 9428272 | 12625320 | 8805435 | 8435404 | 11836935 | 12303096 | 10740062 | 5671492 | 3153711 | 4574006 | 6105888 | 4652963.5 | 2919009.25 | 2566295.5 | 5175667.5 | 1291616.12 | 3383299.5 |
| O35987 | Nsfl1c | NSFL1 cofactor p47 | 266.55 | 16.22 | 40680 | 4 | 6355420 | 6909193.5 | 5941815 | 2 | 3606713 | 3227318.5 | 4036164 | 2835947.25 | 830570.81 | 2516012.5 | 9032443 | 8177956 | 7979113.5 | 6836448 | 6052626 | 11469342 | 9245829 | 7270071.5 | 15120555 | 9119804 |
| Q1HFZ0 | Nsun2 | RNA cytosine C(5)-methyltransferase NSUN2 | 212.94 | 6.21 | 85452 | 4 | 70688.98 | 508599.22 | 1565749.5 | 678984.38 | 313668.34 | 187074.52 | 507282.94 | 277288.81 | 724281.88 | 1521588.38 | 94027.89 | 2 | 499443.62 | 326445.47 | 389439.19 | 2 | 238534.92 | 2 | 94189.2 | 452420.09 |
| Q9JM14 | Nt5c | 5'(3')-deoxyribonucleotidase, cytosolic type | 39.97 | 11 | 23076 | 1 | 2 | 2 | 38501.3 | 2 | 131808.41 | 2 | 239008.5 | 2 | 227387.86 | 2 | 125078.05 | 230021.97 | 2 | 291603 | 569095.19 | 2 | 374204.38 | 2 | 2 | 2 |
| Q3V1L4 | Nt5c2 | Cytosolic purine 5'-nucleotidase | 132.61 | 5.71 | 64809 | 2 | 1515223 | 1672743.12 | 445858.28 | 2331259.25 | 2553917.75 | 383764.66 | 904857.81 | 2 | 1295181.5 | 2087271 | 1004060.75 | 890559 | 1177438.5 | 176463.42 | 791015.62 | 706638.12 | 232781.86 | 2 | 2 | 1526216 |
| Q3UFY7 | Nt5c3b | 7-methylguanosine phosphate-specific 5'-nucleoti | 43.52 | 3.67 | 34425 | 1 | 692091.25 | 767129.38 | 767169.94 | 845662.75 | 711989.56 | 953836.19 | 815112.06 | 1141371.62 | 532642.81 | 347319.62 | 2 | 479040.75 | 562263 | 461222.59 | 2 | 1044255.06 | 680609 | 690358.81 | 648058.19 | 2 |
| P54729 | Nub1 | NEDD8 ultimate buster 1 | 35.28 | 1.3 | 70307 | 1 | 2 | 2 | 2 | 690339.56 | 404921.12 | 571554.19 | 2 | 503101.56 | 359818.84 | 2 | 458851.09 | 2 | 2 | 336611.66 | 2 | 2 | 2 | 2 | 2 | 2 |
| Q9R061 | Nubp2 | Cytosolic Fe-S cluster assembly factor NUBP2 | 139.38 | 11.27 | 29518 | 2 | 2 | 2609028 | 1419225.75 | 2461699.75 | 1813998.5 | 1651500.88 | 1862747.12 | 1830092.75 | 1331251 | 1455296 | 678018.75 | 215459.84 | 932983.06 | 1160249.5 | 1476447 | 2 | 307533.03 | 1584409.12 | 2 | 572229.12 |
| Q9CWD8 | Nubpl | Iron-sulfur cluster transfer protein NUBPL | 48.84 | 5.02 | 34139 | 1 | 308651.25 | 134330.14 | 2 | 2 | 2 | 107290.55 | 2 | 2 | 336047.03 | 257373.09 | 2 | 97184.73 | 277398.06 | 275832.91 | 2 | 2 | 220721.98 | 217370.69 | 121157.77 | 2 |
| Q02819 | Nucb1 | Nucleobindin-1 | 136.9 | 5.01 | 53409 | 2 | 419094.25 | 2 | 2 | 2 | 2 | 2 | 2 | 2 | 2 | 121030.16 | 1441771.62 | 2 | 2 | 766947.69 | 2 | 1236979.38 | 629921.75 | 2 | 2 | 2 |
| Q80XU3 | Nucks1 | Nuclear ubiquitous casein and cyclin-dependent k | 115.59 | 4.7 | 26313 | 2 | 841540.81 | 2946578 | 2 | 6189396 | 6537570 | 6868285.5 | 6795753.5 | 1137072.25 | 7727194.5 | 3014442.5 | 2497962.5 | 3140620.5 | 3996151 | 3434787 | 1131963.5 | 7417674 | 6054184.5 | 1793665.12 | 9797593 | 3555864.25 |
| O35685 | Nudc | Nuclear migration protein nudC | 312.6 | 18.07 | 38358 | 5 | 7788831 | 9908784 | 13028934 | 6661603 | 11592858 | 9113671 | 9007952 | 8627614 | 5559434.5 | 5056893 | 12868993 | 12457629 | 9726518 | 6322334 | 9698883 | 15351891 | 12281149 | 10564420 | 12208166 | 9204473 |
| Q5M823 | Nudcd2 | NudC domain-containing protein 2 | 115.66 | 20.38 | 17674 | 2 | 1051107.75 | 1949999.88 | 1185417.38 | 1841919.62 | 2163650.5 | 1158629.88 | 1329509.38 | 1767121.62 | 1317137.5 | 1677809.75 | 2581384.5 | 2214239.5 | 1947480.12 | 2574095.75 | 505924.25 | 926058.31 | 1646833.38 | 1259198 | 1921894.75 | 1759142.38 |

| | | | | | | | | | | | | | | | | | | | | | | | | | | |
|---|---|---|---|---|---|---|---|---|---|---|---|---|---|---|---|---|---|---|---|---|---|---|---|---|---|---|
| Q9D142 | Nudt14 | Uridine diphosphate glucose pyrophosphatase N | 74.95 | 6.31 | 24444 | 1 | 942405.56 | 1043413.38 | 987169.88 | 1168567.5 | 897904.62 | 999632.75 | 946944.75 | 949707.88 | 384442.94 | 541114.06 | 2 | 1353793.88 | 1035960.44 | 888156.12 | 1061899.5 | 1116241.62 | 684872.06 | 2 | 1560464.12 | 2 |
| Q4KM65 | Nudt21 | Cleavage and polyadenylation specificity factor su | 81.15 | 9.25 | 26240 | 2 | 2 | 2 | 2 | 2 | 705783.44 | 2 | 2 | 2 | 445585.47 | 2 | 894096.19 | 1116572.88 | 1154083 | 2 | 2 | 2 | 2 | 847190.19 | 2 | 2 |
| Q6AY63 | Nudt5 | ADP-sugar pyrophosphatase | 52.15 | 5.94 | 24117 | 1 | 2 | 2 | 2 | 1371334.5 | 1749131.88 | 409057.97 | 1397185.25 | 737787.69 | 2542320.25 | 1340891.75 | 1623968.88 | 2434062.75 | 976682.06 | 1842053.25 | 2 | 494997.94 | 2 | 2 | 2 | 2 |
| E9Q7G0 | Numa1 | Nuclear mitotic apparatus protein 1 | 1414.55 | 14.28 | 235630 | 25 | 3593393.75 | 3429121.5 | 3600044 | 2191039 | 7698624.5 | 4351486 | 3739518.5 | 2412948.75 | 4432301 | 5022292 | 4409390 | 5832565 | 3744047 | 4580105.5 | 2815220.75 | 2295605 | 672297.25 | 975291.75 | 841312.12 | 2791614 |
| Q8R0G9 | Nup133 | Nuclear pore complex protein Nup133 | 310.72 | 6.15 | 128620 | 5 | 2 | 34874.67 | 79018.38 | 64902.46 | 161300.06 | 2 | 206402.48 | 185913.55 | 210596.16 | 172307.77 | 2 | 172493.36 | 88707.09 | 244763.22 | 485677.75 | 2 | 213109.69 | 140635.25 | 2 | 137415.5 |
| P49791 | Nup153 | Nuclear pore complex protein Nup153 | 46.7 | 0.89 | 152824 | 1 | 2 | 2 | 2 | 2 | 2 | 2 | 2 | 793971.19 | 2 | 2 | 2 | 2 | 2 | 2 | 2 | 2 | 2 | 2 | 2 | 2 |
| Q99P88 | Nup155 | Nuclear pore complex protein Nup155 | 426.38 | 6.61 | 155118 | 8 | 2844479.25 | 1925003.5 | 3763610 | 2594049.5 | 2756326.25 | 296809.41 | 1425861.5 | 261522.12 | 2878324.25 | 1750553.5 | 2 | 1449881.62 | 2669610.75 | 1959684.12 | 1893327.5 | 2351410 | 2137700.25 | 2951536.5 | 2 | 3793927 |
| Q6ZQH8 | Nup188 | Nucleoporin NUP188 | 63.07 | 0.97 | 196696 | 1 | 2 | 2 | 57791.71 | 122082.61 | 22660.9 | 2 | 143322.39 | 2 | 46350.59 | 25210.57 | 2 | 2 | 36804.91 | 67958.71 | 2 | 2 | 126347.04 | 77798.52 | 2 | 138241.55 |
| Q9QY81 | Nup210 | Nuclear pore membrane glycoprotein 210 | 439.78 | 7.37 | 204101 | 9 | 2703440.75 | 5567009.5 | 3143422 | 4788574.5 | 3050809.5 | 1240138.88 | 894893.5 | 1861990 | 1444105.25 | 1907624.88 | 2759881.25 | 1597330.38 | 2182414.75 | 2765088.25 | 2090787.5 | 1835450.38 | 3898544 | 2243327.75 | 2196633.25 | 777138.12 |
| Q80U93 | Nup214 | Nuclear pore complex protein Nup214 | 275.65 | 3.65 | 212979 | 5 | 2 | 2 | 2 | 1027783 | 129423.88 | 237449.42 | 294252.06 | 170565.58 | 2 | 109381.37 | 1441501.62 | 2044331.25 | 2 | 1301298.12 | 454294.53 | 877029.06 | 1965822.75 | 615442.38 | 2 | 669554.62 |
| Q8R4R6 | Nup35 | Nucleoporin NUP35 | 198.27 | 12.31 | 34786 | 3 | 2 | 2 | 2 | 2 | 2 | 2 | 2 | 2 | 2 | 2 | 685362.75 | 2 | 2 | 2 | 2 | 2 | 2 | 2 | 2 | 2 |
| P59235 | Nup43 | Nucleoporin Nup43 | 95.43 | 6.32 | 41990 | 2 | 377813.41 | 2 | 766325.75 | 211802.06 | 684267.38 | 2 | 103606.5 | 133534.89 | 2 | 418850.41 | 2 | 2 | 339669.91 | 149786.45 | 416597.59 | 2 | 161382.53 | 219139.92 | 2 | 2 |
| Q9JIH2 | Nup50 | Nuclear pore complex protein Nup50 | 158.03 | 13.52 | 49485 | 3 | 2 | 537340.31 | 2 | 2 | 1687895.88 | 2 | 2 | 2 | 2 | 976208.81 | 999445.94 | 861484.19 | 401762.66 | 541318.12 | 2 | 2 | 2 | 2 | 201037.17 | 2 |
| O08658 | Nup88 | Nuclear pore complex protein Nup88 | 83.72 | 5.26 | 83585 | 2 | 2 | 2 | 2 | 2 | 2 | 2 | 2 | 835670.69 | 398414.03 | 2 | 2 | 2 | 2 | 2 | 2 | 2 | 2 | 2 | 2 | 2 |
| Q8BJ71 | Nup93 | Nuclear pore complex protein Nup93 | 383.94 | 11.11 | 93281 | 8 | 3435012.75 | 3433267.75 | 3356963.5 | 7420783 | 6717233 | 5084028 | 5115349 | 5913426.5 | 3918879.25 | 6161950.5 | 2459735 | 3935301.25 | 6979957 | 4786880.5 | 10054025 | 4663047 | 5508412 | 6056732 | 4628489 | 5841586.5 |
| P49793 | Nup98 | Nuclear pore complex protein Nup98-Nup96 | 83.29 | 1.43 | 197283 | 2 | 149928.2 | 233008.95 | 374504.53 | 1436045.88 | 152137.75 | 1608301.75 | 1200354.75 | 113661.39 | 1053695.25 | 712428.44 | 688025.5 | 861806.75 | 490351.75 | 123439.05 | 2 | 905936.06 | 1011776 | 943336.5 | 1328110.62 | 952671.38 |
| P61971 | Nutf2 | Nuclear transport factor 2 | 40.22 | 11.02 | 14478 | 1 | 2 | 2 | 2 | 2 | 742777.56 | 2 | 2 | 2 | 2 | 291484.06 | 2 | 2 | 2 | 631830.31 | 2 | 2 | 2 | 2 | 98591.01 | 2 |
| Q99JX7 | Nxf1 | Nuclear RNA export factor 1 | 211.5 | 10.19 | 70300 | 5 | 1935480.25 | 2519729 | 4152831.25 | 3981001.75 | 2544743 | 1199254.75 | 1218025.25 | 3867346.25 | 1915627.88 | 2740894.25 | 978031.5 | 3011080.75 | 2406943.75 | 2701610.75 | 1539018.5 | 3131565.75 | 2358416.25 | 5100235.5 | 5380102 | 6372539.5 |
| P11928 | Oas1a | 2'-5'-oligoadenylate synthase 1A | 121.82 | 12.81 | 42429 | 2 | 2 | 2 | 2 | 2 | 45419.31 | 2 | 79485.96 | 1193872.62 | 1272459.5 | 150335.98 | 2 | 2 | 2 | 984487.12 | 2 | 2 | 2 | 2 | 1235007 | 1310099.88 |
| E9Q9A9 | Oas2 | 2'-5'-oligoadenylate synthase 2 | 42.44 | 1.75 | 85051 | 1 | 2 | 2 | 2 | 2 | 102116.46 | 2 | 2 | 158746.62 | 165839.62 | 619241.88 | 2 | 2 | 2 | 2 | 2 | 2 | 2 | 2 | 2 | 2 |
| Q8VI93 | Oas3 | 2'-5'-oligoadenylate synthase 3 | 129.75 | 2.46 | 126333 | 2 | 548387.12 | 886411.19 | 64621.36 | 2328875.25 | 1485094.5 | 2157378 | 3110484 | 225363.62 | 4195360.5 | 3181934.5 | 2272383 | 1723366.25 | 2834228.5 | 2425342.75 | 955641.25 | 2243141.5 | 2823789.75 | 2156515.5 | 2854780.25 | 1920162.12 |
| P29758 | Oat | Ornithine aminotransferase, mitochondrial | 1331.04 | 57.4 | 48355 | 23 | 40842934 | 49537634 | 59307922 | 31138366 | 42665434 | 33315120 | 28286018 | 38821046 | 35151654 | 41498530 | 49162994 | 50614194 | 53098798 | 45763550 | 25636910 | 30946742 | 28035332 | 25835918 | 26516438 | 22135850 |
| Q8VIJ5 | Oga | Protein O-GlcNAcase | 38.27 | 0.76 | 102918 | 1 | 2 | 2 | 2 | 2 | 816248.56 | 681065.5 | 2 | 1117934.25 | 889342.88 | 534552.31 | 274561.31 | 2 | 2 | 2 | 2 | 2 | 2 | 2 | 2 | 2 |
| Q60597 | Ogdh | 2-oxoglutarate dehydrogenase complex compone | 418.18 | 11.24 | 116449 | 11 | 3863028 | 2032116.75 | 2784410.5 | 1504350.88 | 5712724.5 | 1659179.75 | 1513783.75 | 1997737.5 | 5940480.5 | 3106547.75 | 5356550.5 | 5083411.5 | 3839231.75 | 5015638 | 3119476 | 1073300 | 7907402.5 | 2180585.75 | 2954958.75 | 4275856 |
| Q99PG2 | Ogfr | Opioid growth factor receptor | 63.97 | 1.9 | 70679 | 1 | 2 | 2 | 2 | 2 | 2 | 2 | 2 | 2 | 2 | 2 | 2 | 2 | 2 | 2 | 2 | 2 | 2 | 2 | 2 | 2 |
| Q8CGY8 | Ogt | UDP-N-acetylglucosamine--peptide N-acetylglucc | 91.18 | 2.39 | 116952 | 2 | 85326.71 | 1005798.06 | 2 | 2128265.25 | 549972.69 | 212257.67 | 2 | 2 | 1387749.38 | 2 | 2 | 1550652.38 | 752956.31 | 887680.12 | 858372.94 | 2 | 2 | 324923.75 | 2 | 2 |
| Q9CZ30 | Ola1 | Obg-like ATPase 1 | 298.42 | 18.18 | 44730 | 6 | 1014600.31 | 1448584.12 | 1996295.88 | 171192.77 | 1320976.62 | 323194.84 | 1056311.38 | 90898.82 | 189629.33 | 1636018.25 | 926354.38 | 3020114 | 2213006 | 2784907 | 2811680.5 | 110582.14 | 1511674 | 281131.78 | 2224502.75 | 281808.66 |
| Q8CI95 | Osbpl11 | Oxysterol-binding protein-related protein 11 | 52.55 | 1.46 | 83629 | 1 | 416497.78 | 100507.46 | 2 | 175955.98 | 161119.97 | 2 | 63810.15 | 2 | 404223.81 | 262420.81 | 2 | 2 | 2 | 109466.55 | 234078 | 2 | 2 | 58109.6 | 2 | 2 |
| B9EJ86 | Osbpl8 | Oxysterol-binding protein-related protein 8 | 527.66 | 16.76 | 101269 | 9 | 110026.85 | 1779189.12 | 509944.56 | 3054255.5 | 921708.69 | 2569616.75 | 2846752 | 2804412.25 | 3090529 | 4268408 | 1048441.5 | 1400429.25 | 3629905.75 | 4391649.5 | 3579515.5 | 1798870.38 | 3554617.75 | 1584848.62 | 2032112 | 2577781 |
| A2A8Z1 | Osbpl9 | Oxysterol-binding protein-related protein 9 | 38.43 | 1.36 | 83109 | 1 | 2 | 2 | 105419.56 | 2 | 62603.75 | 2 | 120871.22 | 123194.48 | 258036.78 | 193560.72 | 64804.46 | 129445.76 | 149615.69 | 320465.44 | 269441.19 | 2 | 2 | 2 | 2 | 2 |
| B0K025 | Ostc | Oligosaccharyltransferase complex subunit OSTC | 138.72 | 18.12 | 16815 | 3 | 510396.69 | 1112166.5 | 606601.94 | 1259073.75 | 2125414.25 | 1691312.62 | 1437856.5 | 2146398.75 | 2076412.38 | 1578513.38 | 1711155.75 | 1498033.75 | 3220349 | 3443229.5 | 3703091 | 1278403.38 | 2915076 | 4676802.5 | 2997446.5 | 3482259.25 |
| Q62422 | Ostf1 | Osteoclast-stimulating factor 1 | 235.59 | 21.4 | 23783 | 4 | 2962240.5 | 2067645.25 | 2780713.25 | 1662546.75 | 3031474.5 | 213686.64 | 1157247.12 | 472047.72 | 790177.5 | 2053079.62 | 266553.31 | 2 | 39743.21 | 504152.06 | 1129318.38 | 1339886 | 494839.66 | 1297546 | 1015108 | 1461530.25 |
| B2RYG6 | Otub1 | Ubiquitin thioesterase OTUB1 | 160.33 | 12.55 | 31270 | 2 | 496002.34 | 244496.12 | 2803553.25 | 274841.38 | 1304025.25 | 261525.73 | 1977450.62 | 3331470 | 378845.41 | 1983997.38 | 97470.73 | 2 | 2 | 117196.27 | 1719709.12 | 192694.19 | 419221.84 | 2 | 313029.22 | 2 |
| Q8BGA9 | Oxa1l | Mitochondrial inner membrane protein OXA1L | 40.65 | 2.31 | 48220 | 1 | 2 | 2 | 2 | 2 | 2 | 2 | 2 | 2 | 2 | 2 | 2 | 538285.62 | 2 | 2 | 2 | 466840.47 | 2 | 436024.38 | 2 | 2 |
| Q9D0K2 | Oxct1 | Succinyl-CoA:3-ketoacid coenzyme A transferase | 1137.63 | 36.54 | 55989 | 14 | 9349066 | 22640766 | 13953954 | 15495253 | 31694602 | 18183842 | 15185151 | 14632419 | 29883904 | 28349076 | 23323632 | 26162342 | 25094824 | 29055628 | 15562708 | 17313588 | 15166585 | 14664057 | 18256178 | 17612826 |
| Q6P9R2 | Oxsr1 | Serine/threonine-protein kinase OSR1 | 90.8 | 2.85 | 58214 | 2 | 2 | 2 | 2 | 2 | 2 | 2 | 2 | 2 | 2 | 2 | 2 | 2 | 2 | 2 | 2 | 2 | 2 | 2 | 2 | 2 |
| P54001 | P4ha1 | Prolyl 4-hydroxylase subunit alpha-1 | 305.15 | 13.86 | 60898 | 7 | 2 | 440912.5 | 2 | 2 | 740835.44 | 640468.5 | 1851657.12 | 2365452.75 | 4016741 | 1446448.88 | 2177391.25 | 1925226.25 | 2424180 | 2810348.25 | 1091896.88 | 686970.69 | 471876.88 | 663695.25 | 881963.75 | 2 |
| Q60716 | P4ha2 | Prolyl 4-hydroxylase subunit alpha-2 | 84.33 | 4.1 | 61002 | 2 | 2 | 2 | 430129 | 2 | 2 | 2 | 2 | 2 | 2 | 2 | 2 | 2 | 2 | 2 | 2 | 2 | 81738.91 | 2 | 2 | 2 |
| P09103 | P4hb | Protein disulfide-isomerase | 1679.4 | 52.65 | 57058 | 29 | 50577538 | 49967894 | 54359506 | 57374046 | 68329114 | 88081394 | 96825682 | 97847258 | 109065314 | 101177146 | 89166498 | 71966010 | 92112898 | 83260802 | 61708542 | 103315386 | 107427386 | 121451682 | 129311530 | 95977914 |
| P50580 | Pa2g4 | Proliferation-associated protein 2G4 | 334.21 | 19.8 | 43699 | 7 | 5627983.5 | 4079364.75 | 7112141.5 | 2878464.75 | 10353240 | 4727260 | 5161891.5 | 7635042 | 7495598.5 | 6016716 | 14057257 | 12952713 | 10665896 | 9313258 | 2711954 | 15302808 | 5976481 | 1562553.62 | 3886416.25 | 5308534 |
| P29341 | Pabpc1 | Polyadenylate-binding protein 1 | 1133.35 | 27.52 | 70671 | 19 | 17003464 | 23714758 | 26511816 | 19715898 | 36027870 | 43818274 | 37915126 | 46506038 | 27175668 | 26457246 | 45567326 | 46405250 | 41954474 | 41867374 | 35921430 | 35897598 | 34819002 | 31573122 | 29173354 | 36303314 |
| Q8CCS6 | Pabpn1 | Polyadenylate-binding protein 2 | 160.33 | 8.94 | 32297 | 2 | 6218787 | 6635827.5 | 4327797 | 5508554 | 5773923 | 4739317 | 4369262.5 | 4392399.5 | 3885396.5 | 3521761.25 | 4770955 | 5481840.5 | 6685650.5 | 5792645.5 | 3656979 | 7092881.5 | 6812095 | 3907402.5 | 7672989 | 8276457 |
| P63004 | Pafah1b1 | Platelet-activating factor acetylhydrolase IB subui | 109.76 | 8.29 | 46670 | 2 | 2 | 2 | 2 | 2 | 735085.19 | 2 | 2 | 1671655.5 | 1066827.25 | 880180 | 2 | 2 | 2 | 2 | 2 | 2 | 2 | 2 | 2 | 2 |
| O35264 | Pafah1b2 | Platelet-activating factor acetylhydrolase IB subui | 41.97 | 3.93 | 25581 | 1 | 2 | 2 | 2 | 2 | 2 | 2 | 2 | 2 | 2 | 2 | 2 | 2 | 2 | 2 | 2 | 2 | 2 | 2 | 2 | 2 |
| O35263 | Pafah1b3 | Platelet-activating factor acetylhydrolase IB subui | 34.72 | 3.88 | 25863 | 1 | 2 | 2 | 2 | 2 | 2 | 2 | 2 | 2 | 2 | 2 | 2 | 2 | 2 | 2 | 2 | 2 | 2 | 2 | 2 | 2 |
| Q9DCL9 | Paics | Bifunctional phosphoribosylaminoimidazole carb | 469.94 | 21.88 | 47006 | 9 | 14474997 | 19582874 | 11411337 | 13808046 | 16349196 | 8767420 | 7278001 | 11442959 | 9847824 | 5499200 | 9892266 | 11057216 | 10352178 | 10319646 | 9228494 | 10439579 | 8700353 | 10463994 | 10819246 | 13701482 |
| Q8CIN4 | Pak2 | Serine/threonine-protein kinase PAK 2 | 95.66 | 7.06 | 57930 | 3 | 887247.06 | 2 | 594629.69 | 2 | 1795783.75 | 2 | 2 | 2 | 371593.53 | 1222251.12 | 2 | 1258953 | 670903.75 | 2 | 2 | 2 | 2 | 493466.41 | 517970.53 | 2 |
| Q9JLB0 | Pals2 | Protein PALS2 | 365.59 | 15.37 | 62631 | 7 | 2 | 751522.5 | 1003583.88 | 2 | 1091163.5 | 2 | 372507.72 | 656551.31 | 2 | 514773.5 | 1599716 | 794558.19 | 1778416 | 2352290.5 | 1738443.5 | 83466.21 | 900074.81 | 1012600.62 | 234731.72 | 700865.12 |
| Q9CQV1 | Pam16 | Mitochondrial import inner membrane translocase | 111.64 | 24.8 | 13785 | 2 | 1370282.5 | 2 | 2 | 121456.81 | 118599.45 | 2 | 2 | 2 | 2 | 2 | 2 | 2 | 77111.23 | 2 | 2 | 2 | 465493 | 2 | 813241.12 | 2 |
| Q99LX0 | Park7 | Parkinson disease protein 7 homolog | 567.31 | 57.14 | 20021 | 11 | 13988259 | 31861672 | 8393376 | 17572122 | 34176890 | 31814870 | 31744100 | 37452638 | 27940690 | 22504070 | 44167254 | 41496178 | 26291974 | 26917028 | 15373049 | 40106114 | 28422362 | 19758638 | 33037338 | 18605024 |
| P11103 | Parp1 | Poly [ADP-ribose] polymerase 1 | 279.62 | 7.9 | 113100 | 6 | 1574567.25 | 2447632.25 | 2 | 2713060 | 5111700.5 | 1682636.38 | 305041.69 | 2524916.5 | 2717907 | 1632349 | 2312777.5 | 3170863.5 | 1809332.62 | 2 | 1498051 | 2715690.5 | 2 | 1846348.88 | 2025480.38 | 2362292.5 |
| Q2EMV9 | Parp14 | Protein mono-ADP-ribosyltransferase PARP14 | 45.15 | 0.55 | 203802 | 1 | 1133401.75 | 1768877 | 1751671.38 | 2284813.75 | 1500462.75 | 2 | 1212867.62 | 991179 | 925143.56 | 870830.75 | 2 | 645806.12 | 2 | 2 | 2 | 2 | 744057.88 | 2 | 2 | 1335922.75 |
| Q8CAS9 | Parp9 | Protein mono-ADP-ribosyltransferase PARP9 | 91.28 | 2.89 | 96659 | 2 | 576932.44 | 2 | 805663.62 | 570944.38 | 2 | 787104.75 | 1719866.75 | 2 | 2 | 1295412 | 2 | 2 | 2 | 2 | 2 | 888141.94 | 662371.5 | 2 | 2 | 2 |
| A2VD12 | Pbxip1 | Pre-B-cell leukemia transcription factor-interacting | 38.65 | 1.39 | 80276 | 1 | 2 | 2 | 2 | 2 | 2 | 2 | 2 | 2 | 2 | 396457.19 | 2311583.5 | 2422568.75 | 569342.62 | 2478134.5 | 2 | 1014602.56 | 2 | 2 | 2 | 2 |
| Q05920 | Pc | Pyruvate carboxylase, mitochondrial | 79.68 | 2.04 | 129685 | 2 | 720510.25 | 932544.06 | 1013782.81 | 812134.06 | 886388.62 | 2 | 2 | 2 | 577331.12 | 912519.88 | 1184702.5 | 2 | 889193.81 | 513888.62 | 725911.75 | 754898.25 | 1866982 | 1025526.12 | 550400.88 | 2 |
| P60335 | Pcbp1 | Poly(rC)-binding protein 1 | 914.61 | 44.38 | 37498 | 11 | 7189898.5 | 11471905 | 8875188 | 15302410 | 19143796 | 7596127 | 9245277 | 9286498 | 13237682 | 12706422 | 56809302 | 55809998 | 47651510 | 54187330 | 43222330 | 42404454 | 43959934 | 39640782 | 48475118 | 43078566 |
| Q61990 | Pcbp2 | Poly(rC)-binding protein 2 | 645.14 | 33.98 | 38222 | 10 | 8579163 | 10682569 | 7399833.5 | 14527761 | 9665021 | 11758707 | 8830597 | 13741613 | 9730234 | 6417999 | 16856112 | 13912013 | 11695288 | 10930077 | 14838982 | 16321079 | 18842326 | 18154494 | 21377176 | 20113088 |
| Q91ZA3 | Pcca | Propionyl-CoA carboxylase alpha chain, mitochor | 194.45 | 5.25 | 79922 | 3 | 1538820.5 | 1222235.88 | 1071203.5 | 1238230.25 | 2696632.5 | 1313718.25 | 624061.94 | 2 | 856412.88 | 930889.69 | 1738957.75 | 1311485.62 | 1654545.25 | 1773950 | 893092.38 | 1363152.5 | 1536218.12 | 2 | 1425026.25 | 1379804 |
| P07633 | Pccb | Propionyl-CoA carboxylase beta chain, mitochonc | 56.28 | 2.4 | 58626 | 1 | 2 | 2 | 2 | 1030450.5 | 363350.25 | 2 | 822586.88 | 2 | 760958.38 | 574476.5 | 633572.44 | 2 | 2 | 711094.56 | 2 | 2 | 2 | 551342.75 | 2 | 2 |
| Q8BH04 | Pck2 | Phosphoenolpyruvate carboxykinase [GTP], mito | 1020.48 | 27.97 | 70528 | 15 | 5983376.5 | 10774934 | 8102139 | 10479106 | 12178573 | 12170727 | 14978576 | 12404062 | 22569730 | 17559356 | 9991925 | 7453014.5 | 7691604 | 8855700 | 11088532 | 5399380 | 4292352.5 | 7754426 | 10258510 | 5620793 |
| P17918 | Pcna | Proliferating cell nuclear antigen | 696.55 | 61.69 | 28785 | 12 | 32127572 | 50497482 | 24616660 | 44302078 | 52772882 | 22289554 | 16664610 | 30678862 | 23167306 | 22648870 | 38112218 | 30734212 | 42172414 | 43367258 | 31061988 | 19661778 | 25512108 | 28288860 | 26294392 | 27639868 |
| Q6P8I4 | Pcnp | PEST proteolytic signal-containing nuclear protei | 36.26 | 6.18 | 18963 | 1 | 2 | 2 | 2 | 2 | 2 | 2 | 2 | 2 | 2 | 2 | 2 | 2 | 2 | 2 | 2 | 2 | 2 | 2 | 2 | 2 |
| Q99ML5 | Pcyox1 | Prenylcysteine oxidase 1 | 165.6 | 8.93 | 56288 | 3 | 1134100.5 | 2 | 2025641 | 1142585.62 | 843089.12 | 452950.34 | 718323.44 | 2 | 1231011.62 | 1978711.25 | 268631.06 | 913788.62 | 1041919.19 | 2 | 2 | 2 | 1275900.25 | 445714.53 | 271415.84 | 2 |
| P19836 | Pcyt1a | Choline-phosphate cytidylyltransferase A | 40.47 | 3.27 | 41681 | 1 | 905380.12 | 1493997 | 2 | 407133.66 | 809231.94 | 669266.81 | 2 | 2 | 469312.44 | 678692 | 786774.94 | 771517 | 913591.88 | 895578.31 | 2 | 2 | 526509.62 | 772386.12 | 931732.5 | 663776.5 |
| Q3UHX2 | Pdap1 | 28 kDa heat- and acid-stable phosphoprotein | 236.36 | 13.81 | 20605 | 4 | 1219113.25 | 2301653.5 | 1015344.62 | 1045551 | 1038081 | 665262.56 | 480949.59 | 2 | 670040.06 | 1550873 | 3009455.75 | 3891707 | 3327132.25 | 2650756.75 | 2129460.25 | 7038349 | 3477568 | 2639276.75 | 6319377.5 | 5136619.5 |
| Q6NX65 | Pdcd10 | Programmed cell death protein 10 | 41.03 | 4.76 | 24355 | 1 | 1234817.62 | 1291813.25 | 2 | 1809742.62 | 1813365.12 | 1741102.75 | 1269829.75 | 1666072.5 | 1017460.75 | 1650318.5 | 2214501.5 | 1820253.75 | 1602667.62 | 1703013.12 | 1147608.5 | 1979338.12 | 1470884.75 | 1947759.5 | 2108413.25 | 1483962.75 |
| Q6NS46 | Pdcd11 | Protein RRP5 homolog | 244.11 | 5.21 | 207779 | 6 | 654198.62 | 2 | 2 | 2 | 1012081 | 2 | 111178.45 | 230436.89 | 1166886.38 | 571055.5 | 2 | 39326.98 | 465552.75 | 342496.91 | 2 | 2 | 2 | 2 | 2 | 2 |
| P56812 | Pdcd5 | Programmed cell death protein 5 | 283.95 | 45.24 | 14275 | 5 | 4278239 | 5088320 | 4791848 | 3295464 | 4944864.5 | 3720682.25 | 2826819 | 3992262.5 | 2060321.25 | 1899999.25 | 4074997 | 4280372 | 4978882.5 | 4372867 | 3892417.5 | 3421918 | 1751363.88 | 1670969.75 | 3356035 | 1459009.75 |
| Q9WU78 | Pdcd6ip | Programmed cell death 6-interacting protein | 388 | 12.08 | 96024 | 8 | 4298912.5 | 4798319 | 6028320 | 6949594 | 9654048 | 3702239.25 | 4922636.5 | 6090495.5 | 8074658.5 | 7503057 | 5527315 | 6692490 | 5380463 | 5656482 | 9739429 | 4852465 | 6617610 | 4678399 | 5473351.5 | 4866468.5 |
| Q4KLJ8 | Pdcl3 | Phosducin-like protein 3 | 61.3 | 5.42 | 27699 | 1 | 470756.84 | 528354.69 | 2 | 2 | 2 | 894612.81 | 2 | 2 | 2 | 2 | 793380.25 | 848323.56 | 1066526 | 2 | 652343.81 | 2 | 1627104.25 | 809335.12 | 2 | 961803.56 |
| Q3TIU4 | Pde12 | 2',5'-phosphodiesterase 12 | 109.48 | 4.28 | 67514 | 2 | 832866.38 | 1793029.12 | 1431433.62 | 1736068 | 1591067 | 1244856.62 | 870585.38 | 2 | 1284037.5 | 480416.62 | 791495.88 | 2 | 1281273.38 | 1047451.25 | 2 | 1304058.5 | 2 | 1009676.31 | 196359.39 | 2 |
| P26284 | Pdha1 | Pyruvate dehydrogenase E1 component subunit a | 173.88 | 8.21 | 43227 | 4 | 2124065.75 | 2638453.25 | 2264161.5 | 1446405.75 | 3208518.75 | 5669742 | 5143121.5 | 5918887.5 | 5296373 | 3923458.25 | 6045716 | 5452364 | 4275858 | 3547269.5 | 2343097.5 | 5181142 | 2607957 | 821036.75 | 2301765 | 2852910 |
| Q9D051 | Pdhb | Pyruvate dehydrogenase E1 component subunit b | 550.99 | 37.05 | 38937 | 11 | 8545996 | 6415369 | 14682358 | 5464046.5 | 8678856 | 4770312 | 7391243.5 | 14244458 | 9830580 | 14092095 | 5142329.5 | 5269537.5 | 11343797 | 8699269 | 13043490 | 2686067 | 7872977.5 | 10649438 | 6215069.5 | 5642120 |
| Q8BKZ9 | Pdhx | Pyruvate dehydrogenase protein X component, m | 69.61 | 1.8 | 53999 | 1 | 2 | 2 | 2 | 2 | 557057.69 | 2 | 2 | 2 | 2 | 756359.25 | 2 | 2 | 2 | 2 | 2 | 2 | 2 | 2 | 2 | 2 |
| P27773 | Pdia3 | Protein disulfide-isomerase A3 | 1728.59 | 52.08 | 56678 | 30 | 46274058 | 73427762 | 45595626 | 75651218 | 148852002 | 65608842 | 63638754 | 90967306 | 96811570 | 104933458 | 101555154 | 114965562 | 105630002 | 93694866 | 91125274 | 117199362 | 112451498 | 124189410 | 137717346 | 112663154 |
| P08003 | Pdia4 | Protein disulfide-isomerase A4 | 1049.01 | 25.55 | 71982 | 19 | 20205682 | 18948084 | 14686639 | 19286506 | 29865004 | 16360232 | 12233110 | 18367202 | 32563942 | 34374790 | 27668612 | 35178070 | 28807680 | 32836478 | 12656018 | 33193024 | 23862432 | 16769168 | 19662718 | 21352046 |
| Q922R8 | Pdia6 | Protein disulfide-isomerase A6 | 1255.95 | 44.09 | 48100 | 18 | 35555226 | 45085994 | 36261886 | 36415470 | 49349074 | 45506250 | 43939450 | 62271818 | 40898754 | 44832474 | 39008742 | 39781246 | 33905958 | 37203306 | 31194090 | 43177426 | 33021762 | 31654232 | 24836526 | 31549068 |
| Q63065 | Pdk1 | [Pyruvate dehydrogenase (acetyl-transferring)] ki | 105.01 | 7.37 | 49081 | 2 | 2 | 215989.14 | 118728.27 | 517383.81 | 46012.16 | 2 | 370242.38 | 256874.77 | 742087.31 | 325002.75 | 81340.93 | 217886.77 | 213656.47 | 507884.19 | 2 | 2 | 2 | 172437.39 | 263730.88 | 350987.78 |
| Q922H2 | Pdk3 | [Pyruvate dehydrogenase (acetyl-transferring)] ki | 210.05 | 9.4 | 47923 | 3 | 1029635.62 | 225883.28 | 1184284.38 | 678109.88 | 1521280.38 | 228447.53 | 377786.03 | 329558.56 | 764602.81 | 1369336.75 | 223168.39 | 204786.39 | 1268328.62 | 448236.06 | 786206 | 2 | 349364.09 | 353272.06 | 295488.41 | 2 |
| P70271 | Pdlim4 | PDZ and LIM domain protein 4 | 79.11 | 10 | 35556 | 2 | 762013.31 | 2 | 571661.62 | 2 | 2 | 2 | 2 | 2 | 2 | 2 | 2086088.5 | 628761.94 | 1267320.62 | 1663455.5 | 2 | 783845.19 | 964347.69 | 866004.75 | 1115855.62 | 2 |
| Q8CI51 | Pdlim5 | PDZ and LIM domain protein 5 | 315.16 | 14.04 | 63299 | 6 | 1473417.75 | 1395924.5 | 1631361.5 | 1202672.5 | 5529548.5 | 2995039.5 | 2495658.5 | 3045218 | 5045248 | 3244772 | 4311322.5 | 2309663.75 | 1564450.12 | 2302756.25 | 1697356.25 | 2998575.5 | 1135041.38 | 1422901.75 | 915056.5 | 1751461.75 |
| O88484 | Pdp2 | [Pyruvate dehydrogenase [acetyl-transferring]]-ph | 37.79 | 2.45 | 59687 | 1 | 2 | 2 | 2 | 2 | 2 | 1590412.62 | 785558.62 | 2 | 647018 | 2 | 2 | 2 | 676004.81 | 2 | 2 | 2 | 2 | 864280.81 | 2 | 2 |
| Q99K01 | Pdxdc1 | Pyridoxal-dependent decarboxylase domain-conta | 289.24 | 8.13 | 87335 | 6 | 3226718 | 4167231 | 1715916.75 | 5473645 | 5192284.5 | 5924211 | 4386927 | 4647655 | 6299152.5 | 4431192 | 3570890 | 4476066.5 | 3871712 | 4641261 | 2 | 2601960.75 | 3974991 | 3705260.5 | 4733271 | 3521713 |
| Q8K183 | Pdxk | Pyridoxal kinase | 361.95 | 20.51 | 35015 | 6 | 267593.94 | 719516.38 | 1869652.38 | 424204.44 | 1673432.25 | 2164964.5 | 988960 | 1063398.75 | 881271.88 | 1430815.75 | 2423773.25 | 11061347 | 1167466.75 | 1256324.75 | 2671489 | 6290456 | 1380633.75 | 1180816.12 | 1053938.25 | 304939.53 |
| P70296 | Pebp1 | Phosphatidylethanolamine-binding protein 1 | 41.44 | 3.74 | 20830 | 1 | 2 | 2 | 2 | 1103057.12 | 2185235.75 | 1938327 | 676842.25 | 1961779.25 | 2475145.25 | 971388.5 | 7972560 | 7046345 | 2641511.5 | 3576372.5 | 1949398.12 | 7590649.5 | 4176174.5 | 1275887.5 | 5716551 | 940269.69 |
| Q9DBD5 | Pelp1 | Proline-, glutamic acid- and leucine-rich protein 1 | 361.42 | 7.39 | 118069 | 6 | 3627519.75 | 123929.17 | 1082051.88 | 570250.25 | 106464.05 | 268073.62 | 155533.33 | 65398.48 | 753434.56 | 587048 | 679895.75 | 726657.81 | 73933.88 | 175110.66 | 708102.62 | 2 | 2 | 2 | 2 | 2 |
| Q11136 | Pepd | Xaa-Pro dipeptidase | 390.26 | 16.84 | 55029 | 8 | 2358463 | 1234343.75 | 2443930 | 2205081.5 | 2710858 | 3079266 | 1888948 | 4525867.5 | 3159498.25 | 1941895.88 | 9243808 | 11072866 | 9342275 | 9774340 | 5483399 | 12431800 | 7240017 | 7095118.5 | 5210721 | 9175137 |
| O09012 | Pex5 | Peroxisomal targeting signal 1 receptor | 38.24 | 1.56 | 70756 | 1 | 2 | 952320.44 | 2 | 560435.81 | 2 | 2 | 688053.62 | 822632.31 | 451417.59 | 531911.44 | 2 | 421378.53 | 2 | 500278.28 | 2 | 545825.06 | 2 | 2 | 2 | 2 |
| Q99LC9 | Pex6 | Peroxisomal ATPase PEX6 | 37.34 | 0.71 | 104549 | 1 | 2 | 2 | 2 | 2 | 2 | 2 | 2 | 2 | 328657.41 | 2 | 2 | 2 | 2 | 2 | 2 | 2 | 1150442.88 | 2 | 683348.06 | 751982.19 |
| Q5SUR0 | Pfas | Phosphoribosylformylglycinamidine synthase | 308.5 | 6.81 | 144629 | 7 | 409542.81 | 1361604.25 | 1063373.5 | 794951.25 | 2936746.5 | 1114571.25 | 2134927.75 | 3284341 | 3773494.75 | 1932248 | 3886950 | 2747511 | 1646310.25 | 1535130.88 | 1560298.25 | 2008202.12 | 1659472.12 | 1088734.5 | 1808356.5 | 1555307.12 |
| Q9CWM4 | Pfdn1 | Prefoldin subunit 1 | 74.71 | 9.02 | 14255 | 1 | 945997.75 | 1228982 | 1352763.88 | 812419.94 | 1181544.62 | 1068609.88 | 714071.69 | 1588355.62 | 742303.69 | 1051657.88 | 1396878.12 | 1075861.75 | 1441953.5 | 1431832.38 | 1282136.38 | 1735089.75 | 1679341.12 | 1845402.12 | 1604080.38 | 1684234.12 |
| B0BN18 | Pfdn2 | Prefoldin subunit 2 | 193.46 | 22.73 | 16580 | 3 | 4691362 | 8842232 | 5882363 | 4027372 | 11437034 | 7186121.5 | 5647021 | 8829196 | 5167741 | 2742321.75 | 7419561 | 6619264.5 | 6759908.5 | 5031352.5 | 3555067.5 | 4453710 | 3917515 | 3270458.25 | 4893114 | 3032608.75 |
| P12382 | Pfkl | ATP-dependent 6-phosphofructokinase, liver type | 491.1 | 14.23 | 85360 | 9 | 2 | 2 | 419227.91 | 883843.38 | 1533069 | 1004874.19 | 1024813.31 | 1293943.62 | 1781454.62 | 1704755 | 1176579 | 2 | 2 | 1275173.38 | 1224010.38 | 1560763.38 | 202817.47 | 942349.19 | 2 | 265834.34 |

| | | | | | | | | | | | | | | | | | | | | | | | | | | |
|---|---|---|---|---|---|---|---|---|---|---|---|---|---|---|---|---|---|---|---|---|---|---|---|---|---|---|
| P47857 | Pfkm | ATP-dependent 6-phosphofructokinase, muscle t | 222.99 | 7.18 | 85269 | 5 | 2 | 1444813.12 | 732410.38 | 2 | 2 | 2 | 2 | 2 | 1502698 | 2 | 2 | 1844911 | 1619420 | 2 | 795472.94 | 2298566.5 | 2 | 1239101.12 | 2 | 2 |
| Q9WUA3 | Pfkp | ATP-dependent 6-phosphofructokinase, platelet t | 1070.22 | 31.25 | 85455 | 19 | 6368792 | 9035235 | 5461969 | 8769986 | 12159441 | 8320066.5 | 10739398 | 6379278 | 20872934 | 14067134 | 12649078 | 12056666 | 6821727.5 | 12661144 | 7788299 | 10101782 | 11022685 | 9456820 | 7690068.5 | 10463173 |
| P62962 | Pfn1 | Profilin-1 | 804.22 | 90 | 14957 | 13 | 79252314 | 32768326 | 86784618 | 57409862 | 115533490 | 55482106 | 54235882 | 56272182 | 119043978 | 132632346 | 215684194 | 222990082 | 254290322 | 243739762 | 166206930 | 180318818 | 192734306 | 169812578 | 300606722 | 163569986 |
| P25113 | Pgam1 | Phosphoglycerate mutase 1 | 1053.95 | 66.93 | 28832 | 17 | 25626050 | 27544418 | 34826566 | 38242630 | 39976770 | 22999904 | 31231770 | 36848210 | 36508406 | 49136986 | 32595258 | 41437122 | 30644190 | 34632266 | 50150574 | 50392662 | 30572500 | 42648314 | 44714130 | 34438178 |
| Q3UUQ7 | Pgap1 | GPI inositol-deacylase | 33.83 | 0.87 | 104578 | 1 | 402658.84 | 222201.59 | 412622.84 | 501497.34 | 2 | 2 | 2 | 373195.94 | 2 | 253155.67 | 460136.16 | 2 | 2 | 2 | 2 | 2 | 2 | 2 | 2 | 2 |
| Q9DCD0 | Pgd | 6-phosphogluconate dehydrogenase, decarboxyla | 1137.22 | 39.34 | 53247 | 16 | 12782790 | 8395389 | 17129916 | 13820970 | 28444772 | 25679624 | 20704334 | 32937018 | 30716806 | 21129190 | 22021976 | 25148976 | 21734756 | 23684590 | 24244554 | 19074980 | 14195999 | 11166898 | 20008164 | 11695309 |
| P09411 | Pgk1 | Phosphoglycerate kinase 1 | 2165.77 | 80.82 | 44550 | 34 | 227320258 | 397612898 | 239521186 | 371089602 | 347306658 | 325138850 | 391030850 | 399217762 | 503853186 | 416113410 | 437983042 | 508480450 | 374712610 | 436070242 | 363656258 | 420627170 | 424346018 | 407293922 | 375536834 | 438048706 |
| P09041 | Pgk2 | Phosphoglycerate kinase 2 | 375.07 | 20.14 | 44853 | 7 | 2 | 2 | 590142.38 | 167856.48 | 398190.06 | 2 | 134215.17 | 2 | 637237.69 | 2104649.25 | 118372.83 | 126471.13 | 2 | 282662.56 | 847059.81 | 2 | 2 | 2 | 2 | 487173.38 |
| Q9CQ60 | Pgls | 6-phosphogluconolactonase | 553.47 | 52.14 | 27254 | 10 | 2178275.5 | 17589190 | 11362668 | 1139900.75 | 12844700 | 5480172 | 9996777 | 19428564 | 6508263 | 11450004 | 11973291 | 11840086 | 15301210 | 19152070 | 13724952 | 15153770 | 5804323 | 14248824 | 17743362 | 13861800 |
| Q9D0F9 | Pgm1 | Phosphoglucomutase-1 | 591.26 | 26.87 | 61418 | 11 | 7292749 | 4873740 | 5766748.5 | 10976661 | 9745134 | 22886132 | 18026906 | 14385056 | 19969004 | 8312206.5 | 10199055 | 10263712 | 6894621.5 | 4074636 | 7488873 | 12366091 | 13400871 | 10648135 | 12714941 | 7059162.5 |
| Q7TSV4 | Pgm2 | Phosphopentomutase | 36.12 | 1.45 | 68748 | 1 | 2 | 920165.56 | 2 | 619279.62 | 1186159.12 | 1537384.25 | 1121818 | 1582924.25 | 1274710.25 | 707350.19 | 1732883 | 1271538.5 | 724002.75 | 814763.19 | 2 | 700435.06 | 539434.69 | 2 | 662032.88 | 2 |
| Q9CYR6 | Pgm3 | Phosphoacetylglucosamine mutase | 50.45 | 2.58 | 59453 | 1 | 2 | 187308.03 | 2 | 2 | 817742.56 | 290657.03 | 320135.66 | 256368.06 | 997460.06 | 926614.19 | 216766.36 | 188505.92 | 968463.69 | 2 | 344710.56 | 2 | 267701.12 | 164650.34 | 2 | 2 |
| D3ZDK7 | Pgp | Glycerol-3-phosphate phosphatase | 76.91 | 6.23 | 34601 | 2 | 3481012.5 | 5406993 | 2056434.75 | 3974929.25 | 2165823 | 4017818.5 | 3958040.25 | 2540595 | 2909955.75 | 3073525.75 | 4661962.5 | 4358883 | 4297605.5 | 3770949.75 | 3122388.75 | 3913421.75 | 5071420 | 4009659 | 5527487.5 | 5240773.5 |
| O55022 | Pgrmc1 | Membrane-associated progesterone receptor con | 172.23 | 11.79 | 21694 | 3 | 2069017.62 | 3620692.5 | 834708.75 | 2937904.25 | 2881487.75 | 2967627.5 | 1810544 | 2493203.5 | 1959879.25 | 2013591.75 | 2867682.75 | 3456126.75 | 2696025.75 | 2899926 | 1380256.12 | 3801852.5 | 2502810.5 | 1800881.75 | 2532538 | 1791035.25 |
| Q80UU9 | Pgrmc2 | Membrane-associated progesterone receptor con | 365.1 | 30.41 | 23334 | 4 | 4683560.5 | 793483.5 | 2345621.5 | 1896798.75 | 3214997.5 | 850664.62 | 2808237.5 | 912833.06 | 4732881 | 3157420 | 944630.88 | 594664.69 | 2085337.25 | 991570.38 | 1052398.5 | 2133332.25 | 2633810 | 280714.31 | 712978.44 | 993360.25 |
| Q8BHF7 | Pgs1 | CDP-diacylglycerol--glycerol-3-phosphate 3-phos | 34.23 | 1.63 | 62489 | 1 | 2 | 2 | 2 | 2 | 2 | 2 | 2 | 2 | 348425.81 | 2 | 2 | 2 | 2 | 2 | 2 | 2 | 2 | 2 | 2 | 2 |
| P67778 | Phb1 | Prohibitin 1 | 1314.22 | 67.28 | 29820 | 19 | 69330498 | 101909794 | 85966818 | 64631486 | 106739970 | 118057946 | 107962170 | 119755090 | 106861570 | 86689658 | 82583370 | 70436642 | 78123618 | 81685866 | 60771638 | 85444770 | 71023506 | 64287310 | 87940866 | 59971614 |
| O35129 | Phb2 | Prohibitin-2 | 1197.8 | 63.55 | 33296 | 18 | 4204098 | 4630888.5 | 7077039.5 | 5518796.5 | 9586686 | 15832965 | 12552542 | 17125624 | 15482124 | 10304798 | 24869354 | 23461662 | 24523416 | 22303114 | 14611208 | 20986556 | 11144225 | 12650030 | 17765512 | 10430989 |
| A6H5X4 | Phf11 | PHD finger protein 11 | 46.2 | 2.97 | 37728 | 1 | 2 | 2 | 2 | 2 | 27233.97 | 1578457.5 | 1755380.12 | 274260.44 | 2489003.75 | 2 | 2 | 2 | 2 | 2 | 2 | 2 | 2 | 2 | 2 | 2 |
| P83870 | Phf5a | PHD finger-like domain-containing protein 5A | 83.85 | 13.64 | 12405 | 2 | 2 | 2 | 2 | 458022.78 | 1439324.5 | 2 | 2 | 2 | 1202640.5 | 2 | 1954096.88 | 2312702.5 | 2 | 1083397.62 | 2 | 1626706.38 | 2 | 2 | 861274 | 2 |
| Q2TBE6 | Pi4k2a | Phosphatidylinositol 4-kinase type 2-alpha | 102.29 | 3.55 | 54258 | 2 | 2 | 2 | 128004.43 | 2 | 65746.88 | 2 | 357005.31 | 353118.53 | 366551.66 | 253222.5 | 2 | 111073.2 | 154080.5 | 332942.16 | 494701.5 | 2 | 368859.53 | 180020.03 | 2 | 384246.22 |
| O55012 | Picalm | Phosphatidylinositol-binding clathrin assembly pro | 587.71 | 19.53 | 69286 | 10 | 5576203 | 5559786 | 9474070 | 9117667 | 7014852.5 | 6268369 | 8449279 | 10896093 | 8993166 | 6869002 | 6981592 | 6337022.5 | 6181658 | 5641189.5 | 8905162 | 6118449.5 | 6654864.5 | 4995789 | 6444259.5 | 3146984.25 |
| Q9CXY9 | Pigk | GPI-anchor transamidase | 120.01 | 5.82 | 44895 | 2 | 91376.36 | 246563.72 | 305433.28 | 519180.94 | 537278.62 | 224865.5 | 587889.62 | 357579.44 | 508400.47 | 643879.88 | 228424.34 | 2 | 637740.44 | 528467 | 838683 | 2 | 549893.62 | 295672.69 | 2 | 2 |
| Q9JJI6 | Pigo | GPI ethanolamine phosphate transferase 3 | 45.06 | 1.1 | 119156 | 1 | 2 | 2 | 2 | 335902.5 | 2 | 2 | 2 | 512773.72 | 2 | 344777.31 | 2 | 2 | 2 | 2 | 2 | 2 | 2 | 2 | 2 | 2 |
| Q6PD26 | Pigs | GPI transamidase component PIG-S | 267.17 | 12.43 | 61711 | 5 | 2 | 2 | 956972.81 | 2 | 2 | 2 | 2 | 1229785.5 | 2 | 73949.04 | 99135.23 | 2 | 103369.73 | 2 | 1338287.88 | 103240.12 | 2 | 2 | 2 | 2 |
| Q8BXQ2 | Pigt | GPI transamidase component PIG-T | 95.37 | 2.58 | 65705 | 2 | 1574084.88 | 287158.5 | 1815589 | 1718117.75 | 446897.41 | 2 | 1341348.12 | 1646968.75 | 1750767.88 | 1035307.06 | 2404539.75 | 3394499.5 | 2091193.38 | 3099209 | 2369450.5 | 2423161.25 | 2590952 | 4439792.5 | 1535301.5 | 3313195 |
| Q9CQJ2 | Pih1d1 | PIH1 domain-containing protein 1 | 51.72 | 4.14 | 32209 | 1 | 2 | 694856.5 | 2 | 2 | 760660.94 | 2 | 2 | 1319348.38 | 397881.25 | 672580.5 | 2 | 2 | 2 | 2 | 2 | 2 | 2 | 2 | 2 | 2 |
| Q9EQ32 | Pik3ap1 | Phosphoinositide 3-kinase adapter protein 1 | 102.01 | 3.45 | 90928 | 2 | 2 | 2 | 2 | 2 | 2 | 2 | 2 | 2 | 2 | 2 | 820055.31 | 673066 | 599191.69 | 566177.69 | 2 | 927813.81 | 2 | 2 | 641006.12 | 2 |
| Q2YFS3 | Pilra | Paired immunoglobulin-like type 2 receptor alpha | 84.66 | 8.61 | 33064 | 2 | 2955074 | 1496703.12 | 2295763.25 | 1268255.88 | 1850229.38 | 783272.12 | 881346.38 | 1627209 | 818339.94 | 1027893.81 | 2 | 2 | 497421.94 | 894678.12 | 722047 | 2 | 776842.81 | 2 | 2 | 958563.19 |
| Q9QUR7 | Pin1 | Peptidyl-prolyl cis-trans isomerase NIMA-interacti | 172.03 | 25.45 | 18370 | 3 | 3226984.5 | 3352919.5 | 1583141.12 | 4008951 | 3981638.25 | 2786202.5 | 2383043.5 | 3290929.75 | 2115351.25 | 2022535.75 | 2872267.5 | 2305417.75 | 1579895.12 | 1694829.88 | 2408473 | 1671303.88 | 1734993.88 | 2682670.75 | 1001731.31 | 2373756 |
| Q4V888 | Pip4p2 | Type 2 phosphatidylinositol 4,5-bisphosphate 4-p | 61.45 | 3.89 | 28024 | 1 | 302460.62 | 2 | 2 | 380009.09 | 2 | 2 | 2 | 993402.19 | 933750.5 | 2 | 1226501.5 | 2 | 450333.47 | 2 | 2 | 1384108.25 | 2 | 2 | 506409.5 | 2 |
| Q8BWR2 | Pithd1 | PITH domain-containing protein 1 | 55.81 | 4.27 | 24192 | 1 | 2 | 2 | 2 | 2 | 2 | 2 | 2 | 2 | 2 | 2 | 933272.75 | 675917 | 562797 | 530101.88 | 2 | 2 | 2 | 2 | 2 | 2 |
| Q8K411 | Pitrm1 | Presequence protease, mitochondrial | 110.34 | 2.99 | 117372 | 2 | 908125.81 | 2074356.62 | 1208154.12 | 719718.44 | 2553163 | 1729934.12 | 1228628.75 | 2436045.5 | 2108236.5 | 1432979 | 1529431.75 | 1815353.38 | 1346582.5 | 1333338.88 | 668219.38 | 1224011.25 | 971409.81 | 870617.69 | 1240961 | 656224.81 |
| P53657 | Pklr | Pyruvate kinase PKLR | 156.7 | 5.75 | 62309 | 3 | 2 | 2 | 2 | 2 | 359958.94 | 2 | 2 | 2 | 2 | 2 | 2 | 2 | 2 | 2 | 2 | 17672884 | 4663591 | 4463233 | 3083498.5 | 2704733.5 |
| P52480 | Pkm | Pyruvate kinase PKM | 3336.56 | 76.08 | 57845 | 48 | 265479794 | 313056034 | 314399938 | 377684034 | 530355650 | 555021250 | 438330754 | 533714658 | 814649730 | 475482882 | 600597442 | 616932738 | 461517698 | 436891810 | 475698498 | 419376290 | 449027042 | 441798786 | 424795906 | 389210114 |
| P70268 | Pkn1 | Serine/threonine-protein kinase N1 | 79.64 | 2.22 | 104411 | 2 | 2 | 2 | 1488213 | 727953.06 | 1507762.12 | 1810920.38 | 2 | 2 | 1521140.25 | 1026922.88 | 2 | 209705.02 | 2006564.25 | 1293472.12 | 1020926.06 | 2 | 1847208.5 | 1542486.38 | 2 | 2259428 |
| Q8VEB4 | Pla2g15 | Lysosomal phospholipase A and acyltransferase | 40.73 | 2.67 | 47307 | 1 | 1842629.25 | 868697.25 | 1126804 | 467568.41 | 663367.69 | 2 | 399326.03 | 591773.44 | 546696.56 | 1284477.62 | 1051548.75 | 569679.25 | 699566.94 | 528462.25 | 2 | 2 | 2 | 866038.25 | 2 | 2 |
| P47713 | Pla2g4a | Cytosolic phospholipase A2 | 281.84 | 9.09 | 85222 | 5 | 508832.31 | 270046.25 | 158363.16 | 1828703 | 3584506.25 | 1588120.75 | 1409302.88 | 1053498.38 | 3929039.5 | 2927304 | 1049865 | 2511754.5 | 2245282 | 3288934.25 | 1282747.75 | 1431159.5 | 1051587 | 767156.5 | 1670120 | 793289.38 |
| Q62028 | Pla2r1 | Secretory phospholipase A2 receptor | 47.98 | 0.47 | 170512 | 1 | 2 | 2 | 378215 | 2 | 2 | 876966.5 | 327980.75 | 87302.61 | 2 | 2 | 2 | 326282.44 | 2 | 2 | 983210.12 | 2 | 628273.5 | 621692.5 | 2 | 2 |
| P27612 | Plaa | Phospholipase A-2-activating protein | 300.94 | 10.2 | 87221 | 6 | 1874684.12 | 2615806.75 | 108733.91 | 2180039.25 | 5781153 | 2962187.25 | 3927228 | 1499988.25 | 4444860.5 | 3944220 | 1101429.5 | 1997044.25 | 3645714.75 | 2112337 | 2392745 | 1248300.75 | 3384008.5 | 2996645.75 | 1107571.75 | 629622.75 |
| Q3TCN2 | Plbd2 | Putative phospholipase B-like 2 | 268.35 | 11.62 | 66289 | 6 | 4626621 | 5130918 | 3536852.5 | 3207683.5 | 2036128.38 | 1094195 | 2908589 | 4084536.5 | 1889537 | 3595258.5 | 121655.59 | 1982405.62 | 853360 | 1955544.25 | 1479673 | 988797.69 | 1829703.88 | 2424880.5 | 1195682.38 | 1056225.12 |
| P51432 | Plcb3 | 1-phosphatidylinositol 4,5-bisphosphate phospho | 77.57 | 1.62 | 139487 | 2 | 742456.19 | 2 | 473975.16 | 444167.22 | 783945.44 | 2 | 2 | 2 | 935915.69 | 702678.5 | 781814.44 | 201507.27 | 847202.69 | 468566.19 | 317417.66 | 2 | 249369.34 | 165602.25 | 2 | 2 |
| P24135 | Plcg2 | 1-phosphatidylinositol 4,5-bisphosphate phospho | 274.47 | 5.22 | 147735 | 7 | 4303650 | 3823530 | 3894998.25 | 5422433 | 3406758 | 4500865 | 3818338 | 5028178.5 | 3721349.75 | 3889987 | 6335742 | 5138561.5 | 6521321 | 6191456 | 2060000.5 | 6175159.5 | 2938549.5 | 7465787 | 4436234 | 3053894.5 |
| O35405 | Pld3 | 5'-3' exonuclease PLD3 | 175.25 | 7.99 | 54389 | 3 | 1079705.75 | 595689.5 | 1758101.75 | 392022.69 | 143526.12 | 186915.67 | 594245.38 | 1447498.5 | 794675.75 | 1076048 | 1047362.31 | 1030773.69 | 1195501.62 | 1258820.25 | 2503439.75 | 2 | 1165112.38 | 1221350.38 | 322142 | 215404.11 |
| Q8BG07 | Pld4 | 5'-3' exonuclease PLD4 | 943.83 | 38.57 | 56154 | 14 | 56814230 | 54365838 | 48080822 | 49538466 | 42085726 | 28968542 | 28506038 | 28419456 | 26219874 | 40881506 | 17161362 | 14213955 | 16546519 | 14629944 | 27383632 | 21365194 | 19312210 | 24940534 | 18028056 | 20254480 |
| Q9QXS1 | Plec | Plectin | 8475.192 | 35.11 | 534188 | 158 | 153196322 | 204164002 | 174188370 | 155112034 | 181612210 | 172335250 | 185923570 | 200290306 | 184138690 | 186317890 | 305333026 | 306802178 | 221547954 | 235857154 | 181503538 | 277993634 | 201669618 | 196836050 | 215754002 | 184499410 |
| Q9JHK5 | Plek | Pleckstrin | 132.91 | 11.43 | 39901 | 2 | 1936574.12 | 1517567.75 | 2170821.75 | 1248291.75 | 1614609.25 | 1034292.81 | 1114434 | 474391.03 | 1622951.12 | 1507921 | 2032932.25 | 2026564.75 | 2783391 | 2109337.75 | 1255649.12 | 1591819 | 1059582.88 | 923974.12 | 4350481 | 3066091.25 |
| Q9ERS5 | Plekha2 | Pleckstrin homology domain-containing family A | 118.21 | 4.24 | 47380 | 2 | 154474 | 90654.02 | 45472.22 | 2 | 2 | 2 | 84144.96 | 2 | 353514.25 | 65669.27 | 197869.31 | 166311.42 | 117884.27 | 277125.81 | 2 | 87164.17 | 227433.31 | 102783.66 | 193058.98 | 194866 |
| Q8K124 | Plekho2 | Pleckstrin homology domain-containing family O | 178.98 | 8.89 | 53872 | 4 | 2390804 | 1108289.25 | 2673688.5 | 1186334.38 | 1175136.38 | 963586.19 | 1619587.5 | 3500916 | 1999446.25 | 1610797.88 | 4514432 | 1570026.25 | 2586877.5 | 3072915.25 | 3755915 | 3855035.25 | 4550292 | 1526125.38 | 3916472.5 | 2741792.5 |
| P20918 | Plg | Plasminogen | 82.06 | 3.45 | 90808 | 2 | 2 | 2 | 2 | 2997760.25 | 3261242.5 | 2 | 348425.78 | 2 | 2211635.25 | 1750022.88 | 575520.38 | 520224.81 | 740457.06 | 534301.12 | 607399.06 | 2 | 2 | 2 | 2 | 2 |
| D4ACN8 | Plgrkt | Plasminogen receptor (KT) | 38.91 | 6.8 | 17312 | 1 | 2 | 2 | 2 | 2 | 2 | 2 | 2 | 2 | 2 | 2 | 848333.56 | 785997.94 | 851360.81 | 647160.12 | 676889.94 | 679351.12 | 914791.88 | 899959.94 | 2 | 2 |
| P43883 | Plin2 | Perilipin-2 | 693.1 | 30.82 | 46646 | 11 | 7952999.5 | 18255014 | 10925554 | 4999728 | 2460628.75 | 26153950 | 18072512 | 26531128 | 27089402 | 18447142 | 9148442 | 10886130 | 10005744 | 9148190 | 10912864 | 11215546 | 15162728 | 12384024 | 6441545 | 11350985 |
| Q9DBG5 | Plin3 | Perilipin-3 | 855.54 | 45.31 | 47262 | 13 | 4867381 | 10832336 | 5951303 | 6436827 | 8322063.5 | 10788607 | 11496471 | 8265224.5 | 10891878 | 11839705 | 17024450 | 16686097 | 12075924 | 11815219 | 10385306 | 9681559 | 4586707.5 | 6326307 | 7333763 | 5236185 |
| Q63321 | Plod1 | Procollagen-lysine,2-oxoglutarate 5-dioxygenase | 83.38 | 2.47 | 83612 | 2 | 2 | 760481.5 | 2 | 2122437 | 714591.88 | 578943.94 | 1061125.25 | 1354539.38 | 341372.34 | 501073 | 1747974.62 | 553952.88 | 1508950.75 | 772456.62 | 340353.5 | 1133709.38 | 1707257.88 | 2 | 1623814.75 | 1124164.62 |
| Q9R0E1 | Plod3 | Multifunctional procollagen lysine hydroxylase an | 297.73 | 9.58 | 84922 | 6 | 4204454.5 | 2805975.5 | 1545673.75 | 1118815.12 | 2353309 | 1121041.88 | 2 | 2 | 2 | 1630089.38 | 1515509.25 | 876253.62 | 1798774 | 912037.06 | 928830.56 | 1355396.25 | 1333540.88 | 2471295 | 951728.44 | 2115968.5 |
| Q6P742 | Plp2 | Proteolipid protein 2 | 63.86 | 9.93 | 16557 | 1 | 2449469.5 | 596483.69 | 2 | 911144.94 | 409288.03 | 1015453.69 | 3148291.25 | 623732.06 | 2781099 | 3128299.25 | 2 | 2 | 1233851.25 | 1059886.25 | 831151.69 | 2 | 1651425.38 | 606239.81 | 2 | 790728.88 |
| Q9Z2Y8 | Plpbp | Pyridoxal phosphate homeostasis protein | 153.33 | 12.77 | 30049 | 3 | 345823.75 | 1837845.75 | 1652463.25 | 1237484.5 | 2194880 | 8598723 | 2182025.25 | 2195555.5 | 2470676.5 | 1027435.31 | 3763992 | 1197994.25 | 2274218.25 | 1911152.12 | 3086583 | 1995712 | 2510092.75 | 3165717.5 | 1487779.88 | 1993268.25 |
| Q922V4 | Plrg1 | Pleiotropic regulator 1 | 240.81 | 10.33 | 56938 | 4 | 2 | 1244245.25 | 1708232.75 | 1298942.62 | 3967316.5 | 8163664 | 4573766 | 811640.62 | 11399818 | 1950074 | 5454146 | 4511902.5 | 2822520.25 | 4339747 | 2 | 5021983.5 | 1196624.88 | 2 | 1254028.12 | 2 |
| Q99K51 | Pls3 | Plastin-3 | 1016.9 | 30.16 | 70742 | 18 | 10341398 | 6729090 | 10663873 | 5444543.5 | 8387411 | 10619362 | 6979858 | 8863856 | 8689738 | 7294023.5 | 9309210 | 7224744 | 7947178 | 5756852 | 7091422.5 | 3008789.25 | 4436293 | 5091728.5 | 3274246.25 | 3245200.25 |
| Q91ZV7 | Plxdc1 | Plexin domain-containing protein 1 | 39.14 | 3 | 55635 | 1 | 26710.71 | 75077.91 | 40779.4 | 70878.9 | 22222.28 | 2 | 2 | 2 | 128973.68 | 142099.48 | 2 | 47849.65 | 51326.72 | 60369.5 | 194243.91 | 2 | 2 | 2 | 2 | 2 |
| P70206 | Plxna1 | Plexin-A1 | 439.46 | 6.23 | 211099 | 9 | 1883346.38 | 1823904.75 | 568314.38 | 2087547.75 | 3131737 | 3010733.75 | 4093774.75 | 2306569.25 | 5494484 | 4387692 | 1498579.25 | 2053406.5 | 1396579.25 | 1438685.25 | 2462936.25 | 2377422 | 3818821.75 | 4553284 | 948648.06 | 2602914 |
| B2RXS4 | Plxnb2 | Plexin-B2 | 500.43 | 6.03 | 206230 | 9 | 10473187 | 11143706 | 9682170 | 5885522.5 | 8422308 | 2396757 | 632654.44 | 5870254 | 5480084 | 5764672.5 | 3756564 | 3685656 | 4858579 | 3459083 | 3609338.75 | 3586604.25 | 5118156 | 3967404.75 | 3771416.75 | 2162288.75 |
| Q60953 | Pml | Protein PML | 43.25 | 1.02 | 98242 | 1 | 2 | 2 | 2 | 2 | 2 | 2 | 2 | 2 | 2 | 2 | 2 | 2 | 2 | 2 | 2 | 2 | 2 | 2 | 2 | 2 |
| O35621 | Pmm1 | Phosphomannomutase 1 | 38.26 | 2.67 | 29775 | 1 | 619526.06 | 2 | 2 | 2 | 2 | 661993.06 | 2 | 2 | 464868.34 | 2 | 613748 | 2 | 499296.75 | 2 | 494552.5 | 554783.81 | 771198 | 2 | 2 | 2 |
| Q9DC61 | Pmpca | Mitochondrial-processing peptidase subunit alpha | 133.92 | 5.92 | 58279 | 3 | 2249624.5 | 2601690.25 | 1871232.12 | 882627.12 | 1147806.62 | 797229.44 | 930792.31 | 780810.69 | 676214.31 | 1555589 | 2508451.5 | 1597618.25 | 968011 | 1665041.25 | 1346850.12 | 1116874.38 | 956013.69 | 2299214 | 1933918.75 | 1085265.62 |
| Q9CXT8 | Pmpcb | Mitochondrial-processing peptidase subunit beta | 211.93 | 9.82 | 54614 | 5 | 619957.06 | 1742091.5 | 3050756 | 1141721 | 1606759.25 | 2186476.25 | 358152.22 | 1935073.5 | 1211046.88 | 1303711 | 1665646.25 | 2567486 | 1003687.94 | 1922634.25 | 1572201.62 | 6424706 | 2619756 | 2061081.25 | 1795817.25 | 2657669.25 |
| Q9D1G2 | Pmvk | Phosphomevalonate kinase | 95.74 | 15.1 | 21916 | 2 | 2 | 2 | 2 | 2 | 2 | 2 | 2 | 142191.52 | 2 | 96823.24 | 2 | 543631.31 | 2 | 2 | 2228994.25 | 2 | 2 | 2 | 275534.38 | 2 |
| Q9JLV6 | Pnkp | Bifunctional polynucleotide phosphatase/kinase | 184.86 | 7.85 | 57223 | 3 | 670270.62 | 5504760 | 4020498.75 | 680191.31 | 4468375 | 1103074.25 | 6121685.5 | 750444.69 | 2338729.25 | 2523907 | 2865964.5 | 3425538 | 1307580.5 | 4176894.75 | 2 | 152328.73 | 6131788 | 127947.99 | 8544915 | 7797416.5 |
| O35691 | Pnn | Pinin | 167.48 | 4.28 | 82436 | 4 | 2 | 2 | 2 | 2 | 2 | 2 | 2 | 2 | 2 | 2 | 889940.25 | 1730853 | 1276547.75 | 653501.44 | 97283.85 | 414289.41 | 2 | 2 | 2 | 2 |
| Q9CPS7 | Pno1 | RNA-binding protein PNO1 | 34.19 | 3.23 | 27454 | 1 | 870966.19 | 1004988.75 | 2 | 1316330 | 683802.56 | 873642.06 | 2 | 2 | 986574.62 | 2 | 2 | 958736.81 | 1019934.94 | 1029342.62 | 765262.19 | 1039361.69 | 841400.62 | 1070803.38 | 1064584.62 | 2 |
| P23492 | Pnp | Purine nucleoside phosphorylase | 516.38 | 27.34 | 32277 | 7 | 4584734.5 | 8090226.5 | 6241757.5 | 8953700 | 10254576 | 14502430 | 9353715 | 13056587 | 10754311 | 9947476 | 6851702.5 | 7320888 | 3969662.5 | 5085670.5 | 4689308 | 3623757.5 | 4002651 | 2798525.5 | 1567948.25 | 2028840 |
| Q8K1N1 | Pnpla8 | Calcium-independent phospholipase A2-gamma | 51.26 | 1.29 | 87381 | 1 | 2 | 2 | 2 | 2 | 2 | 2 | 2 | 2 | 2 | 2 | 2 | 2 | 2 | 2 | 2 | 2 | 2 | 2 | 2 | 2 |
| Q8K1R3 | Pnpt1 | Polyribonucleotide nucleotidyltransferase 1, mitoc | 282.18 | 8.43 | 85683 | 5 | 1195536.75 | 3766901.75 | 2314816.5 | 972264 | 2835244.5 | 2 | 2 | 4555738 | 5186957 | 2233910.5 | 2568136 | 1354310.88 | 2942513 | 2415482.25 | 357533.09 | 2492633.5 | 731421.5 | 2 | 3704071.5 | 562865.44 |
| Q91ZW2 | Pofut1 | GDP-fucose protein O-fucosyltransferase 1 | 35.79 | 1.78 | 44688 | 1 | 1065915.38 | 1118762.25 | 1171882.75 | 1373548.62 | 822921.62 | 800131.31 | 841705.38 | 1305564.38 | 913714.69 | 649993.69 | 980727 | 2 | 1097461.5 | 721983.75 | 1155974.25 | 2 | 2 | 718511 | 2 | 984626.44 |
| Q8VHI3 | Pofut2 | GDP-fucose protein O-fucosyltransferase 2 | 104.15 | 5.59 | 49429 | 2 | 1024450.75 | 721606.5 | 2 | 2 | 794164.88 | 1599724.5 | 498364.41 | 2 | 441260.75 | 618045 | 289813.62 | 484546.19 | 2 | 195249.16 | 2 | 2 | 2 | 426603.72 | 2 | 973386 |
| Q9JHP7 | Poglut2 | Protein O-glucosyltransferase 2 | 161.98 | 5.18 | 57985 | 3 | 2 | 184431.91 | 2 | 308016.38 | 56898.8 | 2 | 2 | 2 | 521797.5 | 2 | 2 | 300594.38 | 2 | 2 | 92300.77 | 2 | 2 | 2 | 2 | 131641.62 |
| O54747 | Pold1 | DNA polymerase delta catalytic subunit | 37.36 | 1.36 | 123601 | 1 | 2 | 2 | 2 | 2 | 25851.19 | 2 | 2 | 2 | 2 | 2 | 2 | 2 | 24727.25 | 43189.79 | 2 | 2 | 2 | 2 | 2 | 2 |
| P08775 | Polr2a | DNA-directed RNA polymerase II subunit RPB1 | 271.52 | 3.91 | 217176 | 5 | 3824223 | 4547184 | 2507249.5 | 2867118.5 | 2787831.5 | 3549590 | 3053612.25 | 1545788 | 3069191.75 | 589021.31 | 4607134 | 2619110.5 | 3131937.5 | 3171574.25 | 2045139.38 | 844289.19 | 1892053.12 | 1896999.5 | 2354758.75 | 1318019.25 |
| Q8CFI7 | Polr2b | DNA-directed RNA polymerase II subunit RPB2 | 137.96 | 3.58 | 133911 | 3 | 2 | 2 | 2 | 2 | 1637882 | 864054.31 | 80357.77 | 1223832.38 | 1152134.5 | 55622.75 | 1094395.25 | 829454.75 | 2 | 624397.5 | 2 | 557664.38 | 2 | 2 | 2 | 2 |
| B0BNE2 | Polr2e | DNA-directed RNA polymerases I, II, and III subu | 98.06 | 10.48 | 24570 | 2 | 2 | 2 | 2 | 2 | 2 | 2 | 2 | 2 | 359162.84 | 2 | 2 | 257560.91 | 2 | 248701.08 | 2 | 2 | 2 | 2 | 2 | 460785.75 |
| O88828 | Polr2f | DNA-directed RNA polymerases I, II, and III subu | 53.12 | 5.51 | 14464 | 1 | 914052.62 | 829712.5 | 689233.25 | 971178.25 | 1819848.88 | 1240953.25 | 480344.19 | 2 | 766695.25 | 737399.44 | 660511.19 | 2 | 1061637 | 618952.5 | 354712.09 | 978393.94 | 915123.88 | 1078439.38 | 790975.56 | 2 |
| Q923G2 | Polr2h | DNA-directed RNA polymerases I, II, and III subu | 38.82 | 8.67 | 17143 | 1 | 815084.25 | 2 | 2 | 735033.12 | 778811.06 | 2 | 653643.44 | 2 | 557196.5 | 2 | 2 | 833519.56 | 434692.12 | 734763.81 | 2 | 1412045.75 | 2 | 2 | 2 | 651377.19 |
| Q62087 | Pon3 | Serum paraoxonase/lactonase 3 | 106.99 | 7.63 | 39351 | 2 | 396417.75 | 293665.19 | 2 | 2 | 2 | 2 | 308404.81 | 2 | 2 | 2 | 3272392.75 | 2565963.25 | 3093448.25 | 2663525 | 1699117 | 3400310.5 | 3397370 | 3893108 | 1907119.12 | 3518726.25 |
| P37040 | Por | NADPH--cytochrome P450 reductase | 375.16 | 15.34 | 77044 | 7 | 2 | 562183.31 | 790266.62 | 1088527.62 | 590667.19 | 286093.16 | 714452.69 | 326759.44 | 2286038.25 | 740948.25 | 398336.72 | 214818.39 | 2 | 231683.83 | 2 | 1015967.88 | 1212998.25 | 2 | 450412.25 | 221913.64 |
| Q91YU8 | Ppan | Suppressor of SWI4 1 homolog | 150.28 | 6.38 | 52756 | 3 | 325523.5 | 2 | 2 | 2 | 2 | 2 | 2 | 2 | 110003.09 | 2 | 2 | 386207.19 | 2 | 2 | 2 | 2 | 679165.5 | 2 | 2 | 712284.56 |
| P35433 | Ppat | Amidophosphoribosyltransferase | 37.89 | 1.35 | 57437 | 1 | 1384687.88 | 1062445.5 | 918004.81 | 862753.44 | 1043741.19 | 2 | 2 | 2 | 2 | 2 | 2 | 2 | 682479.81 | 586809.25 | 2 | 2 | 580160.31 | 2 | 2 | 2 |
| P17742 | Ppia | Peptidyl-prolyl cis-trans isomerase A | 928.91 | 82.32 | 17971 | 14 | 1135532.38 | 3290772.5 | 4888471 | 468504.22 | 21764478 | 11251121 | 7452503 | 12404534 | 27336840 | 19206822 | 131987122 | 134021498 | 91648154 | 97204114 | 55781694 | 57371822 | 65268026 | 36009546 | 63481854 | 35583158 |
| P24369 | Ppib | Peptidyl-prolyl cis-trans isomerase B | 316.73 | 31.48 | 23713 | 6 | 7769122 | 6933921 | 12863575 | 5568526 | 22130758 | 6136789.5 | 4051082 | 6492822.5 | 9515420 | 9540926 | 15113707 | 17705898 | 19151576 | 4587636.5 | 7607944.5 | 12274179 | 8808842 | 7396565 | 7465452 | 4026972.5 |
| Q6DGG0 | Ppid | Peptidyl-prolyl cis-trans isomerase D | 125.38 | 10.54 | 40766 | 3 | 732442.44 | 710136.88 | 924049.94 | 309959.06 | 64535.71 | 168320.97 | 2 | 2 | 2 | 382523.91 | 189466.89 | 114689.88 | 2 | 233628.78 | 682167.12 | 178091.84 | 472612.78 | 314849.12 | 2 | 148976.67 |
| Q9D868 | Ppih | Peptidyl-prolyl cis-trans isomerase H | 68.2 | 13.3 | 20464 | 1 | 32778.46 | 2 | 70606.97 | 2 | 2 | 2 | 2 | 69799.09 | 2 | 2 | 2 | 2 | 2 | 206352.28 | 2 | 2 | 2 | 86650.09 | 2 | 2 |
| P20650 | Ppm1a | Protein phosphatase 1A | 53.36 | 3.14 | 42417 | 1 | 2 | 2 | 682668.69 | 1824501.88 | 2148924 | 3770942.5 | 2 | 2 | 1408042.38 | 2 | 2391525.25 | 2 | 1670634 | 2 | 989956.06 | 2073483.38 | 1173271.12 | 2 | 2 | 768097.44 |
| Q8CGA0 | Ppm1f | Protein phosphatase 1F | 32.43 | 2.21 | 49611 | 1 | 2 | 2 | 2 | 2 | 2 | 2 | 2 | 2 | 2 | 2 | 482463.47 | 2 | 597660.94 | 492511.5 | 2 | 2 | 2 | 2 | 630505.94 | 606134.81 |
| Q61074 | Ppm1g | Protein phosphatase 1G | 194.93 | 8.49 | 58728 | 4 | 793883.62 | 930971.12 | 1206803 | 2099520 | 3119217 | 1102392.88 | 1595910.12 | 375021.78 | 2101913.75 | 1182253.62 | 1340040.75 | 1493126.38 | 1612928.75 | 400128.03 | 250757.61 | 131248.95 | 1172242.75 | 255132.59 | 905379.5 | 2 |

| | | | | | | | | | | | | | | | | | | | | | | | | | | |
|---|---|---|---|---|---|---|---|---|---|---|---|---|---|---|---|---|---|---|---|---|---|---|---|---|---|---|
| P51175 | Ppox | Protoporphyrinogen oxidase | 70.99 | 4.61 | 50871 | 2 | 1058248.62 | 925482 | 1038461.44 | 637701.81 | 963977.25 | 660151.5 | 876121 | 1445507 | 1811251.88 | 1042888.06 | 813688.88 | 621901.62 | 1316180 | 1480951.12 | 1246317 | 1113153.25 | 1096946.38 | 1272297.5 | 1298906.5 | 999821.31 |
| P62137 | Ppp1ca | Serine/threonine-protein phosphatase PP1-alpha | 409.08 | 22.42 | 37540 | 8 | 2 | 2 | 2 | 139435.06 | 243394.22 | 2 | 124146.74 | 122524.17 | 433840.66 | 2 | 265775.72 | 265987.12 | 897642.12 | 289605.16 | 743040.25 | 247912.31 | 420138.66 | 271423.16 | 240369.12 | 287153.88 |
| P62141 | Ppp1cb | Serine/threonine-protein phosphatase PP1-beta c | 359.73 | 21.1 | 37187 | 7 | 4974943 | 6365495 | 2370274.5 | 3601053.25 | 875571.62 | 4532223 | 3671527.25 | 329918.56 | 3350344.75 | 3161222.75 | 4209329.5 | 4301273 | 3509828.25 | 3266190.25 | 2889539.25 | 3283281.75 | 4870095 | 4578351 | 4829701 | 3859310.75 |
| P63087 | Ppp1cc | Serine/threonine-protein phosphatase PP1-gamm | 350.64 | 19.5 | 36984 | 6 | 657737.69 | 252846.08 | 207217.09 | 1321768.5 | 3345558 | 2679060.75 | 2610676 | 4155477 | 3250498.5 | 2320715 | 2532631.25 | 2813572.75 | 3018874.25 | 2185929.25 | 3052544.25 | 2276027.25 | 2474510.5 | 2381992.75 | 3666731.75 | 2752657.25 |
| Q9DBR7 | Ppp1r12a | Protein phosphatase 1 regulatory subunit 12A | 97.38 | 1.85 | 114996 | 2 | 2 | 2 | 2 | 2 | 1498298.12 | 2 | 2 | 3110476.25 | 1089932 | 1500623.88 | 633698.62 | 2 | 2 | 2 | 2 | 2 | 2 | 2 | 2 | 2 |
| Q8BQ30 | Ppp1r18 | Phostensin | 176.72 | 7.74 | 65630 | 4 | 2578289 | 1995639.5 | 3724923.75 | 3327681.5 | 1651581.75 | 1508743.5 | 2480368.75 | 4723747 | 1340075.62 | 2422675 | 6876542 | 5524388 | 2567715.75 | 2095244.5 | 7132928.5 | 4162518.5 | 2465617 | 3016928.75 | 1382419 | 2 |
| Q3UM45 | Ppp1r7 | Protein phosphatase 1 regulatory subunit 7 | 170.62 | 10.8 | 41292 | 3 | 2 | 2 | 699677.19 | 2 | 2 | 2 | 2 | 2 | 2 | 2 | 2 | 135594.05 | 113437.14 | 109170.34 | 595281.81 | 2 | 239661.44 | 2 | 2 | 442493.47 |
| P62715 | Ppp2cb | Serine/threonine-protein phosphatase 2A catalyti | 427.73 | 34.63 | 35575 | 8 | 2522430.5 | 983004.56 | 4800941.5 | 1887727.25 | 3442438 | 2 | 797975.88 | 1993643.88 | 1401214 | 2811221 | 1491320.62 | 1234816 | 92514.78 | 2623246.25 | 2911970.75 | 798748.75 | 892679.56 | 1251342.5 | 1710935.75 | 1359350.25 |
| Q76MZ3 | Ppp2r1a | Serine/threonine-protein phosphatase 2A 65 kDa | 1052.02 | 34.47 | 65323 | 17 | 12130062 | 23265726 | 16284975 | 16013564 | 19607784 | 15168634 | 16316286 | 12311138 | 29010970 | 12364945 | 13230359 | 14419928 | 15607663 | 15244679 | 10705673 | 8108920 | 10686543 | 11764198 | 11150206 | 8068012 |
| Q4QQT4 | Ppp2r1b | Serine/threonine-protein phosphatase 2A 65 kDa | 510.63 | 14.98 | 66006 | 8 | 2 | 169236.78 | 2 | 94715.71 | 399179.62 | 2 | 110290.14 | 178155.92 | 262933.81 | 320030.06 | 2 | 2 | 2 | 2 | 2 | 2 | 2 | 2 | 2 | 2 |
| P36876 | Ppp2r2a | Serine/threonine-protein phosphatase 2A 55 kDa | 163.13 | 7.16 | 51678 | 3 | 30972.25 | 64315.41 | 255655.59 | 413491.16 | 132906.55 | 2 | 218981.19 | 2 | 2 | 811528.19 | 2 | 2 | 2 | 445524.16 | 623230.69 | 41708.6 | 143834.88 | 50733.91 | 2 | 1272720.75 |
| Q60996 | Ppp2r5c | Serine/threonine-protein phosphatase 2A 56 kDa | 34.06 | 2.1 | 60824 | 1 | 2 | 2 | 168611.61 | 377979.06 | 74774.09 | 2 | 2 | 2 | 204689.67 | 157946.5 | 2 | 2 | 176618.55 | 297256 | 378608.31 | 2 | 338425.09 | 362002.47 | 2 | 269948.31 |
| P63328 | Ppp3ca | Protein phosphatase 3 catalytic subunit alpha | 159.44 | 8.45 | 58644 | 3 | 3197223 | 1438408.5 | 3898545 | 2427517 | 2659301.25 | 2041946.75 | 1370132.25 | 2416171.75 | 1588943 | 3691553 | 1363106.88 | 981646.38 | 1475951.5 | 1234680.75 | 3179824 | 1209625.12 | 2123646.75 | 1943261.38 | 142225.73 | 2216684.5 |
| P53042 | Ppp5c | Serine/threonine-protein phosphatase 5 | 47.5 | 2.61 | 56917 | 1 | 2 | 2 | 453812.62 | 434371.06 | 2 | 2 | 2 | 2 | 2 | 401542.75 | 2 | 2 | 732186.12 | 504711.62 | 2 | 2 | 2 | 571825.94 | 2 | 2 |
| Q64620 | Ppp6c | Serine/threonine-protein phosphatase 6 catalytic | 97.77 | 7.21 | 35159 | 2 | 1084449.75 | 1283511 | 1315750.12 | 1436803.5 | 1555070.12 | 1379307.5 | 1027293.56 | 1365909.5 | 1125501.75 | 987252.06 | 1262517 | 2161311.5 | 1520247.5 | 1551061.25 | 2649245 | 1211231.75 | 1959471.38 | 1489308 | 176199.47 | 1831235.88 |
| Q7TSI3 | Ppp6r1 | Serine/threonine-protein phosphatase 6 regulator | 54.37 | 1.29 | 94527 | 1 | 175429.36 | 2087328.12 | 2 | 2210431.5 | 72048.45 | 240261.97 | 532392.75 | 430605.28 | 1463015.25 | 2 | 130542.39 | 1559547.88 | 2 | 1670539.38 | 2 | 1318470.5 | 698873.94 | 117661.03 | 2 | 1244141.12 |
| O88531 | Ppt1 | Palmitoyl-protein thioesterase 1 | 295.3 | 19.61 | 34490 | 5 | 8838689 | 15477948 | 9727674 | 206665.81 | 8421770 | 4909047 | 9752518 | 10622558 | 3857558.5 | 9830272 | 6617199 | 5117918 | 5362312 | 5001404.5 | 4727743 | 6123915 | 4503606 | 4087956.25 | 4915197.5 | 2964324 |
| Q6PCT5 | Pqbp1 | Polyglutamine-binding protein 1 | 34.37 | 3.42 | 30529 | 1 | 682938.38 | 2 | 599305.81 | 2 | 2 | 634401.31 | 2 | 2 | 287258.22 | 2 | 1703172.12 | 959689.31 | 1314095.88 | 519011.09 | 1546977.62 | 2522625.75 | 1929260.38 | 2 | 2 | 4546424.5 |
| D3ZMK9 | Prag1 | Inactive tyrosine-protein kinase PRAG1 | 45.68 | 0.8 | 147716 | 1 | 431423.69 | 446398.78 | 2 | 881952.94 | 2 | 2 | 501286.12 | 699267.75 | 553602.25 | 661065.69 | 2 | 2 | 2 | 2 | 2 | 2 | 2 | 2 | 2 | 2 |
| P35700 | Prdx1 | Peroxiredoxin-1 | 1103.39 | 53.77 | 22176 | 17 | 13871419 | 28748262 | 13611440 | 37563394 | 34030066 | 61296334 | 56168262 | 72294226 | 55498590 | 39697686 | 41313430 | 50029586 | 37164802 | 45753682 | 27941380 | 39975990 | 33018518 | 22641186 | 28011310 | 23630922 |
| Q61171 | Prdx2 | Peroxiredoxin-2 | 711.26 | 43.94 | 21779 | 11 | 46306174 | 54412038 | 49903114 | 52859890 | 40967898 | 50078134 | 53578998 | 38571750 | 39767258 | 35017250 | 45984970 | 49475654 | 48476778 | 49579678 | 76821658 | 36489774 | 53071706 | 56342466 | 32183974 | 44695654 |
| P20108 | Prdx3 | Thioredoxin-dependent peroxide reductase, mitoc | 202.17 | 18.68 | 28127 | 5 | 8904104 | 6346298 | 7438597 | 5824449 | 3595959.5 | 4081776.5 | 5475390 | 2366655 | 4332271 | 9623880 | 2749267.5 | 3173503.75 | 867063.44 | 4872225 | 10630382 | 2954892 | 5741571 | 1883318.5 | 2055061.12 | 6325908 |
| O08807 | Prdx4 | Peroxiredoxin-4 | 392.13 | 27.01 | 31053 | 7 | 11267086 | 3134515 | 12965255 | 3048096.5 | 8365036 | 593057.62 | 1936787.88 | 2671176.25 | 4499092 | 9121909 | 3177857.5 | 3660511.5 | 3636011.25 | 1692681.5 | 3824756 | 4629682 | 4251336.5 | 4522096 | 4231712 | 3656910 |
| P99029 | Prdx5 | Peroxiredoxin-5, mitochondrial | 794.09 | 68.57 | 21897 | 14 | 9488887 | 18599526 | 14394362 | 8802310 | 26972944 | 44275626 | 48016310 | 57353422 | 40746430 | 63947914 | 23448912 | 22285586 | 21672770 | 23946552 | 19834446 | 22975926 | 16740564 | 16086334 | 20506966 | 12493041 |
| O08709 | Prdx6 | Peroxiredoxin-6 | 573.79 | 56.7 | 24871 | 11 | 19551318 | 26528092 | 21720390 | 11932287 | 23694624 | 16684794 | 15692404 | 18477570 | 16936580 | 21201312 | 19518304 | 20773008 | 21367108 | 20455400 | 19326852 | 19454074 | 17228994 | 16921270 | 19971954 | 18677820 |
| Q9WUQ2 | Preb | Guanine nucleotide-exchange factor SEC12 | 107.26 | 11.03 | 45437 | 3 | 2 | 2 | 1210420.5 | 2 | 393384.28 | 67472.81 | 84113.34 | 87837.76 | 949637.69 | 577237 | 2 | 139560.2 | 2 | 736035.75 | 2 | 2 | 89250.24 | 2 | 2 | 2 |
| O70196 | Prep | Prolyl endopeptidase | 244.1 | 9.01 | 80742 | 5 | 1208404 | 698051.5 | 2378140.5 | 2729698.25 | 3217094.25 | 1973567.38 | 2106008 | 1914418.5 | 2114595 | 1949098.75 | 2374438.5 | 1025739.62 | 2982007.5 | 1016498.81 | 251708.52 | 4285659 | 389494.41 | 2 | 1416477.12 | 2820224 |
| Q69ZK0 | Prex1 | Phosphatidylinositol 3,4,5-trisphosphate-depende | 198.61 | 3.03 | 184935 | 4 | 4671311.5 | 4800441 | 4561890.5 | 4068023.25 | 3112972 | 2852672.25 | 3940321 | 4242052 | 1586398.38 | 3312852.75 | 470543.62 | 911467.69 | 1105735.62 | 1729260.62 | 3009376 | 2 | 2583738 | 1700429.88 | 2876448.75 | 1943441 |
| P05132 | Prkaca | cAMP-dependent protein kinase catalytic subunit | 82.36 | 5.98 | 40571 | 2 | 2 | 2 | 2 | 2 | 37414.03 | 2 | 2 | 2 | 118791.7 | 80893.55 | 2 | 2 | 2 | 2 | 2 | 2 | 2 | 2 | 2 | 2 |
| P68181 | Prkacb | cAMP-dependent protein kinase catalytic subunit | 88.09 | 5.98 | 40708 | 2 | 2 | 2 | 2 | 2 | 128636.17 | 2 | 127924.28 | 2 | 215042.89 | 234720.75 | 2 | 175712.58 | 67886.05 | 2 | 2 | 2 | 2 | 2 | 2 | 2 |
| Q9DBC7 | Prkar1a | cAMP-dependent protein kinase type I-alpha regu | 304.45 | 17.32 | 43185 | 5 | 935364.31 | 1036101.88 | 1780123.62 | 1165090.62 | 2548023 | 504000.62 | 1136338.75 | 169874.97 | 2988839 | 1008506 | 2591991 | 2947374 | 3285255.75 | 2213896.75 | 1111637.5 | 1507018.88 | 740466.25 | 603581.5 | 2 | 1969204.12 |
| P12367 | Prkar2a | cAMP-dependent protein kinase type II-alpha regu | 69.28 | 3.49 | 45389 | 1 | 2 | 666346.06 | 2 | 2 | 767342.69 | 2 | 2 | 887458.69 | 2 | 2 | 2055824.5 | 1152506.88 | 2 | 1478246.12 | 1533211.12 | 1905613.12 | 2214402.5 | 1338215.5 | 1249581 | 2 |
| P28867 | Prkcd | Protein kinase C delta type | 192.93 | 6.53 | 77547 | 4 | 600645.62 | 2 | 2 | 543946.88 | 2565137.75 | 2390181 | 918683.12 | 2725531.5 | 3589544.5 | 1628294.38 | 3479009.25 | 3536567 | 3942442.5 | 3567291 | 2578223 | 3383170 | 1739306.88 | 1814074.5 | 2782766.75 | 2334025.25 |
| O08795 | Prkcsh | Glucosidase 2 subunit beta | 410.27 | 11.9 | 58793 | 7 | 22015980 | 5699820 | 11781647 | 23044178 | 6519366 | 16151913 | 15889345 | 3730790.25 | 6839439 | 6424124 | 12330879 | 11844927 | 14480363 | 11870345 | 13579828 | 14731318 | 10296080 | 25575864 | 9317931 | 24410106 |
| Q63009 | Prmt1 | Protein arginine N-methyltransferase 1 | 78.09 | 8.5 | 40522 | 2 | 772649.25 | 881908.44 | 585384.44 | 3247786.75 | 2540977.5 | 975552.06 | 1478549.88 | 1863897.88 | 982945.5 | 2085439.62 | 2 | 789148.31 | 1251485.5 | 2 | 1824528.75 | 697264.12 | 930394.75 | 1580738 | 2 | 233302.05 |
| Q99KP6 | Prpf19 | Pre-mRNA-processing factor 19 | 259.72 | 21.63 | 55239 | 6 | 1802065.62 | 1367881 | 1225169.75 | 5200482 | 2636250.75 | 1541520.12 | 1627458.62 | 1859515.38 | 2420706.75 | 1142465.88 | 2 | 533287.88 | 2 | 527514.88 | 2 | 44853.05 | 2 | 2 | 2 | 2 |
| Q922U1 | Prpf3 | U4/U6 small nuclear ribonucleoprotein Prp3 | 125.63 | 4.39 | 77455 | 3 | 458294.19 | 194688.42 | 2 | 2 | 2 | 689896.81 | 2 | 546784.06 | 413705.75 | 2 | 2 | 2 | 290459.91 | 2 | 2 | 2 | 2 | 2 | 2 | 2 |
| Q8CCF0 | Prpf31 | U4/U6 small nuclear ribonucleoprotein Prp31 | 178.39 | 12.42 | 55430 | 4 | 981284.94 | 2504893 | 2006073 | 1947242.88 | 3148901.5 | 610942.75 | 1782674.5 | 2484621.25 | 2219221.5 | 1972252 | 1813229.88 | 1302277.88 | 2457180 | 2364413.75 | 1431942.12 | 2 | 2 | 1728739.5 | 1285759.62 | 2 |
| Q4FK66 | Prpf38a | Pre-mRNA-splicing factor 38A | 40.31 | 3.85 | 37437 | 1 | 2 | 2 | 1070568.75 | 2 | 2 | 2 | 2 | 2 | 573930.12 | 2 | 2 | 612862.81 | 2 | 2 | 981088.94 | 2 | 2 | 2 | 2 | 917868.69 |
| Q6AXY7 | Prpf38b | Pre-mRNA-splicing factor 38B | 63.31 | 2.77 | 63932 | 2 | 2892513 | 3558309.75 | 1744432 | 3755884.75 | 3456345.5 | 1941467.38 | 1653914.88 | 2879073.5 | 3461502.25 | 3044738.5 | 2020739.75 | 1078170.62 | 3012580.5 | 2427134.5 | 3332191.75 | 1737453.75 | 4178136 | 1697899.62 | 3587491.25 | 4523266 |
| Q9DAW6 | Prpf4 | U4/U6 small nuclear ribonucleoprotein Prp4 | 46 | 2.5 | 58370 | 1 | 2 | 2 | 2 | 2 | 2 | 2 | 2 | 2 | 2 | 2 | 2 | 2 | 2 | 2 | 2 | 2 | 2 | 2 | 2 | 2 |
| Q9R1C7 | Prpf40a | Pre-mRNA-processing factor 40 homolog A | 86.33 | 2.62 | 108481 | 2 | 2 | 974316.81 | 1479805.5 | 354120.59 | 2 | 2 | 2 | 2 | 2 | 584049.75 | 2 | 2 | 2 | 2 | 2 | 2 | 2 | 445421.41 | 1170409.38 | 952018.56 |
| A1A5S1 | Prpf6 | Pre-mRNA-processing factor 6 | 123.6 | 3.61 | 106764 | 3 | 1658076.12 | 647884.44 | 644235.06 | 2 | 326806.03 | 2 | 350810.62 | 2 | 1022067.94 | 636458.44 | 2 | 2199908.25 | 1720603.25 | 3074311.5 | 2071828.12 | 2138125 | 1103877.38 | 3520685.25 | 3151959 | 3796034.75 |
| Q99PV0 | Prpf8 | Pre-mRNA-processing-splicing factor 8 | 90.4 | 0.81 | 273616 | 2 | 1002048.38 | 2 | 1240882 | 2 | 937611.12 | 2 | 446441.34 | 2 | 573496 | 593829.25 | 1545761.25 | 454546.59 | 1013894.88 | 913230.25 | 2 | 604582.38 | 408962.56 | 2 | 2 | 2 |
| P09330 | Prps2 | Ribose-phosphate pyrophosphokinase 2 | 272.31 | 18.87 | 34813 | 6 | 2590485 | 1014891.12 | 3004717 | 4785663 | 3681532.75 | 1265425.88 | 3457419.25 | 831759.62 | 4320807.5 | 2924963.5 | 1521337.5 | 6033339.5 | 4663941 | 6595203 | 9650472 | 2039929.5 | 5587068 | 1897888.12 | 1177544.75 | 4670373 |
| Q63468 | Prpsap1 | Phosphoribosyl pyrophosphate synthase-associa | 217.02 | 15.73 | 39436 | 4 | 2 | 2 | 1679654.12 | 568597.25 | 2 | 2 | 332654.5 | 176486.55 | 117364.88 | 740392.19 | 2 | 2 | 47747.39 | 297072.03 | 899481.38 | 2 | 2 | 2 | 2 | 2 |
| O08618 | Prpsap2 | Phosphoribosyl pyrophosphate synthase-associa | 152.92 | 9.49 | 40867 | 3 | 2 | 265265.41 | 213786.42 | 596853.38 | 196075.28 | 182729.88 | 546258.75 | 283404.53 | 414847.75 | 272253.62 | 2 | 2 | 105584.05 | 209673.75 | 439803.59 | 2 | 257979.66 | 2 | 2 | 2 |
| Q3T1I4 | Prrc1 | Protein PRRC1 | 36.98 | 2.93 | 46279 | 1 | 1076603.62 | 1291714.25 | 1311545.12 | 1246634.62 | 901832.12 | 737120.5 | 2 | 1732846.75 | 424291.47 | 2 | 957081.69 | 3011616.5 | 1028352.31 | 2 | 1377405.25 | 2 | 2 | 2 | 2 | 2505505 |
| Q3TLH4 | Prrc2c | Protein PRRC2C | 201.9 | 1.51 | 310892 | 5 | 1126060.5 | 1873869.5 | 191439.95 | 3024695.5 | 4849147.5 | 3879694.75 | 2916089 | 1060859.5 | 6428086 | 1750165.25 | 4797656 | 4441691 | 3181832.5 | 4162810.25 | 2865676.75 | 4602226 | 4098282 | 2410838.5 | 5819722 | 606581.38 |
| Q8BIW1 | Prune1 | Exopolyphosphatase PRUNE1 | 41.88 | 3.08 | 50239 | 1 | 2 | 1008074.5 | 2 | 2 | 2 | 2 | 2 | 2 | 410772.69 | 2 | 2 | 2 | 593386.44 | 775039.69 | 2 | 2 | 793045.75 | 738202.5 | 2 | 2 |
| Q61207 | Psap | Prosaposin | 335.95 | 15.62 | 61422 | 7 | 58043146 | 28368834 | 64919290 | 31909894 | 36689478 | 27314566 | 28061566 | 44449598 | 23488012 | 56325506 | 22374538 | 21113362 | 22366956 | 18440558 | 39247638 | 19001262 | 27114490 | 12123280 | 23268838 | 28063704 |
| Q99K85 | Psat1 | Phosphoserine aminotransferase | 625.66 | 40.81 | 40473 | 12 | 9448713 | 8158965 | 14012177 | 10083978 | 12094220 | 10817532 | 10010280 | 17130962 | 10743582 | 11489357 | 15295362 | 14744097 | 17872146 | 16459576 | 13024913 | 15648109 | 16903908 | 18258028 | 12322903 | 15239247 |
| Q812D1 | Psip1 | PC4 and SFRS1-interacting protein | 81.23 | 3.6 | 59639 | 2 | 2 | 2 | 2 | 429349.91 | 352678.88 | 2 | 2 | 107920.57 | 2 | 2 | 129975.93 | 774447.69 | 533386.62 | 2 | 629398.75 | 2 | 2 | 2 | 2 | 374261.88 |
| Q9R1P4 | Psma1 | Proteasome subunit alpha type-1 | 393.23 | 23.95 | 29547 | 8 | 7719415.5 | 5452876.5 | 7548959.5 | 5278407 | 5675505.5 | 4186438.75 | 3189706.25 | 4169066.5 | 6356008.5 | 8814443 | 10520429 | 9292530 | 8091771 | 11528193 | 5012210 | 8554945 | 6097111 | 5720360.5 | 9061453 | 6082500 |
| P17220 | Psma2 | Proteasome subunit alpha type-2 | 383.9 | 32.48 | 25927 | 7 | 7725669.5 | 4468185.5 | 9971947 | 4143552.75 | 7982344.5 | 4499552 | 7900137 | 4275806.5 | 7103348 | 5489035 | 2312023.5 | 5069292 | 3836176.75 | 5722223 | 5552850 | 2318449.75 | 4388279.5 | 2055041.25 | 1513704 | 4683970 |
| O70435 | Psma3 | Proteasome subunit alpha type-3 | 426.61 | 29.02 | 28405 | 7 | 33545350 | 48792914 | 30673740 | 36638322 | 31093562 | 40033010 | 31639274 | 43840402 | 29191530 | 20604120 | 21665390 | 20401898 | 25603076 | 28050738 | 14937402 | 23032304 | 30165126 | 28436302 | 16356142 | 32771928 |
| P21670 | Psma4 | Proteasome subunit alpha type-4 | 236.29 | 15.33 | 29498 | 5 | 12743202 | 19849570 | 17719450 | 24023240 | 33258732 | 23730898 | 19291270 | 13447227 | 20880226 | 21340242 | 14949842 | 12450147 | 13672440 | 20673494 | 18588714 | 18739580 | 15963042 | 19400490 | 21678066 | 15834059 |
| Q9Z2U1 | Psma5 | Proteasome subunit alpha type-5 | 348.08 | 25.73 | 26411 | 6 | 7855501 | 7310222.5 | 12824492 | 8654101 | 23980448 | 13790574 | 12297055 | 22131720 | 13897165 | 20416770 | 11435086 | 9516324 | 14766409 | 13049949 | 13096922 | 9625605 | 10931689 | 11800206 | 11347576 | 7390539.5 |
| P60901 | Psma6 | Proteasome subunit alpha type-6 | 351.95 | 21.54 | 27399 | 6 | 6466325 | 10728180 | 11264161 | 8875845 | 32687574 | 15811203 | 15862194 | 28107632 | 23286244 | 20840626 | 21692750 | 21556704 | 22059018 | 23417578 | 6486013.5 | 16372375 | 12340630 | 6315571 | 21218432 | 5789608.5 |
| Q9Z2U0 | Psma7 | Proteasome subunit alpha type-7 | 289.59 | 29.44 | 27855 | 5 | 31760.02 | 240552.22 | 3915350.5 | 887491.31 | 2624709.75 | 1653525.5 | 2216172 | 1414594.25 | 4113228 | 3484550 | 2686454.75 | 3082809 | 2099897.75 | 3072742.75 | 3594209.5 | 2233349.25 | 2187132.5 | 951497.06 | 638001.25 | 1409119 |
| O09061 | Psmb1 | Proteasome subunit beta type-1 | 461.86 | 48.33 | 26372 | 8 | 7661762.5 | 5470640.5 | 11704808 | 6225250.5 | 9780721 | 8415155 | 8380595.5 | 9470802 | 6501656 | 11328120 | 6318773 | 4384526.5 | 7544734.5 | 5196951.5 | 7608743 | 8563516 | 5770648 | 8902738 | 8453461 | 3784228 |
| O35955 | Psmb10 | Proteasome subunit beta type-10 | 89.07 | 11.36 | 29063 | 2 | 2 | 480559.03 | 2 | 1213084.62 | 3677821.75 | 2761070 | 1660976 | 3328587.25 | 4553782.5 | 2352403.75 | 2786418.75 | 1671894.12 | 1344255.12 | 2124437 | 161939.45 | 2198690.25 | 1299906.88 | 2 | 2430337.75 | 2 |
| P40307 | Psmb2 | Proteasome subunit beta type-2 | 203.03 | 27.36 | 22912 | 4 | 152119.59 | 688955.81 | 2873947 | 3266686.75 | 1401850.38 | 3092324 | 5231155 | 935415.75 | 5125364.5 | 3893060.25 | 1909840 | 2497271 | 4790767 | 5163400 | 7665302 | 1324721.75 | 5297176 | 7486435 | 1246079.38 | 3333940 |
| P40112 | Psmb3 | Proteasome subunit beta type-3 | 401.43 | 40.49 | 22965 | 6 | 7593357.5 | 1751326.75 | 7520347 | 4925593 | 8366525 | 1597245.62 | 4655907.5 | 5786477 | 5255270.5 | 8175808.5 | 1655218.12 | 2716253 | 4106406.25 | 3739480.5 | 5604327.5 | 2 | 1539142.88 | 1361431 | 1892434.38 | 1646391 |
| P99026 | Psmb4 | Proteasome subunit beta type-4 | 367.09 | 38.26 | 29116 | 6 | 7635987 | 5391419 | 15069204 | 7789497 | 10784907 | 11044248 | 10383130 | 12661254 | 12450932 | 11877176 | 10991074 | 6656553 | 10346385 | 11188844 | 10140668 | 6895527 | 10479387 | 10911522 | 12415712 | 6611175 |
| O55234 | Psmb5 | Proteasome subunit beta type-5 | 48.54 | 4.92 | 28532 | 1 | 2 | 943657.25 | 2 | 2 | 2 | 1019992.5 | 651619.5 | 2 | 2 | 2 | 2 | 2 | 2 | 691719.69 | 2 | 2 | 2 | 2 | 1033869.69 | 2 |
| Q60692 | Psmb6 | Proteasome subunit beta type-6 | 193.33 | 16.81 | 25379 | 4 | 5311137.5 | 6450081 | 4340186 | 4557643 | 167663.27 | 2 | 2992026 | 130886.95 | 2218066 | 2801314.75 | 12489238 | 11316821 | 12225833 | 13102580 | 6924246 | 11425199 | 8095560.5 | 9920793 | 9014656 | 9019310 |
| P28063 | Psmb8 | Proteasome subunit beta type-8 | 255.04 | 21.01 | 30260 | 5 | 31144838 | 51451410 | 10795795 | 33815362 | 8968920 | 33222702 | 28340172 | 12520409 | 22704168 | 20291434 | 15978666 | 15938502 | 21304250 | 20727614 | 18064770 | 19571848 | 30607408 | 34422710 | 11692046 | 31032842 |
| O35522 | Psmb9 | Proteasome subunit beta type-9 bactrianus | 116.08 | 10.05 | 23427 | 2 | 11074730 | 12522348 | 7171013 | 14448926 | 7554527 | 13250722 | 10393366 | 11023109 | 13699868 | 14603957 | 7718837 | 7069684 | 9549400 | 8431505 | 6837913.5 | 8768948 | 9543817 | 9458458 | 11767971 | 11868280 |
| P62192 | Psmc1 | 26S proteasome regulatory subunit 4 | 415.89 | 20.91 | 49185 | 9 | 1041212.19 | 718713.38 | 745469.56 | 2152499.25 | 3271364.5 | 1811807.38 | 2035427.88 | 7699082.5 | 5383387.5 | 3663671.5 | 2590398.25 | 3163731 | 2890233 | 3613960.5 | 2840618.75 | 3955490 | 2093432.12 | 4063330 | 2324604 | 1998020.5 |
| P46471 | Psmc2 | 26S proteasome regulatory subunit 7 | 836.74 | 43.65 | 48648 | 16 | 10676790 | 11251473 | 11083330 | 11050570 | 12533602 | 12944468 | 13653607 | 12868969 | 11083633 | 10335644 | 14404920 | 12801256 | 13667678 | 14688781 | 13546005 | 12927290 | 11058307 | 15508287 | 9600467 | 11117361 |
| O88685 | Psmc3 | 26S proteasome regulatory subunit 6A | 706.72 | 29.86 | 49549 | 12 | 12265619 | 12152889 | 6944248.5 | 8783541 | 11440137 | 8644024 | 7522943 | 7846717 | 9920552 | 9168887 | 8385725 | 7376268.5 | 9672807 | 8979169 | 11056055 | 6113232 | 7964434.5 | 8509347 | 8559016 | 9074744 |
| P54775 | Psmc4 | 26S proteasome regulatory subunit 6B | 356.26 | 15.07 | 47408 | 6 | 851727.56 | 213050.41 | 1077189.5 | 1117252.5 | 2979395.5 | 1327975.75 | 2966838.5 | 3553110.5 | 4807502.5 | 3245749.75 | 924845.62 | 1132288.62 | 2079667.75 | 1514058.88 | 3495064 | 967669.94 | 722176 | 399405.88 | 1634287.62 | 770767.62 |
| P62196 | Psmc5 | 26S proteasome regulatory subunit 8 | 469.79 | 26.6 | 45626 | 8 | 5829630.5 | 8187355 | 5373652.5 | 6408341.5 | 9955734 | 7208816 | 3920055.75 | 3574565.5 | 5668170 | 6453209 | 8591783 | 8872737 | 9036572 | 8791815 | 7380224 | 5459301.5 | 6561993.5 | 7774615 | 3728431 | 1183767.62 |
| P62334 | Psmc6 | 26S proteasome regulatory subunit 10B | 482.71 | 30.08 | 44173 | 10 | 2232163.75 | 4643814.5 | 4428600.5 | 5065008 | 8421814 | 4531913.5 | 4141305.5 | 6451144 | 3568733 | 8585544 | 3400808.75 | 4452414 | 5806088.5 | 1088459.25 | 4630109 | 1275975.38 | 1937899 | 2015306.12 | 2019552.5 | 3224395.25 |
| O88761 | Psmd1 | 26S proteasome non-ATPase regulatory subunit | 458.93 | 10.6 | 105748 | 9 | 4903021 | 3541662.75 | 5534060.5 | 8891814 | 7942737.5 | 1346710.25 | 4706455.5 | 8916558 | 10160169 | 6898353.5 | 7473576 | 8015962 | 8066934 | 8954611 | 5834236 | 8326968 | 5040707 | 4219979 | 6465529 | 3762168.25 |
| F1LMZ8 | Psmd11 | 26S proteasome non-ATPase regulatory subunit | 467.88 | 22.51 | 47464 | 9 | 5621725 | 4056830.75 | 7971527 | 2354795.5 | 7270545.5 | 6779526 | 4990028.5 | 6801822 | 5204051.5 | 7564933.5 | 8256473.5 | 5926069 | 6582533 | 7502875 | 6393937 | 8592791 | 6001082.5 | 5971549 | 2 | 5608871 |
| Q9D8W5 | Psmd12 | 26S proteasome non-ATPase regulatory subunit | 444.33 | 19.96 | 52895 | 8 | 7325029 | 5822267 | 8628418 | 4536838 | 4062631 | 2987838.25 | 4707265 | 4195682.5 | 2891901.75 | 4465348 | 3692700 | 3345974 | 5216486 | 4207774.5 | 6663903 | 4490423 | 5606136 | 7836726 | 4150482.75 | 5787133 |
| B0BN93 | Psmd13 | 26S proteasome non-ATPase regulatory subunit | 224.85 | 12.23 | 42817 | 5 | 5409428 | 7316887.5 | 11637260 | 4365922 | 8315481 | 4195695 | 5528016.5 | 7863446.5 | 1578388.62 | 8589324 | 3378196.25 | 3400037.75 | 4855211.5 | 2702906 | 4644788.5 | 3873482.25 | 3555004.75 | 4765436.5 | 4369421 | 4121422.5 |
| O35593 | Psmd14 | 26S proteasome non-ATPase regulatory subunit | 335.78 | 31.61 | 34577 | 6 | 8682156 | 7335195 | 13124154 | 6328385.5 | 11620804 | 7148206 | 5847561.5 | 7420859 | 7323625.5 | 9210600 | 7202651 | 6509749 | 10757460 | 7979681 | 10977517 | 8495962 | 8338983.5 | 13236512 | 9486660 | 10060102 |
| Q8VDM4 | Psmd2 | 26S proteasome non-ATPase regulatory subunit | 779.95 | 18.5 | 100203 | 15 | 11768915 | 19156946 | 13812092 | 16985478 | 16703999 | 16292248 | 16294049 | 13155558 | 9557615 | 19792482 | 15930070 | 15113172 | 19674348 | 18066008 | 15894466 | 12664865 | 14209352 | 12022608 | 13034283 | 10351948 |
| P14685 | Psmd3 | 26S proteasome non-ATPase regulatory subunit | 159.49 | 4.91 | 60718 | 3 | 2 | 2 | 2 | 182490.91 | 62945.85 | 2 | 1635226.62 | 1059899 | 1840489.25 | 403773.41 | 406999 | 592876.31 | 1209183 | 1451740.5 | 707476.94 | 123247.55 | 431152.19 | 660757.06 | 356804.16 | 146259.58 |
| O35226 | Psmd4 | 26S proteasome non-ATPase regulatory subunit | 93.04 | 5.32 | 40704 | 2 | 2779402 | 3644293 | 2304349.5 | 2659288 | 3656623 | 4395527.5 | 2152778.5 | 1369992 | 3663422.5 | 3939181 | 4163676 | 4227300 | 5405960 | 5261857 | 3322323.75 | 3689395.5 | 5902679 | 4678242 | 7824437 | 5675494.5 |
| Q8BJY1 | Psmd5 | 26S proteasome non-ATPase regulatory subunit | 111.3 | 6.75 | 55972 | 2 | 1173755.12 | 434172.72 | 1251657.88 | 460237.78 | 805510.62 | 272924.66 | 1552428.62 | 127152.38 | 1016643.44 | 1202119.75 | 145240.58 | 1216658.62 | 1244552 | 1114356.75 | 1439500.25 | 93295.73 | 1324559.5 | 372902.31 | 2 | 181493.95 |
| Q99JI4 | Psmd6 | 26S proteasome non-ATPase regulatory subunit | 208.05 | 12.85 | 45536 | 5 | 2374235.75 | 2059643.62 | 4248487 | 1152955.62 | 5975977 | 2011993 | 1046940 | 4352494 | 3333668.75 | 4939393 | 4566706.5 | 3659335.5 | 2872957 | 3443555 | 1732260.88 | 2548261 | 2174391 | 3365652 | 2453136.5 | 1584917.75 |
| P26516 | Psmd7 | 26S proteasome non-ATPase regulatory subunit | 258.44 | 16.82 | 36540 | 4 | 1091989.25 | 1059157.5 | 2980458.75 | 1067578 | 1146851 | 2 | 1280344 | 2646433.5 | 1438660.75 | 2900192 | 1889378.5 | 1283174.75 | 1593628 | 1152762.5 | 1882883.88 | 131365.62 | 1848261.25 | 1316565.25 | 133868.47 | 795576.19 |
| Q9CX56 | Psmd8 | 26S proteasome non-ATPase regulatory subunit | 157.69 | 4.82 | 39930 | 2 | 99277.3 | 2 | 310155.56 | 151073.5 | 130774.62 | 2 | 281508.66 | 107737.44 | 137934.55 | 322053.44 | 2 | 52452.28 | 137528.89 | 114939.42 | 1118906 | 2 | 135726.53 | 62318.53 | 2 | 2 |
| Q9CR00 | Psmd9 | 26S proteasome non-ATPase regulatory subunit | 40.85 | 5.41 | 24720 | 1 | 1521758.38 | 1871138.75 | 1186817.38 | 896659.12 | 470612.25 | 2 | 449172.25 | 1210055.88 | 2 | 825797.88 | 1452143.5 | 2 | 1190621.12 | 868576.25 | 1149653.38 | 5760717.5 | 1257586.12 | 3539613 | 6545791.5 | 1083997.88 |
| P97371 | Psme1 | Proteasome activator complex subunit 1 | 240.45 | 17.67 | 28673 | 4 | 14518968 | 10753305 | 12940395 | 15649119 | 15157189 | 17291136 | 18487774 | 18334312 | 13755834 | 17312258 | 4928276.5 | 6329604 | 7259439 | 5337163 | 8145105 | 6145879 | 5158249 | 7575598 | 3609753 | 5843317 |
| P97372 | Psme2 | Proteasome activator complex subunit 2 | 445.84 | 35.15 | 27057 | 8 | 6294491 | 6775690.5 | 11254707 | 12676632 | 16376726 | 9158677 | 9574677 | 10271082 | 8375536 | 18236732 | 3060721.5 | 5525238 | 5917710.5 | 5886359 | 7323923 | 5912933.5 | 5897028.5 | 7707918 | 7710094.5 | 3356980.5 |
| P61290 | Psme3 | Proteasome activator complex subunit 3 | 111.9 | 11.42 | 29506 | 3 | 582673.81 | 1133103 | 3011018.5 | 1035007.38 | 1566607.5 | 2 | 2 | 2 | 2 | 877078.75 | 1160853.62 | 1036260 | 819331.38 | 1149589.38 | 1445069.25 | 1690376.62 | 692036.62 | 2 | 925850.94 | 2 |
| Q9JK23 | Psmg1 | Proteasome assembly chaperone 1 | 44.45 | 3.46 | 33104 | 1 | 2 | 2 | 43251.45 | 2 | 2 | 2 | 2 | 2 | 61870.52 | 62699.6 | 2 | 262477 | 167223.56 | 317175.72 | 440717.97 | 2 | 429849.66 | 179663.09 | 135045.39 | 2 |
| Q9CZH3 | Psmg3 | Proteasome assembly chaperone 3 | 49.75 | 19.67 | 13331 | 1 | 2 | 2 | 2 | 2 | 1428006.25 | 459246.53 | 2 | 310138.75 | 1354664.62 | 694069.25 | 227359.06 | 249336.06 | 2 | 327043.16 | 2 | 224851.14 | 2 | 2 | 173444.59 | 2 |

| | | | | | | | | | | | | | | | | | | | | | | | | | | |
|---|---|---|---|---|---|---|---|---|---|---|---|---|---|---|---|---|---|---|---|---|---|---|---|---|---|---|
| P97814 | Pstpip1 | Proline-serine-threonine phosphatase-interacting | 91.15 | 3.61 | 47590 | 2 | 3940538 | 3511255.5 | 3640446 | 1181537.25 | 3557181 | 5384993.5 | 4490029 | 1121387.62 | 2010283.62 | 1404537.12 | 4179157 | 4451761 | 4221905 | 3390994 | 1878637.5 | 3879561.25 | 5380874 | 1746798.25 | 4389203 | 5016531.5 |
| P17225 | Ptbp1 | Polypyrimidine tract-binding protein 1 | 1343.53 | 43.78 | 59322 | 18 | 6566870.5 | 13413412 | 8587910 | 5314858.5 | 12171742 | 11821474 | 9488993 | 9307988 | 12774328 | 11428248 | 9967194 | 10576291 | 11677013 | 14089212 | 11476751 | 4759626.5 | 10853822 | 8745592 | 5535115.5 | 10159310 |
| Q60866 | Pter | N-acetyltaurine hydrolase | 125.04 | 10.32 | 39218 | 3 | 2546670.75 | 2160554.5 | 2509130.5 | 1176627.38 | 1022597.44 | 2992321 | 2664542.5 | 3434723.5 | 501018.16 | 1066893.38 | 3909899.5 | 3756772 | 876206.06 | 1239880.62 | 3195890.5 | 2734070.75 | 3972678 | 2871501.25 | 4861810.5 | 4246259.5 |
| P83868 | Ptges3 | Prostaglandin E synthase 3 | 51.61 | 6.25 | 18721 | 1 | 2 | 656504.25 | 2 | 1551025.62 | 102868.09 | 2 | 1480454.5 | 219900.84 | 2393672 | 278529.16 | 835522.75 | 1504469.88 | 1545162.38 | 1018009.69 | 2 | 2 | 2 | 2 | 101723.7 | 2 |
| Q91YR9 | Ptgr1 | Prostaglandin reductase 1 | 291.6 | 19.45 | 35560 | 6 | 17711692 | 13418068 | 19829488 | 21647038 | 14866646 | 11929766 | 9361463 | 13955731 | 9984326 | 6961012 | 16566125 | 15422720 | 16921724 | 13869510 | 17321282 | 9986567 | 12523092 | 13666972 | 12464622 | 10465027 |
| Q5BK81 | Ptgr2 | Prostaglandin reductase 2 | 53.63 | 2.56 | 38136 | 1 | 2206260.75 | 2 | 2195399.5 | 774138.06 | 2 | 1865224.25 | 1294260.62 | 2 | 1228187.75 | 1250365.38 | 1923989.38 | 1569614.5 | 1705685.75 | 1611381.75 | 1806320.12 | 1433982 | 1457684.75 | 1425946 | 1736664.38 | 1957654.62 |
| P22437 | Ptgs1 | Prostaglandin G/H synthase 1 | 335.42 | 7.31 | 69042 | 6 | 4254288 | 964790.38 | 894098.75 | 3393250.25 | 6595196 | 796030.88 | 727234.62 | 2 | 2926460.25 | 1157681.12 | 8709288 | 7731776.5 | 7675873 | 7339276 | 5493212.5 | 6829953 | 5848870 | 6737907 | 6119445.5 | 4208349.5 |
| P35355 | Ptgs2 | Prostaglandin G/H synthase 2 | 122.48 | 4.8 | 69164 | 2 | 2 | 2 | 123462.49 | 2 | 2 | 2 | 2 | 2 | 119883.54 | 97958.75 | 2 | 2 | 77647.34 | 2 | 2 | 2 | 2 | 2 | 253339.36 | 222969.05 |
| P70600 | Ptk2b | Protein-tyrosine kinase 2-beta | 236.51 | 5.15 | 115784 | 5 | 5326828.5 | 3691807.25 | 5174757 | 3296552 | 3807096.5 | 3186599.75 | 3570607.5 | 3471881.25 | 2640326 | 4234343 | 2392816 | 1542899.88 | 2736885 | 1869624.12 | 4568742 | 1511583.5 | 2629407.75 | 2716611.5 | 1680626.38 | 3361890.5 |
| P26350 | Ptma | Prothymosin alpha | 381.76 | 36.94 | 12254 | 5 | 7380734.5 | 11752043 | 1342108.62 | 22838258 | 10809984 | 7397657 | 6961656 | 5628428.5 | 9556914 | 6330641.5 | 3121441.5 | 2695824 | 2908587.5 | 2889531 | 3098528 | 15931060 | 9686029 | 2126157.5 | 16743354 | 7331489.5 |
| P04550 | Ptms | Parathymosin | 174.3 | 22.55 | 11559 | 3 | 7954525.5 | 7452682 | 5778362.5 | 10666569 | 4848247 | 3753045 | 4599719.5 | 5010921.5 | 5464039 | 3488119.75 | 366153.81 | 4823141 | 6348166.5 | 4566379.5 | 1966014.5 | 18528574 | 14285151 | 5459394.5 | 27349028 | 11397479 |
| P58389 | Ptpa | Serine/threonine-protein phosphatase 2A activato | 132.85 | 12.38 | 36710 | 3 | 4224244.5 | 5166914 | 5568233 | 3581562 | 3818065.25 | 3062783.25 | 2352178.25 | 153137.73 | 2468467.25 | 2775355.75 | 2148700 | 2399475.75 | 2887048 | 2655521.25 | 2379607.25 | 3122168.25 | 2554408 | 3590522.75 | 2604543.25 | 3703821 |
| P20417 | Ptpn1 | Tyrosine-protein phosphatase non-receptor type 1 | 57.9 | 2.55 | 49674 | 1 | 2 | 2 | 1193777.5 | 162192.77 | 552062.88 | 2 | 158296.48 | 138223.72 | 314122.47 | 578539.31 | 2 | 2 | 425242.84 | 207422.55 | 1114260.75 | 2 | 169738.62 | 180614.91 | 2 | 350115.38 |
| P35831 | Ptpn12 | Tyrosine-protein phosphatase non-receptor type 1 | 50.28 | 1.68 | 86526 | 1 | 2 | 2 | 2 | 627883.81 | 2 | 2 | 2 | 859190.94 | 654312.88 | 2 | 1519860.38 | 982521.12 | 867352.06 | 694732.69 | 775673 | 974136.56 | 2 | 2 | 1126707.62 | 861038.56 |
| P29351 | Ptpn6 | Tyrosine-protein phosphatase non-receptor type ( | 944.89 | 33.95 | 67559 | 19 | 20231650 | 14216926 | 8814796 | 22256474 | 29147748 | 10535582 | 7758245.5 | 6373031.5 | 15217144 | 10443730 | 15756485 | 17622762 | 18024648 | 16523574 | 12507350 | 17870342 | 13281586 | 10245458 | 19283514 | 14715093 |
| P18052 | Ptpra | Receptor-type tyrosine-protein phosphatase alpha | 170.25 | 5.79 | 93698 | 4 | 1966286.25 | 846486 | 447440.03 | 920831.25 | 1626633.12 | 367568.28 | 293277.81 | 1586963.12 | 2068535.88 | 1231232.5 | 2 | 628979.56 | 733117.5 | 135862.77 | 612209.25 | 219235.81 | 2 | 2 | 2 | 2 |
| P06800 | Ptprc | Receptor-type tyrosine-protein phosphatase C | 1081.34 | 17.79 | 144837 | 21 | 16766360 | 12470701 | 15171073 | 16895154 | 14657508 | 10843017 | 15091731 | 12440374 | 23773580 | 25905020 | 20285554 | 16500635 | 23092574 | 21097316 | 22074420 | 21058510 | 20783082 | 14591412 | 16320477 | 13995069 |
| Q8R2Y8 | Ptrh2 | Peptidyl-tRNA hydrolase 2, mitochondrial | 83.79 | 7.73 | 19527 | 1 | 2 | 2 | 2 | 2 | 2 | 2 | 2 | 2 | 2 | 2 | 2 | 2 | 2 | 2 | 2 | 2 | 2 | 2 | 2 | 2 |
| Q8R143 | Pttg1ip | Pituitary tumor-transforming gene 1 protein-intera | 114.5 | 10.34 | 19965 | 3 | 992467.12 | 361268.44 | 275756.28 | 329793.97 | 2 | 969399.69 | 2 | 78254.55 | 1052862.88 | 1153826.25 | 2 | 2 | 573087.62 | 2 | 487931.09 | 342211.91 | 1332688 | 2 | 2 | 1181198.25 |
| Q3UEB3 | Puf60 | Poly(U)-binding-splicing factor PUF60 | 408.66 | 19.86 | 60249 | 7 | 4766328.5 | 3952435.25 | 5714857 | 6682920.5 | 8402830 | 985633.88 | 3320211.25 | 5241498 | 4617472.5 | 5980181.5 | 2536850.5 | 2881044.5 | 3492566 | 3378124.25 | 3738603.5 | 1795852.25 | 661964.38 | 1330883.25 | 1721580.25 | 2 |
| O35295 | Purb | Transcriptional regulator protein Pur-beta | 116.55 | 8.33 | 33901 | 2 | 578366.5 | 2 | 2 | 2 | 2 | 577097.19 | 2 | 2 | 1559987.5 | 2 | 533086.81 | 1509985 | 639035.25 | 741561.31 | 2 | 795318.69 | 2 | 2 | 2 | 2 |
| Q8BU03 | Pwp2 | Periodic tryptophan protein 2 homolog | 81.46 | 2.5 | 102910 | 2 | 2 | 186877.41 | 112737.7 | 220082.55 | 218436.41 | 2 | 224824.81 | 106023.39 | 190055.67 | 649237.25 | 2 | 2 | 133302.95 | 138271.88 | 231957.2 | 1431936 | 2 | 2 | 2 | 2 |
| Q9EPB4 | Pycard | Apoptosis-associated speck-like protein containi | 38.71 | 3.63 | 21459 | 1 | 2 | 2 | 504905.44 | 800896.19 | 1003927 | 2 | 2 | 967621.75 | 594294 | 1253907.75 | 2 | 2 | 2 | 2 | 2 | 2 | 2 | 2 | 2 | 2 |
| Q922Q4 | Pycr2 | Pyrroline-5-carboxylate reductase 2 | 754.87 | 41.56 | 33659 | 13 | 2259140.75 | 11155284 | 6179661.5 | 5320655.5 | 21574370 | 13011068 | 12292829 | 14632193 | 24134154 | 18611184 | 13701379 | 14183985 | 12834390 | 16381444 | 6634786 | 7249359 | 5963582 | 5241608.5 | 10382663 | 4752399 |
| Q5PQJ6 | Pycr3 | Pyrroline-5-carboxylate reductase 3 | 42.11 | 4.01 | 28878 | 1 | 2 | 2 | 2 | 2 | 1221492.62 | 2 | 2 | 2 | 991524.62 | 2 | 2 | 2 | 2 | 2 | 2 | 2 | 2 | 2 | 2 | 2 |
| P53534 | Pygb | Glycogen phosphorylase, brain form (Fragment) | 275.93 | 8.23 | 96174 | 6 | 1405950.75 | 177788.25 | 2451044 | 1308130.12 | 738744.75 | 361628.41 | 1031798.75 | 1914042 | 803341.88 | 1144766.12 | 3126257.5 | 2966767 | 4143328.75 | 3554731 | 3304647 | 2050242.25 | 1429011.88 | 2090705.12 | 1779037.25 | 1517154.5 |
| Q8BML9 | Qars1 | Glutamine--tRNA ligase | 248.89 | 9.16 | 87677 | 6 | 382517.28 | 2 | 1134130.5 | 1931822.38 | 1593221.75 | 2075795.75 | 2063766.88 | 3081453 | 2708368 | 1072202.75 | 2943539.25 | 3539028 | 4154257.5 | 4735130 | 2986355.5 | 3575042.5 | 4044856.25 | 2637241.25 | 5361362.5 | 2 |
| Q8BVI4 | Qdpr | Dihydropteridine reductase | 115.66 | 16.6 | 25570 | 2 | 2593049.25 | 2643115.25 | 1774431 | 1446085.62 | 4633658 | 3444702.75 | 2787089.5 | 4782965.5 | 3264387 | 2270002 | 3616519 | 3362899.5 | 2888641.5 | 3212627 | 1240414 | 1138563.88 | 1366967 | 2 | 1577059.12 | 209278.45 |
| Q91XU1 | Qki | KH domain-containing RNA-binding protein QKI | 251.36 | 22.29 | 37643 | 5 | 2396804.5 | 953609.12 | 4428006 | 1113638.12 | 5037858 | 404374.25 | 1352914.62 | 768709 | 2457637.25 | 3643672.75 | 932696.62 | 816901.88 | 2915621.25 | 2590670 | 2676651 | 250068.88 | 404890.34 | 1104973.75 | 186819.61 | 775000 |
| P61027 | Rab10 | Ras-related protein Rab-10 | 280.56 | 18 | 22541 | 5 | 2 | 2 | 261990.42 | 179260.44 | 143597.61 | 248398.44 | 415250.28 | 2 | 229325.19 | 716744.5 | 2 | 2 | 149845.7 | 207822.39 | 1413203.38 | 2 | 222761.8 | 184438.25 | 2 | 2 |
| O35509 | Rab11b | Ras-related protein Rab-11B | 301.61 | 27.98 | 24489 | 6 | 18020014 | 20952346 | 23874046 | 19838142 | 18319004 | 18860698 | 16583117 | 26296402 | 19928304 | 17078214 | 18678726 | 16859358 | 18736094 | 19840666 | 19575654 | 23916982 | 18327190 | 21517090 | 24642548 | 22944888 |
| P35283 | Rab12 | Ras-related protein Rab-12 | 158.28 | 12.35 | 27329 | 3 | 2626988.25 | 2274646.5 | 1479324.88 | 1650430.88 | 1640128.75 | 2 | 2 | 2 | 2 | 2 | 2 | 2 | 2 | 2 | 2 | 2583009.5 | 2 | 1630908.12 | 2 | 1270759.38 |
| Q91V41 | Rab14 | Ras-related protein Rab-14 | 891.23 | 76.28 | 23897 | 16 | 27776224 | 32426526 | 32192920 | 23327358 | 19961586 | 19227080 | 21799252 | 15683999 | 21269958 | 17139804 | 15968152 | 17232982 | 16460083 | 15416712 | 15020684 | 15331187 | 17008326 | 19564536 | 16123711 | 19719576 |
| P35293 | Rab18 | Ras-related protein Rab-18 | 142.37 | 12.14 | 23035 | 3 | 1982408.88 | 3351281 | 1676044.38 | 3205183.5 | 2742133 | 2368614.25 | 2726549.5 | 3050193.5 | 2495614 | 580883.75 | 1756530.5 | 1128811.88 | 817412.88 | 1914667.12 | 1040340.56 | 1856058.5 | 1291670.25 | 2253934 | 1036195.25 | 1054943.12 |
| P62821 | Rab1a | Ras-related protein Rab-1A | 881.07 | 52.2 | 22678 | 13 | 1085382.5 | 670715.62 | 1800761.62 | 2902218.5 | 2025087.62 | 4882745 | 5294436.5 | 978946.62 | 2742710.75 | 5888267 | 3175549.75 | 2175311.75 | 4424819 | 5082927 | 8261834.5 | 1462007.5 | 3745409 | 3896757.25 | 3015960.75 | 6673385 |
| P10536 | Rab1b | Ras-related protein Rab-1B | 883.33 | 59.2 | 22163 | 12 | 1586494.5 | 3691010.5 | 1542850.88 | 5189332 | 1914249.75 | 1629560 | 1291572 | 1158264 | 4218666 | 4380149.5 | 2595312.5 | 3130302.5 | 2834470 | 4014334 | 2941324 | 3002560.25 | 2421011.5 | 2905456.25 | 774520.12 | 3836961 |
| P35282 | Rab21 | Ras-related protein Rab-21 | 202.18 | 16.22 | 24106 | 3 | 632974.06 | 2 | 2 | 445992.62 | 636770.56 | 1579131.88 | 2124291 | 878925.19 | 2652738.25 | 547475.62 | 560545.5 | 3167582 | 1172710.75 | 1920236.88 | 2 | 3383772 | 1393817.88 | 742670.75 | 2284037 | 2 |
| P53994 | Rab2a | Ras-related protein Rab-2A | 464.87 | 40.57 | 23548 | 7 | 8909061 | 4740250 | 8186096 | 5996357.5 | 4804821 | 4567540 | 7105874 | 6010226.5 | 5734635 | 3378256.5 | 4468920 | 3393213.25 | 5103786 | 4168313.75 | 4240518.5 | 1577990.88 | 5944733 | 5854949.5 | 1125187.88 | 4062867.5 |
| Q921E2 | Rab31 | Ras-related protein Rab-31 | 182.72 | 17.95 | 21463 | 3 | 39678.73 | 1104559.5 | 391929.97 | 2719710.5 | 260693.28 | 2158243.5 | 2667095.75 | 75893.28 | 3631697.5 | 2369412.25 | 1502854.88 | 1411842.88 | 290015.09 | 641526.94 | 818094.25 | 2 | 309356.69 | 464962.22 | 2 | 475731.38 |
| Q5U316 | Rab35 | Ras-related protein Rab-35 | 232.73 | 23.88 | 23025 | 4 | 1522069.38 | 2 | 1414522.38 | 930682.5 | 882623.62 | 131598.12 | 81042.71 | 119486.81 | 779065.56 | 1206140 | 1162394 | 631650.12 | 140025.09 | 2 | 82019.06 | 84472.23 | 184946.48 | 946042.38 | 2 | 2 |
| Q80UJ7 | Rab3gap1 | Rab3 GTPase-activating protein catalytic subunit | 108.01 | 3.87 | 110198 | 2 | 2 | 2 | 59173.42 | 41830.62 | 2 | 109025.05 | 97786.14 | 2 | 144013.31 | 83898.16 | 2 | 2 | 41728.95 | 2 | 2 | 2 | 2 | 2 | 2 | 2 |
| Q8BMG7 | Rab3gap2 | Rab3 GTPase-activating protein non-catalytic sub | 350.93 | 6.3 | 152535 | 8 | 1088695.75 | 836754.75 | 1213048.88 | 1924417.75 | 6609014 | 2289412.75 | 2057793 | 4525728 | 5948276 | 3912778.5 | 3579122 | 2192068.75 | 3925372.25 | 3116820.75 | 1945004.25 | 785122.56 | 1674645.75 | 575894.06 | 1394490.88 | 2 |
| P51146 | Rab4b | Ras-related protein Rab-4B | 129.89 | 11.27 | 23629 | 2 | 2 | 2 | 812492.81 | 2 | 567618.94 | 2 | 2 | 937618.56 | 2 | 2 | 1322533 | 1091398.25 | 1419704.75 | 972347.62 | 2 | 1050800.88 | 1244114.12 | 1530605 | 2 | 1163045.25 |
| Q9CQD1 | Rab5a | Ras-related protein Rab-5A | 358.19 | 32.56 | 23599 | 5 | 4357194 | 3123403 | 3922044.5 | 2583001.75 | 3846518 | 3940149 | 3242784.75 | 5020854.5 | 4242281 | 4016694 | 5644824 | 3456397.25 | 6064181 | 5208051 | 3983280.5 | 5994463 | 6082795 | 5220016.5 | 6034032 | 5862048.5 |
| P61021 | Rab5b | Ras-related protein Rab-5B | 364.8 | 32.56 | 23707 | 5 | 4209256 | 3070636.75 | 3943458.75 | 3187236.5 | 3679796 | 2806360.5 | 2811937.25 | 3310390 | 4091918.25 | 3319951 | 2819535 | 2999933.5 | 4112323 | 4029597.5 | 2630855.5 | 2620722 | 3691745.75 | 2479773.75 | 1976523.5 | 1856057.5 |
| P35278 | Rab5c | Ras-related protein Rab-5C | 621.61 | 52.31 | 23413 | 9 | 16628199 | 12852591 | 16827908 | 12521144 | 17946436 | 13745016 | 11505303 | 15816027 | 17517078 | 14721069 | 15978922 | 14470770 | 18428096 | 17003676 | 10714292 | 14253438 | 14381485 | 12564304 | 8426455 | 14416369 |
| P35279 | Rab6a | Ras-related protein Rab-6A | 546.31 | 52.4 | 23590 | 11 | 2632457 | 3880841.25 | 4454890 | 3601306 | 6016036 | 7392678 | 7431077 | 8761715 | 8401115 | 4769079 | 8062133 | 8483434 | 9266088 | 5584672.5 | 2310469.5 | 5209470 | 2955822 | 2 | 5848074.5 | 873036 |
| P09527 | Rab7a | Ras-related protein Rab-7a | 969.54 | 77.78 | 23504 | 16 | 79725794 | 69026698 | 86186418 | 51292674 | 46882006 | 54870722 | 58315638 | 64479886 | 51150698 | 59932214 | 40090334 | 31785804 | 35977722 | 34465758 | 45270050 | 28003788 | 33135716 | 34217250 | 16345882 | 31797758 |
| Q8VEA8 | Rab7b | Ras-related protein Rab-7b | 53.45 | 7.54 | 22502 | 1 | 2 | 2 | 2 | 2 | 2 | 2 | 2 | 2 | 82797.85 | 84103.37 | 2 | 112739.62 | 2 | 2 | 2 | 166760.34 | 2 | 2 | 2 | 2 |
| P35280 | Rab8a | Ras-related protein Rab-8A | 495.68 | 36.71 | 23668 | 9 | 3405819.5 | 3422811 | 989721 | 3700325.25 | 70418.7 | 2720699 | 2027597.62 | 283630.56 | 1958161.5 | 1011465.38 | 1674592.25 | 1745788.88 | 1723530.62 | 1098320.38 | 2201020.5 | 2519571.75 | 3481492.5 | 3650754.25 | 3051012 | 3385674.75 |
| P61028 | Rab8b | Ras-related protein Rab-8B | 714.04 | 57 | 23603 | 12 | 8390834 | 1955209.62 | 3349147.5 | 3025123 | 3690803 | 1920358.5 | 4776462.5 | 3658205.5 | 4292300 | 6636747 | 897825.25 | 1249371.12 | 4420904.5 | 2133373.5 | 5909894.5 | 810848.5 | 2946356.25 | 3273952.5 | 1184897.75 | 1859450.5 |
| Q99P75 | Rab9a | Ras-related protein Rab-9A | 60.44 | 5.47 | 22896 | 1 | 2 | 2 | 2 | 2 | 738081.88 | 1271576.38 | 1024696.38 | 1192069 | 1538411 | 821994.69 | 476414.12 | 2 | 2 | 2 | 2 | 2 | 2 | 2 | 2 | 2 |
| P63001 | Rac1 | Ras-related C3 botulinum toxin substrate 1 | 478.58 | 43.23 | 21450 | 8 | 8483855 | 12045962 | 9033918 | 10445743 | 14131166 | 12740633 | 14071212 | 17920160 | 15938792 | 15500498 | 6386924 | 9073828 | 10621540 | 8377821.5 | 7650860 | 6102174 | 9070451 | 7672163 | 9242806 | 8459462 |
| Q05144 | Rac2 | Ras-related C3 botulinum toxin substrate 2 | 585.76 | 51.04 | 21441 | 10 | 39023250 | 23433796 | 49348746 | 40920414 | 47268118 | 17107984 | 28423222 | 14257121 | 27814918 | 44500646 | 12744312 | 19105054 | 26331566 | 24491268 | 38453430 | 10082582 | 21694996 | 20610202 | 13271942 | 25109514 |
| Q9WVM1 | Racgap1 | Rac GTPase-activating protein 1 | 44.35 | 1.43 | 70158 | 1 | 848222.5 | 1213314.5 | 781085 | 1432927.25 | 1399658.75 | 15913869 | 15877595 | 897843.12 | 811024.56 | 671306.25 | 13053812 | 849651.81 | 14226790 | 15550331 | 11969865 | 17431116 | 20064176 | 24184836 | 578811.75 | 21840286 |
| P63245 | Rack1 | Small ribosomal subunit protein RACK1 | 845.81 | 49.84 | 35077 | 16 | 10371467 | 12854588 | 18832582 | 19504192 | 29666956 | 17250666 | 16205370 | 28957430 | 22314444 | 29341882 | 41535650 | 36989786 | 40432790 | 36327738 | 40458218 | 27196768 | 21733310 | 21972930 | 24499766 | 17192106 |
| P54728 | Rad23b | UV excision repair protein RAD23 homolog B | 37.99 | 4.57 | 43513 | 1 | 2 | 2 | 2 | 2 | 101839.91 | 2 | 147461.17 | 34322.36 | 2 | 262302.62 | 2 | 2 | 2 | 83056.4 | 2 | 2 | 2 | 2 | 2 | 2 |
| P70388 | Rad50 | DNA repair protein RAD50 | 101.03 | 2.13 | 153488 | 2 | 544858.81 | 2 | 446535.84 | 230770.08 | 892707.31 | 2 | 2 | 2 | 2 | 371288.22 | 2 | 2 | 2 | 153845.45 | 2 | 53016.43 | 2 | 146387.02 | 115283.59 | 2 |
| Q3SWS8 | Rae1 | mRNA export factor | 101.64 | 5.98 | 40919 | 2 | 976807.69 | 1165924.88 | 885898 | 1904314.75 | 1218675.12 | 1401252.12 | 2 | 1883479.88 | 1164723.12 | 976681.31 | 1417738.62 | 1222749.88 | 1354607.62 | 1191767.38 | 1348507.88 | 2 | 2 | 2 | 2 | 1503085.75 |
| Q9JIW9 | Ralb | Ras-related protein Ral-B | 167.89 | 23.3 | 23349 | 4 | 1534514.12 | 1066021.62 | 2 | 576264.44 | 2782438.25 | 2041333.88 | 1907062.25 | 3881014.75 | 3191628.5 | 1321138.75 | 3114538 | 3713156.5 | 3050730.75 | 2465066.5 | 1824183.62 | 2465761 | 3002530 | 3466646.5 | 391160.66 | 2717969 |
| Q64012 | Raly | RNA-binding protein Raly | 376.61 | 23.4 | 33188 | 7 | 1190788.25 | 2747353.25 | 879592.56 | 3567685.25 | 6876947.5 | 2583945 | 2267174 | 2942757.75 | 5645655 | 5213181 | 6815219 | 7450986.5 | 6559574 | 6846773 | 2817028.5 | 6912491 | 5728356.5 | 3043646.25 | 8497124 | 502225.06 |
| P62827 | Ran | GTP-binding nuclear protein Ran | 535.95 | 42.59 | 24423 | 9 | 5505285 | 7219234.5 | 5005873 | 16206653 | 25380134 | 13501386 | 14548178 | 21171520 | 23029562 | 11689674 | 28474954 | 29486334 | 26286854 | 19875018 | 18131204 | 22674796 | 16909250 | 13136974 | 14219792 | 11499051 |
| P34022 | Ranbp1 | Ran-specific GTPase-activating protein | 183.44 | 15.27 | 23596 | 4 | 1420200.5 | 3561573.5 | 1699603.38 | 759469.19 | 1445812.75 | 2 | 1776315.38 | 1102145.88 | 2899010 | 1232289 | 14096292 | 15866154 | 11501342 | 12312088 | 8028302.5 | 13143032 | 8997118 | 8945227 | 14921499 | 9369192 |
| Q9ERU9 | Ranbp2 | E3 SUMO-protein ligase RanBP2 | 826.75 | 7.01 | 341121 | 16 | 831895.88 | 1482306.75 | 938157.75 | 3255237.5 | 4961248 | 127232.87 | 3691200.5 | 2203038.5 | 2554807 | 2814239.75 | 3383760.25 | 4163661.5 | 2226226 | 2904681.5 | 1276552.62 | 2944435.25 | 2 | 3409396.75 | 617533.81 | 1354301 |
| Q9CT10 | Ranbp3 | Ran-binding protein 3 | 143.99 | 5.7 | 52573 | 2 | 2 | 701633.56 | 644010.06 | 527750 | 1570989 | 2 | 2 | 64170.7 | 2 | 2 | 2087794.25 | 2300686 | 1483681.88 | 1747674.12 | 648965.94 | 752009.62 | 2 | 992877.31 | 1144557.75 | 1183675.62 |
| P46061 | Rangap1 | Ran GTPase-activating protein 1 | 481.99 | 20.54 | 63531 | 10 | 9060379 | 6495795 | 7827281 | 7722898 | 6361215 | 1926580.75 | 2586222.5 | 2292656.25 | 2893515 | 4771926 | 2518570.5 | 1173641.75 | 1092664.38 | 2043450.25 | 1185948.38 | 1744415.88 | 2525018.75 | 178639.88 | 1270313.75 | 1076775.12 |
| P62835 | Rap1a | Ras-related protein Rap-1A | 728.78 | 81.52 | 20987 | 12 | 949879.81 | 1459470.62 | 2549508 | 571714.12 | 1267525.75 | 2 | 652047.56 | 877320.56 | 1560718.25 | 1447047.25 | 552164.88 | 2026826 | 1635921 | 2248499.5 | 1162375.62 | 2 | 2 | 1595975.38 | 2 | 2 |
| Q99JI6 | Rap1b | Ras-related protein Rap-1b | 825.45 | 73.37 | 20825 | 12 | 3126311.5 | 4679919 | 4763400 | 2256859.5 | 3122327.25 | 2228273.75 | 1825587.62 | 3047306 | 3337706 | 3260307.25 | 2298165.75 | 2 | 2450010.25 | 2929904.25 | 1754485.25 | 1366606.88 | 3277284 | 1441596.5 | 1783011 | 2 |
| P61226 | Rap2b | Ras-related protein Rap-2b | 37.34 | 6.01 | 20504 | 1 | 2 | 2 | 2 | 2 | 540964.25 | 2213267 | 1847017.75 | 1605740.88 | 2841538.75 | 2 | 2075173.25 | 3586812.75 | 1868532.62 | 2259363.25 | 2 | 2224983.25 | 1511903.12 | 2 | 3421758.25 | 2 |
| Q9D0I9 | Rars1 | Arginine--tRNA ligase, cytoplasmic | 1058.74 | 38.48 | 75674 | 20 | 48614582 | 51505734 | 49053106 | 37385142 | 39725690 | 32282798 | 44173722 | 37117490 | 27210790 | 40540742 | 44280298 | 41805170 | 46465646 | 46212534 | 49696498 | 40325850 | 53214326 | 65404618 | 48113742 | 49671950 |
| Q3U186 | Rars2 | Probable arginine--tRNA ligase, mitochondrial | 58.32 | 2.94 | 65336 | 1 | 2 | 126087.6 | 2 | 166780.78 | 55772.28 | 109724.16 | 277962.31 | 100663.69 | 128551.65 | 110815.06 | 2 | 80438.75 | 85544.65 | 115322.6 | 2 | 2 | 2 | 2 | 2 | 2 |
| Q60790 | Rasa3 | Ras GTPase-activating protein 3 | 40.66 | 1.32 | 95987 | 1 | 2 | 2 | 2 | 2 | 169994.98 | 2 | 2 | 59870.16 | 167690.25 | 2 | 2 | 2 | 2 | 2 | 431506.03 | 2 | 2 | 2 | 2 | 2 |
| Q5SSG5 | Rasl10b | Ras-like protein family member 10B | 38.73 | 3.45 | 23229 | 1 | 2017193.5 | 1597060.88 | 854002.94 | 2010561.88 | 2159457.75 | 1959178.5 | 1477956.62 | 2442007 | 1056055.5 | 1585512 | 1362519.62 | 1317274.12 | 934627 | 1393982.5 | 1028452.94 | 1894398.5 | 2002618.88 | 1523850.38 | 1409352 | 2407032.25 |
| Q99P51 | Rassf3 | Ras association domain-containing protein 3 | 30.36 | 3.88 | 26749 | 1 | 1961580.25 | 11302054 | 5423024.5 | 4656721 | 11678417 | 16641928 | 12031278 | 10359205 | 7780905.5 | 7457339 | 8007595.5 | 7715265 | 8065749 | 8826185 | 7463982 | 9304032 | 7837206.5 | 9300010 | 603970.5 | 8141245 |
| Q5XI28 | Raver1 | Ribonucleoprotein PTB-binding 1 | 75.76 | 2.14 | 79289 | 1 | 2 | 2 | 2 | 371689.03 | 229932.91 | 2 | 2 | 2 | 339767 | 2 | 2 | 2 | 2 | 2 | 2 | 2 | 2 | 2 | 2 | 2 |
| Q60972 | Rbbp4 | Histone-binding protein RBBP4 | 283.69 | 12 | 47656 | 6 | 2 | 2 | 2 | 2 | 2 | 2 | 2 | 2 | 2 | 2 | 719869 | 559240.69 | 2 | 2 | 2 | 960177.19 | 2 | 965841.69 | 2 | 2 |
| Q8BX09 | Rbbp5 | Retinoblastoma-binding protein 5 | 72.3 | 2.23 | 59098 | 1 | 1366367.62 | 1144419 | 2 | 583091.06 | 2 | 713999 | 2 | 2 | 652955.69 | 2 | 1093877.88 | 899126.75 | 856124.94 | 655509.88 | 2 | 1266253.75 | 2 | 510846.97 | 355085.06 | 1065010.5 |
| Q60973 | Rbbp7 | Histone-binding protein RBBP7 | 329.76 | 11.76 | 47790 | 7 | 621361.81 | 2 | 422965 | 465353.84 | 600198.44 | 2 | 2 | 429560 | 574489.19 | 2 | 673837.94 | 590005.94 | 537751.69 | 332902.06 | 2 | 1694900.25 | 2 | 693957.69 | 2 | 678602.88 |
| Q8C2Q3 | Rbm14 | RNA-binding protein 14 | 572.65 | 18.98 | 69449 | 11 | 387683.72 | 1151487 | 805170.56 | 920492.88 | 2944652.5 | 3275644 | 2 | 2 | 900353.44 | 1277022.12 | 7480181 | 6802167 | 2050537.88 | 5843940 | 6195020.5 | 4684507.5 | 7674811.5 | 2659742.5 | 4229229 | 3148073.75 |
| Q0VBL3 | Rbm15 | RNA-binding protein 15 | 87.59 | 2.49 | 105722 | 2 | 2 | 2 | 2 | 2 | 2 | 2 | 2 | 2 | 2 | 2 | 2 | 2 | 2 | 2 | 2 | 2 | 1469389.88 | 2 | 2 | 2 |
| Q8JZX4 | Rbm17 | Splicing factor 45 | 37.05 | 2.22 | 45304 | 1 | 2 | 2 | 2 | 2 | 2 | 2 | 2 | 2 | 2 | 465345.22 | 825575.25 | 688392.06 | 2 | 2 | 2 | 2 | 2 | 2 | 2 | 2 |
| Q8R3C6 | Rbm19 | Probable RNA-binding protein 19 | 53.71 | 1.79 | 106083 | 1 | 2 | 2 | 2 | 90917.68 | 49580.44 | 2 | 2 | 75327.92 | 64514.12 | 37814.43 | 2 | 2 | 2 | 2 | 2 | 2 | 2 | 2 | 2 | 2 |
| Q4V7D7 | Rbm22 | Pre-mRNA-splicing factor RBM22 | 43.97 | 2.86 | 46864 | 1 | 2 | 2 | 2 | 2 | 732087.06 | 756075.62 | 2 | 2 | 566204.56 | 2 | 2 | 2 | 2 | 2 | 2 | 2 | 2 | 2 | 2 | 2 |
| Q8CGC6 | Rbm28 | RNA-binding protein 28 | 424.83 | 12.4 | 84214 | 8 | 1164450.75 | 1656422.25 | 1028358.88 | 2068719.12 | 1294689.12 | 541181.06 | 780168.44 | 195353.33 | 1373370.5 | 403385.22 | 2 | 778303.12 | 1182585.5 | 2 | 2 | 1510568.75 | 1194130.12 | 2 | 1362446.75 | 1212964.88 |
| Q8VH51 | Rbm39 | RNA-binding protein 39 | 457.69 | 17.36 | 59407 | 7 | 5131040.5 | 7073948 | 4663162.5 | 4668773.5 | 30362858 | 20692288 | 5287005.5 | 11131751 | 7572094 | 6633889 | 4519529.5 | 47347234 | 6859152 | 5159838.5 | 5246291.5 | 3723938 | 4432894 | 6085584 | 5890310 | 4648168 |
| P31266 | Rbpj | Recombining binding protein suppressor of hairle | 125.58 | 6.46 | 58537 | 3 | 7877346 | 8637737 | 7622155 | 3414072.25 | 3637919.5 | 2018803.38 | 1837015 | 2845310.75 | 637271.25 | 3875305 | 7891402 | 8270128.5 | 8262607 | 6452590.5 | 7133798 | 6826605.5 | 7133913 | 5037718 | 6247341 | 5758854 |
| P62878 | Rbx1 | E3 ubiquitin-protein ligase RBX1 | 155.78 | 17.59 | 12274 | 2 | 2047950.5 | 3595936.25 | 2 | 4271128.5 | 3192988.5 | 3789163.5 | 2935184.5 | 2225623 | 2655914.5 | 2530382.75 | 2055403 | 2606034 | 2833760 | 2674672.25 | 2342940.5 | 1709691.25 | 2473725.5 | 1524031.62 | 959438.06 | 1571874.25 |
| Q8VE37 | Rcc1 | Regulator of chromosome condensation | 370.7 | 19.24 | 44931 | 6 | 1426123.25 | 3656546.5 | 2 | 821651.75 | 6677530 | 3715638 | 822364.5 | 7640391 | 4812464.5 | 649658.69 | 6512547 | 6425458.5 | 6769793 | 7056964 | 611056 | 6938236 | 4706872.5 | 4192247 | 1250071.75 | 2 |
| Q8BK67 | Rcc2 | Protein RCC2 | 656.84 | 27.88 | 55983 | 15 | 18679902 | 12644512 | 29987248 | 16064125 | 30034124 | 13618611 | 13345007 | 16746831 | 19110470 | 13341436 | 37597250 | 40215586 | 32937096 | 32199360 | 26459280 | 39448246 | 35778902 | 27829702 | 29880774 | 27557246 |
| Q3UZA1 | Rcsd1 | CapZ-interacting protein | 55.4 | 2.91 | 44114 | 1 | 1868649.12 | 1271674.5 | 1164003.5 | 1720249.62 | 1597736.25 | 949748.06 | 2 | 2 | 1048017.81 | 621006.5 | 1710370.25 | 1688982.5 | 1139033.25 | 970324.25 | 1415096 | 2 | 663504 | 1059547.25 | 2207542.5 | 1637489.12 |
| Q8BRE0 | Rd3 | Protein RD3 | 36.3 | 3.59 | 22669 | 1 | 2 | 2 | 2 | 2 | 2 | 2 | 253119.95 | 2 | 2 | 2 | 2 | 2 | 2 | 1117743.75 | 2 | 2 | 2 | 1157420.25 | 2 | 2 |
| Q9QYF1 | Rdh11 | Retinol dehydrogenase 11 | 314.99 | 17.41 | 35148 | 5 | 1543594.5 | 3327363.5 | 2622015.25 | 5415353 | 7413477 | 7415543.5 | 6036989 | 8135877 | 7696217 | 5089878 | 5658060 | 4660717.5 | 5059937 | 5451445.5 | 2524404.25 | 5661622.5 | 4851995.5 | 4977785 | 7023079.5 | 3901835 |
| P26043 | Rdx | Radixin | 650 | 17.5 | 68543 | 12 | 160319.66 | 287694.38 | 2 | 396393.78 | 2 | 780411 | 2 | 167516.27 | 1010395.5 | 2 | 1395009.88 | 2125258.25 | 117358.23 | 2 | 2 | 4789328.5 | 2 | 1970570.5 | 2 | 2 |

| | | | | | | | | | | | | | | | | | | | | | | | | | |
|---|---|---|---|---|---|---|---|---|---|---|---|---|---|---|---|---|---|---|---|---|---|---|---|---|---|---|
| Q75NR7 | Recql4 | ATP-dependent DNA helicase Q4 | 40.39 | 0.82 | 135124 | 1 | 8241386.5 | 5940245 | 2430759 | 8693360 | 1058191 | 5753093.5 | 6230314 | 287843.38 | 3618177.25 | 6031030.5 | 5726326.5 | 6750231.5 | 8173449.5 | 6660905 | 7527639.5 | 3130670.5 | 6398158 | 5499271 | 4534694 | 6033193 |
| B2RZ37 | Reep5 | Receptor expression-enhancing protein 5 | 37.6 | 5.82 | 21431 | 1 | 705024.75 | 1265663.62 | 973521.69 | 2 | 1763215 | 2 | 2 | 2 | 350312.91 | 1697287.75 | 2 | 743364 | 1031915.31 | 1141459.75 | 2 | 2 | 2 | 2 | 2 | 2 |
| P82343 | Renbp | N-acylglucosamine 2-epimerase | 78.85 | 3.58 | 48510 | 2 | 780861.31 | 1794934 | 2 | 2 | 441099.59 | 2 | 2 | 2 | 2 | 2 | 1169777.75 | 1093950.5 | 1279278.88 | 1428477.25 | 1484230.38 | 1019842.94 | 2422603 | 1657333.25 | 1768778 | 1874790.12 |
| Q64FW2 | Retsat | All-trans-retinol 13,14-reductase | 94.68 | 4.11 | 67334 | 2 | 2 | 2 | 185033.25 | 48246.53 | 1295942.12 | 2 | 2 | 2 | 2 | 2 | 2 | 553366.94 | 2 | 2 | 2 | 2 | 2 | 2 | 2 | 2 |
| Q5U1X1 | Rexo2 | Oligoribonuclease, mitochondrial | 166.64 | 8.86 | 26752 | 2 | 2 | 2 | 205243.66 | 465034.88 | 276006.53 | 460183.06 | 455469.75 | 2 | 471860.38 | 543516.31 | 87780.13 | 23232.46 | 103220.99 | 2 | 447958.5 | 2 | 275962.41 | 2 | 2 | 2 |
| P35601 | Rfc1 | Replication factor C subunit 1 | 64.61 | 0.71 | 125985 | 1 | 2 | 2 | 2 | 2 | 251240.94 | 2 | 2 | 2 | 2 | 2 | 519922.62 | 2 | 2 | 2 | 2 | 2 | 2 | 2 | 2 | 2 |
| Q641W4 | Rfc2 | Replication factor C subunit 2 | 131.59 | 10.6 | 38657 | 2 | 1217089.12 | 2 | 1617066.88 | 905994.25 | 1603698 | 2 | 155584.11 | 2 | 2 | 617984.94 | 135684.06 | 167472.16 | 2 | 2 | 1842232.62 | 99827.41 | 2 | 2 | 2 | 2 |
| Q99J62 | Rfc4 | Replication factor C subunit 4 | 69.9 | 6.32 | 39867 | 2 | 1854772.5 | 1711356.25 | 1251112.75 | 1710080.75 | 449712.97 | 793650.38 | 1126382.12 | 1358119.5 | 493849.06 | 1474097.25 | 1254235.75 | 1888428.5 | 1908031.75 | 214794.66 | 1978685.62 | 1700437.25 | 1035430.5 | 1831322.75 | 2 | 2 |
| Q9D0F6 | Rfc5 | Replication factor C subunit 5 | 179.43 | 15.34 | 38096 | 4 | 2342978 | 3776439.75 | 1071435 | 2612855 | 3507526.5 | 747363 | 773547.12 | 112331.38 | 1429829.5 | 1199849.25 | 965527.94 | 843725.25 | 788172.38 | 693701.62 | 1691632.62 | 2 | 2 | 783033.06 | 2 | 2 |
| Q6A0D4 | Rftn1 | Raftlin | 102.33 | 4.51 | 61537 | 2 | 2 | 413806.56 | 2 | 2 | 2 | 965515.75 | 2 | 125259.69 | 2 | 2 | 967382.81 | 1461391.88 | 1106706.88 | 2194061.25 | 2 | 2277511.75 | 2386748 | 1955327.62 | 460469.06 | 1767900 |
| P49806 | Rgs10 | Regulator of G-protein signaling 10 | 44.58 | 6.63 | 21176 | 1 | 49266.01 | 2 | 93207.04 | 2 | 41022.62 | 2 | 2 | 2 | 2 | 2 | 2 | 874732.31 | 2 | 2 | 2 | 2 | 199834.2 | 2 | 2 | 2 |
| Q80WQ6 | Rhbdf2 | Inactive rhomboid protein 2 | 69.64 | 2.18 | 93434 | 2 | 1132098.75 | 565846.81 | 2 | 512975.12 | 706543.06 | 633477.44 | 2 | 2 | 1205046.38 | 2 | 503411.66 | 542657.81 | 728890.56 | 337321.53 | 2 | 2098470 | 2341724.25 | 2250313 | 725566.31 | 3253935.25 |
| Q921J2 | Rheb | GTP-binding protein Rheb | 177.48 | 19.02 | 20451 | 3 | 3294488 | 1678780.38 | 2441696.25 | 942799.31 | 3646993 | 2101896.25 | 897753.94 | 3206874.5 | 1621972.62 | 2300976 | 984709.38 | 1144758.5 | 1620698.25 | 1730263.5 | 854947.38 | 1573475.5 | 1792844.75 | 2165081.5 | 1725838.12 | 2698049 |
| P61589 | Rhoa | Transforming protein RhoA | 297.15 | 31.09 | 21782 | 6 | 2592223 | 4862603.5 | 3271199.5 | 2471043.5 | 5148231 | 3056111.5 | 2792456.5 | 3909784 | 2639045.5 | 3003482.5 | 6349288 | 7087085 | 6154551.5 | 6617157.5 | 4288357.5 | 3418711.25 | 5056042 | 4792141 | 5320417.5 | 3987855.25 |
| Q62159 | Rhoc | Rho-related GTP-binding protein RhoC | 251.62 | 24.87 | 22006 | 5 | 1454328.62 | 2939105.75 | 1155096.75 | 2486241.75 | 2500522.75 | 2479925.75 | 2264504.5 | 3289144.25 | 2833124.25 | 2550940.25 | 5872372.5 | 5251390 | 5997114 | 5358010.5 | 4464569.5 | 6599858.5 | 7317321.5 | 6804928 | 8640946 | 6316176 |
| P84096 | Rhog | Rho-related GTP-binding protein RhoG | 432.66 | 39.79 | 21309 | 6 | 4128968 | 6394076 | 5561855.5 | 5902734 | 11323281 | 14046829 | 11699727 | 21998226 | 17465054 | 13723522 | 5971513.5 | 5890975 | 4942595 | 5697279 | 5187880 | 5104387 | 4806519.5 | 8536686 | 5691649 | 5060319.5 |
| Q8BG51 | Rhot1 | Mitochondrial Rho GTPase 1 | 95.84 | 3.01 | 72242 | 2 | 2 | 2 | 2 | 2 | 2 | 2 | 2 | 2 | 2 | 2 | 2 | 2 | 2 | 2 | 2 | 2 | 2 | 2 | 2 | 2 |
| P52760 | Rida | 2-iminobutanoate/2-iminopropanoate deaminase | 118.75 | 19.26 | 14255 | 2 | 3298926.75 | 5126001 | 2853971.5 | 957054.12 | 2752480 | 2957455.25 | 2780635.25 | 4091654.5 | 2339617.5 | 2174850.75 | 3452192 | 3419717 | 1834671.75 | 3299143.25 | 1139801.88 | 3699918 | 1347103.5 | 1130838.5 | 2597576 | 1312251.5 |
| Q8BR90 | Rimoc1 | RAB7A-interacting MON1-CCZ1 complex subunit | 44.29 | 3.06 | 33471 | 1 | 1085274.25 | 1699759.5 | 924101.81 | 1194533.12 | 690309.06 | 1587977.5 | 1375954.5 | 913492.75 | 720406.94 | 669081 | 356167.5 | 825181.94 | 1139359.88 | 1036506.19 | 979147.94 | 1587285.62 | 1230867 | 1934338.12 | 1194102.88 | 1748436.25 |
| D3ZU57 | Riox1 | Ribosomal oxygenase 1 | 76.64 | 3.35 | 67455 | 2 | 902081.12 | 2 | 816741.88 | 896279.44 | 696328.31 | 589876.31 | 459464.12 | 189728.06 | 484738.84 | 2 | 359236.19 | 2 | 2 | 607413.12 | 2 | 804842.62 | 793188.88 | 547652.12 | 886468.94 | 794954.94 |
| Q9QZL0 | Ripk3 | Receptor-interacting serine/threonine-protein kina | 108.53 | 4.32 | 53322 | 2 | 2297244.5 | 2472215.25 | 740860.88 | 2400400 | 348863.47 | 2169003.25 | 1884278.25 | 291261.38 | 1718475.88 | 1605768.88 | 2086452.25 | 1561678.75 | 1378968.88 | 2043820.12 | 1938328 | 1565331 | 2070935.5 | 1693552.12 | 2225699 | 1817657.5 |
| P13832 | Rlc-a | Myosin regulatory light chain RLC-A | 183.97 | 22.09 | 19895 | 4 | 2 | 2 | 2933549.5 | 330426.88 | 642517.38 | 2 | 2 | 184134.62 | 176772.11 | 2936141.5 | 855099.69 | 1257482.75 | 1548463.88 | 1562829.38 | 4395182 | 2 | 1115633.5 | 1979708.38 | 725527 | 2413586 |
| Q8K3C0 | Rnasek | Ribonuclease kappa | 43.93 | 9.18 | 10992 | 1 | 1830503.62 | 2225980.75 | 1022351.44 | 3190331.75 | 3035003 | 4920525 | 3787895.25 | 7113326.5 | 6150788.5 | 2 | 2590127.75 | 2512287 | 2415973.5 | 2389512.5 | 2520249.25 | 3951180 | 4680946.5 | 5950043 | 6594064 | 4349939.5 |
| Q9ET26 | Rnf114 | E3 ubiquitin-protein ligase RNF114 | 119.15 | 15.72 | 25745 | 3 | 2673515.75 | 3652657.75 | 1842243.5 | 872170.12 | 3477239 | 26633404 | 2426841.5 | 5625907 | 3933111.5 | 3177722.5 | 6119502.5 | 5999011 | 4673621.5 | 5000690 | 960534.88 | 3611543.75 | 4451444.5 | 4400076 | 6381224 | 4288013 |
| E9Q555 | Rnf213 | E3 ubiquitin-protein ligase RNF213 | 665.23 | 2.89 | 584258 | 15 | 6659683 | 7675634.5 | 7031556.5 | 7023087 | 6244417.5 | 5391055 | 6555351 | 8047311.5 | 3924602.75 | 7286084 | 3119949 | 2199626.25 | 2601293 | 3476051.5 | 791133.88 | 3125068.5 | 6071789.5 | 6922080 | 5614230.5 | 4385696.5 |
| Q91VI7 | Rnh1 | Ribonuclease inhibitor | 1368.47 | 61.4 | 49816 | 20 | 36476858 | 21892854 | 47520718 | 17064834 | 29254578 | 14721566 | 13281122 | 21712958 | 11221163 | 38547110 | 14872697 | 7819582.5 | 19635922 | 17172792 | 28620860 | 8330283.5 | 4503126.5 | 14219826 | 7554501 | 10082676 |
| Q5U2U7 | Rnmt | mRNA cap guanine-N(7) methyltransferase | 56.08 | 4.12 | 52817 | 1 | 2 | 2 | 2 | 2 | 2 | 2 | 2 | 2 | 2 | 2 | 2 | 245933.64 | 2 | 2 | 2 | 2 | 2 | 2 | 279426.34 | 2 |
| Q8VCT3 | Rnpep | Aminopeptidase B | 453.8 | 17.08 | 72416 | 8 | 3481206.5 | 5994921 | 2645307.25 | 6817512 | 5621535 | 8526276 | 6316345.5 | 4047655 | 9800315 | 7123265 | 6848093 | 5322457 | 7602846.5 | 6192302 | 1953093 | 6119749 | 5689372 | 5672303 | 7207189.5 | 4359274.5 |
| P60603 | Romo1 | Reactive oxygen species modulator 1 | 195.32 | 77.22 | 8183 | 3 | 2 | 2 | 2 | 2 | 728003.69 | 1422486.12 | 2 | 920095.94 | 1575996 | 2 | 3609320 | 2835384 | 571272.06 | 4708662.5 | 1169851.62 | 2 | 1928126.88 | 1824637.25 | 468633.03 | 1364080.25 |
| Q8VEE4 | Rpa1 | Replication protein A 70 kDa DNA-binding subuni | 45.23 | 2.09 | 69037 | 1 | 2 | 2 | 2 | 2 | 2 | 2 | 2 | 2 | 2 | 2 | 2 | 2 | 2 | 2 | 2 | 2 | 2 | 2 | 2 | 2 |
| P47968 | Rpia | Ribose-5-phosphate isomerase | 179.39 | 15.51 | 32451 | 4 | 4443421 | 6900850 | 1460664.38 | 273751.97 | 2559713 | 2 | 1165409 | 568498.81 | 616514.94 | 1722682.12 | 4655229 | 3565758.5 | 3465761.25 | 4294734 | 675120.81 | 3735296 | 2060600.25 | 2105359 | 1841363 | 1637203 |
| Q6PDV7 | Rpl10 | Large ribosomal subunit protein uL16 | 406.81 | 36.45 | 24604 | 9 | 5287534 | 5781034 | 2638705.5 | 7312081 | 6381104 | 5167812 | 2752492.75 | 5373322 | 11382137 | 5801766.5 | 10032125 | 12601665 | 10344432 | 10621064 | 4174086 | 9519330 | 4212892.5 | 5167509.5 | 5888209 | 3447050.75 |
| P53026 | Rpl10a | Large ribosomal subunit protein uL1 | 168.07 | 9.68 | 24916 | 3 | 2042792.25 | 10195443 | 2701091.75 | 12223213 | 49157990 | 18620134 | 21449658 | 47643570 | 32530308 | 27597346 | 14120695 | 17610034 | 13494737 | 13341631 | 5281745.5 | 14822110 | 10035318 | 7416116 | 10583862 | 9769310 |
| P62914 | Rpl11 | Large ribosomal subunit protein uL5 | 370.02 | 30.9 | 20252 | 7 | 564504.25 | 1457671.25 | 1882088.75 | 10265708 | 23317980 | 7953423.5 | 5468108 | 7542899.5 | 21251954 | 14089294 | 31928682 | 31055308 | 21416070 | 23788338 | 9407910 | 29560686 | 12614142 | 8792854 | 23076994 | 7348003 |
| P35979 | Rpl12 | Large ribosomal subunit protein uL11 | 413.65 | 52.73 | 17805 | 6 | 6611708 | 1426312.62 | 3310987.25 | 1869362.62 | 6159109.5 | 1068614.88 | 3588390.25 | 1533132.38 | 4518973 | 11869186 | 15705313 | 1760379.5 | 3685012.25 | 8986103 | 2462201.25 | 1816798.25 | 1765859.75 | 1332677.5 | 7329650.5 | 2248842.5 |
| P47963 | Rpl13 | Large ribosomal subunit protein eL13 | 447.66 | 33.65 | 24305 | 8 | 491075.75 | 2 | 2 | 2 | 1517240.88 | 2 | 2 | 193586.69 | 3112993 | 2 | 9271817 | 7147376.5 | 2198282.75 | 3235460.5 | 1332002.75 | 6665209 | 2071671.12 | 2 | 1519857.5 | 2 |
| P19253 | Rpl13a | Large ribosomal subunit protein uL13 | 490.61 | 39.9 | 23464 | 10 | 809942.06 | 2065789.75 | 622730.88 | 1155342.62 | 4644144.5 | 16123538 | 12403383 | 21329116 | 19748922 | 2927649.25 | 15885731 | 12933192 | 14460092 | 19538442 | 1195313.25 | 2755117.75 | 3852312.25 | 2144859 | 5852982.5 | 2797487 |
| Q9CR57 | Rpl14 | Large ribosomal subunit protein eL14 | 355.34 | 25.81 | 23564 | 6 | 5217499.5 | 8229449.5 | 5476875 | 10405872 | 15755802 | 6141344.5 | 5037366.5 | 9524234 | 11538332 | 11368322 | 20265922 | 17366108 | 18473098 | 20530322 | 14041842 | 12062041 | 12331687 | 12807103 | 17410970 | 12026771 |
| P61314 | Rpl15 | Large ribosomal subunit protein eL15 | 384.7 | 37.25 | 24146 | 8 | 2062830 | 2737747.75 | 2928194 | 1996129.75 | 5394601.5 | 3095556.25 | 2143672.5 | 6030580.5 | 18013806 | 2837059.75 | 1486693.12 | 1860928.25 | 1739066.88 | 62862.15 | 1124286.12 | 61523.97 | 2 | 2 | 2 | 1701757 |
| P24049 | Rpl17 | Large ribosomal subunit protein uL22 | 218.27 | 24.46 | 21397 | 5 | 4209144.5 | 4409993 | 2911246 | 5371977.5 | 4534420 | 4361944 | 3610781 | 7782272 | 8319668 | 425102.09 | 9014267 | 8900701 | 4902082 | 6392799 | 4662976 | 9602756 | 5680413.5 | 8723828 | 8492576 | 5753117.5 |
| P35980 | Rpl18 | Large ribosomal subunit protein eL18 | 576.32 | 38.83 | 21645 | 9 | 7082105 | 13032912 | 13312382 | 36426102 | 14256378 | 35881826 | 35232498 | 29816846 | 26514570 | 15315222 | 22895662 | 22352914 | 24074952 | 24061846 | 18280846 | 47270978 | 24857418 | 22558386 | 46932114 | 12791116 |
| P62717 | Rpl18a | Large ribosomal subunit protein eL20 | 359.3 | 37.5 | 20732 | 8 | 338509.66 | 963708.38 | 1556835.75 | 2156116.25 | 6282601.5 | 3812477 | 3942445 | 8310582 | 9461891 | 3602769.25 | 5793719 | 11741657 | 5959921 | 8308355 | 2671437.5 | 7141998.5 | 1066030.38 | 410343.47 | 1734294.75 | 2 |
| P84099 | Rpl19 | Large ribosomal subunit protein eL19 | 228.01 | 21.43 | 23466 | 4 | 504970.94 | 2 | 2 | 814919.25 | 4606776.5 | 650645.12 | 2631454.25 | 4386565 | 9034926 | 645770.12 | 5497149.5 | 5014601 | 776755.19 | 1288044.5 | 993107.56 | 3465690.5 | 797889.25 | 2 | 779576.56 | 2 |
| O09167 | Rpl21 | Large ribosomal subunit protein eL21 | 133.56 | 18.12 | 18579 | 2 | 2 | 1269832.5 | 2 | 708284.88 | 4561473.5 | 1533945.12 | 1064753.88 | 2132445.25 | 4078605.5 | 1293983.38 | 2098625.5 | 3585008 | 549013.12 | 786962.75 | 2 | 2 | 2 | 2 | 2 | 2 |
| P67984 | Rpl22 | Large ribosomal subunit protein eL22 | 298.38 | 30.47 | 14759 | 5 | 6735340 | 13132506 | 3531725.75 | 19220804 | 22563198 | 15864269 | 20604938 | 34767366 | 20810448 | 7652512 | 9459652 | 10893612 | 14868383 | 15115904 | 4577790.5 | 24519912 | 14907255 | 6990125.5 | 27123884 | 7449776 |
| Q9D7S7 | Rpl22l1 | Ribosomal protein eL22-like 1 | 127.26 | 19.67 | 14467 | 2 | 1388236.12 | 1285413.62 | 2152286.25 | 1800872.88 | 2716735.25 | 806235.56 | 904295.88 | 1725890.62 | 1619530.75 | 963340.56 | 806782 | 830509.69 | 1229249.38 | 867466.19 | 754241.12 | 1673295.38 | 2 | 628517.62 | 1086537.5 | 2 |
| P62830 | Rpl23 | Large ribosomal subunit protein uL14 | 314.41 | 40.71 | 14865 | 6 | 2378503 | 5021202 | 1291690.62 | 3306389.75 | 5616888 | 3103561.25 | 2444283.5 | 1284367.62 | 4875817 | 2503249 | 13527060 | 12585783 | 8247035 | 8426786 | 3307772.75 | 10219581 | 4829017 | 4047960.25 | 7638212 | 4905355.5 |
| P62751 | Rpl23a | Large ribosomal subunit protein uL23 | 171.19 | 23.72 | 17695 | 4 | 1275482.5 | 1148164.25 | 2 | 2050691 | 3723990 | 870317.5 | 267132.84 | 1550340.38 | 1346716.25 | 1265525.38 | 3483703.75 | 3192825.5 | 4092212.5 | 3718261 | 1668327.88 | 2601844 | 3856431.75 | 4402897 | 4987274.5 | 1072168.25 |
| P83732 | Rpl24 | Large ribosomal subunit protein eL24 | 221.37 | 21.66 | 17779 | 4 | 2 | 694715.19 | 2 | 2 | 2165637.5 | 2 | 2 | 2064352.12 | 5253775.5 | 2 | 3666756.25 | 5445843.5 | 2658988.5 | 3583690.25 | 2 | 1667779.38 | 2 | 935716.06 | 539378.62 | 2 |
| P12749 | Rpl26 | Large ribosomal subunit protein uL24 | 289 | 31.03 | 17277 | 7 | 3747866.25 | 1211685.12 | 2728195.25 | 2946122 | 2225558.5 | 1697964.88 | 3296935.25 | 2280845.75 | 2875285 | 1699378.5 | 4548565.5 | 6304317 | 1720357.5 | 1647247.25 | 5126005 | 4067275.25 | 4216392 | 2572401.25 | 1457490.5 | 3111673.5 |
| P61354 | Rpl27 | Large ribosomal subunit protein eL27 | 123.94 | 20.59 | 15798 | 3 | 731335.44 | 2 | 786127.62 | 1575051.12 | 3195804.5 | 2 | 2 | 2 | 2295598.75 | 2 | 2875305.25 | 2331041.75 | 380921.66 | 2673633.25 | 2 | 570910.69 | 2 | 1161328.62 | 1977222 | 1274524.88 |
| P14115 | Rpl27a | Large ribosomal subunit protein uL15 | 255.94 | 23.65 | 16605 | 4 | 4409057.5 | 10933633 | 3123393.5 | 5540779.5 | 10753978 | 7484739 | 7177433.5 | 10112595 | 7384625 | 4780730.5 | 3837164.5 | 3892746 | 5714408.5 | 6002186.5 | 4232096.5 | 4988958 | 4167458 | 6906814.5 | 6103224.5 | 6746705 |
| P41105 | Rpl28 | Large ribosomal subunit protein eL28 | 325.22 | 28.47 | 15733 | 6 | 1148092.5 | 407991.69 | 881119.62 | 837407.5 | 37646.61 | 669253.56 | 2481803.25 | 196410.45 | 2570499.5 | 1018719.94 | 864297.06 | 2723562 | 2 | 2 | 523827.31 | 2 | 654058.06 | 2 | 2 | 2 |
| P47915 | Rpl29 | Large ribosomal subunit protein eL29 | 104.44 | 11.88 | 17587 | 2 | 2 | 8925586 | 1647330.25 | 12789740 | 2445790.5 | 2875423.5 | 139382.03 | 6257633.5 | 36343150 | 977492.12 | 1858401.62 | 10786735 | 1480253.12 | 13068197 | 238300.05 | 4933737.5 | 2 | 2 | 242372.41 | 1943031.25 |
| P27659 | Rpl3 | Large ribosomal subunit protein uL3 | 496.22 | 23.82 | 46110 | 9 | 1347047 | 1125099.38 | 2912832.25 | 2416240.75 | 4255148 | 2539171 | 1186450.62 | 2001030.5 | 2195698.5 | 2982060 | 6576890 | 6229980.5 | 6310935 | 4664042 | 4682704 | 7646918.5 | 591016.06 | 2705220.25 | 3065118.5 | 3463219.75 |
| P62889 | Rpl30 | Large ribosomal subunit protein eL30 | 336.55 | 42.61 | 12784 | 5 | 4925126 | 3972616.5 | 6226244 | 2632135 | 17975806 | 4366384 | 5006526.5 | 10332006 | 10205503 | 10623449 | 6074456 | 7065765 | 12516958 | 7686627 | 6172660 | 4057268.5 | 5479573 | 3660543 | 5131628.5 | 3523517.5 |
| P62900 | Rpl31 | Large ribosomal subunit protein eL31 | 205.72 | 35.2 | 14463 | 4 | 84752.1 | 74585.77 | 193882.86 | 2 | 3536983 | 2 | 85071.28 | 2392191.25 | 2770180 | 1754531.62 | 4112339.75 | 3370769.75 | 2951323.25 | 1694944.62 | 2548365 | 3186801.5 | 2094227.25 | 2162402.5 | 2919873.25 | 2 |
| P62911 | Rpl32 | Large ribosomal subunit protein eL32 | 179 | 24.44 | 15860 | 4 | 2 | 2 | 2 | 2 | 776113.94 | 706299 | 521685.31 | 880971.62 | 1692547 | 619148.88 | 1266692.12 | 1147056.12 | 1857498.12 | 1476267.25 | 2 | 2 | 2 | 703799.81 | 2 | 2 |
| Q9D1R9 | Rpl34 | Large ribosomal subunit protein eL34 | 177.26 | 25.64 | 13293 | 4 | 482168.97 | 2 | 710385.94 | 608768.5 | 2 | 2 | 2 | 2 | 2 | 400101.88 | 4640684.5 | 2818134.5 | 2332663.75 | 2367315.5 | 2 | 2840980.5 | 549381.88 | 382513.78 | 2 | 460200.88 |
| P17078 | Rpl35 | Large ribosomal subunit protein uL29 | 119.27 | 21.95 | 14553 | 2 | 2 | 2 | 2 | 2 | 2 | 2 | 2 | 2 | 351020.62 | 2 | 2 | 2 | 2 | 2 | 2 | 2 | 2 | 2 | 2 | 2 |
| O55142 | Rpl35a | Large ribosomal subunit protein eL33 | 120.26 | 16.36 | 12554 | 3 | 2 | 723244 | 2 | 630294.75 | 2205865.5 | 1991646.25 | 1484179.12 | 2620347.75 | 1361353.75 | 1747293.5 | 631401.5 | 2 | 923271.5 | 876705.19 | 2 | 2 | 2 | 2 | 2 | 2 |
| P39032 | Rpl36 | Large ribosomal subunit protein eL36 | 84.41 | 21.9 | 12268 | 2 | 2 | 2 | 2 | 2 | 745378.94 | 803147.31 | 2 | 1009456.06 | 835489.31 | 923686.31 | 564581.06 | 489028.81 | 2 | 2 | 2 | 2 | 2 | 2 | 2 | 2 |
| P61514 | Rpl37a | Large ribosomal subunit protein eL43 | 95.21 | 28.26 | 10275 | 2 | 2 | 2 | 2 | 2 | 2 | 2 | 2 | 2 | 764725.31 | 2 | 955486.81 | 2 | 2 | 2 | 2 | 807874.69 | 2 | 2 | 955015.12 | 2 |
| P63174 | Rpl38 | Large ribosomal subunit protein eL38 | 101.67 | 34.29 | 8218 | 2 | 2 | 2 | 2 | 2 | 2 | 2 | 2 | 2 | 2 | 2 | 2 | 2 | 2 | 2 | 2 | 2 | 2 | 2 | 2 | 2 |
| Q9D8E6 | Rpl4 | Large ribosomal subunit protein uL4 | 970.88 | 42.96 | 47154 | 19 | 10151876 | 5969835.5 | 4105612 | 7493339 | 36825042 | 7480062 | 9278391 | 11588010 | 28402476 | 16177793 | 24398930 | 20496580 | 21510620 | 22299876 | 9217665 | 13482940 | 10646552 | 4427411.5 | 6408836.5 | 3329226.5 |
| P09895 | Rpl5 | Large ribosomal subunit protein uL18 | 527.93 | 34.68 | 34459 | 9 | 11769742 | 15005483 | 13847691 | 18785062 | 18236976 | 10841014 | 9251037 | 10807467 | 12444052 | 12230678 | 20817738 | 23253880 | 25268382 | 25805484 | 14022350 | 19914126 | 21938106 | 23360344 | 25767374 | 21353190 |
| P47911 | Rpl6 | Large ribosomal subunit protein eL6 | 545.4 | 29.39 | 33510 | 10 | 1690973.5 | 4263378.5 | 2959971.75 | 5370237.5 | 8640134 | 7396459 | 3762642 | 6789501.5 | 8096470.5 | 4802272.5 | 9030209 | 11028524 | 9189400 | 10575780 | 9013988 | 16998280 | 9950052 | 7349315.5 | 4623155.5 | 9957904 |
| P14148 | Rpl7 | Large ribosomal subunit protein uL30 | 743.45 | 54.44 | 31420 | 16 | 19687090 | 35486650 | 11226459 | 49722802 | 59701798 | 57451126 | 56878398 | 70012250 | 85687890 | 47438034 | 34026062 | 37756454 | 42031298 | 40733586 | 30961386 | 40235122 | 38937802 | 41340338 | 50368394 | 37334990 |
| P12970 | Rpl7a | Large ribosomal subunit protein eL8 | 522.58 | 37.59 | 29977 | 10 | 4794227.5 | 839961.19 | 2765724.75 | 4155681.25 | 8965753 | 1321921.5 | 2354350.5 | 605859 | 5350919 | 4632808.5 | 16169333 | 11910126 | 14721663 | 8898676 | 8638001 | 8710096 | 7644223 | 3051462.5 | 8039036 | 7078602.5 |
| P62918 | Rpl8 | Large ribosomal subunit protein uL2 | 338.46 | 23.74 | 28025 | 7 | 1078003.12 | 3381694.75 | 1118802.25 | 5226617.5 | 12762824 | 3917306.75 | 3557163 | 10628112 | 22101994 | 4575307 | 16562600 | 15995718 | 7047508.5 | 10067494 | 1220076.12 | 15348954 | 7480764.5 | 1385448.12 | 6289394 | 2276846 |
| P51410 | Rpl9 | Large ribosomal subunit protein uL6 | 698.08 | 67.19 | 21881 | 12 | 1874341.5 | 11892226 | 7061566 | 12254184 | 11707654 | 16075208 | 17614160 | 22591100 | 20844160 | 8027666 | 15708552 | 14681894 | 21409452 | 22504406 | 13824138 | 20707986 | 21051774 | 13776298 | 24579566 | 12924292 |
| P14869 | Rplp0 | Large ribosomal subunit protein uL10 | 990.4 | 51.1 | 34216 | 16 | 48307458 | 12308506 | 25287896 | 21277136 | 16746412 | 3802089.25 | 12596474 | 14092201 | 19496268 | 23178490 | 7217573.5 | 9668595 | 15500166 | 20180670 | 36548534 | 1567290.38 | 17168066 | 17842914 | 13768442 | 15183559 |
| P47955 | Rplp1 | Large ribosomal subunit protein P1 | 200.28 | 52.63 | 11475 | 3 | 1925067.12 | 3357469.75 | 4261918 | 8038537.5 | 3168623 | 3110960.5 | 4561911.5 | 1212974.38 | 4451486 | 4739863 | 1956214.38 | 3446322 | 4474671 | 5763386 | 8353837 | 207596.77 | 2496731.5 | 3723773 | 866359.94 | 3664854.5 |
| P99027 | Rplp2 | Large ribosomal subunit protein P2 | 139.34 | 47.83 | 11651 | 3 | 2 | 2 | 2 | 1103889.12 | 1828933.5 | 2739620 | 2 | 1899626.5 | 1096974.38 | 1620542 | 1168143.5 | 140652.11 | 2 | 1298214.12 | 1277829.75 | 2 | 204639.77 | 162649.36 | 2 | 2 |
| Q91YQ5 | Rpn1 | Dolichyl-diphosphooligosaccharide--protein glyco | 1066.64 | 41.28 | 68528 | 20 | 1348725.5 | 1302160.75 | 2578292.25 | 1355679 | 3963686.5 | 739201.5 | 488502.19 | 442613.44 | 2739524.5 | 4920995.5 | 1709030.75 | 1946444.5 | 2681882 | 2805494 | 791136.88 | 3381473.5 | 1933320 | 2038284.12 | 2229125.5 | 3065166.5 |
| Q9DBG6 | Rpn2 | Dolichyl-diphosphooligosaccharide--protein glyco | 924.51 | 25.2 | 69063 | 13 | 6651205.5 | 4438942.5 | 8067472.5 | 8965252 | 3749407 | 8395456 | 6181552 | 9059226 | 3418952 | 2746964.5 | 8209514 | 6210630.5 | 5265312.5 | 4143014.75 | 13753202 | 20098340 | 15274090 | 22187850 | 13830878 | 19179244 |
| O88796 | Rpp30 | Ribonuclease P protein subunit p30 | 72.83 | 4.85 | 29473 | 1 | 2 | 2 | 53594.14 | 2 | 94948.88 | 177717.48 | 2 | 148017.2 | 393510.56 | 81918.53 | 204812.55 | 139676.14 | 166427.78 | 214165.19 | 237648.52 | 2 | 157941.02 | 89414.22 | 355417.72 | 2 |
| P63325 | Rps10 | Small ribosomal subunit protein eS10 | 246.7 | 18.79 | 18916 | 4 | 658517.25 | 2 | 43165.51 | 1258436.88 | 531023.38 | 2 | 2 | 819009.06 | 986111.44 | 2 | 1943700.5 | 327346.91 | 535790.38 | 1427313.5 | 531486.5 | 4304561 | 311223.75 | 1444335.5 | 1801786.88 | 268495.53 |
| P62281 | Rps11 | Small ribosomal subunit protein uS17 | 315.61 | 48.1 | 18431 | 8 | 5109760.5 | 5525068.5 | 3242873.75 | 29637210 | 36105042 | 7373975 | 26378786 | 39904986 | 41044122 | 12394194 | 26676028 | 25593122 | 14304344 | 22519796 | 10045588 | 10764796 | 18046070 | 10795051 | 19969690 | 9704670 |
| P63323 | Rps12 | Small ribosomal subunit protein eS12 | 273.85 | 40.91 | 14515 | 5 | 2718796 | 6007141 | 5163005.5 | 3469842.25 | 6269229.5 | 948849.5 | 1648908.88 | 6252798.5 | 2190993 | 8844248 | 4673705 | 4122271.75 | 3682219.25 | 2800110 | 7954752.5 | 1808468.88 | 2862634.25 | 3270763.25 | 3295513 | 8116051.5 |
| P62278 | Rps13 | Small ribosomal subunit protein uS15 | 213.99 | 19.21 | 17222 | 4 | 6124273 | 7702549.5 | 5221990.5 | 6810850 | 8939779 | 6457356 | 8361031.5 | 16131417 | 8174505.5 | 6235987.5 | 5126831 | 6324910 | 6138979.5 | 5523796 | 6719917 | 7015375 | 5228108 | 6079816.5 | 4166692.75 | 3473495.75 |
| P13471 | Rps14 | Small ribosomal subunit protein uS11 | 455.08 | 33.77 | 16259 | 7 | 6859283 | 5300961.5 | 6420002.5 | 2758912.25 | 3747923.75 | 2482275.75 | 3234807.5 | 4736508 | 4669675.5 | 2667940.5 | 7069506 | 6632261 | 4713949 | 5303863.5 | 5577319.5 | 5123099 | 4334541.5 | 6313370 | 4662193 | 4552236 |
| P62245 | Rps15a | Small ribosomal subunit protein uS8 | 427.9 | 52.31 | 14840 | 9 | 25213590 | 42454954 | 27040442 | 37674474 | 67448370 | 51802082 | 51437754 | 65442934 | 54051074 | 44037442 | 32580194 | 38879254 | 41639158 | 41131722 | 20833476 | 24187974 | 17889716 | 15635472 | 20774716 | 10503055 |
| P14131 | Rps16 | Small ribosomal subunit protein uS9 | 624.1 | 54.79 | 16445 | 10 | 9203700 | 14366713 | 14692505 | 24903804 | 11723616 | 14651200 | 12993121 | 28601722 | 23464372 | 20665994 | 25863874 | 30504374 | 31843082 | 37536246 | 16781038 | 29447054 | 12002095 | 9639003 | 20517298 | 3999179.5 |
| P63276 | Rps17 | Small ribosomal subunit protein eS17 | 243.52 | 39.26 | 15524 | 4 | 2 | 1155955.25 | 2 | 677116.75 | 3516148.75 | 1418239 | 1057548.88 | 2702723 | 3222995 | 2602968.5 | 1410072.25 | 4614113 | 3032207 | 2434076.25 | 3500611.5 | 2755817.75 | 2489012 | 1557589.88 | 1825750.38 | 1931134.88 |
| P62270 | Rps18 | Small ribosomal subunit protein uS13 | 737 | 67.76 | 17719 | 15 | 27574674 | 31603170 | 26621430 | 22139900 | 52244218 | 33216412 | 27635340 | 41537654 | 42880818 | 37218990 | 35087602 | 36714902 | 27844996 | 28038896 | 24599674 | 33114934 | 20410124 | 22594266 | 16471770 | 33959082 |
| P17074 | Rps19 | Small ribosomal subunit protein eS19 | 313.87 | 47.59 | 16085 | 8 | 12565581 | 7701839.5 | 17179902 | 8893324 | 11042064 | 6292594 | 9274432 | 15734296 | 6145635.5 | 10832859 | 6813740.5 | 6734017.5 | 8447786 | 7210108.5 | 14380570 | 3458324 | 5733820.5 | 11349524 | 5463960 | 5666901.5 |
| P25444 | Rps2 | Small ribosomal subunit protein uS5 | 635.34 | 37.2 | 31231 | 13 | 12656460 | 15625446 | 15041970 | 47641946 | 35985338 | 44930970 | 34960430 | 40650530 | 40238774 | 37841610 | 37276634 | 38662254 | 33291866 | 42104022 | 36799566 | 34934310 | 32811232 | 43757498 | 34313386 | 42618970 |
| P60867 | Rps20 | Small ribosomal subunit protein uS10 | 192.59 | 31.09 | 13373 | 5 | 2 | 313986.22 | 584068.81 | 2386269.5 | 2801132.25 | 558513.88 | 500031.59 | 2 | 2032589.12 | 1548877 | 6957512.5 | 8117236 | 3300256 | 2 | 791681.38 | 4651480 | 2576054.25 | 513875.72 | 1718937.62 | 922263.94 |
| P05765 | Rps21 | Small ribosomal subunit protein eS21 | 315.26 | 59.04 | 9127 | 5 | 374543.31 | 1071584.62 | 1777041.5 | 3805792 | 9552724 | 8348942.5 | 6061709.5 | 12740052 | 8698547 | 5700216 | 23094284 | 19644056 | 12806833 | 13403214 | 6946013.5 | 12303262 | 11472316 | 7197595.5 | 15709172 | 6024511 |
| P62267 | Rps23 | Small ribosomal subunit protein uS12 | 147.72 | 23.78 | 15808 | 3 | 2 | 1582395.75 | 2 | 2 | 2 | 1295902.5 | 2 | 2 | 2371359.75 | 880588.38 | 849474.31 | 2242358 | 2 | 2 | 2 | 2565336 | 1372870.75 | 2 | 96570.38 | 2 |
| P62849 | Rps24 | Small ribosomal subunit protein eS24 | 185.11 | 28.57 | 15423 | 4 | 1350819.5 | 1239705.38 | 3000845.75 | 239768.55 | 3265210.5 | 354555.06 | 1327746.38 | 2739097.75 | 2927484 | 4508412 | 7592040 | 6956370 | 9127881 | 9266432 | 6794904 | 4689962 | 5003893.5 | 2685076.75 | 383061.19 | 4067333.25 |
| P62852 | Rps25 | Small ribosomal subunit protein eS25 | 315.4 | 47.2 | 13742 | 7 | 13625144 | 28003198 | 16428976 | 5271170 | 31276958 | 17008478 | 20455518 | 19182512 | 24254298 | 18748666 | 14788684 | 20041110 | 19947910 | 16366809 | 9474466 | 12404740 | 7372780 | 5552487 | 9812838 | 3697821.5 |

| | | | | | | | | | | | | | | | | | | | | | | | | | | |
|---|---|---|---|---|---|---|---|---|---|---|---|---|---|---|---|---|---|---|---|---|---|---|---|---|---|---|
| P62855 | Rps26 | Small ribosomal subunit protein eS26 | 156.29 | 24.35 | 13015 | 3 | 9904573 | 9901352 | 10738054 | 23091446 | 35239962 | 16085287 | 18098556 | 36466286 | 18906138 | 23209786 | 12614103 | 9825643 | 9310006 | 8433979 | 10659129 | 12099105 | 7051116.5 | 8140318.5 | 10385642 | 4908750 |
| P24051 | Rps27l | Small ribosomal subunit protein eS27-like | 120.85 | 25 | 9477 | 2 | 2 | 2 | 2 | 2 | 2 | 1517504.62 | 1607556.12 | 2860443 | 3602225.25 | 2 | 3459366 | 3101743.5 | 1151555.88 | 1696551.62 | 752189.31 | 2 | 2 | 2 | 2 | 2 |
| P62858 | Rps28 | Small ribosomal subunit protein eS28 | 200 | 40.58 | 7841 | 4 | 1741975.38 | 674755.38 | 1317763.38 | 1416598.25 | 3088444 | 1596316.38 | 851072.94 | 147396.55 | 1427506.5 | 2349439 | 10933044 | 9701996 | 8382043 | 5661948.5 | 9986816 | 6828980 | 8204428.5 | 4876772 | 6128217 | 5608006 |
| P62908 | Rps3 | Small ribosomal subunit protein uS3 | 1232.52 | 82.72 | 26674 | 25 | 68962794 | 154523954 | 81353666 | 159125218 | 106005946 | 120093794 | 135252498 | 127594034 | 129426938 | 106618914 | 122771114 | 130850690 | 146253010 | 133049218 | 42212762 | 65604178 | 45851170 | 37471222 | 49419570 | 31856196 |
| P97351 | Rps3a | Small ribosomal subunit protein eS1 | 839.56 | 37.5 | 29885 | 13 | 31643672 | 42477626 | 21390906 | 31620142 | 64734458 | 44745062 | 44785522 | 42286966 | 51444602 | 46374958 | 32761886 | 41032074 | 37924786 | 36369030 | 28732190 | 33676078 | 27498830 | 25007598 | 13447914 | 28636924 |
| P62702 | Rps4x | Small ribosomal subunit protein eS4 | 887.81 | 63.12 | 29598 | 18 | 8850566 | 15992792 | 18780736 | 21787708 | 59851322 | 26947562 | 22876694 | 36154642 | 44602270 | 44060442 | 83222882 | 70571058 | 62999178 | 77136322 | 28657908 | 56885578 | 29326664 | 19345578 | 47767922 | 20613154 |
| P97461 | Rps5 | Small ribosomal subunit protein uS7 | 527.97 | 29.41 | 22876 | 10 | 10262940 | 30387108 | 9392317 | 34141302 | 22647612 | 20601712 | 25160816 | 22309112 | 25138274 | 17613274 | 20462496 | 25571666 | 21012010 | 22538380 | 20354546 | 18069536 | 15308251 | 14765284 | 13085430 | 14345330 |
| P62754 | Rps6 | Small ribosomal subunit protein eS6 | 531.84 | 41.37 | 28681 | 9 | 5425967 | 11169348 | 8941733 | 2227325.25 | 15407159 | 6153274 | 4581802 | 8896789 | 5169472 | 6386200 | 8045388.5 | 7407103.5 | 8696035 | 8042036 | 6825130 | 5391125 | 4886636 | 5203783 | 3180580.75 | 6174853.5 |
| P62082 | Rps7 | Small ribosomal subunit protein eS7 | 219.86 | 22.68 | 22127 | 4 | 3196841.75 | 1258828.38 | 2648984 | 2573534.25 | 4611335.5 | 4220483 | 5077581.5 | 2166527 | 3577204.25 | 7133031 | 1083094.62 | 2126886.5 | 3055168.75 | 6198996 | 10960376 | 278968.94 | 6919696 | 3928491.75 | 2450118 | 4658196 |
| P62242 | Rps8 | Small ribosomal subunit protein eS8 | 751.95 | 53.37 | 24205 | 12 | 25889272 | 35165038 | 22229818 | 18599664 | 54640966 | 29109064 | 31003858 | 44216046 | 33712954 | 29832050 | 25282626 | 20811492 | 47805962 | 34012158 | 33480358 | 39201858 | 28942726 | 26047390 | 35364930 | 21775454 |
| P29314 | Rps9 | Small ribosomal subunit protein uS4 | 353.21 | 29.9 | 22591 | 7 | 7251722 | 12434647 | 8787807 | 7029358 | 13269626 | 13748017 | 11003614 | 15094065 | 15001214 | 10879871 | 15426055 | 14618049 | 16658758 | 16162125 | 4162746.5 | 12780933 | 9383742 | 5331028 | 10454571 | 6988011.5 |
| P14206 | Rpsa | Small ribosomal subunit protein uS2 | 741.75 | 50.85 | 32838 | 12 | 16074994 | 13656222 | 18240308 | 18927790 | 34380682 | 14847563 | 14155545 | 26433574 | 28345000 | 19097098 | 72366498 | 69483690 | 38055890 | 52467102 | 18301042 | 65226098 | 24961334 | 15190722 | 36100678 | 15601563 |
| Q63486 | Rraga | Ras-related GTP-binding protein A | 113.3 | 6.71 | 36566 | 2 | 7256780 | 6091934 | 6348833 | 5655308 | 6408092 | 5148404 | 5582542.5 | 9704875 | 5632498 | 4930169.5 | 3929210.25 | 4747738 | 4477466.5 | 2541026.25 | 2180480.75 | 2 | 2523466.25 | 2560047 | 3546446.25 | 2101292.75 |
| P10833 | Rras | Ras-related protein R-Ras | 96.64 | 13.3 | 23764 | 2 | 581750.25 | 178356.62 | 511308.03 | 223020.72 | 290904.03 | 203569.73 | 633132.56 | 295690 | 611833.19 | 369354.09 | 89391.68 | 131818.06 | 555065.5 | 260514.47 | 784925.75 | 129395.15 | 380165.69 | 178720.17 | 119408.23 | 2 |
| Q99PL5 | Rrbp1 | Ribosome-binding protein 1 | 1111 | 14.08 | 172879 | 22 | 3979221 | 6918687.5 | 4547440.5 | 8420620 | 15452745 | 5847490 | 7034758.5 | 14286051 | 17973906 | 13828141 | 17872964 | 18564328 | 13556286 | 16552179 | 8171393.5 | 19266824 | 10595890 | 9183555 | 11396135 | 6723871.5 |
| P07742 | Rrm1 | Ribonucleoside-diphosphate reductase large sub | 91.15 | 2.9 | 90210 | 2 | 1021867.94 | 917258.75 | 321819.34 | 1130933.12 | 125659.25 | 1121698.12 | 493526.09 | 98487.64 | 605495 | 461545.38 | 2 | 2 | 655641.06 | 589820 | 672344.5 | 2 | 2 | 2 | 2 | 2 |
| P11157 | Rrm2 | Ribonucleoside-diphosphate reductase subunit M | 126.44 | 6.41 | 45096 | 2 | 1332972.5 | 1604896.5 | 1999378.88 | 1425039.75 | 1294339.12 | 2 | 2 | 1568116.88 | 2 | 870748.44 | 3383921.5 | 1591182.75 | 1261983.75 | 2103717.25 | 2 | 1577122 | 1044540.38 | 1006350.75 | 2060814.88 | 2 |
| P56183 | Rrp1 | Ribosomal RNA processing protein 1 homolog A | 63.71 | 1.82 | 54777 | 1 | 2 | 2 | 2 | 2 | 2 | 2 | 2 | 2 | 611820.06 | 2 | 2 | 2 | 51323.12 | 2 | 2 | 2 | 2 | 2 | 2 | 2 |
| Q91YK2 | Rrp1b | Ribosomal RNA processing protein 1 homolog B | 43.69 | 2.07 | 80582 | 1 | 2 | 582354.56 | 2 | 2 | 603557 | 2 | 2 | 2 | 714005.44 | 536386.69 | 2 | 2 | 2 | 2 | 2 | 2 | 2 | 2 | 2 | 2 |
| Q91WM3 | Rrp9 | U3 small nucleolar RNA-interacting protein 2 | 52.94 | 3.16 | 52107 | 1 | 2 | 2 | 2 | 2 | 67577.67 | 2 | 2 | 2 | 2 | 2 | 2 | 2 | 2 | 2 | 2 | 607619.5 | 2 | 2 | 688322.69 | 2 |
| Q9CYH6 | Rrs1 | Ribosome biogenesis regulatory protein homolog | 165.89 | 8.22 | 41552 | 3 | 329215.31 | 387061.75 | 2 | 2 | 15720.29 | 633948.62 | 298616.19 | 2 | 1112548.75 | 2 | 478567.69 | 2 | 2 | 544472.44 | 386475.84 | 426350.12 | 931481 | 1905437 | 758130.81 | 2 |
| Q8BVY0 | Rsl1d1 | Ribosomal L1 domain-containing protein 1 | 715.47 | 21.9 | 50421 | 12 | 1189669.75 | 2693434 | 520062.41 | 1584554.38 | 4599329 | 4596735.5 | 2685922 | 2583015.5 | 4582791 | 1810958.5 | 4534468.5 | 4661116 | 5981649 | 5902865 | 402507.06 | 3916765.75 | 3754095.5 | 4469837.5 | 82716866 | 98478362 |
| Q9D7H3 | RtcA | RNA 3'-terminal phosphate cyclase | 74.68 | 4.92 | 39254 | 2 | 610733.25 | 746454.5 | 255171.48 | 1400058.5 | 809137.56 | 1211951 | 1088710.88 | 152587.12 | 3215506.5 | 2852375.5 | 3159487.75 | 1375361 | 2265405.75 | 1800646.25 | 363878.56 | 2310360.25 | 1873218.75 | 2410383.5 | 2589707.25 | 1658012.5 |
| Q6AYT3 | Rtcb | RNA-splicing ligase RtcB homolog | 234.28 | 15.25 | 55249 | 5 | 3002945 | 4354867 | 2657131.25 | 4815951.5 | 7721308.5 | 4780546.5 | 5072887.5 | 5497237 | 5624365 | 4165929 | 4682753 | 5520467 | 3078869 | 3785244.5 | 2463961 | 4025675 | 1972712.88 | 2304092.25 | 2658077.75 | 2220211.75 |
| Q99P72 | Rtn4 | Reticulon-4 | 417.91 | 11.62 | 126613 | 7 | 3767054 | 7311594.5 | 2678970.5 | 10315450 | 17416918 | 14256043 | 11528687 | 21806970 | 19712886 | 11568924 | 21624424 | 15859654 | 23092026 | 26072818 | 13472121 | 22373326 | 25098900 | 23650134 | 35763274 | 21176258 |
| Q8R4Y8 | Rttn | Rotatin | 58.35 | 0.63 | 248489 | 2 | 2 | 1467835.5 | 2 | 1534728.88 | 4971487.5 | 4011597.5 | 4279878 | 8327476.5 | 7689203 | 3066977 | 1524556.75 | 2140764.75 | 1063920.25 | 1096246.25 | 2 | 2 | 2 | 2 | 2 | 2 |
| P60122 | Ruvbl1 | RuvB-like 1 | 546.91 | 29.39 | 50214 | 11 | 7201930 | 7734591 | 8404010 | 6199657.5 | 15437682 | 5247495 | 6964462.5 | 6630665 | 10715962 | 8675510 | 5501372 | 3646909.5 | 6413640 | 5964866 | 5495590 | 6543438 | 7358740 | 8459851 | 7761210.5 | 6903392 |
| Q9WTM5 | Ruvbl2 | RuvB-like 2 | 987.87 | 44.49 | 51113 | 17 | 7556387 | 4683156 | 8137985 | 9538975 | 17995244 | 159275186 | 8711232 | 123146634 | 10702277 | 12665640 | 16843958 | 151199938 | 12101352 | 71805090 | 8779013 | 14109094 | 55212650 | 7026576.5 | 10886538 | 38922678 |
| P08207 | S100a10 | Protein S100-A10 | 117.74 | 27.84 | 11186 | 2 | 93583.58 | 1689451.25 | 1111943.62 | 2670732 | 2475828.25 | 159492.48 | 2212318.75 | 190588.22 | 1359313.25 | 45148.99 | 43283.77 | 2 | 261936.56 | 1459040.88 | 1806926.25 | 1430898.88 | 2383247.5 | 401827.25 | 592524.62 | 1070585 |
| P50543 | S100a11 | Protein S100-A11 | 246.06 | 39.8 | 11083 | 4 | 15219749 | 17371244 | 5153107.5 | 25062546 | 4688458 | 19048138 | 18094414 | 1020844.06 | 18665508 | 12214254 | 12524276 | 15279072 | 19911266 | 16812930 | 20374420 | 15368758 | 22681674 | 32891918 | 18325370 | 31517654 |
| P07091 | S100a4 | Protein S100-A4 | 284.27 | 51.49 | 11721 | 6 | 11727275 | 12871573 | 32247322 | 25919978 | 3621804.5 | 7954036 | 9642356 | 23705554 | 29206230 | 24651206 | 6266654.5 | 18955276 | 14698336 | 23260300 | 29753746 | 13138616 | 10810436 | 17842584 | 8669528 | 21707678 |
| P14069 | S100a6 | Protein S100-A6 | 331.86 | 53.93 | 10051 | 6 | 164047586 | 84051890 | 130625074 | 93485706 | 82406186 | 102789850 | 81642994 | 14865421 | 76759890 | 93363050 | 76879578 | 66264026 | 77028434 | 58177130 | 93317338 | 84383602 | 116688154 | 151197442 | 87042130 | 129134050 |
| Q9EP69 | Sacm1l | Phosphatidylinositol-3-phosphatase SAC1 | 245.29 | 9.54 | 66944 | 5 | 1548522.88 | 1334235.25 | 1964439.5 | 911555.31 | 897692.06 | 2537751.5 | 2702818 | 227904.67 | 3755421.5 | 3088335 | 1741299.88 | 3663022 | 2825143 | 2002722 | 644080.25 | 1518923.25 | 511431.94 | 1340475.5 | 3019505.25 | 2 |
| Q9R1T2 | Sae1 | SUMO-activating enzyme subunit 1 | 374.18 | 14.29 | 38620 | 7 | 245071.55 | 1057856.5 | 50217.75 | 1258653.88 | 1579477.88 | 526047.94 | 23990.35 | 875560.88 | 1023161.69 | 849572.94 | 3350936.5 | 4146471 | 3359029 | 4374698 | 1584940.88 | 2428026.75 | 2684836.25 | 2340135.5 | 2 | 2253564.75 |
| D3YXK2 | Safb | Scaffold attachment factor B1 | 80.14 | 1.81 | 105104 | 2 | 2 | 469442.88 | 2 | 2 | 297779.84 | 2 | 2 | 2 | 476005.62 | 391045.72 | 565972.88 | 810411.62 | 2 | 509020.62 | 2 | 738975.56 | 2 | 2 | 579286.88 | 2 |
| Q60710 | Samhd1 | Deoxynucleoside triphosphate triphosphohydrol | 188.4 | 6.38 | 75893 | 4 | 1360837.5 | 2 | 4014741.25 | 2 | 789164.81 | 1247917.88 | 2721074.5 | 2598749.75 | 335096.88 | 2439097 | 1671945.5 | 1385785.38 | 2505410 | 2 | 1576997.5 | 2 | 2 | 1835353.88 | 1852753.88 | 2 |
| Q8BGH2 | Samm50 | Sorting and assembly machinery component 50 h | 572.29 | 25.16 | 51864 | 11 | 6890292 | 5856207 | 5157884.5 | 6335069 | 3665821.25 | 5120055.5 | 5004580.5 | 9468936 | 5198136 | 7111819.5 | 1907679.5 | 2297423.25 | 7949634.5 | 4267576 | 2181523.25 | 1055858.5 | 5224338.5 | 4042698.5 | 4057900.5 | 4559874 |
| P36536 | Sar1a | Small COPII coat GTPase SAR1A | 116.02 | 21.21 | 22371 | 3 | 1653043.88 | 2003774.88 | 2779552.5 | 1737587.62 | 2050946.38 | 1860198 | 1709919.5 | 2 | 1683149.12 | 2233457.25 | 1397125.25 | 1976586.25 | 2125363.75 | 1537564 | 1611153 | 893588.25 | 2 | 2 | 1304287.88 | 1245337.88 |
| Q498U4 | Sarnp | SAP domain-containing ribonucleoprotein | 153.92 | 13.81 | 23605 | 3 | 7393550 | 9447349 | 7244052.5 | 3709924 | 2554644.5 | 2559671.25 | 2079082.75 | 2860472 | 1786770.62 | 2064948.38 | 4510213 | 4469940.5 | 2202840.5 | 3116593 | 1348918.75 | 3761498 | 1934064.25 | 1216605.38 | 4858714.5 | 1505475.62 |
| P26638 | Sars1 | Serine--tRNA ligase, cytoplasmic | 893.22 | 34.96 | 58389 | 19 | 18166996 | 13064344 | 20896140 | 19943166 | 25177930 | 21428552 | 20693212 | 22863536 | 30551008 | 31616026 | 51887238 | 50066766 | 51460562 | 46346346 | 58075158 | 47367470 | 60992722 | 62088558 | 55315782 | 49605318 |
| Q9JJL8 | Sars2 | Serine--tRNA ligase, mitochondrial | 39.68 | 2.32 | 58316 | 1 | 609334.44 | 2 | 2 | 552572.31 | 474324.09 | 615668.56 | 1639327.5 | 85243.86 | 406438.94 | 498305.94 | 2 | 2 | 2 | 2 | 1090972.5 | 2 | 2 | 2 | 2 | 2 |
| Q5XIW8 | Sart1 | U4/U6.U5 tri-snRNP-associated protein 1 | 88.09 | 3.6 | 91011 | 2 | 849734.12 | 836952.44 | 2 | 771974.06 | 1186220.5 | 2 | 606813.25 | 637629.19 | 646626.5 | 2 | 2 | 2 | 2 | 351863.19 | 2 | 2 | 2 | 2 | 2 | 2 |
| P70122 | Sbds | Ribosome maturation protein SBDS | 34.33 | 3.2 | 28781 | 1 | 2 | 2 | 2 | 2 | 2 | 2 | 2 | 2 | 2 | 2 | 2 | 2 | 386710.75 | 2 | 2 | 2 | 2 | 2 | 2 | 2 |
| Q6ZPE2 | Sbf1 | Myotubularin-related protein 5 | 85.84 | 1.18 | 208693 | 2 | 2 | 2 | 45938.35 | 2 | 659787.94 | 2 | 582569.31 | 2 | 147110.72 | 163331.73 | 2 | 2 | 2 | 76437.79 | 566720.31 | 2 | 250615.42 | 2 | 2 | 2 |
| Q9ERN0 | Scamp2 | Secretory carrier-associated membrane protein 2 | 172.73 | 9.73 | 36465 | 2 | 2 | 2 | 2 | 2 | 508952.56 | 453632.88 | 2 | 1267678.25 | 717645.75 | 2 | 2 | 2 | 2 | 219791.33 | 2 | 669027 | 2 | 2 | 2 | 2 |
| O35609 | Scamp3 | Secretory carrier-associated membrane protein 3 | 164.37 | 9.17 | 38458 | 2 | 2 | 700288.12 | 536191.88 | 2 | 2135326.5 | 2245838 | 1742991.25 | 3344586 | 2932386.5 | 1106011.62 | 3824814.5 | 2740424.75 | 1387736 | 1261533.62 | 1759677.62 | 4125530.25 | 5068695.5 | 2886236.5 | 4290584 | 2732534 |
| O35114 | Scarb2 | Lysosome membrane protein 2 | 72.38 | 4.18 | 54044 | 2 | 697377.25 | 435945.12 | 1416234.62 | 1143811.12 | 1404471.75 | 2124690 | 2 | 1562534.12 | 1700615.25 | 1205912.12 | 2 | 658694.75 | 2 | 2 | 518552.69 | 2 | 2 | 2 | 2 | 2 |
| Q8R127 | Sccpdh | Saccharopine dehydrogenase-like oxidoreductas | 205.97 | 16.32 | 47129 | 4 | 77433.12 | 691231.62 | 39167.83 | 661971.31 | 45029.38 | 2 | 171809.33 | 2 | 609689.5 | 83609.46 | 631670.38 | 2 | 34169.57 | 669722.06 | 2 | 64467.11 | 2 | 2 | 2 | 2 |
| Q8BRF7 | Scfd1 | Sec1 family domain-containing protein 1 | 255.7 | 16.59 | 72323 | 6 | 570315.5 | 3350454.5 | 1336057.12 | 2642461 | 3791459.25 | 1646316 | 2427121.5 | 2508015.75 | 3369145 | 3292509 | 2457812.5 | 2509046 | 2968613 | 2348849 | 2 | 2 | 188950.78 | 2 | 875668.75 | 2 |
| Q9JLI6 | Scly | Selenocysteine lyase | 263.05 | 18.98 | 47174 | 6 | 2814121.5 | 8484342 | 5573123 | 6661797 | 6849440 | 8811660 | 8853939 | 7242145.5 | 7213990 | 9522834 | 8463603 | 9954658 | 8116566.5 | 7422305 | 5324298.5 | 5801622 | 7080511 | 12644021 | 8427587 | 4671267 |
| P32020 | Scp2 | Sterol carrier protein 2 | 178.23 | 7.13 | 59126 | 4 | 2972118.25 | 3379896.75 | 2921950.5 | 2480197.5 | 5965632.5 | 1706475.25 | 1482144.25 | 3086429 | 1013814.56 | 6404263.5 | 4884856 | 2533797.75 | 3681074.5 | 1732969 | 1667182.38 | 1593662.25 | 1058593.5 | 2 | 1564283.75 | 1067380.5 |
| Q920A5 | Scpep1 | Retinoid-inducible serine carboxypeptidase | 402.73 | 13.27 | 50965 | 6 | 8630537 | 6559299 | 9070526 | 3748161.5 | 3482733 | 6481396 | 3837168 | 4338992.5 | 3530293 | 5711058.5 | 3438308.25 | 2942265.5 | 4193593.25 | 3144805.5 | 6704627 | 3989818.5 | 5348419.5 | 7930216 | 5045063 | 6060700 |
| O08992 | Sdcbp | Syntenin-1 | 308.16 | 17.73 | 32379 | 4 | 2 | 69003.94 | 125932.9 | 2 | 927386.31 | 319883.62 | 124073.13 | 171835.7 | 998658.25 | 1032839.81 | 2 | 240171.73 | 221417.84 | 474154.19 | 815642.81 | 2 | 230156.8 | 77708.84 | 2 | 2 |
| Q8K2B3 | Sdha | Succinate dehydrogenase [ubiquinone] flavoprote | 703.67 | 17.92 | 72585 | 12 | 16874260 | 27653206 | 21553480 | 24631876 | 28255850 | 22674414 | 26339578 | 6669679 | 27039456 | 21757476 | 29489054 | 35114142 | 32660820 | 32388732 | 28423118 | 24610188 | 27459382 | 33809154 | 28162348 | 31167238 |
| Q9CQA3 | Sdhb | Succinate dehydrogenase [ubiquinone] iron-sulfu | 238.19 | 19.5 | 31814 | 5 | 2 | 2 | 3136803.5 | 2097491.25 | 3196573.25 | 2846232 | 2660636.5 | 3134404.25 | 2619772.75 | 2119616.5 | 1879319.25 | 1875036.88 | 1658747.88 | 1570552.25 | 2307380 | 2 | 859980.38 | 1872841.12 | 2 | 2381459.25 |
| Q5M8N4 | Sdr39u1 | Epimerase family protein SDR39U1 | 39.67 | 6.48 | 31458 | 1 | 2 | 2 | 2 | 2 | 2 | 137812.36 | 114798.05 | 2 | 124969.27 | 120088.45 | 2 | 2 | 81450.4 | 111479.01 | 546358.56 | 2 | 234561.3 | 2 | 2 | 2 |
| P42667 | Sec11a | Signal peptidase complex catalytic subunit SEC1 | 109.34 | 11.17 | 20599 | 2 | 441731.5 | 2 | 2 | 950700.25 | 666709.31 | 149942.03 | 95143.32 | 89800.55 | 2392819 | 1100538.62 | 164213.33 | 960075.5 | 773363.81 | 298977.22 | 941225 | 116252.47 | 309061.62 | 268831.88 | 220752.66 | 2 |
| Q5XFW8 | Sec13 | Protein SEC13 homolog | 47.11 | 3.42 | 35548 | 1 | 2 | 2 | 2 | 2 | 697939.81 | 2 | 2 | 2 | 2 | 388520.66 | 1745475.25 | 1459724.38 | 1456768 | 811020.25 | 1312608.5 | 1559792.88 | 1486923.62 | 1096775.25 | 2 | 1180668 |
| O08547 | Sec22b | Vesicle-trafficking protein SEC22b | 391.92 | 33.95 | 24741 | 7 | 1192133.62 | 2240481 | 3076808 | 4410188 | 4755768 | 3921144.75 | 4705524.5 | 5715603 | 6249312.5 | 5700672.5 | 4621488 | 1694853.62 | 4417080 | 4124214.5 | 1802293.5 | 3615059.5 | 2292511 | 2831257 | 2319633.75 | 294953.97 |
| Q9D662 | Sec23b | Protein transport protein Sec23B | 245.29 | 8.74 | 86437 | 5 | 703512.06 | 666809.88 | 855576.25 | 1381709.75 | 3932411.75 | 3519068.75 | 2740050.5 | 5404731.5 | 4867561.5 | 3538348 | 3103802 | 3312073.5 | 867992.25 | 564220.62 | 1270187.25 | 1103050.75 | 304287.19 | 107158.91 | 837865.88 | 729459.56 |
| Q3UPL0 | Sec31a | Protein transport protein Sec31A | 276.12 | 6.59 | 133569 | 6 | 3947556 | 5961529.5 | 4772276.5 | 4218307.5 | 5212131 | 4685077 | 3066338 | 8674258 | 4185006.75 | 4072165.25 | 5391578.5 | 3839286.5 | 3864296.5 | 4153865.5 | 2831087 | 5609006.5 | 4136360.5 | 5450434 | 4450195.5 | 4528916 |
| P61620 | Sec61a1 | Protein transport protein Sec61 subunit alpha isof | 252.84 | 15.55 | 52265 | 4 | 1402717.62 | 1926850.38 | 1740439.38 | 1791927.5 | 1440101.62 | 4170903 | 3322270.25 | 1810671.5 | 1626107.5 | 1599147.5 | 1819699.75 | 1807075.75 | 1723705.12 | 2575991.5 | 3376953.5 | 1524922.12 | 4327380 | 2663520.5 | 3413177.75 | 4598139.5 |
| Q9CQS8 | Sec61b | Protein transport protein Sec61 subunit beta | 164.88 | 37.5 | 9958 | 3 | 9520323 | 13591985 | 4071008.5 | 16225893 | 7640491.5 | 13877716 | 13754185 | 9474208 | 14095850 | 12047508 | 9345631 | 9662060 | 9189460 | 11393982 | 8865702 | 4764826 | 8719901 | 7923220.5 | 6479551.5 | 7497225 |
| P60060 | Sec61g | Protein transport protein Sec61 subunit gamma | 114.04 | 29.41 | 7741 | 2 | 2145020.25 | 4049843.75 | 2268142.75 | 2257530.5 | 2828570.25 | 2963437 | 2644186 | 1561022.62 | 3605219 | 3012158.5 | 1474205 | 1136984.62 | 802669.62 | 2148376.5 | 2 | 1142242.38 | 945471.56 | 819531.19 | 1675090.5 | 2 |
| Q8BU14 | Sec62 | Translocation protein SEC62 | 83.67 | 5.53 | 45581 | 2 | 1776457 | 2031555.88 | 388015 | 1964066.38 | 530956.62 | 3028095.25 | 2 | 837007.88 | 3924183.5 | 2 | 1595504.12 | 1551912.62 | 2228203 | 1924537.38 | 1255539.75 | 1913078.5 | 2991016.5 | 2435707.5 | 3470420.5 | 1897155.62 |
| Q8R2U0 | Seh1l | Nucleoporin SEH1 | 64.48 | 5 | 39775 | 2 | 630559.56 | 502315.38 | 2 | 2 | 2 | 2 | 2 | 2 | 537699.38 | 2 | 610583.06 | 2 | 722777.56 | 588793.88 | 2 | 2 | 2 | 621146.31 | 2 | 891637.06 |
| Q80Z70 | Sel1l | Protein sel-1 homolog 1 | 121.77 | 3.9 | 88683 | 2 | 165631.05 | 836256.94 | 2 | 1378978.88 | 104166.27 | 635670.81 | 766620.94 | 172540.11 | 1583795.88 | 2 | 2 | 1637969.62 | 2 | 2 | 2 | 2 | 581523 | 2 | 2 | 2 |
| Q9DBC0 | Selenoo | Protein adenylyltransferase SelO, mitochondrial | 44.01 | 1.05 | 74221 | 1 | 2 | 2 | 2 | 2 | 2 | 2 | 2 | 2 | 2 | 2 | 629057.69 | 586402.81 | 670048.25 | 864180.44 | 2 | 594293.56 | 757800.31 | 319366.25 | 2 | 2 |
| Q9EP97 | Senp3 | Sentrin-specific protease 3 | 50.71 | 3.52 | 64403 | 1 | 2 | 2 | 2 | 2 | 2 | 2 | 2 | 2 | 419764.31 | 2 | 2 | 2 | 2 | 2 | 2 | 2 | 2 | 2 | 2 | 2 |
| P97364 | Sephs2 | Selenide, water dikinase 2 | 36.13 | 1.55 | 47834 | 1 | 2 | 2 | 2 | 2 | 2 | 2 | 2 | 2 | 2 | 2 | 1082097 | 895160.5 | 471102.22 | 458643.84 | 2 | 2 | 2 | 2 | 654895.25 | 2 |
| Q5EB96 | Septin1 | Septin-1 | 41.99 | 1.91 | 42058 | 1 | 4564255.5 | 2 | 3868124.5 | 2992001 | 712529.44 | 3116162 | 514345.06 | 2 | 2 | 1981077.38 | 1929690.25 | 1698025.5 | 2353164 | 2 | 2486669 | 495290.5 | 3630669.25 | 2968905.75 | 2095686.75 | 397426.81 |
| B3GNI6 | Septin11 | Septin-11 | 109.34 | 6.73 | 49695 | 3 | 969637.19 | 1470152.25 | 992861 | 1649399.75 | 3925346 | 2960330.75 | 3479507.75 | 4204803 | 3271093.5 | 2640684.25 | 4150009 | 4266533.5 | 3491180.75 | 3156219.5 | 2927760.75 | 3784520.75 | 3947603.25 | 3784795 | 5554668.5 | 3902982.75 |
| P42208 | Septin2 | Septin-2 | 473.32 | 32.13 | 41526 | 8 | 3413115.25 | 9212887 | 2160453.5 | 3023925.5 | 6134572.5 | 5360385 | 4537119.5 | 4073613 | 5352461 | 4275302 | 4462783 | 3432128.5 | 3722897 | 3758437 | 3324623.5 | 6856684.5 | 1580541.12 | 4892052 | 9074349 | 6604409 |
| O55131 | Septin7 | Septin-7 | 256.45 | 12.84 | 50550 | 6 | 342020 | 2 | 47517.2 | 907546.62 | 1470197.12 | 1376143.75 | 796497.38 | 2 | 467661.47 | 2 | 2853167.75 | 3972349.75 | 1666445.12 | 1936023.38 | 2215022.5 | 3294660.5 | 1375009.75 | 2271856.25 | 429828.72 | 685594.81 |
| Q8CHH9 | Septin8 | Septin-8 | 289.12 | 12.12 | 49812 | 5 | 3168328.75 | 790347.69 | 49502.61 | 183969.53 | 444451.81 | 2588252 | 2 | 88398.19 | 636710.31 | 2 | 2476879.75 | 3598256.75 | 4063481.5 | 3902357.25 | 4929576 | 3257078.25 | 5464676 | 3725837.75 | 577234.25 | 6313775.5 |
| Q80UG5 | Septin9 | Septin-9 | 48.48 | 1.54 | 65575 | 1 | 2 | 2016806.5 | 1999283.25 | 1004326.94 | 2 | 2079549.75 | 1183993.25 | 2055612.25 | 1304532.5 | 1111424 | 1516689.38 | 895661.56 | 1342151.12 | 1053931.5 | 2 | 455196.12 | 2 | 2 | 2 | 1587684 |
| Q6AXS5 | Serbp1 | SERPINE1 mRNA-binding protein 1 | 208.1 | 9.34 | 44754 | 3 | 336372.62 | 1181235.5 | 453321.28 | 1808714 | 358447.94 | 335241.75 | 938477.25 | 870368.62 | 2982647 | 195396.56 | 1364095.75 | 653713.38 | 1739799.5 | 3768404 | 2702147.75 | 1862686 | 2428350.5 | 1080203.38 | 2 | 3220383.25 |
| Q9EPB5 | Serhl | Serine hydrolase-like protein | 67.84 | 2.89 | 35311 | 1 | 595301.94 | 692404.12 | 2 | 2 | 874627.56 | 1313370.75 | 756807.5 | 2 | 642665.94 | 2 | 2 | 2 | 2 | 2 | 2 | 2 | 2 | 2 | 2 | 2 |
| Q60854 | Serpinb6 | Serpin B6 | 410.59 | 16.93 | 42599 | 6 | 48418850 | 56533778 | 47686474 | 40573434 | 38802646 | 42597970 | 40639030 | 55586614 | 33388056 | 35879454 | 25359872 | 24548882 | 24316266 | 23957408 | 21809752 | 13469201 | 17514252 | 26898156 | 22572436 | 25916026 |
| P32261 | Serpinc1 | Antithrombin-III | 194.37 | 9.03 | 52004 | 4 | 15376298 | 4411473 | 15051972 | 20656882 | 10280195 | 5146518.5 | 5930419 | 13018703 | 16798842 | 9013863 | 2873241.75 | 3827876.5 | 5890942.5 | 3566920.5 | 4130134 | 2261847.75 | 6497517 | 6920275 | 1460917.25 | 7497973 |
| Q63945 | Set | Protein SET | 235.8 | 21.45 | 33406 | 5 | 5081914 | 5875709.5 | 7138754.5 | 4866965 | 6238439.5 | 4204240 | 4302546 | 6106717 | 2825121.75 | 4059759 | 2599681.25 | 2759130.5 | 2734282.25 | 2894488.5 | 2959823.5 | 3547256.25 | 2725510.25 | 1828033.75 | 2538033.5 | 2779915.75 |
| Q64213 | Sf1 | Splicing factor 1 | 364.33 | 13.63 | 70403 | 7 | 2724250.75 | 2156467.25 | 1531398.12 | 4250844 | 1700350 | 2274204.25 | 2646129.5 | 405504.06 | 4725358.5 | 4029427.25 | 6785057 | 7441526.5 | 6169735 | 6316582 | 7599611.5 | 9379657 | 9390071 | 4517537 | 8701181 | 6432799 |
| Q8K4Z5 | Sf3a1 | Splicing factor 3A subunit 1 | 639.38 | 17.57 | 88545 | 14 | 3930070.75 | 4930654 | 3553024 | 1939079.12 | 5616378 | 2 | 2 | 2130397.75 | 1625534 | 7374083.5 | 7709915.5 | 8079193.5 | 7276024 | 6412718 | 3184237.75 | 7809245.5 | 7046720 | 3831667.75 | 3622790 | 4489072.5 |
| Q62203 | Sf3a2 | Splicing factor 3A subunit 2 | 40.36 | 5.68 | 49911 | 1 | 119476.59 | 2 | 2 | 2 | 2 | 2 | 2 | 2 | 2 | 2 | 2 | 2 | 2 | 2 | 2 | 2 | 2 | 2 | 2 | 482936.06 |
| Q6AXT8 | Sf3a2 | Splicing factor 3A subunit 2 | 39.46 | 2.12 | 49890 | 1 | 2 | 2 | 2 | 2 | 26950.09 | 2 | 2 | 2 | 884222 | 2 | 2 | 806166.75 | 2 | 2 | 117137.32 | 2 | 2 | 2 | 2 | 2 |
| Q9D554 | Sf3a3 | Splicing factor 3A subunit 3 | 110.9 | 4.19 | 58842 | 2 | 2 | 2 | 2 | 2 | 2 | 677359.12 | 2 | 2 | 2 | 2 | 1119568.62 | 1966698.25 | 849718.88 | 795634.5 | 2 | 157756.59 | 994636.75 | 2 | 148083.53 | 188295.61 |
| Q99NB9 | Sf3b1 | Splicing factor 3B subunit 1 | 68.15 | 2.07 | 145816 | 2 | 2 | 2 | 793170.38 | 1050710 | 1131471.88 | 2 | 886796.38 | 2 | 1263125.62 | 767382.69 | 1929680.75 | 1538554.75 | 618065.31 | 2229910.75 | 1376404 | 573777.31 | 1435856.75 | 1596900 | 1113056.38 | 2058686.5 |
| Q3UJB0 | Sf3b2 | Splicing factor 3B subunit 2 | 211.72 | 6.61 | 98198 | 5 | 2098247.75 | 1164548.75 | 2841488 | 1621092 | 1729832.25 | 923039.88 | 50424.59 | 3758274 | 47432.8 | 1606369 | 2569586 | 2096565 | 3360779.75 | 2042412 | 2611448.75 | 2227196 | 1827050.75 | 2944913.5 | 1756959.25 | 3210290.5 |
| Q921M3 | Sf3b3 | Splicing factor 3B subunit 3 | 875.09 | 15.12 | 135550 | 14 | 1870964 | 1124345.5 | 756280.88 | 3090305 | 6180642.5 | 3174257.25 | 4273065.5 | 2046937.38 | 6965210.5 | 4384440 | 3469216.25 | 6174332 | 2189580.5 | 2492249.75 | 2244944.5 | 4118052 | 1054244 | 578575.44 | 80715.04 | 2412718 |
| Q6AYL5 | Sf3b4 | Splicing factor 3B subunit 4 | 196.33 | 9.2 | 44356 | 3 | 2380096 | 2 | 3558155 | 1973722 | 1686125 | 1723705.12 | 777305.25 | 2 | 1092180.75 | 855136.94 | 2451252.5 | 2615171 | 1978833.38 | 2134061 | 3042724.75 | 2087197.62 | 2036083.75 | 2829921.5 | 2960554.5 | 2766423 |
| P59708 | Sf3b6 | Splicing factor 3B subunit 6 | 45.59 | 13.6 | 14585 | 1 | 351615.25 | 1130605.5 | 2 | 574880.75 | 730644.81 | 508494.97 | 2 | 2 | 2 | 314895.88 | 2 | 2 | 503307.41 | 2 | 2 | 2 | 2 | 2 | 2 | 2 |
| Q8VIJ6 | Sfpq | Splicing factor, proline- and glutamine-rich | 1081.07 | 22.6 | 75442 | 18 | 13352836 | 14040730 | 14218826 | 25636578 | 27045570 | 22461662 | 21135948 | 23781398 | 22776254 | 24663710 | 18621356 | 19674096 | 12490760 | 12165160 | 13482568 | 16777658 | 12053561 | 13486782 | 11092566 | 9577128 |
| Q8BP27 | Sfr1 | Swi5-dependent recombination DNA repair protei | 37.65 | 3.76 | 35183 | 1 | 2076389.38 | 1937941.88 | 2 | 1763676.75 | 26541.71 | 760002.19 | 1631795.5 | 130650.71 | 1022511 | 2666759.25 | 1135970.88 | 2 | 951618.31 | 938204.06 | 1499435 | 2446070.25 | 1830980.12 | 1924690.25 | 195906.16 | 2 |

| | | | | | | | | | | | | | | | | | | | | | | | | | | |
|---|---|---|---|---|---|---|---|---|---|---|---|---|---|---|---|---|---|---|---|---|---|---|---|---|---|---|
| Q99JR1 | Sfxn1 | Sideroflexin-1 | 560.87 | 39.13 | 35649 | 9 | 4256892.5 | 8779038 | 4886308 | 2733055.5 | 6761225.5 | 6314597 | 4864734 | 6287444.5 | 6950208 | 4216368.5 | 6109861 | 5026741 | 5574662 | 6830980 | 2519197.5 | 4508849 | 5160282 | 4578100.5 | 5933253.5 | 3997711.5 |
| Q91V61 | Sfxn3 | Sideroflexin-3 | 306.56 | 25.23 | 35406 | 5 | 619489 | 1559021.88 | 1461909.5 | 269263.53 | 1108060.62 | 537835.75 | 2754412 | 1372603.62 | 3698677 | 1662431.12 | 124890.12 | 1591299.62 | 1517141.88 | 1860329.5 | 734707.69 | 2 | 1314200 | 787599.88 | 204960.81 | 1278392.12 |
| Q8R0X7 | Sgpl1 | Sphingosine-1-phosphate lyase 1 | 293.64 | 10.39 | 63677 | 6 | 3277166 | 3208282.75 | 1836206.38 | 2110427 | 1801735.38 | 4138422.5 | 7370894.5 | 4896744 | 3698436.5 | 2876605.25 | 2066917.5 | 1893903.25 | 4435955 | 5159214.5 | 3508347 | 2714391.5 | 2811541.5 | 4023402.75 | 2605945.25 | 3136037.25 |
| Q9JJU8 | Sh3bgrl | Adapter Sh3bgrl | 284.55 | 36.84 | 12811 | 5 | 1252862.88 | 3514570.75 | 1335081.38 | 2603994.75 | 9200624 | 3478747.5 | 4149721.5 | 6681299 | 4510806 | 4524245 | 10626893 | 9833746 | 5149711 | 4776162.5 | 7976742 | 11767842 | 6396592 | 2305886.5 | 10296763 | 2115837.25 |
| B2RZ27 | Sh3bgrl3 | SH3 domain-binding glutamic acid-rich-like protei | 210.07 | 43.01 | 10477 | 4 | 3611524.5 | 8669733 | 3520014.25 | 5870936 | 7832764 | 10169950 | 8295098 | 10009271 | 9229872 | 5889823 | 23062958 | 18969366 | 10383930 | 10749836 | 7581504.5 | 13327269 | 6339918 | 5555921 | 9667982 | 6914198.5 |
| O35964 | Sh3gl1 | Endophilin-A2 | 79.44 | 4.35 | 41492 | 2 | 2169741.75 | 2287565.5 | 1702668.75 | 3912312 | 3508092 | 5451887 | 4054999.25 | 1436220 | 668298.5 | 2031117.75 | 4530376 | 5210225.5 | 1030446.94 | 1065948.5 | 3754517 | 986624.88 | 6106159 | 5916669.5 | 6898192.5 | 2487879 |
| Q6AYE2 | Sh3glb1 | Endophilin-B1 | 219.43 | 8.77 | 40787 | 3 | 1555344.25 | 2 | 1647417.75 | 1749817.5 | 1248006.5 | 195956.97 | 1249288.12 | 1173815.25 | 737781.38 | 323551.31 | 644608.62 | 644966.5 | 2 | 168417.84 | 306774.66 | 708073.25 | 2 | 1004678.56 | 2 | 2 |
| A2AAY5 | Sh3pxd2b | SH3 and PX domain-containing protein 2B | 79.31 | 1.98 | 101517 | 2 | 795969.19 | 1494574 | 2623374 | 697958.38 | 1993911.75 | 2 | 1002840.31 | 1948806.75 | 708620.19 | 1550648.62 | 1792292 | 1585049 | 1732618.5 | 1563197.62 | 1957588.62 | 689114.25 | 598436.25 | 1772109.75 | 120274.87 | 1189412.5 |
| Q61120 | Shc3 | SHC-transforming protein 3 | 41.22 | 1.48 | 52121 | 1 | 2 | 2 | 619523.44 | 2 | 847344.69 | 2 | 2 | 2 | 2 | 605646.38 | 1284178.62 | 1397242.88 | 1083866 | 1622399.25 | 1134821 | 1491329 | 1435371.62 | 2 | 2206034.25 | 1553752.88 |
| Q9CZN7 | Shmt2 | Serine hydroxymethyltransferase, mitochondrial | 1446.65 | 49.8 | 55759 | 23 | 18343610 | 50892590 | 35650490 | 23388876 | 59425138 | 50403050 | 43470974 | 55976630 | 64498570 | 44515966 | 43521354 | 41923094 | 40313462 | 43214690 | 51243022 | 48514354 | 58753122 | 76416178 | 59369874 | 78859482 |
| Q8K2Q9 | Shtn1 | Shootin-1 | 277.91 | 5.71 | 71343 | 5 | 1702657.12 | 2 | 1370353.5 | 2 | 1126417.25 | 2 | 2 | 691321.19 | 2 | 1849761.5 | 2 | 2 | 2 | 2 | 2 | 2 | 2 | 2 | 736665.06 | 2 |
| Q9EPK6 | Sil1 | Nucleotide exchange factor SIL1 | 92.6 | 4.52 | 52430 | 2 | 573655.38 | 2 | 747993.75 | 413647.09 | 2 | 2 | 2 | 2 | 2 | 2 | 314497.19 | 2 | 2 | 2 | 2 | 2 | 2 | 2 | 2 | 2 |
| P97710 | Sirpa | Tyrosine-protein phosphatase non-receptor type | 74.25 | 4.91 | 55691 | 1 | 2 | 249888.92 | 2 | 692102.12 | 2 | 2 | 2 | 216977.92 | 752745 | 2 | 2 | 2 | 2 | 510488.25 | 2 | 750515.81 | 2 | 2 | 437325.88 | 2 |
| P97797 | Sirpa | Tyrosine-protein phosphatase non-receptor type | 300.77 | 16.76 | 56414 | 5 | 26574932 | 29065286 | 28924826 | 23409346 | 19369456 | 8636036 | 13319764 | 13294156 | 12952821 | 14680769 | 16087086 | 20368016 | 16670002 | 17609162 | 16556720 | 10280642 | 17884306 | 20104260 | 18982524 | 19682646 |
| Q5RJQ4 | Sirt2 | NAD-dependent protein deacetylase sirtuin-2 | 129.91 | 7.14 | 39319 | 2 | 1639991.12 | 320345.97 | 1919317 | 530474.88 | 1330271.75 | 389628.28 | 494387.91 | 125364.19 | 1529220.25 | 1515910.75 | 212327.7 | 135426.47 | 1072868.75 | 289867.81 | 682940.5 | 239489.92 | 232507.77 | 286765.59 | 276436.78 | 354979.53 |
| Q3UND0 | Skap2 | Src kinase-associated phosphoprotein 2 | 122.18 | 8.1 | 40712 | 3 | 3969949.25 | 3809952.75 | 3530172.5 | 5301210.5 | 3558785.5 | 3133936.75 | 2973437.5 | 3486063 | 2766528.5 | 2947538.5 | 1311216.75 | 1656630.25 | 1974965.38 | 1279712.75 | 1255237.88 | 2672843 | 1643412.12 | 2004220.75 | 2632103 | 2089896.12 |
| Q6PEC4 | Skp1 | S-phase kinase-associated protein 1 | 81.24 | 10.43 | 18672 | 1 | 993053.75 | 815238.06 | 797888.94 | 882430.62 | 679478.5 | 1004131.44 | 870638.88 | 1169333.12 | 639969.12 | 309336.91 | 816683.5 | 734465.12 | 774393.38 | 619257.44 | 416175.72 | 1961582.62 | 1024839.56 | 1459361.88 | 2882711.5 | 1204314.88 |
| Q8CI08 | Slain2 | SLAIN motif-containing protein 2 | 67.53 | 2.93 | 62378 | 1 | 2 | 2 | 2 | 2 | 2 | 2 | 2 | 2 | 2 | 2 | 2 | 2 | 2 | 2 | 2 | 2 | 2 | 2 | 2 | 2 |
| Q924N4 | Slc12a6 | Solute carrier family 12 member 6 | 83.26 | 1.91 | 127527 | 2 | 1774509.25 | 902963.62 | 1696104.5 | 2277592.75 | 850351 | 961463.44 | 1028188.94 | 2 | 160816.53 | 682079 | 761481.75 | 631636.5 | 1176531.5 | 598881.12 | 1363194.5 | 380718.91 | 106127.49 | 1060602.25 | 955858.62 | 1936344.38 |
| Q99MR3 | Slc12a9 | Solute carrier family 12 member 9 | 56.46 | 1.31 | 96330 | 1 | 2 | 992307.69 | 2 | 2 | 80149.61 | 142204.39 | 402492.12 | 211254.14 | 1183422.25 | 46138.3 | 2 | 702805.94 | 133640.88 | 705572.25 | 387867.62 | 2 | 219872.86 | 65489.87 | 2 | 577844.75 |
| Q8BPX9 | Slc15a3 | Solute carrier family 15 member 3 | 388.9 | 14.19 | 64051 | 6 | 6264685.5 | 5042557.5 | 4089030.25 | 3135996 | 3041179.75 | 9781516 | 8555917 | 11783861 | 8614352 | 4212863 | 2144710.25 | 650049.25 | 1938436.38 | 2131668.75 | 3460962 | 4387438 | 5841909 | 7474697.5 | 5858394 | 3776445.5 |
| P53986 | Slc16a1 | Monocarboxylate transporter 1 | 72.52 | 2.64 | 53267 | 1 | 716029.75 | 516522.94 | 2 | 884139.88 | 1131496.12 | 2 | 2 | 1252688.38 | 649073 | 601508 | 586579.88 | 2 | 797938.31 | 2 | 780645.69 | 566229.31 | 925183.62 | 799665.5 | 2 | 669409.25 |
| Q8BN82 | Slc17a5 | Sialin | 39.85 | 2.42 | 54369 | 1 | 2 | 2 | 2 | 2 | 2 | 2 | 2 | 2 | 2 | 2 | 2 | 2 | 2 | 2 | 2 | 404055.69 | 2 | 2 | 312580.78 | 2 |
| P51912 | Slc1a5 | Neutral amino acid transporter B(0) | 455.44 | 20.25 | 58483 | 8 | 88173.43 | 2 | 98026.14 | 137603.83 | 107941.81 | 91752 | 2 | 133919.66 | 192604.41 | 74424.1 | 176309.69 | 158378.81 | 375649.84 | 116905.12 | 1149368.12 | 78287.69 | 263054.19 | 127542.95 | 850940.56 | 152516 |
| Q8JZU2 | Slc25a1 | Tricarboxylate transport protein, mitochondrial | 166.31 | 14.47 | 33931 | 4 | 2 | 331642.69 | 1398626.25 | 301848.97 | 3782764.75 | 401397.22 | 902915 | 597931.5 | 3202898.75 | 4272143 | 228692.94 | 352001.09 | 1771290.25 | 1093936.75 | 878712.19 | 129317.6 | 101221.56 | 160181.58 | 245816.8 | 147370.58 |
| Q9QZD8 | Slc25a10 | Mitochondrial dicarboxylate carrier | 680.98 | 36.93 | 31715 | 12 | 11981910 | 14999013 | 13635671 | 13621286 | 17404118 | 9275321 | 9050245 | 8613553 | 11941318 | 10718740 | 9006348 | 7019183.5 | 8229059.5 | 9334255 | 6532186.5 | 6385108 | 4662828.5 | 8772034 | 5495926 | 7336750 |
| Q9CR62 | Slc25a11 | Mitochondrial 2-oxoglutarate/malate carrier protei | 853.05 | 44.59 | 34155 | 14 | 21739106 | 25860332 | 20349416 | 17345914 | 16462376 | 20744234 | 21366946 | 23169676 | 17319896 | 13133044 | 13109412 | 8640406 | 14279246 | 15695291 | 11719915 | 10152884 | 12657407 | 15273007 | 10374512 | 13551380 |
| Q8BH59 | Slc25a12 | Electrogenic aspartate/glutamate antiporter SLC2 | 953.37 | 31.02 | 74570 | 17 | 119803.46 | 952059.94 | 1295113.12 | 1380939.38 | 3360817 | 2413487.5 | 2094418.38 | 2435527.5 | 5124230.5 | 4818690 | 188094.91 | 619648.44 | 1392152.88 | 1534685.12 | 1287205.5 | 273674.34 | 748840.5 | 632379.44 | 742326 | 1380117.25 |
| Q9QXX4 | Slc25a13 | Electrogenic aspartate/glutamate antiporter SLC2 | 1082.69 | 34.47 | 74467 | 19 | 5555013.5 | 6932500.5 | 11950155 | 10088588 | 11469342 | 7201308 | 9680147 | 10714861 | 8759943 | 12226088 | 6546184.5 | 4548980.5 | 7981231 | 6433673.5 | 8725593 | 6364873.5 | 5128077 | 10233226 | 6995655.5 | 8403664 |
| Q9Z2Z6 | Slc25a20 | Mitochondrial carnitine/acylcarnitine carrier protei | 134.78 | 13.62 | 33027 | 3 | 3043727.25 | 6241431 | 3453561 | 3420881.5 | 4716905 | 3006792 | 1605235.12 | 5599057.5 | 4030237.5 | 3066437.25 | 3029153.5 | 1448916.75 | 2293662.75 | 2819929.25 | 1213180.25 | 2822424.25 | 3049417 | 3645147.5 | 2741627.75 | 3304117.25 |
| A0A0G2 | Slc25a22 | Mitochondrial glutamate carrier 1 | 142.25 | 8.98 | 34662 | 3 | 487279.06 | 2 | 2 | 2 | 871187.25 | 2 | 1395665.88 | 786894.31 | 2389763.25 | 1155772.38 | 2 | 2 | 683781.81 | 1033965.94 | 2 | 2 | 2 | 2 | 351437.12 | 2 |
| Q8BMD8 | Slc25a24 | Mitochondrial adenyl nucleotide antiporter SLC25 | 385.62 | 19.16 | 52902 | 7 | 849659.69 | 638957.75 | 273005.5 | 997179.75 | 530804 | 2500985.75 | 2669683.75 | 756790.44 | 4605725 | 3778015.25 | 1866918.38 | 3077163.5 | 1291616 | 906523.44 | 1404402.62 | 718274.31 | 1482512 | 1457959.25 | 505072.62 | 1259424 |
| Q8VEM8 | Slc25a3 | Solute carrier family 25 member 3 | 655.08 | 25.77 | 39632 | 11 | 20425196 | 13776986 | 22015926 | 20425466 | 35139310 | 21216226 | 21091706 | 34568086 | 38236374 | 39966026 | 22211022 | 19052622 | 21882902 | 25340514 | 25801416 | 16069540 | 16743976 | 17180330 | 20822586 | 14009741 |
| Q8BMG8 | Slc25a32 | Solute carrier family 25 member 32 | 40.17 | 2.53 | 35049 | 1 | 2 | 560190.44 | 2 | 808309.81 | 680785.88 | 2 | 2 | 754161.62 | 411094.25 | 501492.59 | 2 | 302067.75 | 2 | 442612.66 | 2 | 2 | 2 | 2 | 2 | 2 |
| P48962 | Slc25a4 | ADP/ATP translocase 1 | 1434.49 | 65.1 | 32904 | 24 | 1413365.25 | 592426.06 | 2761223 | 623561.56 | 3604893.5 | 852110.12 | 1631571.25 | 5539696.5 | 1505294.75 | 3412593.25 | 557178 | 545710.75 | 1364854.5 | 696256 | 644378.69 | 2 | 2 | 1808978.75 | 1799099.88 | 2 |
| P51881 | Slc25a5 | ADP/ATP translocase 2 | 2195.81 | 87.25 | 32931 | 36 | 61180690 | 50377158 | 77710394 | 63527962 | 42859174 | 65488154 | 63587426 | 62606898 | 45135498 | 41578738 | 41238990 | 21882838 | 41914242 | 43254434 | 73531818 | 43022434 | 54512866 | 78894146 | 47572330 | 68476394 |
| Q91VE0 | Slc27a4 | Long-chain fatty acid transport protein 4 | 91.45 | 2.95 | 72319 | 2 | 2 | 2 | 2 | 899168.62 | 276162.81 | 1644624 | 205906.97 | 2321963.25 | 1523796.75 | 930046.75 | 869142.5 | 137265.69 | 1567213.75 | 413879.94 | 169991.39 | 2 | 410990.38 | 113147.41 | 2 | 191738.89 |
| Q9JIM1 | Slc29a1 | Equilibrative nucleoside transporter 1 | 41.67 | 2.61 | 50192 | 1 | 850270.44 | 1139089.88 | 2 | 297603.16 | 2 | 2 | 698393.19 | 2 | 290410.81 | 2 | 760414.81 | 841274.94 | 793570.38 | 768342.19 | 598226.81 | 715300.75 | 1052073.75 | 1215952.38 | 1062730.38 | 1470197.25 |
| P11167 | Slc2a1 | Solute carrier family 2, facilitated glucose transp | 402.01 | 15.65 | 53963 | 7 | 803247.44 | 722347.81 | 932750.88 | 2 | 288093.06 | 1689649 | 1354138.38 | 2 | 1230539.88 | 1316460.5 | 1332307 | 2 | 1205005.88 | 1145320.25 | 1404022.88 | 2075877 | 2347399 | 3064323.75 | 2075125.75 | 1838276 |
| P97858 | Slc35b1 | Solute carrier family 35 member B1 | 32.55 | 3.11 | 35883 | 1 | 2 | 100178.52 | 2 | 2 | 2 | 2 | 2 | 2 | 130688.76 | 2 | 2 | 2 | 2 | 80484.47 | 2 | 2 | 2 | 132141.25 | 2 | 161195.22 |
| Q91ZN5 | Slc35b2 | Adenosine 3'-phospho 5'-phosphosulfate transpo | 58.37 | 2.32 | 47370 | 1 | 440874.53 | 453175 | 2 | 2 | 2 | 2 | 2 | 2 | 2 | 540816.75 | 2 | 2 | 671666.62 | 840743.25 | 2 | 2 | 2 | 2 | 592325.81 | 2 |
| Q9WU81 | Slc37a2 | Glucose-6-phosphate exchanger SLC37A2 | 114.92 | 4.59 | 55073 | 2 | 2 | 2 | 2 | 112366.64 | 2 | 2 | 760900.25 | 865967.25 | 1548698.12 | 91973.35 | 2 | 161975.66 | 107900.66 | 228440.69 | 261680.81 | 2 | 467682.19 | 156491.64 | 176891.95 | 2 |
| Q6P6S2 | Slc39a11 | Zinc transporter ZIP11 | 73.46 | 6.57 | 34868 | 1 | 2 | 2 | 82907.37 | 2 | 57746.68 | 194698.83 | 89912.07 | 40621.35 | 179938.14 | 62993.93 | 2 | 79965.37 | 70554.09 | 70128.88 | 369621.44 | 2 | 134660.39 | 92932.43 | 2 | 95527.31 |
| Q31125 | Slc39a7 | Zinc transporter SLC39A7 | 50.09 | 2.73 | 50657 | 1 | 897330.19 | 1021811.75 | 2 | 831514.5 | 2 | 2 | 1530696.12 | 173368.66 | 802396.38 | 874210.5 | 681944.38 | 802745.75 | 911709.88 | 677889.56 | 978158.94 | 780758.75 | 1484124.75 | 1090307 | 245727.77 | 944528.31 |
| P10852 | Slc3a2 | Amino acid transporter heavy chain SLC3A2 | 694.44 | 24.33 | 58337 | 12 | 4828608.5 | 869579.75 | 3831578.5 | 3619621 | 7852670 | 4561983.5 | 2641493.5 | 4243965 | 7806160.5 | 7051695.5 | 9361467 | 7813406 | 5911198 | 5812978.5 | 4981272 | 9983170 | 7872445 | 8589901 | 3611284.25 | 6523785 |
| Q8BTY2 | Slc4a7 | Sodium bicarbonate cotransporter 3 | 166.08 | 3.87 | 116514 | 3 | 2 | 2 | 2 | 2 | 2 | 2 | 2 | 2 | 2 | 2 | 2 | 2 | 2 | 2 | 162605.12 | 2 | 2 | 72681.22 | 2 | 2 |
| Q9CXK4 | Slc50a1 | Sugar transporter SWEET1 | 52.73 | 9.05 | 24649 | 1 | 2 | 2 | 377141.88 | 958640.44 | 124974.48 | 2 | 2 | 456797.12 | 1711521 | 679381.06 | 2 | 2 | 67360.34 | 381538.75 | 1789828.5 | 2 | 2 | 575406.12 | 70285.04 | 623564.81 |
| Q63016 | Slc7a5 | Large neutral amino acids transporter small subu | 115.89 | 4.49 | 55903 | 2 | 1183247 | 1416120.12 | 1563635.25 | 2729067.5 | 1800372.25 | 1316543.25 | 980479.94 | 2672236.75 | 1288088.75 | 1307767.38 | 1403804.25 | 1391890 | 1550898.12 | 1650403.88 | 1190342.88 | 1399192 | 1976446.5 | 3298841.5 | 3112213 | 2837092.25 |
| Q9QXW9 | Slc7a8 | Large neutral amino acids transporter small subu | 32.81 | 1.32 | 57873 | 1 | 2 | 2 | 2 | 2 | 2 | 2 | 2 | 2 | 2 | 2 | 1005184.25 | 2 | 2 | 598975.12 | 2 | 553759.62 | 2 | 2 | 2 | 2 |
| Q6PGB8 | Smarca1 | Probable global transcription activator SNF2L1 | 47.93 | 1.05 | 121715 | 1 | 2 | 2 | 2 | 2 | 965414 | 2 | 2 | 2 | 454865.09 | 596406.44 | 2 | 1145250.62 | 956453 | 2 | 2 | 2 | 2 | 2 | 2 | 2 |
| Q3TKT4 | Smarca4 | Transcription activator BRG1 | 77.05 | 1.18 | 181427 | 2 | 2 | 383343.91 | 27911.75 | 2 | 240261.16 | 497291.88 | 2 | 2 | 1461737.5 | 290946.62 | 969621.94 | 1158831 | 585903.31 | 724990.69 | 2 | 908352.19 | 2 | 311756.19 | 590829.44 | 136166.2 |
| Q9Z0H3 | Smarcb1 | SWI/SNF-related matrix-associated actin-depend | 132.03 | 9.61 | 44141 | 3 | 495760.38 | 2 | 73555.66 | 2 | 683836.88 | 1779257.75 | 2 | 785006.25 | 718774.44 | 118784.65 | 2426660.5 | 905298.88 | 1634033.5 | 1458385 | 809023.25 | 2398535.75 | 253067.64 | 748226.94 | 2 | 771251.62 |
| P97496 | Smarcc1 | SWI/SNF complex subunit SMARCC1 | 294.59 | 5.98 | 122890 | 5 | 1770615.62 | 2 | 760948.31 | 868118.31 | 385632.62 | 1769075.88 | 2 | 750918.5 | 495149.91 | 494142.34 | 667283.12 | 984667 | 642064.19 | 526324.5 | 2096605.62 | 2 | 1699843.88 | 715339.25 | 236799.42 | 1793455.25 |
| Q6PDG5 | Smarcc2 | SWI/SNF complex subunit SMARCC2 | 232.15 | 4.7 | 132604 | 5 | 2 | 2 | 49806.61 | 75132.52 | 2 | 2 | 2 | 2 | 2 | 70130.18 | 2 | 82862.34 | 62808.57 | 2 | 157841.92 | 2 | 132027.09 | 78717.39 | 89473.69 | 2 |
| Q61466 | Smarcd1 | SWI/SNF-related matrix-associated actin-depend | 110.97 | 5.83 | 58245 | 2 | 5366536.5 | 1609438.62 | 5080815 | 3137432.25 | 1838794 | 4118106 | 4062068.25 | 2836415.25 | 2 | 3178881.75 | 3761493.25 | 3656161.25 | 5564739 | 2 | 5717346 | 2748828 | 4356600 | 2960584.75 | 2 | 4802026 |
| O54772 | Smarcd2 | SWI/SNF-related matrix-associated actin-depend | 108.9 | 5.65 | 59128 | 2 | 2 | 2 | 2 | 2 | 2 | 2 | 2 | 2 | 2 | 2 | 2 | 2 | 2 | 2 | 2 | 2 | 2 | 2 | 2 | 2 |
| Q9CU62 | Smc1a | Structural maintenance of chromosomes protein 1 | 219.21 | 3.73 | 143235 | 5 | 626623.38 | 3756689.25 | 2 | 2839065.5 | 2249286.25 | 254097.53 | 583464.94 | 1061330.25 | 1893049.5 | 1690507.88 | 2651517.5 | 2657219.5 | 2218868.25 | 1597556.12 | 2 | 2263965 | 1121592.38 | 889892.69 | 1040282.25 | 2 |
| Q8CG48 | Smc2 | Structural maintenance of chromosomes protein 2 | 168.97 | 4.03 | 134239 | 4 | 791365.06 | 642979.38 | 270799.66 | 475365.22 | 1527073.88 | 2 | 135077.23 | 137344.39 | 2 | 2 | 493528.97 | 591335.75 | 2 | 567432.62 | 119463.78 | 133863.14 | 2 | 615069.44 | 2 | 2 |
| P97690 | Smc3 | Structural maintenance of chromosomes protein 3 | 119.76 | 2.69 | 138448 | 2 | 2 | 2 | 1834891 | 1161530.38 | 2173009.5 | 2 | 2 | 2 | 979525.62 | 2 | 2 | 1372183.5 | 1114237.38 | 1286346.88 | 2 | 2 | 2 | 2 | 716985 | 2 |
| Q8CG47 | Smc4 | Structural maintenance of chromosomes protein 4 | 185.01 | 2.95 | 146895 | 4 | 5528178 | 7201362 | 4967845.5 | 7203776 | 6178335 | 72138.85 | 439674.94 | 2917331.25 | 1182578.25 | 2097384.5 | 1343523.38 | 2011011.75 | 4972314 | 3593079 | 3181670.75 | 2678003.5 | 2090249.25 | 3940062 | 5139125 | 4749610 |
| Q6P5D8 | Smchd1 | Structural maintenance of chromosomes flexible h | 125.54 | 1.1 | 225648 | 2 | 2 | 2 | 31257.9 | 78301.5 | 571771.88 | 2 | 141784.7 | 2 | 85945.36 | 61107 | 2 | 2 | 58707.88 | 84701.48 | 2 | 2 | 2 | 2 | 2 | 139995.58 |
| P58242 | Smpdl3b | Acid sphingomyelinase-like phosphodiesterase 3l | 63.35 | 3.51 | 51600 | 1 | 1312011.75 | 2029086 | 1103391.88 | 1974838.38 | 2040837 | 2 | 2 | 1566218.5 | 1284049 | 1208419.75 | 848481.69 | 794241.12 | 1215976.62 | 1116069.75 | 1203743.12 | 738984.5 | 784297.81 | 1698597.75 | 2 | 1254242.62 |
| Q3UKJ7 | Smu1 | WD40 repeat-containing protein SMU1 | 236.28 | 13.06 | 57544 | 5 | 2103904.5 | 2363103.5 | 1655161.75 | 2483611 | 2700870.5 | 1869577.12 | 1529421.75 | 2375077.75 | 2040121.12 | 1618905.38 | 2233271.25 | 2069099.12 | 1038147.56 | 3074681.5 | 1297256.12 | 1090552.75 | 1110463.38 | 1560750 | 1731684 | 529472.62 |
| Q3TYX3 | Smyd5 | Protein-lysine N-trimethyltransferase SMYD5 | 57 | 2.16 | 47095 | 1 | 492967.62 | 774080.44 | 2 | 1028813 | 1041938.81 | 848436.31 | 2 | 735281.56 | 917669.88 | 745113.44 | 323945.69 | 2 | 482670.38 | 2 | 2 | 2 | 2 | 2 | 2 | 2 |
| O09044 | Snap23 | Synaptosomal-associated protein 23 | 39.54 | 6.67 | 23261 | 1 | 622546.44 | 2 | 802851.25 | 159152.31 | 562638.06 | 189548.5 | 2 | 2 | 2 | 571002.12 | 2 | 2 | 2 | 386220.06 | 585487.69 | 2 | 2 | 624273.75 | 2 | 734014.75 |
| Q78PY7 | Snd1 | Staphylococcal nuclease domain-containing prote | 1523.43 | 35.82 | 102088 | 27 | 13557361 | 15999784 | 25431962 | 18797906 | 32657544 | 18450150 | 18719998 | 26071130 | 24718112 | 20430772 | 18391410 | 20532748 | 14767179 | 14193857 | 10407472 | 19399968 | 12966351 | 17094382 | 12546780 | 15188386 |
| Q5RK19 | Snf8 | Vacuolar-sorting protein SNF8 | 47.42 | 5.43 | 28884 | 1 | 2 | 2 | 558141.38 | 2 | 2 | 2 | 2 | 2 | 2 | 2 | 698703.5 | 870320.19 | 759732.56 | 2 | 2 | 2 | 802282.88 | 825523.5 | 884075.56 | 791827.88 |
| F1LNJ2 | Snrnp200 | U5 small nuclear ribonucleoprotein 200 kDa helic | 706.67 | 8.42 | 244875 | 16 | 11609262 | 12076759 | 13563099 | 11279108 | 13219409 | 7241474.5 | 5276201 | 4889992 | 5463917 | 6162063.5 | 6777222 | 4417565 | 9723926 | 10284704 | 7422278 | 6912283 | 6573443 | 11464402 | 4919321 | 7694628.5 |
| Q62376 | Snrnp70 | U1 small nuclear ribonucleoprotein 70 kDa | 194.54 | 9.6 | 51992 | 4 | 225463.02 | 188153.95 | 2 | 515791.28 | 2 | 2 | 2 | 2 | 982891.62 | 2 | 1564352.5 | 1126778 | 2 | 826205.12 | 2 | 1365898.5 | 2 | 2 | 2 | 2 |
| Q62189 | Snrpa | U1 small nuclear ribonucleoprotein A | 225.61 | 20.56 | 31835 | 4 | 2209829 | 3074978.75 | 1437209.38 | 5518651.5 | 4161321.75 | 3973619.75 | 3364080 | 4565974.5 | 3819080 | 3227592.5 | 708733.88 | 2431276.75 | 4053973 | 3792675.75 | 3694833.5 | 2 | 5149061 | 3860767.25 | 2884007.5 | 4551881.5 |
| P57784 | Snrpa1 | U2 small nuclear ribonucleoprotein A' | 397.76 | 29.8 | 28357 | 6 | 2204023.25 | 1025539.94 | 111556.84 | 744483.69 | 2407343.75 | 1917511.38 | 1853503.62 | 733706.12 | 2629605.75 | 994366.38 | 302671.28 | 691414.75 | 1769013.12 | 891828 | 204116.16 | 193688.19 | 1277283.25 | 997372.88 | 247737.48 | 1221074.62 |
| P17136 | Snrpb | Small nuclear ribonucleoprotein-associated protei | 173.16 | 16.02 | 23656 | 4 | 959371.31 | 1442878.12 | 918028.88 | 1791151.12 | 6923009 | 7052762 | 7067983 | 7781420.5 | 9756152 | 8229198 | 10881332 | 13079494 | 12061528 | 11410650 | 6855426.5 | 7027922.5 | 6914523.5 | 4168708.25 | 5564590 | 3979558.25 |
| Q9CQI7 | Snrpb2 | U2 small nuclear ribonucleoprotein B" | 85.25 | 8 | 25323 | 2 | 911526 | 2 | 1240067.12 | 2 | 2344406.25 | 2 | 991691.06 | 183146.78 | 1139041.12 | 1403251.75 | 2 | 769399.75 | 1425667.38 | 1203764 | 714524.88 | 1246021.38 | 2071610.88 | 3659164.5 | 2 | 2406089.75 |
| P62315 | Snrpd1 | Small nuclear ribonucleoprotein Sm D1 | 55.54 | 10.92 | 13282 | 1 | 400947.28 | 812720.31 | 2 | 2 | 1321106.25 | 1614984.5 | 530702.5 | 1017905.88 | 1379937.12 | 837957.12 | 3986705.25 | 3772382.25 | 2456251.75 | 2700423.5 | 2095967 | 3215754.5 | 3325476.75 | 1378213.5 | 2812080 | 2614185.75 |
| P62317 | Snrpd2 | Small nuclear ribonucleoprotein Sm D2 | 384.63 | 55.93 | 13527 | 8 | 1972686.38 | 955544.38 | 1060008 | 969547.75 | 2107569.75 | 1280143 | 1063495.38 | 380113.69 | 797845.38 | 316524.84 | 10602170 | 3380176 | 2357458 | 1657315 | 1295214.12 | 2840360 | 2813428 | 2622497.25 | 2012143.38 | 3001756 |
| P62320 | Snrpd3 | Small nuclear ribonucleoprotein Sm D3 | 135.47 | 32.54 | 13916 | 3 | 1915268.88 | 6486596.5 | 4338117 | 3730228.5 | 9494886 | 12693957 | 9783863 | 13306313 | 14228131 | 10034420 | 20549694 | 17782678 | 19691886 | 16158086 | 14915846 | 21357836 | 21218120 | 18592006 | 24062486 | 17204178 |
| P62305 | Snrpe | Small nuclear ribonucleoprotein E | 146.3 | 32.61 | 10804 | 3 | 19302642 | 5734357 | 1787317.12 | 17627158 | 4382955.5 | 1138073.5 | 7885479.5 | 5407990 | 11690810 | 5639588.5 | 1856147.12 | 4468181 | 3393863.75 | 6406341 | 5239727 | 344180.22 | 5354351.5 | 1555454.75 | 1981640.12 | 1991483.12 |
| P62307 | Snrpf | Small nuclear ribonucleoprotein F | 44.57 | 9.3 | 9725 | 1 | 2 | 2 | 2 | 2 | 880130.06 | 2 | 1243931.5 | 2 | 2 | 1090318.5 | 2615360.25 | 1986050.62 | 2 | 3129813 | 1380557.25 | 3092583 | 2108511.5 | 2667458.75 | 3998551.5 | 1492754 |
| P62309 | Snrpg | Small nuclear ribonucleoprotein G | 127.74 | 17.11 | 8496 | 2 | 2 | 2 | 2151146.5 | 2 | 1338476.38 | 2 | 525273.19 | 451406.72 | 716235.94 | 1733724 | 1763941.5 | 1867547.62 | 2559691.25 | 1540868.25 | 2 | 94344.94 | 2 | 70850.92 | 233817.36 | 275587.19 |
| P55770 | Snu13 | NHP2-like protein 1 | 215.87 | 26.56 | 14174 | 4 | 716985.75 | 5479069 | 3112558.5 | 4750598 | 5763961 | 1333094.62 | 5186628.5 | 5359398.5 | 5487953 | 4348679 | 1849778.25 | 1009489.56 | 3823517.25 | 3571267.75 | 2422290.5 | 2 | 3273322.25 | 3133043 | 2 | 3406690.75 |
| Q9CSN1 | Snw1 | SNW domain-containing protein 1 | 369.86 | 17.54 | 61475 | 8 | 2 | 3225260 | 2 | 748591.12 | 2 | 1584166.88 | 2725309 | 3213320.5 | 2799418.75 | 99503.89 | 3965288 | 3768437 | 3778947 | 4548942 | 1161029.88 | 1799121.25 | 2154485 | 658319.88 | 892098.56 | 769261 |
| Q99N27 | Snx1 | Sorting nexin-1 | 375.92 | 14.94 | 59044 | 6 | 9305294 | 15253178 | 9039620 | 7413306.5 | 11405858 | 15415834 | 12713702 | 17598040 | 9389976 | 12261742 | 10623064 | 9758142 | 10748122 | 10007036 | 9803792 | 9369171 | 8312613 | 10116861 | 9354590 | 12576574 |
| O70493 | Snx12 | Sorting nexin-12 | 32.92 | 5.45 | 19116 | 1 | 945589.62 | 1232910.25 | 694029.69 | 1178855.88 | 1118193.38 | 1521631.12 | 1203373.25 | 2103134.25 | 1280600 | 876341.56 | 937032.38 | 1388668.62 | 1199340.62 | 1004533.19 | 935035.38 | 1778018 | 1688404.38 | 1398538.62 | 1253892.75 | 1512558.5 |
| Q91ZR2 | Snx18 | Sorting nexin-18 | 120.91 | 4.72 | 67904 | 2 | 2 | 2 | 2 | 2 | 1253223.88 | 2 | 615557.31 | 214317.56 | 1445289 | 721114.69 | 1416148.75 | 931362.19 | 1515051.75 | 1376309.5 | 864104 | 2277401 | 1845058.12 | 2413709.5 | 1883922.62 | 2676502.5 |
| Q9CWK8 | Snx2 | Sorting nexin-2 | 441.59 | 20.62 | 58471 | 8 | 15229462 | 26394224 | 16684792 | 8445755 | 19174902 | 14183650 | 10789957 | 16664120 | 14056521 | 13477704 | 23744594 | 21314340 | 19415564 | 19922292 | 13161802 | 21245834 | 14654006 | 14984837 | 18430016 | 17306860 |
| Q91YJ2 | Snx4 | Sorting nexin-4 | 68.7 | 2 | 51778 | 1 | 2 | 2 | 303658.69 | 2 | 312945.34 | 2 | 123837.82 | 2 | 171748.31 | 639657.38 | 2 | 110358.84 | 670487.38 | 271570.91 | 2 | 2 | 236963.61 | 2 | 119267.38 | 2 |
| Q9D8U8 | Snx5 | Sorting nexin-5 | 506.58 | 23.76 | 46797 | 9 | 7200767.5 | 12206003 | 8573238 | 8854966 | 10358931 | 10265806 | 10098172 | 9958779 | 8640344 | 8898369 | 7506580 | 10219492 | 7538589 | 7211974 | 6783055 | 8745164 | 6128160 | 6641857.5 | 9656095 | 7710712.5 |
| Q91VH2 | Snx9 | Sorting nexin-9 | 41.16 | 2.35 | 66546 | 1 | 640972 | 2 | 2 | 920400.5 | 2 | 2 | 2 | 2366741 | 680778.12 | 2 | 2 | 671052.44 | 2761297 | 2723131.75 | 2 | 3946334.5 | 1180095.88 | 4082092.5 | 5129939 | 4206927 |
| O70536 | Soat1 | Sterol O-acyltransferase 1 | 48.49 | 1.65 | 64146 | 1 | 2 | 2 | 2 | 2 | 2 | 2 | 2 | 44647.41 | 57430.05 | 40230.52 | 2 | 2 | 54191.22 | 2 | 148488.78 | 2 | 71317.74 | 137906.39 | 2 | 2 |
| P08228 | Sod1 | Superoxide dismutase [Cu-Zn] | 509.34 | 45.45 | 15943 | 8 | 24150502 | 23647626 | 31878830 | 39941010 | 22859450 | 20888574 | 22749946 | 21974488 | 15163818 | 20919036 | 17676312 | 19743678 | 17052652 | 12425927 | 26402098 | 30140050 | 39744678 | 52103954 | 29906966 | 37129742 |
| P09671 | Sod2 | Superoxide dismutase [Mn], mitochondrial | 181.22 | 16.67 | 24603 | 3 | 792326.25 | 537213.06 | 1285065.5 | 1023283.5 | 2982912.5 | 1471951.5 | 4630082 | 5199713 | 6167571 | 3773316 | 686557.12 | 1009258.38 | 1357944 | 3388065.75 | 1965600.25 | 3392510 | 1064951.88 | 1982810.75 | 3505938.5 | 2238195 |
| Q9QX47 | Son | Protein SON | 149.41 | 1.84 | 265651 | 3 | 2 | 470089.34 | 2 | 2 | 715957.19 | 64406.46 | 2 | 2 | 795468.38 | 2 | 678310 | 976739.44 | 751440.81 | 2 | 507871.09 | 935906.5 | 2 | 2 | 2 | 2 |
| Q64442 | Sord | Sorbitol dehydrogenase | 96.49 | 7 | 38249 | 2 | 2299806.75 | 2 | 1516797.12 | 3737509.5 | 2136200.5 | 2 | 1624842.88 | 2396305.25 | 1568691.12 | 1333642.5 | 4014108.75 | 2826514.5 | 2590321.25 | 2080201 | 2587577.25 | 3444351.5 | 3379877.25 | 4049764.75 | 3887549.25 | 4656626 |

| | | | | | | | | | | | | | | | | | | | | | | | | | | |
|---|---|---|---|---|---|---|---|---|---|---|---|---|---|---|---|---|---|---|---|---|---|---|---|---|---|---|
| O35892 | Sp100 | Nuclear autoantigen Sp-100 | 164.59 | 6.22 | 54727 | 3 | 4150582.75 | 2178992.5 | 4109330.5 | 6343617 | 5014300 | 1640700.5 | 4085536.5 | 6142053.5 | 320638.41 | 463041.81 | 1044814.56 | 926210.31 | 1916685.38 | 2 | 2 | 1704083.62 | 2421518.5 | 3541749 | 1817788.75 | 495062.78 |
| Q8BVK9 | Sp110 | Sp110 nuclear body protein | 243.83 | 14.61 | 50140 | 5 | 51807.29 | 2 | 372559.06 | 100888.59 | 529313.81 | 535467.38 | 124555.94 | 53437.64 | 2 | 678456 | 2 | 1217064 | 65633.95 | 849759.56 | 196357.67 | 2 | 2 | 1418604.62 | 2 | 2 |
| Q58A65 | Spag9 | C-Jun-amino-terminal kinase-interacting protein 4 | 93.83 | 2.35 | 146219 | 2 | 2 | 2 | 2 | 422027.03 | 131188.42 | 2 | 348905.78 | 724417.06 | 2 | 413677.69 | 733878.12 | 2 | 619584.38 | 193169.36 | 1330850.75 | 74805.21 | 179110.8 | 838558.56 | 2 | 2 |
| Q9CYN2 | Spcs2 | Signal peptidase complex subunit 2 | 127.61 | 10.62 | 24978 | 3 | 271372.47 | 2 | 2 | 483714.31 | 1024545.19 | 631177.62 | 579573.12 | 1368770.12 | 1435121.75 | 1434663.25 | 878667.69 | 1109073.75 | 1332435.75 | 1520549.5 | 1505352.38 | 1009167.31 | 1150611 | 1213554.38 | 1059077.75 | 1052337.25 |
| Q6ZWQ7 | Spcs3 | Signal peptidase complex subunit 3 | 247.24 | 21.67 | 20313 | 5 | 4646851 | 1762788.25 | 7690985.5 | 2100074 | 4008491.75 | 1822082.75 | 2394206 | 634932.56 | 5384193.5 | 2785161.5 | 2972906.5 | 4042734.5 | 4070998 | 861336.69 | 3308808.25 | 2938595.5 | 2415084 | 4655039 | 4100491.5 | 3573200.25 |
| Q3ULF4 | Spg7 | Mitochondrial inner membrane m-AAA protease c | 71.91 | 2.56 | 85996 | 2 | 2 | 2 | 34229.25 | 2 | 2 | 2 | 2 | 87784.7 | 197009.62 | 2 | 2 | 53757.48 | 28305.75 | 70845.23 | 2 | 2 | 2 | 2 | 2 | 2 |
| Q9JIA7 | Sphk2 | Sphingosine kinase 2 | 56.32 | 1.78 | 65618 | 1 | 2 | 2 | 65128.19 | 2 | 170671.02 | 568711.69 | 1039425.06 | 529650.56 | 1452776.62 | 601413 | 271837.66 | 392544.69 | 260894.44 | 675179.81 | 413057.09 | 2 | 306869 | 105176.3 | 2 | 2 |
| Q8K1S6 | Spire2 | Protein spire homolog 2 | 38.91 | 1.11 | 80210 | 1 | 608087 | 2 | 2 | 617124.75 | 2 | 2 | 2 | 2 | 2 | 2 | 2 | 705700.25 | 256816.84 | 2 | 2 | 1337545.88 | 2 | 2 | 2 | 2 |
| Q64105 | Spr | Sepiapterin reductase | 314.6 | 26.82 | 27883 | 5 | 1756860.88 | 4826915.5 | 2914771.25 | 1020332.38 | 5380486 | 4306908.5 | 4328696.5 | 4904770 | 3740161.25 | 3970637.5 | 8042656 | 6360110 | 4032527.5 | 4249471.5 | 2 | 4248624 | 175535.39 | 819787.5 | 2399245.5 | 2 |
| Q3TFQ1 | Spryd7 | SPRY domain-containing protein 7 | 37.04 | 5.1 | 21681 | 1 | 2 | 2 | 2 | 2 | 2 | 2 | 2 | 2 | 620371.12 | 2 | 897511.19 | 923243.94 | 747487.88 | 2 | 2 | 2 | 912471.5 | 2 | 2 | 2 |
| Q62261 | Sptbn1 | Spectrin beta chain, non-erythrocytic 1 | 151.45 | 1.82 | 274223 | 3 | 1776399.5 | 3198778.5 | 1722962.12 | 3961297 | 3860236.5 | 1712632.62 | 1626090.12 | 2656560 | 2275858 | 1661269.25 | 2154133.25 | 3211463.75 | 1964291.25 | 2456551 | 2 | 2809568.25 | 1267883.62 | 1500595.62 | 2597931 | 2 |
| P97363 | Sptlc2 | Serine palmitoyltransferase 2 | 53.02 | 3.21 | 62982 | 1 | 2 | 2 | 2 | 2 | 2 | 2 | 215335.08 | 156777.64 | 2 | 151518.81 | 2 | 2 | 2 | 2 | 2 | 2 | 2 | 2 | 2 | 2 |
| Q9R112 | Sqor | Sulfide:quinone oxidoreductase, mitochondrial | 820.09 | 42 | 50282 | 15 | 19509638 | 19066550 | 26872774 | 17902252 | 17786308 | 16576687 | 17994872 | 23049462 | 20474210 | 24643526 | 13989651 | 9522058 | 17076548 | 10679436 | 16951778 | 15351738 | 17389086 | 28320584 | 16780890 | 20864156 |
| Q64337 | Sqstm1 | Sequestosome-1 | 305.58 | 22.62 | 48163 | 6 | 2 | 1353136.38 | 2 | 2 | 2898501.5 | 2598644.5 | 1745524 | 3845600.25 | 3275013.25 | 2580654.25 | 2278712.75 | 2298179.75 | 2086673.12 | 2509225 | 1912027.75 | 2744636.5 | 2665320.5 | 2192270.25 | 3141439 | 2 |
| D4A208 | Srgap2 | SLIT-ROBO Rho GTPase-activating protein 2 | 127.63 | 2.52 | 120878 | 3 | 2733559.5 | 3249441 | 300959.53 | 665654.06 | 1881281 | 2504878.75 | 2105485.25 | 2 | 2238124.5 | 1466275.62 | 1852348 | 1969003.88 | 1517602.38 | 1788551.25 | 1483438.88 | 2194182.75 | 2248705.25 | 2856241.25 | 2149879.25 | 2189539.5 |
| Q6P069 | Sri | Sorcin | 151.35 | 11.62 | 21627 | 3 | 13953588 | 23128646 | 16465425 | 16276046 | 14087524 | 16493951 | 12633662 | 15905589 | 10973786 | 9031470 | 10689832 | 10942434 | 9717686 | 10037392 | 7573880 | 10938448 | 8668127 | 10292616 | 15047740 | 10976065 |
| Q64674 | Srm | Spermidine synthase | 226.76 | 17.22 | 33995 | 4 | 5795107 | 6671999 | 5989491 | 4279145 | 5095767 | 4483482 | 4500024 | 5305569 | 2866326.25 | 2602379.75 | 6210238 | 4012922.5 | 4190058 | 3822285.75 | 3810069.5 | 5327165 | 4046118.25 | 3345587 | 2678923.5 | 2850418.5 |
| Q9D7A6 | Srp19 | Signal recognition particle 19 kDa protein | 38.45 | 8.33 | 16191 | 1 | 2 | 2 | 2 | 2 | 289440.38 | 2 | 2 | 2 | 536586.44 | 309733.25 | 2 | 491367.88 | 748202.38 | 594078.12 | 2 | 2 | 2 | 2 | 2 | 2269178.25 |
| Q8BMA6 | Srp68 | Signal recognition particle subunit SRP68 | 233.02 | 7.36 | 70574 | 4 | 2922054.25 | 2008726.25 | 3156444.5 | 1264743.5 | 615022.81 | 1378025.38 | 781418 | 552089.62 | 1414325.62 | 1834717 | 2088458.62 | 1492189.12 | 3080033.25 | 2705513 | 3279255.5 | 2560013.75 | 1679651.12 | 2568890 | 2140261.25 | 3965783 |
| O70551 | Srpk1 | SRSF protein kinase 1 | 45.33 | 2.16 | 73088 | 1 | 2 | 2 | 2 | 2 | 114211.05 | 2 | 2 | 2 | 110413.32 | 2 | 94980.77 | 2 | 124946.41 | 121024.03 | 166606.69 | 2 | 2 | 2 | 2 | 2 |
| Q9DBG7 | Srpra | Signal recognition particle receptor subunit alpha | 480.48 | 17.3 | 69623 | 9 | 1660162.25 | 753232.06 | 2596801.5 | 1987425.25 | 2964972 | 1323968 | 1851371.38 | 2 | 1001780.94 | 3663901.75 | 3330236.5 | 611218.12 | 1830848.25 | 2552438.25 | 2808913 | 1002619.06 | 2 | 2062577.5 | 194224.28 | 3017602.5 |
| P47758 | Srprb | Signal recognition particle receptor subunit beta | 303.07 | 27.14 | 29579 | 6 | 2 | 2 | 2 | 658349.94 | 676036.94 | 533938.25 | 2 | 931058.38 | 1626036.5 | 1907312.5 | 3268400.75 | 773379 | 2665043.25 | 3583411 | 1684533.62 | 1213347.25 | 2848890.5 | 1224071.25 | 3198947 | 1048925.88 |
| Q52KI8 | Srrm1 | Serine/arginine repetitive matrix protein 1 | 291.52 | 7.19 | 106862 | 4 | 149786.67 | 2 | 167733.22 | 2 | 121300.95 | 370350.66 | 2 | 303522.75 | 2 | 52787.59 | 154417.45 | 2 | 179243.2 | 2 | 2 | 97108.28 | 168012.67 | 97397.55 | 2 | 2 |
| Q8BTI8 | Srrm2 | Serine/arginine repetitive matrix protein 2 | 600.71 | 7.1 | 294840 | 13 | 3911727.5 | 734200.25 | 1002865.62 | 1272737.25 | 1232155.62 | 2016486.38 | 512853.72 | 1673712.75 | 3108491.25 | 2673546.5 | 1725387.12 | 3353251 | 3421615 | 2021872.25 | 1457724 | 3486654.5 | 1847911.12 | 1649529.25 | 2882384.5 | 2549803.5 |
| Q99MR6 | Srrt | Serrate RNA effector molecule homolog | 189.83 | 4.69 | 100452 | 4 | 606827.5 | 726692.38 | 393159.53 | 1191660.25 | 1359905.75 | 2 | 1127229.75 | 172385.58 | 1206897.12 | 1476498.25 | 1176781.88 | 1432236.25 | 989844.62 | 1004270.12 | 2 | 2227919.5 | 1749583.38 | 241190.95 | 2593593.75 | 228488.39 |
| Q6PDM2 | Srsf1 | Serine/arginine-rich splicing factor 1 | 379.45 | 28.63 | 27745 | 8 | 7113532 | 4603593.5 | 7852639.5 | 6933812 | 5263414.5 | 7274896.5 | 6509656.5 | 2929965 | 6543227 | 6368002.5 | 8137249 | 12659371 | 9997577 | 9067511 | 8343594 | 9305208 | 8985922 | 5347154 | 7780919.5 | 19384570 |
| Q9R0U0 | Srsf10 | Serine/arginine-rich splicing factor 10 | 280.07 | 19.47 | 31301 | 5 | 2 | 2 | 826053.94 | 517459.34 | 1683157.12 | 2 | 1704127.75 | 1689385.38 | 1221597 | 2024963.38 | 2 | 2 | 2011844.75 | 477358.34 | 3722128.25 | 2 | 2 | 2 | 2 | 2 |
| Q62093 | Srsf2 | Serine/arginine-rich splicing factor 2 | 194.88 | 17.65 | 25476 | 4 | 1355257.5 | 3860306.5 | 2152554.5 | 2700250 | 9573730 | 4509471 | 3982455 | 6409021 | 5594140 | 4456704.5 | 10386354 | 10697476 | 10291682 | 9767810 | 2748913.5 | 16567955 | 12237522 | 6788755.5 | 15063850 | 7679813.5 |
| P84104 | Srsf3 | Serine/arginine-rich splicing factor 3 | 174.38 | 18.29 | 19330 | 4 | 2697017.25 | 5210512.5 | 3169201.25 | 4226097.5 | 6865514.5 | 9545095 | 11787885 | 12703122 | 10964365 | 6348119.5 | 14239307 | 13984641 | 12153264 | 14070471 | 10311008 | 15509445 | 13494506 | 12329943 | 14724140 | 12374145 |
| Q8VE97 | Srsf4 | Serine/arginine-rich splicing factor 4 | 144.55 | 6.34 | 55979 | 3 | 2 | 2 | 105670 | 2 | 170000.64 | 660920.81 | 2 | 2 | 2 | 2 | 2 | 831860.06 | 787289.31 | 2 | 2 | 1117519.75 | 1158903.62 | 2 | 1222358.62 | 843910.19 |
| O35326 | Srsf5 | Serine/arginine-rich splicing factor 5 | 125.98 | 12.64 | 30891 | 3 | 619255.31 | 548254.5 | 2060893.25 | 1801109 | 3298393.5 | 621729.88 | 516412.44 | 335381.44 | 1693820 | 2153862.5 | 172413.83 | 223181.83 | 2 | 240407.39 | 267358.12 | 2 | 409011.62 | 2 | 1195340.12 | 968392.19 |
| G3V6S8 | Srsf6 | Serine/arginine-rich splicing factor 6 | 210.57 | 11.21 | 39025 | 4 | 5302061.5 | 7472267.5 | 3928843.25 | 6597968.5 | 5789498.5 | 5344790 | 4113453.25 | 5664230 | 5093237 | 3577061.5 | 4175208.25 | 7077858 | 6932576 | 8502267 | 3809532.75 | 10642394 | 9395395 | 5355004.5 | 8569152 | 7704238 |
| Q8BL97 | Srsf7 | Serine/arginine-rich splicing factor 7 | 354.23 | 21.72 | 30818 | 7 | 833677.62 | 1773750 | 1791820.12 | 2612234 | 7085022.5 | 5899477.5 | 2003429.75 | 7935205 | 9005497 | 3014007.5 | 21510646 | 16930490 | 15133388 | 18891270 | 7563547 | 16229724 | 15518916 | 5446773.5 | 17603922 | 10077680 |
| Q9D975 | Srxn1 | Sulfiredoxin-1 | 79.04 | 13.24 | 14149 | 2 | 713716.19 | 1982931.5 | 1814324.62 | 860585.25 | 1568256 | 2509747 | 3147617.5 | 4685241 | 1593850.62 | 1677547.62 | 2346355.75 | 1424765.62 | 558264.62 | 1325061.25 | 301456.78 | 1047395.81 | 2036115.25 | 1475067.25 | 3208798.5 | 1917165.5 |
| Q62280 | Ss18 | Protein SSXT | 45.21 | 2.87 | 45864 | 1 | 898139.31 | 1346007.75 | 1085202.12 | 1014467.69 | 1971914.75 | 1215476.38 | 676902.94 | 207711.33 | 680769.69 | 1136914.38 | 814532.31 | 2 | 679713.12 | 650341.12 | 679850.56 | 2 | 1259934.12 | 941275.75 | 761721.88 | 1464306.38 |
| P32067 | Ssb | Lupus La protein homolog | 308.44 | 12.29 | 47756 | 5 | 47955.12 | 293411.06 | 275794.72 | 184249.03 | 654527.88 | 462011.81 | 957821.56 | 129473.56 | 1108853 | 281666.25 | 354214.59 | 550997.06 | 632012 | 769012.38 | 2095774.5 | 176412.55 | 283954.31 | 1198912.62 | 170901.36 | 1386358.12 |
| Q9CYR0 | Ssbp1 | Single-stranded DNA-binding protein, mitochondr | 182.69 | 28.29 | 17319 | 4 | 3838680 | 10459629 | 2675687 | 7416993 | 2711291 | 968398.5 | 3954957 | 1725719.75 | 6611554.5 | 2582424.5 | 2613522.5 | 7545851 | 4760002 | 6496079 | 9658626 | 5837049 | 9801775 | 6909519 | 3333642 | 3847022 |
| Q08013 | Ssr3 | Translocon-associated protein subunit gamma | 197.4 | 9.73 | 21064 | 2 | 4488552 | 2672782.5 | 4232913 | 2861596 | 3526890.25 | 2371337.75 | 2665232 | 1734542.12 | 3624972 | 4277002 | 1313364.5 | 1021446.44 | 2770101.25 | 2287337 | 2932324 | 1309015 | 1790701.38 | 667552.75 | 1947398.38 | 1667219.88 |
| Q07984 | Ssr4 | Translocon-associated protein subunit delta | 198.54 | 24.86 | 18980 | 3 | 2 | 161350.42 | 68984.69 | 268191.12 | 1025484.75 | 374952.81 | 3634304.75 | 382182.81 | 1541147.25 | 2284500.5 | 1069052.25 | 975028.5 | 1020178.19 | 1075225 | 419809.28 | 378954.84 | 1136756.5 | 628901.5 | 462466.81 | 282169.25 |
| Q04931 | Ssrp1 | FACT complex subunit SSRP1 | 55.65 | 1.69 | 80914 | 1 | 2 | 2 | 858241.81 | 619146.75 | 2 | 2 | 903991.12 | 1221321 | 2 | 511243.75 | 2 | 2 | 308653.19 | 2 | 540845.5 | 2 | 564111.19 | 2 | 2 | 2 |
| Q99L47 | St13 | Hsc70-interacting protein | 309.5 | 15.63 | 41656 | 6 | 9513420 | 5404924.5 | 11355470 | 6867605.5 | 5337390 | 5251625.5 | 4804684.5 | 4888484.5 | 4258696.5 | 6219980 | 6199801.5 | 8629660 | 8555952 | 7345967 | 5172060 | 4188193.5 | 3776593 | 3497335 | 2520118 | 215380.53 |
| Q8R424 | Stambp | STAM-binding protein | 60.51 | 1.65 | 48512 | 1 | 865588.88 | 1223097.62 | 954430.94 | 919080.31 | 967497.81 | 955936.56 | 1034156.12 | 1166924.75 | 582889.5 | 630571.81 | 989312.94 | 552791.25 | 756438.69 | 746163.19 | 2005631.38 | 877894.81 | 592736.25 | 1502846.88 | 1022763.12 | 2136963 |
| Q76N33 | Stambpl1 | AMSH-like protease | 69.44 | 4.59 | 49640 | 1 | 2 | 2 | 2 | 73047.3 | 78517.94 | 2 | 184386.2 | 111515.01 | 84496.65 | 147024.52 | 2 | 2 | 2 | 85599.33 | 136587.03 | 2 | 2 | 2 | 2 | 2 |
| Q9JM90 | Stap1 | Signal-transducing adaptor protein 1 | 327.59 | 24.92 | 34628 | 7 | 148260.69 | 2 | 26142.13 | 1145955.88 | 2656072.25 | 1189623.25 | 988053.5 | 69014.73 | 3713769.5 | 2635952 | 6626667.5 | 5717615.5 | 5421022 | 4879389 | 5134649 | 4244889 | 2497411.25 | 2547818 | 1718836.62 | 2116238.75 |
| P42225 | Stat1 | Signal transducer and activator of transcription 1 | 320.81 | 9.88 | 87197 | 6 | 2 | 455969.56 | 1453364.5 | 1678322 | 1254110 | 898850.38 | 1277740.12 | 192678.2 | 4947940 | 4637011 | 2 | 1715371.62 | 1262843.25 | 962468.38 | 2 | 441520.84 | 2 | 1039440.12 | 2 | 1874678.75 |
| Q9WVL2 | Stat2 | Signal transducer and activator of transcription 2 | 98.6 | 2.17 | 105417 | 2 | 781457.69 | 622287 | 756452.5 | 1258053.25 | 547842.25 | 1101950.62 | 2764309 | 2 | 1203589.62 | 3200031.25 | 2 | 2 | 2 | 2 | 843108.25 | 2 | 2 | 2 | 2 | 491770.81 |
| P42227 | Stat3 | Signal transducer and activator of transcription 3 | 47.14 | 2.34 | 88054 | 1 | 2 | 70824.55 | 87878.54 | 93174.98 | 77254.92 | 2 | 82105.66 | 44864.73 | 300626.25 | 95376.21 | 2 | 2 | 36690.41 | 56175.79 | 2 | 2 | 2 | 58973.91 | 2 | 2 |
| Q5RKL5 | Steap3 | Metalloreductase STEAP3 | 120.4 | 4.92 | 54674 | 2 | 2 | 2 | 2 | 2 | 499463.75 | 2 | 2 | 2417073.25 | 2 | 661823.56 | 2 | 2 | 2 | 284591.03 | 2 | 2 | 2 | 2840437 | 628502.62 | 717135.44 |
| P70302 | Stim1 | Stromal interaction molecule 1 | 36.14 | 2.04 | 77567 | 1 | 33541.41 | 2 | 66852.57 | 2 | 87698.12 | 2 | 2 | 2 | 116400.15 | 103197.49 | 2 | 2 | 2 | 2 | 2 | 2 | 2 | 2 | 2 | 2 |
| F1M391 | Sting1 | Stimulator of interferon genes protein | 40.7 | 2.37 | 42655 | 1 | 2 | 2 | 2 | 2 | 496325.81 | 726729.44 | 495363.28 | 2 | 564327.69 | 481083.28 | 2 | 522873.75 | 2 | 490340.94 | 2 | 2 | 2 | 2 | 2 | 2 |
| Q60864 | Stip1 | Stress-induced-phosphoprotein 1 | 889.59 | 30.39 | 62582 | 18 | 12569840 | 17895654 | 14224676 | 16559270 | 17819090 | 11237760 | 14374830 | 9929908 | 16225873 | 14344894 | 22371744 | 32623344 | 22221482 | 21334350 | 13667905 | 20408486 | 23572866 | 17517056 | 16427634 | 22424688 |
| E9PTG8 | Stk10 | Serine/threonine-protein kinase 10 | 134.53 | 3.93 | 111877 | 3 | 497011.91 | 139250.41 | 2 | 419436.56 | 833639.06 | 112697.41 | 1225894.75 | 242214.62 | 2097024.25 | 639865.25 | 149514.38 | 181775.11 | 2 | 304325.41 | 2 | 2 | 493519.28 | 2 | 2 | 2 |
| B0LT89 | Stk24 | Serine/threonine-protein kinase 24 | 41.25 | 3.94 | 47990 | 1 | 2 | 1285006.5 | 2 | 1008575.5 | 1197370.88 | 1337346 | 987618 | 1051542.38 | 875676.19 | 866495.19 | 939927.31 | 704809.81 | 750532.12 | 2 | 2 | 976957.56 | 2 | 720595 | 668956.62 | 2 |
| O54748 | Stk3 | Serine/threonine-protein kinase 3 | 96.96 | 4.68 | 56122 | 2 | 2 | 2 | 2 | 2 | 872678.12 | 2 | 2 | 854614 | 479792.94 | 729677.56 | 590577.25 | 2 | 2 | 2 | 2 | 2 | 2 | 2 | 2 | 2 |
| Q9JI11 | Stk4 | Serine/threonine-protein kinase 4 | 95.45 | 4.93 | 55541 | 2 | 2 | 125921.51 | 2 | 2 | 2 | 2 | 2 | 2 | 99483.88 | 2 | 2 | 2 | 2 | 114927.02 | 2 | 191202.41 | 2 | 145287.62 | 2 | 2 |
| P54227 | Stmn1 | Stathmin | 243.32 | 25.5 | 17274 | 4 | 1345075 | 1039445.12 | 2 | 1366825.5 | 998115.75 | 2 | 411985.69 | 2 | 782392.44 | 303823.5 | 7008651.5 | 9875368 | 7318129 | 6250941 | 5507581.5 | 6166086 | 4705661.5 | 3624702.5 | 5571385.5 | 5006386 |
| Q6AYD2 | Stn1 | CST complex subunit STN1 | 84.4 | 4.17 | 46783 | 2 | 2 | 672561.5 | 2 | 437047.66 | 3595197.5 | 2 | 549077.44 | 2 | 435880.34 | 2 | 659125.94 | 2 | 428722.94 | 411109.5 | 2 | 2 | 2 | 2 | 2 | 2 |
| P54116 | Stom | Stomatin | 776.15 | 39.44 | 31375 | 12 | 20242630 | 26655502 | 18251922 | 20336976 | 13133069 | 30257302 | 31149056 | 25092226 | 31059586 | 31550270 | 19249702 | 15864378 | 19705082 | 20393460 | 22735194 | 16441495 | 16780034 | 14771437 | 8520973 | 15356280 |
| Q99JB2 | Stoml2 | Stomatin-like protein 2, mitochondrial | 648.78 | 43.91 | 38385 | 12 | 3422600.5 | 1278608.5 | 1523372.62 | 3479683.25 | 2484609.25 | 2027118.38 | 1422915.38 | 1861361.88 | 3613424 | 2357937.5 | 2763539 | 2835641.25 | 3429337.25 | 4439120.5 | 2726408.5 | 2269675 | 2480802.5 | 4130071.75 | 4760967 | 2682759.5 |
| Q5XIG8 | Strap | Serine-threonine kinase receptor-associated prot | 151.4 | 15.14 | 38456 | 3 | 1069741.25 | 2353815.5 | 1317350 | 2120400 | 2515057.25 | 1156853.12 | 841817.38 | 699084.88 | 1414399.38 | 574064.5 | 3399728.5 | 2 | 2588358.5 | 3155579 | 1953442 | 638163.12 | 2462581.25 | 2084777.75 | 2618807 | 1151466.5 |
| Q8VIM6 | Strc | Stereocilin | 40.3 | 0.44 | 196347 | 1 | 2154606 | 3525665.5 | 2 | 1996990.5 | 171278.95 | 2187306.25 | 4929728.5 | 679189.94 | 4668975 | 806825.31 | 4498121 | 8721095 | 5526650 | 4215881 | 2777004.5 | 4511458.5 | 4478902 | 2679500.5 | 176561.31 | 2765789.5 |
| P46978 | Stt3a | Dolichyl-diphosphooligosaccharide--protein glyco | 190.66 | 5.67 | 80598 | 4 | 1740045.25 | 1554315 | 4348640 | 3434757.25 | 1008056.38 | 1568428.12 | 4037981.75 | 2014811.75 | 2764020 | 2398007.5 | 5390157.5 | 4513052.5 | 5798722 | 6521464 | 6090219 | 3426037.5 | 5222915 | 5246175.5 | 3615137 | 4173264 |
| Q3TDQ1 | Stt3b | Dolichyl-diphosphooligosaccharide--protein glyco | 37.15 | 1.09 | 93246 | 1 | 2 | 2 | 2 | 2 | 2 | 2 | 2 | 2 | 2 | 2 | 2076138.62 | 1302558.25 | 1981795.62 | 1268692.25 | 1267586.38 | 2 | 1048808.88 | 2 | 1312271.5 | 2 |
| A6HD62 | Stub1 | E3 ubiquitin-protein ligase CHIP | 87.95 | 7.57 | 34886 | 2 | 928579.75 | 2 | 2 | 701567 | 2082003.5 | 2165575.25 | 2465003.5 | 2472031.5 | 880368.12 | 972992.62 | 1814056.75 | 919844.44 | 1925096.38 | 1222184.5 | 2 | 2 | 863750.38 | 2 | 127751.63 | 2 |
| G3V7P1 | Stx12 | Syntaxin-12 | 86.11 | 11.68 | 31187 | 2 | 2 | 740419.12 | 294883.25 | 340841.47 | 42224.24 | 2 | 307352.34 | 439371.62 | 1392756.5 | 342970.56 | 133308.95 | 530666.75 | 2 | 602546.75 | 1018130.62 | 348796 | 2 | 2 | 2 | 475312.22 |
| P70452 | Stx4 | Syntaxin-4 | 103.5 | 6.71 | 34165 | 2 | 2500431.75 | 1823695.38 | 2729230.5 | 2265172.75 | 2228884.25 | 2009989.25 | 2148183.25 | 4254835.5 | 2094214.62 | 2092548.88 | 5838843 | 3564375.25 | 1089736.38 | 979615 | 1675957.62 | 4330778.5 | 1211835.88 | 2517843.25 | 1444611 | 2542018.5 |
| Q8K1E0 | Stx5 | Syntaxin-5 | 60.36 | 7.04 | 39713 | 1 | 2 | 2 | 2 | 2 | 2 | 2 | 2 | 2 | 2 | 2 | 2 | 2 | 2 | 2 | 2 | 2 | 2 | 2 | 2 | 2 |
| O70439 | Stx7 | Syntaxin-7 | 373.56 | 29.5 | 29821 | 6 | 3905155.5 | 2924744 | 6271025.5 | 2273001.5 | 5719067 | 5223513 | 3312107.5 | 3632767.25 | 4529987 | 4623440.5 | 3735379.5 | 3323118 | 3582894.5 | 1922287.12 | 3949367 | 800612.81 | 2574990.5 | 3585246 | 4373598 | 4784748 |
| O88983 | Stx8 | Syntaxin-8 | 228.09 | 19.07 | 26925 | 4 | 1522677.5 | 319207.38 | 1309632.75 | 469862.19 | 526026.25 | 509244 | 1277725.38 | 297010.81 | 998145.94 | 589675.06 | 2 | 533585.44 | 102655.27 | 866043.94 | 819139.88 | 107881.48 | 1374595 | 317203.56 | 150787.88 | 327636.62 |
| Q64324 | Stxbp2 | Syntaxin-binding protein 2 | 136.14 | 9.78 | 66357 | 3 | 2 | 2 | 118359.12 | 232506.39 | 635084.31 | 2 | 679171 | 203709.16 | 741736.5 | 337839.81 | 127738.26 | 539633.75 | 87991.62 | 106059.29 | 2 | 47061.15 | 2 | 2 | 2 | 2 |
| Q60770 | Stxbp3 | Syntaxin-binding protein 3 | 41.02 | 1.69 | 67942 | 1 | 1176209.62 | 1574669.62 | 1260444 | 2 | 1291387.62 | 1366718.62 | 1434636.25 | 2012085.12 | 1294928.38 | 1421728.88 | 1035146.12 | 2 | 1033748.12 | 1030277.12 | 999171 | 2 | 2 | 2 | 2 | 3839254 |
| Q9Z2I9 | Sucla2 | Succinate--CoA ligase [ADP-forming] subunit bet | 201.96 | 10.8 | 50114 | 5 | 3402934 | 3338923.5 | 3709661.75 | 2673967.75 | 3902184 | 598410.25 | 3084775.75 | 1549907.25 | 6725767 | 5002260 | 3136566.25 | 5784331 | 4171425.5 | 5804383 | 6493630 | 4317742 | 4816782 | 4658299 | 4145888.75 | 4697635 |
| Q9WUM5 | Suclg1 | Succinate--CoA ligase [ADP/GDP-forming] subun | 268.55 | 20.23 | 36155 | 5 | 2153205.75 | 913808.62 | 2 | 215440.64 | 62207.23 | 231678.27 | 588188.56 | 452031.19 | 3963857.75 | 1466597.25 | 4705555.5 | 5208400 | 1198194 | 2327275.5 | 2726630.5 | 2265304.5 | 1073692.38 | 771086.25 | 2 | 1179045.38 |
| Q9Z2I8 | Suclg2 | Succinate--CoA ligase [GDP-forming] subunit bet | 569.31 | 25.4 | 46840 | 10 | 7163420 | 5135851.5 | 10276062 | 4575544.5 | 8032560 | 6142466.5 | 6602538 | 6218233.5 | 7560673 | 8231941 | 6641472 | 5600051.5 | 9377650 | 8693615 | 10535215 | 2325895 | 5018316 | 7137504 | 5586446 | 6670808 |
| B0BN85 | Sugt1 | Protein SGT1 homolog | 70.44 | 3.57 | 38091 | 1 | 3360146.25 | 6291257 | 551511.06 | 6170756.5 | 461183.5 | 6578464.5 | 6413266 | 403669.41 | 4117220.25 | 4552393 | 5479663.5 | 5727676 | 3361440 | 2648124 | 2533754.25 | 7116065.5 | 3442554.5 | 2272389.5 | 5063062.5 | 2413285.25 |
| Q8R0F3 | Sumf1 | Formylglycine-generating enzyme | 71.1 | 4.03 | 40659 | 1 | 2 | 2 | 2 | 1049145.5 | 859623.12 | 2 | 2 | 1174959.62 | 826680.88 | 549532.81 | 2 | 435498.78 | 2 | 2 | 2 | 2 | 2 | 2 | 895240.31 | 764803.38 |
| P61957 | Sumo2 | Small ubiquitin-related modifier 2 | 74.88 | 12.63 | 10871 | 1 | 539168.81 | 589314.38 | 445468.69 | 905354.25 | 1564196.88 | 2 | 629448 | 2 | 1588660.62 | 1053228 | 3907661.25 | 2934246.75 | 1175815.75 | 676922 | 903309.31 | 2 | 2 | 740781.81 | 1737242.62 | 1011631.88 |
| Q9D666 | Sun1 | SUN domain-containing protein 1 | 80.97 | 2.41 | 101976 | 2 | 2658598.5 | 2 | 2 | 1197663.38 | 2 | 750022.69 | 888482.38 | 1064048.62 | 908316.75 | 2001194.12 | 1176547.12 | 2 | 2744923 | 2 | 1131577.75 | 751058.62 | 1514500 | 2 | 2 | 2514671 |
| Q8BJS4 | Sun2 | SUN domain-containing protein 2 | 301.18 | 9.44 | 81605 | 6 | 1270364.5 | 1729638.5 | 2022400.62 | 1784884.38 | 420095.06 | 2 | 2 | 2 | 437306.12 | 2 | 2 | 2 | 148818.41 | 180637.84 | 1551082.5 | 2134424 | 136555.66 | 2 | 2 | 2688659.5 |
| Q920B9 | Supt16h | FACT complex subunit SPT16 | 230.13 | 6.21 | 119825 | 5 | 991666.56 | 766051.31 | 996601.88 | 1894489.38 | 1650562.62 | 1975059.5 | 686785.31 | 1323460.25 | 1488079.5 | 1649183.12 | 2634770 | 1501879.5 | 1762023.75 | 2267510.25 | 2 | 1867357.25 | 1347270.25 | 1218752.62 | 1067529 | 1393607.88 |
| O55201 | Supt5h | Transcription elongation factor SPT5 | 146.03 | 4.44 | 120664 | 3 | 1728923.75 | 1680876.25 | 3477661.25 | 1856451.38 | 1549461 | 1051832.12 | 902829.62 | 1383732.75 | 1507011.62 | 1481699.25 | 1022213.88 | 778070 | 951228.75 | 926730.62 | 1217765.75 | 1436754.38 | 1034174.06 | 1261830.25 | 3883472 | 2392161 |
| Q64310 | Surf4 | Surfeit locus protein 4 | 69.45 | 6.69 | 30381 | 2 | 2 | 595095.19 | 2 | 2186673 | 1684678.25 | 7746673 | 7450136 | 8274980.5 | 7987254 | 3189755 | 1445613.5 | 1173954.62 | 898566.44 | 1287410.5 | 2 | 790746 | 1478189.62 | 2 | 2278103 | 2 |
| P70279 | Surf6 | Surfeit locus protein 6 | 42.7 | 2.82 | 41235 | 1 | 2 | 2 | 2 | 2 | 2 | 2 | 2 | 2 | 2 | 2 | 2 | 52774.89 | 46955.35 | 77252.16 | 2 | 2 | 203078.83 | 109042.1 | 2 | 2 |
| Q6A028 | Swap70 | Switch-associated protein 70 | 87.74 | 3.42 | 68996 | 2 | 2 | 2 | 2 | 2 | 2 | 2 | 2 | 2 | 2 | 50706.56 | 1998248.38 | 1715174.38 | 761978.31 | 2 | 2 | 1768083.25 | 2 | 2 | 2 | 2 |
| Q80X82 | Sympk | Symplekin | 63.78 | 0.85 | 142620 | 1 | 623303.94 | 106012.33 | 583397.56 | 243556.97 | 283120.88 | 132277.42 | 306446.69 | 101432 | 259398.22 | 392781.34 | 2 | 2 | 342735.88 | 564981.06 | 533340.31 | 2 | 2 | 220562.94 | 2 | 2 |
| Q7TMK9 | Syncrip | Heterogeneous nuclear ribonucleoprotein Q | 300.62 | 10.11 | 69633 | 6 | 6656376 | 5594501 | 13198741 | 4490648.5 | 9681693 | 6710556 | 7579261 | 7656460.5 | 6251302.5 | 7621582 | 9518681 | 10104192 | 9791226 | 7006836 | 5230614 | 5185066.5 | 2811431 | 4351424.5 | 4501514 | 2662731.5 |
| O55100 | Syngr1 | Synaptogyrin-1 | 53.26 | 5.13 | 25653 | 1 | 1514054.75 | 1015491.69 | 1007044.69 | 2 | 603652.75 | 1110893.62 | 1114812.12 | 1254791.88 | 652235.81 | 768973.81 | 509426.12 | 2 | 683634.75 | 391213.25 | 2 | 916075.5 | 1443742.62 | 1111210.88 | 110118.95 | 2 |
| O55101 | Syngr2 | Synaptogyrin-2 | 42.51 | 4.46 | 24778 | 1 | 883481.44 | 879555 | 1439341.38 | 1155733.38 | 943681.06 | 2 | 650710.81 | 859400 | 409207.38 | 1110826.88 | 2 | 2 | 2 | 586227.81 | 798507.56 | 2 | 2 | 1309337.38 | 2 | 2 |
| Q62910 | Synj1 | Synaptojanin-1 | 42.91 | 0.57 | 172881 | 1 | 387074.12 | 1415622 | 2 | 695008.31 | 1055083.62 | 2 | 981188.88 | 1150967.62 | 890687.81 | 1079982.38 | 563080.38 | 2 | 477919.25 | 2 | 2 | 2 | 2 | 2 | 2 | 2 |
| O09117 | Sypl1 | Synaptophysin-like protein 1 | 115.01 | 4.21 | 28899 | 2 | 7386843 | 6845195 | 4858624 | 7341161.5 | 4143606.5 | 5914780.5 | 5739564.5 | 125054.38 | 5792159 | 4390265.5 | 3287204 | 3997914 | 5997009.5 | 4554170 | 1709598.38 | 3993807.5 | 5214108 | 6059071 | 4321055 | 7404446 |
| Q9WVA4 | Tagln2 | Transgelin-2 | 1318.76 | 87.44 | 22395 | 20 | 141621986 | 143699874 | 141449090 | 147967874 | 186832498 | 176886002 | 173149778 | 189656994 | 155579330 | 138303394 | 242942866 | 226146626 | 145621858 | 147718978 | 167594562 | 145608034 | 121850066 | 105073586 | 152208770 | 103041842 |
| Q93092 | Taldo1 | Transaldolase | 1134.67 | 54.6 | 37387 | 22 | 41806122 | 69413954 | 52469954 | 56295170 | 60955842 | 86529962 | 78396034 | 79165274 | 56023814 | 18884890 | 86067754 | 79151482 | 58764654 | 58735058 | 72941602 | 84906418 | 83845466 | 73489866 | 89337258 | 67155954 |
| Q53UA7 | Taok3 | Serine/threonine-protein kinase TAO3 | 74.16 | 2.34 | 105474 | 2 | 274142.59 | 2 | 2 | 2 | 1396730.12 | 2 | 2 | 2 | 387301.5 | 816836.94 | 1193553.62 | 2 | 471904.97 | 289498.03 | 2 | 600975.12 | 2 | 2 | 2 | 2 |

| | | | | | | | | | | | | | | | | | | | | | | | | | | |
|---|---|---|---|---|---|---|---|---|---|---|---|---|---|---|---|---|---|---|---|---|---|---|---|---|---|---|
| P21958 | Tap1 | Antigen peptide transporter 1 | 298.66 | 10.08 | 78864 | 5 | 2 | 1090104 | 1342827 | 881340.62 | 2609586.5 | 1399870.25 | 1842805.75 | 2077141.88 | 5910725 | 5567362 | 2 | 2 | 2 | 528483.88 | 327171.12 | 235849.38 | 366826.28 | 1156528 | 496575.03 | 369877.5 |
| P36371 | Tap2 | Antigen peptide transporter 2 | 264.33 | 8.69 | 77445 | 6 | 2360856 | 3662836 | 4193169.75 | 4141515 | 4453851 | 7867523.5 | 7347698 | 10021318 | 8624597 | 9529242 | 889732.62 | 2817701 | 4643617 | 2377934 | 1997989.75 | 954741.25 | 3569631.5 | 1608510.62 | 4439558 | 846160 |
| Q9R233 | Tapbp | Tapasin | 288.71 | 17.42 | 49736 | 6 | 5283079 | 7493800.5 | 75865.82 | 11895044 | 8545807 | 9355808 | 10704515 | 10073298 | 19103320 | 18822134 | 6208656.5 | 7177058.5 | 6474315 | 5831997 | 4352196.5 | 7670488.5 | 5873767 | 7466315 | 6576177.5 | 7576095 |
| Q921F2 | Tardbp | TAR DNA-binding protein 43 | 413.56 | 17.39 | 44548 | 6 | 5051298 | 7201042 | 917203.19 | 8291341.5 | 2688956.5 | 6548520 | 6400967 | 2100354.5 | 7754439 | 5727020.5 | 14065834 | 13013706 | 9388418 | 11387324 | 6781083.5 | 12202236 | 6333363 | 9356212 | 9114005 | 8550262 |
| Q9D0R2 | Tars1 | Threonine--tRNA ligase 1, cytoplasmic | 624.32 | 18.98 | 83356 | 13 | 5774899 | 3711608.5 | 6592564 | 7710436.5 | 9044218 | 2256042.5 | 5431057.5 | 6766872 | 4814316 | 9688358 | 5768144 | 7771356 | 8290885 | 8911304 | 11270032 | 8859193 | 6043169 | 7428632 | 5144101 | 3402348.5 |
| Q3UQ84 | Tars2 | Threonine--tRNA ligase, mitochondrial | 58.7 | 1.38 | 81700 | 1 | 2 | 2 | 552075.62 | 2 | 283924.06 | 2 | 53626.04 | 2 | 2 | 487972.97 | 2 | 2 | 2 | 2 | 2 | 2 | 2 | 2 | 2 | 2 |
| P48428 | Tbca | Tubulin-specific chaperone A | 63.77 | 8.33 | 12758 | 1 | 3631210.25 | 5323165.5 | 3098622.75 | 3899259.25 | 4044558.5 | 3354873.5 | 2392969.25 | 4248079.5 | 2245423.75 | 2687674.5 | 2959594.75 | 2828885.25 | 3176474.25 | 3051736 | 2317306 | 4241948.5 | 3960428 | 3377049 | 4706671.5 | 3695553.25 |
| Q8VCN9 | Tbcc | Tubulin-specific chaperone C | 33.53 | 2.35 | 38125 | 1 | 2378876.75 | 3045797.5 | 1790651.25 | 2820224.75 | 1977941.62 | 2164291 | 1697221.12 | 2402291.75 | 268861.38 | 1211373.25 | 2 | 1478138.38 | 1294026.62 | 1655930.25 | 416088.31 | 2069576.75 | 1360979.75 | 1970274.62 | 2048546.5 | 2231103.75 |
| Q8BYA0 | Tbcd | Tubulin-specific chaperone D | 97.37 | 2.34 | 133321 | 2 | 2 | 899361.5 | 143814.81 | 1159103.75 | 1414475.38 | 2 | 1142233.75 | 2 | 1206191.38 | 127380.15 | 1377238.38 | 1199885.75 | 1078673 | 1378262.25 | 1038822.25 | 2 | 1776679.62 | 928957.06 | 1881545.75 | 1141142.5 |
| Q8C4J7 | Tbl3 | Transducin beta-like protein 3 | 45.25 | 1.62 | 88266 | 1 | 2 | 2 | 118165.28 | 364050.75 | 32864 | 613754.06 | 642745.81 | 134801 | 256418.73 | 392625.78 | 86933.51 | 295321.16 | 547741.31 | 445769.5 | 889565.94 | 2 | 792860.5 | 218055.25 | 2 | 2 |
| Q5M9G9 | Tbrg4 | FAST kinase domain-containing protein 4 | 53.38 | 3.02 | 71181 | 1 | 68865.34 | 464613.12 | 242373.5 | 262112.73 | 75766.34 | 166192.47 | 232357.47 | 67377.05 | 302433.84 | 349576.78 | 114121.66 | 188090.81 | 245323.5 | 289955.06 | 509097.16 | 2 | 415144.19 | 460611.69 | 2 | 306692.53 |
| Q8CGF7 | Tcerg1 | Transcription elongation regulator 1 | 37.31 | 0.73 | 123788 | 1 | 2 | 1099901 | 2 | 1486338.75 | 1645102.25 | 1168682.88 | 836364 | 2 | 1289707 | 1103694.25 | 1683044.25 | 2085849.75 | 983937.19 | 1266612.75 | 843302.56 | 1991968.5 | 2 | 2 | 2 | 1265034.75 |
| O08784 | Tcof1 | Treacle protein | 756.84 | 15 | 135001 | 15 | 16480656 | 19170086 | 8270126 | 14607075 | 12759118 | 12404963 | 12418044 | 9043475 | 19336634 | 7501395 | 9628120 | 11557820 | 7903041 | 9769896 | 8832507 | 11072328 | 5987784 | 8260422 | 4897791.5 | 11148070 |
| P11983 | Tcp1 | T-complex protein 1 subunit alpha | 1427.29 | 53.06 | 60449 | 23 | 15415812 | 26174756 | 24532970 | 40156730 | 52737758 | 42726826 | 30738508 | 46640198 | 44295294 | 32282780 | 32214576 | 32999280 | 26264830 | 30849506 | 29986606 | 28246400 | 22660342 | 25518208 | 30205970 | 20354398 |
| Q64232 | Tecr | Very-long-chain enoyl-CoA reductase | 82.43 | 5.52 | 36123 | 2 | 2 | 2 | 2 | 2 | 2 | 2 | 2 | 2 | 2 | 2 | 2 | 2 | 2 | 2 | 2 | 2 | 2 | 2 | 2 | 2 |
| P47226 | Tes | Testin | 165.59 | 6.15 | 47983 | 3 | 84927.28 | 558819.69 | 2 | 716605.19 | 105603.77 | 847497.62 | 1150454 | 534595 | 1662910.5 | 228829.34 | 1305567.62 | 1510882.75 | 354163.94 | 1037682.94 | 2 | 1807303.88 | 977708.94 | 2 | 2 | 2 |
| Q62351 | Tfrc | Transferrin receptor protein 1 | 480.28 | 12.32 | 85731 | 10 | 4758922 | 3893226 | 8569880 | 3927010.75 | 2413348 | 1272217.12 | 1225314.75 | 4436116.5 | 2246856 | 3604006.25 | 3188943.25 | 2239498.5 | 3882392.75 | 3601248.5 | 4643843.5 | 2808218 | 2318468 | 3907658.5 | 1736293.5 | 786982.44 |
| B1AZI6 | Thoc2 | THO complex subunit 2 | 112.92 | 1.63 | 182773 | 2 | 131701.86 | 146543.44 | 195795.09 | 580313.19 | 429039.38 | 405208.12 | 318888.03 | 39212.32 | 235132.12 | 223981.7 | 73587.22 | 167093.09 | 331119.28 | 337739.09 | 744142.69 | 88975.02 | 100362.19 | 413716.31 | 2 | 448204.5 |
| Q569Z6 | Thrap3 | Thyroid hormone receptor-associated protein 3 | 307.49 | 8.1 | 108178 | 7 | 1289421.5 | 477740.97 | 1670642.12 | 1219053.25 | 1697588.88 | 1540137.75 | 727397.56 | 3491123 | 1962608.25 | 704525.5 | 4812098 | 5544273 | 2563593.25 | 3195089.5 | 1149120 | 2935295.25 | 2277063.5 | 2 | 2231275.25 | 2 |
| Q99J36 | Thumpd1 | THUMP domain-containing protein 1 | 190.12 | 8.57 | 38885 | 3 | 707520.06 | 1900229.12 | 130695.18 | 671655 | 183096.69 | 2021951 | 1847228.88 | 381568.16 | 669157.75 | 794059.06 | 2 | 1172662.75 | 626533.69 | 1233737 | 2 | 565476.81 | 2 | 2 | 787411.19 | 2 |
| P70318 | Tial1 | Nucleolysin TIAR | 107.13 | 6.12 | 43389 | 2 | 2453252 | 445260.62 | 1925120.5 | 2 | 1252623.12 | 2 | 2 | 2 | 2 | 971323.5 | 353129.59 | 1903238.75 | 2036867.12 | 1437889.62 | 1090809.25 | 300700.06 | 1717779 | 2 | 393668.53 | 710953.94 |
| Q80UF7 | Ticam1 | TIR domain-containing adapter molecule 1 | 39.35 | 1.5 | 79230 | 1 | 2 | 2 | 2 | 1657331.75 | 200381.58 | 2 | 2 | 1423614 | 2 | 2 | 2 | 2 | 2 | 2 | 4041150.75 | 2 | 2 | 2 | 2 | 1176554.12 |
| Q8BZA9 | Tigar | Fructose-2,6-bisphosphatase TIGAR | 133.06 | 15.61 | 29191 | 3 | 2 | 105865.96 | 451414.94 | 407488.94 | 273214.72 | 490453.16 | 791137 | 164171.34 | 303104.28 | 803357 | 94460.8 | 215487.69 | 342318.12 | 421701.56 | 1462072.75 | 2 | 987911.88 | 385047.06 | 77744.23 | 835268.88 |
| P62075 | Timm13 | Mitochondrial import inner membrane translocase | 90.99 | 14.74 | 10458 | 1 | 2951467 | 1980725.62 | 2606312 | 1496193 | 1611425.75 | 1285734.62 | 1170344.25 | 1657627 | 846881.69 | 1278633.38 | 1451130.12 | 802565.5 | 1413385.75 | 2 | 1437448.12 | 2 | 2 | 1006918.06 | 2 | 2 |
| O35092 | Timm17a | Mitochondrial import inner membrane translocase | 169.14 | 29.82 | 18038 | 2 | 193351.67 | 399425.12 | 312579.38 | 843746.44 | 327024.34 | 2 | 367968.91 | 536233.56 | 444026.41 | 125637.69 | 135151.38 | 316829.31 | 350742.44 | 554527.44 | 751980.06 | 2 | 2 | 527245.44 | 201277.22 | 392459.75 |
| O35093 | Timm23 | Mitochondrial import inner membrane translocase | 104.3 | 12.92 | 21907 | 2 | 1098240.25 | 2129503 | 1220345.75 | 2297182.75 | 4049270.5 | 2254189 | 1972933.75 | 2824388 | 2455170 | 1661261.25 | 786754.19 | 811343.31 | 1025088.44 | 929792.19 | 352450 | 290347.56 | 2 | 2 | 1095014.38 | 2 |
| O35857 | Timm44 | Mitochondrial import inner membrane translocase | 246.36 | 13.72 | 51091 | 5 | 2131188.5 | 4535974 | 4234552 | 1434525.38 | 2174066.75 | 2836777 | 3106062.75 | 2712767.5 | 2953289.5 | 3118595 | 3242332.25 | 2137304.25 | 3254898.5 | 3359433.25 | 1411272.88 | 3798895 | 2160149.5 | 3828644.5 | 1483876.12 | 1672189.25 |
| Q9D880 | Timm50 | Mitochondrial import inner membrane translocase | 250.77 | 13.6 | 39776 | 5 | 6805235.5 | 7224476 | 852786.75 | 9653416 | 6684182 | 4060657.25 | 3852179 | 6133871 | 8746345 | 4073659 | 7742663 | 3897772 | 7031143.5 | 7977511 | 6795627 | 6779091.5 | 3120233.75 | 8537328 | 9083222 | 4135726.75 |
| Q9WVA1 | Timm8a | Mitochondrial import inner membrane translocase | 106.82 | 22.68 | 11042 | 2 | 956598.81 | 348236 | 1106029.25 | 796088.94 | 342693.19 | 557240 | 1035009 | 98855.83 | 479343.59 | 734625.88 | 1043738.81 | 2 | 2 | 870476.75 | 1509763 | 587479 | 962939.69 | 1052301.88 | 2 | 996833.38 |
| A2VCX1 | Tiprl | TIP41-like protein | 53.98 | 6.64 | 31225 | 1 | 604306.44 | 999283.69 | 582837.5 | 427136.66 | 481055.47 | 491547.62 | 2 | 2 | 307539.94 | 362953.84 | 2 | 2 | 2 | 2 | 2 | 651163.94 | 2 | 2 | 218316.06 | 2 |
| Q9Z0U1 | Tjp2 | Tight junction protein 2 | 332.25 | 7.8 | 131280 | 8 | 535085.88 | 360806.28 | 1547062.38 | 475967.38 | 2796204.75 | 2 | 751093.81 | 785960.5 | 1416962.25 | 2446744.5 | 2 | 1692839 | 1170284.25 | 2 | 394791.16 | 664948.5 | 593523.62 | 2 | 1067661 | 2 |
| P40142 | Tkt | Transketolase | 1674.24 | 49.44 | 67630 | 29 | 39039022 | 78418546 | 33548210 | 33800890 | 89102226 | 84618066 | 88700098 | 55029882 | 110023210 | 53448130 | 163516018 | 182412226 | 126912786 | 138631762 | 77841306 | 187722562 | 143701730 | 129989162 | 151493458 | 135058242 |
| P26039 | Tln1 | Talin-1 | 5070.81 | 40.5 | 269821 | 74 | 107585194 | 159217490 | 98542434 | 161239922 | 160819090 | 131735482 | 126524058 | 116123034 | 159148354 | 143289266 | 111057178 | 127417266 | 108229122 | 118907618 | 115067346 | 117108770 | 124587874 | 133620090 | 127719642 | 128240578 |
| Q6R5N8 | Tlr13 | Toll-like receptor 13 | 101.88 | 4.04 | 114443 | 2 | 2 | 2 | 2 | 2 | 2 | 2 | 2 | 2 | 725805 | 2 | 2 | 2 | 2 | 2 | 2 | 2 | 2 | 2 | 2 | 2 |
| P58681 | Tlr7 | Toll-like receptor 7 | 141.65 | 2.86 | 121837 | 3 | 428290.41 | 2 | 507083.31 | 118351.95 | 33474.23 | 2 | 2 | 2 | 486658.97 | 89491.43 | 2 | 2 | 2 | 117557.74 | 210558.42 | 2 | 2 | 2 | 88617.11 | 2 |
| Q9EQU3 | Tlr9 | Toll-like receptor 9 | 45.18 | 0.87 | 116412 | 1 | 2 | 2 | 2 | 2000980 | 812837.44 | 2 | 583953.12 | 2 | 840786.75 | 541271.38 | 2 | 2 | 2 | 646586.38 | 2 | 1102948.88 | 2 | 1056642.88 | 2 | 778448.06 |
| Q566R2 | Tm2d2 | TM2 domain-containing protein 2 | 38.25 | 5.16 | 22929 | 1 | 785479.69 | 837039.75 | 2 | 998084.88 | 770260.38 | 2 | 665276.88 | 631931.38 | 425573.31 | 2 | 2 | 441790.84 | 583935.75 | 465335.88 | 2 | 2 | 896088.12 | 955161.31 | 2 | 2 |
| P58021 | Tm9sf2 | Transmembrane 9 superfamily member 2 | 39.36 | 2.11 | 75330 | 1 | 2 | 871651.38 | 2 | 935297.62 | 1545780.88 | 1184573.75 | 1506071.62 | 1473577 | 1292848.12 | 961278.19 | 1160930.5 | 1057781.5 | 1158791.25 | 842157.56 | 2 | 1638841.38 | 2 | 2 | 868894.75 | 2 |
| Q9ET30 | Tm9sf3 | Transmembrane 9 superfamily member 3 | 44.7 | 1.87 | 67545 | 1 | 635081 | 145427.31 | 1177670 | 409320.19 | 975139.62 | 2 | 168536.59 | 194576.84 | 341331.16 | 2020690.12 | 2 | 2 | 542498.69 | 2 | 845397.94 | 2 | 157526.56 | 2 | 2 | 2 |
| Q8BH24 | Tm9sf4 | Transmembrane 9 superfamily member 4 | 131.1 | 6.22 | 74693 | 3 | 470460.19 | 2 | 2 | 493679.72 | 100022.19 | 1159737.62 | 1219491.88 | 2 | 621650.75 | 152427.98 | 122035.56 | 648629 | 1183559.12 | 1655342.25 | 2 | 1875003.12 | 1409942 | 2 | 768214.62 | 2 |
| Q5I0H4 | Tmco1 | Calcium load-activated calcium channel | 209.16 | 19.68 | 21175 | 3 | 128589.49 | 2 | 198485.47 | 324438.53 | 141879.17 | 123027.49 | 455134.09 | 1157054.38 | 2294833.5 | 545733.12 | 2261938.75 | 2089909.75 | 1874141.5 | 2501004 | 387632.5 | 1678755 | 1320605.5 | 1044052.25 | 2524544.5 | 1493966.25 |
| Q63584 | Tmed10 | Transmembrane emp24 domain-containing protei | 263.1 | 21.46 | 24858 | 5 | 9823609 | 5335008 | 11301661 | 6110957.5 | 6417275 | 5013357.5 | 6645209.5 | 6387529 | 6052198 | 11142419 | 5004882 | 4122389.5 | 6596607 | 4655075 | 9928546 | 6340163.5 | 7834929 | 11097992 | 5049814 | 7303929.5 |
| Q63524 | Tmed2 | Transmembrane emp24 domain-containing protei | 77.89 | 6.47 | 22733 | 1 | 2 | 2 | 2 | 2 | 2 | 2 | 168006.38 | 151387.86 | 2 | 2 | 250334.98 | 234781.3 | 828570.12 | 298280.31 | 148730.34 | 301292.34 | 260240.67 | 169582.27 | 112005.8 | 155691.83 |
| Q8R1V4 | Tmed4 | Transmembrane emp24 domain-containing protei | 155.42 | 19.82 | 26022 | 4 | 2 | 805752 | 373500.59 | 2 | 3198042.5 | 302291.31 | 1064130.88 | 2182379 | 1994282.12 | 3167605.75 | 1797697.88 | 1738998 | 1999447.25 | 2 | 2 | 285597.5 | 632560.25 | 2 | 2 | 2 |
| Q6AXN3 | Tmed5 | Transmembrane emp24 domain-containing protei | 53.09 | 3.93 | 26136 | 1 | 204427.67 | 2 | 613074.56 | 2 | 842096.5 | 1017520.94 | 714980.88 | 2 | 863124.94 | 894830.12 | 524157.53 | 886835.19 | 675240.75 | 2 | 812550.88 | 1190453 | 2 | 607026.44 | 1284814.25 | 712674.19 |
| D3ZTX0 | Tmed7 | Transmembrane emp24 domain-containing protei | 105.28 | 9.29 | 25479 | 2 | 823601.69 | 2 | 643150.62 | 116711.83 | 28005.08 | 2 | 130152.55 | 2 | 2 | 555958.56 | 61562.72 | 2 | 91300.98 | 117314.84 | 923427.38 | 2 | 2 | 2 | 2 | 2 |
| Q5I0E7 | Tmed9 | Transmembrane emp24 domain-containing protei | 176.46 | 20 | 27028 | 4 | 3573692.75 | 2954117 | 2489207.25 | 4104578.5 | 2578163.75 | 1081017.88 | 2727770.25 | 2694144.5 | 2623489.75 | 2644813.5 | 1290146.75 | 1871314.88 | 1939953.5 | 1746271.12 | 2137977.75 | 1431636.88 | 1862592.5 | 2240060 | 541646.69 | 1244666.38 |
| Q8VC04 | Tmem106a | Transmembrane protein 106A | 175.84 | 14.56 | 29109 | 3 | 1776360.62 | 3250930.5 | 1872025.38 | 3143917 | 2565823.5 | 5899263.5 | 5327733 | 4681551 | 7722462.5 | 7166946 | 554090.06 | 1907583.12 | 1362496.5 | 2428216 | 1583202.88 | 1578631.75 | 2515020.5 | 1758546.5 | 1001731.62 | 756134.56 |
| Q6AYA5 | Tmem106b | Transmembrane protein 106B | 48.09 | 5.82 | 31152 | 1 | 2 | 2 | 2 | 2 | 2 | 2 | 2 | 2 | 2 | 2 | 2 | 2 | 2 | 2 | 2 | 2 | 2 | 2 | 2 | 2 |
| Q3UBX0 | Tmem109 | Voltage-gated monoatomic cation channel TMEM | 196.45 | 13.99 | 26306 | 3 | 2016003.12 | 1639677.75 | 2025053.12 | 2142867.5 | 1607607.88 | 2 | 2 | 1453979.75 | 1961638.5 | 678307.69 | 2 | 836161.94 | 1694435.12 | 2136904 | 1611611.62 | 491747.06 | 811074.81 | 2915747.5 | 801176.44 | 1993885.88 |
| Q9CQN6 | Tmem14c | Transmembrane protein 14C | 161.88 | 29.82 | 11642 | 3 | 1390201.25 | 749258.12 | 1007463.25 | 1182442.25 | 1032098.12 | 111271.62 | 1052268.12 | 1171887.25 | 1153491.75 | 866303.25 | 879663.69 | 175345.89 | 1009948.06 | 606939 | 1335216.12 | 2079246.62 | 2448663.75 | 3910358 | 1326393.25 | 2684085.5 |
| Q5PQM0 | Tmem168 | Transmembrane protein 168 | 83.12 | 4.02 | 79772 | 2 | 849051.38 | 898014.19 | 699093.56 | 912721.88 | 845030.5 | 717084.75 | 622145.94 | 71431.26 | 981628.12 | 813195.25 | 2 | 2 | 62702.82 | 592007.62 | 218857.95 | 2 | 159603.06 | 2 | 2 | 108691.73 |
| Q91XE8 | Tmem205 | Transmembrane protein 205 | 36.58 | 4.76 | 21180 | 1 | 2 | 2 | 2 | 2 | 2 | 938989.12 | 2 | 2 | 975183.62 | 2 | 2 | 754535.44 | 2 | 928668.19 | 2 | 2 | 2 | 960388.25 | 324232.34 | 981606.69 |
| Q8BM55 | Tmem214 | Transmembrane protein 214 | 228.73 | 9.46 | 76430 | 5 | 1180080.38 | 1178703.5 | 961996.56 | 1850109 | 955033.38 | 942145.5 | 1810903.25 | 2 | 715090.75 | 780006.94 | 2 | 121616.54 | 98445.48 | 1103034.62 | 246000.84 | 2 | 208778.92 | 1235214.25 | 2 | 989848.62 |
| P61166 | Tmem258 | Dolichyl-diphosphooligosaccharide--protein glyco | 45.94 | 10.13 | 9079 | 1 | 1769329.75 | 2584479.25 | 2434783 | 2 | 2079371.75 | 1729328.38 | 1368820.25 | 2810771.25 | 1587392.5 | 2061545.25 | 2066926.62 | 2 | 2 | 2562437.5 | 1552600.75 | 1125109.38 | 2 | 2 | 1783921.38 | 3110267.5 |
| Q9CR67 | Tmem33 | Transmembrane protein 33 | 159.4 | 13.36 | 28031 | 3 | 1440172.88 | 673541.94 | 2317869.25 | 1989908.62 | 2560531 | 1582712.25 | 21186962 | 9599845 | 2980248.25 | 2321192.5 | 7211938 | 8409346 | 12201652 | 5366078.5 | 47439698 | 1816937.5 | 15806790 | 5543150 | 2874665 | 12352848 |
| Q9DBS1 | Tmem43 | Transmembrane protein 43 | 214.54 | 14.75 | 44783 | 5 | 924546.81 | 1897611.25 | 837690.38 | 2176375.25 | 3262710 | 2542349.25 | 1312948.25 | 2363838.25 | 3041219 | 5343187 | 2525928.25 | 4333415.5 | 2534167 | 2204699.25 | 2 | 2397508.75 | 1026723.88 | 2259805.5 | 3197377.75 | 5131826.5 |
| Q9JHJ0 | Tmod3 | Tropomodulin-3 | 376.99 | 24.43 | 39503 | 7 | 3208762.75 | 2549301.5 | 3611562.75 | 2048469.75 | 8252077 | 3939786 | 3685842.5 | 4424124 | 6150302.5 | 5461559.5 | 3401916.75 | 2322945.75 | 4133268 | 4872141.5 | 4386238 | 1881879.5 | 3714425.25 | 6148562 | 4339342 | 4846200 |
| Q61033 | Tmpo | Lamina-associated polypeptide 2, isoforms alpha/ | 189.41 | 10.25 | 75168 | 4 | 58229.92 | 201304.25 | 845625.5 | 238661.95 | 909229.38 | 2 | 153800.7 | 57599 | 141460.41 | 49120.76 | 1184644.38 | 102623.46 | 103963.8 | 167869.11 | 807719.25 | 2 | 171589.31 | 2 | 2 | 2 |
| Q61029 | Tmpo | Lamina-associated polypeptide 2, isoforms beta/c | 240.17 | 13.27 | 50373 | 5 | 1365118.62 | 2 | 1625115.12 | 2 | 4636271 | 3238528.5 | 2 | 2 | 3550262.5 | 2 | 3728865.5 | 3237185.5 | 3968876.75 | 4859279.5 | 3007842.5 | 5724932 | 5267971 | 3628334.5 | 6603931.5 | 5742324.5 |
| P20065 | Tmsb4x | Thymosin beta-4 | 99.39 | 38 | 5679 | 2 | 1343639.12 | 1311279.12 | 61893.93 | 989030.12 | 2 | 1055342.25 | 2 | 2 | 542606.94 | 2 | 4041235.25 | 3129924 | 758167.44 | 1098576 | 2496598 | 3183646 | 2 | 1034243.5 | 2467688.5 | 2 |
| Q8VBT0 | Tmx1 | Thioredoxin-related transmembrane protein 1 | 168.15 | 8.27 | 31396 | 3 | 6891471.5 | 6565111 | 6813316 | 5128746 | 4968738 | 3396650.75 | 3988950.75 | 5324447.5 | 5111719.5 | 5736903.5 | 5483595 | 4098067.5 | 8077190 | 8274120 | 4337237 | 8798348 | 9103413 | 9365005 | 11818554 | 11324616 |
| Q8BXZ1 | Tmx3 | Protein disulfide-isomerase TMX3 | 158.26 | 8.77 | 51848 | 4 | 1551647.88 | 989534.62 | 658507.56 | 1091559.62 | 734674.31 | 2 | 576286.88 | 1108651.5 | 2 | 721465 | 2 | 2 | 929329.88 | 2 | 2 | 2 | 2 | 1215249.12 | 2 | 2 |
| Q8BFY9 | Tnpo1 | Transportin-1 | 297.62 | 5.01 | 102357 | 5 | 11038026 | 8185250.5 | 11575319 | 7989010 | 9471328 | 5731841 | 5967353 | 7195839 | 6718364 | 5640400 | 3683115.25 | 4458412.5 | 5869673 | 4904413.5 | 5360765 | 3546715.5 | 4544134 | 2163104 | 2871810.75 | 2622370 |
| Q6P2B1 | Tnpo3 | Transportin-3 | 87.42 | 1.84 | 104170 | 2 | 1951685.5 | 1150480.75 | 872282 | 2804439.5 | 2232083 | 2065495.25 | 1967878.38 | 2201347.25 | 1484934.75 | 1629370 | 1393186.75 | 593446.12 | 1839511.75 | 1613799.62 | 2478598 | 1925763.25 | 980791.88 | 3112571.75 | 2261261.75 | 2789963.25 |
| A2RUW1 | Tollip | Toll-interacting protein | 82.83 | 8.39 | 30315 | 2 | 79016.79 | 163089.5 | 109038.21 | 2 | 1011313.06 | 680354.56 | 2 | 1499901.5 | 1977270 | 1474591.38 | 1583405.5 | 1428239.12 | 694388.81 | 1253121.25 | 2 | 1170849.38 | 797764.75 | 2 | 1616261 | 2 |
| O88746 | Tom1 | Target of Myb1 membrane trafficking protein | 151.78 | 7.32 | 54325 | 3 | 2304783.25 | 2691731.5 | 1811431.38 | 1612652.25 | 718366.38 | 2074843.38 | 2449726.25 | 1647692.62 | 1608750 | 1746831.5 | 1717099.25 | 2028212.25 | 761371 | 1820374.75 | 1946754.5 | 1946736.75 | 2145593.5 | 2338606 | 2995585.75 | 2593314.5 |
| Q62760 | Tomm20 | Mitochondrial import receptor subunit TOM20 hor | 42.23 | 4.83 | 16284 | 1 | 475368.19 | 2 | 881704.81 | 722030.75 | 1167916.5 | 3068184.75 | 1040977.94 | 4584312.5 | 2 | 815893.94 | 644402.94 | 3252681.75 | 745495 | 1436316.38 | 319323.19 | 1332168.12 | 1548076.25 | 762030.19 | 316787.34 | 1989970 |
| Q9CPQ3 | Tomm22 | Mitochondrial import receptor subunit TOM22 hor | 208.64 | 29.58 | 15537 | 3 | 2 | 2 | 312611.38 | 616352 | 166209.55 | 414239.19 | 529844.56 | 559655.81 | 370499.88 | 777695.56 | 338373.31 | 238894.56 | 462608.72 | 702262.56 | 866754.88 | 2 | 558387.25 | 687657.31 | 326239.34 | 300679 |
| Q9CYG7 | Tomm34 | Mitochondrial import receptor subunit TOM34 | 340 | 24.6 | 34278 | 6 | 3941583.25 | 4968733.5 | 1972981.62 | 3836345 | 3722064.5 | 5365718.5 | 5365953.5 | 6517231.5 | 3640912.25 | 3200465 | 6479340.5 | 6504250 | 5319075.5 | 6143359 | 4189747.75 | 5403128 | 5171352 | 4974756.5 | 4855053 | 3802060.25 |
| Q75Q40 | Tomm40 | Mitochondrial import receptor subunit TOM40 hor | 194.55 | 11.91 | 37919 | 3 | 1602244.5 | 2933343 | 2946357 | 4440353 | 3429204 | 3378543 | 3552153.25 | 3590722.25 | 3102092 | 1336639 | 2305578.5 | 2203100 | 3226095.5 | 1985096.75 | 2290278.5 | 1386656.62 | 2306092.75 | 2711042.5 | 945519.38 | 2035124.5 |
| Q9CZW5 | Tomm70 | Mitochondrial import receptor subunit TOM70 | 292.04 | 9.66 | 67590 | 6 | 12400570 | 12977864 | 11430952 | 11422214 | 16287770 | 8822925 | 10124398 | 12283153 | 8627559 | 10502097 | 5700361.5 | 6496184.5 | 5898677 | 5873157 | 7224938.5 | 7384014 | 6515596 | 7827071.5 | 8956943 | 10142102 |
| Q04750 | Top1 | DNA topoisomerase 1 | 148.42 | 3.39 | 90876 | 3 | 2 | 2 | 2 | 81225.91 | 2 | 2 | 2 | 2 | 2 | 2 | 118877.52 | 210962.28 | 2 | 75914.19 | 2 | 92680.42 | 66899.98 | 2 | 2 | 2 |
| Q01320 | Top2a | DNA topoisomerase 2-alpha | 397.87 | 6.15 | 172790 | 8 | 3515953 | 3962353.5 | 3390934.5 | 4701950 | 4132424 | 2 | 759808.94 | 2 | 1798711.75 | 686417.06 | 2489620.75 | 5515715.5 | 7446122 | 6455125 | 3730762.25 | 1189278 | 3645835.5 | 2032580.88 | 4569197 | 1532961.75 |
| Q921T2 | Tor1aip1 | Torsin-1A-interacting protein 1 | 194.77 | 11.6 | 66781 | 5 | 469708.75 | 2 | 2 | 2 | 274803.16 | 2 | 2 | 194161.97 | 5230633 | 455051.12 | 397764.31 | 3967362 | 608338.19 | 2024274.62 | 940521.75 | 2383519.25 | 2 | 1124261.25 | 2 | 852345.12 |
| Q9ER38 | Tor3a | Torsin-3A | 106.69 | 4.94 | 43814 | 2 | 2977942.75 | 2 | 3356036.5 | 4983174.5 | 4943237 | 3649021.5 | 2961192 | 3675563 | 4559045.5 | 5803643.5 | 3562080.5 | 2210031.5 | 2730129.5 | 2672819.25 | 1277746.88 | 3269058.25 | 2793886.75 | 2 | 2075674.12 | 646518.12 |
| Q62393 | Tpd52 | Tumor protein D52 | 233.48 | 21.43 | 24313 | 4 | 6954791 | 10444103 | 8723636 | 7237767.5 | 8918302 | 4363196 | 4983518 | 6663636.5 | 10159531 | 7214435.5 | 4607022 | 10066947 | 5772097 | 8108600 | 7262510 | 8545552 | 8555740 | 7192413.5 | 9952381 | 14124554 |
| Q6PCT3 | Tpd52l2 | Tumor protein D54 | 160.36 | 18.18 | 23992 | 3 | 468399.66 | 1992112.25 | 752950.94 | 1029539.69 | 3107628.5 | 2099995.5 | 2115509 | 2936020 | 2681305.5 | 1457129.25 | 5620047 | 3863176.25 | 2717365.5 | 3184271 | 503415.84 | 3900990 | 2 | 2280054.5 | 4045697.25 | 2 |
| P17751 | Tpi1 | Triosephosphate isomerase | 1025.69 | 79.12 | 26713 | 15 | 49366594 | 141189490 | 99657570 | 100226194 | 79270738 | 78621386 | 99680954 | 78342858 | 141579250 | 115039730 | 199122594 | 284467426 | 156096050 | 207506770 | 174917794 | 225732226 | 212952322 | 129022266 | 151884994 | 183897026 |
| P04692 | Tpm1 | Tropomyosin alpha-1 chain | 590.08 | 21.83 | 32681 | 10 | 3486802.75 | 2456717 | 3724431.75 | 2264649.75 | 1379590.5 | 1795744 | 1818495.5 | 2860182.25 | 1511085.12 | 1176934 | 1646615.88 | 1786489.25 | 2275025 | 1678542.12 | 1075764.5 | 2163061.5 | 2393810 | 2031768.5 | 1518284 | 1952711 |
| P21107 | Tpm3 | Tropomyosin alpha-3 chain | 736.17 | 26.32 | 32994 | 13 | 19450480 | 18463368 | 16331821 | 17673392 | 17797538 | 13533726 | 13052179 | 20647862 | 9767908 | 12073933 | 11375328 | 12365018 | 11207079 | 9592222 | 5648078.5 | 13363112 | 12604268 | 7993012 | 15140918 | 11118335 |
| Q63610 | Tpm3 | Tropomyosin alpha-3 chain | 1215.64 | 53.63 | 29007 | 19 | 24650478 | 31099348 | 15865528 | 25081062 | 25065998 | 20392380 | 14547984 | 27821908 | 25176906 | 15535364 | 30913498 | 23580018 | 16534534 | 16315573 | 6738862.5 | 22962698 | 16193626 | 9419986 | 19651720 | 18244726 |
| Q6IRU2 | Tpm4 | Tropomyosin alpha-4 chain | 916.72 | 46.37 | 28468 | 15 | 10875327 | 5247633.5 | 20122770 | 10397363 | 6021337 | 6174427 | 9122537 | 14567880 | 6300010.5 | 11879345 | 5896682.5 | 6685493.5 | 11759330 | 8656093 | 6077928 | 4918144 | 7590588.5 | 8386581.5 | 3782855.5 | 4404055.5 |
| O89023 | Tpp1 | Tripeptidyl-peptidase 1 | 326.12 | 14.41 | 61342 | 5 | 1064131.38 | 716093.06 | 1706556 | 782679.19 | 474137.03 | 2 | 2 | 529322.62 | 2 | 1143496.88 | 2 | 2 | 2 | 2 | 2 | 2 | 2 | 2 | 2 | 2 |
| Q64514 | Tpp2 | Tripeptidyl-peptidase 2 | 318.74 | 6.97 | 139879 | 8 | 6939734 | 11578578 | 6995595 | 12076338 | 11263846 | 11481146 | 8521381 | 4769609 | 10422061 | 6278698 | 12467583 | 11307493 | 6755059.5 | 11341104 | 7374288 | 11163440 | 10242280 | 9924195 | 11847883 | 11160958 |
| F6ZDS4 | Tpr | Nucleoprotein TPR | 913.86 | 9.38 | 273990 | 18 | 6644318 | 4658031 | 4740075 | 14662877 | 6642258.5 | 18844698 | 17587484 | 17117626 | 8225466 | 6801699 | 11195330 | 12863432 | 10123901 | 6771325.5 | 3578806 | 16144664 | 10390366 | 751893.81 | 7336861.5 | 6150397 |
| P63028 | Tpt1 | Translationally-controlled tumor protein | 159.35 | 21.51 | 19462 | 3 | 632496.94 | 1163805.38 | 2201839.5 | 821358.81 | 1260006.12 | 2 | 2 | 69912.64 | 2 | 278445.5 | 2 | 2 | 888755.5 | 627035 | 690880.12 | 256722.27 | 764707.69 | 788575.75 | 2 | 2 |
| A2APB8 | Tpx2 | Targeting protein for Xklp2 | 35.81 | 1.61 | 85894 | 1 | 2 | 2 | 2 | 2 | 191230.53 | 2 | 2 | 2 | 2 | 2 | 2 | 658713.94 | 506127.25 | 491854.91 | 797042.19 | 504645.53 | 2 | 541120.38 | 2 | 556495.06 |
| Q6PFR5 | Tra2a | Transformer-2 protein homolog alpha | 202.41 | 11.74 | 32316 | 3 | 1070223.25 | 1770915.75 | 1004898.31 | 1376569.12 | 1843319.5 | 1603663.25 | 1973673.12 | 2896975.5 | 1010299.5 | 1236172 | 800958.62 | 595863.69 | 1418288.25 | 730321.12 | 787761.25 | 2 | 1225668.25 | 2 | 1144468.38 | 114308.3 |
| P62996 | Tra2b | Transformer-2 protein homolog beta | 337.06 | 24.65 | 33666 | 6 | 1155427.38 | 2 | 1524222 | 966512.38 | 1741104 | 1191390.12 | 1711636.25 | 1408816.38 | 1028627.56 | 1230759.62 | 2126586.5 | 803188.25 | 2277351.25 | 1143139 | 2237725 | 1539480.75 | 1245242.5 | 1192360.12 | 1321366 | 284378.59 |
| Q99JY4 | Trabd | TraB domain-containing protein | 35.84 | 3.19 | 42189 | 1 | 2 | 2 | 2 | 2 | 2 | 2 | 2 | 2 | 2 | 2 | 2 | 2 | 2 | 2 | 2 | 2 | 2 | 2 | 2 | 2 |
| Q5XI41 | Tram1 | Translocating chain-associated membrane protei | 148.37 | 5.08 | 43030 | 2 | 2 | 438812.84 | 287653.88 | 492305.81 | 2 | 435761.25 | 696673.12 | 230508.08 | 512854.25 | 2 | 158514.61 | 2 | 575900.12 | 788674.5 | 871277.75 | 210422 | 1078246.88 | 412852.53 | 455728.72 | 369438.44 |
| Q9CQN1 | Trap1 | Heat shock protein 75 kDa, mitochondrial | 776.83 | 25.64 | 80209 | 13 | 5324754 | 5963155.5 | 7889784.5 | 8246447 | 7963160 | 3295123.5 | 2360031.5 | 4711253.5 | 6236478 | 7744682 | 1217339.5 | 3922995.25 | 3782413.5 | 2541079 | 4951045 | 2380164.5 | 3701084 | 2673585.5 | 5167504 | 3413228 |

| | | | | | | | | | | | | | | | | | | | | | | | | | | |
|---|---|---|---|---|---|---|---|---|---|---|---|---|---|---|---|---|---|---|---|---|---|---|---|---|---|---|
| B2RXC1 | Trappc11 | Trafficking protein particle complex subunit 11 | 59.89 | 0.71 | 128396 | 1 | 2 | 2 | 2 | 2 | 2 | 2 | 2 | 2 | 2 | 2 | 2 | 2 | 2 | 2 | 2 | 2 | 2 | 2 | 2 | 2 |
| Q9CQA1 | Trappc5 | Trafficking protein particle complex subunit 5 | 40.74 | 5.32 | 20795 | 1 | 2 | 1085125.88 | 1373879.38 | 1106031.12 | 778326.12 | 1178445.75 | 822686.56 | 2 | 595075.06 | 781428.75 | 1001262.12 | 736376.88 | 883864.38 | 577655.31 | 2 | 1273227 | 1304539 | 1331431.25 | 982147.25 | 1171029.62 |
| Q91XB0 | Trex1 | Three-prime repair exonuclease 1 | 190.95 | 14.65 | 33675 | 3 | 2 | 2 | 327759.97 | 112275.16 | 1146648 | 137316.39 | 268779.28 | 166337.39 | 616796.94 | 1137221.12 | 113727.91 | 379023.62 | 396797.72 | 186719.89 | 279014 | 70418.02 | 159725.08 | 194817.19 | 80609.37 | 163041.67 |
| Q62318 | Trim28 | Transcription intermediary factor 1-beta | 741.82 | 21.58 | 88847 | 15 | 5610832.5 | 5626376.5 | 2334451.25 | 7003354 | 15418865 | 6814163.5 | 7182846 | 6676504 | 15319001 | 10847607 | 19209926 | 20177034 | 10219987 | 13639740 | 8054843.5 | 12459017 | 9285698 | 5853285.5 | 8178237 | 9025606 |
| F1LP64 | Trip12 | E3 ubiquitin-protein ligase TRIP12 | 75.56 | 1.23 | 223928 | 2 | 2 | 926422.31 | 2 | 2 | 394338.94 | 2 | 2 | 2 | 593076.69 | 2 | 2 | 2 | 630118.38 | 2 | 2 | 2 | 952046.56 | 2 | 2 | 2 |
| A2RSY6 | Trmt1l | TRMT1-like protein | 79.02 | 3.98 | 80861 | 2 | 81842.43 | 1972818.88 | 1303661.75 | 2727709 | 53617.79 | 1859723 | 64453.75 | 1703613.5 | 1457303.88 | 107947.08 | 1591620.25 | 166001.94 | 1343897.5 | 1250039.5 | 1003014.44 | 2284531.75 | 2170842.25 | 2533143.75 | 2562271.5 | 2 |
| Q8CE96 | Trmt6 | tRNA (adenine(58)-N(1))-methyltransferase non-c | 37.81 | 3.62 | 55518 | 1 | 846834.69 | 320172.72 | 1297057.62 | 247007.08 | 2 | 141931.47 | 352444.41 | 2 | 2 | 2 | 2 | 235731.39 | 2 | 2 | 2 | 209432.8 | 2 | 2 | 2 | 2 |
| Q9WTR1 | Trpv2 | Transient receptor potential cation channel subfa | 74.94 | 3.84 | 85965 | 2 | 2 | 2 | 1220075.88 | 2 | 2 | 2 | 982212.31 | 1641264.38 | 1081364.5 | 863485.94 | 2 | 2 | 2 | 1063143.88 | 2 | 2 | 2 | 2 | 828090.5 | 2 |
| Q61187 | Tsg101 | Tumor susceptibility gene 101 protein | 115.27 | 6.14 | 44124 | 2 | 2258467.25 | 3154123.75 | 2057500 | 2472344.25 | 2376350.25 | 2534412 | 2381729.5 | 2507518.25 | 1999620.12 | 2228298.25 | 2059750.12 | 1654396.5 | 1356621 | 1960267.88 | 2309784 | 2162766 | 2225655.75 | 2439804 | 380078.03 | 2463109.25 |
| Q62348 | Tsn | Translin | 219.63 | 20.61 | 26201 | 4 | 5804024 | 4966648 | 8394774 | 5573997.5 | 7664952 | 6215330 | 5960807 | 6784646 | 4624174 | 5097960.5 | 3406534.5 | 4519130.5 | 5228151 | 3281186 | 6940374 | 5247581 | 5943712.5 | 7939481 | 4279416 | 5723007.5 |
| P50637 | Tspo | Translocator protein | 213.54 | 46.75 | 18841 | 4 | 2 | 2 | 172842.33 | 2518437.25 | 155355.84 | 2 | 258453.05 | 1151412.12 | 14095487 | 141526.88 | 2 | 5991234.5 | 984473.75 | 7337824.5 | 1104948.88 | 7321314 | 2244340.75 | 2 | 1029562.25 | 17269878 |
| Q5SWD9 | Tsr1 | Pre-rRNA-processing protein TSR1 homolog | 66.55 | 1.37 | 92105 | 1 | 980786.06 | 265945.53 | 2 | 321527.47 | 884992.25 | 2 | 953430.81 | 232940.16 | 946177.75 | 395489.41 | 2 | 475918.62 | 595773.25 | 790998.31 | 2 | 2 | 2 | 175180.83 | 2 | 2 |
| Q8CD92 | Ttc27 | Tetratricopeptide repeat protein 27 | 39.46 | 1.06 | 96431 | 1 | 1616956.62 | 549734.44 | 1431277.5 | 2 | 534346.62 | 2 | 457092.16 | 2 | 2 | 2 | 2 | 772410.06 | 668951.75 | 437457.12 | 2359938 | 1477134.38 | 2606656.25 | 2013422.12 | 1547800.12 | 4136664.75 |
| Q3UDE2 | Ttll12 | Tubulin--tyrosine ligase-like protein 12 | 96.15 | 3.13 | 74043 | 2 | 579557.56 | 1079832.38 | 2 | 950216.81 | 2624909.75 | 1883497.5 | 2 | 245446.45 | 1355735.88 | 238950.36 | 2569195.5 | 2390530.25 | 2104270 | 1891769 | 2 | 2709361 | 2297493 | 2102267 | 3047692.5 | 2309476 |
| A2ASS6 | Ttn | Titin | 36.04 | 0.02 | 3906488 | 1 | 2 | 2 | 2 | 2 | 2 | 2 | 2 | 2 | 657788.44 | 2 | 2 | 2 | 2 | 2 | 2 | 2 | 2 | 2 | 2 | 2 |
| P68373 | Tuba1c | Tubulin alpha-1C chain | 1832.37 | 64.59 | 49909 | 27 | 2 | 2 | 2 | 2 | 585939.94 | 272664.97 | 2 | 2 | 2 | 515423.19 | 1130671.62 | 2 | 782119.31 | 870924.25 | 1410365.5 | 2 | 1335472 | 1589970.88 | 630258.31 | 2697449.25 |
| P68368 | Tuba4a | Tubulin alpha-4A chain | 1488.85 | 55.8 | 49924 | 24 | 571617.38 | 2158147 | 89221.08 | 543444 | 381185.34 | 72977.35 | 186093.3 | 119503.62 | 191731.58 | 151869.95 | 116429.66 | 86526.78 | 108316.88 | 203962.94 | 393105.38 | 2 | 229170.61 | 260863.64 | 540562.12 | 165515.52 |
| P85108 | Tubb2a | Tubulin beta-2A chain | 1741.05 | 59.1 | 49907 | 26 | 2150158.5 | 2015168.88 | 1906833.62 | 1018095.44 | 456263.06 | 1063951.12 | 949314.06 | 1505141.88 | 952916.12 | 553944.12 | 2 | 742914.81 | 2 | 1004586.06 | 395686.22 | 861331.81 | 2 | 2 | 860581 | 2 |
| Q3KRE8 | Tubb2b | Tubulin beta-2B chain | 1728.97 | 59.1 | 49953 | 25 | 1154048.12 | 1821816.75 | 1777855.62 | 1222610.75 | 960970.75 | 1392329.88 | 1071612.88 | 1544969.5 | 877473.06 | 953444.25 | 694621.19 | 2 | 955417.19 | 842649 | 717867.81 | 846916.25 | 930017.25 | 913751.81 | 862855.31 | 749012.38 |
| P68372 | Tubb4b | Tubulin beta-4B chain | 1918.74 | 68.99 | 49831 | 28 | 16967294 | 18362810 | 20490194 | 14452172 | 21300686 | 18601680 | 10612326 | 21456926 | 14581788 | 15516207 | 19906986 | 10145814 | 16562093 | 19967836 | 12900074 | 13827706 | 14442539 | 16789958 | 17234050 | 17660984 |
| P69897 | Tubb5 | Tubulin beta-5 chain | 1990.48 | 69.14 | 49671 | 28 | 44639386 | 20875576 | 57374730 | 35556894 | 33452468 | 15226076 | 23965396 | 22910238 | 22234350 | 42714358 | 14741992 | 11209350 | 20266496 | 25008878 | 40266242 | 7729328 | 16077878 | 15082605 | 8151190 | 15312530 |
| Q922F4 | Tubb6 | Tubulin beta-6 chain | 1168.36 | 45.19 | 50090 | 19 | 15889175 | 19201524 | 17351284 | 14801321 | 13201588 | 12685784 | 13338699 | 14150360 | 8979408 | 9314541 | 7642172.5 | 5925416 | 10781049 | 10459778 | 8820867 | 5648359.5 | 10516075 | 10643566 | 10261886 | 10610539 |
| P83887 | Tubg1 | Tubulin gamma-1 chain | 49.08 | 2 | 51101 | 1 | 2 | 580096.94 | 870027.75 | 400360.78 | 584502.75 | 2 | 2 | 582814.62 | 2 | 2 | 2 | 371846.44 | 698151.62 | 725151.5 | 2 | 835689.5 | 2 | 538804.19 | 109405.4 | 2 |
| Q8BFR5 | Tufm | Elongation factor Tu, mitochondrial | 409.73 | 19.91 | 49508 | 8 | 2413480.5 | 6973144.5 | 4475464.5 | 8730422 | 11447985 | 7355173.5 | 8178677 | 10500777 | 11571655 | 4893879.5 | 13151778 | 13015023 | 12089794 | 12788209 | 7456243.5 | 13313812 | 12485534 | 8009733.5 | 9309710 | 10670429 |
| Q91YR1 | Twf1 | Twinfilin-1 | 98.76 | 4.86 | 40079 | 2 | 2014464.88 | 2406623.25 | 1770587.62 | 1655929 | 2186274.5 | 1887858.12 | 2093488.38 | 2439260.25 | 1887440.12 | 2120886.5 | 1183455.38 | 1325746.12 | 1119010 | 1562541.62 | 1089390.88 | 992702.81 | 1256015.62 | 1771850 | 2 | 1019428.31 |
| Q6PAM1 | Txlna | Alpha-taxilin | 116.03 | 7.04 | 62369 | 2 | 159965.03 | 1180602 | 59314.08 | 775747.38 | 98313.29 | 286787.84 | 2 | 1233340.75 | 408326.5 | 2 | 784391.44 | 1133766.25 | 953787.81 | 1348062.12 | 2 | 2 | 2 | 1551074.75 | 1971386.38 | 2 |
| P10639 | Txn | Thioredoxin | 278.19 | 37.14 | 11675 | 4 | 1630619.12 | 931475.38 | 4297961 | 1031324.75 | 70866.51 | 1973540.75 | 1325057.25 | 1891078 | 8265495 | 1453745 | 18068446 | 31003022 | 23280802 | 28040470 | 27170098 | 27343760 | 21295408 | 3147786 | 4503734.5 | 13414000 |
| Q498E0 | Txndc12 | Thioredoxin domain-containing protein 12 | 97.41 | 14.12 | 19019 | 2 | 605492.25 | 2 | 1673868 | 234480.73 | 2647701.25 | 156857.11 | 1377551.62 | 1328356.75 | 2051883.5 | 2294133 | 1421070.88 | 2652927.75 | 1386196.75 | 2791391 | 1573922.25 | 232729.59 | 1488168.88 | 1351290 | 280476.66 | 1265535.5 |
| Q9CQM5 | Txndc17 | Thioredoxin domain-containing protein 17 | 55.83 | 8.94 | 14015 | 2 | 2 | 2 | 2 | 1823332.5 | 2908948.75 | 4543083.5 | 3712258.5 | 5443225 | 3946647.25 | 2856001.5 | 1418021.5 | 996871.75 | 2 | 2 | 2 | 2 | 2 | 2 | 577906.44 | 2 |
| Q91W90 | Txndc5 | Thioredoxin domain-containing protein 5 | 234.36 | 11.51 | 46415 | 4 | 395863.62 | 1122786 | 2 | 1283760.5 | 1253768.88 | 1001219.5 | 990431.12 | 194766.75 | 1204726.12 | 85031.59 | 2 | 106073.95 | 767362.31 | 809436.12 | 2 | 2 | 230834.75 | 2 | 2 | 2 |
| Q8CDN6 | Txnl1 | Thioredoxin-like protein 1 | 173.37 | 18.34 | 32237 | 4 | 340801.19 | 1385897.75 | 980847.62 | 2 | 877980.75 | 1058351.88 | 1215002.25 | 1062080 | 1864510.75 | 1159168 | 7575366 | 6509465.5 | 3940408.5 | 5907890 | 2748887.75 | 7773859 | 4149049.5 | 5284128 | 4080512.5 | 7419201 |
| Q9JMH6 | Txnrd1 | Thioredoxin reductase 1, cytoplasmic | 1055.46 | 31 | 67084 | 17 | 5671443.5 | 3921003 | 9653176 | 8867238 | 11829066 | 7351943 | 8161773.5 | 15499199 | 15774877 | 17937946 | 41038174 | 44365510 | 39475610 | 31023018 | 71183794 | 60260282 | 80854162 | 75358306 | 73789490 | 78949154 |
| Q9JLT4 | Txnrd2 | Thioredoxin reductase 2, mitochondrial | 167 | 5.15 | 56603 | 2 | 2 | 2 | 2 | 2 | 2 | 2 | 2 | 2 | 642741.81 | 626966.5 | 1419506 | 2 | 1071698.25 | 565998.94 | 930757.31 | 1093187.25 | 1109779 | 1460175.75 | 1148533 | 918230.06 |
| Q9D883 | U2af1 | Splicing factor U2AF 35 kDa subunit | 205.42 | 15.06 | 27815 | 4 | 43819.68 | 2 | 150158.34 | 1219042 | 271856.03 | 205363 | 559539.38 | 167092.7 | 1458160.5 | 318990.12 | 374137 | 1241105.25 | 678882.56 | 1667639.5 | 1160852.25 | 79264.78 | 832939.88 | 730252 | 222880.69 | 525472.44 |
| P26369 | U2af2 | Splicing factor U2AF 65 kDa subunit | 501.74 | 33.26 | 53517 | 7 | 817160.56 | 231462.83 | 3125657.5 | 1670443.88 | 2334063.5 | 342061.84 | 203520.94 | 338943.12 | 1569511.25 | 1671336.62 | 1072433.25 | 1263945.5 | 2053447.5 | 542477.25 | 2455430.25 | 752431.75 | 1004791.31 | 575184 | 643605.25 | 1258045 |
| Q6NV83 | U2surp | U2 snRNP-associated SURP motif-containing pro | 69.33 | 1.36 | 118261 | 1 | 1175591.5 | 147707.28 | 792490.06 | 2 | 1182199.75 | 2 | 2 | 2 | 827886.31 | 2 | 2 | 2 | 2 | 2 | 2 | 2 | 2 | 2 | 2 | 2 |
| Q3TW96 | Uap1l1 | UDP-N-acetylhexosamine pyrophosphorylase-like | 595.35 | 23.08 | 56614 | 11 | 4698977 | 6107068.5 | 5704969.5 | 4713651.5 | 10918309 | 9752193 | 7329518 | 11749268 | 7791612 | 7035345.5 | 18799336 | 16035137 | 12304246 | 14502592 | 14716110 | 16319462 | 11120095 | 9030835 | 8932153 | 7711906 |
| Q02053 | Uba1 | Ubiquitin-like modifier-activating enzyme 1 | 1431.37 | 27.41 | 117809 | 22 | 12910365 | 13192582 | 9111129 | 16518201 | 17878580 | 14086806 | 15734254 | 10452510 | 18400642 | 19026422 | 16603296 | 23405408 | 21062928 | 22515174 | 21584850 | 23774020 | 19454994 | 13842178 | 13571138 | 15998115 |
| Q9Z1F9 | Uba2 | SUMO-activating enzyme subunit 2 | 128.04 | 3.13 | 70569 | 2 | 1006333.25 | 1242126.88 | 769803 | 1081954.25 | 1124739 | 813208.31 | 2 | 979645.25 | 2 | 1322103.25 | 1513881.75 | 2326474.5 | 2924663 | 3177004.75 | 938542.56 | 2852374 | 1218270.25 | 3849028.5 | 2385964.25 | 3446494.75 |
| Q8C878 | Uba3 | NEDD8-activating enzyme E1 catalytic subunit | 84.21 | 3.9 | 51737 | 2 | 1503199.75 | 3489808.5 | 2 | 2061882 | 2 | 3450501 | 3474228 | 456969.19 | 2733813 | 2527444.25 | 2297319.25 | 2775811.75 | 2393031.25 | 2663012.5 | 1717269.75 | 2425454.75 | 2541395 | 2322277.5 | 453467.91 | 1905788.38 |
| Q8C7R4 | Uba6 | Ubiquitin-like modifier-activating enzyme 6 | 56.95 | 0.85 | 117966 | 1 | 49065.23 | 2 | 2 | 2 | 34764.3 | 76075.91 | 2 | 103579.14 | 319537.47 | 2 | 41751.72 | 2 | 81240.73 | 2 | 2 | 2 | 2 | 2 | 2 | 2 |
| Q80X50 | Ubap2l | Ubiquitin-associated protein 2-like | 350.23 | 6.68 | 116799 | 5 | 1931714.5 | 3607422 | 289851.91 | 4220809.5 | 4610353.5 | 1368173 | 1766974.25 | 2141159.75 | 4144327.5 | 3118268 | 2136645 | 2900066.25 | 1712850.75 | 2282699 | 2435067 | 3185849.75 | 1912506.12 | 662518.62 | 2574492.25 | 1728168.12 |
| P68037 | Ube2l3 | Ubiquitin-conjugating enzyme E2 L3 | 137.77 | 35.71 | 17862 | 3 | 722612.75 | 80998.12 | 1421456.88 | 105207.96 | 695038.38 | 2 | 2 | 2 | 2 | 929534.88 | 1012053.19 | 111069.36 | 1047158.38 | 935646.56 | 615516 | 1479330.62 | 1513641.12 | 992479.31 | 1900461.62 | 959859.12 |
| P61082 | Ube2m | NEDD8-conjugating enzyme Ubc12 | 138.23 | 16.94 | 20900 | 3 | 4742012.5 | 7947903 | 4337679.5 | 5535159.5 | 10709266 | 11848901 | 6676794.5 | 11055411 | 7472539 | 3728189.5 | 6589180 | 13109207 | 11076121 | 12976171 | 4224475.5 | 8798628 | 11079572 | 8293854 | 10806291 | 5859339.5 |
| Q6ZPJ3 | Ube2o | (E3-independent) E2 ubiquitin-conjugating enzy | 40.76 | 0.78 | 140834 | 1 | 1493602 | 1744390.88 | 1423655 | 1196438 | 1860140.38 | 2 | 1018038.81 | 230029.14 | 758510.19 | 986246.12 | 1353032.25 | 888065.62 | 1169581.38 | 1371126.12 | 761636.12 | 1021396.25 | 1073916.25 | 1202322.38 | 1227189.75 | 1454170.38 |
| Q9CZY3 | Ube2v1 | Ubiquitin-conjugating enzyme E2 variant 1 | 267.63 | 37.41 | 16355 | 5 | 4941600.5 | 7808447 | 6861251 | 5580290 | 8682284 | 7339431 | 4195866 | 12069841 | 8073664 | 9337045 | 14021658 | 12100402 | 10484218 | 12643560 | 6836674 | 7049228 | 5729392 | 6849084.5 | 10052224 | 7020644.5 |
| O08759 | Ube3a | Ubiquitin-protein ligase E3A | 154.2 | 4.25 | 99819 | 3 | 4538032 | 1682012 | 2727902.25 | 1953130 | 2340947.25 | 712224 | 1647487.5 | 3218708.75 | 2488064 | 1820497.25 | 997921.38 | 599528.06 | 2866592.5 | 2093069 | 1621826.25 | 2 | 3622328 | 2 | 1800231.5 | 2 |
| A2AN08 | Ubr4 | E3 ubiquitin-protein ligase UBR4 | 259.09 | 1.16 | 572290 | 5 | 1345374.38 | 3011019.5 | 1455434.12 | 1994360.5 | 2748329.25 | 2832186.5 | 3367964 | 3352660 | 2645126.5 | 2220983.75 | 2949992.25 | 2453716.5 | 2986506.5 | 2708016.25 | 2228317.5 | 2062075.75 | 2007230.12 | 1921059.62 | 2157865.5 | 2467156.25 |
| Q922Y1 | Ubxn1 | UBX domain-containing protein 1 | 149.75 | 14.81 | 33573 | 3 | 517197.91 | 704623.06 | 517041.06 | 862257.19 | 1048461.88 | 2 | 890242.5 | 1189327.38 | 2245112.5 | 898708 | 2479077 | 2337892 | 1729683.25 | 3299714.25 | 2027762.25 | 4058855 | 3126169.75 | 2835307.5 | 3289940.75 | 2322664 |
| Q8VCH8 | Ubxn4 | UBX domain-containing protein 4 | 205.24 | 12.45 | 56472 | 4 | 23049706 | 63928290 | 16709360 | 36582354 | 26852198 | 21990278 | 19101856 | 2406963 | 10070016 | 8346575 | 5193491 | 3302100.5 | 4521407 | 4991237.5 | 2630675.75 | 6785781 | 4565369.5 | 3875161.75 | 5193261 | 3748322.75 |
| Q9JKB1 | Uchl3 | Ubiquitin carboxyl-terminal hydrolase isozyme L3 | 136.93 | 15.65 | 26152 | 3 | 734748.12 | 1743993.25 | 1808517 | 451892.28 | 2932756.75 | 671528.31 | 1041139.56 | 952485.94 | 974674.31 | 1476786.12 | 1546132 | 642729.38 | 1882503.75 | 1424470.5 | 1535665.75 | 857299.56 | 967986.94 | 2081528 | 1028546.31 | 791679.94 |
| Q9WUP7 | Uchl5 | Ubiquitin carboxyl-terminal hydrolase isozyme L5 | 191.14 | 10.03 | 37617 | 4 | 3258122.25 | 1715578.62 | 2282307.75 | 1805412.25 | 2656863.5 | 1551647.25 | 2386165.25 | 2243334 | 941658.44 | 2139150 | 2214144.5 | 1501897.88 | 2366363.75 | 2043823.5 | 2420157.25 | 2167116.25 | 2001086 | 1150485.12 | 1442805.62 | 2 |
| Q91YL3 | Uckl1 | Uridine-cytidine kinase-like 1 | 196.83 | 6.57 | 60842 | 4 | 1774695.62 | 3880068.25 | 2430930.75 | 1290261.25 | 2553645.5 | 4444206 | 2761693 | 2035405.88 | 2892461 | 2249845.25 | 1764713.75 | 1598456.5 | 1728944.75 | 1707290.38 | 1469988.62 | 2499553.5 | 2711577.5 | 951050.94 | 1768204.38 | 1162310.12 |
| B2GV24 | Ufl1 | E3 UFM1-protein ligase 1 | 50 | 1.26 | 89585 | 1 | 2 | 1098790.38 | 990196.75 | 988665.25 | 2 | 2 | 774761.06 | 1114303.75 | 912662.19 | 928007.69 | 2 | 2 | 2 | 2 | 2 | 2 | 2 | 2 | 2 | 2 |
| O70199 | Ugdh | UDP-glucose 6-dehydrogenase | 45.15 | 2.03 | 54892 | 1 | 2 | 1819985.12 | 1926059 | 2 | 2 | 2 | 544803.69 | 2 | 428012.56 | 580923.19 | 2 | 965754.38 | 2 | 593627.12 | 682711.38 | 2 | 2 | 2 | 2 | 2 |
| Q6P5E4 | Uggt1 | UDP-glucose:glycoprotein glucosyltransferase 1 | 1068.2 | 17.92 | 176434 | 19 | 5874343.5 | 4640859 | 6336138.5 | 8477943 | 12424122 | 1172242.25 | 3601360.25 | 6495551 | 6660774.5 | 9347603 | 3568806.25 | 8137765 | 5264757.5 | 7692957 | 7619951.5 | 3412674.25 | 5486898 | 6476444 | 4271130 | 3422755.5 |
| Q91ZJ5 | Ugp2 | UTP--glucose-1-phosphate uridylyltransferase | 132.77 | 6.3 | 56979 | 3 | 2121695.5 | 1910442.75 | 1589530.25 | 2208459.5 | 1949701.62 | 1462363.5 | 1689791.5 | 3163707 | 2275229.25 | 2426424 | 2723114.5 | 2714883.5 | 2726157 | 3042370.25 | 1689267.75 | 4360011.5 | 2392356 | 2533493.75 | 2604402 | 3156463 |
| Q6ZQM8 | Ugt1a7 | UDP-glucuronosyltransferase 1A7 | 502.74 | 24.48 | 59758 | 9 | 10394196 | 10147638 | 8692001 | 10775481 | 8564512 | 4752067 | 4893916 | 1766270.62 | 8216869 | 12364601 | 4064935.5 | 4473645 | 6666035.5 | 7599017.5 | 9448881 | 4967041 | 7571622.5 | 8055529.5 | 5561700 | 6093440.5 |
| Q8VDF2 | Uhrf1 | E3 ubiquitin-protein ligase UHRF1 | 417.62 | 13.94 | 88304 | 8 | 501229.19 | 318297.44 | 2 | 2 | 2 | 2 | 465218.53 | 2 | 321459.38 | 631762.06 | 552963.5 | 2 | 2 | 1014908.12 | 460815.31 | 2 | 677237.44 | 2 | 2 | 2 |
| P13439 | Umps | Uridine 5'-monophosphate synthase | 194.56 | 9.56 | 52292 | 4 | 3413097.5 | 3141355 | 4127764 | 7117542.5 | 5014184 | 7592094 | 6357040.5 | 6166939 | 2682850 | 4328500.5 | 3113839.5 | 5362756.5 | 5169316.5 | 2331315 | 3504308.5 | 1504251.38 | 2871501 | 1344770.12 | 1656161.88 | 2752625.5 |
| Q9EPU0 | Upf1 | Regulator of nonsense transcripts 1 | 290.54 | 8.45 | 123967 | 8 | 2436105 | 1458344.5 | 561055.88 | 2586778.75 | 1420690.5 | 1952966.62 | 2 | 2199038.75 | 849115.75 | 88127.48 | 1557243.25 | 2300660 | 681410.25 | 1683146.5 | 826069.81 | 809195.12 | 1102840 | 2 | 1092105.38 | 1029536.38 |
| Q8R1I1 | Uqcr10 | Cytochrome b-c1 complex subunit 9 | 71.78 | 12.5 | 7446 | 2 | 5766986 | 3071199 | 3000463 | 2156913.75 | 2221980 | 3522641 | 4137132 | 2335787 | 3089473 | 1538639.75 | 88681.15 | 1791798.75 | 1928256.5 | 1318080.25 | 480309.03 | 2093396 | 748677.44 | 1036607.5 | 1071987.88 | 2 |
| Q9CZ13 | Uqcrc1 | Cytochrome b-c1 complex subunit 1, mitochondri | 592.92 | 32.29 | 52852 | 12 | 9092296 | 7421652 | 12121505 | 8502990 | 8993702 | 3629811 | 5893354.5 | 10417681 | 6239692.5 | 9316486 | 5716587 | 5610614.5 | 8983033 | 7041600 | 8807728 | 6660672 | 6767506 | 9945668 | 5845589.5 | 6684926.5 |
| Q9DB77 | Uqcrc2 | Cytochrome b-c1 complex subunit 2, mitochondri | 684.98 | 40.84 | 48235 | 12 | 21428708 | 16732434 | 8237963 | 12520087 | 6114679 | 13852436 | 11869432 | 5845637.5 | 27942916 | 31540846 | 13698238 | 14283418 | 10762057 | 11610451 | 27855942 | 24425538 | 6271483 | 22550546 | 19801378 | 33384994 |
| Q9CR68 | Uqcrfs1 | Cytochrome b-c1 complex subunit Rieske, mitoch | 307.31 | 40.51 | 29368 | 6 | 1607189.38 | 1987266.75 | 792605.75 | 2439204.5 | 7550921.5 | 2355013.75 | 1984827.62 | 5800530 | 3957109.5 | 4127987.25 | 1185013.12 | 2512890.5 | 987781.62 | 611796.75 | 2 | 3136882 | 2062482 | 2 | 360136.56 | 756782.06 |
| P99028 | Uqcrh | Cytochrome b-c1 complex subunit 6, mitochondri | 33.89 | 10.11 | 10435 | 1 | 3752241.25 | 3796178 | 3297227 | 2744369.5 | 2483071.75 | 3371445 | 2737438.5 | 737567.25 | 3013006 | 1990711.25 | 3029071 | 2363139.25 | 3001882.25 | 2480035.5 | 2404919 | 2415295.75 | 2946950 | 3928456.25 | 3222682.5 | 3213813.75 |
| P70697 | Urod | Uroporphyrinogen decarboxylase | 144.71 | 9.54 | 40692 | 2 | 729205.94 | 87417.63 | 959506.44 | 239805.2 | 1114870.62 | 334498.19 | 375606.94 | 89954.34 | 1196149.5 | 1056214 | 268746.62 | 394185.22 | 214891.09 | 570987.38 | 721233.12 | 2 | 286703.56 | 157952.97 | 435148 | 2 |
| Q9Z1Z0 | Uso1 | General vesicular transport factor p115 | 200.73 | 5.53 | 106983 | 4 | 91956.41 | 1282175.12 | 2363185.25 | 222720.27 | 2126943.75 | 2 | 1520967.62 | 2453300 | 1098754.12 | 3962683 | 2208877.5 | 2059899.88 | 3879575 | 3810961.75 | 3460845 | 929548.5 | 1247306 | 1918411.25 | 2467038.25 | 1024101.25 |
| Q9JMA1 | Usp14 | Ubiquitin carboxyl-terminal hydrolase 14 | 574 | 26.17 | 56002 | 12 | 4537725 | 5146324 | 7674032 | 3482739 | 6395692 | 3915662 | 8360373 | 8969435 | 6998561 | 3590243 | 6656504.5 | 5957903.5 | 6436780 | 5410833 | 3859519.25 | 8090497 | 4113633 | 2792867.5 | 3595531.75 | 2579740.5 |
| Q3TIX9 | Usp39 | Ubiquitin carboxyl-terminal hydrolase 39 | 94.87 | 4.08 | 65146 | 2 | 1012559.25 | 1730063.5 | 2 | 2038910.75 | 1515524.75 | 2364807.25 | 1510663.25 | 1667901.62 | 2510950.5 | 1157466.38 | 1306162.88 | 1476448.62 | 1296133.38 | 1352265.38 | 1147010.88 | 2674161.5 | 1472799.75 | 2045846.62 | 1312326.38 | 1607522.62 |
| B2GUZ1 | Usp4 | Ubiquitin carboxyl-terminal hydrolase 4 | 204.77 | 6.14 | 108373 | 5 | 2 | 2 | 2042407 | 2 | 682766.31 | 2 | 87902.69 | 1327681.12 | 2 | 967950.31 | 2 | 841662.12 | 2028738.5 | 1856791.25 | 754535.12 | 2 | 1956147.88 | 2 | 2 | 127144.51 |
| P56399 | Usp5 | Ubiquitin carboxyl-terminal hydrolase 5 | 778.8 | 19.81 | 95833 | 14 | 13070915 | 16924488 | 15965610 | 12209738 | 15600410 | 11185864 | 9842009 | 7456768 | 10658277 | 12083775 | 10102161 | 12295256 | 12003999 | 10301452 | 11631137 | 10105381 | 11188433 | 16016482 | 11688334 | 11780362 |
| Q80U87 | Usp8 | Ubiquitin carboxyl-terminal hydrolase 8 | 37.16 | 1.57 | 122611 | 1 | 2 | 2 | 2 | 2 | 2 | 2 | 2 | 2 | 718498.56 | 235925.64 | 2 | 2 | 2 | 2 | 2 | 2 | 2 | 394118.69 | 2 | 2 |
| Q5SSI6 | Utp18 | U3 small nucleolar RNA-associated protein 18 ho | 211.19 | 9.24 | 61218 | 4 | 329919.22 | 544081.69 | 89515.34 | 629878.25 | 150308.78 | 718340.56 | 584962.19 | 2 | 119076.38 | 164867.22 | 72028.64 | 492073 | 465162.19 | 707519 | 526102.19 | 2 | 2 | 2 | 2 | 2 |
| Q9JI13 | Utp3 | Something about silencing protein 10 | 50.5 | 2.35 | 53399 | 1 | 531038.62 | 2 | 2 | 2 | 2 | 2 | 491409.56 | 738324.25 | 2 | 599313.44 | 2 | 2 | 2 | 2 | 2 | 2 | 2 | 2 | 2 | 2 |
| Q80W92 | Vac14 | Protein VAC14 homolog | 101.98 | 3.19 | 88068 | 2 | 1028617.12 | 1589233.5 | 715809.69 | 1565242.75 | 1616001.62 | 809300.75 | 1990515.5 | 2 | 1180903.5 | 951926.88 | 2 | 2 | 2 | 2 | 2 | 2 | 2 | 2 | 2 | 2 |
| P63024 | Vamp3 | Vesicle-associated membrane protein 3 | 43 | 8.74 | 11480 | 1 | 636520 | 2 | 942776.06 | 842080.62 | 2 | 2 | 737122.5 | 1416665.75 | 860470.31 | 1527755 | 2 | 2 | 2 | 520568.09 | 849556.5 | 2 | 872398.56 | 1508437.12 | 2 | 2 |
| P70280 | Vamp7 | Vesicle-associated membrane protein 7 | 38 | 4.09 | 24967 | 1 | 2 | 2 | 2 | 2 | 2 | 2 | 2 | 2 | 2 | 2 | 2 | 2 | 2 | 443213.44 | 2 | 2 | 2 | 2 | 2 | 2 |
| O70404 | Vamp8 | Vesicle-associated membrane protein 8 | 35.29 | 8.91 | 11451 | 1 | 2 | 2 | 2 | 2 | 92062.99 | 739731.62 | 630138.69 | 199392.16 | 1981702.25 | 2464524.25 | 1465406 | 1959775.38 | 2080573.62 | 2299968.75 | 1480069.5 | 2 | 2250242.75 | 1730968.88 | 731385.5 | 2097928.5 |
| Q9WV55 | Vapa | Vesicle-associated membrane protein-associated | 345.69 | 26.91 | 27855 | 7 | 4226192 | 9751889 | 882824.81 | 6501073 | 2270625.75 | 17022140 | 14835281 | 3925440.75 | 17807618 | 9709440 | 13513602 | 18000004 | 13389884 | 17112306 | 4484620.5 | 20264530 | 13686844 | 12181758 | 18005776 | 6909229 |
| Q9QY76 | Vapb | Vesicle-associated membrane protein-associated | 293.89 | 19.75 | 26946 | 6 | 2 | 1060304.25 | 2 | 1326100.12 | 2 | 2 | 2 | 2 | 2 | 2 | 1328850.75 | 1671131.62 | 1322898 | 3011038 | 2 | 2757720 | 1788616.25 | 1396066.25 | 339210.72 | 1486069.5 |
| Q9Z1Q9 | Vars1 | Valine--tRNA ligase | 1205.42 | 22.41 | 140215 | 22 | 19524336 | 21609394 | 21130346 | 24521910 | 27894410 | 16085711 | 12228056 | 14830964 | 15337555 | 14872723 | 22277502 | 18448728 | 22219588 | 21465782 | 21074276 | 18367782 | 16637139 | 18797420 | 21723972 | 17151604 |
| P70460 | Vasp | Vasodilator-stimulated phosphoprotein | 244.15 | 12.53 | 39667 | 5 | 1923810 | 1188995.38 | 2 | 2 | 35870.67 | 2173413 | 3868925 | 192777.05 | 5663243 | 781271.62 | 6754431 | 6023641 | 2817622.5 | 5124684.5 | 2814825.5 | 2697295.75 | 1448203.62 | 1583815.25 | 1927797.75 | 2 |
| Q62465 | Vat1 | Synaptic vesicle membrane protein VAT-1 homol | 517.22 | 28.08 | 43097 | 8 | 4966907.5 | 13935714 | 6588158 | 1783811.75 | 6689598.5 | 14454152 | 11999288 | 16205762 | 5624263.5 | 4175210 | 10147227 | 10425886 | 6813787 | 6888659.5 | 2635201.5 | 4805264.5 | 4477725 | 4156765 | 5735903.5 | 1772025.12 |
| P27870 | Vav1 | Proto-oncogene vav | 327.13 | 6.86 | 98137 | 5 | 2030993.38 | 123203.7 | 1096799.38 | 483896.84 | 5240591.5 | 2564219.5 | 1273896 | 3815833 | 4057266 | 3686113.5 | 8375784 | 7800287 | 5108799 | 7267738.5 | 2426885 | 7590874 | 3485959.5 | 2086052.25 | 6409591 | 2696153.25 |
| Q9R0C8 | Vav3 | Guanine nucleotide exchange factor VAV3 | 69.02 | 2.13 | 97968 | 2 | 2 | 391021.97 | 2 | 2 | 2 | 2 | 2 | 2 | 2 | 2 | 1263986.38 | 1219317.62 | 1056342.88 | 1083186.5 | 1004115.25 | 914945.25 | 902887.25 | 1268717.38 | 1057711.38 | 915242.56 |
| P85972 | Vcl | Vinculin | 931.9 | 18.86 | 116615 | 17 | 17905044 | 19264266 | 15691141 | 6392768.5 | 10005696 | 12964626 | 13454907 | 14801484 | 6552724.5 | 5082459 | 21437442 | 20363428 | 13098348 | 14210775 | 13591054 | 16175015 | 12960442 | 13653963 | 10652252 | 13336659 |
| P46462 | Vcp | Transitional endoplasmic reticulum ATPase | 1885.57 | 41.32 | 89349 | 33 | 12293020 | 15616319 | 19083164 | 12386286 | 15454222 | 12946910 | 12670348 | 17911234 | 15616610 | 20345178 | 35687142 | 36593034 | 27202898 | 29641356 | 30362014 | 24926682 | 17162066 | 18566426 | 20345566 | 12155621 |
| Q9Z2L0 | Vdac1 | Non-selective voltage-gated ion channel VDAC1 | 1047.27 | 70.67 | 30756 | 15 | 20719656 | 17122736 | 12637907 | 21559252 | 33112320 | 24387052 | 22529228 | 37859574 | 34143266 | 36184926 | 17082604 | 14678875 | 25706552 | 20915906 | 15964904 | 19649004 | 19428222 | 23187606 | 24482472 | 19740802 |
| P81155 | Vdac2 | Non-selective voltage-gated ion channel VDAC2 | 707.41 | 38.31 | 31746 | 11 | 23105672 | 46989590 | 25547970 | 38436154 | 76305778 | 53345186 | 59496650 | 83714178 | 95374634 | 67764914 | 24504088 | 24641670 | 40354502 | 40945450 | 26978060 | 26986850 | 22578252 | 28459086 | 29113230 | 24335104 |
| Q60931 | Vdac3 | Non-selective voltage-gated ion channel VDAC3 | 575.81 | 28.62 | 30753 | 9 | 3412320 | 4777706.5 | 4080942.75 | 6615699.5 | 18338338 | 9830086 | 11261620 | 20417768 | 21442990 | 17631068 | 6498904 | 6998411 | 8499832 | 7745507.5 | 4553143 | 6577643.5 | 4624134 | 5983393 | 8791840 | 5957651 |
| P20152 | Vim | Vimentin | 3751.201 | 80.69 | 53688 | 58 | 292444354 | 261473458 | 320387522 | 246949666 | 257421506 | 395601250 | 419176482 | 605743682 | 420862754 | 458444546 | 605814594 | 542833026 | 570076226 | 563079618 | 409487650 | 522059714 | 462530306 | 352727650 | 392223234 | 342554082 |
| Q6TEK4 | Vkorc1 | Vitamin K epoxide reductase complex subunit 1 | 52.28 | 8.07 | 17783 | 1 | 2 | 481981.34 | 2 | 2 | 2 | 2 | 2 | 530614.94 | 384405.5 | 2 | 476035.5 | 2 | 2 | 2 | 2 | 2 | 915630.31 | 2 | 2 | 2 |

| | | | | | | | | | | | | | | | | | | | | | | | | | | |
|---|---|---|---|---|---|---|---|---|---|---|---|---|---|---|---|---|---|---|---|---|---|---|---|---|---|---|
| Q78T54 | Vma21 | Vacuolar ATPase assembly integral membrane p | 76.61 | 14.85 | 11365 | 2 | 2 | 2 | 2 | 2 | 420753.47 | 2 | 2 | 554734.81 | 2 | 2 | 646886.06 | 2 | 797047.19 | 375663.69 | 2 | 796094.25 | 712968.56 | 556954.12 | 145129.8 | 543691.19 |
| Q8BX70 | Vps13c | Intermembrane lipid transfer protein VPS13C | 479.16 | 3.5 | 420089 | 9 | 4514338 | 5282047.5 | 6020406.5 | 4872832.5 | 3780230 | 3147954.5 | 4148494 | 5005071 | 3372048 | 3378204 | 928694.62 | 980428.62 | 3563590 | 3125135.5 | 1476465.62 | 2935037.25 | 3136278.75 | 3676333.25 | 2859568 | 3263125.25 |
| Q920Q4 | Vps16 | Vacuolar protein sorting-associated protein 16 ho | 41.23 | 1.79 | 94928 | 1 | 1068746.38 | 878823.06 | 443587.72 | 2 | 2 | 2 | 792963.31 | 849896.5 | 2 | 1213726 | 2 | 2 | 606112.69 | 2 | 925876.38 | 2 | 2 | 728922.81 | 765905.56 | 2 |
| Q8C0E2 | Vps26b | Vacuolar protein sorting-associated protein 26B | 38.26 | 3.27 | 39125 | 1 | 2 | 2 | 2 | 2 | 2 | 2 | 2 | 2 | 2 | 2 | 410815.38 | 964041.44 | 1269732.75 | 2 | 755709.19 | 2 | 2 | 2 | 2 | 2002134.12 |
| Q63615 | Vps33a | Vacuolar protein sorting-associated protein 33A | 56.4 | 1.17 | 67514 | 1 | 2 | 2 | 2 | 885821.06 | 2 | 2 | 648177.19 | 2 | 691110.25 | 613854.38 | 801215 | 2 | 2 | 2 | 2 | 2 | 2 | 2 | 2 | 2 |
| Q9EQH3 | Vps35 | Vacuolar protein sorting-associated protein 35 | 223.64 | 6.03 | 91713 | 4 | 4160166.5 | 6535273.5 | 6433009 | 4870731.5 | 7834345 | 6543879 | 5818917 | 8761752 | 5547712.5 | 7842438 | 6586465 | 6561861 | 5284844 | 4408705 | 780376.44 | 5656460 | 3212909.5 | 2 | 2513185.5 | 2173468.25 |
| Q5XI83 | Vps35l | VPS35 endosomal protein-sorting factor-like | 43.91 | 1.28 | 106151 | 1 | 2 | 2 | 2 | 470264.88 | 2 | 2 | 2 | 2 | 307814.94 | 2 | 2 | 2 | 2 | 569865 | 2 | 2 | 2 | 2 | 2 | 2 |
| P0C0A2 | Vps36 | Vacuolar protein-sorting-associated protein 36 | 59.45 | 3.37 | 43718 | 1 | 2 | 724855.06 | 1195357.62 | 2 | 807086.19 | 167936.69 | 2 | 744345.75 | 545118.5 | 848682.88 | 2 | 2 | 555991.06 | 584125.44 | 2 | 2 | 2 | 2 | 2 | 2 |
| P46467 | Vps4b | Vacuolar protein sorting-associated protein 4B | 117.84 | 6.98 | 49419 | 2 | 2 | 2 | 90458.2 | 153711.06 | 62314.73 | 2 | 2 | 2 | 88617.08 | 107935.16 | 105132.84 | 72083.08 | 143555.25 | 252846.44 | 2 | 2 | 257327.25 | 205972.27 | 81412.26 | 173847.72 |
| Q3UVL4 | Vps51 | Vacuolar protein sorting-associated protein 51 ho | 98.49 | 2.05 | 86187 | 2 | 1697669.62 | 767898.19 | 963566.38 | 1374155.38 | 507171.47 | 1712706.5 | 1619790.5 | 66624.07 | 1354809.25 | 394271.75 | 1612219.75 | 2 | 1689434.5 | 876752.69 | 484212 | 2 | 278860.34 | 230234.14 | 147337.56 | 1012739.38 |
| O55166 | Vps52 | Vacuolar protein sorting-associated protein 52 ho | 55 | 0.97 | 82103 | 1 | 1250888.88 | 1759836.38 | 39019202 | 9012660 | 44151610 | 7279955 | 6845841 | 134425.69 | 4970698 | 37150062 | 5150100.5 | 4563665 | 3955081 | 3988011.5 | 38816802 | 4887992 | 5508968 | 6445389.5 | 56389806 | 54406870 |
| Q8BN21 | Vrk2 | Serine/threonine-protein kinase VRK2 | 87.63 | 5.57 | 58119 | 2 | 2 | 2 | 72151.66 | 75994.93 | 22341.23 | 2 | 2 | 32518.95 | 59150.24 | 2 | 2 | 2 | 41439.02 | 54810.65 | 2 | 34829.79 | 2 | 2 | 2 | 2 |
| Q9CR26 | Vta1 | Vacuolar protein sorting-associated protein VTA1 | 36.64 | 4.53 | 33913 | 1 | 2 | 2 | 47218.61 | 2 | 103176.2 | 2 | 2 | 2 | 540592.12 | 62846.04 | 2 | 2 | 2 | 2 | 2 | 2 | 2 | 2 | 2 | 150731.33 |
| O88384 | Vti1b | Vesicle transport through interaction with t-SNAR | 138.29 | 14.22 | 26713 | 3 | 2 | 2 | 2049213.88 | 1671352.75 | 1688848.75 | 2968562 | 2268710.25 | 4892896 | 2754624.25 | 1358659.88 | 2548476.25 | 1558878.62 | 1852908.12 | 1936545.62 | 1718052.62 | 2387651.5 | 2 | 2740612 | 3556625.25 | 2490006 |
| P29788 | Vtn | Vitronectin | 101.99 | 3.77 | 54849 | 1 | 2 | 2 | 140723.03 | 2181160 | 1180703 | 479473.09 | 350494.72 | 2 | 1622391.88 | 1154566.5 | 187975.98 | 2 | 2 | 2 | 385665.97 | 166799.61 | 241459.16 | 2 | 2 | 2 |
| Q99KC8 | Vwa5a | von Willebrand factor A domain-containing protei | 836.66 | 18.41 | 87143 | 15 | 31166170 | 23750970 | 40100890 | 28371514 | 33655334 | 27335032 | 36410474 | 29402400 | 36336762 | 31120206 | 27361118 | 29996660 | 27833720 | 27267988 | 38118198 | 27245976 | 27755876 | 17465592 | 30365554 | 17939906 |
| P32921 | Wars1 | Tryptophan--tRNA ligase, cytoplasmic | 239.9 | 14.55 | 54358 | 5 | 2478380 | 166607.94 | 2427810 | 1017236.12 | 3693346 | 1664800.88 | 1919538.5 | 651562.56 | 2500688 | 4592180 | 3933457.5 | 4300569 | 5039679.5 | 3440914 | 7650628 | 2307206 | 2936172 | 2754301 | 1356623.5 | 4536018 |
| P70315 | Was | Actin nucleation-promoting factor WAS | 38.87 | 2.5 | 54192 | 1 | 1146980.62 | 1414517.62 | 2 | 1313357.25 | 2 | 2 | 1002377.12 | 148664.2 | 2 | 2 | 851555.38 | 2 | 878550.62 | 768603.69 | 1058752.75 | 2 | 1251935.62 | 1308058.75 | 2 | 1405532.12 |
| Q8BH43 | Wasf2 | Actin-binding protein WASF2 | 424.01 | 15.49 | 54074 | 8 | 3370600.5 | 2098519.75 | 2583947 | 2890427.5 | 2282942.5 | 3271613.75 | 3407723.75 | 3043761.5 | 6061664 | 4171569.5 | 8880675 | 9974627 | 6038283.5 | 8691620 | 3442388.5 | 7164465.5 | 6117966.5 | 5017807 | 6840514 | 3204721.75 |
| Q8C2E7 | Washc5 | WASH complex subunit 5 | 249.46 | 5.78 | 134110 | 5 | 801891.56 | 200280.59 | 2134311.75 | 200085.62 | 552541.56 | 2 | 620930.88 | 117899.64 | 534204.5 | 1264833.38 | 2 | 485460 | 417523.94 | 222496.09 | 667274.12 | 2 | 2 | 711206.38 | 388030.12 | 2 |
| P97765 | Wbp2 | WW domain-binding protein 2 | 80.96 | 7.66 | 28032 | 2 | 2104615 | 2897506 | 1275574.75 | 3839242.5 | 2293723.75 | 2083332.75 | 3510466.5 | 3805786.25 | 2005839.5 | 1751749.88 | 1380377 | 2650578.25 | 1530638.5 | 2229338.5 | 2026368.12 | 1309815 | 2004305.75 | 1244937.5 | 3028459.75 | 1501436.75 |
| O88342 | Wdr1 | WD repeat-containing protein 1 | 765.62 | 21.45 | 66407 | 13 | 17124174 | 26997010 | 47996186 | 18269048 | 70113834 | 57356818 | 40543318 | 43077626 | 54174802 | 46679986 | 78924010 | 92804290 | 83290738 | 79082266 | 69264234 | 82308970 | 70261386 | 75680354 | 79777826 | 68370818 |
| Q6ZQL4 | Wdr43 | WD repeat-containing protein 43 | 282.44 | 10.19 | 75381 | 6 | 3837127.5 | 2 | 4017551.75 | 2280892 | 1159179.75 | 3901545.5 | 267946.28 | 3177350.5 | 493760.72 | 1776609.5 | 4267015.5 | 4580015 | 3271147.5 | 2664020 | 4304687.5 | 2527068.25 | 4258164.5 | 3575343.25 | 689558.19 | 2 |
| Q9Z0H1 | Wdr46 | WD repeat-containing protein 46 | 134.04 | 6.27 | 69048 | 3 | 557609.94 | 729993.31 | 2 | 125850.68 | 289050.31 | 2 | 809997.56 | 716635.12 | 1435551.75 | 557953.19 | 912670.25 | 131499.52 | 710846.81 | 96826.81 | 399394.47 | 2 | 215548.3 | 2 | 84829.59 | 271165.78 |
| D4A929 | Wdr81 | WD repeat-containing protein 81 | 35.35 | 0.41 | 212260 | 1 | 2 | 2 | 2 | 558561.81 | 2 | 2 | 329616.09 | 94016.45 | 957912.62 | 514395.31 | 1144185.25 | 1515788.5 | 615683.44 | 979971.44 | 2 | 2 | 2 | 2 | 2 | 2 |
| Q8BFQ4 | Wdr82 | WD repeat-containing protein 82 | 133.47 | 12.14 | 35079 | 3 | 1300087.12 | 1337366.62 | 1196204.88 | 1336498.88 | 814064.56 | 2 | 843349.69 | 757896.94 | 542067.25 | 618958.19 | 694855.19 | 891421.56 | 1082398.62 | 561463.88 | 735788.5 | 947253.5 | 1006026.56 | 1134479.88 | 700055 | 2 |
| B2RYI0 | Wdr91 | WD repeat-containing protein 91 | 38.1 | 1.61 | 83146 | 1 | 2 | 2 | 2 | 2 | 480429.78 | 2 | 2 | 2 | 453154.69 | 2 | 154218.62 | 629994.5 | 2 | 469864.38 | 2 | 2 | 2 | 2 | 2 | 2 |
| Q6IN36 | Wipf1 | WAS/WASL-interacting protein family member 1 | 41.01 | 2.26 | 49751 | 1 | 1370946 | 2250732.5 | 700481.12 | 2038972.5 | 1951090.88 | 2850400.75 | 1944816.75 | 2715911.75 | 1820580.75 | 1157167.62 | 2974309 | 2969445.25 | 2151387 | 2334982.25 | 2234343.75 | 2092726.75 | 2674210.5 | 1970919.62 | 2707127.5 | 2351137.75 |
| P83741 | Wnk1 | Serine/threonine-protein kinase WNK1 | 202.05 | 1.89 | 250934 | 4 | 32873580 | 31837016 | 100448586 | 72371858 | 19596188 | 90396058 | 108785850 | 91150402 | 24001082 | 45253306 | 53945358 | 61499862 | 66028462 | 45362642 | 756910 | 948807.69 | 140757890 | 110021834 | 97187090 | 171914914 |
| Q00519 | Xdh | Xanthine dehydrogenase/oxidase | 292.57 | 4.94 | 146562 | 5 | 1106385 | 2 | 2831955 | 387913.75 | 865343.62 | 2 | 154562.33 | 2 | 982236.62 | 782681.62 | 1424212.38 | 1536609.25 | 2123709.75 | 387795.06 | 2 | 2 | 1551635.5 | 2 | 2 | 204658.69 |
| Q6P1B1 | Xpnpep1 | Xaa-Pro aminopeptidase 1 | 561.21 | 18.62 | 69591 | 9 | 244982.06 | 371159.75 | 1846912.88 | 2 | 2 | 596025.69 | 2 | 2 | 2 | 1408067.75 | 2 | 535811.19 | 2 | 748717.12 | 1659998.75 | 447680.97 | 2 | 818878.19 | 2 | 2 |
| B7ZMP1 | Xpnpep3 | Xaa-Pro aminopeptidase 3 | 35.84 | 1.58 | 56677 | 1 | 2 | 2 | 2 | 2 | 2 | 2 | 2 | 2 | 2 | 2 | 2 | 2 | 2 | 2 | 2 | 2 | 2 | 365209.69 | 2 | 2 |
| Q6P5F9 | Xpo1 | Exportin-1 | 412.12 | 9.24 | 123093 | 7 | 4224522.5 | 999019.31 | 4562625 | 956041.5 | 2420815 | 168133.39 | 957116.88 | 147110.22 | 1333649.88 | 1996166.88 | 1254298.25 | 1083727.5 | 2245244 | 1314772.75 | 1692242 | 666561.81 | 1029138.5 | 1656922 | 1626606.38 | 229519.2 |
| Q924C1 | Xpo5 | Exportin-5 | 58.86 | 1.08 | 136973 | 1 | 1297401.25 | 1870565 | 267253.84 | 1279270.25 | 180101.09 | 833950.56 | 637784.5 | 2 | 407449.28 | 598702.56 | 619725.5 | 665835.12 | 1118584.25 | 947334.62 | 825820.25 | 2 | 2 | 903670.75 | 2 | 1058600.25 |
| Q60596 | Xrcc1 | DNA repair protein XRCC1 | 151.6 | 6.34 | 68971 | 3 | 2 | 2606718.5 | 2 | 2 | 705847 | 2 | 2 | 2 | 1714839.75 | 1113945.5 | 2 | 542971.62 | 656656.25 | 1668957.5 | 2 | 679465.19 | 2 | 2 | 926186.62 | 2 |
| P27641 | Xrcc5 | X-ray repair cross-complementing protein 5 | 59.23 | 1.5 | 83057 | 1 | 896016.38 | 967633.56 | 855055.38 | 2 | 1034765.19 | 790070.38 | 619571.06 | 1377583.62 | 537421.88 | 2 | 652307.62 | 1008125.38 | 1171143 | 1019721.44 | 811720.06 | 1551023 | 854352.56 | 765946.81 | 2 | 2 |
| Q9DBR1 | Xrn2 | 5'-3' exoribonuclease 2 | 501.3 | 14.93 | 108687 | 10 | 1815590.5 | 542564.56 | 792709.88 | 1009718.75 | 1195258.88 | 816235.38 | 836987.25 | 603365.38 | 876907.06 | 1280698.62 | 2880143.75 | 2545775.75 | 2813205.25 | 2690584.75 | 2400325.25 | 280504 | 896167.88 | 731851.62 | 2829287.75 | 1357322.62 |
| Q3U4G3 | Xxylt1 | Xyloside xylosyltransferase 1 | 33.17 | 2.04 | 43839 | 1 | 2 | 2 | 2 | 2 | 2 | 2 | 2 | 2 | 642839.94 | 422723.69 | 2 | 2 | 2 | 2 | 734117.81 | 2 | 2 | 580176.94 | 303192.88 | 783617.62 |
| Q91WQ3 | Yars1 | Tyrosine--tRNA ligase, cytoplasmic | 368.36 | 15.53 | 59105 | 8 | 1139934.25 | 992075.88 | 2459003.25 | 1051232.12 | 6185094 | 1813154.75 | 2 | 5121067 | 1493248.75 | 3302378 | 10694529 | 12851796 | 5117106 | 6350279 | 2141733.5 | 10261517 | 2357200.5 | 1395796.75 | 8829746 | 3969385.5 |
| Q8BYL4 | Yars2 | Tyrosine--tRNA ligase, mitochondrial | 50.72 | 2.54 | 52597 | 1 | 2 | 2 | 2 | 2 | 2 | 2 | 2 | 2 | 2 | 149161.28 | 2 | 2 | 49530.05 | 76769.97 | 184617.41 | 43205.73 | 2 | 2 | 2 | 2 |
| P62960 | Ybx1 | Y-box-binding protein 1 | 102.39 | 5.9 | 35730 | 1 | 766657.31 | 1014144.19 | 636797.25 | 1295894.62 | 1304048.5 | 2 | 2 | 1650829.25 | 1137573.62 | 784355.19 | 3438250 | 2417723.25 | 1900699.75 | 2483723.25 | 3043710.25 | 2601958.75 | 2873486 | 2996822.5 | 2537314.5 | 2327003.75 |
| Q62764 | Ybx3 | Y-box-binding protein 3 | 82.68 | 5.26 | 38852 | 1 | 2 | 2 | 2 | 2 | 2 | 2 | 2 | 2 | 2 | 2 | 986348.56 | 849299.38 | 2 | 756843.06 | 2 | 1396294.88 | 831506.62 | 2 | 799381.31 | 2 |
| Q5EGY4 | Ykt6 | Synaptobrevin homolog YKT6 | 127.31 | 15.15 | 22369 | 2 | 2 | 54795.59 | 106604.62 | 405742.75 | 784542.25 | 2 | 237846.06 | 2 | 952026.88 | 746429 | 284590.38 | 221763.53 | 932757.56 | 176520.34 | 238267.31 | 2 | 231912.47 | 209944.58 | 454962.88 | 274335.44 |
| Q8BYK6 | Ythdf3 | YTH domain-containing family protein 3 | 31.55 | 1.71 | 63961 | 1 | 2 | 2 | 2 | 434283.72 | 2 | 2 | 2 | 2 | 2 | 2 | 2 | 2 | 2 | 283558.91 | 2 | 2 | 2 | 2 | 2 | 2 |
| Q9CQV8 | Ywhab | 14-3-3 protein beta/alpha | 708.81 | 41.46 | 28086 | 12 | 4896747.5 | 3315442.75 | 5708252 | 6802657 | 8317940 | 7370293 | 7398489 | 2326258.75 | 8924247 | 8336892.5 | 12719067 | 18032744 | 16987800 | 16453269 | 13922475 | 8768146 | 9115824 | 12013244 | 10611480 | 8129581.5 |
| P62259 | Ywhae | 14-3-3 protein epsilon | 988.03 | 53.33 | 29174 | 17 | 114669714 | 122373274 | 159577314 | 137712546 | 77329946 | 101115922 | 114434794 | 128815538 | 71749898 | 100758994 | 99199394 | 94093698 | 124206226 | 91268674 | 128254906 | 89204034 | 102284898 | 155913042 | 101566922 | 86468354 |
| P61982 | Ywhag | 14-3-3 protein gamma | 640.05 | 33.2 | 28303 | 12 | 17779614 | 15049505 | 25209882 | 16436428 | 41270894 | 35799838 | 36235546 | 58032390 | 39876250 | 33154942 | 38128778 | 35429578 | 29460106 | 29596796 | 21251668 | 34757330 | 16238926 | 26062562 | 30503304 | 13772258 |
| P68510 | Ywhah | 14-3-3 protein eta | 538.31 | 33.33 | 28212 | 11 | 3971312 | 1535579.12 | 7051150 | 4817204 | 7762490 | 2451334.5 | 6106909 | 10808498 | 6338970 | 8260133.5 | 5070344 | 3285806.5 | 5298485 | 3359560.5 | 8181245 | 1657864.88 | 2427853.5 | 5427459 | 1892001.5 | 1394993.25 |
| P68254 | Ywhaq | 14-3-3 protein theta | 496.68 | 37.55 | 27778 | 9 | 10289632 | 9641549 | 15523674 | 11566198 | 11149067 | 6307019 | 9657434 | 5007231 | 7935234.5 | 6942308 | 8545501 | 6814398.5 | 10516723 | 7285512.5 | 14208925 | 4437912 | 6895788 | 13202230 | 5233097.5 | 5895509.5 |
| P63101 | Ywhaz | 14-3-3 protein zeta/delta | 876.86 | 48.57 | 27771 | 14 | 47919178 | 58831214 | 49123194 | 74235818 | 40057706 | 82341122 | 90425002 | 27328360 | 95403346 | 72581258 | 93968882 | 108277210 | 90550914 | 98461554 | 70538362 | 67332882 | 61610174 | 62501810 | 49181578 | 56384266 |
| Q6NZF1 | Zc3h11a | Zinc finger CCCH domain-containing protein 11A | 95 | 3.41 | 86492 | 2 | 2 | 2 | 2 | 2 | 2 | 2 | 2 | 2 | 2 | 2 | 2 | 2 | 2 | 2 | 2 | 2 | 2 | 2 | 2 | 2 |
| Q3UPF5 | Zc3hav1 | Zinc finger CCCH-type antiviral protein 1 | 102.74 | 2.54 | 106688 | 2 | 2 | 2 | 2 | 2 | 291717.75 | 2 | 2 | 2 | 2 | 2 | 2 | 2 | 2 | 2 | 2 | 2 | 2 | 2 | 2 | 2 |
| O88532 | Zfr | Zinc finger RNA-binding protein | 221.9 | 4.75 | 116859 | 4 | 2634694.75 | 1055543.88 | 579595.94 | 594348.31 | 1705821.25 | 1140386 | 801972 | 431618.44 | 1103626.62 | 953083.06 | 1059644.62 | 1139548.12 | 800224.12 | 1320556 | 448744.69 | 2105222.25 | 1839948.88 | 2641030.75 | 1573500 | 2753850.75 |
| Q80W54 | Zmpste24 | CAAX prenyl protease 1 homolog | 104.35 | 4.84 | 54735 | 2 | 1603798.12 | 1458197.75 | 867637.31 | 771598.88 | 816229 | 679312.81 | 128333.26 | 79235.12 | 63194.96 | 824566.06 | 2 | 51776.03 | 825790.25 | 895249.31 | 71154.18 | 2 | 936863.06 | 1482997 | 1492643.62 | 2097938.75 |
| Q62523 | Zyx | Zyxin | 244.44 | 9.4 | 60546 | 5 | 2 | 2 | 2 | 946016.38 | 2138850 | 1706564.25 | 1544394 | 1459376.5 | 4495694 | 1931477 | 7840169 | 5299049.5 | 2097428.5 | 4544459.5 | 590679.5 | 3398796.25 | 1992089 | 631285.25 | 2674146.5 | 565602.62 |
| Q5U2Q3 | | Ester hydrolase C11orf54 homolog | 85.35 | 5.4 | 34993 | 2 | 849882.38 | 694573.5 | 1038647.5 | 642729.19 | 273197.19 | 479805.91 | 946919.62 | 483768.06 | 1243827.25 | 934624.88 | 279104.12 | 276299.66 | 854451.25 | 391201.69 | 653452.25 | 167286.12 | 669188.25 | 255772.12 | 174940.73 | 354990.09 |
| Q6PIU9 | | Uncharacterized protein FLJ45252 homolog | 34.73 | 3.39 | 37705 | 1 | 1015345.62 | 2102285.5 | 2 | 977497.88 | 1441690.88 | 1523316.5 | 2 | 2183049.25 | 1808990.5 | 1123578 | 1938186.38 | 2120641.5 | 609306.38 | 2 | 2 | 2 | 2 | 2 | 4379947.5 | 2 |
