## supplemental tables 2 to 4 for "Effects of manganese dioxide on macrophages under different exposure schemes"

Supplementary table 2: List of modulated proteins in the recovery condition, selected by the Mann Whitney U test (p<0.05)

| accession number | gene_name | description +MWU PBK14 20 recov | protein_set_score | coverage | MW | #peptides | Ctrl 1 | Ctrl 2 | Ctrl 3 | Ctrl 4 | Ctrl 5 | PBK14 20 recov 1 | PBK14 20 recov 2 | PBK14 20 recov 3 | PBK14 20 recov 4 | PBK14 20 recov 5 | PBK14 4x5 rep1 | PBK14 4x5 rep2 | PBK14 4x5 rep3 | PBK14 4x5 rep4 | PBK14 4x5 rep5 | PBK14 4x10 rep1 | PBK14 4x10 rep2 | PBK14 4x10 rep3 | PBK14 4x10 rep4 | PBK14 4x10 rep5 | ratio PBK14/20 recov/ctrl | U PBK14/20 recov | ratio_PBK14 4x5 repet/ctrl | U_PBK14 4x5 repet | ratio Pbk 14 4x10/ctrl | U Pbk 14 4x10 |
|---|---|---|---|---|---|---|---|---|---|---|---|---|---|---|---|---|---|---|---|---|---|---|---|---|---|---|---|---|---|---|---|---|
| P35283 | Rab12 | Ras-related protein Rab-12 | 158.28 | 12.35 | 27329 | 3 | 2626988.25 | 2274646.5 | 1479324.88 | 1650430.88 | 1640128.75 | 2 | 2 | 2 | 2 | 2 | 2 | 2 | 2 | 2 | 2 | 2583009.5 | 2 | 1630908.12 | 2 | 1270759.38 | 1.03396E-06 | 0 | 1.03396E-06 | 0 | 0.567096115 | 5 |
| P35433 | Ppat | Amidophosphoribosyltransferase | 37.89 | 1.35 | 57437 | 1 | 1384687.88 | 1062445.5 | 918004.81 | 862753.44 | 1043741.19 | 2 | 2 | 2 | 2 | 2 | 2 | 2 | 682479.81 | 586809.25 | 2 | 2 | 580160.31 | 2 | 2 | 2 | 1.89695E-06 | 0 | 0.240778351 | 0 | 0.110054765 | 0 |
| P63028 | Tpt1 | Translationally-controlled tumor protein | 159.35 | 21.51 | 19462 | 3 | 632496.94 | 1163805.38 | 2201839.5 | 821358.81 | 1260006.12 | 2 | 2 | 69912.64 | 2 | 278445.5 | 2 | 2 | 888755.5 | 627035 | 690880.12 | 256722.27 | 764707.69 | 788575.75 | 2 | 2 | 0.057301382 | 0 | 0.362969351 | 3 | 0.297723119 | 2 |
| Q8CG48 | Smc2 | Structural maintenance of chromosomes protein 2 | 168.97 | 4.03 | 134239 | 4 | 791365.06 | 642979.38 | 270799.66 | 475365.22 | 1527073.88 | 2 | 135077.23 | 137344.39 | 2 | 2 | 493528.97 | 591335.75 | 2 | 567432.62 | 119463.78 | 133863.14 | 2 | 615069.44 | 2 | 2 | 0.073478491 | 0 | 0.477875485 | 6 | 0.202001827 | 2 |
| Q8BRT1 | Clasp2 | CLIP-associating protein 2 | 114.8 | 1.79 | 140739 | 3 | 702967.44 | 1062777.38 | 647279.88 | 502298.75 | 780302.75 | 2 | 142333.39 | 62377.88 | 142578.33 | 2 | 2 | 2 | 2 | 237896.53 | 635456.62 | 568163.38 | 2 | 737242.12 | 2 | 670829.88 | 0.093974223 | 0 | 0.236322372 | 1 | 0.534750885 | 6 |
| Q01320 | Top2a | DNA topoisomerase 2-alpha | 397.87 | 6.15 | 172790 | 8 | 3515953 | 3962353.5 | 3390934.5 | 4701950 | 4132424 | 2 | 759808.94 | 2 | 1798711.75 | 686417.06 | 2489620.75 | 5515715.5 | 7446122 | 6455125 | 3730762.25 | 1189278 | 3645835.5 | 2032580.88 | 4569197 | 1532961.75 | 0.164687635 | 0 | 1.301149332 | 8 | 0.658247389 | 6 |
| Q9D338 | Mrpl19 | Large ribosomal subunit protein bL19m | 176.21 | 12.67 | 33578 | 4 | 575289.62 | 1003186.94 | 502012.88 | 769077.5 | 832785.31 | 139775.58 | 2 | 2 | 227855.81 | 277760.75 | 2 | 2 | 100854.18 | 292543.81 | 2 | 640350.12 | 2 | 763403.31 | 702465.81 | 723937.06 | 0.175267355 | 0 | 0.106834969 | 0 | 0.768573485 | 8 |
| Q8CG47 | Smc4 | Structural maintenance of chromosomes protein 4 | 185.01 | 2.95 | 146895 | 4 | 5528178 | 7201362 | 4967845.5 | 7203776 | 6178335 | 72138.85 | 439674.94 | 2917331.25 | 1182578.25 | 2097384.5 | 1343523.38 | 2011011.75 | 4972314 | 3593079 | 3181670.75 | 2678003.5 | 2090249.25 | 3940062 | 5139125 | 4749610 | 0.215869256 | 0 | 0.485902302 | 1 | 0.598370368 | 1 |
| P18395 | Csde1 | Cold shock domain-containing protein E1 | 177.32 | 7.89 | 88895 | 4 | 2532412.75 | 1696915 | 1335742.12 | 1716086 | 2368621.5 | 433744.25 | 506747.19 | 568499.69 | 490966.91 | 790579.75 | 2614241.25 | 3138766.5 | 3546241.5 | 3071236.75 | 1294135 | 1863632.25 | 1967614.12 | 2669362 | 1299263.25 | 2578365.75 | 0.289181572 | 0 | 1.416055571 | 5 | 1.075489824 | 9 |
| Q3UZA1 | Rcsd1 | CapZ-interacting protein | 55.4 | 2.91 | 44114 | 1 | 1868649.12 | 1271674.5 | 1164003.5 | 1720249.62 | 1597736.25 | 949748.06 | 2 | 2 | 1048017.81 | 621006.5 | 1710370.25 | 1688982.5 | 1139033.25 | 970324.25 | 1415096 | 2 | 663504 | 1059547.25 | 2207542.5 | 1637489.12 | 0.343567153 | 0 | 0.908360265 | 8 | 0.730498062 | 8 |
| P49718 | Mcm5 | DNA replication licensing factor MCM5 | 454.31 | 16.21 | 82407 | 11 | 14334609 | 23994526 | 12389125 | 23668398 | 18101840 | 5167847 | 2130634 | 10794580 | 8285995.5 | 7981872 | 8040220 | 7604583 | 8059879 | 10139252 | 6163571.5 | 6093196 | 5124019 | 2573817.75 | 7862198.5 | 9596300 | 0.371515694 | 0 | 0.432567361 | 0 | 0.337874784 | 0 |
| O35737 | Hnrnph1 | Heterogeneous nuclear ribonucleoprotein H | 637.01 | 31.85 | 49199 | 10 | 949454.06 | 1723862.25 | 1234388.62 | 1692359.75 | 2521066 | 879649.06 | 700714.12 | 202986.61 | 618831.38 | 663010.88 | 1988888.75 | 2226868 | 2310739.5 | 1495003.5 | 2023818.75 | 1448498 | 287894.69 | 987214 | 454018.28 | 2 | 0.377434149 | 0 | 1.236935951 | 7 | 0.391278887 | 3 |
| Q9QY81 | Nup210 | Nuclear pore membrane glycoprotein 210 | 439.78 | 7.37 | 204101 | 9 | 2703440.75 | 5567009.5 | 3143422 | 4788574.5 | 3050809.5 | 1240138.88 | 894893.5 | 1861990 | 1444105.25 | 1907624.88 | 2759881.25 | 1597330.38 | 2182414.75 | 2765088.25 | 2090787.5 | 1835450.38 | 3898544 | 2243327.75 | 2196633.25 | 777138.12 | 0.381688812 | 0 | 0.591874018 | 2 | 0.56879176 | 3 |
| P46061 | Rangap1 | Ran GTPase-activating protein 1 | 481.99 | 20.54 | 63531 | 10 | 9060379 | 6495795 | 7827281 | 7722898 | 6361215 | 1926580.75 | 2586222.5 | 2292656.25 | 2893515 | 4771926 | 2518570.5 | 1173641.75 | 1092664.38 | 2043450.25 | 1185948.38 | 1744415.88 | 2525018.75 | 178639.88 | 1270313.75 | 1076775.12 | 0.386224708 | 0 | 0.213898998 | 0 | 0.181361208 | 0 |
| Q6ZPJ3 | Ube2o | (E3-independent) E2 ubiquitin-conjugating enzyme UBE2O | 40.76 | 0.78 | 140834 | 1 | 1493602 | 1744390.88 | 1423655 | 1196438 | 1860140.38 | 2 | 1018038.81 | 230029.14 | 758510.19 | 986246.12 | 1353032.25 | 888065.62 | 1169581.38 | 1371126.12 | 761636.12 | 1021396.25 | 1073916.25 | 1202322.38 | 1227189.75 | 1454170.38 | 0.387760887 | 0 | 0.718227388 | 2 | 0.774659204 | 4 |
| Q9ERK4 | Cse1l | Exportin-2 | 355.02 | 11.84 | 110455 | 9 | 13702212 | 15521284 | 9073197 | 8767833 | 11152760 | 5889202.5 | 6216690 | 1414926 | 3816168.5 | 5343307.5 | 10120087 | 2389382.25 | 9738372 | 9004469 | 10485803 | 3123076 | 9814846 | 15818800 | 10980938 | 12472340 | 0.389580072 | 0 | 0.716936775 | 7 | 0.896812675 | 12 |
| P24547 | Impdh2 | Inosine-5'-monophosphate dehydrogenase 2 | 823.83 | 26.26 | 55815 | 13 | 17446900 | 16179832 | 17789444 | 14903950 | 28514204 | 5449529 | 6618694.5 | 10749625 | 5516218.5 | 10050361 | 15306482 | 14230199 | 14013454 | 13951523 | 12153343 | 10350058 | 6980839.5 | 6417638.5 | 5090295 | 4867009 | 0.404752456 | 0 | 0.734491413 | 1 | 0.355418128 | 0 |
| O35459 | Ech1 | Delta(3,5)-Delta(2,4)-dienoyl-CoA isomerase, mitochondrial | 276.35 | 16.21 | 36118 | 4 | 8338462 | 8295967 | 7909071.5 | 5378734 | 7024098 | 1339023.5 | 2326403.5 | 3594912.5 | 4821097 | 3781058 | 5632368 | 5147330 | 7614112 | 5267336.5 | 4753896 | 4370487 | 4753312 | 5710850.5 | 4788116 | 6006393 | 0.429338812 | 0 | 0.769089665 | 3 | 0.693686132 | 2 |
| B2RXS4 | Plxnb2 | Plexin-B2 | 500.43 | 6.03 | 206230 | 9 | 10473187 | 11143706 | 9682170 | 5885522.5 | 8422308 | 2396757 | 632654.44 | 5870254 | 5480084 | 5764672.5 | 3756564 | 3685656 | 4858579 | 3459083 | 3609338.75 | 3586604.25 | 5118156 | 3967404.75 | 3771416.75 | 2162288.75 | 0.441696866 | 0 | 0.42469941 | 0 | 0.407961803 | 0 |
| Q91YE6 | Ipo9 | Importin-9 | 292.36 | 8.07 | 116052 | 5 | 5661126.5 | 5772475 | 4431324 | 4939674.5 | 5068063.5 | 2326718 | 2362680.5 | 2742037.5 | 2651297.5 | 1563304.5 | 4272772.5 | 3232113.25 | 4295584.5 | 3404548.25 | 2630744.25 | 4011597 | 3580901.75 | 3850355.5 | 3650960.25 | 2961686.75 | 0.450129072 | 0 | 0.68936709 | 0 | 0.697860166 | 0 |
| F1LNJ2 | Snrnp200 | U5 small nuclear ribonucleoprotein 200 kDa helicase | 706.67 | 8.42 | 244875 | 16 | 11609262 | 12076759 | 13563099 | 11279108 | 13219409 | 7241474.5 | 5276201 | 4889992 | 5463917 | 6162063.5 | 6777222 | 4417565 | 9723926 | 10284704 | 7422278 | 6912283 | 6573443 | 11464402 | 4919321 | 7694628.5 | 0.470198528 | 0 | 0.625541266 | 0 | 0.608348421 | 1 |
| Q9DAU1 | Cnpy3 | Protein canopy homolog 3 | 44.65 | 3.99 | 30538 | 1 | 1871318.88 | 1140381.25 | 1410310.88 | 1925868.25 | 1237352.38 | 970706.81 | 483912.25 | 910698 | 665417.5 | 624595.81 | 1629011.12 | 866482 | 1616993.88 | 850250.5 | 2102237.25 | 2045644.38 | 1906464.62 | 2136629.25 | 1598829.88 | 2199510.25 | 0.481900955 | 0 | 0.931411865 | 11 | 1.303464264 | 3 |
| P58389 | Ptpa | Serine/threonine-protein phosphatase 2A activator | 132.85 | 12.38 | 36710 | 3 | 4224244.5 | 5166914 | 5568233 | 3581562 | 3818065.25 | 3062783.25 | 2352178.25 | 153137.73 | 2468467.25 | 2775355.75 | 2148700 | 2399475.75 | 2887048 | 2655521.25 | 2379607.25 | 3122168.25 | 2554408 | 3590522.75 | 2604543.25 | 3703821 | 0.483559782 | 0 | 0.557732537 | 0 | 0.696607638 | 2 |
| P97797 | Sirpa | Tyrosine-protein phosphatase non-receptor type substrate 1 | 300.77 | 16.76 | 56414 | 5 | 26574932 | 29065286 | 28924826 | 23409346 | 19369456 | 8636036 | 13319764 | 13294156 | 12952821 | 14680769 | 16087086 | 20368016 | 16670002 | 17609162 | 16556720 | 10280642 | 17884306 | 20104260 | 18982524 | 19682646 | 0.493809069 | 0 | 0.685474711 | 1 | 0.682674356 | 2 |
| Q922D8 | Mthfd1 | C-1-tetrahydrofolate synthase, cytoplasmic | 429.61 | 12.83 | 101200 | 10 | 10855043 | 12847813 | 7876901.5 | 12099069 | 13685572 | 5068405 | 5026814.5 | 6737143 | 7560533.5 | 4759556.5 | 7925712 | 7086847 | 10204826 | 9285212 | 8244720 | 6200995 | 9569476 | 12682532 | 11670307 | 9005314 | 0.50819765 | 0 | 0.745188969 | 4 | 0.856430561 | 7 |
| Q9D906 | Atg7 | Ubiquitin-like modifier-activating enzyme ATG7 | 141.96 | 4.3 | 77520 | 3 | 4201814.5 | 3761543.75 | 4630190.5 | 3653197.25 | 3896952.5 | 93426.41 | 2989151.25 | 1120287 | 2885658.5 | 3649918.25 | 1759115.75 | 1743334.88 | 2780771.75 | 2936376.25 | 2306131 | 1619819.5 | 3279527.75 | 2674177 | 2116290 | 2440309.5 | 0.533091846 | 0 | 0.572175444 | 0 | 0.602179572 | 0 |
| B5DFN2 | Ahcyl1 | S-adenosylhomocysteine hydrolase-like protein 1 | 279.3 | 12.42 | 53753 | 6 | 3427070 | 5728816 | 3865222 | 4228606.5 | 4206938 | 3360969 | 2228797.5 | 902458.88 | 3149210.75 | 2572746.75 | 4335644.5 | 2856275.75 | 3407000 | 3273964 | 2259869.25 | 2 | 3519145.5 | 2339347.75 | 4373719 | 2 | 0.569249228 | 0 | 0.751876533 | 4 | 0.4768785 | 5 |
| P20060 | Hexb | Beta-hexosaminidase subunit beta | 353.4 | 17.72 | 61116 | 8 | 19787152 | 25215262 | 21984138 | 24970668 | 20180386 | 11218148 | 12726519 | 7381907 | 17365548 | 17356838 | 8912201 | 7231532.5 | 8537964 | 8499397 | 11826519 | 6433805 | 10468561 | 12581139 | 9981005 | 15225826 | 0.588999198 | 0 | 0.40136057 | 0 | 0.487707362 | 0 |
| P97855 | G3bp1 | Ras GTPase-activating protein-binding protein 1 | 416.46 | 21.94 | 51829 | 8 | 2828582.5 | 3678635.5 | 3172076.25 | 2826818.75 | 2906179 | 2091237.88 | 1358502.25 | 1242791.25 | 2259942.5 | 2201344 | 4617295 | 5067650.5 | 5027289 | 4846177.5 | 3790235.5 | 5474398 | 4820392.5 | 2039262.38 | 5806616.5 | 3681098 | 0.593929695 | 0 | 1.514936747 | 0 | 1.415867762 | 5 |
| Q8BG07 | Pld4 | 5'-3' exonuclease PLD4 | 943.83 | 38.57 | 56154 | 14 | 56814230 | 54365838 | 48080822 | 49538466 | 42085726 | 28968542 | 28506038 | 28419456 | 26219874 | 40881506 | 17161362 | 14213955 | 16546519 | 14629944 | 27383632 | 21365194 | 19312210 | 24940534 | 18028056 | 20254480 | 0.609822692 | 0 | 0.358472538 | 0 | 0.41413572 | 0 |
| P29416 | Hexa | Beta-hexosaminidase subunit alpha | 430.36 | 20.45 | 60613 | 9 | 11239542 | 9282127 | 9346251 | 12898934 | 8747906 | 3946593.75 | 5545161.5 | 6881814.5 | 7755726 | 7295286 | 2764089.5 | 2039271.25 | 792291.81 | 2729312.5 | 3822945.5 | 4363559.5 | 3434400 | 6897750 | 4884175 | 4994353.5 | 0.610011223 | 0 | 0.235814174 | 0 | 0.477032951 | 0 |
| O55201 | Supt5h | Transcription elongation factor SPT5 | 146.03 | 4.44 | 120664 | 3 | 1728923.75 | 1680876.25 | 3477661.25 | 1856451.38 | 1549461 | 1051832.12 | 902829.62 | 1383732.75 | 1507011.62 | 1481699.25 | 1022213.88 | 778070 | 951228.75 | 926730.62 | 1217765.75 | 1436754.38 | 1034174.06 | 1261830.25 | 3883472 | 2392161 | 0.614677519 | 0 | 0.475646681 | 0 | 0.972314039 | 9 |
| Q9Z1Q9 | Vars1 | Valine--tRNA ligase | 1205.42 | 22.41 | 140215 | 22 | 19524336 | 21609394 | 21130346 | 24521910 | 27894410 | 16085711 | 12228056 | 14830964 | 15337555 | 14872723 | 22277502 | 18448728 | 22219588 | 21465782 | 21074276 | 18367782 | 16637139 | 18797420 | 21723972 | 17151604 | 0.639647329 | 0 | 0.919824832 | 9 | 0.808140887 | 3 |
| Q8BFY9 | Tnpo1 | Transportin-1 | 297.62 | 5.01 | 102357 | 5 | 11038026 | 8185250.5 | 11575319 | 7989010 | 9471328 | 5731841 | 5967353 | 7195839 | 6718364 | 5640400 | 3683115.25 | 4458412.5 | 5869673 | 4904413.5 | 5360765 | 3546715.5 | 4544134 | 2163104 | 2871810.75 | 2622370 | 0.64762718 | 0 | 0.503044255 | 0 | 0.326325783 | 0 |
| Q9QZD8 | Slc25a10 | Mitochondrial dicarboxylate carrier | 680.98 | 36.93 | 31715 | 12 | 11981910 | 14999013 | 13635671 | 13621286 | 17404118 | 9275321 | 9050245 | 8613553 | 11941318 | 10718740 | 9006348 | 7019183.5 | 8229059.5 | 9334255 | 6532186.5 | 6385108 | 4662828.5 | 8772034 | 5495926 | 7336750 | 0.692319846 | 0 | 0.560021128 | 0 | 0.455775207 | 0 |
| P56399 | Usp5 | Ubiquitin carboxyl-terminal hydrolase 5 | 778.8 | 19.81 | 95833 | 14 | 13070915 | 16924488 | 15965610 | 12209738 | 15600410 | 11185864 | 9842009 | 7456768 | 10658277 | 12083775 | 10102161 | 12295256 | 12003999 | 10301452 | 11631137 | 10105381 | 11188433 | 16016482 | 11688334 | 11780362 | 0.694399984 | 0 | 0.763631807 | 1 | 0.823885529 | 4 |
| Q6DFW4 | Nop58 | Nucleolar protein 58 | 504.86 | 25.37 | 60343 | 9 | 7846229.5 | 10351596 | 10421583 | 8862480 | 7747698.5 | 6737081 | 5246814 | 5930571 | 7066720 | 7190264 | 3015261.75 | 3000159.5 | 6669755 | 4664555.5 | 6073845 | 5083554 | 5514721 | 7353035 | 7480814.5 | 8022738 | 0.711292146 | 0 | 0.51788173 | 0 | 0.739667654 | 2 |
| P08228 | Sod1 | Superoxide dismutase [Cu-Zn] | 509.34 | 45.45 | 15943 | 8 | 24150502 | 23647626 | 31878830 | 39941010 | 22859450 | 20888574 | 22749946 | 21974488 | 15163818 | 20919036 | 17676312 | 19743678 | 17052652 | 12425927 | 26402098 | 30140050 | 39744678 | 52103954 | 29906966 | 37129742 | 0.713768283 | 0 | 0.654845296 | 3 | 1.326704208 | 6 |
| P63017 | Hspa8 | Heat shock cognate 71 kDa protein | 3151.25 | 68.27 | 70871 | 47 | 421192898 | 431482370 | 373439234 | 421580098 | 450136930 | 284628066 | 257253202 | 317597506 | 297581762 | 362076450 | 358839650 | 359159010 | 313883746 | 310589186 | 372030082 | 269497954 | 320537218 | 317056450 | 216600722 | 301734146 | 0.724146322 | 0 | 0.817273289 | 0 | 0.679476149 | 0 |
| Q3UND0 | Skap2 | Src kinase-associated phosphoprotein 2 | 122.18 | 8.1 | 40712 | 3 | 3969949.25 | 3809952.75 | 3530172.5 | 5301210.5 | 3558785.5 | 3133936.75 | 2973437.5 | 3486063 | 2766528.5 | 2947538.5 | 1311216.75 | 1656630.25 | 1974965.38 | 1279712.75 | 1255237.88 | 2672843 | 1643412.12 | 2004220.75 | 2632103 | 2089896.12 | 0.758921703 | 0 | 0.370735591 | 0 | 0.547468339 | 0 |
| P57776 | Eef1d | Elongation factor 1-delta | 669.69 | 36.3 | 31293 | 10 | 55972342 | 56603278 | 49338410 | 54788842 | 53468642 | 42996234 | 41529478 | 47330046 | 34967330 | 43419870 | 29211538 | 30174042 | 34276694 | 32177534 | 27318584 | 26016820 | 27842364 | 25829278 | 29964362 | 31675226 | 0.778183291 | 0 | 0.566893192 | 0 | 0.523104927 | 0 |
| P61804 | Dad1 | Dolichyl-diphosphooligosaccharide--protein glycosyltransferase subunit DAD1 | 233.91 | 28.32 | 12497 | 4 | 2654903.75 | 3176288 | 4159078.5 | 4000526.75 | 3198598.5 | 4849354 | 4303510.5 | 6413080 | 4563749.5 | 4911875 | 2227366.25 | 2716564 | 3881935 | 3760852.5 | 9782006 | 2537953.5 | 5798414 | 5273270.5 | 3806530 | 6960222 | 1.456803353 | 0 | 1.301309505 | 12 | 1.418106297 | 7 |
| Q7TMB8 | Cyfip1 | Cytoplasmic FMR1-interacting protein 1 | 751.51 | 13.01 | 145241 | 15 | 6839314 | 7714240.5 | 8690557 | 7099803 | 8213055.5 | 11880539 | 9639829 | 16722741 | 12318325 | 9600170 | 16534412 | 13017145 | 15349741 | 14831595 | 13903786 | 11246515 | 11660331 | 16479392 | 11029072 | 10603626 | 1.560330182 | 0 | 1.909814983 | 0 | 1.582565642 | 0 |
| Q8VCN5 | Cth | Cystathionine gamma-lyase | 497.8 | 27.14 | 43567 | 10 | 9377385 | 11501992 | 20802036 | 13511600 | 17593928 | 23467510 | 23925028 | 22619538 | 21611462 | 23167656 | 25079238 | 26935692 | 23539544 | 22674670 | 26509684 | 26265474 | 32094046 | 23433328 | 23389266 | 24455162 | 1.577085016 | 0 | 1.713752856 | 0 | 1.781051301 | 0 |
| P20152 | Vim | Vimentin | 3751.201 | 80.69 | 53688 | 58 | 292444354 | 261473458 | 320387522 | 246949666 | 257421506 | 395601250 | 419176482 | 605743682 | 420862754 | 458444546 | 605814594 | 542833026 | 570076226 | 563079618 | 409487650 | 522059714 | 462530306 | 352727650 | 392223234 | 342554082 | 1.668142384 | 0 | 1.952083105 | 0 | 1.502959525 | 0 |
| P09103 | P4hb | Protein disulfide-isomerase | 1679.4 | 52.65 | 57058 | 29 | 50577538 | 49967894 | 54359506 | 57374046 | 68329114 | 88081394 | 96825682 | 97847258 | 109065314 | 101177146 | 89166498 | 71966010 | 92112898 | 83260802 | 61708542 | 103315386 | 107427386 | 121451682 | 129311530 | 95977914 | 1.756887266 | 0 | 1.419113535 | 1 | 1.986699251 | 0 |
| Q9Z110 | Aldh18a1 | Delta-1-pyrroline-5-carboxylate synthase | 937.92 | 22.64 | 87266 | 18 | 2298667.75 | 6509775 | 3835623.75 | 5044323 | 8296683 | 8729280 | 8873910 | 9654278 | 12390760 | 9830626 | 8831375 | 5642840 | 6711214 | 8842139 | 7579126.5 | 9527506 | 2052646.75 | 4290168 | 5805551 | 3643720.75 | 1.904126071 | 0 | 1.447242239 | 4 | 0.974389912 | 11 |
| Q9R0P3 | Esd | S-formylglutathione hydrolase | 944.79 | 56.38 | 31320 | 13 | 25940540 | 26521614 | 25612578 | 20582998 | 29878124 | 51622646 | 40259646 | 62494562 | 48655582 | 44654690 | 56858138 | 55370386 | 65995338 | 55665290 | 47097394 | 97382994 | 77140818 | 65375730 | 94525538 | 61382050 | 1.926988605 | 0 | 2.186055776 | 0 | 3.079351929 | 0 |
| Q9DCN2 | Cyb5r3 | NADH-cytochrome b5 reductase 3 | 616.91 | 32.89 | 34128 | 10 | 7118848 | 9929315 | 4224130 | 7929098 | 13780909 | 14013415 | 13932626 | 14568366 | 25970378 | 14913076 | 11151909 | 9860934 | 9020905 | 8622837 | 7362230 | 5090384.5 | 6342765.5 | 11099776 | 2356297.75 | 7148707 | 1.940283815 | 0 | 1.070645708 | 10 | 0.745374974 | 8 |
| Q8CGK3 | Lonp1 | Lon protease homolog, mitochondrial | 844.89 | 25.08 | 105843 | 17 | 6567353.5 | 4798552.5 | 5443349.5 | 4223956.5 | 7961036.5 | 11063631 | 10650989 | 9479042 | 14065289 | 13852630 | 11698115 | 9505348 | 12599022 | 13572474 | 9914388 | 13300126 | 13773298 | 14880314 | 16567199 | 12039036 | 2.038734717 | 0 | 1.975886597 | 0 | 2.433585164 | 0 |
| P34960 | Mmp12 | Macrophage metalloelastase | 227.05 | 11.21 | 54971 | 5 | 1747290.25 | 1524405.88 | 3815198.75 | 2520478.25 | 2615260 | 5440064 | 4462009 | 7020914.5 | 3938979.25 | 5132736.5 | 7922356 | 5463830 | 8076679 | 8823701 | 10555502 | 8275962 | 3671428 | 11374950 | 9379204 | 6645517 | 2.126767855 | 0 | 3.341511405 | 0 | 3.219196762 | 1 |
| Q62433 | Ndrg1 | Protein NDRG1 | 431.7 | 21.32 | 43009 | 6 | 2195349.25 | 3200801.75 | 4223435.5 | 1512579.75 | 4214666.5 | 7044810 | 5368003.5 | 7987352.5 | 5291346 | 7852961.5 | 12203046 | 9599240 | 7615808 | 4357332 | 4164202 | 6741291.5 | 5685017 | 6320821 | 7349623 | 7718850 | 2.185758719 | 0 | 2.472147095 | 2 | 2.203425492 | 0 |
| P26284 | Pdha1 | Pyruvate dehydrogenase E1 component subunit alpha, somatic form, mitochondrial | 173.88 | 8.21 | 43227 | 4 | 2124065.75 | 2638453.25 | 2264161.5 | 1446405.75 | 3208518.75 | 5669742 | 5143121.5 | 5918887.5 | 5296373 | 3923458.25 | 6045716 | 5452364 | 4275858 | 3547269.5 | 2343097.5 | 5181142 | 2607957 | 821036.75 | 2301765 | 2852910 | 2.221576765 | 0 | 1.85456579 | 2 | 1.178332151 | 10 |
| P35700 | Prdx1 | Peroxiredoxin-1 | 1103.39 | 53.77 | 22176 | 17 | 13871419 | 28748262 | 13611440 | 37563394 | 34030066 | 61296334 | 56168262 | 72294226 | 55498590 | 39697686 | 41313430 | 50029586 | 37164802 | 45753682 | 27941380 | 39975990 | 33018518 | 22641186 | 28011310 | 23630922 | 2.229266826 | 0 | 1.581877902 | 4 | 1.152187825 | 11 |
| O35129 | Phb2 | Prohibitin-2 | 1197.8 | 63.55 | 33296 | 18 | 4204098 | 4630888.5 | 7077039.5 | 5518796.5 | 9586686 | 15832965 | 12552542 | 17125624 | 15482124 | 10304798 | 24869354 | 23461662 | 24523416 | 22303114 | 14611208 | 20986556 | 11144225 | 12650030 | 17765512 | 10430989 | 2.298638945 | 0 | 3.538928796 | 0 | 2.352778012 | 0 |
| P36371 | Tap2 | Antigen peptide transporter 2 | 264.33 | 8.69 | 77445 | 6 | 2360856 | 3662836 | 4193169.75 | 4141515 | 4453851 | 7867523.5 | 7347698 | 10021318 | 8624597 | 9529242 | 889732.62 | 2817701 | 4643617 | 2377934 | 1997989.75 | 954741.25 | 3569631.5 | 1608510.62 | 4439558 | 846160 | 2.306498681 | 0 | 0.676526701 | 7 | 0.606977628 | 5 |
| P62320 | Snrpd3 | Small nuclear ribonucleoprotein Sm D3 | 135.47 | 32.54 | 13916 | 3 | 1915268.88 | 6486596.5 | 4338117 | 3730228.5 | 9494886 | 12693957 | 9783863 | 13306313 | 14228131 | 10034420 | 20549694 | 17782678 | 19691886 | 16158086 | 14915846 | 21357836 | 21218120 | 18592006 | 24062486 | 17204178 | 2.312592334 | 0 | 3.431459948 | 0 | 3.945089305 | 0 |
| Q8BP47 | Nars1 | Asparagine--tRNA ligase, cytoplasmic | 467.25 | 20.57 | 64279 | 9 | 1869360 | 6212728 | 2245978.75 | 3153781.25 | 5512128 | 7168069 | 8152496 | 11061540 | 10577510 | 7196997 | 11909907 | 13311732 | 5311852.5 | 8530780 | 3170412 | 8243874 | 2963372.5 | 3807652 | 6954885 | 1379614 | 2.324769285 | 0 | 2.223583072 | 4 | 1.229305412 | 10 |
| P84096 | Rhog | Rho-related GTP-binding protein RhoG | 432.66 | 39.79 | 21309 | 6 | 4128968 | 6394076 | 5561855.5 | 5902734 | 11323281 | 14046829 | 11699727 | 21998226 | 17465054 | 13723522 | 5971513.5 | 5890975 | 4942595 | 5697279 | 5187880 | 5104387 | 4806519.5 | 8536686 | 5691649 | 5060319.5 | 2.369594446 | 0 | 0.831266356 | 9 | 0.876576385 | 9 |
| Q8VC04 | Tmem106a | Transmembrane protein 106A | 175.84 | 14.56 | 29109 | 3 | 1776360.62 | 3250930.5 | 1872025.38 | 3143917 | 2565823.5 | 5899263.5 | 5327733 | 4681551 | 7722462.5 | 7166946 | 554090.06 | 1907583.12 | 1362496.5 | 2428216 | 1583202.88 | 1578631.75 | 2515020.5 | 1758546.5 | 1001731.62 | 756134.56 | 2.442526511 | 0 | 0.621425421 | 4 | 0.603539577 | 2 |
| P16460 | Ass1 | Argininosuccinate synthase | 297.31 | 16.99 | 46584 | 6 | 419610 | 1442446 | 2260190 | 2549590.25 | 2535476.25 | 4784286 | 5565525 | 5159064.5 | 8680728 | 2981132.5 | 525744.88 | 4577109.5 | 1554590.5 | 4247562 | 1522642.12 | 2709551.5 | 1632722.12 | 1146435.75 | 1927852.88 | 2482161.25 | 2.950995309 | 0 | 1.349758575 | 10 | 1.075093682 | 12 |
| P99029 | Prdx5 | Peroxiredoxin-5, mitochondrial | 794.09 | 68.57 | 21897 | 14 | 9488887 | 18599526 | 14394362 | 8802310 | 26972944 | 44275626 | 48016310 | 57353422 | 40746430 | 63947914 | 23448912 | 22285586 | 21672770 | 23946552 | 19834446 | 22975926 | 16740564 | 16086334 | 20506966 | 12493041 | 3.250014155 | 0 | 1.420790524 | 5 | 1.134744027 | 9 |
| P17136 | Snrpb | Small nuclear ribonucleoprotein-associated protein B | 173.16 | 16.02 | 23656 | 4 | 959371.31 | 1442878.12 | 918028.88 | 1791151.12 | 6923009 | 7052762 | 7067983 | 7781420.5 | 9756152 | 8229198 | 10881332 | 13079494 | 12061528 | 11410650 | 6855426.5 | 7027922.5 | 6914523.5 | 4168708.25 | 5564590 | 3979558.25 | 3.314447594 | 0 | 4.511089638 | 1 | 2.298013543 | 4 |
| B0BNG0 | Emc2 | ER membrane protein complex subunit 2 | 48.06 | 4.04 | 34870 | 1 | 399889.25 | 498415.53 | 2 | 470525.28 | 687011.44 | 710401.25 | 1011525.31 | 3221412 | 910349.75 | 1411488.25 | 808292.56 | 2 | 735148.75 | 678884.75 | 1821959.25 | 2 | 750811.62 | 970587.31 | 559651.81 | 2 | 3.533915184 | 0 | 1.967215554 | 5.5 | 1.109546879 | 10 |
| O09172 | Gclm | Glutamate--cysteine ligase regulatory subunit | 120.51 | 7.66 | 30535 | 2 | 557746.31 | 1343189 | 2 | 1719910.25 | 2426111.25 | 4069312 | 3859213.75 | 5964217 | 4333771.5 | 3143511.25 | 6221283.5 | 5728182 | 5130086.5 | 5111438.5 | 1620347.75 | 8920770 | 3473076.75 | 1971533.38 | 6062396.5 | 1696200.12 | 3.534012083 | 0 | 3.937737795 | 2 | 3.658694799 | 3 |
| Q8BVE3 | Atp6v1h | V-type proton ATPase subunit H | 311.71 | 16.36 | 55855 | 7 | 697950.5 | 592864.12 | 538543.75 | 1240671.38 | 717263.56 | 2976704.75 | 3007390.5 | 3972842 | 2819988 | 1650369.75 | 4564762 | 3849971 | 4224204.5 | 5540296 | 5296571.5 | 4642321 | 4408350 | 4816911 | 5232386 | 3798193 | 3.809394684 | 0 | 6.198570609 | 0 | 6.046049019 | 0 |
| Q9D2Y4 | Mlkl | Mixed lineage kinase domain-like protein | 136.02 | 5.3 | 54317 | 2 | 544209.94 | 762482.5 | 2 | 1179105.12 | 1283127.62 | 3330024.5 | 2616151.5 | 4385413 | 2529525.25 | 1547834.62 | 1221413 | 1961016.12 | 1038784.75 | 1183699.62 | 1026932.38 | 2709914 | 900389.38 | 1170276.62 | 1957982.88 | 2 | 3.823090281 | 0 | 1.70654554 | 6 | 1.787926526 | 8.5 |
| Q8R180 | Ero1a | ERO1-like protein alpha | 859.21 | 31.25 | 54084 | 16 | 3717368.75 | 2176000.5 | 5885350.5 | 4745550.5 | 4152475 | 9543592 | 17777838 | 16568958 | 13682502 | 24260640 | 5008791 | 3599334.75 | 3860243.75 | 4575956 | 8753255 | 5950897.5 | 10286246 | 9318237 | 6567005.5 | 8104663 | 3.957756843 | 0 | 1.247661573 | 10 | 1.945521334 | 0 |
| P62774 | Mtpn | Myotrophin | 245.57 | 26.27 | 12861 | 3 | 2 | 1095913.5 | 2 | 1919900.5 | 2840651 | 4516728.5 | 7197681 | 7915985 | 7837965 | 3481241 | 8185089 | 8744912 | 9448653 | 6625326 | 4733332 | 1573543.62 | 4006832.75 | 3397590.75 | 5039611.5 | 1852668.62 | 5.284686131 | 0 | 6.443697047 | 0 | 2.709866174 | 4 |
| Q9CR60 | Golt1b | Vesicle transport protein GOT1B | 86.94 | 10.87 | 15422 | 2 | 36930.93 | 70095.91 | 52566.57 | 124353.3 | 34206.23 | 402356.62 | 531813.56 | 311920.12 | 545582.62 | 345665.38 | 114545.47 | 265599.25 | 225204.02 | 456010.72 | 477998.69 | 2 | 1052988.25 | 570902.38 | 150759.03 | 384939.88 | 6.717958665 | 0 | 4.838421892 | 1 | 6.787903767 | 5 |
| Q64310 | Surf4 | Surfeit locus protein 4 | 69.45 | 6.69 | 30381 | 2 | 2 | 595095.19 | 2 | 2186673 | 1684678.25 | 7746673 | 7450136 | 8274980.5 | 7987254 | 3189755 | 1445613.5 | 1173954.62 | 898566.44 | 1287410.5 | 2 | 790746 | 1478189.62 | 2 | 2278103 | 2 | 7.75756923 | 0 | 1.075920829 | 12 | 1.018043899 | 12 |
| Q99P75 | Rab9a | Ras-related protein Rab-9A | 60.44 | 5.47 | 22896 | 1 | 2 | 2 | 2 | 2 | 738081.88 | 1271576.38 | 1024696.38 | 1192069 | 1538411 | 821994.69 | 476414.12 | 2 | 2 | 2 | 2 | 2 | 2 | 2 | 2 | 2 | 7.924166973 | 0 | 0.645479816 | 12 | 1.35485E-05 | 10 |
| Q3U5Q7 | Cmpk2 | UMP-CMP kinase 2, mitochondrial | 372.89 | 16.78 | 50036 | 7 | 1301941.75 | 110944.92 | 1007595.31 | 2 | 383668.5 | 4200025 | 6962541 | 3868402.5 | 3923066.25 | 4164663.5 | 709206.31 | 581709 | 1453868 | 416213.47 | 2 | 1451886.75 | 1322394.5 | 2197981.5 | 1498587.75 | 560031.88 | 8.24445119 | 0 | 1.127256382 | 10.5 | 2.507311008 | 2 |
| Q8C0N2 | Gpat3 | Glycerol-3-phosphate acyltransferase 3 | 52.78 | 2.51 | 49000 | 1 | 432245.59 | 2 | 2 | 2 | 2 | 1358715.75 | 990220.5 | 1147605.12 | 643576.81 | 434529.34 | 2 | 2 | 2 | 980599.88 | 703473.88 | 2 | 2 | 943908.75 | 1271862.38 | 1127165.12 | 10.58324934 | 0 | 3.896045745 | 9 | 7.733747798 | 6 |
| Q9BDB7 | Ifi44l | Interferon-induced protein 44-like | 160.87 | 8.5 | 49680 | 3 | 2 | 2 | 2 | 2 | 892866.5 | 3487201.75 | 3250437 | 3870667.75 | 2028524.75 | 1905512.88 | 2 | 2 | 2 | 2 | 2 | 2 | 2 | 130491.77 | 2 | 2 | 16.2871088 | 0 | 1.11998E-05 | 10 | 0.146156901 | 12 |
| Q9JIA7 | Sphk2 | Sphingosine kinase 2 | 56.32 | 1.78 | 65618 | 1 | 2 | 2 | 65128.19 | 2 | 170671.02 | 568711.69 | 1039425.06 | 529650.56 | 1452776.62 | 601413 | 271837.66 | 392544.69 | 260894.44 | 675179.81 | 413057.09 | 2 | 306869 | 105176.3 | 2 | 2 | 17.77728715 | 0 | 8.538885506 | 0 | 1.747422375 | 11.5 |
| O70404 | Vamp8 | Vesicle-associated membrane protein 8 | 35.29 | 8.91 | 11451 | 1 | 2 | 2 | 2 | 2 | 92062.99 | 739731.62 | 630138.69 | 199392.16 | 1981702.25 | 2464524.25 | 1465406 | 1959775.38 | 2080573.62 | 2299968.75 | 1480069.5 | 2 | 2250242.75 | 1730968.88 | 731385.5 | 2097928.5 | 65.33533494 | 0 | 100.8547128 | 0 | 73.97039643 | 3 |
| Q8BJS4 | Sun2 | SUN domain-containing protein 2 | 301.18 | 9.44 | 81605 | 6 | 1270364.5 | 1729638.5 | 2022400.62 | 1784884.38 | 420095.06 | 2 | 2 | 2 | 437306.12 | 2 | 2 | 2 | 148818.41 | 180637.84 | 1551082.5 | 2134424 | 136555.66 | 2 | 2 | 2688659.5 | 0.060507948 | 1 | 0.260196911 | 2 | 0.686229458 | 10 |
| P61290 | Psme3 | Proteasome activator complex subunit 3 | 111.9 | 11.42 | 29506 | 3 | 582673.81 | 1133103 | 3011018.5 | 1035007.38 | 1566607.5 | 2 | 2 | 2 | 2 | 877078.75 | 1160853.62 | 1036260 | 819331.38 | 1149589.38 | 1445069.25 | 1690376.62 | 692036.62 | 2 | 925850.94 | 2 | 0.119683086 | 1 | 0.765664514 | 12 | 0.451430541 | 6 |
| Q61072 | Adam9 | Disintegrin and metalloproteinase domain-containing protein 9 | 164.58 | 7.46 | 92079 | 4 | 1861648.25 | 2290644 | 1780622.75 | 1411145.25 | 873784.44 | 2 | 502634.91 | 120830.77 | 1226207.25 | 173120.5 | 108338.09 | 2 | 881054 | 1160993 | 741829 | 643923 | 429908.94 | 1185651.38 | 2 | 819297.75 | 0.24614671 | 1 | 0.351943387 | 2 | 0.374646052 | 1 |
| Q6IMX7 | Hspbp1 | Hsp70-binding protein 1 | 163.31 | 12.61 | 39191 | 3 | 2276276 | 2767235 | 2667044.25 | 4319962 | 3298623.75 | 336641.5 | 1060189.5 | 602820.38 | 177643.69 | 2344810.5 | 2 | 1270804.12 | 1660076.38 | 1809124 | 2272670.75 | 2 | 955965.06 | 2 | 1231669.25 | 2074187.5 | 0.295000585 | 1 | 0.457473596 | 0 | 0.278021176 | 0 |
| Q62422 | Ostf1 | Osteoclast-stimulating factor 1 | 235.59 | 21.4 | 23783 | 4 | 2962240.5 | 2067645.25 | 2780713.25 | 1662546.75 | 3031474.5 | 213686.64 | 1157247.12 | 472047.72 | 790177.5 | 2053079.62 | 266553.31 | 2 | 39743.21 | 504152.06 | 1129318.38 | 1339886 | 494839.66 | 1297546 | 1015108 | 1461530.25 | 0.374760569 | 1 | 0.15512418 | 0 | 0.448547001 | 0 |
| Q61543 | Glg1 | Golgi apparatus protein 1 | 761.99 | 15.23 | 133734 | 13 | 11023077 | 10381269 | 9165269 | 8765269 | 4318496 | 2416385 | 2097386 | 3033592.5 | 4057770.5 | 5895341.5 | 1991179 | 4741745 | 2408291.25 | 1255900.25 | 2680965.5 | 2015167.5 | 2383715.75 | 3073398.75 | 1749435.12 | 1896130.75 | 0.400896231 | 1 | 0.299589196 | 1 | 0.254684697 | 0 |
| P97310 | Mcm2 | DNA replication licensing factor MCM2 | 466.75 | 12.83 | 102078 | 9 | 5741649 | 7951722.5 | 6668155 | 7418037 | 17108018 | 953887.75 | 1890378 | 4415230 | 5000670.5 | 5865013 | 8469970 | 8531764 | 6544230 | 7781109 | 2907047 | 3600080.75 | 3029859.5 | 2791744.25 | 4273139.5 | 2 | 0.403790506 | 1 | 0.7626635 | 12 | 0.305091643 | 0 |
| Q91WD5 | Ndufs2 | NADH dehydrogenase [ubiquinone] iron-sulfur protein 2, mitochondrial | 369.62 | 15.33 | 52626 | 6 | 2812738.5 | 3707192.75 | 1914599.5 | 3511760 | 2510454.25 | 1251882.88 | 2417935 | 129536.37 | 1333782 | 1083877.75 | 93646.58 | 157165.88 | 1322492.12 | 1456137.88 | 355400.22 | 2 | 326249.25 | 234073.33 | 70574.06 | 645298.69 | 0.430042447 | 1 | 0.234135878 | 0 | 0.088276948 | 0 |
| Q8BFQ4 | Wdr82 | WD repeat-containing protein 82 | 133.47 | 12.14 | 35079 | 3 | 1300087.12 | 1337366.62 | 1196204.88 | 1336498.88 | 814064.56 | 2 | 843349.69 | 757896.94 | 542067.25 | 618958.19 | 694855.19 | 891421.56 | 1082398.62 | 561463.88 | 735788.5 | 947253.5 | 1006026.56 | 1134479.88 | 700055 | 2 | 0.461592842 | 1 | 0.662730713 | 2 | 0.63296731 | 3 |
| Q2EMV9 | Parp14 | Protein mono-ADP-ribosyltransferase PARP14 | 45.15 | 0.55 | 203802 | 1 | 1133401.75 | 1768877 | 1751671.38 | 2284813.75 | 1500462.75 | 2 | 1212867.62 | 991179 | 925143.56 | 870830.75 | 2 | 645806.12 | 2 | 2 | 2 | 2 | 744057.88 | 2 | 2 | 1335922.75 | 0.473979798 | 1 | 0.076525273 | 0 | 0.246466498 | 1 |
| Q61881 | Mcm7 | DNA replication licensing factor MCM7 | 554.39 | 18.5 | 81211 | 11 | 10809012 | 8981728 | 7440702.5 | 17969792 | 20951174 | 4286497.5 | 6001006 | 7453437 | 7134752 | 6537113 | 4556683 | 8747308 | 6959521 | 8936952 | 5406111 | 3390765 | 5366053.5 | 5655188 | 3016416 | 6699942 | 0.474855054 | 1 | 0.523134014 | 2 | 0.364739018 | 0 |
| Q8C147 | Dock8 | Dedicator of cytokinesis protein 8 | 91.29 | 1.1 | 238978 | 2 | 2653438.5 | 2026465.25 | 2961496 | 2843847.5 | 2720138 | 2585858.5 | 1289854.75 | 2 | 1773110.25 | 1349753.75 | 675902.75 | 605989.25 | 2 | 2 | 2003271.75 | 909702 | 2 | 1123261.5 | 1156242.38 | 1304775.38 | 0.52997918 | 1 | 0.248774851 | 0 | 0.340314438 | 0 |
| O70145 | Ncf2 | Neutrophil cytosol factor 2 | 242.78 | 10.29 | 59485 | 5 | 10453740 | 7764297 | 9135245 | 11035356 | 6368168 | 4601927 | 3864192 | 3500713 | 7188726 | 4653514 | 7071908 | 7112322.5 | 6191843 | 4788212.5 | 3495660.25 | 4296729.5 | 1761989 | 1753394.75 | 7163649.5 | 2913393.75 | 0.531965395 | 1 | 0.640348336 | 2 | 0.399696897 | 1 |
| P17918 | Pcna | Proliferating cell nuclear antigen | 696.55 | 61.69 | 28785 | 12 | 32127572 | 50497482 | 24616660 | 44302078 | 52772882 | 22289554 | 16664610 | 30678862 | 23167306 | 22648870 | 38112218 | 30734212 | 42172414 | 43367258 | 31061988 | 19661778 | 25512108 | 28288860 | 26294392 | 27639868 | 0.56505032 | 1 | 0.907650298 | 8 | 0.623527211 | 4 |
| Q9DCL9 | Paics | Bifunctional phosphoribosylaminoimidazole carboxylase/phosphoribosylaminoimidazole succinocarboxamide synthetase | 469.94 | 21.88 | 47006 | 9 | 14474997 | 19582874 | 11411337 | 13808046 | 16349196 | 8767420 | 7278001 | 11442959 | 9847824 | 5499200 | 9892266 | 11057216 | 10352178 | 10319646 | 9228494 | 10439579 | 8700353 | 10463994 | 10819246 | 13701482 | 0.566407705 | 1 | 0.672381158 | 0 | 0.715684182 | 1 |
| Q91YR9 | Ptgr1 | Prostaglandin reductase 1 | 291.6 | 19.45 | 35560 | 6 | 17711692 | 13418068 | 19829488 | 21647038 | 14866646 | 11929766 | 9361463 | 13955731 | 9984326 | 6961012 | 16566125 | 15422720 | 16921724 | 13869510 | 17321282 | 9986567 | 12523092 | 13666972 | 12464622 | 10465027 | 0.596667984 | 1 | 0.915727405 | 9 | 0.675709373 | 1 |
| Q8R5C5 | Actr1b | Beta-centractin | 283.82 | 13.03 | 42281 | 5 | 1050723 | 2179030.5 | 1479127.38 | 1036105.62 | 1085952 | 1009815.06 | 1047446.25 | 157646.72 | 1000024.25 | 904454 | 1042687.69 | 1526150 | 1053810.62 | 1029391.56 | 2 | 922352.62 | 666775.38 | 606604.31 | 916646 | 2 | 0.603048363 | 1 | 0.68102529 | 7 | 0.455629971 | 0 |
| P68254 | Ywhaq | 14-3-3 protein theta | 496.68 | 37.55 | 27778 | 9 | 10289632 | 9641549 | 15523674 | 11566198 | 11149067 | 6307019 | 9657434 | 5007231 | 7935234.5 | 6942308 | 8545501 | 6814398.5 | 10516723 | 7285512.5 | 14208925 | 4437912 | 6895788 | 13202230 | 5233097.5 | 5895509.5 | 0.616282492 | 1 | 0.81435383 | 6 | 0.613107503 | 4 |
| Q9ESY9 | Ifi30 | Gamma-interferon-inducible lysosomal thiol reductase | 248.78 | 20.16 | 27784 | 5 | 8190037.5 | 7046190.5 | 7372462.5 | 5182271 | 4723032.5 | 4666918 | 3878838.5 | 5109326 | 3600058 | 3031614 | 3755892 | 4130874 | 1635717.5 | 4664179.5 | 4194713 | 1341894.12 | 3011742 | 5898206 | 3000273.5 | 3622252.5 | 0.623939172 | 1 | 0.565337374 | 0 | 0.518987859 | 2 |
| Q91VD9 | Ndufs1 | NADH-ubiquinone oxidoreductase 75 kDa subunit, mitochondrial | 643.64 | 22.01 | 79777 | 14 | 6022782.5 | 5873129 | 7168196.5 | 7931593 | 10503735 | 5008654 | 2887670.5 | 5779651 | 4063099.25 | 5965970.5 | 3034090.25 | 1794975.25 | 5258641 | 4598194 | 7806179 | 3246227.75 | 8788156 | 5720888 | 998356.62 | 10603481 | 0.632144047 | 1 | 0.599797808 | 3 | 0.782868024 | 9 |
| P18572 | Bsg | Basigin | 152.76 | 8.74 | 42445 | 3 | 6967153.5 | 6870972.5 | 5594994 | 8548620 | 7067159.5 | 4104970.75 | 3037822.5 | 6394070 | 3848687 | 4788228 | 6456850 | 5771997.5 | 7687915 | 6458344 | 5318201 | 7780640 | 6361833.5 | 11540501 | 9301632 | 8822195 | 0.632652624 | 1 | 0.904259704 | 7 | 1.249876662 | 5 |
| Q64521 | Gpd2 | Glycerol-3-phosphate dehydrogenase, mitochondrial | 534.55 | 14.03 | 80954 | 10 | 3763481 | 3913397.25 | 3148935.25 | 4037330.5 | 2757806.25 | 1220497.62 | 2179101 | 2286991 | 2751843.5 | 2871082.5 | 1070887.25 | 690892.88 | 1388800.62 | 1018378.94 | 4934957 | 2198288 | 5401137 | 3145866 | 4942316 | 905327.75 | 0.641822119 | 1 | 0.516652993 | 5 | 0.941659474 | 11 |
| P11240 | Cox5a | Cytochrome c oxidase subunit 5A, mitochondrial | 289.11 | 27.4 | 16130 | 5 | 65235450 | 70873722 | 64491278 | 75663674 | 51594214 | 47910514 | 41954354 | 57179606 | 38602382 | 46892634 | 29205790 | 27138892 | 34655898 | 34196502 | 42438350 | 34523678 | 44207730 | 66395698 | 42596682 | 60365170 | 0.70926819 | 1 | 0.511304465 | 0 | 0.756695588 | 4 |
| Q91YT0 | Ndufv1 | NADH dehydrogenase [ubiquinone] flavoprotein 1, mitochondrial | 147.55 | 10.13 | 50834 | 3 | 1921741.5 | 1833127.25 | 1783156.75 | 2086493.38 | 1671178.38 | 1571856.75 | 1208416.62 | 1748422.25 | 1242223 | 1251929.12 | 940153.38 | 681869.38 | 1187536.5 | 1028157.44 | 1293136.12 | 561737.31 | 766497.5 | 764194.56 | 796410.06 | 804999.81 | 0.755494456 | 1 | 0.551959974 | 0 | 0.397370863 | 0 |
| Q9EQK5 | Mvp | Major vault protein | 957.51 | 22.76 | 95924 | 16 | 16983506 | 17974464 | 16716660 | 13496643 | 18498924 | 11739071 | 8994401 | 13201104 | 13138908 | 16165552 | 15276967 | 13669997 | 14399027 | 8777938 | 11883016 | 13522547 | 5367795.5 | 11687463 | 10817583 | 7591192 | 0.75581316 | 1 | 0.764990968 | 3 | 0.585472274 | 1 |
| P54116 | Stom | Stomatin | 776.15 | 39.44 | 31375 | 12 | 20242630 | 26655502 | 18251922 | 20336976 | 13133069 | 30257302 | 31149056 | 25092226 | 31059586 | 31550270 | 19249702 | 15864378 | 19705082 | 20393460 | 22735194 | 16441495 | 16780034 | 14771437 | 8520973 | 15356280 | 1.511947783 | 1 | 0.993183104 | 12 | 0.728758334 | 4 |
| Q8JZQ2 | Afg3l2 | Mitochondrial inner membrane m-AAA protease component AFG3L2 | 324.67 | 9.73 | 89519 | 7 | 6055052 | 6920704 | 5515274 | 2532091.5 | 3973665.75 | 7767280 | 7863292 | 6718208 | 9536636 | 6949773.5 | 4596736 | 3443136 | 3800383.5 | 4462960.5 | 4225370.5 | 7364634.5 | 4678547.5 | 4976635 | 5712268.5 | 3025861.5 | 1.553607234 | 1 | 0.821248999 | 8 | 1.030450303 | 12 |

| | | | | | | | | | | | | | | | | | | | | | | | | | | | | | | | | |
|---|---|---|---|---|---|---|---|---|---|---|---|---|---|---|---|---|---|---|---|---|---|---|---|---|---|---|---|---|---|---|---|---|
| P63085 | Mapk1 | Mitogen-activated protein kinase 1 | 88.52 | 5.87 | 41276 | 2 | 1959082.75 | 6205382 | 4095243.25 | 3247900.5 | 4831905 | 6340620 | 5107902.5 | 6437119.5 | 6412637.5 | 7510327 | 10957967 | 10454359 | 6886947 | 8918702 | 5337382.5 | 9006197 | 5355044 | 6571626 | 9020926 | 6046449 | 1.563882366 | 1 | 2.09225051 | 1 | 1.769965737 | 2 |
| P26041 | Msn | Moesin | 2471.44 | 56.67 | 67767 | 41 | 36159634 | 47561142 | 33869290 | 64798106 | 59174970 | 66666686 | 60445490 | 70021162 | 100437402 | 92483298 | 171885314 | 194163906 | 118448938 | 131269890 | 138163378 | 143685234 | 127130450 | 126115922 | 122653026 | 123973450 | 1.614708414 | 1 | 3.121053236 | 0 | 2.664140219 | 0 |
| Q9CY64 | Blvra | Biliverdin reductase A | 288.7 | 23.73 | 33525 | 6 | 4348333 | 3045923.5 | 4052789 | 2906051 | 5641208 | 5624017 | 5893045 | 6941539 | 7059546 | 7843747.5 | 5619433.5 | 5401673.5 | 3834120.25 | 3170150.25 | 4828707 | 5138730 | 3684769.5 | 5466146.5 | 3318415 | 2865863.25 | 1.668569892 | 1 | 1.143029731 | 9 | 1.023987819 | 12 |
| Q8BH04 | Pck2 | Phosphoenolpyruvate carboxykinase [GTP], mitochondrial | 1020.48 | 27.97 | 70528 | 15 | 5983376.5 | 10774934 | 8102139 | 10479106 | 12178573 | 12170727 | 14978576 | 12404062 | 22569730 | 17559356 | 9991925 | 7453014.5 | 7691604 | 8855700 | 11088532 | 5399380 | 4292352.5 | 7754426 | 10258510 | 5620793 | 1.676885297 | 1 | 0.948706881 | 10 | 0.701321002 | 3 |
| P35279 | Rab6a | Ras-related protein Rab-6A | 546.31 | 52.4 | 23590 | 11 | 2632457 | 3880841.25 | 4454890 | 3601306 | 6016036 | 7392678 | 7431077 | 8761715 | 8401115 | 4769079 | 8062133 | 8483434 | 9266088 | 5584672.5 | 2310469.5 | 5209470 | 2955822 | 2 | 5848074.5 | 873036 | 1.785509703 | 1 | 1.637402418 | 6 | 0.723148946 | 9 |
| Q9QUH0 | Glrx | Glutaredoxin-1 | 155.5 | 44.86 | 11871 | 3 | 13159430 | 24741158 | 10146403 | 15153611 | 13293549 | 32381568 | 35286842 | 25786606 | 25500854 | 19932350 | 17560038 | 16522744 | 13678486 | 14231838 | 9859014 | 13904983 | 15139144 | 11632714 | 16845166 | 13664688 | 1.815671109 | 1 | 0.939315216 | 11 | 0.930616185 | 11 |
| P84104 | Srsf3 | Serine/arginine-rich splicing factor 3 | 174.38 | 18.29 | 19330 | 4 | 2697017.25 | 5210512.5 | 3169201.25 | 4226097.5 | 6865514.5 | 9545095 | 11787885 | 12703122 | 10964365 | 6348119.5 | 14239307 | 13984641 | 12153264 | 14070471 | 10311008 | 15509445 | 13494506 | 12329943 | 14724140 | 12374145 | 2.316302418 | 1 | 2.921223792 | 0 | 3.086932524 | 0 |
| P43883 | Plin2 | Perilipin-2 | 693.1 | 30.82 | 46646 | 11 | 7952999.5 | 18255014 | 10925554 | 4999728 | 2460628.75 | 26153950 | 18072512 | 26531128 | 27089402 | 18447142 | 9148442 | 10886130 | 10005744 | 9148190 | 10912864 | 11215546 | 15162728 | 12384024 | 6441545 | 11350985 | 2.607847054 | 1 | 1.123502155 | 10 | 1.268218237 | 7 |
| Q99L13 | Hibadh | 3-hydroxyisobutyrate dehydrogenase, mitochondrial | 322.32 | 28.36 | 35440 | 6 | 2 | 2621118 | 228263.44 | 1843738.75 | 4913798 | 5090706 | 4773189 | 6555788 | 6295202 | 6012854 | 3418446 | 2449322.75 | 1502430.75 | 2331817 | 2065595.25 | 1363039 | 752755 | 309442.44 | 1550039 | 2 | 2.990317233 | 1 | 1.22490991 | 10 | 0.413793116 | 8.5 |
| P09671 | Sod2 | Superoxide dismutase [Mn], mitochondrial | 181.22 | 16.67 | 24603 | 3 | 792326.25 | 537213.06 | 1285065.5 | 1023283.5 | 2982912.5 | 1471951.5 | 4630082 | 5199713 | 6167571 | 3773316 | 686557.12 | 1009258.38 | 1357944 | 3388065.75 | 1965600.25 | 3392510 | 1064951.88 | 1982810.75 | 3505938.5 | 2238195 | 3.208468901 | 1 | 1.26985024 | 9 | 1.840322112 | 4 |
| Q9CQM5 | Txndc17 | Thioredoxin domain-containing protein 17 | 55.83 | 8.94 | 14015 | 2 | 2 | 2 | 2 | 1823332.5 | 2908948.75 | 4543083.5 | 3712258.5 | 5443225 | 3946647.25 | 2856001.5 | 1418021.5 | 996871.75 | 2 | 2 | 2 | 2 | 2 | 2 | 577906.44 | 2 | 4.332200196 | 1 | 0.510302761 | 10.5 | 0.122121589 | 9 |
| P0DOV2 | Ifi204 | Interferon-activable protein 204 | 185.47 | 7.43 | 69438 | 5 | 514680 | 557228 | 2 | 1441750 | 2120659.5 | 2045103.75 | 4366755 | 6472017 | 8597468 | 3732020 | 338369.5 | 1101439.62 | 1280522.5 | 2 | 2 | 523101.03 | 822663.5 | 1012132.81 | 590420 | 2 | 5.440575202 | 1 | 0.586997858 | 8 | 0.636192507 | 11.5 |
| P19253 | Rpl13a | Large ribosomal subunit protein uL13 | 490.61 | 39.9 | 23464 | 10 | 809942.06 | 2065789.75 | 622730.88 | 1155342.62 | 4644144.5 | 16123538 | 12403383 | 21329116 | 19748922 | 2927649.25 | 15885731 | 12933192 | 14460092 | 19538442 | 1195313.25 | 2755117.75 | 3852312.25 | 2144859 | 5852982.5 | 2797487 | 7.800924906 | 1 | 6.884611292 | 2 | 1.871676967 | 4 |
| P54001 | P4ha1 | Prolyl 4-hydroxylase subunit alpha-1 | 305.15 | 13.86 | 60898 | 7 | 2 | 440912.5 | 2 | 2 | 740835.44 | 640468.5 | 1851657.12 | 2365452.75 | 4016741 | 1446448.88 | 2177391.25 | 1925226.25 | 2424180 | 2810348.25 | 1091896.88 | 686970.69 | 471876.88 | 663695.25 | 881963.75 | 2 | 8.733432486 | 1 | 8.825054249 | 0 | 2.288554731 | 6.5 |
| Q9D0S9 | Hint2 | Adenosine 5'-monophosphoramidase HINT2 | 43.74 | 7.98 | 17320 | 1 | 2 | 2 | 2 | 2 | 1039813.69 | 3880632.25 | 1666347.25 | 2713246 | 2764918.5 | 1002608.75 | 3325129.25 | 2115428.5 | 1857201.75 | 1952721.88 | 690175.44 | 2151002 | 1213091 | 2 | 2787937.25 | 1493163.25 | 11.56713008 | 1 | 9.559962939 | 1 | 7.352410104 | 3 |
| Q80UM3 | Naa15 | N-alpha-acetyltransferase 15, NatA auxiliary subunit | 145.59 | 4.16 | 100961 | 3 | 1582372.62 | 72135.76 | 518811.69 | 115409.57 | 1365965.12 | 2 | 152682.05 | 2 | 2 | 2 | 2 | 90276.83 | 109890.77 | 413559.53 | 476779.97 | 2153122.75 | 943140.25 | 2 | 703203.81 | 1627155.88 | 0.041779153 | 2 | 0.298385822 | 6 | 1.484836641 | 9 |
| P13864 | Dnmt1 | DNA (cytosine-5)-methyltransferase 1 | 365.76 | 5.93 | 183189 | 8 | 794419.38 | 1027196.06 | 671121.75 | 1295460.12 | 1833908.38 | 2 | 2 | 2 | 876366 | 2 | 475003 | 2 | 2 | 2 | 2 | 2 | 2 | 2 | 2 | 2 | 0.155880029 | 2 | 0.084489874 | 0 | 1.77869E-06 | 0 |
| Q61033 | Tmpo | Lamina-associated polypeptide 2, isoforms alpha/zeta | 189.41 | 10.25 | 75168 | 4 | 58229.92 | 201304.25 | 845625.5 | 238661.95 | 909229.38 | 2 | 153800.7 | 57599 | 141460.41 | 49120.76 | 1184644.38 | 102623.46 | 103963.8 | 167869.11 | 807719.25 | 2 | 171589.31 | 2 | 2 | 2 | 0.17841712 | 2 | 1.050495528 | 11 | 0.076162195 | 1 |
| Q9DBR3 | Armc8 | Armadillo repeat-containing protein 8 | 44.23 | 1.78 | 75364 | 1 | 387556.09 | 620073.25 | 330131.06 | 361840 | 432933.97 | 2 | 384789.06 | 2 | 2 | 2 | 712553.31 | 731087.81 | 463538.53 | 459387.19 | 2 | 900430.81 | 2 | 632032.69 | 662165.5 | 746739.56 | 0.1804412 | 2 | 1.109744759 | 7 | 1.379284011 | 5 |
| P43247 | Msh2 | DNA mismatch repair protein Msh2 | 55.58 | 1.6 | 104151 | 1 | 1027629 | 256094.42 | 800181.19 | 607052.94 | 569577.81 | 183061.23 | 388186.94 | 66306.38 | 2 | 344784.75 | 207721.11 | 2 | 499754.47 | 482945.34 | 560570.19 | 148123.75 | 2 | 2 | 2 | 2 | 0.30128221 | 2 | 0.53702626 | 3 | 0.045431726 | 0 |
| Q8R1F5 | Hyi | Putative hydroxypyruvate isomerase | 207.87 | 17.69 | 30449 | 4 | 8325826 | 9724301 | 7996059 | 1643181 | 5490223.5 | 1619728.12 | 1407834 | 2940198 | 955364.75 | 3381309 | 7881924 | 6480740.5 | 6322256.5 | 5693454 | 3507237 | 10113052 | 4366669.5 | 4503192 | 4029066 | 4717910.5 | 0.310565432 | 2 | 0.90072275 | 9 | 0.835751424 | 9 |
| Q9D0F6 | Rfc5 | Replication factor C subunit 5 | 179.43 | 15.34 | 38096 | 4 | 2342978 | 3776439.75 | 1071435 | 2612855 | 3507526.5 | 747363 | 773547.12 | 112331.38 | 1429829.5 | 1199849.25 | 965527.94 | 843725.25 | 788172.38 | 693701.62 | 1691632.62 | 2 | 2 | 783033.06 | 2 | 2 | 0.320249811 | 2 | 0.374327408 | 1 | 0.058825579 | 0 |
| Q80W54 | Zmpste24 | CAAX prenyl protease 1 homolog | 104.35 | 4.84 | 54735 | 2 | 1603798.12 | 1458197.75 | 867637.31 | 771598.88 | 816229 | 679312.81 | 128333.26 | 79235.12 | 63194.96 | 824566.06 | 2 | 51776.03 | 825790.25 | 895249.31 | 71154.18 | 2 | 936863.06 | 1482997 | 1492643.62 | 2097938.75 | 0.321641094 | 2 | 0.334206576 | 5 | 1.089349678 | 9 |
| Q9CQQ8 | Lsm7 | U6 snRNA-associated Sm-like protein LSm7 | 99.87 | 15.53 | 11636 | 2 | 2178610.5 | 1354643.62 | 2680846.25 | 606957.31 | 840858.94 | 585085.44 | 315877.31 | 665404.69 | 363822.34 | 684464.06 | 1628829.38 | 1468809.38 | 1283933.25 | 1097639.62 | 1795357.5 | 882415.19 | 1910421.25 | 2 | 1197170.75 | 1654181.5 | 0.341253236 | 2 | 0.949445092 | 12 | 0.736655196 | 10 |
| O88986 | Gcat | 2-amino-3-ketobutyrate coenzyme A ligase, mitochondrial | 237.38 | 17.79 | 44931 | 4 | 515595.69 | 219652.44 | 101391.18 | 241532.59 | 253368.88 | 2 | 204691.16 | 2 | 176482.3 | 88069.7 | 121288.48 | 398855.25 | 217535.27 | 114124.48 | 197662.53 | 2 | 341033.16 | 189321.05 | 2 | 2 | 0.352409154 | 2 | 0.78815912 | 8 | 0.398305646 | 5 |
| P12527 | Alox5 | Polyunsaturated fatty acid 5-lipoxygenase | 241.01 | 5.2 | 78087 | 4 | 1699855 | 1432413 | 2061773.12 | 5336951.5 | 3887440.5 | 276313.5 | 1406784.75 | 398087.81 | 1977974.75 | 1111439.88 | 1165149 | 484581.97 | 2188247.5 | 1599144.62 | 2835594.5 | 166112.67 | 875472 | 2325642.5 | 1407697 | 2657502.5 | 0.358610443 | 2 | 0.57375982 | 7 | 0.515480885 | 6 |
| Q66HD3 | Nasp | Nuclear autoantigenic sperm protein | 146.24 | 5.67 | 84200 | 3 | 1761130.88 | 3454937.5 | 3083415.75 | 2987387.75 | 5566836.5 | 1896204.88 | 899731 | 88893.5 | 1083178.12 | 2085383.75 | 3659502.75 | 7677876.5 | 4409363.5 | 1772932.5 | 1208114.38 | 5535971.5 | 1401823.25 | 1784261.5 | 4943595.5 | 1279893.5 | 0.359172659 | 2 | 1.111196967 | 11 | 0.886780815 | 9 |
| P31266 | Rbpj | Recombining binding protein suppressor of hairless | 125.58 | 6.46 | 58537 | 3 | 7877346 | 8637737 | 7622155 | 3414072.25 | 3637919.5 | 2018803.38 | 1837015 | 2845310.75 | 637271.25 | 3875305 | 7891402 | 8270128.5 | 8262607 | 6452590.5 | 7133798 | 6826605.5 | 7133913 | 5037718 | 6247341 | 5758854 | 0.359537747 | 2 | 1.218706788 | 9 | 0.994074934 | 10 |
| Q8VCH8 | Ubxn4 | UBX domain-containing protein 4 | 205.24 | 12.45 | 56472 | 4 | 23049706 | 63928290 | 16709360 | 36582354 | 26852198 | 21990278 | 19101856 | 2406963 | 10070016 | 8346575 | 5193491 | 3302100.5 | 4521407 | 4991237.5 | 2630675.75 | 6785781 | 4565369.5 | 3875161.75 | 5193261 | 3748322.75 | 0.370482175 | 2 | 0.123496147 | 0 | 0.144612375 | 0 |
| Q498U4 | Sarnp | SAP domain-containing ribonucleoprotein | 153.92 | 13.81 | 23605 | 3 | 7393550 | 9447349 | 7244052.5 | 3709924 | 2554644.5 | 2559671.25 | 2079082.75 | 2860472 | 1786770.62 | 2064948.38 | 4510213 | 4469940.5 | 2202840.5 | 3116593 | 1348918.75 | 3761498 | 1934064.25 | 1216605.38 | 4858714.5 | 1505475.62 | 0.374007398 | 2 | 0.515609662 | 5 | 0.437448689 | 4 |
| A2VCX1 | Tiprl | TIP41-like protein | 53.98 | 6.64 | 31225 | 1 | 604306.44 | 999283.69 | 582837.5 | 427136.66 | 481055.47 | 491547.62 | 2 | 2 | 307539.94 | 362953.84 | 2 | 2 | 2 | 2 | 2 | 651163.94 | 2 | 2 | 218316.06 | 2 | 0.375505067 | 2 | 3.23141E-06 | 0 | 0.28096699 | 4 |
| Q5U2U0 | Clpx | ATP-dependent clpX-like chaperone, mitochondrial | 107.87 | 5.21 | 69208 | 3 | 1263421.62 | 1593959.12 | 1282999.88 | 918913.88 | 1201790.62 | 2 | 1241799.25 | 2 | 611453.06 | 726342.38 | 2 | 811907.25 | 1098744.25 | 893415.31 | 2 | 659967.94 | 2 | 2 | 699500.69 | 2 | 0.41200505 | 2 | 0.447857002 | 1 | 0.217130833 | 0 |
| Q3UBX0 | Tmem109 | Voltage-gated monoatomic cation channel TMEM109 | 196.45 | 13.99 | 26306 | 3 | 2016003.12 | 1639677.75 | 2025053.12 | 2142867.5 | 1607607.88 | 2 | 2 | 1453979.75 | 1961638.5 | 678307.69 | 2 | 836161.94 | 1694435.12 | 2136904 | 1611611.62 | 491747.06 | 811074.81 | 2915747.5 | 801176.44 | 1993885.88 | 0.434083242 | 2 | 0.665780435 | 7 | 0.743661965 | 7 |
| Q04753 | Clns1a | Methylosome subunit pICln | 52.84 | 5.51 | 26092 | 1 | 834791.12 | 1183008.38 | 637328.44 | 1173457.62 | 988731.31 | 2 | 628352.81 | 2 | 557902.5 | 916508.94 | 517390.38 | 2 | 898115.69 | 484148.34 | 2 | 2 | 2 | 762540.38 | 2 | 2 | 0.436501959 | 2 | 0.394339517 | 2 | 0.158293175 | 1 |
| P67871 | Csnk2b | Casein kinase II subunit beta | 54.89 | 5.12 | 24942 | 1 | 1526465.62 | 1589488 | 1894390.75 | 1304467.5 | 1124134 | 2 | 885596.88 | 1457158.75 | 2 | 1096922.62 | 1451257.88 | 1336205.88 | 1477540.38 | 1437904 | 1635350 | 848538.5 | 1274875.5 | 1258747.75 | 1112910.12 | 1121203.5 | 0.462388396 | 2 | 0.986464785 | 12 | 0.754982691 | 2 |
| P25206 | Mcm3 | DNA replication licensing factor MCM3 | 766.74 | 24.51 | 91546 | 15 | 7432625.5 | 5386140.5 | 6225632 | 8960733 | 12890711 | 1398071.75 | 2233366.25 | 4530676 | 6995796.5 | 4720434.5 | 6207410 | 8296383 | 10348992 | 7304144 | 3754682 | 9775681 | 3872019.5 | 3008544 | 3558319.25 | 2308282 | 0.486072521 | 2 | 0.878123771 | 10 | 0.550736815 | 4 |
| P97311 | Mcm6 | DNA replication licensing factor MCM6 | 709.91 | 19.24 | 92867 | 16 | 7016374 | 4972939.5 | 4528599 | 8230403 | 8945009 | 2668162.25 | 2204913.5 | 5255236 | 3732715.5 | 3387898.25 | 5954172.5 | 8142834.5 | 6939933 | 6337963.5 | 7721257 | 6487940.5 | 5803005 | 3423902 | 7762567 | 5478217 | 0.51193896 | 2 | 1.041635428 | 12 | 0.859387785 | 9 |
| Q2YFS3 | Pilra | Paired immunoglobulin-like type 2 receptor alpha | 84.66 | 8.61 | 33064 | 2 | 2955074 | 1496703.12 | 2295763.25 | 1268255.88 | 1850229.38 | 783272.12 | 881346.38 | 1627209 | 818339.94 | 1027893.81 | 2 | 2 | 497421.94 | 894678.12 | 722047 | 2 | 776842.81 | 2 | 2 | 958563.19 | 0.520783286 | 2 | 0.214285989 | 0 | 0.175897779 | 0 |
| P62075 | Timm13 | Mitochondrial import inner membrane translocase subunit Tim13 | 90.99 | 14.74 | 10458 | 1 | 2951467 | 1980725.62 | 2606312 | 1496193 | 1611425.75 | 1285734.62 | 1170344.25 | 1657627 | 846881.69 | 1278633.38 | 1451130.12 | 802565.5 | 1413385.75 | 2 | 1437448.12 | 2 | 2 | 1006918.06 | 2 | 2 | 0.586055668 | 2 | 0.479473261 | 0 | 0.094581476 | 0 |
| P04550 | Ptms | Parathymosin | 174.3 | 22.55 | 11559 | 3 | 7954525.5 | 7452682 | 5778362.5 | 10666569 | 4848247 | 3753045 | 4599719.5 | 5010921.5 | 5464039 | 3488119.75 | 366153.81 | 4823141 | 6348166.5 | 4566379.5 | 1966014.5 | 18528574 | 14285151 | 5459394.5 | 27349028 | 11397479 | 0.60805477 | 2 | 0.492361451 | 2 | 2.098605353 | 4 |
| P56812 | Pdcd5 | Programmed cell death protein 5 | 283.95 | 45.24 | 14275 | 5 | 4278239 | 5088320 | 4791848 | 3295464 | 4944864.5 | 3720682.25 | 2826819 | 3992262.5 | 2060321.25 | 1899999.25 | 4074997 | 4280372 | 4978882.5 | 4372867 | 3892417.5 | 3421918 | 1751363.88 | 1670969.75 | 3356035 | 1459009.75 | 0.647361734 | 2 | 0.964319437 | 10 | 0.520533687 | 2 |
| Q99L47 | St13 | Hsc70-interacting protein | 309.5 | 15.63 | 41656 | 6 | 9513420 | 5404924.5 | 11355470 | 6867605.5 | 5337390 | 5251625.5 | 4804684.5 | 4888484.5 | 4258696.5 | 6219980 | 6199801.5 | 8629660 | 8555952 | 7345967 | 5172060 | 4188193.5 | 3776593 | 3497335 | 2520118 | 215380.53 | 0.660713546 | 2 | 0.933070448 | 11 | 0.36897243 | 0 |
| Q9JL26 | Fmnl1 | Formin-like protein 1 | 875.4 | 18.1 | 122060 | 17 | 11775747 | 13101773 | 15454244 | 14395907 | 17995738 | 9846187 | 8344708.5 | 6498806 | 12363955 | 12501307 | 14271600 | 13471452 | 12837996 | 12017207 | 8079108 | 8110180.5 | 8716334 | 11555913 | 7678054 | 9919609 | 0.681416949 | 2 | 0.834358068 | 6 | 0.632259834 | 0 |
| Q7TMY8 | Huwe1 | E3 ubiquitin-protein ligase HUWE1 | 226.28 | 1.64 | 482635 | 6 | 928049.19 | 1128819.88 | 945011.19 | 1342788.25 | 1402715 | 910651.38 | 544374.38 | 972533.81 | 692609.38 | 808628.5 | 780228.44 | 1364424.62 | 1440775.75 | 1975544.75 | 1398590.75 | 1226988.62 | 1042421.44 | 1789290.5 | 1382888.62 | 2061584.88 | 0.683580179 | 2 | 1.210910025 | 7 | 1.305493891 | 6 |
| Q91V41 | Rab14 | Ras-related protein Rab-14 | 891.23 | 76.28 | 23897 | 16 | 27776224 | 32426526 | 32192920 | 23327358 | 19961586 | 19227080 | 21799252 | 15683999 | 21269958 | 17139804 | 15968152 | 17232982 | 16460083 | 15416712 | 15020684 | 15331187 | 17008326 | 19564536 | 16123711 | 19719576 | 0.701038166 | 2 | 0.590329372 | 0 | 0.646700708 | 0 |
| Q8JZN5 | Acad9 | Complex I assembly factor ACAD9, mitochondrial | 375.79 | 15.04 | 68722 | 8 | 5865435 | 4934719 | 8300346 | 6313488.5 | 7132175 | 4912872 | 2912914 | 5494705.5 | 5289403 | 4681942.5 | 1002076.81 | 2025586.75 | 3740948 | 2678594.5 | 3827852.5 | 690338.5 | 1965555.75 | 2042634.25 | 2621769.5 | 1648101.88 | 0.71565538 | 2 | 0.407883976 | 0 | 0.275559357 | 0 |
| P09895 | Rpl5 | Large ribosomal subunit protein uL18 | 527.93 | 34.68 | 34459 | 9 | 11769742 | 15005483 | 13847691 | 18785062 | 18236976 | 10841014 | 9251037 | 10807467 | 12444052 | 12230678 | 20817738 | 23253880 | 25268382 | 25805484 | 14022350 | 19914126 | 21938106 | 23360344 | 25767374 | 21353190 | 0.715748354 | 2 | 1.40598749 | 3 | 1.4467539 | 0 |
| Q922F4 | Tubb6 | Tubulin beta-6 chain | 1168.36 | 45.19 | 50090 | 19 | 15889175 | 19201524 | 17351284 | 14801321 | 13201588 | 12685784 | 13338699 | 14150360 | 8979408 | 9314541 | 7642172.5 | 5925416 | 10781049 | 10459778 | 8820867 | 5648359.5 | 10516075 | 10643566 | 10261886 | 10610539 | 0.726817956 | 2 | 0.542349942 | 0 | 0.592709174 | 0 |
| P70168 | Kpnb1 | Importin subunit beta-1 | 613.98 | 15.18 | 97184 | 11 | 18024284 | 18542902 | 18828454 | 20989976 | 22986042 | 12895281 | 14158646 | 18709666 | 12434120 | 17518644 | 13247630 | 12490846 | 14452863 | 14233406 | 18420588 | 11314284 | 9490507 | 12631413 | 11150694 | 13587731 | 0.76195123 | 2 | 0.73305945 | 1 | 0.58542476 | 0 |
| Q9CZW5 | Tomm70 | Mitochondrial import receptor subunit TOM70 | 292.04 | 9.66 | 67590 | 6 | 12400570 | 12977864 | 11430952 | 11422214 | 16287770 | 8822925 | 10124398 | 12283153 | 8627559 | 10502097 | 5700361.5 | 6496184.5 | 5898677 | 5873157 | 7224938.5 | 7384014 | 6515596 | 7827071.5 | 8956943 | 10142102 | 0.780542835 | 2 | 0.48347215 | 0 | 0.632766974 | 0 |
| P42932 | Cct8 | T-complex protein 1 subunit theta | 1255.53 | 40.15 | 59555 | 23 | 65590282 | 81012210 | 73498890 | 61645954 | 75480818 | 58934134 | 51355138 | 66749206 | 50582990 | 52395242 | 54396142 | 51981442 | 51978014 | 54507254 | 51336910 | 45046614 | 38367490 | 52167938 | 49501530 | 49083430 | 0.783859578 | 2 | 0.739582698 | 0 | 0.655511049 | 0 |
| P80315 | Cct4 | T-complex protein 1 subunit delta | 1593.52 | 58.63 | 58066 | 27 | 91069810 | 82091754 | 77193618 | 78976786 | 68691130 | 68406170 | 60557294 | 46598494 | 72998562 | 73902290 | 63087982 | 64560522 | 75206482 | 70540538 | 71134722 | 64484190 | 75444570 | 85292042 | 74495482 | 71927938 | 0.810161047 | 2 | 0.865603649 | 3 | 0.933725263 | 7 |
| Q9JKF1 | Iqgap1 | Ras GTPase-activating-like protein IQGAP1 | 2860 | 40.13 | 188742 | 54 | 45486250 | 40236554 | 50791450 | 42160362 | 51043814 | 50010554 | 53459950 | 67535906 | 59378278 | 57945302 | 49686486 | 47107738 | 51163022 | 53226834 | 52519630 | 52345450 | 39549406 | 53170506 | 40452178 | 45854762 | 1.255145223 | 2 | 1.104411649 | 4 | 1.007199562 | 11 |
| P67778 | Phb1 | Prohibitin 1 | 1314.22 | 67.28 | 29820 | 19 | 69330498 | 101909794 | 85966818 | 64631486 | 106739970 | 118057946 | 107962170 | 119755090 | 106861570 | 86689658 | 82583370 | 70436642 | 78123618 | 81685866 | 60771638 | 85444770 | 71023506 | 64287310 | 87940866 | 59971614 | 1.258407388 | 2 | 0.871721462 | 8 | 0.860211161 | 7 |
| P38647 | Hspa9 | Stress-70 protein, mitochondrial | 2285.94 | 52.43 | 73461 | 32 | 72239962 | 83420410 | 65089530 | 65672986 | 90745834 | 99586074 | 82172226 | 98045706 | 114207554 | 110264730 | 140855410 | 128422482 | 117922594 | 127593634 | 112759482 | 135692866 | 136480514 | 125622346 | 116359058 | 110647066 | 1.33700453 | 2 | 1.663853775 | 0 | 1.656557963 | 0 |
| P11762 | Lgals1 | Galectin-1 | 467.38 | 41.48 | 14857 | 6 | 49418446 | 54335358 | 40190382 | 49369270 | 63991886 | 70464026 | 71938010 | 81416442 | 57921298 | 62958006 | 82270634 | 85292090 | 77860538 | 74018418 | 69789218 | 60154234 | 79010378 | 58068374 | 45451058 | 66530626 | 1.339644872 | 2 | 1.512719849 | 0 | 1.201742131 | 6 |
| Q9JLI6 | Scly | Selenocysteine lyase | 263.05 | 18.98 | 47174 | 6 | 2814121.5 | 8484342 | 5573123 | 6661797 | 6849440 | 8811660 | 8853939 | 7242145.5 | 7213990 | 9522834 | 8463603 | 9954658 | 8116566.5 | 7422305 | 5324298.5 | 5801622 | 7080511 | 12644021 | 8427587 | 4671267 | 1.370661568 | 2 | 1.292882836 | 7 | 1.271277767 | 9 |
| P63001 | Rac1 | Ras-related C3 botulinum toxin substrate 1 | 478.58 | 43.23 | 21450 | 8 | 8483855 | 12045962 | 9033918 | 10445743 | 14131166 | 12740633 | 14071212 | 17920160 | 15938792 | 15500498 | 6386924 | 9073828 | 10621540 | 8377821.5 | 7650860 | 6102174 | 9070451 | 7672163 | 9242806 | 8459462 | 1.4069152 | 2 | 0.77780703 | 5 | 0.748920829 | 4 |
| Q9Z2L0 | Vdac1 | Non-selective voltage-gated ion channel VDAC1 | 1047.27 | 70.67 | 30756 | 15 | 20719656 | 17122736 | 12637907 | 21559252 | 33112320 | 24387052 | 22529228 | 37859574 | 34143266 | 36184926 | 17082604 | 14678875 | 25706552 | 20915906 | 15964904 | 19649004 | 19428222 | 23187606 | 24482472 | 19740802 | 1.475047895 | 2 | 0.897262598 | 10 | 1.012707667 | 11 |
| Q00612 | G6pdx | Glucose-6-phosphate 1-dehydrogenase X | 909.04 | 38.64 | 59263 | 18 | 25215466 | 33521372 | 27085950 | 24584892 | 37825638 | 53109550 | 45463218 | 45528342 | 36914922 | 37657726 | 61965706 | 65615970 | 45767062 | 45735122 | 40423842 | 60551838 | 53268030 | 37265554 | 59610586 | 45628554 | 1.475199779 | 2 | 1.750670534 | 0 | 1.729196684 | 1 |
| Q8BH43 | Wasf2 | Actin-binding protein WASF2 | 424.01 | 15.49 | 54074 | 8 | 3370600.5 | 2098519.75 | 2583947 | 2890427.5 | 2282942.5 | 3271613.75 | 3407723.75 | 3043761.5 | 6061664 | 4171569.5 | 8880675 | 9974627 | 6038283.5 | 8691620 | 3442388.5 | 7164465.5 | 6117966.5 | 5017807 | 6840514 | 3204721.75 | 1.50882147 | 2 | 2.79951383 | 0 | 2.143092219 | 1 |
| P23492 | Pnp | Purine nucleoside phosphorylase | 516.38 | 27.34 | 32277 | 7 | 4584734.5 | 8090226.5 | 6241757.5 | 8953700 | 10254576 | 14502430 | 9353715 | 13056587 | 10754311 | 9947476 | 6851702.5 | 7320888 | 3969662.5 | 5085670.5 | 4689308 | 3623757.5 | 4002651 | 2798525.5 | 1567948.25 | 2028840 | 1.511200716 | 2 | 0.732255358 | 6 | 0.367782932 | 0 |
| Q3TRM8 | Hk3 | Hexokinase-3 | 1424.12 | 33.19 | 100101 | 23 | 23562036 | 27690690 | 31346234 | 15946866 | 19989834 | 41054846 | 40949226 | 38174998 | 37476278 | 27504036 | 27184270 | 22011450 | 25849816 | 26634206 | 25436138 | 20303106 | 29758342 | 28309516 | 25037410 | 32954806 | 1.56205638 | 2 | 1.072385137 | 11 | 1.150397948 | 7 |
| P52480 | Pkm | Pyruvate kinase PKM | 3336.56 | 76.08 | 57845 | 48 | 265479794 | 313056034 | 314399938 | 377684034 | 530355650 | 555021250 | 438330754 | 533714658 | 814649730 | 475482882 | 600597442 | 616932738 | 461517698 | 436891810 | 475698498 | 419376290 | 449027042 | 441798786 | 424795906 | 389210114 | 1.564263008 | 2 | 1.439019164 | 3 | 1.179476454 | 5 |
| Q8R1F1 | Niban2 | Protein Niban 2 | 557.52 | 16.42 | 84819 | 11 | 4329201.5 | 3986480 | 6033422.5 | 2105416.5 | 2302049 | 7537920 | 7720119.5 | 6796071.5 | 4388783 | 4490143.5 | 7660116 | 3267919.75 | 6402190 | 5034158.5 | 4183421 | 6566265 | 6023073 | 6086367 | 4788129 | 4795792 | 1.64918417 | 2 | 1.415387033 | 6 | 1.506652163 | 3 |
| P06151 | Ldha | L-lactate dehydrogenase A chain | 1308.42 | 62.95 | 36499 | 21 | 107720930 | 114408562 | 113041034 | 111911970 | 185814290 | 178791042 | 179128546 | 224662658 | 257644754 | 209821682 | 378680290 | 430477794 | 234942530 | 269703106 | 193557730 | 322319906 | 219004674 | 167771970 | 288724514 | 154744930 | 1.659115207 | 2 | 2.381686056 | 0 | 1.821096298 | 2 |
| P08030 | Aprt | Adenine phosphoribosyltransferase | 798.36 | 78.89 | 19724 | 13 | 2989605.75 | 5781460 | 7919466.5 | 9042016 | 9310141 | 13760296 | 12926512 | 12873876 | 7951732.5 | 11989038 | 11692195 | 10646377 | 12062066 | 8760499 | 25475784 | 9417290 | 9771110 | 12319301 | 7725669.5 | 10235680 | 1.697970555 | 2 | 1.958665915 | 2 | 1.411679627 | 3 |
| O35643 | Ap1b1 | AP-1 complex subunit beta-1 | 588.49 | 14.1 | 103935 | 10 | 3084675 | 3088713.25 | 2248709 | 5270204 | 4313089 | 7187583 | 5247875 | 6968444 | 6767137.5 | 4719072.5 | 5888561 | 5311995 | 5200669.5 | 4308224 | 3157457.75 | 5818119 | 5099759.5 | 5622210 | 4680265 | 4369734.5 | 1.715603582 | 2 | 1.325542347 | 5 | 1.421245952 | 3 |
| Q80Z29 | Nampt | Nicotinamide phosphoribosyltransferase | 211.51 | 12.22 | 55438 | 5 | 3552803.5 | 3864766.5 | 3482315 | 5793184.5 | 5809184.5 | 7453220 | 5718451.5 | 8704219 | 8856752 | 7915454 | 6698536.5 | 5358302 | 5385704 | 5975802 | 3938434 | 7612761.5 | 5884829.5 | 7763420.5 | 7915381 | 7467488 | 1.71752112 | 2 | 1.215735033 | 6 | 1.628453776 | 0 |
| F6ZDS4 | Tpr | Nucleoprotein TPR | 913.86 | 9.38 | 273990 | 18 | 6644318 | 4658031 | 4740075 | 14662877 | 6642258.5 | 18844698 | 17587484 | 17117626 | 8225466 | 6801699 | 11195330 | 12863432 | 10123901 | 6771325.5 | 3578806 | 16144664 | 10390366 | 751893.81 | 7336861.5 | 6150397 | 1.836183513 | 2 | 1.192388341 | 9 | 1.091749578 | 10 |
| P48758 | Cbr1 | Carbonyl reductase [NADPH] 1 | 164.41 | 9.03 | 30641 | 3 | 2441443.5 | 1501072.25 | 1516362 | 2428667.5 | 1450550 | 3856318 | 3577368 | 4228679 | 3718460.75 | 1998752.25 | 3401113.5 | 3490770.75 | 1662674.75 | 1962451.5 | 2738676 | 4345914 | 5630500 | 3725519 | 3257181.5 | 9048481 | 1.861148075 | 2 | 1.419527874 | 4 | 2.785107113 | 0 |
| Q8R0X7 | Sgpl1 | Sphingosine-1-phosphate lyase 1 | 293.64 | 10.39 | 63677 | 6 | 3277166 | 3208282.75 | 1836206.38 | 2110427 | 1801735.38 | 4138422.5 | 7370894.5 | 4896744 | 3698436.5 | 2876605.25 | 2066917.5 | 1893903.25 | 4435955 | 5159214.5 | 3508347 | 2714391.5 | 2811541.5 | 4023402.75 | 2605945.25 | 3136037.25 | 1.87848991 | 2 | 1.394849746 | 6 | 1.249922049 | 8 |
| Q9Z0X1 | Aifm1 | Apoptosis-inducing factor 1, mitochondrial | 773.7 | 29.58 | 66765 | 14 | 3788268.75 | 2802908.5 | 3033380.5 | 2710384.75 | 3952886.25 | 5942557 | 5319947.5 | 4914535 | 11501732 | 3378811.5 | 14513321 | 14349574 | 8970858 | 10582814 | 9016713 | 11157832 | 5260577.5 | 7356709.5 | 9470906 | 7630630 | 1.906797 | 2 | 3.526147093 | 0 | 2.509644203 | 0 |
| Q8BPX9 | Slc15a3 | Solute carrier family 15 member 3 | 388.9 | 14.19 | 64051 | 6 | 6264685.5 | 5042557.5 | 4089030.25 | 3135996 | 3041179.75 | 9781516 | 8555917 | 11783861 | 8614352 | 4212863 | 2144710.25 | 650049.25 | 1938436.38 | 2131668.75 | 3460962 | 4387438 | 5841909 | 7474697.5 | 5858394 | 3776445.5 | 1.99080402 | 2 | 0.478635875 | 2 | 1.26724679 | 7 |
| P70206 | Plxna1 | Plexin-A1 | 439.46 | 6.23 | 211099 | 9 | 1883346.38 | 1823904.75 | 568314.38 | 2087547.75 | 3131737 | 3010733.75 | 4093774.75 | 2306569.25 | 5494484 | 4387692 | 1498579.25 | 2053406.5 | 1396579.25 | 1438685.25 | 2462936.25 | 2377422 | 3818821.75 | 4553284 | 948648.06 | 2602914 | 2.031970302 | 2 | 0.932103852 | 10 | 1.506194349 | 6 |
| Q99L45 | Eif2s2 | Eukaryotic translation initiation factor 2 subunit 2 | 244.87 | 8.46 | 38092 | 5 | 4167082.5 | 4643582 | 2449438 | 6060031.5 | 7350135 | 8797467 | 9735414 | 14109189 | 12231153 | 5757658.5 | 11731560 | 9318759 | 10532087 | 9323444 | 6594662.5 | 10411003 | 9204718 | 9059762 | 13275035 | 8546372 | 2.052303585 | 2 | 1.925415264 | 1 | 2.04687229 | 0 |
| Q80X90 | Flnb | Filamin-B | 386.48 | 4.38 | 277825 | 9 | 1131113.5 | 3068476.25 | 1578489.5 | 1299448.12 | 1630904 | 5051342.5 | 3173963.25 | 5835995 | 2369210 | 1863617.75 | 1922865.38 | 1705994.5 | 2372185 | 1040371.56 | 898243.12 | 2723106.5 | 3625364 | 2130807 | 3131857 | 2485479.75 | 2.100737518 | 2 | 0.911720977 | 12 | 1.618731738 | 3 |
| Q9D0F9 | Pgm1 | Phosphoglucomutase-1 | 591.26 | 26.87 | 61418 | 11 | 7292749 | 4873740 | 5766748.5 | 10976661 | 9745134 | 22886132 | 18026906 | 14385056 | 19969004 | 8312206.5 | 10199055 | 10263712 | 6894621.5 | 4074636 | 7488873 | 12366091 | 13400871 | 10648135 | 12714941 | 7059162.5 | 2.162184303 | 2 | 1.006877888 | 12 | 1.453606345 | 4 |
| P06797 | Ctsl | Procathepsin L | 229.11 | 27.54 | 37547 | 5 | 2572353.5 | 4053246 | 4163844.5 | 2752805.5 | 3479982 | 7705782 | 11124732 | 8979567 | 3654417 | 5710457 | 2467147 | 1096804.12 | 2535969.5 | 2235813.5 | 4256049.5 | 584025.62 | 4600990 | 5319623.5 | 2588154 | 3964336.5 | 2.183906088 | 2 | 0.739725789 | 5 | 1.00205015 | 11 |
| P20291 | Alox5ap | Arachidonate 5-lipoxygenase-activating protein | 177.28 | 18.63 | 18070 | 4 | 4542682 | 4309182 | 3095317 | 5150982 | 8571666 | 11633614 | 5258592 | 17239840 | 17905152 | 7383497.5 | 7620363 | 2876846.5 | 3145408 | 2885443.75 | 3378766.25 | 7705565 | 5270311.5 | 7180145.5 | 4226630 | 6673684.5 | 2.31480683 | 2 | 0.775495135 | 6 | 1.209838075 | 8 |
| Q9R0N0 | Galk1 | Galactokinase | 315.15 | 14.54 | 42295 | 7 | 2 | 1757560.62 | 2253764.75 | 1047678.38 | 4286003 | 2638388 | 3953308 | 6729657 | 5417976 | 4555444.5 | 9983930 | 8559553 | 7913805.5 | 5801622.5 | 5493793 | 4574603 | 2306349.5 | 3694550 | 4247227 | 2753370.5 | 2.492750314 | 2 | 4.039878935 | 0 | 1.880800807 | 4 |
| P54775 | Psmc4 | 26S proteasome regulatory subunit 6B | 356.26 | 15.07 | 47408 | 6 | 851727.56 | 213050.41 | 1077189.5 | 1117252.5 | 2979395.5 | 1327975.75 | 2966838.5 | 3553110.5 | 4807502.5 | 3245749.75 | 924845.62 | 1132288.62 | 2079667.75 | 1514058.88 | 3495064 | 967669.94 | 722176 | 399405.88 | 1634287.62 | 770767.62 | 2.548831079 | 2 | 1.466018368 | 6 | 0.720401359 | 9 |
| Q60766 | Irgm1 | Immunity-related GTPase family M protein 1 | 260.48 | 15.16 | 46552 | 5 | 817056.56 | 511029.62 | 502377.16 | 1397889.12 | 1237583.62 | 2139113 | 2168098.75 | 3554083 | 3490261.25 | 896273.75 | 591074.69 | 69662.84 | 744142.5 | 71264.24 | 152039.31 | 2 | 1177696.38 | 2 | 765447.38 | 179397.58 | 2.742500011 | 2 | 0.364578344 | 4 | 0.475274456 | 5 |
| P16110 | Lgals3 | Galectin-3 | 533.1 | 38.64 | 27515 | 9 | 3127265 | 5263680 | 5333368.5 | 4447551 | 19641258 | 20913506 | 11014831 | 27253018 | 28267058 | 18261660 | 104838130 | 112329122 | 53973854 | 64469754 | 27105718 | 112963626 | 55047658 | 31998546 | 130974898 | 27602866 | 2.795592271 | 2 | 9.592346625 | 0 | 9.483152152 | 0 |
| O88351 | Ikbkb | Inhibitor of nuclear factor kappa-B kinase subunit beta | 97.13 | 3.57 | 86690 | 2 | 2 | 873069.06 | 2 | 2 | 888978.56 | 1306493.25 | 1181910.38 | 951296.25 | 1207512.25 | 418248.03 | 1337411.5 | 1062161.25 | 1058185.25 | 1237493.5 | 2 | 981625.88 | 1170010.5 | 716052 | 2 | 1083021.88 | 2.874748023 | 2 | 2.66464848 | 3.5 | 2.242106719 | 5.5 |
| Q9CYN2 | Spcs2 | Signal peptidase complex subunit 2 | 127.61 | 10.62 | 24978 | 3 | 271372.47 | 2 | 2 | 483714.31 | 1024545.19 | 631177.62 | 579573.12 | 1368770.12 | 1435121.75 | 1434663.25 | 878667.69 | 1109073.75 | 1332435.75 | 1520549.5 | 1505352.38 | 1009167.31 | 1150611 | 1213554.38 | 1059077.75 | 1052337.25 | 3.062034007 | 2 | 3.565942236 | 1 | 3.08194922 | 1 |
| O35609 | Scamp3 | Secretory carrier-associated membrane protein 3 | 164.37 | 9.17 | 38458 | 2 | 2 | 700288.12 | 536191.88 | 2 | 2135326.5 | 2245838 | 1742991.25 | 3344586 | 2932386.5 | 1106011.62 | 3824814.5 | 2740424.75 | 1387736 | 1261533.62 | 1759677.62 | 4125530.25 | 5068695.5 | 2886236.5 | 4290584 | 2732534 | 3.372613428 | 2 | 3.254686611 | 3 | 5.665674346 | 0 |
| O08528 | Hk2 | Hexokinase-2 | 717.61 | 18.76 | 102535 | 15 | 512493.81 | 265491.81 | 783772.19 | 522957.19 | 2402984 | 2316517 | 2888890 | 1622379.75 | 4856330.5 | 3623743.75 | 2939062.5 | 2291841.25 | 1815647.25 | 965715.75 | 1405528.5 | 2625344.75 | 4722898 | 4279405 | 3108770.75 | 3588519 | 3.411071242 | 2 | 2.098579974 | 4 | 4.083370453 | 0 |
| P20491 | Fcer1g | High affinity immunoglobulin epsilon receptor subunit gamma | 153.72 | 23.26 | 9652 | 3 | 2 | 931609.12 | 2 | 1123922.88 | 2157792.5 | 3271061.5 | 1921324.88 | 3008396.5 | 5067920.5 | 1606688.75 | 4555421 | 5484753 | 4560800.5 | 4709836 | 2091373.12 | 4432980 | 3293572.75 | 2238744.25 | 4874449 | 2163969.5 | 3.53055598 | 2 | 5.079638015 | 1 | 4.035696599 | 0 |
| Q8C4J7 | Tbl3 | Transducin beta-like protein 3 | 45.25 | 1.62 | 88266 | 1 | 2 | 2 | 118165.28 | 364050.75 | 32864 | 613754.06 | 642745.81 | 134801 | 256418.73 | 392625.78 | 86933.51 | 295321.16 | 547741.31 | 445769.5 | 889565.94 | 2 | 792860.5 | 218055.25 | 2 | 2 | 3.961189362 | 2 | 4.397984189 | 3 | 1.962634621 | 12 |
| Q8VI75 | Ipo4 | Importin-4 | 46.08 | 1.57 | 119275 | 1 | 2 | 2 | 60303.5 | 2 | 33030.78 | 102567.49 | 122518.11 | 58072.23 | 49993.58 | 72882.61 | 2 | 77820.64 | 49839.71 | 61975.25 | 91679.59 | 2 | 2 | 2 | 2 | 2 | 4.350040733 | 2 | 3.013888431 | 4.5 | 0.000107135 | 7.5 |
| Q8BMD8 | Slc25a24 | Mitochondrial adenyl nucleotide antiporter SLC25A24 | 385.62 | 19.16 | 52902 | 7 | 849659.69 | 638957.75 | 273005.5 | 997179.75 | 530804 | 2500985.75 | 2669683.75 | 756790.44 | 4605725 | 3778015.25 | 1866918.38 | 3077163.5 | 1291616 | 906523.44 | 1404402.62 | 718274.31 | 1482512 | 1457959.25 | 505072.62 | 1259424 | 4.350428954 | 2 | 2.598068628 | 1 | 1.648598964 | 6 |
| Q8VDP6 | Cdipt | CDP-diacylglycerol--inositol 3-phosphatidyltransferase | 224.95 | 29.58 | 23599 | 5 | 811432.38 | 1736875.12 | 181852.84 | 2 | 235495.7 | 3642838.5 | 2299277 | 269238.75 | 4303924.5 | 2586288 | 2215709.75 | 147121.11 | 3322537.5 | 3475057.75 | 2991529 | 2138274 | 3933958 | 4637726.5 | 2483439.25 | 2760026.25 | 4.417760434 | 2 | 4.097557758 | 4 | 5.379387571 | 0 |
| P35282 | Rab21 | Ras-related protein Rab-21 | 202.18 | 16.22 | 24106 | 3 | 632974.06 | 2 | 2 | 445992.62 | 636770.56 | 1579131.88 | 2124291 | 878925.19 | 2652738.25 | 547475.62 | 560545.5 | 3167582 | 1172710.75 | 1920236.88 | 2 | 3383772 | 1393817.88 | 742670.75 | 2284037 | 2 | 4.535976497 | 2 | 3.975586161 | 6 | 4.548646059 | 4 |
| Q07984 | Ssr4 | Translocon-associated protein subunit delta | 198.54 | 24.86 | 18980 | 3 | 2 | 161350.42 | 68984.69 | 268191.12 | 1025484.75 | 374952.81 | 3634304.75 | 382182.81 | 1541147.25 | 2284500.5 | 1069052.25 | 975028.5 | 1020178.19 | 1075225 | 419809.28 | 378954.84 | 1136756.5 | 628901.5 | 462466.81 | 282169.25 | 5.391744183 | 2 | 2.991636738 | 3 | 1.895816465 | 4 |
| Q91VE0 | Slc27a4 | Long-chain fatty acid transport protein 4 | 91.45 | 2.95 | 72319 | 2 | 2 | 2 | 2 | 899168.62 | 276162.81 | 1644624 | 205906.97 | 2321963.25 | 1523796.75 | 930046.75 | 869142.5 | 137265.69 | 1567213.75 | 413879.94 | 169991.39 | 2 | 410990.38 | 113147.41 | 2 | 191738.89 | 5.63781732 | 2 | 2.686456833 | 6 | 0.609085239 | 12 |
| Q4LDD4 | Arap1 | Arf-GAP with Rho-GAP domain, ANK repeat and PH domain-containing protein 1 | 151.04 | 2.34 | 162276 | 3 | 2 | 2 | 2 | 2 | 1190503.62 | 860328.5 | 1887876.25 | 1992832 | 1778480.25 | 560440.06 | 860140.56 | 2 | 2 | 2 | 2 | 2 | 2 | 2 | 2 | 2 | 5.946986943 | 2 | 0.722503288 | 12 | 8.39975E-06 | 10 |

Supplementary table 3: List of modulated proteins in the 4x5µg/ml condition, selected by the Mann Whitney U test (p<0.05)

| accession number | gene_name | description | protein_set_score | coverage | MW | #peptides | Ctrl 1 | Ctrl 2 | Ctrl 3 | Ctrl 4 | Ctrl 5 | PBk14 20 recov 1 | PBk14 20 recov 2 | PBk14 20 recov 3 | PBk14 20 recov 4 | PBk14 20 recov 5 | PBk14 4x5 rep1 | PBk14 4x5 rep2 | PBk14 4x5 rep3 | PBk14 4x5 rep4 | PBk14 4x5 rep5 | PBk14 4x10 rep1 | PBk14 4x10 rep2 | PBk14 4x10 rep3 | PBk14 4x10 rep4 | PBk14 4x10 rep5 | ratio PBk14 20 recov/ctrl | U PBk14/20 recov | ratio_PBk14/ ctrl | U_PBk14 4x5 | ratio Pbk 14 4x10/ctrl | U Pbk 14 4x10 |
|---|---|---|---|---|---|---|---|---|---|---|---|---|---|---|---|---|---|---|---|---|---|---|---|---|---|---|---|---|---|---|---|---|
| P35283 | Rab12 | Ras-related protein Rab-12 | 158.28 | 12.35 | 27329 | 3 | 2626988.25 | 2274646.5 | 1479324.88 | 1650430.88 | 1640128.75 | 2 | 2 | 2 | 2 | 2 | 2 | 2 | 2 | 2 | 2 | 2583009.5 | 2 | 1630908.12 | 2 | 1270759.38 | 1.03396E-06 | 0 | 1.03396E-06 | 0 | 0.567096115 | 5 |
| P35433 | Ppat | Amidophosphoribosyltransferase | 37.89 | 1.35 | 57437 | 1 | 1384687.88 | 1062445.5 | 918004.81 | 862753.44 | 1043741.19 | 2 | 2 | 2 | 2 | 2 | 2 | 2 | 682479.81 | 586809.25 | 2 | 2 | 580160.31 | 2 | 2 | 2 | 1.89695E-06 | 0 | 0.240778351 | 0 | 0.110054765 | 0 |
| Q9D338 | Mrpl19 | Large ribosomal subunit protein bL19m | 176.21 | 12.67 | 33578 | 4 | 575289.62 | 1003186.94 | 502012.88 | 769077.5 | 832785.31 | 139775.58 | 2 | 2 | 227855.81 | 277760.75 | 2 | 2 | 100854.18 | 292543.81 | 2 | 640350.12 | 2 | 763403.31 | 702465.81 | 723937.06 | 0.175267355 | 0 | 0.106834969 | 0 | 0.768573485 | 8 |
| P49718 | Mcm5 | DNA replication licensing factor MCM5 | 454.31 | 16.21 | 82407 | 11 | 14334609 | 23994526 | 12389125 | 23668398 | 18101840 | 5167847 | 2130634 | 10794580 | 8285995.5 | 7981872 | 8040220 | 7604583 | 8059879 | 10139252 | 6163571.5 | 6093196 | 5124019 | 2573817.75 | 7862198.5 | 9596300 | 0.371515694 | 0 | 0.432567361 | 0 | 0.337874784 | 0 |
| P46061 | Rangap1 | Ran GTPase-activating protein 1 | 481.99 | 20.54 | 63531 | 10 | 9060379 | 6495795 | 7827281 | 7722898 | 6361215 | 1926580.75 | 2586222.5 | 2292656.25 | 2893515 | 4771926 | 2518570.5 | 1173641.75 | 1092664.38 | 2043450.25 | 1185948.38 | 1744415.88 | 2525018.75 | 178639.88 | 1270313.75 | 1076775.12 | 0.386224708 | 0 | 0.213898998 | 0 | 0.181361208 | 0 |
| B2RXS4 | Plxnb2 | Plexin-B2 | 500.43 | 6.03 | 206230 | 9 | 10473187 | 11143706 | 9682170 | 5885522.5 | 8422308 | 2396757 | 632654.44 | 5870254 | 5480084 | 5764672.5 | 3756564 | 3685656 | 4858579 | 3459083 | 3609338.75 | 3586604.25 | 5118156 | 3967404.75 | 3771416.75 | 2162288.75 | 0.441696866 | 0 | 0.42469941 | 0 | 0.407961803 | 0 |
| Q91YE6 | Ipo9 | Importin-9 | 292.36 | 8.07 | 116052 | 5 | 5661126.5 | 5772475 | 4431324 | 4939674.5 | 5068063.5 | 2326718 | 2362680.5 | 2742037.5 | 2651297.5 | 1563304.5 | 4272772.5 | 3232113.25 | 4295584.5 | 3404548.25 | 2630744.25 | 4011597 | 3580901.75 | 3850355.5 | 3650960.25 | 2961686.75 | 0.450129072 | 0 | 0.68936709 | 0 | 0.697860166 | 0 |
| F1LNJ2 | Snrnp200 | U5 small nuclear ribonucleoprotein 200 kDa helicase | 706.67 | 8.42 | 244875 | 16 | 11609262 | 12076759 | 13563099 | 11279108 | 13219409 | 7241474.5 | 5276201 | 4889992 | 5463917 | 6162063.5 | 6777222 | 4417565 | 9723926 | 10284704 | 7422278 | 6912283 | 6573443 | 11464402 | 4919321 | 7694628.5 | 0.470198528 | 0 | 0.625541266 | 0 | 0.608348421 | 1 |
| P58389 | Ptpa | Serine/threonine-protein phosphatase 2A activator | 132.85 | 12.38 | 36710 | 3 | 4224244.5 | 5166914 | 5568233 | 3581562 | 3818065.25 | 3062783.25 | 2352178.25 | 153137.73 | 2468467.25 | 2775355.75 | 2148700 | 2399475.75 | 2887048 | 2655521.25 | 2379607.25 | 3122168.25 | 2554408 | 3590522.75 | 2604543.25 | 3703821 | 0.483559782 | 0 | 0.557732537 | 0 | 0.696607638 | 2 |
| Q9D906 | Atg7 | Ubiquitin-like modifier-activating enzyme ATG7 | 141.96 | 4.3 | 77520 | 3 | 4201814.5 | 3761543.75 | 4630190.5 | 3653197.25 | 3896952.5 | 93426.41 | 2989151.25 | 1120287 | 2885658.5 | 3649918.25 | 1759115.75 | 1743334.88 | 2780771.75 | 2936376.25 | 2306131 | 1619819.5 | 3279527.75 | 2674177 | 2116290 | 2440309.5 | 0.533091846 | 0 | 0.572175444 | 0 | 0.602179572 | 0 |
| P20060 | Hexb | Beta-hexosaminidase subunit beta | 353.4 | 17.72 | 61116 | 8 | 19787152 | 25215262 | 21984138 | 24970668 | 20180386 | 11218148 | 12726519 | 7381907 | 17365548 | 17356838 | 8912201 | 7231532.5 | 8537964 | 8499397 | 11826519 | 6433805 | 10468561 | 12581139 | 9981005 | 15225826 | 0.588999198 | 0 | 0.40136057 | 0 | 0.487707362 | 0 |
| P97855 | G3bp1 | Ras GTPase-activating protein-binding protein 1 | 416.46 | 21.94 | 51829 | 8 | 2828582.5 | 3678635.5 | 3172076.25 | 2826818.75 | 2906179 | 2091237.88 | 1358502.25 | 1242791.25 | 2259942.5 | 2201344 | 4617295 | 5067650.5 | 5027289 | 4846177.5 | 3790235.5 | 5474398 | 4820392.5 | 2039262.38 | 5806616.5 | 3681098 | 0.593929695 | 0 | 1.514936747 | 0 | 1.415867762 | 5 |
| Q8BG07 | Pld4 | 5'-3' exonuclease PLD4 | 943.83 | 38.57 | 56154 | 14 | 56814230 | 54365838 | 48080822 | 49538466 | 42085726 | 28968542 | 28506038 | 28419456 | 26219874 | 40881506 | 17161362 | 14213955 | 16546519 | 14629944 | 27383632 | 21365194 | 19312210 | 24940534 | 18028056 | 20254480 | 0.609822692 | 0 | 0.358472538 | 0 | 0.41413572 | 0 |
| P29416 | Hexa | Beta-hexosaminidase subunit alpha | 430.36 | 20.45 | 60613 | 9 | 11239542 | 9282127 | 9346251 | 12898934 | 8747906 | 3946593.75 | 5545161.5 | 6881814.5 | 7755726 | 7295286 | 2764089.5 | 2039271.25 | 792291.81 | 2729312.5 | 3822945.5 | 4363559.5 | 3434400 | 6897750 | 4884175 | 4994353.5 | 0.610011223 | 0 | 0.235814174 | 0 | 0.477032951 | 0 |
| O55201 | Supt5h | Transcription elongation factor SPT5 | 146.03 | 4.44 | 120664 | 3 | 1728923.75 | 1680876.25 | 3477661.25 | 1856451.38 | 1549461 | 1051832.12 | 902829.62 | 1383732.75 | 1507011.62 | 1481699.25 | 1022213.88 | 778070 | 951228.75 | 926730.62 | 1217765.75 | 1436754.38 | 1034174.06 | 1261830.25 | 3883472 | 2392161 | 0.614677519 | 0 | 0.475646681 | 0 | 0.972314039 | 9 |
| Q8BFY9 | Tnpo1 | Transportin-1 | 297.62 | 5.01 | 102357 | 5 | 11038026 | 8185250.5 | 11575319 | 7989010 | 9471328 | 5731841 | 5967353 | 7195839 | 6718364 | 5640400 | 3683115.25 | 4458412.5 | 5869673 | 4904413.5 | 5360765 | 3546715.5 | 4544134 | 2163104 | 2871810.75 | 2622370 | 0.64762718 | 0 | 0.503044255 | 0 | 0.326325783 | 0 |
| Q9QZD8 | Slc25a10 | Mitochondrial dicarboxylate carrier | 680.98 | 36.93 | 31715 | 12 | 11981910 | 14999013 | 13635671 | 13621286 | 17404118 | 9275321 | 9050245 | 8613553 | 11941318 | 10718740 | 9006348 | 7019183.5 | 8229059.5 | 9334255 | 6532186.5 | 6385108 | 4662828.5 | 8772034 | 5495926 | 7336750 | 0.692319846 | 0 | 0.560021128 | 0 | 0.455775207 | 0 |
| Q6DFW4 | Nop58 | Nucleolar protein 58 | 504.86 | 25.37 | 60343 | 9 | 7846229.5 | 10351596 | 10421583 | 8862480 | 7747698.5 | 6737081 | 5246814 | 5930571 | 7066720 | 7190264 | 3015261.75 | 3000159.5 | 6669755 | 4664555.5 | 6073845 | 5083554 | 5514721 | 7353035 | 7480814.5 | 8022738 | 0.711292146 | 0 | 0.51788173 | 0 | 0.739667654 | 2 |
| P63017 | Hspa8 | Heat shock cognate 71 kDa protein | 3151.25 | 68.27 | 70871 | 47 | 421192898 | 431482370 | 373439234 | 421580098 | 450136930 | 284628066 | 257253202 | 317597506 | 297581762 | 362076450 | 358839650 | 359159010 | 313883746 | 310589186 | 372030082 | 269497954 | 320537218 | 317056450 | 216600722 | 301734146 | 0.724146322 | 0 | 0.817273289 | 0 | 0.679476149 | 0 |
| Q3UND0 | Skap2 | Src kinase-associated phosphoprotein 2 | 122.18 | 8.1 | 40712 | 3 | 3969949.25 | 3809952.75 | 3530172.5 | 5301210.5 | 3558785.5 | 3133936.75 | 2973437.5 | 3486063 | 2766528.5 | 2947538.5 | 1311216.75 | 1656630.25 | 1974965.38 | 1279712.75 | 1255237.88 | 2672843 | 1643412.12 | 2004220.75 | 2632103 | 2089896.12 | 0.758921703 | 0 | 0.370735591 | 0 | 0.547468339 | 0 |
| P57776 | Eef1d | Elongation factor 1-delta | 669.69 | 36.3 | 31293 | 10 | 55972342 | 56603278 | 49338410 | 54788842 | 53468642 | 42996234 | 41529478 | 47330046 | 34967330 | 43419870 | 29211538 | 30174042 | 34276694 | 32177534 | 27318584 | 26016820 | 27842364 | 25829278 | 29964362 | 31675226 | 0.778183291 | 0 | 0.566893192 | 0 | 0.523104927 | 0 |
| Q7TMB8 | Cyfip1 | Cytoplasmic FMR1-interacting protein 1 | 751.51 | 13.01 | 145241 | 15 | 6839314 | 7714240.5 | 8690557 | 7099803 | 8213055.5 | 11880539 | 9639829 | 16722741 | 12318325 | 9600170 | 16534412 | 13017145 | 15349741 | 14831595 | 13903786 | 11246515 | 11660331 | 16479392 | 11029072 | 10603626 | 1.560330182 | 0 | 1.909814983 | 0 | 1.582565642 | 0 |
| Q8VCN5 | Cth | Cystathionine gamma-lyase | 497.8 | 27.14 | 43567 | 10 | 9377385 | 11501992 | 20802036 | 13511600 | 17593928 | 23467510 | 23925028 | 22619538 | 21611462 | 23167656 | 25079238 | 26935692 | 23539544 | 22674670 | 26509684 | 26265474 | 32094046 | 23433328 | 23389266 | 24455162 | 1.577085016 | 0 | 1.713752856 | 0 | 1.781051301 | 0 |
| P20152 | Vim | Vimentin | 3751.201 | 80.69 | 53688 | 58 | 292444354 | 261473458 | 320387522 | 246949666 | 257421506 | 395601250 | 419176482 | 605743682 | 420862754 | 458444546 | 605814594 | 542833026 | 570076226 | 563079618 | 409487650 | 522059714 | 462530306 | 352727650 | 392223234 | 342554082 | 1.668142384 | 0 | 1.952083105 | 0 | 1.502959525 | 0 |
| Q9R0P3 | Esd | S-formylglutathione hydrolase | 944.79 | 56.38 | 31320 | 13 | 25940540 | 26521614 | 25612578 | 20582998 | 29878124 | 51622646 | 40259646 | 62494562 | 48655582 | 44654690 | 56858138 | 55370386 | 65995338 | 55665290 | 47097394 | 97382994 | 77140818 | 65375730 | 94525538 | 61382050 | 1.926988605 | 0 | 2.186055776 | 0 | 3.079351929 | 0 |
| Q8CGK3 | Lonp1 | Lon protease homolog, mitochondrial | 844.89 | 25.08 | 105843 | 17 | 6567353.5 | 4798552.5 | 5443349.5 | 4223956.5 | 7961036.5 | 11063631 | 10650989 | 9479042 | 14065289 | 13852630 | 11698115 | 9505348 | 12599022 | 13572474 | 9914388 | 13300126 | 13773298 | 14880314 | 16567199 | 12039036 | 2.038734717 | 0 | 1.975886597 | 0 | 2.433585164 | 0 |
| P34960 | Mmp12 | Macrophage metalloelastase | 227.05 | 11.21 | 54971 | 5 | 1747290.25 | 1524405.88 | 3815198.75 | 2520478.25 | 2615260 | 5440064 | 4462009 | 7020914.5 | 3938979.25 | 5132736.5 | 7922356 | 5463830 | 8076679 | 8823701 | 10555502 | 8275962 | 3671428 | 11374950 | 9379204 | 6645517 | 2.126767855 | 0 | 3.341511405 | 0 | 3.219196762 | 1 |
| O35129 | Phb2 | Prohibitin-2 | 1197.8 | 63.55 | 33296 | 18 | 4204098 | 4630888.5 | 7077039.5 | 5518796.5 | 9586686 | 15832965 | 12552542 | 17125624 | 15482124 | 10304798 | 24869354 | 23461662 | 24523416 | 22303114 | 14611208 | 20986556 | 11144225 | 12650030 | 17765512 | 10430989 | 2.298638945 | 0 | 3.538928796 | 0 | 2.352778012 | 0 |
| P62320 | Snrpd3 | Small nuclear ribonucleoprotein Sm D3 | 135.47 | 32.54 | 13916 | 3 | 1915268.88 | 6486596.5 | 4338117 | 3730228.5 | 9494886 | 12693957 | 9783863 | 13306313 | 14228131 | 10034420 | 20549694 | 17782678 | 19691886 | 16158086 | 14915846 | 21357836 | 21218120 | 18592006 | 24062486 | 17204178 | 2.312592334 | 0 | 3.431459948 | 0 | 3.945089305 | 0 |
| Q8BVE3 | Atp6v1h | V-type proton ATPase subunit H | 311.71 | 16.36 | 55855 | 7 | 697950.5 | 592864.12 | 538543.75 | 1240671.38 | 717263.56 | 2976704.75 | 3007390.5 | 3972842 | 2819988 | 1650369.75 | 4564762 | 3849971 | 4224204.5 | 5540296 | 5296571.5 | 4642321 | 4408350 | 4816911 | 5232386 | 3798193 | 3.809394684 | 0 | 6.198570609 | 0 | 6.046049019 | 0 |
| P62774 | Mtpn | Myotrophin | 245.57 | 26.27 | 12861 | 3 | 2 | 1095913.5 | 2 | 1919900.5 | 2840651 | 4516728.5 | 7197681 | 7915985 | 7837965 | 3481241 | 8185089 | 8744912 | 9448653 | 6625326 | 4733332 | 1573543.62 | 4006832.75 | 3397590.75 | 5039611.5 | 1852668.62 | 5.284686131 | 0 | 6.443697047 | 0 | 2.709866174 | 4 |
| Q9JIA7 | Sphk2 | Sphingosine kinase 2 | 56.32 | 1.78 | 65618 | 1 | 2 | 2 | 65128.19 | 2 | 170671.02 | 568711.69 | 1039425.06 | 529650.56 | 1452776.62 | 601413 | 271837.66 | 392544.69 | 260894.44 | 675179.81 | 413057.09 | 2 | 306869 | 105176.3 | 2 | 2 | 17.77728715 | 0 | 8.538885506 | 0 | 1.747422375 | 11.5 |
| O70404 | Vamp8 | Vesicle-associated membrane protein 8 | 35.29 | 8.91 | 11451 | 1 | 2 | 2 | 2 | 2 | 92062.99 | 739731.62 | 630138.69 | 199392.16 | 1981702.25 | 2464524.25 | 1465406 | 1959775.38 | 2080573.62 | 2299968.75 | 1480069.5 | 2 | 2250242.75 | 1730968.88 | 731385.5 | 2097928.5 | 65.33533494 | 0 | 100.8547128 | 0 | 73.97039643 | 3 |
| Q6IMX7 | Hspbp1 | Hsp70-binding protein 1 | 163.31 | 12.61 | 39191 | 3 | 2276276 | 2767235 | 2667044.25 | 4319962 | 3298623.75 | 336641.5 | 1060189.5 | 602820.38 | 177643.69 | 2344810.5 | 2 | 1270804.12 | 1660076.38 | 1809124 | 2272670.75 | 2 | 955965.06 | 2 | 1231669.25 | 2074187.5 | 0.295000585 | 1 | 0.457473596 | 0 | 0.278021176 | 0 |
| Q62422 | Ostf1 | Osteoclast-stimulating factor 1 | 235.59 | 21.4 | 23783 | 4 | 2962240.5 | 2067645.25 | 2780713.25 | 1662546.75 | 3031474.5 | 213686.64 | 1157247.12 | 472047.72 | 790177.5 | 2053079.62 | 266553.31 | 2 | 39743.21 | 504152.06 | 1129318.38 | 1339886 | 494839.66 | 1297546 | 1015108 | 1461530.25 | 0.374760569 | 1 | 0.15512418 | 0 | 0.448547001 | 0 |
| Q91WD5 | Ndufs2 | NADH dehydrogenase [ubiquinone] iron-sulfur protein 2, mitoch | 369.62 | 15.33 | 52626 | 6 | 2812738.5 | 3707192.75 | 1914599.5 | 3511760 | 2510454.25 | 1251882.88 | 2417935 | 129536.37 | 1333782 | 1083877.75 | 93646.58 | 157165.88 | 1322492.12 | 1456137.88 | 355400.22 | 2 | 326249.25 | 234073.33 | 70574.06 | 645298.69 | 0.430042447 | 1 | 0.234135878 | 0 | 0.088276948 | 0 |
| Q2EMV9 | Parp14 | Protein mono-ADP-ribosyltransferase PARP14 | 45.15 | 0.55 | 203802 | 1 | 1133401.75 | 1768877 | 1751671.38 | 2284813.75 | 1500462.75 | 2 | 1212867.62 | 991179 | 925143.56 | 870830.75 | 2 | 645806.12 | 2 | 2 | 2 | 2 | 744057.88 | 2 | 2 | 1335922.75 | 0.473979798 | 1 | 0.076525273 | 0 | 0.246466498 | 1 |
| Q8C147 | Dock8 | Dedicator of cytokinesis protein 8 | 91.29 | 1.1 | 238978 | 2 | 2653438.5 | 2026465.25 | 2961496 | 2843847.5 | 2720138 | 2585858.5 | 1289854.75 | 2 | 1773110.25 | 1349753.75 | 675902.75 | 605989.25 | 2 | 2 | 2003271.75 | 909702 | 2 | 1123261.5 | 1156242.38 | 1304775.38 | 0.52997918 | 1 | 0.248774851 | 0 | 0.340314438 | 0 |
| Q9DCL9 | Paics | Bifunctional phosphoribosylaminoimidazole carboxylase/phospl | 469.94 | 21.88 | 47006 | 9 | 14474997 | 19582874 | 11411337 | 13808046 | 16349196 | 8767420 | 7278001 | 11442959 | 9847824 | 5499200 | 9892266 | 11057216 | 10352178 | 10319646 | 9228494 | 10439579 | 8700353 | 10463994 | 10819246 | 13701482 | 0.566407705 | 1 | 0.672381158 | 0 | 0.715684182 | 1 |
| Q9ESY9 | Ifi30 | Gamma-interferon-inducible lysosomal thiol reductase | 248.78 | 20.16 | 27784 | 5 | 8190037.5 | 7046190.5 | 7372462.5 | 5182271 | 4723032.5 | 4666918 | 3878838.5 | 5109326 | 3600058 | 3031614 | 3755892 | 4130874 | 1635717.5 | 4664179.5 | 4194713 | 1341894.12 | 3011742 | 5898206 | 3000273.5 | 3622252.5 | 0.623939172 | 1 | 0.565337374 | 0 | 0.518987859 | 2 |
| P11240 | Cox5a | Cytochrome c oxidase subunit 5A, mitochondrial | 289.11 | 27.4 | 16130 | 5 | 65235450 | 70873722 | 64491278 | 75663674 | 51594214 | 47910514 | 41954354 | 57179606 | 38602382 | 46892634 | 29205790 | 27138892 | 34655898 | 34196502 | 42438350 | 34523678 | 44207730 | 66395698 | 42596682 | 60365170 | 0.70926819 | 1 | 0.511304465 | 0 | 0.756695588 | 4 |
| Q91YT0 | Ndufv1 | NADH dehydrogenase [ubiquinone] flavoprotein 1, mitochondria | 147.55 | 10.13 | 50834 | 3 | 1921741.5 | 1833127.25 | 1783156.75 | 2086493.38 | 1671178.38 | 1571856.75 | 1208416.62 | 1748422.25 | 1242223 | 1251929.12 | 940153.38 | 681869.38 | 1187536.5 | 1028157.44 | 1293136.12 | 561737.31 | 766497.5 | 764194.56 | 796410.06 | 804999.81 | 0.755494456 | 1 | 0.551959974 | 0 | 0.397370863 | 0 |
| P26041 | Msn | Moesin | 2471.44 | 56.67 | 67767 | 41 | 36159634 | 47561142 | 33869290 | 64798106 | 59174970 | 66666686 | 60445490 | 70021162 | 100437402 | 92483298 | 171885314 | 194163906 | 118448938 | 131269890 | 138163378 | 143685234 | 127130450 | 126115922 | 122653026 | 123973450 | 1.614708414 | 1 | 3.121053236 | 0 | 2.664140219 | 0 |
| P84104 | Srsf3 | Serine/arginine-rich splicing factor 3 | 174.38 | 18.29 | 19330 | 4 | 2697017.25 | 5210512.5 | 3169201.25 | 4226097.5 | 6865514.5 | 9545095 | 11787885 | 12703122 | 10964365 | 6348119.5 | 14239307 | 13984641 | 12153264 | 14070471 | 10311008 | 15509445 | 13494506 | 12329943 | 14724140 | 12374145 | 2.316302418 | 1 | 2.921223792 | 0 | 3.086932524 | 0 |
| P54001 | P4ha1 | Prolyl 4-hydroxylase subunit alpha-1 | 305.15 | 13.86 | 60898 | 7 | 2 | 440912.5 | 2 | 2 | 740835.44 | 640468.5 | 1851657.12 | 2365452.75 | 4016741 | 1446448.88 | 2177391.25 | 1925226.25 | 2424180 | 2810348.25 | 1091896.88 | 686970.69 | 471876.88 | 663695.25 | 881963.75 | 2 | 8.733432486 | 1 | 8.825054249 | 0 | 2.288554731 | 6.5 |
| P13864 | Dnmt1 | DNA (cytosine-5)-methyltransferase 1 | 365.76 | 5.93 | 183189 | 8 | 794419.38 | 1027196.06 | 671121.75 | 1295460.12 | 1833908.38 | 2 | 2 | 2 | 876366 | 2 | 475003 | 2 | 2 | 2 | 2 | 2 | 2 | 2 | 2 | 2 | 0.155880029 | 2 | 0.084489874 | 0 | 1.77869E-06 | 0 |
| Q8VCH8 | Ubxn4 | UBX domain-containing protein 4 | 205.24 | 12.45 | 56472 | 4 | 23049706 | 63928290 | 16709360 | 36582354 | 26852198 | 21990278 | 19101856 | 2406963 | 10070016 | 8346575 | 5193491 | 3302100.5 | 4521407 | 4991237.5 | 2630675.75 | 6785781 | 4565369.5 | 3875161.75 | 5193261 | 3748322.75 | 0.370482175 | 2 | 0.123496147 | 0 | 0.144612375 | 0 |
| A2VCX1 | Tiprl | TIP41-like protein | 53.98 | 6.64 | 31225 | 1 | 604306.44 | 999283.69 | 582837.5 | 427136.66 | 481055.47 | 491547.62 | 2 | 2 | 307539.94 | 362953.84 | 2 | 2 | 2 | 2 | 2 | 651163.94 | 2 | 2 | 218316.06 | 2 | 0.375505067 | 2 | 3.23141E-06 | 0 | 0.28096699 | 4 |
| Q2YFS3 | Pilra | Paired immunoglobulin-like type 2 receptor alpha | 84.66 | 8.61 | 33064 | 2 | 2955074 | 1496703.12 | 2295763.25 | 1268255.88 | 1850229.38 | 783272.12 | 881346.38 | 1627209 | 818339.94 | 1027893.81 | 2 | 2 | 497421.94 | 894678.12 | 722047 | 2 | 776842.81 | 2 | 2 | 958563.19 | 0.520783286 | 2 | 0.214285989 | 0 | 0.175897779 | 0 |
| P62075 | Timm13 | Mitochondrial import inner membrane translocase subunit Tim1 | 90.99 | 14.74 | 10458 | 1 | 2951467 | 1980725.62 | 2606312 | 1496193 | 1611425.75 | 1285734.62 | 1170344.25 | 1657627 | 846881.69 | 1278633.38 | 1451130.12 | 802565.5 | 1413385.75 | 2 | 1437448.12 | 2 | 2 | 1006918.06 | 2 | 2 | 0.586055668 | 2 | 0.479473261 | 0 | 0.094581476 | 0 |
| Q91V41 | Rab14 | Ras-related protein Rab-14 | 891.23 | 76.28 | 23897 | 16 | 27776224 | 32426526 | 32192920 | 23327358 | 19961586 | 19227080 | 21799252 | 15683999 | 21269958 | 17139804 | 15968152 | 17232982 | 16460083 | 15416712 | 15020684 | 15331187 | 17008326 | 19564536 | 16123711 | 19719576 | 0.701038166 | 2 | 0.590329372 | 0 | 0.646700708 | 0 |
| Q8JZN5 | Acad9 | Complex I assembly factor ACAD9, mitochondrial | 375.79 | 15.04 | 68722 | 8 | 5865435 | 4934719 | 8300346 | 6313488.5 | 7132175 | 4912872 | 2912914 | 5494705.5 | 5289403 | 4681942.5 | 1002076.81 | 2025586.75 | 3740948 | 2678594.5 | 3827852.5 | 690338.5 | 1965555.75 | 2042634.25 | 2621769.5 | 1648101.88 | 0.71565538 | 2 | 0.407883976 | 0 | 0.275559357 | 0 |
| Q922F4 | Tubb6 | Tubulin beta-6 chain | 1168.36 | 45.19 | 50090 | 19 | 15889175 | 19201524 | 17351284 | 14801321 | 13201588 | 12685784 | 13338699 | 14150360 | 8979408 | 9314541 | 7642172.5 | 5925416 | 10781049 | 10459778 | 8820867 | 5648359.5 | 10516075 | 10643566 | 10261886 | 10610539 | 0.726817956 | 2 | 0.542349942 | 0 | 0.592709174 | 0 |
| Q9CZW5 | Tomm70 | Mitochondrial import receptor subunit TOM70 | 292.04 | 9.66 | 67590 | 6 | 12400570 | 12977864 | 11430952 | 11422214 | 16287770 | 8822925 | 10124398 | 12283153 | 8627559 | 10502097 | 5700361.5 | 6496184.5 | 5898677 | 5873157 | 7224938.5 | 7384014 | 6515596 | 7827071.5 | 8956943 | 10142102 | 0.780542835 | 2 | 0.48347215 | 0 | 0.632766974 | 0 |
| P42932 | Cct8 | T-complex protein 1 subunit theta | 1255.53 | 40.15 | 59555 | 23 | 65590282 | 81012210 | 73498890 | 61645954 | 75480818 | 58934134 | 51355138 | 66749206 | 50582990 | 52395242 | 54396142 | 51981442 | 51978014 | 54507254 | 51336910 | 45046614 | 38367490 | 52167938 | 49501530 | 49083430 | 0.783859578 | 2 | 0.739582698 | 0 | 0.655511049 | 0 |
| P38647 | Hspa9 | Stress-70 protein, mitochondrial | 2285.94 | 52.43 | 73461 | 32 | 72239962 | 83420410 | 65089530 | 65672986 | 90745834 | 99586074 | 82172226 | 98045706 | 114207554 | 110264730 | 140855410 | 128422482 | 117922594 | 127593634 | 112759482 | 135692866 | 136480514 | 125622346 | 116359058 | 110647066 | 1.33700453 | 2 | 1.663853775 | 0 | 1.656557963 | 0 |
| P11762 | Lgals1 | Galectin-1 | 467.38 | 41.48 | 14857 | 6 | 49418446 | 54335358 | 40190382 | 49369270 | 63991886 | 70464026 | 71938010 | 81416442 | 57921298 | 62958006 | 82270634 | 85292090 | 77860538 | 74018418 | 69789218 | 60154234 | 79010378 | 58068374 | 45451058 | 66530626 | 1.339644872 | 2 | 1.512719849 | 0 | 1.201742131 | 6 |
| Q00612 | G6pdx | Glucose-6-phosphate 1-dehydrogenase X | 909.04 | 38.64 | 59263 | 18 | 25215466 | 33521372 | 27085950 | 24584892 | 37825638 | 53109550 | 45463218 | 45528342 | 36914922 | 37657726 | 61965706 | 65615970 | 45767062 | 45735122 | 40423842 | 60551838 | 53268030 | 37265554 | 59610586 | 45628554 | 1.475199779 | 2 | 1.750670534 | 0 | 1.729196684 | 1 |
| Q8BH43 | Wasf2 | Actin-binding protein WASF2 | 424.01 | 15.49 | 54074 | 8 | 3370600.5 | 2098519.75 | 2583947 | 2890427.5 | 2282942.5 | 3271613.75 | 3407723.75 | 3043761.5 | 6061664 | 4171569.5 | 8880675 | 9974627 | 6038283.5 | 8691620 | 3442388.5 | 7164465.5 | 6117966.5 | 5017807 | 6840514 | 3204721.75 | 1.50882147 | 2 | 2.79951383 | 0 | 2.143092219 | 1 |
| P06151 | Ldha | L-lactate dehydrogenase A chain | 1308.42 | 62.95 | 36499 | 21 | 107720930 | 114408562 | 113041034 | 111911970 | 185814290 | 178791042 | 179128546 | 224662658 | 257644754 | 209821682 | 378680290 | 430477794 | 234942530 | 269703106 | 193557730 | 322319906 | 219004674 | 167771970 | 288724514 | 154744930 | 1.659115207 | 2 | 2.381686056 | 0 | 1.821096298 | 2 |
| Q9Z0X1 | Aifm1 | Apoptosis-inducing factor 1, mitochondrial | 773.7 | 29.58 | 66765 | 14 | 3788268.75 | 2802908.5 | 3033380.5 | 2710384.75 | 3952886.25 | 5942557 | 5319947.5 | 4914535 | 11501732 | 3378811.5 | 14513321 | 14349574 | 8970858 | 10582814 | 9016713 | 11157832 | 5260577.5 | 7356709.5 | 9470906 | 7630630 | 1.906797 | 2 | 3.526147093 | 0 | 2.509644203 | 0 |
| Q9R0N0 | Galk1 | Galactokinase | 315.15 | 14.54 | 42295 | 7 | 2 | 1757560.62 | 2253764.75 | 1047678.38 | 4286003 | 2638388 | 3953308 | 6729657 | 5417976 | 4555444.5 | 9983930 | 8559553 | 7913805.5 | 5801622.5 | 5493793 | 4574603 | 2306349.5 | 3694550 | 4247227 | 2753370.5 | 2.492750314 | 2 | 4.039878935 | 0 | 1.880800807 | 4 |
| P16110 | Lgals3 | Galectin-3 | 533.1 | 38.64 | 27515 | 9 | 3127265 | 5263680 | 5333368.5 | 4447551 | 19641258 | 20913506 | 11014831 | 27253018 | 28267058 | 18261660 | 104838130 | 112329122 | 53973854 | 64469754 | 27105718 | 112963626 | 55047658 | 31998546 | 130974898 | 27602866 | 2.795592271 | 2 | 9.592346625 | 0 | 9.483152152 | 0 |
| P24051 | Rps27l | Small ribosomal subunit protein eS27-like | 120.85 | 25 | 9477 | 2 | 2 | 2 | 2 | 2 | 2 | 1517504.62 | 1607556.12 | 2860443 | 3602225.25 | 2 | 3459366 | 3101743.5 | 1151555.88 | 1696551.62 | 752189.31 | 2 | 2 | 2 | 2 | 2 | 958773.099 | 2.5 | 1016140.631 | 0 | 1 | 12.5 |
| Q60692 | Psmb6 | Proteasome subunit beta type-6 | 193.33 | 16.81 | 25379 | 4 | 5311137.5 | 6450081 | 4340186 | 4557643 | 167663.27 | 2 | 2992026 | 130886.95 | 2218066 | 2801314.75 | 12489238 | 11316821 | 12225833 | 13102580 | 6924246 | 11425199 | 8095560.5 | 9920793 | 9014656 | 9019310 | 0.390954471 | 3 | 2.691674102 | 0 | 2.279549518 | 0 |
| Q9WVJ3 | Cpq | Carboxypeptidase Q | 247.46 | 12.98 | 51813 | 5 | 952888.88 | 1670381.75 | 1302459 | 1054140.5 | 1634993.75 | 2 | 145504.2 | 621619.94 | 529590.56 | 1438468.62 | 2 | 2 | 2 | 2 | 2 | 2 | 2 | 2 | 2 | 2 | 0.413490794 | 3 | 1.51175E-06 | 0 | 1.51175E-06 | 0 |
| Q9JI75 | Nqo2 | Ribosyldihydronicotinamide dehydrogenase [quinone] | 34.95 | 4.33 | 26248 | 1 | 1507739.25 | 1286004.5 | 1514669.88 | 1112830.5 | 1040020.69 | 2 | 1151715 | 1040033.25 | 578664.38 | 810650.25 | 2 | 2 | 765760 | 2 | 2 | 2 | 2 | 2 | 2 | 2 | 0.554235893 | 3 | 0.118516733 | 0 | 1.54768E-06 | 0 |
| O55101 | Syngr2 | Synaptogyrin-2 | 42.51 | 4.46 | 24778 | 1 | 883481.44 | 879555 | 1439341.38 | 1155733.38 | 943681.06 | 2 | 650710.81 | 859400 | 409207.38 | 1110826.88 | 2 | 2 | 2 | 586227.81 | 798507.56 | 2 | 2 | 1309337.38 | 2 | 2 | 0.571532592 | 3 | 0.261183634 | 0 | 0.246962785 | 4 |
| P56391 | Cox6b1 | Cytochrome c oxidase subunit 6B1 | 240.41 | 50 | 10071 | 4 | 4305351 | 4282084.5 | 3941436.5 | 4241617 | 2150465.5 | 1820962.12 | 1739984.5 | 2265527.5 | 2294709 | 3133573.5 | 2 | 2 | 2 | 2 | 1949332.25 | 1294350.62 | 2 | 1680758 | 2238895.25 | 2 | 0.594830278 | 3 | 0.103025471 | 0 | 0.275567909 | 1 |
| P19973 | Lsp1 | Lymphocyte-specific protein 1 | 535.53 | 29.7 | 36714 | 8 | 14110519 | 17164270 | 11376011 | 13671353 | 11952719 | 10026948 | 7991208 | 5507593.5 | 11162857 | 14086948 | 78403722 | 83259778 | 52048194 | 56271646 | 31221914 | 67572890 | 47556506 | 26809206 | 51803542 | 33848426 | 0.714399794 | 3 | 4.411656078 | 0 | 3.333445576 | 0 |
| Q63945 | Set | Protein SET | 235.8 | 21.45 | 33406 | 5 | 5081914 | 5875709.5 | 7138754.5 | 4866965 | 6238439.5 | 4204240 | 4302546 | 6106717 | 2825121.75 | 4059759 | 2599681.25 | 2759130.5 | 2734282.25 | 2894488.5 | 2959823.5 | 3547256.25 | 2725510.25 | 1828033.75 | 2538033.5 | 2779915.75 | 0.736201078 | 3 | 0.477621734 | 0 | 0.459518165 | 0 |
| Q9ERS2 | Ndufa13 | NADH dehydrogenase [ubiquinone] 1 alpha subcomplex subun | 128.28 | 22.92 | 16860 | 3 | 5489364.5 | 5925315.5 | 4368043.5 | 4973944.5 | 3485926.5 | 4244008.5 | 4244567.5 | 3880203 | 2953882.5 | 2774555.75 | 2627483.5 | 1704241 | 2391005.75 | 2071138.25 | 2053243.38 | 1673015.88 | 1649582.38 | 3156083.5 | 391454.22 | 1767813.75 | 0.746504969 | 3 | 0.447440223 | 0 | 0.356312924 | 0 |
| P97371 | Psme1 | Proteasome activator complex subunit 1 | 240.45 | 17.67 | 28673 | 4 | 14518968 | 10753305 | 12940395 | 15649119 | 15157189 | 17291136 | 18487774 | 18334312 | 13755834 | 17312258 | 4928276.5 | 6329604 | 7259439 | 5337163 | 8145105 | 6145879 | 5158249 | 7575598 | 3609753 | 5843317 | 1.234172382 | 3 | 0.463634631 | 0 | 0.410507336 | 0 |
| P51410 | Rpl9 | Large ribosomal subunit protein uL6 | 698.08 | 67.19 | 21881 | 12 | 1874341.5 | 11892226 | 7061566 | 12254184 | 11707654 | 16075208 | 17614160 | 22591100 | 20844160 | 8027666 | 15708552 | 14681894 | 21409452 | 22504406 | 13824138 | 20707986 | 21051774 | 13776298 | 24579566 | 12924292 | 1.901146421 | 3 | 1.96759317 | 0 | 2.077248832 | 0 |
| Q8BML9 | Qars1 | Glutamine--tRNA ligase | 248.89 | 9.16 | 87677 | 6 | 382517.28 | 2 | 1134130.5 | 1931822.38 | 1593221.75 | 2075795.75 | 2063766.88 | 3081453 | 2708368 | 1072202.75 | 2943539.25 | 3539028 | 4154257.5 | 4735130 | 2986355.5 | 3575042.5 | 4044856.25 | 2637241.25 | 5361362.5 | 2 | 2.182121044 | 3 | 3.641298059 | 0 | 3.09786845 | 4.5 |
| P01897 | H2-L | H-2 class I histocompatibility antigen, L-D alpha chain | 327.25 | 21.27 | 40711 | 6 | 552173.75 | 2 | 2 | 553583.5 | 1461362.62 | 1057907.25 | 738171.62 | 1113074.62 | 2279357.5 | 1638222 | 4535230.5 | 3432047.5 | 3455267.25 | 3777174.75 | 3859601.25 | 4108263 | 4436619.5 | 6588569 | 3720116.25 | 3317544.25 | 2.659292397 | 3 | 7.424387063 | 0 | 8.636557144 | 0 |
| P28867 | Prkcd | Protein kinase C delta type | 192.93 | 6.53 | 77547 | 4 | 600645.62 | 2 | 2 | 543946.88 | 2565137.75 | 2390181 | 918683.12 | 2725531.5 | 3589544.5 | 1628294.38 | 3479009.25 | 3536567 | 3942442.5 | 3567291 | 2578223 | 3383170 | 1739306.88 | 1814074.5 | 2782766.75 | 2334025.25 | 3.033164572 | 3 | 4.610446894 | 0 | 3.249112354 | 3 |
| Q8R1Q8 | Dync1li1 | Cytoplasmic dynein 1 light intermediate chain 1 | 196.89 | 7.07 | 56614 | 3 | 2 | 2 | 2 | 2 | 572640.12 | 1033193.12 | 2 | 1105852.12 | 729851.94 | 585884 | 1620736.88 | 1790539 | 1380740.62 | 1500818.12 | 628672.25 | 2 | 384929.12 | 2 | 1479280.62 | 261687.52 | 6.032994887 | 3 | 12.08684117 | 0 | 3.712404155 | 8 |
| P14685 | Psmd3 | 26S proteasome non-ATPase regulatory subunit 3 | 159.49 | 4.91 | 60718 | 3 | 2 | 2 | 2 | 182490.91 | 62945.85 | 2 | 1635226.62 | 1059899 | 1840489.25 | 403773.41 | 406999 | 592876.31 | 1209183 | 1451740.5 | 707476.94 | 123247.55 | 431152.19 | 660757.06 | 356804.16 | 146259.58 | 20.12440815 | 3.5 | 17.79753353 | 0 | 7.000493883 | 2 |
| P18886 | Cpt2 | Carnitine O-palmitoyltransferase 2, mitochondrial | 47.37 | 1.52 | 74110 | 1 | 1313214.62 | 1845231.38 | 1696600.88 | 1516537.25 | 1425697.62 | 2 | 2 | 1824995.12 | 2 | 2 | 546640.69 | 2 | 2 | 2 | 2 | 590470.06 | 1307845.62 | 2 | 2 | 2 | 0.234056326 | 4 | 0.070107597 | 0 | 0.243459418 | 0 |
| P54227 | Stmn1 | Stathmin | 243.32 | 25.5 | 17274 | 4 | 1345075 | 1039445.12 | 2 | 1366825.5 | 998115.75 | 2 | 411985.69 | 2 | 782392.44 | 303823.5 | 7008651.5 | 9875368 | 7318129 | 6250941 | 5507581.5 | 6166086 | 4705661.5 | 3624702.5 | 5571385.5 | 5006386 | 0.315447349 | 4 | 7.57152297 | 0 | 5.27937991 | 0 |
| Q9Z0E6 | Gbp2 | Guanylate-binding protein 2 | 45 | 2.04 | 66739 | 1 | 4477860 | 4877511 | 4182352.75 | 4710503.5 | 6130350 | 741177.5 | 556665.44 | 5856077 | 474604.62 | 402787.31 | 2457598 | 1918367.12 | 2 | 1739172.88 | 2281119.25 | 809392.12 | 1346131.62 | 2245475.25 | 1211270.12 | 2 | 0.329441369 | 4 | 0.344411372 | 0 | 0.230213234 | 0 |
| P42567 | Eps15 | Epidermal growth factor receptor substrate 15 | 57.54 | 1.34 | 98471 | 1 | 478077.12 | 170217.27 | 593845.94 | 251958.88 | 308871.16 | 78573.1 | 2 | 2 | 557743.19 | 2 | 2 | 2 | 2 | 2 | 2 | 2 | 373569.22 | 2 | 2 | 210251.42 | 0.352929976 | 4 | 5.5464E-06 | 0 | 0.323813774 | 4 |
| A2A5R2 | Arfgef2 | Brefeldin A-inhibited guanine nucleotide-exchange protein 2 | 193.33 | 2.85 | 202240 | 4 | 2015456.25 | 1226551.88 | 2048754.25 | 1024803.88 | 1280350.25 | 1358076.12 | 1153857.88 | 67945.49 | 724778.44 | 2 | 2 | 2 | 982825 | 753398.56 | 279739.66 | 2 | 800008.19 | 1149364.62 | 2 | 1405308.75 | 0.435057432 | 4 | 0.265401445 | 0 | 0.44164329 | 4 |
| Q3UIA2 | Arhgap17 | Rho GTPase-activating protein 17 | 45.23 | 1.06 | 92202 | 1 | 1606328.12 | 1761663.62 | 705776.38 | 902074.44 | 840755.69 | 1164809.38 | 2 | 784261.38 | 513321.28 | 463219.31 | 568867.38 | 540131.12 | 461600.47 | 502356.47 | 484372.94 | 2 | 2 | 451081.47 | 2 | 2 | 0.502976693 | 4 | 0.43966048 | 0 | 0.07755211 | 0 |
| P09581 | Csf1r | Macrophage colony-stimulating factor 1 receptor | 77.34 | 1.74 | 109179 | 2 | 1769150.25 | 3220037.5 | 2556959 | 2367380.75 | 4335557 | 1899541.75 | 1447376.88 | 638239.19 | 1873098 | 2488783.25 | 1497542 | 1429790.5 | 1310795.38 | 1335856.62 | 2 | 1212014.12 | 1254697.38 | 1418438.88 | 1076713.12 | 1134834 | 0.585794762 | 4 | 0.391182079 | 0 | 0.42786591 | 0 |
| Q9CQQ7 | Atp5pb | ATP synthase F(0) complex subunit B1, mitochondrial | 219.65 | 20.7 | 28949 | 4 | 6840195.5 | 5844941 | 6928086.5 | 7313789 | 9111476 | 1827050.5 | 2142394.75 | 5548358.5 | 5265286.5 | 7746120 | 3637253 | 3893909.5 | 5252252 | 4264695 | 5193827.5 | 4000270.5 | 1617523.75 | 7188362 | 4962959 | 6273238 | 0.625143048 | 4 | 0.617171758 | 0 | 0.66712991 | 4 |
| Q9D7S7 | Rpl22l1 | Ribosomal protein eL22-like 1 | 127.26 | 19.67 | 14467 | 2 | 1388236.12 | 1285413.62 | 2152286.25 | 1800872.88 | 2716735.25 | 806235.56 | 904295.88 | 1725890.62 | 1619530.75 | 963340.56 | 806782 | 830509.69 | 1229249.38 | 867466.19 | 754241.12 | 1673295.38 | 2 | 628517.62 | 1086537.5 | 2 | 0.644219505 | 4 | 0.480358237 | 0 | 0.362641248 | 2 |
| Q9CQN1 | Trap1 | Heat shock protein 75 kDa, mitochondrial | 776.83 | 25.64 | 80209 | 13 | 5324754 | 5963155.5 | 7889784.5 | 8246447 | 7963160 | 3295123.5 | 2360031.5 | 4711253.5 | 6236478 | 7744682 | 1217339.5 | 3922995.25 | 3782413.5 | 2541079 | 4951045 | 2380164.5 | 3701084 | 2673585.5 | 5167504 | 3413228 | 0.688031238 | 4 | 0.463863357 | 0 | 0.489880989 | 0 |
| P10810 | Cd14 | Monocyte differentiation antigen CD14 | 706.03 | 34.7 | 39204 | 11 | 10942571 | 10061821 | 15048407 | 11770358 | 10768785 | 6218570 | 7596580.5 | 8346176 | 6411814.5 | 14659994 | 1973751.75 | 1787679.5 | 1974995.75 | 3591940.5 | 3960309 | 1611017.12 | 2270443.5 | 2640530.25 | 1191127.88 | 2485951 | 0.737868272 | 4 | 0.226800411 | 0 | 0.174069495 | 0 |
| Q3T1J9 | Mob1a | MOB kinase activator 1A | 68.83 | 8.33 | 25080 | 2 | 2750918.5 | 2277207.25 | 2351624.5 | 3883126.25 | 3112418.25 | 1804570.12 | 2368626 | 2132304 | 2017278.12 | 2473419.5 | 1210335.38 | 1454878.12 | 1510869.38 | 1467715 | 2223966 | 1416230.12 | 2307772.5 | 2924570.5 | 1929644.5 | 2895978.75 | 0.751024444 | 4 | 0.547311482 | 0 | 0.798188599 | 7 |
| Q69ZK0 | Prex1 | Phosphatidylinositol 3,4,5-trisphosphate-dependent Rac exchar | 198.61 | 3.03 | 184935 | 4 | 4671311.5 | 4800441 | 4561890.5 | 4068023.25 | 3112972 | 2852672.25 | 3940321 | 4242052 | 1586398.38 | 3312852.75 | 470543.62 | 911467.69 | 1105735.62 | 1729260.62 | 3009376 | 2 | 2583738 | 1700429.88 | 2876448.75 | 1943441 | 0.751099132 | 4 | 0.340631948 | 0 | 0.429140461 | 0 |
| Q8BX70 | Vps13c | Intermembrane lipid transfer protein VPS13C | 479.16 | 3.5 | 420089 | 9 | 4514338 | 5282047.5 | 6020406.5 | 4872832.5 | 3780230 | 3147954.5 | 4148494 | 5005071 | 3372048 | 3378204 | 928694.62 | 980428.62 | 3563590 | 3125135.5 | 1476465.62 | 2935037.25 | 3136278.75 | 3676333.25 | 2859568 | 3263125.25 | 0.778581315 | 4 | 0.411703076 | 0 | 0.648567097 | 0 |
| P08752 | Gnai2 | Guanine nucleotide-binding protein G(i) subunit alpha-2 | 832.73 | 38.31 | 40489 | 12 | 38154990 | 37151450 | 43610278 | 33984730 | 42021894 | 34239930 | 28910452 | 37368082 | 31825398 | 36520238 | 22027290 | 16357914 | 20013668 | 17289804 | 28910366 | 13794714 | 15245395 | 13513181 | 15860340 | 13804663 | 0.866310306 | 4 | 0.536616297 | 0 | 0.370495869 | 0 |
| P26638 | Sars1 | Serine--tRNA ligase, cytoplasmic | 893.22 | 34.96 | 58389 | 19 | 18166996 | 13064344 | 20896140 | 19943166 | 25177930 | 21428552 | 20693212 | 22863536 | 30551008 | 31616026 | 51887238 | 50066766 | 51460562 | 46346346 | 58075158 | 47367470 | 60992722 | 62088558 | 55315782 | 49605318 | 1.307498158 | 4 | 2.651309465 | 0 | 2.831608043 | 0 |
| Q07797 | Lgals3bp | Galectin-3-binding protein | 1071.16 | 28.77 | 64491 | 15 | 14545500 | 15074563 | 14051970 | 16025310 | 9934597 | 18567562 | 19561450 | 12657042 | 19593790 | 21898162 | 2355002.75 | 6603900.5 | 5928308.5 | 5573111.5 | 4431728 | 4639060 | 6752272.5 | 12554632 | 8493354 | 10972786 | 1.325225263 | 4 | 0.357480364 | 0 | 0.623451027 | 2 |
| Q3TW96 | Uap1l1 | UDP-N-acetylhexosamine pyrophosphorylase-like protein 1 | 595.35 | 23.08 | 56614 | 11 | 4698977 | 6107068.5 | 5704969.5 | 4713651.5 | 10918309 | 9752193 | 7329518 | 11749268 | 7791612 | 7035345.5 | 18799336 | 16035137 | 12304246 | 14502592 | 14716110 | 16319462 | 11120095 | 9030835 | 8932153 | 7711906 | 1.358241912 | 4 | 2.375555462 | 0 | 1.652443502 | 3 |
| Q11011 | Npepps | Puromycin-sensitive aminopeptidase | 391.25 | 8.04 | 103325 | 7 | 2606534.75 | 3479656 | 2591371.25 | 1110200.12 | 3054948 | 3938569.5 | 2762086.5 | 2776830.25 | 5591779 | 3743977 | 7148321 | 6258970.5 | 5569901.5 | 3661506 | 4574184 | 7530916.5 | 4546659 | 2958738 | 5067262.5 | 2472954.75 | 1.464896589 | 4 | 2.11893617 | 0 | 1.757925745 | 6 |
| P17182 | Eno1 | Alpha-enolase | 1996.97 | 65.21 | 47141 | 28 | 18871678 | 20500014 | 31773610 | 13039551 | 31881922 | 24144164 | 30472144 | 38017822 | 39410174 | 51412194 | 53396754 | 57394990 | 46220606 | 38302958 | 50591174 | 65843842 | 43127166 | 35574562 | 41043554 | 31954728 | 1.580611661 | 4 | 2.118663864 | 0 | 1.874299101 | 0 |
| A1L314 | Mpeg1 | Macrophage-expressed gene 1 protein | 665.85 | 19.35 | 78390 | 12 | 4092371.75 | 7790929 | 3064948.5 | 6733613.5 | 6004197.5 | 6303170 | 6677600.5 | 8248892 | 14188936 | 10183316 | 2868876 | 3017573 | 2500374.5 | 2445594.25 | 1998521.88 | 3412769.25 | 3295437 | 2346290.25 | 3529577.25 | 3842626 | 1.647107392 | 4 | 0.463444041 | 0 | 0.593320234 | 4 |
| Q8CDN6 | Txnl1 | Thioredoxin-like protein 1 | 173.37 | 18.34 | 32237 | 4 | 340801.19 | 1385897.75 | 980847.62 | 2 | 877980.75 | 1058351.88 | 1215002.25 | 1062080 | 1864510.75 | 1159168 | 7575366 | 6509465.5 | 3940408.5 | 5907890 | 2748887.75 | 7773859 | 4149049.5 | 5284128 | 4080512.5 | 7419201 | 1.773549267 | 4 | 7.441584057 | 0 | 8.006279553 | 0 |
| Q60875 | Arhgef2 | Rho guanine nucleotide exchange factor 2 | 637.51 | 16.24 | 111974 | 14 | 1510922.88 | 2733868.5 | 2547543.5 | 2325896.75 | 3608813.5 | 4840334.5 | 3543614.25 | 2995796.25 | 9012949 | 2712715.25 | 6179504 | 9439668 | 4405860.5 | 9257214 | 3771255.5 | 5120592.5 | 4070074.75 | 3682908 | 3869355.5 | 11118500 | 1.815457478 | 4 | 2.597107315 | 0 | 2.189151564 | 0 |
| Q9Z1D1 | Eif3g | Eukaryotic translation initiation factor 3 subunit G | 267.82 | 22.5 | 35638 | 6 | 813718.94 | 645834.88 | 1068401.25 | 1489206.38 | 1473741.88 | 2024314.62 | 3544878.5 | 679236.44 | 2623053 | 1785731.62 | 3110392.5 | 3249961.5 | 3699167.5 | 3425830 | 5599095.5 | 2150807.5 | 4024842 | 5127011.5 | 159070.23 | 2042310.25 | 1.940885413 | 4 | 3.475647968 | 0 | 2.459347883 | 5 |
| Q8BL97 | Srsf7 | Serine/arginine-rich splicing factor 7 | 354.23 | 21.72 | 30818 | 7 | 833677.62 | 1773750 | 1791820.12 | 2612234 | 7085022.5 | 5899477.5 | 2003429.75 | 7935205 | 9005497 | 3014007.5 | 21510646 | 16930490 | 15133388 | 18891270 | 7563547 | 16229724 | 15518916 | 5446773.5 | 17603922 | 10077680 | 1.976207454 | 4 | 5.677247326 | 0 | 4.602347816 | 1 |
| P62315 | Snrpd1 | Small nuclear ribonucleoprotein Sm D1 | 55.54 | 10.92 | 13282 | 1 | 400947.28 | 812720.31 | 2 | 2 | 1321106.25 | 1614984.5 | 530702.5 | 1017905.88 | 1379937.12 | 837957.12 | 3986705.25 | 3772382.25 | 2456251.75 | 2700423.5 | 2095967 | 3215754.5 | 3325476.75 | 1378213.5 | 2812080 | 2614185.75 | 2.123060662 | 4 | 5.922305897 | 0 | 5.265041492 | 0 |
| Q61792 | Lasp1 | LIM and SH3 domain protein 1 | 480.51 | 33.46 | 29994 | 9 | 3079353.25 | 2501092.5 | 2487438.75 | 9394012 | 7429395.5 | 19475224 | 17196160 | 4064809 | 14479289 | 5231668.5 | 24225150 | 31162676 | 12979984 | 18550266 | 13604862 | 22630266 | 21510112 | 19416878 | 17571968 | 26931716 | 2.428445679 | 4 | 4.038478115 | 0 | 4.341315027 | 0 |
| P17742 | Ppia | Peptidyl-prolyl cis-trans isomerase A | 928.91 | 82.32 | 17971 | 14 | 1135532.38 | 3290772.5 | 4888471 | 468504.22 | 21764478 | 11251121 | 7452503 | 12404534 | 27336840 | 19206822 | 131987122 | 134021498 | 91648154 | 97204114 | 55781694 | 57371822 | 65268026 | 36009546 | 63481854 | 35583158 | 2.461405332 | 4 | 16.18633503 | 0 | 8.16902441 | 0 |
| P70460 | Vasp | Vasodilator-stimulated phosphoprotein | 244.15 | 12.53 | 39667 | 5 | 1923810 | 1188995.38 | 2 | 2 | 35870.67 | 2173413 | 3868925 | 192777.05 | 5663243 | 781271.62 | 6754431 | 6023641 | 2817622.5 | 5124684.5 | 2814825.5 | 2697295.75 | 1448203.62 | 1583815.25 | 1927797.75 | 2 | 4.026966687 | 4 | 7.474625597 | 0 | 2.431848981 | 6 |
| Q9CR67 | Tmem33 | Transmembrane protein 33 | 159.4 | 13.36 | 28031 | 3 | 1440172.88 | 673541.94 | 2317869.25 | 1989908.62 | 2560531 | 1582712.25 | 21186962 | 9599845 | 2980248.25 | 2321192.5 | 7211938 | 8409346 | 12201652 | 5366078.5 | 47439698 | 1816937.5 | 15806790 | 5543150 | 2874665 | 12352848 | 4.194039261 | 4 | 8.976675556 | 0 | 4.2745813 | 3 |
| Q5I0H4 | Tmco1 | Calcium load-activated calcium channel | 209.16 | 19.68 | 21175 | 3 | 128589.49 | 2 | 198485.47 | 324438.53 | 141879.17 | 123027.49 | 455134.09 | 1157054.38 | 2294833.5 | 545733.12 | 2261938.75 | 2089909.75 | 1874141.5 | 2501004 | 387632.5 | 1678755 | 1320605.5 | 1044052.25 | 2524544.5 | 1493966.25 | 5.767347338 | 4 | 11.48813694 | 0 | 10.16130295 | 0 |
| O88796 | Rpp30 | Ribonuclease P protein subunit p30 | 72.83 | 4.85 | 29473 | 1 | 2 | 2 | 53594.14 | 2 | 94948.88 | 177717.48 | 2 | 148017.2 | 393510.56 | 81918.53 | 204812.55 | 139676.14 | 166427.78 | 214165.19 | 237648.52 | 2 | 157941.02 | 89414.22 | 355417.72 | 2 | 5.393275365 | 4.5 | 6.480892166 | 0 | 4.057764636 | 8 |
| O89023 | Tpp1 | Tripeptidyl-peptidase 1 | 326.12 | 14.41 | 61342 | 5 | 1064131.38 | 716093.06 | 1706556 | 782679.19 | 474137.03 | 2 | 2 | 529322.62 | 2 | 1143496.88 | 2 | 2 | 2 | 2 | 2 | 2 | 2 | 2 | 2 | 2 | 0.352649186 | 5 | 2.10811E-06 | 0 | 2.10811E-06 | 0 |
| P26883 | Fkbp1a | Peptidyl-prolyl cis-trans isomerase FKBP1A | 164 | 34.26 | 11923 | 3 | 1820123.38 | 1370842.75 | 1429936.5 | 1151773.5 | 1995983.62 | 2 | 2 | 2217760 | 707807.06 | 765213.62 | 14284028 | 16163495 | 10632491 | 10729690 | 6113886 | 5814974 | 6636910.5 | 6188723 | 8938636 | 6504553 | 0.475086411 | 5 | 7.456059586 | 0 | 4.387345771 | 0 |
| Q8R146 | Apeh | Acylamino-acid-releasing enzyme | 37.74 | 1.09 | 81581 | 1 | 4977963 | 5092038.5 | 5290853.5 | 1476731.75 | 2520435.5 | 1775069.88 | 1294977.12 | 2529241 | 1609181.12 | 2424875.5 | 1016858.25 | 1153205 | 827822.19 | 1046949.12 | 631601.19 | 1200745.75 | 2283224.75 | 991735.19 | 2503434.75 | 2 | 0.497640952 | 5 | 0.241576112 | 0 | 0.360529725 | 2 |
| O35892 | Sp100 | Nuclear autoantigen Sp-100 | 164.59 | 6.22 | 54727 | 3 | 4150582.75 | 2178992.5 | 4109330.5 | 6343617 | 5014300 | 1640700.5 | 4085536.5 | 6142053.5 | 320638.41 | 463041.81 | 1044814.56 | 926210.31 | 1916685.38 | 2 | 2 | 1704083.62 | 2421518.5 | 3541749 | 1817788.75 | 495062.78 | 0.580450227 | 5 | 0.178361511 | 0 | 0.457874194 | 2 |
| P28271 | Aco1 | Cytoplasmic aconitate hydratase | 41.59 | 1.35 | 98126 | 1 | 1090358 | 1434811.5 | 927537 | 960506.75 | 831564.44 | 913840.81 | 726633 | 1103833.62 | 542521.94 | 2 | 654063.81 | 627699.31 | 800134.31 | 625394 | 2 | 948901.38 | 2 | 2 | 2 | 2 | 0.626686499 | 5 | 0.516188405 | 0 | 0.180924614 | 2 |
| Q8VCN9 | Tbcc | Tubulin-specific chaperone C | 33.53 | 2.35 | 38125 | 1 | 2378876.75 | 3045797.5 | 1790651.25 | 2820224.75 | 1977941.62 | 2164291 | 1697221.12 | 2402291.75 | 268861.38 | 1211373.25 | 2 | 1478138.38 | 1294026.62 | 1655930.25 | 416088.31 | 2069576.75 | 1360979.75 | 1970274.62 | 2048546.5 | 2231103.75 | 0.64461179 | 5 | 0.403228771 | 0 | 0.805800801 | 7 |
| Q6AYK3 | Isyna1 | Inositol-3-phosphate synthase 1 | 326.71 | 10.59 | 60884 | 7 | 7014309 | 8153096 | 4548130.5 | 5811186 | 4295313.5 | 5693363 | 4435100.5 | 4835266.5 | 3437322.25 | 3552967.25 | 14559522 | 12927007 | 9879809 | 10095990 | 9833470 | 12305065 | 11576954 | 3107036 | 9374090 | 7516114.5 | 0.73616772 | 5 | 1.921257151 | 0 | 1.471370398 | 6 |
| P32577 | Csk | Tyrosine-protein kinase CSK | 183.41 | 9.78 | 50746 | 4 | 4502497 | 5576313 | 3487286.5 | 6605307 | 5559377 | 3901235.5 | 3947326 | 5013808.5 | 3559012.5 | 3115808.5 | 2683565.75 | 1975767.88 | 2773500.5 | 2634539 | 2565015.25 | 3493781.5 | 2602728 | 4366455 | 2659443 | 2845839.5 | 0.759292591 | 5 | 0.490944625 | 0 | 0.620589298 | 2 |

| | | | | | | | | | | | | | | | | | | | | | | | | | | | | | | | | |
|---|---|---|---|---|---|---|---|---|---|---|---|---|---|---|---|---|---|---|---|---|---|---|---|---|---|---|---|---|---|---|---|---|
| P21107 | Tpm3 | Tropomyosin alpha-3 chain | 736.17 | 26.32 | 32994 | 13 | 19450480 | 18463368 | 16331821 | 17673392 | 17797538 | 13533726 | 13052179 | 20647862 | 9767908 | 12073933 | 11375328 | 12365018 | 11207079 | 9592222 | 5648078.5 | 13363112 | 12604268 | 7993012 | 15140918 | 11118335 | 0.769931192 | 5 | 0.559402898 | 0 | 0.671220774 | 0 |
| Q63486 | Rraga | Ras-related GTP-binding protein A | 113.3 | 6.71 | 36566 | 2 | 7256780 | 6091934 | 6348833 | 5655308 | 6408092 | 5148404 | 5582542.5 | 9704875 | 5632498 | 4930169.5 | 3929210.25 | 4747738 | 4477466.5 | 2541026.25 | 2180480.75 | 2 | 2523466.25 | 2560047 | 3546446.25 | 2101292.75 | 0.975993852 | 5 | 0.562827102 | 0 | 0.337875765 | 0 |
| Q8CAQ8 | Immt | MICOS complex subunit Mic60 | 1805.59 | 39.1 | 83900 | 29 | 15482612 | 23295006 | 17427990 | 24174128 | 21498916 | 23632562 | 23471872 | 21145300 | 34057606 | 30306002 | 39647094 | 34344882 | 35009886 | 38469806 | 26522260 | 41791870 | 32802866 | 37846578 | 31860216 | 31742002 | 1.301679394 | 5 | 1.707854635 | 0 | 1.727972726 | 0 |
| Q8BGQ7 | Aars1 | Alanine--tRNA ligase, cytoplasmic | 1040.82 | 23.86 | 106909 | 19 | 14857398 | 15724374 | 22310118 | 16816902 | 32632782 | 28528192 | 24486130 | 24159734 | 27561500 | 29517566 | 37923506 | 35735354 | 40622742 | 37853658 | 40428814 | 41443006 | 46812290 | 47575250 | 44113666 | 47680474 | 1.311814122 | 5 | 1.881582103 | 0 | 2.224166359 | 0 |
| P18760 | Cfl1 | Cofilin-1 | 674.92 | 74.1 | 18560 | 10 | 15788739 | 14159229 | 26441510 | 19707402 | 53186534 | 31805398 | 21150924 | 40351130 | 66201578 | 44425730 | 118301770 | 108682354 | 71478322 | 67099610 | 67825754 | 116420698 | 70246474 | 78065674 | 59094290 | 73162114 | 1.577424 | 5 | 3.35223055 | 0 | 3.070689717 | 0 |
| Q9EQ06 | Hsd17b11 | Estradiol 17-beta-dehydrogenase 11 | 125.95 | 9.4 | 32881 | 2 | 985710.25 | 1361469.88 | 824658.75 | 481430.25 | 784571.25 | 1359958.38 | 1593604.62 | 720085.56 | 1831164.75 | 1504905.75 | 345350.59 | 2 | 74769.93 | 2 | 417029.38 | 938852.94 | 2 | 2 | 2 | 282193.94 | 1.579533841 | 5 | 0.18863993 | 0 | 0.275145741 | 3 |
| Q9DCE9 | Igtp | Immunity-related GTPase family M protein 3 | 37.38 | 2.13 | 48494 | 1 | 626401.06 | 114323.32 | 471711.84 | 521948.31 | 738063.31 | 690533.44 | 1650147 | 227845.33 | 2173068 | 1790851.25 | 2 | 2 | 2 | 2 | 2 | 2 | 2 | 2 | 2 | 2 | 2.642096191 | 5 | 4.04457E-06 | 0 | 4.04457E-06 | 0 |
| Q80VA0 | Galnt7 | N-acetylgalactosaminyltransferase 7 | 129 | 4.87 | 75419 | 3 | 824577.31 | 2 | 296602.88 | 692481.56 | 2 | 1729049.5 | 2 | 1033933.62 | 1644264.5 | 775466.94 | 1611618.12 | 1629747.75 | 1179860.75 | 2997367 | 892947.94 | 1080482.75 | 1563489 | 1807969.12 | 1315545.88 | 1481727.12 | 2.857591902 | 5 | 4.582730616 | 0 | 3.996995516 | 0 |
| P31786 | Dbi | Acyl-CoA-binding protein | 193.65 | 47.13 | 10000 | 3 | 2 | 826058.38 | 915903.38 | 2 | 1696471.38 | 3686650 | 769765.12 | 3290002 | 1510455.25 | 1131934.5 | 3593361.75 | 4217579.5 | 3167867.75 | 2436100.25 | 3387703.5 | 4398622.5 | 4887994 | 1963294.38 | 5033773 | 3547265.5 | 3.021374667 | 5 | 4.886700575 | 0 | 5.767431124 | 0 |
| P47758 | Srprb | Signal recognition particle receptor subunit beta | 303.07 | 27.14 | 29579 | 6 | 2 | 2 | 2 | 658349.94 | 676036.94 | 533938.25 | 2 | 931058.38 | 1626036.5 | 1907312.5 | 3268400.75 | 773379 | 2665043.25 | 3583411 | 1684533.62 | 1213347.25 | 2848890.5 | 1224071.25 | 3198947 | 1048925.88 | 3.745784098 | 5.5 | 8.97394448 | 0 | 7.144958597 | 0 |
| P13379 | Cd5 | T-cell surface glycoprotein CD5 | 278.94 | 12.15 | 53849 | 4 | 1926688.25 | 1240782.25 | 820710.69 | 1731334.75 | 3880314 | 2 | 2 | 2 | 1801055.38 | 1791979.5 | 2 | 2 | 2 | 2 | 2 | 2 | 473087.78 | 2 | 2 | 2 | 0.374281722 | 6 | 1.04169E-06 | 0 | 0.049281683 | 0 |
| Q62280 | Ss18 | Protein SSXT | 45.21 | 2.87 | 45864 | 1 | 898139.31 | 1346007.75 | 1085202.12 | 1014467.69 | 1971914.75 | 1215476.38 | 676902.94 | 207711.33 | 680769.69 | 1136914.38 | 814532.31 | 2 | 679713.12 | 650341.12 | 679850.56 | 2 | 1259934.12 | 941275.75 | 761721.88 | 1464306.38 | 0.620320013 | 6 | 0.447206956 | 0 | 0.700986108 | 8 |
| Q3UEB3 | Puf60 | Poly(U)-binding-splicing factor PUF60 | 408.66 | 19.86 | 60249 | 7 | 4766328.5 | 3952435.25 | 5714857 | 6682920.5 | 8402830 | 985633.88 | 3320211.25 | 5241498 | 4617472.5 | 5980181.5 | 2536850.5 | 2881044.5 | 3492566 | 3378124.25 | 3738603.5 | 1795852.25 | 661964.38 | 1330883.25 | 1721580.25 | 2 | 0.682433137 | 6 | 0.542938012 | 0 | 0.18666665 | 0 |
| P60229 | Eif3e | Eukaryotic translation initiation factor 3 subunit E | 369.64 | 11.69 | 52221 | 5 | 16118402 | 19712942 | 17804114 | 22389140 | 34142374 | 12629628 | 12169330 | 20120472 | 11413941 | 20813110 | 10963845 | 9694226 | 11030013 | 11987529 | 9341992 | 8119557 | 8050870 | 10818258 | 7636755.5 | 7956243 | 0.700268689 | 6 | 0.481247728 | 0 | 0.386519505 | 0 |
| Q5RKH0 | Glyr1 | Cytokine-like nuclear factor N-PAC | 237.65 | 10.33 | 60421 | 4 | 1365173.5 | 1882605.25 | 1132758.38 | 1722122.25 | 2907904.75 | 1296961.25 | 1165448 | 1338204.25 | 1249441.88 | 1632180 | 725383.38 | 2 | 962743.19 | 289594.06 | 2 | 973932.44 | 625086.25 | 1969732.38 | 253835.92 | 962293.38 | 0.741600113 | 6 | 0.219489546 | 0 | 0.531030055 | 4 |
| Q6P069 | Sri | Sorcin | 151.35 | 11.62 | 21627 | 3 | 13953588 | 23128646 | 16465425 | 16276046 | 14087524 | 16493951 | 12633662 | 15905589 | 10973786 | 9031470 | 10689832 | 10942434 | 9717686 | 10037392 | 7573880 | 10938448 | 8668127 | 10292616 | 15047740 | 10976065 | 0.775086467 | 6 | 0.583488343 | 0 | 0.66645426 | 2 |
| Q810B6 | Ankfy1 | Rabankyrin-5 | 57.88 | 0.77 | 128653 | 1 | 2355393.5 | 1792635.75 | 2053253 | 2208419.25 | 1463715.62 | 2060796.25 | 1647325.12 | 1949957.88 | 1105956.62 | 993583.19 | 1134123.75 | 1075707.25 | 1349676.75 | 1102966.38 | 1412295.62 | 908798.31 | 1413496.5 | 2187236.5 | 1058798.88 | 1838806.25 | 0.785707619 | 6 | 0.615265179 | 0 | 0.750210018 | 5 |
| P45376 | Akr1b1 | Aldo-keto reductase family 1 member B1 | 583.58 | 42.72 | 35732 | 13 | 36027502 | 22539574 | 35663510 | 17784270 | 41214042 | 21286462 | 17430054 | 21788014 | 31817478 | 34607562 | 66405622 | 67040726 | 50373734 | 44175058 | 46622906 | 53113482 | 36860802 | 37624078 | 50000858 | 35500738 | 0.828365743 | 6 | 1.792207929 | 0 | 1.390729561 | 5 |
| Q62426 | Cstb | Cystatin-B | 155.67 | 44.9 | 11046 | 3 | 7244759 | 6622262 | 7574419 | 5557882.5 | 7828922 | 7157263 | 8995716 | 13235120 | 8885087 | 7127745 | 16487954 | 18527734 | 11732268 | 8017994 | 14924616 | 16760776 | 14373458 | 10242861 | 7963583 | 7664079 | 1.303566449 | 6 | 2.00097843 | 0 | 1.636739314 | 1 |
| P42125 | Eci1 | Enoyl-CoA delta isomerase 1, mitochondrial | 243.58 | 17.99 | 32250 | 4 | 91506.73 | 342071 | 488017.53 | 386377.5 | 493984.72 | 331076.34 | 530816.38 | 471024.25 | 722603.31 | 857499.81 | 1563667.75 | 1875117.38 | 1010825.12 | 1778161.38 | 2323970.25 | 52746.92 | 954491.12 | 790500.75 | 2 | 865943.75 | 1.616586475 | 6 | 4.745806699 | 0 | 1.4782172 | 10 |
| Q9JMH6 | Txnrd1 | Thioredoxin reductase 1, cytoplasmic | 1055.46 | 31 | 67084 | 17 | 5671443.5 | 3921003 | 9653176 | 8867238 | 11829066 | 7351943 | 8161773.5 | 15499199 | 15774877 | 17937946 | 41038174 | 44365510 | 39475610 | 31023018 | 71183794 | 60260282 | 80854162 | 75358306 | 73789490 | 78949154 | 1.62049616 | 6 | 5.685406937 | 0 | 9.243705208 | 0 |
| P10639 | Txn | Thioredoxin | 278.19 | 37.14 | 11675 | 4 | 1630619.12 | 931475.38 | 4297961 | 1031324.75 | 70866.51 | 1973540.75 | 1325057.25 | 1891078 | 8265495 | 1453745 | 18068446 | 31003022 | 23280802 | 28040470 | 27170098 | 27343760 | 21295408 | 3147786 | 4503734.5 | 13414000 | 1.872450886 | 6 | 16.02096014 | 0 | 8.75439943 | 1 |
| A2AQP0 | Myh7b | Myosin-7B | 83.49 | 0.88 | 221497 | 2 | 9182563 | 13507803 | 75768.74 | 15127328 | 2622858.75 | 18483606 | 10243457 | 1694266.62 | 30398210 | 20142752 | 184951922 | 202230274 | 127388786 | 187972418 | 91578922 | 340920322 | 237405666 | 126010874 | 83090978 | 143220674 | 1.998263629 | 6 | 19.60005975 | 0 | 22.96971886 | 0 |
| P61314 | Rpl15 | Large ribosomal subunit protein eL15 | 384.7 | 37.25 | 24146 | 8 | 2062830 | 2737747.75 | 2928194 | 1996129.75 | 5394601.5 | 3095556.25 | 2143672.5 | 6030580.5 | 18013806 | 2837059.75 | 1486693.12 | 1860928.25 | 1739066.88 | 62862.15 | 1124286.12 | 61523.97 | 2 | 2 | 2 | 1701757 | 2.124453099 | 6 | 0.414949917 | 0 | 0.116623342 | 0 |
| Q9ER72 | Cars1 | Cysteine--tRNA ligase, cytoplasmic | 256.06 | 5.9 | 94860 | 4 | 2 | 1678257.12 | 2 | 2 | 3904268.5 | 2056942.75 | 1609134.25 | 2894205.25 | 3116892.5 | 3268230 | 7720486 | 6098886 | 6003927 | 5086881 | 4414969 | 8308785.5 | 6971766 | 5075053 | 9450274 | 4149784.25 | 2.318912929 | 6 | 5.253019776 | 0 | 6.082484625 | 0 |
| Q9Z0H3 | Smarcb1 | SWI/SNF-related matrix-associated actin-dependent regulator o | 132.03 | 9.61 | 44141 | 3 | 495760.38 | 2 | 73555.66 | 2 | 683836.88 | 1779257.75 | 2 | 785006.25 | 718774.44 | 118784.65 | 2426660.5 | 905298.88 | 1634033.5 | 1458385 | 809023.25 | 2398535.75 | 253067.64 | 748226.94 | 2 | 771251.62 | 2.714604241 | 6 | 5.772143149 | 0 | 3.328461012 | 6 |
| Q3TCN2 | Plbd2 | Putative phospholipase B-like 2 | 268.35 | 11.62 | 66289 | 6 | 4626621 | 5130918 | 3536852.5 | 3207683.5 | 2036128.38 | 1094195 | 2908589 | 4084536.5 | 1889537 | 3595258.5 | 121655.59 | 1982405.62 | 853360 | 1955544.25 | 1479673 | 988797.69 | 1829703.88 | 2424880.5 | 1195682.38 | 1056225.12 | 0.732116037 | 7 | 0.344835922 | 0 | 0.404315856 | 1 |
| Q5PQM0 | Tmem168 | Transmembrane protein 168 | 83.12 | 4.02 | 79772 | 2 | 849051.38 | 898014.19 | 699093.56 | 912721.88 | 845030.5 | 717084.75 | 622145.94 | 71431.26 | 981628.12 | 813195.25 | 2 | 2 | 62702.82 | 592007.62 | 218857.95 | 2 | 159603.06 | 2 | 2 | 108691.73 | 0.762500665 | 7 | 0.2077999 | 0 | 0.063821703 | 0 |
| Q7TPR4 | Actn1 | Alpha-actinin-1 | 611.02 | 17.15 | 103068 | 11 | 3943575 | 3686295.75 | 3464208.5 | 2745882.5 | 2673903 | 4004183 | 3343091 | 1429344.5 | 2060494 | 2297425 | 4206290.5 | 4138999 | 4806970 | 5031682 | 5318688 | 3874512.5 | 4417756.5 | 6480856 | 5887459 | 4943698.5 | 0.795364241 | 7 | 1.423205885 | 0 | 1.550471854 | 1 |
| Q3KRE8 | Tubb2b | Tubulin beta-2B chain | 1728.97 | 59.1 | 49953 | 25 | 1154048.12 | 1821816.75 | 1777855.62 | 1222610.75 | 960970.75 | 1392329.88 | 1071612.88 | 1544969.5 | 877473.06 | 953444.25 | 694621.19 | 2 | 955417.19 | 842649 | 717867.81 | 846916.25 | 930017.25 | 913751.81 | 862855.31 | 749012.38 | 0.841801262 | 7 | 0.462796227 | 0 | 0.620205522 | 0 |
| P24668 | M6pr | Cation-dependent mannose-6-phosphate receptor | 506.57 | 37.05 | 31172 | 8 | 24228282 | 26947624 | 27690430 | 26603188 | 23629946 | 21542216 | 19402504 | 34811482 | 15435472 | 25723824 | 12102146 | 12758232 | 15710543 | 12905508 | 19714754 | 13549334 | 16554526 | 20038508 | 16413667 | 17525036 | 0.905623377 | 7 | 0.566936355 | 0 | 0.651289049 | 0 |
| P35564 | Canx | Calnexin | 965.5 | 30.12 | 67278 | 18 | 11680755 | 10880212 | 12214000 | 15951912 | 12449346 | 10106980 | 11162696 | 9945812 | 14933893 | 11800377 | 9814306 | 9192345 | 10559762 | 10239144 | 9637266 | 10700595 | 9853512 | 9222194 | 6904150 | 10544959 | 0.917271616 | 7 | 0.782617559 | 0 | 0.747518707 | 0 |
| P46471 | Psmc2 | 26S proteasome regulatory subunit 7 | 836.74 | 43.65 | 48648 | 16 | 10676790 | 11251473 | 11083330 | 11050570 | 12533602 | 12944468 | 13653607 | 12868969 | 11083633 | 10335644 | 14404920 | 12801256 | 13667678 | 14688781 | 13546005 | 12927290 | 11058307 | 15508287 | 9600467 | 11117361 | 1.075810549 | 7 | 1.221092073 | 0 | 1.06389077 | 10 |
| P46460 | Nsf | Vesicle-fusing ATPase | 565.22 | 18.41 | 82613 | 12 | 7657873.5 | 8285418.5 | 8381081 | 9428272 | 12625320 | 8805435 | 8435404 | 11836935 | 12303096 | 10740062 | 5671492 | 3153711 | 4574006 | 6105888 | 4652963.5 | 2919009.25 | 2566295.5 | 5175667.5 | 1291616.12 | 3383299.5 | 1.123829646 | 7 | 0.520895225 | 0 | 0.330671858 | 0 |
| Q99JX3 | Gorasp2 | Golgi reassembly-stacking protein 2 | 121.45 | 5.32 | 47038 | 2 | 687307.81 | 783291.5 | 766412.56 | 1012279.19 | 1111869.25 | 1096108 | 919737.44 | 1170524.25 | 924634.12 | 827632.75 | 2184248.75 | 1965609.88 | 2278919.5 | 1597432.62 | 1374193.25 | 2693940.25 | 2669931.5 | 3110761.75 | 3844427.75 | 2783130.5 | 1.132413443 | 7 | 2.155482333 | 0 | 3.462883883 | 0 |
| Q9CQV8 | Ywhab | 14-3-3 protein beta/alpha | 708.81 | 41.46 | 28086 | 12 | 4896747.5 | 3315442.75 | 5708252 | 6802657 | 8317940 | 7370293 | 7398489 | 2326258.75 | 8924247 | 8336892.5 | 12719067 | 18032744 | 16987800 | 16453269 | 13922475 | 8768146 | 9115824 | 12013244 | 10611480 | 8129581.5 | 1.183021722 | 7 | 2.689826432 | 0 | 1.67481181 | 1 |
| O70503 | Hsd17b12 | Very-long-chain 3-oxoacyl-CoA reductase | 862.82 | 56.73 | 34742 | 16 | 11685065 | 13341166 | 7106831 | 16014349 | 11538698 | 14953956 | 15093060 | 7045533.5 | 17224804 | 17645932 | 18152814 | 18317658 | 28618138 | 24939064 | 20177930 | 18462868 | 27264064 | 36997754 | 28010094 | 32055632 | 1.205695709 | 7 | 1.846419642 | 0 | 2.392355849 | 0 |
| P48036 | Anxa5 | Annexin A5 | 1841.31 | 77.43 | 35752 | 27 | 74433442 | 65635002 | 83920434 | 60786442 | 38640962 | 100408762 | 97151242 | 65095842 | 60455482 | 108914722 | 97273666 | 98034242 | 106528226 | 93018850 | 89997202 | 55797974 | 103943450 | 94801330 | 85441970 | 92453482 | 1.335820347 | 7 | 1.499158246 | 0 | 1.337094729 | 4 |
| Q922Y1 | Ubxn1 | UBX domain-containing protein 1 | 149.75 | 14.81 | 33573 | 3 | 517197.91 | 704623.06 | 517041.06 | 862257.19 | 1048461.88 | 2 | 890242.5 | 1189327.38 | 2245112.5 | 898708 | 2479077 | 2337892 | 1729683.25 | 3299714.25 | 2027762.25 | 4058855 | 3126169.75 | 2835307.5 | 3289940.75 | 2322664 | 1.431230664 | 7 | 3.253559361 | 0 | 4.283488042 | 0 |
| Q9JM90 | Stap1 | Signal-transducing adaptor protein 1 | 327.59 | 24.92 | 34628 | 7 | 148260.69 | 2 | 26142.13 | 1145955.88 | 2656072.25 | 1189623.25 | 988053.5 | 69014.73 | 3713769.5 | 2635952 | 6626667.5 | 5717615.5 | 5421022 | 4879389 | 5134649 | 4244889 | 2497411.25 | 2547818 | 1718836.62 | 2116238.75 | 2.161840295 | 7 | 6.985995577 | 0 | 3.300745614 | 4 |
| Q63524 | Tmed2 | Transmembrane emp24 domain-containing protein 2 | 77.89 | 6.47 | 22733 | 1 | 2 | 2 | 2 | 2 | 2 | 2 | 168006.38 | 151387.86 | 2 | 2 | 250334.98 | 234781.3 | 828570.12 | 298280.31 | 148730.34 | 301292.34 | 260240.67 | 169582.27 | 112005.8 | 155691.83 | 31940.024 | 7.5 | 176069.705 | 0 | 99881.291 | 0 |
| O35326 | Srsf5 | Serine/arginine-rich splicing factor 5 | 125.98 | 12.64 | 30891 | 3 | 619255.31 | 548254.5 | 2060893.25 | 1801109 | 3298393.5 | 621729.88 | 516412.44 | 335381.44 | 1693820 | 2153862.5 | 172413.83 | 223181.83 | 2 | 240407.39 | 267358.12 | 2 | 409011.62 | 2 | 1195340.12 | 968392.19 | 0.638960927 | 8 | 0.108474233 | 0 | 0.308930968 | 4 |
| Q5I0E7 | Tmed9 | Transmembrane emp24 domain-containing protein 9 | 176.46 | 20 | 27028 | 4 | 3573692.75 | 2954117 | 2489207.25 | 4104578.5 | 2578163.75 | 1081017.88 | 2727770.25 | 2694144.5 | 2623489.75 | 2644813.5 | 1290146.75 | 1871314.88 | 1939953.5 | 1746271.12 | 2137977.75 | 1431636.88 | 1862592.5 | 2240060 | 541646.69 | 1244666.38 | 0.749771744 | 8 | 0.572344063 | 0 | 0.466287561 | 0 |
| Q9R1T2 | Sae1 | SUMO-activating enzyme subunit 1 | 374.18 | 14.29 | 38620 | 7 | 245071.55 | 1057856.5 | 50217.75 | 1258653.88 | 1579477.88 | 526047.94 | 23990.35 | 875560.88 | 1023161.69 | 849572.94 | 3350936.5 | 4146471 | 3359029 | 4374698 | 1584940.88 | 2428026.75 | 2684836.25 | 2340135.5 | 2 | 2253564.75 | 0.786951891 | 8 | 4.01215981 | 0 | 2.315896552 | 5 |
| P48771 | Cox7a2 | Cytochrome c oxidase subunit 7A2, mitochondrial | 63.11 | 12.05 | 9291 | 1 | 4008257.25 | 4249798.5 | 2589541 | 4579443 | 5480141 | 3044305.75 | 2658512.25 | 3710399.75 | 2716044.75 | 4599937 | 1573091.25 | 1710618.75 | 1420000.12 | 1415482.12 | 1837799.38 | 2014566.88 | 1824436.12 | 1582048 | 1746673.75 | 1966623.75 | 0.800165249 | 8 | 0.380586542 | 0 | 0.436900059 | 0 |
| Q9JHI5 | Ivd | Isovaleryl-CoA dehydrogenase, mitochondrial | 169.63 | 10.38 | 46325 | 4 | 3247137.5 | 4195867 | 3298377.75 | 3265066.5 | 3435865.5 | 3498016.25 | 2936365 | 3610277 | 2584817 | 1456980.5 | 1817727.5 | 1643076.88 | 2000313 | 1599960 | 1280573 | 2306199.5 | 2260022.75 | 2661476.25 | 2226282.25 | 2168133 | 0.807602452 | 8 | 0.478242179 | 0 | 0.666317186 | 0 |
| Q9Z2Z6 | Slc25a20 | Mitochondrial carnitine/acylcarnitine carrier protein | 134.78 | 13.62 | 33027 | 3 | 3043727.25 | 6241431 | 3453561 | 3420881.5 | 4716905 | 3006792 | 1605235.12 | 5599057.5 | 4030237.5 | 3066437.25 | 3029153.5 | 1448916.75 | 2293662.75 | 2819929.25 | 1213180.25 | 2822424.25 | 3049417 | 3645147.5 | 2741627.75 | 3304117.25 | 0.82905442 | 8 | 0.517559913 | 0 | 0.745466408 | 5 |
| Q60854 | Serpinb6 | Serpin B6 | 410.59 | 16.93 | 42599 | 6 | 48418850 | 56533778 | 47686474 | 40573434 | 38802646 | 42597970 | 40639030 | 55586614 | 33388056 | 35879454 | 25359872 | 24548882 | 24316266 | 23957408 | 21809752 | 13469201 | 17514252 | 26898156 | 22572436 | 25916026 | 0.896885808 | 8 | 0.517173829 | 0 | 0.458461684 | 0 |
| Q9D7X3 | Dusp3 | Dual specificity protein phosphatase 3 | 338.48 | 28.11 | 20472 | 4 | 6790830 | 11850266 | 6409896.5 | 8638889 | 7945000 | 8845125 | 7341637 | 10241188 | 9318432 | 8444439 | 5625802 | 6012172 | 4370785 | 5446378 | 4121944 | 4924042 | 3329517.75 | 4188902 | 4017292.5 | 3708265.75 | 1.061389379 | 8 | 0.614318573 | 0 | 0.484402003 | 0 |
| P46462 | Vcp | Transitional endoplasmic reticulum ATPase | 1885.57 | 41.32 | 89349 | 33 | 12293020 | 15616319 | 19083164 | 12386286 | 15454222 | 12946910 | 12670348 | 17911234 | 15616610 | 20345178 | 35687142 | 36593034 | 27202898 | 29641356 | 30362014 | 24926682 | 17162066 | 18566426 | 20345566 | 12155621 | 1.062235489 | 8 | 2.131231149 | 0 | 1.244856511 | 7 |
| P26151 | Fcgr1 | High affinity immunoglobulin gamma Fc receptor I | 130.07 | 8.66 | 44888 | 3 | 1108650.38 | 1011444.88 | 1563863.75 | 1690631.62 | 1458674.88 | 1637133.38 | 1676078.12 | 2194248.25 | 1163304.12 | 1290947.12 | 933770.94 | 828623.44 | 2 | 371523.06 | 683253.31 | 2 | 2 | 2 | 2 | 2 | 1.165140002 | 8 | 0.412273275 | 0 | 1.46343E-06 | 0 |
| O35682 | Myadm | Myeloid-associated differentiation marker | 302.69 | 21.25 | 35285 | 4 | 653041.06 | 1371613.12 | 722949.25 | 2607943.5 | 551924.69 | 2252708 | 937192.75 | 1201900.38 | 2129349 | 933837.88 | 2 | 2 | 2 | 2 | 2 | 1874527.75 | 2 | 2 | 2699133.5 | 2 | 1.261959175 | 8 | 1.69277E-06 | 0 | 0.774217388 | 9 |
| Q62159 | Rhoc | Rho-related GTP-binding protein RhoC | 251.62 | 24.87 | 22006 | 5 | 1454328.62 | 2939105.75 | 1155096.75 | 2486241.75 | 2500522.75 | 2479925.75 | 2264504.5 | 3289144.25 | 2833124.25 | 2550940.25 | 5872372.5 | 5251390 | 5997114 | 5358010.5 | 4464569.5 | 6599858.5 | 7317321.5 | 6804928 | 8640946 | 6316176 | 1.273589227 | 8 | 2.557446651 | 0 | 3.386637764 | 0 |
| F1LQ48 | Hnrnpl | Heterogeneous nuclear ribonucleoprotein L | 702.06 | 26.65 | 67903 | 12 | 8078394 | 11682807 | 4391476 | 12318553 | 5053131 | 9879114 | 8760970 | 5351971 | 18972768 | 11996494 | 15704475 | 25198792 | 17523854 | 19698384 | 12949119 | 17124440 | 15093348 | 12263827 | 17884082 | 13788538 | 1.32359212 | 8 | 2.19328177 | 0 | 1.833965248 | 1 |
| Q11136 | Pepd | Xaa-Pro dipeptidase | 390.26 | 16.84 | 55029 | 8 | 2358463 | 1234343.75 | 2443930 | 2205081.5 | 2710858 | 3079266 | 1888948 | 4525867.5 | 3159498.25 | 1941895.88 | 9243808 | 11072866 | 9342275 | 9774340 | 5483399 | 12431800 | 7240017 | 7095118.5 | 5210721 | 9175137 | 1.332594454 | 8 | 4.100978334 | 0 | 3.757327667 | 0 |
| P47911 | Rpl6 | Large ribosomal subunit protein eL6 | 545.4 | 29.39 | 33510 | 10 | 1690973.5 | 4263378.5 | 2959971.75 | 5370237.5 | 8640134 | 7396459 | 3762642 | 6789501.5 | 8096470.5 | 4802272.5 | 9030209 | 11028524 | 9189400 | 10575780 | 9013988 | 16998280 | 9950052 | 7349315.5 | 4623155.5 | 9957904 | 1.345594572 | 8 | 2.13036206 | 0 | 2.132142062 | 3 |
| Q5M9G3 | Caprin1 | Caprin-1 | 128.89 | 4.81 | 78121 | 3 | 943446.62 | 2 | 582356.25 | 2 | 449776.47 | 669854.88 | 498388.97 | 179587.69 | 877046.38 | 701061.62 | 1930102.62 | 1004292.75 | 2471390.5 | 2106798.5 | 1648500.12 | 977938 | 2 | 2 | 2 | 1124780.75 | 1.481050928 | 8 | 4.637154153 | 0 | 1.064356389 | 12 |
| Q8K0D5 | Gfm1 | Elongation factor G, mitochondrial | 345.25 | 10.79 | 83550 | 8 | 1362288.5 | 523055.44 | 1928451.25 | 726433.38 | 1865306 | 1541938.25 | 1179749.75 | 2723652.25 | 1240703.75 | 5314760 | 4159827.5 | 3222339.75 | 5377717 | 2208032.5 | 2313085.5 | 3634126.5 | 2185122.75 | 2274656.75 | 3299990.25 | 1581623 | 1.873505461 | 8 | 2.697823587 | 0 | 2.025673128 | 2 |
| Q61120 | Shc3 | SHC-transforming protein 3 | 41.22 | 1.48 | 52121 | 1 | 2 | 2 | 619523.44 | 2 | 847344.69 | 2 | 2 | 2 | 2 | 605646.38 | 1284178.62 | 1397242.88 | 1083866 | 1622399.25 | 1134821 | 1491329 | 1435371.62 | 2 | 2206034.25 | 1553752.88 | 0.412887764 | 9 | 4.446535402 | 0 | 4.558325499 | 3.5 |
| Q80W92 | Vac14 | Protein VAC14 homolog | 101.98 | 3.19 | 88068 | 2 | 1028617.12 | 1589233.5 | 715809.69 | 1565242.75 | 1616001.62 | 809300.75 | 1990515.5 | 2 | 1180903.5 | 951926.88 | 2 | 2 | 2 | 2 | 2 | 2 | 2 | 2 | 2 | 2 | 0.757132893 | 9 | 1.53494E-06 | 0 | 1.53494E-06 | 0 |
| P31996 | Cd68 | Macrosialin | 117.89 | 8.9 | 34818 | 3 | 165959250 | 207126978 | 138495938 | 95581930 | 82793746 | 124288546 | 107422362 | 147383826 | 98917250 | 82176634 | 80425250 | 76476426 | 74132970 | 77241010 | 63619786 | 120026946 | 131139322 | 108484226 | 3420141.25 | 105867514 | 0.811917169 | 9 | 0.539011834 | 0 | 0.679662033 | 8 |
| Q9CXT6 | Kdm8 | Bifunctional peptidase and arginyl-hydroxylase JMJD5 | 34.27 | 2.17 | 47145 | 1 | 587004.81 | 741062.12 | 233807.59 | 1112596 | 239098.09 | 2 | 2 | 94136.98 | 872458.88 | 1492081.62 | 5061917.5 | 5748963 | 5361124.5 | 5354169 | 3723322.5 | 6701942.5 | 7829952.5 | 8186082 | 7989992 | 8723644 | 0.843872861 | 9 | 8.666175361 | 0 | 13.53378563 | 0 |
| Q9Z0J0 | Npc2 | NPC intracellular cholesterol transporter 2 | 224.79 | 36.91 | 16442 | 5 | 12622767 | 6676157 | 6984917 | 5793348.5 | 3422078 | 5129394 | 4621308 | 5907170 | 4392889 | 10499966 | 887195.88 | 602002.12 | 2319336.5 | 2532777.5 | 996490.06 | 1730604.62 | 2 | 3035582.75 | 1350598.25 | 2546385 | 0.860601617 | 9 | 0.206702915 | 0 | 0.244038067 | 0 |
| P09527 | Rab7a | Ras-related protein Rab-7a | 969.54 | 77.78 | 23504 | 16 | 79725794 | 69026698 | 86186418 | 51292674 | 46882006 | 54870722 | 58315638 | 64479886 | 51150698 | 59932214 | 40090334 | 31785804 | 35977722 | 34465758 | 45270050 | 28003788 | 33135716 | 34217250 | 16345882 | 31797758 | 0.866818907 | 9 | 0.563140243 | 0 | 0.430785169 | 0 |
| O09043 | Napsa | Napsin-A | 169.67 | 13.84 | 45544 | 3 | 16189915 | 18799612 | 13301041 | 13880866 | 11503477 | 13429767 | 10915863 | 15108324 | 14470656 | 12683789 | 10354294 | 9532162 | 11092173 | 9808202 | 4511010.5 | 12049505 | 6660784 | 8497764 | 8948697 | 5883593 | 0.904085232 | 9 | 0.614834017 | 0 | 0.570619529 | 1 |
| Q9D1Q4 | Dpm3 | Dolichol-phosphate mannosyltransferase subunit 3 | 44.45 | 10.87 | 10139 | 1 | 3945883.25 | 2948939.5 | 3786439.5 | 3286844.25 | 2369528.75 | 3319600 | 2792770.75 | 4626073 | 1710608 | 2349878.5 | 1862478.38 | 1529527.75 | 2017991.75 | 1664096.88 | 2142684.25 | 1919664.88 | 3174364 | 4153257 | 2097101 | 3610791.75 | 0.905818377 | 9 | 0.564144007 | 0 | 0.915382086 | 10 |
| P04762 | Cat | Catalase | 335.45 | 13.09 | 59757 | 6 | 3188444.75 | 2783376.5 | 4354390.5 | 183842.34 | 3192236 | 3034477 | 1768582.88 | 4294577 | 1083005.5 | 2674074.75 | 14609217 | 18908598 | 7158840 | 14648034 | 5916564 | 15377904 | 9396479 | 4177789.5 | 13383325 | 5859010 | 0.9381437 | 9 | 4.469417345 | 0 | 3.517259318 | 1 |
| P62627 | Dynlrb1 | Dynein light chain roadblock-type 1 | 86.54 | 21.88 | 10990 | 2 | 638098.94 | 1458284 | 822269.06 | 693453.31 | 1465202.75 | 1371334.38 | 874332.94 | 1647039.25 | 783915.25 | 822466 | 3148254.5 | 2631294.5 | 2535769 | 2852504.25 | 1696800.88 | 2851766 | 2721931.75 | 2206940 | 2968842 | 3380210.75 | 1.083071532 | 9 | 2.533748785 | 0 | 2.782909828 | 0 |
| Q9D8B3 | Chmp4b | Charged multivesicular body protein 4b | 164.67 | 14.73 | 24936 | 3 | 1597388.12 | 2477920.25 | 2048380.5 | 1730587.75 | 4172110.75 | 3856480 | 4818101 | 137097.73 | 2176711 | 2833244.25 | 4628973 | 5113061 | 4478849 | 4513796 | 4442939 | 6408556 | 2395529 | 3169040 | 6794688 | 6147525 | 1.149275635 | 9 | 1.927230288 | 0 | 2.071722558 | 3 |
| Q6MG61 | Clic1 | Chloride intracellular channel protein 1 | 974.04 | 65.56 | 26981 | 16 | 26968158 | 18304856 | 37844354 | 28932872 | 35251702 | 23901558 | 25802932 | 39489206 | 35451110 | 46647030 | 39502898 | 39852202 | 46347422 | 45278302 | 47106122 | 28934484 | 34945310 | 28225012 | 32661868 | 28450174 | 1.162862035 | 9 | 1.48054359 | 0 | 1.040154976 | 12 |
| P63245 | Rack1 | Small ribosomal subunit protein RACK1 | 845.81 | 49.84 | 35077 | 16 | 10371467 | 12854588 | 18832582 | 19504192 | 29666956 | 17250666 | 16205370 | 28957430 | 22314444 | 29341882 | 41535650 | 36989786 | 40432790 | 36327738 | 40458218 | 27196768 | 21733310 | 21972930 | 24499766 | 17192106 | 1.250356909 | 9 | 2.145617048 | 0 | 1.234189908 | 7 |
| P35831 | Ptpn12 | Tyrosine-protein phosphatase non-receptor type 12 | 50.28 | 1.68 | 86526 | 1 | 2 | 2 | 2 | 627883.81 | 2 | 2 | 2 | 859190.94 | 654312.88 | 2 | 1519860.38 | 982521.12 | 867352.06 | 694732.69 | 775673 | 974136.56 | 2 | 2 | 1126707.62 | 861038.56 | 2.410462752 | 9 | 7.708556113 | 0 | 4.717192823 | 6 |
| Q64339 | Isg15 | Ubiquitin-like protein ISG15 | 40.09 | 11.18 | 17898 | 1 | 2 | 2 | 46701.96 | 2 | 2 | 2 | 2 | 2 | 142907.44 | 159444.38 | 810289.06 | 744299.44 | 710402.06 | 1278398.5 | 1048624.62 | 381620.69 | 1937971.62 | 793687.69 | 1464504.88 | 1004523.5 | 6.473090964 | 9 | 98.3090904 | 0 | 119.5100227 | 0 |
| Q9R0C8 | Vav3 | Guanine nucleotide exchange factor VAV3 | 69.02 | 2.13 | 97968 | 2 | 2 | 391021.97 | 2 | 2 | 2 | 2 | 2 | 2 | 2 | 2 | 1263986.38 | 1219317.62 | 1056342.88 | 1083186.5 | 1004115.25 | 914945.25 | 902887.25 | 1268717.38 | 1057711.38 | 915242.56 | 2.55735E-05 | 10 | 14.39006997 | 0 | 12.93891571 | 0 |
| Q62087 | Pon3 | Serum paraoxonase/lactonase 3 | 106.99 | 7.63 | 39351 | 2 | 396417.75 | 293665.19 | 2 | 2 | 2 | 2 | 308404.81 | 2 | 2 | 2 | 3272392.75 | 2565963.25 | 3093448.25 | 2663525 | 1699117 | 3400310.5 | 3397370 | 3893108 | 1907119.12 | 3518726.25 | 0.44691748 | 10 | 19.2648302 | 0 | 23.35443004 | 0 |
| P24452 | Capg | Macrophage-capping protein | 832.16 | 42.33 | 39240 | 14 | 2 | 2 | 1897432.12 | 2 | 797125.56 | 2 | 2 | 2 | 2 | 1754610.12 | 2987200.75 | 2615674.5 | 7843720.5 | 3403464.5 | 1971825.62 | 400574.81 | 1274019.75 | 966181.56 | 2132503.75 | 2 | 0.651169662 | 10 | 6.98513307 | 0 | 1.771448901 | 7.5 |
| Q99KP6 | Prpf19 | Pre-mRNA-processing factor 19 | 259.72 | 21.63 | 55239 | 6 | 1802065.62 | 1367881 | 1225169.75 | 5200482 | 2636250.75 | 1541520.12 | 1627458.62 | 1859515.38 | 2420706.75 | 1142465.88 | 2 | 533287.88 | 2 | 527514.88 | 2 | 44853.05 | 2 | 2 | 2 | 2 | 0.702401302 | 10 | 0.086725134 | 0 | 0.003667561 | 0 |
| P62960 | Ybx1 | Y-box-binding protein 1 | 102.39 | 5.9 | 35730 | 1 | 766657.31 | 1014144.19 | 636797.25 | 1295894.62 | 1304048.5 | 2 | 2 | 1650829.25 | 1137573.62 | 784355.19 | 3438250 | 2417723.25 | 1900699.75 | 2483723.25 | 3043710.25 | 2601958.75 | 2873486 | 2996822.5 | 2537314.5 | 2327003.75 | 0.712054259 | 10 | 2.647532765 | 0 | 2.65799187 | 0 |
| P34022 | Ranbp1 | Ran-specific GTPase-activating protein | 183.44 | 15.27 | 23596 | 4 | 1420200.5 | 3561573.5 | 1699603.38 | 759469.19 | 1445812.75 | 2 | 1776315.38 | 1102145.88 | 2899010 | 1232289 | 14096292 | 15866154 | 11501342 | 12312088 | 8028302.5 | 13143032 | 8997118 | 8945227 | 14921499 | 9369192 | 0.788796105 | 10 | 6.954714508 | 0 | 6.231370643 | 0 |
| Q3U0V1 | Khsrp | Far upstream element-binding protein 2 | 1094.47 | 30.75 | 76775 | 20 | 5712495.5 | 20302098 | 13031439 | 11912410 | 19202644 | 12633669 | 9263024 | 12385738 | 13120324 | 12533748 | 36732502 | 37390714 | 26217434 | 32992780 | 24233652 | 28538248 | 36204358 | 20755536 | 41699170 | 21183100 | 0.854269881 | 10 | 2.245790222 | 0 | 2.114853395 | 0 |
| Q8K2B3 | Sdha | Succinate dehydrogenase [ubiquinone] flavoprotein subunit, mit | 703.67 | 17.92 | 72585 | 12 | 16874260 | 27653206 | 21553480 | 24631876 | 28255850 | 22674414 | 26339578 | 6669679 | 27039456 | 21757476 | 29489054 | 35114142 | 32660820 | 32388732 | 28423118 | 24610188 | 27459382 | 33809154 | 28162348 | 31167238 | 0.878219461 | 10 | 1.328718421 | 0 | 1.220559224 | 6 |
| P09055 | Itgb1 | Integrin beta-1 | 859.43 | 23.68 | 88231 | 17 | 26827254 | 24903076 | 25213880 | 34015998 | 42076114 | 19639934 | 18380142 | 35698314 | 33658262 | 28997768 | 17964546 | 21100814 | 17282256 | 18028760 | 21999884 | 14219216 | 21699568 | 20745810 | 18961220 | 20015696 | 0.891124527 | 10 | 0.629760692 | 0 | 0.624959544 | 0 |
| E9Q555 | Rnf213 | E3 ubiquitin-protein ligase RNF213 | 665.23 | 2.89 | 584258 | 15 | 6659683 | 7675634.5 | 7031556.5 | 7023087 | 6244417.5 | 5391055 | 6555351 | 8047311.5 | 3924602.75 | 7286084 | 3119949 | 2199626.25 | 2601293 | 3476051.5 | 791133.88 | 3125068.5 | 6071789.5 | 6922080 | 5614230.5 | 4385696.5 | 0.900966196 | 10 | 0.351906232 | 0 | 0.754131188 | 2 |
| O70435 | Psma3 | Proteasome subunit alpha type-3 | 426.61 | 29.02 | 28405 | 7 | 33545350 | 48792914 | 30673740 | 36638322 | 31093562 | 40033010 | 31639274 | 43840402 | 29191530 | 20604120 | 21665390 | 20401898 | 25603076 | 28050738 | 14937402 | 23032304 | 30165126 | 28436302 | 16356142 | 32771928 | 0.914599867 | 10 | 0.612239259 | 0 | 0.723464585 | 2 |
| Q6PDM2 | Srsf1 | Serine/arginine-rich splicing factor 1 | 379.45 | 28.63 | 27745 | 8 | 7113532 | 4603593.5 | 7852639.5 | 6933812 | 5263414.5 | 7274896.5 | 6509656.5 | 2929965 | 6543227 | 6368002.5 | 8137249 | 12659371 | 9997577 | 9067511 | 8343594 | 9305208 | 8985922 | 5347154 | 7780919.5 | 19384570 | 0.932595317 | 10 | 1.517465134 | 0 | 1.59926298 | 4 |
| P80314 | Cct2 | T-complex protein 1 subunit beta | 1371.55 | 54.95 | 57477 | 22 | 17484746 | 26218186 | 29614788 | 22148408 | 28373718 | 23819830 | 25087054 | 16678305 | 29117518 | 23540690 | 34388082 | 42108058 | 36619170 | 42793866 | 34957058 | 26175418 | 29551102 | 30537646 | 18817074 | 30194774 | 0.95480898 | 10 | 1.541234426 | 0 | 1.092346433 | 8 |
| Q9CPR5 | Mrpl15 | Large ribosomal subunit protein uL15m | 180.13 | 23.05 | 33542 | 5 | 1027635.94 | 1525119.62 | 716517.38 | 707966.25 | 856651.56 | 587277.06 | 605917.88 | 1541466.62 | 1665088.62 | 470134.16 | 2 | 2 | 63511.32 | 507334.44 | 2 | 2 | 2 | 343751.91 | 2 | 2 | 1.007446091 | 10 | 0.118093641 | 0 | 0.071114539 | 0 |
| O35093 | Timm23 | Mitochondrial import inner membrane translocase subunit Tim2 | 104.3 | 12.92 | 21907 | 2 | 1098240.25 | 2129503 | 1220345.75 | 2297182.75 | 4049270.5 | 2254189 | 1972933.75 | 2824388 | 2455170 | 1661261.25 | 786754.19 | 811343.31 | 1025088.44 | 929792.19 | 352450 | 290347.56 | 2 | 2 | 1095014.38 | 2 | 1.034591532 | 10 | 0.361796549 | 0 | 0.128339665 | 0 |
| Q91YR1 | Twf1 | Twinfilin-1 | 98.76 | 4.86 | 40079 | 2 | 2014464.88 | 2406623.25 | 1770587.62 | 1655929 | 2186274.5 | 1887858.12 | 2093488.38 | 2439260.25 | 1887440.12 | 2120886.5 | 1183455.38 | 1325746.12 | 1119010 | 1562541.62 | 1089390.88 | 992702.81 | 1256015.62 | 1771850 | 2 | 1019428.31 | 1.039372023 | 10 | 0.625893918 | 0 | 0.502298126 | 2 |
| P23116 | Eif3a | Eukaryotic translation initiation factor 3 subunit A | 601.01 | 9.75 | 161936 | 13 | 4492523.5 | 5516413 | 4311230.5 | 3401262 | 4692750 | 2638923.75 | 4322498 | 6232332.5 | 5873450.5 | 4500970.5 | 11102821 | 9795734 | 10696893 | 11979127 | 7383088.5 | 9240019 | 8334341 | 7508762 | 8914103 | 7389175 | 1.0514851 | 10 | 2.273456614 | 0 | 1.846438364 | 0 |
| P46978 | Stt3a | Dolichyl-diphosphooligosaccharide--protein glycosyltransferase | 190.66 | 5.67 | 80598 | 4 | 1740045.25 | 1554315 | 4348640 | 3434757.25 | 1008056.38 | 1568428.12 | 4037981.75 | 2014811.75 | 2764020 | 2398007.5 | 5390157.5 | 4513052.5 | 5798722 | 6521464 | 6090219 | 3426037.5 | 5222915 | 5246175.5 | 3615137 | 4173264 | 1.057706932 | 10 | 2.342714796 | 0 | 1.794130641 | 4 |
| P30204 | Msr1 | Macrophage scavenger receptor types I and II | 152.05 | 9.17 | 50170 | 3 | 6066337 | 7348997 | 4959982.5 | 6351289 | 5461674 | 6122281.5 | 6893375.5 | 8855820 | 6257679 | 4531502 | 4474491.5 | 3600836.5 | 4323919 | 3884632 | 3759890 | 5172127 | 7432259 | 7291486 | 5659761 | 7746067 | 1.081898622 | 10 | 0.66395864 | 0 | 1.10313342 | 8 |
| Q64213 | Sf1 | Splicing factor 1 | 364.33 | 13.63 | 70403 | 7 | 2724250.75 | 2156467.25 | 1531398.12 | 4250844 | 1700350 | 2274204.25 | 2646129.5 | 405504.06 | 4725358.5 | 4029427.25 | 6785057 | 7441526.5 | 6169735 | 6316582 | 7599611.5 | 9379657 | 9390071 | 4517537 | 8701181 | 6432799 | 1.138904017 | 10 | 2.77534994 | 0 | 3.107682702 | 0 |
| P61979 | Hnrnpk | Heterogeneous nuclear ribonucleoprotein K | 996.27 | 37.8 | 50976 | 17 | 7852576 | 18073958 | 17409362 | 30763140 | 28724942 | 18411612 | 19099608 | 25529170 | 28681204 | 27520128 | 37159174 | 39831810 | 37411566 | 43743558 | 31724326 | 33389050 | 30941152 | 33351522 | 42603974 | 32481794 | 1.159668438 | 10 | 1.84655795 | 0 | 1.680225715 | 0 |
| P05555 | Itgam | Integrin alpha-M | 966.76 | 18.39 | 127481 | 16 | 13079176 | 11892058 | 15496410 | 14781898 | 26167896 | 13002111 | 13304308 | 19713490 | 25160576 | 24379834 | 9109539 | 8581599 | 8856685 | 7429529 | 4696345 | 7407742 | 2829743.25 | 2151679.25 | 5894723 | 4603535 | 1.173708254 | 10 | 0.475005084 | 0 | 0.281112045 | 0 |
| Q62418 | Dbnl | Drebrin-like protein | 220.25 | 10.32 | 48700 | 4 | 2 | 935334.19 | 1055620.88 | 592149.06 | 1687568 | 2241108.5 | 2 | 1853245.38 | 1138046.12 | 2 | 3730809.75 | 4507911 | 2232920.25 | 2870302.5 | 1895743.5 | 3501083 | 1248764.62 | 1194395.62 | 138780.22 | 1504003.75 | 1.225193925 | 10 | 3.567981667 | 0 | 1.776540888 | 7 |
| Q6PHZ2 | Camk2d | Calcium/calmodulin-dependent protein kinase type II subunit de | 228.18 | 10.42 | 56369 | 5 | 3551099.5 | 3338733 | 2477870.5 | 1647751.12 | 3625732.5 | 2438797 | 2744452.5 | 2664616 | 5531193.5 | 6317036.5 | 3663084 | 4007589 | 3778555 | 4857742.5 | 4336448 | 1555611.75 | 3377590.25 | 3646485.75 | 1455641.88 | 3270009 | 1.345252677 | 10 | 1.409955288 | 0 | 0.908760948 | 10 |
| O70310 | Nmt1 | Glycylpeptide N-tetradecanoyltransferase 1 | 375.74 | 15.93 | 56888 | 7 | 2 | 2343480.5 | 51883.49 | 1748806 | 1846380.5 | 909011.69 | 2507465.25 | 1907166.5 | 1680328.38 | 1181851.88 | 3194476.5 | 3607014.5 | 5019100 | 3980696.5 | 3052037 | 2328499.75 | 1875573.25 | 1204922.5 | 1308099.88 | 4269907 | 1.36645555 | 10 | 3.147176247 | 0 | 1.834054939 | 8 |
| P84083 | Arf5 | ADP-ribosylation factor 5 | 398.46 | 33.33 | 20530 | 6 | 591706.56 | 735106.5 | 2 | 676942.31 | 2 | 2 | 714756.19 | 2 | 732713.75 | 1488876.38 | 1200409 | 1184836.75 | 995314.81 | 910760 | 1287329.62 | 802542.56 | 1214126.75 | 895252.62 | 487480 | 1029457.81 | 1.465420631 | 10 | 2.784091874 | 0 | 2.210275249 | 3 |
| P49710 | Hcls1 | Hematopoietic lineage cell-specific protein | 331.7 | 12.14 | 54240 | 5 | 2 | 2 | 2 | 2 | 657500.44 | 2 | 2 | 2 | 1656552.25 | 431413.28 | 1713799.5 | 6783646.5 | 1696582.75 | 3450192 | 927273.25 | 1275204.25 | 897277.81 | 2 | 1687206.75 | 1107950.38 | 3.175581336 | 10 | 22.16168358 | 0 | 7.555250834 | 3 |
| Q8BGS2 | Bola2 | BolA-like protein 2 | 58.37 | 18.6 | 10215 | 2 | 2 | 2 | 2 | 2 | 2 | 408182.91 | 2 | 2 | 2 | 2 | 1621187.5 | 1576989.75 | 509885.84 | 1269350.38 | 864614.38 | 2 | 2 | 854409.06 | 454425.66 | 963789.69 | 40819.091 | 10 | 584202.785 | 0 | 227262.841 | 5 |
| Q9DCR2 | Ap3s1 | AP-3 complex subunit sigma-1 | 46.62 | 5.7 | 21732 | 1 | 2 | 2 | 2 | 2 | 2 | 2 | 2 | 2 | 760315.88 | 2 | 949730.12 | 1174466.25 | 1064903.62 | 1068920 | 779012.56 | 1358409.62 | 1371300.75 | 1277988.12 | 883522.12 | 1549241.25 | 76032.388 | 10 | 503703.255 | 0 | 644046.186 | 0 |
| P62858 | Rps28 | Small ribosomal subunit protein eS28 | 200 | 40.58 | 7841 | 4 | 1741975.38 | 674755.38 | 1317763.38 | 1416598.25 | 3088444 | 1596316.38 | 851072.94 | 147396.55 | 1427506.5 | 2349439 | 10933044 | 9701996 | 8382043 | 5661948.5 | 9986816 | 6828980 | 8204428.5 | 4876772 | 6128217 | 5608006 | 0.773311879 | 11 | 5.42091756 | 0 | 3.840799045 | 0 |
| Q9DBR1 | Xrn2 | 5'-3' exoribonuclease 2 | 501.3 | 14.93 | 108687 | 10 | 1815590.5 | 542564.56 | 792709.88 | 1009718.75 | 1195258.88 | 816235.38 | 836987.25 | 603365.38 | 876907.06 | 1280698.62 | 2880143.75 | 2545775.75 | 2813205.25 | 2690584.75 | 2400325.25 | 280504 | 896167.88 | 731851.62 | 2829287.75 | 1357322.62 | 0.824182868 | 11 | 2.488877254 | 0 | 1.138034546 | 12 |
| P60335 | Pcbp1 | Poly(rC)-binding protein 1 | 914.61 | 44.38 | 37498 | 11 | 7189898.5 | 11471905 | 8875188 | 15302410 | 19143796 | 7596127 | 9245277 | 9286498 | 13237682 | 12706422 | 56809302 | 55809998 | 47651510 | 54187330 | 43222330 | 42404454 | 43959934 | 39640782 | 48475118 | 43078566 | 0.840098738 | 11 | 4.157263265 | 0 | 3.509965003 | 0 |
| Q80TL7 | Mon2 | Protein MON2 homolog | 36.5 | 0.41 | 189078 | 1 | 390078.88 | 1069978.12 | 457646.88 | 707330.94 | 539219.62 | 963842.75 | 539615.56 | 922933.81 | 278393.97 | 2 | 330830 | 2 | 2 | 2 | 2 | 2 | 2 | 2 | 674696.94 | 2 | 0.854794752 | 11 | 0.104554803 | 0 | 0.213227145 | 3 |
| P53534 | Pygb | Glycogen phosphorylase, brain form (Fragment) | 275.93 | 8.23 | 96174 | 6 | 1405950.75 | 177788.25 | 2451044 | 1308130.12 | 738744.75 | 361628.41 | 1031798.75 | 1914042 | 803341.88 | 1144766.12 | 3126257.5 | 2966767 | 4143328.75 | 3554731 | 3304647 | 2050242.25 | 1429011.88 | 2090705.12 | 1779037.25 | 1517154.5 | 0.8641685 | 11 | 2.811031402 | 0 | 1.457850999 | 5 |
| Q8R1I1 | Uqcr10 | Cytochrome b-c1 complex subunit 9 | 71.78 | 12.5 | 7446 | 2 | 5766986 | 3071199 | 3000463 | 2156913.75 | 2221980 | 3522641 | 4137132 | 2335787 | 3089473 | 1538639.75 | 88681.15 | 1791798.75 | 1928256.5 | 1318080.25 | 480309.03 | 2093396 | 748677.44 | 1036607.5 | 1071987.88 | 2 | 0.901719445 | 11 | 0.345744489 | 0 | 0.305266414 | 0 |
| O35215 | Ddt | D-dopachrome decarboxylase | 50.82 | 11.02 | 13077 | 1 | 1328448.25 | 2 | 2 | 2 | 2 | 2 | 2 | 2 | 462074.31 | 761013.81 | 2673077.75 | 2518914.75 | 2023755.5 | 2096887.88 | 2026025.62 | 3760333.25 | 2855388.75 | 2154484.75 | 1177419.62 | 2557334.75 | 0.920688295 | 11 | 8.535216346 | 0 | 9.413152386 | 1 |
| Q9CR62 | Slc25a11 | Mitochondrial 2-oxoglutarate/malate carrier protein | 853.05 | 44.59 | 34155 | 14 | 21739106 | 25860332 | 20349416 | 17345914 | 16462376 | 20744234 | 21366946 | 23169676 | 17319896 | 13133044 | 13109412 | 8640406 | 14279246 | 15695291 | 11719915 | 10152884 | 12657407 | 15273007 | 10374512 | 13551380 | 0.940806633 | 11 | 0.623487133 | 0 | 0.609384143 | 0 |
| P62959 | Hint1 | Adenosine 5'-monophosphoramidase HINT1 | 100.33 | 18.25 | 13777 | 2 | 5476761 | 9991302 | 6193847.5 | 5337748 | 15595922 | 5710656 | 5679682.5 | 8666787 | 7546599.5 | 12919569 | 18494190 | 21858152 | 18282924 | 17580766 | 18741080 | 27873396 | 24914500 | 24222898 | 33652446 | 20117758 | 0.95134973 | 11 | 2.229271462 | 0 | 3.070295004 | 0 |
| O55131 | Septin7 | Septin-7 | 256.45 | 12.84 | 50550 | 6 | 342020 | 2 | 47517.2 | 907546.62 | 1470197.12 | 1376143.75 | 796497.38 | 2 | 467661.47 | 2 | 2853167.75 | 3972349.75 | 1666445.12 | 1936023.38 | 2215022.5 | 3294660.5 | 1375009.75 | 2271856.25 | 429828.72 | 685594.81 | 0.954115158 | 11 | 4.568744423 | 0 | 2.911502078 | 5 |
| Q61599 | Arhgdib | Rho GDP-dissociation inhibitor 2 | 315.88 | 41.5 | 22851 | 6 | 12085706 | 8637191 | 14381859 | 10979852 | 17308962 | 9747859 | 11646135 | 16862480 | 13793395 | 9501614 | 21818246 | 27089976 | 24190888 | 20165530 | 19982184 | 14207953 | 21622128 | 14310694 | 25044748 | 13554568 | 0.970942053 | 11 | 1.786408685 | 0 | 1.399827948 | 6 |
| Q8BXQ2 | Pigt | GPI transamidase component PIG-T | 95.37 | 2.58 | 65705 | 2 | 1574084.88 | 287158.5 | 1815589 | 1718117.75 | 446897.41 | 2 | 1341348.12 | 1646968.75 | 1750767.88 | 1035307.06 | 2404539.75 | 3394499.5 | 2091193.38 | 3099209 | 2369450.5 | 2423161.25 | 2590952 | 4439792.5 | 1535301.5 | 3313195 | 0.988453357 | 11 | 2.286758091 | 0 | 2.448266948 | 3 |
| Q3UFY7 | Nt5c3b | 7-methylguanosine phosphate-specific 5'-nucleotidase | 43.52 | 3.67 | 34425 | 1 | 692091.25 | 767129.38 | 767169.94 | 845662.75 | 711989.56 | 953836.19 | 815112.06 | 1141371.62 | 532642.81 | 347319.62 | 2 | 479040.75 | 562263 | 461222.59 | 2 | 1044255.06 | 680609 | 690358.81 | 648058.19 | 2 | 1.001648877 | 11 | 0.397070115 | 0 | 0.809526519 | 5 |
| Q9R112 | Sqor | Sulfide:quinone oxidoreductase, mitochondrial | 820.09 | 42 | 50282 | 15 | 19509638 | 19066550 | 26872774 | 17902252 | 17786308 | 16576687 | 17994872 | 23049462 | 20474210 | 24643526 | 13989651 | 9522058 | 17076548 | 10679436 | 16951778 | 15351738 | 17389086 | 28320584 | 16780890 | 20864156 | 1.015832255 | 11 | 0.674521875 | 0 | 0.975962749 | 9 |
| P11835 | Itgb2 | Integrin beta-2 | 1369.22 | 34.63 | 85026 | 25 | 34598550 | 41012110 | 30867802 | 29820060 | 28389790 | 28077724 | 22246126 | 40234966 | 50593506 | 30388658 | 21049082 | 19726968 | 16605289 | 20482824 | 12470584 | 19564462 | 22706786 | 22723816 | 21980834 | 13978626 | 1.041609923 | 11 | 0.548519478 | 0 | 0.613003575 | 0 |
| D3Z7P3 | Gls | Glutaminase kidney isoform, mitochondrial | 384.18 | 16.91 | 73964 | 7 | 11983941 | 14366878 | 12168808 | 12914043 | 15878416 | 14035675 | 11767126 | 18758262 | 13067364 | 13960429 | 10759921 | 8440191 | 10511498 | 10144089 | 7605875 | 11812026 | 13634850 | 18923092 | 15406768 | 15721714 | 1.063536435 | 11 | 0.705097358 | 0 | 1.12161804 | 9 |
| P62944 | Ap2b1 | AP-2 complex subunit beta | 466.89 | 10.99 | 104553 | 9 | 2411310.25 | 2033997.75 | 1746355.12 | 3509958.5 | 4102000 | 2337627.25 | 2286224.75 | 4070818.25 | 2912285 | 3302223 | 1602881.5 | 1542106.75 | 1089845.88 | 1233824.88 | 2 | 1537738.25 | 2 | 2 | 1063423.5 | 2 | 1.08009178 | 11 | 0.396175812 | 0 | 0.188440963 | 0 |
| P17751 | Tpi1 | Triosephosphate isomerase | 1025.69 | 79.12 | 26713 | 15 | 49366594 | 141189490 | 99657570 | 100226194 | 79270738 | 78621386 | 99680954 | 78342858 | 141579250 | 115039730 | 199122594 | 284467426 | 156096050 | 207506770 | 174917794 | 225732226 | 212952322 | 129022266 | 151884994 | 183897026 | 1.092724314 | 11 | 2.176043429 | 0 | 1.923501111 | 1 |
| P38060 | Hmgcl | Hydroxymethylglutaryl-CoA lyase, mitochondrial | 490.55 | 35.08 | 34239 | 8 | 2713818.75 | 4619631 | 4973925 | 3492790 | 5131860 | 4572965.5 | 4326210.5 | 4349983 | 6684221 | 4779825.5 | 10180738 | 10605768 | 10540576 | 11068789 | 8515280 | 10566430 | 8757616 | 9084900 | 7901771 | 7933410 | 1.180640946 | 11 | 2.432213396 | 0 | 2.113705078 | 0 |
| Q7M0E3 | Dstn | Destrin | 264.71 | 37.58 | 18534 | 5 | 4196600.5 | 3060336.25 | 4590758 | 4310259 | 4519185 | 7666249 | 4288128 | 6777347 | 3951379 | 3282883.75 | 18839586 | 14956345 | 20056462 | 15165702 | 17590094 | 21653850 | 26320282 | 22759698 | 23829482 | 20930826 | 1.255782392 | 11 | 4.188596403 | 0 | 5.585595734 | 0 |

| | | | | | | | | | | | | | | | | | | | | | | | | | | | | | | | | |
|---|---|---|---|---|---|---|---|---|---|---|---|---|---|---|---|---|---|---|---|---|---|---|---|---|---|---|---|---|---|---|---|---|
| P62849 | Rps24 | Small ribosomal subunit protein eS24 | 185.11 | 28.57 | 15423 | 4 | 1350819.5 | 1239705.38 | 3000845.75 | 239768.55 | 3265210.5 | 354555.06 | 1327746.38 | 2739097.75 | 2927484 | 4508412 | 7592040 | 6956370 | 9127881 | 9266432 | 6794904 | 4689962 | 5003893.5 | 2685076.75 | 383061.19 | 4067333.25 | 1.303522359 | 11 | 4.368524562 | 0 | 1.850118705 | 6 |
| Q61425 | Hadh | Hydroxyacyl-coenzyme A dehydrogenase, mitochondrial | 135.32 | 9.24 | 34464 | 3 | 607289.75 | 662230.75 | 1256944.25 | 1385734.38 | 1741114.62 | 1064176 | 1132814 | 87011.98 | 2467141 | 2666716.25 | 2212798 | 3129296.75 | 2467107 | 2316319.5 | 1994959.12 | 1790657.38 | 959847.25 | 1049036 | 1795563.62 | 898702.94 | 1.312125871 | 11 | 2.143960322 | 0 | 1.148672704 | 9 |
| P27546 | Map4 | Microtubule-associated protein 4 | 350.39 | 8 | 117429 | 6 | 947786.44 | 1419360.88 | 1718975.5 | 1831214.75 | 2042578.38 | 1475022.12 | 892676.31 | 1649767.25 | 4878722.5 | 2146993.5 | 6506036 | 6546057.5 | 3477113.5 | 4350250 | 2579878.25 | 2856013.25 | 3470272.5 | 1964331.75 | 3886989 | 2307908.75 | 1.387349031 | 11 | 2.947183789 | 0 | 1.819807563 | 1 |
| Q8K354 | Cbr3 | Carbonyl reductase [NADPH] 3 | 134.38 | 9.75 | 30953 | 3 | 2 | 469415.56 | 2 | 2 | 705827.56 | 587063.81 | 704904.25 | 2 | 407715 | 2 | 1301904.62 | 710783.25 | 979061.56 | 1518611.25 | 903919.31 | 1294165.75 | 733597.44 | 1036044.19 | 1093878.88 | 2 | 1.446235552 | 11 | 4.606921118 | 0 | 3.537708039 | 3.5 |
| P62137 | Ppp1ca | Serine/threonine-protein phosphatase PP1-alpha catalytic subu | 409.08 | 22.42 | 37540 | 8 | 2 | 2 | 2 | 139435.06 | 243394.22 | 2 | 124146.74 | 122524.17 | 433840.66 | 2 | 265775.72 | 265987.12 | 897642.12 | 289605.16 | 743040.25 | 247912.31 | 420138.66 | 271423.16 | 240369.12 | 287153.88 | 1.777567548 | 11 | 6.431095823 | 0 | 3.831927742 | 1 |
| Q923D2 | Blvrb | Flavin reductase (NADPH) | 368.46 | 42.72 | 22197 | 5 | 2 | 744125.31 | 74026.41 | 916532.38 | 909614.75 | 2 | 126051.77 | 276664 | 666382.75 | 1300019.12 | 4655565 | 3433830.25 | 5021616.5 | 4789113 | 3993346.25 | 11786431 | 6163093 | 4988602 | 9568331 | 2956711 | 0.895934228 | 11.5 | 8.279493235 | 0 | 13.41116991 | 0 |
| Q5XFW8 | Sec13 | Protein SEC13 homolog | 47.11 | 3.42 | 35548 | 1 | 2 | 2 | 2 | 2 | 697939.81 | 2 | 2 | 2 | 2 | 388520.66 | 1745475.25 | 1459724.38 | 1456768 | 811020.25 | 1312608.5 | 1559792.88 | 1486923.62 | 1096775.25 | 2 | 1180668 | 0.556672941 | 12 | 9.722211722 | 0 | 7.628309271 | 3 |
| Q8BK64 | Ahsa1 | Activator of 90 kDa heat shock protein ATPase homolog 1 | 103.71 | 7.1 | 38117 | 2 | 2098269.5 | 3661412.5 | 2899471.75 | 4391072.5 | 3526008.75 | 4583782.5 | 3879925 | 1454635.25 | 4062477.5 | 458614.44 | 6156676 | 6984405 | 7003834.5 | 7022895 | 5911192 | 8303389.5 | 9815597 | 11243295 | 7981015 | 10530410 | 0.871092542 | 12 | 1.995567902 | 0 | 2.888092893 | 0 |
| P08556 | Nras | GTPase NRas | 159.74 | 18.52 | 21199 | 3 | 1598634.25 | 920045.88 | 2732995.5 | 1056122.25 | 2307792.5 | 1264861.12 | 1399229.38 | 229032.36 | 2440603 | 2457809 | 6250669 | 6986914.5 | 7202741 | 7389468 | 4437622 | 7290779.5 | 7415979 | 4871740 | 10011688 | 5187667.5 | 0.904352983 | 12 | 3.745235448 | 0 | 4.036618788 | 0 |
| P27659 | Rpl3 | Large ribosomal subunit protein uL3 | 496.22 | 23.82 | 46110 | 9 | 1347047 | 1125099.38 | 2912832.25 | 2416240.75 | 4255148 | 2539171 | 1186450.62 | 2001030.5 | 2195698.5 | 2982060 | 6576890 | 6229980.5 | 6310935 | 4664042 | 4682704 | 7646918.5 | 591016.06 | 2705220.25 | 3065118.5 | 3463219.75 | 0.904452417 | 12 | 2.360955884 | 0 | 1.449150686 | 9 |
| Q05816 | Fabp5 | Fatty acid-binding protein 5 | 670.01 | 64.44 | 15137 | 10 | 34245050 | 34500042 | 50742414 | 9556888 | 10992836 | 39574022 | 37961902 | 48729858 | 5096551 | 6893748 | 79571738 | 91973610 | 85874338 | 68784394 | 83715738 | 54366678 | 84873338 | 70096202 | 73705058 | 54591358 | 0.98728089 | 12 | 2.927220268 | 0 | 2.411020512 | 0 |
| P08508 | Fcgr3 | Low affinity immunoglobulin gamma Fc region receptor III | 105.92 | 8.05 | 30036 | 2 | 2566354.25 | 1890575.12 | 2654166.25 | 2185767.75 | 2148224 | 3068961.5 | 1595041 | 2115634 | 2448457.25 | 2290078.5 | 2947196.75 | 3321333.5 | 2816845.5 | 3080279 | 3746763.75 | 3012151 | 2572532.25 | 2589931.75 | 525771.38 | 4302111 | 1.006385699 | 12 | 1.390327394 | 0 | 1.136076725 | 7 |
| B5DF91 | Elavl1 | ELAV-like protein 1 | 488.16 | 27.3 | 36169 | 9 | 21195206 | 24887008 | 19642276 | 26699298 | 31745138 | 24547870 | 24290250 | 33821718 | 21071814 | 21814548 | 16274388 | 15319762 | 13510757 | 9994121 | 15633048 | 9066060 | 11128088 | 10306440 | 9135048 | 9834043 | 1.011091938 | 12 | 0.569643938 | 0 | 0.398406273 | 0 |
| Q99J93 | Ifitm2 | Interferon-induced transmembrane protein 2 | 252.04 | 17.36 | 15743 | 3 | 3165886 | 3724648.5 | 2567999.25 | 1737459.12 | 1044223.56 | 3241124 | 2254297.75 | 4923385 | 1606223.25 | 626365.62 | 2 | 2 | 511821.19 | 438021.78 | 2 | 2 | 2 | 789722.88 | 916156.06 | 2 | 1.033592477 | 12 | 0.077600668 | 0 | 0.13936722 | 0 |
| Q6NZJ6 | Eif4g1 | Eukaryotic translation initiation factor 4 gamma 1 | 539.48 | 8.56 | 176077 | 11 | 1183375.38 | 4206122.5 | 3073318.25 | 4049755 | 4895371 | 2623924.25 | 3864915.75 | 5631895 | 3101459 | 3806596.25 | 8121578.5 | 8778624 | 7193809 | 9520462 | 5708514.5 | 7098221.5 | 3593569 | 5049929 | 7026229 | 4477652 | 1.093109691 | 12 | 2.258910772 | 0 | 1.565124717 | 4 |
| P62962 | Pfn1 | Profilin-1 | 804.22 | 90 | 14957 | 13 | 79252314 | 32768326 | 86784618 | 57409862 | 115533490 | 55482106 | 54235882 | 56272182 | 119043978 | 132632346 | 215684194 | 222990082 | 254290322 | 243739762 | 166206930 | 180318818 | 192734306 | 169812578 | 300606722 | 163569986 | 1.123518643 | 12 | 2.966820212 | 0 | 2.708933895 | 0 |
| A2ADY9 | Ddi2 | Protein DDI1 homolog 2 | 116.99 | 8.52 | 44591 | 3 | 2 | 559773.69 | 2 | 2 | 2 | 2 | 2 | 2 | 664471.19 | 2 | 1633155.62 | 1699669.62 | 1126665.12 | 1194470.75 | 582048.44 | 2 | 2 | 901158.25 | 2 | 1432063.75 | 1.187032734 | 12 | 11.14007418 | 0 | 4.168103462 | 9 |
| Q9CQW9 | Ifitm3 | Interferon-induced transmembrane protein 3 | 478.05 | 48.91 | 14954 | 5 | 6990885 | 7555690.5 | 6112185.5 | 5423869 | 5122507 | 8250985 | 5130915.5 | 17265762 | 5649892.5 | 3963981.5 | 2 | 2 | 1380408.75 | 463861.12 | 807713.25 | 2014485.25 | 2664617.75 | 5119891.5 | 4826195 | 2118100.5 | 1.290221431 | 12 | 0.084985595 | 0 | 0.536555568 | 0 |
| P01900 | H2-D1 | H-2 class I histocompatibility antigen, D-D alpha chain | 747.34 | 41.92 | 41110 | 14 | 2 | 2996176 | 204206.59 | 2679340.5 | 2912626.5 | 2149102.5 | 1246608.75 | 2651931.75 | 5414471 | 1932304.5 | 13955162 | 8535286 | 5678662 | 5773805 | 5734027.5 | 5464953.5 | 5375568 | 3252325 | 6184366 | 3924042.75 | 1.52341707 | 12 | 4.512665593 | 0 | 2.752534973 | 0 |
| P97379 | G3bp2 | Ras GTPase-activating protein-binding protein 2 | 140.14 | 9.75 | 54088 | 3 | 295010 | 2 | 153760.98 | 68832.12 | 98367.58 | 2 | 2 | 275132.19 | 1735140.38 | 93032.45 | 1518125.75 | 1703441.25 | 1470523.62 | 1702373.12 | 485916.84 | 2053765.25 | 2142269.5 | 185319.78 | 2867176.25 | 2 | 3.414614135 | 12 | 11.16994439 | 0 | 11.76762057 | 5.5 |
| P02802 | Mt1 | Metallothionein-1 | 58.2 | 19.67 | 6018 | 1 | 2 | 2 | 2 | 2 | 50731.34 | 2 | 2 | 2 | 615853.06 | 2 | 403821.53 | 2120836.75 | 90694.78 | 1419229.88 | 717447.25 | 2 | 138730.78 | 2 | 2 | 2 | 12.13774282 | 12 | 93.65573518 | 0 | 2.734343411 | 12 |
| O35143 | Atp5if1 | ATPase inhibitor, mitochondrial | 56.36 | 8.49 | 12159 | 1 | 2 | 2 | 2 | 638228.69 | 1262100 | 921967.94 | 2 | 2 | 877908.44 | 2 | 2098631 | 3314753.75 | 1882607 | 1902794.12 | 1274086.25 | 5901235.5 | 3008104.5 | 1422321.12 | 4440892 | 1494093.12 | 0.947139675 | 12.5 | 5.511067169 | 0 | 8.559884912 | 0 |
| D4ACN8 | Plgrkt | Plasminogen receptor (KT) | 38.91 | 6.8 | 17312 | 1 | 2 | 2 | 2 | 2 | 2 | 2 | 2 | 2 | 2 | 2 | 848333.56 | 785997.94 | 851360.81 | 647160.12 | 676889.94 | 679351.12 | 914791.88 | 899959.94 | 2 | 2 | 1 | 12.5 | 380974.237 | 0 | 249410.694 | 5 |
| O35691 | Pnn | Pinin | 167.48 | 4.28 | 82436 | 4 | 2 | 2 | 2 | 2 | 2 | 2 | 2 | 2 | 2 | 2 | 889940.25 | 1730853 | 1276547.75 | 653501.44 | 97283.85 | 414289.41 | 2 | 2 | 2 | 2 | 1 | 12.5 | 464812.629 | 0 | 41429.741 | 10 |
| Q3TDQ1 | Stt3b | Dolichyl-diphosphooligosaccharide--protein glycosyltransferase | 37.15 | 1.09 | 93246 | 1 | 2 | 2 | 2 | 2 | 2 | 2 | 2 | 2 | 2 | 2 | 2076138.62 | 1302558.25 | 1981795.62 | 1268692.25 | 1267586.38 | 2 | 1048808.88 | 2 | 1312271.5 | 2 | 1 | 12.5 | 789677.112 | 0 | 236108.638 | 7.5 |
| Q9D1J1 | Necap2 | Adaptin ear-binding coat-associated protein 2 | 127.91 | 18.42 | 28598 | 3 | 2 | 2 | 2 | 245752.5 | 322436.03 | 2 | 303895.09 | 2 | 265070.53 | 2 | 812674.5 | 600869.25 | 367753.12 | 351569.84 | 827695.62 | 2 | 826176.56 | 2 | 2 | 197292.52 | 1.001367648 | 12.5 | 5.210473128 | 0 | 1.801275841 | 12.5 |
| Q8BRT1 | Clasp2 | CLIP-associating protein 2 | 114.8 | 1.79 | 140739 | 3 | 702967.44 | 1062777.38 | 647279.88 | 502298.75 | 780302.75 | 2 | 142333.39 | 62377.88 | 142578.33 | 2 | 2 | 2 | 2 | 237896.53 | 635456.62 | 568163.38 | 2 | 737242.12 | 2 | 670829.88 | 0.093974223 | 0 | 0.236322372 | 1 | 0.534750885 | 6 |
| Q8CG47 | Smc4 | Structural maintenance of chromosomes protein 4 | 185.01 | 2.95 | 146895 | 4 | 5528178 | 7201362 | 4967845.5 | 7203776 | 6178335 | 72138.85 | 439674.94 | 2917331.25 | 1182578.25 | 2097384.5 | 1343523.38 | 2011011.75 | 4972314 | 3593079 | 3181670.75 | 2678003.5 | 2090249.25 | 3940062 | 5139125 | 4749610 | 0.215869256 | 0 | 0.485902302 | 1 | 0.598370368 | 1 |
| P24547 | Impdh2 | Inosine-5'-monophosphate dehydrogenase 2 | 823.83 | 26.26 | 55815 | 13 | 17446900 | 16179832 | 17789444 | 14903950 | 28514204 | 5449529 | 6618694.5 | 10749625 | 5516218.5 | 10050361 | 15306482 | 14230199 | 14013454 | 13951523 | 12153343 | 10350058 | 6980839.5 | 6417638.5 | 5090295 | 4867009 | 0.404752456 | 0 | 0.734491413 | 1 | 0.355418128 | 0 |
| P97797 | Sirpa | Tyrosine-protein phosphatase non-receptor type substrate 1 | 300.77 | 16.76 | 56414 | 5 | 26574932 | 29065286 | 28924826 | 23409346 | 19369456 | 8636036 | 13319764 | 13294156 | 12952821 | 14680769 | 16087086 | 20368016 | 16670002 | 17609162 | 16556720 | 10280642 | 17884306 | 20104260 | 18982524 | 19682646 | 0.493809069 | 0 | 0.685474711 | 1 | 0.682674356 | 2 |
| P56399 | Usp5 | Ubiquitin carboxyl-terminal hydrolase 5 | 778.8 | 19.81 | 95833 | 14 | 13070915 | 16924488 | 15965610 | 12209738 | 15600410 | 11185864 | 9842009 | 7456768 | 10658277 | 12083775 | 10102161 | 12295256 | 12003999 | 10301452 | 11631137 | 10105381 | 11188433 | 16016482 | 11688334 | 11780362 | 0.694399984 | 0 | 0.763631807 | 1 | 0.823885529 | 4 |
| P09103 | P4hb | Protein disulfide-isomerase | 1679.4 | 52.65 | 57058 | 29 | 50577538 | 49967894 | 54359506 | 57374046 | 68329114 | 88081394 | 96825682 | 97847258 | 109065314 | 101177146 | 89166498 | 71966010 | 92112898 | 83260802 | 61708542 | 103315386 | 107427386 | 121451682 | 129311530 | 95977914 | 1.756887266 | 0 | 1.419113535 | 1 | 1.986699251 | 0 |
| P17136 | Snrpb | Small nuclear ribonucleoprotein-associated protein B | 173.16 | 16.02 | 23656 | 4 | 959371.31 | 1442878.12 | 918028.88 | 1791151.12 | 6923009 | 7052762 | 7067983 | 7781420.5 | 9756152 | 8229198 | 10881332 | 13079494 | 12061528 | 11410650 | 6855426.5 | 7027922.5 | 6914523.5 | 4168708.25 | 5564590 | 3979558.25 | 3.314447594 | 0 | 4.511089638 | 1 | 2.298013543 | 4 |
| Q9CR60 | Golt1b | Vesicle transport protein GOT1B | 86.94 | 10.87 | 15422 | 2 | 36930.93 | 70095.91 | 52566.57 | 124353.3 | 34206.23 | 402356.62 | 531813.56 | 311920.12 | 545582.62 | 345665.38 | 114545.47 | 265599.25 | 225204.02 | 456010.72 | 477998.69 | 2 | 1052988.25 | 570902.38 | 150759.03 | 384939.88 | 6.717958665 | 0 | 4.838421892 | 1 | 6.787903767 | 5 |
| Q61543 | Glg1 | Golgi apparatus protein 1 | 761.99 | 15.23 | 133734 | 13 | 11023077 | 10381269 | 9165269 | 8765269 | 4318496 | 2416385 | 2097386 | 3033592.5 | 4057770.5 | 5895341.5 | 1991179 | 4741745 | 2408291.25 | 1255900.25 | 2680965.5 | 2015167.5 | 2383715.75 | 3073398.75 | 1749435.12 | 1896130.75 | 0.400896231 | 1 | 0.299589196 | 1 | 0.254684697 | 0 |
| P63085 | Mapk1 | Mitogen-activated protein kinase 1 | 88.52 | 5.87 | 41276 | 2 | 1959082.75 | 6205382 | 4095243.25 | 3247900.5 | 4831905 | 6340620 | 5107902.5 | 6437119.5 | 6412637.5 | 7510327 | 10957967 | 10454359 | 6886947 | 8918702 | 5337382.5 | 9006197 | 5355044 | 6571626 | 9020926 | 6046449 | 1.563882366 | 1 | 2.09225051 | 1 | 1.769965737 | 2 |
| Q9D0S9 | Hint2 | Adenosine 5'-monophosphoramidase HINT2 | 43.74 | 7.98 | 17320 | 1 | 2 | 2 | 2 | 2 | 1039813.69 | 3880632.25 | 1666347.25 | 2713246 | 2764918.5 | 1002608.75 | 3325129.25 | 2115428.5 | 1857201.75 | 1952721.88 | 690175.44 | 2151002 | 1213091 | 2 | 2787937.25 | 1493163.25 | 11.56713008 | 1 | 9.559962939 | 1 | 7.352410104 | 3 |
| Q9D0F6 | Rfc5 | Replication factor C subunit 5 | 179.43 | 15.34 | 38096 | 4 | 2342978 | 3776439.75 | 1071435 | 2612855 | 3507526.5 | 747363 | 773547.12 | 112331.38 | 1429829.5 | 1199849.25 | 965527.94 | 843725.25 | 788172.38 | 693701.62 | 1691632.62 | 2 | 2 | 783033.06 | 2 | 2 | 0.320249811 | 2 | 0.374327408 | 1 | 0.058825579 | 0 |
| Q5U2U0 | Clpx | ATP-dependent clpX-like chaperone, mitochondrial | 107.87 | 5.21 | 69208 | 3 | 1263421.62 | 1593959.12 | 1282999.88 | 918913.88 | 1201790.62 | 2 | 1241799.25 | 2 | 611453.06 | 726342.38 | 2 | 811907.25 | 1098744.25 | 893415.31 | 2 | 659967.94 | 2 | 2 | 699500.69 | 2 | 0.41200505 | 2 | 0.447857002 | 1 | 0.217130833 | 0 |
| P70168 | Kpnb1 | Importin subunit beta-1 | 613.98 | 15.18 | 97184 | 11 | 18024284 | 18542902 | 18828454 | 20989976 | 22986042 | 12895281 | 14158646 | 18709666 | 12434120 | 17518644 | 13247630 | 12490846 | 14452863 | 14233406 | 18420588 | 11314284 | 9490507 | 12631413 | 11150694 | 13587731 | 0.76195123 | 2 | 0.73305945 | 1 | 0.58542476 | 0 |
| Q99L45 | Eif2s2 | Eukaryotic translation initiation factor 2 subunit 2 | 244.87 | 8.46 | 38092 | 5 | 4167082.5 | 4643582 | 2449438 | 6060031.5 | 7350135 | 8797467 | 9735414 | 14109189 | 12231153 | 5757658.5 | 11731560 | 9318759 | 10532087 | 9323444 | 6594662.5 | 10411003 | 9204718 | 9059762 | 13275035 | 8546372 | 2.052303585 | 2 | 1.925415264 | 1 | 2.04687229 | 0 |
| Q9CYN2 | Spcs2 | Signal peptidase complex subunit 2 | 127.61 | 10.62 | 24978 | 3 | 271372.47 | 2 | 2 | 483714.31 | 1024545.19 | 631177.62 | 579573.12 | 1368770.12 | 1435121.75 | 1434663.25 | 878667.69 | 1109073.75 | 1332435.75 | 1520549.5 | 1505352.38 | 1009167.31 | 1150611 | 1213554.38 | 1059077.75 | 1052337.25 | 3.062034007 | 2 | 3.565942236 | 1 | 3.08194922 | 1 |
| P20491 | Fcer1g | High affinity immunoglobulin epsilon receptor subunit gamma | 153.72 | 23.26 | 9652 | 3 | 2 | 931609.12 | 2 | 1123922.88 | 2157792.5 | 3271061.5 | 1921324.88 | 3008396.5 | 5067920.5 | 1606688.75 | 4555421 | 5484753 | 4560800.5 | 4709836 | 2091373.12 | 4432980 | 3293572.75 | 2238744.25 | 4874449 | 2163969.5 | 3.53055598 | 2 | 5.079638015 | 1 | 4.035696599 | 0 |
| Q8BMD8 | Slc25a24 | Mitochondrial adenyl nucleotide antiporter SLC25A24 | 385.62 | 19.16 | 52902 | 7 | 849659.69 | 638957.75 | 273005.5 | 997179.75 | 530804 | 2500985.75 | 2669683.75 | 756790.44 | 4605725 | 3778015.25 | 1866918.38 | 3077163.5 | 1291616 | 906523.44 | 1404402.62 | 718274.31 | 1482512 | 1457959.25 | 505072.62 | 1259424 | 4.350428954 | 2 | 2.598068628 | 1 | 1.648598964 | 6 |
| Q8R5M8 | Cadm1 | Cell adhesion molecule 1 | 31.83 | 1.97 | 49788 | 1 | 1131991.12 | 1296511 | 244147.12 | 764691.44 | 133796.69 | 2 | 2 | 38663.09 | 1071606.12 | 2 | 1288167 | 2115178 | 1716078.38 | 1947441.12 | 1947803.25 | 929565.75 | 2 | 2194545.75 | 1724170.62 | 2845871.25 | 0.310902409 | 3 | 2.524312793 | 1 | 2.154539177 | 7 |
| P05064 | Aldoa | Fructose-bisphosphate aldolase A | 2239.29 | 87.36 | 39356 | 36 | 29702812 | 51170450 | 24810022 | 23877682 | 39785570 | 23068214 | 19911356 | 19880840 | 6046355 | 31403190 | 18647542 | 24111222 | 17180686 | 17314482 | 22092306 | 52906678 | 25868152 | 46411154 | 53371602 | 53132102 | 0.592335441 | 3 | 0.586644642 | 1 | 1.368139517 | 4 |
| Q9DBG5 | Plin3 | Perilipin-3 | 855.54 | 45.31 | 47262 | 13 | 4867381 | 10832336 | 5951303 | 6436827 | 8322063.5 | 10788607 | 11496471 | 8265224.5 | 10891878 | 11839705 | 17024450 | 16686097 | 12075924 | 11815219 | 10385306 | 9681559 | 4586707.5 | 6326307 | 7333763 | 5236185 | 1.463389631 | 3 | 1.867266221 | 1 | 0.91086523 | 10 |
| P29341 | Pabpc1 | Polyadenylate-binding protein 1 | 1133.35 | 27.52 | 70671 | 19 | 17003464 | 23714758 | 26511816 | 19715898 | 36027870 | 43818274 | 37915126 | 46506038 | 27175668 | 26457246 | 45567326 | 46405250 | 41954474 | 41867374 | 35921430 | 35897598 | 34819002 | 31573122 | 29173354 | 36303314 | 1.478951965 | 3 | 1.721633744 | 1 | 1.364244919 | 4 |
| Q8BLN5 | Lss | Lanosterol synthase | 209.45 | 6.41 | 83141 | 4 | 2 | 2 | 103244.32 | 467896.16 | 2 | 827457.12 | 665304 | 95375.59 | 759424.19 | 421871.88 | 215322.97 | 940441.75 | 1142981.5 | 1329781.75 | 1807262.88 | 1230484.88 | 1394468.5 | 1532039.12 | 1531865.88 | 1616098.25 | 4.848901074 | 3 | 9.517332314 | 1 | 12.78998801 | 0 |
| A2A8Z1 | Osbpl9 | Oxysterol-binding protein-related protein 9 | 38.43 | 1.36 | 83109 | 1 | 2 | 2 | 105419.56 | 2 | 62603.75 | 2 | 120871.22 | 123194.48 | 258036.78 | 193560.72 | 64804.46 | 129445.76 | 149615.69 | 320465.44 | 269441.19 | 2 | 2 | 2 | 2 | 2 | 4.140141979 | 3.5 | 5.557200348 | 1 | 5.95134E-05 | 7.5 |
| P22437 | Ptgs1 | Prostaglandin G/H synthase 1 | 335.42 | 7.31 | 69042 | 6 | 4254288 | 964790.38 | 894098.75 | 3393250.25 | 6595196 | 796030.88 | 727234.62 | 2 | 2926460.25 | 1157681.12 | 8709288 | 7731776.5 | 7675873 | 7339276 | 5493212.5 | 6829953 | 5848870 | 6737907 | 6119445.5 | 4208349.5 | 0.348251151 | 4 | 2.29476402 | 1 | 1.84729976 | 4 |
| Q3UHX2 | Pdap1 | 28 kDa heat- and acid-stable phosphoprotein | 236.36 | 13.81 | 20605 | 4 | 1219113.25 | 2301653.5 | 1015344.62 | 1045551 | 1038081 | 665262.56 | 480949.59 | 2 | 670040.06 | 1550873 | 3009455.75 | 3891707 | 3327132.25 | 2650756.75 | 2129460.25 | 7038349 | 3477568 | 2639276.75 | 6319377.5 | 5136619.5 | 0.508649206 | 4 | 2.267234719 | 1 | 3.717846656 | 0 |
| P41498 | Acp1 | Low molecular weight phosphotyrosine protein phosphatase | 168.01 | 13.92 | 18152 | 3 | 7733582 | 5949479 | 6473837 | 4231382.5 | 7086816 | 4303485.5 | 4753398 | 4807112 | 4046285 | 5353144 | 8176747 | 8014165.5 | 10081861 | 7470591.5 | 12888423 | 9088131 | 9813065 | 8269658 | 9215791 | 7692987 | 0.73910574 | 4 | 1.481545513 | 1 | 1.400460583 | 1 |
| P25911 | Lyn | Tyrosine-protein kinase Lyn | 553.23 | 24.02 | 58812 | 9 | 7142423.5 | 6123755.5 | 6448264 | 4608900.5 | 6531477.5 | 4185935.5 | 4722080 | 2879500.5 | 5550315.5 | 6349535.5 | 3081475.25 | 3749627.5 | 5235096 | 4034160 | 3497931 | 3521082.5 | 2951183 | 4583672.5 | 3326545 | 665942 | 0.767703919 | 4 | 0.635177555 | 1 | 0.487717138 | 0 |
| P05063 | Aldoc | Fructose-bisphosphate aldolase C | 1252.38 | 61.16 | 39395 | 21 | 13274122 | 21123936 | 14549677 | 11569550 | 24268408 | 22008330 | 24583246 | 24554790 | 23588642 | 18820232 | 38415966 | 41127062 | 34667174 | 38105294 | 21977542 | 29246642 | 36152190 | 26580400 | 31875758 | 25395772 | 1.339320774 | 4 | 2.055689254 | 1 | 1.760329564 | 0 |
| Q9QUJ7 | Acsl4 | Long-chain-fatty-acid--CoA ligase 4 | 704.09 | 20.25 | 79077 | 13 | 8631226 | 15733784 | 6813302 | 12792210 | 10441894 | 18128884 | 17248188 | 16297322 | 13496159 | 9729802 | 6783487.5 | 6693440 | 6636458.5 | 8391071 | 6725141 | 5313456.5 | 6653838 | 8528710 | 5275641 | 4391894.5 | 1.376530588 | 4 | 0.647455132 | 1 | 0.554350316 | 1 |
| Q9Z0N1 | Eif2s3x | Eukaryotic translation initiation factor 2 subunit 3, X-linked | 751.19 | 33.26 | 51065 | 13 | 3247721 | 4302019 | 6345901 | 5795777.5 | 10736560 | 7081690.5 | 8304664 | 11182894 | 7308629 | 8965900 | 22658074 | 26041514 | 16102489 | 16729072 | 9312756 | 16178032 | 12088381 | 9674222 | 17527564 | 14084660 | 1.408038904 | 4 | 2.985538622 | 1 | 2.28581925 | 1 |
| Q9D6Z1 | Nop56 | Nucleolar protein 56 | 803.37 | 26.03 | 64464 | 14 | 2101565.5 | 4271047.5 | 7835113 | 2829558.5 | 5662746 | 5811211.5 | 6250350 | 7556915 | 8255882 | 6105906 | 7831085 | 8060398.5 | 10093548 | 10671801 | 12949663 | 11062607 | 8238627.5 | 15962203 | 12282738 | 10925965 | 1.49692594 | 4 | 2.185305236 | 1 | 2.575861759 | 0 |
| P68101 | Eif2s1 | Eukaryotic translation initiation factor 2 subunit 1 | 396.54 | 27.3 | 36108 | 8 | 9737491 | 11147962 | 13358570 | 11767718 | 21204608 | 20977982 | 18835206 | 27986656 | 17727166 | 15158821 | 29855744 | 23828778 | 26518606 | 24949358 | 14651443 | 22947946 | 15905980 | 13793750 | 22748670 | 5482553.5 | 1.497936626 | 4 | 1.782362934 | 1 | 1.203262312 | 7 |
| Q8BTM8 | Flna | Filamin-A | 4042.23 | 33.36 | 281222 | 69 | 67233162 | 91182226 | 83583186 | 30546066 | 42653442 | 105562506 | 105359938 | 160628706 | 82822562 | 75044834 | 128851098 | 143336658 | 99317506 | 108401874 | 90491330 | 174092962 | 126538906 | 102280722 | 133019034 | 118139626 | 1.679637587 | 4 | 1.809650815 | 1 | 2.075111771 | 0 |
| P05765 | Rps21 | Small ribosomal subunit protein eS21 | 315.26 | 59.04 | 9127 | 5 | 374543.31 | 1071584.62 | 1777041.5 | 3805792 | 9552724 | 8348942.5 | 6061709.5 | 12740052 | 8698547 | 5700216 | 23094284 | 19644056 | 12806833 | 13403214 | 6946013.5 | 12303262 | 11472316 | 7197595.5 | 15709172 | 6024511 | 2.505744496 | 4 | 4.577001585 | 1 | 3.178618767 | 2 |
| P32233 | Drg1 | Developmentally-regulated GTP-binding protein 1 | 175.45 | 12.81 | 40512 | 4 | 2 | 1028853.94 | 1470820.38 | 985166.31 | 417233.94 | 75580.82 | 130172.73 | 2 | 2 | 526585.62 | 1142197 | 1480585.62 | 2542329 | 2435134.5 | 2250585.25 | 1115609.88 | 1227403 | 1277250.88 | 1265857.75 | 1428072.12 | 0.187680369 | 5 | 2.524510012 | 1 | 1.618162411 | 5 |
| Q8BZM1 | Glmn | Glomulin | 56.34 | 1.85 | 67756 | 1 | 791166.56 | 133686.56 | 439505.53 | 232825.61 | 564132.19 | 2 | 151746.5 | 2 | 257009.48 | 421766.03 | 2 | 2 | 2 | 210056.2 | 2 | 2 | 191477.7 | 2 | 2 | 2 | 0.384268583 | 5 | 0.097192709 | 1 | 0.08859679 | 1 |
| Q9QXT0 | Cnpy2 | Protein canopy homolog 2 | 83.99 | 8.79 | 20767 | 1 | 2061376.12 | 1992621.25 | 1703973.5 | 2076097 | 1623956.38 | 1323680.75 | 1469686.38 | 2620545.75 | 1204640.5 | 2 | 1639381.12 | 1550820.88 | 1499650.62 | 1580990.25 | 1199752.75 | 2695128.25 | 1791855 | 3256960 | 2901415 | 2702003 | 0.699782027 | 5 | 0.78986852 | 1 | 1.411220874 | 3 |
| P48428 | Tbca | Tubulin-specific chaperone A | 63.77 | 8.33 | 12758 | 1 | 3631210.25 | 5323165.5 | 3098622.75 | 3899259.25 | 4044558.5 | 3354873.5 | 2392969.25 | 4248079.5 | 2245423.75 | 2687674.5 | 2959594.75 | 2828885.25 | 3176474.25 | 3051736 | 2317306 | 4241948.5 | 3960428 | 3377049 | 4706671.5 | 3695553.25 | 0.74656987 | 5 | 0.71681392 | 1 | 0.999241579 | 11 |
| Q5U2Q7 | Etf1 | Eukaryotic peptide chain release factor subunit 1 | 227.15 | 11.44 | 49031 | 4 | 3757929 | 3953695.25 | 5297127.5 | 4717134.5 | 6760934.5 | 2989044 | 4771884.5 | 3185173.5 | 3824166 | 3867261.25 | 2439071.25 | 3950169.5 | 3343839.75 | 2787527.75 | 3521739 | 1511522.75 | 2352325.75 | 2378620.5 | 977832.44 | 2471458 | 0.76112491 | 5 | 0.655142103 | 1 | 0.395794927 | 0 |
| Q8VDL4 | Adpgk | ADP-dependent glucokinase | 225.31 | 9.27 | 53902 | 3 | 4469715.5 | 4528430 | 3451894.5 | 4140623 | 4890851 | 4064973.5 | 3164527.25 | 4282185.5 | 3901249.25 | 3951973 | 2485890.25 | 2583490 | 3343616.25 | 4054239.75 | 2907139.75 | 3467294.5 | 3604531 | 4875579 | 1220416.12 | 4360444.5 | 0.901468514 | 5 | 0.715702627 | 1 | 0.815969727 | 8 |
| P62702 | Rps4x | Small ribosomal subunit protein eS4 | 887.81 | 63.12 | 29598 | 18 | 8850566 | 15992792 | 18780736 | 21787708 | 59851322 | 26947562 | 22876694 | 36154642 | 44602270 | 44060442 | 83222882 | 70571058 | 62999178 | 77136322 | 28657908 | 56885578 | 29326664 | 19345578 | 47767922 | 20613154 | 1.394198104 | 5 | 2.575277845 | 1 | 1.388588201 | 7 |
| Q9DC16 | Ergic1 | Endoplasmic reticulum-Golgi intermediate compartment protein | 42.45 | 3.45 | 32562 | 1 | 697680.62 | 671538.06 | 411739.53 | 550431.94 | 794284.56 | 1662543.5 | 1299834.62 | 3129918.25 | 888176 | 2 | 1210114 | 1181084.38 | 1222404.12 | 940654.5 | 734632.19 | 1225969 | 950859.62 | 940519.69 | 961146.62 | 1002520.81 | 2.233269619 | 5 | 1.692079209 | 1 | 1.625574064 | 0 |
| P62821 | Rab1a | Ras-related protein Rab-1A | 881.07 | 52.2 | 22678 | 13 | 1085382.5 | 670715.62 | 1800761.62 | 2902218.5 | 2025087.62 | 4882745 | 5294436.5 | 978946.62 | 2742710.75 | 5888267 | 3175549.75 | 2175311.75 | 4424819 | 5082927 | 8261834.5 | 1462007.5 | 3745409 | 3896757.25 | 3015960.75 | 6673385 | 2.332239397 | 5 | 2.725128478 | 1 | 2.215128725 | 3 |
| Q4KM73 | Cmpk1 | UMP-CMP kinase | 175.47 | 25.51 | 22169 | 4 | 2 | 2 | 2 | 78223.92 | 720241.81 | 2 | 254394.02 | 228135.98 | 877185.12 | 845820.62 | 475871.12 | 1205477.12 | 1268929.75 | 1454895 | 1478276.25 | 244149.39 | 506205.94 | 155628 | 161835.38 | 501818.44 | 2.762198907 | 5.5 | 7.368387657 | 1 | 1.965801782 | 5 |
| P23249 | Mov10 | Putative helicase MOV-10 | 75.68 | 1.79 | 113583 | 2 | 1101826.12 | 911823.44 | 1500767.38 | 1024390.88 | 1307391.5 | 96269.71 | 195562.91 | 1200971.75 | 883958.69 | 1172227.25 | 2 | 2 | 836688.88 | 969424 | 197143.97 | 626249.38 | 1246640.12 | 2 | 733190.69 | 1294621.12 | 0.60705941 | 6 | 0.342660375 | 1 | 0.667220376 | 6 |
| P63101 | Ywhaz | 14-3-3 protein zeta/delta | 876.86 | 48.57 | 27771 | 14 | 47919178 | 58831214 | 49123194 | 74235818 | 40057706 | 82341122 | 90425002 | 27328360 | 95403346 | 72581258 | 93968882 | 108277210 | 90550914 | 98461554 | 70538362 | 67332882 | 61610174 | 62501810 | 49181578 | 56384266 | 1.362412649 | 6 | 1.70930104 | 1 | 1.099359245 | 7 |
| O55125 | Nipsnap1 | Protein NipSnap homolog 1 | 51.35 | 3.17 | 33363 | 1 | 970621.06 | 1782762 | 1352573 | 759483.62 | 2074820.12 | 2043409.38 | 625542.12 | 2271984 | 2905218.75 | 2850992.5 | 6771081.5 | 4350709.5 | 4932520 | 5264809.5 | 1971781.62 | 6145674.5 | 4341445 | 897802.75 | 5653898 | 2991918 | 1.541317913 | 6 | 3.35591214 | 1 | 2.886165479 | 4 |
| P26645 | Marcks | Myristoylated alanine-rich C-kinase substrate | 129.21 | 13.27 | 29661 | 3 | 1038699.94 | 2352934.5 | 662563.19 | 1498296 | 1080071.5 | 3210994.5 | 3774237.5 | 1435101.62 | 4622386.5 | 670658.69 | 2404413.75 | 3860567.5 | 3453738.5 | 3277946.25 | 2209690 | 2727933 | 4614228 | 2069344.88 | 3947676 | 2844894.5 | 2.067582985 | 6 | 2.292680992 | 1 | 2.44310852 | 1 |
| P70296 | Pebp1 | Phosphatidylethanolamine-binding protein 1 | 41.44 | 3.74 | 20830 | 1 | 2 | 2 | 2 | 1103057.12 | 2185235.75 | 1938327 | 676842.25 | 1961779.25 | 2475145.25 | 971388.5 | 7972560 | 7046345 | 2641511.5 | 3576372.5 | 1949398.12 | 7590649.5 | 4176174.5 | 1275887.5 | 5716551 | 940269.69 | 2.440010038 | 6 | 7.051119146 | 1 | 5.99079736 | 3 |
| Q9CSN1 | Snw1 | SNW domain-containing protein 1 | 369.86 | 17.54 | 61475 | 8 | 2 | 3225260 | 2 | 748591.12 | 2 | 1584166.88 | 2725309 | 3213320.5 | 2799418.75 | 99503.89 | 3965288 | 3768437 | 3778947 | 4548942 | 1161029.88 | 1799121.25 | 2154485 | 658319.88 | 892098.56 | 769261 | 2.622570139 | 6 | 4.333986694 | 1 | 1.578638965 | 6 |
| P60603 | Romo1 | Reactive oxygen species modulator 1 | 195.32 | 77.22 | 8183 | 3 | 2 | 2 | 2 | 2 | 728003.69 | 1422486.12 | 2 | 920095.94 | 1575996 | 2 | 3609320 | 2835384 | 571272.06 | 4708662.5 | 1169851.62 | 2 | 1928126.88 | 1824637.25 | 468633.03 | 1364080.25 | 5.382581233 | 6 | 17.71192737 | 1 | 7.672238628 | 4 |
| Q07076 | Anxa7 | Annexin A7 | 543.49 | 19.65 | 49925 | 9 | 8731718 | 10115022 | 8965135 | 7079648 | 6936030 | 5356104.5 | 6052643 | 7622528 | 11759929 | 6648532.5 | 14224084 | 16506326 | 12418379 | 12174210 | 9414593 | 13054605 | 12110024 | 12165517 | 9233605 | 13062747 | 0.895097473 | 7 | 1.547726017 | 1 | 1.425531587 | 1 |
| P14131 | Rps16 | Small ribosomal subunit protein uS9 | 624.1 | 54.79 | 16445 | 10 | 9203700 | 14366713 | 14692505 | 24903804 | 11723616 | 14651200 | 12993121 | 28601722 | 23464372 | 20665994 | 25863874 | 30504374 | 31843082 | 37536246 | 16781038 | 29447054 | 12002095 | 9639003 | 20517298 | 3999179.5 | 1.34031187 | 7 | 1.903164251 | 1 | 1.009537833 | 12 |
| Q3UZ39 | Lrrfip1 | Leucine-rich repeat flightless-interacting protein 1 | 546.07 | 18.52 | 79249 | 8 | 2625397.5 | 5709532.5 | 488388.03 | 6881398 | 5834521 | 4324096 | 6043938 | 8378474.5 | 9619907 | 4985294 | 6705957.5 | 11344484 | 8392904 | 8992784 | 7395230 | 9144341 | 8390963 | 6001216 | 7639786.5 | 6415298 | 1.548416476 | 7 | 1.988527237 | 1 | 1.745261657 | 2 |
| Q8VIM6 | Strc | Stereocilin | 40.3 | 0.44 | 196347 | 1 | 2154606 | 3525665.5 | 2 | 1996990.5 | 171278.95 | 2187306.25 | 4929728.5 | 679189.94 | 4668975 | 806825.31 | 4498121 | 8721095 | 5526650 | 4215881 | 2777004.5 | 4511458.5 | 4478902 | 2679500.5 | 176561.31 | 2765789.5 | 1.691017694 | 7 | 3.279430547 | 1 | 1.861773823 | 5 |
| Q9WUM5 | Suclg1 | Succinate--CoA ligase [ADP/GDP-forming] subunit alpha, mitoc | 268.55 | 20.23 | 36155 | 5 | 2153205.75 | 913808.62 | 2 | 215440.64 | 62207.23 | 231678.27 | 588188.56 | 452031.19 | 3963857.75 | 1466597.25 | 4705555.5 | 5208400 | 1198194 | 2327275.5 | 2726630.5 | 2265304.5 | 1073692.38 | 771086.25 | 2 | 1179045.38 | 2.003894125 | 7 | 4.833386654 | 1 | 1.581363668 | 8.5 |
| Q8CHH9 | Septin8 | Septin-8 | 289.12 | 12.12 | 49812 | 5 | 3168328.75 | 790347.69 | 49502.61 | 183969.53 | 444451.81 | 2588252 | 2 | 88398.19 | 636710.31 | 2 | 2476879.75 | 3598256.75 | 4063481.5 | 3902357.25 | 4929576 | 3257078.25 | 5464676 | 3725837.75 | 577234.25 | 6313775.5 | 0.714610754 | 8 | 4.091478595 | 1 | 4.170857983 | 2 |
| Q60605 | Myl6 | Myosin light polypeptide 6 | 671.06 | 75.5 | 16930 | 11 | 102741986 | 192252562 | 123232466 | 158589266 | 128623970 | 130729938 | 118422634 | 154811186 | 95665810 | 122879106 | 87240922 | 83216290 | 97496970 | 107707954 | 84022106 | 93629010 | 107849362 | 138046114 | 127537986 | 130677962 | 0.882439971 | 8 | 0.651627465 | 1 | 0.847329641 | 9 |
| P47791 | Gsr | Glutathione reductase, mitochondrial | 317.81 | 10.4 | 53663 | 6 | 8307580 | 12517123 | 9198177 | 6652179.5 | 9185014 | 8983948 | 9032163 | 6930154 | 8960204 | 7774349.5 | 21388952 | 20359132 | 15323620 | 14508521 | 9236604 | 28001674 | 14368759 | 13363396 | 4995836 | 10838600 | 0.908869422 | 8 | 1.762248135 | 1 | 1.560578942 | 6 |
| Q02053 | Uba1 | Ubiquitin-like modifier-activating enzyme 1 | 1431.37 | 27.41 | 117809 | 22 | 12910365 | 13192582 | 9111129 | 16518201 | 17878580 | 14086806 | 15734254 | 10452510 | 18400642 | 19026422 | 16603296 | 23405408 | 21062928 | 22515174 | 21584850 | 23774020 | 19454994 | 13842178 | 13571138 | 15998115 | 1.1162143 | 8 | 1.510851332 | 1 | 1.244639827 | 6 |
| Q9CYG7 | Tomm34 | Mitochondrial import receptor subunit TOM34 | 340 | 24.6 | 34278 | 6 | 3941583.25 | 4968733.5 | 1972981.62 | 3836345 | 3722064.5 | 5365718.5 | 5365953.5 | 6517231.5 | 3640912.25 | 3200465 | 6479340.5 | 6504250 | 5319075.5 | 6143359 | 4189747.75 | 5403128 | 5171352 | 4974756.5 | 4855053 | 3802060.25 | 1.306293372 | 8 | 1.552772279 | 1 | 1.312587203 | 4 |
| O88342 | Wdr1 | WD repeat-containing protein 1 | 765.62 | 21.45 | 66407 | 13 | 17124174 | 26997010 | 47996186 | 18269048 | 70113834 | 57356818 | 40543318 | 43077626 | 54174802 | 46679986 | 78924010 | 92804290 | 83290738 | 79082266 | 69264234 | 82308970 | 70261386 | 75680354 | 79777826 | 68370818 | 1.33979065 | 8 | 2.234709002 | 1 | 2.085312069 | 1 |
| D3Z6Q9 | Bin2 | Bridging integrator 2 | 251.4 | 10.22 | 52554 | 4 | 2095120.12 | 930018.38 | 2 | 1553352.75 | 2885104 | 2384462.5 | 2110697.5 | 1218321.75 | 3162429.75 | 1208974.38 | 4481338 | 5000398.5 | 3642050 | 4737144 | 2269586.5 | 4356563 | 4434156 | 3362007 | 4922474.5 | 3540454.75 | 1.351209818 | 8 | 2.69716014 | 1 | 2.762160733 | 0 |
| P22509 | Fbl | rRNA 2'-O-methyltransferase fibrillarin | 507.67 | 39.45 | 34222 | 9 | 939790.69 | 3651883.5 | 1039457.06 | 2869129 | 9344312 | 1652853.25 | 5250978 | 2713877 | 11360373 | 5593778 | 11051714 | 13313134 | 10059814 | 11597807 | 4485952.5 | 12271223 | 9423556 | 4107783.25 | 5634924 | 5668565.5 | 1.489072356 | 8 | 2.830464121 | 1 | 2.079402702 | 3 |
| Q4VAA2 | Cdv3 | Protein CDV3 | 53.5 | 10.68 | 29729 | 1 | 2 | 1712873 | 2 | 2219471.5 | 384129.78 | 2968753 | 1443390.25 | 498945.81 | 1665730.62 | 1206063.5 | 3441231.5 | 2800357.75 | 1721035.5 | 2730475.75 | 2461540.5 | 1078548.5 | 1761052.62 | 1371977.62 | 2941863.5 | 961789.81 | 1.803063209 | 8 | 3.047540181 | 1 | 1.880058586 | 7 |
| P32261 | Serpinc1 | Antithrombin-III | 194.37 | 9.03 | 52004 | 4 | 15376298 | 4411473 | 15051972 | 20656882 | 10280195 | 5146518.5 | 5930419 | 13018703 | 16798842 | 9013863 | 2873241.75 | 3827876.5 | 5890942.5 | 3566920.5 | 4130134 | 2261847.75 | 6497517 | 6920275 | 1460917.25 | 7497973 | 0.758752787 | 9 | 0.308453879 | 1 | 0.374577701 | 3 |
| Q9DBL1 | Acadsb | Short/branched chain specific acyl-CoA dehydrogenase, mitoch | 207.01 | 11.57 | 47874 | 4 | 7018289.5 | 14215144 | 9009966 | 9093174 | 6199821 | 8367582 | 7702696.5 | 8207247 | 6950683 | 7949475 | 5699357 | 3853466 | 4785689.5 | 6292184.5 | 3182772.5 | 4583921.5 | 3738578.75 | 3548650.25 | 5525142 | 3196016.25 | 0.860359805 | 9 | 0.522954656 | 1 | 0.452216496 | 0 |
| P17047 | Lamp2 | Lysosome-associated membrane glycoprotein 2 | 317.8 | 14.22 | 45681 | 7 | 56947722 | 43788750 | 45855514 | 33826986 | 22624090 | 38065446 | 40730018 | 38625830 | 33146262 | 37056942 | 18400476 | 16979212 | 17309448 | 15681099 | 24729204 | 20627916 | 28342646 | 21965152 | 3258490.5 | 23858418 | 0.924062591 | 9 | 0.458520661 | 1 | 0.482915405 | 2 |
| Q99K85 | Psat1 | Phosphoserine aminotransferase | 625.66 | 40.81 | 40473 | 12 | 9448713 | 8158965 | 14012177 | 10083978 | 12094220 | 10817532 | 10010280 | 17130962 | 10743582 | 11489357 | 15295362 | 14744097 | 17872146 | 16459576 | 13024913 | 15648109 | 16903908 | 18258028 | 12322903 | 15239247 | 1.118845565 | 9 | 1.438641172 | 1 | 1.45678497 | 1 |
| Q6NSR8 | Npepl1 | Probable aminopeptidase NPEPL1 | 67.5 | 2.29 | 55940 | 1 | 2 | 2 | 890333.69 | 317294.38 | 486473.44 | 1083593.38 | 619240.38 | 63026.04 | 2 | 700344 | 1203304.5 | 1178328.88 | 1224975.25 | 1205976.75 | 850227.94 | 1212360.88 | 1614224.62 | 913143.62 | 1589625.5 | 1498088.12 | 1.455756909 | 9 | 3.34265681 | 1 | 4.030116601 | 0 |
| P48774 | Gstm5 | Glutathione S-transferase Mu 5 | 123.26 | 11.16 | 26635 | 3 | 2 | 549747.81 | 2 | 1429050 | 1190204.12 | 984227.94 | 717092.12 | 1067435.88 | 1268109.5 | 679128.81 | 2063216.38 | 1471445.38 | 2163423.5 | 1559212.5 | 1296197.5 | 1229201.38 | 2638258.5 | 2185829.25 | 1888519.62 | 2039153.25 | 1.488162015 | 9 | 2.699109894 | 1 | 3.149556113 | 1 |
| P60122 | Ruvbl1 | RuvB-like 1 | 546.91 | 29.39 | 50214 | 11 | 7201930 | 7734591 | 8404010 | 6199657.5 | 15437682 | 5247495 | 6964462.5 | 6630665 | 10715962 | 8675510 | 5501372 | 3646909.5 | 6413640 | 5964866 | 5495590 | 6543438 | 7358740 | 8459851 | 7761210.5 | 6903392 | 0.850064578 | 10 | 0.600792728 | 1 | 0.823218865 | 11 |
| P60060 | Sec61g | Protein transport protein Sec61 subunit gamma | 114.04 | 29.41 | 7741 | 2 | 2145020.25 | 4049843.75 | 2268142.75 | 2257530.5 | 2828570.25 | 2963437 | 2644186 | 1561022.62 | 3605219 | 3012158.5 | 1474205 | 1136984.62 | 802669.62 | 2148376.5 | 2 | 1142242.38 | 945471.56 | 819531.19 | 1675090.5 | 2 | 1.017485699 | 10 | 0.41052429 | 1 | 0.338202175 | 0 |
| Q8CG76 | Akr7a2 | Aflatoxin B1 aldehyde reductase member 2 | 180.37 | 12.81 | 40612 | 4 | 2 | 1342043.38 | 2 | 1417955.75 | 1532417 | 2 | 1836531.88 | 841564.5 | 1245708.5 | 1942688 | 2633762.5 | 2638105.75 | 2029602.25 | 2469918.25 | 1491462.5 | 1890017.25 | 615620.94 | 1384285.88 | 1285391.62 | 1171690.88 | 1.366710318 | 10 | 2.623893027 | 1 | 1.478654553 | 11 |
| Q921T2 | Tor1aip1 | Torsin-1A-interacting protein 1 | 194.77 | 11.6 | 66781 | 5 | 469708.75 | 2 | 2 | 2 | 274803.16 | 2 | 2 | 194161.97 | 5230633 | 455051.12 | 397764.31 | 3967362 | 608338.19 | 2024274.62 | 940521.75 | 2383519.25 | 2 | 1124261.25 | 2 | 852345.12 | 7.897526723 | 10 | 10.66228329 | 1 | 5.856312604 | 7 |
| O88738 | Birc6 | Dual E2 ubiquitin-conjugating enzyme/E3 ubiquitin-protein ligas | 46.51 | 0.23 | 532170 | 1 | 68989.83 | 2 | 155151.41 | 2 | 82147.55 | 2 | 2 | 2 | 144146.98 | 96737.19 | 116538.2 | 219438.5 | 189859.17 | 237339.33 | 478516.47 | 2 | 229710.17 | 284162.19 | 2 | 212691.83 | 0.786470259 | 11 | 4.053936986 | 1 | 2.372136119 | 8 |
| Q8C2Q3 | Rbm14 | RNA-binding protein 14 | 572.65 | 18.98 | 69449 | 11 | 387683.72 | 1151487 | 805170.56 | 920492.88 | 2944652.5 | 3275644 | 2 | 2 | 900353.44 | 1277022.12 | 7480181 | 6802167 | 2050537.88 | 5843940 | 6195020.5 | 4684507.5 | 7674811.5 | 2659742.5 | 4229229 | 3148073.75 | 0.878176226 | 11 | 4.569113026 | 1 | 3.60679803 | 1 |
| P32921 | Wars1 | Tryptophan--tRNA ligase, cytoplasmic | 239.9 | 14.55 | 54358 | 5 | 2478380 | 166607.94 | 2427810 | 1017236.12 | 3693346 | 1664800.88 | 1919538.5 | 651562.56 | 2500688 | 4592180 | 3933457.5 | 4300569 | 5039679.5 | 3440914 | 7650628 | 2307206 | 2936172 | 2754301 | 1356623.5 | 4536018 | 1.157960732 | 11 | 2.49047342 | 1 | 1.419787478 | 8 |
| Q8BFR4 | Gns | N-acetylglucosamine-6-sulfatase | 477.23 | 17.65 | 61175 | 8 | 27891990 | 41338570 | 24056404 | 20303390 | 21449938 | 26083300 | 25711392 | 26993846 | 21171752 | 22315932 | 18783092 | 18641872 | 20313132 | 18097350 | 20170512 | 19881048 | 21256854 | 25787610 | 23036426 | 27720032 | 0.905479544 | 12 | 0.710943057 | 1 | 0.871458202 | 9 |
| P41216 | Acsl1 | Long-chain-fatty-acid--CoA ligase 1 | 748.87 | 20.46 | 77951 | 13 | 15011382 | 7030842.5 | 15467761 | 9452538 | 12973224 | 8044199 | 9666626 | 9527546 | 13460395 | 16206658 | 4069211 | 3346427.75 | 6735819 | 3750970.25 | 7076272.5 | 4907084 | 3824109 | 8155899.5 | 1758339 | 6194244 | 0.949440465 | 12 | 0.416757971 | 1 | 0.414438403 | 1 |
| Q9CR57 | Rpl14 | Large ribosomal subunit protein eL14 | 355.34 | 25.81 | 23564 | 6 | 5217499.5 | 8229449.5 | 5476875 | 10405872 | 15755802 | 6141344.5 | 5037366.5 | 9524234 | 11538332 | 11368322 | 20265922 | 17366108 | 18473098 | 20530322 | 14041842 | 12062041 | 12331687 | 12807103 | 17410970 | 12026771 | 0.967264441 | 12 | 2.011229686 | 1 | 1.478048928 | 4 |
| P18242 | Ctsd | Cathepsin D | 1137.54 | 48.78 | 44954 | 18 | 154405586 | 189911314 | 153709570 | 98533914 | 88569002 | 123585466 | 139359426 | 125183786 | 119250498 | 163430354 | 87023042 | 83809050 | 93645130 | 88233130 | 85334074 | 95319674 | 91113586 | 104785394 | 79968218 | 108342714 | 0.979099049 | 12 | 0.639360149 | 1 | 0.699910989 | 6 |
| P51912 | Slc1a5 | Neutral amino acid transporter B(0) | 455.44 | 20.25 | 58483 | 8 | 88173.43 | 2 | 98026.14 | 137603.83 | 107941.81 | 91752 | 2 | 133919.66 | 192604.41 | 74424.1 | 176309.69 | 158378.81 | 375649.84 | 116905.12 | 1149368.12 | 78287.69 | 263054.19 | 127542.95 | 850940.56 | 152516 | 1.141182059 | 12.5 | 4.578168739 | 1 | 3.410193178 | 5 |
| P47963 | Rpl13 | Large ribosomal subunit protein eL13 | 447.66 | 33.65 | 24305 | 8 | 491075.75 | 2 | 2 | 2 | 1517240.88 | 2 | 2 | 193586.69 | 3112993 | 2 | 9271817 | 7147376.5 | 2198282.75 | 3235460.5 | 1332002.75 | 6665209 | 2071671.12 | 2 | 1519857.5 | 2 | 1.646441483 | 12.5 | 11.54442974 | 1 | 5.107118481 | 7 |
| Q9QY81 | Nup210 | Nuclear pore membrane glycoprotein 210 | 439.78 | 7.37 | 204101 | 9 | 2703440.75 | 5567009.5 | 3143422 | 4788574.5 | 3050809.5 | 1240138.88 | 894893.5 | 1861990 | 1444105.25 | 1907624.88 | 2759881.25 | 1597330.38 | 2182414.75 | 2765088.25 | 2090787.5 | 1835450.38 | 3898544 | 2243327.75 | 2196633.25 | 777138.12 | 0.381688812 | 0 | 0.591874018 | 2 | 0.56879176 | 3 |
| Q6ZPJ3 | Ube2o | (E3-independent) E2 ubiquitin-conjugating enzyme UBE2O | 40.76 | 0.78 | 140834 | 1 | 1493602 | 1744390.88 | 1423655 | 1196438 | 1860140.38 | 2 | 1018038.81 | 230029.14 | 758510.19 | 986246.12 | 1353032.25 | 888065.62 | 1169581.38 | 1371126.12 | 761636.12 | 1021396.25 | 1073916.25 | 1202322.38 | 1227189.75 | 1454170.38 | 0.387760887 | 0 | 0.718227388 | 2 | 0.774659204 | 4 |
| Q62433 | Ndrg1 | Protein NDRG1 | 431.7 | 21.32 | 43009 | 6 | 2195349.25 | 3200801.75 | 4223435.5 | 1512579.75 | 4214666.5 | 7044810 | 5368003.5 | 7987352.5 | 5291346 | 7852961.5 | 12203046 | 9599240 | 7615808 | 4357332 | 4164202 | 6741291.5 | 5685017 | 6320821 | 7349623 | 7718850 | 2.185758719 | 0 | 2.472147095 | 2 | 2.203425492 | 0 |
| P26284 | Pdha1 | Pyruvate dehydrogenase E1 component subunit alpha, somatic | 173.88 | 8.21 | 43227 | 4 | 2124065.75 | 2638453.25 | 2264161.5 | 1446405.75 | 3208518.75 | 5669742 | 5143121.5 | 5918887.5 | 5296373 | 3923458.25 | 6045716 | 5452364 | 4275858 | 3547269.5 | 2343097.5 | 5181142 | 2607957 | 821036.75 | 2301765 | 2852910 | 2.221576765 | 0 | 1.85456579 | 2 | 1.178332151 | 10 |
| O09172 | Gclm | Glutamate--cysteine ligase regulatory subunit | 120.51 | 7.66 | 30535 | 2 | 557746.31 | 1343189 | 2 | 1719910.25 | 2426111.25 | 4069312 | 3859213.75 | 5964217 | 4333771.5 | 3143511.25 | 6221283.5 | 5728182 | 5130086.5 | 5111438.5 | 1620347.75 | 8920770 | 3473076.75 | 1971533.38 | 6062396.5 | 1696200.12 | 3.534012083 | 0 | 3.937737795 | 2 | 3.658694799 | 3 |
| Q8BJS4 | Sun2 | SUN domain-containing protein 2 | 301.18 | 9.44 | 81605 | 6 | 1270364.5 | 1729638.5 | 2022400.62 | 1784884.38 | 420095.06 | 2 | 2 | 2 | 437306.12 | 2 | 2 | 2 | 148818.41 | 180637.84 | 1551082.5 | 2134424 | 136555.66 | 2 | 2 | 2688659.5 | 0.060507948 | 1 | 0.260196911 | 2 | 0.686229458 | 10 |
| Q61072 | Adam9 | Disintegrin and metalloproteinase domain-containing protein 9 | 164.58 | 7.46 | 92079 | 4 | 1861648.25 | 2290644 | 1780622.75 | 1411145.25 | 873784.44 | 2 | 502634.91 | 120830.77 | 1226207.25 | 173120.5 | 108338.09 | 2 | 881054 | 1160993 | 741829 | 643923 | 429908.94 | 1185651.38 | 2 | 819297.75 | 0.24614671 | 1 | 0.351943387 | 2 | 0.374646052 | 1 |
| Q8BFQ4 | Wdr82 | WD repeat-containing protein 82 | 133.47 | 12.14 | 35079 | 3 | 1300087.12 | 1337366.62 | 1196204.88 | 1336498.88 | 814064.56 | 2 | 843349.69 | 757896.94 | 542067.25 | 618958.19 | 694855.19 | 891421.56 | 1082398.62 | 561463.88 | 735788.5 | 947253.5 | 1006026.56 | 1134479.88 | 700055 | 2 | 0.461592842 | 1 | 0.662730713 | 2 | 0.63296731 | 3 |
| Q61881 | Mcm7 | DNA replication licensing factor MCM7 | 554.39 | 18.5 | 81211 | 11 | 10809012 | 8981728 | 7440702.5 | 17969792 | 20951174 | 4286497.5 | 6001006 | 7453437 | 7134752 | 6537113 | 4556683 | 8747308 | 6959521 | 8936952 | 5406111 | 3390765 | 5366053.5 | 5655188 | 3016416 | 6699942 | 0.474855054 | 1 | 0.523134014 | 2 | 0.364739018 | 0 |
| O70145 | Ncf2 | Neutrophil cytosol factor 2 | 242.78 | 10.29 | 59485 | 5 | 10453740 | 7764297 | 9135245 | 11035356 | 6368168 | 4601927 | 3864192 | 3500713 | 7188726 | 4653514 | 7071908 | 7112322.5 | 6191843 | 4788212.5 | 3495660.25 | 4296729.5 | 1761989 | 1753394.75 | 7163649.5 | 2913393.75 | 0.531965395 | 1 | 0.640348336 | 2 | 0.399696897 | 1 |
| P19253 | Rpl13a | Large ribosomal subunit protein uL13 | 490.61 | 39.9 | 23464 | 10 | 809942.06 | 2065789.75 | 622730.88 | 1155342.62 | 4644144.5 | 16123538 | 12403383 | 21329116 | 19748922 | 2927649.25 | 15885731 | 12933192 | 14460092 | 19538442 | 1195313.25 | 2755117.75 | 3852312.25 | 2144859 | 5852982.5 | 2797487 | 7.800924906 | 1 | 6.884611292 | 2 | 1.871676967 | 4 |

| | | | | | | | | | | | | | | | | | | | | | | | | | | | | | | | | |
|---|---|---|---|---|---|---|---|---|---|---|---|---|---|---|---|---|---|---|---|---|---|---|---|---|---|---|---|---|---|---|---|---|
| Q04753 | Clns1a | Methylosome subunit pICln | 52.84 | 5.51 | 26092 | 1 | 834791.12 | 1183008.38 | 637328.44 | 1173457.62 | 988731.31 | 2 | 628352.81 | 2 | 557902.5 | 916508.94 | 517390.38 | 2 | 898115.69 | 484148.34 | 2 | 2 | 2 | 762540.38 | 2 | 2 | 0.436501959 | 2 | 0.394339517 | 2 | 0.158293175 | 1 |
| P04550 | Ptms | Parathymosin | 174.3 | 22.55 | 11559 | 3 | 7954525.5 | 7452682 | 5778362.5 | 10666569 | 4848247 | 3753045 | 4599719.5 | 5010921.5 | 5464039 | 3488119.75 | 366153.81 | 4823141 | 6348166.5 | 4566379.5 | 1966014.5 | 18528574 | 14285151 | 5459394.5 | 27349028 | 11397479 | 0.60805477 | 2 | 0.492361451 | 2 | 2.098605353 | 4 |
| P08030 | Aprt | Adenine phosphoribosyltransferase | 798.36 | 78.89 | 19724 | 13 | 2989605.75 | 5781460 | 7919466.5 | 9042016 | 9310141 | 13760296 | 12926512 | 12873876 | 7951732.5 | 11989038 | 11692195 | 10646377 | 12062066 | 8760499 | 25475784 | 9417290 | 9771110 | 12319301 | 7725669.5 | 10235680 | 1.697970555 | 2 | 1.958665915 | 2 | 1.411679627 | 3 |
| Q8BPX9 | Slc15a3 | Solute carrier family 15 member 3 | 388.9 | 14.19 | 64051 | 6 | 6264685.5 | 5042557.5 | 4089030.25 | 3135996 | 3041179.75 | 9781516 | 8555917 | 11783861 | 8614352 | 4212863 | 2144710.25 | 650049.25 | 1938436.38 | 2131668.75 | 3460962 | 4387438 | 5841909 | 7474697.5 | 5858394 | 3776445.5 | 1.99080402 | 2 | 0.478635875 | 2 | 1.26724679 | 7 |
| Q8BK72 | Mrps27 | Small ribosomal subunit protein mS27 | 76.15 | 5.54 | 47779 | 2 | 2197450.5 | 3423821.75 | 2108926.25 | 1879147 | 1451478.38 | 2 | 163035.25 | 951940.31 | 2 | 2172462.25 | 2 | 1109180.25 | 1561065.38 | 1461012.38 | 431436.41 | 2 | 1772355.88 | 2822044 | 2720152.5 | 2 | 0.297214913 | 3 | 0.412509635 | 2 | 0.661303033 | 9 |
| Q7TNS2 | Micos10 | MICOS complex subunit Mic10 | 39.21 | 9.21 | 8567 | 1 | 3443558.5 | 2122458.75 | 1942349.25 | 2219467.5 | 1415675.75 | 2 | 1602155.38 | 2 | 1478580.75 | 1664193 | 1379795.62 | 1465682 | 1327136.25 | 1273000.5 | 1483594.62 | 1959406 | 2181642 | 2766175.5 | 2067301.38 | 2642807.5 | 0.425802394 | 3 | 0.621815671 | 2 | 1.042520053 | 10 |
| O35864 | Cops5 | COP9 signalosome complex subunit 5 | 122.66 | 8.68 | 37549 | 3 | 4030863.25 | 4898409.5 | 3202776.5 | 2447410.5 | 1245726.75 | 1240920.38 | 1060082.12 | 2 | 1879803.5 | 2950018.25 | 1701907.25 | 1593317.5 | 1201549.38 | 978195.88 | 1175422.88 | 1931377.88 | 2 | 926072.56 | 340138.69 | 864711.12 | 0.450599824 | 3 | 0.420241044 | 2 | 0.256698539 | 1 |
| Q8CGC6 | Rbm28 | RNA-binding protein 28 | 424.83 | 12.4 | 84214 | 8 | 1164450.75 | 1656422.25 | 1028358.88 | 2068719.12 | 1294689.12 | 541181.06 | 780168.44 | 195353.33 | 1373370.5 | 403385.22 | 2 | 778303.12 | 1182585.5 | 2 | 2 | 1510568.75 | 1194130.12 | 2 | 1362446.75 | 1212964.88 | 0.456623164 | 3 | 0.271869189 | 2 | 0.732063768 | 10 |
| Q3TIU4 | Pde12 | 2',5'-phosphodiesterase 12 | 109.48 | 4.28 | 67514 | 2 | 832866.38 | 1793029.12 | 1431433.62 | 1736068 | 1591067 | 1244856.62 | 870585.38 | 2 | 1284037.5 | 480416.62 | 791495.88 | 2 | 1281273.38 | 1047451.25 | 2 | 1304058.5 | 2 | 1009676.31 | 196359.39 | 2 | 0.525413633 | 3 | 0.422539058 | 2 | 0.339916094 | 2 |
| Q66H50 | Far1 | Fatty acyl-CoA reductase 1 | 465.82 | 19.22 | 59268 | 9 | 7513762 | 3707635 | 6519636 | 5714426.5 | 3053391.5 | 2548047.5 | 1694603.5 | 4584293 | 2635502.75 | 3404838.5 | 516316.78 | 2 | 4560184.5 | 2672104.5 | 2263162 | 2 | 1210468.88 | 4050888 | 866485.19 | 1166498.38 | 0.560842311 | 3 | 0.377676489 | 2 | 0.2751663 | 2 |
| P17665 | Cox7c | Cytochrome c oxidase subunit 7C, mitochondrial | 93.59 | 50.79 | 7333 | 2 | 1013006.81 | 862747.12 | 1518389.25 | 1606879.88 | 953151.75 | 2 | 705430.94 | 855514.94 | 759492.12 | 1224121.75 | 2 | 453234.91 | 802563.44 | 680668.94 | 998831.81 | 34012.66 | 101052.56 | 947053.62 | 84409.65 | 95320.48 | 0.595306967 | 3 | 0.492982016 | 2 | 0.211926759 | 1 |
| Q3ULJ0 | Gpd1l | Glycerol-3-phosphate dehydrogenase 1-like protein | 94.81 | 6.55 | 38226 | 2 | 3365666 | 2290730 | 3489163 | 2528300.75 | 1783779 | 1078483.75 | 1731017 | 2416357 | 1146977.75 | 1832222.25 | 1902682 | 672196.75 | 1765201.38 | 1342646 | 2110936.75 | 2 | 1677502 | 1195664.38 | 2093463.25 | 1268181.88 | 0.6096952 | 3 | 0.579125582 | 2 | 0.463291787 | 1 |
| B2RZ27 | Sh3bgrl3 | SH3 domain-binding glutamic acid-rich-like protein 3 | 210.07 | 43.01 | 10477 | 4 | 3611524.5 | 8669733 | 3520014.25 | 5870936 | 7832764 | 10169950 | 8295098 | 10009271 | 9229872 | 5889823 | 23062958 | 18969366 | 10383930 | 10749836 | 7581504.5 | 13327269 | 6339918 | 5555921 | 9667982 | 6914198.5 | 1.477514175 | 3 | 2.397819429 | 2 | 1.41688963 | 7 |
| Q99P72 | Rtn4 | Reticulon-4 | 417.91 | 11.62 | 126613 | 7 | 3767054 | 7311594.5 | 2678970.5 | 10315450 | 17416918 | 14256043 | 11528687 | 21806970 | 19712886 | 11568924 | 21624424 | 15859654 | 23092026 | 26072818 | 13472121 | 22373326 | 25098900 | 23650134 | 35763274 | 21176258 | 1.90102518 | 3 | 2.413137488 | 2 | 3.086573442 | 0 |
| O88668 | Creg1 | Protein CREG1 | 147.04 | 9.55 | 24452 | 3 | 197486.19 | 497987.09 | 478435.28 | 679791 | 302760.25 | 927307.75 | 1772237.88 | 396012.91 | 1298007.38 | 848575.62 | 566873.31 | 941771.75 | 673787 | 1343447.88 | 2703111.75 | 2 | 1775302.75 | 1732937.25 | 132424.36 | 1092411.75 | 2.430901571 | 3 | 2.888526677 | 2 | 2.194837153 | 10 |
| P60710 | Actb | Actin, cytoplasmic 1 | 2631.91 | 91.73 | 41737 | 36 | 149365.02 | 538855.12 | 448502.16 | 2 | 326613.28 | 2 | 2 | 2 | 208345.31 | 2 | 349789.56 | 799870.38 | 860897.06 | 1983560.5 | 1520018.38 | 2 | 1384757.12 | 1591499.75 | 2 | 1315200.38 | 0.142382259 | 4 | 3.76819126 | 2 | 2.932652936 | 9 |
| B5DFC8 | Eif3c | Eukaryotic translation initiation factor 3 subunit C | 373.81 | 8.34 | 105435 | 8 | 7501966 | 4810217 | 7973480.5 | 7757198.5 | 5960281.5 | 1273142.88 | 6716745 | 3186139.25 | 5445673 | 5462474 | 3980322.25 | 5058765.5 | 3837151.5 | 4776934 | 5655003.5 | 3713854.5 | 2466684.5 | 5199586.5 | 3294042.5 | 4888435 | 0.649474485 | 4 | 0.685471234 | 2 | 0.575317485 | 2 |
| Q8BM55 | Tmem214 | Transmembrane protein 214 | 228.73 | 9.46 | 76430 | 5 | 1180080.38 | 1178703.5 | 961996.56 | 1850109 | 955033.38 | 942145.5 | 1810903.25 | 2 | 715090.75 | 780006.94 | 2 | 121616.54 | 98445.48 | 1103034.62 | 246000.84 | 2 | 208778.92 | 1235214.25 | 2 | 989848.62 | 0.693470774 | 4 | 0.256140916 | 2 | 0.397302719 | 6 |
| B0K025 | Ostc | Oligosaccharyltransferase complex subunit OSTC | 138.72 | 18.12 | 16815 | 3 | 510396.69 | 1112166.5 | 606601.94 | 1259073.75 | 2125414.25 | 1691312.62 | 1437856.5 | 2146398.75 | 2076412.38 | 1578513.38 | 1711155.75 | 1498033.75 | 3220349 | 3443229.5 | 3703091 | 1278403.38 | 2915076 | 4676802.5 | 2997446.5 | 3482259.25 | 1.590852413 | 4 | 2.418364421 | 2 | 2.734402585 | 1 |
| Q9JIW9 | Ralb | Ras-related protein Ral-B | 167.89 | 23.3 | 23349 | 4 | 1534514.12 | 1066021.62 | 2 | 576264.44 | 2782438.25 | 2041333.88 | 1907062.25 | 3881014.75 | 3191628.5 | 1321138.75 | 3114538 | 3713156.5 | 3050730.75 | 2465066.5 | 1824183.62 | 2465761 | 3002530 | 3466646.5 | 391160.66 | 2717969 | 2.071099207 | 4 | 2.377429731 | 2 | 2.021074213 | 6 |
| Q9WV55 | Vapa | Vesicle-associated membrane protein-associated protein A | 345.69 | 26.91 | 27855 | 7 | 4226192 | 9751889 | 882824.81 | 6501073 | 2270625.75 | 17022140 | 14835281 | 3925440.75 | 17807618 | 9709440 | 13513602 | 18000004 | 13389884 | 17112306 | 4484620.5 | 20264530 | 13686844 | 12181758 | 18005776 | 6909229 | 2.678499511 | 4 | 2.81392668 | 2 | 3.006360844 | 1 |
| Q8BXZ1 | Tmx3 | Protein disulfide-isomerase TMX3 | 158.26 | 8.77 | 51848 | 4 | 1551647.88 | 989534.62 | 658507.56 | 1091559.62 | 734674.31 | 2 | 576286.88 | 1108651.5 | 2 | 721465 | 2 | 2 | 929329.88 | 2 | 2 | 2 | 2 | 1215249.12 | 2 | 2 | 0.478799 | 5 | 0.184908861 | 2 | 0.241797752 | 4 |
| Q8BPU7 | Elmo1 | Engulfment and cell motility protein 1 | 86.86 | 2.75 | 83936 | 2 | 1261092.5 | 2677358 | 888784.38 | 2019837.5 | 1862457.88 | 569955 | 1050510.88 | 747948.94 | 568839.75 | 2504942 | 734539.94 | 2 | 2 | 762506.5 | 1646194.75 | 2 | 2294865.25 | 2115635.75 | 866796.88 | 1160407.62 | 0.624855349 | 5 | 0.360897212 | 2 | 0.739156683 | 9 |
| Q6ZQM8 | Ugt1a7 | UDP-glucuronosyltransferase 1A7 | 502.74 | 24.48 | 59758 | 9 | 10394196 | 10147638 | 8692001 | 10775481 | 8564512 | 4752067 | 4893916 | 1766270.62 | 8216869 | 12364601 | 4064935.5 | 4473645 | 6666035.5 | 7599017.5 | 9448881 | 4967041 | 7571622.5 | 8055529.5 | 5561700 | 6093440.5 | 0.658661772 | 5 | 0.663989556 | 2 | 0.663924068 | 0 |
| P14069 | S100a6 | Protein S100-A6 | 331.86 | 53.93 | 10051 | 6 | 164047586 | 84051890 | 130625074 | 93485706 | 82406186 | 102789850 | 81642994 | 14865421 | 76759890 | 93363050 | 76879578 | 66264026 | 77028434 | 58177130 | 93317338 | 84383602 | 116688154 | 151197442 | 87042130 | 129134050 | 0.666084121 | 5 | 0.670132506 | 2 | 1.024934234 | 11 |
| Q62093 | Srsf2 | Serine/arginine-rich splicing factor 2 | 194.88 | 17.65 | 25476 | 4 | 1355257.5 | 3860306.5 | 2152554.5 | 2700250 | 9573730 | 4509471 | 3982455 | 6409021 | 5594140 | 4456704.5 | 10386354 | 10697476 | 10291682 | 9767810 | 2748913.5 | 16567955 | 12237522 | 6788755.5 | 15063850 | 7679813.5 | 1.270322084 | 5 | 2.234600112 | 2 | 2.970043959 | 2 |
| O55143 | Atp2a2 | Sarcoplasmic/endoplasmic reticulum calcium ATPase 2 | 1427.38 | 30.65 | 114858 | 25 | 10826347 | 18011260 | 9000194 | 10509625 | 14556440 | 17229276 | 18068524 | 16416002 | 16608979 | 13576443 | 16757226 | 21833882 | 20523892 | 21679550 | 16254305 | 18096568 | 18989466 | 21219398 | 18596494 | 18617870 | 1.301974413 | 5 | 1.542812249 | 2 | 1.518504379 | 0 |
| Q8R1S4 | Mtss1 | Protein MTSS 1 | 317.33 | 8.96 | 82408 | 5 | 1058211.75 | 1119015.88 | 1308424.75 | 475508.38 | 2937052.25 | 2685461.75 | 1191398 | 2005855.5 | 5052270 | 1902276 | 4757602.5 | 4277759.5 | 2236723 | 2733115.75 | 3122340 | 3374362.75 | 3485084.25 | 3877317 | 81927.39 | 3497870 | 1.860954603 | 5 | 2.482895313 | 2 | 2.075401465 | 5 |
| Q9WV85 | Nme3 | Nucleoside diphosphate kinase 3 | 41.05 | 10.06 | 19099 | 1 | 2 | 726311.38 | 678940.88 | 2 | 824940.31 | 2 | 2 | 583023.88 | 2 | 2 | 1487507.75 | 772405.56 | 824459.88 | 883563.06 | 1122998.38 | 826387.12 | 912780.06 | 1063492.5 | 939747.44 | 1005704.38 | 0.261426229 | 6 | 2.282729109 | 2 | 2.129010314 | 0 |
| Q9DCS2 | Mettl26 | Methyltransferase-like 26 | 84.54 | 9.8 | 22687 | 2 | 2 | 515798.03 | 1218049.62 | 953995.69 | 309110.12 | 996580.62 | 2 | 2 | 2 | 2 | 1855916.62 | 2057016.25 | 1703877.25 | 1499359.75 | 860893.19 | 741331.5 | 702133.69 | 806424.62 | 885955.19 | 2 | 0.332533677 | 6 | 2.66172226 | 2 | 1.046344212 | 12.5 |
| Q8R0F6 | Ilkap | Integrin-linked kinase-associated serine/threonine phosphatase | 38.98 | 2.3 | 42774 | 1 | 1338126.12 | 1132639.38 | 1268474.88 | 2045567 | 534302.56 | 2 | 362474.97 | 1190728.5 | 479854.75 | 1423334.38 | 2 | 682640.62 | 2 | 292527.06 | 677218 | 1445874.62 | 1657035.62 | 1485867.62 | 521472.88 | 1588572.38 | 0.546974911 | 6 | 0.26149089 | 2 | 1.060089662 | 9 |
| P24063 | Itgal | Integrin alpha-L | 44.54 | 0.95 | 128314 | 1 | 453825.03 | 667891.94 | 405872.12 | 787520.19 | 884851.75 | 2 | 426510.38 | 2 | 768646.62 | 627547.31 | 469022.25 | 2 | 2 | 2 | 2 | 2 | 703817.06 | 739349.94 | 583936.31 | 599119.06 | 0.569603284 | 6 | 0.146573738 | 2 | 0.82070511 | 10 |
| O08997 | Atox1 | Copper transport protein ATOX1 | 245.32 | 67.65 | 7338 | 5 | 9355946 | 5643894 | 10550204 | 5045698 | 18982864 | 3591144.5 | 2980069.5 | 7162022 | 2933443.5 | 18742262 | 18239776 | 23627698 | 23063174 | 24252604 | 16685044 | 18242782 | 16819030 | 12633327 | 21913490 | 19017740 | 0.714198005 | 6 | 2.135362499 | 2 | 1.787592999 | 3 |
| Q9DC70 | Ndufs7 | NADH dehydrogenase [ubiquinone] iron-sulfur protein 7, mitoch | 150.43 | 10.27 | 24683 | 2 | 2689957.75 | 7029256 | 3418484.75 | 6247232 | 3656251.75 | 7954897 | 6373283 | 3911064.75 | 6434464 | 4593360.5 | 3294419 | 2769830.75 | 2286672.75 | 1853332.5 | 1060118.62 | 2335182 | 1106725.38 | 1030336.56 | 1192718.25 | 2 | 1.270206925 | 6 | 0.488880019 | 2 | 0.245862566 | 0 |
| Q5XJY5 | Arcn1 | Coatomer subunit delta | 531.35 | 19.37 | 57229 | 11 | 2477503 | 1850133.75 | 3494991.5 | 3059532 | 4569527 | 4095419 | 4870941 | 2931953.75 | 5621101 | 3059620.5 | 4136920.5 | 5730033.5 | 3820627 | 6494790 | 5040120.5 | 5253984 | 2856162.75 | 905628 | 4523129 | 1387585.38 | 1.331830946 | 6 | 1.632345458 | 2 | 0.966010306 | 11 |
| P40142 | Tkt | Transketolase | 1674.24 | 49.44 | 67630 | 29 | 39039022 | 78418546 | 33548210 | 33800890 | 89102226 | 84618066 | 88700098 | 55029882 | 110023210 | 53448130 | 163516018 | 182412226 | 126912786 | 138631762 | 77841306 | 187722562 | 143701730 | 129989162 | 151493458 | 135058242 | 1.430473397 | 6 | 2.516581656 | 2 | 2.730707802 | 0 |
| P27870 | Vav1 | Proto-oncogene vav | 327.13 | 6.86 | 98137 | 5 | 2030993.38 | 123203.7 | 1096799.38 | 483896.84 | 5240591.5 | 2564219.5 | 1273896 | 3815833 | 4057266 | 3686113.5 | 8375784 | 7800287 | 5108799 | 7267738.5 | 2426885 | 7590874 | 3485959.5 | 2086052.25 | 6409591 | 2696153.25 | 1.715487056 | 6 | 3.451567708 | 2 | 2.481050383 | 3 |
| Q9CR51 | Atp6v1g1 | V-type proton ATPase subunit G 1 | 295.68 | 34.75 | 13724 | 4 | 2 | 542929.56 | 2 | 995452.44 | 1445024.25 | 2 | 834100.56 | 2136486.75 | 1468424 | 1448848.75 | 2383006.75 | 2891978.5 | 1653574.12 | 1529986.25 | 713232.25 | 2675486 | 1306616.5 | 1342587.12 | 2221189 | 2 | 1.973534166 | 6 | 3.07425969 | 2 | 2.529280249 | 6 |
| Q91ZR2 | Snx18 | Sorting nexin-18 | 120.91 | 4.72 | 67904 | 2 | 2 | 2 | 2 | 2 | 1253223.88 | 2 | 615557.31 | 214317.56 | 1445289 | 721114.69 | 1416148.75 | 931362.19 | 1515051.75 | 1376309.5 | 864104 | 2277401 | 1845058.12 | 2413709.5 | 1883922.62 | 2676502.5 | 2.390842914 | 6 | 4.86979009 | 2 | 8.854381952 | 0 |
| P45377 | Akr1b8 | Aldose reductase-related protein 2 | 599.28 | 32.28 | 36121 | 11 | 1129529 | 3535255.25 | 477839.09 | 1352539.5 | 1671802.38 | 3456705.5 | 3381970.75 | 4171152.25 | 8702617 | 1112073.75 | 13093079 | 14349886 | 3208626.75 | 6125177 | 1847781.5 | 13690011 | 3773392.5 | 3245477.5 | 5282296 | 4849171 | 2.549847916 | 6 | 4.729363871 | 2 | 3.776231093 | 1 |
| Q921N6 | Ddx27 | Probable ATP-dependent RNA helicase DDX27 | 129.44 | 3.82 | 85939 | 3 | 1476115 | 2 | 675239.44 | 480835.78 | 502128.66 | 821956.62 | 4400984 | 1287927.62 | 123141.02 | 6019432 | 671347.44 | 1509798.62 | 1866166 | 1671815.88 | 5873804 | 2 | 1484265 | 1916440.12 | 2 | 2471833.5 | 4.037059939 | 6 | 3.698706158 | 2 | 1.873625211 | 9 |
| P85108 | Tubb2a | Tubulin beta-2A chain | 1741.05 | 59.1 | 49907 | 26 | 2150158.5 | 2015168.88 | 1906833.62 | 1018095.44 | 456263.06 | 1063951.12 | 949314.06 | 1505141.88 | 952916.12 | 553944.12 | 2 | 742914.81 | 2 | 1004586.06 | 395686.22 | 861331.81 | 2 | 2 | 860581 | 2 | 0.665905296 | 7 | 0.28399729 | 2 | 0.228173903 | 2 |
| Q9QXS1 | Plec | Plectin | 8475.192 | 35.11 | 534188 | 158 | 153196322 | 204164002 | 174188370 | 155112034 | 181612210 | 172335250 | 185923570 | 200290306 | 184138690 | 186317890 | 305333026 | 306802178 | 221547954 | 235857154 | 181503538 | 277993634 | 201669618 | 196836050 | 215754002 | 184499410 | 1.069946632 | 7 | 1.440841693 | 2 | 1.240108573 | 3 |
| O35114 | Scarb2 | Lysosome membrane protein 2 | 72.38 | 4.18 | 54044 | 2 | 697377.25 | 435945.12 | 1416234.62 | 1143811.12 | 1404471.75 | 2124690 | 2 | 1562534.12 | 1700615.25 | 1205912.12 | 2 | 658694.75 | 2 | 2 | 518552.69 | 2 | 2 | 2 | 2 | 2 | 1.293440687 | 7 | 0.230931821 | 2 | 1.96162E-06 | 0 |
| Q69ZN7 | Myof | Myoferlin | 581.43 | 6.69 | 233324 | 11 | 5031534 | 4741635 | 2464479.75 | 1675938.75 | 1839702.75 | 4447420 | 4947511 | 7072817 | 3494810 | 4579746.5 | 9010014 | 9206379 | 6817130 | 9112358 | 2766071.5 | 12003277 | 11177555 | 8210976 | 11820012 | 8655194 | 1.557916099 | 7 | 2.343126541 | 2 | 3.292455936 | 0 |
| Q9CR98 | Fam136a | Protein FAM136A | 198.25 | 25.36 | 15674 | 4 | 5526609.5 | 13855725 | 4868453 | 5712001 | 3245322.5 | 14823904 | 13813124 | 4842778 | 11233259 | 7938137 | 13952224 | 15466217 | 10882027 | 14375035 | 7422468 | 10991867 | 15795834 | 13205040 | 13374438 | 12085440 | 1.585492231 | 7 | 1.869963968 | 2 | 1.970982902 | 4 |
| Q9JLB0 | Pals2 | Protein PALS2 | 365.59 | 15.37 | 62631 | 7 | 2 | 751522.5 | 1003583.88 | 2 | 1091163.5 | 2 | 372507.72 | 656551.31 | 2 | 514773.5 | 1599716 | 794558.19 | 1778416 | 2352290.5 | 1738443.5 | 83466.21 | 900074.81 | 1012600.62 | 234731.72 | 700865.12 | 0.542406176 | 8 | 2.903242814 | 2 | 1.030026836 | 12 |
| Q8BK67 | Rcc2 | Protein RCC2 | 656.84 | 27.88 | 55983 | 15 | 18679902 | 12644512 | 29987248 | 16064125 | 30034124 | 13618611 | 13345007 | 16746831 | 19110470 | 13341436 | 37597250 | 40215586 | 32937096 | 32199360 | 26459280 | 39448246 | 35778902 | 27829702 | 29880774 | 27557246 | 0.70908126 | 8 | 1.577215458 | 2 | 1.494227753 | 6 |
| Q9CWZ7 | Napg | Gamma-soluble NSF attachment protein | 92.35 | 4.49 | 34732 | 2 | 2662191 | 1130505.62 | 2707062.25 | 2567666 | 2077869.75 | 889388.75 | 732696 | 935287.88 | 2573530.25 | 2897854.75 | 161494.03 | 227488.69 | 1818851.38 | 1402031.38 | 910754.56 | 2 | 472307.56 | 268797.47 | 249535.89 | 470370.53 | 0.720371951 | 8 | 0.405607944 | 2 | 0.131087916 | 0 |
| P53994 | Rab2a | Ras-related protein Rab-2A | 464.87 | 40.57 | 23548 | 7 | 8909061 | 4740250 | 8186096 | 5996357.5 | 4804821 | 4567540 | 7105874 | 6010226.5 | 5734635 | 3378256.5 | 4468920 | 3393213.25 | 5103786 | 4168313.75 | 4240518.5 | 1577990.88 | 5944733 | 5854949.5 | 1125187.88 | 4062867.5 | 0.821058073 | 8 | 0.654932223 | 2 | 0.568862474 | 4 |
| O55222 | Ilk | Scaffold protein ILK | 72.09 | 4.2 | 51373 | 2 | 1704309.75 | 3210618.75 | 1674992.75 | 2475245 | 2593341 | 2130170 | 2098161.75 | 1719687.12 | 2285113 | 1508791.5 | 1634618.75 | 610386.81 | 1484026.25 | 2035228.25 | 1130897.38 | 680246.62 | 1417336.75 | 1635344.25 | 727841.44 | 855926.44 | 0.835606408 | 8 | 0.591427126 | 2 | 0.456035699 | 0 |
| Q9EQP2 | Ehd4 | EH domain-containing protein 4 | 397.99 | 15.53 | 61481 | 8 | 3857906.5 | 4967847 | 5644242 | 3550824.5 | 10134876 | 4227047 | 7004378.5 | 4080913.5 | 7819204.5 | 11722443 | 943233.81 | 2586093 | 2960140.75 | 4725046 | 3202818.5 | 894422.75 | 1547067.75 | 2744305.75 | 544961.44 | 2734872.75 | 1.237901791 | 8 | 0.512057385 | 2 | 0.300672036 | 0 |
| Q8BFR5 | Tufm | Elongation factor Tu, mitochondrial | 409.73 | 19.91 | 49508 | 8 | 2413480.5 | 6973144.5 | 4475464.5 | 8730422 | 11447985 | 7355173.5 | 8178677 | 10500777 | 11571655 | 4893879.5 | 13151778 | 13015023 | 12089794 | 12788209 | 7456243.5 | 13313812 | 12485534 | 8009733.5 | 9309710 | 10670429 | 1.248517688 | 8 | 1.718572098 | 2 | 1.580153759 | 4 |
| P62827 | Ran | GTP-binding nuclear protein Ran | 535.95 | 42.59 | 24423 | 9 | 5505285 | 7219234.5 | 5005873 | 16206653 | 25380134 | 13501386 | 14548178 | 21171520 | 23029562 | 11689674 | 28474954 | 29486334 | 26286854 | 19875018 | 18131204 | 22674796 | 16909250 | 13136974 | 14219792 | 11499051 | 1.415109766 | 8 | 2.061027935 | 2 | 1.322380188 | 8 |
| P63158 | Hmgb1 | High mobility group protein B1 | 183.08 | 23.26 | 24894 | 4 | 3850828.5 | 5465282 | 2869491.25 | 9230905 | 12269052 | 1990566.62 | 1902141.5 | 5558779 | 5795475.5 | 6177003.5 | 2400657.5 | 3852208.5 | 2761465 | 2077118.62 | 2465305.25 | 9592140 | 4183629.5 | 3578681.5 | 8142885.5 | 5488916 | 0.635998538 | 9 | 0.402450052 | 2 | 0.919867553 | 12 |
| P12970 | Rpl7a | Large ribosomal subunit protein eL8 | 522.58 | 37.59 | 29977 | 10 | 4794227.5 | 839961.19 | 2765724.75 | 4155681.25 | 8965753 | 1321921.5 | 2354350.5 | 605859 | 5350919 | 4632808.5 | 16169333 | 11910126 | 14721663 | 8898676 | 8638001 | 8710096 | 7644223 | 3051462.5 | 8039036 | 7078602.5 | 0.662870128 | 9 | 2.803625492 | 2 | 1.604147682 | 7 |
| P49300 | Clec10a | C-type lectin domain family 10 member A | 308.24 | 23.36 | 34596 | 6 | 5976716.5 | 1849346.75 | 7151406 | 3444021.75 | 3082026.5 | 4734295.5 | 1419452 | 812038.25 | 4667197 | 4163329.25 | 7551285 | 10072427 | 9855357 | 7629128 | 5815012.5 | 9086895 | 7537267 | 7112551.5 | 7413034 | 6110689 | 0.734592003 | 9 | 1.903093738 | 2 | 1.732760071 | 2 |
| B0BN93 | Psmd13 | 26S proteasome non-ATPase regulatory subunit 13 | 224.85 | 12.23 | 42817 | 5 | 5409428 | 7316887.5 | 11637260 | 4365922 | 8315481 | 4195695 | 5528016.5 | 7863446.5 | 1578388.62 | 8589324 | 3378196.25 | 3400037.75 | 4855211.5 | 2702906 | 4644788.5 | 3873482.25 | 3555004.75 | 4765436.5 | 4369421 | 4121422.5 | 0.749220859 | 9 | 0.512380916 | 2 | 0.558368984 | 2 |
| P62830 | Rpl23 | Large ribosomal subunit protein uL14 | 314.41 | 40.71 | 14865 | 6 | 2378503 | 5021202 | 1291690.62 | 3306389.75 | 5616888 | 3103561.25 | 2444283.5 | 1284367.62 | 4875817 | 2503249 | 13527060 | 12585783 | 8247035 | 8426786 | 3307772.75 | 10219581 | 4829017 | 4047960.25 | 7638212 | 4905355.5 | 0.806786369 | 9 | 2.616820408 | 2 | 1.796236869 | 6 |
| P35293 | Rab18 | Ras-related protein Rab-18 | 142.37 | 12.14 | 23035 | 3 | 1982408.88 | 3351281 | 1676044.38 | 3205183.5 | 2742133 | 2368614.25 | 2726549.5 | 3050193.5 | 2495614 | 580883.75 | 1756530.5 | 1128811.88 | 817412.88 | 1914667.12 | 1040340.56 | 1856058.5 | 1291670.25 | 2253934 | 1036195.25 | 1054943.12 | 0.866080963 | 9 | 0.513833207 | 2 | 0.578279831 | 3 |
| P50516 | Atp6v1a | V-type proton ATPase catalytic subunit A | 1285.75 | 49.27 | 68326 | 21 | 40417418 | 63318370 | 55133682 | 46317678 | 52444506 | 52934786 | 55546662 | 66028462 | 47203498 | 51957766 | 45255078 | 41091898 | 23128326 | 37875298 | 25712602 | 43724562 | 40579890 | 26422168 | 40831694 | 47895014 | 1.062257567 | 9 | 0.671746656 | 2 | 0.77418021 | 5 |
| P40124 | Cap1 | Adenylyl cyclase-associated protein 1 | 1697.28 | 55.06 | 51565 | 23 | 63808562 | 74907690 | 60009618 | 94760506 | 69672642 | 87411826 | 85057282 | 104806162 | 65613878 | 60737974 | 104263618 | 103568586 | 91174018 | 84373210 | 103075074 | 82604674 | 94583426 | 81479138 | 84232954 | 87148554 | 1.111433565 | 9 | 1.339508265 | 2 | 1.184188536 | 5 |
| Q8BMK4 | Ckap4 | Cytoskeleton-associated protein 4 | 897.48 | 30.26 | 63692 | 14 | 2045079.12 | 1276170.88 | 2805130.25 | 4041542.75 | 4977199 | 183101.31 | 3188411.25 | 4098052 | 4076896.75 | 7389311.5 | 391244.62 | 1179653.5 | 1516944 | 979984.5 | 1517138.38 | 2 | 2 | 1215795.38 | 2 | 753378.94 | 1.250288562 | 9 | 0.368763289 | 2 | 0.130020763 | 0 |
| O08573 | Lgals9 | Galectin-9 | 54.64 | 2.27 | 40036 | 1 | 1424258.88 | 3599328 | 744543.56 | 5550667.5 | 6586659 | 5355388 | 4426052.5 | 7327019 | 3947723.25 | 2198478.25 | 2 | 350567.38 | 746666.88 | 2 | 1163614.25 | 1121253.5 | 2 | 1235750.12 | 2 | 843971.19 | 1.29874714 | 9 | 0.126266116 | 2 | 0.178771132 | 3 |
| Q99J99 | Mpst | 3-mercaptopyruvate sulfurtransferase | 210.24 | 8.42 | 33097 | 3 | 640321.12 | 1086336 | 2 | 2 | 438552.09 | 161688.08 | 154747.05 | 129216.96 | 1296097.38 | 1546859 | 2398695.25 | 2829877.75 | 2307712.25 | 1839291.75 | 473078.47 | 1950758.75 | 297180.53 | 2 | 243593.41 | 830080.38 | 1.51883817 | 9 | 4.548584603 | 2 | 1.534082212 | 11 |
| P59325 | Eif5 | Eukaryotic translation initiation factor 5 | 161.56 | 8.39 | 48968 | 4 | 2 | 995849.88 | 1382684.5 | 2 | 1492343.38 | 1391790 | 1202094.5 | 1308877.75 | 877351 | 1482653.25 | 1827858.25 | 1831967.5 | 3076929 | 1890178.75 | 1359873.5 | 1275414.88 | 1222210.62 | 1034740.81 | 1568585.5 | 924941.81 | 1.617917283 | 9 | 2.579982448 | 2 | 1.556723763 | 9 |
| Q3TEA8 | Hp1bp3 | Heterochromatin protein 1-binding protein 3 | 695.71 | 26.9 | 60867 | 14 | 7017691 | 8743677 | 4710205 | 8029991 | 12240141 | 5102715 | 8237344.5 | 3382355.5 | 9300695 | 7529281 | 1809590.75 | 5209964.5 | 4447484 | 3023248 | 5152734.5 | 3807166 | 2187949.75 | 1009430.62 | 4658403 | 1720478.5 | 0.823539197 | 10 | 0.482135486 | 2 | 0.328494546 | 0 |
| P07901 | Hsp90aa1 | Heat shock protein HSP 90-alpha | 1417.43 | 33.56 | 84788 | 26 | 23226138 | 22802902 | 31683052 | 19700340 | 27727098 | 23346550 | 21657966 | 24514866 | 20062826 | 22863532 | 38366622 | 39885490 | 42053810 | 40173150 | 27697450 | 26814400 | 28390496 | 29914310 | 24680134 | 19510208 | 0.898562908 | 10 | 1.503733648 | 2 | 1.033322948 | 11 |
| Q5I0G4 | Gars1 | Glycine--tRNA ligase | 1025.97 | 29.81 | 81793 | 18 | 11906081 | 7679214 | 18289832 | 16285279 | 18419070 | 10560660 | 13317587 | 17404582 | 14781104 | 14040758 | 16726976 | 18604314 | 20758266 | 19599232 | 32421080 | 22169148 | 18116376 | 18279184 | 18577508 | 15983027 | 0.965902413 | 10 | 1.489537731 | 2 | 1.283079572 | 7 |
| O35226 | Psmd4 | 26S proteasome non-ATPase regulatory subunit 4 | 93.04 | 5.32 | 40704 | 2 | 2779402 | 3644293 | 2304349.5 | 2659288 | 3656623 | 4395527.5 | 2152778.5 | 1369992 | 3663422.5 | 3939181 | 4163676 | 4227300 | 5405960 | 5261857 | 3322323.75 | 3689395.5 | 5902679 | 4678242 | 7824437 | 5675494.5 | 1.031703497 | 10 | 1.4877149 | 2 | 1.845940584 | 0 |
| P60766 | Cdc42 | Cell division control protein 42 homolog | 397.38 | 52.88 | 21259 | 7 | 3311589 | 9642758 | 2980467 | 9994476 | 6508458.5 | 4819906.5 | 9451364 | 4506138 | 11199434 | 7563479 | 7283191 | 12266770 | 11891869 | 12841441 | 12267923 | 11389411 | 14098203 | 6780438.5 | 4888620 | 11313770 | 1.15730355 | 10 | 1.743376055 | 2 | 1.494260383 | 5 |
| Q6IN36 | Wipf1 | WAS/WASL-interacting protein family member 1 | 41.01 | 2.26 | 49751 | 1 | 1370946 | 2250732.5 | 700481.12 | 2038972.5 | 1951090.88 | 2850400.75 | 1944816.75 | 2715911.75 | 1820580.75 | 1157167.62 | 2974309 | 2969445.25 | 2151387 | 2334982.25 | 2234343.75 | 2092726.75 | 2674210.5 | 1970919.62 | 2707127.5 | 2351137.75 | 1.261861913 | 10 | 1.523595704 | 2 | 1.41912965 | 3 |
| P16858 | Gapdh | Glyceraldehyde-3-phosphate dehydrogenase | 1525.11 | 74.17 | 35810 | 23 | 1654572.88 | 3804337.5 | 48558.84 | 4208651.5 | 686905.19 | 3212006.75 | 2552205.75 | 304902.38 | 4247690 | 3352505 | 4327892 | 5021248 | 4530653.5 | 5400100 | 2362159 | 5685754 | 6131418 | 3387269.5 | 1934048 | 5105123 | 1.313974415 | 10 | 2.080361299 | 2 | 2.138186783 | 4 |
| P28650 | Adss1 | Adenylosuccinate synthetase isozyme 1 | 57.1 | 3.06 | 50254 | 1 | 2 | 129742.62 | 2 | 2 | 696773.38 | 2 | 632175.25 | 98938.09 | 2 | 975611.75 | 244390.44 | 1039360.19 | 730880.75 | 703297.56 | 565399.81 | 339909.19 | 933117 | 298355.16 | 2 | 263731.34 | 2.064953008 | 10 | 3.97246383 | 2 | 2.220285352 | 6.5 |
| P63259 | Actg1 | Actin, cytoplasmic 2 | 2642.15 | 91.73 | 41793 | 36 | 131565.94 | 446769.88 | 449241.94 | 2 | 118483.15 | 339154.06 | 2 | 503080.47 | 345015 | 407441.69 | 206246.73 | 653686.12 | 738996.31 | 2158185.5 | 1957427.38 | 2 | 1238970.88 | 1530309.5 | 2 | 715256 | 1.391453476 | 10.5 | 4.986237658 | 2 | 3.040444246 | 9 |
| P50544 | Acadvl | Very long-chain specific acyl-CoA dehydrogenase, mitochondria | 284.45 | 14.02 | 70875 | 6 | 2 | 70275.31 | 79263.61 | 59675.61 | 810189.5 | 2 | 136818.83 | 63004.89 | 628605.94 | 797179.38 | 1178319.12 | 1252342.5 | 619003.94 | 3071794.75 | 583562.06 | 1103071.62 | 176685.84 | 558532.25 | 886448.75 | 120626 | 1.594664925 | 10.5 | 6.577381507 | 2 | 2.79119838 | 3 |
| P61589 | Rhoa | Transforming protein RhoA | 297.15 | 31.09 | 21782 | 6 | 2592223 | 4862603.5 | 3271199.5 | 2471043.5 | 5148231 | 3056111.5 | 2792456.5 | 3909784 | 2639045.5 | 3003482.5 | 6349288 | 7087085 | 6154551.5 | 6617157.5 | 4288357.5 | 3418711.25 | 5056042 | 4792141 | 5320417.5 | 3987855.25 | 0.839500013 | 11 | 1.662357043 | 2 | 1.230569486 | 7 |
| P80313 | Cct7 | T-complex protein 1 subunit eta | 1455.54 | 54.41 | 59652 | 26 | 37496190 | 48713858 | 39065046 | 51724586 | 55765450 | 43689794 | 47472762 | 50487346 | 41520718 | 39513906 | 31579980 | 37564686 | 36647198 | 38271866 | 27922018 | 35419078 | 26713512 | 34341098 | 34661314 | 41800722 | 0.95669195 | 11 | 0.738881069 | 2 | 0.742962333 | 2 |
| P54311 | Gnb1 | Guanine nucleotide-binding protein G(I)/G(S)/G(T) subunit beta- | 382.11 | 20.88 | 37377 | 7 | 76517.09 | 1188626.38 | 1115604.88 | 2 | 162681.81 | 990081.88 | 231177.19 | 222230.75 | 126882.41 | 889543.31 | 180075.75 | 2276775.25 | 2310264.25 | 2272620.25 | 2285550 | 2901582 | 2648589.25 | 171587.8 | 4178300 | 1190268.12 | 0.967163811 | 11 | 3.666418018 | 2 | 4.360378604 | 2 |
| Q9CRB9 | Chchd3 | MICOS complex subunit Mic19 | 145.24 | 7.49 | 26335 | 2 | 757008.56 | 2 | 2 | 2 | 2469580.5 | 529840.06 | 2 | 2 | 821619.19 | 1906321.88 | 4496131 | 5620043 | 2415838.75 | 3734703 | 1129131.38 | 5602538 | 2721476.25 | 1046669.44 | 3518311.25 | 1537564.12 | 1.009666559 | 11 | 5.391394584 | 2 | 4.471140255 | 2 |
| P14206 | Rpsa | Small ribosomal subunit protein uS2 | 741.75 | 50.85 | 32838 | 12 | 16074994 | 13656222 | 18240308 | 18927790 | 34380682 | 14847563 | 14155545 | 26433574 | 28345000 | 19097098 | 72366498 | 69483690 | 38055890 | 52467102 | 18301042 | 65226098 | 24961334 | 15190722 | 36100678 | 15601563 | 1.015785783 | 11 | 2.475061531 | 2 | 1.550951829 | 9 |
| Q9CQX2 | Cyb5b | Cytochrome b5 type B | 192.13 | 15.75 | 16318 | 2 | 915671.38 | 718192.81 | 1818190 | 585069.75 | 2019306 | 586687 | 2 | 1921275.25 | 1870178.75 | 2225749.75 | 3277430.5 | 1616061.88 | 2948501 | 2703744.5 | 3447611.5 | 2964692 | 2790975 | 4193625 | 4072544.5 | 3269958.75 | 1.09039365 | 11 | 2.310494717 | 2 | 2.855113561 | 0 |
| Q921F2 | Tardbp | TAR DNA-binding protein 43 | 413.56 | 17.39 | 44548 | 6 | 5051298 | 7201042 | 917203.19 | 8291341.5 | 2688956.5 | 6548520 | 6400967 | 2100354.5 | 7754439 | 5727020.5 | 14065834 | 13013706 | 9388418 | 11387324 | 6781083.5 | 12202236 | 6333363 | 9356212 | 9114005 | 8550262 | 1.1814281 | 11 | 2.26239026 | 2 | 1.886392446 | 2 |
| Q6PCT5 | Pqbp1 | Polyglutamine-binding protein 1 | 34.37 | 3.42 | 30529 | 1 | 682938.38 | 2 | 599305.81 | 2 | 2 | 634401.31 | 2 | 2 | 287258.22 | 2 | 1703172.12 | 959689.31 | 1314095.88 | 519011.09 | 1546977.62 | 2522625.75 | 1929260.38 | 2 | 2 | 4546424.5 | 0.718787595 | 11.5 | 4.712766718 | 2 | 7.017596644 | 7 |
| Q8BL66 | Eea1 | Early endosome antigen 1 | 129.17 | 2.2 | 160915 | 3 | 2 | 2 | 2 | 2 | 1178439.62 | 2 | 2 | 2 | 2 | 301107.91 | 2958061.5 | 3319228.5 | 1242859.5 | 639464.38 | 590381.06 | 2924144.5 | 928224.56 | 1139135.62 | 1824110.5 | 824188.31 | 0.255519129 | 12 | 7.425018127 | 2 | 6.482938537 | 3 |
| Q4V8K5 | Brox | BRO1 domain-containing protein BROX | 260.33 | 14.6 | 46192 | 4 | 2546721.25 | 3248494.5 | 798601.38 | 2204525.5 | 950077.12 | 1847293 | 1485935 | 2477972.5 | 2311213.75 | 997439.69 | 259257.86 | 2 | 412646.19 | 1433311.5 | 706209.88 | 2 | 3138416.25 | 2363103 | 964679.75 | 903114.94 | 0.935521261 | 12 | 0.288398274 | 2 | 0.755949798 | 10 |
| P10126 | Eef1a1 | Elongation factor 1-alpha 1 | 1553.77 | 47.62 | 50114 | 25 | 235127010 | 368272994 | 103546554 | 504683650 | 535634114 | 347666114 | 210210274 | 372848194 | 244765074 | 510729762 | 798043010 | 793966850 | 589089090 | 679891202 | 458987650 | 779988098 | 249989506 | 462731010 | 591355650 | 417772162 | 0.965062582 | 12 | 1.900100494 | 2 | 1.431859161 | 7 |
| Q8CBW3 | Abi1 | Abl interactor 1 | 121.21 | 5.61 | 52288 | 2 | 2910481.75 | 3416475.75 | 2762181 | 2067430.5 | 2404614 | 3419500.75 | 2889287.25 | 3350285 | 1423159.88 | 2294113.75 | 2216953.75 | 1897014.12 | 1907361.5 | 1904903.88 | 2364202.25 | 2585467.5 | 3149335.25 | 3517339.5 | 2543449.25 | 3275168.25 | 0.986370188 | 12 | 0.758815474 | 2 | 1.111316008 | 8 |
| Q921G7 | Etfdh | Electron transfer flavoprotein-ubiquinone oxidoreductase, mitoc | 291.02 | 11.85 | 68091 | 6 | 2509600.75 | 1083471 | 642800.88 | 952283.62 | 2 | 694799.88 | 135267.56 | 317937.66 | 1563220.5 | 2498943.5 | 2916740.25 | 2980252.75 | 1763176 | 3357184.5 | 1178410.25 | 2571413.75 | 1573560.12 | 1434587.62 | 180469.5 | 3557995 | 1.004242517 | 12 | 2.350692319 | 2 | 1.796018074 | 6 |
| P63081 | Atp6v0c | V-type proton ATPase 16 kDa proteolipid subunit c | 453.76 | 78.06 | 15808 | 8 | 75299250 | 77029386 | 67346314 | 65527334 | 51232546 | 69549162 | 53896914 | 85451954 | 72418826 | 62634422 | 34678934 | 28890156 | 61342130 | 53516830 | 46498514 | 25316982 | 48405414 | 89260498 | 71275066 | 69942754 | 1.022341468 | 12 | 0.668559091 | 2 | 0.904189123 | 11 |
| P97372 | Psme2 | Proteasome activator complex subunit 2 | 445.84 | 35.15 | 27057 | 8 | 6294491 | 6775690.5 | 11254707 | 12676632 | 16376726 | 9158677 | 9574677 | 10271082 | 8375536 | 18236732 | 3060721.5 | 5525238 | 5917710.5 | 5886359 | 7323923 | 5912933.5 | 5897028.5 | 7707918 | 7710094.5 | 3356980.5 | 1.041935763 | 12 | 0.519199371 | 2 | 0.572985383 | 4 |
| P24049 | Rpl17 | Large ribosomal subunit protein uL22 | 218.27 | 24.46 | 21397 | 5 | 4209144.5 | 4409993 | 2911246 | 5371977.5 | 4534420 | 4361944 | 3610781 | 7782272 | 8319668 | 425102.09 | 9014267 | 8900701 | 4902082 | 6392799 | 4662976 | 9602756 | 5680413.5 | 8723828 | 8492576 | 5753117.5 | 1.14288461 | 12 | 1.580126466 | 2 | 1.784441937 | 0 |
| P06800 | Ptprc | Receptor-type tyrosine-protein phosphatase C | 1081.34 | 17.79 | 144837 | 21 | 16766360 | 12470701 | 15171073 | 16895154 | 14657508 | 10843017 | 15091731 | 12440374 | 23773580 | 25905020 | 20285554 | 16500635 | 23092574 | 21097316 | 22074420 | 21058510 | 20783082 | 14591412 | 16320477 | 13995069 | 1.159199569 | 12 | 1.356627424 | 2 | 1.14201739 | 10 |
| Q8K310 | Matr3 | Matrin-3 | 426.48 | 14.3 | 94630 | 8 | 917155.94 | 129395.91 | 531004.94 | 512322.78 | 1152094.5 | 2 | 157048.72 | 490660.72 | 2254393.5 | 1327784.12 | 1211262.75 | 1810742.88 | 935437.31 | 1177049.88 | 995220.5 | 2422343.5 | 1566792.25 | 717496.69 | 1434027.62 | 592373.88 | 1.30472637 | 12 | 1.890734839 | 2 | 2.076831521 | 4 |

Supplementary table 4: List of modulated proteins in the 4x10µg/ml condition, selected by the Mann Whitney U test (p<0.05)

| accession number | gene_name | description | protein_set_score | coverage | MW | #peptides | Ctrl 1 | Ctrl 2 | Ctrl 3 | Ctrl 4 | Ctrl 5 | PBk14 20 recov 1 | PBk14 20 recov 2 | PBk14 20 recov 3 | PBk14 20 recov 4 | PBk14 20 recov 5 | PBk14 4x5 rep1 | PBk14 4x5 rep2 | PBk14 4x5 rep3 | PBk14 4x5 rep4 | PBk14 4x5 rep5 | PBk14 4x10 rep1 | PBk14 4x10 rep2 | PBk14 4x10 rep3 | PBk14 4x10 rep4 | PBk14 4x10 rep5 | ratio PBk14 20 recov/ctrl | U PBk14/20 recov | ratio_PBk14 4x5/ctrl | U_PBk14 4x5 | ratio Pbk 14 4x10 /ctrl | U Pbk 14 4x10 |
|---|---|---|---|---|---|---|---|---|---|---|---|---|---|---|---|---|---|---|---|---|---|---|---|---|---|---|---|---|---|---|---|
| P35433 | Ppat | Amidophosphoribosyltransferase | 37.89 | 1.35 | 57437 | 1 | 1384687.88 | 1062445.5 | 918004.81 | 862753.44 | 1043741.19 | 2 | 2 | 2 | 2 | 2 | 2 | 2 | 682479.81 | 586809.25 | 2 | 2 | 580160.31 | 2 | 2 | 2 | 1.89695E-06 | 0 | 0.240778351 | 0 | 0.110054765 | 0 |
| P49718 | Mcm5 | DNA replication licensing factor MCM5 | 454.31 | 16.21 | 82407 | 11 | 14334609 | 23994526 | 12389125 | 23668398 | 18101840 | 5167847 | 2130634 | 10794580 | 8285995.5 | 7981872 | 8040220 | 7604583 | 8059879 | 10139252 | 6163571.5 | 6093196 | 5124019 | 2573817.75 | 7862198.5 | 9596300 | 0.371515694 | 0 | 0.432567361 | 0 | 0.337874784 | 0 |
| P46061 | Rangap1 | Ran GTPase-activating protein 1 | 481.99 | 20.54 | 63531 | 10 | 9060379 | 6495795 | 7827281 | 7722898 | 6361215 | 1926580.75 | 2586222.5 | 2292656.25 | 2893515 | 4771926 | 2518570.5 | 1173641.75 | 1092664.38 | 2043450.25 | 1185948.38 | 1744415.88 | 2525018.75 | 178639.88 | 1270313.75 | 1076775.12 | 0.386224708 | 0 | 0.213898998 | 0 | 0.181361208 | 0 |
| B2RXS4 | Plxnb2 | Plexin-B2 | 500.43 | 6.03 | 206230 | 9 | 10473187 | 11143706 | 9682170 | 5885522.5 | 8422308 | 2396757 | 632654.44 | 5870254 | 5480084 | 5764672.5 | 3756564 | 3685656 | 4858579 | 3459083 | 3609338.75 | 3586604.25 | 5118156 | 3967404.75 | 3771416.75 | 2162288.75 | 0.441696866 | 0 | 0.42469941 | 0 | 0.407961803 | 0 |
| Q91YE6 | Ipo9 | Importin-9 | 292.36 | 8.07 | 116052 | 5 | 5661126.5 | 5772475 | 4431324 | 4939674.5 | 5068063.5 | 2326718 | 2362680.5 | 2742037.5 | 2651297.5 | 1563304.5 | 4272772.5 | 3232113.25 | 4295584.5 | 3404548.25 | 2630744.25 | 4011597 | 3580901.75 | 3850355.5 | 3650960.25 | 2961686.75 | 0.450129072 | 0 | 0.68936709 | 0 | 0.697860166 | 0 |
| Q9D906 | Atg7 | Ubiquitin-like modifier-activating enzyme ATG7 | 141.96 | 4.3 | 77520 | 3 | 4201814.5 | 3761543.75 | 4630190.5 | 3653197.25 | 3896952.5 | 93426.41 | 2989151.25 | 1120287 | 2885658.5 | 3649918.25 | 1759115.75 | 1743334.88 | 2780771.75 | 2936376.25 | 2306131 | 1619819.5 | 3279527.75 | 2674177 | 2116290 | 2440309.5 | 0.533091846 | 0 | 0.572175444 | 0 | 0.602179572 | 0 |
| P20060 | Hexb | Beta-hexosaminidase subunit beta | 353.4 | 17.72 | 61116 | 8 | 19787152 | 25215262 | 21984138 | 24970668 | 20180386 | 11218148 | 12726519 | 7381907 | 17365548 | 17356838 | 8912201 | 7231532.5 | 8537964 | 8499397 | 11826519 | 6433805 | 10468561 | 12581139 | 9981005 | 15225826 | 0.588999198 | 0 | 0.40136057 | 0 | 0.487707362 | 0 |
| Q8BG07 | Pld4 | 5'-3' exonuclease PLD4 | 943.83 | 38.57 | 56154 | 14 | 56814230 | 54365838 | 48080822 | 49538466 | 42085726 | 28968542 | 28506038 | 28419456 | 26219874 | 40881506 | 17161362 | 14213955 | 16546519 | 14629944 | 27383632 | 21365194 | 19312210 | 24940534 | 18028056 | 20254480 | 0.609822692 | 0 | 0.358472538 | 0 | 0.41413572 | 0 |
| P29416 | Hexa | Beta-hexosaminidase subunit alpha | 430.36 | 20.45 | 60613 | 9 | 11239542 | 9282127 | 9346251 | 12898934 | 8747906 | 3946593.75 | 5545161.5 | 6881814.5 | 7755726 | 7295286 | 2764089.5 | 2039271.25 | 792291.81 | 2729312.5 | 3822945.5 | 4363559.5 | 3434400 | 6897750 | 4884175 | 4994353.5 | 0.610011223 | 0 | 0.235814174 | 0 | 0.477032951 | 0 |
| Q8BFY9 | Tnpo1 | Transportin-1 | 297.62 | 5.01 | 102357 | 5 | 11038026 | 8185250.5 | 11575319 | 7989010 | 9471328 | 5731841 | 5967353 | 7195839 | 6718364 | 5640400 | 3683115.25 | 4458412.5 | 5869673 | 4904413.5 | 5360765 | 3546715.5 | 4544134 | 2163104 | 2871810.75 | 2622370 | 0.64762718 | 0 | 0.503044255 | 0 | 0.326325783 | 0 |
| Q9QZD8 | Slc25a10 | Mitochondrial dicarboxylate carrier | 680.98 | 36.93 | 31715 | 12 | 11981910 | 14999013 | 13635671 | 13621286 | 17404118 | 9275321 | 9050245 | 8613553 | 11941318 | 10718740 | 9006348 | 7019183.5 | 8229059.5 | 9334255 | 6532186.5 | 6385108 | 4662828.5 | 8772034 | 5495926 | 7336750 | 0.692319846 | 0 | 0.560021128 | 0 | 0.455775207 | 0 |
| P63017 | Hspa8 | Heat shock cognate 71 kDa protein | 3151.25 | 68.27 | 70871 | 47 | 421192898 | 431482370 | 373439234 | 421580098 | 450136930 | 284628066 | 257253202 | 317597506 | 297581762 | 362076450 | 358839650 | 359159010 | 313883746 | 310589186 | 372030082 | 269497954 | 320537218 | 317056450 | 216600722 | 301734146 | 0.724146322 | 0 | 0.817273289 | 0 | 0.679476149 | 0 |
| Q3UND0 | Skap2 | Src kinase-associated phosphoprotein 2 | 122.18 | 8.1 | 40712 | 3 | 3969949.25 | 3809952.75 | 3530172.5 | 5301210.5 | 3558785.5 | 3133936.75 | 2973437.5 | 3486063 | 2766528.5 | 2947538.5 | 1311216.75 | 1656630.25 | 1974965.38 | 1279712.75 | 1255237.88 | 2672843 | 1643412.12 | 2004220.75 | 2632103 | 2089896.12 | 0.758921703 | 0 | 0.370735591 | 0 | 0.547468339 | 0 |
| P57776 | Eef1d | Elongation factor 1-delta | 669.69 | 36.3 | 31293 | 10 | 55972342 | 56603278 | 49338410 | 54788842 | 53468642 | 42996234 | 41529478 | 47330046 | 34967330 | 43419870 | 29211538 | 30174042 | 34276694 | 32177534 | 27318584 | 26016820 | 27842364 | 25829278 | 29964362 | 31675226 | 0.778183291 | 0 | 0.566893192 | 0 | 0.523104927 | 0 |
| Q7TMB8 | Cyfip1 | Cytoplasmic FMR1-interacting protein 1 | 751.51 | 13.01 | 145241 | 15 | 6839314 | 7714240.5 | 8690557 | 7099803 | 8213055.5 | 11880539 | 9639829 | 16722741 | 12318325 | 9600170 | 16534412 | 13017145 | 15349741 | 14831595 | 13903786 | 11246515 | 11660331 | 16479392 | 11029072 | 10603626 | 1.560330182 | 0 | 1.909814983 | 0 | 1.582565642 | 0 |
| Q8VCN5 | Cth | Cystathionine gamma-lyase | 497.8 | 27.14 | 43567 | 10 | 9377385 | 11501992 | 20802036 | 13511600 | 17593928 | 23467510 | 23925028 | 22619538 | 21611462 | 23167656 | 25079238 | 26935692 | 23539544 | 22674670 | 26509684 | 26265474 | 32094046 | 23433328 | 23389266 | 24455162 | 1.577085016 | 0 | 1.713752856 | 0 | 1.781051301 | 0 |
| P20152 | Vim | Vimentin | 3751.201 | 80.69 | 53688 | 58 | 292444354 | 261473458 | 320387522 | 246949666 | 257421506 | 395601250 | 419176482 | 605743682 | 420862754 | 458444546 | 605814594 | 542833026 | 570076226 | 563079618 | 409487650 | 522059714 | 462530306 | 352727650 | 392223234 | 342554082 | 1.668142384 | 0 | 1.952083105 | 0 | 1.502959525 | 0 |
| Q9R0P3 | Esd | S-formylglutathione hydrolase | 944.79 | 56.38 | 31320 | 13 | 25940540 | 26521614 | 25612578 | 20582998 | 29878124 | 51622646 | 40259646 | 62494562 | 48655582 | 44654690 | 56858138 | 55370386 | 65995338 | 55665290 | 47097394 | 97382994 | 77140818 | 65375730 | 94525538 | 61382050 | 1.926988605 | 0 | 2.186055776 | 0 | 3.079351929 | 0 |
| Q8CGK3 | Lonp1 | Lon protease homolog, mitochondrial | 844.89 | 25.08 | 105843 | 17 | 6567353.5 | 4798552.5 | 5443349.5 | 4223956.5 | 7961036.5 | 11063631 | 10650989 | 9479042 | 14065289 | 13852630 | 11698115 | 9505348 | 12599022 | 13572474 | 9914388 | 13300126 | 13773298 | 14880314 | 16567199 | 12039036 | 2.038734717 | 0 | 1.975886597 | 0 | 2.433585164 | 0 |
| O35129 | Phb2 | Prohibitin-2 | 1197.8 | 63.55 | 33296 | 18 | 4204098 | 4630888.5 | 7077039.5 | 5518796.5 | 9586686 | 15832965 | 12552542 | 17125624 | 15482124 | 10304798 | 24869354 | 23461662 | 24523416 | 22303114 | 14611208 | 20986556 | 11144225 | 12650030 | 17765512 | 10430989 | 2.298638945 | 0 | 3.538928796 | 0 | 2.352778012 | 0 |
| P62320 | Snrpd3 | Small nuclear ribonucleoprotein Sm D3 | 135.47 | 32.54 | 13916 | 3 | 1915268.88 | 6486596.5 | 4338117 | 3730228.5 | 9494886 | 12693957 | 9783863 | 13306313 | 14228131 | 10034420 | 20549694 | 17782678 | 19691886 | 16158086 | 14915846 | 21357836 | 21218120 | 18592006 | 24062486 | 17204178 | 2.312592334 | 0 | 3.431459948 | 0 | 3.945089305 | 0 |
| Q8BVE3 | Atp6v1h | V-type proton ATPase subunit H | 311.71 | 16.36 | 55855 | 7 | 697950.5 | 592864.12 | 538543.75 | 1240671.38 | 717263.56 | 2976704.75 | 3007390.5 | 3972842 | 2819988 | 1650369.75 | 4564762 | 3849971 | 4224204.5 | 5540296 | 5296571.5 | 4642321 | 4408350 | 4816911 | 5232386 | 3798193 | 3.809394684 | 0 | 6.198570609 | 0 | 6.046049019 | 0 |
| Q6IMX7 | Hspbp1 | Hsp70-binding protein 1 | 163.31 | 12.61 | 39191 | 3 | 2276276 | 2767235 | 2667044.25 | 4319962 | 3298623.75 | 336641.5 | 1060189.5 | 602820.38 | 177643.69 | 2344810.5 | 2 | 1270804.12 | 1660076.38 | 1809124 | 2272670.75 | 2 | 955965.06 | 2 | 1231669.25 | 2074187.5 | 0.295000585 | 1 | 0.457473596 | 0 | 0.278021176 | 0 |
| Q62422 | Ostf1 | Osteoclast-stimulating factor 1 | 235.59 | 21.4 | 23783 | 4 | 2962240.5 | 2067645.25 | 2780713.25 | 1662546.75 | 3031474.5 | 213686.64 | 1157247.12 | 472047.72 | 790177.5 | 2053079.62 | 266553.31 | 2 | 39743.21 | 504152.06 | 1129318.38 | 1339886 | 494839.66 | 1297546 | 1015108 | 1461530.25 | 0.374760569 | 1 | 0.15512418 | 0 | 0.448547001 | 0 |
| Q91WD5 | Ndufs2 | NADH dehydrogenase [ubiquinone] iron-sulfur protein 2, mitochondrial | 369.62 | 15.33 | 52626 | 6 | 2812738.5 | 3707192.75 | 1914599.5 | 3511760 | 2510454.25 | 1251882.88 | 2417935 | 129536.37 | 1333782 | 1083877.75 | 93646.58 | 157165.88 | 1322492.12 | 1456137.88 | 355400.22 | 2 | 326249.25 | 234073.33 | 70574.06 | 645298.69 | 0.430042447 | 1 | 0.234135878 | 0 | 0.088276948 | 0 |
| Q8C147 | Dock8 | Dedicator of cytokinesis protein 8 | 91.29 | 1.1 | 238978 | 2 | 2653438.5 | 2026465.25 | 2961496 | 2843847.5 | 2720138 | 2585858.5 | 1289854.75 | 2 | 1773110.25 | 1349753.75 | 675902.75 | 605989.25 | 2 | 2 | 2003271.75 | 909702 | 2 | 1123261.5 | 1156242.38 | 1304775.38 | 0.52997918 | 1 | 0.248774851 | 0 | 0.340314438 | 0 |
| Q91YT0 | Ndufv1 | NADH dehydrogenase [ubiquinone] flavoprotein 1, mitochondrial | 147.55 | 10.13 | 50834 | 3 | 1921741.5 | 1833127.25 | 1783156.75 | 2086493.38 | 1671178.38 | 1571856.75 | 1208416.62 | 1748422.25 | 1242223 | 1251929.12 | 940153.38 | 681869.38 | 1187536.5 | 1028157.44 | 1293136.12 | 561737.31 | 766497.5 | 764194.56 | 796410.06 | 804999.81 | 0.755494456 | 1 | 0.551959974 | 0 | 0.397370863 | 0 |
| P26041 | Msn | Moesin | 2471.44 | 56.67 | 67767 | 41 | 36159634 | 47561142 | 33869290 | 64798106 | 59174970 | 66666686 | 60445490 | 70021162 | 100437402 | 92483298 | 171885314 | 194163906 | 118448938 | 131269890 | 138163378 | 143685234 | 127130450 | 126115922 | 122653026 | 123973450 | 1.614708414 | 1 | 3.121053236 | 0 | 2.664140219 | 0 |
| P84104 | Srsf3 | Serine/arginine-rich splicing factor 3 | 174.38 | 18.29 | 19330 | 4 | 2697017.25 | 5210512.5 | 3169201.25 | 4226097.5 | 6865514.5 | 9545095 | 11787885 | 12703122 | 10964365 | 6348119.5 | 14239307 | 13984641 | 12153264 | 14070471 | 10311008 | 15509445 | 13494506 | 12329943 | 14724140 | 12374145 | 2.316302418 | 1 | 2.921223792 | 0 | 3.086932524 | 0 |
| P13864 | Dnmt1 | DNA (cytosine-5)-methyltransferase 1 | 365.76 | 5.93 | 183189 | 8 | 794419.38 | 1027196.06 | 671121.75 | 1295460.12 | 1833908.38 | 2 | 2 | 2 | 876366 | 2 | 475003 | 2 | 2 | 2 | 2 | 2 | 2 | 2 | 2 | 2 | 0.155880029 | 2 | 0.084489874 | 0 | 1.77869E-06 | 0 |
| Q8VCH8 | Ubxn4 | UBX domain-containing protein 4 | 205.24 | 12.45 | 56472 | 4 | 23049706 | 63928290 | 16709360 | 36582354 | 26852198 | 21990278 | 19101856 | 2406963 | 10070016 | 8346575 | 5193491 | 3302100.5 | 4521407 | 4991237.5 | 2630675.75 | 6785781 | 4565369.5 | 3875161.75 | 5193261 | 3748322.75 | 0.370482175 | 2 | 0.123496147 | 0 | 0.144612375 | 0 |
| Q2YFS3 | Pilra | Paired immunoglobulin-like type 2 receptor alpha | 84.66 | 8.61 | 33064 | 2 | 2955074 | 1496703.12 | 2295763.25 | 1268255.88 | 1850229.38 | 783272.12 | 881346.38 | 1627209 | 818339.94 | 1027893.81 | 2 | 2 | 497421.94 | 894678.12 | 722047 | 2 | 776842.81 | 2 | 2 | 958563.19 | 0.520783286 | 2 | 0.214285989 | 0 | 0.175897779 | 0 |
| P62075 | Timm13 | Mitochondrial import inner membrane translocase subunit Tim13 | 90.99 | 14.74 | 10458 | 1 | 2951467 | 1980725.62 | 2606312 | 1496193 | 1611425.75 | 1285734.62 | 1170344.25 | 1657627 | 846881.69 | 1278633.38 | 1451130.12 | 802565.5 | 1413385.75 | 2 | 1437448.12 | 2 | 2 | 1006918.06 | 2 | 2 | 0.586055668 | 2 | 0.479473261 | 0 | 0.094581476 | 0 |
| Q91V41 | Rab14 | Ras-related protein Rab-14 | 891.23 | 76.28 | 23897 | 16 | 27776224 | 32426526 | 32192920 | 23327358 | 19961586 | 19227080 | 21799252 | 15683999 | 21269958 | 17139804 | 15968152 | 17232982 | 16460083 | 15416712 | 15020684 | 15331187 | 17008326 | 19564536 | 16123711 | 19719576 | 0.701038166 | 2 | 0.590329372 | 0 | 0.646700708 | 0 |
| Q8JZN5 | Acad9 | Complex I assembly factor ACAD9, mitochondrial | 375.79 | 15.04 | 68722 | 8 | 5865435 | 4934719 | 8300346 | 6313488.5 | 7132175 | 4912872 | 2912914 | 5494705.5 | 5289403 | 4681942.5 | 1002076.81 | 2025586.75 | 3740948 | 2678594.5 | 3827852.5 | 690338.5 | 1965555.75 | 2042634.25 | 2621769.5 | 1648101.88 | 0.71565538 | 2 | 0.407883976 | 0 | 0.275559357 | 0 |
| Q922F4 | Tubb6 | Tubulin beta-6 chain | 1168.36 | 45.19 | 50090 | 19 | 15889175 | 19201524 | 17351284 | 14801321 | 13201588 | 12685784 | 13338699 | 14150360 | 8979408 | 9314541 | 7642172.5 | 5925416 | 10781049 | 10459778 | 8820867 | 5648359.5 | 10516075 | 10643566 | 10261886 | 10610539 | 0.726817956 | 2 | 0.542349942 | 0 | 0.592709174 | 0 |
| Q9CZW5 | Tomm70 | Mitochondrial import receptor subunit TOM70 | 292.04 | 9.66 | 67590 | 6 | 12400570 | 12977864 | 11430952 | 11422214 | 16287770 | 8822925 | 10124398 | 12283153 | 8627559 | 10502097 | 5700361.5 | 6496184.5 | 5898677 | 5873157 | 7224938.5 | 7384014 | 6515596 | 7827071.5 | 8956943 | 10142102 | 0.780542835 | 2 | 0.48347215 | 0 | 0.632766974 | 0 |
| P42932 | Cct8 | T-complex protein 1 subunit theta | 1255.53 | 40.15 | 59555 | 23 | 65590282 | 81012210 | 73498890 | 61645954 | 75480818 | 58934134 | 51355138 | 66749206 | 50582990 | 52395242 | 54396142 | 51981442 | 51978014 | 54507254 | 51336910 | 45046614 | 38367490 | 52167938 | 49501530 | 49083430 | 0.783859578 | 2 | 0.739582698 | 0 | 0.655511049 | 0 |
| P38647 | Hspa9 | Stress-70 protein, mitochondrial | 2285.94 | 52.43 | 73461 | 32 | 72239962 | 83420410 | 65089530 | 65672986 | 90745834 | 99586074 | 82172226 | 98045706 | 114207554 | 110264730 | 140855410 | 128422482 | 117922594 | 127593634 | 112759482 | 135692866 | 136480514 | 125622346 | 116359058 | 110647066 | 1.33700453 | 2 | 1.663853775 | 0 | 1.656557963 | 0 |
| Q9Z0X1 | Aifm1 | Apoptosis-inducing factor 1, mitochondrial | 773.7 | 29.58 | 66765 | 14 | 3788268.75 | 2802908.5 | 3033380.5 | 2710384.75 | 3952886.25 | 5942557 | 5319947.5 | 4914535 | 11501732 | 3378811.5 | 14513321 | 14349574 | 8970858 | 10582814 | 9016713 | 11157832 | 5260577.5 | 7356709.5 | 9470906 | 7630630 | 1.906797 | 2 | 3.526147093 | 0 | 2.509644203 | 0 |
| P16110 | Lgals3 | Galectin-3 | 533.1 | 38.64 | 27515 | 9 | 3127265 | 5263680 | 5333368.5 | 4447551 | 19641258 | 20913506 | 11014831 | 27253018 | 28267058 | 18261660 | 104838130 | 112329122 | 53973854 | 64469754 | 27105718 | 112963626 | 55047658 | 31998546 | 130974898 | 27602866 | 2.795592271 | 2 | 9.592346625 | 0 | 9.483152152 | 0 |
| Q60692 | Psmb6 | Proteasome subunit beta type-6 | 193.33 | 16.81 | 25379 | 4 | 5311137.5 | 6450081 | 4340186 | 4557643 | 167663.27 | 2 | 2992026 | 130886.95 | 2218066 | 2801314.75 | 12489238 | 11316821 | 12225833 | 13102580 | 6924246 | 11425199 | 8095560.5 | 9920793 | 9014656 | 9019310 | 0.390954471 | 3 | 2.691674102 | 0 | 2.279549518 | 0 |
| Q9WVJ3 | Cpq | Carboxypeptidase Q | 247.46 | 12.98 | 51813 | 5 | 952888.88 | 1670381.75 | 1302459 | 1054140.5 | 1634993.75 | 2 | 145504.2 | 621619.94 | 529590.56 | 1438468.62 | 2 | 2 | 2 | 2 | 2 | 2 | 2 | 2 | 2 | 2 | 0.413490794 | 3 | 1.51175E-06 | 0 | 1.51175E-06 | 0 |
| Q9JI75 | Nqo2 | Ribosyldihydronicotinamide dehydrogenase [quinone] | 34.95 | 4.33 | 26248 | 1 | 1507739.25 | 1286004.5 | 1514669.88 | 1112830.5 | 1040020.69 | 2 | 1151715 | 1040033.25 | 578664.38 | 810650.25 | 2 | 2 | 765760 | 2 | 2 | 2 | 2 | 2 | 2 | 2 | 0.554235893 | 3 | 0.118516733 | 0 | 1.54768E-06 | 0 |
| P19973 | Lsp1 | Lymphocyte-specific protein 1 | 535.53 | 29.7 | 36714 | 8 | 14110519 | 17164270 | 11376011 | 13671353 | 11952719 | 10026948 | 7991208 | 5507593.5 | 11162857 | 14086948 | 78403722 | 83259778 | 52048194 | 56271646 | 31221914 | 67572890 | 47556506 | 26809206 | 51803542 | 33848426 | 0.714399794 | 3 | 4.411656078 | 0 | 3.333445576 | 0 |
| Q63945 | Set | Protein SET | 235.8 | 21.45 | 33406 | 5 | 5081914 | 5875709.5 | 7138754.5 | 4866965 | 6238439.5 | 4204240 | 4302546 | 6106717 | 2825121.75 | 4059759 | 2599681.25 | 2759130.5 | 2734282.25 | 2894488.5 | 2959823.5 | 3547256.25 | 2725510.25 | 1828033.75 | 2538033.5 | 2779915.75 | 0.736201078 | 3 | 0.477621734 | 0 | 0.459518165 | 0 |
| Q9ERS2 | Ndufa13 | NADH dehydrogenase [ubiquinone] 1 alpha subcomplex subunit 13 | 128.28 | 22.92 | 16860 | 3 | 5489364.5 | 5925315.5 | 4368043.5 | 4973944.5 | 3485926.5 | 4244008.5 | 4244567.5 | 3880203 | 2953882.5 | 2774555.75 | 2627483.5 | 1704241 | 2391005.75 | 2071138.25 | 2053243.38 | 1673015.88 | 1649582.38 | 3156083.5 | 391454.22 | 1767813.75 | 0.746504969 | 3 | 0.447440223 | 0 | 0.356312924 | 0 |
| P97371 | Psme1 | Proteasome activator complex subunit 1 | 240.45 | 17.67 | 28673 | 4 | 14518968 | 10753305 | 12940395 | 15649119 | 15157189 | 17291136 | 18487774 | 18334312 | 13755834 | 17312258 | 4928276.5 | 6329604 | 7259439 | 5337163 | 8145105 | 6145879 | 5158249 | 7575598 | 3609753 | 5843317 | 1.234172382 | 3 | 0.463634631 | 0 | 0.410507336 | 0 |
| P51410 | Rpl9 | Large ribosomal subunit protein uL6 | 698.08 | 67.19 | 21881 | 12 | 1874341.5 | 11892226 | 7061566 | 12254184 | 11707654 | 16075208 | 17614160 | 22591100 | 20844160 | 8027666 | 15708552 | 14681894 | 21409452 | 22504406 | 13824138 | 20707986 | 21051774 | 13776298 | 24579566 | 12924292 | 1.901146421 | 3 | 1.96759317 | 0 | 2.077248832 | 0 |
| P01897 | H2-L | H-2 class I histocompatibility antigen, L-D alpha chain | 327.25 | 21.27 | 40711 | 6 | 552173.75 | 2 | 2 | 553583.5 | 1461362.62 | 1057907.25 | 738171.62 | 1113074.62 | 2279357.5 | 1638222 | 4535230.5 | 3432047.5 | 3455267.25 | 3777174.75 | 3859601.25 | 4108263 | 4436619.5 | 6588569 | 3720116.25 | 3317544.25 | 2.659292397 | 3 | 7.424387063 | 0 | 8.636557144 | 0 |
| P18886 | Cpt2 | Carnitine O-palmitoyltransferase 2, mitochondrial | 47.37 | 1.52 | 74110 | 1 | 1313214.62 | 1845231.38 | 1696600.88 | 1516537.25 | 1425697.62 | 2 | 2 | 1824995.12 | 2 | 2 | 546640.69 | 2 | 2 | 2 | 2 | 590470.06 | 1307845.62 | 2 | 2 | 2 | 0.234056326 | 4 | 0.070107597 | 0 | 0.243459418 | 0 |
| P54227 | Stmn1 | Stathmin | 243.32 | 25.5 | 17274 | 4 | 1345075 | 1039445.12 | 2 | 1366825.5 | 998115.75 | 2 | 411985.69 | 2 | 782392.44 | 303823.5 | 7008651.5 | 9875368 | 7318129 | 6250941 | 5507581.5 | 6166086 | 4705661.5 | 3624702.5 | 5571385.5 | 5006386 | 0.315447349 | 4 | 7.57152297 | 0 | 5.27937991 | 0 |
| Q9Z0E6 | Gbp2 | Guanylate-binding protein 2 | 45 | 2.04 | 66739 | 1 | 4477860 | 4877511 | 4182352.75 | 4710503.5 | 6130350 | 741177.5 | 556665.44 | 5856077 | 474604.62 | 402787.31 | 2457598 | 1918367.12 | 2 | 1739172.88 | 2281119.25 | 809392.12 | 1346131.62 | 2245475.25 | 1211270.12 | 2 | 0.329441369 | 4 | 0.344411372 | 0 | 0.230213234 | 0 |
| Q3UIA2 | Arhgap17 | Rho GTPase-activating protein 17 | 45.23 | 1.06 | 92202 | 1 | 1606328.12 | 1761663.62 | 705776.38 | 902074.44 | 840755.69 | 1164809.38 | 2 | 784261.38 | 513321.28 | 463219.31 | 568867.38 | 540131.12 | 461600.47 | 502356.47 | 484372.94 | 2 | 2 | 451081.47 | 2 | 2 | 0.502976693 | 4 | 0.43966048 | 0 | 0.07755211 | 0 |
| P09581 | Csf1r | Macrophage colony-stimulating factor 1 receptor | 77.34 | 1.74 | 109179 | 2 | 1769150.25 | 3220037.5 | 2556959 | 2367380.75 | 4335557 | 1899541.75 | 1447376.88 | 638239.19 | 1873098 | 2488783.25 | 1497542 | 1429790.5 | 1310795.38 | 1335856.62 | 2 | 1212014.12 | 1254697.38 | 1418438.88 | 1076713.12 | 1134834 | 0.585794762 | 4 | 0.391182079 | 0 | 0.42786591 | 0 |
| Q9CQN1 | Trap1 | Heat shock protein 75 kDa, mitochondrial | 776.83 | 25.64 | 80209 | 13 | 5324754 | 5963155.5 | 7889784.5 | 8246447 | 7963160 | 3295123.5 | 2360031.5 | 4711253.5 | 6236478 | 7744682 | 1217339.5 | 3922995.25 | 3782413.5 | 2541079 | 4951045 | 2380164.5 | 3701084 | 2673585.5 | 5167504 | 3413228 | 0.688031238 | 4 | 0.463863357 | 0 | 0.489880989 | 0 |
| P10810 | Cd14 | Monocyte differentiation antigen CD14 | 706.03 | 34.7 | 39204 | 11 | 10942571 | 10061821 | 15048407 | 11770358 | 10768785 | 6218570 | 7596580.5 | 8346176 | 6411814.5 | 14659994 | 1973751.75 | 1787679.5 | 1974995.75 | 3591940.5 | 3960309 | 1611017.12 | 2270443.5 | 2640530.25 | 1191127.88 | 2485951 | 0.737868272 | 4 | 0.226800411 | 0 | 0.174069495 | 0 |
| Q69ZK0 | Prex1 | Phosphatidylinositol 3,4,5-trisphosphate-dependent Rac exchanger 1 protein | 198.61 | 3.03 | 184935 | 4 | 4671311.5 | 4800441 | 4561890.5 | 4068023.25 | 3112972 | 2852672.25 | 3940321 | 4242052 | 1586398.38 | 3312852.75 | 470543.62 | 911467.69 | 1105735.62 | 1729260.62 | 3009376 | 2 | 2583738 | 1700429.88 | 2876448.75 | 1943441 | 0.751099132 | 4 | 0.340631948 | 0 | 0.429140461 | 0 |
| Q8BX70 | Vps13c | Intermembrane lipid transfer protein VPS13C | 479.16 | 3.5 | 420089 | 9 | 4514338 | 5282047.5 | 6020406.5 | 4872832.5 | 3780230 | 3147954.5 | 4148494 | 5005071 | 3372048 | 3378204 | 928694.62 | 980428.62 | 3563590 | 3125135.5 | 1476465.62 | 2935037.25 | 3136278.75 | 3676333.25 | 2859568 | 3263125.25 | 0.778581315 | 4 | 0.411703076 | 0 | 0.648567097 | 0 |
| P08752 | Gnai2 | Guanine nucleotide-binding protein G(i) subunit alpha-2 | 832.73 | 38.31 | 40489 | 12 | 38154990 | 37151450 | 43610278 | 33984730 | 42021894 | 34239930 | 28910452 | 37368082 | 31825398 | 36520238 | 22027290 | 16357914 | 20013668 | 17289804 | 28910366 | 13794714 | 15245395 | 13513181 | 15860340 | 13804663 | 0.866310306 | 4 | 0.536616297 | 0 | 0.370495869 | 0 |
| P26638 | Sars1 | Serine--tRNA ligase, cytoplasmic | 893.22 | 34.96 | 58389 | 19 | 18166996 | 13064344 | 20896140 | 19943166 | 25177930 | 21428552 | 20693212 | 22863536 | 30551008 | 31616026 | 51887238 | 50066766 | 51460562 | 46346346 | 58075158 | 47367470 | 60992722 | 62088558 | 55315782 | 49605318 | 1.307498158 | 4 | 2.651309465 | 0 | 2.831608043 | 0 |
| P17182 | Eno1 | Alpha-enolase | 1996.97 | 65.21 | 47141 | 28 | 18871678 | 20500014 | 31773610 | 13039551 | 31881922 | 24144164 | 30472144 | 38017822 | 39410174 | 51412194 | 53396754 | 57394990 | 46220606 | 38302958 | 50591174 | 65843842 | 43127166 | 35574562 | 41043554 | 31954728 | 1.580611661 | 4 | 2.118663864 | 0 | 1.874299101 | 0 |
| Q8CDN6 | Txnl1 | Thioredoxin-like protein 1 | 173.37 | 18.34 | 32237 | 4 | 340801.19 | 1385897.75 | 980847.62 | 2 | 877980.75 | 1058351.88 | 1215002.25 | 1062080 | 1864510.75 | 1159168 | 7575366 | 6509465.5 | 3940408.5 | 5907890 | 2748887.75 | 7773859 | 4149049.5 | 5284128 | 4080512.5 | 7419201 | 1.773549267 | 4 | 7.441584057 | 0 | 8.006279553 | 0 |
| Q60875 | Arhgef2 | Rho guanine nucleotide exchange factor 2 | 637.51 | 16.24 | 111974 | 14 | 1510922.88 | 2733868.5 | 2547543.5 | 2325896.75 | 3608813.5 | 4840334.5 | 3543614.25 | 2995796.25 | 9012949 | 2712715.25 | 6179504 | 9439668 | 4405860.5 | 9257214 | 3771255.5 | 5120592.5 | 4070074.75 | 3682908 | 3869355.5 | 11118500 | 1.815457478 | 4 | 2.597107315 | 0 | 2.189151564 | 0 |
| P62315 | Snrpd1 | Small nuclear ribonucleoprotein Sm D1 | 55.54 | 10.92 | 13282 | 1 | 400947.28 | 812720.31 | 2 | 2 | 1321106.25 | 1614984.5 | 530702.5 | 1017905.88 | 1379937.12 | 837957.12 | 3986705.25 | 3772382.25 | 2456251.75 | 2700423.5 | 2095967 | 3215754.5 | 3325476.75 | 1378213.5 | 2812080 | 2614185.75 | 2.123060662 | 4 | 5.922305897 | 0 | 5.265041492 | 0 |
| Q61792 | Lasp1 | LIM and SH3 domain protein 1 | 480.51 | 33.46 | 29994 | 9 | 3079353.25 | 2501092.5 | 2487438.75 | 9394012 | 7429395.5 | 19475224 | 17196160 | 4064809 | 14479289 | 5231668.5 | 24225150 | 31162676 | 12979984 | 18550266 | 13604862 | 22630266 | 21510112 | 19416878 | 17571968 | 26931716 | 2.428445679 | 4 | 4.038478115 | 0 | 4.341315027 | 0 |
| P17742 | Ppia | Peptidyl-prolyl cis-trans isomerase A | 928.91 | 82.32 | 17971 | 14 | 1135532.38 | 3290772.5 | 4888471 | 468504.22 | 21764478 | 11251121 | 7452503 | 12404534 | 27336840 | 19206822 | 131987122 | 134021498 | 91648154 | 97204114 | 55781694 | 57371822 | 65268026 | 36009546 | 63481854 | 35583158 | 2.461405332 | 4 | 16.18633503 | 0 | 8.16902441 | 0 |
| Q5I0H4 | Tmco1 | Calcium load-activated calcium channel | 209.16 | 19.68 | 21175 | 3 | 128589.49 | 2 | 198485.47 | 324438.53 | 141879.17 | 123027.49 | 455134.09 | 1157054.38 | 2294833.5 | 545733.12 | 2261938.75 | 2089909.75 | 1874141.5 | 2501004 | 387632.5 | 1678755 | 1320605.5 | 1044052.25 | 2524544.5 | 1493966.25 | 5.767347338 | 4 | 11.48813694 | 0 | 10.16130295 | 0 |
| O89023 | Tpp1 | Tripeptidyl-peptidase 1 | 326.12 | 14.41 | 61342 | 5 | 1064131.38 | 716093.06 | 1706556 | 782679.19 | 474137.03 | 2 | 2 | 529322.62 | 2 | 1143496.88 | 2 | 2 | 2 | 2 | 2 | 2 | 2 | 2 | 2 | 2 | 0.352649186 | 5 | 2.10811E-06 | 0 | 2.10811E-06 | 0 |
| P26883 | Fkbp1a | Peptidyl-prolyl cis-trans isomerase FKBP1A | 164 | 34.26 | 11923 | 3 | 1820123.38 | 1370842.75 | 1429936.5 | 1151773.5 | 1995983.62 | 2 | 2 | 2217760 | 707807.06 | 765213.62 | 14284028 | 16163495 | 10632491 | 10729690 | 6113886 | 5814974 | 6636910.5 | 6188723 | 8938636 | 6504553 | 0.475086411 | 5 | 7.456059586 | 0 | 4.387345771 | 0 |
| P21107 | Tpm3 | Tropomyosin alpha-3 chain | 736.17 | 26.32 | 32994 | 13 | 19450480 | 18463368 | 16331821 | 17673392 | 17797538 | 13533726 | 13052179 | 20647862 | 9767908 | 12073933 | 11375328 | 12365018 | 11207079 | 9592222 | 5648078.5 | 13363112 | 12604268 | 7993012 | 15140918 | 11118335 | 0.769931192 | 5 | 0.559402898 | 0 | 0.671220774 | 0 |
| Q63486 | Rraga | Ras-related GTP-binding protein A | 113.3 | 6.71 | 36566 | 2 | 7256780 | 6091934 | 6348833 | 5655308 | 6408092 | 5148404 | 5582542.5 | 9704875 | 5632498 | 4930169.5 | 3929210.25 | 4747738 | 4477466.5 | 2541026.25 | 2180480.75 | 2 | 2523466.25 | 2560047 | 3546446.25 | 2101292.75 | 0.975993852 | 5 | 0.562827102 | 0 | 0.337875765 | 0 |
| Q8CAQ8 | Immt | MICOS complex subunit Mic60 | 1805.59 | 39.1 | 83900 | 29 | 15482612 | 23295006 | 17427990 | 24174128 | 21498916 | 23632562 | 23471872 | 21145300 | 34057606 | 30306002 | 39647094 | 34344882 | 35009886 | 38469806 | 26522260 | 41791870 | 32802866 | 37846578 | 31860216 | 31742002 | 1.301679394 | 5 | 1.707854635 | 0 | 1.727972726 | 0 |
| Q8BGQ7 | Aars1 | Alanine--tRNA ligase, cytoplasmic | 1040.82 | 23.86 | 106909 | 19 | 14857398 | 15724374 | 22310118 | 16816902 | 32632782 | 28528192 | 24486130 | 24159734 | 27561500 | 29517566 | 37923506 | 35735354 | 40622742 | 37853658 | 40428814 | 41443006 | 46812290 | 47575250 | 44113666 | 47680474 | 1.311814122 | 5 | 1.881582103 | 0 | 2.224166359 | 0 |
| P18760 | Cfl1 | Cofilin-1 | 674.92 | 74.1 | 18560 | 10 | 15788739 | 14159229 | 26441510 | 19707402 | 53186534 | 31805398 | 21150924 | 40351130 | 66201578 | 44425730 | 118301770 | 108682354 | 71478322 | 67099610 | 67825754 | 116420698 | 70246474 | 78065674 | 59094290 | 73162114 | 1.577424 | 5 | 3.35223055 | 0 | 3.070689717 | 0 |
| Q9DCE9 | Igtp | Immunity-related GTPase family M protein 3 | 37.38 | 2.13 | 48494 | 1 | 626401.06 | 114323.32 | 471711.84 | 521948.31 | 738063.31 | 690533.44 | 1650147 | 227845.33 | 2173068 | 1790851.25 | 2 | 2 | 2 | 2 | 2 | 2 | 2 | 2 | 2 | 2 | 2.642096191 | 5 | 4.04457E-06 | 0 | 4.04457E-06 | 0 |
| Q80VA0 | Galnt7 | N-acetylgalactosaminyltransferase 7 | 129 | 4.87 | 75419 | 3 | 824577.31 | 2 | 296602.88 | 692481.56 | 2 | 1729049.5 | 2 | 1033933.62 | 1644264.5 | 775466.94 | 1611618.12 | 1629747.75 | 1179860.75 | 2997367 | 892947.94 | 1080482.75 | 1563489 | 1807969.12 | 1315545.88 | 1481727.12 | 2.857591902 | 5 | 4.582730616 | 0 | 3.996995516 | 0 |
| P31786 | Dbi | Acyl-CoA-binding protein | 193.65 | 47.13 | 10000 | 3 | 2 | 826058.38 | 915903.38 | 2 | 1696471.38 | 3686650 | 769765.12 | 3290002 | 1510455.25 | 1131934.5 | 3593361.75 | 4217579.5 | 3167867.75 | 2436100.25 | 3387703.5 | 4398622.5 | 4887994 | 1963294.38 | 5033773 | 3547265.5 | 3.021374667 | 5 | 4.886700575 | 0 | 5.767431124 | 0 |
| P47758 | Srprb | Signal recognition particle receptor subunit beta | 303.07 | 27.14 | 29579 | 6 | 2 | 2 | 2 | 658349.94 | 676036.94 | 533938.25 | 2 | 931058.38 | 1626036.5 | 1907312.5 | 3268400.75 | 773379 | 2665043.25 | 3583411 | 1684533.62 | 1213347.25 | 2848890.5 | 1224071.25 | 3198947 | 1048925.88 | 3.745784098 | 5.5 | 8.97394448 | 0 | 7.144958597 | 0 |
| P13379 | Cd5 | T-cell surface glycoprotein CD5 | 278.94 | 12.15 | 53849 | 4 | 1926688.25 | 1240782.25 | 820710.69 | 1731334.75 | 3880314 | 2 | 2 | 2 | 1801055.38 | 1791979.5 | 2 | 2 | 2 | 2 | 2 | 2 | 473087.78 | 2 | 2 | 2 | 0.374281722 | 6 | 1.04169E-06 | 0 | 0.049281683 | 0 |
| Q3UEB3 | Puf60 | Poly(U)-binding-splicing factor PUF60 | 408.66 | 19.86 | 60249 | 7 | 4766328.5 | 3952435.25 | 5714857 | 6682920.5 | 8402830 | 985633.88 | 3320211.25 | 5241498 | 4617472.5 | 5980181.5 | 2536850.5 | 2881044.5 | 3492566 | 3378124.25 | 3738603.5 | 1795852.25 | 661964.38 | 1330883.25 | 1721580.25 | 2 | 0.682433137 | 6 | 0.542938012 | 0 | 0.18666665 | 0 |
| P60229 | Eif3e | Eukaryotic translation initiation factor 3 subunit E | 369.64 | 11.69 | 52221 | 5 | 16118402 | 19712942 | 17804114 | 22389140 | 34142374 | 12629628 | 12169330 | 20120472 | 11413941 | 20813110 | 10963845 | 9694226 | 11030013 | 11987529 | 9341992 | 8119557 | 8050870 | 10818258 | 7636755.5 | 7956243 | 0.700268689 | 6 | 0.481247728 | 0 | 0.386519505 | 0 |
| Q9JMH6 | Txnrd1 | Thioredoxin reductase 1, cytoplasmic | 1055.46 | 31 | 67084 | 17 | 5671443.5 | 3921003 | 9653176 | 8867238 | 11829066 | 7351943 | 8161773.5 | 15499199 | 15774877 | 17937946 | 41038174 | 44365510 | 39475610 | 31023018 | 71183794 | 60260282 | 80854162 | 75358306 | 73789490 | 78949154 | 1.62049616 | 6 | 5.685406937 | 0 | 9.243705208 | 0 |
| A2AQP0 | Myh7b | Myosin-7B | 83.49 | 0.88 | 221497 | 2 | 9182563 | 13507803 | 75768.74 | 15127328 | 2622858.75 | 18483606 | 10243457 | 1694266.62 | 30398210 | 20142752 | 184951922 | 202230274 | 127388786 | 187972418 | 91578922 | 340920322 | 237405666 | 126010874 | 83090978 | 143220674 | 1.998263629 | 6 | 19.60005975 | 0 | 22.96971886 | 0 |
| P61314 | Rpl15 | Large ribosomal subunit protein eL15 | 384.7 | 37.25 | 24146 | 8 | 2062830 | 2737747.75 | 2928194 | 1996129.75 | 5394601.5 | 3095556.25 | 2143672.5 | 6030580.5 | 18013806 | 2837059.75 | 1486693.12 | 1860928.25 | 1739066.88 | 62862.15 | 1124286.12 | 61523.97 | 2 | 2 | 2 | 1701757 | 2.124453099 | 6 | 0.414949917 | 0 | 0.116623342 | 0 |
| Q9ER72 | Cars1 | Cysteine--tRNA ligase, cytoplasmic | 256.06 | 5.9 | 94860 | 4 | 2 | 1678257.12 | 2 | 2 | 3904268.5 | 2056942.75 | 1609134.25 | 2894205.25 | 3116892.5 | 3268230 | 7720486 | 6098886 | 6003927 | 5086881 | 4414969 | 8308785.5 | 6971766 | 5075053 | 9450274 | 4149784.25 | 2.318912929 | 6 | 5.253019776 | 0 | 6.082484625 | 0 |
| Q5PQM0 | Tmem168 | Transmembrane protein 168 | 83.12 | 4.02 | 79772 | 2 | 849051.38 | 898014.19 | 699093.56 | 912721.88 | 845030.5 | 717084.75 | 622145.94 | 71431.26 | 981628.12 | 813195.25 | 2 | 2 | 62702.82 | 592007.62 | 218857.95 | 2 | 159603.06 | 2 | 2 | 108691.73 | 0.762500665 | 7 | 0.2077999 | 0 | 0.063821703 | 0 |
| Q3KRE8 | Tubb2b | Tubulin beta-2B chain | 1728.97 | 59.1 | 49953 | 25 | 1154048.12 | 1821816.75 | 1777855.62 | 1222610.75 | 960970.75 | 1392329.88 | 1071612.88 | 1544969.5 | 877473.06 | 953444.25 | 694621.19 | 2 | 955417.19 | 842649 | 717867.81 | 846916.25 | 930017.25 | 913751.81 | 862855.31 | 749012.38 | 0.841801262 | 7 | 0.462796227 | 0 | 0.620205522 | 0 |
| P24668 | M6pr | Cation-dependent mannose-6-phosphate receptor | 506.57 | 37.05 | 31172 | 8 | 24228282 | 26947624 | 27690430 | 26603188 | 23629946 | 21542216 | 19402504 | 34811482 | 15435472 | 25723824 | 12102146 | 12758232 | 15710543 | 12905508 | 19714754 | 13549334 | 16554526 | 20038508 | 16413667 | 17525036 | 0.905623377 | 7 | 0.566936355 | 0 | 0.651289049 | 0 |
| P35564 | Canx | Calnexin | 965.5 | 30.12 | 67278 | 18 | 11680755 | 10880212 | 12214000 | 15951912 | 12449346 | 10106980 | 11162696 | 9945812 | 14933893 | 11800377 | 9814306 | 9192345 | 10559762 | 10239144 | 9637266 | 10700595 | 9853512 | 9222194 | 6904150 | 10544959 | 0.917271616 | 7 | 0.782617559 | 0 | 0.747518707 | 0 |
| P46460 | Nsf | Vesicle-fusing ATPase | 565.22 | 18.41 | 82613 | 12 | 7657873.5 | 8285418.5 | 8381081 | 9428272 | 12625320 | 8805435 | 8435404 | 11836935 | 12303096 | 10740062 | 5671492 | 3153711 | 4574006 | 6105888 | 4652963.5 | 2919009.25 | 2566295.5 | 5175667.5 | 1291616.12 | 3383299.5 | 1.123829646 | 7 | 0.520895225 | 0 | 0.330671858 | 0 |
| Q99JX3 | Gorasp2 | Golgi reassembly-stacking protein 2 | 121.45 | 5.32 | 47038 | 2 | 687307.81 | 783291.5 | 766412.56 | 1012279.19 | 1111869.25 | 1096108 | 919737.44 | 1170524.25 | 924634.12 | 827632.75 | 2184248.75 | 1965609.88 | 2278919.5 | 1597432.62 | 1374193.25 | 2693940.25 | 2669931.5 | 3110761.75 | 3844427.75 | 2783130.5 | 1.132413443 | 7 | 2.155482333 | 0 | 3.462883883 | 0 |
| O70503 | Hsd17b12 | Very-long-chain 3-oxoacyl-CoA reductase | 862.82 | 56.73 | 34742 | 16 | 11685065 | 13341166 | 7106831 | 16014349 | 11538698 | 14953956 | 15093060 | 7045533.5 | 17224804 | 17645932 | 18152814 | 18317658 | 28618138 | 24939064 | 20177930 | 18462868 | 27264064 | 36997754 | 28010094 | 32055632 | 1.205695709 | 7 | 1.846419642 | 0 | 2.392355849 | 0 |
| Q922Y1 | Ubxn1 | UBX domain-containing protein 1 | 149.75 | 14.81 | 33573 | 3 | 517197.91 | 704623.06 | 517041.06 | 862257.19 | 1048461.88 | 2 | 890242.5 | 1189327.38 | 2245112.5 | 898708 | 2479077 | 2337892 | 1729683.25 | 3299714.25 | 2027762.25 | 4058855 | 3126169.75 | 2835307.5 | 3289940.75 | 2322664 | 1.431230664 | 7 | 3.253559361 | 0 | 4.283488042 | 0 |
| Q63524 | Tmed2 | Transmembrane emp24 domain-containing protein 2 | 77.89 | 6.47 | 22733 | 1 | 2 | 2 | 2 | 2 | 2 | 2 | 168006.38 | 151387.86 | 2 | 2 | 250334.98 | 234781.3 | 828570.12 | 298280.31 | 148730.34 | 301292.34 | 260240.67 | 169582.27 | 112005.8 | 155691.83 | 31940.024 | 7.5 | 176069.705 | 0 | 99881.291 | 0 |
| Q5I0E7 | Tmed9 | Transmembrane emp24 domain-containing protein 9 | 176.46 | 20 | 27028 | 4 | 3573692.75 | 2954117 | 2489207.25 | 4104578.5 | 2578163.75 | 1081017.88 | 2727770.25 | 2694144.5 | 2623489.75 | 2644813.5 | 1290146.75 | 1871314.88 | 1939953.5 | 1746271.12 | 2137977.75 | 1431636.88 | 1862592.5 | 2240060 | 541646.69 | 1244666.38 | 0.749771744 | 8 | 0.572344063 | 0 | 0.466287561 | 0 |
| P48771 | Cox7a2 | Cytochrome c oxidase subunit 7A2, mitochondrial | 63.11 | 12.05 | 9291 | 1 | 4008257.25 | 4249798.5 | 2589541 | 4579443 | 5480141 | 3044305.75 | 2658512.25 | 3710399.75 | 2716044.75 | 4599937 | 1573091.25 | 1710618.75 | 1420000.12 | 1415482.12 | 1837799.38 | 2014566.88 | 1824436.12 | 1582048 | 1746673.75 | 1966623.75 | 0.800165249 | 8 | 0.380586542 | 0 | 0.436900059 | 0 |
| Q9JHI5 | Ivd | Isovaleryl-CoA dehydrogenase, mitochondrial | 169.63 | 10.38 | 46325 | 4 | 3247137.5 | 4195867 | 3298377.75 | 3265066.5 | 3435865.5 | 3498016.25 | 2936365 | 3610277 | 2584817 | 1456980.5 | 1817727.5 | 1643076.88 | 2000313 | 1599960 | 1280573 | 2306199.5 | 2260022.75 | 2661476.25 | 2226282.25 | 2168133 | 0.807602452 | 8 | 0.478242179 | 0 | 0.666317186 | 0 |
| Q60854 | Serpinb6 | Serpin B6 | 410.59 | 16.93 | 42599 | 6 | 48418850 | 56533778 | 47686474 | 40573434 | 38802646 | 42597970 | 40639030 | 55586614 | 33388056 | 35879454 | 25359872 | 24548882 | 24316266 | 23957408 | 21809752 | 13469201 | 17514252 | 26898156 | 22572436 | 25916026 | 0.896885808 | 8 | 0.517173829 | 0 | 0.458461684 | 0 |
| Q9D7X3 | Dusp3 | Dual specificity protein phosphatase 3 | 338.48 | 28.11 | 20472 | 4 | 6790830 | 11850266 | 6409896.5 | 8638889 | 7945000 | 8845125 | 7341637 | 10241188 | 9318432 | 8444439 | 5625802 | 6012172 | 4370785 | 5446378 | 4121944 | 4924042 | 3329517.75 | 4188902 | 4017292.5 | 3708265.75 | 1.061389379 | 8 | 0.614318573 | 0 | 0.484402003 | 0 |
| P26151 | Fcgr1 | High affinity immunoglobulin gamma Fc receptor I | 130.07 | 8.66 | 44888 | 3 | 1108650.38 | 1011444.88 | 1563863.75 | 1690631.62 | 1458674.88 | 1637133.38 | 1676078.12 | 2194248.25 | 1163304.12 | 1290947.12 | 933770.94 | 828623.44 | 2 | 371523.06 | 683253.31 | 2 | 2 | 2 | 2 | 2 | 1.165140002 | 8 | 0.412273275 | 0 | 1.46343E-06 | 0 |
| Q62159 | Rhoc | Rho-related GTP-binding protein RhoC | 251.62 | 24.87 | 22006 | 5 | 1454328.62 | 2939105.75 | 1155096.75 | 2486241.75 | 2500522.75 | 2479925.75 | 2264504.5 | 3289144.25 | 2833124.25 | 2550940.25 | 5872372.5 | 5251390 | 5997114 | 5358010.5 | 4464569.5 | 6599858.5 | 7317321.5 | 6804928 | 8640946 | 6316176 | 1.273589227 | 8 | 2.557446651 | 0 | 3.386637764 | 0 |
| Q11136 | Pepd | Xaa-Pro dipeptidase | 390.26 | 16.84 | 55029 | 8 | 2358463 | 1234343.75 | 2443930 | 2205081.5 | 2710858 | 3079266 | 1888948 | 4525867.5 | 3159498.25 | 1941895.88 | 9243808 | 11072866 | 9342275 | 9774340 | 5483399 | 12431800 | 7240017 | 7095118.5 | 5210721 | 9175137 | 1.332594454 | 8 | 4.100978334 | 0 | 3.757327667 | 0 |
| Q80W92 | Vac14 | Protein VAC14 homolog | 101.98 | 3.19 | 88068 | 2 | 1028617.12 | 1589233.5 | 715809.69 | 1565242.75 | 1616001.62 | 809300.75 | 1990515.5 | 2 | 1180903.5 | 951926.88 | 2 | 2 | 2 | 2 | 2 | 2 | 2 | 2 | 2 | 2 | 0.757132893 | 9 | 1.53494E-06 | 0 | 1.53494E-06 | 0 |
| Q9CXT6 | Kdm8 | Bifunctional peptidase and arginyl-hydroxylase JMJD5 | 34.27 | 2.17 | 47145 | 1 | 587004.81 | 741062.12 | 233807.59 | 1112596 | 239098.09 | 2 | 2 | 94136.98 | 872458.88 | 1492081.62 | 5061917.5 | 5748963 | 5361124.5 | 5354169 | 3723322.5 | 6701942.5 | 7829952.5 | 8186082 | 7989992 | 8723644 | 0.843872861 | 9 | 8.666175361 | 0 | 13.53378563 | 0 |
| Q9Z0J0 | Npc2 | NPC intracellular cholesterol transporter 2 | 224.79 | 36.91 | 16442 | 5 | 12622767 | 6676157 | 6984917 | 5793348.5 | 3422078 | 5129394 | 4621308 | 5907170 | 4392889 | 10499966 | 887195.88 | 602002.12 | 2319336.5 | 2532777.5 | 996490.06 | 1730604.62 | 2 | 3035582.75 | 1350598.25 | 2546385 | 0.860601617 | 9 | 0.206702915 | 0 | 0.244038067 | 0 |
| P09527 | Rab7a | Ras-related protein Rab-7a | 969.54 | 77.78 | 23504 | 16 | 79725794 | 69026698 | 86186418 | 51292674 | 46882006 | 54870722 | 58315638 | 64479886 | 51150698 | 59932214 | 40090334 | 31785804 | 35977722 | 34465758 | 45270050 | 28003788 | 33135716 | 34217250 | 16345882 | 31797758 | 0.866818907 | 9 | 0.563140243 | 0 | 0.430785169 | 0 |
| P62627 | Dynlrb1 | Dynein light chain roadblock-type 1 | 86.54 | 21.88 | 10990 | 2 | 638098.94 | 1458284 | 822269.06 | 693453.31 | 1465202.75 | 1371334.38 | 874332.94 | 1647039.25 | 783915.25 | 822466 | 3148254.5 | 2631294.5 | 2535769 | 2852504.25 | 1696800.88 | 2851766 | 2721931.75 | 2206940 | 2968842 | 3380210.75 | 1.083071532 | 9 | 2.533748785 | 0 | 2.782909828 | 0 |
| Q64339 | Isg15 | Ubiquitin-like protein ISG15 | 40.09 | 11.18 | 17898 | 1 | 2 | 2 | 46701.96 | 2 | 2 | 2 | 2 | 2 | 142907.44 | 159444.38 | 810289.06 | 744299.44 | 710402.06 | 1278398.5 | 1048624.62 | 381620.69 | 1937971.62 | 793687.69 | 1464504.88 | 1004523.5 | 6.473090964 | 9 | 98.3090904 | 0 | 119.5100227 | 0 |
| Q9R0C8 | Vav3 | Guanine nucleotide exchange factor VAV3 | 69.02 | 2.13 | 97968 | 2 | 2 | 391021.97 | 2 | 2 | 2 | 2 | 2 | 2 | 2 | 2 | 1263986.38 | 1219317.62 | 1056342.88 | 1083186.5 | 1004115.25 | 914945.25 | 902887.25 | 1268717.38 | 1057711.38 | 915242.56 | 2.55735E-05 | 10 | 14.39006997 | 0 | 12.93891571 | 0 |
| Q62087 | Pon3 | Serum paraoxonase/lactonase 3 | 106.99 | 7.63 | 39351 | 2 | 396417.75 | 293665.19 | 2 | 2 | 2 | 2 | 308404.81 | 2 | 2 | 2 | 3272392.75 | 2565963.25 | 3093448.25 | 2663525 | 1699117 | 3400310.5 | 3397370 | 3893108 | 1907119.12 | 3518726.25 | 0.44691748 | 10 | 19.2648302 | 0 | 23.35443004 | 0 |
| Q99KP6 | Prpf19 | Pre-mRNA-processing factor 19 | 259.72 | 21.63 | 55239 | 6 | 1802065.62 | 1367881 | 1225169.75 | 5200482 | 2636250.75 | 1541520.12 | 1627458.62 | 1859515.38 | 2420706.75 | 1142465.88 | 2 | 533287.88 | 2 | 527514.88 | 2 | 44853.05 | 2 | 2 | 2 | 2 | 0.702401302 | 10 | 0.086725134 | 0 | 0.003667561 | 0 |
| P62960 | Ybx1 | Y-box-binding protein 1 | 102.39 | 5.9 | 35730 | 1 | 766657.31 | 1014144.19 | 636797.25 | 1295894.62 | 1304048.5 | 2 | 2 | 1650829.25 | 1137573.62 | 784355.19 | 3438250 | 2417723.25 | 1900699.75 | 2483723.25 | 3043710.25 | 2601958.75 | 2873486 | 2996822.5 | 2537314.5 | 2327003.75 | 0.712054259 | 10 | 2.647532765 | 0 | 2.65799187 | 0 |
| P34022 | Ranbp1 | Ran-specific GTPase-activating protein | 183.44 | 15.27 | 23596 | 4 | 1420200.5 | 3561573.5 | 1699603.38 | 759469.19 | 1445812.75 | 2 | 1776315.38 | 1102145.88 | 2899010 | 1232289 | 14096292 | 15866154 | 11501342 | 12312088 | 8028302.5 | 13143032 | 8997118 | 8945227 | 14921499 | 9369192 | 0.788796105 | 10 | 6.954714508 | 0 | 6.231370643 | 0 |
| Q3U0V1 | Khsrp | Far upstream element-binding protein 2 | 1094.47 | 30.75 | 76775 | 20 | 5712495.5 | 20302098 | 13031439 | 11912410 | 19202644 | 12633669 | 9263024 | 12385738 | 13120324 | 12533748 | 36732502 | 37390714 | 26217434 | 32992780 | 24233652 | 28538248 | 36204358 | 20755536 | 41699170 | 21183100 | 0.854269881 | 10 | 2.245790222 | 0 | 2.114853395 | 0 |

| | | | | | | | | | | | | | | | | | | | | | | | | | | | | | | | | |
|---|---|---|---|---|---|---|---|---|---|---|---|---|---|---|---|---|---|---|---|---|---|---|---|---|---|---|---|---|---|---|---|---|
| P09055 | Itgb1 | Integrin beta-1 | 859.43 | 23.68 | 88231 | 17 | 26827254 | 24903076 | 25213880 | 34015998 | 42076114 | 19639934 | 18380142 | 35698314 | 33658262 | 28997768 | 17964546 | 21100814 | 17282256 | 18028760 | 21999884 | 14219216 | 21699568 | 20745810 | 18961220 | 20015696 | 0.891124527 | 10 | 0.629760692 | 0 | 0.624959544 | 0 |
| Q9CPR5 | Mrpl15 | Large ribosomal subunit protein uL15m | 180.13 | 23.05 | 33542 | 5 | 1027635.94 | 1525119.62 | 716517.38 | 707966.25 | 856651.56 | 587277.06 | 605917.88 | 1541466.62 | 1665088.62 | 470134.16 | 2 | 2 | 63511.32 | 507334.44 | 2 | 2 | 2 | 343751.91 | 2 | 2 | 1.007446091 | 10 | 0.118093641 | 0 | 0.071114539 | 0 |
| O35093 | Timm23 | Mitochondrial import inner membrane translocase subunit Tim23 | 104.3 | 12.92 | 21907 | 2 | 1098240.25 | 2129503 | 1220345.75 | 2297182.75 | 4049270.5 | 2254189 | 1972933.75 | 2824388 | 2455170 | 1661261.25 | 786754.19 | 811343.31 | 1025088.44 | 929792.19 | 352450 | 290347.56 | 2 | 2 | 1095014.38 | 2 | 1.034591532 | 10 | 0.361796549 | 0 | 0.128339665 | 0 |
| P23116 | Eif3a | Eukaryotic translation initiation factor 3 subunit A | 601.01 | 9.75 | 161936 | 13 | 4492523.5 | 5516413 | 4311230.5 | 3401262 | 4692750 | 2638923.75 | 4322498 | 6232332.5 | 5873450.5 | 4500970.5 | 11102821 | 9795734 | 10696893 | 11979127 | 7383088.5 | 9240019 | 8334341 | 7508762 | 8914103 | 7389175 | 1.0514851 | 10 | 2.273456614 | 0 | 1.846438364 | 0 |
| Q64213 | Sf1 | Splicing factor 1 | 364.33 | 13.63 | 70403 | 7 | 2724250.75 | 2156467.25 | 1531398.12 | 4250844 | 1700350 | 2274204.25 | 2646129.5 | 405504.06 | 4725358.5 | 4029427.25 | 6785057 | 7441526.5 | 6169735 | 6316582 | 7599611.5 | 9379657 | 9390071 | 4517537 | 8701181 | 6432799 | 1.138904017 | 10 | 2.77534994 | 0 | 3.107682702 | 0 |
| P61979 | Hnrnpk | Heterogeneous nuclear ribonucleoprotein K | 996.27 | 37.8 | 50976 | 17 | 7852576 | 18073958 | 17409362 | 30763140 | 28724942 | 18411612 | 19099608 | 25529170 | 28681204 | 27520128 | 37159174 | 39831810 | 37411566 | 43743558 | 31724326 | 33389050 | 30941152 | 33351522 | 42603974 | 32481794 | 1.159668438 | 10 | 1.84655795 | 0 | 1.680225715 | 0 |
| P05555 | Itgam | Integrin alpha-M | 966.76 | 18.39 | 127481 | 16 | 13079176 | 11892058 | 15496410 | 14781898 | 26167896 | 13002111 | 13304308 | 19713490 | 25160576 | 24379834 | 9109539 | 8581599 | 8856685 | 7429529 | 4696345 | 7407742 | 2829743.25 | 2151679.25 | 5894723 | 4603535 | 1.173708254 | 10 | 0.475005084 | 0 | 0.281112045 | 0 |
| Q9DCR2 | Ap3s1 | AP-3 complex subunit sigma-1 | 46.62 | 5.7 | 21732 | 1 | 2 | 2 | 2 | 2 | 2 | 2 | 2 | 2 | 760315.88 | 2 | 949730.12 | 1174466.25 | 1064903.62 | 1068920 | 779012.56 | 1358409.62 | 1371300.75 | 1277988.12 | 883522.12 | 1549241.25 | 76032.388 | 10 | 503703.255 | 0 | 644046.186 | 0 |
| P62858 | Rps28 | Small ribosomal subunit protein eS28 | 200 | 40.58 | 7841 | 4 | 1741975.38 | 674755.38 | 1317763.38 | 1416598.25 | 3088444 | 1596316.38 | 851072.94 | 147396.55 | 1427506.5 | 2349439 | 10933044 | 9701996 | 8382043 | 5661948.5 | 9986816 | 6828980 | 8204428.5 | 4876772 | 6128217 | 5608006 | 0.773311879 | 11 | 5.42091756 | 0 | 3.840799045 | 0 |
| P60335 | Pcbp1 | Poly(rC)-binding protein 1 | 914.61 | 44.38 | 37498 | 11 | 7189898.5 | 11471905 | 8875188 | 15302410 | 19143796 | 7596127 | 9245277 | 9286498 | 13237682 | 12706422 | 56809302 | 55809998 | 47651510 | 54187330 | 43222330 | 42404454 | 43959934 | 39640782 | 48475118 | 43078566 | 0.840098738 | 11 | 4.157263265 | 0 | 3.509965003 | 0 |
| Q8R1I1 | Uqcr10 | Cytochrome b-c1 complex subunit 9 | 71.78 | 12.5 | 7446 | 2 | 5766986 | 3071199 | 3000463 | 2156913.75 | 2221980 | 3522641 | 4137132 | 2335787 | 3089473 | 1538639.75 | 88681.15 | 1791798.75 | 1928256.5 | 1318080.25 | 480309.03 | 2093396 | 748677.44 | 1036607.5 | 1071987.88 | 2 | 0.901719445 | 11 | 0.345744489 | 0 | 0.305266414 | 0 |
| Q9CR62 | Slc25a11 | Mitochondrial 2-oxoglutarate/malate carrier protein | 853.05 | 44.59 | 34155 | 14 | 21739106 | 25860332 | 20349416 | 17345914 | 16462376 | 20744234 | 21366946 | 23169676 | 17319896 | 13133044 | 13109412 | 8640406 | 14279246 | 15695291 | 11719915 | 10152884 | 12657407 | 15273007 | 10374512 | 13551380 | 0.940806633 | 11 | 0.623487133 | 0 | 0.609384143 | 0 |
| P62959 | Hint1 | Adenosine 5'-monophosphoramidase HINT1 | 100.33 | 18.25 | 13777 | 2 | 5476761 | 9991302 | 6193847.5 | 5337748 | 15595922 | 5710656 | 5679682.5 | 8666787 | 7546599.5 | 12919569 | 18494190 | 21858152 | 18282924 | 17580766 | 18741080 | 27873396 | 24914500 | 24222898 | 33652446 | 20117758 | 0.95134973 | 11 | 2.229271462 | 0 | 3.070295004 | 0 |
| P11835 | Itgb2 | Integrin beta-2 | 1369.22 | 34.63 | 85026 | 25 | 34598550 | 41012110 | 30867802 | 29820060 | 28389790 | 28077724 | 22246126 | 40234966 | 50593506 | 30388658 | 21049082 | 19726968 | 16605289 | 20482824 | 12470584 | 19564462 | 22706786 | 22723816 | 21980834 | 13978626 | 1.041609923 | 11 | 0.548519478 | 0 | 0.613003575 | 0 |
| P62944 | Ap2b1 | AP-2 complex subunit beta | 466.89 | 10.99 | 104553 | 9 | 2411310.25 | 2033997.75 | 1746355.12 | 3509958.5 | 4102000 | 2337627.25 | 2286224.75 | 4070818.25 | 2912285 | 3302223 | 1602881.5 | 1542106.75 | 1089845.88 | 1233824.88 | 2 | 1537738.25 | 2 | 2 | 1063423.5 | 2 | 1.08009178 | 11 | 0.396175812 | 0 | 0.188440963 | 0 |
| P38060 | Hmgcl | Hydroxymethylglutaryl-CoA lyase, mitochondrial | 490.55 | 35.08 | 34239 | 8 | 2713818.75 | 4619631 | 4973925 | 3492790 | 5131860 | 4572965.5 | 4326210.5 | 4349983 | 6684221 | 4779825.5 | 10180738 | 10605768 | 10540576 | 11068789 | 8515280 | 10566430 | 8757616 | 9084900 | 7901771 | 7933410 | 1.180640946 | 11 | 2.432213396 | 0 | 2.113705078 | 0 |
| Q7M0E3 | Dstn | Destrin | 264.71 | 37.58 | 18534 | 5 | 4196600.5 | 3060336.25 | 4590758 | 4310259 | 4519185 | 7666249 | 4288128 | 6777347 | 3951379 | 3282883.75 | 18839586 | 14956345 | 20056462 | 15165702 | 17590094 | 21653850 | 26320282 | 22759698 | 23829482 | 20930826 | 1.255782392 | 11 | 4.188596403 | 0 | 5.585595734 | 0 |
| Q923D2 | Blvrb | Flavin reductase (NADPH) | 368.46 | 42.72 | 22197 | 5 | 2 | 744125.31 | 74026.41 | 916532.38 | 909614.75 | 2 | 126051.77 | 276664 | 666382.75 | 1300019.12 | 4655565 | 3433830.25 | 5021616.5 | 4789113 | 3993346.25 | 11786431 | 6163093 | 4988602 | 9568331 | 2956711 | 0.895934228 | 11.5 | 8.279493235 | 0 | 13.41116991 | 0 |
| Q8BK64 | Ahsa1 | Activator of 90 kDa heat shock protein ATPase homolog 1 | 103.71 | 7.1 | 38117 | 2 | 2098269.5 | 3661412.5 | 2899471.75 | 4391072.5 | 3526008.75 | 4583782.5 | 3879925 | 1454635.25 | 4062477.5 | 458614.44 | 6156676 | 6984405 | 7003834.5 | 7022895 | 5911192 | 8303389.5 | 9815597 | 11243295 | 7981015 | 10530410 | 0.871092542 | 12 | 1.995567902 | 0 | 2.888092893 | 0 |
| P08556 | Nras | GTPase NRas | 159.74 | 18.52 | 21199 | 3 | 1598634.25 | 920045.88 | 2732995.5 | 1056122.25 | 2307792.5 | 1264861.12 | 1399229.38 | 229032.36 | 2440603 | 2457809 | 6250669 | 6986914.5 | 7202741 | 7389468 | 4437622 | 7290779.5 | 7415979 | 4871740 | 10011688 | 5187667.5 | 0.904352983 | 12 | 3.745235448 | 0 | 4.036618788 | 0 |
| Q05816 | Fabp5 | Fatty acid-binding protein 5 | 670.01 | 64.44 | 15137 | 10 | 34245050 | 34500042 | 50742414 | 9556888 | 10992836 | 39574022 | 37961902 | 48729858 | 5096551 | 6893748 | 79571738 | 91973610 | 85874338 | 68784394 | 83715738 | 54366678 | 84873338 | 70096202 | 73705058 | 54591358 | 0.98728089 | 12 | 2.927220268 | 0 | 2.411020512 | 0 |
| B5DF91 | Elavl1 | ELAV-like protein 1 | 488.16 | 27.3 | 36169 | 9 | 21195206 | 24887008 | 19642276 | 26699298 | 31745138 | 24547870 | 24290250 | 33821718 | 21071814 | 21814548 | 16274388 | 15319762 | 13510757 | 9994121 | 15633048 | 9066060 | 11128088 | 10306440 | 9135048 | 9834043 | 1.011091938 | 12 | 0.569643938 | 0 | 0.398406273 | 0 |
| Q99J93 | Ifitm2 | Interferon-induced transmembrane protein 2 | 252.04 | 17.36 | 15743 | 3 | 3165886 | 3724648.5 | 2567999.25 | 1737459.12 | 1044223.56 | 3241124 | 2254297.75 | 4923385 | 1606223.25 | 626365.62 | 2 | 2 | 511821.19 | 438021.78 | 2 | 2 | 2 | 789722.88 | 916156.06 | 2 | 1.033592477 | 12 | 0.077600668 | 0 | 0.13936722 | 0 |
| P62962 | Pfn1 | Profilin-1 | 804.22 | 90 | 14957 | 13 | 79252314 | 32768326 | 86784618 | 57409862 | 115533490 | 55482106 | 54235882 | 56272182 | 119043978 | 132632346 | 215684194 | 222990082 | 254290322 | 243739762 | 166206930 | 180318818 | 192734306 | 169812578 | 300606722 | 163569986 | 1.123518643 | 12 | 2.966820212 | 0 | 2.708933895 | 0 |
| Q9CQW9 | Ifitm3 | Interferon-induced transmembrane protein 3 | 478.05 | 48.91 | 14954 | 5 | 6990885 | 7555690.5 | 6112185.5 | 5423869 | 5122507 | 8250985 | 5130915.5 | 17265762 | 5649892.5 | 3963981.5 | 2 | 2 | 1380408.75 | 463861.12 | 807713.25 | 2014485.25 | 2664617.75 | 5119891.5 | 4826195 | 2118100.5 | 1.290221431 | 12 | 0.084985595 | 0 | 0.536555568 | 0 |
| P01900 | H2-D1 | H-2 class I histocompatibility antigen, D-D alpha chain | 747.34 | 41.92 | 41110 | 14 | 2 | 2996176 | 204206.59 | 2679340.5 | 2912626.5 | 2149102.5 | 1246608.75 | 2651931.75 | 5414471 | 1932304.5 | 13955162 | 8535286 | 5678662 | 5773805 | 5734027.5 | 5464953.5 | 5375568 | 3252325 | 6184366 | 3924042.75 | 1.52341707 | 12 | 4.512665593 | 0 | 2.752534973 | 0 |
| O35143 | Atp5if1 | ATPase inhibitor, mitochondrial | 56.36 | 8.49 | 12159 | 1 | 2 | 2 | 2 | 638228.69 | 1262100 | 921967.94 | 2 | 2 | 877908.44 | 2 | 2098631 | 3314753.75 | 1882607 | 1902794.12 | 1274086.25 | 5901235.5 | 3008104.5 | 1422321.12 | 4440892 | 1494093.12 | 0.947139675 | 12.5 | 5.511067169 | 0 | 8.559884912 | 0 |
| P24547 | Impdh2 | Inosine-5'-monophosphate dehydrogenase 2 | 823.83 | 26.26 | 55815 | 13 | 17446900 | 16179832 | 17789444 | 14903950 | 28514204 | 5449529 | 6618694.5 | 10749625 | 5516218.5 | 10050361 | 15306482 | 14230199 | 14013454 | 13951523 | 12153343 | 10350058 | 6980839.5 | 6417638.5 | 5090295 | 4867009 | 0.404752456 | 0 | 0.734491413 | 1 | 0.355418128 | 0 |
| P09103 | P4hb | Protein disulfide-isomerase | 1679.4 | 52.65 | 57058 | 29 | 50577538 | 49967894 | 54359506 | 57374046 | 68329114 | 88081394 | 96825682 | 97847258 | 109065314 | 101177146 | 89166498 | 71966010 | 92112898 | 83260802 | 61708542 | 103315386 | 107427386 | 121451682 | 129311530 | 95977914 | 1.756887266 | 0 | 1.419113535 | 1 | 1.986699251 | 0 |
| Q61543 | Glg1 | Golgi apparatus protein 1 | 761.99 | 15.23 | 133734 | 13 | 11023077 | 10381269 | 9165269 | 8765269 | 4318496 | 2416385 | 2097386 | 3033592.5 | 4057770.5 | 5895341.5 | 1991179 | 4741745 | 2408291.25 | 1255900.25 | 2680965.5 | 2015167.5 | 2383715.75 | 3073398.75 | 1749435.12 | 1896130.75 | 0.400896231 | 1 | 0.299589196 | 1 | 0.254684697 | 0 |
| Q9D0F6 | Rfc5 | Replication factor C subunit 5 | 179.43 | 15.34 | 38096 | 4 | 2342978 | 3776439.75 | 1071435 | 2612855 | 3507526.5 | 747363 | 773547.12 | 112331.38 | 1429829.5 | 1199849.25 | 965527.94 | 843725.25 | 788172.38 | 693701.62 | 1691632.62 | 2 | 2 | 783033.06 | 2 | 2 | 0.320249811 | 2 | 0.374327408 | 1 | 0.058825579 | 0 |
| Q5U2U0 | Clpx | ATP-dependent clpX-like chaperone, mitochondrial | 107.87 | 5.21 | 69208 | 3 | 1263421.62 | 1593959.12 | 1282999.88 | 918913.88 | 1201790.62 | 2 | 1241799.25 | 2 | 611453.06 | 726342.38 | 2 | 811907.25 | 1098744.25 | 893415.31 | 2 | 659967.94 | 2 | 2 | 699500.69 | 2 | 0.41200505 | 2 | 0.447857002 | 1 | 0.217130833 | 0 |
| P70168 | Kpnb1 | Importin subunit beta-1 | 613.98 | 15.18 | 97184 | 11 | 18024284 | 18542902 | 18828454 | 20989976 | 22986042 | 12895281 | 14158646 | 18709666 | 12434120 | 17518644 | 13247630 | 12490846 | 14452863 | 14233406 | 18420588 | 11314284 | 9490507 | 12631413 | 11150694 | 13587731 | 0.76195123 | 2 | 0.73305945 | 1 | 0.58542476 | 0 |
| Q99L45 | Eif2s2 | Eukaryotic translation initiation factor 2 subunit 2 | 244.87 | 8.46 | 38092 | 5 | 4167082.5 | 4643582 | 2449438 | 6060031.5 | 7350135 | 8797467 | 9735414 | 14109189 | 12231153 | 5757658.5 | 11731560 | 9318759 | 10532087 | 9323444 | 6594662.5 | 10411003 | 9204718 | 9059762 | 13275035 | 8546372 | 2.052303585 | 2 | 1.925415264 | 1 | 2.04687229 | 0 |
| P20491 | Fcer1g | High affinity immunoglobulin epsilon receptor subunit gamma | 153.72 | 23.26 | 9652 | 3 | 2 | 931609.12 | 2 | 1123922.88 | 2157792.5 | 3271061.5 | 1921324.88 | 3008396.5 | 5067920.5 | 1606688.75 | 4555421 | 5484753 | 4560800.5 | 4709836 | 2091373.12 | 4432980 | 3293572.75 | 2238744.25 | 4874449 | 2163969.5 | 3.53055598 | 2 | 5.079638015 | 1 | 4.035696599 | 0 |
| Q8BLN5 | Lss | Lanosterol synthase | 209.45 | 6.41 | 83141 | 4 | 2 | 2 | 103244.32 | 467896.16 | 2 | 827457.12 | 665304 | 95375.59 | 759424.19 | 421871.88 | 215322.97 | 940441.75 | 1142981.5 | 1329781.75 | 1807262.88 | 1230484.88 | 1394468.5 | 1532039.12 | 1531865.88 | 1616098.25 | 4.848901074 | 3 | 9.517332314 | 1 | 12.78998801 | 0 |
| Q3UHX2 | Pdap1 | 28 kDa heat- and acid-stable phosphoprotein | 236.36 | 13.81 | 20605 | 4 | 1219113.25 | 2301653.5 | 1015344.62 | 1045551 | 1038081 | 665262.56 | 480949.59 | 2 | 670040.06 | 1550873 | 3009455.75 | 3891707 | 3327132.25 | 2650756.75 | 2129460.25 | 7038349 | 3477568 | 2639276.75 | 6319377.5 | 5136619.5 | 0.508649206 | 4 | 2.267234719 | 1 | 3.717846656 | 0 |
| P25911 | Lyn | Tyrosine-protein kinase Lyn | 553.23 | 24.02 | 58812 | 9 | 7142423.5 | 6123755.5 | 6448264 | 4608900.5 | 6531477.5 | 4185935.5 | 4722080 | 2879500.5 | 5550315.5 | 6349535.5 | 3081475.25 | 3749627.5 | 5235096 | 4034160 | 3497931 | 3521082.5 | 2951183 | 4583672.5 | 3326545 | 665942 | 0.767703919 | 4 | 0.635177555 | 1 | 0.487717138 | 0 |
| P05063 | Aldoc | Fructose-bisphosphate aldolase C | 1252.38 | 61.16 | 39395 | 21 | 13274122 | 21123936 | 14549677 | 11569550 | 24268408 | 22008330 | 24583246 | 24554790 | 23588642 | 18820232 | 38415966 | 41127062 | 34667174 | 38105294 | 21977542 | 29246642 | 36152190 | 26580400 | 31875758 | 25395772 | 1.339320774 | 4 | 2.055689254 | 1 | 1.760329564 | 0 |
| Q9D6Z1 | Nop56 | Nucleolar protein 56 | 803.37 | 26.03 | 64464 | 14 | 2101565.5 | 4271047.5 | 7835113 | 2829558.5 | 5662746 | 5811211.5 | 6250350 | 7556915 | 8255882 | 6105906 | 7831085 | 8060398.5 | 10093548 | 10671801 | 12949663 | 11062607 | 8238627.5 | 15962203 | 12282738 | 10925965 | 1.49692594 | 4 | 2.185305236 | 1 | 2.575861759 | 0 |
| Q8BTM8 | Flna | Filamin-A | 4042.23 | 33.36 | 281222 | 69 | 67233162 | 91182226 | 83583186 | 30546066 | 42653442 | 105562506 | 105359938 | 160628706 | 82822562 | 75044834 | 128851098 | 143336658 | 99317506 | 108401874 | 90491330 | 174092962 | 126538906 | 102280722 | 133019034 | 118139626 | 1.679637587 | 4 | 1.809650815 | 1 | 2.075111771 | 0 |
| Q5U2Q7 | Etf1 | Eukaryotic peptide chain release factor subunit 1 | 227.15 | 11.44 | 49031 | 4 | 3757929 | 3953695.25 | 5297127.5 | 4717134.5 | 6760934.5 | 2989044 | 4771884.5 | 3185173.5 | 3824166 | 3867261.25 | 2439071.25 | 3950169.5 | 3343839.75 | 2787527.75 | 3521739 | 1511522.75 | 2352325.75 | 2378620.5 | 977832.44 | 2471458 | 0.76112491 | 5 | 0.655142103 | 1 | 0.395794927 | 0 |
| Q9DC16 | Ergic1 | Endoplasmic reticulum-Golgi intermediate compartment protein 1 | 42.45 | 3.45 | 32562 | 1 | 697680.62 | 671538.06 | 411739.53 | 550431.94 | 794284.56 | 1662543.5 | 1299834.62 | 3129918.25 | 888176 | 2 | 1210114 | 1181084.38 | 1222404.12 | 940654.5 | 734632.19 | 1225969 | 950859.62 | 940519.69 | 961146.62 | 1002520.81 | 2.233269619 | 5 | 1.692079209 | 1 | 1.625574064 | 0 |
| D3Z6Q9 | Bin2 | Bridging integrator 2 | 251.4 | 10.22 | 52554 | 4 | 2095120.12 | 930018.38 | 2 | 1553352.75 | 2885104 | 2384462.5 | 2110697.5 | 1218321.75 | 3162429.75 | 1208974.38 | 4481338 | 5000398.5 | 3642050 | 4737144 | 2269586.5 | 4356563 | 4434156 | 3362007 | 4922474.5 | 3540454.75 | 1.351209818 | 8 | 2.69716014 | 1 | 2.762160733 | 0 |
| Q9DBL1 | Acadsb | Short/branched chain specific acyl-CoA dehydrogenase, mitochondrial | 207.01 | 11.57 | 47874 | 4 | 7018289.5 | 14215144 | 9009966 | 9093174 | 6199821 | 8367582 | 7702696.5 | 8207247 | 6950683 | 7949475 | 5699357 | 3853466 | 4785689.5 | 6292184.5 | 3182772.5 | 4583921.5 | 3738578.75 | 3548650.25 | 5525142 | 3196016.25 | 0.860359805 | 9 | 0.522954656 | 1 | 0.452216496 | 0 |
| Q6NSR8 | Npepl1 | Probable aminopeptidase NPEPL1 | 67.5 | 2.29 | 55940 | 1 | 2 | 2 | 890333.69 | 317294.38 | 486473.44 | 1083593.38 | 619240.38 | 63026.04 | 2 | 700344 | 1203304.5 | 1178328.88 | 1224975.25 | 1205976.75 | 850227.94 | 1212360.88 | 1614224.62 | 913143.62 | 1589625.5 | 1498088.12 | 1.455756909 | 9 | 3.34265681 | 1 | 4.030116601 | 0 |
| P60060 | Sec61g | Protein transport protein Sec61 subunit gamma | 114.04 | 29.41 | 7741 | 2 | 2145020.25 | 4049843.75 | 2268142.75 | 2257530.5 | 2828570.25 | 2963437 | 2644186 | 1561022.62 | 3605219 | 3012158.5 | 1474205 | 1136984.62 | 802669.62 | 2148376.5 | 2 | 1142242.38 | 945471.56 | 819531.19 | 1675090.5 | 2 | 1.017485699 | 10 | 0.41052429 | 1 | 0.338202175 | 0 |
| Q62433 | Ndrg1 | Protein NDRG1 | 431.7 | 21.32 | 43009 | 6 | 2195349.25 | 3200801.75 | 4223435.5 | 1512579.75 | 4214666.5 | 7044810 | 5368003.5 | 7987352.5 | 5291346 | 7852961.5 | 12203046 | 9599240 | 7615808 | 4357332 | 4164202 | 6741291.5 | 5685017 | 6320821 | 7349623 | 7718850 | 2.185758719 | 0 | 2.472147095 | 2 | 2.203425492 | 0 |
| Q61881 | Mcm7 | DNA replication licensing factor MCM7 | 554.39 | 18.5 | 81211 | 11 | 10809012 | 8981728 | 7440702.5 | 17969792 | 20951174 | 4286497.5 | 6001006 | 7453437 | 7134752 | 6537113 | 4556683 | 8747308 | 6959521 | 8936952 | 5406111 | 3390765 | 5366053.5 | 5655188 | 3016416 | 6699942 | 0.474855054 | 1 | 0.523134014 | 2 | 0.364739018 | 0 |
| Q99P72 | Rtn4 | Reticulon-4 | 417.91 | 11.62 | 126613 | 7 | 3767054 | 7311594.5 | 2678970.5 | 10315450 | 17416918 | 14256043 | 11528687 | 21806970 | 19712886 | 11568924 | 21624424 | 15859654 | 23092026 | 26072818 | 13472121 | 22373326 | 25098900 | 23650134 | 35763274 | 21176258 | 1.90102518 | 3 | 2.413137488 | 2 | 3.086573442 | 0 |
| Q6ZQM8 | Ugt1a7 | UDP-glucuronosyltransferase 1A7 | 502.74 | 24.48 | 59758 | 9 | 10394196 | 10147638 | 8692001 | 10775481 | 8564512 | 4752067 | 4893916 | 1766270.62 | 8216869 | 12364601 | 4064935.5 | 4473645 | 6666035.5 | 7599017.5 | 9448881 | 4967041 | 7571622.5 | 8055529.5 | 5561700 | 6093440.5 | 0.658661772 | 5 | 0.663989556 | 2 | 0.663924068 | 0 |
| O55143 | Atp2a2 | Sarcoplasmic/endoplasmic reticulum calcium ATPase 2 | 1427.38 | 30.65 | 114858 | 25 | 10826347 | 18011260 | 9000194 | 10509625 | 14556440 | 17229276 | 18068524 | 16416002 | 16608979 | 13576443 | 16757226 | 21833882 | 20523892 | 21679550 | 16254305 | 18096568 | 18989466 | 21219398 | 18596494 | 18617870 | 1.301974413 | 5 | 1.542812249 | 2 | 1.518504379 | 0 |
| Q9WV85 | Nme3 | Nucleoside diphosphate kinase 3 | 41.05 | 10.06 | 19099 | 1 | 2 | 726311.38 | 678940.88 | 2 | 824940.31 | 2 | 2 | 583023.88 | 2 | 2 | 1487507.75 | 772405.56 | 824459.88 | 883563.06 | 1122998.38 | 826387.12 | 912780.06 | 1063492.5 | 939747.44 | 1005704.38 | 0.261426229 | 6 | 2.282729109 | 2 | 2.129010314 | 0 |
| Q9DC70 | Ndufs7 | NADH dehydrogenase [ubiquinone] iron-sulfur protein 7, mitochondrial | 150.43 | 10.27 | 24683 | 2 | 2689957.75 | 7029256 | 3418484.75 | 6247232 | 3656251.75 | 7954897 | 6373283 | 3911064.75 | 6434464 | 4593360.5 | 3294419 | 2769830.75 | 2286672.75 | 1853332.5 | 1060118.62 | 2335182 | 1106725.38 | 1030336.56 | 1192718.25 | 2 | 1.270206925 | 6 | 0.488880019 | 2 | 0.245862566 | 0 |
| P40142 | Tkt | Transketolase | 1674.24 | 49.44 | 67630 | 29 | 39039022 | 78418546 | 33548210 | 33800890 | 89102226 | 84618066 | 88700098 | 55029882 | 110023210 | 53448130 | 163516018 | 182412226 | 126912786 | 138631762 | 77841306 | 187722562 | 143701730 | 129989162 | 151493458 | 135058242 | 1.430473397 | 6 | 2.516581656 | 2 | 2.730707802 | 0 |
| Q91ZR2 | Snx18 | Sorting nexin-18 | 120.91 | 4.72 | 67904 | 2 | 2 | 2 | 2 | 2 | 1253223.88 | 2 | 615557.31 | 214317.56 | 1445289 | 721114.69 | 1416148.75 | 931362.19 | 1515051.75 | 1376309.5 | 864104 | 2277401 | 1845058.12 | 2413709.5 | 1883922.62 | 2676502.5 | 2.390842914 | 6 | 4.86979009 | 2 | 8.854381952 | 0 |
| O35114 | Scarb2 | Lysosome membrane protein 2 | 72.38 | 4.18 | 54044 | 2 | 697377.25 | 435945.12 | 1416234.62 | 1143811.12 | 1404471.75 | 2124690 | 2 | 1562534.12 | 1700615.25 | 1205912.12 | 2 | 658694.75 | 2 | 2 | 518552.69 | 2 | 2 | 2 | 2 | 2 | 1.293440687 | 7 | 0.230931821 | 2 | 1.96162E-06 | 0 |
| Q69ZN7 | Myof | Myoferlin | 581.43 | 6.69 | 233324 | 11 | 5031534 | 4741635 | 2464479.75 | 1675938.75 | 1839702.75 | 4447420 | 4947511 | 7072817 | 3494810 | 4579746.5 | 9010014 | 9206379 | 6817130 | 9112358 | 2766071.5 | 12003277 | 11177555 | 8210976 | 11820012 | 8655194 | 1.557916099 | 7 | 2.343126541 | 2 | 3.292455936 | 0 |
| Q9CWZ7 | Napg | Gamma-soluble NSF attachment protein | 92.35 | 4.49 | 34732 | 2 | 2662191 | 1130505.62 | 2707062.25 | 2567666 | 2077869.75 | 889388.75 | 732696 | 935287.88 | 2573530.25 | 2897854.75 | 161494.03 | 227488.69 | 1818851.38 | 1402031.38 | 910754.56 | 2 | 472307.56 | 268797.47 | 249535.89 | 470370.53 | 0.720371951 | 8 | 0.405607944 | 2 | 0.131087916 | 0 |
| O55222 | Ilk | Scaffold protein ILK | 72.09 | 4.2 | 51373 | 2 | 1704309.75 | 3210618.75 | 1674992.75 | 2475245 | 2593341 | 2130170 | 2098161.75 | 1719687.12 | 2285113 | 1508791.5 | 1634618.75 | 610386.81 | 1484026.25 | 2035228.25 | 1130897.38 | 680246.62 | 1417336.75 | 1635344.25 | 727841.44 | 855926.44 | 0.835606408 | 8 | 0.591427126 | 2 | 0.456035699 | 0 |
| Q9EQP2 | Ehd4 | EH domain-containing protein 4 | 397.99 | 15.53 | 61481 | 8 | 3857906.5 | 4967847 | 5644242 | 3550824.5 | 10134876 | 4227047 | 7004378.5 | 4080913.5 | 7819204.5 | 11722443 | 943233.81 | 2586093 | 2960140.75 | 4725046 | 3202818.5 | 894422.75 | 1547067.75 | 2744305.75 | 544961.44 | 2734872.75 | 1.237901791 | 8 | 0.512057385 | 2 | 0.300672036 | 0 |
| Q8BMK4 | Ckap4 | Cytoskeleton-associated protein 4 | 897.48 | 30.26 | 63692 | 14 | 2045079.12 | 1276170.88 | 2805130.25 | 4041542.75 | 4977199 | 183101.31 | 3188411.25 | 4098052 | 4076896.75 | 7389311.5 | 391244.62 | 1179653.5 | 1516944 | 979984.5 | 1517138.38 | 2 | 2 | 1215795.38 | 2 | 753378.94 | 1.250288562 | 9 | 0.368763289 | 2 | 0.130020763 | 0 |
| Q3TEA8 | Hp1bp3 | Heterochromatin protein 1-binding protein 3 | 695.71 | 26.9 | 60867 | 14 | 7017691 | 8743677 | 4710205 | 8029991 | 12240141 | 5102715 | 8237344.5 | 3382355.5 | 9300695 | 7529281 | 1809590.75 | 5209964.5 | 4447484 | 3023248 | 5152734.5 | 3807166 | 2187949.75 | 1009430.62 | 4658403 | 1720478.5 | 0.823539197 | 10 | 0.482135486 | 2 | 0.328494546 | 0 |
| O35226 | Psmd4 | 26S proteasome non-ATPase regulatory subunit 4 | 93.04 | 5.32 | 40704 | 2 | 2779402 | 3644293 | 2304349.5 | 2659288 | 3656623 | 4395527.5 | 2152778.5 | 1369992 | 3663422.5 | 3939181 | 4163676 | 4227300 | 5405960 | 5261857 | 3322323.75 | 3689395.5 | 5902679 | 4678242 | 7824437 | 5675494.5 | 1.031703497 | 10 | 1.4877149 | 2 | 1.845940584 | 0 |
| Q9CQX2 | Cyb5b | Cytochrome b5 type B | 192.13 | 15.75 | 16318 | 2 | 915671.38 | 718192.81 | 1818190 | 585069.75 | 2019306 | 586687 | 2 | 1921275.25 | 1870178.75 | 2225749.75 | 3277430.5 | 1616061.88 | 2948501 | 2703744.5 | 3447611.5 | 2964692 | 2790975 | 4193625 | 4072544.5 | 3269958.75 | 1.09039365 | 11 | 2.310494717 | 2 | 2.855113561 | 0 |
| P24049 | Rpl17 | Large ribosomal subunit protein uL22 | 218.27 | 24.46 | 21397 | 5 | 4209144.5 | 4409993 | 2911246 | 5371977.5 | 4534420 | 4361944 | 3610781 | 7782272 | 8319668 | 425102.09 | 9014267 | 8900701 | 4902082 | 6392799 | 4662976 | 9602756 | 5680413.5 | 8723828 | 8492576 | 5753117.5 | 1.14288461 | 12 | 1.580126466 | 2 | 1.784441937 | 0 |
| Q9JLT4 | Txnrd2 | Thioredoxin reductase 2, mitochondrial | 167 | 5.15 | 56603 | 2 | 2 | 2 | 2 | 2 | 2 | 2 | 2 | 2 | 642741.81 | 626966.5 | 1419506 | 2 | 1071698.25 | 565998.94 | 930757.31 | 1093187.25 | 1109779 | 1460175.75 | 1148533 | 918230.06 | 126971.431 | 7.5 | 398796.25 | 2.5 | 572990.506 | 0 |
| P43247 | Msh2 | DNA mismatch repair protein Msh2 | 55.58 | 1.6 | 104151 | 1 | 1027629 | 256094.42 | 800181.19 | 607052.94 | 569577.81 | 183061.23 | 388186.94 | 66306.38 | 2 | 344784.75 | 207721.11 | 2 | 499754.47 | 482945.34 | 560570.19 | 148123.75 | 2 | 2 | 2 | 2 | 0.30128221 | 2 | 0.53702626 | 3 | 0.045431726 | 0 |
| P09895 | Rpl5 | Large ribosomal subunit protein uL18 | 527.93 | 34.68 | 34459 | 9 | 11769742 | 15005483 | 13847691 | 18785062 | 18236976 | 10841014 | 9251037 | 10807467 | 12444052 | 12230678 | 20817738 | 23253880 | 25268382 | 25805484 | 14022350 | 19914126 | 21938106 | 23360344 | 25767374 | 21353190 | 0.715748354 | 2 | 1.40598749 | 3 | 1.4467539 | 0 |
| O35609 | Scamp3 | Secretory carrier-associated membrane protein 3 | 164.37 | 9.17 | 38458 | 2 | 2 | 700288.12 | 536191.88 | 2 | 2135326.5 | 2245838 | 1742991.25 | 3344586 | 2932386.5 | 1106011.62 | 3824814.5 | 2740424.75 | 1387736 | 1261533.62 | 1759677.62 | 4125530.25 | 5068695.5 | 2886236.5 | 4290584 | 2732534 | 3.372613428 | 2 | 3.254686611 | 3 | 5.665674346 | 0 |
| Q9CWG1 | Glipr1 | Glioma pathogenesis-related protein 1 | 52.43 | 3.92 | 29129 | 1 | 794861.38 | 405147.88 | 615461.5 | 757752.38 | 645906.31 | 2 | 2 | 2 | 653752.5 | 373465.47 | 2 | 2 | 2 | 2 | 743013.88 | 2 | 2 | 2 | 2 | 2 | 0.319099926 | 3 | 0.230814539 | 3 | 3.10643E-06 | 0 |
| Q8C0C7 | Farsa | Phenylalanine--tRNA ligase alpha subunit | 215.64 | 7.68 | 57599 | 4 | 3600885.75 | 2811630.25 | 4019623.25 | 1439490.62 | 3243220 | 1472546 | 991522.06 | 3088438.5 | 1013014.31 | 3128988.75 | 1834197 | 535715.38 | 1501343.38 | 1303359.62 | 2087507.75 | 660146.69 | 360409.69 | 775595.56 | 821682.5 | 2 | 0.641389739 | 5 | 0.480462803 | 3 | 0.173196324 | 0 |
| O35987 | Nsfl1c | NSFL1 cofactor p47 | 266.55 | 16.22 | 40680 | 4 | 6355420 | 6909193.5 | 5941815 | 2 | 3606713 | 3227318.5 | 4036164 | 2835947.25 | 830570.81 | 2516012.5 | 9032443 | 8177956 | 7979113.5 | 6836448 | 6052626 | 11469342 | 9245829 | 7270071.5 | 15120555 | 9119804 | 0.589397645 | 6 | 1.669151229 | 3 | 2.289276859 | 0 |
| Q61207 | Psap | Prosaposin | 335.95 | 15.62 | 61422 | 7 | 58043146 | 28368834 | 64919290 | 31909894 | 36689478 | 27314566 | 28061566 | 44449598 | 23488012 | 56325506 | 22374538 | 21113362 | 22366956 | 18440558 | 39247638 | 19001262 | 27114490 | 12123280 | 23268838 | 28063704 | 0.816799544 | 6 | 0.561736422 | 3 | 0.498209676 | 0 |
| P50396 | Gdi1 | Rab GDP dissociation inhibitor alpha | 355.17 | 12.53 | 50522 | 5 | 1273546.62 | 1611396.38 | 2110220.75 | 1214193 | 1784088.62 | 2180317.25 | 1975787 | 2977043 | 1384478.5 | 1656203.88 | 2770104.25 | 4006699 | 3551606.5 | 1362352.38 | 2128957 | 2553054.25 | 3758165 | 3013912.25 | 4966340 | 2278581 | 1.272771522 | 6 | 1.728881413 | 3 | 2.072954994 | 0 |
| Q08013 | Ssr3 | Translocon-associated protein subunit gamma | 197.4 | 9.73 | 21064 | 2 | 4488552 | 2672782.5 | 4232913 | 2861596 | 3526890.25 | 2371337.75 | 2665232 | 1734542.12 | 3624972 | 4277002 | 1313364.5 | 1021446.44 | 2770101.25 | 2287337 | 2932324 | 1309015 | 1790701.38 | 667552.75 | 1947398.38 | 1667219.88 | 0.825131056 | 7 | 0.580595387 | 3 | 0.415115443 | 0 |
| P28656 | Nap1l1 | Nucleosome assembly protein 1-like 1 | 179.77 | 12.02 | 45345 | 3 | 1951933.5 | 295461.94 | 5139031.5 | 5490414 | 5357791.5 | 507028.94 | 2380314.75 | 765114.94 | 2709621 | 4008197.5 | 373985.09 | 2 | 1087174.38 | 1000198.5 | 2 | 2 | 2 | 2 | 2 | 2 | 0.568713253 | 8 | 0.134982812 | 3 | 5.48407E-07 | 0 |
| P00375 | Dhfr | Dihydrofolate reductase | 41.6 | 3.74 | 21606 | 1 | 2724639 | 2976084.5 | 4816422.5 | 2806103.25 | 3378183 | 2353422.75 | 2690691 | 3733866.25 | 423279.97 | 3794531 | 646227.12 | 433386.78 | 2325262.5 | 2526174.25 | 3332033.25 | 2080717.5 | 1921578.5 | 1766163.75 | 1673888.62 | 2254035 | 0.778124341 | 8 | 0.554628116 | 3 | 0.580571967 | 0 |
| P60843 | Eif4a1 | Eukaryotic initiation factor 4A-I | 1184.98 | 53.94 | 46154 | 18 | 3275947 | 2629167.75 | 6765980 | 3103604.5 | 5576520 | 2458810.5 | 3121775 | 4032822.75 | 1917453.5 | 5094446.5 | 2450592.75 | 1340812.12 | 2173055 | 1289845 | 5445591.5 | 1534033.75 | 1091110.25 | 1629938 | 1270441.25 | 2532110.5 | 0.778658495 | 8 | 0.594808953 | 3 | 0.377385181 | 0 |
| Q7TMF3 | Ndufa12 | NADH dehydrogenase [ubiquinone] 1 alpha subcomplex subunit 12 | 54.8 | 6.21 | 17086 | 1 | 1292617 | 439161.16 | 1355120 | 819242.31 | 1277067.88 | 276542.47 | 1210296.38 | 315515.75 | 1354448 | 1262708.12 | 2 | 606499 | 729440.44 | 684910.94 | 2 | 2 | 2 | 2 | 2 | 2 | 0.852659284 | 8 | 0.389884844 | 3 | 1.92931E-06 | 0 |
| P16675 | Ctsa | Lysosomal protective protein | 900.07 | 30.17 | 53844 | 14 | 60496610 | 63021898 | 65243974 | 33514098 | 53683686 | 45084330 | 43242146 | 54966158 | 45749890 | 60867682 | 36576238 | 32424966 | 35394314 | 34484194 | 31973214 | 22223014 | 20966140 | 27274796 | 27781034 | 23042262 | 0.905602135 | 8 | 0.619121472 | 3 | 0.439509817 | 0 |
| P62307 | Snrpf | Small nuclear ribonucleoprotein F | 44.57 | 9.3 | 9725 | 1 | 2 | 2 | 2 | 2 | 880130.06 | 2 | 1243931.5 | 2 | 2 | 1090318.5 | 2615360.25 | 1986050.62 | 2 | 3129813 | 1380557.25 | 3092583 | 2108511.5 | 2667458.75 | 3998551.5 | 1492754 | 2.652147551 | 9 | 10.35267481 | 3 | 15.17927625 | 0 |
| Q6A0D4 | Rftn1 | Raftlin | 102.33 | 4.51 | 61537 | 2 | 2 | 413806.56 | 2 | 2 | 2 | 965515.75 | 2 | 125259.69 | 2 | 2 | 967382.81 | 1461391.88 | 1106706.88 | 2194061.25 | 2 | 2277511.75 | 2386748 | 1955327.62 | 460469.06 | 1767900 | 2.635918466 | 10 | 13.84568204 | 3 | 21.38145267 | 0 |
| P18052 | Ptpra | Receptor-type tyrosine-protein phosphatase alpha | 170.25 | 5.79 | 93698 | 4 | 1966286.25 | 846486 | 447440.03 | 920831.25 | 1626633.12 | 367568.28 | 293277.81 | 1586963.12 | 2068535.88 | 1231232.5 | 2 | 628979.56 | 733117.5 | 135862.77 | 612209.25 | 219235.81 | 2 | 2 | 2 | 2 | 0.955214611 | 11 | 0.363341695 | 3 | 0.037750692 | 0 |
| Q9JM76 | Arpc3 | Actin-related protein 2/3 complex subunit 3 | 240.64 | 38.2 | 20525 | 5 | 36901054 | 19186378 | 35692566 | 31604164 | 26978390 | 21037770 | 32416528 | 30253432 | 37171606 | 33818910 | 14707474 | 20872154 | 14221682 | 16664045 | 31327250 | 10355558 | 12206244 | 13244689 | 5859636.5 | 10752709 | 1.028834932 | 11 | 0.650378726 | 3 | 0.3486163 | 0 |
| Q61990 | Pcbp2 | Poly(rC)-binding protein 2 | 645.14 | 33.98 | 38222 | 10 | 8579163 | 10682569 | 7399833.5 | 14527761 | 9665021 | 11758707 | 8830597 | 13741613 | 9730234 | 6417999 | 16856112 | 13912013 | 11695288 | 10930077 | 14838982 | 16321079 | 18842326 | 18154494 | 21377176 | 20113088 | 0.992622116 | 12 | 1.341723478 | 3 | 1.864307924 | 0 |
| Q9WTP7 | Ak3 | GTP:AMP phosphotransferase AK3, mitochondrial | 98.74 | 18.5 | 25426 | 2 | 521844.66 | 613491.19 | 297366.12 | 1226404.38 | 968161.75 | 2 | 150859.41 | 1372263.5 | 885752.75 | 1542871.75 | 779254.56 | 2 | 227683.58 | 179257.98 | 2 | 2 | 2 | 2 | 2 | 2 | 1.089456114 | 12 | 0.327023007 | 3 | 2.7569E-06 | 0 |
| Q78JE5 | Fbxo22 | F-box only protein 22 | 56.19 | 2.24 | 44203 | 1 | 513828.25 | 2 | 2 | 2 | 510410.97 | 670892.31 | 655319.5 | 480013.31 | 1239180.25 | 457518.94 | 923267.75 | 747837.06 | 969258.12 | 2 | 821296.38 | 1092166.12 | 1689552.25 | 1506619.62 | 1130237.62 | 1725798.38 | 3.420005524 | 4 | 3.379719273 | 3.5 | 6.975257341 | 0 |
| P59017 | Bcl2l13 | Bcl-2-like protein 13 | 141.13 | 12.21 | 46719 | 3 | 2 | 529414.06 | 2 | 2 | 55342.47 | 2 | 396319.91 | 2 | 1066284.88 | 948859.44 | 1103659.12 | 1323844.88 | 1365154.75 | 1207360.25 | 2 | 1207189.12 | 1128477.12 | 860885.56 | 1117890.12 | 1163538.75 | 4.123841912 | 8 | 8.550515369 | 3.5 | 9.367872237 | 0 |
| Q8R050 | Gspt1 | Eukaryotic peptide chain release factor GTP-binding subunit ERF3A | 177.63 | 8.33 | 68625 | 4 | 361570.44 | 477075.25 | 2 | 2 | 2 | 2 | 2 | 2 | 744064.19 | 321261.47 | 3959867.25 | 4096682.75 | 3440485.5 | 3137756.5 | 2 | 4266507.5 | 2988867.25 | 2672725.75 | 540514.06 | 1097766.38 | 1.27029096 | 12.5 | 17.45038396 | 3.5 | 13.7916385 | 0 |
| P48758 | Cbr1 | Carbonyl reductase [NADPH] 1 | 164.41 | 9.03 | 30641 | 3 | 2441443.5 | 1501072.25 | 1516362 | 2428667.5 | 1450550 | 3856318 | 3577368 | 4228679 | 3718460.75 | 1998752.25 | 3401113.5 | 3490770.75 | 1662674.75 | 1962451.5 | 2738676 | 4345914 | 5630500 | 3725519 | 3257181.5 | 9048481 | 1.861148075 | 2 | 1.419527874 | 4 | 2.785107113 | 0 |
| O08528 | Hk2 | Hexokinase-2 | 717.61 | 18.76 | 102535 | 15 | 512493.81 | 265491.81 | 783772.19 | 522957.19 | 2402984 | 2316517 | 2888890 | 1622379.75 | 4856330.5 | 3623743.75 | 2939062.5 | 2291841.25 | 1815647.25 | 965715.75 | 1405528.5 | 2625344.75 | 4722898 | 4279405 | 3108770.75 | 3588519 | 3.411071242 | 2 | 2.098579974 | 4 | 4.083370453 | 0 |
| Q8VDP6 | Cdipt | CDP-diacylglycerol--inositol 3-phosphatidyltransferase | 224.95 | 29.58 | 23599 | 5 | 811432.38 | 1736875.12 | 181852.84 | 2 | 235495.7 | 3642838.5 | 2299277 | 269238.75 | 4303924.5 | 2586288 | 2215709.75 | 147121.11 | 3322537.5 | 3475057.75 | 2991529 | 2138274 | 3933958 | 4637726.5 | 2483439.25 | 2760026.25 | 4.417760434 | 2 | 4.097557758 | 4 | 5.379387571 | 0 |
| P37996 | Arl3 | ADP-ribosylation factor-like protein 3 | 48.69 | 5.49 | 20457 | 1 | 1035682.69 | 757026.5 | 789468.12 | 726200.62 | 651685.88 | 779741 | 2 | 2 | 587319.19 | 2 | 2 | 916530.25 | 589204.38 | 561637.94 | 2 | 2 | 2 | 2 | 2 | 2 | 0.345213172 | 3 | 0.522056378 | 4 | 2.52521E-06 | 0 |
| P47754 | Capza2 | F-actin-capping protein subunit alpha-2 | 239.39 | 20.28 | 32967 | 5 | 15324306 | 12337470 | 22295166 | 7149418 | 15259538 | 7604127 | 8064184 | 3941744 | 6652318 | 11802642 | 7084455 | 5551150 | 7799050 | 7179145 | 12489584 | 2544906.25 | 4327893 | 6577534 | 3199923 | 4539053.5 | 0.526007637 | 3 | 0.554175172 | 4 | 0.292807943 | 0 |
| P97834 | Gps1 | COP9 signalosome complex subunit 1 | 46.77 | 2.34 | 53428 | 1 | 954119.25 | 955151.38 | 1752671.38 | 1131733.75 | 932492.19 | 909612.5 | 1045888.94 | 37160.68 | 704250.5 | 717611.75 | 611200 | 2 | 709636.44 | 368466.41 | 1406588.75 | 497402.81 | 745334.56 | 2 | 656834.06 | 2 | 0.59630182 | 3 | 0.540657142 | 4 | 0.331735892 | 0 |
| Q5U2X6 | Ccdc47 | PAT complex subunit CCDC47 | 221.74 | 10.14 | 55733 | 5 | 909126.62 | 832468.19 | 171323.95 | 531475.06 | 884316.81 | 2 | 821839.25 | 2 | 528263.75 | 345788.28 | 2 | 403047.19 | 797971.44 | 390353.34 | 2 | 2 | 2 | 2 | 2 | 2 | 0.50947513 | 4 | 0.478075792 | 4 | 3.00417E-06 | 0 |
| Q91VI7 | Rnh1 | Ribonuclease inhibitor | 1368.47 | 61.4 | 49816 | 20 | 36476858 | 21892854 | 47520718 | 17064834 | 29254578 | 14721566 | 13281122 | 21712958 | 11221163 | 38547110 | 14872697 | 7819582.5 | 19635922 | 17172792 | 28620860 | 8330283.5 | 4503126.5 | 14219826 | 7554501 | 10082676 | 0.65359715 | 5 | 0.578949773 | 4 | 0.293610534 | 0 |
| Q5PQN9 | Mrpl38 | Large ribosomal subunit protein mL38 | 45.33 | 2.63 | 44838 | 1 | 2 | 1111864.75 | 943492.88 | 453916.56 | 857129.12 | 2 | 2 | 2 | 630855.31 | 802992 | 1304391.12 | 970550.44 | 1245849.88 | 1205087 | 571852 | 2248741 | 1442691.75 | 1227250.38 | 1439595.38 | 1128540.25 | 0.425930088 | 5.5 | 1.573705467 | 4 | 2.223980202 | 0 |
| Q8BWT1 | Acaa2 | 3-ketoacyl-CoA thiolase, mitochondrial | 215.66 | 9.57 | 41830 | 3 | 1685788.38 | 4060130.5 | 3647722 | 1710022.75 | 2401469.25 | 1965904.12 | 1585725.75 | 2290765.25 | 1949724.12 | 151453.09 | 1479511.38 | 1383788.12 | 1573694.62 | 1789998.62 | 1870887.75 | 1307730.38 | 1633184.62 | 1472161.88 | 1676436.5 | 1608282.38 | 0.588189128 | 6 | 0.599615018 | 4 | 0.569990375 | 0 |
| P69897 | Tubb5 | Tubulin beta-5 chain | 1990.48 | 69.14 | 49671 | 28 | 44639386 | 20875576 | 57374730 | 35556894 | 33452468 | 15226076 | 23965396 | 22910238 | 22234350 | 42714358 | 14741992 | 11209350 | 20266496 | 25008878 | 40266242 | 7729328 | 16077878 | 15082605 | 8151190 | 15312530 | 0.662069017 | 6 | 0.580997955 | 4 | 0.324928809 | 0 |
| Q61656 | Ddx5 | Probable ATP-dependent RNA helicase DDX5 | 1469.85 | 43.97 | 69290 | 26 | 10820591 | 11027891 | 17058738 | 15185129 | 18614352 | 8678098 | 8961564 | 12671051 | 12483630 | 13531575 | 9202171 | 9096006 | 9488320 | 12148721 | 13258819 | 10617940 | 8327026 | 8267685 | 2896182.5 | 6609962.5 | 0.774700505 | 6 | 0.731624957 | 4 | 0.505026297 | 0 |
| Q9CQD1 | Rab5a | Ras-related protein Rab-5A | 358.19 | 32.56 | 23599 | 5 | 4357194 | 3123403 | 3922044.5 | 2583001.75 | 3846518 | 3940149 | 3242784.75 | 5020854.5 | 4242281 | 4016694 | 5644824 | 3456397.25 | 6064181 | 5208051 | 3983280.5 | 5994463 | 6082795 | 5220016.5 | 6034032 | 5862048.5 | 1.147520088 | 6 | 1.365887926 | 4 | 1.637118159 | 0 |
| P10107 | Anxa1 | Annexin A1 | 2096.89 | 77.75 | 38734 | 33 | 68152450 | 53194074 | 68094202 | 57864230 | 62660078 | 46651882 | 62137354 | 58418646 | 53152146 | 66886970 | 58387310 | 56864686 | 52579758 | 47998998 | 55097390 | 34424622 | 48305362 | 40843458 | 30044642 | 39354562 | 0.926707746 | 7 | 0.874060337 | 4 | 0.622562627 | 0 |
| Q9WVL2 | Stat2 | Signal transducer and activator of transcription 2 | 98.6 | 2.17 | 105417 | 2 | 781457.69 | 622287 | 756452.5 | 1258053.25 | 547842.25 | 1101950.62 | 2764309 | 2 | 1203589.62 | 3200031.25 | 2 | 2 | 2 | 2 | 843108.25 | 2 | 2 | 2 | 2 | 491770.81 | 2.085146046 | 7 | 0.212581076 | 4 | 0.123995793 | 0 |
| P30416 | Fkbp4 | Peptidyl-prolyl cis-trans isomerase FKBP4 | 490.44 | 27.73 | 51572 | 11 | 9026196 | 15987230 | 9500416 | 14165276 | 29807284 | 7154258.5 | 8932034 | 16707499 | 11599939 | 11551719 | 7792342.5 | 8838470 | 7062017 | 9838070 | 10124208 | 3712817.75 | 8358066.5 | 8754953 | 7100808 | 5947055 | 0.712804359 | 8 | 0.556212368 | 4 | 0.431586866 | 0 |
| Q9CWM4 | Pfdn1 | Prefoldin subunit 1 | 74.71 | 9.02 | 14255 | 1 | 945997.75 | 1228982 | 1352763.88 | 812419.94 | 1181544.62 | 1068609.88 | 714071.69 | 1588355.62 | 742303.69 | 1051657.88 | 1396878.12 | 1075861.75 | 1441953.5 | 1431832.38 | 1282136.38 | 1735089.75 | 1679341.12 | 1845402.12 | 1604080.38 | 1684234.12 | 0.935398718 | 9 | 1.200473097 | 4 | 1.548098378 | 0 |
| Q91ZJ5 | Ugp2 | UTP--glucose-1-phosphate uridylyltransferase | 132.77 | 6.3 | 56979 | 3 | 2121695.5 | 1910442.75 | 1589530.25 | 2208459.5 | 1949701.62 | 1462363.5 | 1689791.5 | 3163707 | 2275229.25 | 2426424 | 2723114.5 | 2714883.5 | 2726157 | 3042370.25 | 1689267.75 | 4360011.5 | 2392356 | 2533493.75 | 2604402 | 3156463 | 1.126554928 | 9 | 1.318611213 | 4 | 1.53854687 | 0 |
| O08734 | Bak1 | Bcl-2 homologous antagonist/killer | 125.45 | 10.05 | 23295 | 2 | 4226833 | 4227369.5 | 3893848.25 | 3498452.5 | 3758558 | 3361577 | 5627437.5 | 3223317.5 | 4395613.5 | 2 | 3898690 | 3699351.5 | 2193547.5 | 3442582.5 | 2769250.5 | 5951657 | 7392023 | 6606081 | 5576039.5 | 5810183 | 0.847125509 | 10 | 0.816290334 | 4 | 1.598361928 | 0 |
| Q76MZ3 | Ppp2r1a | Serine/threonine-protein phosphatase 2A 65 kDa regulatory subunit A alpha isoform | 1052.02 | 34.47 | 65323 | 17 | 12130062 | 23265726 | 16284975 | 16013564 | 19607784 | 15168634 | 16316286 | 12311138 | 29010970 | 12364945 | 13230359 | 14419928 | 15607663 | 15244679 | 10705673 | 8108920 | 10686543 | 11764198 | 11150206 | 8068012 | 0.975600384 | 11 | 0.792744886 | 4 | 0.570179557 | 0 |
| Q9Z2I9 | Sucla2 | Succinate--CoA ligase [ADP-forming] subunit beta, mitochondrial | 201.96 | 10.8 | 50114 | 5 | 3402934 | 3338923.5 | 3709661.75 | 2673967.75 | 3902184 | 598410.25 | 3084775.75 | 1549907.25 | 6725767 | 5002260 | 3136566.25 | 5784331 | 4171425.5 | 5804383 | 6493630 | 4317742 | 4816782 | 4658299 | 4145888.75 | 4697635 | 0.996091612 | 11 | 1.491122054 | 4 | 1.329385959 | 0 |
| Q921H8 | Acaa1a | 3-ketoacyl-CoA thiolase A, peroxisomal | 873.27 | 46.23 | 43953 | 13 | 12110358 | 10982388 | 13554334 | 14228259 | 20047524 | 7530803 | 14262904 | 9776953 | 19382448 | 14465819 | 10789693 | 13384217 | 10824546 | 13473834 | 8976255 | 3574571.25 | 8340243 | 6947361 | 6106293 | 7106682 | 0.922395462 | 12 | 0.810014466 | 4 | 0.452254025 | 0 |
| Q78IK4 | Apool | MICOS complex subunit Mic27 | 240.83 | 20.75 | 29261 | 4 | 1897305.62 | 1302169.75 | 1464236 | 1635550.5 | 1499614.75 | 2 | 1562425.38 | 872966.44 | 2682531 | 2965031 | 1129232.75 | 1017179 | 845855.31 | 1628721 | 1443955.38 | 1122463.5 | 915354.19 | 2 | 1177567.12 | 2 | 1.036425656 | 12 | 0.777668854 | 4 | 0.412288714 | 0 |

| | | | | | | | | | | | | | | | | | | | | | | | | | | | | | | | | |
|---|---|---|---|---|---|---|---|---|---|---|---|---|---|---|---|---|---|---|---|---|---|---|---|---|---|---|---|---|---|---|---|---|
| P11167 | Slc2a1 | Solute carrier family 2, facilitated glucose transporter member 1 | 402.01 | 15.65 | 53963 | 7 | 803247.44 | 722347.81 | 932750.88 | 2 | 288093.06 | 1689649 | 1354138.38 | 2 | 1230539.88 | 1316460.5 | 1332307 | 2 | 1205005.88 | 1145320.25 | 1404022.88 | 2075877 | 2347399 | 3064323.75 | 2075125.75 | 1838276 | 2.035648817 | 4.5 | 1.852090636 | 4.5 | 4.151190836 | 0 |
| P57759 | Erp29 | Endoplasmic reticulum resident protein 29 | 363.96 | 32.82 | 28823 | 8 | 5431505 | 12407815 | 5713145 | 8786215 | 6681324 | 4454186 | 3798938.75 | 4711707.5 | 6048345 | 5583751.5 | 5509057.5 | 6566905 | 6109407.5 | 5290510 | 3273372 | 5159876 | 3681351.75 | 4101669 | 4248636 | 2532350 | 0.630367151 | 3 | 0.685526634 | 5 | 0.50548131 | 0 |
| Q80Y14 | Glrx5 | Glutaredoxin-related protein 5, mitochondrial | 147.62 | 20.39 | 16292 | 2 | 933031.19 | 983211.12 | 706816.44 | 250712.92 | 289239.72 | 225616.69 | 168701.95 | 117217.07 | 308585.84 | 390916.81 | 277660.88 | 180294.94 | 390664.03 | 2 | 555463.12 | 2 | 188788.33 | 2 | 2 | 2 | 0.382875118 | 4 | 0.443907655 | 5 | 0.059688792 | 0 |
| Q3THK7 | Gmps | GMP synthase [glutamine-hydrolyzing] | 384.7 | 12.7 | 76723 | 7 | 1790518.5 | 1742860 | 2035929 | 2145215.75 | 2654113.25 | 628116 | 1662653.38 | 1179275.25 | 1678558.88 | 2894089.5 | 645404.62 | 994280.5 | 1008670 | 1722025.5 | 3172380.5 | 552581.56 | 891843.19 | 1202097.25 | 2 | 1023859.62 | 0.775675086 | 5 | 0.727459307 | 5 | 0.353989034 | 0 |
| Q501J6 | Ddx17 | Probable ATP-dependent RNA helicase DDX17 | 873.51 | 22.15 | 72399 | 16 | 5375434 | 4466246 | 6413443 | 5826924 | 9431389 | 5613790.5 | 7155625 | 7511244.5 | 6912485 | 8018618 | 6001200.5 | 5181111.5 | 2345533.25 | 3859021.25 | 4644957 | 2544911.5 | 3525124.75 | 3791416 | 2796651.5 | 1642423 | 1.117357149 | 7 | 0.699124764 | 5 | 0.453791416 | 0 |
| Q9D7H3 | RtcA | RNA 3'-terminal phosphate cyclase | 74.68 | 4.92 | 39254 | 2 | 610733.25 | 746454.5 | 255171.48 | 1400058.5 | 809137.56 | 1211951 | 1088710.88 | 152587.12 | 3215506.5 | 2852375.5 | 3159487.75 | 1375361 | 2265405.75 | 1800646.25 | 363878.56 | 2310360.25 | 1873218.75 | 2410383.5 | 2589707.25 | 1658012.5 | 2.229754734 | 7 | 2.34584577 | 5 | 2.836981655 | 0 |
| Q6PEC4 | Skp1 | S-phase kinase-associated protein 1 | 81.24 | 10.43 | 18672 | 1 | 993053.75 | 815238.06 | 797888.94 | 882430.62 | 679478.5 | 1004131.44 | 870638.88 | 1169333.12 | 639969.12 | 309336.91 | 816683.5 | 734465.12 | 774393.38 | 619257.44 | 416175.72 | 1961582.62 | 1024839.56 | 1459361.88 | 2882711.5 | 1204314.88 | 0.958091019 | 12 | 0.806358611 | 5 | 2.047175254 | 0 |
| Q9JL26 | Fmnl1 | Formin-like protein 1 | 875.4 | 18.1 | 122060 | 17 | 11775747 | 13101773 | 15454244 | 14395907 | 17995738 | 9846187 | 8344708.5 | 6498806 | 12363955 | 12501307 | 14271600 | 13471452 | 12837996 | 12017207 | 8079108 | 8110180.5 | 8716334 | 11555913 | 7678054 | 9919609 | 0.681416949 | 2 | 0.834358068 | 6 | 0.632259834 | 0 |
| P23492 | Pnp | Purine nucleoside phosphorylase | 516.38 | 27.34 | 32277 | 7 | 4584734.5 | 8090226.5 | 6241757.5 | 8953700 | 10254576 | 14502430 | 9353715 | 13056587 | 10754311 | 9947476 | 6851702.5 | 7320888 | 3969662.5 | 5085670.5 | 4689308 | 3623757.5 | 4002651 | 2798525.5 | 1567948.25 | 2028840 | 1.511200716 | 2 | 0.732255358 | 6 | 0.367782932 | 0 |
| Q80Z29 | Nampt | Nicotinamide phosphoribosyltransferase | 211.51 | 12.22 | 55438 | 5 | 3552803.5 | 3864766.5 | 3482315 | 5793184.5 | 5809184.5 | 7453220 | 5718451.5 | 8704219 | 8856752 | 7915454 | 6698536.5 | 5358302 | 5385704 | 5975802 | 3938434 | 7612761.5 | 5884829.5 | 7763420.5 | 7915381 | 7467488 | 1.71752112 | 2 | 1.215735033 | 6 | 1.628453776 | 0 |
| P36536 | Sar1a | Small COPII coat GTPase SAR1A | 116.02 | 21.21 | 22371 | 3 | 1653043.88 | 2003774.88 | 2779552.5 | 1737587.62 | 2050946.38 | 1860198 | 1709919.5 | 2 | 1683149.12 | 2233457.25 | 1397125.25 | 1976586.25 | 2125363.75 | 1537564 | 1611153 | 893588.25 | 2 | 2 | 1304287.88 | 1245337.88 | 0.732204913 | 8 | 0.84575769 | 6 | 0.336748158 | 0 |
| O88543 | Cops3 | COP9 signalosome complex subunit 3 | 196.16 | 9.46 | 47832 | 3 | 2440046.75 | 1931010 | 3329929 | 2440862.5 | 4046407 | 1023144.12 | 2803803.25 | 784634 | 3100308.25 | 2501637 | 244537.41 | 1209276.62 | 1678234.25 | 2505780.25 | 2518451.25 | 2 | 535721.62 | 757829.06 | 355072.88 | 1023719.5 | 0.719857829 | 9 | 0.574861365 | 6 | 0.188349097 | 0 |
| P00520 | Abl1 | Tyrosine-protein kinase ABL1 | 55.55 | 0.71 | 122673 | 1 | 1397034.75 | 639634.62 | 2 | 970654.12 | 2 | 963209.31 | 2 | 2 | 313910.03 | 690571.19 | 1234880.38 | 2 | 1220023.62 | 1624548.75 | 3036790.5 | 1916466.5 | 3021854.25 | 2281575.75 | 2655601.5 | 3898689 | 0.654300051 | 10 | 2.366302065 | 6 | 4.580208523 | 0 |
| P07742 | Rrm1 | Ribonucleoside-diphosphate reductase large subunit | 91.15 | 2.9 | 90210 | 2 | 1021867.94 | 917258.75 | 321819.34 | 1130933.12 | 125659.25 | 1121698.12 | 493526.09 | 98487.64 | 605495 | 461545.38 | 2 | 2 | 655641.06 | 589820 | 672344.5 | 2 | 2 | 2 | 2 | 2 | 0.790539267 | 10 | 0.545213539 | 6 | 2.8429E-06 | 0 |
| Q6I7R3 | Isoc1 | Isochorismatase domain-containing protein 1 | 80.87 | 8.42 | 32002 | 2 | 498955.91 | 831354.44 | 305753.94 | 992868 | 971512.31 | 579335.38 | 508858.31 | 1025442.75 | 697822.19 | 821014.38 | 279246.12 | 285199.72 | 669245.12 | 503030.16 | 607500.06 | 2 | 266926.25 | 218329.69 | 127783.14 | 2 | 1.008895682 | 12 | 0.65109214 | 6 | 0.170268716 | 0 |
| Q8R5C5 | Actr1b | Beta-centractin | 283.82 | 13.03 | 42281 | 5 | 1050723 | 2179030.5 | 1479127.38 | 1036105.62 | 1085952 | 1009815.06 | 1047446.25 | 157646.72 | 1000024.25 | 904454 | 1042687.69 | 1526150 | 1053810.62 | 1029391.56 | 2 | 922352.62 | 666775.38 | 606604.31 | 916646 | 2 | 0.603048363 | 1 | 0.68102529 | 7 | 0.455629971 | 0 |
| Q8BT60 | Cpne3 | Copine-3 | 120.47 | 6 | 59585 | 3 | 697019.25 | 680135.62 | 880225.19 | 882320.12 | 986491.25 | 298937.94 | 407248.5 | 184662.38 | 626823.62 | 1119291.5 | 235830.98 | 713199.25 | 248319.69 | 345309.31 | 1023066.44 | 2 | 328182.53 | 259579.23 | 2 | 257776.48 | 0.639079399 | 5 | 0.621814502 | 7 | 0.204920749 | 0 |
| P56960 | Exosc10 | Exosome complex component 10 | 42.58 | 1.47 | 100942 | 1 | 1044002.44 | 840726.56 | 928552.62 | 1003989.94 | 877234.62 | 457106.22 | 1213071.5 | 717949.81 | 415453.72 | 598944.25 | 363504.47 | 575011.06 | 750764.75 | 2127314.75 | 899195 | 2824528.25 | 1938223.88 | 1557320.88 | 3020617 | 1147283.38 | 0.7247888 | 5 | 1.004533778 | 7 | 2.234095129 | 0 |
| Q60866 | Pter | N-acetyltaurine hydrolase | 125.04 | 10.32 | 39218 | 3 | 2546670.75 | 2160554.5 | 2509130.5 | 1176627.38 | 1022597.44 | 2992321 | 2664542.5 | 3434723.5 | 501018.16 | 1066893.38 | 3909899.5 | 3756772 | 876206.06 | 1239880.62 | 3195890.5 | 2734070.75 | 3972678 | 2871501.25 | 4861810.5 | 4246259.5 | 1.132112721 | 9 | 1.378422561 | 8 | 1.984616866 | 0 |
| Q91VH2 | Snx9 | Sorting nexin-9 | 41.16 | 2.35 | 66546 | 1 | 640972 | 2 | 2 | 920400.5 | 2 | 2 | 2 | 2366741 | 680778.12 | 2 | 2 | 671052.44 | 2761297 | 2723131.75 | 2 | 3946334.5 | 1180095.88 | 4082092.5 | 5129939 | 4206927 | 1.951817013 | 11.5 | 3.942340176 | 8 | 11.87757413 | 0 |
| P47802 | Mtx1 | Metaxin-1 | 56.72 | 5.68 | 35624 | 1 | 116579.55 | 91164.46 | 147000.53 | 82939.05 | 103865.27 | 2 | 2 | 55171.68 | 102821.57 | 149783.92 | 2 | 52770.87 | 2 | 136581.08 | 354581.97 | 2 | 2 | 2 | 2 | 2 | 0.568334998 | 7 | 1.004411532 | 9 | 1.84656E-05 | 0 |
| P32020 | Scp2 | Sterol carrier protein 2 | 178.23 | 7.13 | 59126 | 4 | 2972118.25 | 3379896.75 | 2921950.5 | 2480197.5 | 5965632.5 | 1706475.25 | 1482144.25 | 3086429 | 1013814.56 | 6404263.5 | 4884856 | 2533797.75 | 3681074.5 | 1732969 | 1667182.38 | 1593662.25 | 1058593.5 | 2 | 1564283.75 | 1067380.5 | 0.772758724 | 8 | 0.818287075 | 9 | 0.29819317 | 0 |
| P59999 | Arpc4 | Actin-related protein 2/3 complex subunit 4 | 387.53 | 49.4 | 19667 | 8 | 60034538 | 93693834 | 78313666 | 82753746 | 80929082 | 84223346 | 72674538 | 96685178 | 66755446 | 63990694 | 66816402 | 86081090 | 81723338 | 76485266 | 38343918 | 57356946 | 52702762 | 34119898 | 40217626 | 38384474 | 0.971203063 | 12 | 0.883063067 | 9 | 0.562971208 | 0 |
| Q8R180 | Ero1a | ERO1-like protein alpha | 859.21 | 31.25 | 54084 | 16 | 3717368.75 | 2176000.5 | 5885350.5 | 4745550.5 | 4152475 | 9543592 | 17777838 | 16568958 | 13682502 | 24260640 | 5008791 | 3599334.75 | 3860243.75 | 4575956 | 8753255 | 5950897.5 | 10286246 | 9318237 | 6567005.5 | 8104663 | 3.957756843 | 0 | 1.247661573 | 10 | 1.945521334 | 0 |
| Q8K3C0 | Rnasek | Ribonuclease kappa | 43.93 | 9.18 | 10992 | 1 | 1830503.62 | 2225980.75 | 1022351.44 | 3190331.75 | 3035003 | 4920525 | 3787895.25 | 7113326.5 | 6150788.5 | 2 | 2590127.75 | 2512287 | 2415973.5 | 2389512.5 | 2520249.25 | 3951180 | 4680946.5 | 5950043 | 6594064 | 4349939.5 | 1.94375493 | 5 | 1.09943051 | 10 | 2.258119945 | 0 |
| Q7TPV4 | Mybbp1a | Myb-binding protein 1A | 622 | 10.34 | 152037 | 13 | 11748572 | 15640232 | 8310070.5 | 12372192 | 11107584 | 9630485 | 8620172 | 12837765 | 7896032 | 9188104 | 10172424 | 9050800 | 13352510 | 14492735 | 6192767 | 19384078 | 18805010 | 24080896 | 23489408 | 18923574 | 0.814019204 | 7 | 0.900007613 | 10 | 1.76893128 | 0 |
| P13439 | Umps | Uridine 5'-monophosphate synthase | 194.56 | 9.56 | 52292 | 4 | 3413097.5 | 3141355 | 4127764 | 7117542.5 | 5014184 | 7592094 | 6357040.5 | 6166939 | 2682850 | 4328500.5 | 3113839.5 | 5362756.5 | 5169316.5 | 2331315 | 3504308.5 | 1504251.38 | 2871501 | 1344770.12 | 1656161.88 | 2752625.5 | 1.189072139 | 9 | 0.853931124 | 10 | 0.443996458 | 0 |
| P53657 | Pklr | Pyruvate kinase PKLR | 156.7 | 5.75 | 62309 | 3 | 2 | 2 | 2 | 2 | 359958.94 | 2 | 2 | 2 | 2 | 2 | 2 | 2 | 2 | 2 | 2 | 17672884 | 4663591 | 4463233 | 3083498.5 | 2704733.5 | 2.77803E-05 | 10 | 2.77803E-05 | 10 | 90.53036926 | 0 |
| Q91VC3 | Eif4a3 | Eukaryotic initiation factor 4A-III | 616.41 | 26.28 | 46840 | 10 | 3054452.75 | 6760473 | 3182738.5 | 3949801.5 | 4715227.5 | 3386967.5 | 4413072 | 1828835.88 | 10273836 | 2731342.75 | 6137024.5 | 7396773.5 | 3106662.5 | 6151579 | 3090563.5 | 2090045.25 | 2328141 | 2196940 | 1150477.88 | 2755944 | 1.044840264 | 10 | 1.19480079 | 10 | 0.485698985 | 0 |
| Q99K51 | Pls3 | Plastin-3 | 1016.9 | 30.16 | 70742 | 18 | 10341398 | 6729090 | 10663873 | 5444543.5 | 8387411 | 10619362 | 6979858 | 8863856 | 8689738 | 7294023.5 | 9309210 | 7224744 | 7947178 | 5756852 | 7091422.5 | 3008789.25 | 4436293 | 5091728.5 | 3274246.25 | 3245200.25 | 1.021183547 | 11 | 0.89806869 | 10 | 0.458454328 | 0 |
| Q99L47 | St13 | Hsc70-interacting protein | 309.5 | 15.63 | 41656 | 6 | 9513420 | 5404924.5 | 11355470 | 6867605.5 | 5337390 | 5251625.5 | 4804684.5 | 4888484.5 | 4258696.5 | 6219980 | 6199801.5 | 8629660 | 8555952 | 7345967 | 5172060 | 4188193.5 | 3776593 | 3497335 | 2520118 | 215380.53 | 0.660713546 | 2 | 0.933070448 | 11 | 0.36897243 | 0 |
| P70245 | Ebp | 3-beta-hydroxysteroid-Delta(8),Delta(7)-isomerase | 39 | 4.35 | 26215 | 1 | 1458888.75 | 1253762.62 | 1372635.25 | 1448347.38 | 660188.38 | 2340941.25 | 1890605.38 | 3047401.75 | 1556552.62 | 1001341.44 | 1985055.38 | 1303740.62 | 1441539.5 | 1429502.12 | 1062944.12 | 3799011.5 | 2099697.5 | 2112185 | 2574937.25 | 1613916.5 | 1.588169927 | 4 | 1.166126714 | 11 | 1.969663804 | 0 |
| P29758 | Oat | Ornithine aminotransferase, mitochondrial | 1331.04 | 57.4 | 48355 | 23 | 40842934 | 49537634 | 59307922 | 31138366 | 42665434 | 33315120 | 28286018 | 38821046 | 35151654 | 41498530 | 49162994 | 50614194 | 53098798 | 45763550 | 25636910 | 30946742 | 28035332 | 25835918 | 26516438 | 22135850 | 0.792297435 | 5 | 1.003508649 | 11 | 0.597203062 | 0 |
| P62245 | Rps15a | Small ribosomal subunit protein uS8 | 427.9 | 52.31 | 14840 | 9 | 25213590 | 42454954 | 27040442 | 37674474 | 67448370 | 51802082 | 51437754 | 65442934 | 54051074 | 44037442 | 32580194 | 38879254 | 41639158 | 41131722 | 20833476 | 24187974 | 17889716 | 15635472 | 20774716 | 10503055 | 1.334978947 | 5 | 0.876055651 | 11 | 0.44532912 | 0 |
| Q9DBD5 | Pelp1 | Proline-, glutamic acid- and leucine-rich protein 1 | 361.42 | 7.39 | 118069 | 6 | 3627519.75 | 123929.17 | 1082051.88 | 570250.25 | 106464.05 | 268073.62 | 155533.33 | 65398.48 | 753434.56 | 587048 | 679895.75 | 726657.81 | 73933.88 | 175110.66 | 708102.62 | 2 | 2 | 2 | 2 | 2 | 0.332017527 | 10 | 0.428967051 | 11 | 1.81481E-06 | 0 |
| Q91ZV7 | Plxdc1 | Plexin domain-containing protein 1 | 39.14 | 3 | 55635 | 1 | 26710.71 | 75077.91 | 40779.4 | 70878.9 | 22222.28 | 2 | 2 | 2 | 128973.68 | 142099.48 | 2 | 47849.65 | 51326.72 | 60369.5 | 194243.91 | 2 | 2 | 2 | 2 | 2 | 1.150252812 | 10 | 1.501221967 | 11 | 4.24324E-05 | 0 |
| Q9DBG6 | Rpn2 | Dolichyl-diphosphooligosaccharide--protein glycosyltransferase subunit 2 | 924.51 | 25.2 | 69063 | 13 | 6651205.5 | 4438942.5 | 8067472.5 | 8965252 | 3749407 | 8395456 | 6181552 | 9059226 | 3418952 | 2746964.5 | 8209514 | 6210630.5 | 5265312.5 | 4143014.75 | 13753202 | 20098340 | 15274090 | 22187850 | 13830878 | 19179244 | 0.935049233 | 11 | 1.17913354 | 11 | 2.84166691 | 0 |
| P47757 | Capzb | F-actin-capping protein subunit beta | 314.26 | 22.74 | 31345 | 5 | 39950358 | 48412102 | 25477404 | 35014570 | 46382710 | 36411250 | 39819450 | 46205954 | 53296538 | 39834610 | 44858850 | 50486186 | 34958050 | 39342986 | 24532458 | 17643928 | 13944169 | 21228166 | 21062962 | 21257944 | 1.104133146 | 11 | 0.994577804 | 11 | 0.487290313 | 0 |
| Q5SSI6 | Utp18 | U3 small nucleolar RNA-associated protein 18 homolog | 211.19 | 9.24 | 61218 | 4 | 329919.22 | 544081.69 | 89515.34 | 629878.25 | 150308.78 | 718340.56 | 584962.19 | 2 | 119076.38 | 164867.22 | 72028.64 | 492073 | 465162.19 | 707519 | 526102.19 | 2 | 2 | 2 | 2 | 2 | 0.910274339 | 12 | 1.29774661 | 11 | 5.73492E-06 | 0 |
| P97310 | Mcm2 | DNA replication licensing factor MCM2 | 466.75 | 12.83 | 102078 | 9 | 5741649 | 7951722.5 | 6668155 | 7418037 | 17108018 | 953887.75 | 1890378 | 4415230 | 5000670.5 | 5865013 | 8469970 | 8531764 | 6544230 | 7781109 | 2907047 | 3600080.75 | 3029859.5 | 2791744.25 | 4273139.5 | 2 | 0.403790506 | 1 | 0.7626635 | 12 | 0.305091643 | 0 |
| Q8VBT0 | Tmx1 | Thioredoxin-related transmembrane protein 1 | 168.15 | 8.27 | 31396 | 3 | 6891471.5 | 6565111 | 6813316 | 5128746 | 4968738 | 3396650.75 | 3988950.75 | 5324447.5 | 5111719.5 | 5736903.5 | 5483595 | 4098067.5 | 8077190 | 8274120 | 4337237 | 8798348 | 9103413 | 9365005 | 11818554 | 11324616 | 0.775788694 | 5 | 0.996800086 | 12 | 1.660002669 | 0 |
| O70493 | Snx12 | Sorting nexin-12 | 32.92 | 5.45 | 19116 | 1 | 945589.62 | 1232910.25 | 694029.69 | 1178855.88 | 1118193.38 | 1521631.12 | 1203373.25 | 2103134.25 | 1280600 | 876341.56 | 937032.38 | 1388668.62 | 1199340.62 | 1004533.19 | 935035.38 | 1778018 | 1688404.38 | 1398538.62 | 1253892.75 | 1512558.5 | 1.351189415 | 5 | 1.057070678 | 12 | 1.476215474 | 0 |
| P52293 | Kpna2 | Importin subunit alpha-1 | 438.36 | 18.34 | 57928 | 8 | 892339.06 | 1088957.62 | 2143986.75 | 814192.25 | 3481134.75 | 2 | 158659.84 | 894056.69 | 1451458.62 | 1626852.88 | 1143916.25 | 1066026.25 | 852804.62 | 1735035.75 | 1363279.75 | 2 | 2 | 2 | 359489.72 | 2 | 0.490585577 | 8 | 0.731664607 | 12 | 0.042692596 | 0 |
| P08775 | Polr2a | DNA-directed RNA polymerase II subunit RPB1 | 271.52 | 3.91 | 217176 | 5 | 3824223 | 4547184 | 2507249.5 | 2867118.5 | 2787831.5 | 3549590 | 3053612.25 | 1545788 | 3069191.75 | 589021.31 | 4607134 | 2619110.5 | 3131937.5 | 3171574.25 | 2045139.38 | 844289.19 | 1892053.12 | 1896999.5 | 2354758.75 | 1318019.25 | 0.714133562 | 9 | 0.942014413 | 12 | 0.502377978 | 0 |
| P43277 | H1-3 | Histone H1.3 | 800.18 | 41.18 | 22100 | 13 | 2 | 1465921.5 | 2 | 1851447.62 | 1741147.75 | 2 | 2 | 1626947.25 | 1584184 | 459655.03 | 2 | 1576426.5 | 1354392.38 | 2503083.5 | 2 | 7998961 | 4952878.5 | 4361622.5 | 6008979.5 | 4911757 | 0.72566475 | 10 | 1.074208552 | 12 | 5.581512704 | 0 |
| P62908 | Rps3 | Small ribosomal subunit protein uS3 | 1232.52 | 82.72 | 26674 | 25 | 68962794 | 154523954 | 81353666 | 159125218 | 106005946 | 120093794 | 135252498 | 127594034 | 129426938 | 106618914 | 122771114 | 130850690 | 146253010 | 133049218 | 42212762 | 65604178 | 45851170 | 37471222 | 49419570 | 31856196 | 1.085994814 | 10 | 1.009062234 | 12 | 0.403883886 | 0 |
| Q3UMR5 | Mcu | Calcium uniporter protein, mitochondrial | 58.92 | 2.86 | 39682 | 1 | 1130043.5 | 2 | 1072148.38 | 1204563.12 | 1334077.5 | 1261093.38 | 945305.62 | 1170887.25 | 2 | 1430092.12 | 442773.72 | 1117225.25 | 1327553.62 | 1459718.38 | 657223 | 1951256.75 | 1848635.75 | 1809116.75 | 1818235 | 1398138.75 | 1.014036742 | 11.5 | 1.05561457 | 12 | 1.86156741 | 0 |
| P39052 | Dnm2 | Dynamin-2 | 663.16 | 16.21 | 98230 | 14 | 13263781 | 13478282 | 11996636 | 12278042 | 12508248 | 15012535 | 13241766 | 13945679 | 11337590 | 10613720 | 12816476 | 10209760 | 13292963 | 12121189 | 13532649 | 17006294 | 14570324 | 17080540 | 16687280 | 18865232 | 1.009859128 | 12 | 0.975569425 | 12 | 1.325614869 | 0 |
| F1LNJ2 | Snrnp200 | U5 small nuclear ribonucleoprotein 200 kDa helicase | 706.67 | 8.42 | 244875 | 16 | 11609262 | 12076759 | 13563099 | 11279108 | 13219409 | 7241474.5 | 5276201 | 4889992 | 5463917 | 6162063.5 | 6777222 | 4417565 | 9723926 | 10284704 | 7422278 | 6912283 | 6573443 | 11464402 | 4919321 | 7694628.5 | 0.470198528 | 0 | 0.625541266 | 0 | 0.608348421 | 1 |
| P34960 | Mmp12 | Macrophage metalloelastase | 227.05 | 11.21 | 54971 | 5 | 1747290.25 | 1524405.88 | 3815198.75 | 2520478.25 | 2615260 | 5440064 | 4462009 | 7020914.5 | 3938979.25 | 5132736.5 | 7922356 | 5463830 | 8076679 | 8823701 | 10555502 | 8275962 | 3671428 | 11374950 | 9379204 | 6645517 | 2.126767855 | 0 | 3.341511405 | 0 | 3.219196762 | 1 |
| Q2EMV9 | Parp14 | Protein mono-ADP-ribosyltransferase PARP14 | 45.15 | 0.55 | 203802 | 1 | 1133401.75 | 1768877 | 1751671.38 | 2284813.75 | 1500462.75 | 2 | 1212867.62 | 991179 | 925143.56 | 870830.75 | 2 | 645806.12 | 2 | 2 | 2 | 2 | 744057.88 | 2 | 2 | 1335922.75 | 0.473979798 | 1 | 0.076525273 | 0 | 0.246466498 | 1 |
| Q9DCL9 | Paics | Bifunctional phosphoribosylaminoimidazole carboxylase/phosphoribosylaminoimidazole succinocarboxamide synthetase | 469.94 | 21.88 | 47006 | 9 | 14474997 | 19582874 | 11411337 | 13808046 | 16349196 | 8767420 | 7278001 | 11442959 | 9847824 | 5499200 | 9892266 | 11057216 | 10352178 | 10319646 | 9228494 | 10439579 | 8700353 | 10463994 | 10819246 | 13701482 | 0.566407705 | 1 | 0.672381158 | 0 | 0.715684182 | 1 |
| Q00612 | G6pdx | Glucose-6-phosphate 1-dehydrogenase X | 909.04 | 38.64 | 59263 | 18 | 25215466 | 33521372 | 27085950 | 24584892 | 37825638 | 53109550 | 45463218 | 45528342 | 36914922 | 37657726 | 61965706 | 65615970 | 45767062 | 45735122 | 40423842 | 60551838 | 53268030 | 37265554 | 59610586 | 45628554 | 1.475199779 | 2 | 1.750670534 | 0 | 1.729196684 | 1 |
| Q8BH43 | Wasf2 | Actin-binding protein WASF2 | 424.01 | 15.49 | 54074 | 8 | 3370600.5 | 2098519.75 | 2583947 | 2890427.5 | 2282942.5 | 3271613.75 | 3407723.75 | 3043761.5 | 6061664 | 4171569.5 | 8880675 | 9974627 | 6038283.5 | 8691620 | 3442388.5 | 7164465.5 | 6117966.5 | 5017807 | 6840514 | 3204721.75 | 1.50882147 | 2 | 2.79951383 | 0 | 2.143092219 | 1 |
| P56391 | Cox6b1 | Cytochrome c oxidase subunit 6B1 | 240.41 | 50 | 10071 | 4 | 4305351 | 4282084.5 | 3941436.5 | 4241617 | 2150465.5 | 1820962.12 | 1739984.5 | 2265527.5 | 2294709 | 3133573.5 | 2 | 2 | 2 | 2 | 1949332.25 | 1294350.62 | 2 | 1680758 | 2238895.25 | 2 | 0.594830278 | 3 | 0.103025471 | 0 | 0.275567909 | 1 |
| Q8BL97 | Srsf7 | Serine/arginine-rich splicing factor 7 | 354.23 | 21.72 | 30818 | 7 | 833677.62 | 1773750 | 1791820.12 | 2612234 | 7085022.5 | 5899477.5 | 2003429.75 | 7935205 | 9005497 | 3014007.5 | 21510646 | 16930490 | 15133388 | 18891270 | 7563547 | 16229724 | 15518916 | 5446773.5 | 17603922 | 10077680 | 1.976207454 | 4 | 5.677247326 | 0 | 4.602347816 | 1 |
| Q62426 | Cstb | Cystatin-B | 155.67 | 44.9 | 11046 | 3 | 7244759 | 6622262 | 7574419 | 5557882.5 | 7828922 | 7157263 | 8995716 | 13235120 | 8885087 | 7127745 | 16487954 | 18527734 | 11732268 | 8017994 | 14924616 | 16760776 | 14373458 | 10242861 | 7963583 | 7664079 | 1.303566449 | 6 | 2.00097843 | 0 | 1.636739314 | 1 |
| P10639 | Txn | Thioredoxin | 278.19 | 37.14 | 11675 | 4 | 1630619.12 | 931475.38 | 4297961 | 1031324.75 | 70866.51 | 1973540.75 | 1325057.25 | 1891078 | 8265495 | 1453745 | 18068446 | 31003022 | 23280802 | 28040470 | 27170098 | 27343760 | 21295408 | 3147786 | 4503734.5 | 13414000 | 1.872450886 | 6 | 16.02096014 | 0 | 8.75439943 | 1 |
| Q3TCN2 | Plbd2 | Putative phospholipase B-like 2 | 268.35 | 11.62 | 66289 | 6 | 4626621 | 5130918 | 3536852.5 | 3207683.5 | 2036128.38 | 1094195 | 2908589 | 4084536.5 | 1889537 | 3595258.5 | 121655.59 | 1982405.62 | 853360 | 1955544.25 | 1479673 | 988797.69 | 1829703.88 | 2424880.5 | 1195682.38 | 1056225.12 | 0.732116037 | 7 | 0.344835922 | 0 | 0.404315856 | 1 |
| Q7TPR4 | Actn1 | Alpha-actinin-1 | 611.02 | 17.15 | 103068 | 11 | 3943575 | 3686295.75 | 3464208.5 | 2745882.5 | 2673903 | 4004183 | 3343091 | 1429344.5 | 2060494 | 2297425 | 4206290.5 | 4138999 | 4806970 | 5031682 | 5318688 | 3874512.5 | 4417756.5 | 6480856 | 5887459 | 4943698.5 | 0.795364241 | 7 | 1.423205885 | 0 | 1.550471854 | 1 |
| Q9CQV8 | Ywhab | 14-3-3 protein beta/alpha | 708.81 | 41.46 | 28086 | 12 | 4896747.5 | 3315442.75 | 5708252 | 6802657 | 8317940 | 7370293 | 7398489 | 2326258.75 | 8924247 | 8336892.5 | 12719067 | 18032744 | 16987800 | 16453269 | 13922475 | 8768146 | 9115824 | 12013244 | 10611480 | 8129581.5 | 1.183021722 | 7 | 2.689826432 | 0 | 1.67481181 | 1 |
| F1LQ48 | Hnrnpl | Heterogeneous nuclear ribonucleoprotein L | 702.06 | 26.65 | 67903 | 12 | 8078394 | 11682807 | 4391476 | 12318553 | 5053131 | 9879114 | 8760970 | 5351971 | 18972768 | 11996494 | 15704475 | 25198792 | 17523854 | 19698384 | 12949119 | 17124440 | 15093348 | 12263827 | 17884082 | 13788538 | 1.32359212 | 8 | 2.19328177 | 0 | 1.833965248 | 1 |
| O09043 | Napsa | Napsin-A | 169.67 | 13.84 | 45544 | 3 | 16189915 | 18799612 | 13301041 | 13880866 | 11503477 | 13429767 | 10915863 | 15108324 | 14470656 | 12683789 | 10354294 | 9532162 | 11092173 | 9808202 | 4511010.5 | 12049505 | 6660784 | 8497764 | 8948697 | 5883593 | 0.904085232 | 9 | 0.614834017 | 0 | 0.570619529 | 1 |
| P04762 | Cat | Catalase | 335.45 | 13.09 | 59757 | 6 | 3188444.75 | 2783376.5 | 4354390.5 | 183842.34 | 3192236 | 3034477 | 1768582.88 | 4294577 | 1083005.5 | 2674074.75 | 14609217 | 18908598 | 7158840 | 14648034 | 5916564 | 15377904 | 9396479 | 4177789.5 | 13383325 | 5859010 | 0.9381437 | 9 | 4.469417345 | 0 | 3.517259318 | 1 |
| O35215 | Ddt | D-dopachrome decarboxylase | 50.82 | 11.02 | 13077 | 1 | 1328448.25 | 2 | 2 | 2 | 2 | 2 | 2 | 2 | 462074.31 | 761013.81 | 2673077.75 | 2518914.75 | 2023755.5 | 2096887.88 | 2026025.62 | 3760333.25 | 2855388.75 | 2154484.75 | 1177419.62 | 2557334.75 | 0.920688295 | 11 | 8.535216346 | 0 | 9.413152386 | 1 |
| P17751 | Tpi1 | Triosephosphate isomerase | 1025.69 | 79.12 | 26713 | 15 | 49366594 | 141189490 | 99657570 | 100226194 | 79270738 | 78621386 | 99680954 | 78342858 | 141579250 | 115039730 | 199122594 | 284467426 | 156096050 | 207506770 | 174917794 | 225732226 | 212952322 | 129022266 | 151884994 | 183897026 | 1.092724314 | 11 | 2.176043429 | 0 | 1.923501111 | 1 |
| P27546 | Map4 | Microtubule-associated protein 4 | 350.39 | 8 | 117429 | 6 | 947786.44 | 1419360.88 | 1718975.5 | 1831214.75 | 2042578.38 | 1475022.12 | 892676.31 | 1649767.25 | 4878722.5 | 2146993.5 | 6506036 | 6546057.5 | 3477113.5 | 4350250 | 2579878.25 | 2856013.25 | 3470272.5 | 1964331.75 | 3886989 | 2307908.75 | 1.387349031 | 11 | 2.947183789 | 0 | 1.819807563 | 1 |
| P62137 | Ppp1ca | Serine/threonine-protein phosphatase PP1-alpha catalytic subunit | 409.08 | 22.42 | 37540 | 8 | 2 | 2 | 2 | 139435.06 | 243394.22 | 2 | 124146.74 | 122524.17 | 433840.66 | 2 | 265775.72 | 265987.12 | 897642.12 | 289605.16 | 743040.25 | 247912.31 | 420138.66 | 271423.16 | 240369.12 | 287153.88 | 1.777567548 | 11 | 6.431095823 | 0 | 3.831927742 | 1 |
| Q8CG47 | Smc4 | Structural maintenance of chromosomes protein 4 | 185.01 | 2.95 | 146895 | 4 | 5528178 | 7201362 | 4967845.5 | 7203776 | 6178335 | 72138.85 | 439674.94 | 2917331.25 | 1182578.25 | 2097384.5 | 1343523.38 | 2011011.75 | 4972314 | 3593079 | 3181670.75 | 2678003.5 | 2090249.25 | 3940062 | 5139125 | 4749610 | 0.215869256 | 0 | 0.485902302 | 1 | 0.598370368 | 1 |
| Q9CYN2 | Spcs2 | Signal peptidase complex subunit 2 | 127.61 | 10.62 | 24978 | 3 | 271372.47 | 2 | 2 | 483714.31 | 1024545.19 | 631177.62 | 579573.12 | 1368770.12 | 1435121.75 | 1434663.25 | 878667.69 | 1109073.75 | 1332435.75 | 1520549.5 | 1505352.38 | 1009167.31 | 1150611 | 1213554.38 | 1059077.75 | 1052337.25 | 3.062034007 | 2 | 3.565942236 | 1 | 3.08194922 | 1 |
| P41498 | Acp1 | Low molecular weight phosphotyrosine protein phosphatase | 168.01 | 13.92 | 18152 | 3 | 7733582 | 5949479 | 6473837 | 4231382.5 | 7086816 | 4303485.5 | 4753398 | 4807112 | 4046285 | 5353144 | 8176747 | 8014165.5 | 10081861 | 7470591.5 | 12888423 | 9088131 | 9813065 | 8269658 | 9215791 | 7692987 | 0.73910574 | 4 | 1.481545513 | 1 | 1.400460583 | 1 |
| Q9QUJ7 | Acsl4 | Long-chain-fatty-acid--CoA ligase 4 | 704.09 | 20.25 | 79077 | 13 | 8631226 | 15733784 | 6813302 | 12792210 | 10441894 | 18128884 | 17248188 | 16297322 | 13496159 | 9729802 | 6783487.5 | 6693440 | 6636458.5 | 8391071 | 6725141 | 5313456.5 | 6653838 | 8528710 | 5275641 | 4391894.5 | 1.376530588 | 4 | 0.647455132 | 1 | 0.554350316 | 1 |
| Q9Z0N1 | Eif2s3x | Eukaryotic translation initiation factor 2 subunit 3, X-linked | 751.19 | 33.26 | 51065 | 13 | 3247721 | 4302019 | 6345901 | 5795777.5 | 10736560 | 7081690.5 | 8304664 | 11182894 | 7308629 | 8965900 | 22658074 | 26041514 | 16102489 | 16729072 | 9312756 | 16178032 | 12088381 | 9674222 | 17527564 | 14084660 | 1.408038904 | 4 | 2.985538622 | 1 | 2.28581925 | 1 |
| Q8BZM1 | Glmn | Glomulin | 56.34 | 1.85 | 67756 | 1 | 791166.56 | 133686.56 | 439505.53 | 232825.61 | 564132.19 | 2 | 151746.5 | 2 | 257009.48 | 421766.03 | 2 | 2 | 2 | 210056.2 | 2 | 2 | 191477.7 | 2 | 2 | 2 | 0.384268583 | 5 | 0.097192709 | 1 | 0.08859679 | 1 |
| P26645 | Marcks | Myristoylated alanine-rich C-kinase substrate | 129.21 | 13.27 | 29661 | 3 | 1038699.94 | 2352934.5 | 662563.19 | 1498296 | 1080071.5 | 3210994.5 | 3774237.5 | 1435101.62 | 4622386.5 | 670658.69 | 2404413.75 | 3860567.5 | 3453738.5 | 3277946.25 | 2209690 | 2727933 | 4614228 | 2069344.88 | 3947676 | 2844894.5 | 2.067582985 | 6 | 2.292680992 | 1 | 2.44310852 | 1 |
| Q07076 | Anxa7 | Annexin A7 | 543.49 | 19.65 | 49925 | 9 | 8731718 | 10115022 | 8965135 | 7079648 | 6936030 | 5356104.5 | 6052643 | 7622528 | 11759929 | 6648532.5 | 14224084 | 16506326 | 12418379 | 12174210 | 9414593 | 13054605 | 12110024 | 12165517 | 9233605 | 13062747 | 0.895097473 | 7 | 1.547726017 | 1 | 1.425531587 | 1 |
| O88342 | Wdr1 | WD repeat-containing protein 1 | 765.62 | 21.45 | 66407 | 13 | 17124174 | 26997010 | 47996186 | 18269048 | 70113834 | 57356818 | 40543318 | 43077626 | 54174802 | 46679986 | 78924010 | 92804290 | 83290738 | 79082266 | 69264234 | 82308970 | 70261386 | 75680354 | 79777826 | 68370818 | 1.33979065 | 8 | 2.234709002 | 1 | 2.085312069 | 1 |
| Q99K85 | Psat1 | Phosphoserine aminotransferase | 625.66 | 40.81 | 40473 | 12 | 9448713 | 8158965 | 14012177 | 10083978 | 12094220 | 10817532 | 10010280 | 17130962 | 10743582 | 11489357 | 15295362 | 14744097 | 17872146 | 16459576 | 13024913 | 15648109 | 16903908 | 18258028 | 12322903 | 15239247 | 1.118845565 | 9 | 1.438641172 | 1 | 1.45678497 | 1 |
| P48774 | Gstm5 | Glutathione S-transferase Mu 5 | 123.26 | 11.16 | 26635 | 3 | 2 | 549747.81 | 2 | 1429050 | 1190204.12 | 984227.94 | 717092.12 | 1067435.88 | 1268109.5 | 679128.81 | 2063216.38 | 1471445.38 | 2163423.5 | 1559212.5 | 1296197.5 | 1229201.38 | 2638258.5 | 2185829.25 | 1888519.62 | 2039153.25 | 1.488162015 | 9 | 2.699109894 | 1 | 3.149556113 | 1 |
| Q8C2Q3 | Rbm14 | RNA-binding protein 14 | 572.65 | 18.98 | 69449 | 11 | 387683.72 | 1151487 | 805170.56 | 920492.88 | 2944652.5 | 3275644 | 2 | 2 | 900353.44 | 1277022.12 | 7480181 | 6802167 | 2050537.88 | 5843940 | 6195020.5 | 4684507.5 | 7674811.5 | 2659742.5 | 4229229 | 3148073.75 | 0.878176226 | 11 | 4.569113026 | 1 | 3.60679803 | 1 |
| P41216 | Acsl1 | Long-chain-fatty-acid--CoA ligase 1 | 748.87 | 20.46 | 77951 | 13 | 15011382 | 7030842.5 | 15467761 | 9452538 | 12973224 | 8044199 | 9666626 | 9527546 | 13460395 | 16206658 | 4069211 | 3346427.75 | 6735819 | 3750970.25 | 7076272.5 | 4907084 | 3824109 | 8155899.5 | 1758339 | 6194244 | 0.949440465 | 12 | 0.416757971 | 1 | 0.414438403 | 1 |
| Q61072 | Adam9 | Disintegrin and metalloproteinase domain-containing protein 9 | 164.58 | 7.46 | 92079 | 4 | 1861648.25 | 2290644 | 1780622.75 | 1411145.25 | 873784.44 | 2 | 502634.91 | 120830.77 | 1226207.25 | 173120.5 | 108338.09 | 2 | 881054 | 1160993 | 741829 | 643923 | 429908.94 | 1185651.38 | 2 | 819297.75 | 0.24614671 | 1 | 0.351943387 | 2 | 0.374646052 | 1 |
| O70145 | Ncf2 | Neutrophil cytosol factor 2 | 242.78 | 10.29 | 59485 | 5 | 10453740 | 7764297 | 9135245 | 11035356 | 6368168 | 4601927 | 3864192 | 3500713 | 7188726 | 4653514 | 7071908 | 7112322.5 | 6191843 | 4788212.5 | 3495660.25 | 4296729.5 | 1761989 | 1753394.75 | 7163649.5 | 2913393.75 | 0.531965395 | 1 | 0.640348336 | 2 | 0.399696897 | 1 |
| Q04753 | Clns1a | Methylosome subunit pICln | 52.84 | 5.51 | 26092 | 1 | 834791.12 | 1183008.38 | 637328.44 | 1173457.62 | 988731.31 | 2 | 628352.81 | 2 | 557902.5 | 916508.94 | 517390.38 | 2 | 898115.69 | 484148.34 | 2 | 2 | 2 | 762540.38 | 2 | 2 | 0.436501959 | 2 | 0.394339517 | 2 | 0.158293175 | 1 |
| O35864 | Cops5 | COP9 signalosome complex subunit 5 | 122.66 | 8.68 | 37549 | 3 | 4030863.25 | 4898409.5 | 3202776.5 | 2447410.5 | 1245726.75 | 1240920.38 | 1060082.12 | 2 | 1879803.5 | 2950018.25 | 1701907.25 | 1593317.5 | 1201549.38 | 978195.88 | 1175422.88 | 1931377.88 | 2 | 926072.56 | 340138.69 | 864711.12 | 0.450599824 | 3 | 0.420241044 | 2 | 0.256698539 | 1 |
| P17665 | Cox7c | Cytochrome c oxidase subunit 7C, mitochondrial | 93.59 | 50.79 | 7333 | 2 | 1013006.81 | 862747.12 | 1518389.25 | 1606879.88 | 953151.75 | 2 | 705430.94 | 855514.94 | 759492.12 | 1224121.75 | 2 | 453234.91 | 802563.44 | 680668.94 | 998831.81 | 34012.66 | 101052.56 | 947053.62 | 84409.65 | 95320.48 | 0.595306967 | 3 | 0.492982016 | 2 | 0.211926759 | 1 |
| Q3ULJ0 | Gpd1l | Glycerol-3-phosphate dehydrogenase 1-like protein | 94.81 | 6.55 | 38226 | 2 | 3365666 | 2290730 | 3489163 | 2528300.75 | 1783779 | 1078483.75 | 1731017 | 2416357 | 1146977.75 | 1832222.25 | 1902682 | 672196.75 | 1765201.38 | 1342646 | 2110936.75 | 2 | 1677502 | 1195664.38 | 2093463.25 | 1268181.88 | 0.6096952 | 3 | 0.579125582 | 2 | 0.463291787 | 1 |
| B0K025 | Ostc | Oligosaccharyltransferase complex subunit OSTC | 138.72 | 18.12 | 16815 | 3 | 510396.69 | 1112166.5 | 606601.94 | 1259073.75 | 2125414.25 | 1691312.62 | 1437856.5 | 2146398.75 | 2076412.38 | 1578513.38 | 1711155.75 | 1498033.75 | 3220349 | 3443229.5 | 3703091 | 1278403.38 | 2915076 | 4676802.5 | 2997446.5 | 3482259.25 | 1.590852413 | 4 | 2.418364421 | 2 | 2.734402585 | 1 |
| Q9WV55 | Vapa | Vesicle-associated membrane protein-associated protein A | 345.69 | 26.91 | 27855 | 7 | 4226192 | 9751889 | 882824.81 | 6501073 | 2270625.75 | 17022140 | 14835281 | 3925440.75 | 17807618 | 9709440 | 13513602 | 18000004 | 13389884 | 17112306 | 4484620.5 | 20264530 | 13686844 | 12181758 | 18005776 | 6909229 | 2.678499511 | 4 | 2.81392668 | 2 | 3.006360844 | 1 |
| P45377 | Akr1b8 | Aldose reductase-related protein 2 | 599.28 | 32.28 | 36121 | 11 | 1129529 | 3535255.25 | 477839.09 | 1352539.5 | 1671802.38 | 3456705.5 | 3381970.75 | 4171152.25 | 8702617 | 1112073.75 | 13093079 | 14349886 | 3208626.75 | 6125177 | 1847781.5 | 13690011 | 3773392.5 | 3245477.5 | 5282296 | 4849171 | 2.549847916 | 6 | 4.729363871 | 2 | 3.776231093 | 1 |
| Q9EQK5 | Mvp | Major vault protein | 957.51 | 22.76 | 95924 | 16 | 16983506 | 17974464 | 16716660 | 13496643 | 18498924 | 11739071 | 8994401 | 13201104 | 13138908 | 16165552 | 15276967 | 13669997 | 14399027 | 8777938 | 11883016 | 13522547 | 5367795.5 | 11687463 | 10817583 | 7591192 | 0.75581316 | 1 | 0.764990968 | 3 | 0.585472274 | 1 |
| Q5BK63 | Ndufa9 | NADH dehydrogenase [ubiquinone] 1 alpha subcomplex subunit 9, mitochondrial | 190.52 | 11.14 | 42559 | 3 | 1812201.12 | 1321541.12 | 514055.75 | 1666522.25 | 217506.06 | 986630.25 | 405165.53 | 102012.91 | 220452.14 | 147467.5 | 68012.62 | 88906.38 | 250413.31 | 415854.12 | 409671.78 | 2 | 423691.94 | 141351.53 | 113765.64 | 200320.64 | 0.336548588 | 4 | 0.222866399 | 3 | 0.158922515 | 1 |
| P09411 | Pgk1 | Phosphoglycerate kinase 1 | 2165.77 | 80.82 | 44550 | 34 | 227320258 | 397612898 | 239521186 | 371089602 | 347306658 | 325138850 | 391030850 | 399217762 | 503853186 | 416113410 | 437983042 | 508480450 | 374712610 | 436070242 | 363656258 | 420627170 | 424346018 | 407293922 | 375536834 | 438048706 | 1.285878816 | 4 | 1.339925953 | 3 | 1.305146959 | 1 |
| Q9CPT4 | Mydgf | Myeloid-derived growth factor | 111.13 | 16.27 | 17982 | 2 | 1598845.38 | 1729071.88 | 2969832.75 | 2120403 | 2586898.5 | 632000.88 | 665668.62 | 1914178.75 | 637530.94 | 2324775.25 | 4135803.75 | 2894894.75 | 4781744.5 | 3239149 | 2392328.25 | 3348804 | 3571744 | 3118410.5 | 2684700.5 | 3171564.5 | 0.561029127 | 5 | 1.585083017 | 3 | 1.444357029 | 1 |
| Q05144 | Rac2 | Ras-related C3 botulinum toxin substrate 2 | 585.76 | 51.04 | 21441 | 10 | 39023250 | 23433796 | 49348746 | 40920414 | 47268118 | 17107984 | 28423222 | 14257121 | 27814918 | 44500646 | 12744312 | 19105054 | 26331566 | 24491268 | 38453430 | 10082582 | 21694996 | 20610202 | 13271942 | 25109514 | 0.660538201 | 5 | 0.605645338 | 3 | 0.453859061 | 1 |
| P70600 | Ptk2b | Protein-tyrosine kinase 2-beta | 236.51 | 5.15 | 115784 | 5 | 5326828.5 | 3691807.25 | 5174757 | 3296552 | 3807096.5 | 3186599.75 | 3570607.5 | 3471881.25 | 2640326 | 4234343 | 2392816 | 1542899.88 | 2736885 | 1869624.12 | 4568742 | 1511583.5 | 2629407.75 | 2716611.5 | 1680626.38 | 3361890.5 | 0.803104868 | 5 | 0.615623872 | 3 | 0.55876868 | 1 |
| P15651 | Acads | Short-chain specific acyl-CoA dehydrogenase, mitochondrial | 259.74 | 16.75 | 44765 | 5 | 3051880 | 3058870.75 | 4769671 | 1578676.25 | 3866610.5 | 2 | 2209676.25 | 1459848.88 | 2701932 | 2216938.25 | 2022219.12 | 2032165.88 | 1846466.5 | 1558575.38 | 2303273 | 2 | 886844.56 | 2066879.75 | 1369988.75 | 2 | 0.526065829 | 3 | 0.597995479 | 4 | 0.264841006 | 1 |
| Q8BI84 | Mia3 | Transport and Golgi organization protein 1 homolog | 82.41 | 1.14 | 213675 | 2 | 696073.12 | 3275330.5 | 2 | 447391.34 | 441869.88 | 3259741.75 | 2055840.75 | 970834.69 | 2130853 | 2360829 | 5183737 | 5518998 | 4357785.5 | 3536025.75 | 393388.31 | 8258376.5 | 6753916 | 3244943.5 | 7403410 | 6951949 | 2.217411632 | 5 | 3.906857883 | 4 | 6.709489886 | 1 |
| Q9ET26 | Rnf114 | E3 ubiquitin-protein ligase RNF114 | 119.15 | 15.72 | 25745 | 3 | 2673515.75 | 3652657.75 | 1842243.5 | 872170.12 | 3477239 | 26633404 | 2426841.5 | 5625907 | 3933111.5 | 3177722.5 | 6119502.5 | 5999011 | 4673621.5 | 5000690 | 960534.88 | 3611543.75 | 4451444.5 | 4400076 | 6381224 | 4288013 | 3.338997211 | 5 | 1.817676621 | 4 | 1.847948759 | 1 |
| P62317 | Snrpd2 | Small nuclear ribonucleoprotein Sm D2 | 384.63 | 55.93 | 13527 | 8 | 1972686.38 | 955544.38 | 1060008 | 969547.75 | 2107569.75 | 1280143 | 1063495.38 | 380113.69 | 797845.38 | 316524.84 | 10602170 | 3380176 | 2357458 | 1657315 | 1295214.12 | 2840360 | 2813428 | 2622497.25 | 2012143.38 | 3001756 | 0.543231247 | 6 | 2.73055348 | 4 | 1.881035314 | 1 |
| Q9Z1F9 | Uba2 | SUMO-activating enzyme subunit 2 | 128.04 | 3.13 | 70569 | 2 | 1006333.25 | 1242126.88 | 769803 | 1081954.25 | 1124739 | 813208.31 | 2 | 979645.25 | 2 | 1322103.25 | 1513881.75 | 2326474.5 | 2924663 | 3177004.75 | 938542.56 | 2852374 | 1218270.25 | 3849028.5 | 2385964.25 | 3446494.75 | 0.596169725 | 7 | 2.082422468 | 4 | 2.632008911 | 1 |
| Q5RJQ4 | Sirt2 | NAD-dependent protein deacetylase sirtuin-2 | 129.91 | 7.14 | 39319 | 2 | 1639991.12 | 320345.97 | 1919317 | 530474.88 | 1330271.75 | 389628.28 | 494387.91 | 125364.19 | 1529220.25 | 1515910.75 | 212327.7 | 135426.47 | 1072868.75 | 289867.81 | 682940.5 | 239489.92 | 232507.77 | 286765.59 | 276436.78 | 354979.53 | 0.706311559 | 8 | 0.416944974 | 4 | 0.24217466 | 1 |
| O35841 | Api5 | Apoptosis inhibitor 5 | 315.68 | 12.3 | 56785 | 6 | 6659129 | 7436401.5 | 7524829 | 7021067 | 9185429 | 7465720.5 | 5749701 | 9631847 | 6636399.5 | 5840428 | 7585851 | 6494775 | 5612265.5 | 5749233 | 5126585 | 6101737 | 5430676 | 6879302.5 | 5930869 | 4225924.5 | 0.933836438 | 8 | 0.808121878 | 4 | 0.755244088 | 1 |
| Q93092 | Taldo1 | Transaldolase | 1134.67 | 54.6 | 37387 | 22 | 41806122 | 69413954 | 52469954 | 56295170 | 60955842 | 86529962 | 78396034 | 79165274 | 56023814 | 18884890 | 86067754 | 79151482 | 58764654 | 58735058 | 72941602 | 84906418 | 83845466 | 73489866 | 89337258 | 67155954 | 1.135469463 | 8 | 1.265961525 | 4 | 1.419283417 | 1 |
| Q8K1X4 | Nckap1l | Nck-associated protein 1-like | 493.11 | 13.23 | 128905 | 9 | 6614639.5 | 5232776 | 6338283 | 4432460 | 7198865 | 2959159.5 | 5334987 | 8033285.5 | 4792920 | 5951388.5 | 3447525 | 4333870 | 6319404.5 | 5182330.5 | 4593691.5 | 2707175 | 3534296.5 | 4218610.5 | 4997550 | 2021312.25 | 0.907929006 | 10 | 0.80077817 | 4 | 0.586206878 | 1 |
| Q61029 | Tmpo | Lamina-associated polypeptide 2, isoforms beta/delta/epsilon/gamma | 240.17 | 13.27 | 50373 | 5 | 1365118.62 | 2 | 1625115.12 | 2 | 4636271 | 3238528.5 | 2 | 2 | 3550262.5 | 2 | 3728865.5 | 3237185.5 | 3968876.75 | 4859279.5 | 3007842.5 | 5724932 | 5267971 | 3628334.5 | 6603931.5 | 5742324.5 | 0.890157899 | 11 | 2.465354776 | 4 | 3.536020795 | 1 |
| P80317 | Cct6a | T-complex protein 1 subunit zeta | 1122.09 | 38.79 | 58004 | 21 | 19999806 | 12421649 | 20511824 | 24611014 | 22871182 | 15782502 | 20979636 | 22610578 | 20913246 | 18462184 | 20267894 | 23888950 | 27360376 | 25058048 | 31391298 | 25295708 | 27180668 | 34722510 | 23671688 | 28982024 | 0.983395697 | 11 | 1.274370967 | 4 | 1.392739496 | 1 |
| Q91V12 | Acot7 | Cytosolic acyl coenzyme A thioester hydrolase | 238.91 | 16.27 | 42537 | 5 | 2703190 | 3356388 | 3565842 | 2 | 4401150 | 3701512.25 | 3835209.25 | 4236185 | 1242605.62 | 1431446.25 | 6084909.5 | 6206951.5 | 4127295 | 4740934 | 2971209.5 | 4057481.5 | 5491791 | 4560275 | 4650192 | 4416386 | 1.029970713 | 11 | 1.720398933 | 4 | 1.652301468 | 1 |
| A1A5S1 | Prpf6 | Pre-mRNA-processing factor 6 | 123.6 | 3.61 | 106764 | 3 | 1658076.12 | 647884.44 | 644235.06 | 2 | 326806.03 | 2 | 350810.62 | 2 | 1022067.94 | 636458.44 | 2 | 2199908.25 | 1720603.25 | 3074311.5 | 2071828.12 | 2138125 | 1103877.38 | 3520685.25 | 3151959 | 3796034.75 | 0.613164102 | 9 | 2.766750998 | 4.5 | 4.183907876 | 1 |
| Q63413 | Ddx39b | Spliceosome RNA helicase Ddx39b | 563.56 | 22.9 | 49035 | 10 | 5381199 | 2053815.75 | 8287410 | 6378374 | 5546578.5 | 933094.38 | 1392225.25 | 6251334 | 4639227 | 4368186.5 | 4339041 | 3156720.25 | 4329065.5 | 3535299.5 | 3412934.75 | 3723622.25 | 1541691.25 | 1660099.75 | 840798.75 | 2036792.5 | 0.636012124 | 5 | 0.67901779 | 5 | 0.354572675 | 1 |
| Q9ET30 | Tm9sf3 | Transmembrane 9 superfamily member 3 | 44.7 | 1.87 | 67545 | 1 | 635081 | 145427.31 | 1177670 | 409320.19 | 975139.62 | 2 | 168536.59 | 194576.84 | 341331.16 | 2020690.12 | 2 | 2 | 542498.69 | 2 | 845397.94 | 2 | 157526.56 | 2 | 2 | 2 | 0.815265252 | 8 | 0.415211752 | 5 | 0.047128811 | 1 |
| P40112 | Psmb3 | Proteasome subunit beta type-3 | 401.43 | 40.49 | 22965 | 6 | 7593357.5 | 1751326.75 | 7520347 | 4925593 | 8366525 | 1597245.62 | 4655907.5 | 5786477 | 5255270.5 | 8175808.5 | 1655218.12 | 2716253 | 4106406.25 | 3739480.5 | 5604327.5 | 2 | 1539142.88 | 1361431 | 1892434.38 | 1646391 | 0.844599365 | 9 | 0.590960545 | 5 | 0.213528182 | 1 |

| | | | | | | | | | | | | | | | | | | | | | | | | | | | | | | | | |
|---|---|---|---|---|---|---|---|---|---|---|---|---|---|---|---|---|---|---|---|---|---|---|---|---|---|---|---|---|---|---|---|---|
| Q8VCW8 | Acsf2 | Medium-chain acyl-CoA ligase ACSF2, mitochondrial | 124.25 | 9.59 | 67951 | 3 | 1972319.38 | 1903465.5 | 1965640.62 | 936469.06 | 1827733.12 | 1265208.75 | 1153289.75 | 1553528.5 | 2262132.75 | 1651313.12 | 1430726.75 | 1196965.88 | 1894535.12 | 1292320.38 | 415904.62 | 2 | 2 | 1460940.38 | 773044.94 | 2 | 0.91631583 | 9 | 0.72399748 | 5 | 0.259596557 | 1 |
| Q91ZW2 | Pofut1 | GDP-fucose protein O-fucosyltransferase 1 | 35.79 | 1.78 | 44688 | 1 | 1065915.38 | 1118762.25 | 1171882.75 | 1373548.62 | 822921.62 | 800131.31 | 841705.38 | 1305564.38 | 913714.69 | 649993.69 | 980727 | 2 | 1097461.5 | 721983.75 | 1155974.25 | 2 | 2 | 718511 | 2 | 984626.44 | 0.812368913 | 6 | 0.712430521 | 6 | 0.306705213 | 1 |
| P14824 | Anxa6 | Annexin A6 | 607.14 | 20.06 | 75885 | 13 | 12243927 | 5973918.5 | 19104462 | 3273527 | 17450170 | 2381533.5 | 3858216.25 | 6297915 | 5803733.5 | 13511268 | 16981508 | 17358620 | 20450712 | 15774086 | 27271420 | 23973778 | 20022402 | 32962934 | 19579356 | 18819134 | 0.548748644 | 7 | 1.685496648 | 6 | 1.98734788 | 1 |
| Q91XV3 | Basp1 | Brain acid soluble protein 1 | 1220.93 | 84.07 | 22087 | 16 | 4899134.5 | 21497860 | 3050009.5 | 24640508 | 4990265.5 | 13939046 | 13217033 | 12827260 | 47582938 | 11964437 | 15036685 | 31521686 | 16285452 | 33491336 | 16662203 | 39463950 | 32998966 | 22803346 | 52125850 | 43376354 | 1.684740324 | 8 | 1.9126881 | 6 | 3.229107019 | 1 |
| Q6MGD0 | Cuta | Protein CutA | 33.07 | 7.91 | 18659 | 1 | 1373587.75 | 200078.66 | 398775.09 | 312388.62 | 353372.47 | 2 | 2 | 351151.12 | 437511.75 | 490268.69 | 119162.31 | 2 | 173892.3 | 254682.28 | 1480043 | 2 | 2 | 110564.4 | 188906.61 | 289753.66 | 0.484775341 | 10 | 0.768622507 | 6 | 0.223344739 | 1 |
| Q5RJR8 | Lrrc59 | Leucine-rich repeat-containing protein 59 | 210.08 | 19.87 | 34869 | 5 | 1734386.38 | 1295067.75 | 4483953 | 338891.03 | 4258369 | 294676.19 | 1906291 | 4065258.5 | 2688617.75 | 5815032.5 | 1033262.38 | 154654.81 | 169477.89 | 1485401.38 | 4021226 | 2 | 236306.22 | 790257.19 | 2 | 2 | 1.219575746 | 11 | 0.566774924 | 6 | 0.084765719 | 1 |
| Q9CQF0 | Mrpl11 | Large ribosomal subunit protein uL11m | 41.92 | 9.38 | 20680 | 1 | 1462743.5 | 1465318.38 | 856885.69 | 512007.75 | 958565 | 2 | 597271.38 | 1387564.88 | 777314.12 | 870212.88 | 697581.88 | 985976.06 | 669662.56 | 746020.56 | 654610 | 2 | 678664.94 | 2 | 2 | 2 | 0.691152358 | 7 | 0.714268204 | 7 | 0.129135252 | 1 |
| E9Q634 | Myo1e | Unconventional myosin-Ie | 445.54 | 12.56 | 126818 | 10 | 2019042.75 | 2130513.25 | 2325734.25 | 1575068.5 | 2006487.88 | 1591630.75 | 1209055.5 | 224490.03 | 3408237 | 1843765.88 | 2322521.25 | 2138297.75 | 2922257.5 | 2353025.25 | 1531348.5 | 1535794.25 | 536508.62 | 1902320 | 855713.5 | 2 | 0.823039215 | 7 | 1.120376065 | 7 | 0.480303474 | 1 |
| Q8CIE6 | Copa | Coatomer subunit alpha | 590.18 | 11.44 | 138432 | 12 | 9957043 | 11224716 | 12177465 | 13627816 | 20162010 | 10711556 | 13496043 | 15140798 | 14169226 | 16696027 | 9644448 | 8538228 | 12410765 | 13524690 | 10359634 | 8661436 | 10247962 | 5128620 | 6894836 | 5790050.5 | 1.045638769 | 9 | 0.811296139 | 7 | 0.546886434 | 1 |
| Q9CQN6 | Tmem14c | Transmembrane protein 14C | 161.88 | 29.82 | 11642 | 3 | 1390201.25 | 749258.12 | 1007463.25 | 1182442.25 | 1032098.12 | 111271.62 | 1052268.12 | 1171887.25 | 1153491.75 | 866303.25 | 879663.69 | 175345.89 | 1009948.06 | 606939 | 1335216.12 | 2079246.62 | 2448663.75 | 3910358 | 1326393.25 | 2684085.5 | 0.812319697 | 10 | 0.747391667 | 7 | 2.321893696 | 1 |
| Q78T54 | Vma21 | Vacuolar ATPase assembly integral membrane protein Vma21 | 76.61 | 14.85 | 11365 | 2 | 2 | 2 | 2 | 2 | 420753.47 | 2 | 2 | 554734.81 | 2 | 2 | 646886.06 | 2 | 797047.19 | 375663.69 | 2 | 796094.25 | 712968.56 | 556954.12 | 145129.8 | 543691.19 | 1.318425877 | 12 | 4.324542692 | 7 | 6.547267553 | 1 |
| Q3UKJ7 | Smu1 | WD40 repeat-containing protein SMU1 | 236.28 | 13.06 | 57544 | 5 | 2103904.5 | 2363103.5 | 1655161.75 | 2483611 | 2700870.5 | 1869577.12 | 1529421.75 | 2375077.75 | 2040121.12 | 1618905.38 | 2233271.25 | 2069099.12 | 1038147.56 | 3074681.5 | 1297256.12 | 1090552.75 | 1110463.38 | 1560750 | 1731684 | 529472.62 | 0.834296814 | 5 | 0.859003726 | 8 | 0.53268847 | 1 |
| Q68FX0 | Idh3B | Isocitrate dehydrogenase [NAD] subunit beta, mitochondrial | 169.05 | 11.95 | 42354 | 4 | 2 | 605165.38 | 2 | 2 | 2 | 2 | 2 | 2 | 2 | 2 | 2 | 378894.84 | 663085 | 284883.78 | 2 | 1439013.12 | 545379.81 | 721086.25 | 1000910 | 901698.88 | 1.65242E-05 | 10 | 2.192541285 | 8 | 7.614492329 | 1 |
| Q80WQ6 | Rhbdf2 | Inactive rhomboid protein 2 | 69.64 | 2.18 | 93434 | 2 | 1132098.75 | 565846.81 | 2 | 512975.12 | 706543.06 | 633477.44 | 2 | 2 | 1205046.38 | 2 | 503411.66 | 542657.81 | 728890.56 | 337321.53 | 2 | 2098470 | 2341724.25 | 2250313 | 725566.31 | 3253935.25 | 0.630180432 | 9.5 | 0.72401315 | 8.5 | 3.657286755 | 1 |
| Q91YR9 | Ptgr1 | Prostaglandin reductase 1 | 291.6 | 19.45 | 35560 | 6 | 17711692 | 13418068 | 19829488 | 21647038 | 14866646 | 11929766 | 9361463 | 13955731 | 9984326 | 6961012 | 16566125 | 15422720 | 16921724 | 13869510 | 17321282 | 9986567 | 12523092 | 13666972 | 12464622 | 10465027 | 0.596667984 | 1 | 0.915727405 | 9 | 0.675709373 | 1 |
| Q8BVI4 | Qdpr | Dihydropteridine reductase | 115.66 | 16.6 | 25570 | 2 | 2593049.25 | 2643115.25 | 1774431 | 1446085.62 | 4633658 | 3444702.75 | 2787089.5 | 4782965.5 | 3264387 | 2270002 | 3616519 | 3362899.5 | 2888641.5 | 3212627 | 1240414 | 1138563.88 | 1366967 | 2 | 1577059.12 | 209278.45 | 1.264225976 | 6 | 1.094020626 | 9 | 0.32786549 | 1 |
| Q91XU1 | Qki | KH domain-containing RNA-binding protein QKI | 251.36 | 22.29 | 37643 | 5 | 2396804.5 | 953609.12 | 4428006 | 1113638.12 | 5037858 | 404374.25 | 1352914.62 | 768709 | 2457637.25 | 3643672.75 | 932696.62 | 816901.88 | 2915621.25 | 2590670 | 2676651 | 250068.88 | 404890.34 | 1104973.75 | 186819.61 | 775000 | 0.619336687 | 8 | 0.713036671 | 9 | 0.19538902 | 1 |
| P97825 | Jpt1 | Jupiter microtubule associated homolog 1 | 40.54 | 9.74 | 16081 | 1 | 2 | 186149.55 | 2 | 81942.3 | 246853.66 | 2 | 2 | 2 | 90371.71 | 82318.56 | 2 | 90575.14 | 144599.5 | 206767.44 | 893351.06 | 289027 | 269024.91 | 239985.27 | 344037.25 | 1533605.88 | 0.335365442 | 9 | 2.593060318 | 9 | 5.196005158 | 1 |
| Q80X82 | Sympk | Symplekin | 63.78 | 0.85 | 142620 | 1 | 623303.94 | 106012.33 | 583397.56 | 243556.97 | 283120.88 | 132277.42 | 306446.69 | 101432 | 259398.22 | 392781.34 | 2 | 2 | 342735.88 | 564981.06 | 533340.31 | 2 | 2 | 220562.94 | 2 | 2 | 0.648222824 | 9 | 0.783444476 | 9 | 0.119915156 | 1 |
| Q9CWJ9 | Atic | Bifunctional purine biosynthesis protein ATIC | 1178.61 | 46.11 | 64217 | 21 | 20026352 | 17812814 | 49520446 | 30595938 | 31341382 | 19418430 | 20263088 | 21766270 | 22938724 | 24909858 | 18974720 | 18253922 | 23863616 | 19355820 | 47547734 | 14300931 | 15431147 | 19764202 | 12588296 | 16184157 | 0.732073784 | 9 | 0.857323793 | 9 | 0.52424877 | 1 |
| P62334 | Psmc6 | 26S proteasome regulatory subunit 10B | 482.71 | 30.08 | 44173 | 10 | 2232163.75 | 4643814.5 | 4428600.5 | 5065008 | 8421814 | 4531913.5 | 4141305.5 | 6451144 | 3568733 | 8585544 | 3400808.75 | 4452414 | 5806088.5 | 1088459.25 | 4630109 | 1275975.38 | 1937899 | 2015306.12 | 2019552.5 | 3224395.25 | 1.100326693 | 12 | 0.781637137 | 9 | 0.42245004 | 1 |
| Q8BRF7 | Scfd1 | Sec1 family domain-containing protein 1 | 255.7 | 16.59 | 72323 | 6 | 570315.5 | 3350454.5 | 1336057.12 | 2642461 | 3791459.25 | 1646316 | 2427121.5 | 2508015.75 | 3369145 | 3292509 | 2457812.5 | 2509046 | 2968613 | 2348849 | 2 | 2 | 188950.78 | 2 | 875668.75 | 2 | 1.132785341 | 12 | 0.879697608 | 9 | 0.091065652 | 1 |
| Q61033 | Tmpo | Lamina-associated polypeptide 2, isoforms alpha/zeta | 189.41 | 10.25 | 75168 | 4 | 58229.92 | 201304.25 | 845625.5 | 238661.95 | 909229.38 | 2 | 153800.7 | 57599 | 141460.41 | 49120.76 | 1184644.38 | 102623.46 | 103963.8 | 167869.11 | 807719.25 | 2 | 171589.31 | 2 | 2 | 2 | 0.17841712 | 2 | 1.050495528 | 11 | 0.076162195 | 1 |
| P57780 | Actn4 | Alpha-actinin-4 | 968.27 | 23.25 | 104977 | 16 | 12227532 | 14896407 | 17742344 | 10474767 | 13127143 | 10580814 | 9155562 | 10228953 | 11889627 | 14012311 | 11984667 | 13590545 | 12303492 | 10247027 | 18104354 | 8576298 | 8141402 | 11213524 | 8454027 | 6705712.5 | 0.815959419 | 5 | 0.967311712 | 11 | 0.629357394 | 1 |
| Q9JHU4 | Dync1h1 | Cytoplasmic dynein 1 heavy chain 1 | 3268.821 | 15.7 | 532045 | 63 | 48216002 | 51349602 | 72903882 | 47213374 | 57321802 | 46383662 | 41602634 | 56363270 | 38491654 | 52130802 | 49494186 | 47506494 | 55017866 | 48757254 | 56971734 | 40366078 | 40930726 | 47820138 | 36628994 | 32126934 | 0.848260171 | 6 | 0.930480852 | 11 | 0.714330469 | 1 |
| Q03265 | Atp5f1a | ATP synthase subunit alpha, mitochondrial | 2364.39 | 60.94 | 59753 | 36 | 4625405 | 3688032 | 9469738 | 2389386.25 | 11274999 | 3667797.25 | 5017787 | 7191877 | 5053144.5 | 24299220 | 3058582.5 | 2759760.25 | 6352849 | 4288752 | 13676753 | 1572030.88 | 1886899.38 | 2830265 | 1351538.25 | 2107735 | 1.438261836 | 10 | 0.958315892 | 12 | 0.30999125 | 1 |
| P58389 | Ptpa | Serine/threonine-protein phosphatase 2A activator | 132.85 | 12.38 | 36710 | 3 | 4224244.5 | 5166914 | 5568233 | 3581562 | 3818065.25 | 3062783.25 | 2352178.25 | 153137.73 | 2468467.25 | 2775355.75 | 2148700 | 2399475.75 | 2887048 | 2655521.25 | 2379607.25 | 3122168.25 | 2554408 | 3590522.75 | 2604543.25 | 3703821 | 0.483559782 | 0 | 0.557732537 | 0 | 0.696607638 | 2 |
| Q6DFW4 | Nop58 | Nucleolar protein 58 | 504.86 | 25.37 | 60343 | 9 | 7846229.5 | 10351596 | 10421583 | 8862480 | 7747698.5 | 6737081 | 5246814 | 5930571 | 7066720 | 7190264 | 3015261.75 | 3000159.5 | 6669755 | 4664555.5 | 6073845 | 5083554 | 5514721 | 7353035 | 7480814.5 | 8022738 | 0.711292146 | 0 | 0.51788173 | 0 | 0.739667654 | 2 |
| Q9ESY9 | Ifi30 | Gamma-interferon-inducible lysosomal thiol reductase | 248.78 | 20.16 | 27784 | 5 | 8190037.5 | 7046190.5 | 7372462.5 | 5182271 | 4723032.5 | 4666918 | 3878838.5 | 5109326 | 3600058 | 3031614 | 3755892 | 4130874 | 1635717.5 | 4664179.5 | 4194713 | 1341894.12 | 3011742 | 5898206 | 3000273.5 | 3622252.5 | 0.623939172 | 1 | 0.565337374 | 0 | 0.518987859 | 2 |
| P06151 | Ldha | L-lactate dehydrogenase A chain | 1308.42 | 62.95 | 36499 | 21 | 107720930 | 114408562 | 113041034 | 111911970 | 185814290 | 178791042 | 179128546 | 224662658 | 257644754 | 209821682 | 378680290 | 430477794 | 234942530 | 269703106 | 193557730 | 322319906 | 219004674 | 167771970 | 288724514 | 154744930 | 1.659115207 | 2 | 2.381686056 | 0 | 1.821096298 | 2 |
| P14685 | Psmd3 | 26S proteasome non-ATPase regulatory subunit 3 | 159.49 | 4.91 | 60718 | 3 | 2 | 2 | 2 | 182490.91 | 62945.85 | 2 | 1635226.62 | 1059899 | 1840489.25 | 403773.41 | 406999 | 592876.31 | 1209183 | 1451740.5 | 707476.94 | 123247.55 | 431152.19 | 660757.06 | 356804.16 | 146259.58 | 20.12440815 | 3.5 | 17.79753353 | 0 | 7.000493883 | 2 |
| Q9D7S7 | Rpl22l1 | Ribosomal protein eL22-like 1 | 127.26 | 19.67 | 14467 | 2 | 1388236.12 | 1285413.62 | 2152286.25 | 1800872.88 | 2716735.25 | 806235.56 | 904295.88 | 1725890.62 | 1619530.75 | 963340.56 | 806782 | 830509.69 | 1229249.38 | 867466.19 | 754241.12 | 1673295.38 | 2 | 628517.62 | 1086537.5 | 2 | 0.644219505 | 4 | 0.480358237 | 0 | 0.362641248 | 2 |
| Q07797 | Lgals3bp | Galectin-3-binding protein | 1071.16 | 28.77 | 64491 | 15 | 14545500 | 15074563 | 14051970 | 16025310 | 9934597 | 18567562 | 19561450 | 12657042 | 19593790 | 21898162 | 2355002.75 | 6603900.5 | 5928308.5 | 5573111.5 | 4431728 | 4639060 | 6752272.5 | 12554632 | 8493354 | 10972786 | 1.325225263 | 4 | 0.357480364 | 0 | 0.623451027 | 2 |
| Q8R146 | Apeh | Acylamino-acid-releasing enzyme | 37.74 | 1.09 | 81581 | 1 | 4977963 | 5092038.5 | 5290853.5 | 1476731.75 | 2520435.5 | 1775069.88 | 1294977.12 | 2529241 | 1609181.12 | 2424875.5 | 1016858.25 | 1153205 | 827822.19 | 1046949.12 | 631601.19 | 1200745.75 | 2283224.75 | 991735.19 | 2503434.75 | 2 | 0.497640952 | 5 | 0.241576112 | 0 | 0.360529725 | 2 |
| O35892 | Sp100 | Nuclear autoantigen Sp-100 | 164.59 | 6.22 | 54727 | 3 | 4150582.75 | 2178992.5 | 4109330.5 | 6343617 | 5014300 | 1640700.5 | 4085536.5 | 6142053.5 | 320638.41 | 463041.81 | 1044814.56 | 926210.31 | 1916685.38 | 2 | 2 | 1704083.62 | 2421518.5 | 3541749 | 1817788.75 | 495062.78 | 0.580450227 | 5 | 0.178361511 | 0 | 0.457874194 | 2 |
| P28271 | Aco1 | Cytoplasmic aconitate hydratase | 41.59 | 1.35 | 98126 | 1 | 1090358 | 1434811.5 | 927537 | 960506.75 | 831564.44 | 913840.81 | 726633 | 1103833.62 | 542521.94 | 2 | 654063.81 | 627699.31 | 800134.31 | 625394 | 2 | 948901.38 | 2 | 2 | 2 | 2 | 0.626686499 | 5 | 0.516188405 | 0 | 0.180924614 | 2 |
| P32577 | Csk | Tyrosine-protein kinase CSK | 183.41 | 9.78 | 50746 | 4 | 4502497 | 5576313 | 3487286.5 | 6605307 | 5559377 | 3901235.5 | 3947326 | 5013808.5 | 3559012.5 | 3115808.5 | 2683565.75 | 1975767.88 | 2773500.5 | 2634539 | 2565015.25 | 3493781.5 | 2602728 | 4366455 | 2659443 | 2845839.5 | 0.759292591 | 5 | 0.490944625 | 0 | 0.620589298 | 2 |
| Q6P069 | Sri | Sorcin | 151.35 | 11.62 | 21627 | 3 | 13953588 | 23128646 | 16465425 | 16276046 | 14087524 | 16493951 | 12633662 | 15905589 | 10973786 | 9031470 | 10689832 | 10942434 | 9717686 | 10037392 | 7573880 | 10938448 | 8668127 | 10292616 | 15047740 | 10976065 | 0.775086467 | 6 | 0.583488343 | 0 | 0.66645426 | 2 |
| Q8K0D5 | Gfm1 | Elongation factor G, mitochondrial | 345.25 | 10.79 | 83550 | 8 | 1362288.5 | 523055.44 | 1928451.25 | 726433.38 | 1865306 | 1541938.25 | 1179749.75 | 2723652.25 | 1240703.75 | 5314760 | 4159827.5 | 3222339.75 | 5377717 | 2208032.5 | 2313085.5 | 3634126.5 | 2185122.75 | 2274656.75 | 3299990.25 | 1581623 | 1.873505461 | 8 | 2.697823587 | 0 | 2.025673128 | 2 |
| E9Q555 | Rnf213 | E3 ubiquitin-protein ligase RNF213 | 665.23 | 2.89 | 584258 | 15 | 6659683 | 7675634.5 | 7031556.5 | 7023087 | 6244417.5 | 5391055 | 6555351 | 8047311.5 | 3924602.75 | 7286084 | 3119949 | 2199626.25 | 2601293 | 3476051.5 | 791133.88 | 3125068.5 | 6071789.5 | 6922080 | 5614230.5 | 4385696.5 | 0.900966196 | 10 | 0.351906232 | 0 | 0.754131188 | 2 |
| O70435 | Psma3 | Proteasome subunit alpha type-3 | 426.61 | 29.02 | 28405 | 7 | 33545350 | 48792914 | 30673740 | 36638322 | 31093562 | 40033010 | 31639274 | 43840402 | 29191530 | 20604120 | 21665390 | 20401898 | 25603076 | 28050738 | 14937402 | 23032304 | 30165126 | 28436302 | 16356142 | 32771928 | 0.914599867 | 10 | 0.612239259 | 0 | 0.723464585 | 2 |
| Q91YR1 | Twf1 | Twinfilin-1 | 98.76 | 4.86 | 40079 | 2 | 2014464.88 | 2406623.25 | 1770587.62 | 1655929 | 2186274.5 | 1887858.12 | 2093488.38 | 2439260.25 | 1887440.12 | 2120886.5 | 1183455.38 | 1325746.12 | 1119010 | 1562541.62 | 1089390.88 | 992702.81 | 1256015.62 | 1771850 | 2 | 1019428.31 | 1.039372023 | 10 | 0.625893918 | 0 | 0.502298126 | 2 |
| P97797 | Sirpa | Tyrosine-protein phosphatase non-receptor type substrate 1 | 300.77 | 16.76 | 56414 | 5 | 26574932 | 29065286 | 28924826 | 23409346 | 19369456 | 8636036 | 13319764 | 13294156 | 12952821 | 14680769 | 16087086 | 20368016 | 16670002 | 17609162 | 16556720 | 10280642 | 17884306 | 20104260 | 18982524 | 19682646 | 0.493809069 | 0 | 0.685474711 | 1 | 0.682674356 | 2 |
| P63085 | Mapk1 | Mitogen-activated protein kinase 1 | 88.52 | 5.87 | 41276 | 2 | 1959082.75 | 6205382 | 4095243.25 | 3247900.5 | 4831905 | 6340620 | 5107902.5 | 6437119.5 | 6412637.5 | 7510327 | 10957967 | 10454359 | 6886947 | 8918702 | 5337382.5 | 9006197 | 5355044 | 6571626 | 9020926 | 6046449 | 1.563882366 | 1 | 2.09225051 | 1 | 1.769965737 | 2 |
| P05765 | Rps21 | Small ribosomal subunit protein eS21 | 315.26 | 59.04 | 9127 | 5 | 374543.31 | 1071584.62 | 1777041.5 | 3805792 | 9552724 | 8348942.5 | 6061709.5 | 12740052 | 8698547 | 5700216 | 23094284 | 19644056 | 12806833 | 13403214 | 6946013.5 | 12303262 | 11472316 | 7197595.5 | 15709172 | 6024511 | 2.505744496 | 4 | 4.577001585 | 1 | 3.178618767 | 2 |
| Q3UZ39 | Lrrfip1 | Leucine-rich repeat flightless-interacting protein 1 | 546.07 | 18.52 | 79249 | 8 | 2625397.5 | 5709532.5 | 488388.03 | 6881398 | 5834521 | 4324096 | 6043938 | 8378474.5 | 9619907 | 4985294 | 6705957.5 | 11344484 | 8392904 | 8992784 | 7395230 | 9144341 | 8390963 | 6001216 | 7639786.5 | 6415298 | 1.548416476 | 7 | 1.988527237 | 1 | 1.745261657 | 2 |
| Q8CHH9 | Septin8 | Septin-8 | 289.12 | 12.12 | 49812 | 5 | 3168328.75 | 790347.69 | 49502.61 | 183969.53 | 444451.81 | 2588252 | 2 | 88398.19 | 636710.31 | 2 | 2476879.75 | 3598256.75 | 4063481.5 | 3902357.25 | 4929576 | 3257078.25 | 5464676 | 3725837.75 | 577234.25 | 6313775.5 | 0.714610754 | 8 | 4.091478595 | 1 | 4.170857983 | 2 |
| P17047 | Lamp2 | Lysosome-associated membrane glycoprotein 2 | 317.8 | 14.22 | 45681 | 7 | 56947722 | 43788750 | 45855514 | 33826986 | 22624090 | 38065446 | 40730018 | 38625830 | 33146262 | 37056942 | 18400476 | 16979212 | 17309448 | 15681099 | 24729204 | 20627916 | 28342646 | 21965152 | 3258490.5 | 23858418 | 0.924062591 | 9 | 0.458520661 | 1 | 0.482915405 | 2 |
| Q3TIU4 | Pde12 | 2',5'-phosphodiesterase 12 | 109.48 | 4.28 | 67514 | 2 | 832866.38 | 1793029.12 | 1431433.62 | 1736068 | 1591067 | 1244856.62 | 870585.38 | 2 | 1284037.5 | 480416.62 | 791495.88 | 2 | 1281273.38 | 1047451.25 | 2 | 1304058.5 | 2 | 1009676.31 | 196359.39 | 2 | 0.525413633 | 3 | 0.422539058 | 2 | 0.339916094 | 2 |
| Q66H50 | Far1 | Fatty acyl-CoA reductase 1 | 465.82 | 19.22 | 59268 | 9 | 7513762 | 3707635 | 6519636 | 5714426.5 | 3053391.5 | 2548047.5 | 1694603.5 | 4584293 | 2635502.75 | 3404838.5 | 516316.78 | 2 | 4560184.5 | 2672104.5 | 2263162 | 2 | 1210468.88 | 4050888 | 866485.19 | 1166498.38 | 0.560842311 | 3 | 0.377676489 | 2 | 0.2751663 | 2 |
| B5DFC8 | Eif3c | Eukaryotic translation initiation factor 3 subunit C | 373.81 | 8.34 | 105435 | 8 | 7501966 | 4810217 | 7973480.5 | 7757198.5 | 5960281.5 | 1273142.88 | 6716745 | 3186139.25 | 5445673 | 5462474 | 3980322.25 | 5058765.5 | 3837151.5 | 4776934 | 5655003.5 | 3713854.5 | 2466684.5 | 5199586.5 | 3294042.5 | 4888435 | 0.649474485 | 4 | 0.685471234 | 2 | 0.575317485 | 2 |
| Q62093 | Srsf2 | Serine/arginine-rich splicing factor 2 | 194.88 | 17.65 | 25476 | 4 | 1355257.5 | 3860306.5 | 2152554.5 | 2700250 | 9573730 | 4509471 | 3982455 | 6409021 | 5594140 | 4456704.5 | 10386354 | 10697476 | 10291682 | 9767810 | 2748913.5 | 16567955 | 12237522 | 6788755.5 | 15063850 | 7679813.5 | 1.270322084 | 5 | 2.234600112 | 2 | 2.970043959 | 2 |
| P85108 | Tubb2a | Tubulin beta-2A chain | 1741.05 | 59.1 | 49907 | 26 | 2150158.5 | 2015168.88 | 1906833.62 | 1018095.44 | 456263.06 | 1063951.12 | 949314.06 | 1505141.88 | 952916.12 | 553944.12 | 2 | 742914.81 | 2 | 1004586.06 | 395686.22 | 861331.81 | 2 | 2 | 860581 | 2 | 0.665905296 | 7 | 0.28399729 | 2 | 0.228173903 | 2 |
| P49300 | Clec10a | C-type lectin domain family 10 member A | 308.24 | 23.36 | 34596 | 6 | 5976716.5 | 1849346.75 | 7151406 | 3444021.75 | 3082026.5 | 4734295.5 | 1419452 | 812038.25 | 4667197 | 4163329.25 | 7551285 | 10072427 | 9855357 | 7629128 | 5815012.5 | 9086895 | 7537267 | 7112551.5 | 7413034 | 6110689 | 0.734592003 | 9 | 1.903093738 | 2 | 1.732760071 | 2 |
| B0BN93 | Psmd13 | 26S proteasome non-ATPase regulatory subunit 13 | 224.85 | 12.23 | 42817 | 5 | 5409428 | 7316887.5 | 11637260 | 4365922 | 8315481 | 4195695 | 5528016.5 | 7863446.5 | 1578388.62 | 8589324 | 3378196.25 | 3400037.75 | 4855211.5 | 2702906 | 4644788.5 | 3873482.25 | 3555004.75 | 4765436.5 | 4369421 | 4121422.5 | 0.749220859 | 9 | 0.512380916 | 2 | 0.558368984 | 2 |
| P80313 | Cct7 | T-complex protein 1 subunit eta | 1455.54 | 54.41 | 59652 | 26 | 37496190 | 48713858 | 39065046 | 51724586 | 55765450 | 43689794 | 47472762 | 50487346 | 41520718 | 39513906 | 31579980 | 37564686 | 36647198 | 38271866 | 27922018 | 35419078 | 26713512 | 34341098 | 34661314 | 41800722 | 0.95669195 | 11 | 0.738881069 | 2 | 0.742962333 | 2 |
| P54311 | Gnb1 | Guanine nucleotide-binding protein G(I)/G(S)/G(T) subunit beta-1 | 382.11 | 20.88 | 37377 | 7 | 76517.09 | 1188626.38 | 1115604.88 | 2 | 162681.81 | 990081.88 | 231177.19 | 222230.75 | 126882.41 | 889543.31 | 180075.75 | 2276775.25 | 2310264.25 | 2272620.25 | 2285550 | 2901582 | 2648589.25 | 171587.8 | 4178300 | 1190268.12 | 0.967163811 | 11 | 3.666418018 | 2 | 4.360378604 | 2 |
| Q9CRB9 | Chchd3 | MICOS complex subunit Mic19 | 145.24 | 7.49 | 26335 | 2 | 757008.56 | 2 | 2 | 2 | 2469580.5 | 529840.06 | 2 | 2 | 821619.19 | 1906321.88 | 4496131 | 5620043 | 2415838.75 | 3734703 | 1129131.38 | 5602538 | 2721476.25 | 1046669.44 | 3518311.25 | 1537564.12 | 1.009666559 | 11 | 5.391394584 | 2 | 4.471140255 | 2 |
| Q921F2 | Tardbp | TAR DNA-binding protein 43 | 413.56 | 17.39 | 44548 | 6 | 5051298 | 7201042 | 917203.19 | 8291341.5 | 2688956.5 | 6548520 | 6400967 | 2100354.5 | 7754439 | 5727020.5 | 14065834 | 13013706 | 9388418 | 11387324 | 6781083.5 | 12202236 | 6333363 | 9356212 | 9114005 | 8550262 | 1.1814281 | 11 | 2.26239026 | 2 | 1.886392446 | 2 |
| P63028 | Tpt1 | Translationally-controlled tumor protein | 159.35 | 21.51 | 19462 | 3 | 632496.94 | 1163805.38 | 2201839.5 | 821358.81 | 1260006.12 | 2 | 2 | 69912.64 | 2 | 278445.5 | 2 | 2 | 888755.5 | 627035 | 690880.12 | 256722.27 | 764707.69 | 788575.75 | 2 | 2 | 0.057301382 | 0 | 0.362969351 | 3 | 0.297723119 | 2 |
| O35459 | Ech1 | Delta(3,5)-Delta(2,4)-dienoyl-CoA isomerase, mitochondrial | 276.35 | 16.21 | 36118 | 4 | 8338462 | 8295967 | 7909071.5 | 5378734 | 7024098 | 1339023.5 | 2326403.5 | 3594912.5 | 4821097 | 3781058 | 5632368 | 5147330 | 7614112 | 5267336.5 | 4753896 | 4370487 | 4753312 | 5710850.5 | 4788116 | 6006393 | 0.429338812 | 0 | 0.769089665 | 3 | 0.693686132 | 2 |
| O35639 | Anxa3 | Annexin A3 | 969.06 | 45.82 | 36384 | 16 | 27259306 | 26523176 | 25036304 | 23828364 | 23017720 | 31858110 | 25662670 | 30153740 | 23101416 | 27227932 | 41935034 | 39311054 | 30589982 | 29669408 | 25001920 | 37454066 | 32210188 | 33978170 | 38195906 | 26062976 | 1.098189717 | 7 | 1.325011501 | 3 | 1.336103766 | 2 |
| P21619 | Lmnb2 | Lamin-B2 | 379.17 | 13.09 | 67318 | 8 | 4169303.5 | 1628607 | 2943413.5 | 2732309 | 1342188.75 | 2515277.5 | 3509571.75 | 2480076.5 | 1207378.12 | 2 | 753934.44 | 776071 | 1597652.25 | 1182075.25 | 1788158.5 | 2691811.5 | 1302216.38 | 1010086 | 549936.88 | 932836.56 | 0.757837153 | 8 | 0.475809633 | 3 | 0.506162418 | 2 |
| Q8K124 | Plekho2 | Pleckstrin homology domain-containing family O member 2 | 178.98 | 8.89 | 53872 | 4 | 2390804 | 1108289.25 | 2673688.5 | 1186334.38 | 1175136.38 | 963586.19 | 1619587.5 | 3500916 | 1999446.25 | 1610797.88 | 4514432 | 1570026.25 | 2586877.5 | 3072915.25 | 3755915 | 3855035.25 | 4550292 | 1526125.38 | 3916472.5 | 2741792.5 | 1.135932386 | 11 | 1.816230066 | 3 | 1.943898146 | 2 |
| Q8VC04 | Tmem106a | Transmembrane protein 106A | 175.84 | 14.56 | 29109 | 3 | 1776360.62 | 3250930.5 | 1872025.38 | 3143917 | 2565823.5 | 5899263.5 | 5327733 | 4681551 | 7722462.5 | 7166946 | 554090.06 | 1907583.12 | 1362496.5 | 2428216 | 1583202.88 | 1578631.75 | 2515020.5 | 1758546.5 | 1001731.62 | 756134.56 | 2.442526511 | 0 | 0.621425421 | 4 | 0.603539577 | 2 |
| Q64674 | Srm | Spermidine synthase | 226.76 | 17.22 | 33995 | 4 | 5795107 | 6671999 | 5989491 | 4279145 | 5095767 | 4483482 | 4500024 | 5305569 | 2866326.25 | 2602379.75 | 6210238 | 4012922.5 | 4190058 | 3822285.75 | 3810069.5 | 5327165 | 4046118.25 | 3345587 | 2678923.5 | 2850418.5 | 0.709906926 | 4 | 0.792108461 | 4 | 0.655667368 | 2 |
| P00493 | Hprt1 | Hypoxanthine-guanine phosphoribosyltransferase | 126.49 | 8.26 | 24570 | 3 | 6627092 | 15152750 | 14653589 | 13976667 | 13879412 | 8680914 | 7890257 | 9818608 | 7660291.5 | 7672655 | 8157725 | 10190010 | 6757119.5 | 6642590 | 3641746 | 11204917 | 5331092.5 | 4193182 | 9311485 | 4898591 | 0.648981856 | 5 | 0.550466017 | 4 | 0.543467628 | 2 |
| P21956 | Mfge8 | Lactadherin | 259.54 | 14.9 | 51241 | 5 | 8325948 | 8427169 | 11303059 | 10851336 | 9635244 | 5658167 | 5604066 | 14138725 | 7612923 | 9277971 | 8553494 | 6492346.5 | 8446707 | 6773229 | 2761362.75 | 5638457 | 8270068.5 | 8994106 | 7931550 | 7251746.5 | 0.871228902 | 7 | 0.68037215 | 4 | 0.784585202 | 2 |
| G5E829 | Atp2b1 | Plasma membrane calcium-transporting ATPase 1 | 822.81 | 15.57 | 134747 | 15 | 4135646 | 7195983.5 | 4185013.5 | 6333272 | 7697583.5 | 7091795.5 | 5926226 | 8207058 | 7190165 | 9122702 | 5759155 | 3596408.5 | 2503848.25 | 6012841 | 3963900.5 | 6566650 | 8816234 | 14002555 | 8880544 | 10043484 | 1.270427224 | 7 | 0.739018677 | 4 | 1.634976545 | 2 |
| O08784 | Tcof1 | Treacle protein | 756.84 | 15 | 135001 | 15 | 16480656 | 19170086 | 8270126 | 14607075 | 12759118 | 12404963 | 12418044 | 9043475 | 19336634 | 7501395 | 9628120 | 11557820 | 7903041 | 9769896 | 8832507 | 11072328 | 5987784 | 8260422 | 4897791.5 | 11148070 | 0.851550199 | 8 | 0.669004772 | 4 | 0.580279155 | 2 |
| P97429 | Anxa4 | Annexin A4 | 1547.45 | 56.74 | 35916 | 19 | 116795306 | 161584834 | 96054538 | 75794178 | 54846330 | 131658482 | 126236218 | 129608298 | 81397642 | 88745226 | 137044930 | 155682770 | 165856370 | 138351458 | 137834274 | 146368386 | 188628162 | 179039010 | 208931874 | 145825282 | 1.10408486 | 9 | 1.454773116 | 4 | 1.720125514 | 2 |
| Q9QY76 | Vapb | Vesicle-associated membrane protein-associated protein B | 293.89 | 19.75 | 26946 | 6 | 2 | 1060304.25 | 2 | 1326100.12 | 2 | 2 | 2 | 2 | 2 | 2 | 1328850.75 | 1671131.62 | 1322898 | 3011038 | 2 | 2757720 | 1788616.25 | 1396066.25 | 339210.72 | 1486069.5 | 4.19039E-06 | 7.5 | 3.073201685 | 4.5 | 3.254965205 | 2 |
| Q9CQ22 | Lamtor1 | Regulator complex protein LAMTOR1 | 178.38 | 23.6 | 17749 | 3 | 602626.31 | 113806.23 | 672737.12 | 141168.66 | 882906.62 | 286843.78 | 205106.03 | 229184.73 | 373496.31 | 761756.12 | 98325.8 | 88993.02 | 630381.06 | 217988.72 | 2 | 2 | 250080.88 | 2 | 110911.59 | 2 | 0.769249296 | 12 | 0.429169283 | 5 | 0.14959048 | 2 |
| Q9EQH2 | Erap1 | Endoplasmic reticulum aminopeptidase 1 | 429.82 | 10.22 | 106599 | 9 | 4172599.5 | 2289690.75 | 2976944 | 6837048.5 | 8261080 | 3263968.75 | 2421891.25 | 3739057 | 7065165 | 5880967.5 | 1837220.5 | 2720313 | 3605527.25 | 3305518 | 634957.88 | 2152775.75 | 2318369 | 1171257 | 2357924.25 | 642357.25 | 0.911713689 | 12 | 0.493269662 | 5 | 0.352225434 | 2 |
| Q8CG48 | Smc2 | Structural maintenance of chromosomes protein 2 | 168.97 | 4.03 | 134239 | 4 | 791365.06 | 642979.38 | 270799.66 | 475365.22 | 1527073.88 | 2 | 135077.23 | 137344.39 | 2 | 2 | 493528.97 | 591335.75 | 2 | 567432.62 | 119463.78 | 133863.14 | 2 | 615069.44 | 2 | 2 | 0.073478491 | 0 | 0.477875485 | 6 | 0.202001827 | 2 |
| P17426 | Ap2a1 | AP-2 complex subunit alpha-1 | 747.32 | 19.96 | 107664 | 17 | 1874738.75 | 3138201.5 | 1482878.5 | 1485309.88 | 2664180.25 | 3549761.5 | 4617316 | 4699407 | 3368072 | 1837396.5 | 3379391.25 | 1961350 | 3207430.5 | 2260954.25 | 2498707.5 | 2522117.5 | 3746110 | 4523029.5 | 4109693.5 | 3924743.5 | 1.697644775 | 3 | 1.250112481 | 6 | 1.768449766 | 2 |
| Q924T2 | Mrps2 | Small ribosomal subunit protein uS2m | 147.05 | 8.59 | 32313 | 3 | 572693.81 | 157886.75 | 736555.44 | 386538 | 485594.34 | 2 | 2 | 403678.19 | 2 | 460902.09 | 2 | 395461.28 | 2 | 520097.56 | 335320.09 | 2 | 399536.28 | 2 | 2 | 2 | 0.36959688 | 4 | 0.534732552 | 6 | 0.170798823 | 2 |
| P51660 | Hsd17b4 | Peroxisomal multifunctional enzyme type 2 | 823.67 | 24.08 | 79482 | 14 | 9412022 | 12243839 | 4974717.5 | 13527073 | 19139908 | 5706372 | 4344616.5 | 8310890 | 8382862.5 | 7739732 | 9557995 | 10067171 | 6230256 | 6954089 | 4753569.5 | 9694496 | 4688102.5 | 2231365.5 | 2970547.5 | 2730009.5 | 0.581549617 | 4 | 0.633467563 | 6 | 0.376314324 | 2 |
| P80318 | Cct3 | T-complex protein 1 subunit gamma | 1716.17 | 48.99 | 60630 | 29 | 66344106 | 97237906 | 66587698 | 86394674 | 90671194 | 69727362 | 58835178 | 68214314 | 64070782 | 58366122 | 67145322 | 62053666 | 63675350 | 81119754 | 66888022 | 47032462 | 58482254 | 70419626 | 47495442 | 60043898 | 0.78385528 | 4 | 0.837063686 | 6 | 0.696092624 | 2 |
| Q3UJB9 | Edc4 | Enhancer of mRNA-decapping protein 4 | 229.39 | 4.27 | 152484 | 4 | 682731.44 | 1134807.5 | 1302500.25 | 1118666.5 | 2093876.75 | 100394.34 | 1720722.62 | 2531048.5 | 1536827.25 | 2485260.5 | 615591.75 | 2 | 776018.81 | 1558078.5 | 1071635.62 | 751530.12 | 164770.02 | 948482.81 | 604275.38 | 493288.75 | 1.322407294 | 7 | 0.635021607 | 6 | 0.467794475 | 2 |
| Q64442 | Sord | Sorbitol dehydrogenase | 96.49 | 7 | 38249 | 2 | 2299806.75 | 2 | 1516797.12 | 3737509.5 | 2136200.5 | 2 | 1624842.88 | 2396305.25 | 1568691.12 | 1333642.5 | 4014108.75 | 2826514.5 | 2590321.25 | 2080201 | 2587577.25 | 3444351.5 | 3379877.25 | 4049764.75 | 3887549.25 | 4656626 | 0.714474517 | 9.5 | 1.454929121 | 6 | 2.003873662 | 2 |
| P49717 | Mcm4 | DNA replication licensing factor MCM4 | 485.28 | 14.15 | 96736 | 10 | 3512900.25 | 2208974 | 4835831.5 | 7280042 | 8152376.5 | 2129259.5 | 42740.65 | 2061931.38 | 2843059.75 | 3702196 | 3411350 | 2382884 | 2868465.5 | 4836575 | 2269823.75 | 2112699.5 | 2032690 | 2849955.5 | 280835.56 | 2920040.5 | 0.41474166 | 3 | 0.606734239 | 7 | 0.392311363 | 2 |
| Q6Y7W8 | Gigyf2 | GRB10-interacting GYF protein 2 | 85.24 | 2.63 | 149193 | 2 | 750180.5 | 1007115.94 | 703421.56 | 771270.62 | 1046175.19 | 682501.06 | 2 | 2 | 827074 | 542113.38 | 1460790.25 | 1124892.12 | 2 | 1293086.5 | 774791.94 | 522085.53 | 769167.69 | 2 | 2 | 112479.73 | 0.479573137 | 3 | 1.087747692 | 7 | 0.32811669 | 2 |
| Q8VBZ3 | Clptm1 | Putative lipid scramblase CLPTM1 | 296.51 | 12.05 | 75291 | 5 | 1385288.75 | 2203916.5 | 771805.5 | 3569884.75 | 1969743.5 | 2290605.25 | 2470595.75 | 2718709.75 | 3852861.5 | 2740750.75 | 2183536.5 | 1983483.5 | 2975670.5 | 2507126.5 | 3116336.75 | 2421306.5 | 2928435.5 | 4579359.5 | 5038727.5 | 3635243.5 | 1.42147623 | 4 | 1.289427253 | 7 | 1.878976953 | 2 |
| Q62425 | Ndufa4 | Cytochrome c oxidase subunit NDUFA4 | 269.19 | 78.05 | 9327 | 6 | 9717470 | 2558929 | 7266871 | 5583622.5 | 7485388 | 4173039.75 | 3926813.75 | 2853380.25 | 5041305 | 4117846.5 | 3415543 | 2682483.5 | 7207803.5 | 2604042.5 | 6057281 | 1861697.75 | 1706565.25 | 4448245 | 2590307.5 | 1613801.25 | 0.616712016 | 5 | 0.673585323 | 7 | 0.374724385 | 2 |
| Q8VEB4 | Pla2g15 | Lysosomal phospholipase A and acyltransferase | 40.73 | 2.67 | 47307 | 1 | 1842629.25 | 868697.25 | 1126804 | 467568.41 | 663367.69 | 2 | 399326.03 | 591773.44 | 546696.56 | 1284477.62 | 1051548.75 | 569679.25 | 699566.94 | 528462.25 | 2 | 2 | 2 | 866038.25 | 2 | 2 | 0.567968972 | 6 | 0.573399276 | 7 | 0.174287511 | 2 |
| Q8R127 | Sccpdh | Saccharopine dehydrogenase-like oxidoreductase | 205.97 | 16.32 | 47129 | 4 | 77433.12 | 691231.62 | 39167.83 | 661971.31 | 45029.38 | 2 | 171809.33 | 2 | 609689.5 | 83609.46 | 631670.38 | 2 | 34169.57 | 669722.06 | 2 | 64467.11 | 2 | 2 | 2 | 2 | 0.571094069 | 9 | 0.881658758 | 7 | 0.042562513 | 2 |
| Q9D0I8 | Mrto4 | mRNA turnover protein 4 homolog | 117.99 | 14.23 | 27546 | 3 | 757628.06 | 1084293.88 | 1382130 | 1354883.62 | 2210032.5 | 810601.06 | 642132.06 | 1956985.38 | 1604708.25 | 1226849.25 | 1279479 | 567848.5 | 1068001.62 | 1481956.5 | 552462.56 | 846133.75 | 1015396.06 | 2 | 84910.86 | 97440.95 | 0.919326169 | 11 | 0.729086974 | 7 | 0.301059543 | 2 |
| Q9D892 | Itpa | Inosine triphosphate pyrophosphatase | 129.93 | 14.14 | 21897 | 3 | 1457228.38 | 1331999.88 | 450398.84 | 961818.62 | 2143054 | 2252746.5 | 2171167.5 | 1974389.38 | 1377916 | 1043222.69 | 874691.62 | 1648771.38 | 677247 | 810205.06 | 698683.06 | 2 | 601340.19 | 625035.38 | 2 | 2 | 1.390092593 | 6 | 0.742311975 | 8 | 0.193298388 | 2 |
| E9Q7G0 | Numa1 | Nuclear mitotic apparatus protein 1 | 1414.55 | 14.28 | 235630 | 25 | 3593393.75 | 3429121.5 | 3600044 | 2191039 | 7698624.5 | 4351486 | 3739518.5 | 2412948.75 | 4432301 | 5022292 | 4409390 | 5832565 | 3744047 | 4580105.5 | 2815220.75 | 2295605 | 672297.25 | 975291.75 | 841312.12 | 2791614 | 0.973007484 | 8 | 1.042370128 | 8 | 0.369346619 | 2 |
| Q8BU30 | Iars1 | Isoleucine--tRNA ligase, cytoplasmic | 358.88 | 5.31 | 144271 | 8 | 3628220 | 4470157 | 7016193 | 5277355.5 | 6110085 | 4526498 | 5788787.5 | 7741787 | 4927516.5 | 8039221.5 | 2212312.75 | 2340377.5 | 4766107.5 | 4669551.5 | 6190179 | 3706931.5 | 3955375.5 | 3403219 | 3087055.5 | 2975305.75 | 1.170621018 | 8 | 0.761396131 | 8 | 0.646286336 | 2 |
| P68037 | Ube2l3 | Ubiquitin-conjugating enzyme E2 L3 | 137.77 | 35.71 | 17862 | 3 | 722612.75 | 80998.12 | 1421456.88 | 105207.96 | 695038.38 | 2 | 2 | 2 | 2 | 929534.88 | 1012053.19 | 111069.36 | 1047158.38 | 935646.56 | 615516 | 1479330.62 | 1513641.12 | 992479.31 | 1900461.62 | 959859.12 | 0.307255 | 4 | 1.23010153 | 9 | 2.262830102 | 2 |
| O55023 | Impa1 | Inositol monophosphatase 1 | 479.53 | 27.08 | 30436 | 10 | 10867726 | 10868682 | 9837946 | 3703185.25 | 8028560 | 4991466 | 6187442 | 8622216 | 5581928.5 | 4602105 | 9779748 | 9818879 | 8875365 | 9268047 | 7563485 | 11730638 | 10468713 | 14370735 | 14189176 | 12830838 | 0.692400332 | 6 | 1.046169588 | 9 | 1.468386696 | 2 |
| Q9D0M3 | Cyc1 | Cytochrome c1, heme protein, mitochondrial | 220.08 | 21.23 | 35328 | 5 | 1609983.12 | 2611508.5 | 731450.31 | 2735437.75 | 2969240.75 | 2991104.75 | 2533706.75 | 3948920.5 | 2495140.25 | 2801165.25 | 2746327.5 | 2216850.75 | 2 | 3316280.5 | 3307633.5 | 2667254.25 | 3823782 | 4227659.5 | 2981094.25 | 3458064.5 | 1.385866348 | 7 | 1.087212134 | 9 | 1.609914203 | 2 |
| Q8VBW6 | Nae1 | NEDD8-activating enzyme E1 regulatory subunit | 47.09 | 1.87 | 60274 | 1 | 941643.75 | 199879.64 | 172591.36 | 187883.06 | 75477.48 | 104805.47 | 311894.97 | 307387.41 | 389392.97 | 278488.06 | 95502.22 | 195449.42 | 324268.31 | 281874.59 | 443547.88 | 2 | 142548.69 | 108748.75 | 2 | 2 | 0.882402969 | 8 | 0.849865877 | 9 | 0.15930737 | 2 |
| B1H268 | Mcmbp | Mini-chromosome maintenance complex-binding protein | 61.43 | 1.71 | 73007 | 1 | 705141.25 | 1091027.75 | 107445.52 | 1137816.88 | 153762.58 | 766078.06 | 2 | 2 | 791844.25 | 893418.19 | 1029017.62 | 1130249.25 | 920719.44 | 965537.69 | 870311.69 | 1234780.88 | 1166791.12 | 979340.56 | 1169090.62 | 1151531.5 | 0.767197396 | 9 | 1.538509311 | 9 | 1.784409559 | 2 |
| Q8CCF0 | Prpf31 | U4/U6 small nuclear ribonucleoprotein Prp31 | 178.39 | 12.42 | 55430 | 4 | 981284.94 | 2504893 | 2006073 | 1947242.88 | 3148901.5 | 610942.75 | 1782674.5 | 2484621.25 | 2219221.5 | 1972252 | 1813229.88 | 1302277.88 | 2457180 | 2364413.75 | 1431942.12 | 2 | 2 | 1728739.5 | 1285759.62 | 2 | 0.856570965 | 9 | 0.884840748 | 9 | 0.284698959 | 2 |
| P51175 | Ppox | Protoporphyrinogen oxidase | 70.99 | 4.61 | 50871 | 2 | 1058248.62 | 925482 | 1038461.44 | 637701.81 | 963977.25 | 660151.5 | 876121 | 1445507 | 1811251.88 | 1042888.06 | 813688.88 | 621901.62 | 1316180 | 1480951.12 | 1246317 | 1113153.25 | 1096946.38 | 1272297.5 | 1298906.5 | 999821.31 | 1.262128483 | 9 | 1.184946223 | 9 | 1.250278131 | 2 |
| P56812 | Pdcd5 | Programmed cell death protein 5 | 283.95 | 45.24 | 14275 | 5 | 4278239 | 5088320 | 4791848 | 3295464 | 4944864.5 | 3720682.25 | 2826819 | 3992262.5 | 2060321.25 | 1899999.25 | 4074997 | 4280372 | 4978882.5 | 4372867 | 3892417.5 | 3421918 | 1751363.88 | 1670969.75 | 3356035 | 1459009.75 | 0.647361734 | 2 | 0.964319437 | 10 | 0.520533687 | 2 |
| Q8BMS1 | Hadha | Trifunctional enzyme subunit alpha, mitochondrial | 1022.83 | 33.55 | 82670 | 19 | 8553474 | 6586773 | 12173214 | 7117688 | 12133054 | 5131094 | 6313278 | 8996370 | 8415125 | 9746682 | 5014700.5 | 7983497.5 | 7068162.5 | 7483063.5 | 12859570 | 5765869.5 | 7869768 | 4330978.5 | 6133502.5 | 5076732 | 0.829017711 | 8 | 0.867812427 | 10 | 0.626594006 | 2 |
| O70133 | Dhx9 | ATP-dependent RNA helicase A | 966.91 | 15.22 | 149475 | 20 | 12296841 | 25595374 | 14642867 | 22321210 | 19093802 | 16504436 | 14519060 | 13204253 | 18110976 | 15829535 | 16459609 | 19301276 | 18026124 | 19309538 | 10822664 | 15540193 | 10347514 | 11487431 | 9355492 | 9744599 | 0.832018965 | 8 | 0.893231794 | 10 | 0.601119452 | 2 |
| P41105 | Rpl28 | Large ribosomal subunit protein eL28 | 325.22 | 28.47 | 15733 | 6 | 1148092.5 | 407991.69 | 881119.62 | 837407.5 | 37646.61 | 669253.56 | 2481803.25 | 196410.45 | 2570499.5 | 1018719.94 | 864297.06 | 2723562 | 2 | 2 | 523827.31 | 2 | 654058.06 | 2 | 2 | 2 | 2.094247147 | 8 | 1.241355737 | 10 | 0.197468336 | 2 |
| Q7TMK9 | Syncrip | Heterogeneous nuclear ribonucleoprotein Q | 300.62 | 10.11 | 69633 | 6 | 6656376 | 5594501 | 13198741 | 4490648.5 | 9681693 | 6710556 | 7579261 | 7656460.5 | 6251302.5 | 7621582 | 9518681 | 10104192 | 9791226 | 7006836 | 5230614 | 5185066.5 | 2811431 | 4351424.5 | 4501514 | 2662731.5 | 0.904022983 | 11 | 1.051223855 | 10 | 0.492458418 | 2 |
| Q61166 | Mapre1 | Microtubule-associated protein RP/EB family member 1 | 269.41 | 23.88 | 30016 | 5 | 6272556 | 15935866 | 5912440 | 8280878 | 13695462 | 10312074 | 10533020 | 11421521 | 9838128 | 7224735 | 8860180 | 10310043 | 6693497.5 | 7435832 | 3378132.75 | 7717715.5 | 4566370 | 3492483.5 | 4774758.5 | 2585413.25 | 0.984675312 | 11 | 0.732130414 | 10 | 0.461836985 | 2 |
| Q3U5Q7 | Cmpk2 | UMP-CMP kinase 2, mitochondrial | 372.89 | 16.78 | 50036 | 7 | 1301941.75 | 110944.92 | 1007595.31 | 2 | 383668.5 | 4200025 | 6962541 | 3868402.5 | 3923066.25 | 4164663.5 | 709206.31 | 581709 | 1453868 | 416213.47 | 2 | 1451886.75 | 1322394.5 | 2197981.5 | 1498587.75 | 560031.88 | 8.24445119 | 0 | 1.127256382 | 10.5 | 2.507311008 | 2 |
| Q9EQH3 | Vps35 | Vacuolar protein sorting-associated protein 35 | 223.64 | 6.03 | 91713 | 4 | 4160166.5 | 6535273.5 | 6433009 | 4870731.5 | 7834345 | 6543879 | 5818917 | 8761752 | 5547712.5 | 7842438 | 6586465 | 6561861 | 5284844 | 4408705 | 780376.44 | 5656460 | 3212909.5 | 2 | 2513185.5 | 2173468.25 | 1.156909816 | 7 | 0.791802211 | 11 | 0.454388981 | 2 |
| Q6PAR5 | Gapvd1 | GTPase-activating protein and VPS9 domain-containing protein 1 | 109.62 | 2.4 | 162402 | 2 | 1377230 | 292540.06 | 913777.69 | 405275.41 | 903703.88 | 214007.91 | 1063378.75 | 141528.53 | 479522.91 | 861756.75 | 362441.69 | 2 | 1122682.38 | 671313.75 | 958478.06 | 91792.41 | 2 | 214235.33 | 2 | 503630.03 | 0.709101009 | 8 | 0.800230248 | 11 | 0.208004148 | 2 |
| P67871 | Csnk2b | Casein kinase II subunit beta | 54.89 | 5.12 | 24942 | 1 | 1526465.62 | 1589488 | 1894390.75 | 1304467.5 | 1124134 | 2 | 885596.88 | 1457158.75 | 2 | 1096922.62 | 1451257.88 | 1336205.88 | 1477540.38 | 1437904 | 1635350 | 848538.5 | 1274875.5 | 1258747.75 | 1112910.12 | 1121203.5 | 0.462388396 | 2 | 0.986464785 | 12 | 0.754982691 | 2 |
| Q8CD92 | Ttc27 | Tetratricopeptide repeat protein 27 | 39.46 | 1.06 | 96431 | 1 | 1616956.62 | 549734.44 | 1431277.5 | 2 | 534346.62 | 2 | 457092.16 | 2 | 2 | 2 | 2 | 772410.06 | 668951.75 | 437457.12 | 2359938 | 1477134.38 | 2606656.25 | 2013422.12 | 1547800.12 | 4136664.75 | 0.110615943 | 3 | 1.025758369 | 12.5 | 2.851106802 | 2 |
