## supplemental tables 5 to 7 for "Effects of manganese dioxide on macrophages under different exposure schemes"

Supplementary Table 5: Results of the pathway analysis by the David tool on the proteins modulated in the recovery condition

Annotation Cluster 1 — Enrichment Score: 8.23823913508244

| Category | Term | Count | P Value | Genes | FDR |
|---|---|---|---|---|---|
| GOTERM_MF_DIRECT | GO:0005524~ATP binding | 37 | 1.9519E-12 | P54775, Q8CG48, Q8CG47, P38647, Q9R0N0, O08528, Q9D2Y4, Q8R5C5, Q9DCL9, Q922D8, P36371, P49718, P63085, P16460, P25206, P52480, O88351, Q8CGK3, Q9D0F6, P80315, Q9Z1Q9, Q9JIA7, P97310, Q3TRM8, P97311, Q3U5Q7, Q8JZQ2, P58389, P97855, Q01320, Q61881, P42932, Q8BP47, Q9Z110, P63017, P43247, Q6ZPJ3 | 4.8993E-10 |
| GOTERM_MF_DIRECT | GO:0016887~ATP hydrolysis activity | 18 | 8.44552E-10 | P80315, P54775, Q8CG48, Q8CG47, P97310, P97311, P38647, Q8JZQ2, P97855, Q61881, P36371, P49718, P42932, P25206, P63017, P43247, Q8CGK3, Q9D0F6 | 1.4132E-07 |
| UP_KW_LIGAND | KW-0547~Nucleotide-binding | 48 | 7.08749E-08 | Q9D0S9, P54775, Q8CG48, Q8CG47, Q60766, Q922F4, P38647, P35279, Q9R0N0, O08528, Q9D2Y4, Q8R5C5, Q9DCL9, Q922D8, P35283, P63001, P35282, P36371, P49718, P63085, P16460, P25206, P52480, O88351, Q8CGK3, Q9D0F6, P80315, Q9Z1Q9, Q9JIA7, P97310, Q3TRM8, P97311, Q3U5Q7, Q8JZQ2, P58389, P97855, Q91VE0, Q01320, Q61881, P84096, P42932, Q8BP47, Q91V41, Q9Z110, Q8BH04, P63017, P43247, Q6ZPJ3 | 1.5592E-06 |
| UP_KW_LIGAND | KW-0067~ATP-binding | 37 | 9.53822E-06 | P54775, Q8CG48, Q8CG47, P38647, Q9R0N0, O08528, Q9D2Y4, Q8R5C5, Q9DCL9, Q922D8, P36371, P49718, P63085, P16460, P25206, P52480, O88351, Q8CGK3, Q9D0F6, P80315, Q9Z1Q9, Q9JIA7, P97310, Q3TRM8, P97311, Q3U5Q7, Q8JZQ2, P58389, P97855, Q01320, Q61881, P42932, Q8BP47, Q9Z110, P63017, P43247, Q6ZPJ3 | 0.00010492 |

Annotation Cluster 2 — Enrichment Score: 7.423009135113517

| Category | Term | Count | P Value | Genes | FDR |
|---|---|---|---|---|---|
| GOTERM_CC_DIRECT | GO:0005739~mitochondrion | 43 | 1.36339E-14 | P54116, Q9D0S9, Q60766, Q9CZW5, P99029, P38647, O08528, Q91WD5, Q64521, Q8JZN5, Q922D8, Q9DCN2, O35129, P63085, P09671, Q9QUH0, P62075, P16460, Q7TMY8, P52480, Q8CGK3, Q91YT0, P08228, Q9Z1Q9, Q9JIA7, Q9QZD8, Q3TRM8, Q3U5Q7, Q9D338, Q8JZQ2, Q8BMD8, Q99L13, P35700, O35459, P67778, Q91VD9, Q9Z0X1, Q8BP47, O88986, P06151, Q9Z110, O70404, Q8BH04 | 1.4997E-12 |
| GOTERM_CC_DIRECT | GO:0005743~mitochondrial inner membrane | 19 | 4.08575E-09 | Q9JIA7, Q9QZD8, Q9D338, Q8JZQ2, Q8BMD8, Q64521, Q91WD5, Q8JZN5, Q9DCN2, O35129, P67778, Q91VD9, Q9Z0X1, P16110, P09671, O88986, P62075, Q9Z110, Q91YT0 | 3.3707E-07 |
| UP_KW_DOMAIN | KW-0809~Transit peptide | 18 | 5.52203E-07 | Q9D0S9, Q3U5Q7, P99029, Q9D338, P38647, Q8JZQ2, Q99L13, Q64521, Q91WD5, Q8JZN5, O35459, Q91VD9, Q9Z0X1, P09671, O88986, Q8BH04, Q8CGK3, Q91YT0 | 9.9397E-06 |
| UP_SEQ_FEATURE | TRANSIT:Mitochondrion | 18 | 1.2652E-06 | Q9D0S9, Q3U5Q7, P99029, Q9D338, P38647, Q8JZQ2, Q99L13, Q64521, Q91WD5, Q8JZN5, O35459, Q91VD9, Q9Z0X1, P09671, O88986, Q8BH04, Q8CGK3, Q91YT0 | 0.00032052 |
| UP_KW_CELLULAR_COMPONENT | KW-0496~Mitochondrion | 30 | 6.18028E-06 | Q9D0S9, Q60766, Q9CZW5, P99029, P38647, O08528, Q91WD5, Q64521, Q8JZN5, Q9DCN2, O35129, P09671, P62075, Q7TMY8, Q8CGK3, Q91YT0, Q9JIA7, Q9QZD8, Q3U5Q7, Q9D338, Q8JZQ2, Q8BMD8, Q99L13, O35459, P67778, Q91VD9, Q9Z0X1, O88986, Q9Z110, Q8BH04 | 8.0344E-05 |
| UP_KW_CELLULAR_COMPONENT | KW-0999~Mitochondrion inner membrane | 14 | 1.20445E-05 | Q9JIA7, Q9QZD8, Q8JZQ2, Q8BMD8, Q64521, Q91WD5, Q8JZN5, O35129, P67778, Q91VD9, Q9Z0X1, P62075, Q9Z110, Q91YT0 | 0.00010439 |

Annotation Cluster 3 — Enrichment Score: 4.6721207488658845

| Category | Term | Count | P Value | Genes | FDR |
|---|---|---|---|---|---|
| GOTERM_MF_DIRECT | GO:0016887~ATP hydrolysis activity | 18 | 8.44552E-10 | P80315, P54775, Q8CG48, Q8CG47, P97310, P97311, P38647, Q8JZQ2, P97855, Q61881, P36371, P49718, P42932, P25206, P63017, P43247, Q8CGK3, Q9D0F6 | 1.4132E-07 |
| GOTERM_BP_DIRECT | GO:0006279~premeiotic DNA replication | 5 | 1.7939E-08 | P49718, P97310, P97311, P25206, Q61881 | 2.7752E-05 |
| UP_SEQ_FEATURE | MOTIF:Arginine finger | 5 | 5.40813E-08 | P49718, P97310, P97311, P25206, Q61881 | 4.1102E-05 |
| INTERPRO | IPR018525:MCM_CS | 5 | 5.86337E-08 | P49718, P97310, P97311, P25206, Q61881 | 2.2838E-05 |
| INTERPRO | IPR027925:MCM_N | 5 | 5.86337E-08 | P49718, P97310, P97311, P25206, Q61881 | 2.2838E-05 |
| INTERPRO | IPR033762:MCM_OB | 5 | 2.70213E-07 | P49718, P97310, P97311, P25206, Q61881 | 5.2624E-05 |
| INTERPRO | IPR001208:MCM_dom | 5 | 2.70213E-07 | P49718, P97310, P97311, P25206, Q61881 | 5.2624E-05 |
| GOTERM_CC_DIRECT | GO:0071162~CMG complex | 5 | 3.77914E-07 | P49718, P97310, P97311, P25206, Q61881 | 1.5589E-05 |
| GOTERM_CC_DIRECT | GO:0042555~MCM complex | 5 | 3.77914E-07 | P49718, P97310, P97311, P25206, Q61881 | 1.5589E-05 |
| GOTERM_MF_DIRECT | GO:0003697~single-stranded DNA binding | 9 | 3.78876E-07 | P49718, Q8CG48, Q8CG47, P97310, P97311, P25206, Q61881, P43247, Q8CGK3 | 2.7171E-05 |
| UP_SEQ_FEATURE | DOMAIN:MCM | 5 | 4.45987E-07 | P49718, P97310, P97311, P25206, Q61881 | 0.00016947 |
| INTERPRO | IPR031327:MCM | 5 | 4.83343E-07 | P49718, P97310, P97311, P25206, Q61881 | 5.3789E-05 |
| INTERPRO | IPR041562:MCM_lid | 5 | 4.83343E-07 | P49718, P97310, P97311, P25206, Q61881 | 5.3789E-05 |
| SMART | SM00350:MCM | 5 | 5.70628E-07 | P49718, P97310, P97311, P25206, Q61881 | 5.3639E-05 |
| GOTERM_BP_DIRECT | GO:0000727~double-strand break repair via break-induced re | 5 | 5.75677E-07 | P49718, P97310, P97311, P25206, Q61881 | 0.00017811 |
| GOTERM_MF_DIRECT | GO:0061749~forked DNA-dependent helicase activity | 6 | 6.65982E-06 | P49718, P97310, P97311, P25206, P97855, Q61881 | 0.00031191 |
| GOTERM_MF_DIRECT | GO:0017116~single-stranded DNA helicase activity | 5 | 6.97287E-06 | P49718, P97310, P25206, Q61881, Q9D0F6 | 0.00031191 |
| GOTERM_MF_DIRECT | GO:1990518~single-stranded 3'-5' DNA helicase activity | 6 | 7.45599E-06 | P49718, P97310, P97311, P25206, P97855, Q61881 | 0.00031191 |
| GOTERM_MF_DIRECT | GO:0009378~four-way junction helicase activity | 6 | 7.45599E-06 | P49718, P97310, P97311, P25206, P97855, Q61881 | 0.00031191 |
| KEGG_PATHWAY | mmu03030:DNA replication | 7 | 7.82982E-06 | P49718, P97310, P97311, P25206, P17918, Q61881, Q9D0F6 | 0.0018635 |
| GOTERM_BP_DIRECT | GO:0006268~DNA unwinding involved in DNA replication | 5 | 8.12071E-06 | P49718, P97310, P97311, P25206, Q61881 | 0.00157034 |
| GOTERM_MF_DIRECT | GO:0036121~double-stranded DNA helicase activity | 6 | 8.32472E-06 | P49718, P97310, P97311, P25206, P97855, Q61881 | 0.00032146 |
| GOTERM_BP_DIRECT | GO:0006270~DNA replication initiation | 5 | 1.3849E-05 | P49718, P97310, P97311, P25206, Q61881 | 0.00238049 |
| INTERPRO | IPR012340:NA-bd_OB-fold | 8 | 2.96293E-05 | P54775, P49718, P97310, Q8BP47, P97311, P25206, Q61881, P0DOV2 | 0.00230812 |
| GOTERM_MF_DIRECT | GO:0003689~DNA clamp loader activity | 8 | 3.75061E-05 | P49718, P97310, P97311, Q8JZQ2, P25206, P97855, Q61881, Q9D0F6 | 0.00134486 |
| GOTERM_MF_DIRECT | GO:0140584~chromatin extrusion motor activity | 7 | 0.00022554 | P49718, P97310, P97311, Q8JZQ2, P25206, P97855, Q61881 | 0.0063495 |
| GOTERM_MF_DIRECT | GO:0140665~ATP-dependent H3-H4 histone complex chaper | 7 | 0.00022554 | P49718, P97310, P97311, Q8JZQ2, P25206, P97855, Q61881 | 0.0063495 |
| GOTERM_MF_DIRECT | GO:0140849~ATP-dependent H2AZ histone chaperone activit | 7 | 0.00024328 | P49718, P97310, P97311, Q8JZQ2, P25206, P97855, Q61881 | 0.0063495 |
| GOTERM_BP_DIRECT | GO:0140588~chromatin looping | 7 | 0.00024722 | P49718, P97310, P97311, Q8JZQ2, P25206, P97855, Q61881 | 0.02731818 |
| GOTERM_MF_DIRECT | GO:0061775~cohesin loader activity | 7 | 0.00025251 | P49718, P97310, P97311, Q8JZQ2, P25206, P97855, Q61881 | 0.0063495 |
| UP_KW_BIOLOGICAL_PROCESS | KW-0235~DNA replication | 7 | 0.00030489 | P49718, P97310, P97311, P25206, P17918, Q61881, Q9D0F6 | 0.01204298 |
| GOTERM_BP_DIRECT | GO:0032508~DNA duplex unwinding | 5 | 0.00077215 | P49718, P97310, P25206, P97855, Q9D0F6 | 0.06286959 |
| GOTERM_CC_DIRECT | GO:0005694~chromosome | 8 | 0.00133179 | P49718, Q8CG48, Q8CG47, P97310, P97311, P25206, Q61881, P43247 | 0.02197455 |
| UP_KW_CELLULAR_COMPONENT | KW-0158~Chromosome | 13 | 0.0109113 | P46061, Q8CG48, Q8BFQ4, Q8CG47, P97310, P97311, Q61033, Q61881, F6ZDS4, P49718, Q8BRT1, P25206, P43247 | 0.0472823 |
| UP_KW_MOLECULAR_FUNCTION | KW-0347~Helicase | 6 | 0.01128367 | P49718, P97310, P97311, P25206, P97855, Q61881 | 0.11057992 |
| KEGG_PATHWAY | mmu04110:Cell cycle | 7 | 0.02003318 | P49718, P97310, P97311, P25206, P68254, P17918, Q61881 | 0.3925898 |
| GOTERM_MF_DIRECT | GO:0003678~DNA helicase activity | 3 | 0.02192404 | P97311, P97855, Q61881 | 0.24457482 |
| GOTERM_MF_DIRECT | GO:0016787~hydrolase activity | 6 | 0.03050611 | Q9D0S9, P49718, Q60766, P97311, P20060, Q61881 | 0.28359379 |
| UP_KW_BIOLOGICAL_PROCESS | KW-0131~Cell cycle | 12 | 0.06624186 | P49718, Q8CG48, Q8CG47, P97310, P63085, P97311, Q8BRT1, P25206, Q61881, P61290, F6ZDS4, P43247 | 0.49173593 |

| | | | | | |
|---|---|---|---|---|---|
| GOTERM_MF_DIRECT | GO:0004386~helicase activity | 3 | 0.07848544 | P49718, P97310, P25206 | 0.51168428 |
| GOTERM_BP_DIRECT | GO:0006338~chromatin remodeling | 9 | 0.16667913 | P49718, P97311, Q7TMY8, Q8JZQ2, P25206, P52480, P97855, Q61881, O88351 | 0.98723676 |
| Annotation Cluster 4 | Enrichment Score: 3.7998239042725745 | | | | |
| Category | Term | Count | P Value | Genes | FDR |
| GOTERM_MF_DIRECT | GO:0016887~ATP hydrolysis activity | 18 | 8.44552E-10 | P80315, P54775, Q8CG48, Q8CG47, P97310, P97311, P38647, Q8JZQ2, P97855, Q61881, P36371, P49718, P42932, P25206, P63017, P43247, Q8CGK3, Q9D0F6 | 1.4132E-07 |
| GOTERM_MF_DIRECT | GO:0003689~DNA clamp loader activity | 8 | 3.75061E-05 | P49718, P97310, P97311, Q8JZQ2, P25206, P97855, Q61881, Q9D0F6 | 0.00134486 |
| INTERPRO | IPR003593:AAA+_ATPase | 7 | 0.00115859 | P36371, P54775, Q8JZQ2, P25206, Q61881, Q8CGK3, Q9D0F6 | 0.06446743 |
| SMART | SM00382:AAA | 7 | 0.00141426 | P36371, P54775, Q8JZQ2, P25206, Q61881, Q8CGK3, Q9D0F6 | 0.04431343 |
| INTERPRO | IPR003959:ATPase_AAA_core | 4 | 0.00630315 | P54775, Q8JZQ2, Q8CGK3, Q9D0F6 | 0.21348495 |
| UP_SEQ_FEATURE | DOMAIN:AAA+ ATPase | 3 | 0.04856363 | P54775, Q8JZQ2, Q9D0F6 | 0.92270901 |
| Annotation Cluster 5 | Enrichment Score: 3.6228720527730167 | | | | |
| Category | Term | Count | P Value | Genes | FDR |
| UP_KW_BIOLOGICAL_PROCESS | KW-0653~Protein transport | 20 | 9.86745E-06 | Q9ERK4, Q8BFY9, Q9QY81, P35279, Q64310, Q9CR60, P70168, Q8VI75, F6ZDS4, P35283, P36371, P35282, Q91YE6, O35643, P62075, Q9D906, O35609, Q91V41, Q8BPX9, O70404 | 0.00077953 |
| GOTERM_BP_DIRECT | GO:0015031~protein transport | 12 | 0.00013931 | P35283, P35282, P36371, P62075, Q9QY81, Q9D906, O35609, Q91V41, Q64310, Q8BPX9, Q9CR60, O70404 | 0.01657837 |
| UP_KW_BIOLOGICAL_PROCESS | KW-0813~Transport | 32 | 0.00984428 | Q9ERK4, Q8BVE3, P35279, Q3UBX0, Q9CR60, P70168, Q91WD5, F6ZDS4, P35283, P35282, P36371, Q91YE6, O35643, Q9QUH0, P62075, P84104, O35609, Q8BPX9, Q91YT0, Q9QZD8, Q8BFY9, Q9QY81, Q64310, Q8BMD8, P97855, Q91VE0, Q8R180, Q8VI75, Q91VD9, Q9D906, Q91V41, O70404 | 0.1555397 |
| Annotation Cluster 6 | Enrichment Score: 3.48762555579782 | | | | |
| Category | Term | Count | P Value | Genes | FDR |
| GOTERM_CC_DIRECT | GO:0005635~nuclear envelope | 10 | 8.37574E-06 | P46061, Q8VCH8, Q91YE6, Q9ERK4, Q9QY81, Q80W54, P70168, Q8BJS4, Q61033, F6ZDS4 | 0.00025127 |
| GOTERM_BP_DIRECT | GO:0006913~nucleocytoplasmic transport | 6 | 3.73001E-05 | P46061, Q91YE6, Q9ERK4, Q9QY81, Q8VI75, F6ZDS4 | 0.00577033 |
| KEGG_PATHWAY | mmu03013:Nucleocytoplasmic transport | 8 | 0.00110108 | P46061, Q91YE6, Q9ERK4, Q8BFY9, Q9QY81, P70168, Q8VI75, F6ZDS4 | 0.0655145 |
| GOTERM_CC_DIRECT | GO:0031965~nuclear membrane | 8 | 0.00122388 | P46061, Q9QY81, Q3UBX0, Q64310, P70168, Q8BJS4, Q61033, F6ZDS4 | 0.02125686 |
| GOTERM_CC_DIRECT | GO:0005643~nuclear pore | 4 | 0.00866123 | P46061, Q9QY81, P70168, F6ZDS4 | 0.08931896 |
| Annotation Cluster 7 | Enrichment Score: 3.347517746947036 | | | | |
| Category | Term | Count | P Value | Genes | FDR |
| GOTERM_MF_DIRECT | GO:0031267~small GTPase binding | 13 | 3.13546E-07 | P46061, P08228, Q9ERK4, O70145, Q8BFY9, Q9JKF1, P70168, Q8VI75, Q62433, Q9JL26, P63001, Q91YE6, Q7TMB8 | 2.6233E-05 |
| GOTERM_BP_DIRECT | GO:0006606~protein import into nucleus | 9 | 5.58063E-07 | O35129, Q91YE6, Q9ERK4, Q8BFY9, P34960, P70168, Q8VI75, P63017, F6ZDS4 | 0.00017811 |
| SMART | SM00913:IBN_N | 5 | 7.87876E-06 | Q91YE6, Q9ERK4, Q8BFY9, P70168, Q8VI75 | 0.0003703 |
| UP_SEQ_FEATURE | DOMAIN:Importin N-terminal | 5 | 8.02064E-06 | Q91YE6, Q9ERK4, Q8BFY9, P70168, Q8VI75 | 0.00144472 |
| INTERPRO | IPR001494:Importin-beta_N | 5 | 8.68357E-06 | Q91YE6, Q9ERK4, Q8BFY9, P70168, Q8VI75 | 0.00084556 |
| INTERPRO | IPR011989:ARM-like | 11 | 2.28667E-05 | Q9JL26, Q91YE6, O35643, Q9ERK4, Q8BVE3, Q8BFY9, Q7TMY8, Q8BRT1, P70168, Q9DBR3, Q8VI75 | 0.00197924 |
| GOTERM_BP_DIRECT | GO:0006913~nucleocytoplasmic transport | 6 | 3.73001E-05 | P46061, Q91YE6, Q9ERK4, Q9QY81, Q8VI75, F6ZDS4 | 0.00577033 |
| INTERPRO | IPR016024:ARM-type_fold | 12 | 0.00015268 | Q9JL26, Q91YE6, O35643, Q9ERK4, Q8C147, Q8BVE3, Q8BFY9, Q7TMY8, Q8BRT1, P70168, Q9DBR3, Q8VI75 | 0.00991126 |
| GOTERM_MF_DIRECT | GO:0061608~nuclear import signal receptor activity | 4 | 0.00022581 | Q91YE6, Q8BFY9, P70168, Q8VI75 | 0.0063495 |
| INTERPRO | IPR040122:Importin_beta | 3 | 0.00093444 | Q8BFY9, P70168, Q8VI75 | 0.05599446 |
| KEGG_PATHWAY | mmu03013:Nucleocytoplasmic transport | 8 | 0.00110108 | P46061, Q91YE6, Q9ERK4, Q8BFY9, Q9QY81, P70168, Q8VI75, F6ZDS4 | 0.0655145 |
| UP_SEQ_FEATURE | REPEAT:HEAT 6 | 4 | 0.00217383 | Q8BFY9, Q8BRT1, P70168, Q8VI75 | 0.1652109 |
| UP_SEQ_FEATURE | REPEAT:HEAT 5 | 4 | 0.00302564 | Q8BFY9, Q8BRT1, P70168, Q8VI75 | 0.2090443 |
| UP_SEQ_FEATURE | REPEAT:HEAT 4 | 4 | 0.00561469 | Q8BFY9, Q8BRT1, P70168, Q8VI75 | 0.30479743 |
| UP_SEQ_FEATURE | REPEAT:HEAT 3 | 4 | 0.00708606 | Q8BFY9, Q8BRT1, P70168, Q8VI75 | 0.3590269 |
| GOTERM_MF_DIRECT | GO:0008139~nuclear localization sequence binding | 3 | 0.00984441 | Q8BFY9, P70168, Q8VI75 | 0.12925378 |
| UP_SEQ_FEATURE | REPEAT:HEAT 1 | 4 | 0.0111666 | Q8BFY9, Q8BRT1, P70168, Q8VI75 | 0.47147872 |
| UP_SEQ_FEATURE | REPEAT:HEAT 2 | 4 | 0.0111666 | Q8BFY9, Q8BRT1, P70168, Q8VI75 | 0.47147872 |
| UP_SEQ_FEATURE | REPEAT:HEAT 8 | 3 | 0.01628821 | Q8BFY9, Q8BRT1, P70168 | 0.61895185 |
| UP_SEQ_FEATURE | REPEAT:HEAT 7 | 3 | 0.02021137 | Q8BFY9, Q8BRT1, P70168 | 0.71382497 |
| COG_ONTOLOGY | Intracellular trafficking and secretion | 3 | 0.16394992 | Q8BFY9, P70168, Q8VI75 | 0.81974959 |
| Annotation Cluster 8 | Enrichment Score: 2.760926075510565 | | | | |
| Category | Term | Count | P Value | Genes | FDR |
| INTERPRO | IPR036013:Band_7/SPFH_dom_sf | 4 | 0.00013568 | P67778, O35129, P54116, Q9EQK5 | 0.00960853 |
| UP_SEQ_FEATURE | DOMAIN:Band 7 | 3 | 0.00383047 | P67778, O35129, P54116 | 0.24259645 |
| INTERPRO | IPR001107:Band_7 | 3 | 0.00398367 | P67778, O35129, P54116 | 0.17240436 |
| SMART | SM00244:PHB | 3 | 0.00436765 | P67778, O35129, P54116 | 0.06980101 |
| Annotation Cluster 9 | Enrichment Score: 2.6800963334676604 | | | | |
| Category | Term | Count | P Value | Genes | FDR |
| UP_SEQ_FEATURE | CROSSLNK:Glycyl lysine isopeptide (Lys-Gly) (interchain wit | alternate | 6.34146341 | 9.50470872701042E-06 | 0.00145802 |
| UP_SEQ_FEATURE | CROSSLNK:Glycyl lysine isopeptide (Lys-Gly) (interchain wit | 19 | 0.00153764 | P46061, Q9EQK5, P97310, Q9CZW5, Q8C4J7, Q99L45, P13864, Q01320, Q61033, Q61881, P35700, P42932, Q6DFW4, O55201, P52480, P17918, P24547, P20152, O35737 | 0.12984552 |
| UP_KW_PTM | KW-1017~Isopeptide bond | 32 | 0.00508855 | Q9EQK5, Q60766, Q922F4, Q9CZW5, Q99L45, P13864, Q61033, P63001, Q6DFW4, O35609, P52480, P20152, O88351, P46061, Q80X90, P97310, Q8C4J7, P68254, P97855, Q01320, Q61881, P35700, Q9Z0X1, P42932, P06151, O55201, P67871, Q9D906, P17918, P24547, O35737, P63017 | 0.01780992 |
| UP_KW_PTM | KW-0832~Ubl conjugation | 34 | 0.25599183 | Q9EQK5, Q60766, Q9CZW5, Q99L45, P13864, Q61033, P63001, P63085, P09671, Q6DFW4, O35609, P52480, P20152, O88351, P46061, Q80X90, P97310, Q8C4J7, P68254, P97855, Q01320, Q61881, P35700, Q9Z0X1, P42932, P06151, O55201, P67871, Q9D906, P17918, P24547, O35737, P63017, Q6ZPJ3 | 0.44798571 |
| Annotation Cluster 10 | Enrichment Score: 2.5520613867880466 | | | | |
| Category | Term | Count | P Value | Genes | FDR |
| UP_KW_DOMAIN | KW-0676~Redox-active center | 7 | 1.85173E-06 | Q9CQM5, Q9QUH0, P99029, P09103, Q9ESY9, Q8R180, P35700 | 1.6666E-05 |
| UP_SEQ_FEATURE | DISULFID:Redox-active | 6 | 5.39341E-05 | Q9CQM5, Q9QUH0, P99029, P09103, Q9ESY9, Q8R180 | 0.00683165 |
| INTERPRO | IPR036249:Thioredoxin-like_sf | 6 | 0.00465013 | Q8VCH8, Q9CQM5, Q9QUH0, P99029, P09103, P35700 | 0.17249772 |
| UP_SEQ_FEATURE | DOMAIN:Thioredoxin | 4 | 0.00561469 | Q9CQM5, P99029, P09103, P35700 | 0.30479743 |
| GOTERM_MF_DIRECT | GO:0004601~peroxidase activity | 3 | 0.02301914 | Q9CQM5, P99029, P35700 | 0.25120884 |
| INTERPRO | IPR013766:Thioredoxin_domain | 3 | 0.04445694 | P99029, P09103, P35700 | 0.83298369 |
| UP_SEQ_FEATURE | ACT_SITE:Nucleophile | 4 | 0.51204695 | P56399, Q9CQM5, P09103, Q8R180 | 0.9921671 |
| Annotation Cluster 11 | Enrichment Score: 2.4231958091413737 | | | | |
| Category | Term | Count | P Value | Genes | FDR |
| GOTERM_MF_DIRECT | GO:0140662~ATP-dependent protein folding chaperone | 4 | 0.00076945 | P80315, P42932, P38647, P63017 | 0.01839343 |

| | | | | | |
|---|---|---|---|---|---|
| UP_KW_MOLECULAR_FUNCTION | KW-0143~Chaperone | 9 | 0.00101864 | P80315, Q91YE6, P42932, P62075, Q99L47, P38647, P09103, Q9DAU1, P63017 | 0.02495678 |
| GOTERM_BP_DIRECT | GO:0006457~protein folding | 6 | 0.00102933 | P80315, P42932, P38647, P09103, P63017, Q8R180 | 0.07451624 |
| GOTERM_BP_DIRECT | GO:0051085~chaperone cofactor-dependent protein refolding | 4 | 0.0010792 | Q99L47, P38647, P63017, Q8R180 | 0.07451624 |
| GOTERM_MF_DIRECT | GO:0051082~unfolded protein binding | 5 | 0.00233698 | P80315, P42932, Q99L47, P38647, P63017 | 0.04512168 |
| GOTERM_MF_DIRECT | GO:0044183~protein folding chaperone | 4 | 0.00468639 | P80315, P42932, P38647, P63017 | 0.07128992 |
| GOTERM_CC_DIRECT | GO:0005874~microtubule | 7 | 0.01604204 | P80315, P42932, Q922F4, Q8BRT1, Q9JKF1, P63017, Q62433 | 0.14307761 |
| GOTERM_BP_DIRECT | GO:0061077~chaperone-mediated protein folding | 3 | 0.02147211 | P80315, P42932, P63017 | 0.47350907 |
| GOTERM_MF_DIRECT | GO:0051087~protein-folding chaperone binding | 4 | 0.04728846 | P08228, Q99L47, P38647, P63017 | 0.35967893 |

Annotation Cluster 12 — Enrichment Score: 2.397420488885045

| Category | Term | Count | P Value | Genes | FDR |
|---|---|---|---|---|---|
| KEGG_PATHWAY | mmu00520:Amino sugar and nucleotide sugar metabolism | 7 | 6.84953E-05 | Q9DCN2, Q3TRM8, P20060, Q9R0N0, O08528, P29416, Q9D0F9 | 0.00815094 |
| GOTERM_BP_DIRECT | GO:0006006~glucose metabolic process | 5 | 0.00128163 | Q3TRM8, Q00612, P52480, O08528, Q9D0F9 | 0.08261166 |
| KEGG_PATHWAY | mmu00010:Glycolysis / Gluconeogenesis | 6 | 0.002113 | Q3TRM8, P06151, P52480, O08528, Q9D0F9, Q8BH04 | 0.09545399 |
| GOTERM_BP_DIRECT | GO:0046835~carbohydrate phosphorylation | 3 | 0.00818989 | Q3TRM8, Q9R0N0, O08528 | 0.31674388 |
| KEGG_PATHWAY | mmu00052:Galactose metabolism | 4 | 0.00912135 | Q3TRM8, Q9R0N0, O08528, Q9D0F9 | 0.21708819 |
| KEGG_PATHWAY | mmu01250:Biosynthesis of nucleotide sugars | 4 | 0.01462619 | Q3TRM8, Q9R0N0, O08528, Q9D0F9 | 0.31645759 |
| KEGG_PATHWAY | mmu00500:Starch and sucrose metabolism | 3 | 0.08152011 | Q3TRM8, O08528, Q9D0F9 | 0.7185847 |

Annotation Cluster 13 — Enrichment Score: 2.328384335813567

| Category | Term | Count | P Value | Genes | FDR |
|---|---|---|---|---|---|
| GOTERM_CC_DIRECT | GO:0005789~endoplasmic reticulum membrane | 17 | 0.00079931 | P61804, Q60766, Q9QY81, Q80W54, P18572, Q64310, Q91VE0, Q8R180, Q8VDP6, Q9DCN2, Q8VCH8, P36371, P35282, Q8C0N2, Q8R0X7, Q9CYN2, Q8BG07 | 0.01648583 |
| GOTERM_CC_DIRECT | GO:0005783~endoplasmic reticulum | 17 | 0.00413757 | P54116, Q9JIA7, Q60766, Q3UBX0, Q64310, Q9CR60, Q91VE0, P57776, Q8R180, Q9DCN2, Q8VCH8, P36371, Q8C0N2, Q8R0X7, P09103, Q9DAU1, Q8BG07 | 0.05057029 |
| UP_KW_CELLULAR_COMPONENT | KW-0256~Endoplasmic reticulum | 20 | 0.03128851 | P61804, Q9JIA7, Q9QY81, Q80W54, P18572, Q3UBX0, Q64310, Q91VE0, Q8R180, Q8VDP6, Q9DCN2, Q8VCH8, P36371, P35282, Q8C0N2, Q8R0X7, Q9CYN2, P09103, Q9DAU1, Q8BG07 | 0.10168764 |

Annotation Cluster 14 — Enrichment Score: 1.956138756603147

| Category | Term | Count | P Value | Genes | FDR |
|---|---|---|---|---|---|
| GOTERM_CC_DIRECT | GO:0005764~lysosome | 10 | 0.000167 | P35283, P08228, Q60766, P20060, Q91V41, Q9ESY9, O70404, P29416, P63017, P06797 | 0.00423911 |
| GOTERM_CC_DIRECT | GO:0005765~lysosomal membrane | 7 | 0.01073181 | P35283, O35643, Q9JIA7, Q60766, Q8BPX9, O70404, P63017 | 0.10731813 |
| UP_KW_CELLULAR_COMPONENT | KW-0458~Lysosome | 10 | 0.013138 | P35283, Q9JIA7, Q60766, P20060, Q8BPX9, Q9ESY9, O70404, P29416, P63017, P06797 | 0.04879828 |
| GOTERM_CC_DIRECT | GO:0005776~autophagosome | 3 | 0.07976728 | P35283, Q60766, P63017 | 0.39288362 |
| UP_KW_BIOLOGICAL_PROCESS | KW-0072~Autophagy | 5 | 0.08822155 | P35283, Q60766, Q9D906, O70404, P63017 | 0.53611555 |

Annotation Cluster 15 — Enrichment Score: 1.9482435961815707

| Category | Term | Count | P Value | Genes | FDR |
|---|---|---|---|---|---|
| GOTERM_BP_DIRECT | GO:0019430~removal of superoxide radicals | 3 | 0.00450654 | P08228, P09671, P35700 | 0.21126092 |
| GOTERM_BP_DIRECT | GO:0000302~response to reactive oxygen species | 3 | 0.01039869 | P08228, P09671, P35700 | 0.37056932 |
| KEGG_PATHWAY | mmu04146:Peroxisome | 5 | 0.0305104 | O35459, P08228, P09671, P99029, P35700 | 0.39884165 |

Annotation Cluster 16 — Enrichment Score: 1.942813737464382

| Category | Term | Count | P Value | Genes | FDR |
|---|---|---|---|---|---|
| UP_SEQ_FEATURE | MOTIF:Effector region | 6 | 0.00079314 | P35283, P84096, P63001, P35282, P35279, Q91V41 | 0.08611185 |
| UP_SEQ_FEATURE | LIPID:S-geranylgeranyl cysteine | 6 | 0.00137561 | P35283, P84096, P63001, P35282, P35279, Q91V41 | 0.12984552 |
| GOTERM_CC_DIRECT | GO:0005802~trans-Golgi network | 7 | 0.00218216 | P63001, P35282, O35643, Q4LDD4, Q8BRT1, P35279, Q91V41 | 0.0300047 |
| GOTERM_MF_DIRECT | GO:0005525~GTP binding | 9 | 0.00258106 | P35283, P84096, P63001, P35282, Q60766, Q922F4, P35279, Q91V41, Q8BH04 | 0.04627464 |
| GOTERM_CC_DIRECT | GO:0000139~Golgi membrane | 11 | 0.00274296 | P35283, Q61543, P63001, P35282, Q60766, Q7TMY8, O35609, P35279, Q91V41, Q64310, Q9CR60 | 0.03520044 |
| GOTERM_MF_DIRECT | GO:0003924~GTPase activity | 8 | 0.00313536 | P35283, P84096, P63001, P35282, Q60766, Q922F4, P35279, Q91V41 | 0.05246502 |
| SMART | SM00174:RHO | 6 | 0.00399674 | P35283, P84096, P63001, P35282, P35279, Q91V41 | 0.06980101 |
| INTERPRO | IPR001806:Small_GTPase | 6 | 0.00465013 | P35283, P84096, P63001, P35282, P35279, Q91V41 | 0.17249772 |
| SMART | SM00176:RAN | 5 | 0.00518251 | P35283, P84096, P35282, P35279, Q91V41 | 0.06980101 |
| SMART | SM00173:RAS | 6 | 0.00519795 | P35283, P84096, P63001, P35282, P35279, Q91V41 | 0.06980101 |
| SMART | SM00175:RAB | 6 | 0.00879869 | P35283, P84096, P63001, P35282, P35279, Q91V41 | 0.1033846 |
| INTERPRO | IPR005225:Small_GTP-bd_dom | 6 | 0.01115787 | P35283, P84096, P63001, P35282, P35279, Q91V41 | 0.34767933 |
| UP_KW_PTM | KW-0636~Prenylation | 6 | 0.0487131 | P35283, P84096, P63001, P35282, P35279, Q91V41 | 0.0974262 |
| GOTERM_CC_DIRECT | GO:0012505~endomembrane system | 4 | 0.07797903 | P35283, P35282, P35279, Q91V41 | 0.39288362 |
| UP_KW_LIGAND | KW-0342~GTP-binding | 9 | 0.0811441 | P35283, P84096, P63001, P35282, Q60766, Q922F4, P35279, Q91V41, Q8BH04 | 0.19282265 |
| GOTERM_MF_DIRECT | GO:0019003~GDP binding | 3 | 0.09287939 | P35283, P35282, Q91V41 | 0.55506492 |
| GOTERM_CC_DIRECT | GO:0005768~endosome | 4 | 0.34186491 | P35283, P35282, P18572, Q91V41 | 0.92178771 |
| UP_KW_PTM | KW-0449~Lipoprotein | 13 | 0.42084609 | P54116, P08228, Q8R1F1, Q60766, P99029, P35279, P35283, Q9JL26, P84096, P63001, Q9DCN2, P35282, Q91V41 | 0.65464947 |

Annotation Cluster 17 — Enrichment Score: 1.9197149173844907

| Category | Term | Count | P Value | Genes | FDR |
|---|---|---|---|---|---|
| INTERPRO | IPR015422:PyrdxlP-dep_Trfase_small | 4 | 0.0029594 | O88986, Q8R0X7, Q9JLI6, Q8VCN5 | 0.13561006 |
| INTERPRO | IPR015421:PyrdxlP-dep_Trfase_major | 4 | 0.00459445 | O88986, Q8R0X7, Q9JLI6, Q8VCN5 | 0.17249772 |
| INTERPRO | IPR015424:PyrdxlP-dep_Trfase | 4 | 0.00491032 | O88986, Q8R0X7, Q9JLI6, Q8VCN5 | 0.17386996 |
| UP_KW_MOLECULAR_FUNCTION | KW-0456~Lyase | 6 | 0.02673663 | P63001, Q8R0X7, Q9JLI6, Q8BH04, Q8VCN5, Q9DCL9 | 0.16376185 |
| UP_KW_LIGAND | KW-0663~Pyridoxal phosphate | 4 | 0.04050057 | O88986, Q8R0X7, Q9JLI6, Q8VCN5 | 0.1485021 |
| GOTERM_MF_DIRECT | GO:0030170~pyridoxal phosphate binding | 3 | 0.04193688 | O88986, Q8R0X7, Q8VCN5 | 0.33955342 |

Annotation Cluster 18 — Enrichment Score: 1.848831270541089

| Category | Term | Count | P Value | Genes | FDR |
|---|---|---|---|---|---|
| GOTERM_BP_DIRECT | GO:0006006~glucose metabolic process | 5 | 0.00128163 | Q3TRM8, Q00612, P52480, O08528, Q9D0F9 | 0.08261166 |
| KEGG_PATHWAY | mmu00010:Glycolysis / Gluconeogenesis | 6 | 0.002113 | Q3TRM8, P06151, P52480, O08528, Q9D0F9, Q8BH04 | 0.09545399 |
| GOTERM_BP_DIRECT | GO:0051156~glucose 6-phosphate metabolic process | 3 | 0.00223011 | Q3TRM8, Q00612, O08528 | 0.13269129 |
| KEGG_PATHWAY | mmu05230:Central carbon metabolism in cancer | 6 | 0.0024064 | P63085, Q3TRM8, Q00612, P06151, P52480, O08528 | 0.09545399 |
| GOTERM_MF_DIRECT | GO:0005536~D-glucose binding | 3 | 0.00311833 | Q3TRM8, Q00612, O08528 | 0.05246502 |
| KEGG_PATHWAY | mmu04930:Type II diabetes mellitus | 5 | 0.00399108 | P63085, Q3TRM8, P52480, O08528, O88351 | 0.10581893 |
| UP_KW_MOLECULAR_FUNCTION | KW-0021~Allosteric enzyme | 4 | 0.01554823 | Q3TRM8, P52480, O08528, P13864 | 0.12697725 |
| KEGG_PATHWAY | mmu01200:Carbon metabolism | 6 | 0.02517847 | Q3TRM8, Q00612, Q9R0P3, P52480, O08528 | 0.39884165 |
| GOTERM_BP_DIRECT | GO:0006096~glycolytic process | 3 | 0.03572715 | Q3TRM8, P52480, O08528 | 0.5214141 |
| UP_KW_BIOLOGICAL_PROCESS | KW-0324~Glycolysis | 3 | 0.04103304 | Q3TRM8, P52480, O08528 | 0.40520123 |
| KEGG_PATHWAY | mmu04910:Insulin signaling pathway | 5 | 0.12299543 | P63085, Q3TRM8, O08528, Q8BH04, O88351 | 0.76917451 |
| KEGG_PATHWAY | mmu04066:HIF-1 signaling pathway | 4 | 0.20904177 | P63085, Q3TRM8, P06151, O08528 | 0.90458073 |
| UP_KW_MOLECULAR_FUNCTION | KW-0418~Kinase | 9 | 0.34582711 | Q9JIA7, P63085, Q3TRM8, Q3U5Q7, P52480, Q9R0N0, O08528, Q9Z110, O88351 | 0.84727641 |

Annotation Cluster 19 — Enrichment Score: 1.742836636110808

| Category | Term | Count | P Value | Genes | FDR |
|---|---|---|---|---|---|
| GOTERM_BP_DIRECT | GO:0045087~innate immune response | 11 | 0.00110787 | P16110, Q60766, P20491, Q2EMV9, Q9D906, P97855, Q8BPX9, Q9DAU1, Q8VC04, Q8BG07, P0DOV2 | 0.07451624 |
| UP_KW_BIOLOGICAL_PROCESS | KW-0399~Innate immunity | 10 | 0.01766109 | P16110, Q60766, P20491, Q2EMV9, P97855, Q8BPX9, Q9DAU1, Q8VC04, Q8BG07, P0DOV2 | 0.23253769 |

| | | | | | |
|---|---|---|---|---|---|
| UP_KW_BIOLOGICAL_PROCESS | KW-0391~Immunity | 12 | 0.30198415 | P36371, P16110, Q60766, P20491, Q2EMV9, P97855, Q8BPX9, Q9ESY9, Q9DAU1, Q8VC04, Q8BG07, P0DOV2 | 0.97530864 |

Annotation Cluster 20 Enrichment Score: 1.6755968907522143

| Category | Term | Count | P Value | Genes | FDR |
|---|---|---|---|---|---|
| GOTERM_BP_DIRECT | GO:0016601~Rac protein signal transduction | 4 | 0.00054387 | P84096, P63001, Q8BH43, Q7TMB8 | 0.04949185 |
| GOTERM_BP_DIRECT | GO:0010592~positive regulation of lamellipodium assembly | 3 | 0.01370396 | P63001, Q8BH43, Q7TMB8 | 0.41568673 |
| GOTERM_BP_DIRECT | GO:0030032~lamellipodium assembly | 3 | 0.02147211 | P63001, Q8BH43, Q7TMB8 | 0.47350907 |
| GOTERM_CC_DIRECT | GO:0030027~lamellipodium | 4 | 0.12579219 | P63001, Q8BH43, Q7TMB8, P09103 | 0.5188928 |
| KEGG_PATHWAY | mmu04810:Regulation of actin cytoskeleton | 6 | 0.20804094 | P63001, Q8BH43, Q7TMB8, P26041, P63085, Q9JKF1 | 0.90458073 |

Annotation Cluster 21 Enrichment Score: 1.6191795599243954

| Category | Term | Count | P Value | Genes | FDR |
|---|---|---|---|---|---|
| GOTERM_BP_DIRECT | GO:0016601~Rac protein signal transduction | 4 | 0.00054387 | P84096, P63001, Q8BH43, Q7TMB8 | 0.04949185 |
| KEGG_PATHWAY | mmu05135:Yersinia infection | 6 | 0.03770204 | P84096, P63001, Q8BH43, P63085, Q3UND0, O88351 | 0.39884165 |
| GOTERM_BP_DIRECT | GO:0030036~actin cytoskeleton organization | 5 | 0.05782736 | Q9JL26, P84096, P63001, Q8BH43, Q80X90 | 0.57406686 |
| KEGG_PATHWAY | mmu05100:Bacterial invasion of epithelial cells | 3 | 0.2813787 | P84096, P63001, Q8BH43 | 0.97942387 |

Annotation Cluster 22 Enrichment Score: 1.6084534864208497

| Category | Term | Count | P Value | Genes | FDR |
|---|---|---|---|---|---|
| KEGG_PATHWAY | mmu00010:Glycolysis / Gluconeogenesis | 6 | 0.002113 | Q3TRM8, P06151, P52480, O08528, Q9D0F9, Q8BH04 | 0.09545399 |
| GOTERM_BP_DIRECT | GO:0006090~pyruvate metabolic process | 3 | 0.00350656 | P06151, P52480, Q8BH04 | 0.16952047 |
| KEGG_PATHWAY | mmu00620:Pyruvate metabolism | 3 | 0.11995673 | P06151, P52480, Q8BH04 | 0.76917451 |
| KEGG_PATHWAY | mmu04922:Glucagon signaling pathway | 3 | 0.41436127 | P06151, P52480, Q8BH04 | 0.97942387 |

Annotation Cluster 23 Enrichment Score: 1.5834216326475596

| Category | Term | Count | P Value | Genes | FDR |
|---|---|---|---|---|---|
| GOTERM_BP_DIRECT | GO:0032263~GMP salvage | 3 | 0.00153247 | P23492, P08030, P24547 | 0.09482897 |
| KEGG_PATHWAY | mmu01232:Nucleotide metabolism | 4 | 0.10602242 | P23492, Q3U5Q7, P08030, P24547 | 0.76350675 |
| KEGG_PATHWAY | mmu00230:Purine metabolism | 5 | 0.1093822 | P23492, P08030, P24547, Q9D0F9, Q9DCL9 | 0.76350675 |

Annotation Cluster 24 Enrichment Score: 1.5354728951368803

| Category | Term | Count | P Value | Genes | FDR |
|---|---|---|---|---|---|
| GOTERM_CC_DIRECT | GO:0005802~trans-Golgi network | 7 | 0.00218216 | P63001, P35282, O35643, Q4LDD4, Q8BRT1, P35279, Q91V41 | 0.0300047 |
| GOTERM_CC_DIRECT | GO:0005794~Golgi apparatus | 12 | 0.07813769 | Q61543, O35643, Q4LDD4, Q60766, P63085, Q8BRT1, Q9JLI6, P35279, Q91V41, Q64310, Q8VDP6, P06797 | 0.39288362 |
| UP_KW_CELLULAR_COMPONENT | KW-0333~Golgi apparatus | 13 | 0.14515572 | Q61543, Q60766, P35279, Q64310, Q9CR60, Q8R180, P35283, P35282, O35643, Q4LDD4, Q8BRT1, Q91V41, Q8BG07 | 0.37740486 |

Annotation Cluster 25 Enrichment Score: 1.435153092685107

| Category | Term | Count | P Value | Genes | FDR |
|---|---|---|---|---|---|
| UP_KW_LIGAND | KW-0285~Flavoprotein | 6 | 0.02316973 | Q9DCN2, Q9Z0X1, Q64521, Q8R180, Q8JZN5, Q91YT0 | 0.12645123 |
| GOTERM_MF_DIRECT | GO:0071949~FAD binding | 3 | 0.03004398 | Q9DCN2, Q9Z0X1, Q8R180 | 0.28359379 |
| UP_KW_LIGAND | KW-0274~FAD | 5 | 0.07109882 | Q9DCN2, Q9Z0X1, Q64521, Q8R180, Q8JZN5 | 0.19282265 |

Annotation Cluster 26 Enrichment Score: 1.4259625613912825

| Category | Term | Count | P Value | Genes | FDR |
|---|---|---|---|---|---|
| GOTERM_MF_DIRECT | GO:0051015~actin filament binding | 6 | 0.01246518 | Q9JL26, Q80X90, Q7TMB8, Q3UZA1, Q8BRT1, Q9JKF1 | 0.14898858 |
| GOTERM_CC_DIRECT | GO:0005925~focal adhesion | 5 | 0.03536516 | Q80X90, P26041, P63085, Q8BRT1, Q9JKF1 | 0.271678 |
| GOTERM_CC_DIRECT | GO:0005938~cell cortex | 4 | 0.11962945 | P63001, Q80X90, Q8BRT1, Q9JKF1 | 0.5061246 |

Annotation Cluster 27 Enrichment Score: 1.1549475455229936

| Category | Term | Count | P Value | Genes | FDR |
|---|---|---|---|---|---|
| UP_SEQ_FEATURE | REPEAT:3 | 5 | 0.02238528 | P62320, P16110, P13864, Q62433, P0DOV2 | 0.71382497 |
| UP_SEQ_FEATURE | REPEAT:1 | 5 | 0.0426599 | P62320, P16110, P13864, Q62433, P0DOV2 | 0.89899119 |
| UP_SEQ_FEATURE | REPEAT:2 | 5 | 0.0442898 | P62320, P16110, P13864, Q62433, P0DOV2 | 0.89899119 |
| UP_SEQ_FEATURE | REPEAT:5 | 3 | 0.17654404 | P62320, P16110, P13864 | 0.9921671 |
| UP_SEQ_FEATURE | REPEAT:4 | 3 | 0.22496924 | P62320, P16110, P13864 | 0.9921671 |

Annotation Cluster 28 Enrichment Score: 1.1171013651371595

| Category | Term | Count | P Value | Genes | FDR |
|---|---|---|---|---|---|
| KEGG_PATHWAY | mmu05208:Chemical carcinogenesis - reactive oxygen specie | 10 | 0.00396164 | P63001, Q91VD9, P08228, P63085, P09671, O70145, Q91WD5, P48758, O88351, Q91YT0 | 0.10581893 |
| KEGG_PATHWAY | mmu05020:Prion disease | 11 | 0.00400156 | P63001, P54775, Q91VD9, P08228, P63085, Q922F4, O70145, P67871, Q91WD5, P63017, Q91YT0 | 0.10581893 |
| GOTERM_MF_DIRECT | GO:0008137~NADH dehydrogenase (ubiquinone) activity | 3 | 0.00516084 | Q91VD9, Q91WD5, Q91YT0 | 0.07619824 |
| GOTERM_BP_DIRECT | GO:0006120~mitochondrial electron transport, NADH to ubiqu | 3 | 0.01118707 | Q91VD9, Q91WD5, Q91YT0 | 0.37056932 |
| UP_KW_BIOLOGICAL_PROCESS | KW-0249~Electron transport | 5 | 0.02498048 | Q91VD9, Q9QUH0, Q91WD5, Q8R180, Q91YT0 | 0.28192254 |
| UP_KW_LIGAND | KW-0830~Ubiquinone | 3 | 0.02873892 | Q91VD9, Q91WD5, Q91YT0 | 0.12645123 |
| GOTERM_MF_DIRECT | GO:0051539~4 iron, 4 sulfur cluster binding | 3 | 0.02882058 | Q91VD9, Q91WD5, Q91YT0 | 0.28359379 |
| KEGG_PATHWAY | mmu05014:Amyotrophic lateral sclerosis | 11 | 0.03147896 | P63001, P54775, Q91VD9, P08228, Q922F4, Q9QY81, P84104, Q91WD5, F6ZDS4, Q8R5C5, Q91YT0 | 0.39884165 |
| GOTERM_CC_DIRECT | GO:0045271~respiratory chain complex I | 3 | 0.04210353 | Q91VD9, Q91WD5, Q91YT0 | 0.27243459 |
| GOTERM_BP_DIRECT | GO:0042776~proton motive force-driven mitochondrial ATP sy | 3 | 0.05573194 | Q91VD9, Q91WD5, Q91YT0 | 0.57406686 |
| GOTERM_BP_DIRECT | GO:0009060~aerobic respiration | 3 | 0.07020701 | Q91VD9, Q91WD5, Q91YT0 | 0.64648953 |
| UP_KW_MOLECULAR_FUNCTION | KW-1278~Translocase | 4 | 0.07400186 | P36371, Q91VD9, Q91WD5, Q91YT0 | 0.32964465 |
| KEGG_PATHWAY | mmu05415:Diabetic cardiomyopathy | 7 | 0.0751902 | P63001, Q91VD9, Q00612, O70145, P58389, Q91WD5, Q91YT0 | 0.68827951 |
| UP_KW_LIGAND | KW-0004~4Fe-4S | 3 | 0.08764666 | Q91VD9, Q91WD5, Q91YT0 | 0.19282265 |
| KEGG_PATHWAY | mmu05016:Huntington disease | 8 | 0.1260412 | P54775, Q91VD9, P08228, P09671, Q922F4, Q91WD5, Q8R5C5, Q91YT0 | 0.76917451 |
| UP_KW_BIOLOGICAL_PROCESS | KW-0679~Respiratory chain | 3 | 0.13940411 | Q91VD9, Q91WD5, Q91YT0 | 0.73419496 |
| GOTERM_BP_DIRECT | GO:1902600~proton transmembrane transport | 3 | 0.17678533 | Q91VD9, Q91WD5, Q91YT0 | 0.98723676 |
| KEGG_PATHWAY | mmu04932:Non-alcoholic fatty liver disease | 5 | 0.17842293 | P63001, Q91VD9, Q91WD5, O88351, Q91YT0 | 0.90458073 |
| KEGG_PATHWAY | mmu05022:Pathways of neurodegeneration - multiple disease | 10 | 0.20687413 | P63001, P54775, Q91VD9, P08228, P63085, Q922F4, P67871, Q91WD5, Q8R5C5, Q91YT0 | 0.90458073 |
| UP_KW_LIGAND | KW-0411~Iron-sulfur | 3 | 0.22365199 | Q91VD9, Q91WD5, Q91YT0 | 0.44730397 |
| KEGG_PATHWAY | mmu05010:Alzheimer disease | 8 | 0.27752037 | P54775, Q91VD9, P63085, Q922F4, P67871, Q91WD5, O88351, Q91YT0 | 0.97942387 |
| KEGG_PATHWAY | mmu00190:Oxidative phosphorylation | 4 | 0.30016643 | Q91VD9, Q8BVE3, Q91WD5, Q91YT0 | 0.97942387 |
| KEGG_PATHWAY | mmu05012:Parkinson disease | 6 | 0.31598276 | P54775, Q91VD9, P08228, Q922F4, Q91WD5, Q91YT0 | 0.97942387 |
| KEGG_PATHWAY | mmu04723:Retrograde endocannabinoid signaling | 4 | 0.34490833 | Q91VD9, P63085, Q91WD5, Q91YT0 | 0.97942387 |
| KEGG_PATHWAY | mmu04714:Thermogenesis | 3 | 0.83854321 | Q91VD9, Q91WD5, Q91YT0 | 0.97942387 |
| UP_KW_LIGAND | KW-0408~Iron | 3 | 0.96510404 | Q91VD9, Q91WD5, Q91YT0 | 0.96510404 |

Annotation Cluster 29 Enrichment Score: 1.090889831590567

| Category | Term | Count | P Value | Genes | FDR |
|---|---|---|---|---|---|
| GOTERM_CC_DIRECT | GO:0005874~microtubule | 7 | 0.01604204 | P80315, P42932, Q922F4, Q8BRT1, Q9JKF1, P63017, Q62433 | 0.14307761 |
| GOTERM_CC_DIRECT | GO:0015630~microtubule cytoskeleton | 5 | 0.04561708 | Q922F4, Q8BRT1, Q9JKF1, Q62433, Q8R5C5 | 0.27812412 |
| UP_KW_CELLULAR_COMPONENT | KW-0493~Microtubule | 3 | 0.72936277 | Q922F4, Q8BRT1, Q62433 | 0.86666667 |

Annotation Cluster 30 Enrichment Score: 1.0648062888315104

| Category | Term | Count | P Value | Genes | FDR |
|---|---|---|---|---|---|
| KEGG_PATHWAY | mmu04148:Efferocytosis | 7 | 0.0235919 | P63001, Q9JIA7, P63085, P97797, Q61072, P18572, Q91V41 | 0.39884165 |
| KEGG_PATHWAY | mmu04071:Sphingolipid signaling pathway | 5 | 0.09237479 | P63001, Q9JIA7, P63085, P20491, Q8R0X7 | 0.75811036 |
| KEGG_PATHWAY | mmu04666:Fc gamma R-mediated phagocytosis | 4 | 0.13571264 | P63001, Q8BH43, Q9JIA7, P63085 | 0.80749021 |
| KEGG_PATHWAY | mmu04370:VEGF signaling pathway | 3 | 0.18613931 | P63001, Q9JIA7, P63085 | 0.90458073 |

| Annotation Cluster 31 | Enrichment Score: 1.0537412142556406 | | | | |
|---|---|---|---|---|---|
| Category | Term | Count | P Value | Genes | FDR |
| UP_SEQ_FEATURE | REPEAT:TPR | 4 | 0.03018167 | O70145, Q80UM3, Q9CZW5, Q99L47 | 0.71382497 |
| UP_SEQ_FEATURE | REPEAT:TPR 3 | 4 | 0.07682481 | O70145, Q80UM3, Q9CZW5, Q99L47 | 0.9921671 |
| SMART | SM00028:TPR | 4 | 0.0816779 | O70145, Q80UM3, Q9CZW5, Q99L47 | 0.6979748 |
| INTERPRO | IPR019734:TPR_repeat | 4 | 0.08346819 | O70145, Q80UM3, Q9CZW5, Q99L47 | 0.97987421 |
| UP_SEQ_FEATURE | REPEAT:TPR 1 | 4 | 0.09312839 | O70145, Q80UM3, Q9CZW5, Q99L47 | 0.9921671 |
| UP_SEQ_FEATURE | REPEAT:TPR 2 | 4 | 0.09312839 | O70145, Q80UM3, Q9CZW5, Q99L47 | 0.9921671 |
| UP_KW_DOMAIN | KW-0802~TPR repeat | 4 | 0.10517739 | O70145, Q80UM3, Q9CZW5, Q99L47 | 0.47329826 |
| INTERPRO | IPR011990:TPR-like_helical_dom_sf | 4 | 0.25769944 | O70145, Q80UM3, Q9CZW5, Q99L47 | 0.97987421 |

| Annotation Cluster 32 | Enrichment Score: 0.9651374584959795 | | | | |
|---|---|---|---|---|---|
| Category | Term | Count | P Value | Genes | FDR |
| KEGG_PATHWAY | mmu04520:Adherens junction | 5 | 0.03638982 | P63001, Q8BH43, P63085, P67871, Q9JKF1 | 0.39884165 |
| KEGG_PATHWAY | mmu05135:Yersinia infection | 6 | 0.03770204 | P84096, P63001, Q8BH43, P63085, Q3UND0, O88351 | 0.39884165 |
| KEGG_PATHWAY | mmu04666:Fc gamma R-mediated phagocytosis | 4 | 0.13571264 | P63001, Q8BH43, Q9JIA7, P63085 | 0.80749021 |
| KEGG_PATHWAY | mmu04810:Regulation of actin cytoskeleton | 6 | 0.20804094 | P63001, Q8BH43, Q7TMB8, P26041, P63085, Q9JKF1 | 0.90458073 |
| KEGG_PATHWAY | mmu05231:Choline metabolism in cancer | 3 | 0.3856549 | P63001, Q8BH43, P63085 | 0.97942387 |

| Annotation Cluster 33 | Enrichment Score: 0.9414526665392488 | | | | |
|---|---|---|---|---|---|
| Category | Term | Count | P Value | Genes | FDR |
| GOTERM_BP_DIRECT | GO:0016485~protein processing | 4 | 0.02250963 | Q61543, Q61072, Q8JZQ2, Q80W54 | 0.47350907 |
| GOTERM_MF_DIRECT | GO:0004222~metalloendopeptidase activity | 4 | 0.03749763 | Q61072, P34960, Q8JZQ2, Q80W54 | 0.31373021 |
| GOTERM_MF_DIRECT | GO:0005518~collagen binding | 3 | 0.07158019 | Q61072, P34960, P06797 | 0.47844261 |
| GOTERM_BP_DIRECT | GO:0006508~proteolysis | 7 | 0.14713574 | P56399, Q61072, P34960, Q8JZQ2, P61290, P06797, Q8CGK3 | 0.96710553 |
| UP_KW_MOLECULAR_FUNCTION | KW-0482~Metalloprotease | 4 | 0.18727062 | Q61072, P34960, Q8JZQ2, Q80W54 | 0.65397908 |
| UP_KW_MOLECULAR_FUNCTION | KW-0645~Protease | 8 | 0.28912856 | P56399, Q61072, P34960, Q8JZQ2, Q80W54, P61290, P06797, Q8CGK3 | 0.7963536 |
| UP_KW_PTM | KW-0865~Zymogen | 4 | 0.53381092 | Q61072, P34960, P29416, P06797 | 0.74733529 |

| Annotation Cluster 34 | Enrichment Score: 0.9264418966012067 | | | | |
|---|---|---|---|---|---|
| Category | Term | Count | P Value | Genes | FDR |
| GOTERM_CC_DIRECT | GO:0072686~mitotic spindle | 5 | 0.00836474 | P46061, P63085, Q922F4, Q8BRT1, F6ZDS4 | 0.08904398 |
| UP_KW_CELLULAR_COMPONENT | KW-0995~Kinetochore | 3 | 0.24195115 | P46061, Q8BRT1, F6ZDS4 | 0.5242275 |
| GOTERM_CC_DIRECT | GO:0000776~kinetochore | 3 | 0.25301665 | P46061, Q8BRT1, F6ZDS4 | 0.78769333 |
| UP_KW_CELLULAR_COMPONENT | KW-0137~Centromere | 3 | 0.38450616 | P46061, Q8BRT1, F6ZDS4 | 0.62427212 |

| Annotation Cluster 35 | Enrichment Score: 0.9024289997984987 | | | | |
|---|---|---|---|---|---|
| Category | Term | Count | P Value | Genes | FDR |
| GOTERM_BP_DIRECT | GO:0071526~semaphorin-plexin signaling pathway | 3 | 0.02705425 | B2RXS4, P63001, P70206 | 0.48861222 |
| GOTERM_BP_DIRECT | GO:0030334~regulation of cell migration | 3 | 0.16203823 | B2RXS4, P63001, P70206 | 0.98723676 |
| KEGG_PATHWAY | mmu04360:Axon guidance | 4 | 0.44756899 | B2RXS4, P63001, P63085, P70206 | 0.97942387 |

| Annotation Cluster 36 | Enrichment Score: 0.7988137899916837 | | | | |
|---|---|---|---|---|---|
| Category | Term | Count | P Value | Genes | FDR |
| UP_KW_CELLULAR_COMPONENT | KW-0747~Spliceosome | 5 | 0.05272419 | P62320, P16110, Q9CQQ8, O35737, P63017 | 0.15231434 |
| GOTERM_CC_DIRECT | GO:0005681~spliceosomal complex | 4 | 0.05688757 | P62320, P16110, O35737, P63017 | 0.33350086 |
| GOTERM_MF_DIRECT | GO:0003723~RNA binding | 10 | 0.05919502 | P46061, P62320, Q9CQQ8, P84104, P20152, P97855, P13864, P24547, O35737, P63017 | 0.41853378 |
| GOTERM_BP_DIRECT | GO:0000398~mRNA splicing, via spliceosome | 4 | 0.08797666 | P62320, Q9CQQ8, P84104, P63017 | 0.70154584 |
| GOTERM_BP_DIRECT | GO:0008380~RNA splicing | 4 | 0.09943729 | P16110, P84104, O35737, P63017 | 0.76153214 |
| GOTERM_CC_DIRECT | GO:0071013~catalytic step 2 spliceosome | 3 | 0.16492779 | P62320, Q9CQQ8, O35737 | 0.59808981 |
| GOTERM_BP_DIRECT | GO:0006397~mRNA processing | 4 | 0.16924618 | P16110, P84104, O35737, P63017 | 0.98723676 |
| UP_KW_BIOLOGICAL_PROCESS | KW-0508~mRNA splicing | 6 | 0.19250132 | P62320, P16110, Q9CQQ8, P84104, O35737, P63017 | 0.80040023 |
| UP_KW_BIOLOGICAL_PROCESS | KW-0507~mRNA processing | 6 | 0.33915908 | P62320, P16110, Q9CQQ8, P84104, O35737, P63017 | 0.97530864 |
| KEGG_PATHWAY | mmu03040:Spliceosome | 4 | 0.70314524 | P62320, Q9CQQ8, P84104, P63017 | 0.97942387 |
| UP_KW_MOLECULAR_FUNCTION | KW-0694~RNA-binding | 6 | 0.82058605 | P62320, Q9CQQ8, P84104, P97855, P24547, O35737 | 0.9245283 |

| Annotation Cluster 37 | Enrichment Score: 0.6695714920585408 | | | | |
|---|---|---|---|---|---|
| Category | Term | Count | P Value | Genes | FDR |
| KEGG_PATHWAY | mmu04071:Sphingolipid signaling pathway | 5 | 0.09237479 | P63001, Q9JIA7, P63085, P20491, Q8R0X7 | 0.75811036 |
| KEGG_PATHWAY | mmu04664:Fc epsilon RI signaling pathway | 3 | 0.22590508 | P63001, P63085, P20491 | 0.92345126 |
| KEGG_PATHWAY | mmu04650:Natural killer cell mediated cytotoxicity | 3 | 0.46968432 | P63001, P63085, P20491 | 0.97942387 |

| Annotation Cluster 38 | Enrichment Score: 0.6311150555578677 | | | | |
|---|---|---|---|---|---|
| Category | Term | Count | P Value | Genes | FDR |
| KEGG_PATHWAY | mmu04071:Sphingolipid signaling pathway | 5 | 0.09237479 | P63001, Q9JIA7, P63085, P20491, Q8R0X7 | 0.75811036 |
| KEGG_PATHWAY | mmu05152:Tuberculosis | 5 | 0.22892279 | Q9JIA7, P63085, P20491, Q8BVE3, P38647 | 0.92345126 |
| KEGG_PATHWAY | mmu04072:Phospholipase D signaling pathway | 3 | 0.6045223 | Q9JIA7, P63085, P20491 | 0.97942387 |

| Annotation Cluster 39 | Enrichment Score: 0.5910587232304466 | | | | |
|---|---|---|---|---|---|
| Category | Term | Count | P Value | Genes | FDR |
| GOTERM_BP_DIRECT | GO:0032496~response to lipopolysaccharide | 6 | 0.00631618 | P63001, P63085, P09671, O70145, P19253, Q8BH04 | 0.26408446 |
| KEGG_PATHWAY | mmu05135:Yersinia infection | 6 | 0.03770204 | P84096, P63001, Q8BH43, P63085, Q3UND0, O88351 | 0.39884165 |
| GOTERM_BP_DIRECT | GO:0071356~cellular response to tumor necrosis factor | 4 | 0.03875163 | P63085, P16460, Q8BH04, O88351 | 0.53977854 |
| KEGG_PATHWAY | mmu05417:Lipid and atherosclerosis | 7 | 0.0751902 | P63001, P63085, P09671, O70145, P63017, Q8R180, O88351 | 0.68827951 |
| KEGG_PATHWAY | mmu04380:Osteoclast differentiation | 5 | 0.11384085 | P63001, P63085, O70145, P97797, O88351 | 0.76350675 |
| KEGG_PATHWAY | mmu04910:Insulin signaling pathway | 5 | 0.12299543 | P63085, Q3TRM8, O08528, Q8BH04, O88351 | 0.76917451 |
| KEGG_PATHWAY | mmu05169:Epstein-Barr virus infection | 6 | 0.19845294 | P63001, P36371, P54775, P20152, P31266, O88351 | 0.90458073 |
| KEGG_PATHWAY | mmu04068:FoxO signaling pathway | 4 | 0.2705154 | P63085, P09671, Q8BH04, O88351 | 0.97942387 |
| KEGG_PATHWAY | mmu05212:Pancreatic cancer | 3 | 0.27632972 | P63001, P63085, O88351 | 0.97942387 |
| KEGG_PATHWAY | mmu04662:B cell receptor signaling pathway | 3 | 0.31660331 | P63001, P63085, O88351 | 0.97942387 |
| KEGG_PATHWAY | mmu04620:Toll-like receptor signaling pathway | 3 | 0.41436127 | P63001, P63085, O88351 | 0.97942387 |
| KEGG_PATHWAY | mmu05170:Human immunodeficiency virus 1 infection | 5 | 0.41513043 | P63001, P36371, O35643, P63085, O88351 | 0.97942387 |
| KEGG_PATHWAY | mmu04722:Neurotrophin signaling pathway | 3 | 0.49181445 | P63001, P63085, O88351 | 0.97942387 |
| KEGG_PATHWAY | mmu04613:Neutrophil extracellular trap formation | 4 | 0.54340634 | P63001, P63085, O70145, Q9D906 | 0.97942387 |
| KEGG_PATHWAY | mmu05163:Human cytomegalovirus infection | 4 | 0.67545955 | P63001, P36371, P63085, O88351 | 0.97942387 |
| KEGG_PATHWAY | mmu04151:PI3K-Akt signaling pathway | 5 | 0.74029928 | P63001, P63085, P68254, Q8BH04, O88351 | 0.97942387 |
| KEGG_PATHWAY | mmu04062:Chemokine signaling pathway | 3 | 0.74204743 | P63001, P63085, O88351 | 0.97942387 |
| KEGG_PATHWAY | mmu04010:MAPK signaling pathway | 4 | 0.77741372 | P63001, P63085, P63017, O88351 | 0.97942387 |
| KEGG_PATHWAY | mmu05167:Kaposi sarcoma-associated herpesvirus infection | 3 | 0.81068578 | P63001, P63085, O88351 | 0.97942387 |
| KEGG_PATHWAY | mmu04014:Ras signaling pathway | 3 | 0.8350539 | P63001, P63085, O88351 | 0.97942387 |
| KEGG_PATHWAY | mmu05200:Pathways in cancer | 5 | 0.9448643 | P63001, P63085, F6ZDS4, P43247, O88351 | 0.97942387 |

| Annotation Cluster 40 | Enrichment Score: 0.4295005859083525 | | | | |
|---|---|---|---|---|---|
| Category | Term | Count | P Value | Genes | FDR |
| GOTERM_BP_DIRECT | GO:0051321~meiotic cell cycle | 4 | 0.12090182 | Q8CG48, Q8CG47, Q8BRT1, Q8BJS4 | 0.84250052 |
| GOTERM_BP_DIRECT | GO:0051301~cell division | 4 | 0.37743379 | Q8CG48, Q8CG47, Q8BRT1, F6ZDS4 | 0.98723676 |
| UP_KW_BIOLOGICAL_PROCESS | KW-0498~Mitosis | 4 | 0.5488576 | Q8CG48, Q8CG47, Q8BRT1, F6ZDS4 | 0.97530864 |
| UP_KW_BIOLOGICAL_PROCESS | KW-0132~Cell division | 4 | 0.76430087 | Q8CG48, Q8CG47, Q8BRT1, F6ZDS4 | 0.97530864 |

| Annotation Cluster 41 | Enrichment Score: 0.393356561641299 | | | | |
|---|---|---|---|---|---|
| Category | Term | Count | P Value | Genes | FDR |
| KEGG_PATHWAY | mmu05010:Alzheimer disease | 8 | 0.27752037 | P54775, Q91VD9, P63085, Q922F4, P67871, Q91WD5, O88351, Q91YT0 | 0.97942387 |
| KEGG_PATHWAY | mmu05235:PD-L1 expression and PD-1 checkpoint pathway ir | 3 | 0.33657086 | P63085, P67871, O88351 | 0.97942387 |
| GOTERM_MF_DIRECT | GO:0004674~protein serine/threonine kinase activity | 3 | 0.70722718 | P63085, P67871, O88351 | 0.94538606 |

| Annotation Cluster 42 | Enrichment Score: 0.31524459834625407 | | | | |
|---|---|---|---|---|---|
| Category | Term | Count | P Value | Genes | FDR |
| KEGG_PATHWAY | mmu05161:Hepatitis B | 4 | 0.38574173 | P63085, P68254, P17918, O88351 | 0.97942387 |
| KEGG_PATHWAY | mmu05160:Hepatitis C | 4 | 0.3967914 | P63085, P68254, P61290, O88351 | 0.97942387 |
| KEGG_PATHWAY | mmu04151:PI3K-Akt signaling pathway | 5 | 0.74029928 | P63001, P63085, P68254, Q8BH04, O88351 | 0.97942387 |

| Annotation Cluster 43 | Enrichment Score: 0.30263693506379635 | | | | |
|---|---|---|---|---|---|
| Category | Term | Count | P Value | Genes | FDR |
| UP_KW_DOMAIN | KW-0728~SH3 domain | 3 | 0.45919245 | O70145, Q3UND0, Q62422 | 1 |
| UP_SEQ_FEATURE | DOMAIN:SH3 | 3 | 0.49400745 | O70145, Q3UND0, Q62422 | 0.9921671 |
| SMART | SM00326:SH3 | 3 | 0.50110078 | O70145, Q3UND0, Q62422 | 0.97916667 |
| INTERPRO | IPR036028:SH3-like_dom_sf | 3 | 0.50935068 | O70145, Q3UND0, Q62422 | 0.97987421 |
| INTERPRO | IPR001452:SH3_domain | 3 | 0.52984045 | O70145, Q3UND0, Q62422 | 0.97987421 |

| Annotation Cluster 44 | Enrichment Score: 0.24044976820692307 | | | | |
|---|---|---|---|---|---|
| Category | Term | Count | P Value | Genes | FDR |
| INTERPRO | IPR012677:Nucleotide-bd_a/b_plait_sf | 4 | 0.37927321 | Q2EMV9, P84104, P97855, O35737 | 0.97987421 |
| UP_SEQ_FEATURE | DOMAIN:RRM | 3 | 0.55434972 | P84104, P97855, O35737 | 0.9921671 |
| SMART | SM00360:RRM | 3 | 0.56994439 | P84104, P97855, O35737 | 0.97916667 |
| INTERPRO | IPR000504:RRM_dom | 3 | 0.57659962 | P84104, P97855, O35737 | 0.97987421 |
| INTERPRO | IPR035979:RBD_domain_sf | 3 | 0.63640623 | P84104, P97855, O35737 | 0.97987421 |
| UP_KW_MOLECULAR_FUNCTION | KW-0694~RNA-binding | 6 | 0.82058605 | P62320, Q9CQQ8, P84104, P97855, P24547, O35737 | 0.9245283 |

| Annotation Cluster 45 | Enrichment Score: 0.19722416092342535 | | | | |
|---|---|---|---|---|---|
| Category | Term | Count | P Value | Genes | FDR |
| UP_SEQ_FEATURE | DOMAIN:PH | 3 | 0.61941997 | Q8R1F1, Q4LDD4, Q3UND0 | 0.9921671 |
| INTERPRO | IPR001849:PH_domain | 3 | 0.62415516 | Q8R1F1, Q4LDD4, Q3UND0 | 0.97987421 |
| INTERPRO | IPR011993:PH-like_dom_sf | 4 | 0.66229237 | Q8R1F1, Q4LDD4, P26041, Q3UND0 | 0.97987421 |

| Annotation Cluster 46 | Enrichment Score: 0.18169698247799435 | | | | |
|---|---|---|---|---|---|
| Category | Term | Count | P Value | Genes | FDR |
| GOTERM_BP_DIRECT | GO:0030198~extracellular matrix organization | 3 | 0.29842806 | P16110, P34960, Q8R180 | 0.98723676 |
| GOTERM_CC_DIRECT | GO:0005615~extracellular space | 8 | 0.95531847 | P08228, P16110, P20060, Q61072, P34960, P63028, Q8R180, P06797 | 0.95531847 |
| UP_KW_CELLULAR_COMPONENT | KW-0964~Secreted | 6 | 0.99981863 | P63001, P16110, P34960, Q9ESY9, Q8R180, P06797 | 0.99981863 |

| Annotation Cluster 47 | Enrichment Score: 0.10498643458061026 | | | | |
|---|---|---|---|---|---|
| Category | Term | Count | P Value | Genes | FDR |
| KEGG_PATHWAY | mmu04668:TNF signaling pathway | 3 | 0.47860483 | P63085, Q9D2Y4, O88351 | 0.97942387 |
| UP_SEQ_FEATURE | DOMAIN:Protein kinase | 3 | 0.91774185 | P63085, Q9D2Y4, O88351 | 0.9921671 |
| INTERPRO | IPR000719:Prot_kinase_dom | 3 | 0.92056315 | P63085, Q9D2Y4, O88351 | 0.97987421 |
| INTERPRO | IPR011009:Kinase-like_dom_sf | 3 | 0.9403793 | P63085, Q9D2Y4, O88351 | 0.97987421 |

| Annotation Cluster 48 | Enrichment Score: 0.07938357082364626 | | | | |
|---|---|---|---|---|---|
| Category | Term | Count | P Value | Genes | FDR |
| GOTERM_CC_DIRECT | GO:0016020~membrane | 35 | 0.50712181 | Q60766, Q8BVE3, P20060, P19253, P35279, Q3UBX0, Q9CR60, Q61033, Q8VC04, P70206, P63001, P36371, Q00612, Q7TMY8, O35609, Q9JIA7, O70145, P97797, Q8JZQ2, Q9QY81, Q80W54, Q64310, Q8BMD8, Q2YFS3, Q8VDP6, B2RXS4, Q91VD9, Q9Z0X1, P06151, Q8R0X7, Q61072, Q9D906, Q8BJS4, P29416, O70404 | 0.92178771 |
| UP_KW_DOMAIN | KW-1133~Transmembrane helix | 37 | 0.87098992 | P54116, P61804, Q9CZW5, P19253, Q3UBX0, Q9CR60, Q61033, Q8VC04, P70206, P63001, P36371, Q8C0N2, Q7TMY8, O35609, Q9CYN2, Q8BPX9, Q61543, P20491, Q9QZD8, P97797, Q8JZQ2, Q9QY81, Q80W54, P18572, Q64310, Q8BMD8, Q91VE0, Q2YFS3, Q8VDP6, B2RXS4, Q9Z0X1, P06151, Q8R0X7, Q61072, Q8BJS4, O70404, Q8BG07 | 1 |
| UP_KW_CELLULAR_COMPONENT | KW-0472~Membrane | 78 | 0.8777999 | Q8BVE3, Q9CZW5, P19253, Q3UBX0, Q9CR60, Q9D2Y4, Q61033, F6ZDS4, Q8VC04, Q8JZN5, P63001, Q9DCN2, O35129, Q8VCH8, O35643, P63085, P62075, O35609, P20152, Q8BH43, P20491, P97797, Q9QY81, Q8JZQ2, P18572, Q64310, Q2YFS3, Q8R180, P84096, B2RXS4, P67778, Q8C147, P06151, P09103, O70404, P63017, P06797, P54116, P61804, P26041, Q60766, P35279, O08528, Q9JKF1, Q91WD5, Q64521, P70206, P35283, P35282, P36371, Q4LDD4, Q8C0N2, Q00612, Q7TMY8, Q8BRT1, Q9CYN2, Q8BPX9, O88351, Q91YT0, Q61543, Q8R1F1, Q9JIA7, Q9QZD8, Q80W54, Q8BMD8, Q91VE0, Q62433, Q8VDP6, Q9JL26, Q91VD9, Q9Z0X1, Q8R0X7, Q61072, Q91V41, Q9Z110, P43883, Q8BJS4, Q8BG07 | 0.8777999 |
| UP_SEQ_FEATURE | TOPO_DOM:Cytoplasmic | 22 | 0.89193213 | Q61543, P54116, P61804, P20491, Q9CZW5, P97797, Q9QY81, P18572, Q3UBX0, Q9CR60, Q2YFS3, Q8VDP6, P70206, B2RXS4, Q8VCH8, P36371, Q8R0X7, Q61072, O35609, Q9CYN2, O70404, Q8BG07 | 0.9921671 |
| UP_KW_DOMAIN | KW-0812~Transmembrane | 37 | 0.96587291 | P54116, P61804, Q9CZW5, P19253, Q3UBX0, Q9CR60, Q61033, Q8VC04, P70206, P63001, P36371, Q8C0N2, Q7TMY8, O35609, Q9CYN2, Q8BPX9, Q61543, P20491, Q9QZD8, P97797, Q8JZQ2, Q9QY81, Q80W54, P18572, Q64310, Q8BMD8, Q91VE0, Q2YFS3, Q8VDP6, B2RXS4, Q9Z0X1, P06151, Q8R0X7, Q61072, Q8BJS4, O70404, Q8BG07 | 1 |
| UP_SEQ_FEATURE | TRANSMEM:Helical | 33 | 0.99982923 | P54116, P61804, Q9CZW5, P19253, Q3UBX0, Q61033, Q8VC04, P70206, P63001, P36371, Q8C0N2, Q7TMY8, O35609, Q9CYN2, Q8BPX9, Q61543, P20491, P97797, Q8JZQ2, Q9QY81, Q80W54, P18572, Q64310, Q91VE0, Q2YFS3, Q8VDP6, B2RXS4, Q9Z0X1, P06151, Q8R0X7, Q61072, Q8BJS4, O70404 | 0.99982923 |

| Annotation Cluster 49 | Enrichment Score: 0.050885000920844414 | | | | |
|---|---|---|---|---|---|
| Category | Term | Count | P Value | Genes | FDR |
| UP_KW_MOLECULAR_FUNCTION | KW-0678~Repressor | 6 | 0.65118774 | O35129, O55201, P13864, P31266, P63017, P0DOV2 | 0.9245283 |
| UP_KW_MOLECULAR_FUNCTION | KW-0010~Activator | 4 | 0.96258264 | O55201, P13864, P31266, P0DOV2 | 0.96258264 |
| UP_KW_BIOLOGICAL_PROCESS | KW-0805~Transcription regulation | 10 | 0.99904725 | O35129, Q8BFQ4, Q2EMV9, Q80UM3, O55201, P13864, P57776, P31266, P63017, P0DOV2 | 0.99904725 |
| UP_KW_BIOLOGICAL_PROCESS | KW-0804~Transcription | 10 | 0.99937786 | O35129, Q8BFQ4, Q2EMV9, Q80UM3, O55201, P13864, P57776, P31266, P63017, P0DOV2 | 0.99937786 |

| Annotation Cluster 50 | Enrichment Score: 0.020009222607381295 | | | | |
|---|---|---|---|---|---|
| Category | Term | Count | P Value | Genes | FDR |
| UP_SEQ_FEATURE | TRANSMEM:Helical | Name=4 | 1.95121951 | 0.929347080035726 | 1 |
| UP_SEQ_FEATURE | TRANSMEM:Helical | Name=1 | 1.95121951 | 0.933702526630595 | 1 |
| UP_SEQ_FEATURE | TRANSMEM:Helical | Name=2 | 1.95121951 | 0.93559987561138 | 1 |
| UP_SEQ_FEATURE | TRANSMEM:Helical | Name=6 | 1.46341463 | 0.976147152684568 | 1 |
| UP_SEQ_FEATURE | TRANSMEM:Helical | Name=5 | 1.46341463 | 0.97711295627683 | 1 |
| UP_SEQ_FEATURE | TRANSMEM:Helical | Name=3 | 1.46341463 | 0.979507966200704 | 1 |

| Annotation Cluster 51 | Enrichment Score: 0.0072084614403603595 | | | | |
|---|---|---|---|---|---|
| Category | Term | Count | P Value | Genes | FDR |
| INTERPRO | IPR013783:Ig-like_fold | 7 | 0.95323131 | B2RXS4, Q80X90, P97797, P18572, Q2YFS3, P31266, P70206 | 0.97987421 |
| INTERPRO | IPR036179:Ig-like_dom_sf | 3 | 0.99831329 | P97797, P18572, Q2YFS3 | 0.99831329 |
| UP_SEQ_FEATURE | TOPO_DOM:Extracellular | 8 | 0.99979156 | Q61543, B2RXS4, P20491, P97797, Q61072, P18572, Q2YFS3, P70206 | 0.99979156 |

| Annotation Cluster 52 | Enrichment Score: 4.0495934159209303E-4 | | | | |
|---|---|---|---|---|---|
| Category | Term | Count | P Value | Genes | FDR |
| UP_KW_DOMAIN | KW-0732~Signal | 23 | 0.99776009 | Q61543, Q9D0S9, P20491, P20060, P97797, Q9QY81, P34960, P18572, Q3UBX0, Q9ESY9, Q2YFS3, Q8R180, P70206, B2RXS4, P63001, P36371, P16110, Q61072, P09103, Q9DAU1, P29416, P06797, Q8CGK3 | 1 |
| UP_SEQ_FEATURE | CARBOHYD:N-linked (GlcNAc...) asparagine | 17 | 0.99866027 | Q61543, P20060, P97797, Q9QY81, P34960, P18572, Q9ESY9, Q2YFS3, Q8R180, P70206, B2RXS4, Q61072, Q8BPX9, Q9DAU1, P29416, Q8BJS4, Q8BG07 | 0.99866027 |
| UP_KW_PTM | KW-1015~Disulfide bond | 24 | 0.99985326 | P08228, P20491, P56399, Q9CQM5, P20060, P99029, P97797, P34960, P18572, Q9ESY9, Q8R180, P35700, P70206, B2RXS4, P63001, P16110, Q9QUH0, P62075, Q61072, P09103, Q9DAU1, P29416, Q8BJS4, P06797 | 0.99985326 |
| UP_KW_PTM | KW-0325~Glycoprotein | 22 | 0.99999999 | Q61543, P97311, P20060, P97797, Q9QY81, P34960, P18572, Q9ESY9, Q2YFS3, Q61881, Q8R180, P70206, B2RXS4, Q61072, P25206, Q8BPX9, P20152, Q9DAU1, P29416, Q8BJS4, P06797, Q8BG07 | 1 |

Supplementary Table 6: Results of the pathway analysis by the David tool on the proteins modulated in response to the 4x5µg/ml treatment

| Annotation Cluster 1 | Enrichment Score: 10.854896324612293 | | | | |
|---|---|---|---|---|---|
| Category | Term | Count | P Value | Genes | FDR |
| GOTERM_CC_DIRECT | GO:0005739~mitochondrion | 81 | 5.975388E-22 | P10639, Q8CAQ8, Q8BFR5, P07901, Q9CZW5, Q8BK72, Q9CQQ7, Q9JMH6, P48771, Q9CR62, P38647, Q9R112, P16858, P18242, Q8K0D5, Q8JZN5, Q9DBL1, O35129, P63085, Q9CRB9, P62075, Q3TIU4, P45377, P41216, Q8R1I1, O89023, Q9DCE9, Q9WV85, Q9CPR5, Q921G7, P47791, Q8CG76, Q9CQX2, Q9JHI5, Q9CR98, D3Z7P3, Q02053, P50544, Q9CYG7, P06151, P05063, Q9Z2Z6, O70404, Q921F2, Q9D0S9, Q9ERS2, P50516, Q91WD5, Q8K2B3, P63101, P42125, Q9QUJ7, Q7TNS2, P26638, P49710, Q9DC70, Q9JM90, Q8CGK3, Q91YT0, Q9CQN1, P80313, Q9CWZ7, Q61425, Q9JIA7, Q9QZD8, Q9D338, P28271, O35143, Q9WUM5, Q8BMD8, Q60692, P31786, P17665, Q99J99, Q9Z0X1, O55125, P38060, P60603, P28867, P56391, Q8BGQ7 | 6.9464E-20 |
| GOTERM_CC_DIRECT | GO:0005743~mitochondrial inner membrane | 38 | 2.95767E-16 | Q8CAQ8, Q8BFR5, P48771, Q8BK72, Q9CQQ7, Q9CR62, Q9ERS2, Q9R112, Q91WD5, Q8JZN5, Q8K2B3, O35129, P42125, Q9CRB9, P62075, Q9DC70, Q8R1I1, Q91YT0, Q9CQN1, Q9CPR5, Q61425, Q9JIA7, Q921G7, Q9QZD8, Q9D338, Q9WUM5, Q8BMD8, Q9CQX2, P17665, Q99J99, P50544, Q9Z0X1, P16110, O55125, P38060, Q9Z2Z6, P60603, P56391 | 2.7506E-14 |
| UP_KW_DOMAIN | KW-0809~Transit peptide | 34 | 2.070853E-10 | Q8CAQ8, Q9D0S9, Q8BFR5, P48771, Q8BK72, Q9CQQ7, P38647, Q9R112, Q91WD5, Q8K0D5, Q8JZN5, Q8K2B3, Q9DBL1, P42125, Q3TIU4, Q9DC70, Q8CGK3, Q91YT0, Q9CQN1, Q9CPR5, Q61425, Q921G7, P47791, Q9D338, Q8CG76, O35143, Q9WUM5, Q9JHI5, D3Z7P3, P17665, P50544, Q9Z0X1, O55125, P38060 | 4.3488E-09 |
| UP_SEQ_FEATURE | TRANSIT:Mitochondrion | 34 | 3.687019E-10 | Q8CAQ8, Q9D0S9, Q8BFR5, P48771, Q8BK72, Q9CQQ7, P38647, Q9R112, Q91WD5, Q8K0D5, Q8JZN5, Q8K2B3, Q9DBL1, P42125, Q3TIU4, Q9DC70, Q8CGK3, Q91YT0, Q9CQN1, Q9CPR5, Q61425, Q921G7, P47791, Q9D338, Q8CG76, O35143, Q9WUM5, Q9JHI5, D3Z7P3, P17665, P50544, Q9Z0X1, O55125, P38060 | 2.4021E-07 |
| UP_KW_CELLULAR_COMPONENT | KW-0496~Mitochondrion | 62 | 4.266288E-10 | Q8CAQ8, Q8BFR5, P07901, Q9CZW5, Q8BK72, Q9CQQ7, P48771, Q9CR62, P38647, Q9R112, Q8K0D5, Q8JZN5, Q9DBL1, O35129, Q9CRB9, P62075, Q3TIU4, P41216, Q8R1I1, Q9WV85, Q9CPR5, Q921G7, P47791, Q8CG76, P35564, Q9CQX2, Q9JHI5, D3Z7P3, Q02053, P50544, Q9CYG7, Q9Z2Z6, Q921F2, Q9D0S9, Q9ERS2, Q91WD5, Q8K2B3, P42125, Q9QUJ7, Q7TNS2, P49710, Q9DC70, Q8BX70, Q9JM90, Q8CGK3, Q91YT0, Q9CQN1, Q61425, Q9JIA7, Q9QZD8, Q9D338, O35143, Q9WUM5, Q8BMD8, P17665, Q99J99, Q9Z0X1, O55125, P38060, P60603, P28867, P56391 | 6.6127E-09 |
| UP_KW_CELLULAR_COMPONENT | KW-0999~Mitochondrion inner membrane | 26 | 1.864492E-08 | Q8CAQ8, P48771, Q9CQQ7, Q9CR62, Q9ERS2, Q91WD5, Q8JZN5, Q8K2B3, O35129, Q9CRB9, P62075, Q7TNS2, Q9DC70, Q8R1I1, Q91YT0, Q9CQN1, Q9JIA7, Q921G7, Q9QZD8, Q8BMD8, P17665, P50544, Q9Z0X1, Q9Z2Z6, P60603, P56391 | 1.9266E-07 |
| GOTERM_CC_DIRECT | GO:0005759~mitochondrial matrix | 12 | 0.0009659941 | Q9DBL1, Q9D0S9, P42125, Q9CQN1, Q61425, O55125, P38647, Q3TIU4, P38060, Q9JHI5, D3Z7P3, Q8CGK3 | 0.00702292 |

| Annotation Cluster 2 | Enrichment Score: 6.640591655679068 | | | | |
|---|---|---|---|---|---|
| Category | Term | Count | P Value | Genes | FDR |
| GOTERM_BP_DIRECT | GO:0002181~cytoplasmic translation | 16 | 6.299668E-11 | P51410, Q9CR57, Q9D7S7, P62849, P19253, P32233, P14206, P12970, P14131, P62702, P47963, P26638, P27659, P62858, P47911, P62830 | 1.0657E-07 |
| GOTERM_CC_DIRECT | GO:0005840~ribosome | 18 | 7.274771E-11 | P51410, Q9CR57, Q9D7S7, P62849, Q9D338, P19253, P35564, P10126, Q6NZJ6, P14206, P12970, P14131, P62702, P47963, P27659, P62858, P47911, P62830 | 3.9911E-09 |
| GOTERM_CC_DIRECT | GO:0098794~postsynapse | 27 | 7.724721E-11 | P70460, P19253, Q8BTM8, Q61792, P12970, A2A5R2, P42567, P40124, Q7TMB8, P62858, P47911, P62137, P62830, P51410, Q9CR57, P61979, P62849, Q62418, Q6NZJ6, P14206, Q99P72, P14131, P47963, Q8BPU7, P62962, P63017, P60766 | 3.9911E-09 |
| GOTERM_BP_DIRECT | GO:0006412~translation | 20 | 9.042446E-11 | P51410, Q9CR57, Q9CPR5, Q9Z0N1, Q9D7S7, P62849, Q9D338, P19253, P10126, Q6NZJ6, P14206, Q9ER72, P14131, Q8BML9, P62702, P47963, P27659, P62858, P47911, P62830 | 1.0657E-07 |
| GOTERM_CC_DIRECT | GO:0098793~presynapse | 22 | 1.443927E-08 | P51410, Q9CR57, P56399, P62849, Q62418, P19253, P35564, P12970, A2A5R2, Q8CHH9, P40124, P14131, Q9D906, P47963, P62858, Q9DCR2, P47911, P62137, P62962, P63017, P62830, P62821 | 4.1964E-07 |
| GOTERM_CC_DIRECT | GO:0022626~cytosolic ribosome | 13 | 5.744749E-08 | P51410, Q9CR57, P62849, P19253, P10126, P14206, P12970, P14131, P62702, P47963, P27659, P47911, P62830 | 1.4841E-06 |
| GOTERM_BP_DIRECT | GO:0140236~translation at presynapse | 10 | 1.681238E-07 | P51410, P14131, Q9CR57, P62849, P19253, P47963, P62858, P47911, P12970, P62830 | 5.1272E-05 |
| GOTERM_BP_DIRECT | GO:0140242~translation at postsynapse | 10 | 1.928781E-07 | P51410, P14131, Q9CR57, P62849, P19253, P47963, P62858, P47911, P12970, P62830 | 5.1272E-05 |
| GOTERM_MF_DIRECT | GO:0003735~structural constituent of ribosome | 16 | 6.482569E-07 | P51410, Q9CR57, Q9CPR5, Q9D7S7, P62849, Q9D338, P19253, P14206, P12970, P14131, P62702, P47963, P27659, P62858, P47911, P62830 | 3.059E-05 |
| UP_KW_MOLECULAR_FUNCTION | KW-0687~Ribonucleoprotein | 24 | 7.482668E-07 | P51410, Q9CR57, Q9CPR5, P61979, Q9D7S7, P62849, Q8BK72, Q9D338, P19253, Q9D6Z1, P14206, P12970, Q3UEB3, P62320, P14131, Q6DFW4, P60335, P62702, P47963, P27659, P62858, P47911, P62830, P62315 | 4.7889E-05 |
| UP_KW_MOLECULAR_FUNCTION | KW-0689~Ribosomal protein | 17 | 1.202565E-05 | P51410, Q9CR57, Q9CPR5, Q9D7S7, P62849, Q8BK72, Q9D338, P19253, P14206, P12970, P14131, P62702, P47963, P27659, P62858, P47911, P62830 | 0.00038482 |
| GOTERM_CC_DIRECT | GO:0022625~cytosolic large ribosomal subunit | 8 | 0.0010455418 | P51410, Q9CR57, P19253, P47963, P27659, P47911, P12970, P62830 | 0.00736632 |
| KEGG_PATHWAY | mmu03010:Ribosome | 16 | 0.0078070096 | P51410, Q9CR57, Q9CPR5, Q9D7S7, P62849, Q9D338, P19253, P14206, P12970, P14131, P62702, P47963, P27659, P62858, P47911, P62830 | 0.06688005 |
| KEGG_PATHWAY | mmu05171:Coronavirus disease - COVID-19 | 17 | 0.0262388239 | P51410, Q9CR57, Q64339, Q9D7S7, P62849, P08508, P19253, P14206, P12970, P14131, P63085, P62702, P47963, P27659, P62858, P47911, P62830 | 0.14942774 |

| Annotation Cluster 3 | Enrichment Score: 6.3383341882784014 | | | | |
|---|---|---|---|---|---|
| Category | Term | Count | P Value | Genes | FDR |
| GOTERM_MF_DIRECT | GO:0016887~ATP hydrolysis activity | 26 | 5.647519E-10 | P80314, P07901, Q9CQN1, P80313, Q8CG47, P46460, P46471, Q9CQQ7, P38647, Q9CR51, P97855, P50516, Q61881, P49718, P42932, O55143, P60122, P23249, Q921N6, P63017, Q8CGK3, P60710, Q9D0F6, E9Q555 | 6.0913E-08 |
| GOTERM_MF_DIRECT | GO:0005524~ATP binding | 51 | 5.122048E-09 | P07901, Q8CG47, P46471, P38647, Q8BML9, P63085, Q91YR1, P41216, Q8R1Q8, Q9WV85, P46460, P70296, P58389, Q6NZJ6, Q9ER72, Q61881, Q02053, O55222, P42932, O55143, P09581, P63017, E9Q555, Q6ZPJ3, P32921, Q6PHZ2, Q9ERS2, Q9R0N0, P50516, Q9DCL9, P49718, Q9QUJ7, P26638, P23249, Q921N6, Q8CGK3, P60710, Q9D0F6, P80314, Q9CQN1, P80313, Q9JIA7, P97855, Q9EQP2, A2AQP0, P60122, P25911, P28867, Q8BGQ7 | 4.8339E-07 |
| UP_KW_LIGAND | KW-0547~Nucleotide-binding | 78 | 5.28975E-07 | Q8BFR5, P07901, Q8CG47, P46471, Q922F4, P38647, P16858, Q8K0D5, O55131, Q8BML9, P63085, P41216, P47758, Q8R1Q8, Q9DCE9, Q9WV85, P46460, P28650, P47791, P70296, P58389, Q9Z0E6, P10126, Q9ER72, Q61881, Q02053, P35293, O55222, P42932, O55143, Q9JIW9, P09581, P63017, E9Q555, Q6ZPJ3, P32921, Q9D0S9, Q6PHZ2, Q9Z0N1, Q9R0N0, P50516, P59325, Q9DCL9, P35283, P49718, Q9QUJ7, P26638, P23249, Q921N6, Q8CGK3, P60710, Q9D0F6, P53994, Q62159, P80314, Q9CQN1, P80313, Q9JIA7, P62827, Q9WUM5, P97855, P32233, Q9EQP2, A2AQP0, Q8CHH9, P60122, Q91V41, P25911, P08556, P28867, P08752, Q8BGQ7, P60766, Q3UFY7, P62821 | 1.2166E-05 |

| | | | | | |
|---|---|---|---|---|---|
| UP_KW_LIGAND | KW-0067~ATP-binding | 48 | 0.0289685519 | P32921, P07901, Q8CG47, Q6PHZ2, P46471, P38647, Q9R0N0, P50516, Q9DCL9, P49718, Q8BML9, P63085, Q9QUJ7, P26638, P23249, Q921N6, P41216, Q8R1Q8, Q8CGK3, P60710, Q9D0F6, Q9WV85, P80314, Q9CQN1, P80313, Q9JIA7, P46460, P70296, P58389, P97855, Q9EQP2, A2AQP0, Q9ER72, Q61881, Q02053, O55222, P42932, O55143, P60122, P09581, P25911, P28867, P63017, Q8BGQ7, E9Q555, Q6ZPJ3 | 0.08675145 |
| Annotation Cluster 4 | Enrichment Score: 6.18820793899276 | | | | |
| Category | Term | Count | P Value | Genes | FDR |
| UP_SEQ_FEATURE | CROSSLNK:Glycyl lysine isopeptide (Lys-Gly) (interchain with G-C | 48 | 2.264641E-09 | Q921F2, Q3UHX2, Q9CZW5, P16858, Q99L45, P18760, P13864, O35691, P59325, P12970, Q9Z0H3, Q8CGC6, O35226, P40142, Q6DFW4, Q921T2, P60335, P20152, P47911, P27546, Q8K310, P62315, Q9CSN1, P46061, P80314, Q8BK64, Q9CR57, P80313, P61979, Q3TEA8, P62849, P62827, P97379, Q6PDM2, Q9D6Z1, Q61881, Q3UEB3, P42932, P62960, Q8C2Q3, P60122, Q9CYG7, O55201, P62702, P47963, P24547, P27659, E9Q555 | 9.8361E-07 |
| UP_KW_PTM | KW-1017~Isopeptide bond | 79 | 6.176065E-07 | Q3UHX2, Q8BL97, Q922F4, Q9CZW5, Q8BTM8, P16858, Q99L45, P18760, P13864, O35691, P12970, Q9Z0H3, P17751, P40124, P09055, Q6DFW4, Q921T2, P60335, P20152, P47911, P27546, Q8K310, P62315, P46061, Q8BK64, Q9CR57, P62849, Q6PDM2, P10126, P14206, Q61881, P14685, Q9DBG5, P42932, P62960, Q9CYG7, P06151, Q9D906, P05064, P62962, P63017, E9Q555, Q921F2, Q9CQW9, P48036, P59325, Q8CGC6, O35226, P40142, Q9CSN1, P80314, P80313, Q3UZ39, Q9CQV8, Q64339, P61979, Q3TEA8, P62827, P97379, P97855, Q9D6Z1, Q3UEB3, P17742, Q9Z0X1, P63158, P60122, Q8C2Q3, Q3U0V1, O55201, O35892, P62702, P47963, P17182, P24547, P27659, P08556, P62821 | 4.5291E-06 |
| UP_KW_PTM | KW-0832~Ubl conjugation | 97 | 0.0001948341 | Q3UHX2, P07901, Q8BL97, P46471, Q9CZW5, Q9JMH6, Q8BTM8, P16858, P18760, P13864, Q99L45, O35691, P12970, Q9Z0H3, P17751, P40124, O88738, P63085, P09055, Q6DFW4, Q921T2, P60335, A1L314, P20152, P47911, P27546, Q8K310, P62315, P46061, Q9CR57, Q8BK64, P62849, Q6PDM2, P35564, P10126, P14206, Q61881, P14685, Q9DBG5, Q02053, P42932, P62960, Q9CYG7, P06151, P01900, P09581, Q9D906, P05064, P63017, P62962, E9Q555, Q6ZPJ3, Q921F2, Q9CQW9, P48036, P59325, P42567, Q8CGC6, O35226, P40142, Q9D8B3, P23249, P60710, Q9CSN1, P80314, Q3UZ39, P80313, P61979, Q9CQV8, Q3TEA8, P62827, P97379, Q9D6Z1, P97855, P32233, Q3UEB3, P17742, Q9Z0X1, P60122, Q8C2Q3, Q3U0V1, O55201, O35892, O55125, P47963, P62702, P17182, P24547, P25911, P27659, P08556, Q8BGQ7, P62821 | 0.00107159 |
| Annotation Cluster 5 | Enrichment Score: 5.269650994457382 | | | | |
| Category | Term | Count | P Value | Genes | FDR |
| GOTERM_CC_DIRECT | GO:0005764~lysosome | 22 | 4.030076E-09 | P31996, Q9Z0J0, Q8BFR4, P20060, P18242, Q9ESY9, P50516, P35283, P17047, Q9WVJ3, O35114, P02802, O88668, Q91V41, O09043, P45377, Q8BX70, Q3TCN2, P29416, O70404, P63017, O89023 | 1.2493E-07 |
| UP_KW_CELLULAR_COMPONENT | KW-0458~Lysosome | 22 | 9.478346E-05 | P31996, Q9Z0J0, Q8BFR4, Q9JIA7, Q9CQW9, Q99J93, P20060, P18242, Q9ESY9, P50516, P35283, P17047, Q9WVJ3, O35114, Q8BPX9, P24668, Q8BX70, Q3TCN2, P29416, O70404, P63017, O89023 | 0.00058766 |
| KEGG_PATHWAY | mmu04142:Lysosome | 13 | 0.0004064445 | P31996, Q9Z0J0, Q8BFR4, Q8BVE3, P20060, P18242, P17047, O35114, P24668, O09043, Q9DCR2, P29416, O89023 | 0.00652851 |
| Annotation Cluster 6 | Enrichment Score: 4.687840123970711 | | | | |
| Category | Term | Count | P Value | Genes | FDR |
| GOTERM_MF_DIRECT | GO:0051015~actin filament binding | 17 | 7.509388E-08 | Q7TPR4, P24452, P28650, Q62418, Q8BTM8, P18760, A2AQP0, Q9QXS1, Q61792, P21107, Q7TMB8, Q8R1S4, Q8BRT1, P49710, O88342, Q91YR1, P26645 | 5.1542E-06 |
| GOTERM_MF_DIRECT | GO:0003779~actin binding | 19 | 1.139825E-06 | Q8BH43, Q7TPR4, P26041, P70460, Q62418, Q8BTM8, Q9Z0E6, P18760, A2AQP0, Q9QXS1, P19973, P40124, P21107, Q8R1S4, P09055, P09103, O88342, Q91YR1, P62962 | 5.0622E-05 |
| UP_KW_MOLECULAR_FUNCTION | KW-0009~Actin-binding | 18 | 0.0001217498 | Q8BH43, Q7TPR4, P24452, P70460, Q62418, Q8BTM8, P18760, A2AQP0, Q9QXS1, Q61792, P40124, P21107, Q7TMB8, Q8R1S4, O88342, Q91YR1, P62962, P26645 | 0.0015584 |
| GOTERM_CC_DIRECT | GO:0015629~actin cytoskeleton | 13 | 0.0002162229 | Q7TPR4, P21107, P24452, Q8R1S4, Q8BTM8, P05064, Q9Z0E6, P18760, O88342, Q91YR1, P60710 | 0.00193353 |
| GOTERM_BP_DIRECT | GO:0030036~actin cytoskeleton organization | 11 | 0.0016143093 | P40124, Q8BH43, Q7TPR4, P70460, P62827, Q8BTM8, Q8BPU7, O88342, Q9QXS1, P62962, P60766 | 0.05912267 |
| Annotation Cluster 7 | Enrichment Score: 4.368735244034204 | | | | |
| Category | Term | Count | P Value | Genes | FDR |
| GOTERM_MF_DIRECT | GO:0003924~GTPase activity | 25 | 7.728834E-12 | Q62159, P53994, Q9DCE9, Q61599, Q8BFR5, Q9Z0N1, P28650, Q922F4, P62827, Q9Z0E6, Q8VCN9, P32233, P10126, Q8K0D5, P35283, Q8CHH9, P35293, O55131, Q9JIW9, Q91V41, P08556, P08752, P60766, P62821 | 1.1671E-09 |
| GOTERM_MF_DIRECT | GO:0005525~GTP binding | 27 | 1.92078E-11 | Q8BFR5, P07901, Q9Z0N1, Q922F4, P59325, Q8K0D5, P35283, O55131, P47758, Q62159, P53994, Q9DCE9, P28650, P62827, Q9Z0E6, P32233, P10126, Q9EQP2, Q8CHH9, P35293, Q9JIW9, Q91V41, P08556, P08752, P60766, P62821 | 2.417E-09 |
| GOTERM_MF_DIRECT | GO:0019003~GDP binding | 11 | 1.889246E-07 | P35283, P53994, P35293, P62827, Q9JIW9, Q91V41, Q9WUM5, Q9Z0E6, P08556, Q8R1Q8 | 1.0933E-05 |
| INTERPRO | IPR027417:P-loop_NTPase | 39 | 1.855517E-06 | Q8BFR5, Q8CG47, Q9Z0N1, P46471, Q9JLB0, P50516, Q8K0D5, P35283, P49718, O55131, P23249, Q921N6, P47758, Q8R1Q8, Q8CGK3, Q9D0F6, Q62159, P53994, Q9DCE9, P46460, P28650, P62827, Q9Z0E6, P32233, P10126, Q9EQP2, A2AQP0, Q61881, Q8CHH9, P35293, P60122, Q9JIW9, Q91V41, P08556, P08752, P60766, E9Q555, P62821 | 0.00261257 |
| UP_KW_LIGAND | KW-0342~GTP-binding | 25 | 9.418284E-06 | Q62159, P53994, Q9DCE9, Q8BFR5, Q9Z0N1, P28650, Q922F4, P62827, Q9Z0E6, P32233, P10126, P59325, Q8K0D5, P35283, Q8CHH9, P35293, O55131, Q9JIW9, Q91V41, P08556, P47758, P08752, P60766, P62821 | 0.00010831 |
| GOTERM_MF_DIRECT | GO:0003925~G protein activity | 8 | 1.003297E-05 | Q9DCE9, P62827, Q9JIW9, Q9Z0E6, P08556, P60766, P62821 | 0.00032934 |
| UP_SEQ_FEATURE | MOTIF:Effector region | 10 | 3.040213E-05 | P35283, Q62159, P53994, P35293, Q9JIW9, Q91V41, P08556, P60766, P62821 | 0.0079228 |
| INTERPRO | IPR005225:Small_GTP-bd_dom | 13 | 4.704738E-05 | P35283, Q62159, P53994, P35293, P62827, Q9JIW9, Q91V41, P32233, P08556, Q8K0D5, P60766, P62821 | 0.03312136 |
| SMART | SM00174:RHO | 11 | 8.734124E-05 | P35283, Q62159, P53994, P35293, P62827, Q9JIW9, Q91V41, P08556, P60766, P62821 | 0.01297779 |
| SMART | SM00173:RAS | 11 | 0.0001466417 | P35283, Q62159, P53994, P35293, P62827, Q9JIW9, Q91V41, P08556, P60766, P62821 | 0.01297779 |
| INTERPRO | IPR001806:Small_GTPase | 11 | 0.0001555915 | P35283, Q62159, P53994, P35293, P62827, Q9JIW9, Q91V41, P08556, P60766, P62821 | 0.07302426 |
| SMART | SM00175:RAB | 11 | 0.000411058 | P35283, Q62159, P53994, P35293, P62827, Q9JIW9, Q91V41, P08556, P60766, P62821 | 0.01818932 |
| UP_SEQ_FEATURE | LIPID:S-geranylgeranyl cysteine | 9 | 0.0004457424 | P35283, Q62159, P53994, P35293, Q9JIW9, Q91V41, Q9Z0E6, P60766, P62821 | 0.05280021 |
| SMART | SM00176:RAN | 7 | 0.0038668945 | P35283, P53994, P35293, P62827, Q9JIW9, Q91V41, P62821 | 0.11407339 |
| UP_SEQ_FEATURE | PROPEP:Removed in mature form | 13 | 0.0040642809 | Q62159, P35293, Q9Z0X1, Q64339, P10810, Q9JIW9, Q9Z0E6, Q9ESY9, P08556, Q60692, P60766, O89023 | 0.25217895 |
| UP_KW_PTM | KW-0636~Prenylation | 11 | 0.0175059805 | P35283, Q62159, P53994, P35293, Q9JIW9, Q91V41, Q9Z0E6, P08556, P60766, P62821 | 0.0550188 |
| GOTERM_CC_DIRECT | GO:0012505~endomembrane system | 4 | 0.3809098419 | P35283, P35293, Q91V41, P62821 | 0.67863248 |
| UP_KW_PTM | KW-0449~Lipoprotein | 22 | 0.8565950364 | Q62159, P53994, Q99JX3, Q9CQW9, P10810, Q99J93, Q9Z0E6, P35564, P35283, P35293, O35114, Q9CRB9, Q8BMK4, Q9JIW9, Q91V41, P25911, P08556, P08752, P60766, P26645, P62821 | 1 |
| Annotation Cluster 8 | Enrichment Score: 3.961543116303502 | | | | |

| Category | Term | Count | P Value | Genes | FDR |
|---|---|---|---|---|---|
| GOTERM_CC_DIRECT | GO:0002102~podosome | 12 | 3.428491E-11 | D3Z6Q9, P21107, Q60875, P61979, Q62418, Q8BTM8, O88342, Q9QXS1, P35831, P60710 | 2.2775E-09 |
| GOTERM_CC_DIRECT | GO:0005903~brush border | 9 | 2.38764E-05 | P21107, Q7TPR4, Q60605, Q8BTM8, A2AQP0, Q9QXS1, P60710 | 0.00027756 |
| GOTERM_CC_DIRECT | GO:0005884~actin filament | 9 | 8.394886E-05 | P21107, Q7TPR4, Q62418, Q8BTM8, P49710, Q91YR1, P60710 | 0.00083056 |
| GOTERM_CC_DIRECT | GO:0030863~cortical cytoskeleton | 6 | 9.152825E-05 | P21107, Q7TPR4, Q8BTM8, P60710 | 0.00086858 |
| GOTERM_CC_DIRECT | GO:0015629~actin cytoskeleton | 13 | 0.0002162229 | Q7TPR4, P21107, P24452, Q8R1S4, Q8BTM8, P05064, Q9Z0E6, P18760, O88342, Q91YR1, P60710 | 0.00193353 |
| GOTERM_CC_DIRECT | GO:0001725~stress fiber | 8 | 0.0005261638 | O55222, O55131, P21107, Q7TPR4, A2AQP0, P60710 | 0.00436904 |
| UP_KW_MOLECULAR_FUNCTION | KW-0514~Muscle protein | 4 | 0.1464327452 | P21107, Q60605, A2AQP0, P60710 | 0.49324714 |
| KEGG_PATHWAY | mmu04814:Motor proteins | 9 | 0.1937993545 | P21107, Q922F4, Q60605, A2AQP0, P62627, Q8R1Q8, P60710 | 0.60007752 |

Annotation Cluster 9 — Enrichment Score: 3.7898930392007433

| Category | Term | Count | P Value | Genes | FDR |
|---|---|---|---|---|---|
| GOTERM_BP_DIRECT | GO:0006397~mRNA processing | 17 | 1.420215E-07 | Q921F2, Q8BL97, P61979, P29341, O35691, Q3UEB3, O35326, P62960, Q8CGC6, P16110, Q62093, Q3U0V1, P60335, Q3TIU4, P84104, P63017, Q9DBR1 | 5.1272E-05 |
| GOTERM_BP_DIRECT | GO:0008380~RNA splicing | 15 | 2.175324E-07 | Q921F2, Q8BL97, P61979, P29341, Q6PDM2, O35691, Q3UEB3, O35326, P62960, Q8CGC6, P16110, Q62093, Q3U0V1, P84104, P63017 | 5.1272E-05 |
| GOTERM_CC_DIRECT | GO:0005681~spliceosomal complex | 13 | 6.405413E-07 | Q9CSN1, P61979, P29341, Q6PDM2, O35691, Q64213, P62320, Q8CGC6, P16110, Q62093, Q99KP6, P63017, P62315 | 1.241E-05 |
| UP_KW_BIOLOGICAL_PROCESS | KW-0508~mRNA splicing | 20 | 1.481647E-05 | Q9CSN1, Q921F2, Q8BL97, P61979, P29341, Q6PDM2, O35691, Q64213, Q3UEB3, O35326, P62320, P62960, Q8CGC6, P16110, Q62093, Q99KP6, Q3U0V1, P84104, P63017, P62315 | 0.00064452 |
| UP_KW_BIOLOGICAL_PROCESS | KW-0507~mRNA processing | 22 | 3.461787E-05 | Q9CSN1, Q921F2, Q8BL97, P61979, P29341, Q6PDM2, O35691, Q64213, Q3UEB3, O35326, P62320, P62960, Q8CGC6, P16110, Q62093, Q99KP6, Q3U0V1, Q3TIU4, P84104, P63017, Q9DBR1, P62315 | 0.00085892 |
| GOTERM_BP_DIRECT | GO:0000398~mRNA splicing, via spliceosome | 11 | 0.0001298376 | O35326, Q9CSN1, P62320, Q8BL97, Q8C2Q3, Q62093, Q99KP6, P84104, Q64213, P63017, P62315 | 0.01020091 |
| SMART | SM00360:RRM | 14 | 0.0002336008 | Q921F2, Q8BL97, P97379, P29341, Q6PDM2, Q9Z1D1, P97855, Q3UEB3, O35326, Q8CGC6, Q8C2Q3, Q62093, P84104, Q8K310 | 0.01378245 |
| GOTERM_CC_DIRECT | GO:0071013~catalytic step 2 spliceosome | 9 | 0.0002361646 | Q9CSN1, P62320, P61979, Q99KP6, P29341, Q6PDM2, O35691, Q3UEB3, P62315 | 0.00207201 |
| UP_SEQ_FEATURE | DOMAIN:RRM | 14 | 0.0002493431 | Q921F2, Q8BL97, P97379, P29341, Q6PDM2, Q9Z1D1, P97855, Q3UEB3, O35326, Q8CGC6, Q8C2Q3, Q62093, P84104, Q8K310 | 0.03830142 |
| UP_KW_CELLULAR_COMPONENT | KW-0747~Spliceosome | 12 | 0.0002938793 | Q9CSN1, P62320, Q8CGC6, P61979, P16110, Q99KP6, P29341, Q6PDM2, O35691, Q64213, P63017, P62315 | 0.00151838 |
| INTERPRO | IPR000504:RRM_dom | 14 | 0.0003779131 | Q921F2, Q8BL97, P97379, P29341, Q6PDM2, Q9Z1D1, P97855, Q3UEB3, O35326, Q8CGC6, Q8C2Q3, Q62093, P84104, Q8K310 | 0.09270193 |
| INTERPRO | IPR012677:Nucleotide-bd_a/b_plait_sf | 15 | 0.0004608761 | Q921F2, Q8BL97, P97379, P29341, Q6PDM2, Q9Z1D1, P97855, Q3UEB3, O35326, Q8CGC6, Q8C2Q3, Q2EMV9, Q62093, P84104, Q8K310 | 0.09270193 |
| UP_SEQ_FEATURE | DOMAIN:RRM 1 | 8 | 0.0009439206 | O35326, Q921F2, Q8CGC6, Q8C2Q3, P29341, Q6PDM2, Q3UEB3, Q8K310 | 0.08199524 |
| UP_SEQ_FEATURE | DOMAIN:RRM 2 | 8 | 0.0009439206 | O35326, Q921F2, Q8CGC6, Q8C2Q3, P29341, Q6PDM2, Q3UEB3, Q8K310 | 0.08199524 |
| INTERPRO | IPR035979:RBD_domain_sf | 14 | 0.0011358471 | Q921F2, Q8BL97, P97379, P29341, Q6PDM2, Q9Z1D1, P97855, Q3UEB3, O35326, Q8CGC6, Q8C2Q3, Q62093, P84104, Q8K310 | 0.13327273 |
| UP_KW_MOLECULAR_FUNCTION | KW-0694~RNA-binding | 28 | 0.0055374829 | Q921F2, Q8BL97, Q8BK72, Q9Z1D1, O35326, Q8CGC6, Q62093, P84104, P60335, P23249, Q8K310, P61979, P97379, P29341, Q6PDM2, P28271, P97855, Q6NZJ6, Q64213, Q3UEB3, P62320, P62960, Q8C2Q3, Q3U0V1, P62702, P23116, P24547, Q8BGQ7 | 0.03937766 |
| GOTERM_CC_DIRECT | GO:0016607~nuclear speck | 13 | 0.0165271547 | Q9CSN1, Q921F2, Q8BL97, Q8CG47, Q3TEA8, Q6PDM2, O35691, O35326, Q8C2Q3, Q62093, Q99KP6, P60335, P84104 | 0.06351345 |
| KEGG_PATHWAY | mmu03040:Spliceosome | 13 | 0.0704240948 | Q9CSN1, Q8BL97, P61979, Q6PDM2, Q3UEB3, O35326, P62320, Q62093, Q99KP6, P60335, P84104, P63017, P62315 | 0.30164987 |

Annotation Cluster 10 — Enrichment Score: 3.7503100647827416

| Category | Term | Count | P Value | Genes | FDR |
|---|---|---|---|---|---|
| GOTERM_BP_DIRECT | GO:0046166~glyceraldehyde-3-phosphate biosynthetic process | 6 | 4.724053E-07 | Q9QZD8, P40142, P16858, P17182, P17751 | 0.00010122 |
| KEGG_PATHWAY | mmu01200:Carbon metabolism | 15 | 7.738876E-06 | Q99K85, P28271, Q8VDL4, Q9R0P3, P16858, Q9WUM5, P17751, Q8K2B3, P40142, Q00612, P05063, P05064, P17182 | 0.00066296 |
| UP_KW_BIOLOGICAL_PROCESS | KW-0324~Glycolysis | 7 | 3.949055E-05 | P05063, Q8VDL4, P16858, P05064, P17182, P17751 | 0.00085892 |
| GOTERM_BP_DIRECT | GO:0006096~glycolytic process | 7 | 5.093582E-05 | P05063, Q8VDL4, P16858, P05064, P17182, P17751 | 0.00480223 |
| GOTERM_BP_DIRECT | GO:0006094~gluconeogenesis | 7 | 7.11663E-05 | Q9QZD8, Q9CR62, P05063, P16858, P17182, P17751 | 0.0064515 |
| GOTERM_BP_DIRECT | GO:0061621~canonical glycolysis | 5 | 8.056168E-05 | P16858, P05064, P17182, P17751 | 0.00655422 |
| KEGG_PATHWAY | mmu01230:Biosynthesis of amino acids | 10 | 0.0003519316 | Q99K85, P40142, P28271, P05063, P16858, P05064, P17182, P17751, Q8VCN5 | 0.00646046 |
| KEGG_PATHWAY | mmu00010:Glycolysis / Gluconeogenesis | 8 | 0.002636855 | P06151, P05063, Q8VDL4, P16858, P05064, P17182, P17751 | 0.03080326 |
| UP_KW_MOLECULAR_FUNCTION | KW-0456~Lyase | 10 | 0.0179220257 | O35215, P28271, P05063, P05064, P38060, P17182, P17751, Q8VCN5, Q9DCL9 | 0.09558414 |
| KEGG_PATHWAY | mmu04066:HIF-1 signaling pathway | 8 | 0.0447770686 | Q6PHZ2, P63085, P06151, P05063, P16858, P05064, P17182 | 0.22130205 |

Annotation Cluster 11 — Enrichment Score: 3.5539206122034943

| Category | Term | Count | P Value | Genes | FDR |
|---|---|---|---|---|---|
| GOTERM_BP_DIRECT | GO:0045087~innate immune response | 21 | 2.286132E-05 | Q9DCE9, Q60875, Q9CR67, Q64339, P20491, P26151, P10810, Q99J93, P97379, P16858, P97855, P63158, Q8C2Q3, P16110, Q2EMV9, P09581, Q9D906, Q8BPX9, P25911, Q8K310, Q8BG07 | 0.00256591 |
| UP_KW_BIOLOGICAL_PROCESS | KW-0399~Innate immunity | 22 | 5.490612E-05 | Q9DCE9, Q60875, Q9CR67, Q9CQW9, P20491, P26151, P10810, Q99J93, P97379, P16858, P97855, Q9Z0E6, P63158, Q8C2Q3, P16110, Q2EMV9, P09581, A1L314, Q8BPX9, P25911, Q8K310, Q8BG07 | 0.00095537 |
| UP_KW_BIOLOGICAL_PROCESS | KW-0391~Immunity | 29 | 0.0173586575 | Q9CQW9, Q9CR67, P10810, Q8R5M8, P16858, Q9ESY9, Q2EMV9, O08573, A1L314, P01897, Q8BPX9, Q8K310, Q9DCE9, Q60875, P20491, P26151, Q99J93, P97379, Q62418, Q9Z0E6, P97855, P63158, Q8C2Q3, P16110, P09581, P01900, P25911, E9Q555, Q8BG07 | 0.16780036 |

Annotation Cluster 12 — Enrichment Score: 3.1674992455601916

| Category | Term | Count | P Value | Genes | FDR |
|---|---|---|---|---|---|
| KEGG_PATHWAY | mmu04670:Leukocyte transendothelial migration | 14 | 2.791595E-05 | P05555, Q7TPR4, P27870, P26041, P70460, O70145, P24063, P11835, P09055, Q9R0C8, P08752, P60766, P60710 | 0.00143488 |
| KEGG_PATHWAY | mmu04810:Regulation of actin cytoskeleton | 19 | 9.234356E-05 | P05555, Q8BH43, Q7TPR4, P27870, P26041, P24063, P11835, P18760, Q7TMB8, P63085, P09055, Q9R0C8, P08556, P62137, P62962, P60766, P60710 | 0.00263692 |
| KEGG_PATHWAY | mmu04015:Rap1 signaling pathway | 17 | 0.0003441737 | P05555, P27870, P70460, P24063, P11835, P63085, P09055, Q9JIW9, P09581, Q9R0C8, P08556, P62962, P08752, P60766, P60710 | 0.00646046 |
| KEGG_PATHWAY | mmu04510:Focal adhesion | 14 | 0.0045627905 | Q7TPR4, P27870, P70460, Q8BTM8, Q61120, O55222, P63085, P09055, Q9R0C8, P08556, P62137, P60766, P60710 | 0.04510143 |
| KEGG_PATHWAY | mmu05135:Yersinia infection | 11 | 0.004932506 | Q8BH43, P27870, P63085, P09055, P08508, Q3UND0, Q9R0C8, Q8BPU7, P60766, P60710 | 0.04544067 |
| KEGG_PATHWAY | mmu05205:Proteoglycans in cancer | 14 | 0.0049507349 | P27870, Q6PHZ2, P26041, P63085, P09055, Q8BTM8, Q9R0C8, P49710, P08556, P62137, P60766, P60710 | 0.04544067 |

Annotation Cluster 13 — Enrichment Score: 3.115468715233612

| Category | Term | Count | P Value | Genes | FDR |
|---|---|---|---|---|---|
| UP_KW_DOMAIN | KW-0728~SH3 domain | 13 | 0.0002218122 | Q91ZR2, P27870, Q9JLB0, O70145, Q62418, Q3UND0, Q9QXS1, Q61792, Q62422, Q8CBW3, Q9R0C8, P49710, P25911 | 0.00155269 |
| INTERPRO | IPR001452:SH3_domain | 13 | 0.0006377135 | Q91ZR2, P27870, Q9JLB0, O70145, Q62418, Q3UND0, Q9QXS1, Q61792, Q62422, Q8CBW3, Q9R0C8, P49710, P25911 | 0.09976673 |
| SMART | SM00326:SH3 | 12 | 0.0009847898 | Q91ZR2, P27870, O70145, Q9JLB0, Q62418, Q3UND0, Q9R0C8, P49710, P25911, Q61792, Q62422, Q8CBW3 | 0.03486156 |
| UP_SEQ_FEATURE | DOMAIN:SH3 | 12 | 0.0011983771 | Q91ZR2, P27870, O70145, Q9JLB0, Q62418, Q3UND0, P49710, P25911, Q9QXS1, Q61792, Q62422, Q8CBW3 | 0.09759283 |

| | | | | | |
|---|---|---|---|---|---|
| INTERPRO | IPR036028:SH3-like_dom_sf | 12 | 0.0015852887 | Q91ZR2, P27870, O70145, Q9JLB0, Q62418, Q3UND0, Q9R0C8, P49710, P25911, Q61792, Q62422, Q8CBW3 | 0.15482308 |
| Annotation Cluster 14 | Enrichment Score: 3.071569698865659 | | | | |
| Category | Term | Count | P Value | Genes | FDR |
| KEGG_PATHWAY | mmu05415:Diabetic cardiomyopathy | 20 | 1.090687E-05 | Q6PHZ2, O70145, P47791, P48771, Q9CQQ7, Q9ERS2, P16858, P58389, P18242, Q91WD5, Q8K2B3, P17665, O55143, Q00612, Q9DC70, P28867, P62137, Q8R1I1, P56391, Q91YT0 | 0.00070077 |
| KEGG_PATHWAY | mmu05014:Amyotrophic lateral sclerosis | 26 | 5.261862E-05 | Q921F2, Q07076, O70435, Q8BL97, P46471, Q922F4, P48771, Q9CQQ7, Q9ERS2, Q91WD5, Q8K2B3, O35226, P84104, Q9DC70, Q8R1I1, Q8K310, P60710, Q91YT0, Q9QY81, Q60692, P14685, P17665, P62962, P56391, P62821 | 0.00225383 |
| KEGG_PATHWAY | mmu05012:Parkinson disease | 21 | 8.858128E-05 | P10639, Q9CQN1, O70435, Q6PHZ2, P46471, Q922F4, P48771, Q9CQQ7, Q9ERS2, Q60692, Q91WD5, P14685, Q8K2B3, P17665, Q02053, O35226, Q9DC70, Q8R1I1, P56391, P08752, Q91YT0 | 0.00263692 |
| KEGG_PATHWAY | mmu05020:Prion disease | 20 | 0.0002463795 | O70435, P46471, Q922F4, O70145, P48771, Q9CQQ7, Q9ERS2, Q60692, Q91WD5, P14685, Q8K2B3, P17665, O35226, P63085, Q9DC70, P28867, Q8R1I1, P56391, P63017, Q91YT0 | 0.00575632 |
| KEGG_PATHWAY | mmu04714:Thermogenesis | 18 | 0.0003862299 | P48771, Q9CQQ7, Q9ERS2, Q9Z0H3, Q91WD5, Q8K2B3, P17665, Q9QUJ7, Q9Z2Z6, Q9DC70, P08556, P41216, Q8R1I1, P56391, P60710, Q91YT0 | 0.00652851 |
| KEGG_PATHWAY | mmu00190:Oxidative phosphorylation | 13 | 0.000603231 | Q8BVE3, P48771, Q9CQQ7, Q9ERS2, Q9CR51, P50516, Q91WD5, Q8K2B3, P17665, Q9DC70, Q8R1I1, P56391, Q91YT0 | 0.0086128 |
| GOTERM_BP_DIRECT | GO:0042776~proton motive force-driven mitochondrial ATP synthes | 6 | 0.0016078581 | Q9CQQ7, Q9ERS2, Q9DC70, Q91WD5, Q8K2B3, Q91YT0 | 0.05912267 |
| KEGG_PATHWAY | mmu05208:Chemical carcinogenesis - reactive oxygen species | 16 | 0.0020093965 | O70145, P48771, Q9CQQ7, Q9ERS2, Q91WD5, Q8K2B3, P17665, P63085, P48774, Q9DC70, P08556, P28867, Q8R1I1, P56391, Q91YT0 | 0.02459119 |
| KEGG_PATHWAY | mmu05010:Alzheimer disease | 22 | 0.0031094331 | O70435, P46471, Q922F4, P48771, Q9CQQ7, Q9ERS2, P16858, Q60692, Q99P72, Q91WD5, P14685, Q8K2B3, P17665, O55143, O35226, P63085, Q9DC70, P08556, Q8R1I1, P56391, Q91YT0 | 0.03474454 |
| KEGG_PATHWAY | mmu05022:Pathways of neurodegeneration - multiple diseases | 25 | 0.0044785468 | Q921F2, O70435, Q9CQN1, Q6PHZ2, P46471, Q922F4, P48771, Q9CQQ7, Q9ERS2, Q91WD5, Q60692, P14685, Q8K2B3, P17665, Q02053, O55143, O35226, P63085, Q9DC70, P08556, Q8R1I1, P56391, Q91YT0, P62821 | 0.04510143 |
| KEGG_PATHWAY | mmu05016:Huntington disease | 16 | 0.0270472942 | O70435, P46471, Q922F4, P48771, Q9CQQ7, Q9ERS2, Q60692, Q91WD5, P14685, Q8K2B3, P17665, O35226, Q9DC70, Q8R1I1, P56391, Q91YT0 | 0.14942774 |
| KEGG_PATHWAY | mmu04932:Non-alcoholic fatty liver disease | 10 | 0.0389755438 | P48771, Q9ERS2, Q9DC70, Q91WD5, P56391, Q8R1I1, P60766, Q8K2B3, P17665, Q91YT0 | 0.2003343 |
| Annotation Cluster 15 | Enrichment Score: 2.8473752138704427 | | | | |
| Category | Term | Count | P Value | Genes | FDR |
| UP_KW_BIOLOGICAL_PROCESS | KW-0648~Protein biosynthesis | 16 | 1.617591E-07 | P32921, Q8BFR5, Q9Z0N1, Q9Z1D1, Q99L45, P10126, Q6NZJ6, P59325, P57776, Q9ER72, Q8K0D5, Q8BML9, P23116, P26638, P60229, Q8BGQ7 | 1.4073E-05 |
| GOTERM_BP_DIRECT | GO:0001732~formation of cytoplasmic translation initiation complex | 5 | 4.893329E-05 | P23116, Q9Z1D1, Q99L45, P59325, P60229 | 0.00480223 |
| GOTERM_MF_DIRECT | GO:0003743~translation initiation factor activity | 7 | 0.0002823095 | Q9Z0N1, P23116, Q9Z1D1, Q99L45, Q6NZJ6, P59325, P60229 | 0.00734978 |
| UP_KW_MOLECULAR_FUNCTION | KW-0396~Initiation factor | 7 | 0.0012691146 | Q9Z0N1, P23116, Q9Z1D1, Q99L45, Q6NZJ6, P59325, P60229 | 0.01160333 |
| GOTERM_BP_DIRECT | GO:0006413~translational initiation | 6 | 0.0014983086 | Q9Z0N1, Q9Z1D1, Q99L45, Q6NZJ6, P59325, P60229 | 0.05912267 |
| BIOCARTA | m_eifPathway:Eukaryotic protein translation | 4 | 0.0083230224 | Q9Z0N1, P23116, Q99L45, P59325 | 0.43775475 |
| GOTERM_BP_DIRECT | GO:0001731~formation of translation preinitiation complex | 3 | 0.0089201105 | Q9Z0N1, Q99L45, P59325 | 0.19288716 |
| BIOCARTA | m_eif2Pathway:Regulation of eIF2 | 4 | 0.0114641682 | Q9Z0N1, Q99L45, P59325, P62137 | 0.43775475 |
| GOTERM_CC_DIRECT | GO:0005852~eukaryotic translation initiation factor 3 complex | 3 | 0.0184544992 | P23116, Q9Z1D1, P60229 | 0.07033887 |
| GOTERM_CC_DIRECT | GO:0033290~eukaryotic 48S preinitiation complex | 3 | 0.0231257374 | P23116, Q9Z1D1, P60229 | 0.08672152 |
| GOTERM_CC_DIRECT | GO:0016282~eukaryotic 43S preinitiation complex | 3 | 0.0309300124 | P23116, Q9Z1D1, P60229 | 0.10461011 |
| Annotation Cluster 16 | Enrichment Score: 2.731596997906237 | | | | |
| Category | Term | Count | P Value | Genes | FDR |
| GOTERM_MF_DIRECT | GO:0050660~flavin adenine dinucleotide binding | 7 | 0.0003855345 | Q9DBL1, P47791, Q9JMH6, Q9JHI5, Q8JZN5, Q8K2B3, P50544 | 0.00938963 |
| INTERPRO | IPR006089:Acyl-CoA_DH_CS | 4 | 0.0004337968 | Q9DBL1, Q9JHI5, Q8JZN5, P50544 | 0.09270193 |
| COG_ONTOLOGY | Lipid metabolism | 9 | 0.0005693856 | Q9DBL1, P42125, Q61425, O70310, Q9QUJ7, Q9JHI5, P41216, Q8JZN5, P50544 | 0.0079714 |
| INTERPRO | IPR013786:AcylCoA_DH/ox_N | 4 | 0.0010918452 | Q9DBL1, Q9JHI5, Q8JZN5, P50544 | 0.13327273 |
| INTERPRO | IPR009075:AcylCo_DH/oxidase_C | 4 | 0.0010918452 | Q9DBL1, Q9JHI5, Q8JZN5, P50544 | 0.13327273 |
| UP_KW_LIGAND | KW-0274~FAD | 11 | 0.0011022652 | Q9DBL1, Q9Z0X1, Q921G7, P47791, Q9JMH6, Q9R112, Q9JI75, Q9JHI5, Q8JZN5, Q8K2B3, P50544 | 0.00633802 |
| GOTERM_BP_DIRECT | GO:0006635~fatty acid beta-oxidation | 6 | 0.0011142519 | Q9DBL1, P42125, Q61425, Q9JHI5, Q8JZN5, P50544 | 0.04955268 |
| INTERPRO | IPR037069:AcylCoA_DH/ox_N_sf | 4 | 0.0017593532 | Q9DBL1, Q9JHI5, Q8JZN5, P50544 | 0.15482308 |
| INTERPRO | IPR006091:Acyl-CoA_Oxase/DH_mid-dom | 4 | 0.0021703382 | Q9DBL1, Q9JHI5, Q8JZN5, P50544 | 0.17975507 |
| INTERPRO | IPR036250:AcylCo_DH-like_C | 4 | 0.0026361712 | Q9DBL1, Q9JHI5, Q8JZN5, P50544 | 0.18558646 |
| INTERPRO | IPR009100:AcylCoA_DH/oxidase_NM_dom_sf | 4 | 0.0026361712 | Q9DBL1, Q9JHI5, Q8JZN5, P50544 | 0.18558646 |
| INTERPRO | IPR046373:Acyl-CoA_Oxase/DH_mid-dom_sf | 4 | 0.0026361712 | Q9DBL1, Q9JHI5, Q8JZN5, P50544 | 0.18558646 |
| GOTERM_MF_DIRECT | GO:0003995~acyl-CoA dehydrogenase activity | 3 | 0.0073188094 | Q9DBL1, Q8JZN5, P50544 | 0.09527071 |
| UP_SEQ_FEATURE | DOMAIN:Acyl-CoA dehydrogenase/oxidase C-terminal | 3 | 0.012196291 | Q9DBL1, Q8JZN5, P50544 | 0.54799197 |
| UP_SEQ_FEATURE | DOMAIN:Acyl-CoA dehydrogenase/oxidase N-terminal | 3 | 0.012196291 | Q9DBL1, Q8JZN5, P50544 | 0.54799197 |
| Annotation Cluster 17 | Enrichment Score: 2.723344946282976 | | | | |
| Category | Term | Count | P Value | Genes | FDR |
| UP_KW_LIGAND | KW-0285~Flavoprotein | 12 | 0.0003888893 | Q9DBL1, Q9Z0X1, Q921G7, P47791, Q9JMH6, Q9R112, Q9JI75, Q9JHI5, Q8JZN5, Q8K2B3, P50544, Q91YT0 | 0.00298148 |
| UP_SEQ_FEATURE | DOMAIN:FAD/NAD(P)-binding | 4 | 0.0003964664 | Q9Z0X1, P47791, Q9JMH6, Q9R112 | 0.05165957 |
| GOTERM_MF_DIRECT | GO:0016491~oxidoreductase activity | 11 | 0.0005399094 | Q9Z0X1, Q921G7, P48771, Q8CG76, Q9JMH6, P45376, Q9R112, Q9JI75, O70503, Q8K354, Q8K2B3 | 0.01235247 |
| UP_KW_LIGAND | KW-0274~FAD | 11 | 0.0011022652 | Q9DBL1, Q9Z0X1, Q921G7, P47791, Q9JMH6, Q9R112, Q9JI75, Q9JHI5, Q8JZN5, Q8K2B3, P50544 | 0.00633802 |
| INTERPRO | IPR023753:FAD/NAD-binding_dom | 4 | 0.0014007431 | Q9Z0X1, P47791, Q9JMH6, Q9R112 | 0.15171125 |
| INTERPRO | IPR036188:FAD/NAD-bd_sf | 6 | 0.0056162456 | Q9Z0X1, Q921G7, P47791, Q9JMH6, Q9R112, Q8K2B3 | 0.30570487 |
| INTERPRO | IPR016156:FAD/NAD-linked_Rdtase_dimer_sf | 3 | 0.0104687858 | Q9Z0X1, P47791, Q9JMH6 | 0.45796501 |
| GOTERM_MF_DIRECT | GO:0071949~FAD binding | 4 | 0.0216222149 | Q9Z0X1, Q9JMH6, Q9R112, Q9JI75 | 0.20929195 |
| Annotation Cluster 18 | Enrichment Score: 2.7149128697695986 | | | | |
| Category | Term | Count | P Value | Genes | FDR |
| GOTERM_BP_DIRECT | GO:0006898~receptor-mediated endocytosis | 8 | 3.007371E-05 | P05555, P40124, P30204, Q9CQW9, P26151, P10810, O35114, P11835 | 0.00322199 |
| GOTERM_MF_DIRECT | GO:0038024~cargo receptor activity | 4 | 0.0127892994 | P05555, P30204, O35114, P11835 | 0.14411822 |
| GOTERM_BP_DIRECT | GO:0097242~amyloid-beta clearance | 3 | 0.0186306355 | P05555, P30204, P11835 | 0.29081065 |
| Annotation Cluster 19 | Enrichment Score: 2.6482652443058377 | | | | |
| Category | Term | Count | P Value | Genes | FDR |
| GOTERM_BP_DIRECT | GO:0006897~endocytosis | 13 | 9.297061E-06 | Q8BH43, Q91ZR2, Q8BVE3, Q9D1J1, Q62418, Q9EQP2, P42567, O88668, P08556, Q8BL66, Q810B6, P60766, P62821 | 0.00128901 |
| GOTERM_BP_DIRECT | GO:0016197~endosomal transport | 5 | 0.0098229512 | P42567, Q91ZR2, Q9EQP2, Q810B6, P60766 | 0.2067205 |
| UP_KW_BIOLOGICAL_PROCESS | KW-0254~Endocytosis | 6 | 0.1243416356 | P42567, P30204, Q91ZR2, Q62418, Q9D1J1, Q810B6 | 0.44673127 |
| Annotation Cluster 20 | Enrichment Score: 2.6204671509795356 | | | | |
| Category | Term | Count | P Value | Genes | FDR |
| GOTERM_BP_DIRECT | GO:0006457~protein folding | 13 | 1.497693E-07 | P80314, Q8BK64, Q9CQN1, P80313, P07901, P38647, P35564, Q8VCN9, P17742, P42932, P09103, P48428, P63017 | 5.1272E-05 |
| GOTERM_MF_DIRECT | GO:0140662~ATP-dependent protein folding chaperone | 7 | 2.336911E-06 | P80314, P80313, P42932, Q9CQN1, P07901, P38647, P63017 | 9.2861E-05 |
| GOTERM_MF_DIRECT | GO:0044183~protein folding chaperone | 7 | 9.566149E-05 | P80314, P80313, P42932, P07901, Q3U0V1, P38647, P63017 | 0.00288898 |
| UP_KW_MOLECULAR_FUNCTION | KW-0143~Chaperone | 16 | 9.774191E-05 | P80314, Q8BK64, Q9CQN1, P80313, P07901, P38647, P35564, Q8VCN9, O08997, Q91YE6, P42932, Q9CYG7, P62075, P09103, P48428, P63017 | 0.0015584 |
| GOTERM_MF_DIRECT | GO:0051082~unfolded protein binding | 8 | 0.0002238336 | P80314, P80313, P42932, Q9CQN1, P07901, P38647, P35564, P63017 | 0.00603551 |

| | | | | | |
|---|---|---|---|---|---|
| GOTERM_BP_DIRECT | GO:0051086~chaperone mediated protein folding independent of cc | 3 | 0.0035275421 | P80314, P80313, P42932 | 0.10524578 |
| GOTERM_CC_DIRECT | GO:0044297~cell body | 7 | 0.0061685006 | P80314, P80313, P42932, P09581, P20152, P08752, P60710 | 0.02987867 |
| GOTERM_CC_DIRECT | GO:0005832~chaperonin-containing T-complex | 3 | 0.0072909753 | P80314, P80313, P42932 | 0.03424549 |
| GOTERM_BP_DIRECT | GO:1904851~positive regulation of establishment of protein localiza | 3 | 0.0073624957 | P80314, P80313, P42932 | 0.17021731 |
| INTERPRO | IPR002194:Chaperonin_TCP-1_CS | 3 | 0.0129330966 | P80314, P80313, P42932 | 0.49215676 |
| GOTERM_BP_DIRECT | GO:0061077~chaperone-mediated protein folding | 4 | 0.0138659495 | P80314, P80313, P42932, P63017 | 0.24555001 |
| GOTERM_BP_DIRECT | GO:0032212~positive regulation of telomere maintenance via telom | 4 | 0.0138659495 | P80314, P80313, P42932, P63085 | 0.24555001 |
| INTERPRO | IPR017998:Chaperone_TCP-1 | 3 | 0.015622718 | P80314, P80313, P42932 | 0.536507 |
| INTERPRO | IPR027413:GROEL-like_equatorial_sf | 3 | 0.0249570596 | P80314, P80313, P42932 | 0.77604599 |
| GOTERM_BP_DIRECT | GO:0007339~binding of sperm to zona pellucida | 4 | 0.0272308538 | P80314, P80313, P42932, P05064 | 0.35460289 |
| INTERPRO | IPR027409:GroEL-like_apical_dom_sf | 3 | 0.0284627091 | P80314, P80313, P42932 | 0.77604599 |
| INTERPRO | IPR027410:TCP-1-like_intermed_sf | 3 | 0.0284627091 | P80314, P80313, P42932 | 0.77604599 |
| INTERPRO | IPR002423:Cpn60/GroEL/TCP-1 | 3 | 0.0284627091 | P80314, P80313, P42932 | 0.77604599 |
| GOTERM_BP_DIRECT | GO:0042026~protein refolding | 3 | 0.0312184753 | P07901, P38647, P63017 | 0.38524579 |
| Annotation Cluster 21 | Enrichment Score: 2.391426578792554 | | | | |
| Category | Term | Count | P Value | Genes | FDR |
| UP_SEQ_FEATURE | REPEAT:1-2 | 5 | 0.0007057396 | P31996, Q9CR57, P61979, P35564, Q02053 | 0.07355438 |
| UP_SEQ_FEATURE | REPEAT:2-1 | 4 | 0.0069886198 | P31996, Q9CR57, P61979, P35564 | 0.36424686 |
| UP_SEQ_FEATURE | REPEAT:2-2 | 4 | 0.0069886198 | P31996, Q9CR57, P61979, P35564 | 0.36424686 |
| UP_SEQ_FEATURE | REPEAT:1-1 | 4 | 0.0078861741 | P31996, P61979, P35564, Q02053 | 0.39521865 |
| Annotation Cluster 22 | Enrichment Score: 2.375146495678034 | | | | |
| Category | Term | Count | P Value | Genes | FDR |
| GOTERM_BP_DIRECT | GO:0042590~antigen processing and presentation of exogenous pe | 5 | 1.388077E-05 | P26151, P20491, P08508, A1L314, Q9ESY9 | 0.00172195 |
| GOTERM_BP_DIRECT | GO:0001805~positive regulation of type III hypersensitivity | 3 | 0.001034862 | P26151, P20491, P08508 | 0.04690711 |
| GOTERM_BP_DIRECT | GO:0050766~positive regulation of phagocytosis | 6 | 0.0027071157 | P06800, P26151, P20491, P08508, P97797, P63017 | 0.08986862 |
| GOTERM_MF_DIRECT | GO:0019864~IgG binding | 3 | 0.0046343038 | P26151, P20491, P08508 | 0.06479443 |
| GOTERM_BP_DIRECT | GO:0001798~positive regulation of type IIa hypersensitivity | 3 | 0.0046622253 | P26151, P20491, P08508 | 0.12928077 |
| GOTERM_BP_DIRECT | GO:0038094~Fc-gamma receptor signaling pathway | 3 | 0.0059418854 | P26151, P20491, P08508 | 0.14898961 |
| GOTERM_BP_DIRECT | GO:0007166~cell surface receptor signaling pathway | 7 | 0.1404069803 | P27870, P26151, P63085, P20491, P08508, P24063, Q8BZM1 | 0.86182097 |
| GOTERM_BP_DIRECT | GO:0050776~regulation of immune response | 3 | 0.1422768997 | P26151, P20491, P08508 | 0.86652882 |
| Annotation Cluster 23 | Enrichment Score: 2.268407338677204 | | | | |
| Category | Term | Count | P Value | Genes | FDR |
| UP_KW_DOMAIN | KW-0676~Redox-active center | 8 | 2.031767E-05 | P10639, P47791, Q9JMH6, P09103, Q8BXZ1, Q9ESY9, Q8CDN6, Q99J99 | 0.00021334 |
| UP_SEQ_FEATURE | DISULFID:Redox-active | 7 | 0.0002645531 | P10639, P47791, Q9JMH6, P09103, Q8BXZ1, Q9ESY9, Q8CDN6 | 0.03830142 |
| GOTERM_MF_DIRECT | GO:0015035~protein-disulfide reductase activity | 4 | 0.0029327991 | P10639, P09103, Q8BXZ1, Q8CDN6 | 0.04920585 |
| INTERPRO | IPR017937:Thioredoxin_CS | 4 | 0.0037413687 | P10639, P09103, Q8BXZ1, Q8CDN6 | 0.2394476 |
| INTERPRO | IPR013766:Thioredoxin_domain | 4 | 0.0385133398 | P10639, P09103, Q8BXZ1, Q8CDN6 | 0.82161792 |
| UP_SEQ_FEATURE | DOMAIN:Thioredoxin | 4 | 0.0448584319 | P10639, P09103, Q8BXZ1, Q8CDN6 | 0.88561419 |
| GOTERM_BP_DIRECT | GO:0045454~cell redox homeostasis | 3 | 0.0745834959 | P10639, P47791, Q9JMH6 | 0.634656 |
| INTERPRO | IPR036249:Thioredoxin-like_sf | 6 | 0.0937440377 | Q8VCH8, P10639, P09103, Q8BXZ1, P48774, Q8CDN6 | 0.99085151 |
| Annotation Cluster 24 | Enrichment Score: 2.239113065777496 | | | | |
| Category | Term | Count | P Value | Genes | FDR |
| GOTERM_BP_DIRECT | GO:0048870~cell motility | 8 | 1.46437E-06 | Q8BH43, P97797, P18760, Q8BPU7, Q9QXS1, P60710 | 0.00028763 |
| GOTERM_CC_DIRECT | GO:0016363~nuclear matrix | 11 | 1.582253E-05 | Q9CSN1, O35129, P60122, P20152, P18760, Q9Z0H3, P28867, Q8K310, P60710 | 0.00019885 |
| GOTERM_CC_DIRECT | GO:0071564~npBAF complex | 5 | 4.798376E-05 | Q62280, Q9Z0H3, P60710 | 0.0005071 |
| GOTERM_CC_DIRECT | GO:0030863~cortical cytoskeleton | 6 | 9.152825E-05 | P21107, Q7TPR4, Q8BTM8, P60710 | 0.00086858 |
| GOTERM_MF_DIRECT | GO:0098973~structural constituent of postsynaptic actin cytoskeletc | 4 | 0.0001817414 | Q62418, P60710 | 0.00527749 |
| GOTERM_MF_DIRECT | GO:0030957~Tat protein binding | 4 | 0.0003500278 | Q9Z0H3, P60710 | 0.00880903 |
| GOTERM_CC_DIRECT | GO:0140092~bBAF complex | 4 | 0.0004595279 | Q9Z0H3, P60710 | 0.0038851 |
| GOTERM_BP_DIRECT | GO:0072749~cellular response to cytochalasin B | 3 | 0.0005220198 | P60710 | 0.03237896 |
| GOTERM_CC_DIRECT | GO:0001725~stress fiber | 8 | 0.0005261638 | O55222, O55131, P21107, Q7TPR4, A2AQP0, P60710 | 0.00436904 |
| GOTERM_CC_DIRECT | GO:0016514~SWI/SNF complex | 5 | 0.0006243946 | Q62280, Q9Z0H3, P60710 | 0.00509375 |
| GOTERM_BP_DIRECT | GO:2000819~regulation of nucleotide-excision repair | 5 | 0.0006362806 | P63158, Q9Z0H3, P60710 | 0.03657837 |
| GOTERM_BP_DIRECT | GO:1903076~regulation of protein localization to plasma membrane | 5 | 0.000722954 | Q6PHZ2, O70404, P60710 | 0.03847493 |
| GOTERM_BP_DIRECT | GO:0051621~regulation of norepinephrine uptake | 3 | 0.001034862 | P60710 | 0.04690711 |
| GOTERM_CC_DIRECT | GO:0140288~GBAF complex | 4 | 0.0011247109 | Q62280, P60710 | 0.00769104 |
| GOTERM_CC_DIRECT | GO:0035060~brahma complex | 4 | 0.0011247109 | Q9Z0H3, P60710 | 0.00769104 |
| GOTERM_BP_DIRECT | GO:0045176~apical protein localization | 4 | 0.0013721363 | Q91V41, P60710 | 0.0576576 |
| GOTERM_BP_DIRECT | GO:2000045~regulation of G1/S transition of mitotic cell cycle | 6 | 0.0013943501 | P97372, Q9Z0H3, P97371, P60710 | 0.0576576 |
| GOTERM_CC_DIRECT | GO:0000776~kinetochore | 9 | 0.0015707005 | P46061, O55131, Q9D8B3, Q8BRT1, Q9Z0H3, Q8R1Q8, P60710 | 0.01023698 |
| GOTERM_CC_DIRECT | GO:0016586~RSC-type complex | 4 | 0.0016070957 | Q9Z0H3, P60710 | 0.01023698 |
| GOTERM_CC_DIRECT | GO:0071565~nBAF complex | 4 | 0.0016070957 | Q9Z0H3, P60710 | 0.01023698 |
| GOTERM_BP_DIRECT | GO:0071257~cellular response to electrical stimulus | 4 | 0.0016304512 | P28650, P60710 | 0.05912267 |
| GOTERM_BP_DIRECT | GO:0150111~regulation of transepithelial transport | 3 | 0.0017096252 | P60710 | 0.06105434 |
| GOTERM_BP_DIRECT | GO:0051726~regulation of cell cycle | 11 | 0.0018199366 | Q921F2, P06800, O35864, P60122, P09055, P08556, P63017, P60710 | 0.06402374 |
| GOTERM_BP_DIRECT | GO:1904030~negative regulation of cyclin-dependent protein kinase | 3 | 0.0025419445 | P60710 | 0.0855909 |
| GOTERM_BP_DIRECT | GO:0007163~establishment or maintenance of cell polarity | 5 | 0.0032075428 | Q8BRT1, P60766, P60710 | 0.10216457 |
| GOTERM_BP_DIRECT | GO:0045582~positive regulation of T cell differentiation | 5 | 0.0032075428 | P06800, Q9Z0H3, P60710 | 0.10216457 |
| GOTERM_BP_DIRECT | GO:0071896~protein localization to adherens junction | 3 | 0.0035275421 | P60710 | 0.10524578 |
| GOTERM_MF_DIRECT | GO:0050998~nitric-oxide synthase binding | 4 | 0.0042537308 | Q6PHZ2, P60710 | 0.0617609 |
| GOTERM_CC_DIRECT | GO:0098685~Schaffer collateral - CA1 synapse | 8 | 0.0043286838 | P09055, Q8R5M8, P62962, Q8BL66, P60766, P60710 | 0.02340509 |
| GOTERM_BP_DIRECT | GO:1902459~positive regulation of stem cell population maintenanc | 5 | 0.0043429859 | Q62280, Q9Z0H3, P60710 | 0.12483436 |
| GOTERM_BP_DIRECT | GO:0001738~morphogenesis of a polarized epithelium | 3 | 0.0046622253 | P60710 | 0.12928077 |
| GOTERM_BP_DIRECT | GO:0070316~regulation of G0 to G1 transition | 4 | 0.0053510221 | Q9Z0H3, P60710 | 0.1417119 |
| GOTERM_CC_DIRECT | GO:0097433~dense body | 3 | 0.0058839055 | P60710 | 0.02904689 |
| GOTERM_BP_DIRECT | GO:0045663~positive regulation of myoblast differentiation | 5 | 0.0061075902 | O55222, Q9Z0H3, P60710 | 0.15153253 |
| GOTERM_BP_DIRECT | GO:2000779~regulation of double-strand break repair | 4 | 0.007908442 | P60122, P60710 | 0.17752569 |
| GOTERM_CC_DIRECT | GO:0035267~NuA4 histone acetyltransferase complex | 4 | 0.008524011 | P60122, P60710 | 0.03924421 |
| GOTERM_BP_DIRECT | GO:0030071~regulation of mitotic metaphase/anaphase transition | 4 | 0.0119657172 | Q9Z0H3, P60710 | 0.23820967 |
| GOTERM_BP_DIRECT | GO:0034333~adherens junction assembly | 3 | 0.0143767085 | P60710 | 0.24555001 |
| INTERPRO | IPR004001:Actin_CS | 3 | 0.0185288926 | P60710 | 0.62115906 |
| GOTERM_BP_DIRECT | GO:2000781~positive regulation of double-strand break repair | 4 | 0.0205410112 | Q9Z0H3, P60710 | 0.29862181 |
| INTERPRO | IPR020902:Actin/actin-like_CS | 3 | 0.0216431016 | P60710 | 0.70868574 |
| GOTERM_BP_DIRECT | GO:1905168~positive regulation of double-strand break repair via | 4 | 0.0217968576 | P60122, P60710 | 0.30580472 |
| INTERPRO | IPR043129:ATPase_NBD | 5 | 0.0258431108 | P38647, P63017, P60710 | 0.77604599 |
| GOTERM_CC_DIRECT | GO:0098871~postsynaptic actin cytoskeleton | 3 | 0.0282265578 | P60710 | 0.09902917 |
| GOTERM_MF_DIRECT | GO:0019894~kinesin binding | 4 | 0.0330598612 | P70168, P60710 | 0.2683892 |
| GOTERM_CC_DIRECT | GO:0000786~nucleosome | 6 | 0.0480805009 | Q9JIA7, Q3TEA8, P60122, P60710 | 0.14708837 |
| GOTERM_BP_DIRECT | GO:1900242~regulation of synaptic vesicle endocytosis | 3 | 0.0495470015 | P60710 | 0.52398509 |
| UP_KW_PTM | KW-0558~Oxidation | 4 | 0.05540648 | P63158, P60710 | 0.13543806 |
| GOTERM_CC_DIRECT | GO:0043296~apical junction complex | 3 | 0.0593132526 | P60710 | 0.170251 |
| GOTERM_BP_DIRECT | GO:0045597~positive regulation of cell differentiation | 4 | 0.0596166866 | Q9Z0H3, P60710 | 0.56889284 |
| GOTERM_BP_DIRECT | GO:0032091~negative regulation of protein binding | 4 | 0.066212917 | Q9JIW9, P60710 | 0.58892017 |
| GOTERM_BP_DIRECT | GO:0042981~regulation of apoptotic process | 7 | 0.0790053485 | Q921F2, P07901, P60122, Q9CZW5, P60710 | 0.63991617 |
| GOTERM_BP_DIRECT | GO:0045596~negative regulation of cell differentiation | 5 | 0.0815472805 | P63085, Q62280, P60710 | 0.65824295 |
| INTERPRO | IPR004000:Actin | 3 | 0.0828373997 | P60710 | 0.99085151 |
| SMART | SM00268:ACTIN | 3 | 0.0851156197 | P60710 | 0.80189084 |
| GOTERM_CC_DIRECT | GO:0044305~calyx of Held | 3 | 0.0855707566 | P60710 | 0.22480453 |
| GOTERM_CC_DIRECT | GO:0070160~tight junction | 3 | 0.0895646522 | P60710 | 0.22622239 |
| GOTERM_CC_DIRECT | GO:0000785~chromatin | 12 | 0.0900024548 | Q9CSN1, Q921F2, O55222, Q8BFQ4, O35864, P61979, Q62280, P62827, Q9Z0H3, P60710 | 0.22622239 |
| GOTERM_BP_DIRECT | GO:0007409~axonogenesis | 4 | 0.2275063503 | P54227, P60710 | 0.97195876 |
| KEGG_PATHWAY | mmu04971:Gastric acid secretion | 4 | 0.3527266913 | Q6PHZ2, P08752, P60710 | 0.7471419 |
| KEGG_PATHWAY | mmu03082:ATP-dependent chromatin remodeling | 5 | 0.5342536509 | P60122, Q62280, Q9Z0H3, P60710 | 0.86825994 |

| | | | | | |
|---|---|---|---|---|---|
| GOTERM_BP_DIRECT | GO:0045893~positive regulation of DNA-templated transcription | 8 | 0.8971271946 | O55222, Q7TPR4, P60122, Q9CXT6, P63085, P60710 | 0.97195876 |

Annotation Cluster 25 Enrichment Score: 2.2121127878273623

| Category | Term | Count | P Value | Genes | FDR |
|---|---|---|---|---|---|
| GOTERM_CC_DIRECT | GO:0015935~small ribosomal subunit | 4 | 0.0033217469 | P14131, P62849, P62702, P14206 | 0.01860979 |
| GOTERM_CC_DIRECT | GO:0032040~small-subunit processome | 6 | 0.0062539284 | P14131, Q6DFW4, P62849, P62702, Q9D6Z1, P62858 | 0.02998017 |
| GOTERM_BP_DIRECT | GO:0042274~ribosomal small subunit biogenesis | 6 | 0.0066935011 | P14131, Q6DFW4, P62849, P62702, Q9D6Z1, P62858 | 0.15776582 |
| GOTERM_CC_DIRECT | GO:0022627~cytosolic small ribosomal subunit | 5 | 0.0101947379 | P14131, P62849, P62702, P14206, P62858 | 0.04514813 |

Annotation Cluster 26 Enrichment Score: 2.131297877074714

| Category | Term | Count | P Value | Genes | FDR |
|---|---|---|---|---|---|
| GOTERM_CC_DIRECT | GO:0005643~nuclear pore | 6 | 0.0024917046 | P46061, P62827, P34022, Q9D8B3, Q9QY81, P70168 | 0.01485439 |
| BIOCARTA | m_ranMSPathway:Role of Ran in mitotic spindle regulation | 4 | 0.0114641682 | P46061, P62827, P34022, P70168 | 0.43775475 |
| BIOCARTA | m_ranPathway:Cycling of Ran in nucleocytoplasmic transport | 3 | 0.0141340709 | P46061, P62827, P34022 | 0.43775475 |

Annotation Cluster 27 Enrichment Score: 2.0637951654512547

| Category | Term | Count | P Value | Genes | FDR |
|---|---|---|---|---|---|
| UP_KW_CELLULAR_COMPONENT | KW-0647~Proteasome | 8 | 7.788567E-05 | P97372, O70435, O35226, P46471, Q60692, Q8CDN6, P14685, P97371 | 0.00058766 |
| GOTERM_CC_DIRECT | GO:0000502~proteasome complex | 6 | 0.0007904178 | O70435, O35226, P46471, Q60692, Q8CDN6, P14685 | 0.00612574 |
| KEGG_PATHWAY | mmu03050:Proteasome | 7 | 0.0018986507 | P97372, O70435, O35226, P46471, Q60692, P14685, P97371 | 0.02439766 |
| GOTERM_CC_DIRECT | GO:0022624~proteasome accessory complex | 3 | 0.0207347772 | O35226, P46471, P14685 | 0.07838757 |
| KEGG_PATHWAY | mmu05017:Spinocerebellar ataxia | 7 | 0.2052871746 | O70435, O35226, O55143, P46471, Q9D6Z1, Q60692, P14685 | 0.60754938 |
| GOTERM_BP_DIRECT | GO:0043161~proteasome-mediated ubiquitin-dependent protein cat | 4 | 0.8325463446 | O70435, O35226, P46471, Q60692 | 0.97195876 |

Annotation Cluster 28 Enrichment Score: 1.94261152112871

| Category | Term | Count | P Value | Genes | FDR |
|---|---|---|---|---|---|
| INTERPRO | IPR009000:Transl_B-barrel_sf | 6 | 0.0002358443 | Q8BFR5, Q9Z0N1, P10126, P27659, Q8K0D5, Q8BGQ7 | 0.08301721 |
| INTERPRO | IPR004161:EFTu-like_2 | 4 | 0.0017593532 | Q8BFR5, Q9Z0N1, P10126, Q8K0D5 | 0.15482308 |
| UP_SEQ_FEATURE | DOMAIN:tr-type G | 4 | 0.0040235953 | Q8BFR5, Q9Z0N1, P10126, Q8K0D5 | 0.25217895 |
| GOTERM_MF_DIRECT | GO:0003746~translation elongation factor activity | 4 | 0.0042537308 | Q8BFR5, P10126, P57776, Q8K0D5 | 0.0617609 |
| INTERPRO | IPR000795:T_Tr_GTP-bd_dom | 4 | 0.0050912089 | Q8BFR5, Q9Z0N1, P10126, Q8K0D5 | 0.29868425 |
| INTERPRO | IPR031157:G_TR_CS | 3 | 0.0104687858 | Q8BFR5, P10126, Q8K0D5 | 0.45796501 |
| UP_KW_MOLECULAR_FUNCTION | KW-0251~Elongation factor | 4 | 0.0125752354 | Q8BFR5, P10126, P57776, Q8K0D5 | 0.07316501 |
| INTERPRO | IPR009001:Transl_elong_EF1A/Init_IF2_C | 3 | 0.0129330966 | Q8BFR5, Q9Z0N1, P10126 | 0.49215676 |
| UP_SEQ_FEATURE | REGION:G1 | 3 | 0.037876874 | Q8BFR5, Q9Z0N1, P10126 | 0.82255945 |
| UP_SEQ_FEATURE | REGION:G2 | 3 | 0.037876874 | Q8BFR5, Q9Z0N1, P10126 | 0.82255945 |
| UP_SEQ_FEATURE | REGION:G4 | 3 | 0.037876874 | Q8BFR5, Q9Z0N1, P10126 | 0.82255945 |
| UP_SEQ_FEATURE | REGION:G5 | 3 | 0.037876874 | Q8BFR5, Q9Z0N1, P10126 | 0.82255945 |
| UP_SEQ_FEATURE | REGION:G3 | 3 | 0.0418596127 | Q8BFR5, Q9Z0N1, P10126 | 0.87972702 |
| GOTERM_BP_DIRECT | GO:0006414~translational elongation | 3 | 0.1199080465 | Q8BFR5, P10126, P57776 | 0.78506463 |

Annotation Cluster 29 Enrichment Score: 1.9360558208221548

| Category | Term | Count | P Value | Genes | FDR |
|---|---|---|---|---|---|
| INTERPRO | IPR029006:ADF-H/Gelsolin-like_dom_sf | 4 | 0.008577118 | P24452, Q62418, P18760, Q91YR1 | 0.41643387 |
| UP_SEQ_FEATURE | DOMAIN:ADF-H | 3 | 0.012196291 | Q62418, P18760, Q91YR1 | 0.54799197 |
| INTERPRO | IPR002108:ADF-H | 3 | 0.0129330966 | Q62418, P18760, Q91YR1 | 0.49215676 |
| SMART | SM00102:ADF | 3 | 0.0133200248 | Q62418, P18760, Q91YR1 | 0.33680634 |

Annotation Cluster 30 Enrichment Score: 1.9124528205314761

| Category | Term | Count | P Value | Genes | FDR |
|---|---|---|---|---|---|
| GOTERM_MF_DIRECT | GO:0005178~integrin binding | 13 | 1.406419E-06 | P05555, Q07076, Q7TPR4, Q64339, P24063, P11835, P17742, O55222, P63158, P09055, Q61072, P09103, P25911 | 5.8991E-05 |
| GOTERM_BP_DIRECT | GO:0007229~integrin-mediated signaling pathway | 11 | 2.001198E-06 | P05555, O55222, P27870, Q64339, P20491, P09055, P24063, P11835, Q9R0C8, P8BK67, P60766 | 0.00033692 |
| GOTERM_BP_DIRECT | GO:0006909~phagocytosis | 8 | 8.064163E-05 | P05555, P27870, P09055, O70145, P24063, P11835, Q8BPU7, Q8BG07 | 0.00655422 |
| GOTERM_BP_DIRECT | GO:0007160~cell-matrix adhesion | 8 | 0.0003162727 | P05555, O55222, P09055, Q8VIM6, P24063, P97797, P11835, Q61072 | 0.02130733 |
| GOTERM_BP_DIRECT | GO:0007159~leukocyte cell-cell adhesion | 5 | 0.0006362806 | P05555, P06800, P09055, P24063, P11835 | 0.03657837 |
| GOTERM_BP_DIRECT | GO:0033627~cell adhesion mediated by integrin | 5 | 0.0019118375 | P05555, P09055, P24063, P11835, Q61072 | 0.06626766 |
| GOTERM_CC_DIRECT | GO:0008305~integrin complex | 4 | 0.0064649728 | P05555, P09055, P24063, P11835 | 0.03067564 |
| GOTERM_BP_DIRECT | GO:0034113~heterotypic cell-cell adhesion | 4 | 0.007908442 | P05555, P06800, P09055, P11835 | 0.17752569 |
| INTERPRO | IPR032695:Integrin_dom_sf | 4 | 0.0119185614 | P05555, P09055, P24063, P11835 | 0.49215676 |
| GOTERM_BP_DIRECT | GO:0050798~activated T cell proliferation | 3 | 0.0143767085 | P05555, P24063, P11835 | 0.24555001 |
| BIOCARTA | m_monocytePathway:Monocyte and its Surface Molecules | 4 | 0.0151998178 | P05555, P09055, P24063, P11835 | 0.43775475 |
| GOTERM_MF_DIRECT | GO:0050839~cell adhesion molecule binding | 5 | 0.0183296811 | P26041, P09055, Q8R5M8, P24063, P11835 | 0.18701229 |
| GOTERM_BP_DIRECT | GO:0098609~cell-cell adhesion | 8 | 0.0205568083 | P05555, P09055, P24063, P11835, P14206, O35691, P62137, P60766 | 0.29862181 |
| KEGG_PATHWAY | mmu05133:Pertussis | 7 | 0.020904456 | P05555, P10810, P63085, P09055, P11835, P18760, P08752 | 0.12494059 |
| BIOCARTA | m_neutrophilPathway:Neutrophil and Its Surface Molecules | 3 | 0.0578445442 | P05555, P24063, P11835 | 1 |
| UP_SEQ_FEATURE | DOMAIN:VWFA | 5 | 0.0704272188 | P05555, O35226, P09055, P24063, P11835 | 0.99162861 |
| BIOCARTA | m_lymphocytePathway:Adhesion Molecules on Lymphocyte | 3 | 0.0719935504 | P09055, P24063, P11835 | 1 |
| UP_KW_MOLECULAR_FUNCTION | KW-0401~Integrin | 4 | 0.0971875336 | P05555, P09055, P24063, P11835 | 0.36588248 |
| INTERPRO | IPR036465:vWFA_dom_sf | 5 | 0.1177918927 | P05555, O35226, P09055, P24063, P11835 | 0.99085151 |
| KEGG_PATHWAY | mmu04514:Cell adhesion molecules | 9 | 0.1314415077 | Q61543, P05555, P06800, P09055, Q8R5M8, P01900, P24063, P11835, P01897 | 0.45040623 |
| SMART | SM00327:VWA | 3 | 0.3883335149 | P05555, O35226, P24063 | 0.98882682 |
| INTERPRO | IPR002035:VWF_A | 3 | 0.4239977846 | P05555, O35226, P24063 | 0.99085151 |
| KEGG_PATHWAY | mmu05150:Staphylococcus aureus infection | 5 | 0.4865170998 | P05555, P26151, P08508, P24063, P11835 | 0.85252108 |
| GOTERM_BP_DIRECT | GO:0007155~cell adhesion | 8 | 0.4913054202 | P05555, P09055, Q8R5M8, P24063, P11835, Q61072, P14206, Q07797 | 0.97195876 |
| UP_KW_BIOLOGICAL_PROCESS | KW-0130~Cell adhesion | 6 | 0.9706261819 | P05555, P09055, Q8R5M8, P24063, P11835, Q07797 | 0.97062618 |

Annotation Cluster 31 Enrichment Score: 1.8763934421528101

| Category | Term | Count | P Value | Genes | FDR |
|---|---|---|---|---|---|
| COG_ONTOLOGY | Lipid metabolism | 9 | 0.0005693856 | Q9DBL1, P42125, Q61425, O70310, Q9QUJ7, Q9JHI5, P41216, Q8JZN5, P50544 | 0.0079714 |
| GOTERM_BP_DIRECT | GO:0006635~fatty acid beta-oxidation | 6 | 0.0011142519 | Q9DBL1, P42125, Q61425, Q9JHI5, Q8JZN5, P50544 | 0.04955268 |
| KEGG_PATHWAY | mmu00071:Fatty acid degradation | 6 | 0.0149455642 | Q9DBL1, P42125, Q61425, Q9QUJ7, P41216, P50544 | 0.10669472 |
| KEGG_PATHWAY | mmu01212:Fatty acid metabolism | 6 | 0.0297314159 | Q9DBL1, Q61425, Q9QUJ7, O70503, P41216, P50544 | 0.15918696 |
| UP_KW_BIOLOGICAL_PROCESS | KW-0276~Fatty acid metabolism | 7 | 0.1049248674 | Q9DBL1, P42125, Q61425, P22437, Q9QUJ7, P41216, P50544 | 0.39688972 |
| GOTERM_BP_DIRECT | GO:0006631~fatty acid metabolic process | 4 | 0.1864846525 | Q9DBL1, Q9QUJ7, P41216, P31786 | 0.97195876 |

Annotation Cluster 32 Enrichment Score: 1.859950694893972

| Category | Term | Count | P Value | Genes | FDR |
|---|---|---|---|---|---|
| INTERPRO | IPR023210:NADP_OxRdtase_dom | 4 | 0.0058622382 | Q8CG76, P45376, O09172, P45377 | 0.30570487 |
| INTERPRO | IPR036812:NADP_OxRdtase_dom_sf | 4 | 0.0058622382 | Q8CG76, P45376, O09172, P45377 | 0.30570487 |
| GOTERM_MF_DIRECT | GO:0004032~aldose reductase (NADPH) activity | 3 | 0.0310425738 | Q8CG76, P45376, P45377 | 0.25755102 |
| UP_SEQ_FEATURE | SITE:Lowers pKa of active site Tyr | 3 | 0.034049644 | Q8CG76, P45376, P45377 | 0.82255945 |

Annotation Cluster 33 Enrichment Score: 1.8062434061726127

| Category | Term | Count | P Value | Genes | FDR |
|---|---|---|---|---|---|
| INTERPRO | IPR000594:ThiF_NAD_FAD-bd | 3 | 0.015622718 | Q9R1T2, Q9D906, Q02053 | 0.536507 |
| INTERPRO | IPR035985:Ubiquitin-activating_enz | 3 | 0.015622718 | Q9R1T2, Q9D906, Q02053 | 0.536507 |
| INTERPRO | IPR045886:ThiF/MoeB/HesA | 3 | 0.015622718 | Q9R1T2, Q9D906, Q02053 | 0.536507 |

Annotation Cluster 34 Enrichment Score: 1.7882324386304533

| Category | Term | Count | P Value | Genes | FDR |
|---|---|---|---|---|---|
| UP_KW_MOLECULAR_FUNCTION | KW-0436~Ligase | 12 | 0.0003137393 | P32921, Q8BML9, Q9R1T2, P28650, Q9QUJ7, P26638, Q9WUM5, Q9ER72, P41216, Q8BGQ7, Q02053, Q9DCL9 | 0.00334655 |
| UP_KW_MOLECULAR_FUNCTION | KW-0030~Aminoacyl-tRNA synthetase | 5 | 0.0110361324 | P32921, Q8BML9, P26638, Q9ER72, Q8BGQ7 | 0.07063125 |

| | | | | | |
|---|---|---|---|---|---|
| KEGG_PATHWAY | mmu00970:Aminoacyl-tRNA biosynthesis | 5 | 0.1141567629 | P32921, Q8BML9, P26638, Q9ER72, Q8BGQ7 | 0.42190789 |
| INTERPRO | IPR014729:Rossmann-like_a/b/a_fold | 3 | 0.1779027959 | P32921, Q8BML9, Q9ER72 | 0.99085151 |
| Annotation Cluster 35 | Enrichment Score: 1.7296673425020153 | | | | |
| Category | Term | Count | P Value | Genes | FDR |
| GOTERM_MF_DIRECT | GO:0003689~DNA clamp loader activity | 9 | 0.0009370101 | P49718, P60122, P46460, P97855, P23249, Q921N6, Q61881, E9Q555, Q9D0F6 | 0.01912007 |
| GOTERM_MF_DIRECT | GO:0140584~chromatin extrusion motor activity | 8 | 0.0028996328 | P49718, P60122, P46460, P97855, P23249, Q921N6, Q61881, E9Q555 | 0.04920585 |
| GOTERM_MF_DIRECT | GO:0140665~ATP-dependent H3-H4 histone complex chaperone a | 8 | 0.0028996328 | P49718, P60122, P46460, P97855, P23249, Q921N6, Q61881, E9Q555 | 0.04920585 |
| GOTERM_MF_DIRECT | GO:0140849~ATP-dependent H2AZ histone chaperone activity | 8 | 0.0031339974 | P49718, P60122, P46460, P97855, P23249, Q921N6, Q61881, E9Q555 | 0.05143844 |
| GOTERM_MF_DIRECT | GO:0061775~cohesin loader activity | 8 | 0.0032564727 | P49718, P60122, P46460, P97855, P23249, Q921N6, Q61881, E9Q555 | 0.05231142 |
| GOTERM_BP_DIRECT | GO:0140588~chromatin looping | 8 | 0.0034418013 | P49718, P60122, P46460, P97855, P23249, Q921N6, Q61881, E9Q555 | 0.10524578 |
| GOTERM_MF_DIRECT | GO:0061749~forked DNA-dependent helicase activity | 4 | 0.0203758138 | P49718, P60122, P97855, Q61881 | 0.20511653 |
| GOTERM_MF_DIRECT | GO:1990518~single-stranded 3'-5' DNA helicase activity | 4 | 0.0216222149 | P49718, P60122, P97855, Q61881 | 0.20929195 |
| GOTERM_MF_DIRECT | GO:0009378~four-way junction helicase activity | 4 | 0.0216222149 | P49718, P60122, P97855, Q61881 | 0.20929195 |
| GOTERM_MF_DIRECT | GO:0036121~double-stranded DNA helicase activity | 4 | 0.0229094449 | P49718, P60122, P97855, Q61881 | 0.21371116 |
| INTERPRO | IPR003593:AAA+_ATPase | 7 | 0.0467461835 | P60122, P46460, P46471, Q61881, Q8CGK3, E9Q555, Q9D0F6 | 0.95672708 |
| INTERPRO | IPR012340:NA-bd_OB-fold | 6 | 0.0468837252 | P49718, P62960, P60122, P46471, P62858, Q61881 | 0.95672708 |
| SMART | SM00382:AAA | 7 | 0.0493071572 | P60122, P46460, P46471, Q61881, Q8CGK3, E9Q555, Q9D0F6 | 0.67133591 |
| INTERPRO | IPR003959:ATPase_AAA_core | 4 | 0.0510178496 | P46460, P46471, Q8CGK3, Q9D0F6 | 0.95672708 |
| GOTERM_BP_DIRECT | GO:0032508~DNA duplex unwinding | 4 | 0.0639782918 | P49718, P60122, P97855, Q9D0F6 | 0.57892868 |
| GOTERM_MF_DIRECT | GO:0003678~DNA helicase activity | 3 | 0.0898666168 | P60122, P97855, Q61881 | 0.49525033 |
| UP_SEQ_FEATURE | DOMAIN:AAA+ ATPase | 3 | 0.1820109594 | P60122, P46471, Q9D0F6 | 0.99162861 |
| UP_KW_MOLECULAR_FUNCTION | KW-0347~Helicase | 6 | 0.2107981888 | P49718, P60122, P97855, P23249, Q921N6, Q61881 | 0.5621285 |
| Annotation Cluster 36 | Enrichment Score: 1.5982351084014745 | | | | |
| Category | Term | Count | P Value | Genes | FDR |
| GOTERM_BP_DIRECT | GO:0051301~cell division | 14 | 0.0015539826 | Q91ZR2, Q60875, Q8CG47, P62827, O88738, P60122, Q9JIW9, Q8BRT1, P27546, P62137, Q8BK67, Q8R1Q8, P08752, P60766 | 0.05912267 |
| GOTERM_CC_DIRECT | GO:0030496~midbody | 9 | 0.01075624 | O55131, O88738, P62827, Q9D8B3, Q9JIW9, O70404, Q8BK67, P08752, P60766 | 0.04588671 |
| UP_KW_BIOLOGICAL_PROCESS | KW-0132~Cell division | 14 | 0.0606572953 | Q91ZR2, Q60875, Q8CG47, P62827, O55131, O88738, P60122, Q9JIW9, Q8BRT1, P62137, Q8BK67, Q8R1Q8, P08752, P60766 | 0.3284828 |
| UP_KW_BIOLOGICAL_PROCESS | KW-0131~Cell cycle | 20 | 0.0784093197 | Q91ZR2, Q60875, Q8CG47, Q9CXT6, P62827, Q9Z0H3, Q61881, P49718, O55131, O88738, P60122, Q9JIW9, Q8BRT1, P28867, P62137, Q8BK67, Q8R1Q8, P08752, P60766 | 0.37897838 |
| UP_KW_BIOLOGICAL_PROCESS | KW-0498~Mitosis | 10 | 0.1283710538 | O55131, Q91ZR2, Q60875, Q8CG47, O88738, P60122, P62827, Q8BRT1, Q8BK67, Q8R1Q8 | 0.44673127 |
| Annotation Cluster 37 | Enrichment Score: 1.5903499541627624 | | | | |
| Category | Term | Count | P Value | Genes | FDR |
| UP_KW_BIOLOGICAL_PROCESS | KW-0249~Electron transport | 10 | 0.0004102493 | P10639, Q921G7, Q9ERS2, Q9CQX2, Q9DC70, Q91WD5, Q8CDN6, Q8R1I1, Q8K2B3, Q91YT0 | 0.00509881 |
| GOTERM_MF_DIRECT | GO:0048038~quinone binding | 4 | 0.0011310078 | Q921G7, Q9R112, Q9DC70, Q91WD5 | 0.02189515 |
| GOTERM_BP_DIRECT | GO:0042776~proton motive force-driven mitochondrial ATP synthes | 6 | 0.0016078581 | Q9CQQ7, Q9ERS2, Q9DC70, Q91WD5, Q8K2B3, Q91YT0 | 0.05912267 |
| GOTERM_MF_DIRECT | GO:0051539~4 iron, 4 sulfur cluster binding | 5 | 0.0026938118 | Q921G7, P28271, Q9DC70, Q91WD5, Q91YT0 | 0.04920585 |
| GOTERM_BP_DIRECT | GO:0032981~mitochondrial respiratory chain complex I assembly | 5 | 0.0127574171 | Q9Z0X1, Q9ERS2, Q9DC70, Q91WD5, Q8JZN5 | 0.2386447 |
| UP_KW_LIGAND | KW-0830~Ubiquinone | 4 | 0.0153814898 | Q921G7, Q9DC70, Q91WD5, Q91YT0 | 0.06385646 |
| UP_KW_LIGAND | KW-0004~4Fe-4S | 5 | 0.016658207 | Q921G7, P28271, Q9DC70, Q91WD5, Q91YT0 | 0.06385646 |
| GOTERM_MF_DIRECT | GO:0008137~NADH dehydrogenase (ubiquinone) activity | 3 | 0.0232110132 | Q9DC70, Q91WD5, Q91YT0 | 0.21371116 |
| GOTERM_CC_DIRECT | GO:0045271~respiratory chain complex I | 4 | 0.0361455987 | Q9ERS2, Q9DC70, Q91WD5, Q91YT0 | 0.11836411 |
| KEGG_PATHWAY | mmu04932:Non-alcoholic fatty liver disease | 10 | 0.0389755438 | P48771, Q9ERS2, Q9DC70, Q91WD5, P56391, Q8R1I1, P60766, Q8K2B3, P17665, Q91YT0 | 0.2003343 |
| UP_KW_BIOLOGICAL_PROCESS | KW-0679~Respiratory chain | 5 | 0.0436169043 | Q9ERS2, Q9DC70, Q91WD5, Q8R1I1, Q91YT0 | 0.27104791 |
| GOTERM_BP_DIRECT | GO:0006120~mitochondrial electron transport, NADH to ubiquinone | 3 | 0.0495470015 | Q9DC70, Q91WD5, Q91YT0 | 0.52398509 |
| GOTERM_BP_DIRECT | GO:0009060~aerobic respiration | 4 | 0.0731274073 | Q9ERS2, Q9DC70, Q91WD5, Q91YT0 | 0.634656 |
| GOTERM_BP_DIRECT | GO:1902600~proton transmembrane transport | 5 | 0.0911757859 | Q9CQQ7, P50516, Q9DC70, Q91WD5, Q91YT0 | 0.68878631 |
| UP_KW_LIGAND | KW-0411~Iron-sulfur | 5 | 0.1022323113 | Q921G7, P28271, Q9DC70, Q91WD5, Q91YT0 | 0.21375847 |
| UP_KW_MOLECULAR_FUNCTION | KW-1278~Translocase | 5 | 0.1964157964 | O55143, P50516, Q9DC70, Q91WD5, Q91YT0 | 0.5465483 |
| KEGG_PATHWAY | mmu04723:Retrograde endocannabinoid signaling | 6 | 0.4304425606 | P63085, Q9ERS2, Q9DC70, Q91WD5, P08752, Q91YT0 | 0.82555028 |
| UP_KW_LIGAND | KW-0408~Iron | 8 | 0.883750222 | P22437, Q921G7, Q9CXT6, P28271, Q9CQX2, Q9DC70, Q91WD5, Q91YT0 | 0.88375022 |
| Annotation Cluster 38 | Enrichment Score: 1.551499068349253 | | | | |
| Category | Term | Count | P Value | Genes | FDR |
| GOTERM_BP_DIRECT | GO:0006606~protein import into nucleus | 8 | 0.0010244711 | O35129, Q91YE6, P62827, Q8BFY9, P34960, P18760, P70168, P63017 | 0.04690711 |
| GOTERM_MF_DIRECT | GO:0061608~nuclear import signal receptor activity | 3 | 0.0283297338 | Q91YE6, Q8BFY9, P70168 | 0.24305624 |
| SMART | SM00913:IBN_N | 3 | 0.0330866333 | Q91YE6, Q8BFY9, P70168 | 0.52891302 |
| UP_SEQ_FEATURE | DOMAIN:Importin N-terminal | 3 | 0.034049644 | Q91YE6, Q8BFY9, P70168 | 0.82255945 |
| INTERPRO | IPR001494:Importin-beta_N | 3 | 0.0360179573 | Q91YE6, Q8BFY9, P70168 | 0.78020437 |
| INTERPRO | IPR011989:ARM-like | 6 | 0.4168493881 | Q91YE6, Q8BVE3, Q8BFY9, Q8BRT1, P70168, Q8BPU7 | 0.99085151 |
| Annotation Cluster 39 | Enrichment Score: 1.5161608443579047 | | | | |
| Category | Term | Count | P Value | Genes | FDR |
| GOTERM_BP_DIRECT | GO:0007159~leukocyte cell-cell adhesion | 5 | 0.0006362806 | P05555, P06800, P09055, P24063, P11835 | 0.03657837 |
| BIOCARTA | m_blymphocytePathway:B Lymphocyte Cell Surface Molecules | 3 | 0.0719935504 | P06800, P24063, P11835 | 1 |
| BIOCARTA | m_tcytotoxicPathway:T Cytotoxic Cell Surface Molecules | 3 | 0.1371535907 | P06800, P24063, P11835 | 1 |
| BIOCARTA | m_thelperPathway:T Helper Cell Surface Molecules | 3 | 0.1371535907 | P06800, P24063, P11835 | 1 |
| Annotation Cluster 40 | Enrichment Score: 1.4968670615077464 | | | | |
| Category | Term | Count | P Value | Genes | FDR |
| UP_SEQ_FEATURE | REPEAT:2 | 11 | 0.0007338503 | P42567, P62320, Q8BL97, P70460, P16110, Q62280, Q3U0V1, P23116, P13864, Q9Z0H3, Q62433 | 0.07355438 |
| UP_SEQ_FEATURE | REPEAT:1 | 10 | 0.0025349974 | P42567, P62320, Q8BL97, P70460, P16110, Q62280, Q3U0V1, P13864, Q9Z0H3, Q62433 | 0.1943001 |
| UP_SEQ_FEATURE | REPEAT:4 | 7 | 0.014259312 | P42567, P62320, Q8BL97, P16110, Q3U0V1, P23116, P13864 | 0.61932945 |
| UP_SEQ_FEATURE | REPEAT:3 | 7 | 0.0308132611 | P42567, P62320, Q8BL97, P16110, Q3U0V1, P13864, Q62433 | 0.82255945 |
| UP_SEQ_FEATURE | REPEAT:5 | 5 | 0.0871989899 | P42567, P62320, P16110, P23116, P13864 | 0.99162861 |
| UP_SEQ_FEATURE | REPEAT:6 | 4 | 0.1438663937 | P42567, P16110, P23116, P13864 | 0.99162861 |
| UP_SEQ_FEATURE | REPEAT:8 | 3 | 0.302969169 | P42567, P16110, P23116 | 0.99162861 |
| UP_SEQ_FEATURE | REPEAT:7 | 3 | 0.3410146266 | P42567, P16110, P23116 | 0.99162861 |
| Annotation Cluster 41 | Enrichment Score: 1.4218504984114433 | | | | |
| Category | Term | Count | P Value | Genes | FDR |
| UP_SEQ_FEATURE | REGION:G1 motif | 4 | 0.0378572882 | Q8CHH9, O55131, Q9EQP2, P08752 | 0.82255945 |
| UP_SEQ_FEATURE | REGION:G3 motif | 4 | 0.0378572882 | Q8CHH9, O55131, Q9EQP2, P08752 | 0.82255945 |
| UP_SEQ_FEATURE | REGION:G4 motif | 4 | 0.0378572882 | Q8CHH9, O55131, Q9EQP2, P08752 | 0.82255945 |
| Annotation Cluster 42 | Enrichment Score: 1.3158780389433715 | | | | |
| Category | Term | Count | P Value | Genes | FDR |
| GOTERM_CC_DIRECT | GO:0030018~Z disc | 7 | 0.0162722518 | P26883, Q7TPR4, P48036, Q8BTM8, P05064, Q9QXS1, Q6P069 | 0.06305498 |
| SMART | SM00033:CH | 5 | 0.031382657 | P27870, Q7TPR4, Q8BTM8, Q9R0C8, Q9QXS1 | 0.52891302 |
| UP_SEQ_FEATURE | DOMAIN:Calponin-homology (CH) | 5 | 0.0442168064 | P27870, Q7TPR4, Q8BTM8, Q9R0C8, Q9QXS1 | 0.88561419 |
| INTERPRO | IPR001715:CH_dom | 5 | 0.0586183561 | P27870, Q7TPR4, Q8BTM8, Q9R0C8, Q9QXS1 | 0.95672708 |
| UP_SEQ_FEATURE | DOMAIN:Calponin-homology (CH) 1 | 3 | 0.0592187423 | Q7TPR4, Q8BTM8, Q9QXS1 | 0.95261755 |
| UP_SEQ_FEATURE | DOMAIN:Calponin-homology (CH) 2 | 3 | 0.0592187423 | Q7TPR4, Q8BTM8, Q9QXS1 | 0.95261755 |
| INTERPRO | IPR001589:Actinin_actin-bd_CS | 3 | 0.0625128112 | Q7TPR4, Q8BTM8, Q9QXS1 | 0.95672708 |

| | | | | | |
|---|---|---|---|---|---|
| INTERPRO | IPR036872:CH_dom_sf | 5 | 0.062945432 | P27870, Q7TPR4, Q8BTM8, Q9R0C8, Q9QXS1 | 0.95672708 |
| UP_SEQ_FEATURE | REGION:Actin-binding | 4 | 0.0786117874 | Q7TPR4, Q8BTM8, A2AQP0, Q9QXS1 | 0.99162861 |
| Annotation Cluster 43 | Enrichment Score: 1.315051272651153 | | | | |
| Category | Term | Count | P Value | Genes | FDR |
| UP_SEQ_FEATURE | DOMAIN:KH 3 | 3 | 0.020431745 | P61979, Q3U0V1, P60335 | 0.82255945 |
| INTERPRO | IPR004087:KH_dom | 4 | 0.0297987866 | P61979, Q3U0V1, P60335, Q64213 | 0.77604599 |
| SMART | SM00322:KH | 4 | 0.0309444996 | P61979, Q3U0V1, P60335, Q64213 | 0.52891302 |
| UP_SEQ_FEATURE | DOMAIN:KH 1 | 3 | 0.0546769925 | P61979, Q3U0V1, P60335 | 0.90182432 |
| UP_SEQ_FEATURE | DOMAIN:KH 2 | 3 | 0.0546769925 | P61979, Q3U0V1, P60335 | 0.90182432 |
| GOTERM_MF_DIRECT | GO:0003730~mRNA 3'-UTR binding | 5 | 0.05851715 | Q921F2, P61979, Q3U0V1, P29341, P60335 | 0.38754779 |
| INTERPRO | IPR036612:KH_dom_type_1_sf | 4 | 0.0652343204 | P61979, Q3U0V1, P60335, Q64213 | 0.95672708 |
| INTERPRO | IPR004088:KH_dom_type_1 | 3 | 0.1403245858 | P61979, Q3U0V1, P60335 | 0.99085151 |
| Annotation Cluster 44 | Enrichment Score: 1.3130333348466086 | | | | |
| Category | Term | Count | P Value | Genes | FDR |
| KEGG_PATHWAY | mmu05140:Leishmaniasis | 8 | 0.0033850647 | P05555, P26151, P63085, P09055, O70145, P08508, P11835, P10126 | 0.0362484 |
| KEGG_PATHWAY | mmu04613:Neutrophil extracellular trap formation | 11 | 0.0698610181 | P05555, P63158, P26151, P63085, O70145, P08508, P24063, P11835, Q9D906, P60710 | 0.30164987 |
| KEGG_PATHWAY | mmu05150:Staphylococcus aureus infection | 5 | 0.4865170998 | P05555, P26151, P08508, P24063, P11835 | 0.85252108 |
| Annotation Cluster 45 | Enrichment Score: 1.3106172520033494 | | | | |
| Category | Term | Count | P Value | Genes | FDR |
| SMART | SM00088:PINT | 3 | 0.0292939549 | P23116, P60229, P14685 | 0.52891302 |
| INTERPRO | IPR000717:PCI_dom | 3 | 0.0625128112 | P23116, P60229, P14685 | 0.95672708 |
| UP_SEQ_FEATURE | DOMAIN:PCI | 3 | 0.0638853394 | P23116, P60229, P14685 | 0.99162861 |
| Annotation Cluster 46 | Enrichment Score: 1.3039113459762344 | | | | |
| Category | Term | Count | P Value | Genes | FDR |
| GOTERM_CC_DIRECT | GO:1904949~ATPase complex | 4 | 0.0007455329 | Q8BVE3, Q9CR51, P58389, P50516 | 0.00587581 |
| GOTERM_CC_DIRECT | GO:0033180~proton-transporting V-type ATPase, V1 domain | 3 | 0.0058839055 | Q8BVE3, Q9CR51, P50516 | 0.02904689 |
| GOTERM_CC_DIRECT | GO:0000221~vacuolar proton-transporting V-type ATPase, V1 dom | 3 | 0.010508712 | Q8BVE3, Q9CR51, P50516 | 0.04588671 |
| GOTERM_CC_DIRECT | GO:0033176~proton-transporting V-type ATPase complex | 3 | 0.0123118323 | Q8BVE3, Q9CR51, P50516 | 0.05111609 |
| GOTERM_BP_DIRECT | GO:0097401~synaptic vesicle lumen acidification | 3 | 0.0284910324 | Q8BVE3, Q9CR51, P50516 | 0.36103959 |
| GOTERM_CC_DIRECT | GO:0098850~extrinsic component of synaptic vesicle membrane | 3 | 0.042692425 | Q8BVE3, Q9CR51, P50516 | 0.13147005 |
| GOTERM_MF_DIRECT | GO:0046961~proton-transporting ATPase activity, rotational mecha | 3 | 0.0428450652 | Q8BVE3, Q9CR51, P50516 | 0.31103869 |
| UP_KW_BIOLOGICAL_PROCESS | KW-0375~Hydrogen ion transport | 4 | 0.0937493271 | Q8BVE3, Q9CQQ7, Q9CR51, P50516 | 0.39688972 |
| GOTERM_CC_DIRECT | GO:1902495~transmembrane transporter complex | 3 | 0.1060594142 | Q8BVE3, Q9CR51, P50516 | 0.25956646 |
| KEGG_PATHWAY | mmu05323:Rheumatoid arthritis | 5 | 0.2282773041 | Q8BVE3, P24063, P11835, Q9CR51, P50516 | 0.61755018 |
| KEGG_PATHWAY | mmu04721:Synaptic vesicle cycle | 4 | 0.3681447753 | P46460, Q8BVE3, Q9CR51, P50516 | 0.75089847 |
| KEGG_PATHWAY | mmu04150:mTOR signaling pathway | 6 | 0.4580512413 | P63085, Q8BVE3, Q9CR51, P50516, P08556 | 0.82614513 |
| UP_KW_BIOLOGICAL_PROCESS | KW-0406~Ion transport | 7 | 0.9891796559 | O55143, Q8BVE3, Q9CQQ7, Q9CR51, P50516, Q61792, O08997 | 0.98917966 |
| Annotation Cluster 47 | Enrichment Score: 1.29479762583336 | | | | |
| Category | Term | Count | P Value | Genes | FDR |
| GOTERM_BP_DIRECT | GO:0009615~response to virus | 7 | 0.0023078805 | Q9Z0J0, Q64339, Q9CQW9, Q99J93, P18760, P54227, P60710 | 0.07883586 |
| GOTERM_BP_DIRECT | GO:0045071~negative regulation of viral genome replication | 3 | 0.1607487063 | Q64339, Q9CQW9, Q99J93 | 0.92410903 |
| UP_KW_BIOLOGICAL_PROCESS | KW-0051~Antiviral defense | 4 | 0.3517598858 | Q64339, Q9CQW9, Q99J93, O70404 | 0.94565217 |
| Annotation Cluster 48 | Enrichment Score: 1.2749022570474697 | | | | |
| Category | Term | Count | P Value | Genes | FDR |
| UP_KW_MOLECULAR_FUNCTION | KW-0064~Aspartyl protease | 3 | 0.0332874135 | P18242, O09043, A2ADY9 | 0.1420263 |
| GOTERM_MF_DIRECT | GO:0004190~aspartic-type endopeptidase activity | 3 | 0.0667190473 | P18242, O09043, A2ADY9 | 0.4163048 |
| INTERPRO | IPR021109:Peptidase_aspartic_dom_sf | 3 | 0.067416056 | P18242, O09043, A2ADY9 | 0.95672708 |
| Annotation Cluster 49 | Enrichment Score: 1.2457730325231868 | | | | |
| Category | Term | Count | P Value | Genes | FDR |
| GOTERM_CC_DIRECT | GO:0000776~kinetochore | 9 | 0.0015707005 | P46061, O55131, Q9D8B3, Q8BRT1, Q9Z0H3, Q8R1Q8, P60710 | 0.01023698 |
| UP_KW_CELLULAR_COMPONENT | KW-0137~Centromere | 5 | 0.3391868187 | P46061, O55131, Q8BRT1, Q8BK67, Q8R1Q8 | 0.56073367 |
| UP_KW_CELLULAR_COMPONENT | KW-0995~Kinetochore | 4 | 0.3436754763 | P46061, O55131, Q8BRT1, Q8R1Q8 | 0.56073367 |
| Annotation Cluster 50 | Enrichment Score: 1.2145700725486657 | | | | |
| Category | Term | Count | P Value | Genes | FDR |
| GOTERM_MF_DIRECT | GO:0001784~phosphotyrosine residue binding | 5 | 0.0037048101 | P27870, P63085, Q61120, Q9JM90, P62962 | 0.05708432 |
| GOTERM_BP_DIRECT | GO:0002768~immune response-regulating cell surface receptor sig | 3 | 0.0089201105 | P27870, Q9R0C8, P25911 | 0.19288716 |
| KEGG_PATHWAY | mmu04662:B cell receptor signaling pathway | 6 | 0.0871595071 | P27870, P63085, Q9R0C8, P25911, P08556 | 0.35195169 |
| UP_KW_DOMAIN | KW-0727~SH2 domain | 5 | 0.1063296685 | P27870, Q9R0C8, Q61120, P25911, Q9JM90 | 0.37215384 |
| UP_SEQ_FEATURE | DOMAIN:SH2 | 5 | 0.1141522075 | P27870, Q9R0C8, Q61120, P25911, Q9JM90 | 0.99162861 |
| SMART | SM00252:SH2 | 5 | 0.119507927 | P27870, Q9R0C8, Q61120, P25911, Q9JM90 | 0.82968295 |
| INTERPRO | IPR000980:SH2 | 5 | 0.1300225844 | P27870, Q9R0C8, Q61120, P25911, Q9JM90 | 0.99085151 |
| INTERPRO | IPR036860:SH2_dom_sf | 5 | 0.1460127122 | P27870, Q9R0C8, Q61120, P25911, Q9JM90 | 0.99085151 |
| GOTERM_BP_DIRECT | GO:0007169~cell surface receptor protein tyrosine kinase signaling | 4 | 0.1477316133 | P09581, Q61120, P25911, Q9JM90 | 0.88150947 |
| Annotation Cluster 51 | Enrichment Score: 1.1154209839299127 | | | | |
| Category | Term | Count | P Value | Genes | FDR |
| GOTERM_BP_DIRECT | GO:0006694~steroid biosynthetic process | 4 | 0.0159277518 | Q9D0S9, Q8BLN5, Q9EQ06, O70503 | 0.26625327 |
| UP_KW_BIOLOGICAL_PROCESS | KW-0752~Steroid biosynthesis | 4 | 0.0604012956 | Q9D0S9, Q8BLN5, Q9EQ06, O70503 | 0.3284828 |
| UP_KW_BIOLOGICAL_PROCESS | KW-0444~Lipid biosynthesis | 5 | 0.4683131096 | Q9D0S9, P22437, Q8BLN5, Q9EQ06, O70503 | 0.94565217 |
| Annotation Cluster 52 | Enrichment Score: 1.0890806573353309 | | | | |
| Category | Term | Count | P Value | Genes | FDR |
| GOTERM_CC_DIRECT | GO:0033017~sarcoplasmic reticulum membrane | 4 | 0.0136806363 | P26883, O55143, Q6PHZ2, Q6P069 | 0.05484048 |
| UP_KW_CELLULAR_COMPONENT | KW-0703~Sarcoplasmic reticulum | 4 | 0.0909623341 | P26883, O55143, Q6PHZ2, Q6P069 | 0.18798882 |
| GOTERM_BP_DIRECT | GO:0006816~calcium ion transport | 3 | 0.4343003049 | O55143, Q6PHZ2, Q6P069 | 0.97195876 |
| Annotation Cluster 53 | Enrichment Score: 1.08583236803452 | | | | |
| Category | Term | Count | P Value | Genes | FDR |
| GOTERM_MF_DIRECT | GO:0140839~RNA polymerase II CTD heptapeptide repeat P3 isom | 3 | 0.0526135423 | P26883, P58389, P17742 | 0.35153296 |
| GOTERM_MF_DIRECT | GO:0140840~RNA polymerase II CTD heptapeptide repeat P6 isom | 3 | 0.0526135423 | P26883, P58389, P17742 | 0.35153296 |
| UP_KW_MOLECULAR_FUNCTION | KW-0413~Isomerase | 8 | 0.0709621399 | P42125, P26883, Q8BLN5, P09103, Q8BXZ1, P58389, P17751, P17742 | 0.28384856 |
| GOTERM_MF_DIRECT | GO:0003755~peptidyl-prolyl cis-trans isomerase activity | 3 | 0.0980387741 | P26883, P58389, P17742 | 0.53251277 |
| UP_KW_MOLECULAR_FUNCTION | KW-0697~Rotamase | 3 | 0.1932940562 | P26883, P58389, P17742 | 0.5465483 |
| Annotation Cluster 54 | Enrichment Score: 1.0575233564121682 | | | | |
| Category | Term | Count | P Value | Genes | FDR |
| UP_SEQ_FEATURE | DOMAIN:SRCR | 3 | 0.0735706884 | P30204, Q07797, P13379 | 0.99162861 |
| INTERPRO | IPR001190:SRCR | 3 | 0.088195989 | P30204, Q07797, P13379 | 0.99085151 |
| SMART | SM00202:SR | 3 | 0.0906091348 | P30204, Q07797, P13379 | 0.80189084 |
| INTERPRO | IPR036772:SRCR-like_dom_sf | 3 | 0.0936545303 | P30204, Q07797, P13379 | 0.99085151 |
| INTERPRO | IPR017448:SRCR-like_dom | 3 | 0.0936545303 | P30204, Q07797, P13379 | 0.99085151 |
| Annotation Cluster 55 | Enrichment Score: 0.9433267847010083 | | | | |
| Category | Term | Count | P Value | Genes | FDR |
| GOTERM_MF_DIRECT | GO:0005085~guanyl-nucleotide exchange factor activity | 9 | 0.0118968981 | P27870, Q60875, Q8C147, Q69ZK0, Q9R0C8, Q8BPU7, P57776, A2A5R2, Q8BK67 | 0.13818705 |
| INTERPRO | IPR001331:GDS_CDC24_CS | 3 | 0.067416056 | P27870, Q69ZK0, Q9R0C8 | 0.95672708 |
| GOTERM_BP_DIRECT | GO:0045785~positive regulation of cell adhesion | 4 | 0.0755010957 | P27870, Q69ZK0, P09103, Q9R0C8 | 0.634656 |
| INTERPRO | IPR011993:PH-like_dom_sf | 13 | 0.0780200068 | P27870, Q60875, P26041, P70460, Q9D1J1, Q69ZK0, Q3UND0, Q61120, A2A8Z1, P34022, Q9R0C8, Q8BPU7, Q9JM90 | 0.99085151 |

| | | | | | |
|---|---|---|---|---|---|
| UP_SEQ_FEATURE | DOMAIN:DH | 4 | 0.1129903464 | P27870, Q60875, Q69ZK0, Q9R0C8 | 0.99162861 |
| SMART | SM00109:C1 | 4 | 0.1131928259 | P27870, Q60875, Q9R0C8, P28867 | 0.82968295 |
| SMART | SM00325:RhoGEF | 4 | 0.1131928259 | P27870, Q60875, Q69ZK0, Q9R0C8 | 0.82968295 |
| INTERPRO | IPR002219:PE/DAG-bd | 4 | 0.1169977068 | P27870, Q60875, Q9R0C8, P28867 | 0.99085151 |
| INTERPRO | IPR000219:DH-domain | 4 | 0.1208745084 | P27870, Q60875, Q69ZK0, Q9R0C8 | 0.99085151 |
| INTERPRO | IPR035899:DBL_dom_sf | 4 | 0.1208745084 | P27870, Q60875, Q69ZK0, Q9R0C8 | 0.99085151 |
| UP_SEQ_FEATURE | ZN_FING:Phorbol-ester/DAG-type | 3 | 0.1574233832 | P27870, Q60875, Q9R0C8 | 0.99162861 |
| UP_KW_MOLECULAR_FUNCTION | KW-0344~Guanine-nucleotide releasing factor | 7 | 0.1737440747 | P27870, Q60875, Q8C147, Q69ZK0, Q9R0C8, A2A5R2, Q8BK67 | 0.52950575 |
| UP_SEQ_FEATURE | DOMAIN:PH | 8 | 0.1799568551 | P27870, Q60875, A2A8Z1, Q69ZK0, Q3UND0, Q9R0C8, Q8BPU7, Q9JM90 | 0.99162861 |
| INTERPRO | IPR001849:PH_domain | 8 | 0.1898031925 | P27870, Q60875, A2A8Z1, Q69ZK0, Q3UND0, Q9R0C8, Q8BPU7, Q9JM90 | 0.99085151 |
| UP_SEQ_FEATURE | DOMAIN:Phorbol-ester/DAG-type | 3 | 0.2325409363 | P27870, Q60875, P28867 | 0.99162861 |
| SMART | SM00233:PH | 7 | 0.3176425095 | P27870, Q60875, A2A8Z1, Q69ZK0, Q3UND0, Q9R0C8, Q9JM90 | 0.98882682 |
| Annotation Cluster 56 | Enrichment Score: 0.9269560238567052 | | | | |
| Category | Term | Count | P Value | Genes | FDR |
| GOTERM_BP_DIRECT | GO:0006099~tricarboxylic acid cycle | 3 | 0.0745834959 | P28271, Q9WUM5, Q8K2B3 | 0.634656 |
| UP_KW_BIOLOGICAL_PROCESS | KW-0816~Tricarboxylic acid cycle | 3 | 0.0980425572 | P28271, Q9WUM5, Q8K2B3 | 0.39688972 |
| KEGG_PATHWAY | mmu00020:Citrate cycle (TCA cycle) | 3 | 0.2265033149 | P28271, Q9WUM5, Q8K2B3 | 0.61755018 |
| Annotation Cluster 57 | Enrichment Score: 0.8290636123242175 | | | | |
| Category | Term | Count | P Value | Genes | FDR |
| KEGG_PATHWAY | mmu05221:Acute myeloid leukemia | 7 | 0.0135833593 | P05555, P26151, P10810, P63085, P09581, P08556 | 0.09974067 |
| KEGG_PATHWAY | mmu04640:Hematopoietic cell lineage | 5 | 0.2713227044 | P05555, P26151, P10810, P09581, P13379 | 0.67048014 |
| KEGG_PATHWAY | mmu05202:Transcriptional misregulation in cancer | 5 | 0.8837223451 | P05555, P26151, P10810, Q62280, P09581 | 0.88372235 |
| Annotation Cluster 58 | Enrichment Score: 0.7775359201665435 | | | | |
| Category | Term | Count | P Value | Genes | FDR |
| GOTERM_MF_DIRECT | GO:0017116~single-stranded DNA helicase activity | 3 | 0.0310425738 | P49718, Q61881, Q9D0F6 | 0.25755102 |
| KEGG_PATHWAY | mmu03030:DNA replication | 3 | 0.2684358345 | P49718, Q61881, Q9D0F6 | 0.67048014 |
| UP_KW_BIOLOGICAL_PROCESS | KW-0235~DNA replication | 3 | 0.5579483869 | P49718, Q61881, Q9D0F6 | 0.94565217 |
| Annotation Cluster 59 | Enrichment Score: 0.7759046479806261 | | | | |
| Category | Term | Count | P Value | Genes | FDR |
| GOTERM_BP_DIRECT | GO:0006869~lipid transport | 6 | 0.0107972924 | Q05816, Q9QZD8, A2A8Z1, Q9CR62, Q9Z2Z6, Q8BX70 | 0.22129755 |
| GOTERM_BP_DIRECT | GO:0006839~mitochondrial transport | 3 | 0.0164444521 | Q9QZD8, Q9Z2Z6, Q8BMD8 | 0.2673074 |
| UP_SEQ_FEATURE | REPEAT:Solcar 3 | 4 | 0.0448584319 | Q9QZD8, Q9CR62, Q9Z2Z6, Q8BMD8 | 0.88561419 |
| UP_SEQ_FEATURE | REPEAT:Solcar 1 | 4 | 0.0524538437 | Q9QZD8, Q9CR62, Q9Z2Z6, Q8BMD8 | 0.88762803 |
| UP_SEQ_FEATURE | REPEAT:Solcar 2 | 4 | 0.0524538437 | Q9QZD8, Q9CR62, Q9Z2Z6, Q8BMD8 | 0.88762803 |
| INTERPRO | IPR018108:Mitochondrial_sb/sol_carrier | 4 | 0.0652343204 | Q9QZD8, Q9CR62, Q9Z2Z6, Q8BMD8 | 0.95672708 |
| INTERPRO | IPR023395:Mt_carrier_dom_sf | 4 | 0.0652343204 | Q9QZD8, Q9CR62, Q9Z2Z6, Q8BMD8 | 0.95672708 |
| UP_KW_BIOLOGICAL_PROCESS | KW-0445~Lipid transport | 7 | 0.1049248674 | Q9Z0J0, Q05816, Q9QZD8, A2A8Z1, Q9CR62, Q9Z2Z6, Q8BX70 | 0.39688972 |
| UP_SEQ_FEATURE | REPEAT:Solcar | 3 | 0.1574233832 | Q9CR62, Q9Z2Z6, Q8BMD8 | 0.99162861 |
| GOTERM_MF_DIRECT | GO:0015297~antiporter activity | 3 | 0.1787809263 | Q9QZD8, Q9CR62, P51912 | 0.75949047 |
| UP_KW_BIOLOGICAL_PROCESS | KW-0050~Antiport | 4 | 0.2458174604 | Q9QZD8, Q9CR62, Q8BMD8, P51912 | 0.79207848 |
| UP_SEQ_FEATURE | TRANSMEM:Helical | Name=4 | 1.4989293362 | 0.988005548471695 | 1 |
| UP_SEQ_FEATURE | TRANSMEM:Helical | Name=1 | 1.4989293362 | 0.989445406858943 | 1 |
| UP_SEQ_FEATURE | TRANSMEM:Helical | Name=2 | 1.4989293362 | 0.990044667126893 | 1 |
| UP_SEQ_FEATURE | TRANSMEM:Helical | Name=6 | 1.2847965739 | 0.994219888327866 | 1 |
| UP_SEQ_FEATURE | TRANSMEM:Helical | Name=5 | 1.2847965739 | 0.99464289626655 | 1 |
| UP_SEQ_FEATURE | TRANSMEM:Helical | Name=3 | 1.2847965739 | 0.995631168862764 | 1 |
| Annotation Cluster 60 | Enrichment Score: 0.7756926427176729 | | | | |
| Category | Term | Count | P Value | Genes | FDR |
| UP_SEQ_FEATURE | REPEAT:HEAT 8 | 3 | 0.0686711479 | Q8BFY9, Q8BRT1, P70168 | 0.99162861 |
| UP_SEQ_FEATURE | REPEAT:HEAT 7 | 3 | 0.0836898073 | Q8BFY9, Q8BRT1, P70168 | 0.99162861 |
| UP_SEQ_FEATURE | REPEAT:HEAT 6 | 3 | 0.1106228698 | Q8BFY9, Q8BRT1, P70168 | 0.99162861 |
| UP_SEQ_FEATURE | REPEAT:HEAT 5 | 3 | 0.1335506916 | Q8BFY9, Q8BRT1, P70168 | 0.99162861 |
| UP_SEQ_FEATURE | REPEAT:HEAT 4 | 3 | 0.1882451652 | Q8BFY9, Q8BRT1, P70168 | 0.99162861 |
| UP_SEQ_FEATURE | REPEAT:HEAT 3 | 3 | 0.2134430971 | Q8BFY9, Q8BRT1, P70168 | 0.99162861 |
| UP_SEQ_FEATURE | REPEAT:HEAT 2 | 3 | 0.2709702315 | Q8BFY9, Q8BRT1, P70168 | 0.99162861 |
| UP_SEQ_FEATURE | REPEAT:HEAT 1 | 3 | 0.2709702315 | Q8BFY9, Q8BRT1, P70168 | 0.99162861 |
| INTERPRO | IPR011989:ARM-like | 6 | 0.4168493881 | Q91YE6, Q8BVE3, Q8BFY9, Q8BRT1, P70168, Q8BPU7 | 0.99085151 |
| Annotation Cluster 61 | Enrichment Score: 0.6856443181646703 | | | | |
| Category | Term | Count | P Value | Genes | FDR |
| GOTERM_MF_DIRECT | GO:0003925~G protein activity | 8 | 1.003297E-05 | Q9DCE9, P62827, Q9JIW9, Q9Z0E6, P08556, P60766, P62821 | 0.00032934 |
| KEGG_PATHWAY | mmu04664:Fc epsilon RI signaling pathway | 7 | 0.0103246262 | P27870, P63085, P20491, Q9R0C8, P25911, P08556 | 0.08559448 |
| KEGG_PATHWAY | mmu05230:Central carbon metabolism in cancer | 7 | 0.0127092198 | P63085, Q00612, P06151, P51912, P08556, D3Z7P3 | 0.09606675 |
| KEGG_PATHWAY | mmu05221:Acute myeloid leukemia | 7 | 0.0135833593 | P05555, P26151, P10810, P63085, P09581, P08556 | 0.09974067 |
| KEGG_PATHWAY | mmu04650:Natural killer cell mediated cytotoxicity | 9 | 0.0163417964 | P27870, P63085, P20491, P24063, P11835, Q9R0C8, Q61120, P08556 | 0.11350923 |
| GOTERM_BP_DIRECT | GO:0050853~B cell receptor signaling pathway | 4 | 0.019326232 | P06800, P63085, Q9R0C8, P25911 | 0.29772502 |
| KEGG_PATHWAY | mmu04062:Chemokine signaling pathway | 12 | 0.0202338716 | P27870, P63085, Q69ZK0, Q9R0C8, Q61120, Q8BPU7, P25911, P08556, P28867, P08752, P60766 | 0.12494059 |
| KEGG_PATHWAY | mmu04915:Estrogen signaling pathway | 9 | 0.0350139147 | P07901, P63085, P18242, Q61120, P08556, P28867, P63017, P08752 | 0.18364441 |
| KEGG_PATHWAY | mmu05417:Lipid and atherosclerosis | 12 | 0.0424279437 | P27870, P07901, Q6PHZ2, P10810, P63085, O70145, Q9R0C8, P25911, P08556, P63017, P60766 | 0.21380356 |
| KEGG_PATHWAY | mmu04722:Neurotrophin signaling pathway | 8 | 0.0540133893 | Q61599, Q6PHZ2, P63085, Q61120, P08556, P28867, P60766 | 0.25706372 |
| KEGG_PATHWAY | mmu04921:Oxytocin signaling pathway | 9 | 0.0665481588 | Q6PHZ2, P63085, Q60605, P08556, P62137, P08752, P60710 | 0.30164987 |
| INTERPRO | IPR020849:Small_GTPase_Ras-type | 3 | 0.067416056 | Q9JIW9, P08556 | 0.95672708 |
| KEGG_PATHWAY | mmu04360:Axon guidance | 10 | 0.0690750979 | B2RXS4, O55222, Q6PHZ2, P63085, P09055, P18760, P08556, P08752, P60766 | 0.30164987 |
| KEGG_PATHWAY | mmu04370:VEGF signaling pathway | 5 | 0.0795551657 | Q9JIA7, P63085, P08556, P60766 | 0.33517504 |
| KEGG_PATHWAY | mmu05216:Thyroid cancer | 4 | 0.0846945944 | P21107, P63085, P08556 | 0.35107275 |
| UP_KW_DISEASE | KW-0656~Proto-oncogene | 6 | 0.0853313784 | P27870, Q60875, P09581, P25911, P08556 | 0.25599414 |
| KEGG_PATHWAY | mmu04662:B cell receptor signaling pathway | 6 | 0.0871595071 | P27870, P63085, Q9R0C8, P25911, P08556 | 0.35195169 |
| KEGG_PATHWAY | mmu04730:Long-term depression | 5 | 0.0876455579 | P63085, P25911, P08556, P08752 | 0.35195169 |
| KEGG_PATHWAY | mmu04912:GnRH signaling pathway | 6 | 0.1088349384 | Q6PHZ2, P63085, P08556, P28867, P60766 | 0.41747133 |
| KEGG_PATHWAY | mmu04720:Long-term potentiation | 5 | 0.1188847897 | Q6PHZ2, P63085, P08556, P62137 | 0.42190789 |
| KEGG_PATHWAY | mmu05225:Hepatocellular carcinoma | 9 | 0.1198415398 | P63085, Q9JMH6, P48774, Q61120, P08556, Q9Z0H3, P60710 | 0.42190789 |
| KEGG_PATHWAY | mmu04660:T cell receptor signaling pathway | 7 | 0.1287580277 | P27870, P06800, P63085, Q9R0C8, P08556, P60766 | 0.44717315 |
| BIOCARTA | m_fcer1Pathway:Fc Epsilon Receptor I Signaling in Mast Cells | 5 | 0.1390996321 | P27870, P63085, P20491, P25911, P08556 | 1 |
| KEGG_PATHWAY | mmu04071:Sphingolipid signaling pathway | 7 | 0.1435570659 | Q9JIA7, P63085, P20491, P18242, P08556, P08752 | 0.48544955 |
| KEGG_PATHWAY | mmu05214:Glioma | 5 | 0.1542135597 | Q6PHZ2, P63085, Q61120, P08556 | 0.51471279 |
| GOTERM_BP_DIRECT | GO:0050852~T cell receptor signaling pathway | 4 | 0.1765520017 | P06800, P63085, P08556 | 0.97000715 |
| KEGG_PATHWAY | mmu04210:Apoptosis | 7 | 0.1795038572 | Q9Z0X1, P63085, P18242, P08556, P60710 | 0.57665614 |
| KEGG_PATHWAY | mmu04625:C-type lectin receptor signaling pathway | 6 | 0.2068394999 | P63085, P20491, P08556, P28867, P19973 | 0.60754938 |
| KEGG_PATHWAY | mmu04012:ErbB signaling pathway | 5 | 0.2103964769 | Q6PHZ2, P63085, Q61120, P08556 | 0.60754938 |
| KEGG_PATHWAY | mmu04540:Gap junction | 5 | 0.2222733335 | P63085, Q922F4, P08556, P08752 | 0.61755018 |
| KEGG_PATHWAY | mmu05213:Endometrial cancer | 4 | 0.2225429234 | O55222, P63085, P08556 | 0.61755018 |
| GOTERM_BP_DIRECT | GO:0008286~insulin receptor signaling pathway | 4 | 0.2415479021 | P63085, P09581, P84104, P08556 | 0.97195876 |
| KEGG_PATHWAY | mmu04072:Phospholipase D signaling pathway | 7 | 0.2415609416 | Q9JIA7, P63085, P20491, Q9JIW9, Q61120, P08556 | 0.63348124 |
| KEGG_PATHWAY | mmu04919:Thyroid hormone signaling pathway | 6 | 0.24802421 | O55143, P63085, P08556, P60710 | 0.64386083 |
| KEGG_PATHWAY | mmu05166:Human T-cell leukemia virus 1 infection | 10 | 0.2565784798 | P63085, P62827, P34022, P01900, P24063, P11835, P01897, P35564, P08556 | 0.65940669 |
| GOTERM_BP_DIRECT | GO:0007265~Ras protein signal transduction | 3 | 0.2623611714 | Q9JIW9, P08556 | 0.97195876 |
| BIOCARTA | m_malPathway:Role of MAL in Rho-Mediated Activation of SRF | 3 | 0.2681449744 | P63085, P08556, P60766 | 1 |
| GOTERM_BP_DIRECT | GO:0048009~insulin-like growth factor receptor signaling pathway | 3 | 0.2918542799 | P63085, P09581, P08556 | 0.97195876 |

| | | | | | |
|---|---|---|---|---|---|
| KEGG_PATHWAY | mmu04926:Relaxin signaling pathway | 6 | 0.3020757547 | P63085, Q61120, P08556, P08752, P60710 | 0.73239122 |
| KEGG_PATHWAY | mmu04916:Melanogenesis | 5 | 0.3092454769 | Q6PHZ2, P63085, P08556, P08752 | 0.7358897 |
| KEGG_PATHWAY | mmu05211:Renal cell carcinoma | 4 | 0.3140444579 | P63085, P08556, P60766 | 0.737413 |
| KEGG_PATHWAY | mmu04933:AGE-RAGE signaling pathway in diabetic complications | 5 | 0.3156242425 | P63085, P08556, P28867, P60766 | 0.737413 |
| KEGG_PATHWAY | mmu05219:Bladder cancer | 3 | 0.3207543721 | P63085, P08556 | 0.74264751 |
| KEGG_PATHWAY | mmu04917:Prolactin signaling pathway | 4 | 0.3449988699 | P63085, Q61120, P08556 | 0.7471419 |
| KEGG_PATHWAY | mmu04371:Apelin signaling pathway | 6 | 0.352131397 | Q9JIA7, P63085, P08556, P08752, P60710 | 0.7471419 |
| KEGG_PATHWAY | mmu05220:Chronic myeloid leukemia | 4 | 0.3604419215 | P63085, Q61120, P08556 | 0.7471419 |
| KEGG_PATHWAY | mmu01521:EGFR tyrosine kinase inhibitor resistance | 4 | 0.3911465356 | P63085, Q61120, P08556 | 0.77600071 |
| KEGG_PATHWAY | mmu04725:Cholinergic synapse | 5 | 0.392529543 | Q6PHZ2, P63085, P08556, P08752 | 0.77600071 |
| KEGG_PATHWAY | mmu04935:Growth hormone synthesis, secretion and action | 5 | 0.4180000309 | P63085, Q61120, P08556, P08752 | 0.80841582 |
| KEGG_PATHWAY | mmu04261:Adrenergic signaling in cardiomyocytes | 6 | 0.447047426 | P21107, O55143, Q6PHZ2, P63085, P62137, P08752 | 0.82614513 |
| KEGG_PATHWAY | mmu05210:Colorectal cancer | 4 | 0.4512829364 | P63085, Q9JIW9, P08556 | 0.82614513 |
| KEGG_PATHWAY | mmu04114:Oocyte meiosis | 5 | 0.4557152139 | P63101, Q9CQV8, Q6PHZ2, P63085, P62137 | 0.82614513 |
| KEGG_PATHWAY | mmu04150:mTOR signaling pathway | 6 | 0.4580512413 | P63085, Q8BVE3, Q9CR51, P50516, P08556 | 0.82614513 |
| KEGG_PATHWAY | mmu05207:Chemical carcinogenesis - receptor activation | 8 | 0.4602560215 | P07901, P63085, P48774, P70168, P08556, Q6ZQM8, P08752 | 0.82614513 |
| KEGG_PATHWAY | mmu04010:MAPK signaling pathway | 10 | 0.4636656424 | P10810, P63085, P09581, Q8BTM8, P08556, P63017, Q9D7X3, P60766, P54227 | 0.82614513 |
| BIOCARTA | m_keratinocytePathway:Keratinocyte Differentiation | 4 | 0.4825157868 | P63085, Q60854, P08556, P28867 | 1 |
| KEGG_PATHWAY | mmu01522:Endocrine resistance | 4 | 0.4876287869 | P63085, Q61120, P08556 | 0.85252108 |
| KEGG_PATHWAY | mmu05160:Hepatitis C | 6 | 0.5067140345 | P63101, Q9CQV8, P63085, P08556, P60229 | 0.86825994 |
| BIOCARTA | m_bcrPathway:BCR Signaling Pathway | 3 | 0.5210640629 | P27870, P25911, P08556 | 1 |
| KEGG_PATHWAY | mmu05231:Choline metabolism in cancer | 4 | 0.5227771633 | Q8BH43, P63085, P08556 | 0.86825994 |
| KEGG_PATHWAY | mmu04929:GnRH secretion | 3 | 0.5321717669 | P63085, P08556 | 0.86825994 |
| KEGG_PATHWAY | mmu04726:Serotonergic synapse | 5 | 0.5342536509 | P22437, P63085, P08556, P08752 | 0.86825994 |
| KEGG_PATHWAY | mmu05215:Prostate cancer | 4 | 0.5364655508 | P07901, P63085, P08556 | 0.86825994 |
| KEGG_PATHWAY | mmu05165:Human papillomavirus infection | 11 | 0.5571086589 | Q64339, P63085, Q8BVE3, P09055, P01900, Q9CR51, P01897, P50516, P08556, P60766 | 0.88013699 |
| KEGG_PATHWAY | mmu04910:Insulin signaling pathway | 5 | 0.5572857388 | P63085, Q61120, P08556, P62137 | 0.88013699 |
| KEGG_PATHWAY | mmu04218:Cellular senescence | 6 | 0.5835777163 | P63085, P01900, P01897, P08556, P62137 | 0.88013699 |
| KEGG_PATHWAY | mmu05167:Kaposi sarcoma-associated herpesvirus infection | 7 | 0.6008193949 | P63085, P01900, Q69ZK0, P01897, P25911, P08556 | 0.88013699 |
| KEGG_PATHWAY | mmu05218:Melanoma | 3 | 0.6049929599 | P63085, P08556 | 0.88013699 |
| KEGG_PATHWAY | mmu05223:Non-small cell lung cancer | 3 | 0.6049929599 | P63085, P08556 | 0.88013699 |
| KEGG_PATHWAY | mmu04014:Ras signaling pathway | 7 | 0.6556244364 | P63085, P09581, Q9JIW9, Q61120, P08556, P60766 | 0.88013699 |
| KEGG_PATHWAY | mmu05170:Human immunodeficiency virus 1 infection | 7 | 0.6754315712 | P63085, P01900, P01897, P18760, P08556, P08752 | 0.88013699 |
| KEGG_PATHWAY | mmu05161:Hepatitis B | 5 | 0.6813891512 | P63101, Q9CQV8, P63085, P08556 | 0.88013699 |
| KEGG_PATHWAY | mmu05034:Alcoholism | 6 | 0.6958664925 | P63085, Q61120, P08556, P62137, P08752 | 0.88013699 |
| KEGG_PATHWAY | mmu05235:PD-L1 expression and PD-1 checkpoint pathway in canc | 3 | 0.712683984 | P63085, P08556 | 0.88013699 |
| KEGG_PATHWAY | mmu05224:Breast cancer | 4 | 0.7866230614 | P63085, Q61120, P08556 | 0.88013699 |
| KEGG_PATHWAY | mmu05226:Gastric cancer | 4 | 0.7979456433 | P63085, Q61120, P08556 | 0.88013699 |
| KEGG_PATHWAY | mmu05163:Human cytomegalovirus infection | 6 | 0.846345235 | P63085, P01900, P01897, P08556, P08752 | 0.88013699 |
| KEGG_PATHWAY | mmu05200:Pathways in cancer | 13 | 0.8511948126 | P21107, P07901, Q6PHZ2, P63085, P09055, Q9JIW9, P09581, Q9JMH6, P48774, P08556, P08752, P60766 | 0.88013699 |
| KEGG_PATHWAY | mmu04068:FoxO signaling pathway | 3 | 0.892155924 | P63085, P08556 | 0.89215592 |
| KEGG_PATHWAY | mmu04151:PI3K-Akt signaling pathway | 8 | 0.8960858553 | P63101, P07901, Q9CQV8, P63085, P09055, P09581, P08556 | 0.89608586 |
| KEGG_PATHWAY | mmu04550:Signaling pathways regulating pluripotency of stem cells | 3 | 0.9081639885 | P63085, P08556 | 0.90816399 |

Annotation Cluster 62 Enrichment Score: 0.6832718872377764

| Category | Term | Count | P Value | Genes | FDR |
|---|---|---|---|---|---|
| SMART | SM00054:EFh | 6 | 0.1279921056 | P42567, Q7TPR4, Q60605, Q8BMD8, Q6P069, P14069 | 0.82968295 |
| UP_SEQ_FEATURE | DOMAIN:EF-hand 1 | 6 | 0.1574789605 | P42567, Q7TPR4, Q60605, Q8BMD8, Q6P069, P14069 | 0.99162861 |
| UP_SEQ_FEATURE | DOMAIN:EF-hand 2 | 6 | 0.1602436731 | P42567, Q7TPR4, Q60605, Q8BMD8, Q6P069, P14069 | 0.99162861 |
| GOTERM_MF_DIRECT | GO:0005509~calcium ion binding | 14 | 0.1793460045 | Q07076, Q7TPR4, P48036, P34960, Q8BMD8, P35564, Q9EQP2, P14069, P42567, O55143, P40142, P09055, Q60605, Q6P069 | 0.75949047 |
| UP_SEQ_FEATURE | DOMAIN:EF-hand 3 | 4 | 0.2078073358 | P42567, Q60605, Q8BMD8, Q6P069 | 0.99162861 |
| INTERPRO | IPR018247:EF_Hand_1_Ca_BS | 6 | 0.2157802912 | P42567, Q7TPR4, Q8BMD8, Q9EQP2, Q6P069, P14069 | 0.99085151 |
| INTERPRO | IPR002048:EF_hand_dom | 7 | 0.2158518858 | P42567, Q7TPR4, Q60605, Q8BMD8, Q9EQP2, Q6P069, P14069 | 0.99085151 |
| UP_SEQ_FEATURE | DOMAIN:EF-hand | 6 | 0.2624904214 | P42567, Q7TPR4, Q60605, Q8BMD8, Q9EQP2, P14069 | 0.99162861 |
| UP_SEQ_FEATURE | DOMAIN:EF-hand 4 | 3 | 0.2837875011 | P42567, Q8BMD8, Q6P069 | 0.99162861 |
| INTERPRO | IPR011992:EF-hand-dom_pair | 7 | 0.3519391549 | P42567, Q7TPR4, Q60605, Q8BMD8, Q9EQP2, Q6P069, P14069 | 0.99085151 |

Annotation Cluster 63 Enrichment Score: 0.6328403744307034

| Category | Term | Count | P Value | Genes | FDR |
|---|---|---|---|---|---|
| GOTERM_BP_DIRECT | GO:0006508~proteolysis | 14 | 0.0584294691 | Q11136, Q9CXT6, P56399, Q6NSR8, P34960, P18242, O35864, Q9WVJ3, Q61072, O09043, Q8R146, Q11011, Q8CGK3, O89023 | 0.56441909 |
| UP_KW_MOLECULAR_FUNCTION | KW-0482~Metalloprotease | 6 | 0.2733815456 | Q11136, O35864, Q9WVJ3, Q61072, P34960, Q11011 | 0.69985676 |
| UP_KW_MOLECULAR_FUNCTION | KW-0645~Protease | 15 | 0.3801564559 | Q11136, Q9CXT6, P56399, Q6NSR8, P34960, P18242, Q60692, O35864, Q9WVJ3, Q61072, O09043, Q11011, A2ADY9, Q8CGK3, O89023 | 0.83896597 |
| UP_KW_PTM | KW-0865~Zymogen | 8 | 0.4844790452 | Q9WVJ3, Q61072, P34960, P18242, O09043, Q60692, P29416, O89023 | 0.96895809 |

Annotation Cluster 64 Enrichment Score: 0.6109763694942051

| Category | Term | Count | P Value | Genes | FDR |
|---|---|---|---|---|---|
| GOTERM_MF_DIRECT | GO:0004721~phosphoprotein phosphatase activity | 4 | 0.0397459808 | P58389, P62137, Q9D7X3, P35831 | 0.29142823 |
| GOTERM_BP_DIRECT | GO:0016311~dephosphorylation | 3 | 0.0634252737 | P06800, P62137, Q9D7X3 | 0.57892868 |
| GOTERM_BP_DIRECT | GO:0006470~protein dephosphorylation | 4 | 0.0755010957 | P06800, P62137, Q9D7X3, P35831 | 0.634656 |
| GOTERM_MF_DIRECT | GO:0140793~histone H2AXY142 phosphatase activity | 3 | 0.3150400222 | P06800, Q9D7X3, P35831 | 0.93905473 |
| GOTERM_MF_DIRECT | GO:0030946~protein tyrosine phosphatase activity, metal-depender | 3 | 0.3150400222 | P06800, Q9D7X3, P35831 | 0.93905473 |
| GOTERM_MF_DIRECT | GO:0004726~non-membrane spanning protein tyrosine phosphatas | 3 | 0.3150400222 | P06800, Q9D7X3, P35831 | 0.93905473 |
| GOTERM_MF_DIRECT | GO:0004725~protein tyrosine phosphatase activity | 3 | 0.3344239736 | P06800, Q9D7X3, P35831 | 0.93905473 |
| UP_KW_MOLECULAR_FUNCTION | KW-0904~Protein phosphatase | 5 | 0.354756158 | P06800, Q8R0F6, P62137, Q9D7X3, P35831 | 0.81087122 |
| INTERPRO | IPR016130:Tyr_Pase_AS | 3 | 0.3803150637 | P06800, Q9D7X3, P35831 | 0.99085151 |
| UP_SEQ_FEATURE | DOMAIN:Tyrosine-protein phosphatase | 3 | 0.3906556995 | P06800, Q9D7X3, P35831 | 0.99162861 |
| UP_SEQ_FEATURE | ACT_SITE:Phosphocysteine intermediate | 3 | 0.4385615944 | P06800, Q9D7X3, P35831 | 0.99162861 |
| INTERPRO | IPR000387:Tyr_Pase_dom | 3 | 0.4601389133 | P06800, Q9D7X3, P35831 | 0.99085151 |
| INTERPRO | IPR029021:Prot-tyrosine_phosphatase-like | 3 | 0.5390345321 | P06800, Q9D7X3, P35831 | 0.99085151 |

Annotation Cluster 65 Enrichment Score: 0.5959684646038045

| Category | Term | Count | P Value | Genes | FDR |
|---|---|---|---|---|---|
| GOTERM_MF_DIRECT | GO:1990439~MAP kinase serine/threonine phosphatase activity | 3 | 0.2416347721 | Q8R0F6, P62137, Q9D7X3 | 0.88560317 |
| GOTERM_MF_DIRECT | GO:0140791~histone H2AXS140 phosphatase activity | 3 | 0.2416347721 | Q8R0F6, P62137, Q9D7X3 | 0.88560317 |
| GOTERM_MF_DIRECT | GO:0180004~RNA polymerase II CTD heptapeptide repeat Y1 phos | 3 | 0.2416347721 | Q8R0F6, P62137, Q9D7X3 | 0.88560317 |
| GOTERM_MF_DIRECT | GO:0180005~RNA polymerase II CTD heptapeptide repeat T4 phos | 3 | 0.2416347721 | Q8R0F6, P62137, Q9D7X3 | 0.88560317 |
| GOTERM_MF_DIRECT | GO:0180006~RNA polymerase II CTD heptapeptide repeat S2 phos | 3 | 0.2416347721 | Q8R0F6, P62137, Q9D7X3 | 0.88560317 |
| GOTERM_MF_DIRECT | GO:0033192~calmodulin-dependent protein phosphatase activity | 3 | 0.2416347721 | Q8R0F6, P62137, Q9D7X3 | 0.88560317 |
| GOTERM_MF_DIRECT | GO:0180007~RNA polymerase II CTD heptapeptide repeat S5 phos | 3 | 0.2416347721 | Q8R0F6, P62137, Q9D7X3 | 0.88560317 |
| GOTERM_MF_DIRECT | GO:0180008~RNA polymerase II CTD heptapeptide repeat S7 phos | 3 | 0.2416347721 | Q8R0F6, P62137, Q9D7X3 | 0.88560317 |
| GOTERM_MF_DIRECT | GO:0017018~myosin phosphatase activity | 3 | 0.2661398841 | Q8R0F6, P62137, Q9D7X3 | 0.93895146 |
| UP_KW_MOLECULAR_FUNCTION | KW-0904~Protein phosphatase | 5 | 0.354756158 | P06800, Q8R0F6, P62137, Q9D7X3, P35831 | 0.81087122 |

Annotation Cluster 66 Enrichment Score: 0.5385608086580478

| Category | Term | Count | P Value | Genes | FDR |
|---|---|---|---|---|---|
| GOTERM_BP_DIRECT | GO:0045596~negative regulation of cell differentiation | 5 | 0.0815472805 | P63085, Q62280, P60710 | 0.65824295 |
| KEGG_PATHWAY | mmu04520:Adherens junction | 6 | 0.1165941781 | Q8BH43, Q7TPR4, P63085, P60766, P60710 | 0.42190789 |
| KEGG_PATHWAY | mmu04919:Thyroid hormone signaling pathway | 6 | 0.24802421 | O55143, P63085, P08556, P60710 | 0.64386083 |
| GOTERM_BP_DIRECT | GO:0045893~positive regulation of DNA-templated transcription | 8 | 0.8971271946 | O55222, Q7TPR4, P60122, Q9CXT6, P63085, P60710 | 0.97195876 |
| KEGG_PATHWAY | mmu05164:Influenza A | 3 | 0.9588739328 | P63085, P60710 | 0.95887393 |

Annotation Cluster 67 Enrichment Score: 0.529081836825233

| Category | Term | Count | P Value | Genes | FDR |
|---|---|---|---|---|---|

| | | | | | |
|---|---|---|---|---|---|
| KEGG_PATHWAY | mmu04820:Cytoskeleton in muscle cells | 10 | 0.2057953333 | P21107, P09055, P20152, P17182, A2AQP0, Q8BJS4, Q9QXS1, P60710 | 0.60754938 |
| KEGG_PATHWAY | mmu04919:Thyroid hormone signaling pathway | 6 | 0.24802421 | O55143, P63085, P08556, P60710 | 0.64386083 |
| KEGG_PATHWAY | mmu05410:Hypertrophic cardiomyopathy | 5 | 0.3092454769 | P21107, O55143, P09055, P60710 | 0.7358897 |
| KEGG_PATHWAY | mmu05414:Dilated cardiomyopathy | 5 | 0.3284139779 | P21107, O55143, P09055, P60710 | 0.7471419 |
| KEGG_PATHWAY | mmu05412:Arrhythmogenic right ventricular cardiomyopathy | 4 | 0.4364517398 | O55143, P09055, P60710 | 0.82614513 |
| Annotation Cluster 68 | Enrichment Score: 0.4677724967473608 | | | | |
| Category | Term | Count | P Value | Genes | FDR |
| GOTERM_BP_DIRECT | GO:0042752~regulation of circadian rhythm | 3 | 0.2279927087 | Q921F2, Q9D906, P62137 | 0.97195876 |
| GOTERM_BP_DIRECT | GO:0048511~rhythmic process | 3 | 0.2869452187 | Q921F2, Q9D906, P62137 | 0.97195876 |
| UP_KW_BIOLOGICAL_PROCESS | KW-0090~Biological rhythms | 4 | 0.6038965642 | Q921F2, Q9CXT6, Q9D906, P62137 | 0.94565217 |
| Annotation Cluster 69 | Enrichment Score: 0.350458683570693 | | | | |
| Category | Term | Count | P Value | Genes | FDR |
| BIOCARTA | m_ecmPathway:Erk and PI-3 Kinase Are Necessary for Collagen Bi | 4 | 0.1284025806 | P63085, P09055, P08556, P62962 | 1 |
| BIOCARTA | m_tffPathway:Trefoil Factors Initiate Mucosal Healing | 3 | 0.4003881873 | P63085, P09055, P08556 | 1 |
| BIOCARTA | m_erkPathway:Erk1/Erk2 Mapk Signaling pathway | 3 | 0.5047906191 | P63085, P09055, P08556 | 1 |
| BIOCARTA | m_metPathway:Signaling of Hepatocyte Growth Factor Receptor | 3 | 0.5825939849 | P63085, P09055, P08556 | 1 |
| BIOCARTA | m_integrinPathway:Integrin Signaling Pathway | 3 | 0.5825939849 | P63085, P09055, P08556 | 1 |
| KEGG_PATHWAY | mmu04151:PI3K-Akt signaling pathway | 8 | 0.8960858553 | P63101, P07901, Q9CQV8, P63085, P09055, P09581, P08556 | 0.89608586 |
| Annotation Cluster 70 | Enrichment Score: 0.3438014119984769 | | | | |
| Category | Term | Count | P Value | Genes | FDR |
| UP_KW_MOLECULAR_FUNCTION | KW-0646~Protease inhibitor | 5 | 0.3037516107 | Q62426, O88738, Q60854, P32261, P70296 | 0.74769627 |
| GOTERM_MF_DIRECT | GO:0004867~serine-type endopeptidase inhibitor activity | 3 | 0.5349330248 | Q60854, P32261, P70296 | 0.93905473 |
| UP_KW_MOLECULAR_FUNCTION | KW-0722~Serine protease inhibitor | 3 | 0.5725029963 | Q60854, P32261, P70296 | 0.87671233 |
| Annotation Cluster 71 | Enrichment Score: 0.3152903410365088 | | | | |
| Category | Term | Count | P Value | Genes | FDR |
| UP_SEQ_FEATURE | ACT_SITE:Glycyl thioester intermediate | 4 | 0.1438663937 | O88738, Q9D906, Q02053, Q6ZPJ3 | 0.99162861 |
| GOTERM_MF_DIRECT | GO:0004842~ubiquitin-protein transferase activity | 4 | 0.6145302418 | O88738, Q99KP6, E9Q555, Q6ZPJ3 | 0.93905473 |
| KEGG_PATHWAY | mmu04120:Ubiquitin mediated proteolysis | 5 | 0.6327877207 | O88738, Q9R1T2, Q99KP6, Q02053, Q6ZPJ3 | 0.88013699 |
| UP_KW_BIOLOGICAL_PROCESS | KW-0833~Ubl conjugation pathway | 9 | 0.979665864 | O88738, Q64339, Q9R1T2, P56399, Q99KP6, Q9D906, E9Q555, Q02053, Q6ZPJ3 | 0.97966586 |
| Annotation Cluster 72 | Enrichment Score: 0.31245760467866507 | | | | |
| Category | Term | Count | P Value | Genes | FDR |
| GOTERM_BP_DIRECT | GO:0070534~protein K63-linked ubiquitination | 3 | 0.2231040827 | Q99KP6, E9Q555, Q6ZPJ3 | 0.97195876 |
| GOTERM_MF_DIRECT | GO:0004842~ubiquitin-protein transferase activity | 4 | 0.6145302418 | O88738, Q99KP6, E9Q555, Q6ZPJ3 | 0.93905473 |
| GOTERM_MF_DIRECT | GO:0061630~ubiquitin protein ligase activity | 4 | 0.8425126073 | Q99KP6, P60766, E9Q555, Q6ZPJ3 | 0.93905473 |
| Annotation Cluster 73 | Enrichment Score: 0.2774770410873437 | | | | |
| Category | Term | Count | P Value | Genes | FDR |
| UP_SEQ_FEATURE | LIPID:S-palmitoyl cysteine | 8 | 0.2492183172 | P35293, Q9CQW9, Q99J93, P35564, P25911, P08556, P08752 | 0.99162861 |
| UP_KW_PTM | KW-0564~Palmitate | 11 | 0.6889907066 | P35293, Q99JX3, Q9CQW9, O35114, Q99J93, Q8BMK4, P35564, P25911, P08556, P08752 | 1 |
| UP_KW_PTM | KW-0449~Lipoprotein | 22 | 0.8565950364 | Q62159, P53994, Q99JX3, Q9CQW9, P10810, Q99J93, Q9Z0E6, P35564, P35283, P35293, O35114, Q9CRB9, Q8BMK4, Q9JIW9, Q91V41, P25911, P08556, P08752, P60766, P26645, P62821 | 1 |
| Annotation Cluster 74 | Enrichment Score: 0.27432167923108325 | | | | |
| Category | Term | Count | P Value | Genes | FDR |
| KEGG_PATHWAY | mmu04912:GnRH signaling pathway | 6 | 0.1088349384 | Q6PHZ2, P63085, P08556, P28867, P60766 | 0.41747133 |
| GOTERM_BP_DIRECT | GO:0018107~peptidyl-threonine phosphorylation | 3 | 0.133223183 | Q6PHZ2, P63085, P28867 | 0.83512511 |
| GOTERM_BP_DIRECT | GO:0018105~peptidyl-serine phosphorylation | 4 | 0.1667736468 | O55222, Q6PHZ2, P63085, P28867 | 0.95040496 |
| GOTERM_BP_DIRECT | GO:0006468~protein phosphorylation | 8 | 0.1683609261 | P63101, O55222, Q6PHZ2, P63085, P18760, P25911, P28867, P60766 | 0.95040496 |
| GOTERM_BP_DIRECT | GO:0046777~protein autophosphorylation | 4 | 0.3169339497 | P06800, Q6PHZ2, P09581, P25911 | 0.97195876 |
| GOTERM_MF_DIRECT | GO:0140801~histone H2AXY142 kinase activity | 3 | 0.4506520058 | P09581, P25911, P28867 | 0.93905473 |
| GOTERM_MF_DIRECT | GO:0035401~histone H3Y41 kinase activity | 3 | 0.4506520058 | P09581, P25911, P28867 | 0.93905473 |
| GOTERM_BP_DIRECT | GO:0006974~DNA damage response | 6 | 0.4688937798 | P63085, Q9R0C8, P25911, P28867, Q61881, Q02053 | 0.97195876 |
| GOTERM_MF_DIRECT | GO:0004672~protein kinase activity | 5 | 0.6181647044 | O55222, Q6PHZ2, P63085, P25911, P28867 | 0.93905473 |
| INTERPRO | IPR001245:Ser-Thr/Tyr_kinase_cat_dom | 3 | 0.7109437353 | O55222, P09581, P25911 | 0.99085151 |
| GOTERM_MF_DIRECT | GO:0106310~protein serine kinase activity | 4 | 0.8646601159 | O55222, Q6PHZ2, P63085, P28867 | 0.93905473 |
| GOTERM_MF_DIRECT | GO:0004674~protein serine/threonine kinase activity | 4 | 0.9088915826 | O55222, Q6PHZ2, P63085, P28867 | 0.93905473 |
| INTERPRO | IPR017441:Protein_kinase_ATP_BS | 5 | 0.9215903675 | Q6PHZ2, P63085, P09581, P25911, P28867 | 0.99085151 |
| UP_SEQ_FEATURE | DOMAIN:Protein kinase | 6 | 0.9481448699 | O55222, Q6PHZ2, P63085, P09581, P25911, P28867 | 0.99162861 |
| INTERPRO | IPR000719:Prot_kinase_dom | 6 | 0.9540898384 | O55222, Q6PHZ2, P63085, P09581, P25911, P28867 | 0.99085151 |
| INTERPRO | IPR011009:Kinase-like_dom_sf | 6 | 0.9721901361 | O55222, Q6PHZ2, P63085, P09581, P25911, P28867 | 0.99085151 |
| UP_KW_MOLECULAR_FUNCTION | KW-0418~Kinase | 10 | 0.9770416418 | Q9WV85, O55222, Q9JIA7, Q6PHZ2, P63085, P09581, Q8VDL4, Q9R0N0, P25911, P28867 | 0.97704164 |
| INTERPRO | IPR008271:Ser/Thr_kinase_AS | 3 | 0.9824952494 | Q6PHZ2, P63085, P28867 | 0.99085151 |
| SMART | SM00220:S_TKc | 3 | 0.9930393354 | Q6PHZ2, P63085, P28867 | 0.99303934 |
| UP_KW_MOLECULAR_FUNCTION | KW-0723~Serine/threonine-protein kinase | 4 | 0.9950929625 | O55222, Q6PHZ2, P63085, P28867 | 0.99509296 |
| Annotation Cluster 75 | Enrichment Score: 0.27123397778238434 | | | | |
| Category | Term | Count | P Value | Genes | FDR |
| UP_SEQ_FEATURE | REPEAT:ANK | 5 | 0.4003944551 | O55222, P62774, D3Z7P3, Q810B6, Q62422 | 0.99162861 |
| UP_SEQ_FEATURE | REPEAT:ANK 1 | 5 | 0.4885349065 | O55222, P62774, D3Z7P3, Q810B6, Q62422 | 0.99162861 |
| UP_SEQ_FEATURE | REPEAT:ANK 2 | 5 | 0.4885349065 | O55222, P62774, D3Z7P3, Q810B6, Q62422 | 0.99162861 |
| UP_KW_DOMAIN | KW-0040~ANK repeat | 5 | 0.5593246663 | O55222, P62774, D3Z7P3, Q810B6, Q62422 | 0.95454545 |
| SMART | SM00248:ANK | 5 | 0.5806805325 | O55222, P62774, D3Z7P3, Q810B6, Q62422 | 0.98882682 |
| UP_SEQ_FEATURE | REPEAT:ANK 3 | 4 | 0.5841346909 | O55222, P62774, Q810B6, Q62422 | 0.99162861 |
| INTERPRO | IPR002110:Ankyrin_rpt | 5 | 0.5976433827 | O55222, P62774, D3Z7P3, Q810B6, Q62422 | 0.99085151 |
| INTERPRO | IPR036770:Ankyrin_rpt-contain_sf | 5 | 0.624155167 | O55222, P62774, D3Z7P3, Q810B6, Q62422 | 0.99085151 |
| Annotation Cluster 76 | Enrichment Score: 0.24227664131015783 | | | | |
| Category | Term | Count | P Value | Genes | FDR |
| GOTERM_BP_DIRECT | GO:0006910~phagocytosis, recognition | 3 | 0.0209317687 | P26151, P08508, P97797 | 0.29862181 |
| KEGG_PATHWAY | mmu04380:Osteoclast differentiation | 7 | 0.1837068643 | O55143, P26151, P63085, O70145, P09581, P08508, P97797 | 0.5828724 |
| UP_SEQ_FEATURE | DOMAIN:Ig-like C2-type 2 | 4 | 0.4627633477 | P26151, Q8R5M8, P09581, P08508 | 0.99162861 |
| UP_SEQ_FEATURE | DOMAIN:Ig-like C2-type 1 | 4 | 0.4672149607 | P26151, Q8R5M8, P09581, P08508 | 0.99162861 |
| INTERPRO | IPR003006:Ig/MHC_CS | 3 | 0.5173146016 | P01900, P97797, P01897 | 0.99085151 |
| INTERPRO | IPR003597:Ig_C1-set | 3 | 0.6143338758 | P01900, P97797, P01897 | 0.99085151 |
| SMART | SM00407:IGc1 | 3 | 0.6146363868 | P01900, P97797, P01897 | 0.98882682 |
| INTERPRO | IPR003598:Ig_sub2 | 3 | 0.9322113041 | P26151, Q8R5M8, P09581 | 0.99085151 |
| SMART | SM00408:IGc2 | 3 | 0.9371878007 | P26151, Q8R5M8, P09581 | 0.98882682 |
| UP_KW_DOMAIN | KW-0393~Immunoglobulin domain | 5 | 0.9508508511 | P26151, Q8R5M8, P09581, P08508, P97797 | 0.95454545 |
| INTERPRO | IPR003599:Ig_sub | 5 | 0.9952198119 | P26151, Q8R5M8, P09581, P08508, P97797 | 0.99521981 |
| UP_SEQ_FEATURE | DOMAIN:Ig-like | 7 | 0.9956954667 | P26151, Q8R5M8, P09581, P08508, P01900, P97797, P01897 | 0.99569547 |
| SMART | SM00409:IG | 5 | 0.9962166787 | P26151, Q8R5M8, P09581, P08508, P97797 | 0.99621668 |
| INTERPRO | IPR013783:Ig-like_fold | 12 | 0.9987387796 | B2RXS4, Q9WV55, P06800, P26151, Q8R5M8, P09581, P08508, P01900, P97797, Q8BTM8, P01897, Q2YFS3 | 0.99873878 |
| INTERPRO | IPR036179:Ig-like_dom_sf | 8 | 0.9995508767 | P26151, Q8R5M8, P09581, P08508, P01900, P97797, P01897, Q2YFS3 | 0.99955088 |
| INTERPRO | IPR007110:Ig-like_dom | 7 | 0.9997230491 | P26151, Q8R5M8, P09581, P08508, P01900, P97797, P01897 | 0.99972305 |
| Annotation Cluster 77 | Enrichment Score: 0.226411215733629 | | | | |
| Category | Term | Count | P Value | Genes | FDR |
| KEGG_PATHWAY | mmu04916:Melanogenesis | 5 | 0.3092454769 | Q6PHZ2, P63085, P08556, P08752 | 0.7358897 |

| | | | | | |
|---|---|---|---|---|---|
| KEGG_PATHWAY | mmu04725:Cholinergic synapse | 5 | 0.392529543 | Q6PHZ2, P63085, P08556, P08752 | 0.77600071 |
| KEGG_PATHWAY | mmu04261:Adrenergic signaling in cardiomyocytes | 6 | 0.447047426 | P21107, O55143, Q6PHZ2, P63085, P62137, P08752 | 0.82614513 |
| KEGG_PATHWAY | mmu04024:cAMP signaling pathway | 7 | 0.6051969857 | P27870, O55143, Q6PHZ2, P63085, Q9R0C8, P62137, P08752 | 0.88013699 |
| KEGG_PATHWAY | mmu04022:cGMP-PKG signaling pathway | 5 | 0.7169177617 | O55143, P70460, P63085, P62137, P08752 | 0.88013699 |
| KEGG_PATHWAY | mmu04713:Circadian entrainment | 3 | 0.7716008735 | Q6PHZ2, P63085, P08752 | 0.88013699 |
| KEGG_PATHWAY | mmu04728:Dopaminergic synapse | 3 | 0.8993100489 | Q6PHZ2, P62137, P08752 | 0.89931005 |
| KEGG_PATHWAY | mmu04934:Cushing syndrome | 3 | 0.9451760691 | Q6PHZ2, P63085, P08752 | 0.94517607 |

Annotation Cluster 78 — Enrichment Score: 0.21024061946845682

| Category | Term | Count | P Value | Genes | FDR |
|---|---|---|---|---|---|
| INTERPRO | IPR011990:TPR-like_helical_dom_sf | 6 | 0.3510821268 | Q9CWZ7, Q9CYG7, O70145, Q9CZW5, Q8BK72, P14685 | 0.99085151 |
| UP_SEQ_FEATURE | REPEAT:TPR 3 | 3 | 0.6424850451 | Q9CYG7, O70145, Q9CZW5 | 0.99162861 |
| SMART | SM00028:TPR | 3 | 0.6470458151 | Q9CYG7, O70145, Q9CZW5 | 0.98882682 |
| INTERPRO | IPR019734:TPR_repeat | 3 | 0.6675275315 | Q9CYG7, O70145, Q9CZW5 | 0.99085151 |
| UP_SEQ_FEATURE | REPEAT:TPR 1 | 3 | 0.6872526332 | Q9CYG7, O70145, Q9CZW5 | 0.99162861 |
| UP_SEQ_FEATURE | REPEAT:TPR 2 | 3 | 0.6872526332 | Q9CYG7, O70145, Q9CZW5 | 0.99162861 |
| UP_KW_DOMAIN | KW-0802~TPR repeat | 3 | 0.73351142 | Q9CYG7, O70145, Q9CZW5 | 0.95454545 |

Annotation Cluster 79 — Enrichment Score: 0.05662323057152441

| Category | Term | Count | P Value | Genes | FDR |
|---|---|---|---|---|---|
| INTERPRO | IPR020472:G-protein_beta_WD-40_rep | 3 | 0.4892272039 | Q8BFQ4, Q99KP6, O88342 | 0.99085151 |
| UP_SEQ_FEATURE | REPEAT:WD 6 | 3 | 0.8359623406 | Q8BFQ4, Q99KP6, O88342 | 0.99162861 |
| UP_SEQ_FEATURE | REPEAT:WD 5 | 3 | 0.9039554341 | Q8BFQ4, Q99KP6, O88342 | 0.99162861 |
| INTERPRO | IPR036322:WD40_repeat_dom_sf | 4 | 0.9149665921 | Q8BFQ4, O88738, Q99KP6, O88342 | 0.99085151 |
| UP_SEQ_FEATURE | REPEAT:WD 4 | 3 | 0.9222710168 | Q8BFQ4, Q99KP6, O88342 | 0.99162861 |
| UP_SEQ_FEATURE | REPEAT:WD 3 | 3 | 0.9335582315 | Q8BFQ4, Q99KP6, O88342 | 0.99162861 |
| UP_SEQ_FEATURE | REPEAT:WD 1 | 3 | 0.9372645042 | Q8BFQ4, Q99KP6, O88342 | 0.99162861 |
| UP_SEQ_FEATURE | REPEAT:WD 2 | 3 | 0.9372645042 | Q8BFQ4, Q99KP6, O88342 | 0.99162861 |
| UP_KW_DOMAIN | KW-0853~WD repeat | 3 | 0.9410760013 | Q8BFQ4, Q99KP6, O88342 | 0.95454545 |
| INTERPRO | IPR015943:WD40/YVTN_repeat-like_dom_sf | 4 | 0.9527566811 | B2RXS4, Q8BFQ4, Q99KP6, O88342 | 0.99085151 |
| INTERPRO | IPR001680:WD40_rpt | 3 | 0.9532935053 | Q8BFQ4, Q99KP6, O88342 | 0.99085151 |
| SMART | SM00320:WD40 | 3 | 0.9565639637 | Q8BFQ4, Q99KP6, O88342 | 0.98882682 |

Annotation Cluster 80 — Enrichment Score: 0.038734058351749895

| Category | Term | Count | P Value | Genes | FDR |
|---|---|---|---|---|---|
| GOTERM_BP_DIRECT | GO:0007283~spermatogenesis | 6 | 0.8061060897 | O55131, Q99JX3, Q61425, Q8R5M8, P47791, Q9DBR1 | 0.97195876 |
| UP_KW_BIOLOGICAL_PROCESS | KW-0744~Spermatogenesis | 4 | 0.9318958466 | O55131, Q99JX3, Q61425, Q8R5M8 | 0.94565217 |
| GOTERM_BP_DIRECT | GO:0030154~cell differentiation | 6 | 0.9394453795 | O55222, Q99JX3, Q60875, Q61425, P16110, Q8R5M8 | 0.97195876 |
| UP_KW_BIOLOGICAL_PROCESS | KW-0221~Differentiation | 9 | 0.9918195252 | O55131, Q99JX3, Q60875, Q61425, Q7TMB8, P16110, Q8R5M8, P60766, P54227 | 0.99181953 |

Annotation Cluster 81 — Enrichment Score: 0.0322857676846713

| Category | Term | Count | P Value | Genes | FDR |
|---|---|---|---|---|---|
| UP_KW_MOLECULAR_FUNCTION | KW-0678~Repressor | 12 | 0.7536374232 | Q921F2, O35129, Q3UZ39, P62960, Q8BL97, P61979, Q9CRB9, O55201, P13864, Q64213, Q3UEB3, P63017 | 0.87671233 |
| UP_KW_MOLECULAR_FUNCTION | KW-0010~Activator | 9 | 0.9880479972 | P10639, P62960, P61979, P60122, Q62280, O55201, P13864, O35691, Q9Z0H3 | 0.988048 |
| UP_KW_BIOLOGICAL_PROCESS | KW-0805~Transcription regulation | 26 | 0.9983849034 | Q921F2, P10639, P13864, O35691, Q9Z0H3, O35326, O35129, Q9CRB9, Q2EMV9, Q9DBR1, Q9CSN1, Q3UZ39, Q8BFQ4, P61979, Q9CXT6, Q64213, P57776, Q3UEB3, P62960, Q8C2Q3, P60122, Q62280, O35892, Q3U0V1, O55201, P63017 | 0.9983849 |
| UP_KW_BIOLOGICAL_PROCESS | KW-0804~Transcription | 26 | 0.9991217785 | Q921F2, P10639, P13864, O35691, Q9Z0H3, O35326, O35129, Q9CRB9, Q2EMV9, Q9DBR1, Q9CSN1, Q3UZ39, Q8BFQ4, P61979, Q9CXT6, Q64213, P57776, Q3UEB3, P62960, Q8C2Q3, P60122, Q62280, O35892, Q3U0V1, O55201, P63017 | 0.99912178 |

Annotation Cluster 82 — Enrichment Score: 0.015018386156453812

| Category | Term | Count | P Value | Genes | FDR |
|---|---|---|---|---|---|
| UP_SEQ_FEATURE | TOPO_DOM:Cytoplasmic | 51 | 0.8848660764 | P30204, Q9CR67, P49300, Q9CZW5, Q8R5M8, Q9CR60, Q8VCH8, P09055, Q9DC16, Q8BXZ1, P41216, P20491, P26151, P97797, Q9QY81, P35564, P51912, Q99P72, Q2YFS3, B2RXS4, P46978, P17047, O55143, Q8BXQ2, P60060, O35114, Q8BMK4, P09581, P01900, Q9Z2Z6, O70404, P31996, Q9CQW9, Q9WV55, P24063, Q9QUJ7, Q80VA0, Q9CYN2, P01897, Q3TDQ1, P05555, Q61543, Q69ZN7, P06800, Q99J93, P11835, P08508, P13379, Q61072, P24668, Q8BG07 | 0.99162861 |
| UP_KW_DOMAIN | KW-1133~Transmembrane helix | 79 | 0.9847783847 | P30204, Q8CAQ8, Q9CR67, P49300, Q9CZW5, Q8R5M8, Q9CQQ7, P48771, Q69ZK0, Q9CR62, P19253, Q9CR60, Q9DBL1, P09055, Q921T2, Q9DC16, Q8BXZ1, A1L314, P41216, P47758, Q8R1I1, P20491, P26151, P97797, Q9QY81, O35682, P35564, P51912, Q9CQX2, Q99P72, Q2YFS3, B2RXS4, P46978, O55101, P17047, O55143, Q8BXQ2, P60060, O35114, P06151, Q8BMK4, P09581, P01900, Q9Z2Z6, O70404, P31996, Q9D1Q4, Q9CQW9, Q9WV55, P24063, Q9ERS2, Q9QUJ7, Q80VA0, Q9CYN2, Q7TNS2, P01897, Q8BPX9, Q3TDQ1, Q6ZQM8, Q8BM55, P05555, Q61543, Q69ZN7, P06800, Q9QZD8, Q99J93, P11835, P08508, Q8BMD8, P13379, P17665, Q80TL7, Q9Z0X1, Q61072, P24668, O70503, Q8BJS4, P60603, Q8BG07 | 0.98477838 |
| UP_KW_DOMAIN | KW-0812~Transmembrane | 79 | 0.999333497 | P30204, Q8CAQ8, Q9CR67, P49300, Q9CZW5, Q8R5M8, Q9CQQ7, P48771, Q69ZK0, Q9CR62, P19253, Q9CR60, Q9DBL1, P09055, Q921T2, Q9DC16, Q8BXZ1, A1L314, P41216, P47758, Q8R1I1, P20491, P26151, P97797, Q9QY81, O35682, P35564, P51912, Q9CQX2, Q99P72, Q2YFS3, B2RXS4, P46978, O55101, P17047, O55143, Q8BXQ2, P60060, O35114, P06151, Q8BMK4, P09581, P01900, Q9Z2Z6, O70404, P31996, Q9D1Q4, Q9CQW9, Q9WV55, P24063, Q9ERS2, Q9QUJ7, Q80VA0, Q9CYN2, Q7TNS2, P01897, Q8BPX9, Q3TDQ1, Q6ZQM8, Q8BM55, P05555, Q61543, Q69ZN7, P06800, Q9QZD8, Q99J93, P11835, P08508, Q8BMD8, P13379, P17665, Q80TL7, Q9Z0X1, Q61072, P24668, O70503, Q8BJS4, P60603, Q8BG07 | 0.9993335 |
| UP_SEQ_FEATURE | TRANSMEM:Helical | 72 | 0.9999999017 | P30204, Q8CAQ8, Q9CR67, P49300, Q9CZW5, Q8R5M8, Q9CQQ7, P48771, Q69ZK0, P19253, Q9DBL1, P09055, Q921T2, Q9DC16, Q8BXZ1, A1L314, P41216, P47758, Q8R1I1, P20491, P26151, P97797, Q9QY81, O35682, P35564, P51912, Q9CQX2, Q99P72, Q2YFS3, B2RXS4, P46978, O55101, P17047, O55143, Q8BXQ2, P60060, O35114, P06151, Q8BMK4, P09581, P01900, Q9Z2Z6, O70404, P31996, Q9D1Q4, Q9CQW9, Q9WV55, P24063, Q9ERS2, Q9CYN2, Q7TNS2, P01897, Q8BPX9, Q3TDQ1, Q6ZQM8, Q8BM55, P05555, Q61543, Q69ZN7, P06800, Q99J93, P11835, P08508, P13379, P17665, Q80TL7, Q9Z0X1, Q61072, P24668, O70503, Q8BJS4, P60603 | 0.9999999 |

Annotation Cluster 83 — Enrichment Score: 0.009513705297499635

| Category | Term | Count | P Value | Genes | FDR |
|---|---|---|---|---|---|
| UP_SEQ_FEATURE | CARBOHYD:N-linked (GlcNAc...) asparagine | 56 | 0.916756822 | P30204, Q8BFR4, P49300, Q8R5M8, P34960, P18242, Q62087, Q07797, P09055, Q921T2, Q9DC16, Q8BXZ1, O09043, A1L314, O89023, P26151, P97797, Q9QY81, P51912, Q2YFS3, B2RXS4, P46978, P17047, Q8BXQ2, O35114, P32261, P09581, P01900, P29416, P31996, Q9Z0J0, P10810, P20060, P24063, Q9ESY9, P22437, Q9WVJ3, P01897, Q8BPX9, Q3TDQ1, Q6ZQM8, Q8BM55, P05555, Q61543, P06800, P11835, P08508, P13379, P17742, Q8VIM6, O88668, Q61072, P24668, Q3TCN2, Q8BJS4, Q8BG07 | 0.99162861 |

| | | | | | |
|---|---|---|---|---|---|
| UP_KW_DOMAIN | KW-0732~Signal | 58 | 0.9992908513 | Q8BFR4, Q9CR67, Q8R5M8, P34960, P18242, Q62087, Q07797, O88738, P09055, Q8BXZ1, O09043, A1L314, O89023, Q9WV85, P20491, P26151, P97797, Q9QY81, P35564, Q2YFS3, B2RXS4, P17047, Q8BXQ2, P16110, P32261, P09581, P01900, P09103, P05064, P29416, Q9D0S9, P31996, Q9Z0J0, Q6PHZ2, P10810, Q9EQ06, P20060, P24063, Q9ESY9, Q9QXT0, P22437, Q9WVJ3, P01897, Q6ZQM8, Q8CGK3, P05555, Q61543, P06800, P11835, P08508, Q8VDL4, Q9D6Z1, P13379, Q8VIM6, O88668, Q61072, P24668, Q3TCN2 | 0.99929085 |
| UP_KW_PTM | KW-1015~Disulfide bond | 61 | 0.9999979668 | P10639, P30204, P49300, P56399, Q8R5M8, Q9JMH6, P34960, Q9R112, P18242, Q62087, Q07797, Q9CRB9, P09055, P62075, Q8BXZ1, O09043, A1L314, O89023, P20491, P26151, P47791, P70296, P97797, P35564, B2RXS4, P17047, O55143, Q8BXQ2, P16110, O35114, P32261, P09581, P01900, P09103, P29416, Q8CDN6, P31996, Q9Z0J0, Q9WV55, P10810, P20060, P24063, Q9ESY9, Q9QXT0, P22437, Q80VA0, P01897, P05555, Q64339, P11835, P08508, P13379, Q99J99, Q05816, P63158, Q61072, P38060, P24668, Q3TCN2, Q8BJS4, P56391 | 1 |
| UP_KW_PTM | KW-0325~Glycoprotein | 60 | 1 | P30204, Q8BFR4, P49300, Q8R5M8, P34960, P18242, Q62087, Q07797, P09055, Q921T2, Q9DC16, Q8BXZ1, O09043, A1L314, P20152, P41216, O89023, P26151, P97797, Q9QY81, P51912, Q2YFS3, Q61881, B2RXS4, P46978, P17047, Q8BXQ2, O35114, P32261, P09581, P01900, P29416, P31996, Q9Z0J0, P10810, P20060, P24063, Q9ESY9, P22437, Q9WVJ3, P01897, Q8BPX9, Q3TDQ1, Q6ZQM8, Q8BM55, P05555, Q61543, P61979, P06800, P11835, P08508, P13379, P17742, Q8VIM6, O88668, Q61072, P24668, Q3TCN2, Q8BJS4, Q8BG07 | 1 |

Annotation Cluster 84 Enrichment Score: 0.001189229701544587

| Category | Term | Count | P Value | Genes | FDR |
|---|---|---|---|---|---|
| UP_KW_BIOLOGICAL_PROCESS | KW-0221~Differentiation | 9 | 0.9918195252 | O55131, Q99JX3, Q60875, Q61425, Q7TMB8, P16110, Q8R5M8, P60766, P54227 | 0.99181953 |
| UP_KW_MOLECULAR_FUNCTION | KW-0217~Developmental protein | 6 | 0.9999996053 | B2RXS4, P35293, Q60875, Q7TMB8, Q8R5M8, P54227 | 0.99999961 |
| UP_KW_MOLECULAR_FUNCTION | KW-9996~Developmental protein | 6 | 0.9999996053 | B2RXS4, P35293, Q60875, Q7TMB8, Q8R5M8, P54227 | 0.99999961 |

Supplementary Table 7: Results of the pathway analysis by the David tool on the proteins modulated in response to the 4x10µg/ml treatment

Annotation Cluster 1 — Enrichment Score: 11.68179045913022

| Category | Term | Count | P Value | Genes | FDR |
|---|---|---|---|---|---|
| GOTERM_CC_DIRECT | GO:0005743~mitochondrial inner membrane | 37 | 1.9564336068E-15 | Q8CAQ8, Q9D0M3, Q62425, P48771, Q9CR62, Q9ERS2, Q7TMF3, Q91WD5, Q8JZN5, O35129, Q3UMR5, Q9CRB9, Q8BWT1, P62075, Q9DC70, Q924T2, Q9CQF0, Q8R1I1, Q91YT0, P62908, Q9CQN1, Q9CPR5, Q9QZD8, Q9CQN6, P51175, Q03265, Q9CQX2, P17665, Q9Z0X1, P16110, P38060, Q78IK4, Q8BMS1, P47802, P56391, P00375 | 2.303701E-13 |
| UP_KW_CELLULAR_COMPONENT | KW-0496~Mitochondrion | 67 | 1.3525086012E-12 | Q9D0M3, Q8CAQ8, Q9CZW5, P48771, Q9CR62, P38647, Q9WTP7, Q7TMF3, Q8K0D5, Q8JZN5, Q9DBL1, O35129, Q3UMR5, Q9CRB9, Q8BWT1, P62075, Q3TIU4, P41216, Q8R1I1, Q9WV85, Q9CPR5, Q3U5Q7, Q03265, P35564, Q9CQX2, Q9JHI5, Q9Z2I9, P32020, Q8BMS1, P00375, Q8VCW8, Q921F2, Q62425, Q9ERS2, O08528, Q64442, Q91WD5, Q9JLT4, Q9QUJ7, Q9DC70, Q8BX70, Q924T2, Q9CQF0, P00520, Q8CGK3, Q91YT0, P62908, Q9CQN1, P30416, Q9QZD8, Q9CQN6, P51175, O35143, P59017, O08734, P17665, O35459, Q9Z0X1, Q80Y14, P38060, Q78IK4, Q8VBT0, P29758, P47802, P56391 | 2.231639E-11 |
| UP_KW_CELLULAR_COMPONENT | KW-0999~Mitochondrion inner membrane | 27 | 3.4042658847E-09 | Q8CAQ8, Q9D0M3, Q62425, P48771, Q9CR62, Q9ERS2, Q7TMF3, Q91WD5, Q8JZN5, O35129, Q3UMR5, Q9CRB9, P62075, Q9DC70, Q8R1I1, Q91YT0, P62908, Q9CQN1, Q9QZD8, P51175, Q03265, P17665, Q9Z0X1, Q78IK4, Q8BMS1, P56391 | 3.744692E-08 |

Annotation Cluster 2 — Enrichment Score: 9.988715455079534

| Category | Term | Count | P Value | Genes | FDR |
|---|---|---|---|---|---|
| UP_KW_CELLULAR_COMPONENT | KW-0496~Mitochondrion | 67 | 1.3525086012E-12 | Q9D0M3, Q8CAQ8, Q9CZW5, P48771, Q9CR62, P38647, Q9WTP7, Q7TMF3, Q8K0D5, Q8JZN5, Q9DBL1, O35129, Q3UMR5, Q9CRB9, Q8BWT1, P62075, Q3TIU4, P41216, Q8R1I1, Q9WV85, Q9CPR5, Q3U5Q7, Q03265, P35564, Q9CQX2, Q9JHI5, Q9Z2I9, P32020, Q8BMS1, P00375, Q8VCW8, Q921F2, Q62425, Q9ERS2, O08528, Q64442, Q91WD5, Q9JLT4, Q9QUJ7, Q9DC70, Q8BX70, Q924T2, Q9CQF0, P00520, Q8CGK3, Q91YT0, P62908, Q9CQN1, P30416, Q9QZD8, Q9CQN6, P51175, O35143, P59017, O08734, P17665, O35459, Q9Z0X1, Q80Y14, P38060, Q78IK4, Q8VBT0, P29758, P47802, P56391 | 2.231639E-11 |
| UP_KW_DOMAIN | KW-0809~Transit peptide | 34 | 6.1088418445E-11 | Q8VCW8, Q8CAQ8, Q9D0M3, Q921H8, P48771, P38647, Q91WD5, Q8K0D5, Q8JZN5, Q9DBL1, Q3UMR5, Q9JLT4, Q8BWT1, Q3TIU4, Q9DC70, Q9CQF0, Q8CGK3, Q91YT0, Q9CQN1, Q9CPR5, Q3U5Q7, O35143, Q03265, Q9JHI5, P17665, O35459, Q9Z0X1, Q9Z2I9, Q80Y14, P38060, Q78IK4, P29758, Q8BMS1 | 1.343945E-09 |
| UP_SEQ_FEATURE | TRANSIT:Mitochondrion | 31 | 1.3084427271E-08 | Q8VCW8, Q8CAQ8, Q9D0M3, P48771, P38647, Q91WD5, Q8K0D5, Q8JZN5, Q9DBL1, Q3UMR5, Q9JLT4, Q3TIU4, Q9DC70, Q9CQF0, Q8CGK3, Q91YT0, Q9CQN1, Q9CPR5, Q3U5Q7, O35143, Q03265, Q9JHI5, P17665, O35459, Q9Z0X1, Q9Z2I9, Q80Y14, P38060, Q78IK4, P29758, Q8BMS1 | 4.298234E-06 |

Annotation Cluster 3 — Enrichment Score: 9.362601053239956

| Category | Term | Count | P Value | Genes | FDR |
|---|---|---|---|---|---|
| UP_SEQ_FEATURE | CROSSLNK:Glycyl lysine isopeptide (Lys-Gly) (interchain with G-Cter in SUMO2) | 55 | 2.6394323685E-13 | Q3UHX2, Q9EQK5, O70133, Q9CZW5, Q99L45, P18760, P13864, Q61033, Q61990, Q9JM76, Q6DFW4, Q9Z1F9, P60335, P62317, P20152, P27546, Q6Y7W8, P62315, P46061, Q8BK64, P97310, P56960, Q61881, Q91VC3, P42932, P62960, P21619, Q5SSI6, E9Q555, Q921F2, Q61656, Q7TPV4, Q501J6, Q80X82, P49717, O35226, P40142, Q91XV3, P62908, P80313, P61979, Q61029, E9Q7G0, Q3TEA8, Q9D6Z1, Q3UKJ7, Q7TMK9, Q3UEB3, O08784, Q8C2Q3, P24547, Q8CCF0, P80318, P80317, P41105, P60843 | 1.734107E-10 |
| UP_KW_PTM | KW-1017~Isopeptide bond | 89 | 3.4112818727E-11 | Q3UHX2, Q9EQK5, Q8BL97, O70133, Q922F4, Q9CZW5, Q8BTM8, Q99L45, P18760, P13864, Q61033, Q61990, P17751, Q9QY76, Q9JM76, P09055, Q6DFW4, Q9Z1F9, P60335, O35609, P62317, P20152, P27546, Q6Y7W8, P62315, P46061, Q8BK64, P97310, P56960, Q61881, Q91VC3, Q9DBG6, P14685, P42932, P62960, P21619, P06151, Q9D906, Q5SSI6, P62962, P63017, P08775, E9Q555, P00493, Q921F2, Q61656, Q9CQW9, Q7TPV4, Q501J6, Q9CQ22, Q80X82, P49717, O35226, P40142, Q91XV3, P28656, P62908, P80313, Q3UZ39, Q9CQV8, Q64339, P61979, Q61029, P30416, E9Q7G0, Q3TEA8, P10107, Q9D6Z1, Q3UKJ7, Q7TMK9, Q3UEB3, Q05144, P17742, O08784, Q9Z0X1, Q8C2Q3, Q3U0V1, O35892, P67871, P17182, P24547, P08556, Q8CCF0, P47802, P80318, P80317, P41105, P60843 | 4.093538E-10 |
| UP_KW_PTM | KW-0832~Ubl conjugation | 100 | 9.0733715526E-06 | Q8BL97, O70133, Q9CZW5, Q9JMH6, Q9WVL2, P18760, P13864, Q9DBD5, Q61033, Q9JM76, Q6DFW4, P60335, P20152, Q8BK64, P14685, Q91VC3, P06151, P01900, Q9D906, P52293, Q5SSI6, E9Q555, Q921F2, Q61656, Q7TPV4, Q9ET26, Q9CQ22, P40142, Q91XV3, P61979, Q61029, Q3TEA8, P10107, Q9D6Z1, Q3UKJ7, Q8C2Q3, Q3U0V1, O35892, P08556, P41105, Q3UHX2, Q9EQK5, Q91VH2, Q8VBW6, Q8BTM8, Q99L45, Q61990, P17751, Q9QY76, P63085, P09055, Q9Z1F9, O35609, P62317, P27546, Q6Y7W8, P62315, P46061, P97310, P56960, P35564, Q61881, Q9DBG6, P42932, P62960, P21619, P09581, P63017, P62962, P08775, P00493, Q9CQW9, Q501J6, Q80X82, P49717, O35226, P68037, P00520, P62908, Q3UZ39, P80313, Q9CQV8, E9Q7G0, P30416, Q7TMK9, Q3UEB3, P17742, O08784, Q9Z0X1, P67871, P17182, P24547, P25911, Q8CCF0, P47802, P80318, P60843, P80317, Q8BGQ7 | 5.444023E-05 |

Annotation Cluster 4 — Enrichment Score: 8.477049976856629

| Category | Term | Count | P Value | Genes | FDR |
|---|---|---|---|---|---|
| GOTERM_MF_DIRECT | GO:0005524~ATP binding | 62 | 1.6269100193E-14 | Q8CG48, Q8CG47, Q8C0C7, O70133, P38647, Q9WTP7, P63085, Q9Z1F9, P09411, Q8BU30, Q91YR1, P41216, Q9WV85, P97310, P46460, Q3U5Q7, P58389, Q03265, Q9ER72, P53657, Q61881, Q91VC3, O55222, P42932, O55143, Q9Z2I9, P09581, Q9JHU4, E9Q634, P63017, E9Q555, Q8VCW8, Q61656, Q501J6, Q9ERS2, O08528, Q8R5C5, Q9DCL9, P49718, P49717, P68037, Q9QUJ7, P26638, G5E829, P62334, Q9D7H3, P00520, Q8CGK3, Q9D0F6, Q9CQN1, P80313, P30416, Q9EQP2, A2AQP0, Q3THK7, P25911, P80318, P43247, P80317, Q8BGQ7, P60843, P07742 | 4.251658E-12 |
| GOTERM_MF_DIRECT | GO:0016887~ATP hydrolysis activity | 29 | 3.3236377954E-12 | Q8CG48, Q8CG47, Q61656, O70133, Q501J6, P38647, P49718, P49717, G5E829, P62334, Q8CGK3, Q9D0F6, Q9CQN1, P80313, P97310, P46460, Q03265, Q61881, Q91VC3, P42932, O55143, Q9JHU4, P80318, E9Q634, P63017, P43247, P60843, P80317, E9Q555 | 3.948975E-10 |
| UP_KW_LIGAND | KW-0547~Nucleotide-binding | 78 | 4.5302722355E-06 | Q8CG48, Q8CG47, Q8C0C7, O70133, Q922F4, P38647, Q9WTP7, Q8K0D5, P63085, Q9Z1F9, Q9CQD1, P09411, Q8BU30, P41216, P47758, Q9DCE9, Q9WV85, P97310, P46460, Q3U5Q7, P58389, Q03265, Q9Z0E6, Q9ER72, P53657, Q61881, P36536, Q91VC3, O55222, P42932, Q8R050, O55143, Q9Z2I9, P09581, Q9JHU4, E9Q634, P63017, E9Q555, Q8VCW8, P00493, Q61656, Q9Z0N1, Q501J6, O08528, Q8R5C5, Q9DCL9, P49718, P49717, P68037, Q9JLT4, Q9QUJ7, P26638, G5E829, P62334, Q9D7H3, P00520, Q8CGK3, Q9D0F6, Q62159, Q9CQN1, P80313, Q9EQP2, A2AQP0, Q3THK7, Q05144, Q9D892, Q8CHH9, Q91V41, P25911, P08556, P80318, P43247, P80317, P08752, Q8BGQ7, P60843, P07742 | 0.0001223174 |
| UP_KW_LIGAND | KW-0067~ATP-binding | 58 | 0.00050431114877 | Q8CG48, Q8CG47, Q8C0C7, O70133, P38647, P63085, Q9Z1F9, P09411, Q8BU30, P41216, Q9WV85, P97310, P46460, Q3U5Q7, P58389, Q03265, Q9ER72, P53657, Q61881, Q91VC3, O55222, P42932, O55143, Q9Z2I9, P09581, Q9JHU4, E9Q634, P63017, E9Q555, Q8VCW8, Q61656, Q501J6, O08528, Q8R5C5, Q9DCL9, P49718, P49717, P68037, Q9QUJ7, P26638, G5E829, P62334, Q9D7H3, P00520, Q8CGK3, Q9D0F6, Q9CQN1, P80313, Q9EQP2, A2AQP0, Q3THK7, P25911, P80318, P43247, P80317, Q8BGQ7, P60843, P07742 | 0.0045388003 |

Annotation Cluster 5 — Enrichment Score: 5.37181629330274

| Category | Term | Count | P Value | Genes | FDR |
|---|---|---|---|---|---|
| GOTERM_MF_DIRECT | GO:0051015~actin filament binding | 19 | 1.7214428145E-09 | Q99K51, Q7TPR4, Q8BTM8, P18760, A2AQP0, P59999, Q61792, Q9JL26, P21107, Q7TMB8, Q9JM76, P57780, P47754, O88342, P47757, Q91YR1, E9Q634, P00520, P26645 | 1.499568E-07 |
| GOTERM_MF_DIRECT | GO:0003779~actin binding | 20 | 2.4272166728E-07 | Q8BH43, Q7TPR4, P26041, Q8BTM8, Q9Z0E6, P18760, A2AQP0, P19973, Q9JL26, P21107, Q9JM76, P57780, P09055, P09103, P47754, O88342, P47757, Q91YR1, P62962, E9Q634 | 1.463798E-05 |
| UP_KW_MOLECULAR_FUNCTION | KW-0009~Actin-binding | 19 | 2.3571732151E-05 | Q99K51, Q8BH43, Q7TPR4, Q8BTM8, P18760, A2AQP0, P59999, Q61792, P21107, Q7TMB8, Q9JM76, P57780, P47754, O88342, P47757, Q91YR1, P62962, E9Q634, P26645 | 0.0005107209 |
| GOTERM_CC_DIRECT | GO:0005903~brush border | 8 | 0.00017730692947 | P21107, Q7TPR4, P57780, Q8BTM8, P47754, A2AQP0, P47757, E9Q634 | 0.0022570693 |
| GOTERM_CC_DIRECT | GO:0015629~actin cytoskeleton | 12 | 0.00079213678272 | P21107, Q7TPR4, P57780, O70133, Q8BTM8, Q9CQD1, Q9Z0E6, P18760, O88342, Q91YR1, E9Q634, P00520 | 0.0066624362 |

Annotation Cluster 6 — Enrichment Score: 4.934097146321651

| Category | Term | Count | P Value | Genes | FDR |
|---|---|---|---|---|---|
| GOTERM_MF_DIRECT | GO:0003723~RNA binding | 36 | 1.2498599033E-09 | Q921F2, Q3UHX2, Q8BL97, Q61656, Q9D7S7, O70133, P13864, Q61990, O35841, Q62093, P84104, P60335, P62317, P20152, P62315, P46061, P62908, Q9CQN1, P61979, P28271, Q9D6Z1, Q7TMK9, Q64213, Q3UEB3, Q91VC3, P62320, P62960, Q8C2Q3, Q3U0V1, P23116, P17182, P24547, P62307, P63017, P60843 | 1.224863E-07 |
| GOTERM_CC_DIRECT | GO:0005681~spliceosomal complex | 14 | 8.3145965858E-08 | Q61656, P61979, Q7TMK9, Q64213, Q91VC3, P62320, O35841, P16110, Q62093, Q99KP6, P62317, P62307, P63017, P62315 | 2.797268E-06 |
| GOTERM_MF_DIRECT | GO:0003729~mRNA binding | 19 | 1.3938020244E-07 | P62908, Q8BL97, Q61656, P61979, O70133, Q501J6, Q99L45, Q64213, Q61990, Q91VC3, Q8CIE6, P62960, Q8C2Q3, Q3U0V1, P60335, P84104, P23116, P60843, P00375 | 9.106173E-06 |
| UP_KW_BIOLOGICAL_PROCESS | KW-0508~mRNA splicing | 23 | 4.3627851102E-07 | Q921F2, Q8BL97, Q61656, P61979, O70133, Q501J6, Q3UKJ7, Q7TMK9, Q64213, Q3UEB3, Q91VC3, P62320, P62960, P16110, Q62093, Q99KP6, Q3U0V1, P84104, P62317, P62307, Q8CCF0, P63017, P62315 | 3.926507E-05 |
| GOTERM_BP_DIRECT | GO:0000398~mRNA splicing, via spliceosome | 14 | 6.7154659413E-07 | Q8BL97, Q3UKJ7, Q64213, Q91VC3, P62320, Q8C2Q3, Q62093, Q99KP6, P84104, P62317, P62307, Q8CCF0, P63017, P62315 | 0.0003188503 |
| GOTERM_BP_DIRECT | GO:0008380~RNA splicing | 14 | 1.2570416472E-06 | Q921F2, Q8BL97, P61979, Q3UKJ7, Q7TMK9, Q3UEB3, Q91VC3, P62960, P16110, Q62093, Q3U0V1, P84104, P62307, P63017 | 0.0004973695 |
| UP_KW_BIOLOGICAL_PROCESS | KW-0507~mRNA processing | 25 | 1.5787617575E-06 | Q921F2, Q8BL97, Q61656, O70133, Q501J6, Q80X82, Q62093, Q99KP6, P84104, Q3TIU4, P62317, P62315, P61979, Q3UKJ7, Q7TMK9, Q64213, Q3UEB3, Q91VC3, P62320, P62960, P16110, Q3U0V1, P62307, Q8CCF0, P63017 | 7.104428E-05 |
| GOTERM_CC_DIRECT | GO:0005687~U4 snRNP | 5 | 9.0767868826E-06 | P62320, P62317, P62307, Q8CCF0, P62315 | 0.0002464707 |
| UP_KW_CELLULAR_COMPONENT | KW-0747~Spliceosome | 14 | 1.3140000304E-05 | Q61656, P61979, Q3UKJ7, Q7TMK9, Q64213, Q91VC3, P62320, P16110, Q99KP6, P62317, P62307, Q8CCF0, P63017, P62315 | 0.000108405 |
| GOTERM_CC_DIRECT | GO:0071013~catalytic step 2 spliceosome | 10 | 3.8126400968E-05 | P62320, Q61656, P61979, Q99KP6, P62317, P62307, Q7TMK9, Q3UEB3, Q91VC3, P62315 | 0.0007807624 |
| GOTERM_CC_DIRECT | GO:0016607~nuclear speck | 16 | 0.00097136599997 | Q921F2, Q8BL97, Q8CG47, Q61656, Q3TEA8, Q501J6, Q3UKJ7, Q91VC3, O35841, Q8C2Q3, Q62093, Q99KP6, P60335, P84104, Q8CCF0, Q91XV3 | 0.0080265506 |
| GOTERM_CC_DIRECT | GO:0046540~U4/U6 x U5 tri-snRNP complex | 5 | 0.001386376195 | P62320, P62317, P62307, Q8CCF0, P62315 | 0.0100458952 |
| GOTERM_CC_DIRECT | GO:0071007~U2-type catalytic step 2 spliceosome | 5 | 0.00337728719737 | P62320, Q99KP6, P62317, P62307, P62315 | 0.020658471 |
| UP_KW_MOLECULAR_FUNCTION | KW-0694~RNA-binding | 28 | 0.00355475363187 | Q921F2, Q8BL97, Q61656, O70133, Q501J6, Q61990, Q62093, P84104, P60335, P62908, P61979, P56960, P28271, Q7TMK9, Q64213, Q3UEB3, Q91VC3, P62320, P62960, Q8C2Q3, Q3U0V1, P23116, P24547, P62307, Q8CCF0, Q8BGQ7, P60843, P00375 | 0.0288823733 |
| KEGG_PATHWAY | mmu03040:Spliceosome | 15 | 0.01573034118537 | Q8BL97, Q61656, P61979, Q3UEB3, Q91VC3, P62320, Q62093, Q99KP6, P60335, P84104, P62317, P62307, Q8CCF0, P63017, P62315 | 0.1182022781 |

Annotation Cluster 7 — Enrichment Score: 4.40814615645226

| Category | Term | Count | P Value | Genes | FDR |
|---|---|---|---|---|---|
| GOTERM_CC_DIRECT | GO:0005764~lysosome | 20 | 1.0483849734E-07 | Q9Z0J0, Q9EQH3, Q61207, P20060, Q9ESY9, Q9CQ22, P16675, P17047, Q9WVJ3, O35114, Q91V41, O09043, Q78T54, P45377, Q8BX70, Q3TCN2, P29416, Q8VEB4, P63017, O89023 | 3.291929E-06 |
| KEGG_PATHWAY | mmu04142:Lysosome | 13 | 0.00036803873951 | Q9Z0J0, Q61207, Q8BVE3, P20060, P16675, P17047, O35114, P24668, O09043, Q9DCR2, P29416, Q8VEB4, O89023 | 0.0060496368 |
| UP_KW_CELLULAR_COMPONENT | KW-0458~Lysosome | 19 | 0.00154577830794 | Q9Z0J0, Q9CQW9, Q99J93, Q61207, P20060, Q9ESY9, Q9CQ22, P36536, P16675, P17047, Q9WVJ3, O35114, P24668, Q8BX70, Q3TCN2, P29416, Q8VEB4, P63017, O89023 | 0.0072872406 |

Annotation Cluster 8 — Enrichment Score: 4.379021761271706

| Category | Term | Count | P Value | Genes | FDR |
|---|---|---|---|---|---|
| UP_KW_CELLULAR_COMPONENT | KW-0999~Mitochondrion inner membrane | 27 | 3.4042658847E-09 | Q8CAQ8, Q9D0M3, Q62425, P48771, Q9CR62, Q9ERS2, Q7TMF3, Q91WD5, Q8JZN5, O35129, Q3UMR5, Q9CRB9, P62075, Q9DC70, Q8R1I1, Q91YT0, P62908, Q9CQN1, Q9QZD8, P51175, Q03265, P17665, Q9Z0X1, Q78IK4, Q8BMS1, P56391 | 3.744692E-08 |
| KEGG_PATHWAY | mmu05020:Prion disease | 24 | 2.0313547357E-06 | Q9D0M3, O70435, Q62425, Q922F4, O70145, P48771, Q9ERS2, Q03265, Q7TMF3, Q60692, Q91WD5, P14685, Q05144, P17665, Q3UMR5, O35226, P63085, P67871, Q9DC70, P62334, Q8R1I1, P56391, P63017, Q91YT0 | 0.0004911376 |
| UP_KW_BIOLOGICAL_PROCESS | KW-0249~Electron transport | 13 | 3.2545255013E-06 | Q9D0M3, P10639, Q62425, Q9ERS2, Q7TMF3, Q9CQX2, Q91WD5, Q8R180, Q8VBT0, Q9DC70, Q8CDN6, Q8R1I1, Q91YT0 | 9.763577E-05 |
| KEGG_PATHWAY | mmu05012:Parkinson disease | 23 | 7.4697727525E-06 | Q9D0M3, P10639, Q9CQN1, O70435, Q62425, Q922F4, P48771, Q9ERS2, Q03265, Q7TMF3, Q60692, Q91WD5, P14685, P17665, Q3UMR5, O35226, P68037, Q9DC70, P62334, Q8R1I1, P56391, P08752, Q91YT0 | 0.0004911376 |
| KEGG_PATHWAY | mmu05014:Amyotrophic lateral sclerosis | 27 | 1.5562545594E-05 | Q921F2, Q9D0M3, Q07076, O70435, Q8BL97, Q62425, Q922F4, P48771, Q9ERS2, Q7TMF3, Q91WD5, Q9QY76, Q8R5C5, Q3UMR5, O35226, P84104, Q9CQD1, Q9DC70, P62334, Q8R1I1, Q91YT0, Q03265, Q60692, P14685, P17665, P62962, P56391 | 0.0008185899 |
| KEGG_PATHWAY | mmu05022:Pathways of neurodegeneration - multiple diseases | 31 | 2.6521784692E-05 | Q921F2, Q9D0M3, O70435, Q62425, Q922F4, P48771, Q9ERS2, Q7TMF3, Q91WD5, Q9QY76, Q8R5C5, Q3UMR5, O35226, P63085, P68037, Q9CQD1, Q9DC70, P62334, Q8R1I1, Q91YT0, Q9CQN1, Q03265, Q60692, O08734, P14685, P17665, O55143, P67871, P08556, P56391 | 0.0009964613 |
| KEGG_PATHWAY | mmu05208:Chemical carcinogenesis - reactive oxygen species | 19 | 6.7217756618E-05 | Q9D0M3, Q62425, O70145, P48771, Q9ERS2, Q03265, Q7TMF3, Q91WD5, P17665, P63085, P48774, Q9DC70, P08556, P48758, Q8R1I1, P56391, P00520, Q91YT0 | 0.0017649765 |
| KEGG_PATHWAY | mmu05010:Alzheimer disease | 26 | 8.0531248699E-05 | Q9D0M3, O70435, Q62425, Q922F4, P48771, Q8VBW6, Q9ERS2, Q7TMF3, Q91WD5, Q3UMR5, O35226, P63085, Q9DC70, P62334, Q8R1I1, Q91YT0, Q03265, Q60692, Q99P72, P14685, P17665, O55143, P67871, P08556, P56391 | 0.0017649765 |
| KEGG_PATHWAY | mmu05415:Diabetic cardiomyopathy | 18 | 0.00011187152033 | Q9D0M3, Q62425, O70145, P48771, Q9ERS2, P58389, Q03265, Q7TMF3, Q91WD5, Q05144, P17665, O55143, Q00612, Q9DC70, P62137, Q8R1I1, P56391, Q91YT0 | 0.0022632469 |
| KEGG_PATHWAY | mmu05016:Huntington disease | 21 | 0.00036496908335 | Q9D0M3, O70435, Q62425, Q922F4, P48771, Q9ERS2, Q03265, Q7TMF3, Q60692, Q91WD5, P17426, P14685, Q8R5C5, P17665, O35226, Q9DC70, P62334, Q8R1I1, P56391, Q91YT0, P08775 | 0.0060496368 |
| UP_KW_BIOLOGICAL_PROCESS | KW-0679~Respiratory chain | 8 | 0.00041268523348 | Q9D0M3, Q62425, Q9ERS2, Q7TMF3, Q9DC70, Q91WD5, Q8R1I1, Q91YT0 | 0.005305953 |
| KEGG_PATHWAY | mmu00190:Oxidative phosphorylation | 13 | 0.0005471428077 | Q9D0M3, Q62425, Q8BVE3, P48771, Q9ERS2, Q03265, Q7TMF3, Q91WD5, P17665, Q9DC70, Q8R1I1, P56391, Q91YT0 | 0.0084646211 |
| KEGG_PATHWAY | mmu04714:Thermogenesis | 16 | 0.00260442250111 | Q9D0M3, Q62425, P48771, Q9ERS2, Q03265, Q7TMF3, Q91WD5, P17665, Q9QUJ7, Q9DC70, P08556, P41216, Q8R1I1, P56391, Q91YT0 | 0.0311346872 |
| KEGG_PATHWAY | mmu04932:Non-alcoholic fatty liver disease | 12 | 0.00547712334262 | Q9D0M3, Q62425, P48771, Q9ERS2, Q7TMF3, Q9DC70, P53657, Q91WD5, P56391, Q8R1I1, P17665, Q91YT0 | 0.0554032092 |

Annotation Cluster 9 — Enrichment Score: 4.024500793213954

| Category | Term | Count | P Value | Genes | FDR |
|---|---|---|---|---|---|
| GOTERM_BP_DIRECT | GO:0046166~glyceraldehyde-3-phosphate biosynthetic process | 6 | 4.5495571687E-07 | Q9QZD8, P40142, P09411, P17182, P17751 | 0.0002700162 |
| KEGG_PATHWAY | mmu01200:Carbon metabolism | 15 | 6.8232416014E-06 | Q99K85, P28271, Q9R0P3, O08528, P53657, P17751, Q93092, P40142, Q00612, Q9Z2I9, P05063, P09411, P17182 | 0.0004911376 |
| UP_KW_BIOLOGICAL_PROCESS | KW-0324~Glycolysis | 7 | 4.6742405541E-05 | P05063, O08528, P09411, P17182, P53657, P17751 | 0.0008413633 |
| GOTERM_BP_DIRECT | GO:0006096~glycolytic process | 7 | 4.8829490861E-05 | P05063, O08528, P09411, P17182, P53657, P17751 | 0.0072450757 |
| KEGG_PATHWAY | mmu01230:Biosynthesis of amino acids | 11 | 6.1754569397E-05 | Q99K85, P40142, P28271, P05063, P09411, P17182, P53657, P17751, Q93092, Q8VCN5 | 0.0017649765 |
| GOTERM_BP_DIRECT | GO:0006094~gluconeogenesis | 7 | 6.8240879363E-05 | Q9QZD8, Q9CR62, P05063, P09411, P17182, P17751 | 0.0095296381 |
| GOTERM_BP_DIRECT | GO:0061621~canonical glycolysis | 5 | 7.8221498173E-05 | P09411, P17182, P59017, P17751 | 0.0103165465 |
| KEGG_PATHWAY | mmu00010:Glycolysis / Gluconeogenesis | 8 | 0.00247947254089 | P06151, P05063, O08528, P09411, P17182, P53657, P17751 | 0.0310524418 |
| KEGG_PATHWAY | mmu04066:HIF-1 signaling pathway | 7 | 0.10392991807348 | P63085, P06151, P05063, O08528, P09411, P17182 | 0.3961386732 |

Annotation Cluster 10 — Enrichment Score: 4.012171247968376

| Category | Term | Count | P Value | Genes | FDR |
|---|---|---|---|---|---|
| GOTERM_MF_DIRECT | GO:0001786~phosphatidylserine binding | 9 | 8.1140208016E-06 | P21956, Q07076, P10107, O35114, P14824, O35639, P97429, P63017, P26645 | 0.0002859082 |
| UP_KW_DOMAIN | KW-0041~Annexin | 5 | 2.8076885987E-05 | Q07076, P10107, P14824, O35639, P97429 | 0.0002058972 |
| UP_SEQ_FEATURE | REPEAT:Annexin 1 | 5 | 3.5664651856E-05 | Q07076, P10107, P14824, O35639, P97429 | 0.0042603048 |
| UP_SEQ_FEATURE | REPEAT:Annexin 2 | 5 | 3.5664651856E-05 | Q07076, P10107, P14824, O35639, P97429 | 0.0042603048 |
| UP_SEQ_FEATURE | REPEAT:Annexin 3 | 5 | 3.5664651856E-05 | Q07076, P10107, P14824, O35639, P97429 | 0.0042603048 |
| UP_SEQ_FEATURE | REPEAT:Annexin 4 | 5 | 3.5664651856E-05 | Q07076, P10107, P14824, O35639, P97429 | 0.0042603048 |
| SMART | SM00335:ANX | 5 | 3.8271313451E-05 | Q07076, P10107, P14824, O35639, P97429 | 0.0067740225 |
| INTERPRO | IPR018252:Annexin_repeat_CS | 5 | 4.2810493764E-05 | Q07076, P10107, P14824, O35639, P97429 | 0.0076951863 |
| INTERPRO | IPR018502:Annexin_repeat | 5 | 4.2810493764E-05 | Q07076, P10107, P14824, O35639, P97429 | 0.0076951863 |
| INTERPRO | IPR037104:Annexin_sf | 5 | 4.2810493764E-05 | Q07076, P10107, P14824, O35639, P97429 | 0.0076951863 |
| INTERPRO | IPR001464:Annexin | 5 | 4.2810493764E-05 | Q07076, P10107, P14824, O35639, P97429 | 0.0076951863 |
| GOTERM_CC_DIRECT | GO:0012506~vesicle membrane | 6 | 0.00011885096865 | Q07076, P10107, P14824, Q9Z0E6, O35639, P97429 | 0.0016464355 |
| UP_KW_LIGAND | KW-0111~Calcium/phospholipid-binding | 5 | 0.00028830430807 | Q07076, P10107, P14824, O35639, P97429 | 0.0038921082 |
| GOTERM_MF_DIRECT | GO:0005544~calcium-dependent phospholipid binding | 6 | 0.0007240062237 | Q07076, P10107, P14824, O35639, P97429, Q8BT60 | 0.0129004745 |
| GOTERM_MF_DIRECT | GO:0048306~calcium-dependent protein binding | 7 | 0.00155570404991 | Q07076, Q61656, P10107, P14824, O35639, P97429, Q8BT60 | 0.0259504676 |
| GOTERM_CC_DIRECT | GO:0062023~collagen-containing extracellular matrix | 10 | 0.03492412767491 | B2RXS4, P21956, Q07076, Q62426, P16110, Q60854, P10107, P14824, O35639, P97429 | 0.1359443317 |

Annotation Cluster 11 — Enrichment Score: 3.4309799212895293

| Category | Term | Count | P Value | Genes | FDR |
|---|---|---|---|---|---|
| GOTERM_BP_DIRECT | GO:0006457~protein folding | 13 | 1.3798536961E-07 | Q8BK64, Q9CQN1, P80313, P38647, P35564, Q8R180, P17742, P42932, Q9CWM4, P09103, P80318, P63017, P80317 | 0.0001091924 |
| GOTERM_MF_DIRECT | GO:0140662~ATP-dependent protein folding chaperone | 7 | 2.302127862E-06 | P80313, P42932, Q9CQN1, P38647, P63017, P80318, P80317 | 0.0001061687 |
| GOTERM_MF_DIRECT | GO:0051082~unfolded protein binding | 10 | 3.6487784674E-06 | P80313, P42932, Q9CQN1, Q99L47, P38647, Q9CWM4, P35564, P63017, P80318, P80317 | 0.0001505601 |
| GOTERM_MF_DIRECT | GO:0044183~protein folding chaperone | 8 | 8.752291515E-06 | P80313, P42932, Q3U0V1, P38647, Q9CWM4, P63017, P80318, P80317 | 0.0002859082 |
| GOTERM_BP_DIRECT | GO:0061077~chaperone-mediated protein folding | 6 | 0.00013535640158 | P80313, P42932, P30416, P63017, P80318, P80317 | 0.0139711347 |
| UP_KW_MOLECULAR_FUNCTION | KW-0143~Chaperone | 15 | 0.00024299185241 | Q8BK64, Q9CQN1, P80313, P30416, P38647, Q99L47, P35564, Q91YE6, P42932, P62075, Q9CWM4, P09103, P80318, P63017, P80317 | 0.0031588941 |
| GOTERM_BP_DIRECT | GO:1904851~positive regulation of establishment of protein localization to telomere | 4 | 0.00025369771128 | P80313, P42932, P80318, P80317 | 0.0192673144 |
| GOTERM_CC_DIRECT | GO:0005832~chaperonin-containing T-complex | 4 | 0.00025372950459 | P80313, P42932, P80318, P80317 | 0.0029876649 |
| INTERPRO | IPR002194:Chaperonin_TCP-1_CS | 4 | 0.00060258717144 | P80313, P42932, P80318, P80317 | 0.0509717854 |
| INTERPRO | IPR017998:Chaperone_TCP-1 | 4 | 0.00081770312501 | P80313, P42932, P80318, P80317 | 0.0605979211 |
| GOTERM_CC_DIRECT | GO:0044297~cell body | 8 | 0.00130946225656 | P80313, P42932, P09581, P20152, P80318, P80317, P08752, P41105 | 0.0096368238 |
| GOTERM_BP_DIRECT | GO:0032212~positive regulation of telomere maintenance via telomerase | 5 | 0.00153328879461 | P80313, P42932, P63085, P80318, P80317 | 0.0769098136 |
| INTERPRO | IPR027413:GROEL-like_equatorial_sf | 4 | 0.00173405477336 | P80313, P42932, P80318, P80317 | 0.1038987818 |
| INTERPRO | IPR002423:Cpn60/GroEL/TCP-1 | 4 | 0.00213927332929 | P80313, P42932, P80318, P80317 | 0.1139361129 |
| INTERPRO | IPR027409:GroEL-like_apical_dom_sf | 4 | 0.00213927332929 | P80313, P42932, P80318, P80317 | 0.1139361129 |
| INTERPRO | IPR027410:TCP-1-like_intermed_sf | 4 | 0.00213927332929 | P80313, P42932, P80318, P80317 | 0.1139361129 |
| GOTERM_BP_DIRECT | GO:0051086~chaperone mediated protein folding independent of cofactor | 3 | 0.00347526732748 | P80313, P42932, P80318 | 0.1269274559 |
| INTERPRO | IPR053374:TCP-1_chaperonin | 3 | 0.00815861902832 | P80313, P80318, P80317 | 0.2793355753 |
| GOTERM_CC_DIRECT | GO:0002199~zona pellucida receptor complex | 3 | 0.00879088192962 | P42932, P80318, P80317 | 0.0450054934 |
| GOTERM_BP_DIRECT | GO:0007339~binding of sperm to zona pellucida | 4 | 0.0267042473669 | P80313, P42932, P80318, P80317 | 0.3913326126 |

Annotation Cluster 12 — Enrichment Score: 3.399939886102026

| Category | Term | Count | P Value | Genes | FDR |
|---|---|---|---|---|---|
| GOTERM_MF_DIRECT | GO:0016887~ATP hydrolysis activity | 29 | 3.3236377954E-12 | Q8CG48, Q8CG47, Q61656, O70133, Q501J6, P38647, P49718, P49717, G5E829, P62334, Q8CGK3, Q9D0F6, Q9CQN1, P80313, P97310, P46460, Q03265, Q61881, Q91VC3, P42932, O55143, Q9JHU4, P80318, E9Q634, P63017, P43247, P60843, P80317, E9Q555 | 3.948975E-10 |
| GOTERM_MF_DIRECT | GO:0003697~single-stranded DNA binding | 14 | 2.8003590521E-09 | Q8CG48, Q8CG47, P61979, P97310, P10107, O70133, Q61990, Q61881, P49718, P49717, P62960, P60335, P43247, Q8CGK3 | 2.195481E-07 |
| GOTERM_BP_DIRECT | GO:0140588~chromatin looping | 12 | 4.0073455456E-06 | P49718, P49717, Q61656, P97310, P46460, O70133, Q501J6, Q61990, Q61881, P60843, Q91VC3, E9Q555 | 0.0011891798 |
| GOTERM_MF_DIRECT | GO:0003689~DNA clamp loader activity | 12 | 5.6377955888E-06 | P49718, P49717, Q61656, P97310, P46460, O70133, Q501J6, Q61881, P60843, Q91VC3, E9Q555, Q9D0F6 | 0.0002104777 |
| GOTERM_MF_DIRECT | GO:0140584~chromatin extrusion motor activity | 11 | 2.0144501199E-05 | P49718, P49717, Q61656, P97310, P46460, O70133, Q501J6, Q61881, P60843, Q91VC3, E9Q555 | 0.0006074342 |
| GOTERM_MF_DIRECT | GO:0140665~ATP-dependent H3-H4 histone complex chaperone activity | 11 | 2.0144501199E-05 | P49718, P49717, Q61656, P97310, P46460, O70133, Q501J6, Q61881, P60843, Q91VC3, E9Q555 | 0.0006074342 |
| GOTERM_MF_DIRECT | GO:0140849~ATP-dependent H2AZ histone chaperone activity | 11 | 2.2740443871E-05 | P49718, P49717, Q61656, P97310, P46460, O70133, Q501J6, Q61881, P60843, Q91VC3, E9Q555 | 0.0006367324 |
| GOTERM_MF_DIRECT | GO:0061775~cohesin loader activity | 11 | 2.4142601135E-05 | P49718, P49717, Q61656, P97310, P46460, O70133, Q501J6, Q61881, P60843, Q91VC3, E9Q555 | 0.0006526827 |
| GOTERM_BP_DIRECT | GO:0006279~premeiotic DNA replication | 4 | 4.3976787303E-05 | P49718, P49717, P97310, Q61881 | 0.0069600595 |
| UP_SEQ_FEATURE | MOTIF:Arginine finger | 4 | 9.2155783642E-05 | P49718, P49717, P97310, Q61881 | 0.0100910583 |
| INTERPRO | IPR018525:MCM_CS | 4 | 0.0001058831112 | P49718, P49717, P97310, Q61881 | 0.0138418104 |
| INTERPRO | IPR027925:MCM_N | 4 | 0.0001058831112 | P49718, P49717, P97310, Q61881 | 0.0138418104 |
| GOTERM_MF_DIRECT | GO:0017116~single-stranded DNA helicase activity | 5 | 0.00014936364599 | P49718, P49717, P97310, Q61881, Q9D0F6 | 0.0035485181 |
| INTERPRO | IPR001208:MCM_dom | 4 | 0.00028873248688 | P49718, P49717, P97310, Q61881 | 0.0296569512 |
| INTERPRO | IPR033762:MCM_OB | 4 | 0.00028873248688 | P49718, P49717, P97310, Q61881 | 0.0296569512 |
| GOTERM_CC_DIRECT | GO:0071162~CMG complex | 4 | 0.00034547427375 | P49718, P49717, P97310, Q61881 | 0.0036159641 |
| GOTERM_CC_DIRECT | GO:0042555~MCM complex | 4 | 0.00034547427375 | P49718, P49717, P97310, Q61881 | 0.0036159641 |
| UP_SEQ_FEATURE | DOMAIN:MCM | 4 | 0.00037268467498 | P49718, P49717, P97310, Q61881 | 0.0361779491 |
| SMART | SM00350:MCM | 4 | 0.00039442123062 | P49718, P49717, P97310, Q61881 | 0.0349062789 |
| INTERPRO | IPR041562:MCM_lid | 4 | 0.00042741513576 | P49718, P49717, P97310, Q61881 | 0.0384139353 |
| INTERPRO | IPR031327:MCM | 4 | 0.00042741513576 | P49718, P49717, P97310, Q61881 | 0.0384139353 |
| GOTERM_BP_DIRECT | GO:0000727~double-strand break repair via break-induced replication | 4 | 0.00045608460934 | P49718, P49717, P97310, Q61881 | 0.0292633747 |
| GOTERM_MF_DIRECT | GO:0016787~hydrolase activity | 13 | 0.00048636207948 | Q9DCE9, Q61656, O70133, P20060, Q501J6, Q61881, Q91VC3, P49718, P49717, Q8R146, Q8VEB4, P60843, P08775 | 0.009300192 |
| GOTERM_BP_DIRECT | GO:0006268~DNA unwinding involved in DNA replication | 4 | 0.00289566328027 | P49718, P49717, P97310, Q61881 | 0.1185224936 |
| GOTERM_MF_DIRECT | GO:1990518~single-stranded 3'-5' DNA helicase activity | 5 | 0.0028990356171 | P49718, P49717, P97310, O70133, Q61881 | 0.0398744548 |
| GOTERM_MF_DIRECT | GO:0036121~double-stranded DNA helicase activity | 5 | 0.00314255964971 | P49718, P49717, P97310, P10107, Q61881 | 0.0424787373 |

| | | | | | |
|---|---|---|---|---|---|
| GOTERM_BP_DIRECT | GO:0006270~DNA replication initiation | 4 | 0.00420040931689 | P49718, P49717, P97310, Q61881 | 0.140447489 |
| GOTERM_CC_DIRECT | GO:0005694~chromosome | 10 | 0.01138679593305 | P49718, Q8CG48, P49717, Q8CG47, P97310, Q3TEA8, Q9CXT6, E9Q7G0, Q61881, P43247 | 0.0558664675 |
| UP_KW_MOLECULAR_FUNCTION | KW-0347~Helicase | 9 | 0.01181473207071 | P49718, P49717, Q61656, P97310, O70133, Q501J6, Q61881, P60843, Q91VC3 | 0.0698143259 |
| GOTERM_BP_DIRECT | GO:0032508~DNA duplex unwinding | 5 | 0.01306712957422 | P49718, P97310, P10107, O70133, Q9D0F6 | 0.2820124146 |
| INTERPRO | IPR012340:NA-bd_OB-fold | 7 | 0.01360877553256 | P49718, P62960, P49717, P97310, P62858, P62334, Q61881 | 0.3837141023 |
| GOTERM_MF_DIRECT | GO:0043138~3'-5' DNA helicase activity | 3 | 0.01422262523218 | P49718, P97310, O70133 | 0.1343438335 |
| GOTERM_MF_DIRECT | GO:0003688~DNA replication origin binding | 3 | 0.01626901959438 | P49718, P97310, O70133 | 0.1483129228 |
| KEGG_PATHWAY | mmu03030:DNA replication | 5 | 0.01708282151808 | P49718, P49717, P97310, Q61881, Q9D0F6 | 0.1214265421 |
| GOTERM_MF_DIRECT | GO:0061749~forked DNA-dependent helicase activity | 4 | 0.02023803194619 | P49718, P49717, P97310, Q61881 | 0.1743584291 |
| GOTERM_MF_DIRECT | GO:0009378~four-way junction helicase activity | 4 | 0.02147653168368 | P49718, P49717, P97310, Q61881 | 0.1810494714 |
| GOTERM_BP_DIRECT | GO:0006338~chromatin remodeling | 18 | 0.08665180672096 | Q61656, Q9CXT6, P46460, O70133, Q7TPV4, P18052, Q501J6, Q61881, Q9D7X3, Q91VC3, P49718, P49717, P09581, P25911, P62137, P00520, P60843, E9Q555 | 0.7017410113 |
| UP_KW_BIOLOGICAL_PROCESS | KW-0235~DNA replication | 5 | 0.1251668831209 | P49718, P49717, P97310, Q61881, Q9D0F6 | 0.5635422367 |
| KEGG_PATHWAY | mmu04110:Cell cycle | 7 | 0.27131848044109 | P49718, Q9CQV8, P49717, P97310, Q61881, P00520, Q76MZ3 | 0.7219590359 |

Annotation Cluster 13 — Enrichment Score: 3.2127755876441295

| Category | Term | Count | P Value | Genes | FDR |
|---|---|---|---|---|---|
| GOTERM_BP_DIRECT | GO:0006635~fatty acid beta-oxidation | 11 | 4.9382731739E-09 | Q9DBL1, P51660, O35459, Q921H8, Q8BWT1, P32020, Q9JHI5, Q8BMS1, Q8JZN5 | 1.172346E-05 |
| GOTERM_MF_DIRECT | GO:0003988~acetyl-CoA C-acyltransferase activity | 6 | 2.1096803933E-08 | Q921H8, Q8BWT1, P32020, Q8BMS1 | 1.503627E-06 |
| INTERPRO | IPR020615:Thiolase_acyl_enz_int_AS | 5 | 3.2478277762E-06 | Q921H8, Q8BWT1, P32020 | 0.0023351882 |
| UP_SEQ_FEATURE | DOMAIN:Thiolase N-terminal | 5 | 9.4512911377E-06 | Q921H8, Q8BWT1, P32020 | 0.0017741424 |
| INTERPRO | IPR020613:Thiolase_CS | 5 | 1.1367189125E-05 | Q921H8, Q8BWT1, P32020 | 0.0040865045 |
| INTERPRO | IPR020616:Thiolase_N | 5 | 1.1367189125E-05 | Q921H8, Q8BWT1, P32020 | 0.0040865045 |
| GOTERM_MF_DIRECT | GO:0050633~acetyl-CoA C-myristoyltransferase activity | 4 | 2.2332891959E-05 | Q921H8, P32020 | 0.0006367324 |
| KEGG_PATHWAY | mmu01212:Fatty acid metabolism | 10 | 4.829220906E-05 | Q9DBL1, P51660, Q921H8, Q8BWT1, P32020, Q9QUJ7, O70503, Q8BMS1, P41216 | 0.0015876064 |
| INTERPRO | IPR016039:Thiolase-like | 5 | 6.0974126622E-05 | Q921H8, Q8BWT1, P32020 | 0.0097423105 |
| GOTERM_CC_DIRECT | GO:0005777~peroxisome | 10 | 7.887525339E-05 | P51660, O35459, Q921H8, P40142, P32020, Q9QUJ7, P38060, P20152 | 0.0012810429 |
| GOTERM_BP_DIRECT | GO:0006631~fatty acid metabolic process | 9 | 0.00012141489881 | Q9DBL1, Q8VCW8, Q921H8, Q9QUJ7, Q8BMS1, P41216, Q8VEB4, P31786 | 0.0137256652 |
| UP_KW_BIOLOGICAL_PROCESS | KW-0276~Fatty acid metabolism | 13 | 0.00012524838477 | Q9DBL1, Q8VCW8, P51660, O35459, Q921H8, Q91YR9, Q8BWT1, Q9QUJ7, Q8BMS1, P41216, Q91V12, Q8VEB4 | 0.0018787258 |
| GOTERM_MF_DIRECT | GO:0016747~acyltransferase activity, transferring groups other than amino-acyl grou | 5 | 0.00018066245577 | Q921H8, Q8BWT1, P32020 | 0.0039344268 |
| GOTERM_CC_DIRECT | GO:0005782~peroxisomal matrix | 5 | 0.001122075218 | P51660, Q921H8, P32020 | 0.0085241521 |
| GOTERM_BP_DIRECT | GO:0008206~bile acid metabolic process | 4 | 0.00218569102406 | Q921H8, P32020 | 0.0997852018 |
| UP_KW_CELLULAR_COMPONENT | KW-0576~Peroxisome | 9 | 0.00225938042613 | P51660, O35459, Q921H8, P32020, Q9QUJ7, P38060, P41216 | 0.0082843949 |
| KEGG_PATHWAY | mmu01040:Biosynthesis of unsaturated fatty acids | 6 | 0.00229488712046 | P51660, Q921H8, P32020, O70503, Q91V12 | 0.0301777656 |
| GOTERM_BP_DIRECT | GO:0000038~very long-chain fatty acid metabolic process | 4 | 0.00252525856371 | P51660, Q921H8, P41216 | 0.1110178487 |
| UP_KW_BIOLOGICAL_PROCESS | KW-0443~Lipid metabolism | 28 | 0.00256745025956 | Q8VCW8, P51660, Q9Z0J0, Q921H8, Q91YR9, P20060, P70245, Q9DBL1, Q8BLN5, Q8BWT1, Q9QUJ7, Q6ZQM8, P41216, Q91V12, Q8VEB4, Q8VCN5, Q8VDP6, O35459, P32020, P38060, Q3TCN2, O70503, P29416, P48758, Q8BMS1, E9Q555 | 0.0210064112 |
| KEGG_PATHWAY | mmu00071:Fatty acid degradation | 7 | 0.00303356593591 | Q9DBL1, Q921H8, Q8BWT1, Q9QUJ7, Q8BMS1, P41216 | 0.034688167 |
| GOTERM_MF_DIRECT | GO:0003985~acetyl-CoA C-acetyltransferase activity | 3 | 0.00348850477151 | Q921H8, Q8BWT1 | 0.0448358646 |
| UP_SEQ_FEATURE | DOMAIN:SCP2 | 3 | 0.00408326967299 | P51660, P32020 | 0.2235590146 |
| UP_SEQ_FEATURE | DOMAIN:Thiolase C-terminal | 3 | 0.00408326967299 | Q921H8, Q8BWT1 | 0.2235590146 |
| INTERPRO | IPR036527:SCP2_sterol-bd_dom_sf | 3 | 0.00447449373053 | P51660, P32020 | 0.201072562 |
| INTERPRO | IPR003033:SCP2_sterol-bd_dom | 3 | 0.00447449373053 | P51660, P32020 | 0.201072562 |
| KEGG_PATHWAY | mmu00280:Valine, leucine and isoleucine degradation | 7 | 0.00481872585506 | Q9DBL1, Q921H8, Q8BWT1, P38060, Q9JHI5, Q8BMS1 | 0.0508461285 |
| INTERPRO | IPR020610:Thiolase_AS | 3 | 0.00619114739297 | Q921H8, Q8BWT1 | 0.2344570807 |
| INTERPRO | IPR002155:Thiolase | 3 | 0.00619114739297 | Q921H8, Q8BWT1 | 0.2344570807 |
| INTERPRO | IPR020617:Thiolase_C | 3 | 0.00619114739297 | Q921H8, Q8BWT1 | 0.2344570807 |
| UP_SEQ_FEATURE | MOTIF:Microbody targeting signal | 5 | 0.00692449763009 | P51660, O35459, P32020, P38060 | 0.3032929962 |
| KEGG_PATHWAY | mmu04146:Peroxisome | 8 | 0.01043816900083 | P51660, O35459, Q921H8, P32020, Q9QUJ7, P38060, P41216 | 0.0885560789 |
| KEGG_PATHWAY | mmu03320:PPAR signaling pathway | 8 | 0.01174888450984 | O55222, Q05816, Q921H8, P32020, Q9QUJ7, P41216, P31786 | 0.0965611446 |
| PIR_SUPERFAMILY | PIRSF000429:Ac-CoA_Ac_transf | 3 | 0.01800040335885 | Q921H8, Q8BWT1 | 1 |
| UP_SEQ_FEATURE | ACT_SITE:Acyl-thioester intermediate | 3 | 0.03275703768838 | Q921H8, Q8BWT1 | 0.7101571319 |
| GOTERM_BP_DIRECT | GO:0007584~response to nutrient | 4 | 0.06502821944465 | Q921H8, P53657, P41216 | 0.6379214585 |
| UP_KW_MOLECULAR_FUNCTION | KW-0012~Acyltransferase | 6 | 0.34266261903208 | Q921H8, Q8BWT1, P32020, Q8VEB4 | 0.7971617661 |

Annotation Cluster 14 — Enrichment Score: 3.0433972177367177

| Category | Term | Count | P Value | Genes | FDR |
|---|---|---|---|---|---|
| UP_SEQ_FEATURE | DOMAIN:Nop | 3 | 0.00084465369535 | Q6DFW4, Q9D6Z1, Q8CCF0 | 0.0739916637 |
| SMART | SM00931:NOSIC | 3 | 0.00087993770349 | Q6DFW4, Q9D6Z1, Q8CCF0 | 0.0519163245 |
| INTERPRO | IPR036070:Nop_dom_sf | 3 | 0.00092708919628 | Q6DFW4, Q9D6Z1, Q8CCF0 | 0.0605979211 |
| INTERPRO | IPR042239:Nop_C | 3 | 0.00092708919628 | Q6DFW4, Q9D6Z1, Q8CCF0 | 0.0605979211 |
| INTERPRO | IPR012976:NOSIC | 3 | 0.00092708919628 | Q6DFW4, Q9D6Z1, Q8CCF0 | 0.0605979211 |
| INTERPRO | IPR002687:Nop_dom | 3 | 0.00092708919628 | Q6DFW4, Q9D6Z1, Q8CCF0 | 0.0605979211 |

Annotation Cluster 15 — Enrichment Score: 2.794504731229608

| Category | Term | Count | P Value | Genes | FDR |
|---|---|---|---|---|---|
| GOTERM_CC_DIRECT | GO:0005687~U4 snRNP | 5 | 9.0767868826E-06 | P62320, P62317, P62307, Q8CCF0, P62315 | 0.0002464707 |
| GOTERM_CC_DIRECT | GO:0071011~precatalytic spliceosome | 6 | 2.9533968176E-05 | P62320, P62317, Q3UKJ7, Q8CCF0, Q3UEB3, P62315 | 0.0006322954 |
| GOTERM_CC_DIRECT | GO:0071013~catalytic step 2 spliceosome | 10 | 3.8126400968E-05 | P62320, Q61656, P61979, Q99KP6, P62317, P62307, Q7TMK9, Q3UEB3, Q91VC3, P62315 | 0.0007807624 |
| GOTERM_CC_DIRECT | GO:0034715~pICln-Sm protein complex | 4 | 7.6216355916E-05 | P62320, P62317, P62307, P62315 | 0.0012810429 |
| GOTERM_CC_DIRECT | GO:0005689~U12-type spliceosomal complex | 5 | 0.00054162202701 | P62320, P62960, P62317, P62307, P62315 | 0.0052062036 |
| GOTERM_CC_DIRECT | GO:0034709~methylosome | 4 | 0.00074007530275 | P62320, P62317, P62307, P62315 | 0.0065768956 |
| GOTERM_CC_DIRECT | GO:0071005~U2-type precatalytic spliceosome | 6 | 0.00116288155045 | P62320, P62317, P62307, Q3UKJ7, Q8CCF0, P62315 | 0.008693924 |
| GOTERM_CC_DIRECT | GO:0046540~U4/U6 x U5 tri-snRNP complex | 5 | 0.001386376195 | P62320, P62317, P62307, Q8CCF0, P62315 | 0.0100458952 |
| GOTERM_CC_DIRECT | GO:0034719~SMN-Sm protein complex | 4 | 0.00159548999021 | P62320, P62317, P62307, P62315 | 0.0112160565 |
| GOTERM_CC_DIRECT | GO:0005685~U1 snRNP | 4 | 0.00187625589074 | P62320, P62317, P62307, P62315 | 0.0126245218 |
| GOTERM_CC_DIRECT | GO:0097526~spliceosomal tri-snRNP complex | 3 | 0.00250428713437 | P62320, Q8CCF0, P62315 | 0.0161577978 |
| GOTERM_CC_DIRECT | GO:0071007~U2-type catalytic step 2 spliceosome | 5 | 0.00337728719737 | P62320, Q99KP6, P62317, P62307, P62315 | 0.020658471 |
| SMART | SM00651:Sm | 4 | 0.00400634016827 | P62320, P62317, P62307, P62315 | 0.1185974106 |
| INTERPRO | IPR001163:Sm_dom_euk/arc | 4 | 0.00432318646596 | P62320, P62317, P62307, P62315 | 0.201072562 |
| GOTERM_BP_DIRECT | GO:0000387~spliceosomal snRNP assembly | 4 | 0.00641929928032 | P62320, P62317, P62307, P62315 | 0.1881409443 |
| INTERPRO | IPR010920:LSM_dom_sf | 4 | 0.00660654537582 | P62320, P62317, P62307, P62315 | 0.2435951859 |
| GOTERM_CC_DIRECT | GO:0005684~U2-type spliceosomal complex | 4 | 0.00706463830739 | P62320, P62317, P62307, P62315 | 0.0396124362 |
| GOTERM_MF_DIRECT | GO:1990446~U1 snRNP binding | 3 | 0.00728212507669 | P62320, P62317, P62315 | 0.0782080282 |
| UP_SEQ_FEATURE | DOMAIN:Sm | 4 | 0.01036291874938 | P62320, P62317, P62307, P62315 | 0.368023655 |
| INTERPRO | IPR047575:Sm | 4 | 0.01175729031283 | P62320, P62317, P62307, P62315 | 0.3505373999 |
| GOTERM_CC_DIRECT | GO:0005682~U5 snRNP | 3 | 0.01836675342321 | P62320, P62317, P62315 | 0.0839877754 |
| GOTERM_CC_DIRECT | GO:0005686~U2 snRNP | 3 | 0.03078628443311 | P62320, P62317, P62315 | 0.1250029308 |

Annotation Cluster 16 — Enrichment Score: 2.7476650903875375

| Category | Term | Count | P Value | Genes | FDR |
|---|---|---|---|---|---|
| UP_KW_DOMAIN | KW-0676~Redox-active center | 9 | 1.2970204742E-06 | P10639, Q9JLT4, Q80Y14, Q9JMH6, P09103, Q8VBT0, Q9ESY9, Q8CDN6, Q8R180 | 1.426723E-05 |
| UP_SEQ_FEATURE | DISULFID:Redox-active | 9 | 2.4247694936E-06 | P10639, Q9JLT4, Q9JMH6, P09103, Q8VBT0, Q9ESY9, Q8CDN6, Q8R180, P07742 | 0.0005310245 |
| GOTERM_MF_DIRECT | GO:0015035~protein-disulfide reductase activity | 5 | 0.00018066245577 | P10639, P09103, Q8VBT0, Q8CDN6, Q8R180 | 0.0039344268 |
| INTERPRO | IPR017937:Thioredoxin_CS | 4 | 0.00368855500674 | P10639, P09103, Q8VBT0, Q8CDN6 | 0.1894336464 |
| GOTERM_BP_DIRECT | GO:0045454~cell redox homeostasis | 4 | 0.01001658675106 | P10639, Q9JLT4, Q9JMH6, Q8R180 | 0.2499104562 |
| INTERPRO | IPR036249:Thioredoxin-like_sf | 8 | 0.01036113489548 | P57759, Q8VCH8, P10639, Q80Y14, P09103, Q8VBT0, P48774, Q8CDN6 | 0.3172024363 |
| INTERPRO | IPR013766:Thioredoxin_domain | 4 | 0.03802887079225 | P10639, P09103, Q8VBT0, Q8CDN6 | 0.6433590141 |
| UP_SEQ_FEATURE | DOMAIN:Thioredoxin | 4 | 0.04257835413549 | P10639, P09103, Q8VBT0, Q8CDN6 | 0.7992565333 |
| UP_SEQ_FEATURE | ACT_SITE:Nucleophile | 7 | 0.52996029992866 | P10639, Q9WVJ3, P09103, Q8VBT0, Q3THK7, Q60692, Q8R180 | 0.9909502262 |

Annotation Cluster 17 — Enrichment Score: 2.660332277557125

| Category | Term | Count | P Value | Genes | FDR |
|---|---|---|---|---|---|
| GOTERM_BP_DIRECT | GO:0006177~GMP biosynthetic process | 4 | 0.00017933911234 | Q9CWJ9, P24547, Q3THK7, Q9DCL9 | 0.0170300421 |
| UP_KW_BIOLOGICAL_PROCESS | KW-0658~Purine biosynthesis | 4 | 0.00225266358818 | Q9CWJ9, P24547, Q3THK7, Q9DCL9 | 0.020893617 |
| GOTERM_BP_DIRECT | GO:0097294~'de novo' XMP biosynthetic process | 3 | 0.00459344349612 | Q9CWJ9, P24547, Q9DCL9 | 0.1476179791 |
| KEGG_PATHWAY | mmu00230:Purine metabolism | 10 | 0.01230725378618 | Q9WV85, Q9CWJ9, P23492, P00493, Q9WTP7, P24547, Q3THK7, Q9D892, P07742, Q9DCL9 | 0.0980850832 |

Annotation Cluster 18 — Enrichment Score: 2.6172431801905596

| Category | Term | Count | P Value | Genes | FDR |
|---|---|---|---|---|---|
| GOTERM_BP_DIRECT | GO:0032263~GMP salvage | 4 | 0.00025369771128 | P23492, P00493, P24547, Q3THK7 | 0.0192673144 |
| KEGG_PATHWAY | mmu01232:Nucleotide metabolism | 9 | 0.00225064720096 | Q9WV85, P23492, P00493, Q3U5Q7, Q9WTP7, P24547, Q3THK7, Q9D892, P07742 | 0.0301777656 |
| KEGG_PATHWAY | mmu00983:Drug metabolism - other enzymes | 9 | 0.00483328217848 | Q9WV85, P00493, P13439, P48774, P24547, Q3THK7, Q6ZQM8, Q9D892, P07742 | 0.0508461285 |
| KEGG_PATHWAY | mmu00230:Purine metabolism | 10 | 0.01230725378618 | Q9WV85, Q9CWJ9, P23492, P00493, Q9WTP7, P24547, Q3THK7, Q9D892, P07742, Q9DCL9 | 0.0980850832 |

Annotation Cluster 19 — Enrichment Score: 2.486968194134315

| Category | Term | Count | P Value | Genes | FDR |
|---|---|---|---|---|---|
| KEGG_PATHWAY | mmu04810:Regulation of actin cytoskeleton | 19 | 7.955775429E-05 | P05555, Q8BH43, Q7TPR4, P26041, P11835, P18760, P59999, Q05144, Q7TMB8, P63085, Q9JM76, P57780, P09055, Q8K1X4, Q9R0C8, P08556, P62137, P62962 | 0.0017649765 |
| KEGG_PATHWAY | mmu04670:Leukocyte transendothelial migration | 10 | 0.00583443333599 | P05555, Q7TPR4, P26041, P57780, P09055, O70145, P11835, Q9R0C8, P08752, Q05144 | 0.0568317025 |
| KEGG_PATHWAY | mmu04015:Rap1 signaling pathway | 11 | 0.07454426208388 | P05555, P63085, P09055, P09581, P11835, Q9R0C8, P08556, P62962, P08752, Q05144 | 0.3380196712 |

Annotation Cluster 20 — Enrichment Score: 2.4631977642938847

| Category | Term | Count | P Value | Genes | FDR |
|---|---|---|---|---|---|
| GOTERM_BP_DIRECT | GO:0140588~chromatin looping | 12 | 4.0073455456E-06 | P49718, P49717, Q61656, P97310, P46460, O70133, Q501J6, Q61990, Q61881, P60843, Q91VC3, E9Q555 | 0.0011891798 |
| GOTERM_MF_DIRECT | GO:0003689~DNA clamp loader activity | 12 | 5.6377955888E-06 | P49718, P49717, Q61656, P97310, P46460, O70133, Q501J6, Q61881, P60843, Q91VC3, E9Q555, Q9D0F6 | 0.0002104777 |
| GOTERM_MF_DIRECT | GO:0140584~chromatin extrusion motor activity | 11 | 2.0144501199E-05 | P49718, P49717, Q61656, P97310, P46460, O70133, Q501J6, Q61881, P60843, Q91VC3, E9Q555 | 0.0006074342 |
| GOTERM_MF_DIRECT | GO:0140665~ATP-dependent H3-H4 histone complex chaperone activity | 11 | 2.0144501199E-05 | P49718, P49717, Q61656, P97310, P46460, O70133, Q501J6, Q61881, P60843, Q91VC3, E9Q555 | 0.0006074342 |
| GOTERM_MF_DIRECT | GO:0140849~ATP-dependent H2AZ histone chaperone activity | 11 | 2.2740443871E-05 | P49718, P49717, Q61656, P97310, P46460, O70133, Q501J6, Q61881, P60843, Q91VC3, E9Q555 | 0.0006367324 |

| | | | | | |
|---|---|---|---|---|---|
| GOTERM_MF_DIRECT | GO:0061775~cohesin loader activity | 11 | 2.4142601135E-05 | P49718, P49717, Q61656, P97310, P46460, O70133, Q501J6, Q61881, P60843, Q91VC3, E9Q555 | 0.0006526827 |
| GOTERM_MF_DIRECT | GO:0043021~ribonucleoprotein complex binding | 7 | 8.514254752E-05 | Q61656, P61979, P67871, Q501J6, Q8CCF0, P62137, Q91VC3 | 0.0021532825 |
| GOTERM_MF_DIRECT | GO:0016787~hydrolase activity | 13 | 0.00048636207948 | Q9DCE9, Q61656, O70133, P20060, Q501J6, Q61881, Q91VC3, P49718, P49717, Q8R146, Q8VEB4, P60843, P08775 | 0.009300192 |
| GOTERM_BP_DIRECT | GO:0000381~regulation of alternative mRNA splicing, via spliceosome | 6 | 0.00178414863169 | Q61656, Q62093, Q501J6, Q3UKJ7, Q3UEB3, Q91VC3 | 0.0864401806 |
| GOTERM_BP_DIRECT | GO:0000380~alternative mRNA splicing, via spliceosome | 4 | 0.00581125371636 | Q61656, O70133, Q501J6, Q3UEB3 | 0.1772459373 |
| UP_SEQ_FEATURE | DOMAIN:DEAD-box RNA helicase Q | 4 | 0.01146349421322 | Q61656, Q501J6, P60843, Q91VC3 | 0.3963955631 |
| UP_KW_MOLECULAR_FUNCTION | KW-0347~Helicase | 9 | 0.01181473207071 | P49718, P49717, Q61656, P97310, O70133, Q501J6, Q61881, P60843, Q91VC3 | 0.0698143259 |
| GOTERM_MF_DIRECT | GO:0003724~RNA helicase activity | 5 | 0.0138062497185 | Q61656, O70133, Q501J6, P60843, Q91VC3 | 0.1336308615 |
| INTERPRO | IPR000629:RNA-helicase_DEAD-box_CS | 4 | 0.01569668012004 | Q61656, Q501J6, P60843, Q91VC3 | 0.4030683217 |
| UP_SEQ_FEATURE | MOTIF:DEAD box | 4 | 0.01651850950622 | Q61656, Q501J6, P60843, Q91VC3 | 0.5462800932 |
| INTERPRO | IPR014014:RNA_helicase_DEAD_Q_motif | 4 | 0.02372100907282 | Q61656, Q501J6, P60843, Q91VC3 | 0.5501743717 |
| UP_SEQ_FEATURE | MOTIF:Q motif | 4 | 0.02982041926909 | Q61656, Q501J6, P60843, Q91VC3 | 0.7101571319 |
| INTERPRO | IPR011545:DEAD/DEAH_box_helicase_dom | 5 | 0.04056964677409 | Q61656, O70133, Q501J6, P60843, Q91VC3 | 0.6783622333 |
| UP_SEQ_FEATURE | DOMAIN:Helicase C-terminal | 5 | 0.11903308304295 | Q61656, O70133, Q501J6, P60843, Q91VC3 | 0.9909502262 |
| SMART | SM00490:HELICc | 5 | 0.1255986510267 | Q61656, O70133, Q501J6, P60843, Q91VC3 | 0.9943820225 |
| SMART | SM00487:DEXDc | 5 | 0.13163496089115 | Q61656, O70133, Q501J6, P60843, Q91VC3 | 0.9943820225 |
| INTERPRO | IPR001650:Helicase_C-like | 5 | 0.1345543882944 | Q61656, O70133, Q501J6, P60843, Q91VC3 | 0.9862825789 |
| UP_SEQ_FEATURE | DOMAIN:Helicase ATP-binding | 5 | 0.13667356111813 | Q61656, O70133, Q501J6, P60843, Q91VC3 | 0.9909502262 |
| INTERPRO | IPR014001:Helicase_ATP-bd | 5 | 0.14413947566537 | Q61656, O70133, Q501J6, P60843, Q91VC3 | 0.9862825789 |
| GOTERM_MF_DIRECT | GO:0003676~nucleic acid binding | 7 | 0.16770804297942 | P62960, Q8BL97, Q61656, P56960, Q3U0V1, Q501J6, Q91VC3 | 0.6718819814 |
| Annotation Cluster 21 | Enrichment Score: 2.417150664128563 | | | | |
| Category | Term | Count | P Value | Genes | FDR |
| UP_KW_BIOLOGICAL_PROCESS | KW-0648~Protein biosynthesis | 13 | 3.86751513E-05 | Q9Z0N1, Q8C0C7, Q99L45, P57776, Q9ER72, Q8K0D5, Q8R050, P23116, P26638, Q8BU30, P60229, Q8BGQ7, P60843 | 0.0008413633 |
| UP_KW_MOLECULAR_FUNCTION | KW-0436~Ligase | 12 | 0.00023575410257 | Q8VCW8, Q8C0C7, Q9Z2I9, Q9QUJ7, P26638, Q8BU30, Q3THK7, Q9ER72, P41216, Q9D7H3, Q8BGQ7, Q9DCL9 | 0.0031588941 |
| GOTERM_MF_DIRECT | GO:0000049~tRNA binding | 6 | 0.00439370132513 | Q9Z0N1, Q8C0C7, P26638, Q8BU30, Q9ER72, Q8BGQ7 | 0.0529947975 |
| UP_KW_MOLECULAR_FUNCTION | KW-0030~Aminoacyl-tRNA synthetase | 5 | 0.00985256668031 | Q8C0C7, P26638, Q8BU30, Q9ER72, Q8BGQ7 | 0.0640416834 |
| INTERPRO | IPR045864:aa-tRNA-synth_II/BPL/LPL | 3 | 0.07180770516355 | Q8C0C7, P26638, Q8BGQ7 | 0.9862825789 |
| KEGG_PATHWAY | mmu00970:Aminoacyl-tRNA biosynthesis | 5 | 0.11082909149998 | Q8C0C7, P26638, Q8BU30, Q9ER72, Q8BGQ7 | 0.4105359305 |
| Annotation Cluster 22 | Enrichment Score: 2.391986299322439 | | | | |
| Category | Term | Count | P Value | Genes | FDR |
| GOTERM_CC_DIRECT | GO:0032040~small-subunit processome | 7 | 0.00109581069534 | P62245, P56960, Q6DFW4, Q9D6Z1, Q5SSI6, P62858 | 0.0084610957 |
| GOTERM_BP_DIRECT | GO:0042274~ribosomal small subunit biogenesis | 7 | 0.00116187969802 | P62245, P56960, Q6DFW4, Q9D6Z1, Q5SSI6, P62858 | 0.062688691 |
| GOTERM_CC_DIRECT | GO:0022627~cytosolic small ribosomal subunit | 4 | 0.05237745639055 | P62908, P62245, P62858 | 0.1800714012 |
| Annotation Cluster 23 | Enrichment Score: 2.2631150785913117 | | | | |
| Category | Term | Count | P Value | Genes | FDR |
| UP_KW_DOMAIN | KW-0728~SH3 domain | 11 | 0.00207426797115 | Q91ZR2, Q91VH2, O70145, Q8BI84, Q3UND0, Q9R0C8, P25911, Q61792, E9Q634, P00520, Q62422 | 0.0114084738 |
| INTERPRO | IPR036028:SH3-like_dom_sf | 11 | 0.00481667978835 | Q91ZR2, Q91VH2, O70145, Q8BI84, Q3UND0, Q9R0C8, P25911, Q61792, E9Q634, P00520, Q62422 | 0.2098904708 |
| INTERPRO | IPR001452:SH3_domain | 11 | 0.00619566694387 | Q91ZR2, Q91VH2, O70145, Q8BI84, Q3UND0, Q9R0C8, P25911, Q61792, E9Q634, P00520, Q62422 | 0.2344570807 |
| SMART | SM00326:SH3 | 10 | 0.00760580120429 | Q91ZR2, Q91VH2, O70145, Q3UND0, Q9R0C8, P25911, Q61792, E9Q634, P00520, Q62422 | 0.1923181162 |
| UP_SEQ_FEATURE | DOMAIN:SH3 | 10 | 0.01027020083969 | Q91ZR2, Q91VH2, O70145, Q8BI84, Q3UND0, P25911, Q61792, E9Q634, P00520, Q62422 | 0.368023655 |
| Annotation Cluster 24 | Enrichment Score: 2.2613742383779885 | | | | |
| Category | Term | Count | P Value | Genes | FDR |
| UP_KW_MOLECULAR_FUNCTION | KW-0687~Ribonucleoprotein | 23 | 1.5763258102E-06 | P62908, P51410, Q9EQK5, Q9CPR5, P61979, Q9D7S7, Q9D6Z1, Q7TMK9, Q61990, Q3UEB3, P62320, P62245, Q6DFW4, P60335, P62317, P62307, P62858, Q924T2, Q9CQF0, Q8CCF0, P41105, P62315 | 0.0001024612 |
| GOTERM_BP_DIRECT | GO:0006412~translation | 13 | 2.9752332864E-05 | P62908, P51410, Q9CPR5, Q8R050, P62245, Q9Z0N1, Q9D7S7, P62858, Q924T2, Q9CQF0, Q9ER72, P41105 | 0.0050451456 |
| GOTERM_BP_DIRECT | GO:0002181~cytoplasmic translation | 8 | 0.00102999565942 | P62908, P51410, P62245, Q9D7S7, P26638, P62858, P41105 | 0.0582192785 |
| GOTERM_CC_DIRECT | GO:0005840~ribosome | 9 | 0.00154748134323 | P62908, P51410, P62245, Q9D7S7, P35564, Q9CQF0, P62858, P41105 | 0.0110433896 |
| GOTERM_MF_DIRECT | GO:0003735~structural constituent of ribosome | 10 | 0.00421455650825 | P62908, P51410, Q9CPR5, P62245, Q9D7S7, Q924T2, Q9CQF0, P62858, P41105 | 0.0516283172 |
| GOTERM_BP_DIRECT | GO:0140236~translation at presynapse | 5 | 0.01010589881833 | P51410, P62245, P62858, P41105 | 0.2499104562 |
| GOTERM_BP_DIRECT | GO:0140242~translation at postsynapse | 5 | 0.01065928828997 | P51410, P62245, P62858, P41105 | 0.2556075798 |
| UP_KW_MOLECULAR_FUNCTION | KW-0689~Ribosomal protein | 10 | 0.03192615428345 | P62908, P51410, Q9CPR5, P62245, Q9D7S7, Q924T2, Q9CQF0, P62858, P41105 | 0.1482285735 |
| GOTERM_CC_DIRECT | GO:0022627~cytosolic small ribosomal subunit | 4 | 0.05237745639055 | P62908, P62245, P62858 | 0.1800714012 |
| GOTERM_CC_DIRECT | GO:0022626~cytosolic ribosome | 5 | 0.07089594812559 | P62908, P51410, Q8R050, P62245, P41105 | 0.230289597 |
| KEGG_PATHWAY | mmu03010:Ribosome | 10 | 0.30984969340841 | P62908, P51410, Q9CPR5, P62245, Q9D7S7, Q924T2, Q9CQF0, P62858, P41105 | 0.7803282278 |
| KEGG_PATHWAY | mmu05171:Coronavirus disease - COVID-19 | 10 | 0.58581809144776 | P62908, P51410, Q64339, P62245, Q9D7S7, P63085, Q9WVL2, P62858, P41105 | 0.8855218855 |
| Annotation Cluster 25 | Enrichment Score: 2.2055366246284573 | | | | |
| Category | Term | Count | P Value | Genes | FDR |
| UP_KW_LIGAND | KW-0285~Flavoprotein | 11 | 0.00209811669065 | Q9DBL1, Q9Z0X1, Q9JLT4, P51175, Q9JMH6, Q9JI75, Q9JHI5, Q8R180, Q8JZN5, Q91YT0 | 0.0141622877 |
| UP_KW_LIGAND | KW-0274~FAD | 10 | 0.00536532878456 | Q9DBL1, Q9Z0X1, Q9JLT4, P51175, Q9JMH6, Q9JI75, Q9JHI5, Q8R180, Q8JZN5 | 0.0289727754 |
| GOTERM_MF_DIRECT | GO:0071949~FAD binding | 4 | 0.02147653168368 | Q9Z0X1, Q9JMH6, Q9JI75, Q8R180 | 0.1810494714 |
| Annotation Cluster 26 | Enrichment Score: 2.159025131170111 | | | | |
| Category | Term | Count | P Value | Genes | FDR |
| UP_SEQ_FEATURE | DOMAIN:KH 3 | 4 | 0.00120666058037 | P61979, Q3U0V1, P60335, Q61990 | 0.0932677649 |
| GOTERM_MF_DIRECT | GO:0003730~mRNA 3'-UTR binding | 7 | 0.00343488251913 | Q921F2, Q61656, P61979, Q3U0V1, Q501J6, P60335, Q7TMK9 | 0.0448358646 |
| SMART | SM00322:KH | 5 | 0.0040202512075 | P61979, Q3U0V1, P60335, Q61990, Q64213 | 0.1185974106 |
| INTERPRO | IPR004087:KH_dom | 5 | 0.00444143964649 | P61979, Q3U0V1, P60335, Q61990, Q64213 | 0.201072562 |
| UP_SEQ_FEATURE | DOMAIN:KH 1 | 4 | 0.00580583148685 | P61979, Q3U0V1, P60335, Q61990 | 0.2825504657 |
| UP_SEQ_FEATURE | DOMAIN:KH 2 | 4 | 0.00580583148685 | P61979, Q3U0V1, P60335, Q61990 | 0.2825504657 |
| INTERPRO | IPR036612:KH_dom_type_1_sf | 5 | 0.01343145037253 | P61979, Q3U0V1, P60335, Q61990, Q64213 | 0.3837141023 |
| INTERPRO | IPR004088:KH_dom_type_1 | 4 | 0.0274447514299 | P61979, Q3U0V1, P60335, Q61990 | 0.5979629175 |
| UP_SEQ_FEATURE | DOMAIN:K Homology | 3 | 0.04028921301399 | P61979, Q3U0V1, Q61990 | 0.7901496403 |
| Annotation Cluster 27 | Enrichment Score: 2.1564529809244193 | | | | |
| Category | Term | Count | P Value | Genes | FDR |
| GOTERM_CC_DIRECT | GO:0070937~CRD-mediated mRNA stability complex | 3 | 0.00250428713437 | P62960, O70133, Q7TMK9 | 0.0161577978 |
| GOTERM_BP_DIRECT | GO:1900152~negative regulation of nuclear-transcribed mRNA catabolic process, de | 3 | 0.00585461617691 | P62960, O70133, Q7TMK9 | 0.1772459373 |
| GOTERM_BP_DIRECT | GO:0070934~CRD-mediated mRNA stabilization | 3 | 0.00879027125586 | P62960, O70133, Q7TMK9 | 0.2293198238 |
| GOTERM_BP_DIRECT | GO:2000767~positive regulation of cytoplasmic translation | 3 | 0.01836552080711 | P62960, O70133, Q7TMK9 | 0.3182463241 |
| Annotation Cluster 28 | Enrichment Score: 2.153441736200704 | | | | |
| Category | Term | Count | P Value | Genes | FDR |
| UP_KW_BIOLOGICAL_PROCESS | KW-0249~Electron transport | 13 | 3.2545255013E-06 | Q9D0M3, P10639, Q62425, Q9ERS2, Q7TMF3, Q9CQX2, Q91WD5, Q8R180, Q8VBT0, Q9DC70, Q8CDN6, Q8R1I1, Q91YT0 | 9.763577E-05 |
| KEGG_PATHWAY | mmu05415:Diabetic cardiomyopathy | 18 | 0.00011187152033 | Q9D0M3, Q62425, O70145, P48771, Q9ERS2, P58389, Q03265, Q7TMF3, Q91WD5, Q05144, P17665, O55143, Q00612, Q9DC70, P62137, Q8R1I1, P56391, Q91YT0 | 0.0022632469 |
| UP_KW_BIOLOGICAL_PROCESS | KW-0679~Respiratory chain | 8 | 0.00041268523348 | Q9D0M3, Q62425, Q9ERS2, Q7TMF3, Q9DC70, Q91WD5, Q8R1I1, Q91YT0 | 0.005305953 |
| KEGG_PATHWAY | mmu00190:Oxidative phosphorylation | 13 | 0.0005471428077 | Q9D0M3, Q62425, Q8BVE3, P48771, Q9ERS2, Q03265, Q7TMF3, Q91WD5, P17665, Q9DC70, Q8R1I1, P56391, Q91YT0 | 0.0084646211 |
| GOTERM_CC_DIRECT | GO:0045271~respiratory chain complex I | 6 | 0.0007814947495 | Q62425, Q9ERS2, Q7TMF3, Q9DC70, Q91WD5, Q91YT0 | 0.0066624362 |
| GOTERM_BP_DIRECT | GO:0042776~proton motive force-driven mitochondrial ATP synthesis | 6 | 0.00155504256708 | Q9ERS2, Q03265, Q7TMF3, Q9DC70, Q91WD5, Q91YT0 | 0.0769098136 |
| GOTERM_BP_DIRECT | GO:0032981~mitochondrial respiratory chain complex I assembly | 6 | 0.00203704834335 | Q9Z0X1, Q9ERS2, Q7TMF3, Q9DC70, Q91WD5, Q8JZN5 | 0.0949339139 |
| KEGG_PATHWAY | mmu04714:Thermogenesis | 16 | 0.00260442250111 | Q9D0M3, Q62425, P48771, Q9ERS2, Q03265, Q7TMF3, Q91WD5, P17665, Q9QUJ7, Q9DC70, P08556, P41216, Q8R1I1, P56391, Q91YT0 | 0.0311346872 |
| KEGG_PATHWAY | mmu04932:Non-alcoholic fatty liver disease | 12 | 0.00547712334262 | Q9D0M3, Q62425, P48771, Q9ERS2, Q7TMF3, Q9DC70, P53657, Q91WD5, P56391, Q8R1I1, P17665, Q91YT0 | 0.0554032092 |
| GOTERM_BP_DIRECT | GO:0042775~mitochondrial ATP synthesis coupled electron transport | 3 | 0.01417026039962 | Q7TMF3, Q91WD5, Q91YT0 | 0.2850864253 |
| GOTERM_BP_DIRECT | GO:0009060~aerobic respiration | 5 | 0.01579614937339 | Q9ERS2, Q7TMF3, Q9DC70, Q91WD5, Q91YT0 | 0.2937486193 |
| UP_KW_LIGAND | KW-0830~Ubiquinone | 4 | 0.01736400561535 | Q7TMF3, Q9DC70, Q91WD5, Q91YT0 | 0.058603519 |
| GOTERM_MF_DIRECT | GO:0051539~4 iron, 4 sulfur cluster binding | 4 | 0.02023803194619 | P28271, Q9DC70, Q91WD5, Q91YT0 | 0.1743584291 |
| GOTERM_MF_DIRECT | GO:0008137~NADH dehydrogenase (ubiquinone) activity | 3 | 0.02309883154104 | Q9DC70, Q91WD5, Q91YT0 | 0.1906261466 |
| GOTERM_BP_DIRECT | GO:1902600~proton transmembrane transport | 6 | 0.02742462306386 | Q9D0M3, Q03265, Q9DC70, Q8K3C0, Q91WD5, Q91YT0 | 0.3945821524 |
| GOTERM_BP_DIRECT | GO:0006120~mitochondrial electron transport, NADH to ubiquinone | 3 | 0.04887680461073 | Q9DC70, Q91WD5, Q91YT0 | 0.5606242819 |
| UP_KW_MOLECULAR_FUNCTION | KW-1278~Translocase | 6 | 0.07120450044328 | Q9D0M3, O55143, G5E829, Q9DC70, Q91WD5, Q91YT0 | 0.2722525017 |
| UP_KW_LIGAND | KW-0004~4Fe-4S | 4 | 0.08452172653744 | P28271, Q9DC70, Q91WD5, Q91YT0 | 0.2074624197 |
| UP_KW_LIGAND | KW-0411~Iron-sulfur | 5 | 0.11568075328343 | Q80Y14, P28271, Q9DC70, Q91WD5, Q91YT0 | 0.2602816949 |
| KEGG_PATHWAY | mmu04723:Retrograde endocannabinoid signaling | 8 | 0.13393124858824 | Q62425, P63085, Q9ERS2, Q7TMF3, Q9DC70, Q91WD5, P08752, Q91YT0 | 0.4825194298 |
| UP_KW_LIGAND | KW-0408~Iron | 8 | 0.90920849160961 | Q9D0M3, Q9CXT6, Q80Y14, P28271, Q9CQX2, Q9DC70, Q91WD5, Q91YT0 | 0.9310344828 |
| Annotation Cluster 29 | Enrichment Score: 2.152684861825073 | | | | |
| Category | Term | Count | P Value | Genes | FDR |
| GOTERM_MF_DIRECT | GO:0008641~ubiquitin-like modifier activating enzyme activity | 3 | 0.00169053352331 | Q9Z1F9, Q8VBW6, Q9D906 | 0.0270485364 |
| UP_SEQ_FEATURE | DOMAIN:THIF-type NAD/FAD binding fold | 3 | 0.00275289837639 | Q9Z1F9, Q8VBW6, Q9D906 | 0.1903846561 |
| INTERPRO | IPR035985:Ubiquitin-activating_enz | 3 | 0.01547345304337 | Q9Z1F9, Q8VBW6, Q9D906 | 0.4030683217 |
| INTERPRO | IPR000594:ThiF_NAD_FAD-bd | 3 | 0.01547345304337 | Q9Z1F9, Q8VBW6, Q9D906 | 0.4030683217 |
| INTERPRO | IPR045886:ThiF/MoeB/HesA | 3 | 0.01547345304337 | Q9Z1F9, Q8VBW6, Q9D906 | 0.4030683217 |
| Annotation Cluster 30 | Enrichment Score: 2.1511749226139703 | | | | |
| Category | Term | Count | P Value | Genes | FDR |
| GOTERM_BP_DIRECT | GO:0045087~innate immune response | 18 | 0.00056423736531 | Q9DCE9, Q60875, Q64339, P20491, P26151, P10810, Q99J93, P10107, O70133, Q61990, Q8VC04, Q8C2Q3, P16110, Q2EMV9, P09581, Q9D906, P25911, Q8BG07 | 0.0343461412 |
| UP_KW_BIOLOGICAL_PROCESS | KW-0399~Innate immunity | 18 | 0.00394797537024 | Q9DCE9, Q60875, Q9CQW9, P20491, P26151, P10810, Q99J93, P10107, O70133, Q9Z0E6, Q61990, Q8VC04, Q8C2Q3, P16110, Q2EMV9, P09581, P25911, Q8BG07 | 0.0296098153 |
| GOTERM_BP_DIRECT | GO:0032760~positive regulation of tumor necrosis factor production | 7 | 0.00563694140137 | Q60875, P26151, P10810, P20491, O70133, P09581, Q8VC04 | 0.1760802485 |

| | | | | | |
|---|---|---|---|---|---|
| UP_KW_BIOLOGICAL_PROCESS | KW-0391~Immunity | 24 | 0.19788847963169 | Q9DCE9, Q9EQH2, Q60875, Q9CQW9, P20491, P26151, P10810, Q99J93, P10107, O70133, Q501J6, Q9Z0E6, Q9ESY9, Q61990, Q8VC04, Q8C2Q3, P16110, Q2EMV9, P09581, P01900, P01897, P25911, E9Q555, Q8BG07 | 0.6596282654 |
| Annotation Cluster 31 | Enrichment Score: 2.0642876743395173 | | | | |
| Category | Term | Count | P Value | Genes | FDR |
| GOTERM_BP_DIRECT | GO:0006979~response to oxidative stress | 8 | 0.00411862393358 | Q9Z0X1, Q9JLT4, Q00612, Q9JMH6, Q7TMF3, Q91WD5, P00520, Q8R180 | 0.1396801888 |
| INTERPRO | IPR036188:FAD/NAD-bd_sf | 6 | 0.00549910899094 | P50396, Q9Z0X1, Q9JLT4, P51175, Q9JMH6 | 0.2325799626 |
| UP_SEQ_FEATURE | DOMAIN:FAD/NAD(P)-binding | 3 | 0.00947633438952 | Q9Z0X1, Q9JLT4, Q9JMH6 | 0.368023655 |
| INTERPRO | IPR016156:FAD/NAD-linked_Rdtase_dimer_sf | 3 | 0.01036753442644 | Q9Z0X1, Q9JLT4, Q9JMH6 | 0.3172024363 |
| INTERPRO | IPR023753:FAD/NAD-binding_dom | 3 | 0.02143884077013 | Q9Z0X1, Q9JLT4, Q9JMH6 | 0.5053943119 |
| Annotation Cluster 32 | Enrichment Score: 2.0416830594670943 | | | | |
| Category | Term | Count | P Value | Genes | FDR |
| UP_KW_MOLECULAR_FUNCTION | KW-0413~Isomerase | 11 | 0.00270475916253 | P51660, O35459, P26883, P30416, Q8BLN5, P09103, P58389, Q8VBT0, P70245, P17751, P17742 | 0.0251156208 |
| GOTERM_MF_DIRECT | GO:0140839~RNA polymerase II CTD heptapeptide repeat P3 isomerase activity | 4 | 0.00584215661938 | P26883, P30416, P58389, P17742 | 0.0645105745 |
| GOTERM_MF_DIRECT | GO:0140840~RNA polymerase II CTD heptapeptide repeat P6 isomerase activity | 4 | 0.00584215661938 | P26883, P30416, P58389, P17742 | 0.0645105745 |
| GOTERM_MF_DIRECT | GO:0003755~peptidyl-prolyl cis-trans isomerase activity | 4 | 0.01568936356152 | P26883, P30416, P58389, P17742 | 0.1447113063 |
| UP_KW_MOLECULAR_FUNCTION | KW-0697~Rotamase | 4 | 0.04272719512998 | P26883, P30416, P58389, P17742 | 0.1851511789 |
| Annotation Cluster 33 | Enrichment Score: 1.955590366453885 | | | | |
| Category | Term | Count | P Value | Genes | FDR |
| GOTERM_MF_DIRECT | GO:0016491~oxidoreductase activity | 11 | 0.00052847372987 | P51660, Q9Z0X1, P48771, Q9JMH6, Q8R127, Q8BVI4, Q9JI75, P48758, O70503, Q8BMS1, Q8R180 | 0.009864843 |
| INTERPRO | IPR020904:Sc_DH/Rdtase_CS | 4 | 0.0274447514299 | P51660, Q8BVI4, P48758, O70503 | 0.5979629175 |
| INTERPRO | IPR002347:SDR_fam | 4 | 0.09370142492905 | P51660, Q8BVI4, P48758, O70503 | 0.9862825789 |
| Annotation Cluster 34 | Enrichment Score: 1.9390047987405785 | | | | |
| Category | Term | Count | P Value | Genes | FDR |
| SMART | SM00088:PINT | 4 | 0.00197917070802 | P23116, O88543, P60229, P14685 | 0.0875783038 |
| UP_SEQ_FEATURE | DOMAIN:PCI | 4 | 0.00744294619914 | P23116, O88543, P60229, P14685 | 0.3154848808 |
| INTERPRO | IPR000717:PCI_dom | 4 | 0.00749906707879 | P23116, O88543, P60229, P14685 | 0.26301606 |
| INTERPRO | IPR036390:WH_DNA-bd_sf | 7 | 0.15876158725873 | Q3TEA8, Q8C0C7, Q69ZK0, O88543, P43277, P60229, P14685 | 0.9862825789 |
| Annotation Cluster 35 | Enrichment Score: 1.9009365450809925 | | | | |
| Category | Term | Count | P Value | Genes | FDR |
| UP_KW_CELLULAR_COMPONENT | KW-0647~Proteasome | 7 | 0.00056424511728 | O70435, O35226, P62334, Q60692, Q8CDN6, P14685, P97371 | 0.0037240178 |
| GOTERM_CC_DIRECT | GO:0000502~proteasome complex | 6 | 0.0007814947495 | O70435, O35226, P62334, Q60692, Q8CDN6, P14685 | 0.0066624362 |
| KEGG_PATHWAY | mmu03050:Proteasome | 6 | 0.00944461130903 | O70435, O35226, P62334, Q60692, P14685, P97371 | 0.0860300618 |
| GOTERM_CC_DIRECT | GO:0022624~proteasome accessory complex | 3 | 0.02063663863165 | O35226, P62334, P14685 | 0.0916967622 |
| KEGG_PATHWAY | mmu05017:Spinocerebellar ataxia | 8 | 0.09783492586958 | Q3UMR5, O70435, O35226, O55143, Q9D6Z1, P62334, Q60692, P14685 | 0.3841276933 |
| GOTERM_BP_DIRECT | GO:0043161~proteasome-mediated ubiquitin-dependent protein catabolic process | 6 | 0.46737974413481 | Q78JE5, O70435, O35226, P62334, Q60692, Q61990 | 0.9757501028 |
| Annotation Cluster 36 | Enrichment Score: 1.8824675102379829 | | | | |
| Category | Term | Count | P Value | Genes | FDR |
| GOTERM_MF_DIRECT | GO:0005525~GTP binding | 20 | 1.2966318021E-06 | Q62159, Q9DCE9, P30416, Q9Z0N1, Q922F4, Q9WTP7, P14824, Q9Z0E6, Q9EQP2, Q8K0D5, P36536, Q05144, Q8CHH9, Q8R050, Q9CQD1, Q91V41, P08556, P47758, P08752 | 6.353496E-05 |
| GOTERM_MF_DIRECT | GO:0003924~GTPase activity | 15 | 8.9183377336E-05 | Q62159, Q9DCE9, Q9Z0N1, Q922F4, Q9Z0E6, Q8K0D5, P36536, Q05144, Q8CHH9, Q8R050, Q9CQD1, Q91V41, P08556, P08752 | 0.0021849927 |
| GOTERM_MF_DIRECT | GO:0003925~G protein activity | 6 | 0.00092816469384 | Q9DCE9, Q9CQD1, Q9Z0E6, P08556, P36536 | 0.0161706916 |
| INTERPRO | IPR005225:Small_GTP-bd_dom | 9 | 0.00998101231736 | Q62159, Q8R050, Q9CQD1, Q91V41, P08556, Q8K0D5, P36536, Q05144 | 0.3172024363 |
| UP_SEQ_FEATURE | MOTIF:Effector region | 6 | 0.01994136018318 | Q62159, Q9CQD1, Q91V41, P08556, Q05144 | 0.6238796972 |
| GOTERM_BP_DIRECT | GO:0042832~defense response to protozoan | 4 | 0.02013655583186 | Q9DCE9, Q9CQD1, Q9Z0E6, P08556 | 0.337849069 |
| GOTERM_MF_DIRECT | GO:0019003~GDP binding | 5 | 0.02618171503441 | Q9CQD1, Q91V41, Q9Z0E6, P08556 | 0.1994946763 |
| UP_KW_LIGAND | KW-0342~GTP-binding | 17 | 0.02676684053863 | Q62159, Q9DCE9, Q9Z0N1, Q922F4, Q9WTP7, Q9Z0E6, Q8K0D5, P36536, Q05144, Q8CHH9, Q8R050, Q9CQD1, Q91V41, P08556, P47758, P08752 | 0.0803005216 |
| SMART | SM00174:RHO | 6 | 0.06532651926472 | Q62159, Q9CQD1, Q91V41, P08556, Q05144 | 0.9943820225 |
| UP_KW_PTM | KW-0636~Prenylation | 9 | 0.07819760479967 | Q62159, P21619, Q9CQD1, Q91V41, Q9Z0E6, P08556, Q05144, P28656 | 0.2345928144 |
| SMART | SM00173:RAS | 6 | 0.08043285835009 | Q62159, Q9CQD1, Q91V41, P08556, Q05144 | 0.9943820225 |
| INTERPRO | IPR001806:Small_GTPase | 6 | 0.09223718799107 | Q62159, Q9CQD1, Q91V41, P08556, Q05144 | 0.9862825789 |
| UP_SEQ_FEATURE | LIPID:S-geranylgeranyl cysteine | 5 | 0.09981495059857 | Q62159, Q9CQD1, Q91V41, Q9Z0E6, Q05144 | 0.9909502262 |
| SMART | SM00175:RAB | 6 | 0.12054891956549 | Q62159, Q9CQD1, Q91V41, P08556, Q05144 | 0.9943820225 |
| SMART | SM00176:RAN | 3 | 0.4214737292681 | Q9CQD1, Q91V41, Q05144 | 0.9943820225 |
| Annotation Cluster 37 | Enrichment Score: 1.7133833672581156 | | | | |
| Category | Term | Count | P Value | Genes | FDR |
| GOTERM_BP_DIRECT | GO:0006606~protein import into nucleus | 9 | 0.00017447258026 | O35129, Q91YE6, Q61656, Q8BFY9, P34960, P52293, P18760, P70168, P63017 | 0.0170300421 |
| GOTERM_MF_DIRECT | GO:0061608~nuclear import signal receptor activity | 4 | 0.00219762253873 | Q91YE6, Q8BFY9, P52293, P70168 | 0.0319062235 |
| COG_ONTOLOGY | Intracellular trafficking and secretion | 5 | 0.01358244801043 | Q8BRF7, Q8BFY9, P52293, P70168, Q8BX70 | 0.1510429872 |
| SMART | SM00913:IBN_N | 3 | 0.03037031098822 | Q91YE6, Q8BFY9, P70168 | 0.5972827828 |
| UP_SEQ_FEATURE | DOMAIN:Importin N-terminal | 3 | 0.03275703768838 | Q91YE6, Q8BFY9, P70168 | 0.7101571319 |
| INTERPRO | IPR001494:Importin-beta_N | 3 | 0.03568635658712 | Q91YE6, Q8BFY9, P70168 | 0.6109164378 |
| GOTERM_MF_DIRECT | GO:0008139~nuclear localization sequence binding | 3 | 0.04264459076711 | Q8BFY9, P52293, P70168 | 0.280952598 |
| INTERPRO | IPR000225:Armadillo | 3 | 0.2086652582386 | Q8BFY9, P52293, P70168 | 0.9862825789 |
| KEGG_PATHWAY | mmu03013:Nucleocytoplasmic transport | 6 | 0.23084538169575 | P46061, Q91YE6, Q8BFY9, P52293, P70168, Q91VC3 | 0.6528208106 |
| Annotation Cluster 38 | Enrichment Score: 1.685895514879543 | | | | |
| Category | Term | Count | P Value | Genes | FDR |
| GOTERM_BP_DIRECT | GO:0008380~RNA splicing | 14 | 1.2570416472E-06 | Q921F2, Q8BL97, P61979, Q3UKJ7, Q7TMK9, Q3UEB3, Q91VC3, P62960, P16110, Q62093, Q3U0V1, P84104, P62307, P63017 | 0.0004973695 |
| GOTERM_BP_DIRECT | GO:0006397~mRNA processing | 14 | 1.8421236325E-05 | Q921F2, Q8BL97, P61979, Q7TMK9, Q3UEB3, Q80X82, P62960, P16110, Q62093, Q3U0V1, P60335, Q3TIU4, P84104, P63017 | 0.0039756377 |
| UP_SEQ_FEATURE | DOMAIN:RRM 1 | 4 | 0.19506128669307 | Q921F2, Q8C2Q3, Q7TMK9, Q3UEB3 | 0.9909502262 |
| UP_SEQ_FEATURE | DOMAIN:RRM 2 | 4 | 0.19506128669307 | Q921F2, Q8C2Q3, Q7TMK9, Q3UEB3 | 0.9909502262 |
| SMART | SM00360:RRM | 7 | 0.19665882320174 | Q921F2, Q8BL97, Q8C2Q3, Q62093, P84104, Q7TMK9, Q3UEB3 | 0.9943820225 |
| UP_SEQ_FEATURE | DOMAIN:RRM | 7 | 0.21084444317845 | Q921F2, Q8BL97, Q8C2Q3, Q62093, P84104, Q7TMK9, Q3UEB3 | 0.9909502262 |
| INTERPRO | IPR012677:Nucleotide-bd_a/b_plait_sf | 8 | 0.21718353695117 | Q921F2, Q8BL97, Q8C2Q3, Q2EMV9, Q62093, P84104, Q7TMK9, Q3UEB3 | 0.9862825789 |
| INTERPRO | IPR000504:RRM_dom | 7 | 0.24799807072959 | Q921F2, Q8BL97, Q8C2Q3, Q62093, P84104, Q7TMK9, Q3UEB3 | 0.9862825789 |
| INTERPRO | IPR035979:RBD_domain_sf | 7 | 0.34117637972034 | Q921F2, Q8BL97, Q8C2Q3, Q62093, P84104, Q7TMK9, Q3UEB3 | 0.9862825789 |
| Annotation Cluster 39 | Enrichment Score: 1.658885910815123 | | | | |
| Category | Term | Count | P Value | Genes | FDR |
| INTERPRO | IPR006089:Acyl-CoA_DH_CS | 3 | 0.01036753442644 | Q9DBL1, Q9JHI5, Q8JZN5 | 0.3172024363 |
| GOTERM_MF_DIRECT | GO:0050660~flavin adenine dinucleotide binding | 5 | 0.015178730542 | Q9DBL1, Q9JLT4, Q9JMH6, Q9JHI5, Q8JZN5 | 0.1416681517 |
| INTERPRO | IPR009075:AcylCo_DH/oxidase_C | 3 | 0.01835294351816 | Q9DBL1, Q9JHI5, Q8JZN5 | 0.4473141149 |
| INTERPRO | IPR013786:AcylCoA_DH/ox_N | 3 | 0.01835294351816 | Q9DBL1, Q9JHI5, Q8JZN5 | 0.4473141149 |
| COG_ONTOLOGY | Lipid metabolism | 6 | 0.02157756960411 | Q9DBL1, O35459, Q9QUJ7, Q9JHI5, P41216, Q8JZN5 | 0.1510429872 |
| INTERPRO | IPR037069:AcylCoA_DH/ox_N_sf | 3 | 0.02472297176974 | Q9DBL1, Q9JHI5, Q8JZN5 | 0.555494272 |
| INTERPRO | IPR006091:Acyl-CoA_Oxase/DH_mid-dom | 3 | 0.02819738691537 | Q9DBL1, Q9JHI5, Q8JZN5 | 0.6051916774 |
| INTERPRO | IPR009100:AcylCoA_DH/oxidase_NM_dom_sf | 3 | 0.03185435471419 | Q9DBL1, Q9JHI5, Q8JZN5 | 0.6080878132 |
| INTERPRO | IPR036250:AcylCo_DH-like_C | 3 | 0.03185435471419 | Q9DBL1, Q9JHI5, Q8JZN5 | 0.6080878132 |
| INTERPRO | IPR046373:Acyl-CoA_Oxase/DH_mid-dom_sf | 3 | 0.03185435471419 | Q9DBL1, Q9JHI5, Q8JZN5 | 0.6080878132 |
| Annotation Cluster 40 | Enrichment Score: 1.6400610534314646 | | | | |
| Category | Term | Count | P Value | Genes | FDR |
| GOTERM_BP_DIRECT | GO:0002183~cytoplasmic translational initiation | 3 | 0.00879027125586 | Q9Z0N1, Q99L45, P60843 | 0.2293198238 |
| GOTERM_MF_DIRECT | GO:0003743~translation initiation factor activity | 5 | 0.01251532515782 | Q9Z0N1, P23116, Q99L45, P60229, P60843 | 0.1242027206 |
| GOTERM_BP_DIRECT | GO:0001732~formation of cytoplasmic translation initiation complex | 3 | 0.01836552080711 | P23116, Q99L45, P60229 | 0.3182463241 |
| UP_KW_MOLECULAR_FUNCTION | KW-0396~Initiation factor | 5 | 0.02947013565955 | Q9Z0N1, P23116, Q99L45, P60229, P60843 | 0.1473506783 |
| BIOCARTA | m_eifPathway:Eukaryotic protein translation | 3 | 0.04813353256637 | Q9Z0N1, P23116, Q99L45 | 1 |
| GOTERM_BP_DIRECT | GO:0006413~translational initiation | 4 | 0.0503913676249 | Q9Z0N1, Q99L45, P60229, P60843 | 0.5606242819 |
| Annotation Cluster 41 | Enrichment Score: 1.608971323991155 | | | | |
| Category | Term | Count | P Value | Genes | FDR |
| GOTERM_MF_DIRECT | GO:0015485~cholesterol binding | 5 | 0.00490122136185 | Q9Z0J0, O35114, P32020, P14824 | 0.0556892398 |
| GOTERM_BP_DIRECT | GO:0032367~intracellular cholesterol transport | 3 | 0.01620938042716 | Q9Z0J0, P32020 | 0.2937486193 |
| GOTERM_MF_DIRECT | GO:0120020~cholesterol transfer activity | 3 | 0.0395708609332 | Q9Z0J0, P32020 | 0.2651585895 |
| UP_KW_BIOLOGICAL_PROCESS | KW-0445~Lipid transport | 7 | 0.11659192479365 | Q9Z0J0, Q05816, Q9QZD8, P32020, Q9CR62, Q8BX70 | 0.5635422367 |
| Annotation Cluster 42 | Enrichment Score: 1.549007696788112 | | | | |
| Category | Term | Count | P Value | Genes | FDR |
| GOTERM_CC_DIRECT | GO:0030863~cortical cytoskeleton | 6 | 9.044534189E-05 | P21107, Q7TPR4, P57780, Q8BTM8, P47754, P47757 | 0.0013312424 |
| GOTERM_CC_DIRECT | GO:0005903~brush border | 8 | 0.00017730692947 | P21107, Q7TPR4, P57780, Q8BTM8, P47754, A2AQP0, P47757, E9Q634 | 0.0022570693 |
| GOTERM_CC_DIRECT | GO:0005884~actin filament | 7 | 0.00283266730421 | Q99K51, P21107, Q7TPR4, P10107, Q8BTM8, Q91YR1, Q05144 | 0.0180295446 |
| UP_SEQ_FEATURE | DOMAIN:Calponin-homology (CH) 1 | 4 | 0.00659409529303 | Q99K51, Q7TPR4, P57780, Q8BTM8 | 0.2987807316 |
| UP_SEQ_FEATURE | DOMAIN:Calponin-homology (CH) 2 | 4 | 0.00659409529303 | Q99K51, Q7TPR4, P57780, Q8BTM8 | 0.2987807316 |
| INTERPRO | IPR001589:Actinin_actin-bd_CS | 4 | 0.00749906707879 | Q99K51, Q7TPR4, P57780, Q8BTM8 | 0.26301606 |
| UP_SEQ_FEATURE | DOMAIN:Calponin-homology (CH) | 6 | 0.00981484311697 | Q99K51, Q7TPR4, P57780, Q8BTM8, Q9R0C8, Q61166 | 0.368023655 |
| INTERPRO | IPR001715:CH_dom | 6 | 0.01522757705677 | Q99K51, Q7TPR4, P57780, Q8BTM8, Q9R0C8, Q61166 | 0.4030683217 |
| GOTERM_BP_DIRECT | GO:0051017~actin filament bundle assembly | 4 | 0.01560962208353 | Q99K51, Q7TPR4, P57780, P26645 | 0.2937486193 |
| GOTERM_CC_DIRECT | GO:0030018~Z disc | 7 | 0.01609083050794 | P26883, Q7TPR4, Q78JE5, P57780, Q8BTM8, P47757, Q6P069 | 0.0755953876 |
| UP_SEQ_FEATURE | REGION:Actin-binding | 5 | 0.01662953099496 | Q7TPR4, P57780, Q8BTM8, A2AQP0, E9Q634 | 0.5462800932 |
| INTERPRO | IPR036872:CH_dom_sf | 6 | 0.01674763512848 | Q99K51, Q7TPR4, P57780, Q8BTM8, Q9R0C8, Q61166 | 0.4225105143 |
| SMART | SM00033:CH | 5 | 0.02707307277796 | Q99K51, Q7TPR4, P57780, Q8BTM8, Q9R0C8 | 0.5972827828 |
| GOTERM_CC_DIRECT | GO:0001725~stress fiber | 5 | 0.05171513244205 | O55222, P21107, Q7TPR4, P57780, A2AQP0 | 0.1791016719 |
| INTERPRO | IPR018247:EF_Hand_1_Ca_BS | 5 | 0.38797538320585 | Q99K51, Q7TPR4, P57780, Q9EQP2, Q6P069 | 0.9862825789 |
| SMART | SM00054:EFh | 4 | 0.46816097029 | Q99K51, Q7TPR4, P57780, Q6P069 | 0.9943820225 |
| UP_SEQ_FEATURE | DOMAIN:EF-hand 1 | 4 | 0.52996154119726 | Q99K51, Q7TPR4, P57780, Q6P069 | 0.9909502262 |

| | | | | | |
|---|---|---|---|---|---|
| UP_SEQ_FEATURE | DOMAIN:EF-hand 2 | 4 | 0.53403612397781 | Q99K51, Q7TPR4, P57780, Q6P069 | 0.9909502262 |
| INTERPRO | IPR002048:EF_hand_dom | 5 | 0.56954878675301 | Q99K51, Q7TPR4, P57780, Q9EQP2, Q6P069 | 0.9862825789 |
| UP_SEQ_FEATURE | DOMAIN:EF-hand | 4 | 0.65641979047531 | Q99K51, Q7TPR4, P57780, Q9EQP2 | 0.9909502262 |
| INTERPRO | IPR011992:EF-hand-dom_pair | 5 | 0.70959826535652 | Q99K51, Q7TPR4, P57780, Q9EQP2, Q6P069 | 0.9862825789 |
| Annotation Cluster 43 | Enrichment Score: 1.5118739836232185 | | | | |
| Category | Term | Count | P Value | Genes | FDR |
| INTERPRO | IPR009000:Transl_B-barrel_sf | 4 | 0.0186863252893 | Q8R050, Q9Z0N1, Q8K0D5, Q8BGQ7 | 0.4478489294 |
| INTERPRO | IPR004161:EFTu-like_2 | 3 | 0.02472297176974 | Q8R050, Q9Z0N1, Q8K0D5 | 0.555494272 |
| UP_SEQ_FEATURE | DOMAIN:tr-type G | 3 | 0.04028921301399 | Q8R050, Q9Z0N1, Q8K0D5 | 0.7901496403 |
| INTERPRO | IPR000795:T_Tr_GTP-bd_dom | 3 | 0.04816046856332 | Q8R050, Q9Z0N1, Q8K0D5 | 0.7736119135 |
| Annotation Cluster 44 | Enrichment Score: 1.4360462274619736 | | | | |
| Category | Term | Count | P Value | Genes | FDR |
| GOTERM_BP_DIRECT | GO:0097320~plasma membrane tubulation | 3 | 0.01620938042716 | D3Z6Q9, Q91ZR2, Q91VH2 | 0.2937486193 |
| UP_SEQ_FEATURE | DOMAIN:BAR | 4 | 0.01944321439964 | D3Z6Q9, Q91ZR2, Q91VH2, Q3UIA2 | 0.6231313103 |
| INTERPRO | IPR027267:AH/BAR_dom_sf | 4 | 0.15607266183814 | D3Z6Q9, Q91ZR2, Q91VH2, Q3UIA2 | 0.9862825789 |
| Annotation Cluster 45 | Enrichment Score: 1.4081143822190803 | | | | |
| Category | Term | Count | P Value | Genes | FDR |
| UP_SEQ_FEATURE | REPEAT:4 | 8 | 0.0032613582204 | P62320, O08784, Q8BL97, P16110, Q3U0V1, P23116, P13864, P08775 | 0.2141615333 |
| UP_SEQ_FEATURE | REPEAT:3 | 8 | 0.00846169284963 | P62320, O08784, Q8BL97, P16110, Q3U0V1, P13864, Q62433, P08775 | 0.3474582626 |
| UP_SEQ_FEATURE | REPEAT:1 | 8 | 0.02369844126078 | P62320, O08784, Q8BL97, P16110, Q3U0V1, P13864, Q62433, P08775 | 0.6830397234 |
| UP_SEQ_FEATURE | REPEAT:5 | 6 | 0.02443140563006 | P62320, O08784, P16110, P23116, P13864, P08775 | 0.6830397234 |
| UP_SEQ_FEATURE | REPEAT:2 | 8 | 0.02508378128718 | P62320, O08784, Q8BL97, P16110, Q3U0V1, P23116, P13864, Q62433 | 0.6866685127 |
| UP_SEQ_FEATURE | REPEAT:6 | 5 | 0.03981557558655 | O08784, P16110, P23116, P13864, P08775 | 0.7901496403 |
| UP_SEQ_FEATURE | REPEAT:8 | 4 | 0.0940304974473 | O08784, P16110, P23116, P08775 | 0.9909502262 |
| UP_SEQ_FEATURE | REPEAT:7 | 4 | 0.11500655162178 | O08784, P16110, P23116, P08775 | 0.9909502262 |
| UP_SEQ_FEATURE | REPEAT:10 | 3 | 0.21925395254483 | O08784, P23116, P08775 | 0.9909502262 |
| UP_SEQ_FEATURE | REPEAT:9 | 3 | 0.21925395254483 | O08784, P23116, P08775 | 0.9909502262 |
| Annotation Cluster 46 | Enrichment Score: 1.371090541206241 | | | | |
| Category | Term | Count | P Value | Genes | FDR |
| GOTERM_BP_DIRECT | GO:1905820~positive regulation of chromosome separation | 3 | 0.00725484579349 | Q8CG48, Q8CG47, E9Q7G0 | 0.1979655622 |
| GOTERM_BP_DIRECT | GO:0051984~positive regulation of chromosome segregation | 3 | 0.01417026039962 | Q8CG48, Q8CG47, E9Q7G0 | 0.2850864253 |
| GOTERM_BP_DIRECT | GO:0051321~meiotic cell cycle | 3 | 0.7494149621768 | Q8CG48, Q8CG47, E9Q7G0 | 0.9757501028 |
| Annotation Cluster 47 | Enrichment Score: 1.3610955067730057 | | | | |
| Category | Term | Count | P Value | Genes | FDR |
| INTERPRO | IPR011989:ARM-like | 11 | 0.01010086431473 | Q9JL26, Q91YE6, O35841, Q8BVE3, Q8BFY9, P52293, P70168, Q9DBD5, P17426, Q80X82, Q76MZ3 | 0.3172024363 |
| UP_SEQ_FEATURE | REPEAT:HEAT 15 | 3 | 0.02265788462179 | Q8BFY9, P70168, Q76MZ3 | 0.6766468271 |
| UP_SEQ_FEATURE | REPEAT:HEAT 5 | 4 | 0.02433130205737 | Q8BFY9, P70168, Q80X82, Q76MZ3 | 0.6830397234 |
| UP_SEQ_FEATURE | REPEAT:HEAT 14 | 3 | 0.02922429429459 | Q8BFY9, P70168, Q76MZ3 | 0.7101571319 |
| UP_SEQ_FEATURE | REPEAT:HEAT 13 | 3 | 0.03275703768838 | Q8BFY9, P70168, Q76MZ3 | 0.7101571319 |
| UP_SEQ_FEATURE | REPEAT:HEAT 12 | 3 | 0.03644744590428 | Q8BFY9, P70168, Q76MZ3 | 0.760189586 |
| UP_SEQ_FEATURE | REPEAT:HEAT 11 | 3 | 0.04028921301399 | Q8BFY9, P70168, Q76MZ3 | 0.7901496403 |
| UP_SEQ_FEATURE | REPEAT:HEAT 4 | 4 | 0.04257835413549 | Q8BFY9, P70168, Q80X82, Q76MZ3 | 0.7992565333 |
| UP_SEQ_FEATURE | REPEAT:HEAT 10 | 3 | 0.04840243179279 | Q8BFY9, P70168, Q76MZ3 | 0.8849956187 |
| UP_SEQ_FEATURE | REPEAT:HEAT 3 | 4 | 0.05236585167208 | Q8BFY9, P70168, Q80X82, Q76MZ3 | 0.9174497213 |
| UP_SEQ_FEATURE | REPEAT:HEAT 9 | 3 | 0.05704952006599 | Q8BFY9, P70168, Q76MZ3 | 0.9610649919 |
| UP_SEQ_FEATURE | REPEAT:HEAT 8 | 3 | 0.06618579827663 | Q8BFY9, P70168, Q76MZ3 | 0.9893860137 |
| UP_SEQ_FEATURE | REPEAT:HEAT 1 | 4 | 0.07790518842958 | Q8BFY9, P70168, Q80X82, Q76MZ3 | 0.9909502262 |
| UP_SEQ_FEATURE | REPEAT:HEAT 2 | 4 | 0.07790518842958 | Q8BFY9, P70168, Q80X82, Q76MZ3 | 0.9909502262 |
| UP_SEQ_FEATURE | REPEAT:HEAT 7 | 3 | 0.08071554270985 | Q8BFY9, P70168, Q76MZ3 | 0.9909502262 |
| UP_SEQ_FEATURE | REPEAT:HEAT 6 | 3 | 0.10681030145286 | Q8BFY9, P70168, Q76MZ3 | 0.9909502262 |
| Annotation Cluster 48 | Enrichment Score: 1.327523684757782 | | | | |
| Category | Term | Count | P Value | Genes | FDR |
| UP_SEQ_FEATURE | LIPID:N-myristoyl glycine | 10 | 0.00376915265763 | Q9JL26, Q99JX3, Q9CRB9, Q6A0D4, P25911, P00520, Q9CQ22, P08752, P26645, Q91XV3 | 0.2235590146 |
| UP_KW_PTM | KW-0519~Myristate | 10 | 0.0650906256269 | Q9JL26, Q99JX3, Q9CRB9, Q6A0D4, P25911, P00520, Q9CQ22, P08752, P26645, Q91XV3 | 0.2231678593 |
| UP_KW_PTM | KW-0449~Lipoprotein | 27 | 0.4242938846807 | Q9CQW9, P10810, Q6A0D4, Q9CQ22, Q9CRB9, Q9CQD1, P00520, Q91XV3, P28656, Q62159, Q99JX3, E9Q7G0, Q99J93, Q9Z0E6, P35564, Q05144, Q9JL26, P21619, O35114, Q8BMK4, Q91V41, Q8VBT0, P25911, P08556, P08752, P26645 | 0.8158465471 |
| Annotation Cluster 49 | Enrichment Score: 1.2272274884220515 | | | | |
| Category | Term | Count | P Value | Genes | FDR |
| UP_SEQ_FEATURE | REPEAT:2-1 | 3 | 0.05704952006599 | P61979, P35564, Q7TMK9 | 0.9610649919 |
| UP_SEQ_FEATURE | REPEAT:2-2 | 3 | 0.05704952006599 | P61979, P35564, Q7TMK9 | 0.9610649919 |
| UP_SEQ_FEATURE | REPEAT:1-2 | 3 | 0.06155920859774 | P61979, P35564, Q7TMK9 | 0.9893860137 |
| UP_SEQ_FEATURE | REPEAT:1-1 | 3 | 0.06155920859774 | P61979, P35564, Q7TMK9 | 0.9893860137 |
| Annotation Cluster 50 | Enrichment Score: 1.2080601339753034 | | | | |
| Category | Term | Count | P Value | Genes | FDR |
| GOTERM_BP_DIRECT | GO:0006909~phagocytosis | 9 | 8.8229469913E-06 | P05555, P10107, P09055, O70145, P11835, Q9CQD1, O35639, P00520, Q8BG07 | 0.0023272974 |
| GOTERM_BP_DIRECT | GO:0007229~integrin-mediated signaling pathway | 9 | 0.00010037697088 | P05555, O55222, Q64339, P20491, P09055, P18052, P11835, Q9R0C8, P00520 | 0.0119625 |
| GOTERM_BP_DIRECT | GO:0007160~cell-matrix adhesion | 6 | 0.00925350736477 | P05555, O55222, P09055, P97797, P11835, Q61072 | 0.236213188 |
| GOTERM_BP_DIRECT | GO:0050798~activated T cell proliferation | 3 | 0.01417026039962 | P05555, P11835, P00520 | 0.2850864253 |
| GOTERM_BP_DIRECT | GO:0033627~cell adhesion mediated by integrin | 4 | 0.01560962208353 | P05555, P09055, P11835, Q61072 | 0.2937486193 |
| KEGG_PATHWAY | mmu05133:Pertussis | 7 | 0.01992352449359 | P05555, P10810, P63085, P09055, P11835, P18760, P08752 | 0.1343560754 |
| KEGG_PATHWAY | mmu05140:Leishmaniasis | 6 | 0.04474856469044 | P05555, P26151, P63085, P09055, O70145, P11835 | 0.2374874645 |
| GOTERM_CC_DIRECT | GO:0008305~integrin complex | 3 | 0.05558333131035 | P05555, P09055, P11835 | 0.1883435183 |
| GOTERM_BP_DIRECT | GO:0007159~leukocyte cell-cell adhesion | 3 | 0.05904536193303 | P05555, P09055, P11835 | 0.6175052389 |
| GOTERM_BP_DIRECT | GO:0034113~heterotypic cell-cell adhesion | 3 | 0.06258329051497 | P05555, P09055, P11835 | 0.6318014784 |
| UP_SEQ_FEATURE | DOMAIN:VWFA | 5 | 0.0662602505394 | P05555, O35226, P09055, P11835, Q8BT60 | 0.9893860137 |
| BIOCARTA | m_monocytePathway:Monocyte and its Surface Molecules | 3 | 0.06984796354927 | P05555, P09055, P11835 | 1 |
| INTERPRO | IPR032695:Integrin_dom_sf | 3 | 0.08212154546752 | P05555, P09055, P11835 | 0.9862825789 |
| INTERPRO | IPR036465:vWFA_dom_sf | 5 | 0.11621627612515 | P05555, O35226, P09055, P11835, Q8BT60 | 0.9862825789 |
| GOTERM_BP_DIRECT | GO:0098609~cell-cell adhesion | 5 | 0.28772454822415 | P05555, P09055, P11835, P62137, P00520 | 0.9757501028 |
| UP_KW_MOLECULAR_FUNCTION | KW-0401~Integrin | 3 | 0.28814336629059 | P05555, P09055, P11835 | 0.7803882837 |
| SMART | SM00327:VWA | 3 | 0.3665247081522 | P05555, O35226, Q8BT60 | 0.9943820225 |
| INTERPRO | IPR002035:VWF_A | 3 | 0.42148222728719 | P05555, O35226, Q8BT60 | 0.9862825789 |
| UP_SEQ_FEATURE | DOMAIN:EGF-like 2 | 3 | 0.46215995803872 | P21956, P09055, P11835 | 0.9909502262 |
| GOTERM_BP_DIRECT | GO:0007155~cell adhesion | 8 | 0.47795104519333 | P21956, P05555, P09055, P18052, P11835, Q61072, Q07797, Q80X82 | 0.9757501028 |
| KEGG_PATHWAY | mmu04514:Cell adhesion molecules | 6 | 0.55887432766039 | Q61543, P05555, P09055, P01900, P11835, P01897 | 0.8855218855 |
| UP_SEQ_FEATURE | DOMAIN:EGF-like 1 | 3 | 0.60448099290159 | P21956, P09055, P11835 | 0.9909502262 |
| UP_KW_DOMAIN | KW-0245~EGF-like domain | 4 | 0.7686576218091 | P21956, P09055, P11835, Q61072 | 1 |
| UP_KW_BIOLOGICAL_PROCESS | KW-0130~Cell adhesion | 7 | 0.94252735952127 | P21956, P05555, P09055, P11835, Q07797, P00520, Q80X82 | 0.9425273595 |
| Annotation Cluster 51 | Enrichment Score: 1.194793082414913 | | | | |
| Category | Term | Count | P Value | Genes | FDR |
| SMART | SM00382:AAA | 7 | 0.04087880067412 | P46460, Q9JHU4, P62334, Q61881, Q8CGK3, E9Q555, Q9D0F6 | 0.7235547719 |
| INTERPRO | IPR003593:AAA+_ATPase | 7 | 0.04581166708484 | P46460, Q9JHU4, P62334, Q61881, Q8CGK3, E9Q555, Q9D0F6 | 0.7572089341 |
| INTERPRO | IPR003959:ATPase_AAA_core | 4 | 0.05039192594997 | P46460, P62334, Q8CGK3, Q9D0F6 | 0.7736119135 |
| UP_SEQ_FEATURE | DOMAIN:AAA+ ATPase | 3 | 0.17619455025228 | Q9JHU4, P62334, Q9D0F6 | 0.9909502262 |
| Annotation Cluster 52 | Enrichment Score: 1.0771315179578915 | | | | |
| Category | Term | Count | P Value | Genes | FDR |
| GOTERM_CC_DIRECT | GO:0005694~chromosome | 10 | 0.01138679593305 | P49718, Q8CG48, P49717, Q8CG47, P97310, Q3TEA8, Q9CXT6, E9Q7G0, Q61881, P43247 | 0.0558664675 |
| UP_KW_BIOLOGICAL_PROCESS | KW-0131~Cell cycle | 19 | 0.15040586432652 | P62908, Q91ZR2, Q8CG48, Q60875, Q8CG47, P97310, Q9CXT6, E9Q7G0, Q91VH2, Q8VBW6, Q61881, Q61166, P49718, P49717, P63085, Q9JHU4, P62137, P43247, P08752 | 0.5635422367 |
| UP_KW_CELLULAR_COMPONENT | KW-0158~Chromosome | 15 | 0.34271975510772 | P46061, Q8CG48, Q8CG47, Q61029, P97310, Q9CXT6, E9Q7G0, Q3TEA8, Q61033, Q61881, Q76MZ3, P49718, P49717, P43277, P43247, P08775 | 0.5654875959 |
| Annotation Cluster 53 | Enrichment Score: 1.069404356850949 | | | | |
| Category | Term | Count | P Value | Genes | FDR |
| INTERPRO | IPR020845:AMP-binding_CS | 3 | 0.07180770516355 | Q8VCW8, Q9QUJ7, P41216 | 0.9862825789 |
| INTERPRO | IPR042099:ANL_N_sf | 3 | 0.0928555658462 | Q8VCW8, Q9QUJ7, P41216 | 0.9862825789 |
| INTERPRO | IPR000873:AMP-dep_Synth/Lig_com | 3 | 0.0928555658462 | Q8VCW8, Q9QUJ7, P41216 | 0.9862825789 |
| Annotation Cluster 54 | Enrichment Score: 1.059425108598835 | | | | |
| Category | Term | Count | P Value | Genes | FDR |
| GOTERM_BP_DIRECT | GO:0050853~B cell receptor signaling pathway | 6 | 0.00024596044811 | P63085, Q8K1X4, Q9R0C8, Q6A0D4, P25911, P00520 | 0.0192673144 |
| UP_KW_DOMAIN | KW-0727~SH2 domain | 4 | 0.24715326527137 | Q9WVL2, Q9R0C8, P25911, P00520 | 0.7767674051 |
| SMART | SM00252:SH2 | 4 | 0.26754926218582 | Q9WVL2, Q9R0C8, P25911, P00520 | 0.9943820225 |
| UP_SEQ_FEATURE | DOMAIN:SH2 | 4 | 0.27091618265527 | Q9WVL2, Q9R0C8, P25911, P00520 | 0.9909502262 |
| INTERPRO | IPR000980:SH2 | 4 | 0.30411980939524 | Q9WVL2, Q9R0C8, P25911, P00520 | 0.9862825789 |
| INTERPRO | IPR036860:SH2_dom_sf | 4 | 0.3283479471781 | Q9WVL2, Q9R0C8, P25911, P00520 | 0.9862825789 |
| Annotation Cluster 55 | Enrichment Score: 1.034825508296877 | | | | |
| Category | Term | Count | P Value | Genes | FDR |
| KEGG_PATHWAY | mmu05100:Bacterial invasion of epithelial cells | 6 | 0.0626701870061 | Q8CHH9, Q8BH43, O55222, Q9JM76, P09055, P59999 | 0.3109860223 |
| KEGG_PATHWAY | mmu05135:Yersinia infection | 8 | 0.08459890183977 | Q8BH43, Q9JM76, P63085, P09055, Q3UND0, Q9R0C8, P59999, Q05144 | 0.3476486122 |
| BIOCARTA | m_actinYPathway:Y branching of actin filaments | 3 | 0.14828518348147 | Q8BH43, Q9JM76, P59999 | 1 |
| Annotation Cluster 56 | Enrichment Score: 0.9594428183361571 | | | | |

| Category | Term | Count | P Value | Genes | FDR |
|---|---|---|---|---|---|
| GOTERM_BP_DIRECT | GO:0051607~defense response to virus | 8 | 0.04846847494053 | Q64339, Q9CQW9, Q99J93, Q501J6, Q9D906, Q9WVL2, Q9Z0E6, Q61990 | 0.5606242819 |
| GOTERM_BP_DIRECT | GO:0045071~negative regulation of viral genome replication | 3 | 0.15883919606353 | Q64339, Q9CQW9, Q99J93 | 0.9021154341 |
| UP_KW_BIOLOGICAL_PROCESS | KW-0051~Antiviral defense | 5 | 0.17189154289767 | Q64339, Q9CQW9, Q99J93, Q501J6, Q61990 | 0.595009187 |
| Annotation Cluster 57 | Enrichment Score: 0.896871904351465 | | | | |
| Category | Term | Count | P Value | Genes | FDR |
| GOTERM_BP_DIRECT | GO:0048511~rhythmic process | 4 | 0.08887916461738 | Q921F2, Q61656, Q9D906, P62137 | 0.7017410113 |
| UP_KW_BIOLOGICAL_PROCESS | KW-0090~Biological rhythms | 7 | 0.10174148554595 | Q921F2, Q61656, Q9CXT6, O70133, Q7TPV4, Q9D906, P62137 | 0.5386313941 |
| GOTERM_BP_DIRECT | GO:0042752~regulation of circadian rhythm | 3 | 0.22546869356687 | Q921F2, Q9D906, P62137 | 0.9757501028 |
| Annotation Cluster 58 | Enrichment Score: 0.8745465208990642 | | | | |
| Category | Term | Count | P Value | Genes | FDR |
| GOTERM_BP_DIRECT | GO:0051301~cell division | 11 | 0.02654928945412 | P62908, Q91ZR2, Q8CG48, Q60875, Q8CG47, E9Q7G0, Q9JHU4, P27546, Q61166, P62137, P08752 | 0.3913326126 |
| UP_KW_BIOLOGICAL_PROCESS | KW-0131~Cell cycle | 19 | 0.15040586432652 | P62908, Q91ZR2, Q8CG48, Q60875, Q8CG47, P97310, Q9CXT6, E9Q7G0, Q91VH2, Q8VBW6, Q61881, Q61166, P49718, P49717, P63085, Q9JHU4, P62137, P43247, P08752 | 0.5635422367 |
| UP_KW_BIOLOGICAL_PROCESS | KW-0498~Mitosis | 9 | 0.25023291495691 | P62908, Q91ZR2, Q8CG48, Q60875, Q8CG47, Q91VH2, E9Q7G0, Q9JHU4, Q61166 | 0.7731060609 |
| UP_KW_BIOLOGICAL_PROCESS | KW-0132~Cell division | 11 | 0.31779847526033 | P62908, Q91ZR2, Q8CG48, Q60875, Q8CG47, Q91VH2, E9Q7G0, Q9JHU4, Q61166, P62137, P08752 | 0.841231258 |
| Annotation Cluster 59 | Enrichment Score: 0.861986333181995 | | | | |
| Category | Term | Count | P Value | Genes | FDR |
| KEGG_PATHWAY | mmu05221:Acute myeloid leukemia | 7 | 0.01292356287368 | P05555, P26151, P10810, P63085, P09581, P08556 | 0.0999675599 |
| KEGG_PATHWAY | mmu04640:Hematopoietic cell lineage | 5 | 0.26493037338599 | P05555, P26151, P10810, P09581, P13379 | 0.7219590359 |
| KEGG_PATHWAY | mmu05202:Transcriptional misregulation in cancer | 6 | 0.75775182123993 | P05555, Q61656, P26151, P10810, P09581, O08734 | 0.8855218855 |
| Annotation Cluster 60 | Enrichment Score: 0.8097452402425509 | | | | |
| Category | Term | Count | P Value | Genes | FDR |
| GOTERM_MF_DIRECT | GO:0035091~phosphatidylinositol binding | 5 | 0.0628575377254 | Q91ZR2, Q91VH2, E9Q7G0, O70493, E9Q634 | 0.3559773998 |
| SMART | SM00312:PX | 3 | 0.16360067595342 | Q91ZR2, Q91VH2, O70493 | 0.9943820225 |
| UP_SEQ_FEATURE | DOMAIN:PX | 3 | 0.18837188531427 | Q91ZR2, Q91VH2, O70493 | 0.9909502262 |
| INTERPRO | IPR001683:PX_dom | 3 | 0.20218218340164 | Q91ZR2, Q91VH2, O70493 | 0.9862825789 |
| INTERPRO | IPR036871:PX_dom_sf | 3 | 0.22822981261271 | Q91ZR2, Q91VH2, O70493 | 0.9862825789 |
| Annotation Cluster 61 | Enrichment Score: 0.7871105885433825 | | | | |
| Category | Term | Count | P Value | Genes | FDR |
| GOTERM_MF_DIRECT | GO:0008233~peptidase activity | 6 | 0.05026758513477 | Q11136, Q9EQH2, Q6NSR8, Q9CYN2, O09043, O89023 | 0.3103132815 |
| GOTERM_MF_DIRECT | GO:0004175~endopeptidase activity | 5 | 0.05964415284071 | Q9EQH2, Q9CXT6, O09043, Q60692, O89023 | 0.3542501199 |
| GOTERM_BP_DIRECT | GO:0006508~proteolysis | 13 | 0.10073411269104 | Q11136, Q9EQH2, Q9CXT6, Q6NSR8, P34960, P16675, O35864, Q9WVJ3, Q61072, O09043, Q8R146, Q8CGK3, O89023 | 0.7335668206 |
| UP_KW_MOLECULAR_FUNCTION | KW-0482~Metalloprotease | 6 | 0.25161398969515 | Q11136, Q9EQH2, O35864, Q9WVJ3, Q61072, P34960 | 0.7110830144 |
| UP_KW_PTM | KW-0865~Zymogen | 8 | 0.44191687967594 | P16675, Q9WVJ3, Q61072, P34960, O09043, Q60692, P29416, O89023 | 0.8158465471 |
| UP_KW_MOLECULAR_FUNCTION | KW-0645~Protease | 13 | 0.56393445833315 | Q11136, Q9EQH2, Q9CXT6, Q6NSR8, P34960, Q60692, P16675, O35864, Q9WVJ3, Q61072, O09043, Q8CGK3, O89023 | 0.8783783784 |
| Annotation Cluster 62 | Enrichment Score: 0.7847583569320772 | | | | |
| Category | Term | Count | P Value | Genes | FDR |
| UP_SEQ_FEATURE | REGION:G1 motif | 3 | 0.16415028574604 | Q8CHH9, Q9EQP2, P08752 | 0.9909502262 |
| UP_SEQ_FEATURE | REGION:G3 motif | 3 | 0.16415028574604 | Q8CHH9, Q9EQP2, P08752 | 0.9909502262 |
| UP_SEQ_FEATURE | REGION:G4 motif | 3 | 0.16415028574604 | Q8CHH9, Q9EQP2, P08752 | 0.9909502262 |
| Annotation Cluster 63 | Enrichment Score: 0.7837796090283677 | | | | |
| Category | Term | Count | P Value | Genes | FDR |
| UP_SEQ_FEATURE | REPEAT:TPR | 5 | 0.05795638313639 | P30416, O70145, Q9CZW5, Q99L47, Q8CD92 | 0.9639833853 |
| SMART | SM00028:TPR | 5 | 0.15692298724557 | P30416, O70145, Q9CZW5, Q99L47, Q8CD92 | 0.9943820225 |
| UP_SEQ_FEATURE | REPEAT:TPR 3 | 5 | 0.16494643612601 | P30416, O70145, Q9CZW5, Q99L47, Q8CD92 | 0.9909502262 |
| INTERPRO | IPR019734:TPR_repeat | 5 | 0.19206796504008 | P30416, O70145, Q9CZW5, Q99L47, Q8CD92 | 0.9862825789 |
| INTERPRO | IPR011990:TPR-like_helical_dom_sf | 7 | 0.19492316707926 | Q9CWZ7, P30416, O70145, Q9CZW5, Q99L47, Q8CD92, P14685 | 0.9862825789 |
| UP_SEQ_FEATURE | REPEAT:TPR 1 | 5 | 0.2020212908586 | P30416, O70145, Q9CZW5, Q99L47, Q8CD92 | 0.9909502262 |
| UP_SEQ_FEATURE | REPEAT:TPR 2 | 5 | 0.2020212908586 | P30416, O70145, Q9CZW5, Q99L47, Q8CD92 | 0.9909502262 |
| UP_KW_DOMAIN | KW-0802~TPR repeat | 5 | 0.2341637539524 | P30416, O70145, Q9CZW5, Q99L47, Q8CD92 | 0.7767674051 |
| Annotation Cluster 64 | Enrichment Score: 0.7032632848961203 | | | | |
| Category | Term | Count | P Value | Genes | FDR |
| INTERPRO | IPR015422:PyrdxlP-dep_Trfase_small | 3 | 0.13314833139114 | Q99K85, P29758, Q8VCN5 | 0.9862825789 |
| GOTERM_MF_DIRECT | GO:0030170~pyridoxal phosphate binding | 3 | 0.15932811094634 | Q99K85, P29758, Q8VCN5 | 0.6438826752 |
| INTERPRO | IPR015421:PyrdxlP-dep_Trfase_major | 3 | 0.17018618912575 | Q99K85, P29758, Q8VCN5 | 0.9862825789 |
| INTERPRO | IPR015424:PyrdxlP-dep_Trfase | 3 | 0.17651911035972 | Q99K85, P29758, Q8VCN5 | 0.9862825789 |
| UP_KW_LIGAND | KW-0663~Pyridoxal phosphate | 3 | 0.47790260384775 | Q99K85, P29758, Q8VCN5 | 0.806460644 |
| Annotation Cluster 65 | Enrichment Score: 0.6544059307143841 | | | | |
| Category | Term | Count | P Value | Genes | FDR |
| UP_SEQ_FEATURE | LIPID:S-palmitoyl cysteine | 10 | 0.06415053774879 | Q9CQW9, Q99J93, Q8VBT0, P35564, Q6A0D4, P25911, P08556, Q9CQ22, P08752 | 0.9893860137 |
| UP_KW_PTM | KW-0564~Palmitate | 13 | 0.39986660395419 | Q99JX3, Q9CQW9, O35114, Q99J93, Q8BMK4, Q8VBT0, P35564, P25911, Q6A0D4, P08556, Q9CQ22, P08752 | 0.8158465471 |
| UP_KW_PTM | KW-0449~Lipoprotein | 27 | 0.4242938846807 | Q9CQW9, P10810, Q6A0D4, Q9CQ22, Q9CRB9, Q9CQD1, P00520, Q91XV3, P28656, Q62159, Q99JX3, E9Q7G0, Q99J93, Q9Z0E6, P35564, Q05144, Q9JL26, P21619, O35114, Q8BMK4, Q91V41, Q8VBT0, P25911, P08556, P08752, P26645 | 0.8158465471 |
| Annotation Cluster 66 | Enrichment Score: 0.6244696329949984 | | | | |
| Category | Term | Count | P Value | Genes | FDR |
| GOTERM_BP_DIRECT | GO:0006511~ubiquitin-dependent protein catabolic process | 9 | 0.01432974485657 | Q78JE5, O70435, Q64339, P68037, Q9ET26, Q8VBW6, O88543, P14685, E9Q555 | 0.285872389 |
| GOTERM_BP_DIRECT | GO:0016567~protein ubiquitination | 4 | 0.94877566993353 | P68037, Q9ET26, Q8VBW6, E9Q555 | 0.9757501028 |
| UP_KW_BIOLOGICAL_PROCESS | KW-0833~Ubl conjugation pathway | 9 | 0.98443966501232 | Q78JE5, Q64339, P68037, Q99KP6, Q9Z1F9, Q9ET26, Q8VBW6, Q9D906, E9Q555 | 0.984439665 |
| Annotation Cluster 67 | Enrichment Score: 0.5701233647874511 | | | | |
| Category | Term | Count | P Value | Genes | FDR |
| KEGG_PATHWAY | mmu04664:Fc epsilon RI signaling pathway | 7 | 0.00981331503821 | P63085, P20491, Q9R0C8, P25911, P08556, Q05144 | 0.0860300618 |
| KEGG_PATHWAY | mmu05221:Acute myeloid leukemia | 7 | 0.01292356287368 | P05555, P26151, P10810, P63085, P09581, P08556 | 0.0999675599 |
| GOTERM_BP_DIRECT | GO:0051726~regulation of cell cycle | 9 | 0.01706559740874 | Q921F2, O35864, P09055, P08556, O08734, P00520, P63017, P43247 | 0.3046144981 |
| KEGG_PATHWAY | mmu05216:Thyroid cancer | 5 | 0.01874622977335 | P21107, P63085, P08556, O08734 | 0.1297436429 |
| KEGG_PATHWAY | mmu05203:Viral carcinogenesis | 13 | 0.02510499842202 | Q7TPR4, Q9CQV8, P61979, P63085, P57780, P34022, O35892, P01900, P01897, P25911, P08556, O08734 | 0.1539845188 |
| KEGG_PATHWAY | mmu04730:Long-term depression | 6 | 0.02517617607032 | P63085, P25911, P08556, P08752, Q76MZ3 | 0.1539845188 |
| GOTERM_BP_DIRECT | GO:0032729~positive regulation of type II interferon production | 5 | 0.02803584266289 | Q64339, P10810, P08556, P00520 | 0.3993573069 |
| UP_KW_DISEASE | KW-0656~Proto-oncogene | 6 | 0.0298469000567 | Q60875, P09581, P25911, P08556, P00520 | 0.0596938001 |
| KEGG_PATHWAY | mmu05230:Central carbon metabolism in cancer | 6 | 0.0424753502756 | P63085, Q00612, P06151, O08528, P08556 | 0.2327295234 |
| GOTERM_BP_DIRECT | GO:0050852~T cell receptor signaling pathway | 5 | 0.05620090441228 | P63085, Q6A0D4, P08556, P00520 | 0.592981987 |
| KEGG_PATHWAY | mmu04360:Axon guidance | 10 | 0.06542867214693 | B2RXS4, O55222, P63085, P09055, P18760, P08556, P00520, P08752, Q05144 | 0.3156598657 |
| KEGG_PATHWAY | mmu04015:Rap1 signaling pathway | 11 | 0.07454426208388 | P05555, P63085, P09055, P09581, P11835, Q9R0C8, P08556, P62962, P08752, Q05144 | 0.3380196712 |
| KEGG_PATHWAY | mmu05213:Endometrial cancer | 5 | 0.07710149358778 | O55222, P63085, P08556, O08734 | 0.3392357758 |
| GOTERM_BP_DIRECT | GO:0001932~regulation of protein phosphorylation | 4 | 0.07894409085361 | B2RXS4, Q9WVL2, P08556 | 0.6915618881 |
| GOTERM_BP_DIRECT | GO:0048146~positive regulation of fibroblast proliferation | 4 | 0.08137999104732 | O70133, P08556, P00520 | 0.7017410113 |
| KEGG_PATHWAY | mmu04662:B cell receptor signaling pathway | 6 | 0.08408992713362 | P63085, Q9R0C8, P25911, P08556, Q05144 | 0.3476486122 |
| GOTERM_BP_DIRECT | GO:0048009~insulin-like growth factor receptor signaling pathway | 4 | 0.09144126262631 | P63085, P09581, P08556, Q6Y7W8 | 0.7164407837 |
| KEGG_PATHWAY | mmu05210:Colorectal cancer | 6 | 0.09785762529988 | P63085, P08556, O08734, P43247, Q05144 | 0.3841276933 |
| KEGG_PATHWAY | mmu04650:Natural killer cell mediated cytotoxicity | 7 | 0.10392991807348 | P63085, P20491, P11835, Q9R0C8, P08556, Q05144 | 0.3961386732 |
| KEGG_PATHWAY | mmu04071:Sphingolipid signaling pathway | 7 | 0.13840304138139 | P63085, P20491, P08556, P08752, Q05144, Q76MZ3 | 0.4918918903 |
| KEGG_PATHWAY | mmu05220:Chronic myeloid leukemia | 5 | 0.16049491743555 | P63085, P08556, O08734, P00520 | 0.5628021771 |
| KEGG_PATHWAY | mmu04062:Chemokine signaling pathway | 9 | 0.16735009637428 | P63085, Q69ZK0, Q9WVL2, Q9R0C8, P25911, P08556, P08752, Q05144 | 0.5676776311 |
| KEGG_PATHWAY | mmu05160:Hepatitis C | 8 | 0.1831331647159 | Q9CQV8, P63085, Q9WVL2, P08556, O08734, P60229, Q76MZ3 | 0.5835850346 |
| KEGG_PATHWAY | mmu04625:C-type lectin receptor signaling pathway | 6 | 0.20077470349854 | P63085, P20491, Q9WVL2, P08556, P19973 | 0.6131460358 |
| KEGG_PATHWAY | mmu05205:Proteoglycans in cancer | 9 | 0.20515912983397 | Q61656, P26041, P63085, P09055, Q8BTM8, Q9R0C8, P08556, P62137 | 0.6131460358 |
| KEGG_PATHWAY | mmu04540:Gap junction | 5 | 0.21669403049407 | P63085, Q922F4, P08556, P08752 | 0.629669723 |
| BIOCARTA | m_fcer1Pathway:Fc Epsilon Receptor I Signaling in Mast Cells | 4 | 0.21724053551078 | P63085, P20491, P25911, P08556 | 1 |
| KEGG_PATHWAY | mmu04370:VEGF signaling pathway | 4 | 0.21787051253182 | P63085, P08556, Q05144 | 0.629669723 |
| KEGG_PATHWAY | mmu05417:Lipid and atherosclerosis | 9 | 0.25390595122072 | P10810, P63085, O70145, Q9R0C8, P25911, P08556, P63017, Q8R180 | 0.710396438 |
| KEGG_PATHWAY | mmu04611:Platelet activation | 6 | 0.27275027959906 | P63085, P20491, P09055, P25911, P62137, P08752 | 0.7219590359 |
| KEGG_PATHWAY | mmu05167:Kaposi sarcoma-associated herpesvirus infection | 9 | 0.27725423051104 | P63085, P01900, Q69ZK0, Q9WVL2, P01897, P25911, P08556, O08734 | 0.7219590359 |
| KEGG_PATHWAY | mmu04720:Long-term potentiation | 4 | 0.28524182654815 | P63085, P08556, P62137 | 0.7354764743 |
| KEGG_PATHWAY | mmu05231:Choline metabolism in cancer | 5 | 0.28976690681991 | Q8BH43, P63085, P08556, Q05144 | 0.7398902572 |
| KEGG_PATHWAY | mmu05219:Bladder cancer | 3 | 0.31609465045572 | P63085, P08556 | 0.7803282278 |
| KEGG_PATHWAY | mmu04915:Estrogen signaling pathway | 6 | 0.31610684098916 | P30416, P63085, P08556, P63017, P08752 | 0.7803282278 |
| KEGG_PATHWAY | mmu04210:Apoptosis | 6 | 0.321595502907 | Q9Z0X1, P21619, P63085, P08556, O08734 | 0.7803282278 |
| KEGG_PATHWAY | mmu05218:Melanoma | 4 | 0.32344517477961 | P63085, P08556, O08734 | 0.7803282278 |
| KEGG_PATHWAY | mmu05223:Non-small cell lung cancer | 4 | 0.32344517477961 | P63085, P08556, O08734 | 0.7803282278 |
| KEGG_PATHWAY | mmu05214:Glioma | 4 | 0.3387540215542 | P63085, P08556, O08734 | 0.787523842 |
| KEGG_PATHWAY | mmu04910:Insulin signaling pathway | 6 | 0.34918174945817 | P63085, O08528, P08556, P53657, P62137 | 0.787523842 |
| KEGG_PATHWAY | mmu05170:Human immunodeficiency virus 1 infection | 9 | 0.35034330615339 | P63085, P01900, P01897, P18760, P08556, O08734, P08752, Q05144 | 0.787523842 |
| KEGG_PATHWAY | mmu05225:Hepatocellular carcinoma | 7 | 0.3593219150146 | P63085, Q9JLT4, Q9JMH6, P48774, P08556, O08734 | 0.7941316273 |
| KEGG_PATHWAY | mmu01524:Platinum drug resistance | 4 | 0.39202756271996 | P63085, P48774, O08734, P43247 | 0.8309757224 |
| KEGG_PATHWAY | mmu04218:Cellular senescence | 7 | 0.39423348710405 | Q3UMR5, P63085, P01900, P01897, P08556, P62137 | 0.8309757224 |
| KEGG_PATHWAY | mmu04012:ErbB signaling pathway | 4 | 0.41455806592095 | P63085, P08556, P00520 | 0.8458103253 |
| KEGG_PATHWAY | mmu04660:T cell receptor signaling pathway | 5 | 0.44114070852034 | P63085, Q9R0C8, P08556, Q76MZ3 | 0.877308973 |
| KEGG_PATHWAY | mmu05207:Chemical carcinogenesis - receptor activation | 8 | 0.44271173187406 | P63085, P52293, P48774, P70168, P08556, Q6ZQM8, P08752 | 0.877308973 |
| KEGG_PATHWAY | mmu05235:PD-L1 expression and PD-1 checkpoint pathway in cancer | 4 | 0.44417074985834 | P63085, P67871, P08556 | 0.877308973 |
| KEGG_PATHWAY | mmu04010:MAPK signaling pathway | 10 | 0.4508902410179 | P10810, P63085, P09581, Q8BTM8, P08556, P63017, Q9D7X3, Q05144, P54227 | 0.8784009881 |
| KEGG_PATHWAY | mmu05206:MicroRNAs in cancer | 10 | 0.46962664969495 | P61979, P63085, P20152, P13864, P08556, O08734, P00520, P54227, P26645 | 0.8855218855 |

| | | | | | |
|---|---|---|---|---|---|
| KEGG_PATHWAY | mmu04140:Autophagy - animal | 6 | 0.49698500153735 | P17047, Q99JX3, P63085, Q9D906, P08556 | 0.8855218855 |
| KEGG_PATHWAY | mmu05166:Human T-cell leukemia virus 1 infection | 8 | 0.52585186623809 | P63085, P34022, P01900, P11835, P01897, P35564, P08556 | 0.8855218855 |
| KEGG_PATHWAY | mmu04929:GnRH secretion | 3 | 0.5262211326949 | P63085, P08556 | 0.8855218855 |
| KEGG_PATHWAY | mmu04916:Melanogenesis | 4 | 0.52900755513884 | P63085, P08556, P08752 | 0.8855218855 |
| KEGG_PATHWAY | mmu05165:Human papillomavirus infection | 11 | 0.54295747111094 | Q64339, P63085, Q8BVE3, P09055, P01900, Q9WVL2, P01897, P08556, O08734, Q76MZ3 | 0.8855218855 |
| KEGG_PATHWAY | mmu05163:Human cytomegalovirus infection | 8 | 0.56309113452227 | P63085, P01900, P01897, P08556, O08734, P08752, Q05144 | 0.8855218855 |
| KEGG_PATHWAY | mmu05211:Renal cell carcinoma | 3 | 0.58354607203801 | P63085, P08556 | 0.8855218855 |
| KEGG_PATHWAY | mmu04725:Cholinergic synapse | 4 | 0.61222770468822 | P63085, P08556, P08752 | 0.8855218855 |
| KEGG_PATHWAY | mmu04917:Prolactin signaling pathway | 3 | 0.613943190429 | P63085, P08556 | 0.8855218855 |
| KEGG_PATHWAY | mmu04921:Oxytocin signaling pathway | 5 | 0.61902839480774 | P63085, P08556, P62137, P08752 | 0.8855218855 |
| KEGG_PATHWAY | mmu05212:Pancreatic cancer | 3 | 0.62849554179651 | P63085, O08734, Q05144 | 0.8855218855 |
| KEGG_PATHWAY | mmu04935:Growth hormone synthesis, secretion and action | 4 | 0.6357718217724 | P63085, P08556, P08752 | 0.8855218855 |
| KEGG_PATHWAY | mmu04150:mTOR signaling pathway | 5 | 0.64423856586623 | P63085, Q8BVE3, P08556, Q9CQ22 | 0.8855218855 |
| KEGG_PATHWAY | mmu04014:Ras signaling pathway | 7 | 0.64513797566283 | P63085, P09581, Q9CQD1, P08556, P00520, Q05144 | 0.8855218855 |
| KEGG_PATHWAY | mmu04919:Thyroid hormone signaling pathway | 4 | 0.65276776692108 | O55143, P63085, P08556 | 0.8855218855 |
| KEGG_PATHWAY | mmu01521:EGFR tyrosine kinase inhibitor resistance | 3 | 0.65631777989802 | P63085, P08556 | 0.8855218855 |
| KEGG_PATHWAY | mmu04722:Neurotrophin signaling pathway | 4 | 0.6583061316852 | P63085, P08556, P00520 | 0.8855218855 |
| KEGG_PATHWAY | mmu05200:Pathways in cancer | 15 | 0.66701998821177 | Q9JMH6, Q9WVL2, O08734, Q05144, P21107, P63085, Q9JLT4, P09055, P09581, P48774, P08556, P00520, P43247, P08752 | 0.8855218855 |
| KEGG_PATHWAY | mmu05161:Hepatitis B | 5 | 0.67301713251657 | Q9CQV8, P63085, Q9WVL2, P08556 | 0.8855218855 |
| KEGG_PATHWAY | mmu04926:Relaxin signaling pathway | 4 | 0.70529150390023 | P63085, P08556, P08752 | 0.8855218855 |
| KEGG_PATHWAY | mmu04022:cGMP-PKG signaling pathway | 5 | 0.70883746081408 | O55143, P63085, G5E829, P62137, P08752 | 0.8855218855 |
| KEGG_PATHWAY | mmu04912:GnRH signaling pathway | 3 | 0.71859658150703 | P63085, P08556 | 0.8855218855 |
| KEGG_PATHWAY | mmu04726:Serotonergic synapse | 4 | 0.73378202783741 | P63085, P08556, P08752 | 0.8855218855 |
| KEGG_PATHWAY | mmu01522:Endocrine resistance | 3 | 0.73533689645308 | P63085, P08556 | 0.8855218855 |
| KEGG_PATHWAY | mmu04371:Apelin signaling pathway | 4 | 0.7471953089087 | P63085, P08556, P08752 | 0.8855218855 |
| KEGG_PATHWAY | mmu05215:Prostate cancer | 3 | 0.77112940280333 | P63085, P08556 | 0.8855218855 |
| KEGG_PATHWAY | mmu04933:AGE-RAGE signaling pathway in diabetic complications | 3 | 0.77588444770076 | P63085, P08556 | 0.8855218855 |
| KEGG_PATHWAY | mmu05224:Breast cancer | 4 | 0.78033733703771 | P63085, P08556, O08734 | 0.8855218855 |
| KEGG_PATHWAY | mmu05142:Chagas disease | 3 | 0.78513675379459 | P63085, P08752, Q76MZ3 | 0.8855218855 |
| KEGG_PATHWAY | mmu04072:Phospholipase D signaling pathway | 4 | 0.78804349482387 | P63085, P20491, P08556 | 0.8855218855 |
| KEGG_PATHWAY | mmu05226:Gastric cancer | 4 | 0.79181212523259 | P63085, P08556, O08734 | 0.8855218855 |
| KEGG_PATHWAY | mmu05034:Alcoholism | 5 | 0.83205217823286 | P63085, P08556, P62137, P08752 | 0.8855218855 |
| KEGG_PATHWAY | mmu04068:FoxO signaling pathway | 3 | 0.88850967699058 | P63085, P08556 | 0.888509677 |
| KEGG_PATHWAY | mmu04550:Signaling pathways regulating pluripotency of stem cells | 3 | 0.90485516634526 | P63085, P08556 | 0.9048551663 |
| KEGG_PATHWAY | mmu04151:PI3K-Akt signaling pathway | 7 | 0.94576965928179 | Q9CQV8, P63085, P09055, P09581, P08556, Q76MZ3 | 0.9457696593 |
| Annotation Cluster 68 | Enrichment Score: 0.5640160483155834 | | | | |
| Category | Term | Count | P Value | Genes | FDR |
| KEGG_PATHWAY | mmu05140:Leishmaniasis | 6 | 0.04474856469044 | P05555, P26151, P63085, P09055, O70145, P11835 | 0.2374874645 |
| KEGG_PATHWAY | mmu04613:Neutrophil extracellular trap formation | 7 | 0.51882814500154 | P05555, P26151, P63085, O70145, P11835, Q9D906, Q05144 | 0.8855218855 |
| KEGG_PATHWAY | mmu05150:Staphylococcus aureus infection | 3 | 0.87528445971954 | P05555, P26151, P11835 | 0.8855218855 |
| Annotation Cluster 69 | Enrichment Score: 0.5566188189024477 | | | | |
| Category | Term | Count | P Value | Genes | FDR |
| GOTERM_BP_DIRECT | GO:0050853~B cell receptor signaling pathway | 6 | 0.00024596044811 | P63085, Q8K1X4, Q9R0C8, Q6A0D4, P25911, P00520 | 0.0192673144 |
| GOTERM_BP_DIRECT | GO:0001934~positive regulation of protein phosphorylation | 9 | 0.02204524887384 | Q9DCE9, P57759, O55222, P09581, P25911, P08556, P00520, Q9CPT4, P17742 | 0.3560232709 |
| UP_KW_DISEASE | KW-0656~Proto-oncogene | 6 | 0.0298469000567 | Q60875, P09581, P25911, P08556, P00520 | 0.0596938001 |
| GOTERM_MF_DIRECT | GO:0004713~protein tyrosine kinase activity | 4 | 0.13084527319795 | P09581, P25911, Q91YR1, P00520 | 0.5914045932 |
| GOTERM_BP_DIRECT | GO:0018108~peptidyl-tyrosine phosphorylation | 3 | 0.30828317906006 | P09581, P25911, P00520 | 0.9757501028 |
| SMART | SM00219:TyrKc | 3 | 0.4154829407819 | P09581, P25911, P00520 | 0.9943820225 |
| INTERPRO | IPR020635:Tyr_kinase_cat_dom | 3 | 0.42757714655698 | P09581, P25911, P00520 | 0.9862825789 |
| INTERPRO | IPR001245:Ser-Thr/Tyr_kinase_cat_dom | 4 | 0.44758558546068 | O55222, P09581, P25911, P00520 | 0.9862825789 |
| GOTERM_MF_DIRECT | GO:0140801~histone H2AXY142 kinase activity | 3 | 0.44932586658479 | P09581, P25911, P00520 | 0.9234393404 |
| GOTERM_MF_DIRECT | GO:0035401~histone H3Y41 kinase activity | 3 | 0.44932586658479 | P09581, P25911, P00520 | 0.9234393404 |
| GOTERM_BP_DIRECT | GO:0006468~protein phosphorylation | 6 | 0.46292592591572 | O55222, P63085, P18052, P18760, P25911, P00520 | 0.9757501028 |
| INTERPRO | IPR008266:Tyr_kinase_AS | 3 | 0.53093365695147 | P09581, P25911, P00520 | 0.9862825789 |
| GOTERM_BP_DIRECT | GO:0046777~protein autophosphorylation | 3 | 0.58593192323367 | P09581, P25911, P00520 | 0.9757501028 |
| UP_KW_MOLECULAR_FUNCTION | KW-0829~Tyrosine-protein kinase | 3 | 0.70333897007661 | P09581, P25911, P00520 | 0.8783783784 |
| GOTERM_MF_DIRECT | GO:0004672~protein kinase activity | 4 | 0.79950700597895 | O55222, P63085, P25911, P00520 | 0.9234393404 |
| UP_KW_MOLECULAR_FUNCTION | KW-0418~Kinase | 11 | 0.94057223537226 | Q9WV85, O55222, P63085, Q3U5Q7, P09581, Q9WTP7, O08528, P09411, P25911, P53657, P00520 | 0.9405722354 |
| INTERPRO | IPR017441:Protein_kinase_ATP_BS | 4 | 0.97140777843338 | P63085, P09581, P25911, P00520 | 0.9862825789 |
| UP_SEQ_FEATURE | DOMAIN:Protein kinase | 5 | 0.9775836709996 | O55222, P63085, P09581, P25911, P00520 | 0.9909502262 |
| INTERPRO | IPR000719:Prot_kinase_dom | 5 | 0.98240413599727 | O55222, P63085, P09581, P25911, P00520 | 0.9862825789 |
| INTERPRO | IPR011009:Kinase-like_dom_sf | 5 | 0.99002440172172 | O55222, P63085, P09581, P25911, P00520 | 0.9900244017 |
| Annotation Cluster 70 | Enrichment Score: 0.4580148370059564 | | | | |
| Category | Term | Count | P Value | Genes | FDR |
| BIOCARTA | m_ecmPathway:Erk and PI-3 Kinase Are Necessary for Collagen Binding in Corneal | 4 | 0.07660192374464 | P63085, P09055, P08556, P62962 | 1 |
| BIOCARTA | m_tffPathway:Trefoil Factors Initiate Mucosal Healing | 3 | 0.29879795890462 | P63085, P09055, P08556 | 1 |
| BIOCARTA | m_erkPathway:Erk1/Erk2 Mapk Signaling pathway | 3 | 0.38908131356647 | P63085, P09055, P08556 | 1 |
| BIOCARTA | m_integrinPathway:Integrin Signaling Pathway | 3 | 0.46050458866124 | P63085, P09055, P08556 | 1 |
| BIOCARTA | m_metPathway:Signaling of Hepatocyte Growth Factor Receptor | 3 | 0.46050458866124 | P63085, P09055, P08556 | 1 |
| KEGG_PATHWAY | mmu04151:PI3K-Akt signaling pathway | 7 | 0.94576965928179 | Q9CQV8, P63085, P09055, P09581, P08556, Q76MZ3 | 0.9457696593 |
| Annotation Cluster 71 | Enrichment Score: 0.3969782338382424 | | | | |
| Category | Term | Count | P Value | Genes | FDR |
| GOTERM_MF_DIRECT | GO:0005085~guanyl-nucleotide exchange factor activity | 7 | 0.08504391384258 | Q60875, Q8C147, Q69ZK0, Q9R0C8, P57776, Q9CQ22, Q6PAR5 | 0.4357805781 |
| SMART | SM00325:RhoGEF | 3 | 0.30971049088837 | Q60875, Q69ZK0, Q9R0C8 | 0.9943820225 |
| UP_SEQ_FEATURE | DOMAIN:DH | 3 | 0.31938263240036 | Q60875, Q69ZK0, Q9R0C8 | 0.9909502262 |
| INTERPRO | IPR000219:DH-domain | 3 | 0.33956465518454 | Q60875, Q69ZK0, Q9R0C8 | 0.9862825789 |
| INTERPRO | IPR035899:DBL_dom_sf | 3 | 0.33956465518454 | Q60875, Q69ZK0, Q9R0C8 | 0.9862825789 |
| UP_KW_MOLECULAR_FUNCTION | KW-0344~Guanine-nucleotide releasing factor | 5 | 0.49288957669382 | Q60875, Q8C147, Q69ZK0, Q9R0C8, Q6PAR5 | 0.8783783784 |
| SMART | SM00233:PH | 5 | 0.6462103803798 | Q60875, Q69ZK0, Q3UND0, Q9R0C8, Q8K124 | 0.9943820225 |
| UP_SEQ_FEATURE | DOMAIN:PH | 5 | 0.66043821884683 | Q60875, Q69ZK0, Q3UND0, Q9R0C8, Q8K124 | 0.9909502262 |
| INTERPRO | IPR001849:PH_domain | 5 | 0.68735669914689 | Q60875, Q69ZK0, Q3UND0, Q9R0C8, Q8K124 | 0.9862825789 |
| INTERPRO | IPR011993:PH-like_dom_sf | 7 | 0.76440641172123 | Q60875, P26041, P34022, Q69ZK0, Q3UND0, Q9R0C8, Q8K124 | 0.9862825789 |
| Annotation Cluster 72 | Enrichment Score: 0.3648267096222603 | | | | |
| Category | Term | Count | P Value | Genes | FDR |
| GOTERM_BP_DIRECT | GO:0006281~DNA repair | 8 | 0.08772344277176 | Q9WV85, P23492, P62908, P56960, P00520, P43247, P07742, Q9D0F6 | 0.7017410113 |
| UP_KW_BIOLOGICAL_PROCESS | KW-0234~DNA repair | 5 | 0.93672242235268 | P62908, P56960, Q99KP6, P00520, P43247 | 0.9367224224 |
| UP_KW_BIOLOGICAL_PROCESS | KW-0227~DNA damage | 5 | 0.97902388545753 | P62908, P56960, Q99KP6, P00520, P43247 | 0.9790238855 |
| Annotation Cluster 73 | Enrichment Score: 0.3296763113064824 | | | | |
| Category | Term | Count | P Value | Genes | FDR |
| INTERPRO | IPR020472:G-protein_beta_WD-40_rep | 4 | 0.22305095756981 | Q8CIE6, Q99KP6, Q3UKJ7, O88342 | 0.9862825789 |
| UP_SEQ_FEATURE | REPEAT:WD 4 | 6 | 0.38580278382223 | Q3UJB9, Q8CIE6, Q99KP6, Q5SSI6, Q3UKJ7, O88342 | 0.9909502262 |
| UP_SEQ_FEATURE | REPEAT:WD 6 | 5 | 0.38933039960653 | Q8CIE6, Q99KP6, Q5SSI6, Q3UKJ7, O88342 | 0.9909502262 |
| UP_KW_DOMAIN | KW-0853~WD repeat | 6 | 0.41887718603353 | Q3UJB9, Q8CIE6, Q99KP6, Q5SSI6, Q3UKJ7, O88342 | 1 |
| UP_SEQ_FEATURE | REPEAT:WD 3 | 6 | 0.42256827115408 | Q3UJB9, Q8CIE6, Q99KP6, Q5SSI6, Q3UKJ7, O88342 | 0.9909502262 |
| UP_SEQ_FEATURE | REPEAT:WD 1 | 6 | 0.43580554464648 | Q3UJB9, Q8CIE6, Q99KP6, Q5SSI6, Q3UKJ7, O88342 | 0.9909502262 |
| UP_SEQ_FEATURE | REPEAT:WD 2 | 6 | 0.43580554464648 | Q3UJB9, Q8CIE6, Q99KP6, Q5SSI6, Q3UKJ7, O88342 | 0.9909502262 |
| INTERPRO | IPR019775:WD40_repeat_CS | 4 | 0.45220730021851 | Q8CIE6, Q99KP6, Q3UKJ7, O88342 | 0.9862825789 |
| SMART | SM00320:WD40 | 6 | 0.49162330362325 | Q3UJB9, Q8CIE6, Q99KP6, Q5SSI6, Q3UKJ7, O88342 | 0.9943820225 |
| INTERPRO | IPR001680:WD40_rpt | 6 | 0.51067467584419 | Q3UJB9, Q8CIE6, Q99KP6, Q5SSI6, Q3UKJ7, O88342 | 0.9862825789 |
| UP_SEQ_FEATURE | REPEAT:WD 5 | 5 | 0.53315223343125 | Q8CIE6, Q99KP6, Q5SSI6, Q3UKJ7, O88342 | 0.9909502262 |
| INTERPRO | IPR015943:WD40/YVTN_repeat-like_dom_sf | 7 | 0.59959252043632 | Q3UJB9, B2RXS4, Q8CIE6, Q99KP6, Q5SSI6, Q3UKJ7, O88342 | 0.9862825789 |
| UP_SEQ_FEATURE | REPEAT:WD | 4 | 0.60379950961758 | Q3UJB9, Q8CIE6, Q5SSI6, O88342 | 0.9909502262 |
| INTERPRO | IPR036322:WD40_repeat_dom_sf | 6 | 0.64161867002413 | Q3UJB9, Q8CIE6, Q99KP6, Q5SSI6, Q3UKJ7, O88342 | 0.9862825789 |
| UP_SEQ_FEATURE | REPEAT:WD 7 | 3 | 0.71648471696161 | Q99KP6, Q3UKJ7, O88342 | 0.9909502262 |
| Annotation Cluster 74 | Enrichment Score: 0.31650528829363206 | | | | |
| Category | Term | Count | P Value | Genes | FDR |
| KEGG_PATHWAY | mmu04261:Adrenergic signaling in cardiomyocytes | 7 | 0.26657718035113 | P21107, O55143, P63085, G5E829, P62137, P08752, Q76MZ3 | 0.7219590359 |
| KEGG_PATHWAY | mmu04024:cAMP signaling pathway | 7 | 0.59444886738164 | O55143, P63085, Q9R0C8, G5E829, P62137, P08752, Q05144 | 0.8855218855 |
| KEGG_PATHWAY | mmu04022:cGMP-PKG signaling pathway | 5 | 0.70883746081408 | O55143, P63085, G5E829, P62137, P08752 | 0.8855218855 |
| Annotation Cluster 75 | Enrichment Score: 0.13501818871364404 | | | | |
| Category | Term | Count | P Value | Genes | FDR |
| KEGG_PATHWAY | mmu04380:Osteoclast differentiation | 7 | 0.1775002186131 | O55143, P26151, P63085, O70145, P09581, P97797, Q9WVL2 | 0.5835319687 |
| INTERPRO | IPR003006:Ig/MHC_CS | 3 | 0.51460047989865 | P01900, P97797, P01897 | 0.9862825789 |
| SMART | SM00407:IGc1 | 3 | 0.58917099404249 | P01900, P97797, P01897 | 0.9943820225 |
| INTERPRO | IPR003597:Ig_C1-set | 3 | 0.61156812604316 | P01900, P97797, P01897 | 0.9862825789 |
| UP_KW_DOMAIN | KW-0393~Immunoglobulin domain | 3 | 0.99520519590962 | P26151, P09581, P97797 | 1 |
| UP_SEQ_FEATURE | DOMAIN:Ig-like | 5 | 0.99956971542535 | P26151, P09581, P01900, P97797, P01897 | 0.9995697154 |
| SMART | SM00409:IG | 3 | 0.999779975369 | P26151, P09581, P97797 | 0.9997799754 |
| INTERPRO | IPR003599:Ig_sub | 3 | 0.99980450151256 | P26151, P09581, P97797 | 0.9998045015 |
| INTERPRO | IPR013783:Ig-like_fold | 10 | 0.99981637301634 | B2RXS4, Q9WV55, P26151, P09581, P01900, P97797, Q8BTM8, P01897, Q2YFS3, Q9QY76 | 0.999816373 |
| INTERPRO | IPR036179:Ig-like_dom_sf | 6 | 0.99996704424647 | P26151, P09581, P01900, P97797, P01897, Q2YFS3 | 0.9999670442 |
| INTERPRO | IPR007110:Ig-like_dom | 5 | 0.99998495712939 | P26151, P09581, P01900, P97797, P01897 | 0.9999849571 |
| Annotation Cluster 76 | Enrichment Score: 0.07318509722915362 | | | | |
| Category | Term | Count | P Value | Genes | FDR |
| UP_KW_MOLECULAR_FUNCTION | KW-0678~Repressor | 13 | 0.60356478892118 | Q921F2, Q3UZ39, Q8BL97, P61979, Q7TPV4, P13864, Q9DBD5, Q64213, Q3UEB3, O35129, P62960, Q9CRB9, P63017 | 0.8783783784 |

| | | | | | |
|---|---|---|---|---|---|
| UP_KW_BIOLOGICAL_PROCESS | KW-0804~Transcription | 26 | 0.99955751459311 | Q921F2, P10639, Q61656, O70133, Q7TPV4, Q501J6, Q9WVL2, P13864, Q9DBD5, O35129, P68037, Q9CRB9, Q2EMV9, P62908, Q3UZ39, P61979, Q9CXT6, Q64213, P57776, Q3UEB3, P62960, Q8C2Q3, O35892, Q3U0V1, P63017, P08775 | 0.9995575146 |
| UP_KW_BIOLOGICAL_PROCESS | KW-0805~Transcription regulation | 24 | 0.99979975056818 | Q921F2, P62908, P10639, Q3UZ39, Q61656, P61979, Q9CXT6, O70133, Q7TPV4, Q501J6, Q9WVL2, P13864, P57776, Q64213, Q3UEB3, O35129, P62960, P68037, Q8C2Q3, Q9CRB9, Q2EMV9, Q3U0V1, O35892, P63017 | 0.9997997506 |
| Annotation Cluster 77 | Enrichment Score: 0.0639077820181816 | | | | |
| Category | Term | Count | P Value | Genes | FDR |
| UP_KW_BIOLOGICAL_PROCESS | KW-0109~Calcium transport | 3 | 0.65238038051472 | Q3UMR5, O55143, G5E829 | 0.9278350515 |
| UP_KW_BIOLOGICAL_PROCESS | KW-0406~Ion transport | 7 | 0.99170848502641 | Q3UMR5, O55143, Q8BVE3, Q03265, G5E829, Q8K3C0, Q61792 | 0.991708485 |
| KEGG_PATHWAY | mmu04020:Calcium signaling pathway | 3 | 0.99401224215905 | Q3UMR5, O55143, G5E829 | 0.9940122422 |
| Annotation Cluster 78 | Enrichment Score: 0.03142638002151806 | | | | |
| Category | Term | Count | P Value | Genes | FDR |
| UP_SEQ_FEATURE | REPEAT:LRR 6 | 3 | 0.85239742719732 | P46061, P10810, Q91VI7 | 0.9909502262 |
| UP_SEQ_FEATURE | REPEAT:LRR 5 | 3 | 0.89738682373574 | P46061, P10810, Q91VI7 | 0.9909502262 |
| UP_SEQ_FEATURE | REPEAT:LRR 4 | 3 | 0.91765859134793 | P46061, P10810, Q91VI7 | 0.9909502262 |
| UP_SEQ_FEATURE | REPEAT:LRR 3 | 3 | 0.93319569886029 | P46061, P10810, Q91VI7 | 0.9909502262 |
| UP_SEQ_FEATURE | REPEAT:LRR 2 | 3 | 0.94278260265385 | P46061, P10810, Q91VI7 | 0.9909502262 |
| UP_SEQ_FEATURE | REPEAT:LRR 1 | 3 | 0.94358624248693 | P46061, P10810, Q91VI7 | 0.9909502262 |
| INTERPRO | IPR001611:Leu-rich_rpt | 3 | 0.94786860533173 | P46061, P10810, Q91VI7 | 0.9862825789 |
| UP_KW_DOMAIN | KW-0433~Leucine-rich repeat | 3 | 0.94884587568796 | P46061, P10810, Q91VI7 | 1 |
| INTERPRO | IPR032675:LRR_dom_sf | 3 | 0.9948324075302 | P46061, P10810, Q91VI7 | 0.9948324075 |
| Annotation Cluster 79 | Enrichment Score: 0.018318498356488053 | | | | |
| Category | Term | Count | P Value | Genes | FDR |
| UP_KW_BIOLOGICAL_PROCESS | KW-0524~Neurogenesis | 5 | 0.84534195033915 | Q60875, Q7TMB8, Q99P72, P54227, P28656 | 0.9278350515 |
| UP_KW_BIOLOGICAL_PROCESS | KW-0221~Differentiation | 7 | 0.99929627286397 | Q99JX3, Q60875, Q7TMB8, P16110, Q9ET26, Q8VBZ3, P54227 | 0.9992962729 |
| UP_KW_MOLECULAR_FUNCTION | KW-0217~Developmental protein | 6 | 0.99999921421415 | B2RXS4, Q60875, Q7TMB8, Q9ET26, Q8VBZ3, P54227 | 0.9999992142 |
| UP_KW_MOLECULAR_FUNCTION | KW-9996~Developmental protein | 6 | 0.99999921421415 | B2RXS4, Q60875, Q7TMB8, Q9ET26, Q8VBZ3, P54227 | 0.9999992142 |
| Annotation Cluster 80 | Enrichment Score: 0.011695586694326186 | | | | |
| Category | Term | Count | P Value | Genes | FDR |
| UP_SEQ_FEATURE | TOPO_DOM:Cytoplasmic | 48 | 0.9314062525905 | Q91ZV7, Q9CQW9, Q9WV55, P49300, Q9CZW5, Q9QY76, Q8VCH8, P09055, Q80WQ6, Q9DC16, Q9QUJ7, O35609, Q80VA0, Q9CYN2, P01897, G5E829, Q78T54, P41216, P05555, Q61543, Q9EQH2, Q69ZN7, P20491, P26151, Q99J93, Q8BI84, P18052, P11835, P97797, P35564, Q99P72, Q2YFS3, P13379, Q9DBG6, Q8VDP6, B2RXS4, P17047, O55143, P60060, O35114, Q8BMK4, Q61072, P09581, P01900, Q8VBZ3, P24668, Q8VBT0, Q8BG07 | 0.9909502262 |
| UP_KW_DOMAIN | KW-1133~Transmembrane helix | 79 | 0.94941957003643 | Q9D0M3, Q8CAQ8, Q921H8, P49300, Q9CZW5, P48771, Q69ZK0, Q9CR62, Q61033, Q8VC04, Q9QY76, Q9DBL1, Q3UMR5, P09055, Q80WQ6, Q9DC16, O35609, Q78T54, P41216, P47758, Q8R1I1, P20491, P26151, P18052, P97797, Q9ET30, P35564, Q9CQX2, Q99P72, Q2YFS3, Q9DBG6, B2RXS4, P17047, O55143, P60060, O35114, P06151, Q8BMK4, P09581, P01900, Q8VBZ3, Q9CWG1, Q91ZV7, Q62425, Q9CQW9, Q9WV55, Q9ERS2, P70245, Q8R127, Q9QUJ7, Q80VA0, Q9CYN2, Q8K1X4, P01897, G5E829, Q6ZQM8, P05555, Q61543, Q9EQH2, Q61029, Q69ZN7, Q9QZD8, Q9CQN6, Q99J93, Q8BI84, P11835, P59017, Q8K3C0, O08734, P13379, Q8VDP6, P17665, Q9Z0X1, Q61072, Q78IK4, Q8VBT0, P24668, O70503, P47802, Q8BG07 | 1 |
| UP_KW_CELLULAR_COMPONENT | KW-0472~Membrane | 162 | 0.99222233905541 | Q8CAQ8, Q921H8, Q8BVE3, P49300, Q9CZW5, Q9CR62, P18760, O70493, Q6A0D4, Q61033, P19973, P17426, Q8JZN5, Q6PAR5, Q9DBL1, O35129, D3Z6Q9, P62075, Q9CQD1, P20152, P47758, Q9DCE9, P26151, Q9ET30, Q03265, Q99P72, P17047, P60060, O55143, O35114, P06151, P01900, Q8BMK4, Q8VBZ3, Q8BMS1, Q9CWG1, Q91ZR2, P26883, Q9WV55, Q91WD5, Q9CQ22, Q80VA0, Q8R127, Q8K1X4, P01897, Q8BX70, Q91XV3, P05555, Q61543, Q69ZN7, Q61029, P10107, Q99J93, Q8BI84, P11835, A2AQP0, P59017, Q8K3C0, O08734, Q05144, P17665, Q76MZ3, Q9JL26, Q78IK4, P08556, O70503, P56391, P26645, Q6P069, Q8BT60, Q9D0M3, Q91VH2, P48771, Q69ZK0, Q8VBW6, Q3UIA2, Q7TMF3, Q8VC04, P97429, Q9QY76, Q8VCH8, Q3UMR5, Q8CIE6, P63085, Q8BLN5, Q9CRB9, P09055, Q80WQ6, Q9DC16, O35609, Q78T54, Q9DCR2, P41216, Q8R1I1, Q9WV85, Q8BH43, Q99JX3, P20491, P18052, P97797, P35564, Q9Z0E6, Q9CQX2, Q2YFS3, Q8R180, P36536, Q9DBG6, B2RXS4, O55222, Q8C147, P09581, P09103, P63017, E9Q634, Q91ZV7, Q7TPR4, Q62425, Q9CQW9, P26041, P10810, Q9ERS2, O08528, Q64442, P70245, Q80X82, Q00612, Q9QUJ7, Q9CYN2, G5E829, Q9DC70, Q6ZQM8, Q8VEB4, Q91YT0, Q62159, P62908, Q9EQH2, Q9CWZ7, Q9CQN1, Q9EQH3, E9Q7G0, Q8BRF7, Q9QZD8, Q9CQN6, P51175, Q9EQP2, P13379, Q62433, Q8VDP6, Q8CHH9, P21956, Q9Z0X1, Q61072, Q91V41, P17182, Q8VBT0, P24668, P25911, P47802, P08752, Q8BG07 | 0.9922223391 |
| UP_KW_DOMAIN | KW-0812~Transmembrane | 79 | 0.99613000932157 | Q9D0M3, Q8CAQ8, Q921H8, P49300, Q9CZW5, P48771, Q69ZK0, Q9CR62, Q61033, Q8VC04, Q9QY76, Q9DBL1, Q3UMR5, P09055, Q80WQ6, Q9DC16, O35609, Q78T54, P41216, P47758, Q8R1I1, P20491, P26151, P18052, P97797, Q9ET30, P35564, Q9CQX2, Q99P72, Q2YFS3, Q9DBG6, B2RXS4, P17047, O55143, P60060, O35114, P06151, Q8BMK4, P09581, P01900, Q8VBZ3, Q9CWG1, Q91ZV7, Q62425, Q9CQW9, Q9WV55, Q9ERS2, P70245, Q8R127, Q9QUJ7, Q80VA0, Q9CYN2, Q8K1X4, P01897, G5E829, Q6ZQM8, P05555, Q61543, Q9EQH2, Q61029, Q69ZN7, Q9QZD8, Q9CQN6, Q99J93, Q8BI84, P11835, P59017, Q8K3C0, O08734, P13379, Q8VDP6, P17665, Q9Z0X1, Q61072, Q78IK4, Q8VBT0, P24668, O70503, P47802, Q8BG07 | 1 |
| UP_SEQ_FEATURE | TRANSMEM:Helical | 73 | 0.99999934065362 | Q9D0M3, Q8CAQ8, Q921H8, P49300, Q9CZW5, P48771, Q69ZK0, Q61033, Q8VC04, Q9QY76, Q9DBL1, Q3UMR5, P09055, Q80WQ6, Q9DC16, O35609, Q78T54, P41216, P47758, Q8R1I1, P20491, P26151, P18052, P97797, Q9ET30, P35564, Q9CQX2, Q99P72, Q2YFS3, Q9DBG6, B2RXS4, P17047, O55143, P60060, O35114, P06151, Q8BMK4, P09581, P01900, Q8VBZ3, Q9CWG1, Q91ZV7, Q62425, Q9CQW9, Q9WV55, Q9ERS2, P70245, Q8R127, Q9CYN2, Q8K1X4, P01897, G5E829, Q6ZQM8, P05555, Q61543, Q61029, Q69ZN7, Q9CQN6, Q99J93, Q8BI84, P11835, P59017, Q8K3C0, O08734, P13379, Q8VDP6, P17665, Q9Z0X1, Q61072, Q78IK4, Q8VBT0, P24668, O70503, P47802 | 0.9999993407 |
| Annotation Cluster 81 | Enrichment Score: 3.719665980094597E-4 | | | | |
| Category | Term | Count | P Value | Genes | FDR |
| UP_SEQ_FEATURE | CARBOHYD:N-linked (GlcNAc...) asparagine | 45 | 0.99775247508044 | Q91ZV7, Q9Z0J0, P10810, P49300, P20060, P34960, Q9ESY9, Q62087, Q07797, Q9WVJ3, P09055, Q9DC16, P01897, O09043, Q6ZQM8, Q8VEB4, O89023, P05555, Q61543, Q9EQH2, P26151, Q61207, Q8BI84, P18052, P11835, P97797, Q9ET30, Q2YFS3, P13379, Q8R180, Q9DBG6, P17742, B2RXS4, P21956, P17047, O35114, Q61072, P09581, P01900, Q8VBZ3, P24668, Q3TCN2, P29416, Q91ZW2, Q8BG07 | 0.9977524751 |
| UP_KW_DOMAIN | KW-0732~Signal | 56 | 0.99882501451688 | Q9D0M3, P34960, Q62087, Q07797, Q6PAR5, Q3UMR5, P09055, O09043, O89023, Q9WV85, P20491, P26151, Q61207, P18052, P97797, Q9ET30, P35564, Q2YFS3, Q8R180, Q9DBG6, B2RXS4, P16675, P17047, P16110, P09581, P01900, P09103, P29416, Q9CWG1, Q91ZW2, Q91ZV7, Q9Z0J0, P10810, P20060, Q9ESY9, Q9WVJ3, P01897, G5E829, Q6ZQM8, Q8VEB4, Q8CGK3, P05555, Q61543, P57759, Q9EQH2, Q8BI84, P51175, P11835, Q9D6Z1, P13379, Q9CPT4, P21956, Q61072, Q8VBT0, P24668, Q3TCN2 | 1 |
| UP_KW_PTM | KW-1015~Disulfide bond | 55 | 0.99999979173956 | P10639, P49300, Q9JMH6, P34960, Q62087, Q07797, Q9CRB9, P09055, P62075, O09043, O89023, P20491, P26151, Q61207, P97797, P35564, Q8R180, B2RXS4, P16675, P17047, O55143, P16110, O35114, P09581, P01900, P09103, P29416, Q8CDN6, Q91ZW2, Q9Z0J0, Q9WV55, P10810, P20060, Q9ESY9, Q9JLT4, Q80VA0, P01897, Q8VEB4, P05555, Q9EQH2, Q64339, P10107, P51175, P11835, P13379, P21956, Q05816, Q61072, P38060, Q8VBT0, P24668, Q3TCN2, P80318, P56391, P07742 | 1 |
| UP_KW_PTM | KW-0325~Glycoprotein | 53 | 1 | P49300, P34960, Q62087, Q07797, P09055, Q9DC16, O09043, P20152, P41216, O89023, P26151, Q61207, P18052, P97797, Q9ET30, Q03265, Q2YFS3, Q61881, Q8R180, Q9DBG6, B2RXS4, P16675, P17047, P21619, O35114, P09581, P01900, Q8VBZ3, P29416, Q91ZW2, Q91ZV7, Q9Z0J0, P10810, P20060, Q9ESY9, Q9WVJ3, P01897, Q6ZQM8, Q8VEB4, P05555, Q61543, Q9EQH2, P61979, E9Q7G0, Q8BI84, P11835, P13379, P17742, P21956, Q61072, P24668, Q3TCN2, Q8BG07 | 1 |
| Annotation Cluster 82 | Enrichment Score: 2.0677550881510817E-5 | | | | |
| Category | Term | Count | P Value | Genes | FDR |
| UP_SEQ_FEATURE | TRANSMEM:Helical | Name=6 | 0.65217391304348 | 0.999933757454269 | 1 |
| UP_SEQ_FEATURE | TRANSMEM:Helical | Name=5 | 0.65217391304348 | 0.999939913815654 | 1 |
| UP_SEQ_FEATURE | TRANSMEM:Helical | Name=3 | 0.65217391304348 | 0.999953696109178 | 1 |
| UP_SEQ_FEATURE | TRANSMEM:Helical | Name=4 | 0.65217391304348 | 0.999956620407172 | 1 |
| UP_SEQ_FEATURE | TRANSMEM:Helical | Name=1 | 0.65217391304348 | 0.999963751854307 | 1 |
| UP_SEQ_FEATURE | TRANSMEM:Helical | Name=2 | 0.65217391304348 | 0.999966596664148 | 1 |
