## supplemental tables 8 to 13 for "Effects of manganese dioxide on macrophages under different exposure schemes"

Supplementary Table 8: List of mitochondrial proteins modulated in response to at least one treatment
Modulated proteins appear with their name in color. Un-modulated but related proteins (e.g. other subunits of the same complex) appear in black

| accession | name | MWU 4x5 | MWU 4x10 | MWU 20R | ratio 4x5 | ratio 4x10 | ratio 20R |
|---|---|---|---|---|---|---|---|
| Q80W89 | NADH dehydrogenase [ubiquinone] 1 alpha subcomplex subunit 11 | 8 | 8 | 12 | 0.62 | 0.59 | 0.95 |
| Q7TMF3 | NADH dehydrogenase [ubiquinone] 1 alpha subcomplex subunit 12 | 3 | 0 | 8 | 0.39 | 1.93E-06 | 0.85 |
| Q9ERS2 | NADH dehydrogenase [ubiquinone] 1 alpha subcomplex subunit 13 | 0 | 0 | 3 | 0.45 | 0.36 | 0.75 |
| Q9CQ75 | NADH dehydrogenase [ubiquinone] 1 alpha subcomplex subunit 2 | 6.5 | 2.5 | 10 | 0.64 | 1.63E-06 | 1.18 |
| Q5BK63 | NADH dehydrogenase [ubiquinone] 1 alpha subcomplex subunit 9, mitochondrial | 3 | 1 | 4 | 0.22 | 0.16 | 0.34 |
| Q9DCS9 | NADH dehydrogenase [ubiquinone] 1 beta subcomplex subunit 10 | 6 | 8 | 11 | 0.46 | 0.60 | 1.18 |
| Q9CQH3 | NADH dehydrogenase [ubiquinone] 1 beta subcomplex subunit 5, mitochondrial | 9 | 6 | 9 | 0.53 | 0.19 | 0.47 |
| Q91VD9 | NADH-ubiquinone oxidoreductase 75 kDa subunit, mitochondrial | 3 | 9 | 1 | 0.60 | 0.78 | 0.63 |
| Q91WD5 | NADH dehydrogenase [ubiquinone] iron-sulfur protein 2, mitochondrial | 0 | 0 | 1 | 0.23 | 0.09 | 0.43 |
| Q9DCT2 | NADH dehydrogenase [ubiquinone] iron-sulfur protein 3, mitochondrial | 10 | 8 | 12 | 0.69 | 1.23 | 0.79 |
| Q9DC70 | NADH dehydrogenase [ubiquinone] iron-sulfur protein 7, mitochondrial | 2 | 0 | 6 | 0.49 | 0.25 | 1.27 |
| Q91YT0 | NADH dehydrogenase [ubiquinone] flavoprotein 1, mitochondrial | 0 | 0 | 1 | 0.55 | 0.40 | 0.76 |
| Q8K2B3 | Succinate dehydrogenase [ubiquinone] flavoprotein subunit, mitochondrial | 0 | 6 | 10 | 1.33 | 1.22 | 0.88 |
| Q9CQA3 | Succinate dehydrogenase [ubiquinone] iron-sulfur subunit, mitochondrial | 11 | 9 | 10 | 1.10 | 0.61 | 1.59 |
| Q8R1I1 | Cytochrome b-c1 complex subunit 9 | 0 | 0 | 11 | 0.35 | 0.31 | 0.90 |
| Q9CZ13 | Cytochrome b-c1 complex subunit 1, mitochondrial | 4 | 4 | 8 | 0.78 | 0.78 | 0.77 |
| Q9DB77 | Cytochrome b-c1 complex subunit 2, mitochondrial | 10 | 5 | 10 | 1.20 | 1.64 | 1.40 |

| | | | | | | | |
|---|---|---|---|---|---|---|---|
| Q9CR68 | Cytochrome b-c1 complex subunit Rieske, mitochondrial | 6 | 7 | 8 | 0.37 | 0.44 | 1.27 |
| P99028 | Cytochrome b-c1 complex subunit 6, mitochondrial | 4 | 11 | 6 | 0.83 | 0.98 | 0.74 |
| Q62425 | Cytochrome c oxidase subunit NDUFA4 | 7 | 2 | 5 | 0.67 | 0.37 | 0.62 |
| P19783 | Cytochrome c oxidase subunit 4 isoform 1, mitochondrial | 9 | 8 | 8 | 0.90 | 0.94 | 0.87 |
| P11240 | Cytochrome c oxidase subunit 5A, mitochondrial | 0 | 4 | 1 | 0.51 | 0.76 | 0.71 |
| P19536 | Cytochrome c oxidase subunit 5B, mitochondrial | 12 | 9 | 3 | 0.99 | 0.96 | 1.65 |
| P10818 | Cytochrome c oxidase subunit 6A1, mitochondrial | 10 | 12.5 | 12.5 | 5526.64 | 1.00 | 1.00 |
| P56391 | Cytochrome c oxidase subunit 6B1 | 0 | 1 | 3 | 0.10 | 0.28 | 0.59 |
| P11951 | Cytochrome c oxidase subunit 6C-2 | 5 | 8.5 | 5 | 1.95 | 1.57 | 2.05 |
| P48771 | Cytochrome c oxidase subunit 7A2, mitochondrial | 0 | 0 | 8 | 0.38 | 0.44 | 0.80 |
| P17665 | Cytochrome c oxidase subunit 7C, mitochondrial | 2 | 1 | 3 | 0.49 | 0.21 | 0.60 |
| Q03265 | ATP synthase subunit alpha, mitochondrial | 12 | 1 | 10 | 0.96 | 0.31 | 1.44 |
| P56480 | ATP synthase subunit beta, mitochondrial | 5 | 6 | 10 | 0.78 | 0.82 | 1.14 |
| Q91VR2 | ATP synthase subunit gamma, mitochondrial | 9 | 9 | 10 | 1.03 | 0.85 | 1.31 |
| P35434 | ATP synthase subunit delta, mitochondrial | 8 | 8 | 4 | 0.89 | 0.94 | 0.87 |
| P29418 | ATP synthase subunit epsilon, mitochondrial | 9 | 11 | 4 | 0.82 | 0.83 | 3.36 |
| Q06185 | ATP synthase subunit e, mitochondrial | 12 | 11.5 | 8 | 1.06 | 1.46 | 2.00 |
| Q6PDU7 | ATP synthase subunit g, mitochondrial | 8.5 | 3 | 12.5 | 0.71 | 0.06 | 1.09 |
| Q78IK2 | ATP synthase membrane subunit K, mitochondrial | 6 | 8 | 4 | 0.61 | 0.68 | 1.62 |
| Q9CQQ7 | ATP synthase F(0) complex subunit B1, mitochondrial | 0 | 4 | 4 | 0.62 | 0.67 | 0.63 |
| Q9DCX2 | ATP synthase subunit d, mitochondrial | 5 | 10 | 11 | 1.32 | 1.15 | 1.09 |
| Q9DB20 | ATP synthase subunit O, mitochondrial | 11 | 8 | 12 | 0.92 | 0.77 | 1.14 |
| O35143 | ATPase inhibitor, mitochondrial | 0 | 0 | 12.5 | 5.51 | 8.56 | 0.95 |
| O88986 | 2-amino-3-ketobutyrate coenzyme A ligase, mitochondrial | 8 | 5 | 2 | 0.79 | 0.40 | 0.35 |
| Q3TIU4 | 2',5'-phosphodiesterase 12 | 2 | 2 | 3 | 0.42 | 0.34 | 0.53 |

| | | | | | | | |
|---|---|---|---|---|---|---|---|
| Q99L13 | 3-hydroxyisobutyrate dehydrogenase, mitochondrial | 10 | 8.5 | 1 | 1.22 | 0.41 | 2.99 |
| Q8BWT1 | 3-ketoacyl-CoA thiolase, mitochondrial | 4 | 0 | 6 | 0.60 | 0.57 | 0.59 |
| Q99J99 | 3-mercaptopyruvate sulfurtransferase | 2 | 11 | 9 | 4.55 | 1.53 | 1.52 |
| Q9D0S9 | Adenosine 5'-monophosphoramidase HINT2 | 1 | 3 | 1 | 9.56 | 7.35 | 11.57 |
| Q8CG76 | Aflatoxin B1 aldehyde reductase member 2 | 1 | 11 | 10 | 2.62 | 1.48 | 1.37 |
| Q9Z0X1 | Apoptosis-inducing factor 1, mitochondrial | 0 | 0 | 2 | 3.53 | 2.51 | 1.91 |
| Q03265 | ATP synthase subunit alpha, mitochondrial | 12 | 1 | 10 | 0.96 | 0.31 | 1.44 |
| O08734 | Bcl-2 homologous antagonist/killer | 4 | 0 | 10 | 0.82 | 1.60 | 0.85 |
| P59017 | Bcl-2-like protein 13 | 3.5 | 0 | 8 | 8.55 | 9.37 | 4.12 |
| Q3UMR5 | Calcium uniporter protein, mitochondrial | 12 | 0 | 11.5 | 1.06 | 1.86 | 1.01 |
| P35564 | Calnexin | 0 | 0 | 7 | 0.78 | 0.75 | 0.92 |
| Q8JZN5 | Complex I assembly factor ACAD9, mitochondrial | 0 | 0 | 2 | 0.41 | 0.28 | 0.72 |
| Q9CQX2 | Cytochrome b5 type B | 2 | 0 | 11 | 2.31 | 2.86 | 1.09 |
| Q9D0M3 | Cytochrome c1, heme protein, mitochondrial | 9 | 2 | 7 | 1.09 | 1.61 | 1.39 |
| Q9Z110 | Delta-1-pyrroline-5-carboxylate synthase | 4 | 11 | 0 | 1.45 | 0.97 | 1.90 |
| O35459 | Delta(3,5)-Delta(2,4)-dienoyl-CoA isomerase, mitochondrial | 3 | 2 | 0 | 0.77 | 0.69 | 0.43 |
| P00375 | Dihydrofolate reductase | 3 | 0 | 8 | 0.55 | 0.58 | 0.78 |
| Q7TMY8 | E3 ubiquitin-protein ligase HUWE1 | 7 | 6 | 2 | 1.21 | 1.31 | 0.68 |
| Q921G7 | Electron transfer flavoprotein-ubiquinone oxidoreductase, mitochondrial | 2 | 6 | 12 | 2.35 | 1.80 | 1.00 |
| Q8K0D5 | Elongation factor G, mitochondrial | 0 | 2 | 8 | 2.70 | 2.03 | 1.87 |
| Q8BFR5 | Elongation factor Tu, mitochondrial | 2 | 4 | 8 | 1.72 | 1.58 | 1.25 |
| P42125 | Enoyl-CoA delta isomerase 1, mitochondrial | 0 | 10 | 6 | 4.75 | 1.48 | 1.62 |
| D3Z7P3 | Glutaminase kidney isoform, mitochondrial | 0 | 9 | 11 | 0.71 | 1.12 | 1.06 |
| Q80Y14 | Glutaredoxin-related protein 5, mitochondrial | 5 | 0 | 4 | 0.44 | 0.06 | 0.38 |
| P47791 | Glutathione reductase, mitochondrial | 1 | 6 | 8 | 1.76 | 1.56 | 0.91 |
| Q64521 | Glycerol-3-phosphate dehydrogenase, mitochondrial | 5 | 11 | 1 | 0.52 | 0.94 | 0.64 |
| Q9WTP7 | GTP:AMP phosphotransferase AK3, mitochondrial | 3 | 0 | 12 | 0.33 | 0.00 | 1.09 |
| Q9CQN1 | Heat shock protein 75 kDa, mitochondrial | 0 | 0 | 4 | 0.46 | 0.49 | 0.69 |
| P49710 | Hematopoietic lineage cell-specific protein | 0 | 3 | 10 | 22.16 | 7.56 | 3.18 |
| O08528 | Hexokinase-2 | 4 | 0 | 2 | 2.10 | 4.08 | 3.41 |
| Q61425 | Hydroxyacyl-coenzyme A dehydrogenase, mitochondrial | 0 | 9 | 11 | 2.14 | 1.15 | 1.31 |
| P38060 | Hydroxymethylglutaryl-CoA lyase, mitochondrial | 0 | 0 | 11 | 2.43 | 2.11 | 1.18 |

| | | | | | | | |
|---|---|---|---|---|---|---|---|
| Q60766 | Immunity-related GTPase family M protein 1 | 4 | 5 | 2 | 0.36 | 0.48 | 2.74 |
| Q9DCE9 | Immunity-related GTPase family M protein 3 | 0 | 0 | 5 | 0.00 | 0.00 | 2.64 |
| Q8BX70 | Intermembrane lipid transfer protein VPS13C | 0 | 0 | 4 | 0.41 | 0.65 | 0.78 |
| Q9JHI5 | Isovaleryl-CoA dehydrogenase, mitochondrial | 0 | 0 | 8 | 0.48 | 0.67 | 0.81 |
| Q9D338 | Large ribosomal subunit protein bL19m | 0 | 8 | 0 | 0.11 | 0.77 | 0.18 |
| Q9CQF0 | Large ribosomal subunit protein uL11m | 7 | 1 | 7 | 0.71 | 0.13 | 0.69 |
| Q9CPR5 | Large ribosomal subunit protein uL15m | 0 | 0 | 10 | 0.12 | 0.07 | 1.01 |
| Q8CGK3 | Lon protease homolog, mitochondrial | 0 | 0 | 0 | 1.98 | 2.43 | 2.04 |
| P41216 | Long-chain-fatty-acid--CoA ligase 1 | 1 | 1 | 12 | 0.42 | 0.41 | 0.95 |
| Q9QUJ7 | Long-chain-fatty-acid--CoA ligase 4 | 1 | 1 | 4 | 0.65 | 0.55 | 1.38 |
| Q8VCW8 | Medium-chain acyl-CoA ligase ACSF2, mitochondrial | 5 | 1 | 9 | 0.72 | 0.26 | 0.92 |
| P47802 | Metaxin-1 | 9 | 0 | 7 | 1.00 | 0.00 | 0.57 |
| Q7TNS2 | MICOS complex subunit Mic10 | 2 | 10 | 3 | 0.62 | 1.04 | 0.43 |
| Q9CRB9 | MICOS complex subunit Mic19 | 2 | 2 | 11 | 5.39 | 4.47 | 1.01 |
| D4A7N1 | MICOS complex subunit Mic25 | 9 | 8 | 12 | 0.82 | 2.06 | 1.15 |
| Q78IK4 | MICOS complex subunit Mic27 | 4 | 0 | 12 | 0.78 | 0.41 | 1.04 |
| Q8CAQ8 | MICOS complex subunit Mic60 | 0 | 0 | 5 | 1.71 | 1.73 | 1.30 |
| Q9CR62 | Mitochondrial 2-oxoglutarate/malate carrier protein | 0 | 0 | 11 | 0.62 | 0.61 | 0.94 |
| Q8BMD8 | Mitochondrial adenyl nucleotide antiporter SLC25A24 | 1 | 6 | 2 | 2.60 | 1.65 | 4.35 |
| Q9Z2Z6 | Mitochondrial carnitine/acylcarnitine carrier protein | 0 | 5 | 8 | 0.52 | 0.75 | 0.83 |
| Q9QZD8 | Mitochondrial dicarboxylate carrier | 0 | 0 | 0 | 0.56 | 0.46 | 0.69 |
| P62075 | Mitochondrial import inner membrane translocase subunit Tim13 | 0 | 0 | 2 | 0.48 | 0.09 | 0.59 |
| Q9CYG7 | Mitochondrial import receptor subunit TOM34 | 1 | 4 | 8 | 1.55 | 1.31 | 1.31 |
| Q9CZW5 | Mitochondrial import receptor subunit TOM70 | 0 | 0 | 2 | 0.48 | 0.63 | 0.78 |
| Q920A7 | Mitochondrial inner membrane m-AAA protease component AFG3L1 | 11 | 6 | 9 | 0.88 | 1.38 | 1.22 |
| Q8JZQ2 | Mitochondrial inner membrane m-AAA protease component AFG3L2 | 8 | 12 | 1 | 0.82 | 1.03 | 1.55 |
| Q9DCN2 | NADH-cytochrome b5 reductase 3 | 10 | 8 | 0 | 1.07 | 0.75 | 1.94 |
| Q9WV85 | Nucleoside diphosphate kinase 3 | 2 | 0 | 6 | 2.28 | 2.13 | 0.26 |
| P29758 | Ornithine aminotransferase, mitochondrial | 11 | 0 | 5 | 1.00 | 0.60 | 0.79 |
| P30416 | Peptidyl-prolyl cis-trans isomerase FKBP4 | 4 | 0 | 8 | 0.56 | 0.43 | 0.71 |
| P99029 | Peroxiredoxin-5, mitochondrial | 5 | 9 | 0 | 1.42 | 1.13 | 3.25 |

| | | | | | | | |
|---|---|---|---|---|---|---|---|
| Q8BH04 | Phosphoenolpyruvate carboxykinase [GTP], mitochondrial | 10 | 3 | 1 | 0.95 | 0.70 | 1.68 |
| P67778 | Prohibitin 1 | 8 | 7 | 2 | 0.87 | 0.86 | 1.26 |
| O35129 | Prohibitin-2 | 0 | 0 | 0 | 3.54 | 2.35 | 2.30 |
| Q9CR98 | Protein FAM136A | 2 | 4 | 7 | 1.87 | 1.97 | 1.59 |
| P28867 | Protein kinase C delta type | 0 | 3 | 3 | 4.61 | 3.25 | 3.03 |
| O55125 | Protein NipSnap homolog 1 | 1 | 4 | 6 | 3.36 | 2.89 | 1.54 |
| P51175 | Protoporphyrinogen oxidase | 9 | 2 | 9 | 1.18 | 1.25 | 1.26 |
| P60603 | Reactive oxygen species modulator 1 | 1 | 4 | 6 | 17.71 | 7.67 | 5.38 |
| P15651 | Short-chain specific acyl-CoA dehydrogenase, mitochondrial | 4 | 1 | 3 | 0.60 | 0.26 | 0.53 |
| Q9DBL1 | Short/branched chain specific acyl-CoA dehydrogenase, mitochondrial | 1 | 0 | 9 | 0.52 | 0.45 | 0.86 |
| Q9JM90 | Signal-transducing adaptor protein 1 | 0 | 4 | 7 | 6.99 | 3.30 | 2.16 |
| Q8BK72 | Small ribosomal subunit protein mS27 | 2 | 9 | 3 | 0.41 | 0.66 | 0.30 |
| Q924T2 | Small ribosomal subunit protein uS2m | 6 | 2 | 4 | 0.53 | 0.17 | 0.37 |
| Q64442 | Sorbitol dehydrogenase | 6 | 2 | 9.5 | 1.45 | 2.00 | 0.71 |
| Q9JIA7 | Sphingosine kinase 2 | 0 | 11.5 | 0 | 8.54 | 1.75 | 17.78 |
| P32020 | Sterol carrier protein 2 | 9 | 0 | 8 | 0.82 | 0.30 | 0.77 |
| P38647 | Stress-70 protein, mitochondrial | 0 | 0 | 2 | 1.66 | 1.66 | 1.34 |
| Q9Z2I9 | Succinate--CoA ligase [ADP-forming] subunit beta, mitochondrial | 4 | 0 | 11 | 1.49 | 1.33 | 1.00 |
| Q9WUM5 | Succinate--CoA ligase [ADP/GDP-forming] subunit alpha, mitochondrial | 1 | 8.5 | 7 | 4.83 | 1.58 | 2.00 |
| Q9Z2I8 | Succinate--CoA ligase [GDP-forming] subunit beta, mitochondrial | 8 | 7 | 12 | 1.16 | 0.76 | 0.99 |
| Q9R112 | Sulfide:quinone oxidoreductase, mitochondrial | 0 | 9 | 11 | 0.67 | 0.98 | 1.02 |
| P09671 | Superoxide dismutase [Mn], mitochondrial | 9 | 4 | 1 | 1.27 | 1.84 | 3.21 |
| Q921F2 | TAR DNA-binding protein 43 | 2 | 2 | 11 | 2.26 | 1.89 | 1.18 |
| Q9JMH6 | Thioredoxin reductase 1, cytoplasmic | 0 | 0 | 6 | 5.69 | 9.24 | 1.62 |
| Q9JLT4 | Thioredoxin reductase 2, mitochondrial | 2.5 | 0 | 7.5 | 398796.25 | 572990.51 | 126971.43 |
| Q8VBT0 | Thioredoxin-related transmembrane protein 1 | 12 | 0 | 5 | 1.00 | 1.66 | 0.78 |
| Q9CQN6 | Transmembrane protein 14C | 7 | 1 | 10 | 0.75 | 2.32 | 0.81 |
| Q8BMS1 | Trifunctional enzyme subunit alpha, mitochondrial | 10 | 2 | 8 | 0.87 | 0.63 | 0.83 |
| O89023 | Tripeptidyl-peptidase 1 | 0 | 0 | 5 | 0.00 | 0.00 | 0.35 |
| P00520 | Tyrosine-protein kinase ABL1 | 6 | 0 | 10 | 2.37 | 4.58 | 0.65 |
| Q02053 | Ubiquitin-like modifier-activating enzyme 1 | 1 | 6 | 8 | 1.51 | 1.24 | 1.12 |
| Q3U5Q7 | UMP-CMP kinase 2, mitochondrial | 10.5 | 2 | 0 | 1.13 | 2.51 | 8.24 |

| | | | | | | | |
|---|---|---|---|---|---|---|---|
| P50544 | Very long-chain specific acyl-CoA dehydrogenase, mitochondrial | 2 | 3 | 10.5 | 6.58 | 2.79 | 1.59 |

Supplementary Table 9: List of glycolytic and pentose phosphate pathway proteins modulated in response to at least one treatment
Modulated proteins appear with their name in color. Un-modulated but related proteins (e.g. other proteins of the same pathway) appear in black

| accession number | name | MWU 4x5 | MWU 4x10 | MWU 20R | ratio 4x5 | ratio 4x10 | ratio 20R |
|---|---|---|---|---|---|---|---|
| O08528 | Hexokinase-2 | 4 | 0 | 2 | 2.10 | 4.08 | 3.41 |
| P05064 | Fructose-bisphosphate aldolase A | 1 | 4 | 3 | 0.59 | 1.37 | 0.59 |
| P05063 | Fructose-bisphosphate aldolase C | 1 | 0 | 4 | 2.06 | 1.76 | 1.34 |
| P06151 | L-lactate dehydrogenase A chain | 0 | 2 | 2 | 2.38 | 1.82 | 1.66 |
| P09411 | Phosphoglycerate kinase 1 | 3 | 1 | 4 | 1.34 | 1.31 | 1.29 |
| P16858 | Glyceraldehyde-3-phosphate dehydrogenase | 2 | 4 | 10 | 2.08 | 2.14 | 1.31 |
| P17182 | Alpha-enolase | 0 | 0 | 4 | 2.12 | 1.87 | 1.58 |
| P17751 | Triosephosphate isomerase | 0 | 1 | 11 | 2.18 | 1.92 | 1.09 |
| P28271 | Cytoplasmic aconitate hydratase | 0 | 2 | 5 | 0.52 | 0.18 | 0.63 |
| P40142 | Transketolase | 2 | 0 | 6 | 2.52 | 2.73 | 1.43 |
| P52480 | Pyruvate kinase PKM | 3 | 5 | 2 | 1.44 | 1.18 | 1.56 |
| P53657 | Pyruvate kinase PKLR | 10 | 0 | 10 | 0.00003 | 90.53 | 0.00003 |
| Q00612 | Glucose-6-phosphate 1-dehydrogenase X | 0 | 1 | 2 | 1.75 | 1.73 | 1.48 |
| Q3TRM8 | Hexokinase-3 | 11 | 7 | 2 | 1.07 | 1.15 | 1.56 |
| Q8VDL4 | ADP-dependent glucokinase | 1 | 8 | 5 | 0.72 | 0.82 | 0.90 |
| Q93092 | Transaldolase | 4 | 1 | 8 | 1.27 | 1.42 | 1.14 |
| Q9D0F9 | Phosphoglucomutase-1 | 12 | 4 | 2 | 1.01 | 1.45 | 2.16 |
| P09041 | Phosphoglycerate kinase 2 | 12 | 8 | 11 | 1.19 | 0.42 | 2.49 |
| P17710 | Hexokinase-1 | 10 | 7 | 11 | 0.89 | 0.71 | 0.99 |
| Q9CQ60 | 6-phosphogluconolactonase | 6 | 6 | 11 | 1.60 | 1.48 | 1.17 |
| Q9DCD0 | 6-phosphogluconate dehydrogenase, decarboxylating | 5 | 12 | 3 | 1.45 | 0.94 | 1.63 |
| P12382 | ATP-dependent 6-phosphofructokinase, liver type | 11 | 11 | 3 | 1.30 | 1.05 | 2.40 |
| P47857 | ATP-dependent 6-phosphofructokinase, muscle type | 8 | 11.5 | 11 | 1.96 | 1.62 | 0.69 |
| Q9WUA3 | ATP-dependent 6-phosphofructokinase, platelet type | 7 | 7 | 7 | 1.24 | 1.17 | 1.44 |
| P06745 | Glucose-6-phosphate isomerase | 6 | 7 | 9 | 1.31 | 1.16 | 1.10 |
| P47968 | Ribose-5-phosphate isomerase | 11 | 11 | 5 | 1.07 | 0.73 | 0.26 |

Supplementary Table 10: List of  proteinsimplicate in oxidative stress response and  modulated in response to at least one treatment
Modulated proteins appear with their name in color. Un-modulated but related proteins (e.g. other proteins of the same pathway) appear in black

| accession number | Name | MWU 4x5 | MWU 4x10 | MWU 20R | ratio 4x5 | ratio 4x10 | ratio 20R |
|---|---|---|---|---|---|---|---|
| P08228 | Superoxide dismutase [Cu-Zn] | 3 | 6 | 0 | 0.65 | 1.33 | 0.71 |
| P09671 | Superoxide dismutase [Mn], mitochondrial | 9 | 4 | 1 | 1.27 | 1.84 | 3.21 |
| P04762 | Catalase | 0 | 1 | 9 | 4.47 | 3.52 | 0.94 |
| P35700 | Peroxiredoxin-1 | 4 | 11 | 0 | 1.58 | 1.15 | 2.23 |
| Q61171 | Peroxiredoxin-2 | 12 | 10 | 7 | 1.11 | 0.91 | 0.89 |
| P20108 | Peroxiredoxin-3 | 6 | 3 | 8 | 0.69 | 0.59 | 0.81 |
| O08807 | Peroxiredoxin-4 | 8 | 10 | 5 | 0.41 | 0.55 | 0.49 |
| P99029 | Peroxiredoxin-5, mitochondrial | 5 | 9 | 0 | 1.42 | 1.13 | 3.25 |
| O08709 | Peroxiredoxin-6 | 8 | 6 | 6 | 0.98 | 0.89 | 0.86 |
| P47791 | Glutathione reductase, mitochondrial | 1 | 6 | 8 | 1.76 | 1.56 | 0.91 |
| Q9QUH0 | Glutaredoxin-1 | 11 | 11 | 1 | 0.94 | 0.93 | 1.82 |
| Q80Y14 | Glutaredoxin-related protein 5, mitochondrial | 5 | 0 | 4 | 0.44 | 0.06 | 0.38 |
| Q8VBT0 | Thioredoxin-related transmembrane protein 1 | 12 | 0 | 5 | 1.00 | 1.66 | 0.78 |
| P10639 | Thioredoxin | 0 | 1 | 6 | 16.02 | 8.75 | 1.87 |
| Q498E0 | Thioredoxin domain-containing protein 12 | 6 | 12 | 10 | 1.90 | 0.89 | 1.40 |
| Q9CQM5 | Thioredoxin domain-containing protein 17 | 10.5 | 9 | 1 | 0.51 | 0.12 | 4.33 |
| Q91W90 | Thioredoxin domain-containing protein 5 | 6 | 3 | 9 | 0.41 | 0.06 | 0.86 |
| Q8CDN6 | Thioredoxin-like protein 1 | 0 | 0 | 4 | 7.44 | 8.01 | 1.77 |
| Q9JMH6 | Thioredoxin reductase 1, cytoplasmic | 0 | 0 | 6 | 5.69 | 9.24 | 1.62 |
| Q9JLT4 | Thioredoxin reductase 2, mitochondrial | 2.5 | 0 | 7.5 | 398796.25 | 572990.51 | 126971.43 |
| P11352 | Glutathione peroxidase 1 | 9 | 6 | 9 | 1.13 | 0.84 | 0.89 |
| O70325 | Phospholipid hydroperoxide glutathione peroxidase GPX4 | 7 | 10 | 5 | 1.55 | 1.21 | 1.54 |
| P19468 | Glutamate--cysteine ligase catalytic subunit | 8 | 10 | 9 | 1.95 | 0.00 | 3.12 |
| O09172 | Glutamate--cysteine ligase regulatory subunit | 2 | 3 | 0 | 3.94 | 3.66 | 3.53 |
| P46413 | Glutathione synthetase | 3.5 | 11.5 | 9 | 3.39 | 1.30 | 2.06 |

Supplementary Table 11: List of lysosomal proteins modulated in response to at least one treatment
Modulated proteins appear with their name in color. Un-modulated but related proteins (e.g. other proteins of the same family) appear in black

| accession | name | MWU 4x5 | MWU 4x10 | MWU 20R | ratio 4x5 | ratio 4x10 | ratio 20R |
|---|---|---|---|---|---|---|---|
| P45377 | Aldose reductase-related protein 2 | 2 | 1 | 6 | 4.73 | 3.78 | 2.55 |
| P29416 | Beta-hexosaminidase subunit alpha | 0 | 0 | 0 | 0.24 | 0.48 | 0.61 |
| P20060 | Beta-hexosaminidase subunit beta | 0 | 0 | 0 | 0.40 | 0.49 | 0.59 |
| Q9WVJ3 | Carboxypeptidase Q | 0 | 0 | 3 | 1.51E-06 | 1.51E-06 | 0.41 |
| Q9ESY9 | Gamma-interferon-inducible lysosomal thiol reductase | 0 | 2 | 1 | 0.57 | 0.52 | 0.62 |
| P63017 | Heat shock cognate 71 kDa protein | 0 | 0 | 0 | 0.82 | 0.68 | 0.72 |
| Q60766 | Immunity-related GTPase family M protein 1 | 4 | 5 | 2 | 0.36 | 0.48 | 2.74 |
| Q8BX70 | Intermembrane lipid transfer protein VPS13C | 0 | 0 | 4 | 0.41 | 0.65 | 0.78 |
| Q8VEB4 | Lysosomal phospholipase A and acyltransferase | 7 | 2 | 6 | 0.57 | 0.17 | 0.57 |
| P16675 | Lysosomal protective protein | 3 | 0 | 8 | 0.62 | 0.44 | 0.91 |
| O35114 | Lysosome membrane protein 2 | 2 | 0 | 7 | 0.23 | 1.96E-06 | 1.29 |
| P11438 | Lysosome-associated membrane glycoprotein 1 | 7 | 9 | 8 | 0.55 | 0.86 | 0.85 |
| P17047 | Lysosome-associated membrane glycoprotein 2 | 1 | 2 | 9 | 0.46 | 0.48 | 0.92 |
| P31996 | Macrosialin | 0 | 8 | 9 | 0.54 | 0.68 | 0.81 |
| P02802 | Metallothionein-1 | 0 | 12 | 12 | 93.66 | 2.73 | 12.14 |
| Q8BFR4 | N-acetylglucosamine-6-sulfatase | 1 | 9 | 12 | 0.71 | 0.87 | 0.91 |
| O09043 | Napsin-A | 0 | 1 | 9 | 0.61 | 0.57 | 0.90 |
| Q9Z0J0 | NPC intracellular cholesterol transporter 2 | 0 | 0 | 9 | 0.21 | 0.24 | 0.86 |
| Q61207 | Prosaposin | 3 | 0 | 6 | 0.56 | 0.50 | 0.82 |
| O88668 | Protein CREG1 | 2 | 10 | 3 | 2.89 | 2.19 | 2.43 |
| Q3TCN2 | Putative phospholipase B-like 2 | 0 | 1 | 7 | 0.34 | 0.40 | 0.73 |
| Q9CQ22 | Ragulator complex protein LAMTOR1 | 5 | 2 | 12 | 0.43 | 0.15 | 0.77 |
| Q9JHS3 | Ragulator complex protein LAMTOR2 | 9 | 7 | 11 | 1.46 | 0.27 | 1.13 |
| P35283 | Ras-related protein Rab-12 | 0 | 5 | 0 | 0.00 | 0.57 | 0.00 |
| Q91V41 | Ras-related protein Rab-14 | 0 | 0 | 2 | 0.59 | 0.65 | 0.70 |
| P08228 | Superoxide dismutase [Cu-Zn] | 3 | 6 | 0 | 0.65 | 1.33 | 0.71 |
| O89023 | Tripeptidyl-peptidase 1 | 0 | 0 | 5 | 0.00 | 0.00 | 0.35 |
| Q78T54 | Vacuolar ATPase assembly integral membrane protein Vma21 | 7 | 1 | 12 | 4.32 | 6.55 | 1.32 |
| Q9EQH3 | Vacuolar protein sorting-associated protein 35 | 11 | 2 | 7 | 0.79 | 0.45 | 1.16 |
| O70404 | Vesicle-associated membrane protein 8 | 0 | 3 | 0 | 100.85 | 73.97 | 65.34 |
| P10605 | Cathepsin B | 10 | 8 | 11 | 0.69 | 0.51 | 0.90 |
| P18242 | Cathepsin D | 1 | 6 | 12 | 0.64 | 0.70 | 0.98 |

| | | | | | | | |
|---|---|---|---|---|---|---|---|
| O70370 | Cathepsin S | 3 | 7 | 12 | 0.48 | 0.58 | 0.88 |
| Q9WUU7 | Cathepsin Z | 10 | 9 | 11 | 1.06 | 1.15 | 0.94 |
| P97821 | Dipeptidyl peptidase 1 | 5 | 4 | 11 | 0.43 | 0.27 | 1.02 |
| P06797 | Procathepsin L | 5 | 11 | 2 | 0.74 | 1.00 | 2.18 |
| P25286 | V-type proton ATPase 116 kDa subunit a 1 | 8 | 7 | 3 | 0.61 | 0.34 | 2.14 |
| P63081 | V-type proton ATPase 16 kDa proteolipid subunit c | 2 | 11 | 12 | 0.67 | 0.90 | 1.02 |
| P51863 | V-type proton ATPase subunit d 1 | 3 | 3 | 6 | 0.68 | 0.77 | 0.83 |
| P50516 | V-type proton ATPase catalytic subunit A | 2 | 5 | 9 | 0.67 | 0.77 | 1.06 |
| P62814 | V-type proton ATPase subunit B, brain isoform | 10 | 7 | 9 | 0.92 | 1.19 | 1.10 |
| Q5FVI6 | V-type proton ATPase subunit C 1 | 9 | 3 | 4 | 0.83 | 0.52 | 1.55 |
| P57746 | V-type proton ATPase subunit D | 3 | 8 | 9 | 5.08 | 2.60 | 2.43 |
| P50518 | V-type proton ATPase subunit E 1 | 5 | 5 | 4 | 1.48 | 1.48 | 1.85 |
| Q9CR51 | V-type proton ATPase subunit G 1 | 2 | 6 | 6 | 3.07 | 2.53 | 1.97 |
| Q8BVE3 | V-type proton ATPase subunit H | 0 | 0 | 0 | 6.20 | 6.05 | 3.81 |

Supplementary Table 12: List of proteins associated with immunity and modulated in response to at least one treatment

| accession | nom | MWU 4x5 | MWU 4x10 | MWU 20R | ratio 4x5 | ratio 4x10 | Ratio 20R |
|---|---|---|---|---|---|---|---|
| A1L314 | Macrophage-expressed gene 1 protein | 0 | 4 | 4 | 0.46 | 0.59 | 1.65 |
| O70133 | ATP-dependent RNA helicase A | 10 | 2 | 8 | 0.89 | 0.60 | 0.83 |
| P09581 | Macrophage colony-stimulating factor 1 receptor | 0 | 0 | 4 | 0.39 | 0.43 | 0.59 |
| P0DOV2 | Interferon-activable protein 204 | 8 | 11.5 | 1 | 0.59 | 0.64 | 5.44 |
| P10107 | Annexin A1 | 4 | 0 | 7 | 0.87 | 0.62 | 0.93 |
| P10810 | Monocyte differentiation antigen CD14 | 0 | 0 | 4 | 0.23 | 0.17 | 0.74 |
| P16110 | Galectin-3 | 0 | 0 | 2 | 9.59 | 9.48 | 2.80 |
| P16858 | Glyceraldehyde-3-phosphate dehydrogenase | 2 | 4 | 10 | 2.08 | 2.14 | 1.31 |
| P20491 | High affinity immunoglobulin epsilon receptor subunit gamma | 1 | 0 | 2 | 5.08 | 4.04 | 3.53 |
| P25911 | Tyrosine-protein kinase Lyn | 1 | 0 | 4 | 0.64 | 0.49 | 0.77 |
| P26151 | High affinity immunoglobulin gamma Fc receptor I | 0 | 0 | 8 | 0.41 | 0.00 | 1.17 |
| P63158 | High mobility group protein B1 | 2 | 12 | 9 | 0.40 | 0.92 | 0.64 |
| P97379 | Ras GTPase-activating protein-binding protein 2 | 0 | 5.5 | 12 | 11.17 | 11.77 | 3.41 |
| P97855 | Ras GTPase-activating protein-binding protein 1 | 0 | 5 | 0 | 1.51 | 1.42 | 0.59 |
| Q2EMV9 | Protein mono-ADP-ribosyltransferase PARP14 | 0 | 1 | 1 | 0.08 | 0.25 | 0.47 |
| Q60766 | Immunity-related GTPase family M protein 1 | 4 | 5 | 2 | 0.36 | 0.48 | 2.74 |
| Q60875 | Rho guanine nucleotide exchange factor 2 | 0 | 0 | 4 | 2.60 | 2.19 | 1.82 |
| Q61990 | Poly(rC)-binding protein 2 | 3 | 0 | 12 | 1.34 | 1.86 | 0.99 |
| Q64339 | Ubiquitin-like protein ISG15 | 0 | 0 | 9 | 98.31 | 119.51 | 6.47 |
| Q8BG07 | 5'-3' exonuclease PLD4 | 0 | 0 | 0 | 0.36 | 0.41 | 0.61 |
| Q8BPX9 | Solute carrier family 15 member 3 | 2 | 7 | 2 | 0.48 | 1.27 | 1.99 |
| Q8C2Q3 | RNA-binding protein 14 | 1 | 1 | 11 | 4.57 | 3.61 | 0.88 |
| Q8K310 | Matrin-3 | 2 | 4 | 12 | 1.89 | 2.08 | 1.30 |
| Q8VC04 | Transmembrane protein 106A | 4 | 2 | 0 | 0.62 | 0.60 | 2.44 |
| Q99J93 | Interferon-induced transmembrane protein 2 | 0 | 0 | 12 | 0.08 | 0.14 | 1.03 |
| Q9CQW9 | Interferon-induced transmembrane protein 3 | 0 | 0 | 12 | 0.08 | 0.54 | 1.29 |
| Q9CR67 | Transmembrane protein 33 | 0 | 3 | 4 | 8.98 | 4.27 | 4.19 |
| Q9D906 | Ubiquitin-like modifier-activating enzyme ATG7 | 0 | 0 | 0 | 0.57 | 0.60 | 0.53 |
| Q9DAU1 | Protein canopy homolog 3 | 11 | 3 | 0 | 0.93 | 1.30 | 0.48 |
| Q9DCE9 | Immunity-related GTPase family M protein 3 | 0 | 0 | 5 | 0.00 | 0.00 | 2.64 |
| Q9Z0E6 | Guanylate-binding protein 2 | 0 | 0 | 4 | 0.34 | 0.23 | 0.33 |
| P31996 | Macrosialin | 0 | 8 | 9 | 0.54 | 0.68 | 0.81 |

| | | | | | | | |
|---|---|---|---|---|---|---|---|
| P24063 | Integrin alpha-L | 2 | 10 | 6 | 0.15 | 0.82 | 0.57 |
| P05555 | Integrin alpha-M | 0 | 0 | 10 | 0.48 | 0.28 | 1.17 |
| P09055 | Integrin beta-1 | 0 | 0 | 10 | 0.63 | 0.62 | 0.89 |
| P11835 | Integrin beta-2 | 0 | 0 | 11 | 0.55 | 0.61 | 1.04 |
| P34960 | Macrophage metalloelastase | 0 | 1 | 0 | 3.34 | 3.22 | 2.13 |
| Q07797 | Galectin-3-binding protein | 0 | 2 | 4 | 0.36 | 0.62 | 1.33 |
| P18572 | Basigin | 7 | 5 | 1 | 0.90 | 1.25 | 0.63 |
| Q9EQP2 | EH domain-containing protein 4 | 2 | 0 | 8 | 0.51 | 0.30 | 1.24 |
| O08573 | Galectin-9 | 2 | 3 | 9 | 0.13 | 0.18 | 1.30 |
| P46460 | Vesicle-fusing ATPase | 0 | 0 | 7 | 0.52 | 0.33 | 1.12 |
| Q62422 | Osteoclast-stimulating factor 1 | 0 | 0 | 1 | 0.16 | 0.45 | 0.37 |
| Q2YFS3 | Paired immunoglobulin-like type 2 receptor alpha | 0 | 0 | 2 | 0.21 | 0.18 | 0.52 |
| Q91ZR2 | Sorting nexin-18 | 2 | 0 | 6 | 4.87 | 8.85 | 2.39 |
| Q91VH2 | Sorting nexin-9 | 8 | 0 | 11.5 | 3.94 | 11.88 | 1.95 |
| O70404 | Vesicle-associated membrane protein 8 | 0 | 3 | 0 | 100.85 | 73.97 | 65.34 |

Supplementary Table 13: List of proteinsimplicated to the response to chemical stess and  modulated in response to at least one treatment
Modulated proteins appear with their name in color. Un-modulated but related proteins (e.g. other proteins of the same family) appear in black

| accession | nom | MWU 4x5 | MWU 4x10 | MWU 20 R | ratio 4x5 | ratio 4x10 | Ratio 20 R |
|---|---|---|---|---|---|---|---|
| P48758 | Carbonyl reductase [NADPH] 1 | 4 | 0 | 2 | 1.42 | 2.79 | 1.86 |
| Q8K354 | Carbonyl reductase [NADPH] 3 | 0 | 3.5 | 11 | 4.61 | 3.54 | 1.45 |
| Q9R0P3 | S-formylglutathione hydrolase | 0 | 0 | 0 | 2.19 | 3.08 | 1.93 |
| Q9CPU0 | Lactoylglutathione lyase | 7 | 10 | 11 | 2.46 | 0.32 | 0.73 |
| P10649 | Glutathione S-transferase Mu 1 | 6 | 8 | 9 | 1.69 | 0.25 | 0.60 |
| P48774 | Glutathione S-transferase Mu 5 | 1 | 1 | 9 | 2.70 | 3.15 | 1.49 |
| O09131 | Glutathione S-transferase omega-1 | 5 | 12.5 | 10 | 36385.49 | 1.00 | 7450.05 |
| Q9CPU4 | Glutathione S-transferase 3, mitochondrial | 7 | 3.5 | 11.5 | 0.46 | 0.18 | 0.82 |
| Q9R112 | Sulfide:quinone oxidoreductase, mitochondrial | 0 | 9 | 11 | 0.67 | 0.98 | 1.02 |
| P47199 | Quinone oxidoreductase | 8 | 10.5 | 9 | 1.37 | 1.19 | 1.29 |
| Q99L04 | Dehydrogenase/reductase SDR family member 1 | 9 | 12 | 7 | 1.19 | 0.87 | 1.71 |
| Q9JII6 | Aldo-keto reductase family 1 member A1 | 4 | 6 | 10 | 1.36 | 1.17 | 1.10 |
| P45376 | Aldo-keto reductase family 1 member B1 | 0 | 5 | 6 | 1.79 | 1.39 | 0.83 |
| P45377 | Aldose reductase-related protein 2 | 2 | 1 | 6 | 4.73 | 3.78 | 2.55 |
| Q8CG76 | Aflatoxin B1 aldehyde reductase member 2 | 1 | 11 | 10 | 2.62 | 1.48 | 1.37 |
| Q3T1L0 | Aldehyde dehydrogenase family 16 member A1 | 11 | 10 | 8 | 1.60 | 1.39E-05 | 1.61 |
| P47740 | Aldehyde dehydrogenase family 3 member A2 | 10 | 5 | 5 | 1.22 | 6.24E-06 | 2.24 |
| Q5XI42 | Aldehyde dehydrogenase family 3 member B1 | 12 | 6 | 10 | 0.92 | 0.59 | 1.01 |
| Q91Z53 | Glyoxylate reductase/hydroxypyruvate reductase | 12 | 9 | 11 | 1.16 | 1.19 | 0.88 |
| Q64105 | Sepiapterin reductase | 9 | 5 | 9 | 1.43 | 0.48 | 1.34 |
| O35379 | Multidrug resistance-associated protein 1 | 6 | 12 | 4 | 1.72 | 1.80 | 1.86 |
| B2RX12 | ATP-binding cassette sub-family C member 3 | 10.5 | 12 | 7 | 0.90 | 0.76 | 1.56 |
| O88269 | ATP-binding cassette sub-family C member 6 | 5 | 10 | 5 | 155762.95 | 59008.43 | 138037.70 |
| P55096 | ATP-binding cassette sub-family D member 3 | 8 | 4 | 6 | 0.56 | 1.50 | 1.14 |
| P61222 | ATP-binding cassette sub-family E member 1 | 11 | 11 | 12 | 1.70 | 0.77 | 1.29 |
| Q6MG08 | ATP-binding cassette sub-family F member 1 | 12 | 9 | 12 | 0.91 | 2.20 | 0.73 |

| | | | | | | | |
|---|---|---|---|---|---|---|---|
| P02802 | Metallothionein-1 | 0 | 12 | 12 | 93.66 | 2.73 | 12.14 |
| P09528 | Ferritin heavy chain | 12 | 11 | 6 | 0.73 | 0.77 | 1.51 |
| P29391 | Ferritin light chain 1 | 9 | 10 | 9 | 0.50 | 0.60 | 0.75 |
| P14901 | Heme oxygenase 1 | 5 | 5 | 7.5 | 231535.76 | 285077.11 | 225178.00 |
| O70252 | Heme oxygenase 2 | 12 | 5 | 11 | 0.99 | 0.71 | 1.01 |
| Q9CY64 | Biliverdin reductase A | 9 | 12 | 1 | 1.14 | 1.02 | 1.67 |
| Q923D2 | Flavin reductase (NADPH) | 0 | 0 | 11.5 | 8.28 | 13.41 | 0.90 |
| Q922D8 | C-1-tetrahydrofolate synthase, cytoplasmic | 4 | 7 | 0 | 0.75 | 0.86 | 0.51 |
| P00375 | Dihydrofolate reductase | 3 | 0 | 8 | 0.55 | 0.58 | 0.78 |
